# KDM4A promotes the progression of neuroendocrine prostate cancer

**DOI:** 10.1101/2022.05.14.491739

**Authors:** Celia Sze Ling Mak, Ming Zhu, Xin Liang, Feng Wang, Fei Yuan, Anh G Hoang, Xingzhi Song, Peter Shepherd, Derek Liang, Jessica Suh, Bijeta Pradhan, Jiwon Park, Miao Zhang, Eric Metzger, Roland Schüle, Abhinav K. Jain, Ellen Karasik, Barbara A. Foster, Min Gyu Lee, Paul Corn, Christopher J. Logothetis, Ana Aparicio, Nora Navone, Patricia Troncoso, Zhi Tan, Jianhua Zhang, Sue-Hwa Lin, Guocan Wang

## Abstract

Neuroendocrine prostate cancer (NEPC) represents one of the most lethal forms of prostate cancer (PCa) and lacks life-prolonging treatment. The incidence of NEPC is increased due to the widespread use of AR pathway inhibitors (ARPIs) in the treatment of non-metastatic CRPC and hormone-sensitive metastatic tumors. Here, we identified histone lysine demethylase KDM4A as a key player in NEPC progression and an effective therapeutic target. We found that KDM4A mRNA and protein are overexpressed in human and mouse NEPC compared to prostate adenocarcinoma. Knockdown (KD) or knockout (KO) of *KDM4A* in NEPC cell lines suppressed cancer cell growth *in vitro* and *in vivo*. Mechanistically, we found that KDM4A promotes NEPC progression, in part, through direct transcriptional regulation of *MYC*. We showed that *MYC* is hyper-activated in human and mouse NEPC. *KDM4A* KD led to suppression of MYC signaling. *MYC* KD or inhibition profoundly suppressed NEPC cell proliferation. Furthermore, a potent pan-KDM4 inhibitor QC6352 significantly reduced NEPC cell growth *in vitro* and *in vivo*. Taken together, we demonstrated that KDM4A is an important regulator of NEPC progression and targeting KDM4A may potentially be an effective therapeutic strategy for NEPC.

**Significance:** Neuroendocrine prostate cancer (NEPC) is a highly aggressive prostate cancer subtype that is resistant to potent androgen receptor pathway inhibitors (ARPIs) and currently lacks effective therapeutic options. Histone lysine demethylase KDM4A is an important epigenetic regulator of gene expression in development and cancer. In this study, we show that KDM4A is highly expressed in NEPC and is required for NEPC proliferation, anchorage-independent growth, and *in vivo* growth, which is in part mediated through the regulation of MYC expression. Importantly, we demonstrate that inhibition of KDM4A significantly impairs NEPC growth in preclinical models. Thus, our findings provide valuable insights into the molecular mechanisms underlying NEPC progression and offer a rationale for clinical trials with KDM4 inhibitor in NEPC patients.

## INTRODUCTION

Neuroendocrine prostate cancer (NEPC) is a highly lethal subtype of castration-resistant prostate cancer (CRPC) with a median survival of seven months after initial diagnosis,^1-4^ as compared to a median survival of 13 to 31 months in castration-resistant prostate adenocarcinoma, the more common subtype of CRPC.^5^ NEPC is characterized by attenuated androgen receptor (AR) signaling, the expression of neuroendocrine lineage markers (e.g., synaptophysin), uncontrolled hyperproliferation (e.g., high Ki67 index), and widespread metastasis (e.g., bone, liver, and lung). De novo NEPCs are rare (2%-5%). The widespread use of AR pathway inhibitors (ARPIs) in non-metastatic CRPC and hormone-sensitive metastatic tumors has led to an increase in the incidence of NEPC.^6^ Due to the lack of life-prolonging systemic therapies, there is an *urgent need* to better understand the mechanisms underlying the pathogenesis of NEPC.

Recent evidence suggests that epigenetic dysregulation is a hallmark of NEPC.^4,6^ Aberrantly expressed transcription factors (TFs) and regulators of DNA methylation and histone modification have been found in NEPC preclinical models and patient samples.^4,6^ Among the various epigenetic regulatory mechanisms, histone lysine methylation, which is balanced by writers (histone lysine methyltransferase [KMT]) and erasers (histone lysine demethylases [KDM]), plays an important role in development and cancer, including prostate.^7^ Multiple lysine methylation modifiers (e.g., EZH2, WHSC1, DOT1L, LSD1/KDM1A, KDM4A, KDM4B) have been implicated in prostate cancer progression.^8-12^ However, their roles in NEPC are just starting to emerge. For example, histone lysine methyltransferase EZH2 has recently been implicated as a key player in NEPC.^6,13-18^ In contrast, it is currently unknown whether KDMs play a role in NEPC progression.

KDM4A is one of the KDM family genes and has been shown to regulate cell cycle, replication, and DNA damage response in cancers. KDM4A is amplified or overexpressed in multiple cancer types, including breast, lung, and prostate.^19^ In prostate cancer, KDM4A overexpression is not sufficient to drive the formation of prostate adenocarcinoma in a genetically engineered mouse model (GEMM).^20^ However, KDM4A cooperates with the loss of tumor suppressor PTEN and overexpression of oncogenic ETV1 to enhance the development of primary prostate cancer.^20^ Although overexpression or shRNA KD of KDM4A affects prostate cancer cell proliferation,^20^ whether KDM4A plays a role in NEPC is unknown. Here, we identified KDM4A as a key regulator of NEPC progression and established alternative epigenetic pathways driving NEPC progression. We also showed that targeting KDM4A may be a promising therapeutic approach for treating NEPC.

## RESULTS

### KDM4A is overexpressed in neuroendocrine prostate cancer (NEPC)

Although EZH2, a KMT that regulates H3K27me3, has been implicated in NEPC progression, the role of KDMs, the erasers of histone lysine methylation, has not been examined in NEPC. To identify KDMs that play a role in NEPC, we analyzed a well-annotated RNA-seq dataset,^21^ which includes five pathologically defined subtypes based on a proposed classification approach:^22^ adenocarcinoma (n=34), large cell neuroendocrine carcinoma (NEPC) (n=7), mixed NEPC-adenocarcinoma (n=2), small cell prostate carcinoma (SCPC) (n=4), and mixed SCPC-adenocarcinoma (n=2). Although it has been reported that NEPC and SCPC share some genetic and molecular features (e.g., expression of NE markers),^23,24^ we observed that SCPC and adeno mixed with SCPC samples from this cohort have lower expression of NE markers such as CHGA and CHGB compared to NEPC and mixed NEPC-adenocarcinoma, even though the expression of these NE markers was higher than adeno-CRPC (**Suppl. Fig. 1A**). This observation is consistent with the existing heterogeneity in NEPC.^25,26^ Thus, we did not include the SCPC in the NEPC group in the DEG analysis. Due to the small sample size of the non-adenocarcinoma samples, we grouped these samples into adenocarcinoma, large cell neuroendocrine carcinoma (NEPC and mixed NEPC-adenocarcinoma), and SCPC (SCPC and mixed SCPC-adenocarcinoma) for differentially expressed gene analysis (DEGs). We compared the genome-wide gene expression of NEPC (NEPC and mixed NEPC-adenocarcinoma) to adenocarcinoma and SCPC (SCPC and mixed SCPC-adenocarcinoma) and identified 4669 and 1834 upregulated genes in NEPC compared to adenocarcinoma and SCPC, respectively (**Suppl. Table 1-2**). Also, we identified 1525 and 33 downregulated genes in NEPC compared to adenocarcinoma and SCPC, respectively (**Suppl. Table 1-2**). Among the histone lysine demethylases, KDM4A and KDM5D were uniquely upregulated in NEPC compared to adeno-CRPC and SCPC (**Figure 1A-B**). To determine whether KDM4A or KDM5D mRNA is also upregulated in mouse NEPC, we took advantage of a publicly available RNA-seq dataset from Pb-Cre+;**<u>P</u>**ten^f/f^;**<u>N</u>**MYC/NMYC;**<u>R</u>**b1^f/f^ NEPC model (referred to as **PNR** hereafter), we found that KDM4A, but not KDM5D, is upregulated in advanced tumors from older mice (16 weeks) compared to tumors from younger mice (12-13 weeks) (**Figure 1C**), suggesting KDM4A overexpression may play a role in NEPC progression. Also, KDM4A is overexpressed in the more aggressive NEPCs from PNR mice compared to tumors from Pb-Cre+;**<u>P</u>**ten^f/f^;**<u>N</u>**MYC/NMYC (**PN**) and Pb-Cre+;**<u>P</u>**ten^f/f^;NMYC/+;**<u>R</u>**b1^f/+^ (**PN^het^ R^het^**) mice with comparable age (16-17 weeks) (**Figure 1C**), which develop prostate adenocarcinoma. In addition, KDM4A, but not KDM5D (also named JARID1D),^27^ was shown to play an oncogenic role in prostate adenocarcinoma.^20^ Thus, we decided to focus on the functions of KDM4A in NEPC progression.

**Figure 1.**
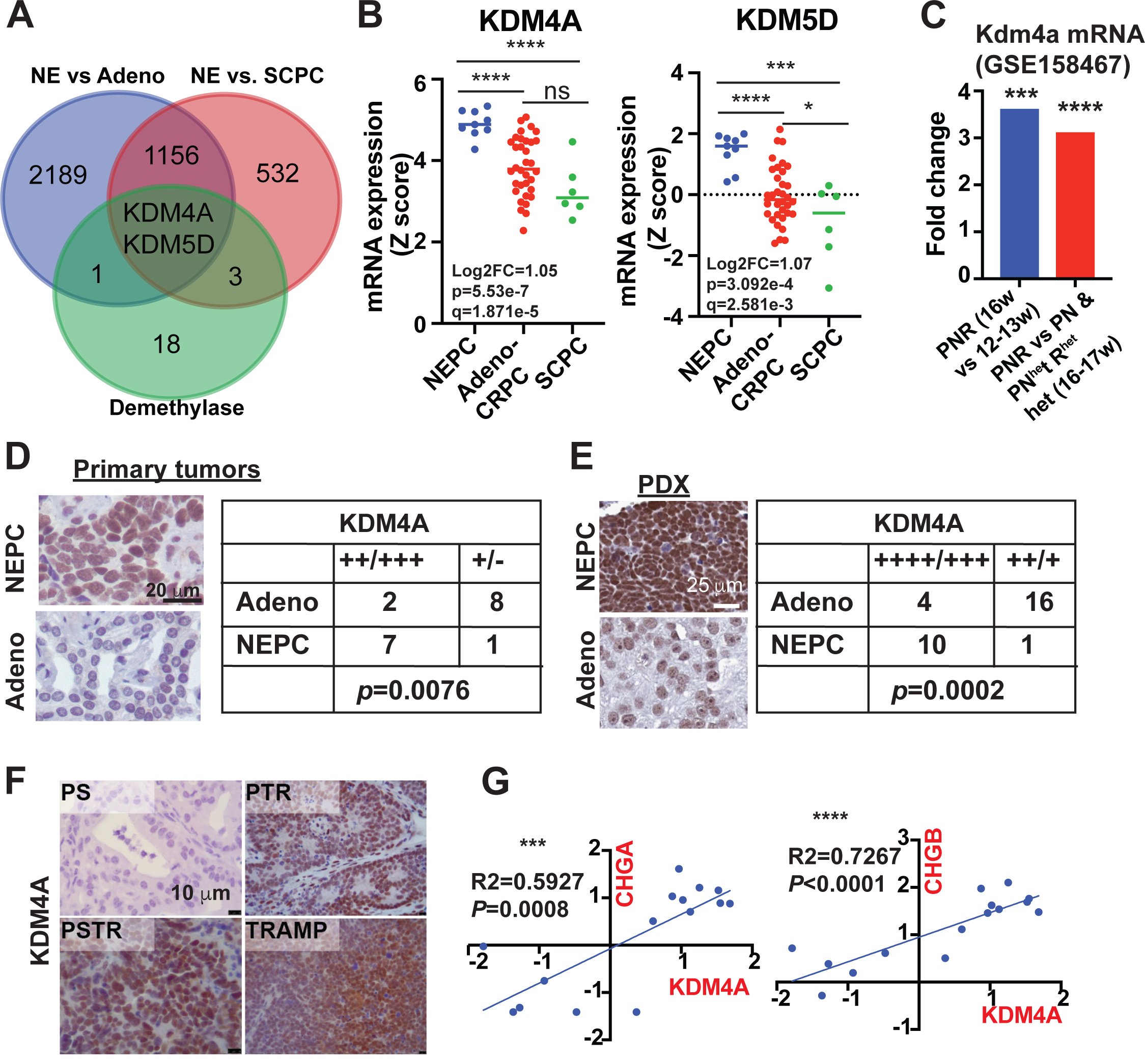
KDM4A is overexpressed in human and mouse NEPC. (**A**) Venn Diagram using DEGs (NEPC vs adeno & NEPC vs SCPC) showed that KDM4A and KDM5D are the two histone lysine demethylases overexpressing in NEPC using the Beltran et al. RNA-seq dataset. (**B**) The expression of *KDM4A* and *KDM5D* in NEPC compared to adeno-CRPC and SCPC using the Beltran et al. RNA-seq dataset. (**C**) *Kdm4a* mRNA is upregulated in prostate tumors from older PNR mice than tumors from young PNR mice, PN mice, and PN^het^R^het^ mice (PNR 16w vs 12-13w: FDR adj. P value=0.000479969; 16-17w PNR vs 16-17w PN/ PN^het^R^het^: FDR adj. P value=0.0000725). (**D**) KDM4A IHC in 8 cases of primary NEPC and 10 cases of primary adenocarcinoma-CRPC. Tumor samples were separated into KDM4A-high (+++/++) and KDM4A-low (-/+) groups and Fisher’s exact test was performed using Prism 5 software. (**E**) KDM4A IHC staining in prostate cancer PDXs. Tumor samples were separated into KDM4A-high (++++/+++) and KDM4A-low (++/+) groups and Fisher’s exact test was performed using Prism 5 software. (**F**) KDM4A IHC in primary tumors from PS, PTR, PSTR and TRAMP mice. (**G**) Correlation of KDM4A mRNA expression with NE markers *CHGA* and *CHGB* in the Beltran et al. RNA-seq dataset.

To confirm the findings from the human and mouse RNA-seq data, we performed IHC staining of KDM4A in human and mouse primary prostate adenocarcinoma and NEPC. The KDM4A IHC results were scored based on the staining intensity (−: negative; 1:+; 2: ++; 3: +++; 4: ++++). We found that KDM4A is overexpressed in human primary NEPC but not in primary prostate adenocarcinoma (**Figure 1D**) and the majority of NEPCs are moderately or strongly positive for KDM4A. We also examined KDM4A protein expression in a well-characterized prostate cancer patient-derived xenograft (PDX) tissue microarray (TMA).^28^ We found that KDM4A protein is overexpressed in NEPC PDXs compared to adenocarcinoma PDXs (**Figure 1E**) and the majority of NEPC PDXs are strongly positive for KDM4A. To determine the expression of KDM4A protein in murine NEPC, we examined its expression in three genetically engineered mouse models (GEMMs) that generate tumors with NEPC characteristics: TRAMP,^29^ prostate-specific conditional knockout (cKO) of *Pten*/*Trp53*/*Rb1* (PTR),^14^ and a recently generated prostate-specific cKO of *Pten*/*Smad4*/*Trp53*/*Rb1* (PSTR). We found that KDM4A protein is highly expressed in the primary NEPC tumors from all three NEPC GEMMs compared to two well-established prostate adenocarcinoma models, i.e., the prostate-specific cKO of *Pten*/*Smad4* (PS model) and cKO of *Pten* (**Figure 1F & Suppl. Figure 1B**), consistent with our findings in human prostate tumor tissues. Of note, the specificity of the KDM4A antibody was validated using *Kdm4a*-KO PSTR cells, a cell line derived from PSTR primary tumors (**Suppl. Figure 1C**).

We next determined whether KDM4A overexpression is associated with neuroendocrine markers. Using the Beltran et al. RNA-seq dataset,^21^ we observed a strong positive correlation between *KDM4A* mRNA and multiple neuroendocrine (NE) markers, including *CHGA*, *CHGB*, *ENO2,* and *DLL3* (**Figure 1G & Suppl. Figure 1D**). We also examined the expression of *KDM4A* in the Abida et al. RNA-seq dataset,^30^ which also contains a small subset of NEPC. We found that *KDM4A* mRNA is significantly higher in mCRPC tumor samples with NE features compared to those without NE features (**Suppl. Figure 1E**). Collectively, our data strongly suggest that KDM4A may play a role in NEPC progression.

### KDM4A KD or KO impairs the growth of NEPC *in vitro* and *in vivo*

To examine whether KDM4A plays a role in NEPC progression, we used shRNA, siRNA, and sgRNA to generate *KDM4A* knockdown (KD) and knockout (KO) cells. Mouse prostate cancer cell lines derived from PSTR and PTR primary tumors, which were confirmed to have upregulated expression of NE markers such as *Ncam1*, *Chga*, *Syp*, *Foxa2*, *Prox1*, and *Sox2* compared to Myc-CaP adenocarcinoma using qRT-PCR (**Suppl. Figure 2A-B**), were used in this study. We confirmed the efficient KD of *Kdm4a* using Western blot in PSTR cells (**Figure 2A**). Given the highly proliferative nature of NEPC cells, we first examined whether KDM4A is required for the proliferation of NEPC cell lines using *Kdm4a*-KD cells and control cells. We found that *Kdm4a* KD using shRNA or siRNA significantly reduced cell proliferation in PSTR cells as shown by foci-formation assay (**Figure 2B & Suppl. Figure 2C**). *Kdm4a* KD in PTR cells also reduced cell proliferation (**Figure 2C-D**). We also used the CRISPR/Cas9 system to generate *Kdm4a* KO and control PSTR cells. We confirmed the complete KO of *Kdm4a* using Western blot (**Figure 2E**). We found that *Kdm4a* KO in PSTR cells significantly reduced cell proliferation (**Figure 2F**). To determine the effect of KDM4A on human NEPC cells, we generated *KDM4A* KD and control human NEPC cells using lentiviral shRNA in 144-13 and LASCPC-1 models (**Figure 2G & Suppl. Figure 2D**). The human NEPC cell line 144-13 was derived from an NEPC PDX (MDA PCa 144-13)^31^ and LASCPC-01 was derived from Myr-AKT/N-MYC transformed PCa.^17^ We found that KD of *KDM4A* also significantly reduced 144-13 and LASCPC-1 cell proliferation (**Figure 2H & Suppl. Figure 2E**), consistent with our findings using mouse NEPC cell lines. Together, our data suggest that KDM4A plays a role in the growth of NEPC cells *in vitro*.

**Figure 2.**
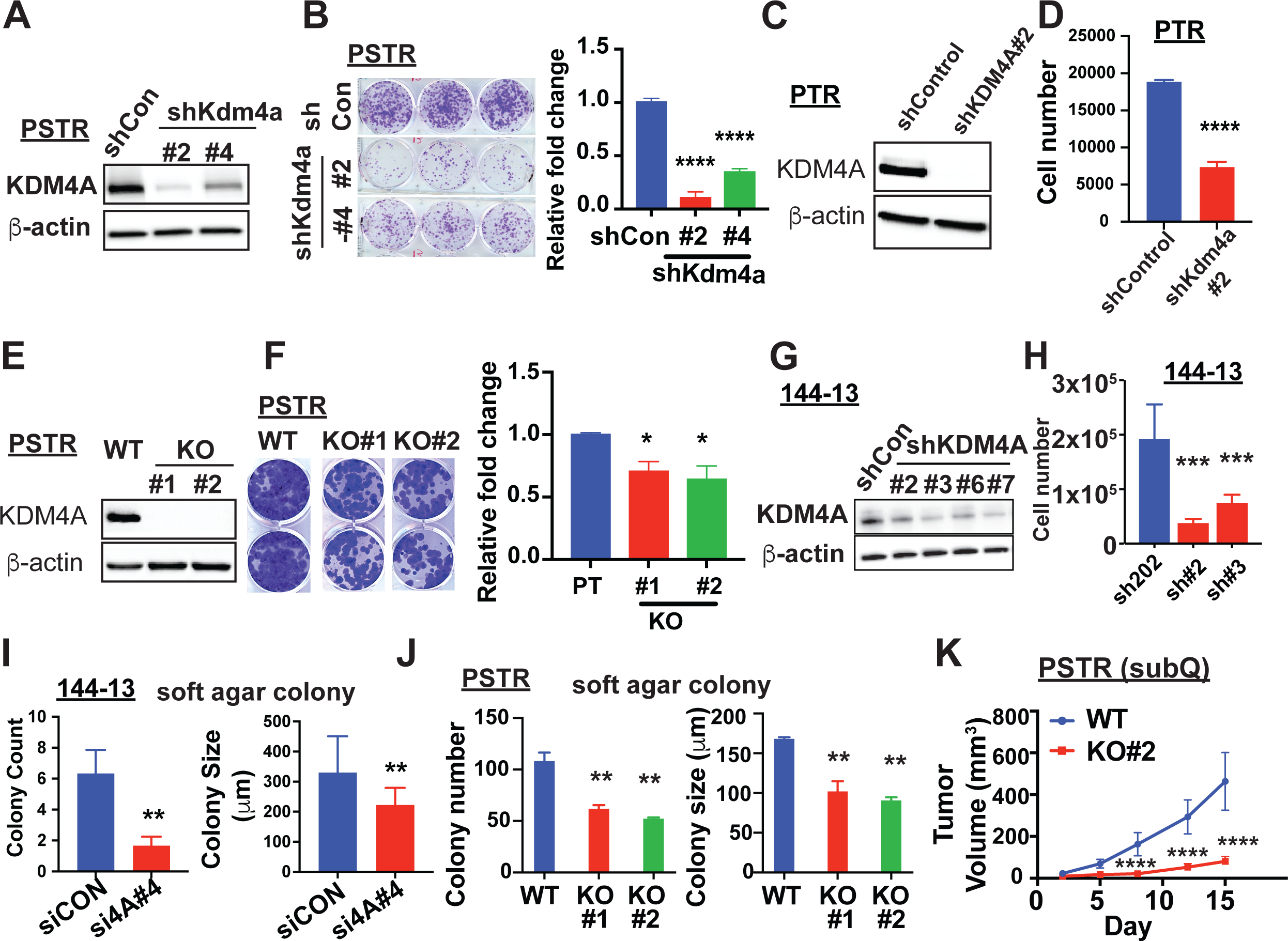
KDM4A knockdown (KD) or knockout (KO) suppresses NEPC growth *in vitro*. (**A**) Western blot was used to confirm the *Kdm4a* KD efficiency in PSTR cells infected with two independent *Kdm4a shRNA* or control shRNA (shCon). (**B**) Foci-forming assay was used to determine the effect of *Kdm4a* KD on the PSTR growth *in vitro* and the quantification of crystal violet staining intensity was performed using ImageJ. (**C**) Western blot was used to confirm the *Kdm4a* KD efficiency in PTR cells. (**D**) Direct cell counting was performed using Cellometer K2 to determine the effect of *Kdm4a* KD on PTR cell growth *in vitro*. (**E**) Western blot was used to confirm *Kdm4a* KO efficiency in PSTR cells. (**F**) Foci-forming assay was used to determine the effect of *Kdm4a* KO on the growth of PSTR cells and the quantification of crystal violet staining intensity was performed using ImageJ. (**G**) Western blot confirmed *KDM4A* KD efficiency in 144-13 cells. (**H**) Direct cell counting was performed using Cellometer K2 to determine the effect of *KDM4A* KD on the growth of 144-13 cells *in vitro.* (**I-J**) The effect of *KDM4A* KD in 144-13 cells (**I**) or *Kdm4a* KO in PSTR cells (**J**) on the anchorage-independent growth was determined by soft agar colony formation assay. (**K**) The subQ tumor volumes of *Kdm4a-*KO and *Kdm4a*-WT cells immune-deficient nude mice.

We also tested the impact of *KDM4A* KD and KO in NEPC cells on anchorage-independent growth, an assay that measured the tumorigenicity of cancer cells *in vitro*, using a soft agar colony formation assay. Because human NEPC 144-13 and LASCPC-01 infected with shKDM4A lentivirus generated above failed to expand, we switched to use small interference RNA (siRNA) to knock down *KDM4A* in human NEPC cells. We observed that *KDM4A* KD in 144-13 cells led to a reduction in colony numbers and sizes compared to control siRNA (**Figure 2I**). Similar results were observed in PSTR *Kdm4a-*KO and control cells (**Figure 2J**). To examine the effect of *Kdm4a* KO on NEPC cell growth *in vivo*, we subcutaneously implanted *Kdm4a-KO* PSTR cells and control cells in nude mice. We found that *Kdm4a* KO significantly reduced tumor growth *in vivo* (**Figure 2K**).

Collectively, our data suggest that KDM4A plays an important role in NEPC growth both *in vitro* and *in vivo*.

### KDM4A regulates MYC expression in NEPC cells

KDM4A is an epigenetic modifier, and its dysregulation may alter the expression of many genes. To identify genes and pathways that were regulated by KDM4A and may play a role in NEPC progression, we performed RNA-seq of *Kdm4a*-KD and control PTR cells. We identified 1270 downregulated genes and 2448 upregulated genes in *Kdm4a*-KD cells compared to control PTR cells (FDR≤0.05, fold change≥1.5) (**Suppl. Table 3**). Gene Set Enrichment Analysis (GSEA) showed that *Kdm4a* KD led to reduced activation of E2F signaling (**Suppl. Figure 3A**), which is consistent with previous findings that KDM4A interacts with E2F1 to regulate transcription.^32^ Also, mTORC1 signaling is suppressed in *Kdm4a*-KD PTR cells as shown by GSEA (**Suppl. Figure 3B**), which is consistent with a previous report that KDM4A is required for the activation of the HIF1α/DDIT4/mTOR1 axis.^33^ Interestingly, we identified MYC signatures (HALLMARK MYC TARGETS V1 and HALLMARK MYC TARGETS V2) as the top downregulated pathways in *Kdm4a* KD cells (**Figure 3A**). Multiple MYC target genes (e.g., *Ldha*, *Nme1*, and *Srsf2*, etc.) were significantly downregulated in *Kdm4a*-KD cells (**Figure 3B**). Interestingly, MYC has been implicated to play a functional role in NEPC, as overexpression of MYC, AKT1, BCL2, and RB1-KD/p53-KD transformed normal prostate epithelial cells into NEPC.^34^ We also found that MYC is the top hallmark pathway activated in both human and mouse NEPC as shown by GSEA analyses of publicly available RNA-seq data of adenocarcinoma and NEPC from patients^21^ and mice^14^ (**Figure 3C-D**). We performed qRT-PCR and Western blot analyses to examine the effect of *KDM4A* KD on MYC expression in PSTR, PTR, and 144-13 cell lines. We found that *KDM4A* KD reduced *MYC* mRNA and protein expression in all three cell lines (**Figure 3E-G & Suppl. Figure 3C**). Additionally, MYC expression was downregulated in QC6352-treated 144-13 NEPC cells compared to control cells (**Figure 3H)**. Collectively, our data show that KDM4A regulates MYC expression in NEPC, and targeting KDM4A can lead to reduced MYC expression.

**Figure 3.**
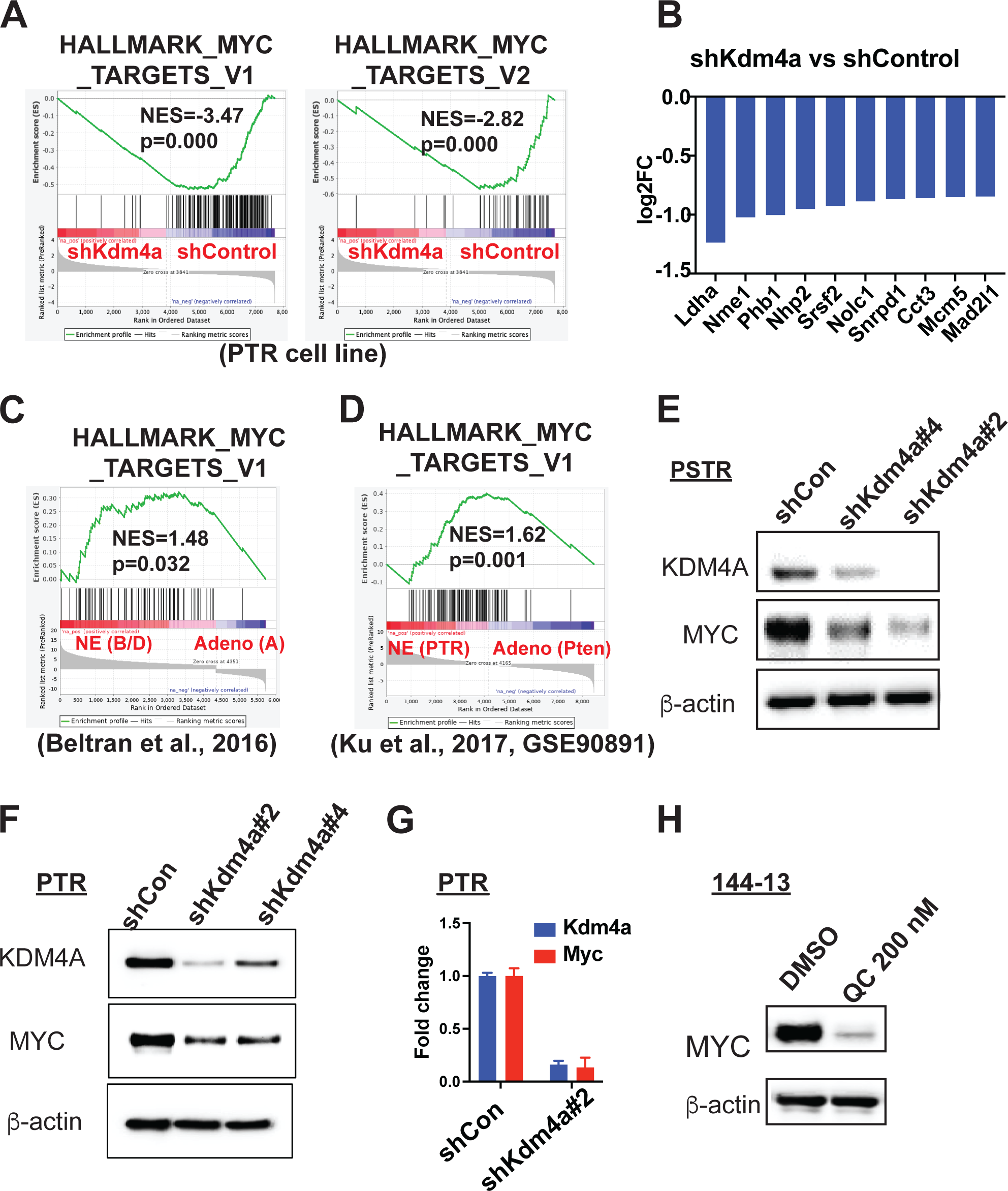
KDM4A regulates *MYC* transcription in NEPC. (**A**) GSEA analysis of RNA-seq on *Kdm4a*-KD and control PTR cells showed that MYC signaling is suppressed in *Kdm4a*-KD cells. (**B**) Fold changes in the expression of selective MYC-target genes comparing *Kdm4a*-KD cells to control cells. (**C-D**) GSEA analysis of Beltran et al. RNA-seq data (**C**) and Ku et al. RNA-seq data (**D**) showed that MYC signaling is activated in NEPC compared to prostate adenocarcinoma. (**E-F**) Western blot showed that KDM4A KD led to reduced MYC expression in PSTR (**E**) and PTR (**F**) cells. (**G**) qRT-PCR showed that *Kdm4a* KD led to reduced *Myc* mRNA expression in PTR cells. (**H**) KDM4 inhibitor QC6352 treatment led to reduced MYC expression as shown by Western blot.

### KDM4A directly regulates the transcription of MYC which is required for NEPC proliferation

Given the well-characterized function of KDM4A in epigenetic regulation, we aimed to determine whether it directly regulates the transcription of MYC. We performed ChIP-seq to identify KDM4A target genes. Of note, the ChIP-seq grade antibody from Schüle lab did not work on mouse cells, and several other KDM4A antibodies also failed to produce good quality ChIP-seq data (**data not shown**). Consequently, we performed KDM4A ChIP-seq analysis in human NEPC cells. We found that KDM4A preferentially binds to promoter/transcription starting site (TSS) regions of its target genes (**Figure 4A-B**). Importantly, we found that KDM4A binds to MYC locus in our ChIP-seq data (**Figure 4C**). Since KDM4A KD and inhibition lead to reduced MYC mRNA expression, our data suggest that MYC is a direct target gene of KDM4A and requires KDM4A to maintain its active transcription. To determine the role of MYC in NEPC, we first determined the effect of *MYC* KD or MYC inhibitor on NEPC cell proliferation in 144-13 cells. We found that *MYC* KD or MYC inhibitor MYCi975 indeed significantly reduced cell proliferation in 144-13 (**Figure 4D-E**). Collectively, our data suggest that KDM4A promotes NEPC progression in part through direct regulation of MYC transcription.

**Figure 4.**
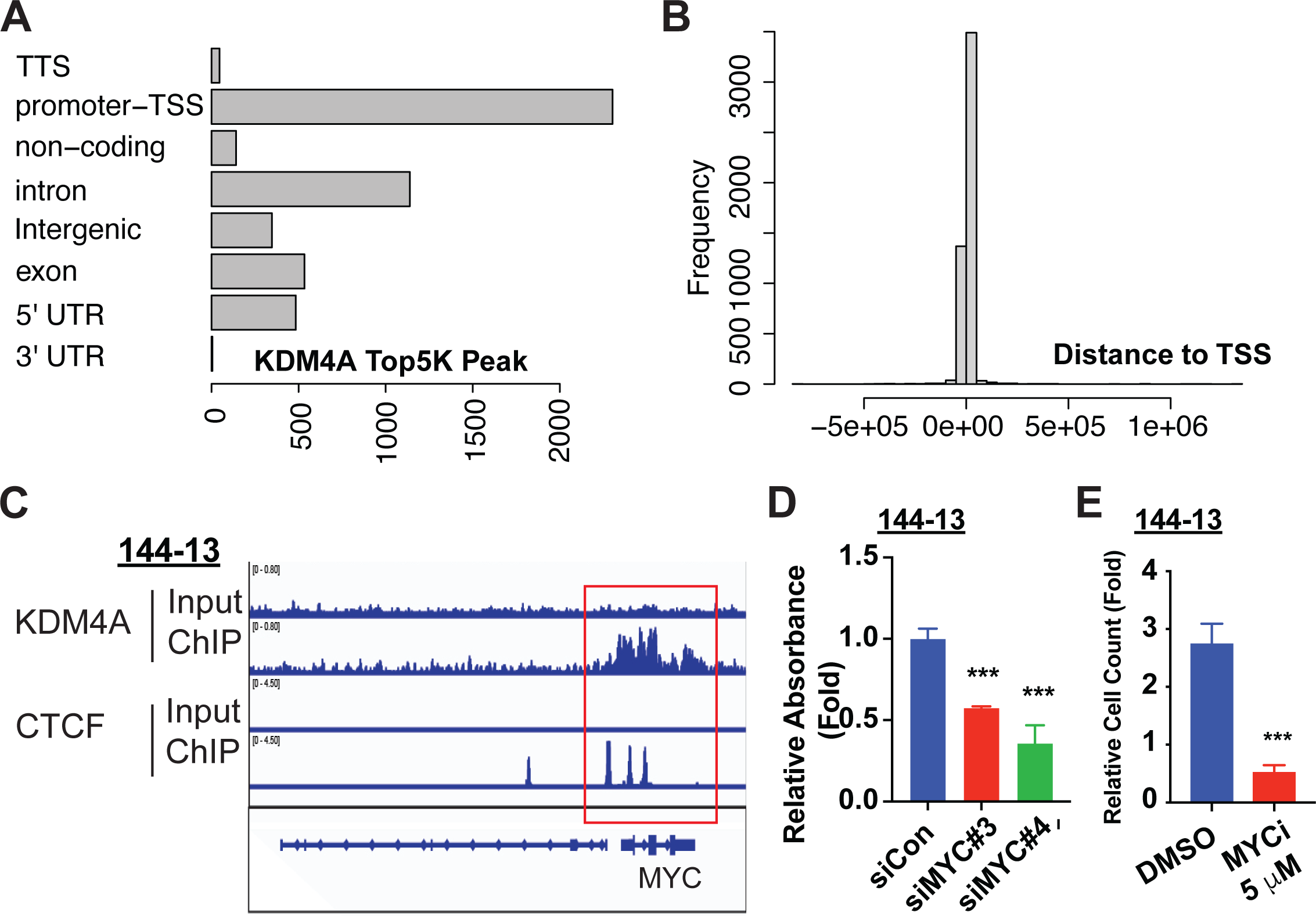
MYC is a direct transcriptional target gene of KDM4A. (**A**) Classification of the distribution of top 5000 KDM4A binding peaks (TSS: transcription starting sites) from KDM4A ChIP-seq. (**B**) The distances of the KDM4A binding peaks to the TSS. (**C**) KDM4A binding to MYC loci as shown by ChIP-seq. (**D**). *MYC* KD led to reduced cell proliferation in 144-13 cells. (**E**) MYC inhibitor MYCi975 led to reduced cell proliferation in 144-13 cells.

### KDM4 inhibitor suppresses cell proliferation in NEPC *in vitro*

Because our genetic data showed that KDM4A plays an important role in NEPC progression, we examined whether targeting KDM4A can be an effective therapeutic strategy for NEPC. Although there are no KDM4A-specific inhibitors, several pan-KDM4 inhibitors (KDM4i) [e.g., QC6352,^35,36^ NSC636819,^37^ NCGC00244536,^38^ IOX1,^39^ SD70^40^] have been shown to suppress the growth of cancer cells. For example, QC6352 was shown to suppress the growth of colorectal cancer (CRC) and triple-negative breast cancer (TNBC), and NCGC00244536 was shown to suppress the growth of prostate cancer cell lines PC3 and LNCaP. Among them, QC6352 is a highly selective inhibitor against the KDM4 family and rigorously characterized (e.g., cocrystal structure with KDM4A) with potent anti-tumor activities, favorable pharmacokinetics, and low toxicity in mice.^35,36^ We found that treatment of PSTR cells with QC6352 resulted in a significant increase in the global level of H3K9me3 and H3K36me3 expression, two of KDM4’s substrates **(Figure 5A**). We then examined the effect of QC6352 on the growth of PSTR cells *in vitro*, which are largely resistant to AR inhibitor enzalutamide (ENZ) (**Suppl. Figure 4A**). We found that QC6352 treatment led to a dramatic decrease in the proliferation of PSTR cells in a dose-dependent manner (**Figure 5B**). QC6352 treatment also dramatically suppressed the proliferation of PTR cells (**Figure 5C**) and 144-13 cells (**Figure 5D**). Furthermore, NCGC00244536, another potent inhibitor for KDM4 histone lysine demethylase,^38^ suppressed the proliferation of 144-13 and PSTR cells (**Figure 5D & Suppl. Figure 4B**). To determine whether KDM4A inhibition is responsible for the effect of QC6352 on cell proliferation, we compared the response of *Kdm4a*-WT and -KO PSTR cells to QC6352 treatment. We found that *Kdm4a* KO largely abrogated the response of PSTR cells to QC6352 (**Figure 5E**), supporting the notion that KDM4A is the critical target of QC6352 in NEPC cells.

**Figure 5.**
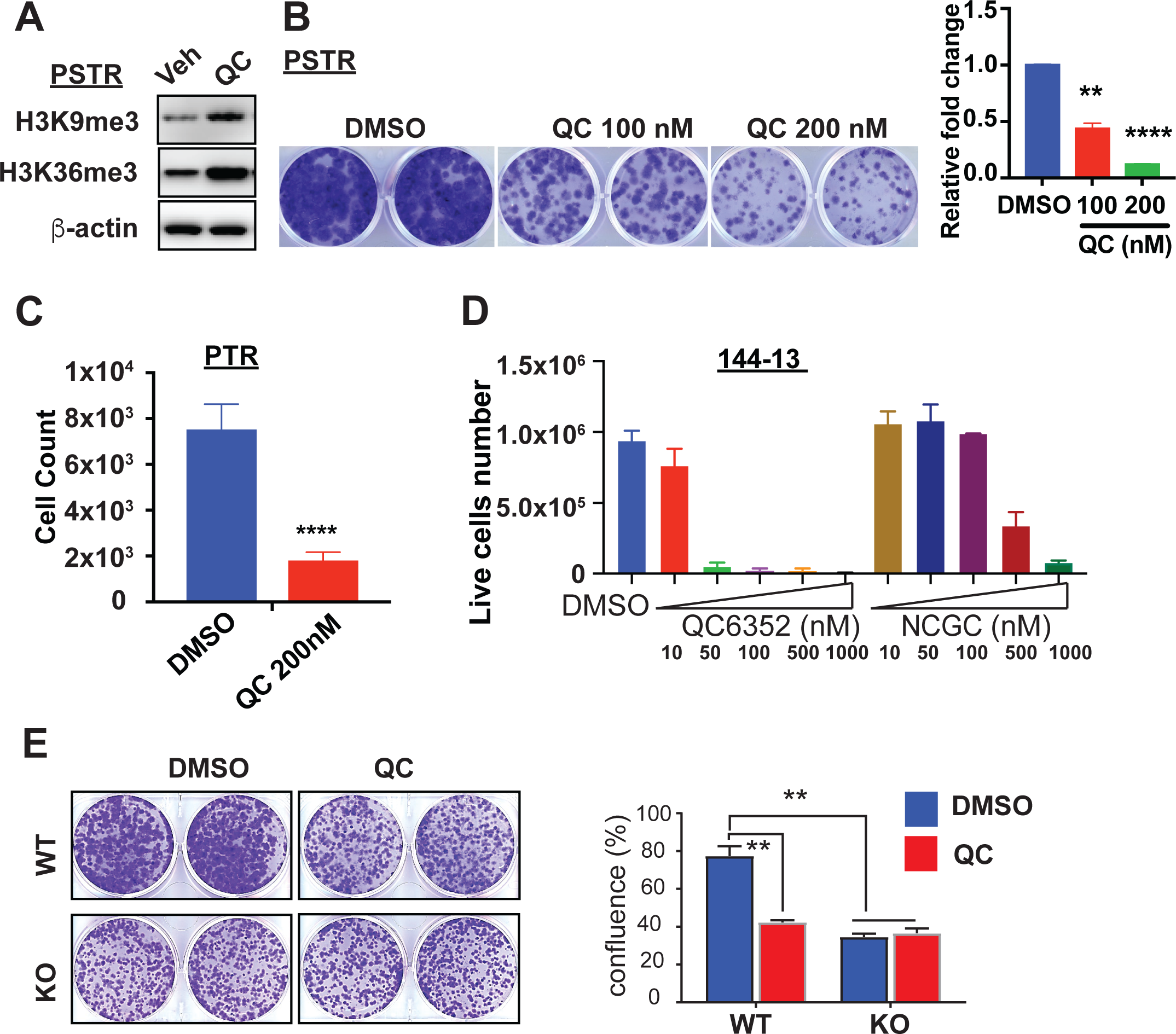
KDM4 inhibitors suppress NEPC cell growth *in vitro*. (**A**) KDM4 inhibitor QC6352 treatment led to increased total H3K9me3 and H3K36me3 in PSTR cells. (**B**) QC6352 suppressed PSTR growth *in vitro* as shown by foci-forming assay. (**C**) QC6352 suppressed the growth of PTR cells. (**D**) QC6352 and NCGC00244536 suppressed the growth of 144-13 cells *in vitro*. (**E**) *Kdm4a*-KO blunts the effect of QC6352 treatment on cell growth in *vitro*.

### KDM4 inhibitor suppresses NEPC progression *in vivo*

To determine whether QC6352 could suppress NEPC growth *in vivo*, we treated immune-deficient mice bearing subcutaneous PSTR tumors and 144-13 tumors with QC6352. We found that QC6352 treatment dramatically suppressed the growth of PSTR cells (**Figure 6A-B**) and 144-13 cells (**Figure 6C-D**), as shown by reduced tumor sizes and tumor weights. To ensure QC6352 has similar effects on tumor progression in immune-competent hosts, we treated PSTR GEMM with QC6352. Similarly, QC6352 treatment in the PSTR GEMM model led to a significant reduction in tumor burden as shown by tumor weights (**Figure 6E).** We then examined the effects of QC6352 treatment on tumor cell proliferation. We found that QC6352 treatment led to a significantly reduced proliferation in PSTR GEMM as shown by Ki67 IHC staining (**Figure 6F**). We did not observe noticeable toxicity as shown by the measurement of peripheral blood count and the measurement of weight of major organs **(Suppl. Figure 5A-B)**, consistent with the findings in the breast cancer models.^35,36^ Together, our data suggest that KDM4 inhibitor QC6352 can significantly delay NEPC progression *in vivo*.

**Figure 6.**
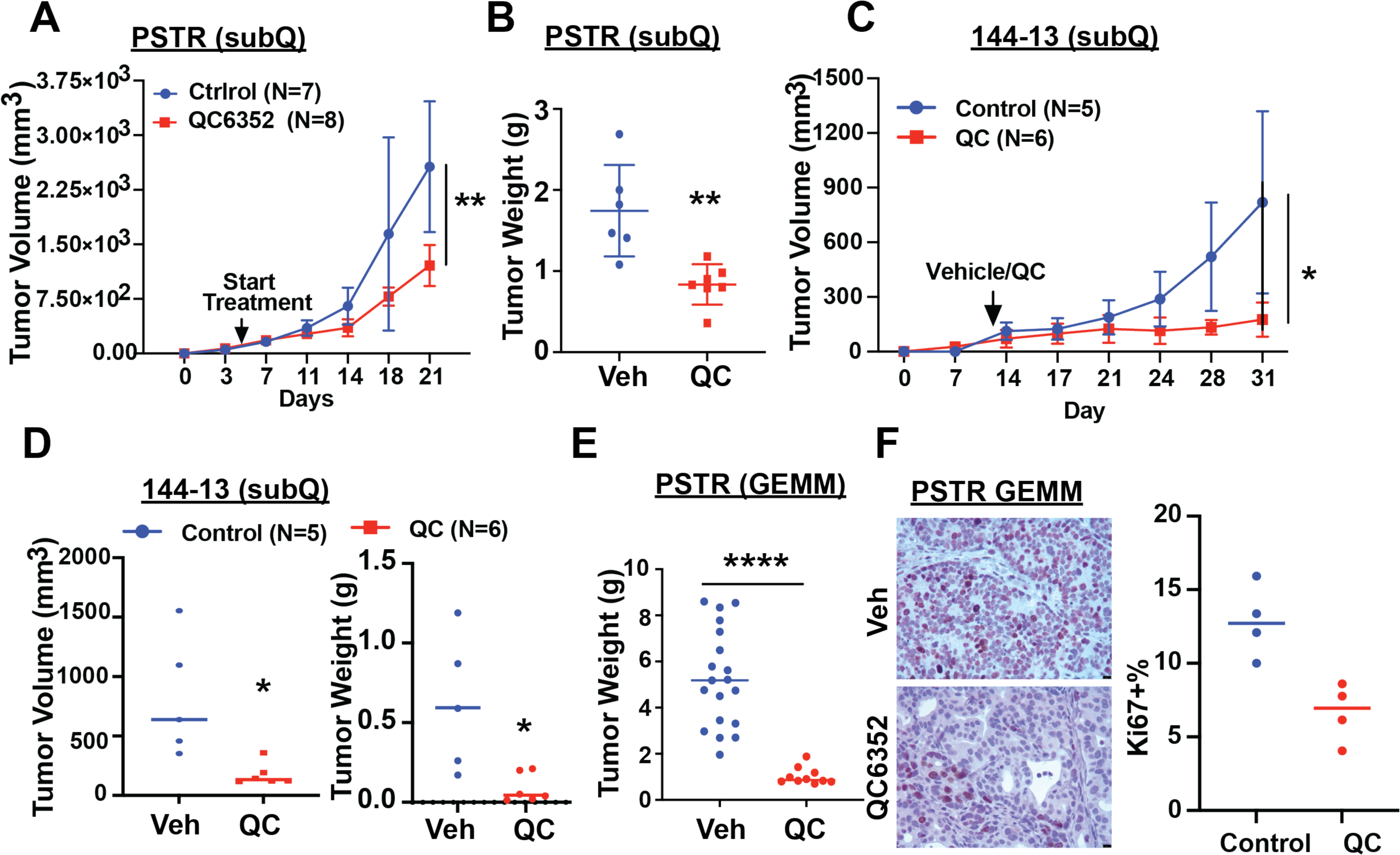
KDM4 inhibitor suppresses NEPC cell growth *in vivo*. (**A-B**) Weekly measurement of tumor volumes (**A**) and endpoint tumor weights (**B**) of subQ implanted PSTR cells treated with QC6352 (50 mg/kg daily) or vehicle control. (**C-D**) Weekly measurement of tumor volumes (**C**) and endpoint tumor volumes and tumor weights (**D**) of subQ implanted 144-13 cells treated with QC6352 (50 mg/kg daily) or vehicle control. (**E**) Endpoint tumor weights of primary tumors from PSTR GEMM treated with QC6352 (50 mg/kg daily) or vehicle control. (**F**) Ki67 staining of QC6352-treated and vehicle-treated primary tumors from PSTR mice.

## DISCUSSIONS

The incidence of NEPC increases due to the widespread use of AR pathway inhibitors (ARPIs) in the treatment of non-metastatic CRPC and hormone-sensitive metastatic tumors. We showed that KDM4A is highly upregulated in NEPC and is functionally involved in the tumor progression of NEPC. We also demonstrated that targeting KDM4A, by genetic deletion, RNA interference, or small-molecule inhibitors, suppresses NEPC tumor growth. Together, our studies revealed a novel role of KDM4A in mediating NEPC progression and provide a therapeutic target for the treatment of NEPC.

Our finding that KDM4A regulates MYC expression in NEPC provides one of the downstream genes regulated by KDM4A. KDM4A is an epigenetic modifier, and its dysregulation may alter the expression of many genes. We identified MYC as a KDM4A-target gene through RNA-seq analysis of *KDM4A* KD and control PTR cells and ChIP-seq of KDM4A in 144-13 human NEPC cells, and thus provided a novel mechanism for how KDM4A exerts its effect on NEPC progression. MYC has previously been shown to be a key player in prostate cancer progression, and overexpression of MYC in the prostate was sufficient to induce prostate cancer and promote tumor progression in mouse models.^41-43^ MYC was also shown to play an important role in NEPC.^34^ However, how MYC is upregulated in NEPC was not clear. We found that MYC signaling is activated in human and mouse NEPC and MYC is required for NEPC proliferation *in vitro*. Our study reveals KDM4A as an upstream regulator of MYC activation and provides a novel insight into how an epigenetic modifier regulates oncogene expression. This novel finding also led us to examine whether such regulation occurs in other cancer types. We analyzed a publicly available RNA-seq dataset^44^ and found that *Myc* mRNA was downregulated in mouse squamous cell carcinoma (SCC) cell lines upon *Kdm4a* KO compared to control cells (**Suppl. Figure 6A**). Also, by analyzing a publicly available RNA-seq dataset,^36^ we found that QC6352 treatment led to reduced expression of *MYC* mRNA in triple-negative breast cancer (TNBC) (**Suppl. Figure 6A**). Furthermore, we also examined a publicly available ChIP-seq dataset (KDM4A) in TNBC^36^ and found that KDM4A binds to MYC promoter/gene body in TNBC (**Suppl. Figure 6B**). These findings suggest that KDM4A may also upregulate MYC expression in other cancer types. Inhibitors that directly target MYC (e.g., MYCi975^45^) are available, but they are still in preclinical studies.^46^ Inhibition of KDM4A may be one novel therapy strategy for targeting tumors overexpressing both KDM4A and MYC. A clinical trial using TACH101 (also named QC8222), a more potent second-generation pan-KDM4 inhibitor similar to QC6352, is ongoing.^47^ Our study provides one mechanism for targeting KDM4 for cancer treatment.

While we showed that the decrease in the expression of MYC plays a role in suppressing NEPC proliferation in *KDM4A* KD or KO cells, other KDM4A-regulated pathways (e.g., E2F, mTORC1) (**Suppl. Figure 3A-B)** may also contribute to the phenotypes observed in *KDM4A-KD/KO* cells. For example, the activation of E2F signaling, which plays a critical role in cancer progression, was frequently observed in NEPC due to RB1-deficiency.^48^ Also, mTORC1 signaling, another crucial regulator of tumor initiation, progression and therapy responses, has been implicated in the regulation of NEPC pathogenesis.^49^ Thus, MYC is one of the mediators of KDM4A’s functions in NEPC.

Although we showed that the effects of KDM4 inhibitor QC6352 on NEPC cells are in large part due to the inhibition of KDM4A, QC6352 can inhibit other KDM4 family members besides KDM4A^35,36^. KDM4A shares structural similarities with KDM4B and KDM4C. Indeed, we found that KDM4 inhibitor QC6352 induces more pronounced apoptosis in PTR, PSTR, and 144-13 cells compared to *Kdm4a* KD. Treatment of PTR, PSTR, and 144-13 cells with QC6352 led to increased levels of cleaved caspase 3 and cleaved PARP1 by Western blot analysis, and green fluorescent signal from IncuCyte Caspase-3/7 Green Apoptosis Assay Reagent (**Suppl. Figure 6C-F**). In contrast, *Kdm4a* KD led to a minor increase in apoptosis in *PSTR cells* by Western blot for cleaved caspase 3 and cleaved PARP1 (**Suppl. Figure 6G**). Although apoptosis signature in *Kdm4a*-KD PTR cells was increased compared to control cells in GSEA analysis (**Suppl. Figure 6H**), it does not rank as the top pathway, suggesting that KDM4A also regulates cell apoptosis but to a minor extent. KDM4B and KDM4D have been shown to play a role in DNA damage response.^50,51^ Thus, QC6352 may also have an effect on DNA damage response. It will be interesting to analyze whether these pathways are affected in specimens from QC6352 clinical trial.

Besides playing a role in the progression of NEPC, KDM4A may also play a role in the development of CRPC. Shin et al.^52^ showed that KDM4A interacts with AR and activates AR’s transcriptional activities. Also, Xue et al.^53^ showed that p300-mediated acetylation of KDM4A recruits the BET family member BRD4 to stabilize KDM4A and promote its recruitment to AR targets. Furthermore, Zhang et al^54^ showed that KDM4A-AS1, a long non-coding RNA that is located near the 3’UTR of KDM4A, represses AR/AR-Vs degradation and enhances resistance to enzalutamide treatment. Since natural antisense long non-coding RNAs (lncRNAs), which are transcribed from the opposite strand of either protein-coding or non-coding genes, play a regulatory role in modulating their own sense gene expression,^55-57^ it will be interesting to determine whether KDM4A-AS1 lncRNA modulate the expression of KDM4A.

In summary, our study established KDM4A as an important epigenetic regulator that drives NEPC progression and can serve as an effective therapeutic target.

## MATERIALS and METHODS

We obtained human prostate tissues through an MD Anderson Cancer Center Institutional Review Board (IRB)-approved protocol. All animal experiments were performed under an Institutional Animal Care and Use Committee (IACUC)-approved protocol (00001713-RN01). The datasets generated during the current study are available from the corresponding author upon reasonable request. The data generated in this study have been deposited into the publicly available Gene Expression Omnibus (GEO) (GSE227717 and GSE227688). Details of model systems (mouse models, cell lines), reagents (antibodies, chemicals, siRNA, shRNA, sgRNA), experimental procedures, immunohistochemistry, cell culture, transfection, qRT-PCR, cell-viability assays, Western blotting, IP, *in vivo* experiments, microarray, RNA-seq, ChIP-seq, pathway-enrichment analysis, and statistical analyses are included in **SI Materials and Methods**.

## Acknowledgments

G.W. is supported by funding from MDACC (Moon Shot, IRG, PCRP, TDD drug development award), UT STARs Award (The University of Texas System Board of Regents), NIH [R00 CA194289 (GW), P50 CA140388 (C. Logothetis, S.-H. Lin)], and DoD [DOD-PCRP-Idea W81XWH-21-1-0522 (GW). S.-H. Lin is supported by R01CA174798 (S.-H. Lin)], Cancer Prevention Research Institute of Texas grants RP230247 & RP190252 (S.-H. Lin). M.G.L. is supported by NIH [R01 CA207109; R01 CA207098; R01 CA262324 (M.G.L). This study is supported by NIH P30CA016672 for the use of the Research Animal Support Facility, Flow Cytometry and Cellular Imaging Core Facility, and Functional Genomics Core at MD Anderson Cancer Center. We thank the support from the Cancer Prevention and Research Institute of Texas (CPRIT RP180734) for the RNA-seq service provided at the Cancer Genomics Center, The University of Texas Health Science Center at Houston. We thank the support from Epigenomics Profiling Core, Center for Cancer Epigenetics and the Department of Epigenetics and Molecular Carcinogenesis for the ChIP-seq services. We acknowledge the Movember GAP1 PDX Collaboration with thanks for use of the PDX TMA.

