## Supplementary material for "KDM4A promotes the progression of neuroendocrine prostate cancer": Suppl. information

### SI MATERIALS and METHODS

**Mouse strains.** *Pb-Cre4; Pten*$^{loxP/loxP}$ (*Pten*$^{pc-/-}$), *Pb-Cre4; Pten*$^{loxP/loxP}$*;Smad4*$^{loxP/loxP}$ (aka *Pten*$^{pc-/-}$ *Smad4*$^{pc-/-}$, PS), and *Pb-Cre4;Pten*$^{loxP/loxP}$*;Trp53*$^{loxP/loxP}$ (aka *Pten*$^{pc-/-}$ *Trp53*$^{pc-/-}$, PT) models were developed previously[1]. *Rb1*$^{loxP/loxP}$ strain (FVB;129-Rb1tm2Brn/Nci) was obtained from NCI Mouse Repository. *Pten*$^{pc-/-}$*Smad4*$^{pc-/-}$ and *Pten*$^{pc-/-}$*Trp53*$^{pc-/-}$ mice were crossed with *Rb1*$^{loxP/loxP}$ mice to generate *Pb-Cre4;Pten*$^{loxP/loxP}$*;Smad4*$^{loxP/loxP}$*;Trp53*$^{loxP/loxP}$*;Rb1*$^{loxP/loxP}$ (*Pten*$^{pc-/-}$*Smad4*$^{pc-/-}$*Trp53*$^{pc-/-}$ *Rb1*$^{pc-/-}$, PSTR) and *Pb-Cre4;Pten*$^{loxP/loxP}$*;Trp53*$^{loxP/loxP}$*;Rb1*$^{loxP/loxP}$ mice (*Pten*$^{pc-/-}$*Trp53*$^{pc-/-}$*Rb1*$^{pc-/-}$, PTR). The genotyping of the Pb-Cre4 transgene and all the conditional alleles were performed using conventional PCR as described previously.[1-4] The copy number of TRAMP transgene will be determined by quantitative PCR[5] and only mice that are homozygous for the transgene were used in this study. Mice were maintained in pathogen-free conditions at M.D. Anderson Cancer Center. All manipulations were approved under MD Anderson Cancer Center (MDACC) Institutional Animal Care and Use Committee (IACUC) under protocol number 00001713-RN01.

**Human prostate tumor tissues and patient-derived xenografts (PDX).** FFPE Primary castration-resistant prostate tumor (CRPC) tissues, including prostate adenocarcinoma and NEPC, were obtained from Dr. Patricia Troncoso and Dr. Miao Zhang (MD Anderson Tissue Bank). FFPE and fresh tumor tissues from PDXs were obtained from Dr. Nora Navone (GU PDX core). The Movember PDX TMA has been described previously.[6]

**Cell lines and cell cultures**. PTR and PSTR cell lines were isolated from *Pten*$^{pc-/-}$*Trp53*$^{pc-/}$ *Rb1*$^{pc-/-}$ and isolated from *Pten*$^{pc-/-}$ *Smad4*$^{c-/-}$*Trp53*$^{pc-/-}$ *Rb1*$^{pc-/-}$ mice. All cell lines tested for mycoplasma were negative within 6 months of performing the experiments. Cell line authentication was not performed. 293T, LASCPC-01, and MYC-CaP cells were obtained from ATCC. 144-13 was generated from PDX MDA PCa 144-13.[7] 293FT cells were obtained from Thermo Fisher Scientific Inc. 144-13 and LASCPC-01 cells were cultured in HITES medium supplemented with 5% fetal bovine serum using ATCC-formulated RPMI-1640 Medium (Catalog No.30-2001) with the following components to the base medium: 0.005 mg/ml Insulin, 0.01 mg/ml Transferrin, 30 nM Sodium selenite (final conc.), 10 nM Hydrocortisone (final conc.), 10 nM beta-estradiol (final conc.), extra 2mM L-glutamine (final conc. of 4 mM).

**Small interfering RNAs (siRNAs)/shRNA knockdown, CRISPR/Cas9 genome editing, and ORF overexpression**. siRNAs were ordered from Dharmacon Inc (human KDM4A: GUAUGAUCUUCCAGACUUA; GUGCGGAGUCUACCAAUUU; mouse Kdm4a: GAACAUCCUACGACGAUAU; GUUCGUGAGUUCCGCAAGA; human MYC: AACGUUAGCUUCACCAACA; AACGUUAGCUUCACCAACA; mouse Myc: GGACACACAACGUCUUYGGA; UCGAAACUCUGGUGCAUAA). Lentiviral shRNA plasmids were obtained from Sigma Aldrich (Human KDM4A: shKDM4A#1 TRCN0000013493, shKDM4A#2 TRCN0000013494, shKDM4A#3 TRCN0000013495; Mouse Kdm4a: shKdm4a#2 TRCN0000103526). Mouse shKdm4a#4 plasmid was obtained from VectorBuilder of which sequence of siKdm4a#4 (GUUCGUGAGUUCCGCAAGA) was cloned into the lentiviral vector (VB210619-1017gjm). Lentiviruses were packaged in 293FT cells using second-generation packaging vectors, psPAX2 (Addgene plasmid 12260) and pMD2.G (Addgene plasmid 12259). Synthetic guided RNA (sgRNA) was ordered from Synthego Inc. Recombinant cas9 was obtained from Thermo Fisher Scientific Inc. sgRNAs and recombinant cas9 were transfected into cells according to the protocol from the manufacturer. Single clones were selected, and *Kdm4a* KO clones were confirmed by western blot analysis. MYC_pLX307 (Plasmid #98363, Addgene)) was used to overexpress MYC.

**Chemicals and inhibitors.** QC6352 (HY-104048), NCGC00247743 (HY-112308), Enzalutamide (HY-70002), and MYCi975 (HY-129601) were ordered from MedChemExpress Inc. Incucyte Caspase 3/7 Dye for Apoptosis (Cat#4440) was ordered from Sartorius.

**Western Blot Analysis**. Cells were lysed on ice using RIPA buffer (Boston BioProducts) supplemented with Protease and Phosphatase Inhibitor Cocktail (Thermo Fisher Scientific). Proteins with loading buffer (20 μg) were subjected to SDS-PAGE and transferred onto a nitrocellulose membrane. Membranes were blocked in 2% non-fat dry milk for an hour before being incubated with primary antibodies prepared in Tris-buffered saline (TBS) containing 0.1% Tween 20 (TBST) consisting of 0.5% BSA overnight at 4°C. Next, the membrane was washed 3 times with TBS containing 0.1% Tween 20 (TBST) and was incubated with HRP- conjugated secondary antibody for 2 h at room temperature. After washing 3 times with TBST, the membrane was exposed to Clarity Western ECL Substrate (Bio-Rad) according to the protocol and imaged with Azure Biosystems c600. The following antibodies were used in this study: KDM4A (ab191433, Abcam), beta-Actin (MA5-15739, Invitrogen), cleaved-PARP (Mouse specific #9544; human specific #9546, Cell Signaling Technology) cleaved-caspase 3 (#9662, Cell Signaling Technology), cMyc (ab32072, Abcam), c-Anti-Rabbit IgG HRP- linked antibody (#7074, Cell Signaling Technology), Anti-Mouse IgG HRP-linked antibody (#7076, Cell Signaling Technology).

**Immunohistochemistry analyses.** Tissues were fixed in 10% formalin overnight and embedded in paraffin. Immunohistochemical (IHC) analysis was performed as described earlier (Aguirre et al., 2003). Primary antibodies used for immunohistochemistry are as follows: KDM4A (ab191433, Abcam), KDM4A (3393S, CST), Ki-67 (GTX16667, GeneTex), Synaptophysin (M7315, Dako), AR (ab133273, Abcam), Chromogranin A (20086, ImmunoStar). Secondary antibodies used are as follows: anti-rabbit (RMR622L, Biocare), and anti-mouse (8125S, CST). The immunohistochemistry signals were developed with DAB Quanto (TA-125-QHDX, Epredia). The staining intensity of KDM4A was scored as negative (-), weakly positive (+), moderately positive (++), and strongly positive (+++ or ++++).

**Cell proliferation, foci-formation, soft-agar colony forming, and apoptosis assays.** For cell proliferation assay, $5x10^2$-$2x10^3$ cells per well were seeded in 96-well plates with or without corresponding treatment. After 5 days, absorbance was measured using Cell Counting Kit-8 (B34034, Bimake) according to the manufacturer's manual. For foci-formation assay, $1\times10^3$ cells were seeded in 6-well plates and cultured for 5 to 7 days before they were fixed and stained with crystal violet as described.[8] For soft-agar colony forming assay, $2.5x10^3$ cells were seeded in 6-well. Cells were cultured for 3 weeks. Colony sizes and numbers were measured as described.[9] Apoptosis was detected by Western blot analysis of cleaved caspase 3 and cleaved PARP or by Cytation 5 using Caspase3/7 dye for apoptosis (Cat#4440, Sartorius).

**Xenograft Studies.** PSTR ($1x10^5$ cells/100ul per mouse) and 144-13 ($2x10^6$/100ul per mouse) cells were prepared in 1:1 PBS and Matrigel (BD Biosciences) and injected into flanks of 6-week-old nude mice (n = 5-10 per group). Mice were treated with 50 mg/kg QC6352 (once a day) when tumors reached $100mm^3$. Tumor growth was monitored by measuring tumor sizes twice a week. Tumor Volume was calculated according to the formula: Volume = length × $width^2$/2. All xenograft experiments were approved by the MD Anderson IACUC under protocol number #00001713-RN01.

**Quantitative RT-PCR and RNA-sequencing**. Cell mRNA was isolated by Direct-zol RNA MiniPrep (50ug) (#11-331, ZymoResearch). iScript cDNA Synthesis Kit (Bio-Rad) was used to generate cDNA for quantitative PCR (qPCR) analysis using PerfeCTa reg SYBR reg Green SuperMix Reaction Mixes (QuantaBio). The primers used for qPCR analysis were designed using Primer3 and ordered from Sigma Aldrich. The primers used were listed in Suppl. Table 4. RNA-seq of PTR cells with shRNA control and shKdm4a#2 and shKdm4a#4 was performed on total RNA (3 replicates) using NEBNext® Ultra II Directional RNA Library Prep Kit for Illumina NextSeq at the Cancer Genomics Center, The University of Texas Health Science Center at Houston. The sequences were aligned to mouse genome mm10. The raw data will be submitted to the NCBI Gene Expression Omnibus (GEO) with the accession number GSE227717. Processed human RNA-seq datasets from previous studies (Grasso et al.,[10] Abida et al.,[11] and Beltran et al.[12]) were downloaded from cBioportal.[13,14] Publicly available RNA-seq datasets (GSE158467,[15] GSE90891,[16] GSE95293,[17] and GSE137953[18]) were downloaded from NCBI GEO. DeSeq2 was used to identify differentially expressed genes (DEGs). Gene Set Enrichment Analysis (GSEA) was performed as described.[2]

**Chromatin-immunoprecipitation (ChIP)-sequencing.** Chromatin immunoprecipitation (ChIP) assays were performed at the MD Anderson Cancer Center, Epigenomics Profiling Core. 144-13 cells were crosslinked with formaldehyde and subjected to chromatin immunoprecipitation with antibodies specific KDM4A (#5766, lot 021110, Schuele Laboratory). ChIPs were performed as described previously[8,19,20] with modifications. ChIP DNA and corresponding input libraries were prepared using the NEBNext Ultra II DNA library prep kit (New England Biolabs) and were sequenced using Illumina HiSeq 2500 instrument. Reads were aligned to a reference genome Hg19 as described.[21] The raw data has been deposited to the NCBI Gene Expression Omnibus (GEO) with the accession number GSE227688. The publicly available ChIP-seq dataset GSE95190[17] was downloaded from NCBI GEO.

**Statistical analysis.** All the experiments were replicated at least twice in the laboratory except for microarray and ChIP. Data are presented as mean ± SD unless indicated otherwise. Fisher's exact test was used to analyze 2X2 contingency table on the IHC staining of KDM4A in human primary prostate tumors and PDXs. Student's t-test assuming two-tailed distributions was used to calculate statistical significance between two groups and one-way ANOVA Turkey's multiple comparisons test was used for data with three or more groups. **** $P < 0.0001$, *** $P < 0.001$; ** $P < 0.005$; * $P < 0.05$.

**Data Availability Statement**. The datasets generated during the current study are available from the corresponding author upon reasonable request. The data generated in this study have been deposited into the publicly available Gene Expression Omnibus (GEO) (GSE227717 and GSE227688)

**Author Contributions**. G.W. contributed to the study's conception and design of this study. G.W., C.M., M.Z., X.L., F.W., A.G.H., X.S., D.L., J.S., J.P., M.Z., P.T., and J.Z. performed the experiments and acquired, analyzed, and interpreted the data (e.g., statistical analysis, biostatistics, computational analysis). P.S., N.N., E.M., E.K., B.A.F., and R.S. provided key reagents. A.K.J., M.G.L, P.C., C.J.L., A.A., provided scientific inputs for the development of the project. C.M., M.Z., S.H.L and G.W. contributed to the writing and editing the manuscript.

**Competing Interests statement**. No potential conflicts of interest were disclosed by the other authors.

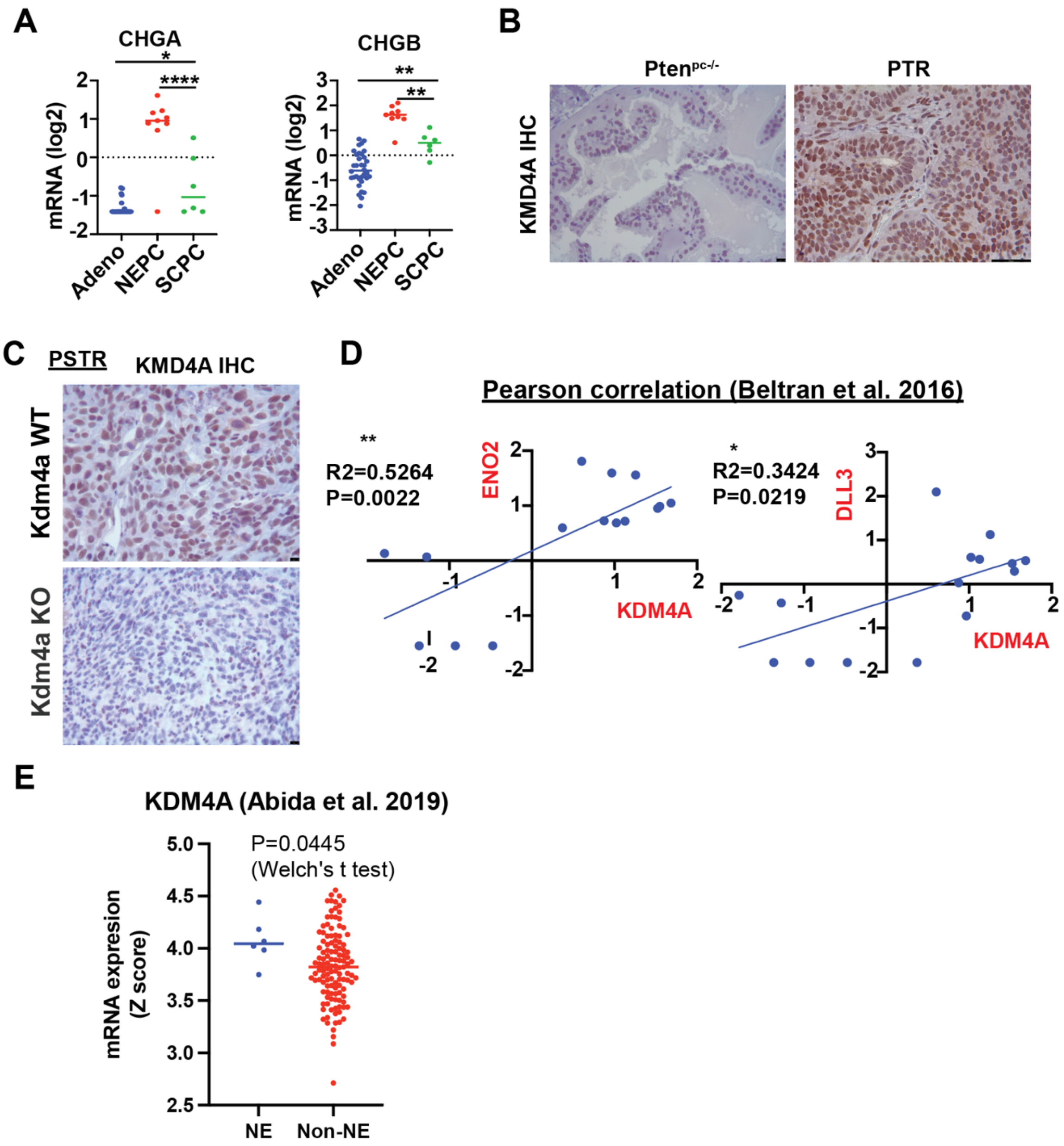


**Fig. S1**. (**A**) *CHGA* and *CHGB* mRNA expression in adeno, NEPC and SCPC from the Beltran et al. RNA-seq dataset. (**B**) KDM4A IHC in primary tumors from $Pten^{pc-/-}$ and PTR mice. (**C**) Validation of the specificity of anti-KDM4A antibody in IHC using *Kdm4a*-KO PSTR cells and parental cells. (**D**) Correlation of *KDM4A* mRNA expression with the mRNA expression of NE markers ENO2 and DLL3 in the Beltran et al. RNA-seq dataset. (**E**) *KDM4A* mRNA expression in mCRPC with NE features and without NE features in the Abida et al. RNA-seq dataset.

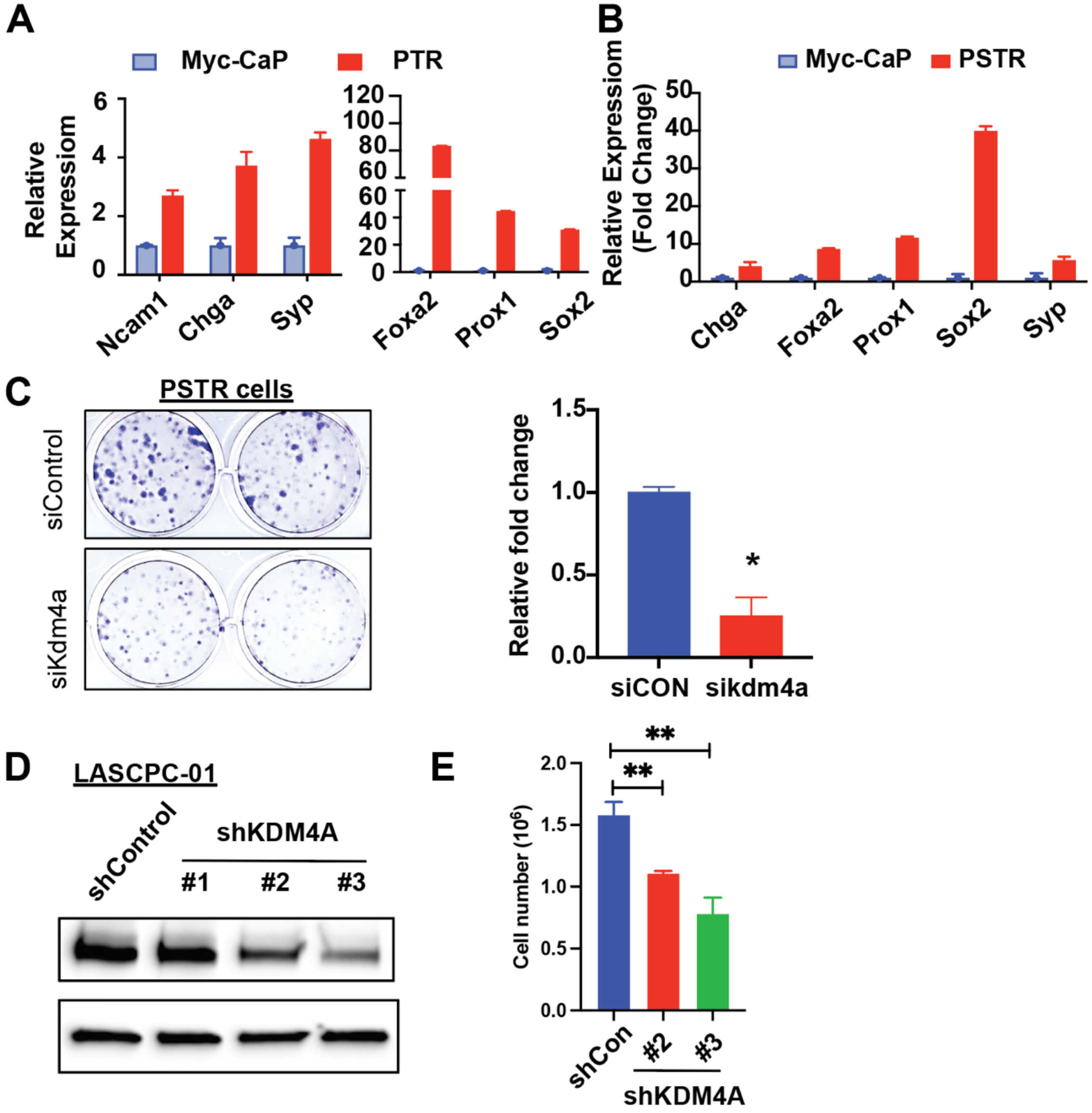


**Fig. S2**. (**A-B**) qRT-PCR analysis of NE markers in PTR, PSTR and MYC-CaP cells. (**C**) The effect of siRNA KD of *Kdm4a* in PSTR cells on cell proliferation as shown by foci-forming assay. (**D**) WB analysis of KDM4A expression in *KDM4A*-KD and control LASCPC-01 cells. (**E**) The effect of *KDM4A* KD on cell proliferation in LASCPC-01 cells measured by direct cell counting.

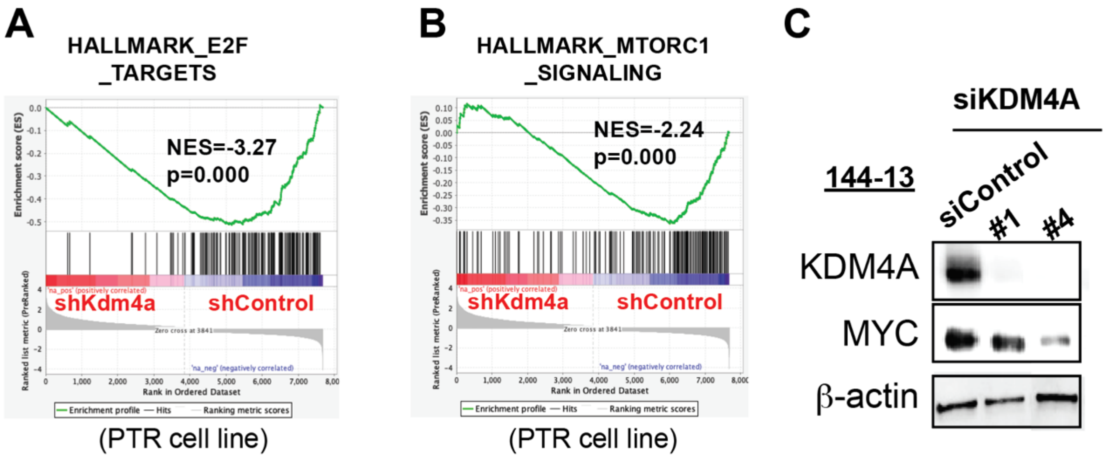


**Fig. S3**. (**A-B**) GSEA analysis of RNA-seq data from *Kdm4a*-KD PTR cells and control cells identified E2F (**A**) and mTORC1 (**B**) pathways among the top pathways downregulated in *Kdm4a*-KD cells. (**C**) WB analysis of MYC and KDM4A expression in *KDM4A*-KD and control 144-13 cells.

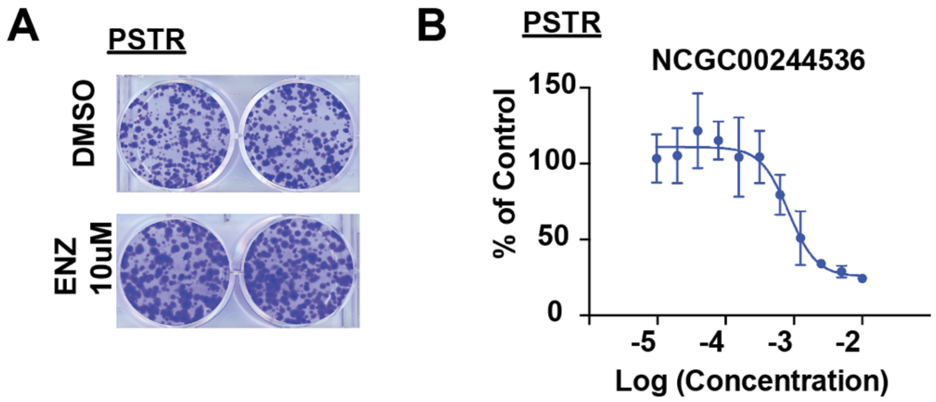


**Fig. S4**. (**A**) Foci-forming assay was used to determine the effect of enzalutamide (ENZ) on the proliferation of PSTR cells. (**B**) Dosage-response of KDM4 inhibitor NCGC00244536 in PSTR cells as measured by cell proliferation assay.

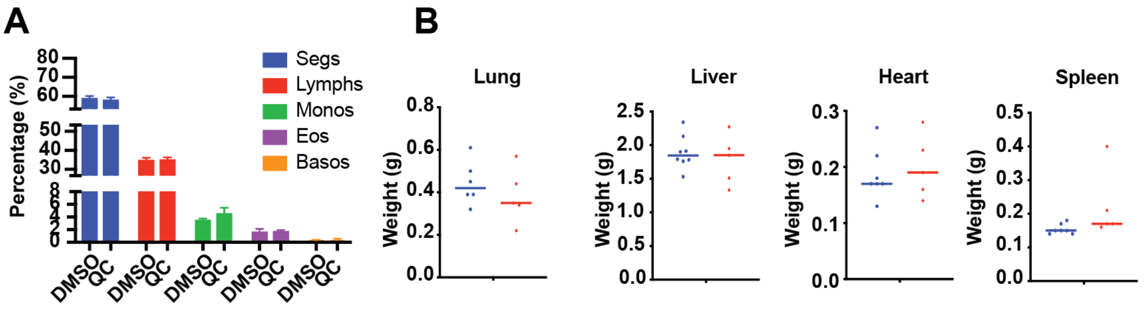


**Fig. S5**. (**A**) Peripheral blood counts from mice treated with QC6352 or vehicle control (n=3). (**B**) Weights of major organs from mice treated with QC6352 and vehicle control.

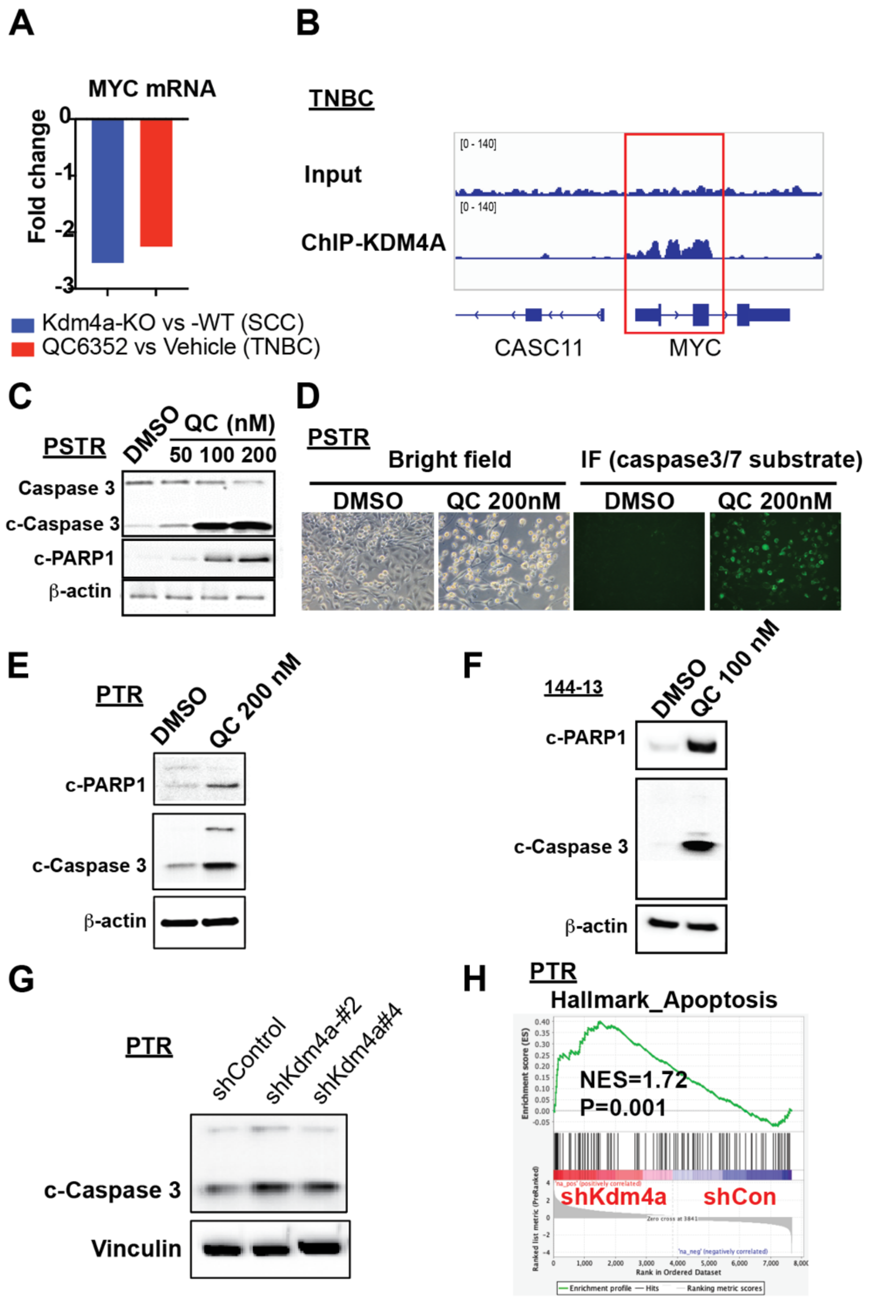


**Fig. S6**. (**A**) The effect of *Kdm4a* KO in SCC or QC6352 treatment in TNBC on MYC mRNA expression compared to *Kdm4a* WT cells or vehicle treatment using publicly available RNA-seq datasets. (**B**) KDM4A ChIP-seq peaks at *MYC* locus in TNBC cells using a publicly available RNA-seq dataset. (**C**) Western blot analysis of cleaved caspase 3 (c-Caspase 3) and cleaved PARP1 (c-PARP1) in PSTR cells treated with different concentrations of QC6352. (**D**) Fluorescence signal detected by live cell imaging with Cytation 5 in PSTR cells incubated with caspase3/7 substrates. (**E-F**) Western blot analysis of c-Caspase 3 and c-PARP1 in PTR cells (**E**) and 144-13 cells (**F**) treated with QC6352. (**G**) Western blot analysis of c-Caspase 3 in PTR cells infected control shRNA and Kdm4a-specific shRNAs. (**H**) GSEA analysis identified apoptosis as a pathway that is induced in *Kdm4a*-KD PTR cells compared to control cells.

| Primer ID | Species | Forward | Reverse |
|---|---|---|---|
| Gapdh | Mouse | AGGTCGGTGTGAACGGATTTG | TGTAGACCATGTAGTTGAGGTCA |
| Kdm4a | | ACCATCAGGTGGAATTTGGA | CTAGGTACGGGGTGGACAGA |
| Chga | | TTCCATGCAGGCTACAAAG | GTCTTTCCATCTCCATCCAC |
| cMyc | | CGGACACACAACGTCTTGGAA | AGGATGTAGGCGGTGGCTTTT |
| Syp | | CAGTTCCGGGTGGTCAAGG | ACTCTCCGTCTTGTTGGCAC |
| beta-actin | | CTAAGGCCAACCGTGAAAG | ACCAGAGGCATACAGGGACA |
| Ncam | | GACAGAACCCGAAAAGGGC | GTTGGGGACCGTCTTGACTT |
| GAPDH | Human | CAGGAGGCATTGCTGATGAT | GAAGGCTGGGGCTCATTT |
| KDM4A | | AGAGTTCCGCAAGATAGCCAA | CCACCAAGTCCAGGATTGTTCT |
| CHGA | | TAAAGGGGATACCGAGGTGATG | TCGGAGTGTCTCAAAACATTCC |
| CMYC | | | |
| SYP | | CTGAGGTCACTCTCGGTCTTG | CTCGGCTTTGTGAAGGTGCT |
| BETA-ACTIN | | CGCTCAGGAGGAGCAATG | TGACAGGATCGAGAAGGAGA |
| NCAM | | CGGCATTTACAAGTGTGTGG | CACACAATCACGGCATCTTC |

**Table S1**. Primers sequences used in this study.

**Dataset S1**. Differentially expressed genes comparing NEPC to adeno-PCa in Beltra et al. RNA-seq dataset obtained from cBioportal.

**Dataset S2**. Differentially expressed genes comparing NEPC to SCPC in Beltra et al. RNA-seq dataset obtained from cBioportal.

**Dataset S3**. Differentially expressed genes comparing shKdm4a-PTR cells to shControl-PTR cells (FDR ≤0.05, Fold change≥1.5)
