## Supplementary material for "KDM4A promotes the progression of neuroendocrine prostate cancer": Suppl. Table 1

**Suppl.Table 1_DE_KDM4A_class A_vs_B-D_mRNA expression z-Scores relative to diploid samples**

| Gene | (A) NE (B/D) | (B) Pathology | (A) NE (B/D) | (B) Pathology | Log Ratio | p-Value | q-Value | Higher expression in |
|---|---|---|---|---|---|---|---|---|
| CHGB | 16.11 | 6.62 | 2.03 | 3.05 | 9.5 | 1.09E-09 | 1.44E-07 | (A) NE (B/D) |
| SLC26A9 | 7.66 | 0.45 | 3.59 | 0.82 | 7.21 | 2.95E-04 | 2.49E-03 | (A) NE (B/D) |
| RNA5SP111 | 11.83 | 4.77 | 2.25 | 3.4 | 7.06 | 5.03E-07 | 1.73E-05 | (A) NE (B/D) |
| GPC5-AS1 | 8.95 | 2.37 | 2.12 | 2.52 | 6.58 | 1.16E-06 | 3.35E-05 | (A) NE (B/D) |
| FOXA2 | 6.92 | 0.39 | 2.65 | 0.63 | 6.53 | 6.99E-05 | 8.22E-04 | (A) NE (B/D) |
| MEG3 | 10.81 | 4.35 | 4.38 | 2.06 | 6.46 | 2.00E-03 | 0.0107 | (A) NE (B/D) |
| DDC | 9.19 | 2.84 | 2.84 | 2.45 | 6.35 | 6.40E-05 | 7.67E-04 | (A) NE (B/D) |
| MAST1 | 8.01 | 1.83 | 1.35 | 1.37 | 6.18 | 2.14E-08 | 1.39E-06 | (A) NE (B/D) |
| UCN3 | 6.29 | 0.17 | 3.08 | 0.36 | 6.12 | 3.29E-04 | 2.71E-03 | (A) NE (B/D) |
| MARCH11 | 6.25 | 0.14 | 1.37 | 0.31 | 6.11 | 7.41E-07 | 2.35E-05 | (A) NE (B/D) |
| CHGA | 6.31 | 0.26 | 2.48 | 0.57 | 6.06 | 7.51E-05 | 8.70E-04 | (A) NE (B/D) |
| DLK1 | 6.12 | 0.06 | 4.62 | 0.34 | 6.06 | 4.33E-03 | 0.0193 | (A) NE (B/D) |
| SCG2 | 5.99 | 0.02 | 2.35 | 0.1 | 5.98 | 6.18E-05 | 7.46E-04 | (A) NE (B/D) |
| LINC00261 | 6.8 | 0.85 | 3.05 | 1.15 | 5.96 | 3.23E-04 | 2.67E-03 | (A) NE (B/D) |
| C1ORF127 | 7.71 | 1.82 | 3.81 | 1.19 | 5.89 | 1.59E-03 | 8.99E-03 | (A) NE (B/D) |
| XKR7 | 6.64 | 0.89 | 0.31 | 1.8 | 5.75 | 3.69E-20 | 6.10E-16 | (A) NE (B/D) |
| INSM1 | 5.95 | 0.2 | 1.51 | 0.47 | 5.75 | 2.32E-06 | 5.77E-05 | (A) NE (B/D) |
| NKX2-1-AS1 | 6.83 | 1.1 | 3.62 | 1.25 | 5.73 | 1.34E-03 | 7.90E-03 | (A) NE (B/D) |
| SPAG11A | 5.78 | 0.16 | 4.37 | 0.5 | 5.62 | 4.77E-03 | 0.0208 | (A) NE (B/D) |
| LHX2 | 6.32 | 0.71 | 2.08 | 0.95 | 5.61 | 2.64E-05 | 3.80E-04 | (A) NE (B/D) |
| NR0B1 | 5.8 | 0.2 | 3.8 | 0.5 | 5.6 | 2.22E-03 | 0.0116 | (A) NE (B/D) |
| ATCAY | 5.8 | 0.24 | 2.57 | 0.61 | 5.56 | 1.77E-04 | 1.68E-03 | (A) NE (B/D) |
| TMEM61 | 9.01 | 3.48 | 1.37 | 1.82 | 5.53 | 2.10E-08 | 1.37E-06 | (A) NE (B/D) |
| KCNB2 | 6.2 | 0.69 | 2.89 | 0.98 | 5.51 | 3.95E-04 | 3.12E-03 | (A) NE (B/D) |
| CSAG2 | 7.11 | 1.63 | 4.05 | 1.84 | 5.49 | 3.42E-03 | 0.0161 | (A) NE (B/D) |
| RIMBP2 | 7.28 | 1.83 | 1.05 | 1.64 | 5.45 | 1.31E-10 | 3.06E-08 | (A) NE (B/D) |
| SCN3A | 8.14 | 2.7 | 3.4 | 1.47 | 5.44 | 1.20E-03 | 7.27E-03 | (A) NE (B/D) |
| ELAVL4 | 5.96 | 0.57 | 2.27 | 0.8 | 5.4 | 7.98E-05 | 9.08E-04 | (A) NE (B/D) |
| LINC00404 | 5.46 | 0.08 | 2.22 | 0.35 | 5.38 | 8.44E-05 | 9.53E-04 | (A) NE (B/D) |
| RNF183 | 6.05 | 0.8 | 2.25 | 0.72 | 5.25 | 9.68E-05 | 1.06E-03 | (A) NE (B/D) |
| TFF3 | 10.75 | 5.58 | 1.31 | 2.31 | 5.18 | 9.18E-09 | 7.41E-07 | (A) NE (B/D) |
| AOC1 | 6.99 | 1.84 | 3.53 | 1.4 | 5.15 | 2.22E-03 | 0.0116 | (A) NE (B/D) |
| CEACAM5 | 5.13 | 0.07 | 2.12 | 0.37 | 5.06 | 9.24E-05 | 1.02E-03 | (A) NE (B/D) |
| MEG9 | 5.42 | 0.35 | 3.68 | 0.8 | 5.06 | 3.25E-03 | 0.0155 | (A) NE (B/D) |
| CCDC33 | 5.32 | 0.29 | 2.19 | 0.57 | 5.03 | 1.15E-04 | 1.20E-03 | (A) NE (B/D) |
| OGDHL | 7.79 | 2.83 | 0.78 | 1.85 | 4.96 | 1.47E-13 | 1.58E-10 | (A) NE (B/D) |
| TCEAL5 | 6 | 1.11 | 4.01 | 1.37 | 4.89 | 6.23E-03 | 0.0253 | (A) NE (B/D) |
| SOX1-OT | 4.87 | 0.03 | 1.97 | 0.15 | 4.85 | 7.58E-05 | 8.76E-04 | (A) NE (B/D) |
| DLL3 | 4.89 | 0.08 | 2.24 | 0.25 | 4.81 | 1.97E-04 | 1.82E-03 | (A) NE (B/D) |
| ATP1A3 | 4.84 | 0.12 | 1.35 | 0.29 | 4.72 | 5.19E-06 | 1.09E-04 | (A) NE (B/D) |
| ANKRD65 | 6.41 | 1.7 | 0.79 | 1.56 | 4.71 | 1.14E-12 | 8.61E-10 | (A) NE (B/D) |
| KRT8P30 | 5 | 0.35 | 0.99 | 0.73 | 4.65 | 7.43E-08 | 3.81E-06 | (A) NE (B/D) |
| BRINP1 | 6.08 | 1.5 | 2.44 | 1.4 | 4.57 | 3.70E-04 | 2.97E-03 | (A) NE (B/D) |
| MAGEA2 | 4.94 | 0.39 | 2.81 | 0.96 | 4.55 | 1.18E-03 | 7.16E-03 | (A) NE (B/D) |
| FABP6 | 5.53 | 1.06 | 2.28 | 1.23 | 4.47 | 2.81E-04 | 2.39E-03 | (A) NE (B/D) |
| MIR-429/429 | 6.83 | 2.39 | 1.04 | 2.41 | 4.44 | 2.62E-09 | 2.97E-07 | (A) NE (B/D) |
| SLC4A11 | 5.89 | 1.46 | 1.67 | 1.3 | 4.44 | 1.58E-05 | 2.59E-04 | (A) NE (B/D) |
| MIR-642/642 | 5.3 | 0.86 | 2.56 | 1.02 | 4.44 | 7.29E-04 | 4.99E-03 | (A) NE (B/D) |
| MIR-642A/3P | 5.3 | 0.86 | 2.56 | 1.02 | 4.44 | 7.29E-04 | 4.99E-03 | (A) NE (B/D) |
| MIR-642A/5P | 5.3 | 0.86 | 2.56 | 1.02 | 4.44 | 7.29E-04 | 4.99E-03 | (A) NE (B/D) |
| MIR-642A/64 | 5.3 | 0.86 | 2.56 | 1.02 | 4.44 | 7.29E-04 | 4.99E-03 | (A) NE (B/D) |
| MIR-642A/64 | 5.3 | 0.86 | 2.56 | 1.02 | 4.44 | 7.29E-04 | 4.99E-03 | (A) NE (B/D) |
| TOX | 5.73 | 1.3 | 2.9 | 1.1 | 4.43 | 1.67E-03 | 9.34E-03 | (A) NE (B/D) |
| TMEM176B | 5.79 | 1.38 | 2.27 | 1.54 | 4.41 | 2.57E-04 | 2.22E-03 | (A) NE (B/D) |
| PROM1 | 6.28 | 1.87 | 2.7 | 1.56 | 4.41 | 9.72E-04 | 6.19E-03 | (A) NE (B/D) |
| SLITRK6 | 7.66 | 3.26 | 2.61 | 2.11 | 4.4 | 6.99E-04 | 4.85E-03 | (A) NE (B/D) |
| BAALC-AS2 | 4.5 | 0.12 | 2.42 | 0.23 | 4.38 | 6.14E-04 | 4.38E-03 | (A) NE (B/D) |
| DDX25 | 4.6 | 0.22 | 2.63 | 0.36 | 4.38 | 1.05E-03 | 6.56E-03 | (A) NE (B/D) |
| MIR-200B/20 | 5.89 | 1.55 | 1.38 | 1.75 | 4.34 | 7.75E-07 | 2.43E-05 | (A) NE (B/D) |
| MIR-200B/20 | 5.89 | 1.55 | 1.38 | 1.75 | 4.34 | 7.75E-07 | 2.43E-05 | (A) NE (B/D) |
| MIR-200B/3P | 5.89 | 1.55 | 1.38 | 1.75 | 4.34 | 7.75E-07 | 2.43E-05 | (A) NE (B/D) |
| MIR-200B/5P | 5.89 | 1.55 | 1.38 | 1.75 | 4.34 | 7.75E-07 | 2.43E-05 | (A) NE (B/D) |
| KIAA0408 | 7 | 2.67 | 2.47 | 1.35 | 4.34 | 6.08E-04 | 4.35E-03 | (A) NE (B/D) |

| | | | | | | | | |
|---|---|---|---|---|---|---|---|---|
| MIAT | 8.53 | 4.2 | 3.26 | 1.43 | 4.33 | 3.83E-03 | 0.0176 | (A) NE (B/D) |
| PCSK1 | 4.36 | 0.07 | 2.19 | 0.26 | 4.29 | 3.64E-04 | 2.93E-03 | (A) NE (B/D) |
| RN7SKP106 | 4.99 | 0.74 | 3.2 | 1.4 | 4.25 | 3.85E-03 | 0.0176 | (A) NE (B/D) |
| GABRA3 | 5.93 | 1.68 | 3.46 | 1.92 | 4.25 | 5.91E-03 | 0.0243 | (A) NE (B/D) |
| WDR62 | 8.72 | 4.5 | 1.22 | 1.49 | 4.22 | 2.52E-07 | 9.98E-06 | (A) NE (B/D) |
| DAZ2 | 4.43 | 0.23 | 3.35 | 0.71 | 4.2 | 5.51E-03 | 0.0231 | (A) NE (B/D) |
| KCNK17 | 4.82 | 0.63 | 2.47 | 0.91 | 4.19 | 8.47E-04 | 5.59E-03 | (A) NE (B/D) |
| CPLX2 | 4.32 | 0.16 | 0.72 | 0.45 | 4.16 | 1.94E-08 | 1.30E-06 | (A) NE (B/D) |
| C6ORF141 | 4.81 | 0.65 | 1.01 | 1.19 | 4.16 | 3.73E-08 | 2.20E-06 | (A) NE (B/D) |
| MS4A8 | 4.38 | 0.22 | 1.98 | 0.56 | 4.16 | 2.10E-04 | 1.91E-03 | (A) NE (B/D) |
| PTCHD3P3 | 5.13 | 0.99 | 3.01 | 1.19 | 4.14 | 3.15E-03 | 0.0152 | (A) NE (B/D) |
| NELL1 | 4.44 | 0.33 | 2.28 | 0.94 | 4.11 | 5.51E-04 | 4.04E-03 | (A) NE (B/D) |
| TEX101 | 4.46 | 0.35 | 2.4 | 0.75 | 4.11 | 8.20E-04 | 5.46E-03 | (A) NE (B/D) |
| GOLGA7B | 4.99 | 0.9 | 0.58 | 0.89 | 4.1 | 8.76E-13 | 7.27E-10 | (A) NE (B/D) |
| RPS23P3 | 4.31 | 0.27 | 3.17 | 0.64 | 4.04 | 5.06E-03 | 0.0217 | (A) NE (B/D) |
| FBLL1 | 6.45 | 2.41 | 0.6 | 1.87 | 4.03 | 3.40E-13 | 3.50E-10 | (A) NE (B/D) |
| SYP-AS1 | 4.9 | 0.87 | 0.59 | 0.9 | 4.03 | 1.41E-12 | 9.30E-10 | (A) NE (B/D) |
| LINC00958 | 4.72 | 0.7 | 2.56 | 1.05 | 4.02 | 1.37E-03 | 8.04E-03 | (A) NE (B/D) |
| EPOR | 5.08 | 1.07 | 1.74 | 1.61 | 4.01 | 4.49E-05 | 5.80E-04 | (A) NE (B/D) |
| PTTG1 | 9.72 | 5.72 | 1.17 | 2.12 | 4 | 1.07E-07 | 5.05E-06 | (A) NE (B/D) |
| PHF19 | 4.46 | 0.48 | 0.77 | 0.76 | 3.98 | 7.03E-09 | 5.96E-07 | (A) NE (B/D) |
| CNTNAP5 | 4.5 | 0.54 | 2.3 | 0.87 | 3.96 | 7.67E-04 | 5.19E-03 | (A) NE (B/D) |
| KLHDC8A | 5.92 | 1.97 | 2.35 | 1.53 | 3.95 | 7.66E-04 | 5.19E-03 | (A) NE (B/D) |
| GCK | 3.97 | 0.03 | 2.28 | 0.1 | 3.94 | 8.35E-04 | 5.54E-03 | (A) NE (B/D) |
| SNORD114-2 | 4.59 | 0.66 | 2.92 | 1.67 | 3.93 | 3.43E-03 | 0.0162 | (A) NE (B/D) |
| UCHL1 | 4.07 | 0.15 | 2.28 | 0.72 | 3.91 | 8.09E-04 | 5.41E-03 | (A) NE (B/D) |
| GIPR | 4.12 | 0.23 | 2.12 | 0.39 | 3.89 | 5.62E-04 | 4.10E-03 | (A) NE (B/D) |
| MIR-4793/3P | 8.1 | 4.27 | 0.57 | 2.12 | 3.84 | 1.02E-11 | 3.98E-09 | (A) NE (B/D) |
| MIR-4793/5P | 8.1 | 4.27 | 0.57 | 2.12 | 3.84 | 1.02E-11 | 3.98E-09 | (A) NE (B/D) |
| MYT1 | 4.87 | 1.03 | 0.94 | 1.53 | 3.84 | 6.42E-09 | 5.66E-07 | (A) NE (B/D) |
| BPIFB6 | 4.22 | 0.4 | 2.19 | 0.82 | 3.83 | 6.95E-04 | 4.83E-03 | (A) NE (B/D) |
| KIFC1 | 7.77 | 3.95 | 0.66 | 2.09 | 3.82 | 3.14E-11 | 9.40E-09 | (A) NE (B/D) |
| HES6 | 4.16 | 0.34 | 1.67 | 0.65 | 3.82 | 1.06E-04 | 1.14E-03 | (A) NE (B/D) |
| FOXD3 | 5.15 | 1.34 | 2.35 | 1.28 | 3.81 | 1.03E-03 | 6.47E-03 | (A) NE (B/D) |
| GUCY1B2 | 4.35 | 0.55 | 1.3 | 1 | 3.8 | 6.78E-06 | 1.34E-04 | (A) NE (B/D) |
| RGL3 | 9.13 | 5.33 | 2.08 | 1.51 | 3.8 | 3.94E-04 | 3.12E-03 | (A) NE (B/D) |
| FGF14 | 5.19 | 1.39 | 2.85 | 1.11 | 3.8 | 3.77E-03 | 0.0173 | (A) NE (B/D) |
| MIR-130B/13 | 4.27 | 0.47 | 1.02 | 1.1 | 3.79 | 1.91E-07 | 8.06E-06 | (A) NE (B/D) |
| MIR-130B/13 | 4.27 | 0.47 | 1.02 | 1.1 | 3.79 | 1.91E-07 | 8.06E-06 | (A) NE (B/D) |
| MIR-130B/3P | 4.27 | 0.47 | 1.02 | 1.1 | 3.79 | 1.91E-07 | 8.06E-06 | (A) NE (B/D) |
| MIR-130B/5P | 4.27 | 0.47 | 1.02 | 1.1 | 3.79 | 1.91E-07 | 8.06E-06 | (A) NE (B/D) |
| C2ORF70 | 6.12 | 2.35 | 1.18 | 1.7 | 3.77 | 4.14E-07 | 1.48E-05 | (A) NE (B/D) |
| LINC00599 | 4 | 0.23 | 2.79 | 0.37 | 3.77 | 3.62E-03 | 0.0168 | (A) NE (B/D) |
| MAGEA12 | 4.07 | 0.31 | 2.32 | 0.77 | 3.76 | 1.15E-03 | 7.04E-03 | (A) NE (B/D) |
| SFTA3 | 3.76 | 0.02 | 2.19 | 0.07 | 3.74 | 9.01E-04 | 5.84E-03 | (A) NE (B/D) |
| LINC00951 | 3.92 | 0.19 | 2.49 | 0.42 | 3.73 | 1.99E-03 | 0.0107 | (A) NE (B/D) |
| GPR148 | 4.24 | 0.52 | 0.52 | 1 | 3.72 | 2.08E-14 | 3.06E-11 | (A) NE (B/D) |
| ONECUT2 | 7.03 | 3.31 | 0.24 | 1.75 | 3.72 | 2.63E-14 | 3.56E-11 | (A) NE (B/D) |
| HAGLR | 6.03 | 2.31 | 1.78 | 1.55 | 3.72 | 1.13E-04 | 1.19E-03 | (A) NE (B/D) |
| IQGAP3 | 9.03 | 5.32 | 0.91 | 1.75 | 3.71 | 4.32E-09 | 4.13E-07 | (A) NE (B/D) |
| RASSF6 | 4.35 | 0.64 | 1.8 | 0.99 | 3.71 | 1.86E-04 | 1.74E-03 | (A) NE (B/D) |
| C6ORF183 | 4.01 | 0.3 | 2.66 | 0.59 | 3.71 | 2.96E-03 | 0.0144 | (A) NE (B/D) |
| SPTBN4 | 4.13 | 0.43 | 1.12 | 0.73 | 3.7 | 2.94E-06 | 6.90E-05 | (A) NE (B/D) |
| CXXC4 | 6.58 | 2.89 | 2.25 | 1.15 | 3.69 | 1.01E-03 | 6.34E-03 | (A) NE (B/D) |
| PTPRN2 | 8.12 | 4.44 | 0.38 | 1.69 | 3.68 | 1.65E-14 | 2.65E-11 | (A) NE (B/D) |
| FABP5P1 | 6.15 | 2.47 | 1.74 | 2.17 | 3.68 | 7.85E-05 | 8.97E-04 | (A) NE (B/D) |
| ATP8A2 | 5.56 | 1.88 | 1.26 | 1.84 | 3.67 | 1.51E-06 | 4.15E-05 | (A) NE (B/D) |
| GRM4 | 4.25 | 0.59 | 1.1 | 0.74 | 3.67 | 2.78E-06 | 6.61E-05 | (A) NE (B/D) |
| ETV5 | 6.01 | 2.34 | 2.12 | 0.94 | 3.67 | 7.00E-04 | 4.85E-03 | (A) NE (B/D) |
| SCG3 | 3.66 | 0.01 | 1.61 | 0.08 | 3.65 | 1.36E-04 | 1.37E-03 | (A) NE (B/D) |
| PHGR1 | 4.47 | 0.83 | 0.9 | 1.36 | 3.63 | 1.23E-08 | 9.21E-07 | (A) NE (B/D) |
| FOXD2-AS1 | 4.73 | 1.1 | 1.62 | 0.77 | 3.63 | 1.07E-04 | 1.14E-03 | (A) NE (B/D) |
| PI4KAP1 | 7 | 3.39 | 1.36 | 1.43 | 3.61 | 9.03E-06 | 1.68E-04 | (A) NE (B/D) |
| MIR-4321/43 | 6 | 2.39 | 1.47 | 1.62 | 3.61 | 1.89E-05 | 2.95E-04 | (A) NE (B/D) |
| MIR-656/3P | 3.71 | 0.12 | 2.87 | 0.41 | 3.6 | 5.56E-03 | 0.0232 | (A) NE (B/D) |

| | | | | | | | | |
|---|---|---|---|---|---|---|---|---|
| MIR-656/5P | 3.71 | 0.12 | 2.87 | 0.41 | 3.6 | 5.56E-03 | 0.0232 | (A) NE (B/D) |
| MIR-656/656 | 3.71 | 0.12 | 2.87 | 0.41 | 3.6 | 5.56E-03 | 0.0232 | (A) NE (B/D) |
| ITPR3 | 8.22 | 4.64 | 0.39 | 1.16 | 3.59 | 1.18E-17 | 7.58E-14 | (A) NE (B/D) |
| UNC13A | 3.84 | 0.25 | 0.42 | 0.69 | 3.59 | 4.85E-15 | 1.01E-11 | (A) NE (B/D) |
| SGSM1 | 5.4 | 1.8 | 0.69 | 1.12 | 3.59 | 7.99E-11 | 1.94E-08 | (A) NE (B/D) |
| EIF3KP1 | 4.19 | 0.61 | 1.16 | 0.97 | 3.59 | 3.13E-06 | 7.27E-05 | (A) NE (B/D) |
| STMN1 | 5.93 | 2.35 | 1.64 | 1.96 | 3.59 | 5.47E-05 | 6.79E-04 | (A) NE (B/D) |
| MIR-3917/39 | 4.93 | 1.36 | 1.07 | 1.58 | 3.58 | 2.31E-07 | 9.34E-06 | (A) NE (B/D) |
| RASD1 | 4.02 | 0.44 | 1.72 | 0.82 | 3.58 | 1.92E-04 | 1.79E-03 | (A) NE (B/D) |
| LINC00342 | 7.33 | 3.79 | 1.14 | 1.24 | 3.54 | 1.47E-06 | 4.06E-05 | (A) NE (B/D) |
| MIR-412/3P | 3.66 | 0.12 | 2.88 | 0.53 | 3.54 | 6.14E-03 | 0.025 | (A) NE (B/D) |
| MIR-412/412 | 3.66 | 0.12 | 2.88 | 0.53 | 3.54 | 6.14E-03 | 0.025 | (A) NE (B/D) |
| MIR-412/5P | 3.66 | 0.12 | 2.88 | 0.53 | 3.54 | 6.14E-03 | 0.025 | (A) NE (B/D) |
| PRC1-AS1 | 8.12 | 4.59 | 0.98 | 1.55 | 3.53 | 5.50E-08 | 2.99E-06 | (A) NE (B/D) |
| RIMKLA | 4.98 | 1.47 | 0.71 | 1.09 | 3.51 | 3.37E-10 | 6.14E-08 | (A) NE (B/D) |
| LINC00460 | 3.82 | 0.31 | 1.83 | 1.03 | 3.51 | 3.18E-04 | 2.64E-03 | (A) NE (B/D) |
| SNAI1P1 | 4.25 | 0.78 | 0.98 | 1.02 | 3.48 | 3.49E-07 | 1.28E-05 | (A) NE (B/D) |
| MIR-1178/11 | 4.63 | 1.15 | 1.12 | 1.13 | 3.48 | 1.86E-06 | 4.87E-05 | (A) NE (B/D) |
| MIR-1178/3P | 4.63 | 1.15 | 1.12 | 1.13 | 3.48 | 1.86E-06 | 4.87E-05 | (A) NE (B/D) |
| MIR-1178/5P | 4.63 | 1.15 | 1.12 | 1.13 | 3.48 | 1.86E-06 | 4.87E-05 | (A) NE (B/D) |
| FBXO43 | 5.9 | 2.44 | 1.42 | 1.28 | 3.47 | 2.76E-05 | 3.94E-04 | (A) NE (B/D) |
| CNTNAP2 | 8.21 | 4.76 | 1.22 | 2.04 | 3.45 | 1.99E-06 | 5.12E-05 | (A) NE (B/D) |
| C2CD4B | 7.12 | 3.68 | 2.19 | 1.89 | 3.45 | 1.13E-03 | 6.91E-03 | (A) NE (B/D) |
| TNFRSF25 | 4.01 | 0.57 | 2.06 | 0.84 | 3.44 | 9.15E-04 | 5.91E-03 | (A) NE (B/D) |
| RIPPLY3 | 4.2 | 0.77 | 0.87 | 1.06 | 3.43 | 4.78E-08 | 2.67E-06 | (A) NE (B/D) |
| SCGB2A1 | 7.45 | 4.02 | 1.55 | 2.21 | 3.43 | 4.64E-05 | 5.95E-04 | (A) NE (B/D) |
| C14ORF132 | 7.01 | 3.59 | 0.64 | 1.7 | 3.42 | 2.67E-11 | 8.34E-09 | (A) NE (B/D) |
| BCAM | 4.97 | 1.56 | 0.8 | 1.95 | 3.41 | 3.21E-09 | 3.39E-07 | (A) NE (B/D) |
| CDC25B | 4.56 | 1.16 | 1.61 | 1.17 | 3.4 | 1.25E-04 | 1.28E-03 | (A) NE (B/D) |
| APLP1 | 3.55 | 0.16 | 1.32 | 0.75 | 3.39 | 3.20E-05 | 4.48E-04 | (A) NE (B/D) |
| SCGN | 3.56 | 0.17 | 1.4 | 0.42 | 3.39 | 7.67E-05 | 8.82E-04 | (A) NE (B/D) |
| SYT11 | 4 | 0.62 | 1.62 | 0.83 | 3.38 | 1.81E-04 | 1.70E-03 | (A) NE (B/D) |
| HOXD8 | 5.69 | 2.31 | 1.94 | 1.27 | 3.38 | 6.01E-04 | 4.32E-03 | (A) NE (B/D) |
| LRP4 | 5.76 | 2.38 | 2.09 | 1.19 | 3.38 | 1.05E-03 | 6.53E-03 | (A) NE (B/D) |
| STMN3 | 4.83 | 1.48 | 0.29 | 1.21 | 3.35 | 6.73E-18 | 5.77E-14 | (A) NE (B/D) |
| DNALI1 | 4.29 | 0.94 | 1.51 | 1.38 | 3.35 | 6.44E-05 | 7.70E-04 | (A) NE (B/D) |
| TAS2R38 | 3.67 | 0.32 | 2.36 | 0.66 | 3.35 | 2.69E-03 | 0.0135 | (A) NE (B/D) |
| KIF2C | 6.69 | 3.35 | 1.2 | 1.76 | 3.34 | 2.69E-06 | 6.43E-05 | (A) NE (B/D) |
| HOXD4 | 3.66 | 0.32 | 1.53 | 0.6 | 3.34 | 1.48E-04 | 1.47E-03 | (A) NE (B/D) |
| NAV2-AS1 | 5.03 | 1.69 | 1.9 | 1.44 | 3.34 | 5.11E-04 | 3.80E-03 | (A) NE (B/D) |
| BCRP9 | 3.44 | 0.1 | 2.47 | 0.28 | 3.34 | 3.65E-03 | 0.0169 | (A) NE (B/D) |
| TPBGL | 4.58 | 1.24 | 0.67 | 1.14 | 3.33 | 1.36E-10 | 3.13E-08 | (A) NE (B/D) |
| FAM3B | 5.8 | 2.47 | 0.61 | 2.11 | 3.33 | 6.62E-10 | 1.05E-07 | (A) NE (B/D) |
| DIRAS2 | 3.8 | 0.46 | 1.62 | 1.14 | 3.33 | 1.61E-04 | 1.56E-03 | (A) NE (B/D) |
| CDCA8 | 4.45 | 1.13 | 1.24 | 1.34 | 3.32 | 7.41E-06 | 1.44E-04 | (A) NE (B/D) |
| NKX2-2 | 4.89 | 1.57 | 1.51 | 1.85 | 3.32 | 5.29E-05 | 6.61E-04 | (A) NE (B/D) |
| LINC00491 | 3.55 | 0.23 | 1.48 | 0.4 | 3.32 | 1.34E-04 | 1.36E-03 | (A) NE (B/D) |
| PROX1 | 3.5 | 0.19 | 1.44 | 0.53 | 3.31 | 1.01E-04 | 1.09E-03 | (A) NE (B/D) |
| VGF | 3.35 | 0.05 | 1.37 | 0.31 | 3.3 | 8.20E-05 | 9.30E-04 | (A) NE (B/D) |
| ETV1 | 3.88 | 0.57 | 1.65 | 1.68 | 3.3 | 1.52E-04 | 1.50E-03 | (A) NE (B/D) |
| TUBB2B | 3.64 | 0.34 | 1.56 | 0.49 | 3.3 | 1.98E-04 | 1.83E-03 | (A) NE (B/D) |
| FABP5P7 | 13.5 | 10.2 | 1.96 | 2.04 | 3.3 | 6.54E-04 | 4.60E-03 | (A) NE (B/D) |
| NCAM1-AS1 | 4.06 | 0.77 | 2.04 | 0.89 | 3.3 | 1.12E-03 | 6.90E-03 | (A) NE (B/D) |
| PTPRN | 3.3 | 0.01 | 2 | 0.05 | 3.29 | 1.14E-03 | 6.99E-03 | (A) NE (B/D) |
| SEC11C | 3.55 | 0.27 | 1.3 | 0.68 | 3.28 | 4.17E-05 | 5.48E-04 | (A) NE (B/D) |
| SIX2 | 3.29 | 0.01 | 0.74 | 0.07 | 3.27 | 9.89E-07 | 2.96E-05 | (A) NE (B/D) |
| SPATA46 | 4.39 | 1.12 | 1.4 | 1.64 | 3.27 | 2.74E-05 | 3.92E-04 | (A) NE (B/D) |
| SPRY2 | 7.41 | 4.14 | 1.41 | 0.99 | 3.27 | 5.92E-05 | 7.22E-04 | (A) NE (B/D) |
| HOXD10 | 3.65 | 0.38 | 2.59 | 0.81 | 3.27 | 5.23E-03 | 0.0222 | (A) NE (B/D) |
| MAGEL2 | 3.84 | 0.57 | 2.89 | 0.93 | 3.27 | 9.32E-03 | 0.0339 | (A) NE (B/D) |
| CPHL1P | 7.79 | 4.53 | 1.72 | 1.98 | 3.26 | 2.30E-04 | 2.04E-03 | (A) NE (B/D) |
| MIR-5707/57 | 3.47 | 0.22 | 1.58 | 0.92 | 3.25 | 1.84E-04 | 1.73E-03 | (A) NE (B/D) |
| FABP5P2 | 7.45 | 4.21 | 1.85 | 1.64 | 3.25 | 4.99E-04 | 3.74E-03 | (A) NE (B/D) |
| MIR-200A/20 | 4.22 | 0.98 | 1.07 | 1.38 | 3.24 | 1.11E-06 | 3.23E-05 | (A) NE (B/D) |
| MIR-200A/20 | 4.22 | 0.98 | 1.07 | 1.38 | 3.24 | 1.11E-06 | 3.23E-05 | (A) NE (B/D) |

| | | | | | | | | |
|---|---|---|---|---|---|---|---|---|
| MIR-200A/3P | 4.22 | 0.98 | 1.07 | 1.38 | 3.24 | 1.11E-06 | 3.23E-05 | (A) NE (B/D) |
| MIR-200A/5P | 4.22 | 0.98 | 1.07 | 1.38 | 3.24 | 1.11E-06 | 3.23E-05 | (A) NE (B/D) |
| ACOT11 | 5.13 | 1.89 | 1.8 | 1.15 | 3.24 | 4.83E-04 | 3.64E-03 | (A) NE (B/D) |
| SV2B | 3.7 | 0.46 | 2.16 | 0.62 | 3.24 | 1.93E-03 | 0.0104 | (A) NE (B/D) |
| FOXD1 | 6.5 | 3.28 | 0.53 | 1.53 | 3.22 | 2.64E-12 | 1.42E-09 | (A) NE (B/D) |
| TYMS | 4.57 | 1.36 | 0.99 | 1.49 | 3.21 | 2.92E-07 | 1.13E-05 | (A) NE (B/D) |
| CCDC154 | 5.98 | 2.78 | 1.6 | 1.18 | 3.2 | 1.90E-04 | 1.77E-03 | (A) NE (B/D) |
| SEZ6L2 | 8.19 | 5 | 0.71 | 2.32 | 3.19 | 2.92E-08 | 1.81E-06 | (A) NE (B/D) |
| CNKSR3 | 4.1 | 0.91 | 1.07 | 1.19 | 3.19 | 2.13E-06 | 5.38E-05 | (A) NE (B/D) |
| SHC3 | 3.62 | 0.43 | 1.78 | 0.8 | 3.19 | 5.73E-04 | 4.16E-03 | (A) NE (B/D) |
| DDX53 | 3.47 | 0.29 | 2.49 | 1.08 | 3.18 | 4.77E-03 | 0.0208 | (A) NE (B/D) |
| STXBP1 | 6.95 | 3.78 | 0.37 | 1.28 | 3.17 | 1.55E-15 | 3.99E-12 | (A) NE (B/D) |
| B3GALT5 | 3.98 | 0.8 | 1.78 | 1.21 | 3.17 | 4.82E-04 | 3.64E-03 | (A) NE (B/D) |
| ABLIM2 | 4.13 | 0.96 | 1.89 | 0.82 | 3.17 | 8.95E-04 | 5.81E-03 | (A) NE (B/D) |
| SLC44A5 | 3.82 | 0.66 | 2.2 | 1.41 | 3.16 | 2.23E-03 | 0.0116 | (A) NE (B/D) |
| PGBD5 | 3.76 | 0.61 | 0.6 | 0.87 | 3.15 | 1.93E-10 | 4.08E-08 | (A) NE (B/D) |
| MIR-4648/46 | 4.86 | 1.71 | 0.87 | 1.43 | 3.15 | 4.33E-08 | 2.47E-06 | (A) NE (B/D) |
| SOX2 | 3.33 | 0.18 | 1.54 | 0.38 | 3.15 | 2.64E-04 | 2.28E-03 | (A) NE (B/D) |
| H2BFXP | 6.19 | 3.04 | 2.17 | 1.67 | 3.15 | 2.04E-03 | 0.0109 | (A) NE (B/D) |
| PLEKHB1 | 3.52 | 0.37 | 2.15 | 0.68 | 3.15 | 2.16E-03 | 0.0113 | (A) NE (B/D) |
| HMX3 | 3.19 | 0.03 | 2.46 | 0.12 | 3.15 | 4.87E-03 | 0.0211 | (A) NE (B/D) |
| RN7SL176P | 5.66 | 2.52 | 1.15 | 1.56 | 3.14 | 4.17E-06 | 9.19E-05 | (A) NE (B/D) |
| MIR-210/210 | 7.48 | 4.34 | 0.82 | 2.77 | 3.13 | 1.18E-06 | 3.39E-05 | (A) NE (B/D) |
| MIR-210/3P | 7.48 | 4.34 | 0.82 | 2.77 | 3.13 | 1.18E-06 | 3.39E-05 | (A) NE (B/D) |
| MIR-210/5P | 7.48 | 4.34 | 0.82 | 2.77 | 3.13 | 1.18E-06 | 3.39E-05 | (A) NE (B/D) |
| DMPK | 8.79 | 5.66 | 1.07 | 1.34 | 3.13 | 2.04E-06 | 5.22E-05 | (A) NE (B/D) |
| SDK1 | 4.76 | 1.63 | 1.1 | 1.37 | 3.13 | 2.76E-06 | 6.57E-05 | (A) NE (B/D) |
| SKINT1L | 3.64 | 0.5 | 2.75 | 0.6 | 3.13 | 9.01E-03 | 0.0331 | (A) NE (B/D) |
| FOXO6 | 5.72 | 2.6 | 1.02 | 1.12 | 3.12 | 1.76E-06 | 4.68E-05 | (A) NE (B/D) |
| SCGB1B2P | 5.33 | 2.21 | 1.5 | 1.36 | 3.12 | 1.15E-04 | 1.21E-03 | (A) NE (B/D) |
| RTBDN | 3.16 | 0.04 | 1.51 | 0.2 | 3.12 | 2.51E-04 | 2.18E-03 | (A) NE (B/D) |
| TAAR1 | 3.15 | 0.04 | 2.42 | 0.14 | 3.11 | 4.84E-03 | 0.021 | (A) NE (B/D) |
| MIR-10B/10B | 3.9 | 0.81 | 0.99 | 0.98 | 3.09 | 1.91E-06 | 4.97E-05 | (A) NE (B/D) |
| MIR-10B/10B | 3.9 | 0.81 | 0.99 | 0.98 | 3.09 | 1.91E-06 | 4.97E-05 | (A) NE (B/D) |
| MIR-10B/3P | 3.9 | 0.81 | 0.99 | 0.98 | 3.09 | 1.91E-06 | 4.97E-05 | (A) NE (B/D) |
| MIR-10B/5P | 3.9 | 0.81 | 0.99 | 0.98 | 3.09 | 1.91E-06 | 4.97E-05 | (A) NE (B/D) |
| SYT13 | 3.11 | 0.02 | 1.19 | 0.14 | 3.09 | 5.25E-05 | 6.57E-04 | (A) NE (B/D) |
| MIR-3911/39 | 6.4 | 3.32 | 0.74 | 1.4 | 3.08 | 2.69E-09 | 2.99E-07 | (A) NE (B/D) |
| CDC20P1 | 6.05 | 2.97 | 1.11 | 1.71 | 3.08 | 2.65E-06 | 6.36E-05 | (A) NE (B/D) |
| NKX2-2-AS1 | 4.07 | 0.99 | 1.52 | 1.49 | 3.08 | 1.37E-04 | 1.38E-03 | (A) NE (B/D) |
| NAV2 | 4.72 | 1.64 | 1.84 | 1.48 | 3.08 | 7.36E-04 | 5.02E-03 | (A) NE (B/D) |
| UBE2S | 3.85 | 0.78 | 0.75 | 0.9 | 3.07 | 3.41E-08 | 2.05E-06 | (A) NE (B/D) |
| PLXNA2 | 6.97 | 3.92 | 1.02 | 1.35 | 3.05 | 1.30E-06 | 3.67E-05 | (A) NE (B/D) |
| TMEM63C | 4.21 | 1.16 | 1 | 0.81 | 3.05 | 4.17E-06 | 9.18E-05 | (A) NE (B/D) |
| NUSAP1 | 10.22 | 7.17 | 1.49 | 2.14 | 3.05 | 1.08E-04 | 1.15E-03 | (A) NE (B/D) |
| PRSS46P | 4.28 | 1.23 | 1.72 | 1.48 | 3.05 | 4.63E-04 | 3.53E-03 | (A) NE (B/D) |
| CALCB | 4.2 | 1.15 | 1.95 | 1.17 | 3.05 | 1.29E-03 | 7.66E-03 | (A) NE (B/D) |
| FMN2 | 3.54 | 0.5 | 2.36 | 0.91 | 3.04 | 4.58E-03 | 0.0201 | (A) NE (B/D) |
| ELOVL2 | 4.6 | 1.57 | 2.1 | 1.8 | 3.03 | 2.14E-03 | 0.0113 | (A) NE (B/D) |
| FOXP4 | 7.45 | 4.43 | 0.62 | 1.19 | 3.02 | 1.21E-10 | 2.86E-08 | (A) NE (B/D) |
| UBE2SP1 | 10.02 | 7 | 0.7 | 1.4 | 3.02 | 1.25E-09 | 1.60E-07 | (A) NE (B/D) |
| TROAP | 4.55 | 1.53 | 0.76 | 1.54 | 3.02 | 6.94E-09 | 5.91E-07 | (A) NE (B/D) |
| MAP6 | 3.34 | 0.32 | 1.22 | 0.65 | 3.02 | 4.69E-05 | 6.01E-04 | (A) NE (B/D) |
| GPC6 | 5.83 | 2.83 | 0.94 | 1.79 | 3 | 3.38E-07 | 1.26E-05 | (A) NE (B/D) |
| MIR-4641/45 | 5.5 | 2.5 | 1.07 | 1.6 | 3 | 2.36E-06 | 5.85E-05 | (A) NE (B/D) |
| MIR-4641/46 | 5.5 | 2.5 | 1.07 | 1.6 | 3 | 2.36E-06 | 5.85E-05 | (A) NE (B/D) |
| MIR-4641/5P | 5.5 | 2.5 | 1.07 | 1.6 | 3 | 2.36E-06 | 5.85E-05 | (A) NE (B/D) |
| CAMK2N2 | 3.14 | 0.15 | 0.4 | 0.53 | 2.99 | 2.80E-12 | 1.47E-09 | (A) NE (B/D) |
| IL17RB | 4.69 | 1.7 | 0.84 | 1.52 | 2.99 | 5.33E-08 | 2.92E-06 | (A) NE (B/D) |
| RHEBL1 | 6.05 | 3.05 | 0.93 | 1.55 | 2.99 | 3.13E-07 | 1.19E-05 | (A) NE (B/D) |
| MEG8 | 3.62 | 0.63 | 2.74 | 0.86 | 2.99 | 0.0114 | 0.0391 | (A) NE (B/D) |
| HJURP | 5.38 | 2.4 | 0.83 | 1.65 | 2.98 | 5.21E-08 | 2.88E-06 | (A) NE (B/D) |
| HGFAC | 7.55 | 4.57 | 1.27 | 1.77 | 2.98 | 2.34E-05 | 3.45E-04 | (A) NE (B/D) |
| MIR9-3HG | 5.61 | 2.63 | 0.93 | 1.72 | 2.97 | 3.47E-07 | 1.28E-05 | (A) NE (B/D) |
| BEX2 | 10.14 | 7.17 | 0.95 | 1.82 | 2.97 | 4.99E-07 | 1.72E-05 | (A) NE (B/D) |

| | | | | | | | | |
|---|---|---|---|---|---|---|---|---|
| NNT-AS1 | 4.31 | 1.34 | 0.96 | 1.18 | 2.97 | 1.04E-06 | 3.07E-05 | (A) NE (B/D) |
| VWA7 | 4.64 | 1.68 | 0.59 | 0.82 | 2.96 | 7.07E-10 | 1.09E-07 | (A) NE (B/D) |
| PLP2 | 8.22 | 5.25 | 0.59 | 2.05 | 2.96 | 5.61E-09 | 5.10E-07 | (A) NE (B/D) |
| TRIM46 | 6.19 | 3.23 | 1.21 | 1.29 | 2.96 | 2.09E-05 | 3.17E-04 | (A) NE (B/D) |
| CFHR3 | 3.3 | 0.34 | 2.39 | 1.07 | 2.96 | 5.71E-03 | 0.0237 | (A) NE (B/D) |
| ABCC8 | 3.68 | 0.74 | 1.16 | 1.12 | 2.94 | 1.75E-05 | 2.79E-04 | (A) NE (B/D) |
| PLEKHG5 | 3.19 | 0.26 | 1.6 | 0.43 | 2.93 | 5.38E-04 | 3.95E-03 | (A) NE (B/D) |
| LNX1-AS2 | 3.63 | 0.7 | 2.12 | 1.05 | 2.93 | 2.92E-03 | 0.0143 | (A) NE (B/D) |
| BUB1 | 7.93 | 5.01 | 1.01 | 1.86 | 2.92 | 1.64E-06 | 4.40E-05 | (A) NE (B/D) |
| GPRC5C | 6.46 | 3.54 | 1.16 | 1.88 | 2.92 | 9.99E-06 | 1.81E-04 | (A) NE (B/D) |
| TSPEAR-AS1 | 4.32 | 1.4 | 1.87 | 1.1 | 2.92 | 1.34E-03 | 7.87E-03 | (A) NE (B/D) |
| ENO2 | 3.71 | 0.8 | 0.54 | 1.13 | 2.91 | 9.75E-12 | 3.98E-09 | (A) NE (B/D) |
| CDC20 | 4.22 | 1.31 | 0.97 | 1.39 | 2.91 | 1.01E-06 | 3.00E-05 | (A) NE (B/D) |
| KIF20A | 7 | 4.1 | 1.15 | 1.86 | 2.91 | 9.02E-06 | 1.67E-04 | (A) NE (B/D) |
| ICA1L | 4.74 | 1.83 | 1.38 | 1.09 | 2.91 | 1.16E-04 | 1.21E-03 | (A) NE (B/D) |
| DHPS | 5.46 | 2.56 | 0.71 | 1.29 | 2.9 | 5.16E-09 | 4.76E-07 | (A) NE (B/D) |
| SMOC1 | 3.27 | 0.36 | 1.52 | 1.01 | 2.9 | 2.98E-04 | 2.51E-03 | (A) NE (B/D) |
| HOXD13 | 4.09 | 1.19 | 1.87 | 1.46 | 2.9 | 1.29E-03 | 7.66E-03 | (A) NE (B/D) |
| RAB3A | 4.32 | 1.42 | 1.9 | 1.52 | 2.9 | 1.46E-03 | 8.46E-03 | (A) NE (B/D) |
| ABHD8 | 3.2 | 0.31 | 0.48 | 0.55 | 2.89 | 3.05E-10 | 5.77E-08 | (A) NE (B/D) |
| ADAMTS14 | 5.11 | 2.23 | 1.51 | 1.71 | 2.89 | 2.15E-04 | 1.94E-03 | (A) NE (B/D) |
| MIR-559/559 | 7.55 | 4.67 | 1.55 | 2.43 | 2.89 | 3.25E-04 | 2.68E-03 | (A) NE (B/D) |
| E2F7 | 5.33 | 2.44 | 1.83 | 1.59 | 2.89 | 1.10E-03 | 6.80E-03 | (A) NE (B/D) |
| TRI-TAT2-2 | 5.88 | 3 | 1.08 | 1.93 | 2.88 | 5.25E-06 | 1.10E-04 | (A) NE (B/D) |
| NEK2 | 6.37 | 3.49 | 1.15 | 2.08 | 2.88 | 1.30E-05 | 2.25E-04 | (A) NE (B/D) |
| RAB3B | 8.07 | 5.21 | 0.44 | 2.08 | 2.86 | 5.06E-09 | 4.68E-07 | (A) NE (B/D) |
| KLHL14 | 4.21 | 1.35 | 2.45 | 1.41 | 2.86 | 7.80E-03 | 0.0297 | (A) NE (B/D) |
| CKMT1A | 5.91 | 3.06 | 0.34 | 1.93 | 2.85 | 6.66E-10 | 1.05E-07 | (A) NE (B/D) |
| CDK18 | 3.69 | 0.84 | 0.99 | 0.99 | 2.85 | 4.52E-06 | 9.75E-05 | (A) NE (B/D) |
| IGFBP2 | 4.23 | 1.39 | 1.52 | 1.48 | 2.85 | 2.73E-04 | 2.34E-03 | (A) NE (B/D) |
| PRELID1P2 | 3.55 | 0.71 | 1.93 | 1.25 | 2.85 | 1.89E-03 | 0.0103 | (A) NE (B/D) |
| SMPD3 | 2.95 | 0.11 | 0.69 | 0.31 | 2.84 | 8.51E-07 | 2.61E-05 | (A) NE (B/D) |
| UBE2C | 3.3 | 0.46 | 0.93 | 0.78 | 2.84 | 3.83E-06 | 8.56E-05 | (A) NE (B/D) |
| LGSN | 4.42 | 1.59 | 2.73 | 1.57 | 2.84 | 0.0146 | 0.0469 | (A) NE (B/D) |
| SCAMP5 | 2.97 | 0.14 | 0.44 | 0.33 | 2.83 | 2.72E-09 | 2.99E-07 | (A) NE (B/D) |
| PCSK1N | 2.85 | 0.02 | 1.5 | 0.09 | 2.83 | 4.79E-04 | 3.62E-03 | (A) NE (B/D) |
| FAAH | 5.48 | 2.67 | 0.27 | 1.83 | 2.82 | 2.07E-10 | 4.16E-08 | (A) NE (B/D) |
| AMIGO2 | 3.35 | 0.52 | 1.45 | 0.66 | 2.82 | 3.06E-04 | 2.56E-03 | (A) NE (B/D) |
| SEMA6D | 3.81 | 0.99 | 1.99 | 1.01 | 2.82 | 2.54E-03 | 0.0129 | (A) NE (B/D) |
| DLX6-AS2 | 3.52 | 0.7 | 2.13 | 1.01 | 2.82 | 3.93E-03 | 0.0179 | (A) NE (B/D) |
| DLX6-AS1 | 3.74 | 0.91 | 2.25 | 1.1 | 2.82 | 5.31E-03 | 0.0225 | (A) NE (B/D) |
| CCDC151 | 3.15 | 0.34 | 0.7 | 0.51 | 2.81 | 3.56E-07 | 1.30E-05 | (A) NE (B/D) |
| MIR-496/496 | 3.08 | 0.27 | 2.55 | 0.92 | 2.81 | 0.0107 | 0.0374 | (A) NE (B/D) |
| HCN3 | 3.13 | 0.33 | 0.37 | 0.52 | 2.8 | 7.71E-13 | 6.84E-10 | (A) NE (B/D) |
| BAIAP3 | 3.51 | 0.71 | 0.86 | 0.88 | 2.8 | 9.87E-07 | 2.96E-05 | (A) NE (B/D) |
| USP27X | 7.59 | 4.79 | 1 | 1.5 | 2.8 | 2.53E-06 | 6.15E-05 | (A) NE (B/D) |
| CDC25C | 6.17 | 3.37 | 1.2 | 1.67 | 2.8 | 2.47E-05 | 3.60E-04 | (A) NE (B/D) |
| GCGR | 3.07 | 0.27 | 2.1 | 0.81 | 2.8 | 3.83E-03 | 0.0176 | (A) NE (B/D) |
| KIF23 | 6.2 | 3.41 | 1.44 | 1.53 | 2.79 | 1.95E-04 | 1.81E-03 | (A) NE (B/D) |
| NRGN | 7 | 4.21 | 1.63 | 2.57 | 2.79 | 7.24E-04 | 4.97E-03 | (A) NE (B/D) |
| TMEM198 | 4.3 | 1.52 | 0.34 | 0.93 | 2.78 | 1.51E-16 | 5.54E-13 | (A) NE (B/D) |
| UQCC2 | 6.15 | 3.37 | 0.28 | 1.23 | 2.78 | 5.10E-15 | 1.01E-11 | (A) NE (B/D) |
| OCIAD2 | 5.55 | 2.77 | 0.56 | 1.7 | 2.78 | 8.82E-10 | 1.27E-07 | (A) NE (B/D) |
| MAST2 | 7.43 | 4.64 | 0.71 | 1.13 | 2.78 | 1.29E-08 | 9.48E-07 | (A) NE (B/D) |
| TFAP2A-AS1 | 6.47 | 3.69 | 1.36 | 1.82 | 2.78 | 1.12E-04 | 1.18E-03 | (A) NE (B/D) |
| GNG4 | 3.25 | 0.48 | 0.25 | 0.79 | 2.77 | 4.74E-20 | 6.10E-16 | (A) NE (B/D) |
| PPFIA3 | 2.95 | 0.18 | 0.62 | 0.4 | 2.77 | 1.89E-07 | 8.04E-06 | (A) NE (B/D) |
| PAX5 | 5.73 | 2.95 | 1.03 | 1.98 | 2.77 | 5.12E-06 | 1.08E-04 | (A) NE (B/D) |
| GAD1 | 2.77 | 0 | 1.12 | 0 | 2.77 | 7.49E-05 | 8.68E-04 | (A) NE (B/D) |
| GADD45A | 6.13 | 3.36 | 1.66 | 1.36 | 2.77 | 7.61E-04 | 5.16E-03 | (A) NE (B/D) |
| MAPK13 | 4.45 | 1.69 | 0.26 | 1.1 | 2.76 | 1.95E-16 | 6.27E-13 | (A) NE (B/D) |
| NR1D1 | 3.89 | 1.12 | 0.81 | 1.18 | 2.76 | 1.72E-07 | 7.47E-06 | (A) NE (B/D) |
| CHRNB2 | 3.19 | 0.43 | 0.89 | 1.24 | 2.76 | 6.99E-07 | 2.23E-05 | (A) NE (B/D) |
| CLDN9 | 5.11 | 2.35 | 0.99 | 1.81 | 2.76 | 2.73E-06 | 6.53E-05 | (A) NE (B/D) |
| TNFAIP8L1 | 7.29 | 4.53 | 1.16 | 0.9 | 2.76 | 4.35E-05 | 5.66E-04 | (A) NE (B/D) |

| | | | | | | | | |
|---|---|---|---|---|---|---|---|---|
| ZNF853 | 5.02 | 2.26 | 1.42 | 1.27 | 2.76 | 2.11E-04 | 1.91E-03 | (A) NE (B/D) |
| GALNT13 | 3.15 | 0.39 | 1.9 | 0.95 | 2.76 | 2.21E-03 | 0.0115 | (A) NE (B/D) |
| HOXA-AS3 | 5.83 | 3.08 | 2.24 | 1.36 | 2.76 | 5.86E-03 | 0.0242 | (A) NE (B/D) |
| MAPK8IP2 | 7.52 | 4.77 | 0.38 | 1.86 | 2.75 | 7.64E-10 | 1.14E-07 | (A) NE (B/D) |
| KLRC2 | 4.51 | 1.77 | 2.68 | 1.59 | 2.75 | 0.0154 | 0.049 | (A) NE (B/D) |
| TUBA1A | 4.1 | 1.36 | 0.57 | 1.32 | 2.74 | 1.94E-10 | 4.08E-08 | (A) NE (B/D) |
| SAPCD2 | 5.29 | 2.55 | 0.83 | 1.15 | 2.74 | 2.93E-07 | 1.13E-05 | (A) NE (B/D) |
| SPC24 | 6.81 | 4.07 | 1.19 | 1.34 | 2.74 | 3.52E-05 | 4.81E-04 | (A) NE (B/D) |
| QPCT | 6.29 | 3.55 | 2.43 | 1.86 | 2.74 | 9.63E-03 | 0.0347 | (A) NE (B/D) |
| STX1A | 2.74 | 0.01 | 0.31 | 0.06 | 2.73 | 3.66E-09 | 3.68E-07 | (A) NE (B/D) |
| SYS1-DBNDD | 7.54 | 4.81 | 0.78 | 1.3 | 2.73 | 8.43E-08 | 4.19E-06 | (A) NE (B/D) |
| TYMSOS | 4.4 | 1.68 | 0.94 | 1.47 | 2.73 | 1.48E-06 | 4.08E-05 | (A) NE (B/D) |
| TYRP1 | 3.34 | 0.61 | 1.92 | 1.04 | 2.73 | 2.46E-03 | 0.0125 | (A) NE (B/D) |
| ARL5AP3 | 4.67 | 1.94 | 2.65 | 1.67 | 2.73 | 0.0153 | 0.0487 | (A) NE (B/D) |
| SSBP4 | 3.85 | 1.12 | 0.51 | 0.89 | 2.72 | 2.64E-11 | 8.34E-09 | (A) NE (B/D) |
| GGT7 | 5.89 | 3.17 | 1.13 | 1.4 | 2.72 | 1.98E-05 | 3.06E-04 | (A) NE (B/D) |
| ZFHX2 | 5.88 | 3.15 | 1.1 | 0.8 | 2.72 | 3.31E-05 | 4.57E-04 | (A) NE (B/D) |
| INPPL1 | 4.92 | 2.19 | 1.26 | 1.2 | 2.72 | 7.97E-05 | 9.08E-04 | (A) NE (B/D) |
| WFDC2 | 2.8 | 0.08 | 1.21 | 0.3 | 2.72 | 1.35E-04 | 1.37E-03 | (A) NE (B/D) |
| UTS2 | 7.57 | 4.85 | 1.98 | 1.95 | 2.72 | 2.97E-03 | 0.0145 | (A) NE (B/D) |
| CPT1C | 2.89 | 0.18 | 2.04 | 0.42 | 2.72 | 3.88E-03 | 0.0177 | (A) NE (B/D) |
| CAMK1D | 7.51 | 4.78 | 2.17 | 0.98 | 2.72 | 5.27E-03 | 0.0223 | (A) NE (B/D) |
| GADD45G | 4.13 | 1.42 | 0.88 | 1.43 | 2.71 | 5.63E-07 | 1.90E-05 | (A) NE (B/D) |
| PTTG3P | 4.68 | 1.97 | 0.94 | 1.13 | 2.71 | 2.54E-06 | 6.17E-05 | (A) NE (B/D) |
| ATP5MC1P4 | 4.21 | 1.5 | 1.08 | 1.09 | 2.71 | 1.65E-05 | 2.67E-04 | (A) NE (B/D) |
| ZNF165 | 6.75 | 4.04 | 1.09 | 0.93 | 2.71 | 2.65E-05 | 3.82E-04 | (A) NE (B/D) |
| TUBB7P | 3.91 | 1.22 | 0.46 | 1 | 2.7 | 1.09E-12 | 8.47E-10 | (A) NE (B/D) |
| DDX39A | 4.37 | 1.67 | 0.38 | 1.49 | 2.7 | 7.66E-12 | 3.34E-09 | (A) NE (B/D) |
| HYI | 6.75 | 4.04 | 0.65 | 1.46 | 2.7 | 4.15E-09 | 4.03E-07 | (A) NE (B/D) |
| GNA14 | 6.55 | 3.85 | 0.77 | 1.48 | 2.7 | 6.65E-08 | 3.47E-06 | (A) NE (B/D) |
| TUBB2A | 3.71 | 1.01 | 1.37 | 1.16 | 2.7 | 1.99E-04 | 1.83E-03 | (A) NE (B/D) |
| KCNMB2 | 2.87 | 0.17 | 1.99 | 0.69 | 2.7 | 3.40E-03 | 0.0161 | (A) NE (B/D) |
| FKBP9P1 | 3.4 | 0.7 | 2.12 | 1.04 | 2.7 | 4.86E-03 | 0.021 | (A) NE (B/D) |
| TUBB3 | 6.43 | 3.74 | 1.1 | 1.49 | 2.69 | 1.44E-05 | 2.43E-04 | (A) NE (B/D) |
| IGHVII-40-1 | 2.7 | 0.01 | 1.81 | 0.05 | 2.69 | 2.08E-03 | 0.011 | (A) NE (B/D) |
| BPIFA1 | 2.7 | 0.01 | 2.01 | 0.08 | 2.69 | 3.92E-03 | 0.0179 | (A) NE (B/D) |
| MCF2L-AS1 | 5.26 | 2.58 | 0.68 | 1.35 | 2.68 | 8.66E-09 | 7.05E-07 | (A) NE (B/D) |
| BEND5 | 4.03 | 1.35 | 0.94 | 1.05 | 2.68 | 3.77E-06 | 8.47E-05 | (A) NE (B/D) |
| CC2D1A | 3.13 | 0.45 | 0.94 | 0.61 | 2.68 | 1.11E-05 | 1.98E-04 | (A) NE (B/D) |
| SLC36A4 | 2.74 | 0.06 | 2.1 | 0.24 | 2.68 | 5.00E-03 | 0.0215 | (A) NE (B/D) |
| RAB11B | 7.98 | 5.31 | 0.44 | 1.3 | 2.67 | 3.44E-12 | 1.73E-09 | (A) NE (B/D) |
| HOOK2 | 3.44 | 0.77 | 0.52 | 0.73 | 2.67 | 3.32E-10 | 6.11E-08 | (A) NE (B/D) |
| TACC3 | 4.76 | 2.09 | 0.66 | 1.73 | 2.67 | 1.86E-08 | 1.26E-06 | (A) NE (B/D) |
| TCEAL6 | 6.06 | 3.38 | 1.08 | 2.96 | 2.67 | 1.21E-04 | 1.25E-03 | (A) NE (B/D) |
| SLC7A11-AS1 | 5.36 | 2.69 | 1.59 | 1.81 | 2.67 | 6.83E-04 | 4.76E-03 | (A) NE (B/D) |
| EPB41L4B | 5.67 | 3.01 | 0.98 | 1.35 | 2.66 | 4.06E-06 | 8.99E-05 | (A) NE (B/D) |
| CENPA | 3.64 | 0.98 | 0.98 | 1.27 | 2.66 | 4.36E-06 | 9.50E-05 | (A) NE (B/D) |
| FREM1 | 4.76 | 2.1 | 1.72 | 1.75 | 2.66 | 1.25E-03 | 7.46E-03 | (A) NE (B/D) |
| NEURL1B | 3.38 | 0.73 | 1.01 | 1.1 | 2.65 | 9.57E-06 | 1.76E-04 | (A) NE (B/D) |
| MTMR9LP | 7.2 | 4.55 | 1.26 | 1.55 | 2.65 | 7.86E-05 | 8.98E-04 | (A) NE (B/D) |
| UBE2QL1 | 3.38 | 0.73 | 2.05 | 1.08 | 2.65 | 4.47E-03 | 0.0198 | (A) NE (B/D) |
| MICAL1 | 4.85 | 2.21 | 0.84 | 1.54 | 2.64 | 4.10E-07 | 1.47E-05 | (A) NE (B/D) |
| ROR2 | 5.49 | 2.86 | 1.82 | 1.77 | 2.64 | 2.07E-03 | 0.011 | (A) NE (B/D) |
| GAPDHP72 | 6.39 | 3.76 | 1.96 | 1.51 | 2.64 | 3.40E-03 | 0.0161 | (A) NE (B/D) |
| DCAF15 | 3.61 | 0.98 | 0.37 | 1.04 | 2.63 | 2.14E-14 | 3.06E-11 | (A) NE (B/D) |
| FAM83E | 3.36 | 0.73 | 1.12 | 1.13 | 2.63 | 3.49E-05 | 4.77E-04 | (A) NE (B/D) |
| DPYSL3 | 2.99 | 0.35 | 1.42 | 0.61 | 2.63 | 4.42E-04 | 3.41E-03 | (A) NE (B/D) |
| PPM1N | 6.01 | 3.39 | 0.27 | 1.15 | 2.62 | 5.12E-15 | 1.01E-11 | (A) NE (B/D) |
| MT1DP | 4.01 | 1.39 | 1.4 | 1.56 | 2.62 | 2.67E-04 | 2.29E-03 | (A) NE (B/D) |
| SIGLEC15 | 5.15 | 2.53 | 1.86 | 1.25 | 2.62 | 2.57E-03 | 0.013 | (A) NE (B/D) |
| EZH2P1 | 6.12 | 3.5 | 1.99 | 2.11 | 2.62 | 4.15E-03 | 0.0187 | (A) NE (B/D) |
| KLHL41 | 2.79 | 0.17 | 2.16 | 0.57 | 2.62 | 6.51E-03 | 0.0261 | (A) NE (B/D) |
| VAV2 | 6.13 | 3.51 | 0.55 | 1.07 | 2.61 | 1.74E-10 | 3.83E-08 | (A) NE (B/D) |
| NPM2 | 5.25 | 2.64 | 0.84 | 1.4 | 2.61 | 5.02E-07 | 1.73E-05 | (A) NE (B/D) |
| APOBEC3B-A | 5.93 | 3.33 | 1.19 | 1.88 | 2.61 | 5.39E-05 | 6.70E-04 | (A) NE (B/D) |

| | | | | | | | | |
|---|---|---|---|---|---|---|---|---|
| CEL | 5.3 | 2.69 | 1.57 | 1.14 | 2.61 | 8.25E-04 | 5.49E-03 | (A) NE (B/D) |
| TCL6 | 4.87 | 2.25 | 2.02 | 2.11 | 2.61 | 4.59E-03 | 0.0202 | (A) NE (B/D) |
| PLEKHJ1 | 3.1 | 0.5 | 0.36 | 0.74 | 2.6 | 1.46E-14 | 2.50E-11 | (A) NE (B/D) |
| ZDHHC11B | 8.05 | 5.45 | 1.08 | 1.69 | 2.6 | 1.83E-05 | 2.88E-04 | (A) NE (B/D) |
| RNU7-31P | 5.8 | 3.2 | 2.34 | 1.97 | 2.6 | 0.0107 | 0.0374 | (A) NE (B/D) |
| RTL8A | 7.06 | 4.47 | 0.31 | 1.77 | 2.59 | 7.41E-10 | 1.13E-07 | (A) NE (B/D) |
| FAM183A | 3.34 | 0.75 | 1.45 | 0.93 | 2.59 | 5.03E-04 | 3.76E-03 | (A) NE (B/D) |
| KCNH2 | 2.76 | 0.18 | 0.4 | 0.69 | 2.58 | 8.46E-13 | 7.26E-10 | (A) NE (B/D) |
| SMOC2 | 5.65 | 3.07 | 0.42 | 1.61 | 2.58 | 2.44E-10 | 4.76E-08 | (A) NE (B/D) |
| ASNS | 4.38 | 1.8 | 0.76 | 1.41 | 2.58 | 1.29E-07 | 5.96E-06 | (A) NE (B/D) |
| LYPD1 | 3.85 | 1.27 | 0.78 | 1.21 | 2.58 | 2.10E-07 | 8.66E-06 | (A) NE (B/D) |
| SNORA56 | 8.93 | 6.36 | 0.82 | 1.57 | 2.57 | 4.45E-07 | 1.58E-05 | (A) NE (B/D) |
| TIMELESS | 4.55 | 1.99 | 0.46 | 1.54 | 2.56 | 2.15E-10 | 4.29E-08 | (A) NE (B/D) |
| SERGEF | 9 | 6.43 | 0.62 | 1.09 | 2.56 | 4.26E-09 | 4.10E-07 | (A) NE (B/D) |
| SPIRE2 | 5.76 | 3.2 | 0.64 | 1.13 | 2.56 | 6.84E-09 | 5.87E-07 | (A) NE (B/D) |
| GUSB | 3.96 | 1.4 | 0.69 | 1.32 | 2.56 | 2.41E-08 | 1.52E-06 | (A) NE (B/D) |
| SLC1A5 | 9.88 | 7.32 | 0.77 | 1.38 | 2.56 | 1.63E-07 | 7.16E-06 | (A) NE (B/D) |
| KIF4A | 6.27 | 3.7 | 0.93 | 1.97 | 2.56 | 5.48E-06 | 1.13E-04 | (A) NE (B/D) |
| CDKN2B-AS1 | 3.27 | 0.71 | 1.25 | 0.89 | 2.56 | 1.63E-04 | 1.57E-03 | (A) NE (B/D) |
| ZNF414 | 4.47 | 1.92 | 0.91 | 1.24 | 2.55 | 2.88E-06 | 6.79E-05 | (A) NE (B/D) |
| ALG1L3P | 2.57 | 0.02 | 1.16 | 0.08 | 2.55 | 1.70E-04 | 1.63E-03 | (A) NE (B/D) |
| MAGEA3 | 2.7 | 0.15 | 1.56 | 0.44 | 2.55 | 1.11E-03 | 6.86E-03 | (A) NE (B/D) |
| RPS6KL1 | 5.06 | 2.51 | 1.76 | 1.37 | 2.55 | 2.02E-03 | 0.0108 | (A) NE (B/D) |
| HSD17B3 | 4.35 | 1.8 | 2.01 | 1.17 | 2.55 | 4.97E-03 | 0.0214 | (A) NE (B/D) |
| GEMIN8P4 | 5.53 | 3 | 0.42 | 1.12 | 2.54 | 1.51E-12 | 9.39E-10 | (A) NE (B/D) |
| TMEM240 | 4.12 | 1.58 | 1.26 | 0.96 | 2.54 | 1.79E-04 | 1.69E-03 | (A) NE (B/D) |
| SOX5P1 | 3.16 | 0.62 | 2.04 | 1.09 | 2.54 | 5.52E-03 | 0.0231 | (A) NE (B/D) |
| NXF2B | 2.74 | 0.2 | 2.04 | 0.66 | 2.54 | 5.53E-03 | 0.0232 | (A) NE (B/D) |
| RPS4XP11 | 7.67 | 5.14 | 0.93 | 2.47 | 2.53 | 2.65E-05 | 3.82E-04 | (A) NE (B/D) |
| DOCK9-DT | 5.03 | 2.5 | 1.27 | 1.48 | 2.53 | 1.39E-04 | 1.39E-03 | (A) NE (B/D) |
| SLC38A11 | 2.73 | 0.19 | 1.6 | 0.68 | 2.53 | 1.31E-03 | 7.76E-03 | (A) NE (B/D) |
| GPR173 | 3 | 0.47 | 1.66 | 0.89 | 2.53 | 1.64E-03 | 9.21E-03 | (A) NE (B/D) |
| MARK2P9 | 3.37 | 0.84 | 1.69 | 0.66 | 2.53 | 1.88E-03 | 0.0102 | (A) NE (B/D) |
| WDR34 | 8.69 | 6.17 | 0.53 | 1.54 | 2.52 | 1.47E-09 | 1.83E-07 | (A) NE (B/D) |
| SPAG4 | 6.73 | 4.21 | 0.57 | 1.48 | 2.52 | 2.41E-09 | 2.75E-07 | (A) NE (B/D) |
| FAM24B | 3.93 | 1.42 | 0.76 | 1.16 | 2.52 | 2.26E-07 | 9.21E-06 | (A) NE (B/D) |
| AURKB | 3.76 | 1.24 | 0.8 | 1.28 | 2.52 | 4.34E-07 | 1.54E-05 | (A) NE (B/D) |
| CCDC24 | 6.31 | 3.79 | 0.8 | 1.19 | 2.52 | 4.94E-07 | 1.71E-05 | (A) NE (B/D) |
| PCDHA12 | 6.58 | 4.06 | 0.85 | 1.99 | 2.52 | 2.90E-06 | 6.82E-05 | (A) NE (B/D) |
| CA8 | 2.53 | 0.01 | 1.72 | 0.04 | 2.52 | 2.30E-03 | 0.0119 | (A) NE (B/D) |
| RPS28 | 5.12 | 2.6 | 1.82 | 1.65 | 2.52 | 2.83E-03 | 0.0139 | (A) NE (B/D) |
| DACH2 | 2.9 | 0.38 | 1.82 | 0.89 | 2.52 | 3.03E-03 | 0.0147 | (A) NE (B/D) |
| C7ORF13 | 4.58 | 2.07 | 0.87 | 0.92 | 2.51 | 3.70E-06 | 8.35E-05 | (A) NE (B/D) |
| CFAP57 | 5.71 | 3.2 | 1.1 | 0.89 | 2.51 | 5.88E-05 | 7.18E-04 | (A) NE (B/D) |
| PRC1 | 3.04 | 0.53 | 1.14 | 0.84 | 2.51 | 8.72E-05 | 9.78E-04 | (A) NE (B/D) |
| LINC01023 | 4.16 | 1.65 | 1.32 | 1.26 | 2.51 | 2.35E-04 | 2.08E-03 | (A) NE (B/D) |
| TMEM59L | 4.79 | 2.28 | 1.72 | 2 | 2.51 | 2.07E-03 | 0.011 | (A) NE (B/D) |
| HMGB2 | 4.83 | 2.32 | 1.01 | 1.85 | 2.5 | 1.47E-05 | 2.47E-04 | (A) NE (B/D) |
| KLC3 | 6.54 | 4.04 | 1.58 | 0.92 | 2.5 | 1.20E-03 | 7.24E-03 | (A) NE (B/D) |
| RIC3 | 4.56 | 2.06 | 1.77 | 1.9 | 2.5 | 2.58E-03 | 0.013 | (A) NE (B/D) |
| TUBB8P8 | 3.31 | 0.82 | 0.48 | 0.82 | 2.49 | 5.66E-11 | 1.47E-08 | (A) NE (B/D) |
| MANEAL | 2.88 | 0.39 | 0.51 | 0.76 | 2.49 | 7.02E-10 | 1.09E-07 | (A) NE (B/D) |
| PCDHA7 | 6.94 | 4.45 | 0.8 | 1.95 | 2.49 | 1.63E-06 | 4.40E-05 | (A) NE (B/D) |
| LGI2 | 3.22 | 0.73 | 1.14 | 0.77 | 2.49 | 9.85E-05 | 1.07E-03 | (A) NE (B/D) |
| MYLK2 | 3.11 | 0.62 | 1.55 | 0.8 | 2.49 | 1.15E-03 | 7.01E-03 | (A) NE (B/D) |
| FGF17 | 2.9 | 0.42 | 1.7 | 0.69 | 2.49 | 2.15E-03 | 0.0113 | (A) NE (B/D) |
| SNORD114-1 | 2.61 | 0.13 | 2.01 | 0.38 | 2.49 | 5.84E-03 | 0.0241 | (A) NE (B/D) |
| MAP2K7 | 7.64 | 5.16 | 0.6 | 1.03 | 2.48 | 5.19E-09 | 4.77E-07 | (A) NE (B/D) |
| FZD8 | 6.77 | 4.29 | 1.13 | 1.52 | 2.48 | 5.40E-05 | 6.72E-04 | (A) NE (B/D) |
| CSAG1 | 2.51 | 0.02 | 1.43 | 0.13 | 2.48 | 7.99E-04 | 5.37E-03 | (A) NE (B/D) |
| MIR-135B/13 | 2.73 | 0.26 | 1.64 | 0.71 | 2.48 | 1.75E-03 | 9.67E-03 | (A) NE (B/D) |
| MIR-135B/13 | 2.73 | 0.26 | 1.64 | 0.71 | 2.48 | 1.75E-03 | 9.67E-03 | (A) NE (B/D) |
| MIR-135B/3P | 2.73 | 0.26 | 1.64 | 0.71 | 2.48 | 1.75E-03 | 9.67E-03 | (A) NE (B/D) |
| MIR-135B/5P | 2.73 | 0.26 | 1.64 | 0.71 | 2.48 | 1.75E-03 | 9.67E-03 | (A) NE (B/D) |
| MIR-1909/19 | 4.85 | 2.39 | 0.31 | 1.3 | 2.47 | 1.36E-12 | 9.18E-10 | (A) NE (B/D) |

| | | | | | | | | |
|---|---|---|---|---|---|---|---|---|
| MIR-1909/19 | 4.85 | 2.39 | 0.31 | 1.3 | 2.47 | 1.36E-12 | 9.18E-10 | (A) NE (B/D) |
| MIR-1909/3P | 4.85 | 2.39 | 0.31 | 1.3 | 2.47 | 1.36E-12 | 9.18E-10 | (A) NE (B/D) |
| MIR-1909/5P | 4.85 | 2.39 | 0.31 | 1.3 | 2.47 | 1.36E-12 | 9.18E-10 | (A) NE (B/D) |
| SNORD41 | 3.45 | 0.98 | 0.75 | 1.11 | 2.47 | 2.41E-07 | 9.61E-06 | (A) NE (B/D) |
| SOBP | 2.72 | 0.25 | 1.3 | 0.65 | 2.47 | 3.62E-04 | 2.92E-03 | (A) NE (B/D) |
| SPRY4 | 2.69 | 0.22 | 1.44 | 0.69 | 2.47 | 7.42E-04 | 5.06E-03 | (A) NE (B/D) |
| IGHV7-40 | 3.56 | 1.09 | 2.38 | 2.09 | 2.47 | 0.0156 | 0.0495 | (A) NE (B/D) |
| MELTF-AS1 | 7.02 | 4.56 | 0.46 | 1.32 | 2.46 | 7.10E-11 | 1.79E-08 | (A) NE (B/D) |
| AKR7A3 | 5.47 | 3.01 | 1.71 | 1.18 | 2.46 | 2.16E-03 | 0.0113 | (A) NE (B/D) |
| ZNF711 | 2.87 | 0.41 | 1.85 | 0.89 | 2.46 | 3.81E-03 | 0.0175 | (A) NE (B/D) |
| INPP5E | 6.16 | 3.71 | 0.65 | 1.12 | 2.45 | 1.97E-08 | 1.31E-06 | (A) NE (B/D) |
| TM4SF1 | 3.84 | 1.39 | 0.66 | 1.48 | 2.45 | 3.81E-08 | 2.23E-06 | (A) NE (B/D) |
| CCDC74B | 5.72 | 3.27 | 0.68 | 2.25 | 2.45 | 2.64E-06 | 6.36E-05 | (A) NE (B/D) |
| CENPF | 5.75 | 3.3 | 1.08 | 1.74 | 2.45 | 3.88E-05 | 5.18E-04 | (A) NE (B/D) |
| CALB2 | 3.25 | 0.8 | 1.69 | 1.31 | 2.45 | 2.11E-03 | 0.0112 | (A) NE (B/D) |
| GPC4 | 4.84 | 2.39 | 1.88 | 1.57 | 2.45 | 4.12E-03 | 0.0186 | (A) NE (B/D) |
| HOXD9 | 2.77 | 0.31 | 2.21 | 0.89 | 2.45 | 0.0103 | 0.0365 | (A) NE (B/D) |
| HSD11B1L | 5.24 | 2.8 | 0.51 | 1.14 | 2.44 | 1.47E-10 | 3.33E-08 | (A) NE (B/D) |
| CKMT1B | 4.2 | 1.76 | 0.46 | 1.53 | 2.44 | 7.55E-10 | 1.14E-07 | (A) NE (B/D) |
| PCDHA3 | 6.27 | 3.83 | 0.84 | 1.92 | 2.44 | 3.46E-06 | 7.91E-05 | (A) NE (B/D) |
| ARHGAP33 | 3.48 | 1.04 | 0.92 | 1.04 | 2.44 | 7.63E-06 | 1.47E-04 | (A) NE (B/D) |
| SYCE3 | 3.43 | 0.99 | 1.46 | 1.2 | 2.44 | 7.49E-04 | 5.10E-03 | (A) NE (B/D) |
| CPE | 8.05 | 5.62 | 1.48 | 1.92 | 2.44 | 8.25E-04 | 5.49E-03 | (A) NE (B/D) |
| CEP126 | 2.58 | 0.15 | 1.73 | 0.29 | 2.44 | 2.84E-03 | 0.014 | (A) NE (B/D) |
| LSM7 | 6.63 | 4.2 | 0.33 | 1.28 | 2.43 | 2.10E-12 | 1.23E-09 | (A) NE (B/D) |
| CACFD1 | 4.44 | 2.01 | 0.73 | 1.52 | 2.43 | 2.18E-07 | 8.96E-06 | (A) NE (B/D) |
| POLR2I | 3.87 | 1.44 | 0.85 | 1.36 | 2.43 | 1.92E-06 | 4.97E-05 | (A) NE (B/D) |
| PCDHA2 | 5.77 | 3.34 | 0.81 | 1.83 | 2.43 | 2.03E-06 | 5.19E-05 | (A) NE (B/D) |
| MUTYH | 6.79 | 4.37 | 0.93 | 1.14 | 2.43 | 8.44E-06 | 1.60E-04 | (A) NE (B/D) |
| HSPB9 | 2.92 | 0.48 | 0.85 | 0.58 | 2.43 | 9.92E-06 | 1.81E-04 | (A) NE (B/D) |
| MFSD2A | 3.32 | 0.88 | 1.45 | 0.95 | 2.43 | 7.92E-04 | 5.33E-03 | (A) NE (B/D) |
| DCDC2 | 5.29 | 2.87 | 2 | 1.89 | 2.43 | 6.61E-03 | 0.0264 | (A) NE (B/D) |
| RPS4XP3 | 7.77 | 5.34 | 2.08 | 2.35 | 2.43 | 9.00E-03 | 0.0331 | (A) NE (B/D) |
| OR7E111P | 3.03 | 0.59 | 2.28 | 1.34 | 2.43 | 0.0128 | 0.0426 | (A) NE (B/D) |
| ZNF358 | 7.19 | 4.78 | 0.62 | 1.18 | 2.42 | 1.17E-08 | 8.90E-07 | (A) NE (B/D) |
| CENPW | 4.05 | 1.63 | 0.89 | 1.45 | 2.42 | 3.74E-06 | 8.43E-05 | (A) NE (B/D) |
| NUF2 | 3.09 | 0.67 | 1.87 | 1.11 | 2.42 | 4.33E-03 | 0.0193 | (A) NE (B/D) |
| OBSL1 | 3.19 | 0.78 | 0.46 | 1.07 | 2.41 | 1.97E-11 | 6.68E-09 | (A) NE (B/D) |
| SOX13 | 5.84 | 3.43 | 0.46 | 1.28 | 2.41 | 7.41E-11 | 1.85E-08 | (A) NE (B/D) |
| RIPK4 | 2.71 | 0.3 | 0.47 | 0.57 | 2.41 | 1.78E-09 | 2.14E-07 | (A) NE (B/D) |
| LINC00265 | 5.63 | 3.22 | 0.62 | 1.24 | 2.41 | 1.06E-08 | 8.34E-07 | (A) NE (B/D) |
| CAMK2N1 | 7.56 | 5.15 | 0.5 | 2.17 | 2.41 | 5.51E-07 | 1.87E-05 | (A) NE (B/D) |
| TPSP2 | 2.61 | 0.2 | 1.01 | 0.7 | 2.41 | 5.00E-05 | 6.31E-04 | (A) NE (B/D) |
| XCL1 | 2.45 | 0.04 | 1.16 | 0.24 | 2.41 | 2.35E-04 | 2.08E-03 | (A) NE (B/D) |
| GPR153 | 2.71 | 0.3 | 1.38 | 0.96 | 2.41 | 5.78E-04 | 4.19E-03 | (A) NE (B/D) |
| C6ORF136 | 2.98 | 0.59 | 0.27 | 0.83 | 2.4 | 4.12E-17 | 1.77E-13 | (A) NE (B/D) |
| PTPN3 | 4.49 | 2.09 | 0.34 | 1.04 | 2.4 | 5.87E-14 | 6.86E-11 | (A) NE (B/D) |
| C11ORF52 | 6.64 | 4.24 | 0.47 | 1.94 | 2.4 | 7.93E-08 | 3.99E-06 | (A) NE (B/D) |
| SNAP25 | 2.64 | 0.24 | 0.79 | 0.56 | 2.4 | 5.46E-06 | 1.13E-04 | (A) NE (B/D) |
| PVRIG2P | 5.91 | 3.51 | 1.18 | 1.06 | 2.4 | 1.39E-04 | 1.40E-03 | (A) NE (B/D) |
| SOX11 | 3.12 | 0.72 | 2.25 | 1.27 | 2.4 | 0.0126 | 0.0422 | (A) NE (B/D) |
| APP | 7.74 | 5.35 | 0.51 | 1.12 | 2.39 | 2.02E-10 | 4.12E-08 | (A) NE (B/D) |
| BSPRY | 5.36 | 2.96 | 0.58 | 1.28 | 2.39 | 4.55E-09 | 4.32E-07 | (A) NE (B/D) |
| TMEM125 | 9.27 | 6.88 | 0.75 | 1.97 | 2.39 | 1.92E-06 | 4.97E-05 | (A) NE (B/D) |
| NEBL | 5.55 | 3.16 | 1.1 | 1.55 | 2.39 | 5.53E-05 | 6.85E-04 | (A) NE (B/D) |
| DYNC1I1 | 2.67 | 0.29 | 1.23 | 0.5 | 2.39 | 3.40E-04 | 2.77E-03 | (A) NE (B/D) |
| RPL9P18 | 4.08 | 1.69 | 1.9 | 1.8 | 2.39 | 5.28E-03 | 0.0224 | (A) NE (B/D) |
| QTRT1 | 4.08 | 1.7 | 0.44 | 1.52 | 2.38 | 8.65E-10 | 1.25E-07 | (A) NE (B/D) |
| BAMBI | 6.44 | 4.06 | 0.76 | 1.57 | 2.38 | 5.80E-07 | 1.94E-05 | (A) NE (B/D) |
| PCDHA5 | 5.77 | 3.39 | 0.86 | 1.87 | 2.38 | 5.55E-06 | 1.14E-04 | (A) NE (B/D) |
| PCDHA9 | 10.16 | 7.78 | 0.94 | 2.18 | 2.38 | 3.01E-05 | 4.25E-04 | (A) NE (B/D) |
| SPOCK1 | 7.01 | 4.63 | 1.03 | 2.69 | 2.38 | 2.04E-04 | 1.86E-03 | (A) NE (B/D) |
| PROX1-AS1 | 2.64 | 0.26 | 1.2 | 0.65 | 2.38 | 2.47E-04 | 2.16E-03 | (A) NE (B/D) |
| TESC | 4.95 | 2.57 | 1.89 | 1.75 | 2.38 | 5.11E-03 | 0.0219 | (A) NE (B/D) |
| RCCD1 | 7.68 | 5.31 | 0.56 | 0.88 | 2.37 | 3.53E-09 | 3.60E-07 | (A) NE (B/D) |

| | | | | | | | | |
|---|---|---|---|---|---|---|---|---|
| EPCAM | 8.08 | 5.7 | 0.55 | 1.87 | 2.37 | 1.20E-07 | 5.56E-06 | (A) NE (B/D) |
| ADPRHL1 | 6.03 | 3.66 | 0.95 | 1.55 | 2.37 | 1.16E-05 | 2.06E-04 | (A) NE (B/D) |
| HMGB3P6 | 9.85 | 7.48 | 1.16 | 1.98 | 2.37 | 1.43E-04 | 1.43E-03 | (A) NE (B/D) |
| TOP2A | 4.91 | 2.54 | 1.22 | 2.06 | 2.37 | 2.33E-04 | 2.07E-03 | (A) NE (B/D) |
| MIR-3074/30 | 3.86 | 1.49 | 1.29 | 1.78 | 2.37 | 3.22E-04 | 2.66E-03 | (A) NE (B/D) |
| MIR-3074/3P | 3.86 | 1.49 | 1.29 | 1.78 | 2.37 | 3.22E-04 | 2.66E-03 | (A) NE (B/D) |
| MIR-3074/5P | 3.86 | 1.49 | 1.29 | 1.78 | 2.37 | 3.22E-04 | 2.66E-03 | (A) NE (B/D) |
| FANCL | 3.92 | 1.54 | 1.59 | 1.23 | 2.37 | 1.72E-03 | 9.58E-03 | (A) NE (B/D) |
| SPPL2B | 6.86 | 4.49 | 0.45 | 0.92 | 2.36 | 1.59E-11 | 5.69E-09 | (A) NE (B/D) |
| MYO7A | 5.05 | 2.69 | 0.66 | 1.04 | 2.36 | 5.59E-08 | 3.02E-06 | (A) NE (B/D) |
| RNY5P8 | 6.78 | 4.42 | 1.12 | 1.86 | 2.36 | 9.37E-05 | 1.03E-03 | (A) NE (B/D) |
| POU5F1P6 | 3.3 | 0.94 | 1.11 | 0.88 | 2.36 | 1.12E-04 | 1.18E-03 | (A) NE (B/D) |
| PCDHB10 | 2.63 | 0.27 | 1.36 | 0.74 | 2.36 | 6.40E-04 | 4.53E-03 | (A) NE (B/D) |
| THEMIS3P | 3.13 | 0.76 | 2.12 | 1.17 | 2.36 | 9.98E-03 | 0.0356 | (A) NE (B/D) |
| ZNF618 | 3.41 | 1.06 | 0.54 | 0.67 | 2.35 | 1.14E-08 | 8.85E-07 | (A) NE (B/D) |
| ARHGEF16 | 4.71 | 2.36 | 0.64 | 1.25 | 2.35 | 3.39E-08 | 2.05E-06 | (A) NE (B/D) |
| PSRC1 | 2.56 | 0.22 | 0.95 | 0.65 | 2.35 | 3.83E-05 | 5.14E-04 | (A) NE (B/D) |
| RPL39L | 3.36 | 1.01 | 1.03 | 1.15 | 2.35 | 3.87E-05 | 5.17E-04 | (A) NE (B/D) |
| CEMIP | 2.51 | 0.15 | 1.19 | 0.52 | 2.35 | 2.85E-04 | 2.42E-03 | (A) NE (B/D) |
| PARM1 | 4.11 | 1.76 | 1.26 | 1.21 | 2.35 | 2.86E-04 | 2.43E-03 | (A) NE (B/D) |
| RTL1 | 3.12 | 0.76 | 2.15 | 1.22 | 2.35 | 0.011 | 0.0381 | (A) NE (B/D) |
| TLE5 | 3.49 | 1.15 | 0.18 | 1.15 | 2.34 | 8.09E-14 | 9.05E-11 | (A) NE (B/D) |
| LINC00479 | 3.26 | 0.92 | 0.61 | 0.95 | 2.34 | 2.22E-08 | 1.43E-06 | (A) NE (B/D) |
| PAF1 | 5.01 | 2.67 | 0.57 | 1.66 | 2.34 | 3.58E-08 | 2.13E-06 | (A) NE (B/D) |
| SUGP1 | 2.82 | 0.47 | 0.93 | 0.58 | 2.34 | 3.29E-05 | 4.56E-04 | (A) NE (B/D) |
| CEP55 | 5.77 | 3.43 | 1.04 | 1.63 | 2.34 | 3.85E-05 | 5.16E-04 | (A) NE (B/D) |
| ARHGAP11B | 3.87 | 1.52 | 1.1 | 1.1 | 2.34 | 8.63E-05 | 9.71E-04 | (A) NE (B/D) |
| SCNN1A | 3.07 | 0.73 | 1.42 | 1 | 2.34 | 8.46E-04 | 5.59E-03 | (A) NE (B/D) |
| PRSS45P | 3.41 | 1.07 | 1.55 | 1.19 | 2.34 | 1.60E-03 | 9.05E-03 | (A) NE (B/D) |
| LINC00331 | 2.68 | 0.34 | 2.02 | 0.61 | 2.34 | 8.12E-03 | 0.0306 | (A) NE (B/D) |
| CUEDC1 | 5.72 | 3.38 | 0.72 | 1.06 | 2.33 | 3.55E-07 | 1.30E-05 | (A) NE (B/D) |
| DMAP1 | 7.88 | 5.55 | 0.82 | 1.19 | 2.33 | 2.11E-06 | 5.33E-05 | (A) NE (B/D) |
| HOXC5 | 3.9 | 1.57 | 0.87 | 1.38 | 2.33 | 4.35E-06 | 9.48E-05 | (A) NE (B/D) |
| SHISA2 | 2.4 | 0.07 | 0.86 | 0.37 | 2.33 | 2.58E-05 | 3.75E-04 | (A) NE (B/D) |
| URB1-AS1 | 4.97 | 2.65 | 0.39 | 1.43 | 2.32 | 2.01E-10 | 4.12E-08 | (A) NE (B/D) |
| TECRP1 | 8.17 | 5.85 | 0.42 | 1.41 | 2.32 | 2.94E-10 | 5.60E-08 | (A) NE (B/D) |
| BMP8B | 3.63 | 1.32 | 0.59 | 0.97 | 2.32 | 1.16E-08 | 8.90E-07 | (A) NE (B/D) |
| ATP6V1B1 | 2.85 | 0.53 | 0.55 | 0.66 | 2.32 | 2.14E-08 | 1.39E-06 | (A) NE (B/D) |
| LFNG | 2.74 | 0.42 | 0.57 | 0.48 | 2.32 | 2.09E-07 | 8.64E-06 | (A) NE (B/D) |
| SPAG5 | 3.01 | 0.69 | 1.26 | 0.9 | 2.32 | 3.72E-04 | 2.99E-03 | (A) NE (B/D) |
| DLL1 | 3.09 | 0.76 | 1.39 | 0.76 | 2.32 | 8.49E-04 | 5.60E-03 | (A) NE (B/D) |
| RPS12P21 | 3.72 | 1.4 | 1.98 | 1.02 | 2.32 | 7.67E-03 | 0.0294 | (A) NE (B/D) |
| DAPK1 | 7.3 | 4.99 | 0.59 | 1.47 | 2.31 | 2.37E-08 | 1.50E-06 | (A) NE (B/D) |
| CNKSR1 | 3.78 | 1.47 | 0.73 | 1.28 | 2.31 | 3.85E-07 | 1.39E-05 | (A) NE (B/D) |
| TCF19 | 7.06 | 4.75 | 0.91 | 1.75 | 2.31 | 1.26E-05 | 2.20E-04 | (A) NE (B/D) |
| HAUS5 | 3.29 | 0.98 | 0.92 | 1.01 | 2.31 | 1.39E-05 | 2.37E-04 | (A) NE (B/D) |
| CKS2 | 8.33 | 6.02 | 1.01 | 2.11 | 2.31 | 6.67E-05 | 7.92E-04 | (A) NE (B/D) |
| ZNF367 | 5.73 | 3.42 | 1.17 | 1.58 | 2.31 | 1.54E-04 | 1.51E-03 | (A) NE (B/D) |
| ATP1B1 | 7.33 | 5.02 | 1.36 | 1.41 | 2.31 | 6.13E-04 | 4.38E-03 | (A) NE (B/D) |
| PCDHB14 | 3.4 | 1.1 | 1.4 | 0.77 | 2.31 | 9.28E-04 | 5.96E-03 | (A) NE (B/D) |
| SOX21 | 2.57 | 0.26 | 1.55 | 0.86 | 2.31 | 1.83E-03 | 0.01 | (A) NE (B/D) |
| SNORD113-4 | 2.43 | 0.12 | 1.87 | 0.36 | 2.31 | 5.96E-03 | 0.0245 | (A) NE (B/D) |
| HMSD | 5.65 | 3.35 | 1.95 | 1 | 2.31 | 7.34E-03 | 0.0285 | (A) NE (B/D) |
| ACAP3 | 8.25 | 5.94 | 0.53 | 0.78 | 2.3 | 3.31E-09 | 3.47E-07 | (A) NE (B/D) |
| ATG4D | 5.67 | 3.37 | 0.56 | 1.11 | 2.3 | 3.79E-09 | 3.75E-07 | (A) NE (B/D) |
| ICA1 | 3.62 | 1.32 | 0.6 | 1.24 | 2.3 | 1.50E-08 | 1.06E-06 | (A) NE (B/D) |
| GUCA1B | 3.31 | 1.01 | 1.23 | 0.89 | 2.3 | 3.31E-04 | 2.72E-03 | (A) NE (B/D) |
| ZSCAN12P1 | 4.17 | 1.87 | 1.24 | 0.83 | 2.3 | 3.66E-04 | 2.94E-03 | (A) NE (B/D) |
| HOXB-AS1 | 5.83 | 3.54 | 1.69 | 1.36 | 2.3 | 3.16E-03 | 0.0152 | (A) NE (B/D) |
| HNRNPA1P47 | 2.76 | 0.46 | 1.9 | 0.83 | 2.3 | 6.41E-03 | 0.0258 | (A) NE (B/D) |
| GALNT12 | 6.7 | 4.4 | 1.87 | 2.17 | 2.3 | 6.63E-03 | 0.0265 | (A) NE (B/D) |
| PCDHA8 | 5.33 | 3.04 | 0.89 | 1.77 | 2.29 | 1.12E-05 | 1.99E-04 | (A) NE (B/D) |
| BPIFB4 | 3.47 | 1.18 | 1.13 | 1.18 | 2.29 | 1.33E-04 | 1.35E-03 | (A) NE (B/D) |
| C21ORF58 | 4.16 | 1.87 | 1.31 | 1.24 | 2.29 | 5.01E-04 | 3.75E-03 | (A) NE (B/D) |
| SP9 | 4.22 | 1.92 | 1.66 | 1.92 | 2.29 | 3.10E-03 | 0.015 | (A) NE (B/D) |

| | | | | | | | | |
|---|---|---|---|---|---|---|---|---|
| CASC9 | 2.96 | 0.66 | 1.78 | 1.08 | 2.29 | 4.47E-03 | 0.0198 | (A) NE (B/D) |
| TMED1 | 4.68 | 2.4 | 0.53 | 1.44 | 2.28 | 6.94E-09 | 5.91E-07 | (A) NE (B/D) |
| NRM | 2.84 | 0.56 | 0.55 | 0.77 | 2.28 | 9.72E-09 | 7.79E-07 | (A) NE (B/D) |
| DTNA | 3.31 | 1.03 | 0.59 | 1 | 2.28 | 1.45E-08 | 1.03E-06 | (A) NE (B/D) |
| ATP13A1 | 3.75 | 1.47 | 0.6 | 1.13 | 2.28 | 1.91E-08 | 1.29E-06 | (A) NE (B/D) |
| USP11 | 3.06 | 0.78 | 1 | 1.01 | 2.28 | 4.54E-05 | 5.86E-04 | (A) NE (B/D) |
| C1ORF226 | 3.91 | 1.64 | 1.16 | 1.27 | 2.28 | 1.66E-04 | 1.60E-03 | (A) NE (B/D) |
| C1ORF210 | 4.22 | 1.94 | 1.16 | 1.67 | 2.28 | 1.68E-04 | 1.61E-03 | (A) NE (B/D) |
| MIR-4697/3P | 3.72 | 1.44 | 1.34 | 1.22 | 2.28 | 6.14E-04 | 4.38E-03 | (A) NE (B/D) |
| MIR-4697/5P | 3.72 | 1.44 | 1.34 | 1.22 | 2.28 | 6.14E-04 | 4.38E-03 | (A) NE (B/D) |
| SMAD9-IT1 | 3.56 | 1.29 | 1.95 | 0.91 | 2.28 | 7.81E-03 | 0.0297 | (A) NE (B/D) |
| RPSAP41 | 5.47 | 3.2 | 0.65 | 1.26 | 2.27 | 8.39E-08 | 4.19E-06 | (A) NE (B/D) |
| CLPSL2 | 3.17 | 0.9 | 1.63 | 1.07 | 2.27 | 2.75E-03 | 0.0137 | (A) NE (B/D) |
| ROBO1 | 8.36 | 6.09 | 1.71 | 2.04 | 2.27 | 4.19E-03 | 0.0188 | (A) NE (B/D) |
| LRRC61 | 6.19 | 3.93 | 0.54 | 0.88 | 2.26 | 3.72E-09 | 3.69E-07 | (A) NE (B/D) |
| MIR-639/639 | 3.34 | 1.08 | 0.58 | 0.85 | 2.26 | 1.92E-08 | 1.29E-06 | (A) NE (B/D) |
| ULK3 | 8.75 | 6.5 | 0.76 | 1.18 | 2.26 | 1.14E-06 | 3.30E-05 | (A) NE (B/D) |
| SMG9 | 3.53 | 1.27 | 0.87 | 1.1 | 2.26 | 8.07E-06 | 1.54E-04 | (A) NE (B/D) |
| PGPEP1 | 2.55 | 0.29 | 1.09 | 0.52 | 2.26 | 1.95E-04 | 1.81E-03 | (A) NE (B/D) |
| LAPTM4B | 12.99 | 10.73 | 1.18 | 1.37 | 2.26 | 2.13E-04 | 1.93E-03 | (A) NE (B/D) |
| DNAJC3-DT | 3.64 | 1.38 | 1.2 | 0.72 | 2.26 | 3.53E-04 | 2.86E-03 | (A) NE (B/D) |
| MIR-3118-1/ | 4.43 | 2.17 | 1.2 | 2.52 | 2.26 | 6.24E-04 | 4.43E-03 | (A) NE (B/D) |
| MAGEA1 | 2.28 | 0.02 | 1.47 | 0.11 | 2.26 | 1.76E-03 | 9.72E-03 | (A) NE (B/D) |
| LINC00928 | 2.4 | 0.14 | 1.83 | 0.54 | 2.26 | 5.83E-03 | 0.0241 | (A) NE (B/D) |
| C19ORF73 | 5.67 | 3.42 | 0.38 | 1.14 | 2.25 | 8.43E-12 | 3.54E-09 | (A) NE (B/D) |
| NDUFA13 | 4.44 | 2.19 | 0.52 | 1.52 | 2.25 | 1.40E-08 | 1.01E-06 | (A) NE (B/D) |
| SVBP | 7.59 | 5.35 | 0.83 | 1.59 | 2.25 | 4.85E-06 | 1.03E-04 | (A) NE (B/D) |
| TSPYL2 | 8.63 | 6.37 | 1.28 | 1.66 | 2.25 | 4.52E-04 | 3.46E-03 | (A) NE (B/D) |
| PGAP1 | 4.64 | 2.39 | 1.47 | 0.81 | 2.25 | 1.56E-03 | 8.90E-03 | (A) NE (B/D) |
| STMND1 | 2.76 | 0.52 | 1.79 | 0.86 | 2.25 | 5.21E-03 | 0.0221 | (A) NE (B/D) |
| SLC17A9 | 4.01 | 1.77 | 1.9 | 1.46 | 2.25 | 7.41E-03 | 0.0287 | (A) NE (B/D) |
| MIR-1233-1/ | 3.52 | 1.28 | 0.91 | 1.23 | 2.24 | 1.47E-05 | 2.46E-04 | (A) NE (B/D) |
| MIR-1233-1/ | 3.52 | 1.28 | 0.91 | 1.23 | 2.24 | 1.47E-05 | 2.46E-04 | (A) NE (B/D) |
| MIR-1233-1/ | 3.52 | 1.28 | 0.91 | 1.23 | 2.24 | 1.47E-05 | 2.46E-04 | (A) NE (B/D) |
| RNA5SP283 | 5.47 | 3.23 | 1.19 | 2.14 | 2.24 | 4.01E-04 | 3.16E-03 | (A) NE (B/D) |
| ZDHHC8P1 | 4 | 1.76 | 1.38 | 1.89 | 2.24 | 9.72E-04 | 6.19E-03 | (A) NE (B/D) |
| BTBD3 | 5.73 | 3.49 | 1.38 | 0.78 | 2.24 | 1.06E-03 | 6.59E-03 | (A) NE (B/D) |
| RBM20 | 2.36 | 0.12 | 1.92 | 0.31 | 2.24 | 8.11E-03 | 0.0306 | (A) NE (B/D) |
| FZD1 | 3.37 | 1.14 | 0.52 | 1.09 | 2.23 | 1.38E-09 | 1.74E-07 | (A) NE (B/D) |
| NDUFA7 | 4.75 | 2.52 | 0.27 | 1.73 | 2.23 | 1.28E-08 | 9.48E-07 | (A) NE (B/D) |
| IRS2 | 7.83 | 5.6 | 0.62 | 1.46 | 2.23 | 8.80E-08 | 4.34E-06 | (A) NE (B/D) |
| EMC9 | 6.34 | 4.11 | 0.75 | 1.46 | 2.23 | 1.09E-06 | 3.21E-05 | (A) NE (B/D) |
| MAU2 | 4.65 | 2.42 | 0.8 | 0.81 | 2.23 | 6.43E-06 | 1.29E-04 | (A) NE (B/D) |
| ZNF266 | 7.36 | 5.13 | 1.04 | 1.26 | 2.23 | 7.07E-05 | 8.28E-04 | (A) NE (B/D) |
| PI4KAP2 | 3.02 | 0.79 | 1 | 0.63 | 2.23 | 8.87E-05 | 9.91E-04 | (A) NE (B/D) |
| UHRF1 | 5.74 | 3.52 | 1.08 | 1.45 | 2.23 | 9.69E-05 | 1.06E-03 | (A) NE (B/D) |
| SLC35G2 | 4.23 | 2 | 1.28 | 0.78 | 2.23 | 6.26E-04 | 4.44E-03 | (A) NE (B/D) |
| TBXT | 2.47 | 0.24 | 1.49 | 0.94 | 2.23 | 1.77E-03 | 9.76E-03 | (A) NE (B/D) |
| GLRA2 | 2.58 | 0.35 | 1.85 | 0.61 | 2.23 | 6.69E-03 | 0.0266 | (A) NE (B/D) |
| TEDC2 | 6.93 | 4.71 | 0.39 | 1.5 | 2.22 | 1.57E-09 | 1.92E-07 | (A) NE (B/D) |
| CDK3 | 8.61 | 6.39 | 0.64 | 1.54 | 2.22 | 1.90E-07 | 8.06E-06 | (A) NE (B/D) |
| FN3K | 3.05 | 0.83 | 0.84 | 1.01 | 2.22 | 7.48E-06 | 1.45E-04 | (A) NE (B/D) |
| MIA-RAB4B | 4.79 | 2.57 | 0.92 | 1.49 | 2.22 | 1.74E-05 | 2.77E-04 | (A) NE (B/D) |
| PILRB | 6.3 | 4.07 | 1.07 | 1.04 | 2.22 | 1.08E-04 | 1.15E-03 | (A) NE (B/D) |
| KLHL35 | 4.3 | 2.08 | 1.09 | 1.26 | 2.22 | 1.19E-04 | 1.24E-03 | (A) NE (B/D) |
| GRID1 | 4.2 | 1.97 | 1.11 | 1.25 | 2.22 | 1.36E-04 | 1.37E-03 | (A) NE (B/D) |
| CTSV | 2.35 | 0.13 | 1.17 | 0.67 | 2.22 | 3.32E-04 | 2.72E-03 | (A) NE (B/D) |
| TIAM1 | 7.39 | 5.17 | 1.25 | 1.24 | 2.22 | 4.20E-04 | 3.27E-03 | (A) NE (B/D) |
| MRPL23-AS1 | 3.04 | 0.82 | 1.82 | 1 | 2.22 | 6.24E-03 | 0.0253 | (A) NE (B/D) |
| NIPA2P4 | 2.62 | 0.41 | 2.12 | 0.73 | 2.22 | 0.0139 | 0.0454 | (A) NE (B/D) |
| MRPS26 | 5.99 | 3.78 | 0.46 | 1.11 | 2.21 | 1.94E-10 | 4.08E-08 | (A) NE (B/D) |
| MIR-1914/19 | 3.51 | 1.3 | 0.53 | 1.2 | 2.21 | 3.35E-09 | 3.47E-07 | (A) NE (B/D) |
| MIR-1914/19 | 3.51 | 1.3 | 0.53 | 1.2 | 2.21 | 3.35E-09 | 3.47E-07 | (A) NE (B/D) |
| MIR-1914/3P | 3.51 | 1.3 | 0.53 | 1.2 | 2.21 | 3.35E-09 | 3.47E-07 | (A) NE (B/D) |
| MIR-1914/5P | 3.51 | 1.3 | 0.53 | 1.2 | 2.21 | 3.35E-09 | 3.47E-07 | (A) NE (B/D) |

| | | | | | | | | |
|---|---|---|---|---|---|---|---|---|
| COQ10A | 3.46 | 1.25 | 0.68 | 1.24 | 2.21 | 2.32E-07 | 9.34E-06 | (A) NE (B/D) |
| CBFA2T2 | 8.02 | 5.81 | 0.7 | 0.96 | 2.21 | 6.07E-07 | 2.00E-05 | (A) NE (B/D) |
| PCDHA6 | 5.18 | 2.97 | 0.87 | 1.78 | 2.21 | 1.58E-05 | 2.59E-04 | (A) NE (B/D) |
| PCDH8P1 | 2.84 | 0.64 | 1.24 | 1.07 | 2.21 | 4.30E-04 | 3.33E-03 | (A) NE (B/D) |
| CCDC85A | 5.26 | 3.05 | 2.07 | 1.32 | 2.21 | 0.0126 | 0.0423 | (A) NE (B/D) |
| PAXX | 5.83 | 3.63 | 0.3 | 1.3 | 2.2 | 3.08E-11 | 9.33E-09 | (A) NE (B/D) |
| DUS3L | 3.38 | 1.17 | 0.57 | 1.22 | 2.2 | 1.33E-08 | 9.72E-07 | (A) NE (B/D) |
| MTURN | 7.37 | 5.17 | 0.88 | 1.18 | 2.2 | 1.15E-05 | 2.04E-04 | (A) NE (B/D) |
| LINC00654 | 3.14 | 0.95 | 0.97 | 0.73 | 2.2 | 7.13E-05 | 8.34E-04 | (A) NE (B/D) |
| PKMP4 | 7.05 | 4.85 | 0.97 | 2.06 | 2.2 | 7.78E-05 | 8.91E-04 | (A) NE (B/D) |
| TSPEAR-AS2 | 3.9 | 1.7 | 1.81 | 1.36 | 2.2 | 6.40E-03 | 0.0258 | (A) NE (B/D) |
| FFAR4 | 3.15 | 0.95 | 2.02 | 1.32 | 2.2 | 0.0117 | 0.0399 | (A) NE (B/D) |
| PEG10 | 2.24 | 0.05 | 0.98 | 0.21 | 2.19 | 1.49E-04 | 1.47E-03 | (A) NE (B/D) |
| MIR924HG | 2.53 | 0.34 | 1.08 | 0.66 | 2.19 | 2.05E-04 | 1.87E-03 | (A) NE (B/D) |
| TM6SF2 | 3.99 | 1.8 | 1.17 | 1.32 | 2.19 | 2.59E-04 | 2.24E-03 | (A) NE (B/D) |
| EVA1A | 2.2 | 0 | 1.44 | 0.02 | 2.19 | 1.86E-03 | 0.0101 | (A) NE (B/D) |
| CNGA4 | 3.36 | 1.17 | 1.82 | 1.02 | 2.19 | 6.57E-03 | 0.0263 | (A) NE (B/D) |
| ACD | 2.49 | 0.31 | 0.49 | 0.58 | 2.18 | 1.40E-08 | 1.01E-06 | (A) NE (B/D) |
| OVOL2 | 3.03 | 0.85 | 0.82 | 0.65 | 2.18 | 1.43E-05 | 2.42E-04 | (A) NE (B/D) |
| SELENOW | 9.27 | 7.09 | 0.92 | 1.23 | 2.18 | 2.02E-05 | 3.10E-04 | (A) NE (B/D) |
| FAM226A | 6.21 | 4.04 | 1.02 | 1.62 | 2.18 | 7.26E-05 | 8.47E-04 | (A) NE (B/D) |
| BEST4 | 4.55 | 2.37 | 1.23 | 1.19 | 2.18 | 4.18E-04 | 3.26E-03 | (A) NE (B/D) |
| GSTM3P1 | 2.83 | 0.64 | 1.69 | 1.14 | 2.18 | 4.37E-03 | 0.0195 | (A) NE (B/D) |
| CFAP65 | 4.59 | 2.42 | 0.41 | 1.3 | 2.17 | 3.22E-10 | 5.95E-08 | (A) NE (B/D) |
| AP1M2 | 4 | 1.84 | 0.27 | 1.68 | 2.17 | 1.38E-08 | 1.00E-06 | (A) NE (B/D) |
| DMWD | 2.91 | 0.74 | 0.66 | 0.65 | 2.17 | 1.04E-06 | 3.08E-05 | (A) NE (B/D) |
| CASTOR3 | 7.06 | 4.89 | 0.85 | 1.15 | 2.17 | 9.30E-06 | 1.72E-04 | (A) NE (B/D) |
| CDKN2D | 2.89 | 0.72 | 0.94 | 1.08 | 2.17 | 3.11E-05 | 4.36E-04 | (A) NE (B/D) |
| ENOSF1 | 3.08 | 0.91 | 1.08 | 1.14 | 2.17 | 1.39E-04 | 1.40E-03 | (A) NE (B/D) |
| TPBG | 3.23 | 1.06 | 1.44 | 1.11 | 2.17 | 1.59E-03 | 9.01E-03 | (A) NE (B/D) |
| PON2 | 5.93 | 3.76 | 1.5 | 1.05 | 2.17 | 2.10E-03 | 0.0111 | (A) NE (B/D) |
| LRP4-AS1 | 2.43 | 0.27 | 1.48 | 0.43 | 2.17 | 2.18E-03 | 0.0114 | (A) NE (B/D) |
| B3GAT1 | 5.96 | 3.79 | 1.55 | 1.77 | 2.17 | 2.77E-03 | 0.0138 | (A) NE (B/D) |
| WDR83OS | 4.39 | 2.23 | 0.5 | 1.83 | 2.16 | 3.53E-07 | 1.29E-05 | (A) NE (B/D) |
| RNA5SP244 | 6.26 | 4.1 | 0.38 | 1.98 | 2.16 | 5.20E-07 | 1.77E-05 | (A) NE (B/D) |
| SPRED2 | 8.31 | 6.15 | 0.73 | 0.88 | 2.16 | 1.86E-06 | 4.88E-05 | (A) NE (B/D) |
| SDR39U1 | 3.94 | 1.78 | 0.81 | 1.54 | 2.16 | 5.99E-06 | 1.22E-04 | (A) NE (B/D) |
| HMGN2P4 | 8.41 | 6.25 | 0.88 | 1.97 | 2.16 | 3.56E-05 | 4.84E-04 | (A) NE (B/D) |
| TMEM151B | 2.6 | 0.45 | 0.99 | 0.22 | 2.16 | 1.76E-04 | 1.67E-03 | (A) NE (B/D) |
| NRCAM | 3.38 | 1.22 | 1.17 | 1.32 | 2.16 | 2.80E-04 | 2.39E-03 | (A) NE (B/D) |
| LINC00888 | 6.04 | 3.88 | 1.18 | 1.22 | 2.16 | 3.29E-04 | 2.70E-03 | (A) NE (B/D) |
| MIR-3153/31 | 5.36 | 3.2 | 1.26 | 1.58 | 2.16 | 5.57E-04 | 4.07E-03 | (A) NE (B/D) |
| TCERG1L | 2.3 | 0.14 | 1.26 | 0.31 | 2.16 | 8.40E-04 | 5.56E-03 | (A) NE (B/D) |
| HOXC-AS1 | 2.21 | 0.05 | 1.32 | 0.17 | 2.16 | 1.20E-03 | 7.24E-03 | (A) NE (B/D) |
| TMEM255B | 2.96 | 0.8 | 1.38 | 0.86 | 2.16 | 1.28E-03 | 7.63E-03 | (A) NE (B/D) |
| DLGAP1-AS2 | 2.83 | 0.67 | 1.39 | 0.64 | 2.16 | 1.48E-03 | 8.56E-03 | (A) NE (B/D) |
| TGFBR3L | 4.01 | 1.85 | 1.56 | 1.19 | 2.16 | 2.84E-03 | 0.014 | (A) NE (B/D) |
| AGPAT4 | 5.41 | 3.25 | 1.96 | 1.22 | 2.16 | 0.0108 | 0.0376 | (A) NE (B/D) |
| NCAPH2 | 3.9 | 1.74 | 0.67 | 1.25 | 2.15 | 2.75E-07 | 1.07E-05 | (A) NE (B/D) |
| LPIN3 | 7.98 | 5.83 | 0.81 | 1.67 | 2.15 | 8.61E-06 | 1.62E-04 | (A) NE (B/D) |
| CRIPAK | 6.73 | 4.58 | 0.88 | 1.45 | 2.15 | 1.48E-05 | 2.48E-04 | (A) NE (B/D) |
| GYG1 | 3.97 | 1.82 | 1.3 | 1.21 | 2.15 | 7.58E-04 | 5.14E-03 | (A) NE (B/D) |
| RNF208 | 7.58 | 5.43 | 1.34 | 1.52 | 2.15 | 9.69E-04 | 6.18E-03 | (A) NE (B/D) |
| CHRM4 | 2.23 | 0.08 | 1.32 | 0.29 | 2.15 | 1.20E-03 | 7.29E-03 | (A) NE (B/D) |
| SYN2 | 3.49 | 1.34 | 1.76 | 1.08 | 2.15 | 5.98E-03 | 0.0246 | (A) NE (B/D) |
| MBD3 | 3.46 | 1.32 | 0.44 | 1.2 | 2.14 | 3.98E-10 | 7.05E-08 | (A) NE (B/D) |
| GPR108 | 3.49 | 1.35 | 0.45 | 1.27 | 2.14 | 9.93E-10 | 1.36E-07 | (A) NE (B/D) |
| CCDC40 | 6.18 | 4.04 | 0.49 | 1.45 | 2.14 | 1.30E-08 | 9.59E-07 | (A) NE (B/D) |
| PHF1 | 6.98 | 4.84 | 0.66 | 1.37 | 2.14 | 3.32E-07 | 1.25E-05 | (A) NE (B/D) |
| SNORD56 | 9.49 | 7.35 | 0.56 | 2.12 | 2.14 | 5.44E-06 | 1.13E-04 | (A) NE (B/D) |
| FAM166A | 4.5 | 2.35 | 0.94 | 1.47 | 2.14 | 3.32E-05 | 4.58E-04 | (A) NE (B/D) |
| CATSPER2 | 3.55 | 1.41 | 0.97 | 1.1 | 2.14 | 5.66E-05 | 6.97E-04 | (A) NE (B/D) |
| ZSCAN16-AS | 4.07 | 1.93 | 1 | 1.29 | 2.14 | 6.68E-05 | 7.93E-04 | (A) NE (B/D) |
| BRD8 | 3.24 | 1.1 | 1.14 | 1.22 | 2.14 | 2.67E-04 | 2.30E-03 | (A) NE (B/D) |
| LINC00346 | 3.22 | 1.09 | 1.38 | 1.12 | 2.14 | 1.32E-03 | 7.80E-03 | (A) NE (B/D) |

| | | | | | | | | |
|---|---|---|---|---|---|---|---|---|
| DTX3 | 5.43 | 3.3 | 0.44 | 1.52 | 2.13 | 1.16E-08 | 8.90E-07 | (A) NE (B/D) |
| CD320 | 3.69 | 1.56 | 0.57 | 1.42 | 2.13 | 5.95E-08 | 3.16E-06 | (A) NE (B/D) |
| MDP1 | 5.33 | 3.2 | 0.81 | 1.38 | 2.13 | 6.21E-06 | 1.25E-04 | (A) NE (B/D) |
| KPNA2 | 5.27 | 3.14 | 0.87 | 1.49 | 2.13 | 1.60E-05 | 2.61E-04 | (A) NE (B/D) |
| ACYP1 | 5.66 | 3.53 | 0.93 | 1.07 | 2.13 | 3.82E-05 | 5.14E-04 | (A) NE (B/D) |
| AP1G2 | 3.11 | 0.98 | 0.98 | 1.08 | 2.13 | 6.77E-05 | 8.01E-04 | (A) NE (B/D) |
| EXTL3 | 3.51 | 1.38 | 1.01 | 0.92 | 2.13 | 1.01E-04 | 1.09E-03 | (A) NE (B/D) |
| ACTG1P4 | 2.34 | 0.21 | 1.73 | 0.42 | 2.13 | 6.04E-03 | 0.0247 | (A) NE (B/D) |
| HERC2P8 | 3.82 | 1.69 | 1.8 | 0.98 | 2.13 | 7.21E-03 | 0.0282 | (A) NE (B/D) |
| MAP3K9 | 7.93 | 5.81 | 0.85 | 1.18 | 2.12 | 1.12E-05 | 2.00E-04 | (A) NE (B/D) |
| STAG3L5P-PV | 4.42 | 2.3 | 1.1 | 0.95 | 2.12 | 2.28E-04 | 2.03E-03 | (A) NE (B/D) |
| CAPN13 | 5.64 | 3.52 | 1.28 | 2 | 2.12 | 9.69E-04 | 6.18E-03 | (A) NE (B/D) |
| MAGEA6 | 2.21 | 0.09 | 1.3 | 0.38 | 2.12 | 1.11E-03 | 6.86E-03 | (A) NE (B/D) |
| MARK1 | 6.83 | 4.71 | 1.53 | 1.18 | 2.12 | 2.78E-03 | 0.0138 | (A) NE (B/D) |
| RPL36AP26 | 3.23 | 1.11 | 1.57 | 1.46 | 2.12 | 3.30E-03 | 0.0157 | (A) NE (B/D) |
| PTCSC3 | 2.31 | 0.19 | 2.04 | 0.43 | 2.12 | 0.0143 | 0.0463 | (A) NE (B/D) |
| PSD | 2.42 | 0.31 | 0.57 | 0.59 | 2.11 | 2.38E-07 | 9.55E-06 | (A) NE (B/D) |
| E2F2 | 6.4 | 4.29 | 0.81 | 2.22 | 2.11 | 6.12E-05 | 7.41E-04 | (A) NE (B/D) |
| RBM38 | 4.71 | 2.6 | 0.92 | 2.04 | 2.11 | 8.67E-05 | 9.74E-04 | (A) NE (B/D) |
| WNT9A | 2.34 | 0.23 | 1.14 | 0.52 | 2.11 | 4.49E-04 | 3.44E-03 | (A) NE (B/D) |
| PER3 | 2.82 | 0.71 | 1.33 | 0.97 | 2.11 | 1.12E-03 | 6.86E-03 | (A) NE (B/D) |
| TEX30 | 5.89 | 3.78 | 1.36 | 1.46 | 2.11 | 1.24E-03 | 7.41E-03 | (A) NE (B/D) |
| CRIP1 | 5.93 | 3.82 | 1.49 | 1.9 | 2.11 | 2.75E-03 | 0.0137 | (A) NE (B/D) |
| C2CD4C | 5.73 | 3.61 | 1.51 | 2.17 | 2.11 | 3.38E-03 | 0.016 | (A) NE (B/D) |
| FAM102A | 4.78 | 2.68 | 0.21 | 1.28 | 2.1 | 3.75E-11 | 1.01E-08 | (A) NE (B/D) |
| LMNB2 | 3.04 | 0.94 | 0.5 | 0.93 | 2.1 | 2.33E-09 | 2.69E-07 | (A) NE (B/D) |
| HSPB2-C11O | 5.93 | 3.83 | 0.41 | 1.76 | 2.1 | 1.43E-07 | 6.42E-06 | (A) NE (B/D) |
| H2AFZ | 5.11 | 3.01 | 0.69 | 1.46 | 2.1 | 1.18E-06 | 3.39E-05 | (A) NE (B/D) |
| LINC00482 | 2.71 | 0.61 | 1.35 | 0.68 | 2.1 | 1.39E-03 | 8.11E-03 | (A) NE (B/D) |
| ARHGAP11A | 2.93 | 0.83 | 1.46 | 0.98 | 2.1 | 2.19E-03 | 0.0114 | (A) NE (B/D) |
| C4ORF48 | 6.82 | 4.72 | 1.5 | 1.19 | 2.1 | 2.63E-03 | 0.0132 | (A) NE (B/D) |
| PNMA2 | 3.83 | 1.72 | 1.83 | 1.43 | 2.1 | 8.80E-03 | 0.0325 | (A) NE (B/D) |
| NDUFS7 | 2.92 | 0.84 | 0.4 | 0.96 | 2.09 | 2.72E-11 | 8.34E-09 | (A) NE (B/D) |
| VARS2 | 9.14 | 7.04 | 0.48 | 1.19 | 2.09 | 2.22E-09 | 2.60E-07 | (A) NE (B/D) |
| PPP5C | 3.04 | 0.95 | 0.51 | 0.99 | 2.09 | 4.07E-09 | 3.98E-07 | (A) NE (B/D) |
| PTMAP3 | 5.18 | 3.09 | 0.59 | 1.68 | 2.09 | 6.07E-07 | 2.00E-05 | (A) NE (B/D) |
| TTC22 | 3.01 | 0.92 | 0.89 | 1 | 2.09 | 2.84E-05 | 4.04E-04 | (A) NE (B/D) |
| MAD2L1 | 7.07 | 4.98 | 1.09 | 1.7 | 2.09 | 2.48E-04 | 2.17E-03 | (A) NE (B/D) |
| YBX1P6 | 5.44 | 3.36 | 1.65 | 1.08 | 2.09 | 4.99E-03 | 0.0215 | (A) NE (B/D) |
| AGAP2-AS1 | 7.21 | 5.12 | 0.48 | 1.09 | 2.08 | 1.41E-09 | 1.76E-07 | (A) NE (B/D) |
| ILF3 | 9.86 | 7.78 | 0.51 | 1.15 | 2.08 | 4.63E-09 | 4.38E-07 | (A) NE (B/D) |
| FAM71E1 | 7.43 | 5.35 | 0.7 | 1.16 | 2.08 | 1.06E-06 | 3.13E-05 | (A) NE (B/D) |
| ORC1 | 5.05 | 2.98 | 0.8 | 1.59 | 2.08 | 9.55E-06 | 1.76E-04 | (A) NE (B/D) |
| S100A13 | 3.15 | 1.07 | 1.18 | 1.34 | 2.08 | 4.31E-04 | 3.34E-03 | (A) NE (B/D) |
| MIR-1302-8/ | 2.59 | 0.51 | 1.25 | 0.78 | 2.08 | 8.46E-04 | 5.59E-03 | (A) NE (B/D) |
| MCIDAS | 3.82 | 1.74 | 1.4 | 1.9 | 2.08 | 2.01E-03 | 0.0108 | (A) NE (B/D) |
| NXPH1 | 2.56 | 0.48 | 1.49 | 1.12 | 2.08 | 2.60E-03 | 0.0131 | (A) NE (B/D) |
| RN7SL268P | 4.01 | 1.94 | 0.41 | 1.2 | 2.07 | 3.62E-10 | 6.51E-08 | (A) NE (B/D) |
| SH3GL1P3 | 2.88 | 0.81 | 0.47 | 0.84 | 2.07 | 1.28E-09 | 1.63E-07 | (A) NE (B/D) |
| SLC29A1 | 6.05 | 3.98 | 0.56 | 1.38 | 2.07 | 7.79E-08 | 3.94E-06 | (A) NE (B/D) |
| PPIP5K1 | 3.83 | 1.75 | 0.75 | 1.07 | 2.07 | 3.23E-06 | 7.45E-05 | (A) NE (B/D) |
| BFSP1 | 2.59 | 0.52 | 0.9 | 0.58 | 2.07 | 6.86E-05 | 8.10E-04 | (A) NE (B/D) |
| WDR76 | 4.25 | 2.19 | 0.97 | 1.34 | 2.07 | 6.96E-05 | 8.20E-04 | (A) NE (B/D) |
| ASIC2 | 2.11 | 0.04 | 0.96 | 0.13 | 2.07 | 1.97E-04 | 1.82E-03 | (A) NE (B/D) |
| ITGA3 | 2.89 | 0.82 | 1.14 | 0.97 | 2.07 | 3.90E-04 | 3.10E-03 | (A) NE (B/D) |
| DLK2 | 4.63 | 2.55 | 1.29 | 1.15 | 2.07 | 9.40E-04 | 6.03E-03 | (A) NE (B/D) |
| ESPN | 5.15 | 3.08 | 1.59 | 1.57 | 2.07 | 4.34E-03 | 0.0193 | (A) NE (B/D) |
| RPS11P6 | 2.42 | 0.34 | 1.7 | 0.6 | 2.07 | 6.33E-03 | 0.0256 | (A) NE (B/D) |
| FLOT1 | 5.35 | 3.29 | 0.31 | 1.2 | 2.06 | 3.34E-11 | 9.66E-09 | (A) NE (B/D) |
| MIR-637/637 | 3.27 | 1.21 | 0.46 | 0.81 | 2.06 | 8.98E-10 | 1.27E-07 | (A) NE (B/D) |
| FBL | 4.45 | 2.39 | 0.43 | 1.78 | 2.06 | 2.94E-07 | 1.13E-05 | (A) NE (B/D) |
| KIF1A | 2.23 | 0.17 | 0.52 | 0.37 | 2.06 | 4.72E-07 | 1.65E-05 | (A) NE (B/D) |
| WRB | 2.99 | 0.93 | 0.66 | 1.09 | 2.06 | 4.77E-07 | 1.66E-05 | (A) NE (B/D) |
| UBE2T | 3.44 | 1.39 | 0.67 | 1.39 | 2.06 | 8.86E-07 | 2.70E-05 | (A) NE (B/D) |
| ZNF133 | 2.86 | 0.8 | 0.78 | 1 | 2.06 | 6.03E-06 | 1.22E-04 | (A) NE (B/D) |

| | | | | | | | | |
|---|---|---|---|---|---|---|---|---|
| GPS2 | 3.33 | 1.26 | 0.8 | 1.21 | 2.06 | 7.35E-06 | 1.43E-04 | (A) NE (B/D) |
| RPS12P28 | 2.64 | 0.58 | 0.88 | 0.72 | 2.06 | 4.49E-05 | 5.80E-04 | (A) NE (B/D) |
| MAGIX | 4.29 | 2.23 | 1 | 1.21 | 2.06 | 1.07E-04 | 1.14E-03 | (A) NE (B/D) |
| KCTD14 | 2.81 | 0.75 | 1 | 1.02 | 2.06 | 1.18E-04 | 1.23E-03 | (A) NE (B/D) |
| DCLRE1CP1 | 2.84 | 0.77 | 1.26 | 0.89 | 2.06 | 9.27E-04 | 5.96E-03 | (A) NE (B/D) |
| SNORD114-2 | 2.41 | 0.35 | 1.43 | 1.04 | 2.06 | 2.19E-03 | 0.0114 | (A) NE (B/D) |
| TMSB15B | 3.15 | 1.09 | 1.59 | 1.22 | 2.06 | 4.18E-03 | 0.0188 | (A) NE (B/D) |
| BPIFB2 | 2.1 | 0.04 | 1.58 | 0.25 | 2.06 | 4.44E-03 | 0.0197 | (A) NE (B/D) |
| HMGN3P1 | 6.58 | 4.52 | 1.69 | 2.05 | 2.06 | 7.44E-03 | 0.0288 | (A) NE (B/D) |
| ZSWIM4 | 3.4 | 1.35 | 0.3 | 0.76 | 2.05 | 4.16E-14 | 5.10E-11 | (A) NE (B/D) |
| SGTA | 3.74 | 1.69 | 0.15 | 1.25 | 2.05 | 3.50E-11 | 9.80E-09 | (A) NE (B/D) |
| TECR | 3.27 | 1.21 | 0.4 | 0.98 | 2.05 | 4.91E-11 | 1.30E-08 | (A) NE (B/D) |
| GDAP1 | 3.28 | 1.23 | 0.49 | 1.32 | 2.05 | 1.20E-08 | 9.09E-07 | (A) NE (B/D) |
| SNORD86 | 5.6 | 3.55 | 0.48 | 1.4 | 2.05 | 1.92E-08 | 1.29E-06 | (A) NE (B/D) |
| TMEM94 | 3.17 | 1.13 | 0.62 | 1 | 2.05 | 2.10E-07 | 8.66E-06 | (A) NE (B/D) |
| E2F1 | 2.96 | 0.92 | 0.65 | 0.94 | 2.05 | 5.85E-07 | 1.94E-05 | (A) NE (B/D) |
| PRAF2 | 8.64 | 6.59 | 0.51 | 1.86 | 2.05 | 1.25E-06 | 3.52E-05 | (A) NE (B/D) |
| NNT | 5.15 | 3.1 | 0.72 | 1.13 | 2.05 | 1.68E-06 | 4.49E-05 | (A) NE (B/D) |
| SLC25A6 | 12.3 | 10.24 | 0.76 | 1.33 | 2.05 | 4.17E-06 | 9.18E-05 | (A) NE (B/D) |
| ANKRD10 | 4.61 | 2.56 | 0.78 | 0.97 | 2.05 | 6.76E-06 | 1.34E-04 | (A) NE (B/D) |
| HMGN2 | 9.55 | 7.51 | 0.88 | 1.7 | 2.05 | 4.05E-05 | 5.35E-04 | (A) NE (B/D) |
| RINL | 5.82 | 3.77 | 1.15 | 1.36 | 2.05 | 3.87E-04 | 3.07E-03 | (A) NE (B/D) |
| C2ORF74 | 5.85 | 3.8 | 1.12 | 1.88 | 2.05 | 4.46E-04 | 3.43E-03 | (A) NE (B/D) |
| MIF | 5.53 | 3.48 | 1.32 | 1.14 | 2.05 | 1.29E-03 | 7.66E-03 | (A) NE (B/D) |
| SOX21-AS1 | 2.23 | 0.18 | 1.44 | 0.68 | 2.05 | 2.51E-03 | 0.0127 | (A) NE (B/D) |
| CFHR5 | 2.61 | 0.55 | 1.88 | 1.27 | 2.05 | 0.0112 | 0.0387 | (A) NE (B/D) |
| DOT1L | 4.73 | 2.69 | 0.29 | 0.67 | 2.04 | 1.01E-14 | 1.85E-11 | (A) NE (B/D) |
| EPHA10 | 2.81 | 0.76 | 0.4 | 0.8 | 2.04 | 3.44E-11 | 9.80E-09 | (A) NE (B/D) |
| TNPO2 | 7.32 | 5.29 | 0.41 | 1.01 | 2.04 | 9.83E-11 | 2.36E-08 | (A) NE (B/D) |
| CCHCR1 | 7.31 | 5.27 | 0.42 | 1.23 | 2.04 | 9.16E-10 | 1.28E-07 | (A) NE (B/D) |
| ALDOC | 5.63 | 3.59 | 0.6 | 1.18 | 2.04 | 1.38E-07 | 6.26E-06 | (A) NE (B/D) |
| FN3KRP | 3.79 | 1.75 | 0.51 | 1.65 | 2.04 | 3.06E-07 | 1.17E-05 | (A) NE (B/D) |
| DDR1 | 8.16 | 6.12 | 0.42 | 1.77 | 2.04 | 3.39E-07 | 1.26E-05 | (A) NE (B/D) |
| THSD7A | 3.91 | 1.87 | 0.82 | 1.61 | 2.04 | 1.63E-05 | 2.64E-04 | (A) NE (B/D) |
| RNU2-35P | 2.46 | 0.42 | 0.89 | 0.8 | 2.04 | 4.66E-05 | 5.97E-04 | (A) NE (B/D) |
| ZGLP1 | 3.89 | 1.84 | 0.95 | 1.41 | 2.04 | 6.78E-05 | 8.02E-04 | (A) NE (B/D) |
| SNRNP70 | 6.16 | 4.12 | 1.06 | 1.59 | 2.04 | 2.18E-04 | 1.96E-03 | (A) NE (B/D) |
| CCNA2 | 3.13 | 1.09 | 1.25 | 1.32 | 2.04 | 8.44E-04 | 5.58E-03 | (A) NE (B/D) |
| CDKL4 | 3.3 | 1.26 | 1.96 | 0.9 | 2.04 | 0.0144 | 0.0464 | (A) NE (B/D) |
| CEP131 | 5.14 | 3.11 | 0.35 | 0.98 | 2.03 | 4.85E-12 | 2.31E-09 | (A) NE (B/D) |
| CELSR2 | 6.83 | 4.8 | 0.47 | 1.3 | 2.03 | 7.23E-09 | 6.10E-07 | (A) NE (B/D) |
| MAPRE3 | 4 | 1.97 | 0.59 | 1.01 | 2.03 | 9.60E-08 | 4.64E-06 | (A) NE (B/D) |
| KLC4 | 3.64 | 1.61 | 0.6 | 1.12 | 2.03 | 1.46E-07 | 6.54E-06 | (A) NE (B/D) |
| C6ORF226 | 5.63 | 3.6 | 0.69 | 1.36 | 2.03 | 1.47E-06 | 4.06E-05 | (A) NE (B/D) |
| MIR-5581/3P | 7.13 | 5.1 | 0.73 | 1.44 | 2.03 | 3.63E-06 | 8.23E-05 | (A) NE (B/D) |
| MIR-5581/5P | 7.13 | 5.1 | 0.73 | 1.44 | 2.03 | 3.63E-06 | 8.23E-05 | (A) NE (B/D) |
| LCN12 | 5.56 | 3.53 | 0.56 | 1.95 | 2.03 | 4.30E-06 | 9.43E-05 | (A) NE (B/D) |
| MIR-4750/3P | 3.63 | 1.6 | 0.83 | 1.25 | 2.03 | 1.49E-05 | 2.49E-04 | (A) NE (B/D) |
| MIR-4750/47 | 3.63 | 1.6 | 0.83 | 1.25 | 2.03 | 1.49E-05 | 2.49E-04 | (A) NE (B/D) |
| MIR-4750/5P | 3.63 | 1.6 | 0.83 | 1.25 | 2.03 | 1.49E-05 | 2.49E-04 | (A) NE (B/D) |
| CLASRP | 2.47 | 0.43 | 0.95 | 0.55 | 2.03 | 1.31E-04 | 1.33E-03 | (A) NE (B/D) |
| HMGN2P6 | 8.32 | 6.28 | 0.91 | 2.2 | 2.03 | 1.98E-04 | 1.83E-03 | (A) NE (B/D) |
| PLXND1 | 2.43 | 0.4 | 1.01 | 0.37 | 2.03 | 2.64E-04 | 2.28E-03 | (A) NE (B/D) |
| STRC | 3.78 | 1.75 | 1.24 | 1.11 | 2.03 | 8.12E-04 | 5.43E-03 | (A) NE (B/D) |
| RGS3 | 2.73 | 0.7 | 0.49 | 0.79 | 2.02 | 4.89E-09 | 4.59E-07 | (A) NE (B/D) |
| TEN1-CDK3 | 8.25 | 6.24 | 0.54 | 1.32 | 2.02 | 5.91E-08 | 3.15E-06 | (A) NE (B/D) |
| NR2F1 | 3.73 | 1.71 | 0.66 | 1.32 | 2.02 | 8.33E-07 | 2.58E-05 | (A) NE (B/D) |
| RAB4B | 7.75 | 5.72 | 0.64 | 1.55 | 2.02 | 1.20E-06 | 3.42E-05 | (A) NE (B/D) |
| TIMM29 | 8.1 | 6.08 | 0.65 | 0.68 | 2.02 | 1.51E-06 | 4.15E-05 | (A) NE (B/D) |
| SEPTIN5 | 4.74 | 2.72 | 0.81 | 1.02 | 2.02 | 1.31E-05 | 2.26E-04 | (A) NE (B/D) |
| NPHP4 | 3.11 | 1.09 | 0.75 | 0.63 | 2.02 | 1.32E-05 | 2.28E-04 | (A) NE (B/D) |
| NTHL1 | 2.81 | 0.79 | 0.84 | 0.93 | 2.02 | 2.22E-05 | 3.32E-04 | (A) NE (B/D) |
| KHDC1 | 2.59 | 0.57 | 0.88 | 0.84 | 2.02 | 4.42E-05 | 5.74E-04 | (A) NE (B/D) |
| GATAD1 | 3.91 | 1.89 | 0.98 | 1.08 | 2.02 | 1.04E-04 | 1.12E-03 | (A) NE (B/D) |
| RPL37AP1 | 8 | 5.97 | 1.53 | 1.4 | 2.02 | 3.79E-03 | 0.0174 | (A) NE (B/D) |

| | | | | | | | | |
|---|---|---|---|---|---|---|---|---|
| RNA5SP271 | 2.47 | 0.45 | 1.98 | 1.13 | 2.02 | 0.0157 | 0.0496 | (A) NE (B/D) |
| C19ORF57 | 2.52 | 0.51 | 0.55 | 0.48 | 2.01 | 5.60E-07 | 1.89E-05 | (A) NE (B/D) |
| CCNL2 | 8.66 | 6.65 | 0.69 | 1.22 | 2.01 | 1.31E-06 | 3.68E-05 | (A) NE (B/D) |
| SNORD62B | 6.95 | 4.94 | 0.71 | 0.85 | 2.01 | 3.22E-06 | 7.45E-05 | (A) NE (B/D) |
| GLS2 | 2.73 | 0.72 | 0.68 | 0.67 | 2.01 | 3.49E-06 | 7.97E-05 | (A) NE (B/D) |
| ARTN | 2.83 | 0.82 | 0.73 | 0.9 | 2.01 | 3.94E-06 | 8.76E-05 | (A) NE (B/D) |
| NUDT14 | 8.09 | 6.07 | 0.75 | 1.5 | 2.01 | 6.10E-06 | 1.23E-04 | (A) NE (B/D) |
| CIT | 2.24 | 0.23 | 0.85 | 0.61 | 2.01 | 4.90E-05 | 6.21E-04 | (A) NE (B/D) |
| ARMCX2 | 4.41 | 2.4 | 0.89 | 1.78 | 2.01 | 6.44E-05 | 7.70E-04 | (A) NE (B/D) |
| MTFR2 | 5.95 | 3.94 | 1 | 1.6 | 2.01 | 1.46E-04 | 1.45E-03 | (A) NE (B/D) |
| FABP5 | 2.18 | 0.17 | 1.26 | 0.54 | 2.01 | 1.27E-03 | 7.59E-03 | (A) NE (B/D) |
| CUL9 | 2.85 | 0.85 | 0.38 | 0.81 | 2 | 2.12E-11 | 7.09E-09 | (A) NE (B/D) |
| KLHDC3 | 3.48 | 1.48 | 0.31 | 1.33 | 2 | 6.61E-10 | 1.05E-07 | (A) NE (B/D) |
| MAGEF1 | 3.28 | 1.28 | 0.41 | 1.34 | 2 | 4.18E-09 | 4.04E-07 | (A) NE (B/D) |
| CYP4F59P | 4.99 | 2.98 | 0.58 | 1.58 | 2 | 6.70E-07 | 2.16E-05 | (A) NE (B/D) |
| PPP2R3B | 3.3 | 1.3 | 0.72 | 0.79 | 2 | 5.38E-06 | 1.12E-04 | (A) NE (B/D) |
| PTOV1 | 5.44 | 3.44 | 0.77 | 1.07 | 2 | 7.63E-06 | 1.47E-04 | (A) NE (B/D) |
| ARHGEF2 | 4.33 | 2.33 | 0.87 | 0.98 | 2 | 3.66E-05 | 4.95E-04 | (A) NE (B/D) |
| PLXDC1 | 5.06 | 3.07 | 0.89 | 1.9 | 2 | 9.30E-05 | 1.03E-03 | (A) NE (B/D) |
| CEACAM19 | 4.84 | 2.83 | 1.05 | 1.52 | 2 | 2.30E-04 | 2.05E-03 | (A) NE (B/D) |
| PCDHGA1 | 8.95 | 6.94 | 1.3 | 1.56 | 2 | 1.38E-03 | 8.06E-03 | (A) NE (B/D) |
| CEP152 | 2.11 | 0.11 | 1.65 | 0.39 | 2 | 6.57E-03 | 0.0263 | (A) NE (B/D) |
| MIR-23B/23B | 3.35 | 1.34 | 1.79 | 1.68 | 2 | 0.0105 | 0.0368 | (A) NE (B/D) |
| MIR-23B/23B | 3.35 | 1.34 | 1.79 | 1.68 | 2 | 0.0105 | 0.0368 | (A) NE (B/D) |
| MIR-23B/3P | 3.35 | 1.34 | 1.79 | 1.68 | 2 | 0.0105 | 0.0368 | (A) NE (B/D) |
| MIR-23B/5P | 3.35 | 1.34 | 1.79 | 1.68 | 2 | 0.0105 | 0.0368 | (A) NE (B/D) |
| SAMD1 | 4.77 | 2.78 | 0.21 | 1.15 | 1.99 | 1.02E-11 | 3.98E-09 | (A) NE (B/D) |
| TRIR | 3.8 | 1.81 | 0.21 | 1.17 | 1.99 | 1.49E-11 | 5.40E-09 | (A) NE (B/D) |
| STRN4 | 4.6 | 2.61 | 0.37 | 0.96 | 1.99 | 2.24E-11 | 7.21E-09 | (A) NE (B/D) |
| SNHG6 | 9.82 | 7.83 | 0.27 | 1.49 | 1.99 | 7.13E-09 | 6.04E-07 | (A) NE (B/D) |
| PHC2 | 4.38 | 2.39 | 0.5 | 1 | 1.99 | 8.03E-09 | 6.68E-07 | (A) NE (B/D) |
| CCDC28B | 4.36 | 2.36 | 0.49 | 1.17 | 1.99 | 8.35E-09 | 6.82E-07 | (A) NE (B/D) |
| H2AFV | 4.83 | 2.85 | 0.44 | 1.42 | 1.99 | 2.01E-08 | 1.33E-06 | (A) NE (B/D) |
| SNRPD2 | 6.13 | 4.14 | 0.36 | 1.91 | 1.99 | 1.23E-06 | 3.49E-05 | (A) NE (B/D) |
| TRAF1 | 5.95 | 3.96 | 0.84 | 1.26 | 1.99 | 2.19E-05 | 3.28E-04 | (A) NE (B/D) |
| ISYNA1 | 2.15 | 0.16 | 0.89 | 0.43 | 1.99 | 1.14E-04 | 1.20E-03 | (A) NE (B/D) |
| SLC22A23 | 5.79 | 3.8 | 1.11 | 0.95 | 1.99 | 4.20E-04 | 3.28E-03 | (A) NE (B/D) |
| RNA5SP392 | 6.95 | 4.95 | 0.63 | 2.9 | 1.99 | 6.63E-04 | 4.65E-03 | (A) NE (B/D) |
| REC8 | 7.3 | 5.31 | 1.26 | 1.5 | 1.99 | 1.13E-03 | 6.91E-03 | (A) NE (B/D) |
| HNRNPA3P5 | 8.2 | 6.21 | 1.15 | 2.31 | 1.99 | 1.27E-03 | 7.59E-03 | (A) NE (B/D) |
| RPL7AP26 | 2.59 | 0.6 | 1.32 | 0.71 | 1.99 | 1.71E-03 | 9.51E-03 | (A) NE (B/D) |
| RPL12P37 | 2.34 | 0.36 | 1.44 | 0.64 | 1.99 | 3.02E-03 | 0.0147 | (A) NE (B/D) |
| LOXL4 | 2.31 | 0.33 | 1.63 | 0.87 | 1.99 | 6.20E-03 | 0.0253 | (A) NE (B/D) |
| AP4M1 | 5.51 | 3.53 | 0.16 | 1.17 | 1.98 | 1.69E-11 | 5.88E-09 | (A) NE (B/D) |
| ZMIZ2 | 3.57 | 1.6 | 0.27 | 1.17 | 1.98 | 3.25E-11 | 9.50E-09 | (A) NE (B/D) |
| HID1 | 2.85 | 0.87 | 0.35 | 1.06 | 1.98 | 3.51E-11 | 9.80E-09 | (A) NE (B/D) |
| GTF3C5 | 6.37 | 4.39 | 0.12 | 1.24 | 1.98 | 7.58E-11 | 1.86E-08 | (A) NE (B/D) |
| PLXNB1 | 2.55 | 0.56 | 0.45 | 0.84 | 1.98 | 1.10E-09 | 1.45E-07 | (A) NE (B/D) |
| MIR-1233-2/ | 2.78 | 0.8 | 0.47 | 1.02 | 1.98 | 2.66E-09 | 2.98E-07 | (A) NE (B/D) |
| MIR-1233-2/ | 2.78 | 0.8 | 0.47 | 1.02 | 1.98 | 2.66E-09 | 2.98E-07 | (A) NE (B/D) |
| MIR-1233-2/ | 2.78 | 0.8 | 0.47 | 1.02 | 1.98 | 2.66E-09 | 2.98E-07 | (A) NE (B/D) |
| EGFL8 | 7.62 | 5.64 | 0.61 | 1.63 | 1.98 | 1.64E-06 | 4.40E-05 | (A) NE (B/D) |
| SLCO3A1 | 2.03 | 0.05 | 1.11 | 0.21 | 1.98 | 6.59E-04 | 4.63E-03 | (A) NE (B/D) |
| SLC25A27 | 6.48 | 4.5 | 1.47 | 1.69 | 1.98 | 3.72E-03 | 0.0172 | (A) NE (B/D) |
| LINC00618 | 4.13 | 2.15 | 1.81 | 1.47 | 1.98 | 0.0113 | 0.039 | (A) NE (B/D) |
| LIN28B-AS1 | 2.33 | 0.35 | 1.83 | 0.71 | 1.98 | 0.0118 | 0.0401 | (A) NE (B/D) |
| BRAT1 | 3.55 | 1.58 | 0.34 | 0.82 | 1.97 | 1.53E-12 | 9.39E-10 | (A) NE (B/D) |
| NT5C | 4.42 | 2.46 | 0.6 | 1.14 | 1.97 | 1.94E-07 | 8.18E-06 | (A) NE (B/D) |
| HNRNPA1P10 | 6.24 | 4.26 | 0.65 | 1.18 | 1.97 | 8.36E-07 | 2.58E-05 | (A) NE (B/D) |
| YJEFN3 | 9.91 | 7.94 | 0.62 | 1.52 | 1.97 | 1.12E-06 | 3.25E-05 | (A) NE (B/D) |
| MAP3K10 | 2.21 | 0.24 | 0.55 | 0.37 | 1.97 | 1.35E-06 | 3.79E-05 | (A) NE (B/D) |
| ATP5F1D | 4.72 | 2.75 | 0.69 | 1.28 | 1.97 | 1.99E-06 | 5.12E-05 | (A) NE (B/D) |
| ARF4P2 | 3.08 | 1.11 | 0.72 | 1.07 | 1.97 | 3.65E-06 | 8.27E-05 | (A) NE (B/D) |
| DDX11 | 3.71 | 1.73 | 0.7 | 0.71 | 1.97 | 5.17E-06 | 1.09E-04 | (A) NE (B/D) |
| ADGRL1 | 9.27 | 7.3 | 0.81 | 1.04 | 1.97 | 1.70E-05 | 2.75E-04 | (A) NE (B/D) |

| | | | | | | | | |
|---|---|---|---|---|---|---|---|---|
| RNU4-8P | 4.93 | 2.96 | 0.96 | 1.81 | 1.97 | 1.82E-04 | 1.71E-03 | (A) NE (B/D) |
| HOXC-AS2 | 2.39 | 0.42 | 1.23 | 0.69 | 1.97 | 1.11E-03 | 6.85E-03 | (A) NE (B/D) |
| RNU6-554P | 2.52 | 0.56 | 1.28 | 1.16 | 1.97 | 1.33E-03 | 7.84E-03 | (A) NE (B/D) |
| PCDHGA2 | 9.49 | 7.52 | 1.34 | 1.55 | 1.97 | 1.90E-03 | 0.0103 | (A) NE (B/D) |
| B3GALT2 | 2.08 | 0.11 | 1.37 | 0.23 | 1.97 | 2.47E-03 | 0.0126 | (A) NE (B/D) |
| GKN2 | 2.08 | 0.11 | 1.56 | 0.34 | 1.97 | 5.23E-03 | 0.0222 | (A) NE (B/D) |
| SLC17A4 | 2.67 | 0.7 | 1.65 | 1.15 | 1.97 | 7.11E-03 | 0.0279 | (A) NE (B/D) |
| FAM95B1 | 3.61 | 1.65 | 0.46 | 1.12 | 1.96 | 3.55E-09 | 3.61E-07 | (A) NE (B/D) |
| ODF2 | 8.47 | 6.51 | 0.68 | 0.98 | 1.96 | 1.90E-06 | 4.96E-05 | (A) NE (B/D) |
| LRRC73 | 3.98 | 2.02 | 0.71 | 1.11 | 1.96 | 3.13E-06 | 7.27E-05 | (A) NE (B/D) |
| ABR | 3.41 | 1.45 | 0.82 | 0.75 | 1.96 | 3.22E-05 | 4.50E-04 | (A) NE (B/D) |
| TMEM17 | 5.15 | 3.19 | 1.1 | 1.04 | 1.96 | 4.19E-04 | 3.27E-03 | (A) NE (B/D) |
| RPS14P3 | 6.05 | 4.08 | 1.34 | 1.59 | 1.96 | 2.07E-03 | 0.011 | (A) NE (B/D) |
| TCTE1 | 2.04 | 0.08 | 1.38 | 0.34 | 1.96 | 2.63E-03 | 0.0132 | (A) NE (B/D) |
| TP53TG3D | 2.11 | 0.16 | 1.6 | 0.4 | 1.96 | 6.25E-03 | 0.0254 | (A) NE (B/D) |
| PABPC1L | 4.48 | 2.53 | 0.47 | 1.27 | 1.95 | 1.51E-08 | 1.06E-06 | (A) NE (B/D) |
| KDF1 | 4.75 | 2.8 | 0.34 | 1.6 | 1.95 | 7.00E-08 | 3.62E-06 | (A) NE (B/D) |
| WDR83 | 3.05 | 1.1 | 0.63 | 1.19 | 1.95 | 6.03E-07 | 2.00E-05 | (A) NE (B/D) |
| TUBB4B | 11.65 | 9.71 | 0.59 | 1.71 | 1.95 | 2.65E-06 | 6.36E-05 | (A) NE (B/D) |
| SCNN1D | 5.88 | 3.93 | 0.74 | 1.46 | 1.95 | 7.58E-06 | 1.47E-04 | (A) NE (B/D) |
| ALG1L | 2.72 | 0.77 | 0.75 | 1.4 | 1.95 | 8.62E-06 | 1.62E-04 | (A) NE (B/D) |
| RPL12P1 | 4.49 | 2.54 | 1.1 | 1.31 | 1.95 | 4.30E-04 | 3.33E-03 | (A) NE (B/D) |
| TFCP2L1 | 2.41 | 0.45 | 1.1 | 0.95 | 1.95 | 4.52E-04 | 3.46E-03 | (A) NE (B/D) |
| ATP2A3 | 2.15 | 0.21 | 1.25 | 0.45 | 1.95 | 1.46E-03 | 8.42E-03 | (A) NE (B/D) |
| STAG3 | 2.48 | 0.54 | 1.29 | 0.64 | 1.95 | 1.69E-03 | 9.42E-03 | (A) NE (B/D) |
| MIR-3064/3P | 8.64 | 6.69 | 1.45 | 1.44 | 1.95 | 3.45E-03 | 0.0163 | (A) NE (B/D) |
| MIR-3064/5P | 8.64 | 6.69 | 1.45 | 1.44 | 1.95 | 3.45E-03 | 0.0163 | (A) NE (B/D) |
| AGT | 5.74 | 3.79 | 1.55 | 2.52 | 1.95 | 8.75E-03 | 0.0323 | (A) NE (B/D) |
| SMAD9 | 3.01 | 1.06 | 1.78 | 1 | 1.95 | 0.0111 | 0.0384 | (A) NE (B/D) |
| BORCS8-MEF | 4.41 | 2.47 | 0.35 | 1.27 | 1.94 | 1.12E-09 | 1.47E-07 | (A) NE (B/D) |
| TMEM30B | 3.61 | 1.66 | 0.26 | 1.39 | 1.94 | 2.72E-09 | 2.99E-07 | (A) NE (B/D) |
| PCDHB5 | 2.52 | 0.58 | 0.52 | 0.86 | 1.94 | 3.00E-08 | 1.85E-06 | (A) NE (B/D) |
| COMMD4 | 3.43 | 1.49 | 0.55 | 1.24 | 1.94 | 1.07E-07 | 5.05E-06 | (A) NE (B/D) |
| STARD10 | 5.23 | 3.29 | 0.73 | 1.31 | 1.94 | 5.83E-06 | 1.19E-04 | (A) NE (B/D) |
| MIR-4690/3P | 5.13 | 3.19 | 0.72 | 1.67 | 1.94 | 1.18E-05 | 2.08E-04 | (A) NE (B/D) |
| MIR-4690/5P | 5.13 | 3.19 | 0.72 | 1.67 | 1.94 | 1.18E-05 | 2.08E-04 | (A) NE (B/D) |
| NPDC1 | 7.61 | 5.68 | 0.79 | 1.57 | 1.94 | 2.07E-05 | 3.16E-04 | (A) NE (B/D) |
| PPM1E | 4.69 | 2.75 | 0.74 | 1.83 | 1.94 | 2.55E-05 | 3.71E-04 | (A) NE (B/D) |
| NOS1AP | 5.79 | 3.84 | 1.1 | 1.46 | 1.94 | 4.36E-04 | 3.36E-03 | (A) NE (B/D) |
| KIF5C | 4.87 | 2.92 | 1.1 | 1.61 | 1.94 | 4.75E-04 | 3.59E-03 | (A) NE (B/D) |
| RN7SL622P | 2.72 | 0.78 | 1.29 | 1.33 | 1.94 | 1.59E-03 | 9.02E-03 | (A) NE (B/D) |
| PHACTR3 | 2.81 | 0.87 | 1.51 | 1.31 | 1.94 | 4.64E-03 | 0.0203 | (A) NE (B/D) |
| CDCA2 | 7 | 5.06 | 1.55 | 1.8 | 1.94 | 6.05E-03 | 0.0248 | (A) NE (B/D) |
| SYMPK | 5.89 | 3.95 | 0.47 | 0.94 | 1.93 | 3.38E-09 | 3.48E-07 | (A) NE (B/D) |
| ZBED6CL | 3.17 | 1.24 | 0.5 | 0.81 | 1.93 | 1.68E-08 | 1.16E-06 | (A) NE (B/D) |
| PCNA | 9.39 | 7.46 | 0.64 | 1.68 | 1.93 | 5.21E-06 | 1.09E-04 | (A) NE (B/D) |
| ZC3HAV1L | 3.28 | 1.35 | 0.72 | 0.8 | 1.93 | 7.57E-06 | 1.47E-04 | (A) NE (B/D) |
| CD99L2 | 2.99 | 1.05 | 1.12 | 1.16 | 1.93 | 5.27E-04 | 3.88E-03 | (A) NE (B/D) |
| HMGN2P3 | 10.96 | 9.03 | 1.05 | 2.01 | 1.93 | 5.90E-04 | 4.25E-03 | (A) NE (B/D) |
| FABP5P11 | 2.7 | 0.77 | 1.15 | 1.11 | 1.93 | 6.71E-04 | 4.69E-03 | (A) NE (B/D) |
| MYEF2 | 2.62 | 0.7 | 1.16 | 0.67 | 1.93 | 8.83E-04 | 5.76E-03 | (A) NE (B/D) |
| DIPK1B | 2.7 | 0.77 | 1.2 | 1 | 1.93 | 1.01E-03 | 6.34E-03 | (A) NE (B/D) |
| ACBD7 | 2.24 | 0.31 | 1.22 | 0.66 | 1.93 | 1.22E-03 | 7.36E-03 | (A) NE (B/D) |
| TMPRSS7 | 2.41 | 0.48 | 1.27 | 0.67 | 1.93 | 1.64E-03 | 9.21E-03 | (A) NE (B/D) |
| ZNF835 | 3.71 | 1.78 | 1.35 | 1.17 | 1.93 | 2.27E-03 | 0.0118 | (A) NE (B/D) |
| RPS4XP7 | 4.11 | 2.18 | 1.42 | 1.59 | 1.93 | 3.36E-03 | 0.0159 | (A) NE (B/D) |
| STARD9 | 2.9 | 0.97 | 1.64 | 0.92 | 1.93 | 7.69E-03 | 0.0294 | (A) NE (B/D) |
| DTYMK | 4 | 2.08 | 0.42 | 1.14 | 1.92 | 1.53E-09 | 1.88E-07 | (A) NE (B/D) |
| MDC1 | 6.78 | 4.87 | 0.42 | 1.16 | 1.92 | 2.31E-09 | 2.69E-07 | (A) NE (B/D) |
| STK25 | 3.75 | 1.83 | 0.48 | 1.24 | 1.92 | 2.02E-08 | 1.34E-06 | (A) NE (B/D) |
| BCAT2 | 4.61 | 2.7 | 0.54 | 1.54 | 1.92 | 5.82E-07 | 1.94E-05 | (A) NE (B/D) |
| ARVCF | 8.89 | 6.97 | 0.67 | 1.14 | 1.92 | 1.80E-06 | 4.75E-05 | (A) NE (B/D) |
| SPINT2 | 3.85 | 1.93 | 0.78 | 1.36 | 1.92 | 1.55E-05 | 2.56E-04 | (A) NE (B/D) |
| NASP | 3.3 | 1.38 | 0.79 | 1.21 | 1.92 | 1.67E-05 | 2.69E-04 | (A) NE (B/D) |
| POMT1 | 4.21 | 2.29 | 0.81 | 1.05 | 1.92 | 2.16E-05 | 3.25E-04 | (A) NE (B/D) |

| | | | | | | | | |
|---|---|---|---|---|---|---|---|---|
| NR6A1 | 2.18 | 0.26 | 0.74 | 0.42 | 1.92 | 3.21E-05 | 4.49E-04 | (A) NE (B/D) |
| PAK6 | 2.3 | 0.38 | 0.91 | 0.69 | 1.92 | 1.21E-04 | 1.25E-03 | (A) NE (B/D) |
| ABTB1 | 4.29 | 2.37 | 0.92 | 1.67 | 1.92 | 1.31E-04 | 1.33E-03 | (A) NE (B/D) |
| LINC00680 | 4.72 | 2.81 | 1 | 0.99 | 1.92 | 2.21E-04 | 1.98E-03 | (A) NE (B/D) |
| PLCD4 | 2.49 | 0.56 | 1.17 | 0.63 | 1.92 | 9.84E-04 | 6.24E-03 | (A) NE (B/D) |
| CHD7 | 3.29 | 1.37 | 1.25 | 0.79 | 1.92 | 1.46E-03 | 8.42E-03 | (A) NE (B/D) |
| DNAJC6 | 1.97 | 0.05 | 1.24 | 0.22 | 1.92 | 1.67E-03 | 9.34E-03 | (A) NE (B/D) |
| COL6A4P1 | 1.99 | 0.07 | 1.39 | 0.28 | 1.92 | 3.10E-03 | 0.015 | (A) NE (B/D) |
| PCDH9-AS1 | 2.26 | 0.34 | 1.52 | 0.69 | 1.92 | 5.20E-03 | 0.0221 | (A) NE (B/D) |
| RASSF10 | 3.37 | 1.45 | 1.57 | 1.69 | 1.92 | 6.65E-03 | 0.0265 | (A) NE (B/D) |
| SIX3-AS1 | 1.94 | 0.02 | 1.81 | 0.11 | 1.92 | 0.013 | 0.0431 | (A) NE (B/D) |
| ZNF540 | 5.72 | 3.8 | 1.87 | 0.99 | 1.92 | 0.0156 | 0.0493 | (A) NE (B/D) |
| HNRNPA1P4 | 4.47 | 2.56 | 0.44 | 1.66 | 1.91 | 5.15E-07 | 1.76E-05 | (A) NE (B/D) |
| NDUFS5 | 6.06 | 4.14 | 0.51 | 1.63 | 1.91 | 7.43E-07 | 2.35E-05 | (A) NE (B/D) |
| C9ORF40 | 2.76 | 0.84 | 0.66 | 1.16 | 1.91 | 1.62E-06 | 4.37E-05 | (A) NE (B/D) |
| RPS16 | 3.58 | 1.67 | 0.56 | 1.67 | 1.91 | 1.95E-06 | 5.04E-05 | (A) NE (B/D) |
| TPX2 | 2.87 | 0.96 | 0.81 | 1 | 1.91 | 2.34E-05 | 3.45E-04 | (A) NE (B/D) |
| MIR-25/25 | 8.19 | 6.28 | 0.42 | 2.23 | 1.91 | 3.28E-05 | 4.55E-04 | (A) NE (B/D) |
| MIR-25/25* | 8.19 | 6.28 | 0.42 | 2.23 | 1.91 | 3.28E-05 | 4.55E-04 | (A) NE (B/D) |
| MIR-25/3P | 8.19 | 6.28 | 0.42 | 2.23 | 1.91 | 3.28E-05 | 4.55E-04 | (A) NE (B/D) |
| MIR-25/5P | 8.19 | 6.28 | 0.42 | 2.23 | 1.91 | 3.28E-05 | 4.55E-04 | (A) NE (B/D) |
| APOD | 3.09 | 1.18 | 0.91 | 1.31 | 1.91 | 8.73E-05 | 9.79E-04 | (A) NE (B/D) |
| PRDM8 | 4.76 | 2.85 | 1.54 | 1.71 | 1.91 | 6.23E-03 | 0.0253 | (A) NE (B/D) |
| PFKP | 9.03 | 7.12 | 0.45 | 1.28 | 1.9 | 1.64E-08 | 1.15E-06 | (A) NE (B/D) |
| CUTA | 8.06 | 6.16 | 0.31 | 1.56 | 1.9 | 6.14E-08 | 3.24E-06 | (A) NE (B/D) |
| ATG4B | 7.49 | 5.59 | 0.54 | 0.99 | 1.9 | 6.74E-08 | 3.50E-06 | (A) NE (B/D) |
| RACGAP1 | 3.59 | 1.69 | 0.79 | 1.4 | 1.9 | 2.14E-05 | 3.23E-04 | (A) NE (B/D) |
| GAB2 | 6.45 | 4.56 | 0.86 | 0.84 | 1.9 | 6.29E-05 | 7.57E-04 | (A) NE (B/D) |
| PDZD4 | 2.18 | 0.28 | 0.84 | 0.49 | 1.9 | 8.67E-05 | 9.74E-04 | (A) NE (B/D) |
| PLK4 | 3.62 | 1.73 | 1.09 | 1.36 | 1.9 | 5.13E-04 | 3.81E-03 | (A) NE (B/D) |
| LONRF2 | 4.19 | 2.29 | 1.18 | 1.29 | 1.9 | 9.63E-04 | 6.15E-03 | (A) NE (B/D) |
| FAM111B | 6.72 | 4.82 | 1.21 | 1.91 | 1.9 | 1.62E-03 | 9.12E-03 | (A) NE (B/D) |
| RNA5SP118 | 2.13 | 0.23 | 1.57 | 0.47 | 1.9 | 6.65E-03 | 0.0265 | (A) NE (B/D) |
| SDK2 | 3.28 | 1.38 | 1.59 | 1.14 | 1.9 | 7.12E-03 | 0.0279 | (A) NE (B/D) |
| RMST | 2.72 | 0.82 | 1.69 | 1.05 | 1.9 | 9.66E-03 | 0.0348 | (A) NE (B/D) |
| MIR-1180/11 | 3.43 | 1.53 | 1.79 | 1.8 | 1.9 | 0.0147 | 0.0472 | (A) NE (B/D) |
| MIR-1180/3P | 3.43 | 1.53 | 1.79 | 1.8 | 1.9 | 0.0147 | 0.0472 | (A) NE (B/D) |
| MIR-1180/5P | 3.43 | 1.53 | 1.79 | 1.8 | 1.9 | 0.0147 | 0.0472 | (A) NE (B/D) |
| GOLGA6L10 | 4.26 | 2.37 | 0.41 | 0.93 | 1.89 | 4.94E-10 | 8.58E-08 | (A) NE (B/D) |
| RPP38-DT | 5.89 | 4.01 | 0.23 | 1.33 | 1.89 | 1.53E-09 | 1.88E-07 | (A) NE (B/D) |
| VEGFB | 10.2 | 8.31 | 0.29 | 1.46 | 1.89 | 1.66E-08 | 1.15E-06 | (A) NE (B/D) |
| MRPL34 | 2.32 | 0.43 | 0.55 | 0.58 | 1.89 | 4.53E-07 | 1.59E-05 | (A) NE (B/D) |
| PPIE | 2.91 | 1.02 | 0.69 | 1.07 | 1.89 | 3.21E-06 | 7.43E-05 | (A) NE (B/D) |
| HNRNPA1P27 | 4.41 | 2.52 | 0.71 | 1.31 | 1.89 | 5.45E-06 | 1.13E-04 | (A) NE (B/D) |
| C17ORF100 | 5.41 | 3.52 | 0.85 | 0.9 | 1.89 | 5.35E-05 | 6.66E-04 | (A) NE (B/D) |
| PAXBP1-AS1 | 4.96 | 3.07 | 0.89 | 0.59 | 1.89 | 1.24E-04 | 1.28E-03 | (A) NE (B/D) |
| C1QTNF3 | 3.23 | 1.35 | 0.93 | 1.71 | 1.89 | 1.78E-04 | 1.68E-03 | (A) NE (B/D) |
| TENT5A | 2.32 | 0.43 | 0.93 | 0.58 | 1.89 | 1.99E-04 | 1.83E-03 | (A) NE (B/D) |
| ELF3 | 6.28 | 4.39 | 1.16 | 1.78 | 1.89 | 1.10E-03 | 6.77E-03 | (A) NE (B/D) |
| SCHIP1 | 3.86 | 1.97 | 1.49 | 1.35 | 1.89 | 5.01E-03 | 0.0215 | (A) NE (B/D) |
| MIR-27B/27B | 3.09 | 1.2 | 1.53 | 1.72 | 1.89 | 6.34E-03 | 0.0256 | (A) NE (B/D) |
| MIR-27B/27B | 3.09 | 1.2 | 1.53 | 1.72 | 1.89 | 6.34E-03 | 0.0256 | (A) NE (B/D) |
| MIR-27B/3P | 3.09 | 1.2 | 1.53 | 1.72 | 1.89 | 6.34E-03 | 0.0256 | (A) NE (B/D) |
| MIR-27B/5P | 3.09 | 1.2 | 1.53 | 1.72 | 1.89 | 6.34E-03 | 0.0256 | (A) NE (B/D) |
| SF3A2 | 2.45 | 0.57 | 0.17 | 0.69 | 1.88 | 1.69E-17 | 8.70E-14 | (A) NE (B/D) |
| BABAM1 | 9.58 | 7.69 | 0.44 | 1.24 | 1.88 | 1.12E-08 | 8.73E-07 | (A) NE (B/D) |
| COPE | 3.37 | 1.48 | 0.45 | 1.36 | 1.88 | 4.29E-08 | 2.45E-06 | (A) NE (B/D) |
| PPT2 | 7.98 | 6.1 | 0.51 | 1.38 | 1.88 | 1.76E-07 | 7.60E-06 | (A) NE (B/D) |
| EMC10 | 10.17 | 8.29 | 0.6 | 1.01 | 1.88 | 4.17E-07 | 1.48E-05 | (A) NE (B/D) |
| SNHG32 | 6.85 | 4.97 | 0.59 | 1.32 | 1.88 | 6.30E-07 | 2.06E-05 | (A) NE (B/D) |
| HAPLN2 | 2.55 | 0.67 | 0.61 | 0.9 | 1.88 | 6.62E-07 | 2.13E-05 | (A) NE (B/D) |
| CD24P4 | 13.88 | 12.01 | 0.49 | 1.67 | 1.88 | 1.22E-06 | 3.47E-05 | (A) NE (B/D) |
| TOE1 | 6.14 | 4.25 | 0.64 | 1.23 | 1.88 | 1.32E-06 | 3.72E-05 | (A) NE (B/D) |
| ADORA2A-AS | 4.26 | 2.39 | 0.72 | 0.88 | 1.88 | 8.31E-06 | 1.57E-04 | (A) NE (B/D) |
| EXO5 | 6.78 | 4.91 | 0.76 | 1.29 | 1.88 | 1.39E-05 | 2.37E-04 | (A) NE (B/D) |

| | | | | | | | | |
|---|---|---|---|---|---|---|---|---|
| LARGE2 | 3.19 | 1.32 | 0.77 | 1.39 | 1.88 | 1.81E-05 | 2.86E-04 | (A) NE (B/D) |
| RNF31 | 3.45 | 1.57 | 0.8 | 1.44 | 1.88 | 2.91E-05 | 4.12E-04 | (A) NE (B/D) |
| UPF3A | 8.6 | 6.72 | 0.88 | 1.3 | 1.88 | 7.06E-05 | 8.28E-04 | (A) NE (B/D) |
| NT5M | 5.73 | 3.84 | 0.9 | 1.62 | 1.88 | 1.16E-04 | 1.21E-03 | (A) NE (B/D) |
| IVNS1ABP | 2.33 | 0.45 | 1.09 | 0.53 | 1.88 | 7.06E-04 | 4.88E-03 | (A) NE (B/D) |
| GCH1 | 8.37 | 6.49 | 1.16 | 1.05 | 1.88 | 8.88E-04 | 5.78E-03 | (A) NE (B/D) |
| MIR-3188/31 | 2.61 | 0.74 | 1.19 | 1.31 | 1.88 | 1.12E-03 | 6.86E-03 | (A) NE (B/D) |
| JAKMIP2 | 1.89 | 0.01 | 1.43 | 0.03 | 1.88 | 4.28E-03 | 0.0192 | (A) NE (B/D) |
| RANP4 | 3.58 | 1.7 | 1.63 | 0.96 | 1.88 | 8.47E-03 | 0.0316 | (A) NE (B/D) |
| COLGALT1 | 4.62 | 2.75 | 0.25 | 1.05 | 1.87 | 8.54E-12 | 3.54E-09 | (A) NE (B/D) |
| MIR-3934/39 | 5.77 | 3.91 | 0.53 | 1.15 | 1.87 | 7.69E-08 | 3.90E-06 | (A) NE (B/D) |
| MIR-3934/3P | 5.77 | 3.91 | 0.53 | 1.15 | 1.87 | 7.69E-08 | 3.90E-06 | (A) NE (B/D) |
| MIR-3934/5P | 5.77 | 3.91 | 0.53 | 1.15 | 1.87 | 7.69E-08 | 3.90E-06 | (A) NE (B/D) |
| PYCR2 | 5.06 | 3.2 | 0.31 | 1.61 | 1.87 | 1.70E-07 | 7.42E-06 | (A) NE (B/D) |
| FAAH2 | 5.85 | 3.98 | 0.65 | 1.32 | 1.87 | 2.19E-06 | 5.50E-05 | (A) NE (B/D) |
| TPPP | 7.57 | 5.71 | 0.69 | 0.83 | 1.87 | 5.53E-06 | 1.14E-04 | (A) NE (B/D) |
| MESTP1 | 4.62 | 2.75 | 0.76 | 1.23 | 1.87 | 1.35E-05 | 2.32E-04 | (A) NE (B/D) |
| GAPDHP1 | 5.06 | 3.18 | 0.89 | 1.5 | 1.87 | 9.65E-05 | 1.06E-03 | (A) NE (B/D) |
| FOXM1 | 2.74 | 0.87 | 0.97 | 1.18 | 1.87 | 1.89E-04 | 1.77E-03 | (A) NE (B/D) |
| PEX5L-AS1 | 2.43 | 0.56 | 1.06 | 0.8 | 1.87 | 5.17E-04 | 3.83E-03 | (A) NE (B/D) |
| DPYSL5 | 1.95 | 0.08 | 1.11 | 0.46 | 1.87 | 8.90E-04 | 5.79E-03 | (A) NE (B/D) |
| ETV5-AS1 | 3.42 | 1.55 | 1.33 | 1.34 | 1.87 | 2.50E-03 | 0.0127 | (A) NE (B/D) |
| NOVA1-AS1 | 2.21 | 0.34 | 1.53 | 0.6 | 1.87 | 6.30E-03 | 0.0255 | (A) NE (B/D) |
| RANBP3 | 2.29 | 0.43 | 0.33 | 0.5 | 1.86 | 5.35E-11 | 1.41E-08 | (A) NE (B/D) |
| CCDC124 | 4.28 | 2.41 | 0.43 | 1.17 | 1.86 | 5.78E-09 | 5.22E-07 | (A) NE (B/D) |
| H2AFZP3 | 4.91 | 3.05 | 0.55 | 1 | 1.86 | 1.20E-07 | 5.57E-06 | (A) NE (B/D) |
| RTN2 | 1.87 | 0.01 | 0.37 | 0.03 | 1.86 | 3.36E-07 | 1.25E-05 | (A) NE (B/D) |
| CENPM | 6.99 | 5.14 | 0.6 | 1.67 | 1.86 | 4.97E-06 | 1.05E-04 | (A) NE (B/D) |
| KIF21B | 3.01 | 1.14 | 0.77 | 1.31 | 1.86 | 1.82E-05 | 2.87E-04 | (A) NE (B/D) |
| ULBP2 | 2.47 | 0.61 | 0.8 | 1.07 | 1.86 | 2.73E-05 | 3.91E-04 | (A) NE (B/D) |
| NEURL4 | 7.09 | 5.23 | 0.88 | 1.09 | 1.86 | 7.98E-05 | 9.08E-04 | (A) NE (B/D) |
| TMEM169 | 2.43 | 0.58 | 0.87 | 0.72 | 1.86 | 1.06E-04 | 1.14E-03 | (A) NE (B/D) |
| AURKA | 2.16 | 0.3 | 1.14 | 0.72 | 1.86 | 9.52E-04 | 6.08E-03 | (A) NE (B/D) |
| RNU7-123P | 4.31 | 2.44 | 1.26 | 1.6 | 1.86 | 1.98E-03 | 0.0106 | (A) NE (B/D) |
| PCDHGA3 | 5.21 | 3.35 | 1.28 | 1.43 | 1.86 | 2.03E-03 | 0.0108 | (A) NE (B/D) |
| ZNF878 | 3.36 | 1.5 | 1.46 | 0.65 | 1.86 | 4.80E-03 | 0.0208 | (A) NE (B/D) |
| FAM90A20P | 3.52 | 1.67 | 1.53 | 1.39 | 1.86 | 6.51E-03 | 0.0261 | (A) NE (B/D) |
| TYK2 | 4.85 | 3.01 | 0.33 | 0.91 | 1.85 | 1.08E-11 | 4.08E-09 | (A) NE (B/D) |
| MIR-4292/42 | 4.16 | 2.31 | 0.35 | 0.79 | 1.85 | 1.49E-11 | 5.40E-09 | (A) NE (B/D) |
| TIMM50 | 3.94 | 2.08 | 0.47 | 1.02 | 1.85 | 1.10E-08 | 8.60E-07 | (A) NE (B/D) |
| USP20 | 8.66 | 6.81 | 0.55 | 1.17 | 1.85 | 1.68E-07 | 7.35E-06 | (A) NE (B/D) |
| SEMA4B | 8.68 | 6.83 | 0.65 | 0.91 | 1.85 | 2.18E-06 | 5.49E-05 | (A) NE (B/D) |
| DNASE2 | 3.33 | 1.47 | 0.65 | 1.5 | 1.85 | 4.93E-06 | 1.04E-04 | (A) NE (B/D) |
| STK11IP | 2.95 | 1.09 | 0.72 | 0.89 | 1.85 | 8.62E-06 | 1.62E-04 | (A) NE (B/D) |
| H2AFX | 9.64 | 7.79 | 0.71 | 1.48 | 1.85 | 1.19E-05 | 2.10E-04 | (A) NE (B/D) |
| FHP1 | 4.71 | 2.86 | 0.94 | 1.56 | 1.85 | 1.98E-04 | 1.83E-03 | (A) NE (B/D) |
| SH3BP4 | 3.42 | 1.56 | 0.98 | 1.21 | 1.85 | 2.27E-04 | 2.02E-03 | (A) NE (B/D) |
| CEP192 | 2.37 | 0.52 | 1.01 | 0.79 | 1.85 | 3.74E-04 | 3.00E-03 | (A) NE (B/D) |
| ALDH16A1 | 2.2 | 0.35 | 1.11 | 0.53 | 1.85 | 8.93E-04 | 5.80E-03 | (A) NE (B/D) |
| FUT2 | 4.54 | 2.68 | 1.16 | 0.98 | 1.85 | 1.01E-03 | 6.38E-03 | (A) NE (B/D) |
| PCDHGB1 | 9.42 | 7.57 | 1.23 | 1.53 | 1.85 | 1.65E-03 | 9.26E-03 | (A) NE (B/D) |
| PCDHGA7 | 8.22 | 6.37 | 1.25 | 1.54 | 1.85 | 1.93E-03 | 0.0104 | (A) NE (B/D) |
| CUTALP | 3.91 | 2.07 | 1.62 | 1.21 | 1.85 | 9.28E-03 | 0.0338 | (A) NE (B/D) |
| DEPDC1 | 2.74 | 0.89 | 1.65 | 1.03 | 1.85 | 9.77E-03 | 0.0351 | (A) NE (B/D) |
| GPI | 4.64 | 2.8 | 0.35 | 1.13 | 1.84 | 5.54E-10 | 9.25E-08 | (A) NE (B/D) |
| AGAP3 | 5.33 | 3.49 | 0.41 | 0.98 | 1.84 | 9.98E-10 | 1.36E-07 | (A) NE (B/D) |
| RABAC1 | 2.84 | 0.99 | 0.47 | 0.94 | 1.84 | 7.50E-09 | 6.31E-07 | (A) NE (B/D) |
| PRICKLE3 | 4.49 | 2.65 | 0.41 | 1.35 | 1.84 | 3.01E-08 | 1.85E-06 | (A) NE (B/D) |
| TMEM44 | 2.24 | 0.4 | 0.5 | 0.57 | 1.84 | 1.46E-07 | 6.53E-06 | (A) NE (B/D) |
| CPSF1 | 8.74 | 6.9 | 0.56 | 1.25 | 1.84 | 3.13E-07 | 1.18E-05 | (A) NE (B/D) |
| POMGNT1 | 7.11 | 5.27 | 0.6 | 1.16 | 1.84 | 7.60E-07 | 2.40E-05 | (A) NE (B/D) |
| HLA-T | 2.82 | 0.98 | 0.62 | 0.86 | 1.84 | 1.37E-06 | 3.83E-05 | (A) NE (B/D) |
| NPFF | 2.96 | 1.11 | 0.75 | 1.32 | 1.84 | 1.43E-05 | 2.42E-04 | (A) NE (B/D) |
| VPS37D | 3.08 | 1.25 | 0.84 | 1.15 | 1.84 | 5.55E-05 | 6.86E-04 | (A) NE (B/D) |
| PCDHB8 | 2.47 | 0.63 | 1.08 | 0.76 | 1.84 | 6.50E-04 | 4.58E-03 | (A) NE (B/D) |

| | | | | | | | | |
|---|---|---|---|---|---|---|---|---|
| PAM | 2.59 | 0.74 | 1.09 | 0.78 | 1.84 | 7.07E-04 | 4.89E-03 | (A) NE (B/D) |
| LINC01022 | 3.02 | 1.17 | 1.14 | 1.38 | 1.84 | 9.14E-04 | 5.91E-03 | (A) NE (B/D) |
| PCDHGA6 | 6.71 | 4.86 | 1.23 | 1.54 | 1.84 | 1.79E-03 | 9.87E-03 | (A) NE (B/D) |
| NUP62CL | 4.71 | 2.87 | 1.3 | 1.41 | 1.84 | 2.43E-03 | 0.0124 | (A) NE (B/D) |
| TCP11X2 | 1.89 | 0.06 | 1.43 | 0.2 | 1.84 | 4.81E-03 | 0.0209 | (A) NE (B/D) |
| VCX3A | 2.14 | 0.3 | 1.75 | 0.87 | 1.84 | 0.0135 | 0.0444 | (A) NE (B/D) |
| POLRMT | 7.83 | 6 | 0.3 | 0.94 | 1.83 | 5.04E-12 | 2.36E-09 | (A) NE (B/D) |
| CCDC130 | 8.97 | 7.14 | 0.35 | 1.01 | 1.83 | 1.35E-10 | 3.13E-08 | (A) NE (B/D) |
| HNRNPL | 4.7 | 2.87 | 0.57 | 1.41 | 1.83 | 1.12E-06 | 3.25E-05 | (A) NE (B/D) |
| ATP6V0B | 4.39 | 2.56 | 0.56 | 1.47 | 1.83 | 1.35E-06 | 3.78E-05 | (A) NE (B/D) |
| MIR-1238/12 | 3.33 | 1.5 | 0.66 | 1.12 | 1.83 | 2.63E-06 | 6.33E-05 | (A) NE (B/D) |
| MIR-1238/3P | 3.33 | 1.5 | 0.66 | 1.12 | 1.83 | 2.63E-06 | 6.33E-05 | (A) NE (B/D) |
| MIR-1238/5P | 3.33 | 1.5 | 0.66 | 1.12 | 1.83 | 2.63E-06 | 6.33E-05 | (A) NE (B/D) |
| SLFNL1 | 4.21 | 2.38 | 0.73 | 0.71 | 1.83 | 1.93E-05 | 2.99E-04 | (A) NE (B/D) |
| CAPS | 3.03 | 1.2 | 0.92 | 1.46 | 1.83 | 1.72E-04 | 1.64E-03 | (A) NE (B/D) |
| TMEM14A | 3.14 | 1.32 | 0.94 | 1.41 | 1.83 | 1.97E-04 | 1.82E-03 | (A) NE (B/D) |
| RNA5SP203 | 10.5 | 8.66 | 0.66 | 2.53 | 1.83 | 5.22E-04 | 3.86E-03 | (A) NE (B/D) |
| ZNF700 | 2.87 | 1.04 | 1.13 | 0.93 | 1.83 | 9.20E-04 | 5.94E-03 | (A) NE (B/D) |
| C10ORF62 | 2.73 | 0.9 | 1.18 | 1.13 | 1.83 | 1.22E-03 | 7.35E-03 | (A) NE (B/D) |
| PCDHGC3 | 4.89 | 3.06 | 1.26 | 1.49 | 1.83 | 2.13E-03 | 0.0112 | (A) NE (B/D) |
| NR4A2 | 6.96 | 5.13 | 1.55 | 1.37 | 1.83 | 7.64E-03 | 0.0293 | (A) NE (B/D) |
| PCDHGC4 | 2.98 | 1.15 | 1.59 | 1.26 | 1.83 | 8.59E-03 | 0.0319 | (A) NE (B/D) |
| KHSRP | 6.06 | 4.24 | 0.15 | 0.95 | 1.82 | 5.47E-13 | 5.21E-10 | (A) NE (B/D) |
| PIN1 | 5.18 | 3.36 | 0.32 | 1.12 | 1.82 | 3.10E-10 | 5.79E-08 | (A) NE (B/D) |
| HNRNPM | 4.96 | 3.14 | 0.26 | 1.24 | 1.82 | 9.95E-10 | 1.36E-07 | (A) NE (B/D) |
| DPH7 | 8.37 | 6.55 | 0.44 | 1.2 | 1.82 | 1.82E-08 | 1.24E-06 | (A) NE (B/D) |
| TMEM191A | 3.27 | 1.45 | 0.5 | 0.94 | 1.82 | 4.21E-08 | 2.42E-06 | (A) NE (B/D) |
| TBCB | 2.4 | 0.58 | 0.51 | 0.75 | 1.82 | 8.68E-08 | 4.29E-06 | (A) NE (B/D) |
| ENDOG | 8.85 | 7.03 | 0.64 | 1.29 | 1.82 | 2.75E-06 | 6.56E-05 | (A) NE (B/D) |
| ILF3-DT | 3.53 | 1.71 | 0.81 | 1.2 | 1.82 | 4.22E-05 | 5.54E-04 | (A) NE (B/D) |
| RN7SL573P | 5.36 | 3.54 | 0.9 | 1.21 | 1.82 | 1.19E-04 | 1.24E-03 | (A) NE (B/D) |
| CRTC1 | 1.84 | 0.02 | 0.94 | 0.11 | 1.82 | 4.06E-04 | 3.19E-03 | (A) NE (B/D) |
| L3MBTL1 | 4.88 | 3.06 | 1.04 | 0.8 | 1.82 | 5.20E-04 | 3.85E-03 | (A) NE (B/D) |
| LRRC10B | 3.15 | 1.33 | 1.06 | 1.33 | 1.82 | 5.72E-04 | 4.16E-03 | (A) NE (B/D) |
| STAG3L5P | 5.49 | 3.67 | 1.07 | 0.8 | 1.82 | 6.88E-04 | 4.79E-03 | (A) NE (B/D) |
| PCDHGB4 | 7.93 | 6.11 | 1.21 | 1.54 | 1.82 | 1.77E-03 | 9.76E-03 | (A) NE (B/D) |
| EEF1A1P9 | 6.69 | 4.87 | 1.55 | 1.48 | 1.82 | 8.04E-03 | 0.0304 | (A) NE (B/D) |
| RNA5SP401 | 1.98 | 0.16 | 1.63 | 0.63 | 1.82 | 9.91E-03 | 0.0355 | (A) NE (B/D) |
| ACTR1B | 3.88 | 2.07 | 0.26 | 1.07 | 1.81 | 3.56E-11 | 9.84E-09 | (A) NE (B/D) |
| RTL8C | 8.16 | 6.35 | 0.26 | 1.58 | 1.81 | 1.87E-07 | 7.96E-06 | (A) NE (B/D) |
| MIR-647/647 | 7.2 | 5.39 | 0.52 | 1.29 | 1.81 | 2.63E-07 | 1.04E-05 | (A) NE (B/D) |
| LY6G5B | 7.88 | 6.06 | 0.32 | 1.62 | 1.81 | 3.38E-07 | 1.26E-05 | (A) NE (B/D) |
| EFHC1 | 5.9 | 4.09 | 0.62 | 1.03 | 1.81 | 1.20E-06 | 3.42E-05 | (A) NE (B/D) |
| SMYD2 | 4.64 | 2.83 | 0.73 | 1.31 | 1.81 | 1.37E-05 | 2.35E-04 | (A) NE (B/D) |
| UPF3AP1 | 7.78 | 5.97 | 0.71 | 1.45 | 1.81 | 1.40E-05 | 2.37E-04 | (A) NE (B/D) |
| LENG8 | 5.22 | 3.4 | 0.78 | 1.13 | 1.81 | 2.63E-05 | 3.80E-04 | (A) NE (B/D) |
| HYLS1 | 3.15 | 1.34 | 0.84 | 1.01 | 1.81 | 6.18E-05 | 7.46E-04 | (A) NE (B/D) |
| ANKZF1 | 3.62 | 1.81 | 0.89 | 0.98 | 1.81 | 1.20E-04 | 1.24E-03 | (A) NE (B/D) |
| CAPN10-DT | 6.94 | 5.12 | 0.89 | 0.71 | 1.81 | 1.53E-04 | 1.50E-03 | (A) NE (B/D) |
| SYK | 3.7 | 1.89 | 0.92 | 1.51 | 1.81 | 1.90E-04 | 1.77E-03 | (A) NE (B/D) |
| CDC45 | 3.92 | 2.11 | 0.99 | 1.61 | 1.81 | 3.98E-04 | 3.14E-03 | (A) NE (B/D) |
| PCDHGA4 | 8.02 | 6.21 | 1.26 | 1.51 | 1.81 | 2.37E-03 | 0.0122 | (A) NE (B/D) |
| USP9YP28 | 2.42 | 0.61 | 1.36 | 0.87 | 1.81 | 3.61E-03 | 0.0168 | (A) NE (B/D) |
| EEF1A1P11 | 5.36 | 3.56 | 1.44 | 1.48 | 1.81 | 5.64E-03 | 0.0235 | (A) NE (B/D) |
| SERP2 | 4.05 | 2.24 | 1.53 | 1.12 | 1.81 | 7.34E-03 | 0.0285 | (A) NE (B/D) |
| HYI-AS1 | 5.07 | 3.26 | 1.62 | 1.25 | 1.81 | 0.0101 | 0.036 | (A) NE (B/D) |
| DAPK3 | 2.1 | 0.3 | 0.41 | 0.5 | 1.8 | 1.26E-08 | 9.37E-07 | (A) NE (B/D) |
| MIR-939/939 | 9.11 | 7.31 | 0.37 | 1.33 | 1.8 | 2.17E-08 | 1.40E-06 | (A) NE (B/D) |
| IFI27L1 | 4.97 | 3.17 | 0.43 | 1.27 | 1.8 | 3.51E-08 | 2.09E-06 | (A) NE (B/D) |
| DNLZ | 3.78 | 1.98 | 0.52 | 0.96 | 1.8 | 8.17E-08 | 4.10E-06 | (A) NE (B/D) |
| RPP21 | 4.26 | 2.47 | 0.38 | 1.48 | 1.8 | 1.41E-07 | 6.37E-06 | (A) NE (B/D) |
| GIGYF1 | 4.33 | 2.53 | 0.54 | 1.16 | 1.8 | 2.07E-07 | 8.60E-06 | (A) NE (B/D) |
| SIGIRR | 3.19 | 1.39 | 0.62 | 1.31 | 1.8 | 2.07E-06 | 5.26E-05 | (A) NE (B/D) |
| ATP11A | 3.88 | 2.08 | 0.7 | 1.12 | 1.8 | 6.94E-06 | 1.36E-04 | (A) NE (B/D) |
| SNORD23 | 6.12 | 4.32 | 0.72 | 1.23 | 1.8 | 1.22E-05 | 2.14E-04 | (A) NE (B/D) |

| | | | | | | | | |
|---|---|---|---|---|---|---|---|---|
| ARNT2 | 2.02 | 0.22 | 0.68 | 0.36 | 1.8 | 2.76E-05 | 3.94E-04 | (A) NE (B/D) |
| LINC00847 | 5.43 | 3.64 | 0.82 | 1.33 | 1.8 | 5.72E-05 | 7.03E-04 | (A) NE (B/D) |
| NDUFB2-AS1 | 4.3 | 2.5 | 0.84 | 1.15 | 1.8 | 6.53E-05 | 7.78E-04 | (A) NE (B/D) |
| HOXC-AS3 | 2.27 | 0.48 | 0.94 | 0.6 | 1.8 | 3.06E-04 | 2.56E-03 | (A) NE (B/D) |
| ABCA7 | 2.67 | 0.88 | 0.99 | 1.18 | 1.8 | 3.54E-04 | 2.87E-03 | (A) NE (B/D) |
| HLA-Z | 3.97 | 2.18 | 1.18 | 1.92 | 1.8 | 2.20E-03 | 0.0115 | (A) NE (B/D) |
| CTHRC1 | 3.44 | 1.63 | 1.31 | 1.62 | 1.8 | 3.35E-03 | 0.0159 | (A) NE (B/D) |
| IGFBP5 | 3.97 | 2.17 | 1.65 | 1.88 | 1.8 | 0.0133 | 0.0439 | (A) NE (B/D) |
| ELAVL1 | 5.75 | 3.96 | 0.38 | 1.05 | 1.79 | 1.17E-09 | 1.51E-07 | (A) NE (B/D) |
| CAMSAP3 | 2.62 | 0.83 | 0.43 | 0.95 | 1.79 | 3.36E-09 | 3.47E-07 | (A) NE (B/D) |
| SUPT5H | 9.16 | 7.37 | 0.53 | 1.03 | 1.79 | 1.28E-07 | 5.90E-06 | (A) NE (B/D) |
| GDI1 | 9.07 | 7.28 | 0.49 | 1.36 | 1.79 | 2.70E-07 | 1.05E-05 | (A) NE (B/D) |
| GYG1P3 | 2.94 | 1.15 | 0.64 | 0.92 | 1.79 | 2.41E-06 | 5.94E-05 | (A) NE (B/D) |
| MIR-4647/46 | 10.29 | 8.49 | 0.46 | 1.71 | 1.79 | 2.90E-06 | 6.83E-05 | (A) NE (B/D) |
| TUBA1B | 12.19 | 10.41 | 0.5 | 1.67 | 1.79 | 3.13E-06 | 7.27E-05 | (A) NE (B/D) |
| RASSF8-AS1 | 4.95 | 3.15 | 0.89 | 1.33 | 1.79 | 1.34E-04 | 1.35E-03 | (A) NE (B/D) |
| RN7SL521P | 3.06 | 1.27 | 0.92 | 0.94 | 1.79 | 1.85E-04 | 1.73E-03 | (A) NE (B/D) |
| RNASEH2A | 2.01 | 0.21 | 0.91 | 0.55 | 1.79 | 2.50E-04 | 2.17E-03 | (A) NE (B/D) |
| PCDHGB6 | 8.62 | 6.83 | 1.25 | 1.51 | 1.79 | 2.40E-03 | 0.0123 | (A) NE (B/D) |
| RPSAP14 | 3.43 | 1.64 | 1.41 | 0.93 | 1.79 | 4.78E-03 | 0.0208 | (A) NE (B/D) |
| ALDH8A1 | 1.89 | 0.1 | 1.45 | 0.45 | 1.79 | 6.01E-03 | 0.0246 | (A) NE (B/D) |
| RPL34P17 | 2.07 | 0.28 | 1.54 | 0.5 | 1.79 | 8.06E-03 | 0.0305 | (A) NE (B/D) |
| VN1R90P | 2.04 | 0.25 | 1.54 | 0.57 | 1.79 | 8.26E-03 | 0.031 | (A) NE (B/D) |
| MCM7 | 5.33 | 3.55 | 0.27 | 1.34 | 1.78 | 8.26E-09 | 6.79E-07 | (A) NE (B/D) |
| SPOUT1 | 7.03 | 5.25 | 0.39 | 1.32 | 1.78 | 3.47E-08 | 2.08E-06 | (A) NE (B/D) |
| SNHG5 | 4.45 | 2.67 | 0.42 | 1.6 | 1.78 | 9.00E-07 | 2.73E-05 | (A) NE (B/D) |
| TIMM44 | 2.08 | 0.31 | 0.66 | 0.49 | 1.78 | 1.59E-05 | 2.60E-04 | (A) NE (B/D) |
| TARDBP | 5.6 | 3.82 | 0.72 | 1.49 | 1.78 | 2.31E-05 | 3.43E-04 | (A) NE (B/D) |
| SSR4 | 8.75 | 6.97 | 0.72 | 1.57 | 1.78 | 2.81E-05 | 4.01E-04 | (A) NE (B/D) |
| VAMP2 | 6.66 | 4.87 | 0.83 | 1.42 | 1.78 | 7.76E-05 | 8.91E-04 | (A) NE (B/D) |
| STXBP5 | 2.12 | 0.34 | 0.89 | 0.65 | 1.78 | 1.89E-04 | 1.77E-03 | (A) NE (B/D) |
| ARHGEF1 | 6.48 | 4.71 | 1.11 | 1.35 | 1.78 | 9.96E-04 | 6.30E-03 | (A) NE (B/D) |
| HOTAIRM1 | 2.75 | 0.97 | 1.17 | 0.92 | 1.78 | 1.47E-03 | 8.50E-03 | (A) NE (B/D) |
| DACT2 | 1.98 | 0.21 | 1.31 | 0.48 | 1.78 | 3.46E-03 | 0.0163 | (A) NE (B/D) |
| TRIB2 | 2.54 | 0.76 | 1.49 | 1.07 | 1.78 | 6.94E-03 | 0.0274 | (A) NE (B/D) |
| FOXG1 | 1.89 | 0.11 | 1.55 | 0.45 | 1.78 | 8.85E-03 | 0.0326 | (A) NE (B/D) |
| ADCY2 | 2.29 | 0.52 | 1.7 | 0.86 | 1.78 | 0.014 | 0.0456 | (A) NE (B/D) |
| ABCA5 | 6.17 | 4.39 | 1.71 | 0.83 | 1.78 | 0.0142 | 0.0461 | (A) NE (B/D) |
| RAP1GAP2 | 3.24 | 1.47 | 0.33 | 0.75 | 1.77 | 1.06E-11 | 4.08E-09 | (A) NE (B/D) |
| DDX51 | 3.45 | 1.68 | 0.36 | 1.08 | 1.77 | 9.07E-10 | 1.27E-07 | (A) NE (B/D) |
| PIH1D1 | 3.18 | 1.41 | 0.52 | 1.19 | 1.77 | 1.85E-07 | 7.91E-06 | (A) NE (B/D) |
| MIR-3198-2/ | 7.18 | 5.41 | 0.46 | 1.42 | 1.77 | 2.93E-07 | 1.13E-05 | (A) NE (B/D) |
| NDUFB7 | 3.84 | 2.07 | 0.41 | 1.5 | 1.77 | 3.25E-07 | 1.23E-05 | (A) NE (B/D) |
| COX7A2 | 5.34 | 3.56 | 0.48 | 1.61 | 1.77 | 1.88E-06 | 4.92E-05 | (A) NE (B/D) |
| MIR-4746/3F | 6.17 | 4.4 | 0.62 | 1.05 | 1.77 | 2.06E-06 | 5.24E-05 | (A) NE (B/D) |
| MIR-4746/5F | 6.17 | 4.4 | 0.62 | 1.05 | 1.77 | 2.06E-06 | 5.24E-05 | (A) NE (B/D) |
| TMEM53 | 7.03 | 5.25 | 0.63 | 0.9 | 1.77 | 2.55E-06 | 6.18E-05 | (A) NE (B/D) |
| SNORD48 | 6.54 | 4.77 | 0.67 | 0.79 | 1.77 | 7.12E-06 | 1.39E-04 | (A) NE (B/D) |
| SAFB2 | 7.36 | 5.59 | 0.69 | 0.72 | 1.77 | 1.20E-05 | 2.10E-04 | (A) NE (B/D) |
| AKR7L | 2.62 | 0.85 | 0.96 | 0.95 | 1.77 | 3.09E-04 | 2.58E-03 | (A) NE (B/D) |
| EIF4EBP1 | 9.5 | 7.73 | 1.01 | 1.16 | 1.77 | 4.59E-04 | 3.50E-03 | (A) NE (B/D) |
| HERC2P3 | 7.49 | 5.72 | 1.06 | 0.94 | 1.77 | 7.17E-04 | 4.94E-03 | (A) NE (B/D) |
| RPS19P3 | 8.23 | 6.46 | 1.18 | 1.06 | 1.77 | 1.59E-03 | 8.99E-03 | (A) NE (B/D) |
| RTL5 | 8.13 | 6.37 | 1.13 | 1.88 | 1.77 | 1.83E-03 | 0.01 | (A) NE (B/D) |
| GPX2 | 1.8 | 0.02 | 1.33 | 0.08 | 1.77 | 3.86E-03 | 0.0177 | (A) NE (B/D) |
| CHRM5 | 2.69 | 0.92 | 1.62 | 0.79 | 1.77 | 0.0111 | 0.0384 | (A) NE (B/D) |
| IMPA1P1 | 1.91 | 0.14 | 1.7 | 0.41 | 1.77 | 0.0143 | 0.0464 | (A) NE (B/D) |
| REPIN1 | 3.01 | 1.25 | 0.33 | 0.71 | 1.76 | 1.28E-11 | 4.76E-09 | (A) NE (B/D) |
| LLGL2 | 2.51 | 0.75 | 0.49 | 0.89 | 1.76 | 3.75E-08 | 2.21E-06 | (A) NE (B/D) |
| DPP7 | 9.06 | 7.3 | 0.62 | 1.17 | 1.76 | 2.02E-06 | 5.18E-05 | (A) NE (B/D) |
| UPF3AP2 | 3.21 | 1.45 | 0.66 | 1.11 | 1.76 | 4.89E-06 | 1.04E-04 | (A) NE (B/D) |
| MARCKSL1 | 5.71 | 3.95 | 0.55 | 1.79 | 1.76 | 1.56E-05 | 2.57E-04 | (A) NE (B/D) |
| CHDH | 4.86 | 3.1 | 0.61 | 1.7 | 1.76 | 1.61E-05 | 2.63E-04 | (A) NE (B/D) |
| CDKN2C | 1.82 | 0.06 | 1.03 | 0.28 | 1.76 | 8.45E-04 | 5.58E-03 | (A) NE (B/D) |
| MIR-3940/39 | 3.41 | 1.65 | 0.24 | 0.93 | 1.75 | 2.42E-12 | 1.32E-09 | (A) NE (B/D) |

| | | | | | | | | |
|---|---|---|---|---|---|---|---|---|
| MIR-3940/3P | 3.41 | 1.65 | 0.24 | 0.93 | 1.75 | 2.42E-12 | 1.32E-09 | (A) NE (B/D) |
| MIR-3940/5P | 3.41 | 1.65 | 0.24 | 0.93 | 1.75 | 2.42E-12 | 1.32E-09 | (A) NE (B/D) |
| CNNM3 | 2.21 | 0.46 | 0.41 | 0.63 | 1.75 | 3.84E-09 | 3.79E-07 | (A) NE (B/D) |
| C1ORF53 | 4.89 | 3.15 | 0.36 | 1.38 | 1.75 | 5.86E-08 | 3.13E-06 | (A) NE (B/D) |
| URAHP | 5.49 | 3.74 | 0.63 | 1.07 | 1.75 | 3.01E-06 | 7.02E-05 | (A) NE (B/D) |
| KRTAP5-AS1 | 4.77 | 3.02 | 0.53 | 1.82 | 1.75 | 1.73E-05 | 2.77E-04 | (A) NE (B/D) |
| INCENP | 2.56 | 0.81 | 0.74 | 0.95 | 1.75 | 2.42E-05 | 3.55E-04 | (A) NE (B/D) |
| HNRNPA3P10 | 4.08 | 2.34 | 0.58 | 1.94 | 1.75 | 5.01E-05 | 6.32E-04 | (A) NE (B/D) |
| PEMT | 1.98 | 0.23 | 0.83 | 0.49 | 1.75 | 1.48E-04 | 1.47E-03 | (A) NE (B/D) |
| CCDC14 | 2.58 | 0.83 | 1.27 | 0.72 | 1.75 | 3.03E-03 | 0.0147 | (A) NE (B/D) |
| SGCB | 4.21 | 2.46 | 1.29 | 1.21 | 1.75 | 3.30E-03 | 0.0157 | (A) NE (B/D) |
| F7 | 2.53 | 0.78 | 1.3 | 1.09 | 1.75 | 3.34E-03 | 0.0158 | (A) NE (B/D) |
| HMGN3 | 4.19 | 2.43 | 1.49 | 1.43 | 1.75 | 8.02E-03 | 0.0303 | (A) NE (B/D) |
| MIR-4444-1/ | 2.41 | 0.66 | 1.51 | 0.92 | 1.75 | 8.33E-03 | 0.0312 | (A) NE (B/D) |
| CLPTM1 | 4.12 | 2.38 | 0.21 | 1.23 | 1.74 | 1.66E-09 | 2.01E-07 | (A) NE (B/D) |
| NOP53 | 6.82 | 5.08 | 0.31 | 1.2 | 1.74 | 2.70E-09 | 2.99E-07 | (A) NE (B/D) |
| BRICD5 | 3.43 | 1.69 | 0.36 | 1.14 | 1.74 | 2.90E-09 | 3.12E-07 | (A) NE (B/D) |
| NELFB | 9.77 | 8.03 | 0.4 | 1.12 | 1.74 | 7.86E-09 | 6.57E-07 | (A) NE (B/D) |
| HNRNPA1 | 4.04 | 2.29 | 0.21 | 1.55 | 1.74 | 2.16E-07 | 8.89E-06 | (A) NE (B/D) |
| TBCAP1 | 4.24 | 2.5 | 0.49 | 1.63 | 1.74 | 3.70E-06 | 8.35E-05 | (A) NE (B/D) |
| SAPCD1-AS1 | 6.23 | 4.49 | 0.51 | 1.69 | 1.74 | 6.06E-06 | 1.23E-04 | (A) NE (B/D) |
| PKD1 | 3.46 | 1.73 | 0.61 | 0.55 | 1.74 | 6.68E-06 | 1.33E-04 | (A) NE (B/D) |
| MEA1 | 4.19 | 2.45 | 0.59 | 1.58 | 1.74 | 8.29E-06 | 1.57E-04 | (A) NE (B/D) |
| FAM13A-AS1 | 7.25 | 5.51 | 0.75 | 1.14 | 1.74 | 2.74E-05 | 3.92E-04 | (A) NE (B/D) |
| NPIPA8 | 3.01 | 1.26 | 0.73 | 0.64 | 1.74 | 3.30E-05 | 4.57E-04 | (A) NE (B/D) |
| PSMG4 | 3.22 | 1.48 | 0.95 | 0.79 | 1.74 | 3.61E-04 | 2.91E-03 | (A) NE (B/D) |
| TSPAN12 | 1.77 | 0.02 | 0.89 | 0.14 | 1.74 | 3.66E-04 | 2.94E-03 | (A) NE (B/D) |
| TOP3BP1 | 5.1 | 3.36 | 1.01 | 0.93 | 1.74 | 5.55E-04 | 4.06E-03 | (A) NE (B/D) |
| ANKRD13B | 6.1 | 4.37 | 1.04 | 1.46 | 1.74 | 7.95E-04 | 5.35E-03 | (A) NE (B/D) |
| SUGP2 | 1.94 | 0.2 | 1.15 | 0.29 | 1.74 | 1.81E-03 | 9.93E-03 | (A) NE (B/D) |
| PLEKHH2 | 1.83 | 0.09 | 1.25 | 0.31 | 1.74 | 3.01E-03 | 0.0146 | (A) NE (B/D) |
| PCDHB18P | 2.97 | 1.23 | 1.27 | 1 | 1.74 | 3.08E-03 | 0.0149 | (A) NE (B/D) |
| EIF3LP2 | 5.02 | 3.28 | 1.38 | 1.61 | 1.74 | 5.81E-03 | 0.024 | (A) NE (B/D) |
| HCG4P7 | 7.07 | 5.33 | 1.46 | 1.42 | 1.74 | 7.43E-03 | 0.0288 | (A) NE (B/D) |
| ZNF585B | 3.13 | 1.4 | 1.49 | 0.79 | 1.74 | 7.80E-03 | 0.0297 | (A) NE (B/D) |
| TJAP1 | 2.51 | 0.78 | 0.26 | 0.87 | 1.73 | 1.45E-12 | 9.30E-10 | (A) NE (B/D) |
| PIAS4 | 1.93 | 0.2 | 0.27 | 0.33 | 1.73 | 6.77E-11 | 1.72E-08 | (A) NE (B/D) |
| GNL1 | 7.2 | 5.47 | 0.34 | 0.98 | 1.73 | 2.72E-10 | 5.26E-08 | (A) NE (B/D) |
| DDX41 | 3.73 | 2.01 | 0.28 | 1.11 | 1.73 | 4.99E-10 | 8.61E-08 | (A) NE (B/D) |
| RPS10P7 | 3.31 | 1.58 | 0.38 | 0.92 | 1.73 | 5.79E-10 | 9.60E-08 | (A) NE (B/D) |
| LRRC45 | 2.81 | 1.08 | 0.33 | 1.08 | 1.73 | 6.80E-10 | 1.07E-07 | (A) NE (B/D) |
| PPT2-EGFL8 | 8.59 | 6.86 | 0.26 | 1.22 | 1.73 | 2.60E-09 | 2.96E-07 | (A) NE (B/D) |
| NOXA1 | 4.2 | 2.47 | 0.32 | 1.24 | 1.73 | 6.22E-09 | 5.52E-07 | (A) NE (B/D) |
| BACE1 | 5.29 | 3.57 | 0.43 | 1 | 1.73 | 9.35E-09 | 7.51E-07 | (A) NE (B/D) |
| MPV17L2 | 5.65 | 3.92 | 0.38 | 1.33 | 1.73 | 5.86E-08 | 3.13E-06 | (A) NE (B/D) |
| DDA1 | 2.13 | 0.4 | 0.46 | 0.51 | 1.73 | 1.15E-07 | 5.40E-06 | (A) NE (B/D) |
| TONSL | 3.59 | 1.85 | 0.53 | 0.99 | 1.73 | 2.02E-07 | 8.40E-06 | (A) NE (B/D) |
| GPX4 | 11.8 | 10.07 | 0.41 | 1.43 | 1.73 | 2.66E-07 | 1.04E-05 | (A) NE (B/D) |
| BSN | 2.02 | 0.29 | 0.5 | 0.52 | 1.73 | 4.80E-07 | 1.67E-05 | (A) NE (B/D) |
| ERAL1 | 4 | 2.27 | 0.38 | 1.56 | 1.73 | 6.96E-07 | 2.22E-05 | (A) NE (B/D) |
| SLC35B2 | 8.72 | 6.99 | 0.5 | 1.4 | 1.73 | 8.48E-07 | 2.61E-05 | (A) NE (B/D) |
| ARID3B | 2.95 | 1.21 | 0.67 | 0.99 | 1.73 | 6.34E-06 | 1.28E-04 | (A) NE (B/D) |
| TMEM44-AS1 | 8.79 | 7.06 | 0.56 | 1.64 | 1.73 | 8.89E-06 | 1.66E-04 | (A) NE (B/D) |
| MINK1 | 3.37 | 1.63 | 0.75 | 1.04 | 1.73 | 2.99E-05 | 4.22E-04 | (A) NE (B/D) |
| PCDHA1 | 2.79 | 1.06 | 0.85 | 0.95 | 1.73 | 1.28E-04 | 1.31E-03 | (A) NE (B/D) |
| NFKBID | 4.67 | 2.94 | 0.89 | 1.24 | 1.73 | 1.80E-04 | 1.70E-03 | (A) NE (B/D) |
| CDT1 | 2.7 | 0.97 | 0.9 | 0.89 | 1.73 | 2.19E-04 | 1.97E-03 | (A) NE (B/D) |
| PCDHB9 | 1.97 | 0.23 | 0.92 | 0.39 | 1.73 | 3.96E-04 | 3.13E-03 | (A) NE (B/D) |
| EPHB3 | 2.58 | 0.85 | 1.06 | 1.11 | 1.73 | 8.00E-04 | 5.38E-03 | (A) NE (B/D) |
| NAP1L1P1 | 4.09 | 2.35 | 1.06 | 1.08 | 1.73 | 8.31E-04 | 5.52E-03 | (A) NE (B/D) |
| NT5DC4 | 3.14 | 1.42 | 1.08 | 1.31 | 1.73 | 1.03E-03 | 6.45E-03 | (A) NE (B/D) |
| TSHZ2 | 3.14 | 1.42 | 1.24 | 1.11 | 1.73 | 2.75E-03 | 0.0137 | (A) NE (B/D) |
| RNU7-61P | 3.2 | 1.47 | 1.35 | 1.65 | 1.73 | 5.32E-03 | 0.0225 | (A) NE (B/D) |
| RNU6-431P | 3.51 | 1.77 | 1.46 | 1.31 | 1.73 | 7.50E-03 | 0.029 | (A) NE (B/D) |
| MIR-196B/19 | 2.35 | 0.62 | 1.56 | 0.93 | 1.73 | 0.0103 | 0.0365 | (A) NE (B/D) |

| | | | | | | | | |
|---|---|---|---|---|---|---|---|---|
| MIR-196B/19 | 2.35 | 0.62 | 1.56 | 0.93 | 1.73 | 0.0103 | 0.0365 | (A) NE (B/D) |
| MIR-196B/3P | 2.35 | 0.62 | 1.56 | 0.93 | 1.73 | 0.0103 | 0.0365 | (A) NE (B/D) |
| MIR-196B/5P | 2.35 | 0.62 | 1.56 | 0.93 | 1.73 | 0.0103 | 0.0365 | (A) NE (B/D) |
| UCP2 | 8.88 | 7.15 | 1.41 | 2.28 | 1.73 | 0.0103 | 0.0365 | (A) NE (B/D) |
| ANKS6 | 4.2 | 2.48 | 1.58 | 1.03 | 1.73 | 0.0114 | 0.0391 | (A) NE (B/D) |
| DDRGK1 | 6.22 | 4.5 | 0.32 | 0.99 | 1.72 | 1.78E-10 | 3.84E-08 | (A) NE (B/D) |
| FAM174B | 2.62 | 0.9 | 0.29 | 1.11 | 1.72 | 6.36E-10 | 1.03E-07 | (A) NE (B/D) |
| CENPO | 3.64 | 1.92 | 0.34 | 1.06 | 1.72 | 8.90E-10 | 1.27E-07 | (A) NE (B/D) |
| TMEM178B | 2.63 | 0.91 | 0.39 | 0.74 | 1.72 | 1.18E-09 | 1.51E-07 | (A) NE (B/D) |
| SNRNP200 | 3.33 | 1.62 | 0.41 | 1.07 | 1.72 | 8.83E-09 | 7.17E-07 | (A) NE (B/D) |
| CDK11B | 4.85 | 3.13 | 0.33 | 1.32 | 1.72 | 2.63E-08 | 1.64E-06 | (A) NE (B/D) |
| MCRIP1 | 8.06 | 6.34 | 0.48 | 1.24 | 1.72 | 1.77E-07 | 7.61E-06 | (A) NE (B/D) |
| RNF5P1 | 4.87 | 3.15 | 0.43 | 1.53 | 1.72 | 1.04E-06 | 3.08E-05 | (A) NE (B/D) |
| DUSP4 | 1.89 | 0.17 | 0.59 | 0.39 | 1.72 | 8.99E-06 | 1.67E-04 | (A) NE (B/D) |
| FUZ | 3.44 | 1.72 | 0.7 | 0.73 | 1.72 | 1.83E-05 | 2.89E-04 | (A) NE (B/D) |
| GCDH | 3.16 | 1.44 | 0.72 | 1.2 | 1.72 | 2.00E-05 | 3.09E-04 | (A) NE (B/D) |
| METTL17 | 4.37 | 2.65 | 0.76 | 1.31 | 1.72 | 4.24E-05 | 5.56E-04 | (A) NE (B/D) |
| RNU1-123P | 2.15 | 0.43 | 0.78 | 0.64 | 1.72 | 7.67E-05 | 8.83E-04 | (A) NE (B/D) |
| BOLA3-AS1 | 2.68 | 0.96 | 0.86 | 0.77 | 1.72 | 1.65E-04 | 1.59E-03 | (A) NE (B/D) |
| GP1BB | 2.04 | 0.32 | 0.94 | 0.6 | 1.72 | 4.23E-04 | 3.29E-03 | (A) NE (B/D) |
| DLGAP1-AS1 | 1.8 | 0.08 | 1 | 0.25 | 1.72 | 8.10E-04 | 5.41E-03 | (A) NE (B/D) |
| ABCD4 | 2.29 | 0.57 | 1.24 | 0.73 | 1.72 | 2.93E-03 | 0.0143 | (A) NE (B/D) |
| PCDHGA5 | 8.17 | 6.46 | 1.25 | 1.57 | 1.72 | 3.30E-03 | 0.0157 | (A) NE (B/D) |
| HERC2P4 | 2.07 | 0.35 | 1.44 | 0.5 | 1.72 | 6.91E-03 | 0.0273 | (A) NE (B/D) |
| ESRRG | 3.96 | 2.24 | 1.55 | 1.51 | 1.72 | 0.0113 | 0.0388 | (A) NE (B/D) |
| SNORD114-1 | 1.99 | 0.26 | 1.6 | 0.78 | 1.72 | 0.0122 | 0.0411 | (A) NE (B/D) |
| TOLLIP-AS1 | 3.85 | 2.15 | 0.47 | 1 | 1.71 | 5.10E-08 | 2.83E-06 | (A) NE (B/D) |
| RNF207 | 4.14 | 2.44 | 0.47 | 1.26 | 1.71 | 2.01E-07 | 8.40E-06 | (A) NE (B/D) |
| APBB1 | 5.25 | 3.55 | 0.65 | 1.32 | 1.71 | 9.01E-06 | 1.67E-04 | (A) NE (B/D) |
| PKD1P1 | 9.35 | 7.64 | 0.73 | 0.69 | 1.71 | 3.76E-05 | 5.07E-04 | (A) NE (B/D) |
| ADGRB2 | 1.77 | 0.06 | 0.73 | 0.2 | 1.71 | 1.03E-04 | 1.11E-03 | (A) NE (B/D) |
| WHRN | 1.86 | 0.15 | 0.86 | 0.48 | 1.71 | 2.48E-04 | 2.17E-03 | (A) NE (B/D) |
| INPP5B | 3.82 | 2.12 | 0.91 | 0.66 | 1.71 | 3.30E-04 | 2.71E-03 | (A) NE (B/D) |
| PIF1 | 1.88 | 0.18 | 1 | 0.36 | 1.71 | 8.10E-04 | 5.41E-03 | (A) NE (B/D) |
| TIMP1 | 4.26 | 2.55 | 1.14 | 1.46 | 1.71 | 1.74E-03 | 9.64E-03 | (A) NE (B/D) |
| TECTA | 2.35 | 0.64 | 1.16 | 0.57 | 1.71 | 1.99E-03 | 0.0107 | (A) NE (B/D) |
| PPIAP6 | 5.9 | 4.19 | 1.42 | 1.26 | 1.71 | 6.74E-03 | 0.0268 | (A) NE (B/D) |
| A1BG-AS1 | 3.13 | 1.42 | 1.45 | 0.97 | 1.71 | 7.50E-03 | 0.029 | (A) NE (B/D) |
| MIR-3607/3P | 2.19 | 0.48 | 1.61 | 0.77 | 1.71 | 0.0127 | 0.0425 | (A) NE (B/D) |
| MIR-3607/5P | 2.19 | 0.48 | 1.61 | 0.77 | 1.71 | 0.0127 | 0.0425 | (A) NE (B/D) |
| FAM222B | 3.21 | 1.51 | 0.31 | 1.01 | 1.7 | 2.17E-10 | 4.30E-08 | (A) NE (B/D) |
| UCK1 | 6.56 | 4.85 | 0.28 | 1.24 | 1.7 | 6.01E-09 | 5.37E-07 | (A) NE (B/D) |
| KCTD17 | 3.4 | 1.7 | 0.39 | 1.1 | 1.7 | 6.64E-09 | 5.77E-07 | (A) NE (B/D) |
| RUVBL2 | 4.55 | 2.85 | 0.47 | 1.13 | 1.7 | 9.25E-08 | 4.51E-06 | (A) NE (B/D) |
| SLC2A8 | 5.38 | 3.68 | 0.37 | 1.36 | 1.7 | 1.18E-07 | 5.48E-06 | (A) NE (B/D) |
| UQCRH | 5.14 | 3.44 | 0.28 | 1.56 | 1.7 | 4.91E-07 | 1.70E-05 | (A) NE (B/D) |
| CIRBP | 5.19 | 3.49 | 0.65 | 1.52 | 1.7 | 1.96E-05 | 3.03E-04 | (A) NE (B/D) |
| SLC25A5 | 6.1 | 4.41 | 0.54 | 1.83 | 1.7 | 3.04E-05 | 4.27E-04 | (A) NE (B/D) |
| STK36 | 5.16 | 3.46 | 0.78 | 1.15 | 1.7 | 5.27E-05 | 6.59E-04 | (A) NE (B/D) |
| GAPDHP15 | 1.92 | 0.22 | 0.82 | 0.39 | 1.7 | 1.87E-04 | 1.75E-03 | (A) NE (B/D) |
| SNORD38A | 3.54 | 1.85 | 0.88 | 1.23 | 1.7 | 2.00E-04 | 1.84E-03 | (A) NE (B/D) |
| EXOSC10 | 2.56 | 0.86 | 0.89 | 0.93 | 1.7 | 2.36E-04 | 2.09E-03 | (A) NE (B/D) |
| CRIP2 | 3.06 | 1.36 | 0.92 | 1.16 | 1.7 | 2.80E-04 | 2.39E-03 | (A) NE (B/D) |
| TMX4 | 4.34 | 2.63 | 0.96 | 0.78 | 1.7 | 4.67E-04 | 3.54E-03 | (A) NE (B/D) |
| LINC00312 | 2.42 | 0.72 | 1.04 | 0.72 | 1.7 | 9.07E-04 | 5.88E-03 | (A) NE (B/D) |
| XAB2 | 2.84 | 1.15 | 0.38 | 0.82 | 1.69 | 7.49E-10 | 1.13E-07 | (A) NE (B/D) |
| RFXANK | 4.69 | 3.01 | 0.31 | 1.25 | 1.69 | 1.16E-08 | 8.90E-07 | (A) NE (B/D) |
| UQCR11 | 7.62 | 5.93 | 0.16 | 1.36 | 1.69 | 2.55E-08 | 1.60E-06 | (A) NE (B/D) |
| TEN1 | 7.88 | 6.19 | 0.43 | 1.16 | 1.69 | 4.43E-08 | 2.51E-06 | (A) NE (B/D) |
| GOT1 | 5.73 | 4.05 | 0.33 | 1.41 | 1.69 | 1.45E-07 | 6.49E-06 | (A) NE (B/D) |
| LINC00526 | 4.49 | 2.8 | 0.53 | 0.84 | 1.69 | 4.29E-07 | 1.52E-05 | (A) NE (B/D) |
| FDX2 | 2.39 | 0.7 | 0.65 | 0.97 | 1.69 | 6.42E-06 | 1.29E-04 | (A) NE (B/D) |
| TMC4 | 4.43 | 2.74 | 0.68 | 1.32 | 1.69 | 1.61E-05 | 2.62E-04 | (A) NE (B/D) |
| SLC25A29 | 9.53 | 7.84 | 0.64 | 1.52 | 1.69 | 1.73E-05 | 2.77E-04 | (A) NE (B/D) |
| BEX3 | 4.26 | 2.57 | 0.65 | 1.92 | 1.69 | 1.18E-04 | 1.23E-03 | (A) NE (B/D) |

| | | | | | | | | |
|---|---|---|---|---|---|---|---|---|
| UPF3B | 8 | 6.31 | 0.86 | 1.3 | 1.69 | 1.68E-04 | 1.61E-03 | (A) NE (B/D) |
| RNU6-353P | 3.12 | 1.42 | 1.11 | 1.44 | 1.69 | 1.53E-03 | 8.74E-03 | (A) NE (B/D) |
| FAM200B | 8.57 | 6.88 | 1.26 | 1.03 | 1.69 | 3.36E-03 | 0.0159 | (A) NE (B/D) |
| CEND1 | 1.79 | 0.1 | 1.25 | 0.3 | 1.69 | 3.58E-03 | 0.0167 | (A) NE (B/D) |
| CDK2AP2P2 | 2.44 | 0.75 | 1.28 | 0.86 | 1.69 | 3.90E-03 | 0.0178 | (A) NE (B/D) |
| RP9P | 3.3 | 1.61 | 1.38 | 0.94 | 1.69 | 5.95E-03 | 0.0245 | (A) NE (B/D) |
| MIR-922/922 | 5.87 | 4.19 | 0.3 | 0.84 | 1.68 | 1.67E-11 | 5.88E-09 | (A) NE (B/D) |
| TLNRD1 | 4.68 | 2.99 | 0.4 | 0.56 | 1.68 | 8.16E-09 | 6.75E-07 | (A) NE (B/D) |
| CDC34 | 3.69 | 2.01 | 0.37 | 1.18 | 1.68 | 1.50E-08 | 1.06E-06 | (A) NE (B/D) |
| CPSF7 | 3.48 | 1.8 | 0.25 | 1.34 | 1.68 | 3.11E-08 | 1.90E-06 | (A) NE (B/D) |
| CNPY4 | 3.87 | 2.19 | 0.45 | 1.04 | 1.68 | 4.08E-08 | 2.36E-06 | (A) NE (B/D) |
| UBAC1 | 6.15 | 4.46 | 0.44 | 1.21 | 1.68 | 1.01E-07 | 4.86E-06 | (A) NE (B/D) |
| MAN2A2 | 7.54 | 5.85 | 0.56 | 1.04 | 1.68 | 8.37E-07 | 2.58E-05 | (A) NE (B/D) |
| ZFAND2B | 5.44 | 3.76 | 0.56 | 1.16 | 1.68 | 1.20E-06 | 3.42E-05 | (A) NE (B/D) |
| COG1 | 2.77 | 1.09 | 0.58 | 1.02 | 1.68 | 1.37E-06 | 3.82E-05 | (A) NE (B/D) |
| DDX11-AS1 | 2.04 | 0.36 | 0.67 | 0.58 | 1.68 | 2.32E-05 | 3.44E-04 | (A) NE (B/D) |
| ZBED8 | 3.86 | 2.18 | 0.72 | 0.96 | 1.68 | 2.43E-05 | 3.56E-04 | (A) NE (B/D) |
| RPL18AP3 | 13.68 | 12 | 0.78 | 1.55 | 1.68 | 1.10E-04 | 1.16E-03 | (A) NE (B/D) |
| FBXO41 | 2.05 | 0.37 | 0.78 | 0.49 | 1.68 | 1.17E-04 | 1.23E-03 | (A) NE (B/D) |
| DNAJC12 | 2.02 | 0.33 | 0.92 | 0.92 | 1.68 | 3.25E-04 | 2.68E-03 | (A) NE (B/D) |
| RADIL | 1.89 | 0.21 | 0.96 | 0.32 | 1.68 | 6.89E-04 | 4.79E-03 | (A) NE (B/D) |
| SNORD87 | 8.1 | 6.42 | 1.11 | 1.49 | 1.68 | 1.66E-03 | 9.29E-03 | (A) NE (B/D) |
| TRNP1 | 5.13 | 3.45 | 1.21 | 1.27 | 1.68 | 2.94E-03 | 0.0144 | (A) NE (B/D) |
| PRR3 | 2.6 | 0.93 | 0.27 | 0.93 | 1.67 | 2.23E-11 | 7.21E-09 | (A) NE (B/D) |
| HDAC10 | 6.46 | 4.79 | 0.34 | 0.99 | 1.67 | 8.04E-10 | 1.19E-07 | (A) NE (B/D) |
| EHMT2 | 5.97 | 4.31 | 0.31 | 1.12 | 1.67 | 2.07E-09 | 2.43E-07 | (A) NE (B/D) |
| ZMYM3 | 2.62 | 0.95 | 0.39 | 1.01 | 1.67 | 4.29E-09 | 4.11E-07 | (A) NE (B/D) |
| WRAP73 | 5.62 | 3.96 | 0.27 | 1.2 | 1.67 | 4.53E-09 | 4.32E-07 | (A) NE (B/D) |
| NFKBIL1 | 7.75 | 6.07 | 0.38 | 1.06 | 1.67 | 5.04E-09 | 4.68E-07 | (A) NE (B/D) |
| YIPF3 | 10.18 | 8.51 | 0.31 | 1.27 | 1.67 | 2.18E-08 | 1.40E-06 | (A) NE (B/D) |
| HNRNPUL1 | 3.91 | 2.24 | 0.46 | 1.05 | 1.67 | 5.66E-08 | 3.05E-06 | (A) NE (B/D) |
| TFEB | 4.27 | 2.6 | 0.44 | 1.2 | 1.67 | 9.31E-08 | 4.53E-06 | (A) NE (B/D) |
| ENC1 | 2.82 | 1.15 | 0.38 | 1.34 | 1.67 | 1.44E-07 | 6.49E-06 | (A) NE (B/D) |
| HTR5BP | 4.25 | 2.57 | 0.48 | 0.52 | 1.67 | 3.91E-07 | 1.41E-05 | (A) NE (B/D) |
| TXLNA | 5.85 | 4.18 | 0.61 | 0.77 | 1.67 | 4.45E-06 | 9.62E-05 | (A) NE (B/D) |
| SMTN | 4.99 | 3.32 | 0.63 | 1.18 | 1.67 | 5.43E-06 | 1.12E-04 | (A) NE (B/D) |
| TMEM238 | 4.13 | 2.46 | 0.64 | 1.4 | 1.67 | 1.31E-05 | 2.27E-04 | (A) NE (B/D) |
| RAB17 | 1.78 | 0.1 | 0.63 | 0.31 | 1.67 | 2.79E-05 | 3.98E-04 | (A) NE (B/D) |
| PLXNA3 | 6.22 | 4.55 | 0.72 | 1.21 | 1.67 | 3.03E-05 | 4.26E-04 | (A) NE (B/D) |
| XRCC1 | 6.4 | 4.73 | 0.82 | 1.33 | 1.67 | 1.33E-04 | 1.35E-03 | (A) NE (B/D) |
| TTC28-AS1 | 2.47 | 0.8 | 0.83 | 0.8 | 1.67 | 1.47E-04 | 1.46E-03 | (A) NE (B/D) |
| PERP | 9.65 | 7.98 | 0.76 | 1.79 | 1.67 | 2.02E-04 | 1.85E-03 | (A) NE (B/D) |
| RGS12 | 1.73 | 0.06 | 0.84 | 0.21 | 1.67 | 3.03E-04 | 2.54E-03 | (A) NE (B/D) |
| GNRH2 | 1.91 | 0.24 | 1.48 | 0.45 | 1.67 | 9.74E-03 | 0.035 | (A) NE (B/D) |
| SDHAF1 | 3.03 | 1.37 | 0.36 | 0.95 | 1.66 | 1.02E-09 | 1.38E-07 | (A) NE (B/D) |
| INTS1 | 5.43 | 3.76 | 0.42 | 0.61 | 1.66 | 1.67E-08 | 1.15E-06 | (A) NE (B/D) |
| FAM193B | 6.79 | 5.13 | 0.46 | 0.96 | 1.66 | 4.54E-08 | 2.56E-06 | (A) NE (B/D) |
| MARCH9 | 5.51 | 3.85 | 0.41 | 1.23 | 1.66 | 8.66E-08 | 4.29E-06 | (A) NE (B/D) |
| SZT2-AS1 | 2.69 | 1.03 | 0.48 | 0.89 | 1.66 | 1.02E-07 | 4.88E-06 | (A) NE (B/D) |
| ASNA1 | 8.83 | 7.17 | 0.37 | 1.39 | 1.66 | 2.30E-07 | 9.33E-06 | (A) NE (B/D) |
| MAFF | 8.23 | 6.57 | 0.47 | 1.29 | 1.66 | 4.16E-07 | 1.48E-05 | (A) NE (B/D) |
| SLC12A9 | 4.43 | 2.77 | 0.53 | 1.26 | 1.66 | 1.21E-06 | 3.45E-05 | (A) NE (B/D) |
| YIPF2 | 3.71 | 2.05 | 0.59 | 0.87 | 1.66 | 2.18E-06 | 5.49E-05 | (A) NE (B/D) |
| NENF | 6.01 | 4.36 | 0.36 | 1.61 | 1.66 | 2.22E-06 | 5.55E-05 | (A) NE (B/D) |
| GOLGA6L9 | 7.12 | 5.47 | 0.68 | 1.05 | 1.66 | 1.48E-05 | 2.48E-04 | (A) NE (B/D) |
| LSR | 3.08 | 1.41 | 0.77 | 1.08 | 1.66 | 6.13E-05 | 7.41E-04 | (A) NE (B/D) |
| MIR-4683/46 | 2.45 | 0.79 | 1.06 | 1.12 | 1.66 | 1.18E-03 | 7.17E-03 | (A) NE (B/D) |
| TRAPPC5 | 5.08 | 3.42 | 1.21 | 1.01 | 1.66 | 2.94E-03 | 0.0144 | (A) NE (B/D) |
| NUAK2 | 2.21 | 0.55 | 1.29 | 1 | 1.66 | 4.59E-03 | 0.0202 | (A) NE (B/D) |
| LINC00431 | 3.56 | 1.9 | 1.3 | 0.78 | 1.66 | 4.63E-03 | 0.0203 | (A) NE (B/D) |
| ZNF135 | 4.06 | 2.39 | 1.49 | 0.77 | 1.66 | 0.0102 | 0.0361 | (A) NE (B/D) |
| CMAHP | 3.34 | 1.68 | 1.61 | 1.01 | 1.66 | 0.015 | 0.0479 | (A) NE (B/D) |
| TMUB1 | 2 | 0.35 | 0.43 | 0.55 | 1.65 | 4.38E-08 | 2.49E-06 | (A) NE (B/D) |
| HIGD2A | 3.11 | 1.46 | 0.32 | 1.31 | 1.65 | 5.55E-08 | 3.01E-06 | (A) NE (B/D) |
| C7ORF26 | 9.53 | 7.88 | 0.47 | 1.01 | 1.65 | 8.72E-08 | 4.31E-06 | (A) NE (B/D) |

| | | | | | | | | |
|---|---|---|---|---|---|---|---|---|
| ANAPC11 | 2.97 | 1.32 | 0.42 | 1.31 | 1.65 | 2.26E-07 | 9.21E-06 | (A) NE (B/D) |
| SMIM29 | 8.11 | 6.46 | 0.47 | 1.21 | 1.65 | 2.81E-07 | 1.09E-05 | (A) NE (B/D) |
| PSMC5 | 4.42 | 2.77 | 0.21 | 1.67 | 1.65 | 2.25E-06 | 5.62E-05 | (A) NE (B/D) |
| NOL4L | 3.24 | 1.59 | 0.65 | 0.71 | 1.65 | 1.38E-05 | 2.36E-04 | (A) NE (B/D) |
| ATP6AP1 | 4.18 | 2.52 | 0.63 | 1.47 | 1.65 | 1.79E-05 | 2.84E-04 | (A) NE (B/D) |
| ING5 | 7.51 | 5.86 | 0.74 | 1.04 | 1.65 | 4.51E-05 | 5.82E-04 | (A) NE (B/D) |
| NCAPG2 | 2.6 | 0.94 | 0.75 | 0.94 | 1.65 | 5.15E-05 | 6.46E-04 | (A) NE (B/D) |
| KIF11 | 3.8 | 2.15 | 0.97 | 1.57 | 1.65 | 8.07E-04 | 5.40E-03 | (A) NE (B/D) |
| TAPT1 | 5.18 | 3.53 | 1.05 | 0.87 | 1.65 | 1.13E-03 | 6.94E-03 | (A) NE (B/D) |
| CDKN3 | 2.05 | 0.4 | 1.1 | 0.82 | 1.65 | 1.60E-03 | 9.03E-03 | (A) NE (B/D) |
| GAS2L3 | 2.25 | 0.6 | 1.36 | 0.8 | 1.65 | 6.20E-03 | 0.0252 | (A) NE (B/D) |
| NAP1L5 | 6.63 | 4.98 | 1.44 | 1.54 | 1.65 | 9.79E-03 | 0.0352 | (A) NE (B/D) |
| SHOX2 | 1.73 | 0.09 | 1.56 | 0.26 | 1.65 | 0.0135 | 0.0443 | (A) NE (B/D) |
| PRKCSH | 2.37 | 0.73 | 0.27 | 0.84 | 1.64 | 7.91E-12 | 3.39E-09 | (A) NE (B/D) |
| NIPA1 | 2.94 | 1.3 | 0.43 | 0.95 | 1.64 | 2.13E-08 | 1.39E-06 | (A) NE (B/D) |
| UNK | 8.55 | 6.91 | 0.45 | 0.84 | 1.64 | 3.92E-08 | 2.28E-06 | (A) NE (B/D) |
| TRIM52-AS1 | 3.59 | 1.94 | 0.38 | 1.24 | 1.64 | 6.43E-08 | 3.36E-06 | (A) NE (B/D) |
| EIF3G | 4.19 | 2.55 | 0.49 | 1.25 | 1.64 | 6.89E-07 | 2.21E-05 | (A) NE (B/D) |
| DSN1 | 5.95 | 4.31 | 0.71 | 1.29 | 1.64 | 3.34E-05 | 4.60E-04 | (A) NE (B/D) |
| MCM3AP-AS | 8.56 | 6.93 | 0.73 | 0.91 | 1.64 | 4.16E-05 | 5.47E-04 | (A) NE (B/D) |
| SNORA41 | 4.71 | 3.07 | 0.74 | 1.05 | 1.64 | 4.35E-05 | 5.66E-04 | (A) NE (B/D) |
| MAP1S | 1.75 | 0.11 | 0.64 | 0.24 | 1.64 | 4.64E-05 | 5.95E-04 | (A) NE (B/D) |
| SIN3B | 2.31 | 0.67 | 0.75 | 0.85 | 1.64 | 5.57E-05 | 6.89E-04 | (A) NE (B/D) |
| TMEM184A | 4.16 | 2.51 | 0.77 | 1.23 | 1.64 | 7.71E-05 | 8.87E-04 | (A) NE (B/D) |
| ARX | 3.18 | 1.54 | 0.79 | 1.42 | 1.64 | 1.31E-04 | 1.33E-03 | (A) NE (B/D) |
| CBARP | 2.56 | 0.92 | 0.99 | 0.92 | 1.64 | 7.27E-04 | 4.98E-03 | (A) NE (B/D) |
| RPAIN | 2.26 | 0.62 | 0.98 | 0.83 | 1.64 | 7.45E-04 | 5.07E-03 | (A) NE (B/D) |
| ANKRD10-IT1 | 7.44 | 5.79 | 1.01 | 0.71 | 1.64 | 9.27E-04 | 5.96E-03 | (A) NE (B/D) |
| IQCN | 5.57 | 3.93 | 1.06 | 1.29 | 1.64 | 1.32E-03 | 7.80E-03 | (A) NE (B/D) |
| ID2 | 1.88 | 0.24 | 1.06 | 0.69 | 1.64 | 1.39E-03 | 8.11E-03 | (A) NE (B/D) |
| SNORD113-2 | 1.94 | 0.3 | 1.52 | 0.68 | 1.64 | 0.0119 | 0.0404 | (A) NE (B/D) |
| ZNF224 | 2.87 | 1.23 | 1.56 | 0.94 | 1.64 | 0.0139 | 0.0452 | (A) NE (B/D) |
| NELFE | 4.21 | 2.58 | 0.23 | 1.07 | 1.63 | 4.24E-10 | 7.48E-08 | (A) NE (B/D) |
| MLLT11 | 3.26 | 1.64 | 0.35 | 0.87 | 1.63 | 5.48E-10 | 9.21E-08 | (A) NE (B/D) |
| SAFB | 7.51 | 5.88 | 0.36 | 0.72 | 1.63 | 5.87E-10 | 9.61E-08 | (A) NE (B/D) |
| ANKRD23 | 2.38 | 0.75 | 0.37 | 0.72 | 1.63 | 1.02E-09 | 1.38E-07 | (A) NE (B/D) |
| DDX39B | 8.07 | 6.44 | 0.28 | 1.14 | 1.63 | 2.83E-09 | 3.05E-07 | (A) NE (B/D) |
| CSNK2B | 10.16 | 8.53 | 0.27 | 1.23 | 1.63 | 1.15E-08 | 8.87E-07 | (A) NE (B/D) |
| UCKL1 | 3.91 | 2.28 | 0.44 | 0.96 | 1.63 | 3.64E-08 | 2.16E-06 | (A) NE (B/D) |
| TFPT | 5.01 | 3.38 | 0.47 | 1.08 | 1.63 | 1.48E-07 | 6.59E-06 | (A) NE (B/D) |
| TP53INP2 | 6.35 | 4.72 | 0.47 | 1.11 | 1.63 | 1.57E-07 | 6.90E-06 | (A) NE (B/D) |
| ZFYVE19 | 6.68 | 5.05 | 0.53 | 0.92 | 1.63 | 6.16E-07 | 2.02E-05 | (A) NE (B/D) |
| SS18L1 | 3 | 1.36 | 0.58 | 0.89 | 1.63 | 2.05E-06 | 5.24E-05 | (A) NE (B/D) |
| ZMYND10-AS | 5.19 | 3.55 | 0.61 | 1.49 | 1.63 | 1.78E-05 | 2.82E-04 | (A) NE (B/D) |
| ZDHHC23 | 4.81 | 3.19 | 0.67 | 1.21 | 1.63 | 1.79E-05 | 2.84E-04 | (A) NE (B/D) |
| PITPNA-AS1 | 9.56 | 7.92 | 0.67 | 1.42 | 1.63 | 2.91E-05 | 4.12E-04 | (A) NE (B/D) |
| ANP32B | 3.7 | 2.08 | 0.47 | 1.83 | 1.63 | 3.74E-05 | 5.04E-04 | (A) NE (B/D) |
| FAM241B | 5.07 | 3.44 | 0.55 | 1.87 | 1.63 | 7.52E-05 | 8.70E-04 | (A) NE (B/D) |
| TRIM2 | 3.7 | 2.06 | 0.67 | 1.73 | 1.63 | 9.75E-05 | 1.06E-03 | (A) NE (B/D) |
| TMEM81 | 3.98 | 2.35 | 0.87 | 0.77 | 1.63 | 2.88E-04 | 2.44E-03 | (A) NE (B/D) |
| CHEK1 | 2.31 | 0.68 | 0.91 | 0.79 | 1.63 | 4.14E-04 | 3.23E-03 | (A) NE (B/D) |
| P2RY11 | 2.3 | 0.67 | 0.92 | 0.94 | 1.63 | 4.28E-04 | 3.32E-03 | (A) NE (B/D) |
| KCNC1 | 1.65 | 0.02 | 0.86 | 0.09 | 1.63 | 4.73E-04 | 3.58E-03 | (A) NE (B/D) |
| CTC1 | 3.23 | 1.6 | 0.94 | 1.18 | 1.63 | 5.26E-04 | 3.88E-03 | (A) NE (B/D) |
| RUNX1T1 | 1.65 | 0.02 | 1.01 | 0.12 | 1.63 | 1.31E-03 | 7.74E-03 | (A) NE (B/D) |
| RN7SL608P | 4.71 | 3.08 | 1.09 | 0.76 | 1.63 | 1.73E-03 | 9.59E-03 | (A) NE (B/D) |
| RPL26P37 | 3.11 | 1.49 | 1.11 | 1.22 | 1.63 | 1.95E-03 | 0.0105 | (A) NE (B/D) |
| SPP1 | 2.9 | 1.27 | 1.18 | 1.25 | 1.63 | 2.82E-03 | 0.0139 | (A) NE (B/D) |
| RN7SL262P | 3.53 | 1.9 | 1.19 | 1.12 | 1.63 | 3.16E-03 | 0.0152 | (A) NE (B/D) |
| RNF224 | 4.37 | 2.74 | 1.16 | 1.64 | 1.63 | 3.19E-03 | 0.0153 | (A) NE (B/D) |
| ANOS2P | 3.52 | 1.89 | 1.24 | 1.4 | 1.63 | 4.30E-03 | 0.0192 | (A) NE (B/D) |
| SLC24A5 | 1.77 | 0.14 | 1.28 | 0.31 | 1.63 | 5.01E-03 | 0.0215 | (A) NE (B/D) |
| DUSP6 | 1.75 | 0.12 | 1.31 | 0.39 | 1.63 | 5.52E-03 | 0.0231 | (A) NE (B/D) |
| PHLDA2 | 2.3 | 0.66 | 1.51 | 0.85 | 1.63 | 0.0118 | 0.0402 | (A) NE (B/D) |
| SYNGAP1 | 3.89 | 2.27 | 0.3 | 0.84 | 1.62 | 3.21E-11 | 9.48E-09 | (A) NE (B/D) |

| | | | | | | | | |
|---|---|---|---|---|---|---|---|---|
| INTS11 | 2.82 | 1.2 | 0.44 | 0.88 | 1.62 | 3.22E-08 | 1.96E-06 | (A) NE (B/D) |
| MDC1-AS1 | 8.02 | 6.4 | 0.46 | 0.87 | 1.62 | 6.33E-08 | 3.34E-06 | (A) NE (B/D) |
| TBC1D8 | 5.67 | 4.05 | 0.39 | 1.25 | 1.62 | 1.04E-07 | 4.98E-06 | (A) NE (B/D) |
| PPP1R26 | 7.98 | 6.36 | 0.52 | 1.16 | 1.62 | 1.03E-06 | 3.06E-05 | (A) NE (B/D) |
| KLC2 | 3.43 | 1.81 | 0.6 | 1.19 | 1.62 | 5.28E-06 | 1.10E-04 | (A) NE (B/D) |
| NPIPA7 | 10.12 | 8.5 | 0.58 | 1.4 | 1.62 | 9.17E-06 | 1.69E-04 | (A) NE (B/D) |
| SURF2 | 5.59 | 3.97 | 0.64 | 1.59 | 1.62 | 4.65E-05 | 5.95E-04 | (A) NE (B/D) |
| KIF4B | 2.68 | 1.06 | 0.75 | 1.14 | 1.62 | 6.14E-05 | 7.42E-04 | (A) NE (B/D) |
| H1FX-AS1 | 2.36 | 0.74 | 0.77 | 0.81 | 1.62 | 8.74E-05 | 9.80E-04 | (A) NE (B/D) |
| MIB2 | 3.31 | 1.69 | 0.76 | 0.64 | 1.62 | 9.64E-05 | 1.06E-03 | (A) NE (B/D) |
| ELP1 | 8.77 | 7.15 | 0.79 | 0.93 | 1.62 | 1.09E-04 | 1.16E-03 | (A) NE (B/D) |
| OSGEP | 2.31 | 0.7 | 0.81 | 0.77 | 1.62 | 1.68E-04 | 1.61E-03 | (A) NE (B/D) |
| LYPD5 | 2.37 | 0.75 | 0.83 | 0.74 | 1.62 | 2.12E-04 | 1.92E-03 | (A) NE (B/D) |
| TEAD1 | 2.27 | 0.65 | 0.88 | 0.73 | 1.62 | 3.62E-04 | 2.91E-03 | (A) NE (B/D) |
| MIR-631/631 | 7.47 | 5.85 | 0.91 | 1.19 | 1.62 | 4.23E-04 | 3.29E-03 | (A) NE (B/D) |
| H2AFY2 | 7.95 | 6.33 | 0.86 | 1.67 | 1.62 | 4.60E-04 | 3.51E-03 | (A) NE (B/D) |
| BIRC5 | 2.03 | 0.41 | 0.93 | 0.85 | 1.62 | 4.91E-04 | 3.69E-03 | (A) NE (B/D) |
| PLCG1 | 9.25 | 7.63 | 0.94 | 1.21 | 1.62 | 5.38E-04 | 3.95E-03 | (A) NE (B/D) |
| NDRG1 | 5.33 | 3.7 | 0.89 | 1.77 | 1.62 | 7.31E-04 | 5.00E-03 | (A) NE (B/D) |
| TMEM160 | 1.77 | 0.15 | 1 | 0.29 | 1.62 | 1.19E-03 | 7.23E-03 | (A) NE (B/D) |
| RPL17P1 | 1.64 | 0.02 | 1.02 | 0.07 | 1.62 | 1.39E-03 | 8.12E-03 | (A) NE (B/D) |
| CCDC180 | 3.85 | 2.23 | 1.11 | 0.93 | 1.62 | 1.91E-03 | 0.0104 | (A) NE (B/D) |
| HOXA5 | 1.76 | 0.13 | 1.1 | 0.49 | 1.62 | 2.04E-03 | 0.0109 | (A) NE (B/D) |
| CAPRIN2 | 2.19 | 0.57 | 1.14 | 0.84 | 1.62 | 2.37E-03 | 0.0122 | (A) NE (B/D) |
| ARGLU1 | 2.76 | 1.13 | 1.2 | 0.98 | 1.62 | 3.26E-03 | 0.0155 | (A) NE (B/D) |
| DDO | 5.57 | 3.95 | 1.18 | 1.68 | 1.62 | 3.81E-03 | 0.0175 | (A) NE (B/D) |
| TMTC1 | 1.63 | 0.02 | 1.28 | 0.1 | 1.62 | 5.28E-03 | 0.0224 | (A) NE (B/D) |
| CCNO | 3.92 | 2.3 | 1.22 | 1.81 | 1.62 | 5.33E-03 | 0.0225 | (A) NE (B/D) |
| PANK4 | 5.66 | 4.05 | 0.31 | 0.63 | 1.61 | 2.71E-11 | 8.34E-09 | (A) NE (B/D) |
| NEURL2 | 5.29 | 3.68 | 0.36 | 0.77 | 1.61 | 7.11E-10 | 1.09E-07 | (A) NE (B/D) |
| SKIV2L | 6.96 | 5.36 | 0.36 | 1 | 1.61 | 3.70E-09 | 3.69E-07 | (A) NE (B/D) |
| IDH3G | 2.91 | 1.31 | 0.24 | 1.26 | 1.61 | 2.07E-08 | 1.36E-06 | (A) NE (B/D) |
| PRMT1 | 8.2 | 6.59 | 0.39 | 1.17 | 1.61 | 5.20E-08 | 2.88E-06 | (A) NE (B/D) |
| GTPBP6 | 5.09 | 3.48 | 0.33 | 1.26 | 1.61 | 5.42E-08 | 2.95E-06 | (A) NE (B/D) |
| WDR90 | 5.44 | 3.83 | 0.49 | 1 | 1.61 | 2.40E-07 | 9.61E-06 | (A) NE (B/D) |
| NPIPA2 | 7.32 | 5.71 | 0.42 | 1.35 | 1.61 | 5.60E-07 | 1.89E-05 | (A) NE (B/D) |
| NCALD | 2.62 | 1 | 0.51 | 1.3 | 1.61 | 1.56E-06 | 4.25E-05 | (A) NE (B/D) |
| MRPL54 | 4.56 | 2.96 | 0.28 | 1.56 | 1.61 | 1.58E-06 | 4.29E-05 | (A) NE (B/D) |
| COX5BP6 | 4.67 | 3.06 | 0.59 | 1.3 | 1.61 | 6.94E-06 | 1.36E-04 | (A) NE (B/D) |
| CBX8 | 2.48 | 0.87 | 0.68 | 0.96 | 1.61 | 2.04E-05 | 3.13E-04 | (A) NE (B/D) |
| MAPRE2 | 2.11 | 0.5 | 0.69 | 0.73 | 1.61 | 3.30E-05 | 4.57E-04 | (A) NE (B/D) |
| ABHD17AP3 | 3.83 | 2.22 | 0.78 | 1.06 | 1.61 | 1.06E-04 | 1.14E-03 | (A) NE (B/D) |
| MSANTD3-TI | 3.76 | 2.15 | 0.93 | 1.25 | 1.61 | 5.52E-04 | 4.04E-03 | (A) NE (B/D) |
| RPS4XP1 | 7.81 | 6.21 | 0.8 | 2.08 | 1.61 | 9.48E-04 | 6.06E-03 | (A) NE (B/D) |
| KCNMB3 | 2.92 | 1.3 | 1.16 | 0.74 | 1.61 | 2.77E-03 | 0.0138 | (A) NE (B/D) |
| ERFE | 2.49 | 0.88 | 1.24 | 0.99 | 1.61 | 4.31E-03 | 0.0193 | (A) NE (B/D) |
| VKORC1 | 2.97 | 1.36 | 1.4 | 1.46 | 1.61 | 9.51E-03 | 0.0344 | (A) NE (B/D) |
| CLCN4 | 5.06 | 3.45 | 1.44 | 1.2 | 1.61 | 0.0105 | 0.0368 | (A) NE (B/D) |
| ZNF302 | 7.36 | 5.75 | 1.56 | 1.17 | 1.61 | 0.0155 | 0.0491 | (A) NE (B/D) |
| AGPAT1 | 2.56 | 0.96 | 0.26 | 0.97 | 1.6 | 1.45E-10 | 3.30E-08 | (A) NE (B/D) |
| STK19 | 7 | 5.4 | 0.31 | 0.98 | 1.6 | 5.82E-10 | 9.60E-08 | (A) NE (B/D) |
| GTF2F1 | 6.79 | 5.19 | 0.38 | 0.97 | 1.6 | 6.12E-09 | 5.45E-07 | (A) NE (B/D) |
| PIDD1 | 1.94 | 0.34 | 0.43 | 0.57 | 1.6 | 7.25E-08 | 3.73E-06 | (A) NE (B/D) |
| SPINT1 | 2.74 | 1.14 | 0.51 | 0.97 | 1.6 | 5.12E-07 | 1.75E-05 | (A) NE (B/D) |
| RING1 | 5.52 | 3.92 | 0.45 | 1.33 | 1.6 | 9.33E-07 | 2.81E-05 | (A) NE (B/D) |
| FOXK1 | 2.27 | 0.67 | 0.51 | 0.51 | 1.6 | 1.52E-06 | 4.17E-05 | (A) NE (B/D) |
| FAM50A | 10.05 | 8.45 | 0.55 | 1.36 | 1.6 | 5.82E-06 | 1.19E-04 | (A) NE (B/D) |
| PARP2 | 3.66 | 2.05 | 0.67 | 1.12 | 1.6 | 1.86E-05 | 2.92E-04 | (A) NE (B/D) |
| ANP32BP1 | 9.53 | 7.93 | 0.48 | 1.86 | 1.6 | 6.00E-05 | 7.28E-04 | (A) NE (B/D) |
| ZWINT | 3.15 | 1.55 | 0.83 | 1.54 | 1.6 | 3.35E-04 | 2.75E-03 | (A) NE (B/D) |
| AMH | 1.75 | 0.15 | 0.93 | 0.48 | 1.6 | 7.12E-04 | 4.91E-03 | (A) NE (B/D) |
| DRP2 | 1.89 | 0.29 | 1.2 | 0.61 | 1.6 | 3.64E-03 | 0.0169 | (A) NE (B/D) |
| LCORL | 2.27 | 0.68 | 1.41 | 0.76 | 1.6 | 9.21E-03 | 0.0336 | (A) NE (B/D) |
| SLC18A3 | 1.76 | 0.16 | 1.56 | 0.39 | 1.6 | 0.0154 | 0.0488 | (A) NE (B/D) |
| AKAP17A | 2.45 | 0.86 | 0.34 | 0.83 | 1.59 | 4.91E-10 | 8.58E-08 | (A) NE (B/D) |

| | | | | | | | | |
|---|---|---|---|---|---|---|---|---|
| ATF6B | 5.36 | 3.77 | 0.3 | 1.3 | 1.59 | 8.95E-08 | 4.38E-06 | (A) NE (B/D) |
| CTPS1 | 6.82 | 5.24 | 0.47 | 1.05 | 1.59 | 2.32E-07 | 9.34E-06 | (A) NE (B/D) |
| PRKACA | 3.96 | 2.37 | 0.26 | 1.4 | 1.59 | 2.41E-07 | 9.61E-06 | (A) NE (B/D) |
| PLD3 | 8.75 | 7.16 | 0.48 | 1.05 | 1.59 | 3.08E-07 | 1.17E-05 | (A) NE (B/D) |
| LSM4 | 3.35 | 1.76 | 0.5 | 0.92 | 1.59 | 3.53E-07 | 1.29E-05 | (A) NE (B/D) |
| TRMT1 | 7.76 | 6.18 | 0.52 | 1.26 | 1.59 | 2.09E-06 | 5.30E-05 | (A) NE (B/D) |
| RNF167 | 3.73 | 2.14 | 0.55 | 1.6 | 1.59 | 2.39E-05 | 3.51E-04 | (A) NE (B/D) |
| FAM86JP | 4.51 | 2.92 | 0.56 | 1.65 | 1.59 | 3.49E-05 | 4.77E-04 | (A) NE (B/D) |
| HNRNPA1P48 | 8.4 | 6.82 | 0.55 | 1.75 | 1.59 | 5.80E-05 | 7.10E-04 | (A) NE (B/D) |
| EMILIN3 | 3.51 | 1.93 | 0.72 | 1.28 | 1.59 | 6.50E-05 | 7.75E-04 | (A) NE (B/D) |
| TMEM181 | 3 | 1.41 | 0.78 | 0.53 | 1.59 | 1.76E-04 | 1.67E-03 | (A) NE (B/D) |
| ALKBH7 | 1.86 | 0.27 | 0.84 | 0.55 | 1.59 | 3.18E-04 | 2.64E-03 | (A) NE (B/D) |
| HOXD-AS2 | 1.7 | 0.11 | 0.9 | 0.23 | 1.59 | 6.76E-04 | 4.72E-03 | (A) NE (B/D) |
| PPIAP29 | 9.3 | 7.71 | 0.91 | 1.67 | 1.59 | 8.54E-04 | 5.62E-03 | (A) NE (B/D) |
| ITGB3BP | 3.04 | 1.45 | 1 | 0.89 | 1.59 | 1.06E-03 | 6.59E-03 | (A) NE (B/D) |
| DCAF12L2 | 1.7 | 0.11 | 1.02 | 0.44 | 1.59 | 1.41E-03 | 8.20E-03 | (A) NE (B/D) |
| RN7SL221P | 2.62 | 1.02 | 1.06 | 1.08 | 1.59 | 1.55E-03 | 8.83E-03 | (A) NE (B/D) |
| SLC25A5P5 | 3.59 | 1.99 | 1.05 | 1.34 | 1.59 | 1.57E-03 | 8.93E-03 | (A) NE (B/D) |
| BPHL | 6.86 | 5.27 | 1.09 | 1.19 | 1.59 | 2.02E-03 | 0.0108 | (A) NE (B/D) |
| PCDHGA9 | 3.62 | 2.04 | 1.2 | 1.19 | 1.59 | 3.80E-03 | 0.0175 | (A) NE (B/D) |
| TEKT3 | 1.65 | 0.06 | 1.26 | 0.21 | 1.59 | 5.45E-03 | 0.0229 | (A) NE (B/D) |
| ZIC2 | 3.49 | 1.9 | 1.32 | 1.27 | 1.59 | 6.82E-03 | 0.0271 | (A) NE (B/D) |
| PCDHGA11 | 4.9 | 3.32 | 1.31 | 1.48 | 1.59 | 7.43E-03 | 0.0288 | (A) NE (B/D) |
| RPS11P7 | 2.33 | 0.74 | 1.37 | 1.26 | 1.59 | 8.62E-03 | 0.032 | (A) NE (B/D) |
| RNPEPL1 | 3.71 | 2.13 | 0.12 | 0.9 | 1.58 | 6.43E-12 | 2.90E-09 | (A) NE (B/D) |
| TRIM27 | 13.89 | 12.32 | 0.39 | 0.67 | 1.58 | 6.58E-09 | 5.76E-07 | (A) NE (B/D) |
| RTEL1 | 4.84 | 3.25 | 0.4 | 0.73 | 1.58 | 8.13E-09 | 6.74E-07 | (A) NE (B/D) |
| MIR-1181/11 | 9.53 | 7.95 | 0.48 | 1.07 | 1.58 | 3.97E-07 | 1.43E-05 | (A) NE (B/D) |
| MRPS5 | 3.13 | 1.54 | 0.52 | 1.07 | 1.58 | 8.54E-07 | 2.62E-05 | (A) NE (B/D) |
| GOLGA6L5P | 6.2 | 4.61 | 0.46 | 1.45 | 1.58 | 3.27E-06 | 7.52E-05 | (A) NE (B/D) |
| FBXW5 | 8.24 | 6.66 | 0.58 | 0.91 | 1.58 | 3.96E-06 | 8.80E-05 | (A) NE (B/D) |
| FUT5 | 4.45 | 2.87 | 0.56 | 1.32 | 1.58 | 6.02E-06 | 1.22E-04 | (A) NE (B/D) |
| YBX1P2 | 6.29 | 4.72 | 0.65 | 1.21 | 1.58 | 1.95E-05 | 3.02E-04 | (A) NE (B/D) |
| SSNA1 | 4.25 | 2.68 | 0.68 | 1.43 | 1.58 | 5.99E-05 | 7.28E-04 | (A) NE (B/D) |
| KIAA1841 | 2.67 | 1.09 | 0.76 | 0.99 | 1.58 | 9.23E-05 | 1.02E-03 | (A) NE (B/D) |
| HNRNPA3 | 9.14 | 7.56 | 0.79 | 1.12 | 1.58 | 1.43E-04 | 1.42E-03 | (A) NE (B/D) |
| PTRHD1 | 6.18 | 4.6 | 0.79 | 1.55 | 1.58 | 2.48E-04 | 2.17E-03 | (A) NE (B/D) |
| PKD1P5 | 2.93 | 1.35 | 0.82 | 0.65 | 1.58 | 2.54E-04 | 2.20E-03 | (A) NE (B/D) |
| FOXS1 | 4.31 | 2.73 | 0.79 | 1.89 | 1.58 | 6.27E-04 | 4.45E-03 | (A) NE (B/D) |
| RPL10P1 | 1.88 | 0.3 | 1.13 | 0.5 | 1.58 | 2.77E-03 | 0.0138 | (A) NE (B/D) |
| TGM3 | 4.41 | 2.83 | 1.22 | 1.97 | 1.58 | 7.21E-03 | 0.0282 | (A) NE (B/D) |
| IZUMO4 | 1.93 | 0.35 | 1.34 | 0.64 | 1.58 | 7.58E-03 | 0.0291 | (A) NE (B/D) |
| LEMD2 | 5.15 | 3.58 | 0.25 | 0.71 | 1.57 | 7.40E-13 | 6.80E-10 | (A) NE (B/D) |
| SPATA25 | 5.45 | 3.88 | 0.36 | 0.86 | 1.57 | 1.89E-09 | 2.24E-07 | (A) NE (B/D) |
| SCMH1 | 2.5 | 0.93 | 0.38 | 0.99 | 1.57 | 1.39E-08 | 1.00E-06 | (A) NE (B/D) |
| POLR2A | 4.01 | 2.44 | 0.38 | 1.15 | 1.57 | 6.43E-08 | 3.36E-06 | (A) NE (B/D) |
| FLII | 8.61 | 7.04 | 0.48 | 0.9 | 1.57 | 2.59E-07 | 1.03E-05 | (A) NE (B/D) |
| EHMT2-AS1 | 7.81 | 6.25 | 0.46 | 1.28 | 1.57 | 1.05E-06 | 3.11E-05 | (A) NE (B/D) |
| PKM | 5.11 | 3.54 | 0.57 | 1.06 | 1.57 | 3.78E-06 | 8.49E-05 | (A) NE (B/D) |
| MAD2L2 | 2.66 | 1.08 | 0.6 | 1.05 | 1.57 | 6.21E-06 | 1.25E-04 | (A) NE (B/D) |
| AKR1A1 | 4.81 | 3.24 | 0.45 | 1.58 | 1.57 | 8.55E-06 | 1.61E-04 | (A) NE (B/D) |
| ARAP1-AS1 | 6.18 | 4.61 | 0.46 | 1.58 | 1.57 | 9.82E-06 | 1.79E-04 | (A) NE (B/D) |
| PABPN1 | 5.94 | 4.37 | 0.61 | 1.44 | 1.57 | 2.38E-05 | 3.50E-04 | (A) NE (B/D) |
| SRSF4 | 5.77 | 4.19 | 0.66 | 1.25 | 1.57 | 2.55E-05 | 3.71E-04 | (A) NE (B/D) |
| NAPB | 8.05 | 6.47 | 0.65 | 0.6 | 1.57 | 2.99E-05 | 4.22E-04 | (A) NE (B/D) |
| ZDHHC13 | 3.66 | 2.1 | 0.7 | 0.98 | 1.57 | 4.15E-05 | 5.46E-04 | (A) NE (B/D) |
| TRNAU1AP | 7.12 | 5.55 | 0.76 | 0.97 | 1.57 | 9.56E-05 | 1.05E-03 | (A) NE (B/D) |
| PPP1R37 | 1.64 | 0.07 | 0.72 | 0.16 | 1.57 | 1.70E-04 | 1.63E-03 | (A) NE (B/D) |
| CLDN12 | 4.2 | 2.62 | 0.85 | 1.35 | 1.57 | 3.59E-04 | 2.90E-03 | (A) NE (B/D) |
| PRSS44P | 1.78 | 0.21 | 0.96 | 0.5 | 1.57 | 1.04E-03 | 6.47E-03 | (A) NE (B/D) |
| TK1 | 3.26 | 1.69 | 0.98 | 1.72 | 1.57 | 1.67E-03 | 9.33E-03 | (A) NE (B/D) |
| FST | 1.7 | 0.13 | 1.12 | 0.5 | 1.57 | 2.74E-03 | 0.0137 | (A) NE (B/D) |
| ZNF311 | 2.98 | 1.41 | 1.23 | 1.13 | 1.57 | 4.78E-03 | 0.0208 | (A) NE (B/D) |
| SCML2 | 5.75 | 4.18 | 1.25 | 1.27 | 1.57 | 5.52E-03 | 0.0231 | (A) NE (B/D) |
| SMIM22 | 2.89 | 1.32 | 1.28 | 1.48 | 1.57 | 6.75E-03 | 0.0269 | (A) NE (B/D) |

| | | | | | | | | |
|---|---|---|---|---|---|---|---|---|
| ZNF334 | 2.43 | 0.87 | 1.43 | 0.98 | 1.57 | 0.0111 | 0.0383 | (A) NE (B/D) |
| PNMA8B | 4.26 | 2.69 | 1.45 | 1.08 | 1.57 | 0.012 | 0.0406 | (A) NE (B/D) |
| RPL5P30 | 5.32 | 3.75 | 1.51 | 0.97 | 1.57 | 0.0146 | 0.0469 | (A) NE (B/D) |
| CDH22 | 1.83 | 0.26 | 1.53 | 0.72 | 1.57 | 0.0154 | 0.0489 | (A) NE (B/D) |
| RASA4CP | 7.68 | 6.12 | 0.35 | 0.95 | 1.56 | 3.05E-09 | 3.25E-07 | (A) NE (B/D) |
| SNORD110 | 5.93 | 4.37 | 0.27 | 1.16 | 1.56 | 1.12E-08 | 8.72E-07 | (A) NE (B/D) |
| IDH3B | 8.37 | 6.81 | 0.36 | 1.15 | 1.56 | 3.97E-08 | 2.31E-06 | (A) NE (B/D) |
| C11ORF94 | 3.61 | 2.05 | 0.55 | 1.17 | 1.56 | 3.81E-06 | 8.55E-05 | (A) NE (B/D) |
| ME3 | 3.81 | 2.25 | 0.49 | 1.46 | 1.56 | 6.64E-06 | 1.32E-04 | (A) NE (B/D) |
| MRPS6 | 4.65 | 3.09 | 0.61 | 1.23 | 1.56 | 1.31E-05 | 2.26E-04 | (A) NE (B/D) |
| GOLGA6L3 | 2.55 | 0.99 | 0.72 | 1 | 1.56 | 6.08E-05 | 7.37E-04 | (A) NE (B/D) |
| HNRNPA1P10 | 8.41 | 6.86 | 0.67 | 1.58 | 1.56 | 1.02E-04 | 1.11E-03 | (A) NE (B/D) |
| PTGES3L | 2.7 | 1.14 | 0.76 | 0.84 | 1.56 | 1.13E-04 | 1.19E-03 | (A) NE (B/D) |
| CCDC163 | 5.51 | 3.95 | 0.79 | 1.04 | 1.56 | 1.57E-04 | 1.53E-03 | (A) NE (B/D) |
| PRELID1P4 | 3.95 | 2.4 | 0.75 | 1.81 | 1.56 | 4.35E-04 | 3.36E-03 | (A) NE (B/D) |
| INPP5J | 3.43 | 1.87 | 0.92 | 1.43 | 1.56 | 7.85E-04 | 5.29E-03 | (A) NE (B/D) |
| GNAI1 | 2.11 | 0.55 | 0.96 | 0.83 | 1.56 | 9.11E-04 | 5.89E-03 | (A) NE (B/D) |
| CRLF1 | 6.14 | 4.58 | 1.06 | 0.87 | 1.56 | 1.82E-03 | 9.99E-03 | (A) NE (B/D) |
| FAM163B | 1.6 | 0.04 | 1.15 | 0.23 | 1.56 | 3.55E-03 | 0.0166 | (A) NE (B/D) |
| TUBA4A | 4.5 | 2.94 | 1.23 | 1.39 | 1.56 | 5.28E-03 | 0.0224 | (A) NE (B/D) |
| DLEU7-AS1 | 2.02 | 0.47 | 1.28 | 0.76 | 1.56 | 6.26E-03 | 0.0254 | (A) NE (B/D) |
| FBLN7 | 4.93 | 3.37 | 1.32 | 1.46 | 1.56 | 8.35E-03 | 0.0312 | (A) NE (B/D) |
| CHST2 | 4.15 | 2.59 | 1.33 | 1.52 | 1.56 | 8.71E-03 | 0.0322 | (A) NE (B/D) |
| HOOK1 | 4.61 | 3.05 | 1.37 | 1.08 | 1.56 | 9.01E-03 | 0.0331 | (A) NE (B/D) |
| KRT8P3 | 5.55 | 3.99 | 1.42 | 1.7 | 1.56 | 0.0135 | 0.0444 | (A) NE (B/D) |
| BUD23 | 3.52 | 1.98 | 0.12 | 0.96 | 1.55 | 6.50E-11 | 1.67E-08 | (A) NE (B/D) |
| QSOX2 | 7.85 | 6.3 | 0.28 | 1.13 | 1.55 | 8.26E-09 | 6.79E-07 | (A) NE (B/D) |
| ATP6V1G2-D | 9.93 | 8.37 | 0.21 | 1.21 | 1.55 | 1.66E-08 | 1.15E-06 | (A) NE (B/D) |
| LINC01003 | 3.96 | 2.42 | 0.4 | 0.95 | 1.55 | 2.12E-08 | 1.38E-06 | (A) NE (B/D) |
| ZSCAN16 | 3.22 | 1.68 | 0.37 | 1.2 | 1.55 | 1.20E-07 | 5.57E-06 | (A) NE (B/D) |
| FAAP20 | 4.44 | 2.88 | 0.46 | 1.02 | 1.55 | 1.96E-07 | 8.21E-06 | (A) NE (B/D) |
| ALKBH6 | 2.67 | 1.12 | 0.49 | 0.86 | 1.55 | 3.39E-07 | 1.26E-05 | (A) NE (B/D) |
| CALM1 | 8.21 | 6.66 | 0.5 | 0.92 | 1.55 | 4.47E-07 | 1.58E-05 | (A) NE (B/D) |
| MAN2C1 | 3.12 | 1.57 | 0.51 | 0.85 | 1.55 | 6.98E-07 | 2.23E-05 | (A) NE (B/D) |
| ATP6V1F | 3.29 | 1.74 | 0.45 | 1.45 | 1.55 | 4.50E-06 | 9.70E-05 | (A) NE (B/D) |
| SDHDP6 | 2.6 | 1.04 | 0.59 | 0.75 | 1.55 | 6.46E-06 | 1.29E-04 | (A) NE (B/D) |
| TCEA2 | 2.59 | 1.04 | 0.56 | 1.26 | 1.55 | 7.13E-06 | 1.40E-04 | (A) NE (B/D) |
| UXT | 4.47 | 2.91 | 0.59 | 1.82 | 1.55 | 1.47E-04 | 1.46E-03 | (A) NE (B/D) |
| LRRFIP1P1 | 6.86 | 5.31 | 0.84 | 0.77 | 1.55 | 3.14E-04 | 2.61E-03 | (A) NE (B/D) |
| OR8Q1P | 1.58 | 0.04 | 0.83 | 0.11 | 1.55 | 5.12E-04 | 3.80E-03 | (A) NE (B/D) |
| HDAC1 | 6.53 | 4.99 | 0.91 | 1.44 | 1.55 | 8.16E-04 | 5.45E-03 | (A) NE (B/D) |
| SLC26A11 | 6.14 | 4.59 | 0.96 | 1.12 | 1.55 | 8.82E-04 | 5.75E-03 | (A) NE (B/D) |
| PTOV1-AS1 | 7.1 | 5.55 | 1.04 | 0.84 | 1.55 | 1.74E-03 | 9.63E-03 | (A) NE (B/D) |
| LGALS3BP | 2.62 | 1.07 | 1.07 | 1.2 | 1.55 | 2.15E-03 | 0.0113 | (A) NE (B/D) |
| THOP1 | 4.6 | 3.06 | 0.18 | 1.06 | 1.54 | 8.40E-10 | 1.23E-07 | (A) NE (B/D) |
| CACTIN-AS1 | 4.07 | 2.53 | 0.35 | 0.81 | 1.54 | 1.06E-09 | 1.42E-07 | (A) NE (B/D) |
| RTEL1-TNFRS | 3.02 | 1.48 | 0.36 | 0.69 | 1.54 | 1.87E-09 | 2.23E-07 | (A) NE (B/D) |
| ZNF317 | 4.98 | 3.44 | 0.42 | 0.68 | 1.54 | 4.43E-08 | 2.51E-06 | (A) NE (B/D) |
| DENND6B | 5.92 | 4.39 | 0.34 | 1.22 | 1.54 | 9.44E-08 | 4.58E-06 | (A) NE (B/D) |
| IFNGR2 | 4.27 | 2.72 | 0.45 | 0.97 | 1.54 | 1.39E-07 | 6.30E-06 | (A) NE (B/D) |
| MIR-636/636 | 2.77 | 1.23 | 0.47 | 0.89 | 1.54 | 2.31E-07 | 9.34E-06 | (A) NE (B/D) |
| SLC25A4 | 5.13 | 3.59 | 0.45 | 1.26 | 1.54 | 9.97E-07 | 2.98E-05 | (A) NE (B/D) |
| UBL5 | 3.77 | 2.23 | 0.24 | 1.52 | 1.54 | 1.69E-06 | 4.52E-05 | (A) NE (B/D) |
| YPEL1 | 1.98 | 0.44 | 0.55 | 0.71 | 1.54 | 3.53E-06 | 8.04E-05 | (A) NE (B/D) |
| TUBB | 11 | 9.47 | 0.48 | 1.48 | 1.54 | 8.68E-06 | 1.63E-04 | (A) NE (B/D) |
| ILVBL | 1.6 | 0.06 | 0.51 | 0.27 | 1.54 | 9.61E-06 | 1.76E-04 | (A) NE (B/D) |
| SAE1 | 4.7 | 3.15 | 0.57 | 1.29 | 1.54 | 9.81E-06 | 1.79E-04 | (A) NE (B/D) |
| LSM2 | 9.17 | 7.63 | 0.51 | 1.46 | 1.54 | 9.99E-06 | 1.81E-04 | (A) NE (B/D) |
| GRB7 | 2.24 | 0.7 | 0.61 | 0.76 | 1.54 | 1.11E-05 | 1.98E-04 | (A) NE (B/D) |
| HNRNPA3P6 | 12.06 | 10.52 | 0.57 | 1.36 | 1.54 | 1.23E-05 | 2.16E-04 | (A) NE (B/D) |
| ZSWIM5 | 1.91 | 0.38 | 0.57 | 0.37 | 1.54 | 1.88E-05 | 2.94E-04 | (A) NE (B/D) |
| HNRNPD | 4.61 | 3.07 | 0.64 | 1.33 | 1.54 | 3.26E-05 | 4.54E-04 | (A) NE (B/D) |
| LINC01006 | 3.01 | 1.47 | 0.69 | 0.94 | 1.54 | 4.42E-05 | 5.74E-04 | (A) NE (B/D) |
| ATAD3C | 2.77 | 1.23 | 0.73 | 1.05 | 1.54 | 8.45E-05 | 9.53E-04 | (A) NE (B/D) |
| GRAMD2A | 1.71 | 0.16 | 0.7 | 0.3 | 1.54 | 1.25E-04 | 1.29E-03 | (A) NE (B/D) |

| | | | | | | | | |
|---|---|---|---|---|---|---|---|---|
| PCGF2 | 8.22 | 6.68 | 0.63 | 1.71 | 1.54 | 1.35E-04 | 1.36E-03 | (A) NE (B/D) |
| ZNF223 | 5.5 | 3.96 | 0.9 | 1.18 | 1.54 | 6.02E-04 | 4.32E-03 | (A) NE (B/D) |
| CDRT4 | 2.46 | 0.92 | 1.15 | 1.16 | 1.54 | 3.48E-03 | 0.0163 | (A) NE (B/D) |
| C12ORF75 | 8.14 | 6.6 | 1.42 | 1.56 | 1.54 | 0.0137 | 0.0449 | (A) NE (B/D) |
| SLC25A5P1 | 5.96 | 4.42 | 0.26 | 1.08 | 1.53 | 3.20E-09 | 3.39E-07 | (A) NE (B/D) |
| FZR1 | 1.78 | 0.26 | 0.32 | 0.38 | 1.53 | 3.62E-09 | 3.67E-07 | (A) NE (B/D) |
| NDUFB2 | 4.49 | 2.96 | 0.34 | 1.14 | 1.53 | 3.85E-08 | 2.25E-06 | (A) NE (B/D) |
| THAP8 | 2.23 | 0.7 | 0.44 | 0.82 | 1.53 | 9.26E-08 | 4.51E-06 | (A) NE (B/D) |
| CARM1 | 5.47 | 3.93 | 0.46 | 0.83 | 1.53 | 1.56E-07 | 6.90E-06 | (A) NE (B/D) |
| AKAP8L | 2.41 | 0.88 | 0.48 | 0.85 | 1.53 | 3.45E-07 | 1.27E-05 | (A) NE (B/D) |
| NECTIN2 | 2.97 | 1.44 | 0.25 | 1.4 | 1.53 | 4.77E-07 | 1.66E-05 | (A) NE (B/D) |
| CCDC107 | 6.88 | 5.35 | 0.37 | 1.54 | 1.53 | 5.24E-06 | 1.09E-04 | (A) NE (B/D) |
| SNHG11 | 7.3 | 5.77 | 0.53 | 1.41 | 1.53 | 1.12E-05 | 2.00E-04 | (A) NE (B/D) |
| CDCA4 | 2.63 | 1.1 | 0.61 | 1.26 | 1.53 | 1.77E-05 | 2.82E-04 | (A) NE (B/D) |
| MIR-621/621 | 3.62 | 2.08 | 0.65 | 0.99 | 1.53 | 2.19E-05 | 3.28E-04 | (A) NE (B/D) |
| SAPCD1 | 9.01 | 7.47 | 0.39 | 1.8 | 1.53 | 4.40E-05 | 5.72E-04 | (A) NE (B/D) |
| CHAF1A | 3.63 | 2.11 | 0.69 | 1.19 | 1.53 | 5.68E-05 | 7.00E-04 | (A) NE (B/D) |
| ECHDC2 | 4.85 | 3.32 | 0.74 | 1.21 | 1.53 | 1.10E-04 | 1.17E-03 | (A) NE (B/D) |
| SNORD55 | 6.16 | 4.63 | 0.68 | 1.62 | 1.53 | 1.71E-04 | 1.63E-03 | (A) NE (B/D) |
| RPL18A | 3.68 | 2.15 | 0.76 | 1.54 | 1.53 | 2.84E-04 | 2.42E-03 | (A) NE (B/D) |
| PCDHA11 | 1.97 | 0.44 | 0.97 | 0.7 | 1.53 | 1.13E-03 | 6.91E-03 | (A) NE (B/D) |
| TSPYL5 | 2.49 | 0.96 | 1.09 | 0.92 | 1.53 | 2.49E-03 | 0.0127 | (A) NE (B/D) |
| CHST1 | 2.83 | 1.31 | 1.16 | 1.06 | 1.53 | 3.99E-03 | 0.0181 | (A) NE (B/D) |
| MIR-5695/56 | 2.87 | 1.34 | 1.2 | 1.43 | 1.53 | 5.33E-03 | 0.0225 | (A) NE (B/D) |
| RPL21P54 | 1.54 | 0.01 | 1.22 | 0.04 | 1.53 | 5.59E-03 | 0.0233 | (A) NE (B/D) |
| LINC01011 | 6.17 | 4.63 | 1.31 | 0.8 | 1.53 | 7.87E-03 | 0.0299 | (A) NE (B/D) |
| MIR-5047/50 | 10.88 | 9.35 | 1.31 | 1.19 | 1.53 | 8.37E-03 | 0.0313 | (A) NE (B/D) |
| EEF1A1P13 | 10.06 | 8.53 | 1.42 | 1.49 | 1.53 | 0.0134 | 0.0442 | (A) NE (B/D) |
| BAG6 | 2.59 | 1.08 | 0.23 | 0.95 | 1.52 | 2.06E-10 | 4.16E-08 | (A) NE (B/D) |
| NAA40 | 2.24 | 0.72 | 0.41 | 0.84 | 1.52 | 3.28E-08 | 1.99E-06 | (A) NE (B/D) |
| TENT4A | 1.95 | 0.43 | 0.54 | 0.57 | 1.52 | 5.25E-06 | 1.10E-04 | (A) NE (B/D) |
| CHPF | 5.18 | 3.66 | 0.59 | 1.34 | 1.52 | 2.06E-05 | 3.15E-04 | (A) NE (B/D) |
| SUN1 | 2.19 | 0.66 | 0.64 | 0.71 | 1.52 | 2.42E-05 | 3.56E-04 | (A) NE (B/D) |
| SHKBP1 | 5.22 | 3.7 | 0.58 | 1.42 | 1.52 | 2.72E-05 | 3.91E-04 | (A) NE (B/D) |
| FAM83D | 2.99 | 1.48 | 0.81 | 1.33 | 1.52 | 3.23E-04 | 2.67E-03 | (A) NE (B/D) |
| SYTL1 | 5.12 | 3.6 | 0.81 | 1.46 | 1.52 | 3.98E-04 | 3.14E-03 | (A) NE (B/D) |
| ZFP30 | 6.76 | 5.24 | 1.16 | 0.7 | 1.52 | 4.01E-03 | 0.0182 | (A) NE (B/D) |
| CKAP2 | 2.74 | 1.21 | 1.24 | 1.15 | 1.52 | 6.22E-03 | 0.0253 | (A) NE (B/D) |
| DSCAM | 2.29 | 0.77 | 1.38 | 1.25 | 1.52 | 0.0114 | 0.0392 | (A) NE (B/D) |
| MVB12A | 3.42 | 1.91 | 0.23 | 0.82 | 1.51 | 6.69E-12 | 2.97E-09 | (A) NE (B/D) |
| PAXIP1-AS1 | 7.57 | 6.06 | 0.31 | 1.04 | 1.51 | 6.49E-09 | 5.70E-07 | (A) NE (B/D) |
| WASH4P | 2.61 | 1.09 | 0.35 | 1.11 | 1.51 | 4.56E-08 | 2.57E-06 | (A) NE (B/D) |
| CCNF | 4.07 | 2.55 | 0.56 | 1.15 | 1.51 | 5.79E-06 | 1.18E-04 | (A) NE (B/D) |
| TUBGCP3 | 3.25 | 1.74 | 0.58 | 0.8 | 1.51 | 7.20E-06 | 1.41E-04 | (A) NE (B/D) |
| ATP5PF | 4.26 | 2.75 | 0.21 | 1.65 | 1.51 | 8.30E-06 | 1.57E-04 | (A) NE (B/D) |
| NAT9 | 3.77 | 2.26 | 0.4 | 1.55 | 1.51 | 8.73E-06 | 1.63E-04 | (A) NE (B/D) |
| KMT2B | 4.02 | 2.52 | 0.6 | 0.76 | 1.51 | 1.07E-05 | 1.92E-04 | (A) NE (B/D) |
| SARS2 | 2.16 | 0.65 | 0.6 | 0.76 | 1.51 | 1.08E-05 | 1.94E-04 | (A) NE (B/D) |
| SLC35D2 | 6.07 | 4.57 | 0.59 | 1.15 | 1.51 | 1.19E-05 | 2.09E-04 | (A) NE (B/D) |
| LINC00106 | 3.64 | 2.13 | 0.58 | 1.31 | 1.51 | 1.52E-05 | 2.51E-04 | (A) NE (B/D) |
| RTL8B | 5.59 | 4.07 | 0.47 | 1.74 | 1.51 | 5.31E-05 | 6.62E-04 | (A) NE (B/D) |
| CEP164 | 7.25 | 5.74 | 0.7 | 0.82 | 1.51 | 6.51E-05 | 7.76E-04 | (A) NE (B/D) |
| SH3GL2 | 1.68 | 0.18 | 0.85 | 0.48 | 1.51 | 5.50E-04 | 4.03E-03 | (A) NE (B/D) |
| OSGEPL1-AS | 4.68 | 3.17 | 0.92 | 0.77 | 1.51 | 8.23E-04 | 5.48E-03 | (A) NE (B/D) |
| RSPH1 | 3 | 1.49 | 0.85 | 1.66 | 1.51 | 8.91E-04 | 5.80E-03 | (A) NE (B/D) |
| ZNF284 | 2.82 | 1.31 | 1.06 | 0.88 | 1.51 | 2.40E-03 | 0.0123 | (A) NE (B/D) |
| XRCC2 | 2.61 | 1.1 | 1.18 | 1.12 | 1.51 | 4.64E-03 | 0.0203 | (A) NE (B/D) |
| NUPR2 | 2.34 | 0.83 | 1.31 | 1.25 | 1.51 | 9.25E-03 | 0.0337 | (A) NE (B/D) |
| ADAMTSL1 | 1.7 | 0.19 | 1.41 | 0.79 | 1.51 | 0.0126 | 0.0422 | (A) NE (B/D) |
| NUP188 | 6.6 | 5.1 | 0.31 | 0.92 | 1.5 | 1.25E-09 | 1.60E-07 | (A) NE (B/D) |
| DUSP8 | 1.99 | 0.49 | 0.39 | 0.75 | 1.5 | 1.17E-08 | 8.91E-07 | (A) NE (B/D) |
| ASPSCR1 | 2.61 | 1.11 | 0.32 | 1.06 | 1.5 | 1.26E-08 | 9.37E-07 | (A) NE (B/D) |
| CPT1B | 3.16 | 1.66 | 0.38 | 1.28 | 1.5 | 5.71E-07 | 1.91E-05 | (A) NE (B/D) |
| IRF2BP1 | 4.84 | 3.34 | 0.48 | 0.94 | 1.5 | 5.84E-07 | 1.94E-05 | (A) NE (B/D) |
| MAPK8IP3 | 1.88 | 0.37 | 0.47 | 0.59 | 1.5 | 6.62E-07 | 2.13E-05 | (A) NE (B/D) |

| | | | | | | | | |
|---|---|---|---|---|---|---|---|---|
| PAQR7 | 6.15 | 4.65 | 0.44 | 1.19 | 1.5 | 7.31E-07 | 2.32E-05 | (A) NE (B/D) |
| HIP1R | 7.01 | 5.51 | 0.5 | 1.02 | 1.5 | 1.15E-06 | 3.30E-05 | (A) NE (B/D) |
| FARSA | 2.73 | 1.24 | 0.51 | 1.18 | 1.5 | 2.76E-06 | 6.58E-05 | (A) NE (B/D) |
| SLC38A1 | 6.89 | 5.38 | 0.52 | 1.29 | 1.5 | 6.26E-06 | 1.26E-04 | (A) NE (B/D) |
| ERRFI1 | 6.11 | 4.61 | 0.49 | 1.53 | 1.5 | 1.93E-05 | 3.00E-04 | (A) NE (B/D) |
| RANGRF | 2.81 | 1.31 | 0.59 | 1.44 | 1.5 | 3.50E-05 | 4.78E-04 | (A) NE (B/D) |
| CCDC149 | 5.39 | 3.89 | 0.7 | 0.83 | 1.5 | 6.75E-05 | 8.00E-04 | (A) NE (B/D) |
| FOXJ3 | 9.23 | 7.73 | 0.71 | 0.7 | 1.5 | 9.47E-05 | 1.04E-03 | (A) NE (B/D) |
| ANKRD6 | 2.85 | 1.34 | 0.74 | 0.84 | 1.5 | 1.27E-04 | 1.30E-03 | (A) NE (B/D) |
| PLK3 | 5.92 | 4.42 | 0.85 | 0.8 | 1.5 | 4.32E-04 | 3.34E-03 | (A) NE (B/D) |
| RBM23 | 3.16 | 1.66 | 0.85 | 1.22 | 1.5 | 5.02E-04 | 3.75E-03 | (A) NE (B/D) |
| CCT8P1 | 7.94 | 6.44 | 0.87 | 1.12 | 1.5 | 5.47E-04 | 4.01E-03 | (A) NE (B/D) |
| LRRC20 | 6.5 | 5 | 0.85 | 1.49 | 1.5 | 6.62E-04 | 4.64E-03 | (A) NE (B/D) |
| LINC00111 | 2.12 | 0.62 | 1.15 | 0.84 | 1.5 | 4.15E-03 | 0.0187 | (A) NE (B/D) |
| ATG16L2 | 2.66 | 1.17 | 1.15 | 1.34 | 1.5 | 4.52E-03 | 0.02 | (A) NE (B/D) |
| MIR-1184-1/ | 10.15 | 8.65 | 0.99 | 2.17 | 1.5 | 5.14E-03 | 0.022 | (A) NE (B/D) |
| GJA4 | 9.35 | 7.84 | 1.09 | 2.43 | 1.5 | 0.0109 | 0.0379 | (A) NE (B/D) |
| RAVER1 | 1.76 | 0.27 | 0.24 | 0.44 | 1.49 | 1.70E-12 | 1.02E-09 | (A) NE (B/D) |
| PDCD2L | 2.77 | 1.28 | 0.33 | 0.9 | 1.49 | 2.41E-09 | 2.75E-07 | (A) NE (B/D) |
| TOMM40 | 5.13 | 3.64 | 0.29 | 1.1 | 1.49 | 1.36E-08 | 9.92E-07 | (A) NE (B/D) |
| SRRT | 8.76 | 7.27 | 0.17 | 1.17 | 1.49 | 1.79E-08 | 1.23E-06 | (A) NE (B/D) |
| GMPPA | 3.49 | 2 | 0.23 | 1.22 | 1.49 | 5.52E-08 | 3.00E-06 | (A) NE (B/D) |
| DDX49 | 6.01 | 4.53 | 0.41 | 1.1 | 1.49 | 2.37E-07 | 9.50E-06 | (A) NE (B/D) |
| WASH7P | 9.28 | 7.8 | 0.25 | 1.38 | 1.49 | 6.50E-07 | 2.11E-05 | (A) NE (B/D) |
| CAPN10 | 1.78 | 0.29 | 0.42 | 0.35 | 1.49 | 7.94E-07 | 2.48E-05 | (A) NE (B/D) |
| XRCC3 | 5.34 | 3.85 | 0.49 | 0.78 | 1.49 | 8.22E-07 | 2.56E-05 | (A) NE (B/D) |
| CNTROB | 6.62 | 5.13 | 0.51 | 0.98 | 1.49 | 1.69E-06 | 4.52E-05 | (A) NE (B/D) |
| ST14 | 3.53 | 2.05 | 0.54 | 1.05 | 1.49 | 3.56E-06 | 8.08E-05 | (A) NE (B/D) |
| MCM3AP | 5.86 | 4.37 | 0.57 | 0.69 | 1.49 | 7.39E-06 | 1.44E-04 | (A) NE (B/D) |
| UQCRBP1 | 6.2 | 4.71 | 0.55 | 1.39 | 1.49 | 2.04E-05 | 3.13E-04 | (A) NE (B/D) |
| HCG17 | 4.62 | 3.13 | 0.64 | 0.83 | 1.49 | 2.47E-05 | 3.61E-04 | (A) NE (B/D) |
| DUT | 2.27 | 0.78 | 0.66 | 0.87 | 1.49 | 3.84E-05 | 5.15E-04 | (A) NE (B/D) |
| DPYSL2 | 2.97 | 1.48 | 0.68 | 1 | 1.49 | 5.45E-05 | 6.77E-04 | (A) NE (B/D) |
| C14ORF93 | 6.41 | 4.92 | 0.7 | 1.06 | 1.49 | 7.41E-05 | 8.61E-04 | (A) NE (B/D) |
| FARP1-AS1 | 5.02 | 3.53 | 0.68 | 1.45 | 1.49 | 1.20E-04 | 1.24E-03 | (A) NE (B/D) |
| MIR-3650/36 | 7.05 | 5.56 | 0.6 | 1.7 | 1.49 | 1.52E-04 | 1.49E-03 | (A) NE (B/D) |
| MC1R | 5.08 | 3.59 | 0.86 | 1.15 | 1.49 | 5.66E-04 | 4.11E-03 | (A) NE (B/D) |
| MCM4 | 3.88 | 2.39 | 1.04 | 1.56 | 1.49 | 3.10E-03 | 0.015 | (A) NE (B/D) |
| NT5DC2 | 3.33 | 1.84 | 1.06 | 1.48 | 1.49 | 3.13E-03 | 0.0151 | (A) NE (B/D) |
| GGH | 1.72 | 0.23 | 1.09 | 0.53 | 1.49 | 3.24E-03 | 0.0155 | (A) NE (B/D) |
| TMEM38B | 6.83 | 5.33 | 1.11 | 1.02 | 1.49 | 3.46E-03 | 0.0163 | (A) NE (B/D) |
| KRT8P48 | 2.37 | 0.87 | 1.2 | 0.85 | 1.49 | 5.41E-03 | 0.0228 | (A) NE (B/D) |
| MAP7D2 | 3.04 | 1.55 | 1.44 | 1.36 | 1.49 | 0.0158 | 0.05 | (A) NE (B/D) |
| GTPBP2 | 1.73 | 0.25 | 0.33 | 0.52 | 1.48 | 1.75E-09 | 2.11E-07 | (A) NE (B/D) |
| STXBP2 | 2.63 | 1.15 | 0.3 | 1.1 | 1.48 | 2.11E-08 | 1.38E-06 | (A) NE (B/D) |
| ST20-MTHFS | 9.25 | 7.76 | 0.39 | 1.12 | 1.48 | 1.73E-07 | 7.50E-06 | (A) NE (B/D) |
| CDK9 | 7.1 | 5.62 | 0.36 | 1.2 | 1.48 | 2.29E-07 | 9.31E-06 | (A) NE (B/D) |
| MED16 | 2.03 | 0.55 | 0.46 | 0.71 | 1.48 | 3.12E-07 | 1.18E-05 | (A) NE (B/D) |
| ZNF76 | 1.74 | 0.25 | 0.5 | 0.43 | 1.48 | 4.13E-06 | 9.12E-05 | (A) NE (B/D) |
| SNORA51 | 2.67 | 1.19 | 0.56 | 0.96 | 1.48 | 5.43E-06 | 1.12E-04 | (A) NE (B/D) |
| RBM10 | 3.46 | 1.98 | 0.46 | 1.43 | 1.48 | 9.14E-06 | 1.69E-04 | (A) NE (B/D) |
| VN1R108P | 3.77 | 2.29 | 0.61 | 0.94 | 1.48 | 1.51E-05 | 2.51E-04 | (A) NE (B/D) |
| PODXL2 | 2.24 | 0.76 | 0.74 | 0.98 | 1.48 | 1.41E-04 | 1.41E-03 | (A) NE (B/D) |
| MCRIP2 | 4.44 | 2.96 | 0.74 | 1.23 | 1.48 | 1.73E-04 | 1.64E-03 | (A) NE (B/D) |
| SH3GLB1 | 8.38 | 6.9 | 0.77 | 0.72 | 1.48 | 2.41E-04 | 2.12E-03 | (A) NE (B/D) |
| SGO1-AS1 | 4.84 | 3.36 | 0.75 | 1.51 | 1.48 | 3.30E-04 | 2.71E-03 | (A) NE (B/D) |
| PMS2P1 | 6.8 | 5.32 | 0.86 | 0.79 | 1.48 | 5.43E-04 | 3.99E-03 | (A) NE (B/D) |
| IFNLR1 | 3.25 | 1.77 | 0.91 | 1.31 | 1.48 | 1.03E-03 | 6.45E-03 | (A) NE (B/D) |
| PCDHB12 | 1.5 | 0.03 | 0.92 | 0.13 | 1.48 | 1.27E-03 | 7.59E-03 | (A) NE (B/D) |
| KLHL32 | 1.69 | 0.22 | 0.93 | 0.34 | 1.48 | 1.29E-03 | 7.65E-03 | (A) NE (B/D) |
| LINC00535 | 1.57 | 0.09 | 1.01 | 0.23 | 1.48 | 2.26E-03 | 0.0117 | (A) NE (B/D) |
| CADM2 | 1.49 | 0.01 | 1.02 | 0.08 | 1.48 | 2.37E-03 | 0.0122 | (A) NE (B/D) |
| KRT8P11 | 1.8 | 0.32 | 1.09 | 0.56 | 1.48 | 3.25E-03 | 0.0155 | (A) NE (B/D) |
| TRAF3IP2-AS | 2.24 | 0.76 | 1.09 | 0.6 | 1.48 | 3.41E-03 | 0.0161 | (A) NE (B/D) |
| UAP1L1 | 8.08 | 6.61 | 1.07 | 1.46 | 1.48 | 3.50E-03 | 0.0164 | (A) NE (B/D) |

| | | | | | | | | |
|---|---|---|---|---|---|---|---|---|
| NCAPG | 1.88 | 0.4 | 1.15 | 0.85 | 1.48 | 4.42E-03 | 0.0196 | (A) NE (B/D) |
| CTBP2P7 | 1.8 | 0.32 | 1.21 | 0.49 | 1.48 | 6.32E-03 | 0.0256 | (A) NE (B/D) |
| NLRP1 | 1.84 | 0.35 | 1.22 | 0.54 | 1.48 | 6.47E-03 | 0.026 | (A) NE (B/D) |
| RXRB | 9.19 | 7.72 | 0.14 | 1.15 | 1.47 | 1.36E-08 | 9.92E-07 | (A) NE (B/D) |
| ARRDC1-AS1 | 5.02 | 3.56 | 0.4 | 0.9 | 1.47 | 4.65E-08 | 2.61E-06 | (A) NE (B/D) |
| YRDC | 3.73 | 2.26 | 0.25 | 1.21 | 1.47 | 7.86E-08 | 3.97E-06 | (A) NE (B/D) |
| TSPAN17 | 2.52 | 1.05 | 0.41 | 0.96 | 1.47 | 8.88E-08 | 4.35E-06 | (A) NE (B/D) |
| CDK5RAP2 | 6.93 | 5.47 | 0.42 | 0.6 | 1.47 | 1.17E-07 | 5.47E-06 | (A) NE (B/D) |
| ENTPD6 | 3.74 | 2.26 | 0.42 | 1.21 | 1.47 | 9.16E-07 | 2.77E-05 | (A) NE (B/D) |
| RPS15 | 6.92 | 5.45 | 0.49 | 1.34 | 1.47 | 7.85E-06 | 1.51E-04 | (A) NE (B/D) |
| TSTD2 | 2.87 | 1.4 | 0.58 | 0.91 | 1.47 | 8.67E-06 | 1.63E-04 | (A) NE (B/D) |
| SLC25A35 | 5.92 | 4.45 | 0.35 | 1.63 | 1.47 | 1.80E-05 | 2.85E-04 | (A) NE (B/D) |
| ZDHHC12 | 6.86 | 5.39 | 0.53 | 1.43 | 1.47 | 2.44E-05 | 3.57E-04 | (A) NE (B/D) |
| HNRNPA1L2 | 2.99 | 1.52 | 0.54 | 1.43 | 1.47 | 2.57E-05 | 3.73E-04 | (A) NE (B/D) |
| MEST | 2.02 | 0.54 | 0.6 | 0.45 | 1.47 | 3.33E-05 | 4.59E-04 | (A) NE (B/D) |
| RFC4 | 2.01 | 0.54 | 0.75 | 0.85 | 1.47 | 1.60E-04 | 1.55E-03 | (A) NE (B/D) |
| UBE2SP2 | 1.56 | 0.09 | 0.78 | 0.36 | 1.47 | 3.93E-04 | 3.11E-03 | (A) NE (B/D) |
| DBN1 | 2.05 | 0.58 | 0.9 | 0.87 | 1.47 | 8.60E-04 | 5.64E-03 | (A) NE (B/D) |
| FSD1L | 1.5 | 0.03 | 0.87 | 0.11 | 1.47 | 9.35E-04 | 6.00E-03 | (A) NE (B/D) |
| CACNA2D2 | 1.63 | 0.16 | 0.94 | 0.36 | 1.47 | 1.40E-03 | 8.17E-03 | (A) NE (B/D) |
| STRCP1 | 2.71 | 1.24 | 0.97 | 0.86 | 1.47 | 1.44E-03 | 8.35E-03 | (A) NE (B/D) |
| ENPP7P2 | 2.9 | 1.43 | 1.04 | 1 | 1.47 | 2.39E-03 | 0.0123 | (A) NE (B/D) |
| PDLIM1 | 7.35 | 5.88 | 1.06 | 1.22 | 1.47 | 3.01E-03 | 0.0146 | (A) NE (B/D) |
| DOCK9 | 3.93 | 2.46 | 1.12 | 0.87 | 1.47 | 4.06E-03 | 0.0184 | (A) NE (B/D) |
| RAB11FIP1 | 2.54 | 1.07 | 1.13 | 0.96 | 1.47 | 4.21E-03 | 0.0189 | (A) NE (B/D) |
| ZNF846 | 4.82 | 3.34 | 1.14 | 0.86 | 1.47 | 4.46E-03 | 0.0197 | (A) NE (B/D) |
| ZNF461 | 4.59 | 3.12 | 1.28 | 0.79 | 1.47 | 8.33E-03 | 0.0312 | (A) NE (B/D) |
| RNU6-611P | 2.86 | 1.39 | 1.29 | 1 | 1.47 | 9.24E-03 | 0.0337 | (A) NE (B/D) |
| PCDHGA8 | 2.46 | 0.99 | 1.35 | 1.02 | 1.47 | 0.0117 | 0.0399 | (A) NE (B/D) |
| DPP9 | 1.85 | 0.39 | 0.19 | 0.49 | 1.46 | 5.02E-16 | 1.44E-12 | (A) NE (B/D) |
| ATAD3B | 7.02 | 5.56 | 0.31 | 0.97 | 1.46 | 4.92E-09 | 4.61E-07 | (A) NE (B/D) |
| ABHD17A | 2.49 | 1.03 | 0.29 | 1.12 | 1.46 | 3.49E-08 | 2.09E-06 | (A) NE (B/D) |
| ROM1 | 4.01 | 2.54 | 0.47 | 0.69 | 1.46 | 5.58E-07 | 1.89E-05 | (A) NE (B/D) |
| P4HTM | 7.37 | 5.91 | 0.47 | 1.09 | 1.46 | 1.40E-06 | 3.89E-05 | (A) NE (B/D) |
| GADD45GIP1 | 5.67 | 4.22 | 0.45 | 1.33 | 1.46 | 4.43E-06 | 9.61E-05 | (A) NE (B/D) |
| TRAF3IP2 | 6.69 | 5.23 | 0.59 | 0.86 | 1.46 | 1.26E-05 | 2.19E-04 | (A) NE (B/D) |
| USP2 | 3.56 | 2.1 | 0.58 | 1.11 | 1.46 | 1.32E-05 | 2.28E-04 | (A) NE (B/D) |
| ARHGAP39 | 5.78 | 4.33 | 0.63 | 1.45 | 1.46 | 9.30E-05 | 1.03E-03 | (A) NE (B/D) |
| ADM5 | 5.21 | 3.75 | 0.73 | 1.27 | 1.46 | 1.86E-04 | 1.74E-03 | (A) NE (B/D) |
| DLG4 | 1.93 | 0.47 | 0.75 | 0.62 | 1.46 | 2.15E-04 | 1.94E-03 | (A) NE (B/D) |
| RAB11B-AS1 | 5.2 | 3.74 | 0.89 | 0.9 | 1.46 | 8.44E-04 | 5.58E-03 | (A) NE (B/D) |
| HMGN2P41 | 2.64 | 1.18 | 0.91 | 1.07 | 1.46 | 9.90E-04 | 6.27E-03 | (A) NE (B/D) |
| MYBPHL | 1.48 | 0.01 | 1.04 | 0.05 | 1.46 | 2.89E-03 | 0.0142 | (A) NE (B/D) |
| GLRA4 | 1.82 | 0.36 | 1.37 | 0.52 | 1.46 | 0.0125 | 0.0421 | (A) NE (B/D) |
| MAN2B1 | 3.85 | 2.4 | 0.22 | 0.97 | 1.45 | 7.01E-10 | 1.09E-07 | (A) NE (B/D) |
| ILKAP | 4.74 | 3.29 | 0.29 | 0.89 | 1.45 | 8.29E-10 | 1.22E-07 | (A) NE (B/D) |
| SPIRE1 | 7.24 | 5.79 | 0.39 | 1.01 | 1.45 | 1.06E-07 | 5.02E-06 | (A) NE (B/D) |
| SLC26A10 | 2.32 | 0.87 | 0.39 | 1.15 | 1.45 | 3.76E-07 | 1.37E-05 | (A) NE (B/D) |
| COA8 | 4.7 | 3.26 | 0.45 | 0.95 | 1.45 | 4.46E-07 | 1.58E-05 | (A) NE (B/D) |
| CPNE1 | 5.45 | 4 | 0.55 | 1.01 | 1.45 | 6.53E-06 | 1.31E-04 | (A) NE (B/D) |
| PDCL3P4 | 6.13 | 4.68 | 0.4 | 1.43 | 1.45 | 6.65E-06 | 1.32E-04 | (A) NE (B/D) |
| SSBP3 | 3.63 | 2.18 | 0.52 | 1.25 | 1.45 | 9.01E-06 | 1.67E-04 | (A) NE (B/D) |
| PHTF1 | 4.88 | 3.43 | 0.59 | 0.66 | 1.45 | 2.00E-05 | 3.08E-04 | (A) NE (B/D) |
| TIMM13 | 3.5 | 2.05 | 0.55 | 1.37 | 1.45 | 2.68E-05 | 3.86E-04 | (A) NE (B/D) |
| SEC14L1 | 5.49 | 4.05 | 0.64 | 0.84 | 1.45 | 3.51E-05 | 4.79E-04 | (A) NE (B/D) |
| C19ORF47 | 5.3 | 3.85 | 0.65 | 0.86 | 1.45 | 4.43E-05 | 5.74E-04 | (A) NE (B/D) |
| CCNJL | 1.57 | 0.13 | 0.62 | 0.48 | 1.45 | 5.25E-05 | 6.57E-04 | (A) NE (B/D) |
| HSPA1B | 12.45 | 10.99 | 0.66 | 1.13 | 1.45 | 5.50E-05 | 6.81E-04 | (A) NE (B/D) |
| SPCS2P4 | 5.2 | 3.75 | 0.29 | 1.79 | 1.45 | 6.23E-05 | 7.50E-04 | (A) NE (B/D) |
| COQ4 | 1.93 | 0.47 | 0.68 | 0.71 | 1.45 | 7.98E-05 | 9.09E-04 | (A) NE (B/D) |
| TCEAL1 | 4.75 | 3.3 | 0.73 | 1.46 | 1.45 | 3.17E-04 | 2.63E-03 | (A) NE (B/D) |
| EBF4 | 1.65 | 0.2 | 0.9 | 0.4 | 1.45 | 1.14E-03 | 6.96E-03 | (A) NE (B/D) |
| LMO7 | 2.12 | 0.67 | 0.95 | 0.79 | 1.45 | 1.46E-03 | 8.42E-03 | (A) NE (B/D) |
| TMEM59 | 3.59 | 2.14 | 0.95 | 1.46 | 1.45 | 1.93E-03 | 0.0104 | (A) NE (B/D) |
| CEP295 | 3.58 | 2.13 | 1.1 | 0.77 | 1.45 | 3.93E-03 | 0.0179 | (A) NE (B/D) |

| | | | | | | | | |
|---|---|---|---|---|---|---|---|---|
| FGFR1 | 4.28 | 2.82 | 1.17 | 1.33 | 1.45 | 6.13E-03 | 0.025 | (A) NE (B/D) |
| ASXL1 | 2.44 | 1 | 0.31 | 0.8 | 1.44 | 6.62E-10 | 1.05E-07 | (A) NE (B/D) |
| RNF220 | 2.71 | 1.27 | 0.34 | 0.77 | 1.44 | 1.98E-09 | 2.34E-07 | (A) NE (B/D) |
| EIF4A2P4 | 3.61 | 2.17 | 0.42 | 1.09 | 1.44 | 4.94E-07 | 1.71E-05 | (A) NE (B/D) |
| MED9 | 3.92 | 2.49 | 0.47 | 1.02 | 1.44 | 1.01E-06 | 3.01E-05 | (A) NE (B/D) |
| POLD3 | 1.96 | 0.52 | 0.55 | 0.75 | 1.44 | 6.30E-06 | 1.27E-04 | (A) NE (B/D) |
| HIRIP3 | 6.78 | 5.35 | 0.46 | 1.42 | 1.44 | 1.32E-05 | 2.28E-04 | (A) NE (B/D) |
| PSMG2 | 7.43 | 5.99 | 0.56 | 1.21 | 1.44 | 1.56E-05 | 2.56E-04 | (A) NE (B/D) |
| DHRS4 | 2.65 | 1.21 | 0.58 | 1.21 | 1.44 | 2.09E-05 | 3.18E-04 | (A) NE (B/D) |
| ACTN4P1 | 3.67 | 2.24 | 0.6 | 1.11 | 1.44 | 2.20E-05 | 3.29E-04 | (A) NE (B/D) |
| FBXW9 | 5.51 | 4.07 | 0.65 | 1.16 | 1.44 | 5.48E-05 | 6.79E-04 | (A) NE (B/D) |
| RFNG | 4.76 | 3.31 | 0.65 | 1.14 | 1.44 | 5.88E-05 | 7.18E-04 | (A) NE (B/D) |
| FKBP8 | 4.24 | 2.8 | 0.45 | 1.7 | 1.44 | 7.60E-05 | 8.77E-04 | (A) NE (B/D) |
| TRIT1 | 2.77 | 1.33 | 0.69 | 0.92 | 1.44 | 9.39E-05 | 1.04E-03 | (A) NE (B/D) |
| MFSD10 | 3.03 | 1.6 | 0.78 | 0.95 | 1.44 | 2.91E-04 | 2.46E-03 | (A) NE (B/D) |
| SFPQ | 6.3 | 4.87 | 0.78 | 1.04 | 1.44 | 3.14E-04 | 2.62E-03 | (A) NE (B/D) |
| C4ORF46 | 6.36 | 4.91 | 0.81 | 1.1 | 1.44 | 4.27E-04 | 3.31E-03 | (A) NE (B/D) |
| SNX18P7 | 3.95 | 2.5 | 0.62 | 1.9 | 1.44 | 5.90E-04 | 4.25E-03 | (A) NE (B/D) |
| ECSIT | 1.67 | 0.23 | 0.9 | 0.49 | 1.44 | 1.13E-03 | 6.92E-03 | (A) NE (B/D) |
| KIF5A | 1.45 | 0.01 | 1.02 | 0.04 | 1.44 | 2.81E-03 | 0.0139 | (A) NE (B/D) |
| HSFX2 | 2.9 | 1.46 | 1.04 | 0.98 | 1.44 | 2.84E-03 | 0.014 | (A) NE (B/D) |
| DPM3 | 9.57 | 8.14 | 1.01 | 1.5 | 1.44 | 3.20E-03 | 0.0154 | (A) NE (B/D) |
| GCOM2 | 6.04 | 4.6 | 1.07 | 0.81 | 1.44 | 3.34E-03 | 0.0159 | (A) NE (B/D) |
| B4GALNT4 | 4.03 | 2.59 | 1.11 | 1.46 | 1.44 | 5.29E-03 | 0.0224 | (A) NE (B/D) |
| MED15P9 | 2.67 | 1.23 | 1.2 | 1.15 | 1.44 | 7.02E-03 | 0.0276 | (A) NE (B/D) |
| LINC00593 | 1.63 | 0.19 | 1.24 | 0.51 | 1.44 | 8.09E-03 | 0.0305 | (A) NE (B/D) |
| LTBP4 | 2.33 | 0.9 | 1.24 | 1.36 | 1.44 | 9.44E-03 | 0.0342 | (A) NE (B/D) |
| HMMR | 1.76 | 0.32 | 1.31 | 0.82 | 1.44 | 0.0109 | 0.038 | (A) NE (B/D) |
| MCOLN1 | 2.68 | 1.25 | 0.2 | 0.85 | 1.43 | 3.66E-11 | 1.00E-08 | (A) NE (B/D) |
| DNM2 | 3.51 | 2.08 | 0.31 | 0.71 | 1.43 | 3.40E-10 | 6.15E-08 | (A) NE (B/D) |
| ALX3 | 4.12 | 2.69 | 0.26 | 0.99 | 1.43 | 2.74E-09 | 3.00E-07 | (A) NE (B/D) |
| KEAP1 | 1.98 | 0.55 | 0.35 | 0.62 | 1.43 | 5.73E-09 | 5.19E-07 | (A) NE (B/D) |
| PCIF1 | 8.08 | 6.65 | 0.24 | 1.07 | 1.43 | 1.21E-08 | 9.09E-07 | (A) NE (B/D) |
| MAP2K2 | 4.02 | 2.58 | 0.38 | 0.87 | 1.43 | 2.60E-08 | 1.62E-06 | (A) NE (B/D) |
| COPS6 | 3.3 | 1.88 | 0.15 | 1.29 | 1.43 | 2.82E-07 | 1.09E-05 | (A) NE (B/D) |
| RMND5B | 2.8 | 1.37 | 0.37 | 1.2 | 1.43 | 5.78E-07 | 1.93E-05 | (A) NE (B/D) |
| MSL3P1 | 6.93 | 5.51 | 0.34 | 1.25 | 1.43 | 6.58E-07 | 2.13E-05 | (A) NE (B/D) |
| AMPD3 | 2.52 | 1.09 | 0.42 | 1.13 | 1.43 | 8.37E-07 | 2.58E-05 | (A) NE (B/D) |
| CLSTN3 | 6.38 | 4.96 | 0.43 | 1.2 | 1.43 | 1.61E-06 | 4.35E-05 | (A) NE (B/D) |
| SIX1 | 2.93 | 1.51 | 0.48 | 1.09 | 1.43 | 2.29E-06 | 5.70E-05 | (A) NE (B/D) |
| ZNF511 | 2.01 | 0.58 | 0.51 | 0.73 | 1.43 | 2.88E-06 | 6.79E-05 | (A) NE (B/D) |
| GPS1 | 5.07 | 3.65 | 0.39 | 1.37 | 1.43 | 3.72E-06 | 8.38E-05 | (A) NE (B/D) |
| POGLUT2 | 3.96 | 2.53 | 0.53 | 0.93 | 1.43 | 4.57E-06 | 9.83E-05 | (A) NE (B/D) |
| ZNF322 | 7.43 | 6 | 0.54 | 0.81 | 1.43 | 4.75E-06 | 1.02E-04 | (A) NE (B/D) |
| THAP3 | 4.46 | 3.03 | 0.53 | 1.09 | 1.43 | 6.69E-06 | 1.33E-04 | (A) NE (B/D) |
| ZNF713 | 4.36 | 2.93 | 0.55 | 0.69 | 1.43 | 7.30E-06 | 1.42E-04 | (A) NE (B/D) |
| SMYD3 | 1.48 | 0.05 | 0.51 | 0.31 | 1.43 | 1.50E-05 | 2.50E-04 | (A) NE (B/D) |
| SNORD35B | 4.46 | 3.03 | 0.61 | 1.13 | 1.43 | 3.15E-05 | 4.41E-04 | (A) NE (B/D) |
| MIR-3605/3P | 6.92 | 5.48 | 0.66 | 0.51 | 1.43 | 9.79E-05 | 1.07E-03 | (A) NE (B/D) |
| MIR-3605/5P | 6.92 | 5.48 | 0.66 | 0.51 | 1.43 | 9.79E-05 | 1.07E-03 | (A) NE (B/D) |
| MAD2L1BP | 7.88 | 6.46 | 0.64 | 1.38 | 1.43 | 1.05E-04 | 1.13E-03 | (A) NE (B/D) |
| SOGA1 | 2.8 | 1.37 | 0.71 | 0.9 | 1.43 | 1.20E-04 | 1.24E-03 | (A) NE (B/D) |
| SCAND1 | 2.49 | 1.06 | 0.78 | 0.89 | 1.43 | 3.19E-04 | 2.65E-03 | (A) NE (B/D) |
| C12ORF57 | 2.95 | 1.52 | 0.78 | 1.4 | 1.43 | 5.07E-04 | 3.78E-03 | (A) NE (B/D) |
| SNORD124 | 4.9 | 3.47 | 0.85 | 0.79 | 1.43 | 6.58E-04 | 4.62E-03 | (A) NE (B/D) |
| MAFG-DT | 3.55 | 2.12 | 0.87 | 1.2 | 1.43 | 9.17E-04 | 5.92E-03 | (A) NE (B/D) |
| PCDHGA10 | 3.23 | 1.79 | 1.15 | 1.06 | 1.43 | 5.46E-03 | 0.0229 | (A) NE (B/D) |
| IGHVIII-38-1 | 1.48 | 0.05 | 1.22 | 0.24 | 1.43 | 7.95E-03 | 0.0301 | (A) NE (B/D) |
| NECAB1 | 2.06 | 0.63 | 1.24 | 0.93 | 1.43 | 8.46E-03 | 0.0316 | (A) NE (B/D) |
| OR5BH1P | 1.74 | 0.31 | 1.29 | 0.63 | 1.43 | 0.0104 | 0.0368 | (A) NE (B/D) |
| ANKRD39 | 5.85 | 4.43 | 0.25 | 0.65 | 1.42 | 5.49E-12 | 2.52E-09 | (A) NE (B/D) |
| NSUN5 | 7.94 | 6.53 | 0.32 | 1.1 | 1.42 | 7.65E-08 | 3.90E-06 | (A) NE (B/D) |
| WASH3P | 4.91 | 3.49 | 0.33 | 1.25 | 1.42 | 6.44E-07 | 2.10E-05 | (A) NE (B/D) |
| PKP4 | 6.65 | 5.23 | 0.51 | 0.74 | 1.42 | 2.80E-06 | 6.65E-05 | (A) NE (B/D) |
| NPIPA3 | 4.27 | 2.85 | 0.46 | 1.18 | 1.42 | 2.83E-06 | 6.71E-05 | (A) NE (B/D) |

| | | | | | | | | |
|---|---|---|---|---|---|---|---|---|
| MIR-4640/3P | 2.47 | 1.06 | 0.53 | 0.9 | 1.42 | 4.40E-06 | 9.56E-05 | (A) NE (B/D) |
| MIR-4640/5P | 2.47 | 1.06 | 0.53 | 0.9 | 1.42 | 4.40E-06 | 9.56E-05 | (A) NE (B/D) |
| SPRY1 | 1.84 | 0.42 | 0.52 | 0.63 | 1.42 | 4.64E-06 | 9.95E-05 | (A) NE (B/D) |
| ELP5 | 7.95 | 6.53 | 0.55 | 1.18 | 1.42 | 1.33E-05 | 2.28E-04 | (A) NE (B/D) |
| ACTN4 | 10.25 | 8.82 | 0.58 | 0.99 | 1.42 | 1.63E-05 | 2.65E-04 | (A) NE (B/D) |
| HES1 | 4.35 | 2.93 | 0.44 | 1.57 | 1.42 | 3.32E-05 | 4.58E-04 | (A) NE (B/D) |
| HSPA1A | 10.98 | 9.56 | 0.66 | 1.08 | 1.42 | 7.48E-05 | 8.68E-04 | (A) NE (B/D) |
| HEXD | 2.04 | 0.63 | 0.69 | 0.81 | 1.42 | 9.94E-05 | 1.08E-03 | (A) NE (B/D) |
| ZACN | 9.47 | 8.05 | 0.68 | 1.26 | 1.42 | 1.39E-04 | 1.40E-03 | (A) NE (B/D) |
| SYT17 | 5.26 | 3.84 | 0.65 | 1.55 | 1.42 | 2.19E-04 | 1.97E-03 | (A) NE (B/D) |
| SNHG20 | 4.66 | 3.24 | 0.77 | 0.93 | 1.42 | 2.88E-04 | 2.44E-03 | (A) NE (B/D) |
| RN7SL736P | 2.3 | 0.89 | 0.79 | 0.84 | 1.42 | 3.77E-04 | 3.01E-03 | (A) NE (B/D) |
| PNMA1 | 6.3 | 4.88 | 0.75 | 1.46 | 1.42 | 4.77E-04 | 3.60E-03 | (A) NE (B/D) |
| COX8A | 9.31 | 7.89 | 0.8 | 1.32 | 1.42 | 5.64E-04 | 4.11E-03 | (A) NE (B/D) |
| SETP20 | 1.6 | 0.18 | 0.89 | 0.34 | 1.42 | 1.28E-03 | 7.62E-03 | (A) NE (B/D) |
| RN7SKP150 | 3.35 | 1.93 | 0.95 | 0.87 | 1.42 | 1.66E-03 | 9.32E-03 | (A) NE (B/D) |
| MBOAT1 | 2.16 | 0.73 | 0.99 | 0.87 | 1.42 | 2.19E-03 | 0.0114 | (A) NE (B/D) |
| FAM83C-AS1 | 2.71 | 1.29 | 1.01 | 1.13 | 1.42 | 2.62E-03 | 0.0132 | (A) NE (B/D) |
| STPG3 | 3.57 | 2.15 | 1 | 1.22 | 1.42 | 2.64E-03 | 0.0133 | (A) NE (B/D) |
| PCP2 | 1.88 | 0.46 | 1.05 | 0.61 | 1.42 | 3.33E-03 | 0.0158 | (A) NE (B/D) |
| YBX2 | 1.46 | 0.04 | 1.06 | 0.26 | 1.42 | 3.85E-03 | 0.0176 | (A) NE (B/D) |
| HSF2 | 2.25 | 0.83 | 1.09 | 0.87 | 1.42 | 4.16E-03 | 0.0187 | (A) NE (B/D) |
| HEYL | 3.59 | 2.16 | 1.05 | 1.49 | 1.42 | 4.34E-03 | 0.0193 | (A) NE (B/D) |
| PTMAP1 | 4.28 | 2.86 | 1.07 | 1.41 | 1.42 | 4.39E-03 | 0.0195 | (A) NE (B/D) |
| TMA16P1 | 2.3 | 0.88 | 1.13 | 1.18 | 1.42 | 5.69E-03 | 0.0237 | (A) NE (B/D) |
| SLC25A1P5 | 6.62 | 5.2 | 1.24 | 0.72 | 1.42 | 8.56E-03 | 0.0318 | (A) NE (B/D) |
| MIR-1225/3P | 4.97 | 3.55 | 1.15 | 1.81 | 1.42 | 9.34E-03 | 0.0339 | (A) NE (B/D) |
| MIR-1225/5P | 4.97 | 3.55 | 1.15 | 1.81 | 1.42 | 9.34E-03 | 0.0339 | (A) NE (B/D) |
| KRT18 | 10.1 | 8.67 | 1.19 | 1.9 | 1.42 | 0.012 | 0.0406 | (A) NE (B/D) |
| GRAMD1A | 5.29 | 3.88 | 0.3 | 0.83 | 1.41 | 1.06E-09 | 1.42E-07 | (A) NE (B/D) |
| PICK1 | 2.7 | 1.29 | 0.23 | 1.08 | 1.41 | 1.55E-08 | 1.09E-06 | (A) NE (B/D) |
| MIR-1229/12 | 6.03 | 4.62 | 0.34 | 1.03 | 1.41 | 5.31E-08 | 2.91E-06 | (A) NE (B/D) |
| MIR-1229/3P | 6.03 | 4.62 | 0.34 | 1.03 | 1.41 | 5.31E-08 | 2.91E-06 | (A) NE (B/D) |
| MIR-1229/5P | 6.03 | 4.62 | 0.34 | 1.03 | 1.41 | 5.31E-08 | 2.91E-06 | (A) NE (B/D) |
| PPP2R5D | 3.08 | 1.67 | 0.33 | 1.08 | 1.41 | 7.89E-08 | 3.97E-06 | (A) NE (B/D) |
| CSRP1 | 2.76 | 1.35 | 0.39 | 1.07 | 1.41 | 2.85E-07 | 1.10E-05 | (A) NE (B/D) |
| JUND | 7.09 | 5.68 | 0.44 | 0.59 | 1.41 | 5.05E-07 | 1.73E-05 | (A) NE (B/D) |
| EIF3K | 2.84 | 1.43 | 0.22 | 1.32 | 1.41 | 6.63E-07 | 2.13E-05 | (A) NE (B/D) |
| CFAP157 | 4.91 | 3.5 | 0.45 | 1.09 | 1.41 | 1.42E-06 | 3.94E-05 | (A) NE (B/D) |
| SRSF7 | 7.34 | 5.93 | 0.49 | 1.3 | 1.41 | 1.24E-05 | 2.16E-04 | (A) NE (B/D) |
| FGD1 | 6.23 | 4.82 | 0.35 | 1.64 | 1.41 | 3.94E-05 | 5.24E-04 | (A) NE (B/D) |
| KLHL23 | 4.25 | 2.85 | 0.64 | 1.32 | 1.41 | 1.02E-04 | 1.10E-03 | (A) NE (B/D) |
| OVCA2 | 3.07 | 1.67 | 0.65 | 1.3 | 1.41 | 1.14E-04 | 1.20E-03 | (A) NE (B/D) |
| KLHDC9 | 3.53 | 2.12 | 0.5 | 1.66 | 1.41 | 1.22E-04 | 1.26E-03 | (A) NE (B/D) |
| IQCD | 4.77 | 3.35 | 0.76 | 1.14 | 1.41 | 2.96E-04 | 2.50E-03 | (A) NE (B/D) |
| GOLM1 | 4.85 | 3.44 | 0.56 | 1.81 | 1.41 | 3.73E-04 | 2.99E-03 | (A) NE (B/D) |
| NRL | 4.44 | 3.03 | 0.88 | 0.78 | 1.41 | 9.66E-04 | 6.16E-03 | (A) NE (B/D) |
| TVP23C-CDRT | 5.34 | 3.94 | 0.89 | 1.18 | 1.41 | 1.24E-03 | 7.45E-03 | (A) NE (B/D) |
| SNHG12 | 2.96 | 1.55 | 0.94 | 1.03 | 1.41 | 1.60E-03 | 9.05E-03 | (A) NE (B/D) |
| ZNF227 | 6.59 | 5.18 | 0.99 | 0.79 | 1.41 | 2.44E-03 | 0.0125 | (A) NE (B/D) |
| PSMG3 | 3.34 | 1.93 | 1.02 | 1.41 | 1.41 | 3.61E-03 | 0.0168 | (A) NE (B/D) |
| AMY1B | 2.02 | 0.61 | 1.2 | 0.93 | 1.41 | 7.74E-03 | 0.0295 | (A) NE (B/D) |
| HDAC6 | 1.76 | 0.35 | 1.29 | 0.55 | 1.41 | 0.0112 | 0.0386 | (A) NE (B/D) |
| HCG25 | 2.23 | 0.82 | 0.27 | 0.81 | 1.4 | 2.74E-10 | 5.26E-08 | (A) NE (B/D) |
| PRKD2 | 7.7 | 6.3 | 0.32 | 1.02 | 1.4 | 3.77E-08 | 2.22E-06 | (A) NE (B/D) |
| ARFGAP1 | 3.57 | 2.17 | 0.34 | 1.02 | 1.4 | 5.94E-08 | 3.16E-06 | (A) NE (B/D) |
| MAPK12 | 1.81 | 0.41 | 0.4 | 0.58 | 1.4 | 9.59E-08 | 4.64E-06 | (A) NE (B/D) |
| CFAP298 | 2.43 | 1.03 | 0.36 | 1.1 | 1.4 | 2.44E-07 | 9.72E-06 | (A) NE (B/D) |
| ZSCAN21 | 1.88 | 0.48 | 0.46 | 0.64 | 1.4 | 8.38E-07 | 2.58E-05 | (A) NE (B/D) |
| B9D2 | 1.54 | 0.14 | 0.4 | 0.25 | 1.4 | 2.09E-06 | 5.30E-05 | (A) NE (B/D) |
| TMCO3 | 5.72 | 4.32 | 0.58 | 0.83 | 1.4 | 1.61E-05 | 2.63E-04 | (A) NE (B/D) |
| ZBTB22 | 6.99 | 5.6 | 0.55 | 1.16 | 1.4 | 1.73E-05 | 2.77E-04 | (A) NE (B/D) |
| RNASEK | 5.26 | 3.86 | 0.15 | 1.65 | 1.4 | 2.32E-05 | 3.43E-04 | (A) NE (B/D) |
| LMCD1 | 2.41 | 1.01 | 0.7 | 1.02 | 1.4 | 1.36E-04 | 1.37E-03 | (A) NE (B/D) |
| KIAA1211L | 1.44 | 0.04 | 0.63 | 0.12 | 1.4 | 1.55E-04 | 1.52E-03 | (A) NE (B/D) |

| | | | | | | | | |
|---|---|---|---|---|---|---|---|---|
| MTX2 | 2.48 | 1.08 | 0.71 | 1.26 | 1.4 | 2.37E-04 | 2.10E-03 | (A) NE (B/D) |
| CNTRL | 2.72 | 1.32 | 0.75 | 0.76 | 1.4 | 2.66E-04 | 2.29E-03 | (A) NE (B/D) |
| ACIN1 | 4.02 | 2.62 | 0.79 | 0.97 | 1.4 | 4.26E-04 | 3.30E-03 | (A) NE (B/D) |
| EIF4HP2 | 5.53 | 4.14 | 0.8 | 1.03 | 1.4 | 5.04E-04 | 3.76E-03 | (A) NE (B/D) |
| AMZ2P1 | 4.29 | 2.89 | 0.82 | 1.39 | 1.4 | 9.02E-04 | 5.85E-03 | (A) NE (B/D) |
| MRAP2 | 1.49 | 0.09 | 0.86 | 0.33 | 1.4 | 1.10E-03 | 6.77E-03 | (A) NE (B/D) |
| PFN2 | 5.59 | 4.19 | 0.82 | 1.58 | 1.4 | 1.18E-03 | 7.19E-03 | (A) NE (B/D) |
| DTL | 3.26 | 1.86 | 0.93 | 1.38 | 1.4 | 1.92E-03 | 0.0104 | (A) NE (B/D) |
| KRTCAP2 | 4.39 | 2.99 | 0.92 | 1.51 | 1.4 | 2.12E-03 | 0.0112 | (A) NE (B/D) |
| BIVM | 6.74 | 5.35 | 1.01 | 0.89 | 1.4 | 2.76E-03 | 0.0137 | (A) NE (B/D) |
| ELOVL2-AS1 | 1.66 | 0.26 | 1.04 | 0.47 | 1.4 | 3.55E-03 | 0.0166 | (A) NE (B/D) |
| ATP5F1AP2 | 4.51 | 3.12 | 1.01 | 1.8 | 1.4 | 5.53E-03 | 0.0232 | (A) NE (B/D) |
| RUFY3 | 1.65 | 0.25 | 1.16 | 0.33 | 1.4 | 6.56E-03 | 0.0263 | (A) NE (B/D) |
| GPC5 | 1.53 | 0.14 | 1.18 | 0.54 | 1.4 | 7.47E-03 | 0.0289 | (A) NE (B/D) |
| RPL34P31 | 6.72 | 5.32 | 1.13 | 1.55 | 1.4 | 7.51E-03 | 0.029 | (A) NE (B/D) |
| ZNF671 | 2.42 | 1.02 | 1.18 | 0.93 | 1.4 | 7.56E-03 | 0.0291 | (A) NE (B/D) |
| SNPH | 1.52 | 0.12 | 1.36 | 0.29 | 1.4 | 0.015 | 0.0479 | (A) NE (B/D) |
| TMEM201 | 3.67 | 2.28 | 0.4 | 0.9 | 1.39 | 1.49E-07 | 6.62E-06 | (A) NE (B/D) |
| RAD23A | 4.62 | 3.23 | 0.16 | 1.23 | 1.39 | 1.84E-07 | 7.90E-06 | (A) NE (B/D) |
| CENPS | 3.12 | 1.73 | 0.39 | 1.21 | 1.39 | 1.36E-06 | 3.80E-05 | (A) NE (B/D) |
| NUP85 | 3.02 | 1.63 | 0.42 | 1.3 | 1.39 | 5.04E-06 | 1.06E-04 | (A) NE (B/D) |
| LSM10 | 3.19 | 1.8 | 0.3 | 1.51 | 1.39 | 1.21E-05 | 2.13E-04 | (A) NE (B/D) |
| RNASEK-C17 | 7.89 | 6.5 | 0.17 | 1.61 | 1.39 | 1.84E-05 | 2.89E-04 | (A) NE (B/D) |
| MRPL55 | 3.42 | 2.04 | 0.45 | 1.41 | 1.39 | 1.93E-05 | 3.00E-04 | (A) NE (B/D) |
| TMPO-AS1 | 3.68 | 2.29 | 0.6 | 1.07 | 1.39 | 3.33E-05 | 4.58E-04 | (A) NE (B/D) |
| UBA52 | 7.35 | 5.96 | 0.41 | 1.68 | 1.39 | 8.24E-05 | 9.33E-04 | (A) NE (B/D) |
| CIRBP-AS1 | 2.36 | 0.97 | 0.67 | 0.85 | 1.39 | 1.01E-04 | 1.10E-03 | (A) NE (B/D) |
| COX7C | 4.55 | 3.15 | 0.39 | 1.9 | 1.39 | 2.90E-04 | 2.45E-03 | (A) NE (B/D) |
| KLC1 | 2.41 | 1.02 | 0.76 | 0.54 | 1.39 | 3.75E-04 | 3.00E-03 | (A) NE (B/D) |
| ZMYND12 | 4.54 | 3.15 | 0.78 | 1.14 | 1.39 | 4.41E-04 | 3.40E-03 | (A) NE (B/D) |
| SAP25 | 7.84 | 6.45 | 0.71 | 1.57 | 1.39 | 5.19E-04 | 3.84E-03 | (A) NE (B/D) |
| TRA2A | 2.63 | 1.24 | 0.84 | 1.16 | 1.39 | 8.69E-04 | 5.69E-03 | (A) NE (B/D) |
| RHOJ | 4.6 | 3.21 | 0.95 | 1.66 | 1.39 | 3.50E-03 | 0.0164 | (A) NE (B/D) |
| GAPDHP65 | 2.52 | 1.14 | 1.04 | 1.19 | 1.39 | 3.81E-03 | 0.0175 | (A) NE (B/D) |
| DDX5 | 8.2 | 6.81 | 1.12 | 1.18 | 1.39 | 5.92E-03 | 0.0244 | (A) NE (B/D) |
| PIGS | 4.92 | 3.54 | 0.22 | 1.12 | 1.38 | 4.63E-08 | 2.60E-06 | (A) NE (B/D) |
| SIRT5 | 2.57 | 1.18 | 0.38 | 0.86 | 1.38 | 6.42E-08 | 3.36E-06 | (A) NE (B/D) |
| CSNK1D | 3 | 1.62 | 0.37 | 1.27 | 1.38 | 2.46E-06 | 6.03E-05 | (A) NE (B/D) |
| YTHDF3-AS1 | 6.48 | 5.11 | 0.33 | 1.37 | 1.38 | 4.50E-06 | 9.70E-05 | (A) NE (B/D) |
| TBC1D13 | 4.82 | 3.44 | 0.44 | 1.24 | 1.38 | 4.62E-06 | 9.92E-05 | (A) NE (B/D) |
| HMGB3P24 | 4.53 | 3.15 | 0.57 | 1.14 | 1.38 | 2.77E-05 | 3.96E-04 | (A) NE (B/D) |
| TXNDC12 | 5.66 | 4.28 | 0.64 | 1.1 | 1.38 | 7.41E-05 | 8.61E-04 | (A) NE (B/D) |
| UBE2E3 | 6.54 | 5.16 | 0.55 | 1.47 | 1.38 | 8.79E-05 | 9.84E-04 | (A) NE (B/D) |
| ENOPH1 | 3.26 | 1.88 | 0.61 | 1.36 | 1.38 | 1.04E-04 | 1.12E-03 | (A) NE (B/D) |
| TEKT4P2 | 8.21 | 6.82 | 0.66 | 1.29 | 1.38 | 1.47E-04 | 1.46E-03 | (A) NE (B/D) |
| HABP4 | 4.73 | 3.34 | 0.71 | 1.14 | 1.38 | 1.98E-04 | 1.83E-03 | (A) NE (B/D) |
| ARPC5L | 3.43 | 2.05 | 0.67 | 1.37 | 1.38 | 2.20E-04 | 1.98E-03 | (A) NE (B/D) |
| FDPSP8 | 1.49 | 0.11 | 0.73 | 0.25 | 1.38 | 4.09E-04 | 3.21E-03 | (A) NE (B/D) |
| THAP5P1 | 4.6 | 3.22 | 0.81 | 0.98 | 1.38 | 6.24E-04 | 4.43E-03 | (A) NE (B/D) |
| HNRNPDL | 5.58 | 4.2 | 0.88 | 1.3 | 1.38 | 1.46E-03 | 8.45E-03 | (A) NE (B/D) |
| CFL1P6 | 1.59 | 0.21 | 0.93 | 0.46 | 1.38 | 1.82E-03 | 9.99E-03 | (A) NE (B/D) |
| CDH24 | 1.69 | 0.31 | 1.09 | 0.76 | 1.38 | 5.01E-03 | 0.0215 | (A) NE (B/D) |
| RNU6-945P | 1.48 | 0.1 | 1.14 | 0.26 | 1.38 | 6.42E-03 | 0.0259 | (A) NE (B/D) |
| NPAS2 | 2.42 | 1.04 | 1.14 | 1.08 | 1.38 | 6.73E-03 | 0.0268 | (A) NE (B/D) |
| ZNF607 | 5.84 | 4.45 | 1.27 | 0.84 | 1.38 | 0.0114 | 0.0391 | (A) NE (B/D) |
| TAF6 | 1.69 | 0.32 | 0.19 | 0.58 | 1.37 | 3.44E-14 | 4.42E-11 | (A) NE (B/D) |
| TARBP2 | 1.99 | 0.62 | 0.36 | 0.81 | 1.37 | 2.02E-08 | 1.34E-06 | (A) NE (B/D) |
| KHSRPP1 | 4.81 | 3.44 | 0.4 | 0.93 | 1.37 | 2.16E-07 | 8.88E-06 | (A) NE (B/D) |
| SLC37A1 | 6.93 | 5.56 | 0.3 | 1.26 | 1.37 | 9.70E-07 | 2.92E-05 | (A) NE (B/D) |
| RBMX2 | 2.19 | 0.83 | 0.46 | 0.94 | 1.37 | 1.40E-06 | 3.88E-05 | (A) NE (B/D) |
| LAGE3 | 6.17 | 4.8 | 0.18 | 1.36 | 1.37 | 1.78E-06 | 4.70E-05 | (A) NE (B/D) |
| SNRPC | 3 | 1.63 | 0.27 | 1.38 | 1.37 | 3.31E-06 | 7.62E-05 | (A) NE (B/D) |
| SNRPA | 5.49 | 4.12 | 0.41 | 1.24 | 1.37 | 3.36E-06 | 7.71E-05 | (A) NE (B/D) |
| BCL2L2-PABP | 4.15 | 2.78 | 0.45 | 1.35 | 1.37 | 1.31E-05 | 2.26E-04 | (A) NE (B/D) |
| B4GALT2 | 7.6 | 6.23 | 0.55 | 1.14 | 1.37 | 2.01E-05 | 3.09E-04 | (A) NE (B/D) |

| | | | | | | | | |
|---|---|---|---|---|---|---|---|---|
| ANKRD19P | 4.3 | 2.92 | 0.62 | 0.84 | 1.37 | 4.50E-05 | 5.81E-04 | (A) NE (B/D) |
| MED10 | 2.84 | 1.47 | 0.55 | 1.34 | 1.37 | 4.82E-05 | 6.13E-04 | (A) NE (B/D) |
| NSUN5P1 | 7.6 | 6.22 | 0.68 | 1.01 | 1.37 | 1.25E-04 | 1.29E-03 | (A) NE (B/D) |
| PKN1 | 1.59 | 0.22 | 0.66 | 0.46 | 1.37 | 1.56E-04 | 1.52E-03 | (A) NE (B/D) |
| BRD3OS | 4.04 | 2.68 | 0.69 | 1.14 | 1.37 | 1.80E-04 | 1.70E-03 | (A) NE (B/D) |
| CCDC159 | 7.35 | 5.98 | 0.68 | 1.54 | 1.37 | 4.60E-04 | 3.51E-03 | (A) NE (B/D) |
| ARMC9 | 5.63 | 4.26 | 0.79 | 1.12 | 1.37 | 5.82E-04 | 4.21E-03 | (A) NE (B/D) |
| CCM2L | 3.48 | 2.11 | 0.8 | 1.35 | 1.37 | 8.12E-04 | 5.43E-03 | (A) NE (B/D) |
| REEP4 | 5.27 | 3.9 | 0.91 | 1.35 | 1.37 | 2.07E-03 | 0.011 | (A) NE (B/D) |
| MIR-4512/45 | 2.88 | 1.51 | 0.98 | 1.28 | 1.37 | 3.10E-03 | 0.015 | (A) NE (B/D) |
| TEX21P | 4.09 | 2.71 | 1.17 | 0.83 | 1.37 | 7.52E-03 | 0.029 | (A) NE (B/D) |
| MIR-4741/47 | 7.17 | 5.8 | 1.11 | 2.09 | 1.37 | 0.0136 | 0.0446 | (A) NE (B/D) |
| C1ORF122 | 4.47 | 3.11 | 0.33 | 0.89 | 1.36 | 1.86E-08 | 1.26E-06 | (A) NE (B/D) |
| SIPA1L3 | 3.58 | 2.22 | 0.3 | 1.01 | 1.36 | 3.40E-08 | 2.05E-06 | (A) NE (B/D) |
| CIZ1 | 2.12 | 0.76 | 0.37 | 0.77 | 1.36 | 4.21E-08 | 2.42E-06 | (A) NE (B/D) |
| CSNK1E | 5.49 | 4.13 | 0.35 | 0.95 | 1.36 | 6.86E-08 | 3.56E-06 | (A) NE (B/D) |
| IPO13 | 3.56 | 2.2 | 0.36 | 0.95 | 1.36 | 8.87E-08 | 4.35E-06 | (A) NE (B/D) |
| CMTR1 | 7.72 | 6.37 | 0.36 | 0.96 | 1.36 | 9.73E-08 | 4.69E-06 | (A) NE (B/D) |
| ERI3 | 6.04 | 4.68 | 0.38 | 1.09 | 1.36 | 4.97E-07 | 1.72E-05 | (A) NE (B/D) |
| MLLT6 | 2.48 | 1.12 | 0.34 | 1.16 | 1.36 | 5.53E-07 | 1.87E-05 | (A) NE (B/D) |
| RELL2 | 2.02 | 0.66 | 0.43 | 0.64 | 1.36 | 5.64E-07 | 1.90E-05 | (A) NE (B/D) |
| GBA2 | 7.46 | 6.1 | 0.29 | 1.21 | 1.36 | 5.65E-07 | 1.90E-05 | (A) NE (B/D) |
| PMPCA | 8.01 | 6.65 | 0.4 | 1.18 | 1.36 | 1.78E-06 | 4.71E-05 | (A) NE (B/D) |
| KLHL17 | 4.09 | 2.73 | 0.5 | 0.91 | 1.36 | 3.92E-06 | 8.74E-05 | (A) NE (B/D) |
| ADPRHL2 | 7.75 | 6.38 | 0.38 | 1.35 | 1.36 | 6.89E-06 | 1.36E-04 | (A) NE (B/D) |
| CIART | 1.82 | 0.46 | 0.55 | 0.83 | 1.36 | 1.13E-05 | 2.01E-04 | (A) NE (B/D) |
| DUTP6 | 5.1 | 3.74 | 0.52 | 1.13 | 1.36 | 1.31E-05 | 2.26E-04 | (A) NE (B/D) |
| MIR-4517/45 | 4.38 | 3.02 | 0.5 | 1.26 | 1.36 | 1.82E-05 | 2.87E-04 | (A) NE (B/D) |
| TEF | 7.64 | 6.28 | 0.44 | 1.39 | 1.36 | 1.89E-05 | 2.95E-04 | (A) NE (B/D) |
| AK7 | 2.89 | 1.52 | 0.57 | 1.05 | 1.36 | 2.31E-05 | 3.43E-04 | (A) NE (B/D) |
| TMED9 | 3.74 | 2.38 | 0.5 | 1.43 | 1.36 | 5.14E-05 | 6.46E-04 | (A) NE (B/D) |
| RPL18 | 4.83 | 3.47 | 0.54 | 1.43 | 1.36 | 8.09E-05 | 9.20E-04 | (A) NE (B/D) |
| NUDT3 | 4.13 | 2.77 | 0.64 | 1 | 1.36 | 8.49E-05 | 9.58E-04 | (A) NE (B/D) |
| MIR-1302-3/ | 2.9 | 1.54 | 0.59 | 1.4 | 1.36 | 1.18E-04 | 1.23E-03 | (A) NE (B/D) |
| ABCC10 | 1.42 | 0.06 | 0.61 | 0.19 | 1.36 | 1.34E-04 | 1.36E-03 | (A) NE (B/D) |
| SMIM20 | 4.55 | 3.19 | 0.66 | 1.22 | 1.36 | 1.48E-04 | 1.47E-03 | (A) NE (B/D) |
| SEC61B | 3.68 | 2.31 | 0.36 | 1.79 | 1.36 | 1.74E-04 | 1.65E-03 | (A) NE (B/D) |
| FUS | 3.32 | 1.96 | 0.22 | 1.87 | 1.36 | 2.05E-04 | 1.87E-03 | (A) NE (B/D) |
| SLX1B | 2.9 | 1.53 | 0.73 | 0.89 | 1.36 | 2.62E-04 | 2.26E-03 | (A) NE (B/D) |
| SLC4A3 | 1.39 | 0.03 | 0.95 | 0.1 | 1.36 | 2.64E-03 | 0.0133 | (A) NE (B/D) |
| MCOLN3 | 1.39 | 0.03 | 0.96 | 0.16 | 1.36 | 2.80E-03 | 0.0139 | (A) NE (B/D) |
| ZNF558 | 1.56 | 0.19 | 1.02 | 0.45 | 1.36 | 3.69E-03 | 0.0171 | (A) NE (B/D) |
| USP27X-AS1 | 4.06 | 2.71 | 1.08 | 1.3 | 1.36 | 6.12E-03 | 0.025 | (A) NE (B/D) |
| RAB12 | 2.51 | 1.15 | 1.24 | 0.86 | 1.36 | 0.0113 | 0.0388 | (A) NE (B/D) |
| EEF1A1P12 | 4.13 | 2.76 | 1.28 | 1.28 | 1.36 | 0.0144 | 0.0465 | (A) NE (B/D) |
| AP3D1 | 6.6 | 5.25 | 0.16 | 0.71 | 1.35 | 1.05E-12 | 8.41E-10 | (A) NE (B/D) |
| CARMIL1 | 3.41 | 2.07 | 0.24 | 0.69 | 1.35 | 1.85E-11 | 6.35E-09 | (A) NE (B/D) |
| PTBP1 | 10.09 | 8.74 | 0.23 | 0.87 | 1.35 | 6.24E-10 | 1.02E-07 | (A) NE (B/D) |
| USP22 | 3.77 | 2.42 | 0.29 | 1.07 | 1.35 | 9.56E-08 | 4.63E-06 | (A) NE (B/D) |
| AACS | 3.52 | 2.17 | 0.37 | 1.03 | 1.35 | 3.35E-07 | 1.25E-05 | (A) NE (B/D) |
| NFYC | 5.1 | 3.75 | 0.47 | 0.96 | 1.35 | 2.19E-06 | 5.50E-05 | (A) NE (B/D) |
| CCDC120 | 6.67 | 5.32 | 0.31 | 1.31 | 1.35 | 2.48E-06 | 6.08E-05 | (A) NE (B/D) |
| IFT27 | 7.09 | 5.74 | 0.5 | 1.01 | 1.35 | 5.51E-06 | 1.13E-04 | (A) NE (B/D) |
| PER2 | 1.63 | 0.28 | 0.49 | 0.43 | 1.35 | 8.72E-06 | 1.63E-04 | (A) NE (B/D) |
| ARHGEF7 | 2.5 | 1.14 | 0.66 | 0.72 | 1.35 | 1.16E-04 | 1.21E-03 | (A) NE (B/D) |
| MAFK | 3.36 | 2.01 | 0.69 | 0.93 | 1.35 | 1.73E-04 | 1.65E-03 | (A) NE (B/D) |
| RNU6-30P | 4.07 | 2.72 | 0.43 | 1.71 | 1.35 | 1.80E-04 | 1.70E-03 | (A) NE (B/D) |
| TUSC3 | 4.8 | 3.45 | 0.66 | 1.45 | 1.35 | 2.99E-04 | 2.52E-03 | (A) NE (B/D) |
| SLC25A5-AS1 | 3.74 | 2.38 | 0.66 | 1.45 | 1.35 | 3.00E-04 | 2.52E-03 | (A) NE (B/D) |
| GUSBP4 | 2.29 | 0.95 | 0.8 | 0.7 | 1.35 | 7.06E-04 | 4.88E-03 | (A) NE (B/D) |
| DHCR24 | 12.45 | 11.1 | 0.64 | 1.78 | 1.35 | 8.45E-04 | 5.58E-03 | (A) NE (B/D) |
| RBBP7 | 4.11 | 2.76 | 0.74 | 1.58 | 1.35 | 9.70E-04 | 6.18E-03 | (A) NE (B/D) |
| HERC2P7 | 6.03 | 4.68 | 0.84 | 0.98 | 1.35 | 9.72E-04 | 6.19E-03 | (A) NE (B/D) |
| P4HA2-AS1 | 3.43 | 2.08 | 0.75 | 1.6 | 1.35 | 1.08E-03 | 6.70E-03 | (A) NE (B/D) |
| STIL | 2.06 | 0.71 | 0.85 | 0.69 | 1.35 | 1.09E-03 | 6.71E-03 | (A) NE (B/D) |

| | | | | | | | | |
|---|---|---|---|---|---|---|---|---|
| GAPDH | 7.65 | 6.3 | 0.86 | 1.44 | 1.35 | 1.85E-03 | 0.0101 | (A) NE (B/D) |
| YWHAQP6 | 3.97 | 2.62 | 0.89 | 1.58 | 1.35 | 2.66E-03 | 0.0134 | (A) NE (B/D) |
| ZNF674-AS1 | 5.9 | 4.55 | 0.95 | 1.25 | 1.35 | 2.76E-03 | 0.0137 | (A) NE (B/D) |
| IGBP1-AS1 | 5.74 | 4.39 | 0.88 | 1.62 | 1.35 | 2.80E-03 | 0.0139 | (A) NE (B/D) |
| SMARCA1 | 7.34 | 5.99 | 0.88 | 1.72 | 1.35 | 3.16E-03 | 0.0152 | (A) NE (B/D) |
| SYT1 | 1.55 | 0.19 | 1.19 | 0.52 | 1.35 | 8.85E-03 | 0.0326 | (A) NE (B/D) |
| UBL5P2 | 9.53 | 8.19 | 1 | 2.07 | 1.35 | 9.91E-03 | 0.0355 | (A) NE (B/D) |
| DPY19L2 | 2.04 | 0.69 | 1.21 | 1.12 | 1.35 | 0.0107 | 0.0374 | (A) NE (B/D) |
| IRF9 | 2.31 | 0.96 | 1.22 | 1.05 | 1.35 | 0.0111 | 0.0385 | (A) NE (B/D) |
| EEF1A1P8 | 2.34 | 1 | 1.28 | 0.89 | 1.35 | 0.0139 | 0.0454 | (A) NE (B/D) |
| PPP1R35 | 1.59 | 0.25 | 0.31 | 0.47 | 1.34 | 5.01E-09 | 4.67E-07 | (A) NE (B/D) |
| EVI5L | 3.08 | 1.74 | 0.33 | 0.77 | 1.34 | 8.30E-09 | 6.80E-07 | (A) NE (B/D) |
| AGAP9 | 4.16 | 2.82 | 0.29 | 0.98 | 1.34 | 2.26E-08 | 1.44E-06 | (A) NE (B/D) |
| NSD2 | 1.63 | 0.3 | 0.37 | 0.67 | 1.34 | 3.70E-08 | 2.19E-06 | (A) NE (B/D) |
| HECTD3 | 4.44 | 3.11 | 0.34 | 1.01 | 1.34 | 1.30E-07 | 5.99E-06 | (A) NE (B/D) |
| LRRC56 | 3.19 | 1.85 | 0.38 | 0.99 | 1.34 | 2.94E-07 | 1.13E-05 | (A) NE (B/D) |
| TMEM80 | 6.5 | 5.15 | 0.37 | 1.05 | 1.34 | 4.04E-07 | 1.45E-05 | (A) NE (B/D) |
| DENND4B | 2.57 | 1.23 | 0.35 | 1.1 | 1.34 | 4.53E-07 | 1.59E-05 | (A) NE (B/D) |
| PSMC4 | 7.29 | 5.95 | 0.32 | 1.2 | 1.34 | 8.73E-07 | 2.67E-05 | (A) NE (B/D) |
| PLCXD1 | 6.74 | 5.4 | 0.43 | 1.02 | 1.34 | 1.29E-06 | 3.63E-05 | (A) NE (B/D) |
| MANBAL | 3.77 | 2.43 | 0.17 | 1.34 | 1.34 | 1.72E-06 | 4.58E-05 | (A) NE (B/D) |
| CCT6P1 | 8.68 | 7.33 | 0.47 | 0.85 | 1.34 | 1.86E-06 | 4.88E-05 | (A) NE (B/D) |
| UBALD2 | 1.86 | 0.52 | 0.43 | 0.39 | 1.34 | 2.36E-06 | 5.85E-05 | (A) NE (B/D) |
| HNRNPA0 | 5.18 | 3.84 | 0.32 | 1.29 | 1.34 | 2.54E-06 | 6.17E-05 | (A) NE (B/D) |
| BTN2A1 | 4.45 | 3.1 | 0.27 | 1.35 | 1.34 | 3.23E-06 | 7.45E-05 | (A) NE (B/D) |
| PPP6R1 | 6.64 | 5.3 | 0.49 | 0.65 | 1.34 | 3.70E-06 | 8.35E-05 | (A) NE (B/D) |
| ABHD17AP1 | 3.59 | 2.25 | 0.5 | 0.99 | 1.34 | 5.75E-06 | 1.18E-04 | (A) NE (B/D) |
| LINC00938 | 2.76 | 1.42 | 0.53 | 0.99 | 1.34 | 1.18E-05 | 2.08E-04 | (A) NE (B/D) |
| OGFOD3 | 5 | 3.66 | 0.55 | 0.96 | 1.34 | 1.63E-05 | 2.65E-04 | (A) NE (B/D) |
| INAFM1 | 5.46 | 4.13 | 0.53 | 1.08 | 1.34 | 1.68E-05 | 2.72E-04 | (A) NE (B/D) |
| LINC00174 | 5.88 | 4.54 | 0.62 | 1.06 | 1.34 | 7.31E-05 | 8.52E-04 | (A) NE (B/D) |
| BCYRN1 | 3.93 | 2.59 | 0.61 | 1.23 | 1.34 | 9.70E-05 | 1.06E-03 | (A) NE (B/D) |
| NFRKB | 4.69 | 3.35 | 0.65 | 0.76 | 1.34 | 1.11E-04 | 1.17E-03 | (A) NE (B/D) |
| FANCG | 4.83 | 3.49 | 0.65 | 1.21 | 1.34 | 1.62E-04 | 1.57E-03 | (A) NE (B/D) |
| SUMO2P1 | 7.27 | 5.93 | 0.54 | 1.55 | 1.34 | 1.70E-04 | 1.63E-03 | (A) NE (B/D) |
| TMEM50B | 7.56 | 6.22 | 0.69 | 1.09 | 1.34 | 2.11E-04 | 1.92E-03 | (A) NE (B/D) |
| STAMBPL1 | 3.59 | 2.25 | 0.71 | 0.95 | 1.34 | 2.32E-04 | 2.06E-03 | (A) NE (B/D) |
| NAIF1 | 4.54 | 3.2 | 0.71 | 0.94 | 1.34 | 2.35E-04 | 2.08E-03 | (A) NE (B/D) |
| ARL6IP4 | 3.47 | 2.12 | 0.61 | 1.5 | 1.34 | 2.47E-04 | 2.16E-03 | (A) NE (B/D) |
| SNHG10 | 5.7 | 4.36 | 0.73 | 0.95 | 1.34 | 2.93E-04 | 2.48E-03 | (A) NE (B/D) |
| GTF2IP1 | 10.4 | 9.06 | 0.78 | 0.79 | 1.34 | 5.88E-04 | 4.25E-03 | (A) NE (B/D) |
| TMEM175 | 2.58 | 1.24 | 0.88 | 0.8 | 1.34 | 1.38E-03 | 8.09E-03 | (A) NE (B/D) |
| PCDHB19P | 1.51 | 0.17 | 1.09 | 0.4 | 1.34 | 5.88E-03 | 0.0242 | (A) NE (B/D) |
| FNDC5 | 3.73 | 2.39 | 1.09 | 1.06 | 1.34 | 6.07E-03 | 0.0248 | (A) NE (B/D) |
| TRIM55 | 1.36 | 0.02 | 1.11 | 0.1 | 1.34 | 6.68E-03 | 0.0266 | (A) NE (B/D) |
| FAM72B | 3.86 | 2.52 | 1.05 | 1.67 | 1.34 | 7.64E-03 | 0.0293 | (A) NE (B/D) |
| CELF6 | 2.11 | 0.78 | 1.14 | 0.97 | 1.34 | 7.91E-03 | 0.03 | (A) NE (B/D) |
| BORA | 4.77 | 3.43 | 1.2 | 1.1 | 1.34 | 0.0107 | 0.0374 | (A) NE (B/D) |
| MIR-200C/20 | 2.37 | 1.03 | 1.2 | 1.31 | 1.34 | 0.0117 | 0.0399 | (A) NE (B/D) |
| MIR-200C/20 | 2.37 | 1.03 | 1.2 | 1.31 | 1.34 | 0.0117 | 0.0399 | (A) NE (B/D) |
| MIR-200C/3P | 2.37 | 1.03 | 1.2 | 1.31 | 1.34 | 0.0117 | 0.0399 | (A) NE (B/D) |
| MIR-200C/5P | 2.37 | 1.03 | 1.2 | 1.31 | 1.34 | 0.0117 | 0.0399 | (A) NE (B/D) |
| COMTD1 | 8.39 | 7.05 | 1.27 | 0.86 | 1.34 | 0.0138 | 0.0451 | (A) NE (B/D) |
| HLA-K | 3.48 | 2.13 | 1.29 | 1.27 | 1.34 | 0.0157 | 0.0496 | (A) NE (B/D) |
| HCFC1 | 5.26 | 3.93 | 0.23 | 0.61 | 1.33 | 3.28E-12 | 1.69E-09 | (A) NE (B/D) |
| CDC37 | 5.86 | 4.52 | 0.16 | 1.09 | 1.33 | 4.20E-08 | 2.42E-06 | (A) NE (B/D) |
| TPD52L2 | 5.61 | 4.28 | 0.37 | 0.95 | 1.33 | 1.51E-07 | 6.71E-06 | (A) NE (B/D) |
| PAK4 | 1.88 | 0.55 | 0.41 | 0.79 | 1.33 | 2.68E-07 | 1.05E-05 | (A) NE (B/D) |
| ARF5 | 3.08 | 1.75 | 0.29 | 1.15 | 1.33 | 3.71E-07 | 1.35E-05 | (A) NE (B/D) |
| NAA10 | 4.24 | 2.91 | 0.29 | 1.27 | 1.33 | 1.66E-06 | 4.44E-05 | (A) NE (B/D) |
| KLHL25 | 7 | 5.67 | 0.46 | 0.87 | 1.33 | 1.68E-06 | 4.49E-05 | (A) NE (B/D) |
| NOP56 | 5.14 | 3.8 | 0.36 | 1.23 | 1.33 | 2.19E-06 | 5.50E-05 | (A) NE (B/D) |
| ZNF32-AS1 | 6.99 | 5.66 | 0.29 | 1.38 | 1.33 | 6.36E-06 | 1.28E-04 | (A) NE (B/D) |
| PCED1A | 5.31 | 3.98 | 0.39 | 1.32 | 1.33 | 8.52E-06 | 1.61E-04 | (A) NE (B/D) |
| RHEBP1 | 3.99 | 2.66 | 0.48 | 1.22 | 1.33 | 1.47E-05 | 2.47E-04 | (A) NE (B/D) |

| | | | | | | | | |
|---|---|---|---|---|---|---|---|---|
| ZNF337 | 2.49 | 1.16 | 0.59 | 1.06 | 1.33 | 4.61E-05 | 5.93E-04 | (A) NE (B/D) |
| CLDN7 | 4.74 | 3.41 | 0.43 | 1.74 | 1.33 | 2.46E-04 | 2.16E-03 | (A) NE (B/D) |
| NPPA | 4.6 | 3.26 | 0.68 | 1.34 | 1.33 | 3.29E-04 | 2.70E-03 | (A) NE (B/D) |
| MIR-4751/47 | 7.7 | 6.37 | 0.59 | 1.62 | 1.33 | 4.04E-04 | 3.18E-03 | (A) NE (B/D) |
| RAB15 | 1.49 | 0.16 | 0.73 | 0.32 | 1.33 | 4.93E-04 | 3.70E-03 | (A) NE (B/D) |
| CEP95 | 7.9 | 6.57 | 0.8 | 0.88 | 1.33 | 7.68E-04 | 5.20E-03 | (A) NE (B/D) |
| ATP6V0E2-A | 6.52 | 5.19 | 0.8 | 1.21 | 1.33 | 9.26E-04 | 5.96E-03 | (A) NE (B/D) |
| WBP1LP2 | 4.46 | 3.12 | 0.91 | 0.9 | 1.33 | 1.88E-03 | 0.0102 | (A) NE (B/D) |
| LINC00667 | 4.49 | 3.16 | 0.93 | 0.95 | 1.33 | 2.25E-03 | 0.0117 | (A) NE (B/D) |
| HMGB3 | 1.9 | 0.57 | 0.99 | 0.7 | 1.33 | 3.35E-03 | 0.0159 | (A) NE (B/D) |
| ZNF510 | 2.57 | 1.24 | 1.02 | 0.76 | 1.33 | 4.17E-03 | 0.0188 | (A) NE (B/D) |
| SIRT2 | 1.76 | 0.43 | 1.03 | 0.6 | 1.33 | 4.51E-03 | 0.0199 | (A) NE (B/D) |
| KIFC3 | 5.07 | 3.73 | 1.06 | 1.05 | 1.33 | 5.31E-03 | 0.0225 | (A) NE (B/D) |
| FAM90A14P | 1.71 | 0.38 | 1.2 | 0.59 | 1.33 | 0.0102 | 0.0362 | (A) NE (B/D) |
| RNU2-22P | 2.8 | 1.47 | 1.19 | 1.06 | 1.33 | 0.0104 | 0.0368 | (A) NE (B/D) |
| SNORD54 | 3.39 | 2.06 | 1.21 | 1.04 | 1.33 | 0.0112 | 0.0387 | (A) NE (B/D) |
| ACTBP7 | 1.56 | 0.24 | 1.27 | 0.44 | 1.33 | 0.0141 | 0.0459 | (A) NE (B/D) |
| TMEM184B | 2.68 | 1.36 | 0.26 | 0.74 | 1.32 | 3.07E-10 | 5.77E-08 | (A) NE (B/D) |
| LY6G5C | 2.37 | 1.05 | 0.18 | 0.93 | 1.32 | 1.84E-09 | 2.20E-07 | (A) NE (B/D) |
| KLHL42 | 8.44 | 7.12 | 0.38 | 0.83 | 1.32 | 1.34E-07 | 6.14E-06 | (A) NE (B/D) |
| RNF25 | 3.81 | 2.49 | 0.24 | 1.18 | 1.32 | 3.83E-07 | 1.39E-05 | (A) NE (B/D) |
| ATPAF2 | 3.23 | 1.9 | 0.34 | 1.21 | 1.32 | 1.56E-06 | 4.25E-05 | (A) NE (B/D) |
| CENPV | 1.81 | 0.49 | 0.49 | 0.9 | 1.32 | 4.33E-06 | 9.46E-05 | (A) NE (B/D) |
| PKIG | 5.04 | 3.71 | 0.46 | 1.2 | 1.32 | 9.15E-06 | 1.69E-04 | (A) NE (B/D) |
| GOLGA6L4 | 3.32 | 2 | 0.5 | 1.07 | 1.32 | 9.74E-06 | 1.78E-04 | (A) NE (B/D) |
| AHSA2P | 4.72 | 3.4 | 0.51 | 1.09 | 1.32 | 1.36E-05 | 2.33E-04 | (A) NE (B/D) |
| LINC01002 | 2.26 | 0.94 | 0.41 | 1.42 | 1.32 | 2.68E-05 | 3.86E-04 | (A) NE (B/D) |
| ELOCP21 | 2.12 | 0.8 | 0.58 | 0.79 | 1.32 | 3.38E-05 | 4.65E-04 | (A) NE (B/D) |
| TTC25 | 4.34 | 3.02 | 0.54 | 1.24 | 1.32 | 4.77E-05 | 6.07E-04 | (A) NE (B/D) |
| LPGAT1 | 4.75 | 3.42 | 0.59 | 1.06 | 1.32 | 4.84E-05 | 6.15E-04 | (A) NE (B/D) |
| COX7A2P2 | 2.63 | 1.31 | 0.56 | 1.29 | 1.32 | 7.45E-05 | 8.65E-04 | (A) NE (B/D) |
| CDK10 | 4.33 | 3.01 | 0.65 | 0.95 | 1.32 | 1.25E-04 | 1.29E-03 | (A) NE (B/D) |
| MRPS15 | 8.14 | 6.82 | 0.64 | 1.2 | 1.32 | 1.56E-04 | 1.53E-03 | (A) NE (B/D) |
| TM9SF1 | 3.52 | 2.2 | 0.67 | 1.3 | 1.32 | 2.80E-04 | 2.39E-03 | (A) NE (B/D) |
| TPPP3 | 1.56 | 0.23 | 0.73 | 0.72 | 1.32 | 3.75E-04 | 3.00E-03 | (A) NE (B/D) |
| RHOBTB3 | 3.2 | 1.88 | 0.74 | 0.84 | 1.32 | 4.00E-04 | 3.15E-03 | (A) NE (B/D) |
| FAM92A | 1.49 | 0.17 | 0.86 | 0.4 | 1.32 | 1.58E-03 | 8.98E-03 | (A) NE (B/D) |
| SLC25A42 | 1.58 | 0.26 | 0.86 | 0.48 | 1.32 | 1.59E-03 | 9.00E-03 | (A) NE (B/D) |
| FAM171B | 2.25 | 0.93 | 0.91 | 0.88 | 1.32 | 2.03E-03 | 0.0109 | (A) NE (B/D) |
| HMGA1P8 | 2.5 | 1.19 | 0.94 | 0.81 | 1.32 | 2.57E-03 | 0.013 | (A) NE (B/D) |
| ARHGAP5-AS | 2.55 | 1.23 | 0.96 | 0.83 | 1.32 | 2.89E-03 | 0.0142 | (A) NE (B/D) |
| HNRNPH1 | 7.03 | 5.72 | 0.96 | 0.94 | 1.32 | 2.94E-03 | 0.0144 | (A) NE (B/D) |
| CCDC183 | 4.12 | 2.8 | 0.97 | 1.38 | 1.32 | 4.09E-03 | 0.0185 | (A) NE (B/D) |
| KLHL2 | 2.35 | 1.03 | 1.04 | 0.82 | 1.32 | 4.91E-03 | 0.0212 | (A) NE (B/D) |
| KIAA0319 | 1.35 | 0.03 | 1.24 | 0.17 | 1.32 | 0.0125 | 0.0419 | (A) NE (B/D) |
| VRK2 | 3.84 | 2.52 | 1.25 | 0.84 | 1.32 | 0.0133 | 0.044 | (A) NE (B/D) |
| GAPDHP70 | 5.82 | 4.5 | 1.25 | 1.24 | 1.32 | 0.0153 | 0.0486 | (A) NE (B/D) |
| PHF2 | 1.87 | 0.56 | 0.35 | 0.57 | 1.31 | 3.12E-08 | 1.90E-06 | (A) NE (B/D) |
| ABHD16A | 8.24 | 6.93 | 0.12 | 1.16 | 1.31 | 1.86E-07 | 7.96E-06 | (A) NE (B/D) |
| POLG | 3.79 | 2.48 | 0.42 | 0.67 | 1.31 | 5.73E-07 | 1.92E-05 | (A) NE (B/D) |
| AARS2 | 1.55 | 0.24 | 0.39 | 0.42 | 1.31 | 5.89E-07 | 1.95E-05 | (A) NE (B/D) |
| PNKP | 1.63 | 0.32 | 0.42 | 0.51 | 1.31 | 9.02E-07 | 2.73E-05 | (A) NE (B/D) |
| CYB5R1 | 7.31 | 6 | 0.39 | 1.22 | 1.31 | 4.24E-06 | 9.32E-05 | (A) NE (B/D) |
| PBX2 | 8.09 | 6.78 | 0.34 | 1.39 | 1.31 | 1.30E-05 | 2.26E-04 | (A) NE (B/D) |
| CAMSAP1 | 1.77 | 0.46 | 0.52 | 0.57 | 1.31 | 1.35E-05 | 2.31E-04 | (A) NE (B/D) |
| RN7SL481P | 2.42 | 1.11 | 0.56 | 0.85 | 1.31 | 2.14E-05 | 3.23E-04 | (A) NE (B/D) |
| MXD3 | 8.74 | 7.42 | 0.55 | 1.01 | 1.31 | 2.48E-05 | 3.62E-04 | (A) NE (B/D) |
| SDHAP3 | 7.85 | 6.54 | 0.56 | 0.97 | 1.31 | 2.73E-05 | 3.91E-04 | (A) NE (B/D) |
| NDUFA10 | 2.63 | 1.32 | 0.56 | 1.1 | 1.31 | 3.95E-05 | 5.24E-04 | (A) NE (B/D) |
| RPL10 | 10.3 | 8.98 | 0.29 | 1.64 | 1.31 | 7.79E-05 | 8.92E-04 | (A) NE (B/D) |
| C17ORF80 | 2.66 | 1.35 | 0.63 | 0.96 | 1.31 | 9.19E-05 | 1.02E-03 | (A) NE (B/D) |
| ARMC6 | 1.73 | 0.41 | 0.64 | 0.53 | 1.31 | 1.36E-04 | 1.37E-03 | (A) NE (B/D) |
| SMPD4 | 4.66 | 3.35 | 0.66 | 1.14 | 1.31 | 1.97E-04 | 1.82E-03 | (A) NE (B/D) |
| CRACR2B | 9 | 7.69 | 0.65 | 1.43 | 1.31 | 3.72E-04 | 2.98E-03 | (A) NE (B/D) |
| SDC3 | 5.61 | 4.29 | 0.73 | 1.18 | 1.31 | 4.52E-04 | 3.46E-03 | (A) NE (B/D) |

| | | | | | | | | |
|---|---|---|---|---|---|---|---|---|
| RHEBP2 | 9.58 | 8.27 | 0.7 | 1.34 | 1.31 | 4.63E-04 | 3.53E-03 | (A) NE (B/D) |
| RNU1-103P | 5.08 | 3.77 | 0.79 | 1.08 | 1.31 | 8.26E-04 | 5.49E-03 | (A) NE (B/D) |
| HMGN3-AS1 | 5.38 | 4.07 | 0.9 | 0.75 | 1.31 | 1.95E-03 | 0.0105 | (A) NE (B/D) |
| PET100 | 4.45 | 3.14 | 0.96 | 1.28 | 1.31 | 3.59E-03 | 0.0167 | (A) NE (B/D) |
| DOP1A | 1.88 | 0.57 | 1 | 0.63 | 1.31 | 3.98E-03 | 0.0181 | (A) NE (B/D) |
| FTH1P12 | 4.47 | 3.16 | 0.93 | 1.62 | 1.31 | 4.45E-03 | 0.0197 | (A) NE (B/D) |
| CCNB1 | 1.72 | 0.41 | 1.02 | 0.67 | 1.31 | 4.67E-03 | 0.0204 | (A) NE (B/D) |
| LINC01063 | 2.23 | 0.92 | 1.05 | 0.9 | 1.31 | 5.57E-03 | 0.0233 | (A) NE (B/D) |
| EIF3LP3 | 1.98 | 0.68 | 1.19 | 0.94 | 1.31 | 0.0114 | 0.0391 | (A) NE (B/D) |
| TUBGCP6 | 6.58 | 5.28 | 0.25 | 0.91 | 1.3 | 6.40E-09 | 5.66E-07 | (A) NE (B/D) |
| GGA3 | 2.16 | 0.86 | 0.37 | 0.92 | 1.3 | 2.04E-07 | 8.49E-06 | (A) NE (B/D) |
| UBL7 | 7.07 | 5.77 | 0.26 | 1.19 | 1.3 | 6.82E-07 | 2.19E-05 | (A) NE (B/D) |
| THAP4 | 5.63 | 4.32 | 0.41 | 0.92 | 1.3 | 7.03E-07 | 2.24E-05 | (A) NE (B/D) |
| ETV2 | 2.19 | 0.9 | 0.46 | 0.72 | 1.3 | 2.09E-06 | 5.31E-05 | (A) NE (B/D) |
| POLR2J4 | 7.78 | 6.48 | 0.46 | 0.87 | 1.3 | 2.18E-06 | 5.49E-05 | (A) NE (B/D) |
| AURKAIP1 | 2.89 | 1.59 | 0.4 | 1.12 | 1.3 | 2.43E-06 | 5.97E-05 | (A) NE (B/D) |
| CLIP2 | 2.45 | 1.14 | 0.47 | 0.75 | 1.3 | 2.93E-06 | 6.89E-05 | (A) NE (B/D) |
| CAMKK1 | 3.03 | 1.73 | 0.47 | 0.93 | 1.3 | 3.34E-06 | 7.67E-05 | (A) NE (B/D) |
| HNRNPLP2 | 6.61 | 5.3 | 0.32 | 1.27 | 1.3 | 3.44E-06 | 7.88E-05 | (A) NE (B/D) |
| CRB2 | 3.16 | 1.86 | 0.49 | 0.96 | 1.3 | 7.67E-06 | 1.48E-04 | (A) NE (B/D) |
| MSH5-SAPCD | 6.98 | 5.68 | 0.33 | 1.38 | 1.3 | 1.18E-05 | 2.07E-04 | (A) NE (B/D) |
| EPS8L2 | 2.86 | 1.56 | 0.52 | 1.12 | 1.3 | 2.36E-05 | 3.48E-04 | (A) NE (B/D) |
| DIS3L | 6.01 | 4.7 | 0.58 | 0.79 | 1.3 | 4.01E-05 | 5.31E-04 | (A) NE (B/D) |
| PITPNM1 | 5.69 | 4.39 | 0.6 | 0.99 | 1.3 | 6.14E-05 | 7.43E-04 | (A) NE (B/D) |
| CLBA1 | 7.09 | 5.79 | 0.57 | 1.21 | 1.3 | 6.91E-05 | 8.15E-04 | (A) NE (B/D) |
| RABGGTA | 1.73 | 0.43 | 0.61 | 0.49 | 1.3 | 1.12E-04 | 1.18E-03 | (A) NE (B/D) |
| C9ORF16 | 7.85 | 6.55 | 0.69 | 1.04 | 1.3 | 2.52E-04 | 2.19E-03 | (A) NE (B/D) |
| SMKR1 | 2.57 | 1.27 | 0.7 | 1.34 | 1.3 | 5.01E-04 | 3.75E-03 | (A) NE (B/D) |
| PRKRA | 8.1 | 6.79 | 0.77 | 0.96 | 1.3 | 6.30E-04 | 4.47E-03 | (A) NE (B/D) |
| FAN1 | 4.23 | 2.94 | 1.01 | 0.73 | 1.3 | 4.50E-03 | 0.0199 | (A) NE (B/D) |
| PITX1 | 3 | 1.7 | 1.11 | 1.14 | 1.3 | 8.45E-03 | 0.0315 | (A) NE (B/D) |
| SPTAN1 | 6.5 | 5.21 | 0.24 | 0.82 | 1.29 | 9.06E-10 | 1.27E-07 | (A) NE (B/D) |
| ZNF524 | 3.07 | 1.78 | 0.31 | 0.78 | 1.29 | 7.73E-09 | 6.48E-07 | (A) NE (B/D) |
| PREP | 1.97 | 0.68 | 0.32 | 0.8 | 1.29 | 1.15E-08 | 8.90E-07 | (A) NE (B/D) |
| CRB3 | 2.34 | 1.05 | 0.44 | 0.8 | 1.29 | 1.08E-06 | 3.19E-05 | (A) NE (B/D) |
| MRPL4 | 1.74 | 0.45 | 0.44 | 0.71 | 1.29 | 1.23E-06 | 3.48E-05 | (A) NE (B/D) |
| VPS16 | 9.39 | 8.1 | 0.47 | 0.92 | 1.29 | 4.62E-06 | 9.92E-05 | (A) NE (B/D) |
| PFKL | 4.97 | 3.68 | 0.49 | 0.75 | 1.29 | 4.66E-06 | 9.98E-05 | (A) NE (B/D) |
| PRPF6 | 4.9 | 3.61 | 0.38 | 1.26 | 1.29 | 6.89E-06 | 1.36E-04 | (A) NE (B/D) |
| PIGL | 2.13 | 0.84 | 0.51 | 0.98 | 1.29 | 1.39E-05 | 2.37E-04 | (A) NE (B/D) |
| ZNF90 | 2.83 | 1.54 | 0.22 | 1.47 | 1.29 | 1.77E-05 | 2.82E-04 | (A) NE (B/D) |
| MIS18A | 7.84 | 6.55 | 0.47 | 1.23 | 1.29 | 2.05E-05 | 3.14E-04 | (A) NE (B/D) |
| LPAR2 | 8.04 | 6.75 | 0.31 | 1.47 | 1.29 | 2.87E-05 | 4.08E-04 | (A) NE (B/D) |
| PTMS | 5.35 | 4.06 | 0.29 | 1.65 | 1.29 | 1.06E-04 | 1.14E-03 | (A) NE (B/D) |
| SLC16A6P1 | 3.42 | 2.12 | 0.64 | 0.88 | 1.29 | 1.15E-04 | 1.20E-03 | (A) NE (B/D) |
| PPIL6 | 5.14 | 3.85 | 0.64 | 0.91 | 1.29 | 1.18E-04 | 1.23E-03 | (A) NE (B/D) |
| BACE2 | 3.05 | 1.76 | 0.63 | 0.98 | 1.29 | 1.19E-04 | 1.24E-03 | (A) NE (B/D) |
| SLC2A6 | 5.81 | 4.52 | 0.64 | 0.98 | 1.29 | 1.28E-04 | 1.31E-03 | (A) NE (B/D) |
| LIN37 | 7.1 | 5.81 | 0.64 | 1.09 | 1.29 | 1.59E-04 | 1.55E-03 | (A) NE (B/D) |
| STAG3L3 | 3.43 | 2.15 | 0.65 | 0.63 | 1.29 | 1.73E-04 | 1.64E-03 | (A) NE (B/D) |
| UVSSA | 4.71 | 3.41 | 0.69 | 0.86 | 1.29 | 2.41E-04 | 2.12E-03 | (A) NE (B/D) |
| MRPL41 | 5.95 | 4.67 | 0.67 | 1.19 | 1.29 | 2.90E-04 | 2.46E-03 | (A) NE (B/D) |
| COQ8A | 3.06 | 1.77 | 0.63 | 1.44 | 1.29 | 4.14E-04 | 3.24E-03 | (A) NE (B/D) |
| SNORD36C | 7.51 | 6.22 | 0.52 | 1.72 | 1.29 | 5.26E-04 | 3.88E-03 | (A) NE (B/D) |
| RBAKDN | 1.48 | 0.2 | 0.76 | 0.74 | 1.29 | 6.14E-04 | 4.38E-03 | (A) NE (B/D) |
| NEO1 | 6.7 | 5.41 | 0.74 | 1.31 | 1.29 | 7.73E-04 | 5.22E-03 | (A) NE (B/D) |
| ARHGEF40 | 1.53 | 0.24 | 0.78 | 0.55 | 1.29 | 8.21E-04 | 5.47E-03 | (A) NE (B/D) |
| MNX1-AS2 | 3.13 | 1.84 | 0.8 | 1.11 | 1.29 | 1.03E-03 | 6.47E-03 | (A) NE (B/D) |
| PTGES2 | 4.96 | 3.67 | 0.85 | 0.96 | 1.29 | 1.53E-03 | 8.74E-03 | (A) NE (B/D) |
| GUCY1B1 | 1.7 | 0.41 | 0.91 | 0.53 | 1.29 | 2.59E-03 | 0.0131 | (A) NE (B/D) |
| LINC00240 | 2.14 | 0.85 | 0.93 | 0.58 | 1.29 | 2.84E-03 | 0.014 | (A) NE (B/D) |
| TTLL7 | 1.4 | 0.11 | 1 | 0.26 | 1.29 | 4.61E-03 | 0.0202 | (A) NE (B/D) |
| RN7SL404P | 2.53 | 1.24 | 0.97 | 1.35 | 1.29 | 4.70E-03 | 0.0205 | (A) NE (B/D) |
| AHI1 | 3.15 | 1.86 | 1.12 | 0.81 | 1.29 | 8.18E-03 | 0.0307 | (A) NE (B/D) |
| OSTN | 1.49 | 0.21 | 1.16 | 0.46 | 1.29 | 0.0106 | 0.0371 | (A) NE (B/D) |

| | | | | | | | | |
|---|---|---|---|---|---|---|---|---|
| RALYL | 1.41 | 0.12 | 1.2 | 0.37 | 1.29 | 0.0118 | 0.0402 | (A) NE (B/D) |
| RAB4B-EGLN | 6.12 | 4.84 | 0.23 | 0.98 | 1.28 | 2.27E-08 | 1.44E-06 | (A) NE (B/D) |
| DLL4 | 1.6 | 0.33 | 0.36 | 0.49 | 1.28 | 1.05E-07 | 5.02E-06 | (A) NE (B/D) |
| NARF | 1.49 | 0.21 | 0.37 | 0.53 | 1.28 | 1.55E-07 | 6.86E-06 | (A) NE (B/D) |
| PPME1 | 2.54 | 1.26 | 0.29 | 1.04 | 1.28 | 1.75E-07 | 7.58E-06 | (A) NE (B/D) |
| DCAF7 | 2.81 | 1.53 | 0.36 | 1.02 | 1.28 | 4.64E-07 | 1.62E-05 | (A) NE (B/D) |
| TEPSIN | 4.08 | 2.79 | 0.31 | 1.12 | 1.28 | 6.49E-07 | 2.11E-05 | (A) NE (B/D) |
| ZMYND8 | 1.47 | 0.19 | 0.39 | 0.38 | 1.28 | 1.12E-06 | 3.26E-05 | (A) NE (B/D) |
| TRIM28 | 2.64 | 1.36 | 0.43 | 0.99 | 1.28 | 2.15E-06 | 5.43E-05 | (A) NE (B/D) |
| TSEN34 | 4.3 | 3.02 | 0.45 | 0.99 | 1.28 | 3.80E-06 | 8.52E-05 | (A) NE (B/D) |
| SUPT7L | 5.5 | 4.22 | 0.49 | 0.94 | 1.28 | 8.51E-06 | 1.61E-04 | (A) NE (B/D) |
| WASH2P | 2.28 | 1 | 0.49 | 1.06 | 1.28 | 1.38E-05 | 2.36E-04 | (A) NE (B/D) |
| FASTK | 2.49 | 1.21 | 0.49 | 1.1 | 1.28 | 1.73E-05 | 2.77E-04 | (A) NE (B/D) |
| DNAJB11 | 5.96 | 4.68 | 0.48 | 1.17 | 1.28 | 1.84E-05 | 2.89E-04 | (A) NE (B/D) |
| TSNAXIP1 | 5.97 | 4.7 | 0.52 | 0.99 | 1.28 | 1.95E-05 | 3.02E-04 | (A) NE (B/D) |
| AMN | 3.6 | 2.32 | 0.41 | 1.39 | 1.28 | 3.87E-05 | 5.17E-04 | (A) NE (B/D) |
| MIR-877/3P | 1.74 | 0.47 | 0.58 | 0.63 | 1.28 | 5.96E-05 | 7.25E-04 | (A) NE (B/D) |
| MIR-877/5P | 1.74 | 0.47 | 0.58 | 0.63 | 1.28 | 5.96E-05 | 7.25E-04 | (A) NE (B/D) |
| MIR-877/877 | 1.74 | 0.47 | 0.58 | 0.63 | 1.28 | 5.96E-05 | 7.25E-04 | (A) NE (B/D) |
| MIR-877/877 | 1.74 | 0.47 | 0.58 | 0.63 | 1.28 | 5.96E-05 | 7.25E-04 | (A) NE (B/D) |
| PLEKHM1P1 | 7.53 | 6.25 | 0.59 | 0.72 | 1.28 | 6.21E-05 | 7.48E-04 | (A) NE (B/D) |
| DUS1L | 4.22 | 2.94 | 0.57 | 1.32 | 1.28 | 1.47E-04 | 1.46E-03 | (A) NE (B/D) |
| NCDN | 3.46 | 2.18 | 0.65 | 1.06 | 1.28 | 1.82E-04 | 1.71E-03 | (A) NE (B/D) |
| TSTD1 | 5.92 | 4.64 | 0.45 | 1.63 | 1.28 | 2.42E-04 | 2.13E-03 | (A) NE (B/D) |
| GYG1P1 | 1.79 | 0.51 | 0.71 | 0.67 | 1.28 | 3.93E-04 | 3.11E-03 | (A) NE (B/D) |
| GHDC | 3.78 | 2.5 | 0.67 | 1.35 | 1.28 | 4.66E-04 | 3.54E-03 | (A) NE (B/D) |
| KRBA2 | 3.86 | 2.58 | 0.74 | 0.86 | 1.28 | 5.18E-04 | 3.84E-03 | (A) NE (B/D) |
| C9ORF163 | 2.61 | 1.33 | 0.77 | 0.9 | 1.28 | 7.56E-04 | 5.14E-03 | (A) NE (B/D) |
| MSH2 | 1.72 | 0.44 | 0.78 | 0.76 | 1.28 | 8.10E-04 | 5.41E-03 | (A) NE (B/D) |
| MAGEA2B | 1.29 | 0.01 | 0.75 | 0.04 | 1.28 | 9.15E-04 | 5.91E-03 | (A) NE (B/D) |
| TRIM73 | 5.06 | 3.79 | 0.83 | 0.89 | 1.28 | 1.31E-03 | 7.78E-03 | (A) NE (B/D) |
| RFWD3 | 5.45 | 4.18 | 0.84 | 0.89 | 1.28 | 1.53E-03 | 8.74E-03 | (A) NE (B/D) |
| LURAP1 | 4.47 | 3.18 | 0.9 | 1.44 | 1.28 | 3.49E-03 | 0.0164 | (A) NE (B/D) |
| AMY1C | 2.11 | 0.83 | 0.99 | 0.87 | 1.28 | 4.44E-03 | 0.0197 | (A) NE (B/D) |
| CLSTN1 | 2.53 | 1.25 | 0.98 | 1.13 | 1.28 | 4.52E-03 | 0.0199 | (A) NE (B/D) |
| ZNF230 | 5.71 | 4.43 | 1 | 0.83 | 1.28 | 4.79E-03 | 0.0208 | (A) NE (B/D) |
| ZNF419 | 1.59 | 0.31 | 1.01 | 0.55 | 1.28 | 5.06E-03 | 0.0217 | (A) NE (B/D) |
| LAT2 | 7.66 | 6.38 | 0.95 | 1.59 | 1.28 | 5.95E-03 | 0.0244 | (A) NE (B/D) |
| ZNF546 | 1.44 | 0.16 | 1.11 | 0.36 | 1.28 | 8.56E-03 | 0.0318 | (A) NE (B/D) |
| CDK1 | 2.64 | 1.36 | 1.07 | 1.34 | 1.28 | 8.68E-03 | 0.0322 | (A) NE (B/D) |
| GPSM1 | 2.58 | 1.3 | 1.15 | 1.07 | 1.28 | 0.0108 | 0.0376 | (A) NE (B/D) |
| ENHO | 5.66 | 4.37 | 1.08 | 1.87 | 1.28 | 0.0141 | 0.0459 | (A) NE (B/D) |
| DEK | 3.82 | 2.54 | 1.21 | 1.21 | 1.28 | 0.0144 | 0.0464 | (A) NE (B/D) |
| GAPDHP73 | 1.89 | 0.6 | 1.25 | 0.8 | 1.28 | 0.0158 | 0.0498 | (A) NE (B/D) |
| NUP62 | 8.62 | 7.35 | 0.34 | 0.87 | 1.27 | 7.18E-08 | 3.70E-06 | (A) NE (B/D) |
| WDR73 | 8.31 | 7.04 | 0.43 | 0.76 | 1.27 | 9.41E-07 | 2.83E-05 | (A) NE (B/D) |
| ERGIC3 | 5.59 | 4.32 | 0.4 | 1.11 | 1.27 | 2.99E-06 | 7.01E-05 | (A) NE (B/D) |
| RBMXP2 | 3.65 | 2.38 | 0.48 | 0.8 | 1.27 | 4.44E-06 | 9.62E-05 | (A) NE (B/D) |
| ITFG2 | 2.67 | 1.4 | 0.46 | 1.09 | 1.27 | 9.38E-06 | 1.73E-04 | (A) NE (B/D) |
| H1FX | 2.69 | 1.42 | 0.56 | 0.97 | 1.27 | 3.90E-05 | 5.19E-04 | (A) NE (B/D) |
| LINC00896 | 1.74 | 0.47 | 0.66 | 0.84 | 1.27 | 1.85E-04 | 1.73E-03 | (A) NE (B/D) |
| CYFIP2 | 2.42 | 1.15 | 0.65 | 1.06 | 1.27 | 2.02E-04 | 1.85E-03 | (A) NE (B/D) |
| IFT52 | 7.33 | 6.07 | 0.61 | 1.27 | 1.27 | 2.21E-04 | 1.98E-03 | (A) NE (B/D) |
| NCKIPSD | 1.89 | 0.62 | 0.68 | 0.73 | 1.27 | 2.81E-04 | 2.39E-03 | (A) NE (B/D) |
| WDR6 | 9.34 | 8.08 | 0.56 | 1.54 | 1.27 | 3.73E-04 | 2.99E-03 | (A) NE (B/D) |
| LYSMD2 | 3.26 | 1.99 | 0.71 | 0.9 | 1.27 | 3.80E-04 | 3.03E-03 | (A) NE (B/D) |
| CYP2D7 | 4.42 | 3.14 | 0.72 | 1.09 | 1.27 | 5.13E-04 | 3.81E-03 | (A) NE (B/D) |
| RAD51 | 2.96 | 1.68 | 0.72 | 1.15 | 1.27 | 5.26E-04 | 3.88E-03 | (A) NE (B/D) |
| UFC1 | 5.18 | 3.91 | 0.56 | 1.65 | 1.27 | 6.25E-04 | 4.44E-03 | (A) NE (B/D) |
| NDUFA8 | 8.85 | 7.58 | 0.53 | 1.72 | 1.27 | 6.54E-04 | 4.60E-03 | (A) NE (B/D) |
| NGDN | 6.6 | 5.33 | 0.74 | 1.14 | 1.27 | 6.64E-04 | 4.65E-03 | (A) NE (B/D) |
| MEAF6 | 2.35 | 1.07 | 0.73 | 1.27 | 1.27 | 7.81E-04 | 5.26E-03 | (A) NE (B/D) |
| CES3 | 1.45 | 0.18 | 0.84 | 0.48 | 1.27 | 1.65E-03 | 9.26E-03 | (A) NE (B/D) |
| MIR-4467/44 | 1.62 | 0.35 | 0.91 | 0.58 | 1.27 | 2.85E-03 | 0.0141 | (A) NE (B/D) |
| RNU4-50P | 1.31 | 0.05 | 1.06 | 0.24 | 1.27 | 6.98E-03 | 0.0275 | (A) NE (B/D) |

| | | | | | | | | |
|---|---|---|---|---|---|---|---|---|
| CALM2P3 | 7.56 | 6.3 | 1.08 | 1.31 | 1.27 | 9.47E-03 | 0.0343 | (A) NE (B/D) |
| CYFIP1 | 3.61 | 2.35 | 0.26 | 0.71 | 1.26 | 5.18E-10 | 8.83E-08 | (A) NE (B/D) |
| MAP3K12 | 3.39 | 2.12 | 0.34 | 0.76 | 1.26 | 2.99E-08 | 1.84E-06 | (A) NE (B/D) |
| MGAT4B | 2.44 | 1.19 | 0.32 | 0.87 | 1.26 | 5.69E-08 | 3.06E-06 | (A) NE (B/D) |
| MRI1 | 2.31 | 1.05 | 0.37 | 0.87 | 1.26 | 2.85E-07 | 1.10E-05 | (A) NE (B/D) |
| MAN1B1 | 3.36 | 2.1 | 0.38 | 0.97 | 1.26 | 7.60E-07 | 2.40E-05 | (A) NE (B/D) |
| USF2 | 1.81 | 0.55 | 0.43 | 0.67 | 1.26 | 1.20E-06 | 3.42E-05 | (A) NE (B/D) |
| LZTR1 | 2.3 | 1.04 | 0.37 | 1.08 | 1.26 | 1.55E-06 | 4.22E-05 | (A) NE (B/D) |
| BTBD2 | 2.36 | 1.1 | 0.44 | 0.76 | 1.26 | 1.64E-06 | 4.42E-05 | (A) NE (B/D) |
| POLR2H | 3.54 | 2.28 | 0.31 | 1.27 | 1.26 | 5.36E-06 | 1.11E-04 | (A) NE (B/D) |
| CALR | 12.95 | 11.69 | 0.43 | 1.11 | 1.26 | 6.12E-06 | 1.24E-04 | (A) NE (B/D) |
| DNMT1 | 1.35 | 0.09 | 0.44 | 0.31 | 1.26 | 9.98E-06 | 1.81E-04 | (A) NE (B/D) |
| SDF2 | 2.85 | 1.59 | 0.42 | 1.35 | 1.26 | 3.27E-05 | 4.55E-04 | (A) NE (B/D) |
| EXOC7 | 6.2 | 4.94 | 0.55 | 1.08 | 1.26 | 5.25E-05 | 6.57E-04 | (A) NE (B/D) |
| ASMTL-AS1 | 7.79 | 6.53 | 0.51 | 1.23 | 1.26 | 5.33E-05 | 6.65E-04 | (A) NE (B/D) |
| KIF22 | 2.81 | 1.55 | 0.46 | 1.46 | 1.26 | 1.09E-04 | 1.16E-03 | (A) NE (B/D) |
| MIR-671/3P | 8.43 | 7.17 | 0.51 | 1.42 | 1.26 | 1.43E-04 | 1.43E-03 | (A) NE (B/D) |
| MIR-671/5P | 8.43 | 7.17 | 0.51 | 1.42 | 1.26 | 1.43E-04 | 1.43E-03 | (A) NE (B/D) |
| MIR-671/671 | 8.43 | 7.17 | 0.51 | 1.42 | 1.26 | 1.43E-04 | 1.43E-03 | (A) NE (B/D) |
| NUCB2 | 4.71 | 3.44 | 0.51 | 1.45 | 1.26 | 1.65E-04 | 1.59E-03 | (A) NE (B/D) |
| IER5L | 6.83 | 5.57 | 0.53 | 1.42 | 1.26 | 1.72E-04 | 1.64E-03 | (A) NE (B/D) |
| VWA5A | 1.41 | 0.15 | 0.74 | 0.28 | 1.26 | 8.21E-04 | 5.47E-03 | (A) NE (B/D) |
| SDCBP2 | 2.08 | 0.82 | 0.76 | 1.28 | 1.26 | 1.16E-03 | 7.05E-03 | (A) NE (B/D) |
| GBE1 | 1.54 | 0.28 | 0.85 | 0.58 | 1.26 | 1.84E-03 | 0.0101 | (A) NE (B/D) |
| PCDHAC1 | 1.72 | 0.47 | 0.92 | 0.61 | 1.26 | 3.22E-03 | 0.0154 | (A) NE (B/D) |
| LTBP3 | 1.87 | 0.61 | 1.04 | 0.82 | 1.26 | 6.46E-03 | 0.026 | (A) NE (B/D) |
| AGBL4 | 1.4 | 0.14 | 1.13 | 0.37 | 1.26 | 0.0101 | 0.0359 | (A) NE (B/D) |
| ANP32E | 5.15 | 3.89 | 1.09 | 1.43 | 1.26 | 0.0109 | 0.0378 | (A) NE (B/D) |
| PIMREG | 1.34 | 0.08 | 1.18 | 0.47 | 1.26 | 0.0125 | 0.0419 | (A) NE (B/D) |
| TP53TG5 | 3.85 | 2.61 | 0.28 | 0.57 | 1.25 | 5.06E-10 | 8.67E-08 | (A) NE (B/D) |
| C19ORF44 | 1.7 | 0.45 | 0.32 | 0.51 | 1.25 | 1.49E-08 | 1.06E-06 | (A) NE (B/D) |
| STK40 | 8.21 | 6.96 | 0.29 | 0.87 | 1.25 | 1.95E-08 | 1.30E-06 | (A) NE (B/D) |
| MED12 | 1.63 | 0.37 | 0.38 | 0.56 | 1.25 | 2.32E-07 | 9.34E-06 | (A) NE (B/D) |
| AGAP6 | 5.71 | 4.46 | 0.3 | 1.08 | 1.25 | 5.67E-07 | 1.90E-05 | (A) NE (B/D) |
| MCUR1 | 5.07 | 3.82 | 0.39 | 0.86 | 1.25 | 6.25E-07 | 2.05E-05 | (A) NE (B/D) |
| PRRC2B | 2.91 | 1.66 | 0.4 | 0.87 | 1.25 | 8.48E-07 | 2.61E-05 | (A) NE (B/D) |
| RN7SL657P | 4.53 | 3.29 | 0.44 | 0.85 | 1.25 | 2.44E-06 | 6.00E-05 | (A) NE (B/D) |
| ZNF691 | 7.43 | 6.18 | 0.41 | 1.22 | 1.25 | 1.31E-05 | 2.26E-04 | (A) NE (B/D) |
| CALM3 | 2.41 | 1.15 | 0.5 | 0.95 | 1.25 | 1.33E-05 | 2.29E-04 | (A) NE (B/D) |
| STAG3L2 | 2.61 | 1.36 | 0.5 | 0.51 | 1.25 | 2.00E-05 | 3.09E-04 | (A) NE (B/D) |
| NUMA1 | 5.92 | 4.67 | 0.52 | 0.96 | 1.25 | 2.36E-05 | 3.48E-04 | (A) NE (B/D) |
| KMT5C | 1.3 | 0.05 | 0.48 | 0.13 | 1.25 | 4.32E-05 | 5.65E-04 | (A) NE (B/D) |
| MOAP1 | 6.92 | 5.67 | 0.51 | 1.23 | 1.25 | 5.85E-05 | 7.15E-04 | (A) NE (B/D) |
| ATP5MC3 | 7.85 | 6.6 | 0.39 | 1.46 | 1.25 | 6.62E-05 | 7.87E-04 | (A) NE (B/D) |
| TMEM107 | 2.19 | 0.94 | 0.58 | 0.56 | 1.25 | 7.87E-05 | 8.98E-04 | (A) NE (B/D) |
| CLN6 | 2.62 | 1.36 | 0.6 | 1.01 | 1.25 | 1.10E-04 | 1.17E-03 | (A) NE (B/D) |
| EEF1DP2 | 2.64 | 1.39 | 0.63 | 0.89 | 1.25 | 1.61E-04 | 1.56E-03 | (A) NE (B/D) |
| PRDX2 | 7.84 | 6.58 | 0.49 | 1.5 | 1.25 | 2.04E-04 | 1.86E-03 | (A) NE (B/D) |
| SLC26A6 | 2.05 | 0.8 | 0.65 | 0.9 | 1.25 | 2.06E-04 | 1.88E-03 | (A) NE (B/D) |
| MAGOH | 9.41 | 8.16 | 0.5 | 1.49 | 1.25 | 2.20E-04 | 1.97E-03 | (A) NE (B/D) |
| ZNF692 | 4.54 | 3.28 | 0.65 | 1.04 | 1.25 | 2.27E-04 | 2.02E-03 | (A) NE (B/D) |
| COX6B1 | 5.65 | 4.39 | 0.56 | 1.59 | 1.25 | 5.33E-04 | 3.92E-03 | (A) NE (B/D) |
| EDF1 | 4.02 | 2.77 | 0.53 | 1.66 | 1.25 | 6.25E-04 | 4.44E-03 | (A) NE (B/D) |
| SNHG7 | 8.39 | 7.14 | 0.77 | 1.03 | 1.25 | 1.00E-03 | 6.32E-03 | (A) NE (B/D) |
| DUSP23 | 3 | 1.75 | 0.79 | 1.17 | 1.25 | 1.35E-03 | 7.96E-03 | (A) NE (B/D) |
| UCN | 2.78 | 1.52 | 0.84 | 0.85 | 1.25 | 1.71E-03 | 9.50E-03 | (A) NE (B/D) |
| MCM2 | 2.22 | 0.97 | 0.88 | 1.17 | 1.25 | 2.79E-03 | 0.0138 | (A) NE (B/D) |
| GTF2IRD2B | 1.6 | 0.35 | 0.94 | 0.55 | 1.25 | 3.63E-03 | 0.0168 | (A) NE (B/D) |
| EPM2A | 5.63 | 4.38 | 0.96 | 1.12 | 1.25 | 4.67E-03 | 0.0204 | (A) NE (B/D) |
| CALM2 | 2.48 | 1.23 | 0.98 | 1.06 | 1.25 | 5.17E-03 | 0.022 | (A) NE (B/D) |
| BGN | 3.13 | 1.88 | 0.95 | 1.57 | 1.25 | 6.79E-03 | 0.027 | (A) NE (B/D) |
| USP14 | 3.47 | 2.22 | 0.97 | 1.64 | 1.25 | 8.07E-03 | 0.0305 | (A) NE (B/D) |
| NUDT1 | 2.81 | 1.56 | 0.99 | 1.56 | 1.25 | 8.08E-03 | 0.0305 | (A) NE (B/D) |
| MIR-3122/31 | 1.86 | 0.61 | 1.06 | 1.14 | 1.25 | 8.40E-03 | 0.0314 | (A) NE (B/D) |
| RPS4Y1 | 5.72 | 4.46 | 0.95 | 1.79 | 1.25 | 9.00E-03 | 0.0331 | (A) NE (B/D) |

| | | | | | | | | |
|---|---|---|---|---|---|---|---|---|
| EFCAB7 | 1.98 | 0.73 | 1.14 | 0.52 | 1.25 | 0.0111 | 0.0383 | (A) NE (B/D) |
| CLPTM1LP1 | 1.34 | 0.09 | 1.22 | 0.28 | 1.25 | 0.015 | 0.048 | (A) NE (B/D) |
| HDAC5 | 2.25 | 1.01 | 0.33 | 0.76 | 1.24 | 3.68E-08 | 2.18E-06 | (A) NE (B/D) |
| ZNF688 | 2.59 | 1.35 | 0.32 | 0.81 | 1.24 | 3.79E-08 | 2.22E-06 | (A) NE (B/D) |
| DPH2 | 2.87 | 1.63 | 0.23 | 1.09 | 1.24 | 2.66E-07 | 1.05E-05 | (A) NE (B/D) |
| ZNF322P1 | 5.35 | 4.11 | 0.4 | 0.67 | 1.24 | 5.84E-07 | 1.94E-05 | (A) NE (B/D) |
| CHTOP | 7.88 | 6.64 | 0.31 | 1.09 | 1.24 | 9.02E-07 | 2.73E-05 | (A) NE (B/D) |
| ZNF767P | 7.68 | 6.44 | 0.43 | 0.75 | 1.24 | 1.39E-06 | 3.88E-05 | (A) NE (B/D) |
| TAF11 | 9.39 | 8.15 | 0.35 | 1.11 | 1.24 | 2.03E-06 | 5.19E-05 | (A) NE (B/D) |
| RBM22 | 8.42 | 7.18 | 0.47 | 0.93 | 1.24 | 8.19E-06 | 1.56E-04 | (A) NE (B/D) |
| PLA2G4B | 8.01 | 6.77 | 0.47 | 1.06 | 1.24 | 1.25E-05 | 2.18E-04 | (A) NE (B/D) |
| LOXL1-AS1 | 5.13 | 3.9 | 0.49 | 1.09 | 1.24 | 2.65E-05 | 3.82E-04 | (A) NE (B/D) |
| LRATD2 | 2.92 | 1.69 | 0.61 | 1.01 | 1.24 | 1.47E-04 | 1.46E-03 | (A) NE (B/D) |
| RAC1 | 3.68 | 2.45 | 0.57 | 1.25 | 1.24 | 1.60E-04 | 1.56E-03 | (A) NE (B/D) |
| NPIPA5 | 2.36 | 1.13 | 0.6 | 1.12 | 1.24 | 1.62E-04 | 1.57E-03 | (A) NE (B/D) |
| LNP1 | 6.04 | 4.79 | 0.73 | 0.97 | 1.24 | 6.23E-04 | 4.43E-03 | (A) NE (B/D) |
| FLVCR1-DT | 3.41 | 2.17 | 0.8 | 1.09 | 1.24 | 1.45E-03 | 8.40E-03 | (A) NE (B/D) |
| THTPA | 1.99 | 0.75 | 0.85 | 0.91 | 1.24 | 2.04E-03 | 0.0109 | (A) NE (B/D) |
| RNU4-14P | 1.93 | 0.69 | 0.92 | 1.08 | 1.24 | 3.61E-03 | 0.0168 | (A) NE (B/D) |
| MCM8 | 1.89 | 0.65 | 0.99 | 0.72 | 1.24 | 5.40E-03 | 0.0227 | (A) NE (B/D) |
| PCDHGB2 | 2.65 | 1.41 | 0.99 | 0.86 | 1.24 | 5.51E-03 | 0.0231 | (A) NE (B/D) |
| WDHD1 | 4.49 | 3.25 | 0.97 | 1.21 | 1.24 | 5.51E-03 | 0.0231 | (A) NE (B/D) |
| SNORD73A | 4.44 | 3.2 | 0.99 | 1.27 | 1.24 | 6.23E-03 | 0.0253 | (A) NE (B/D) |
| MCUB | 4.33 | 3.08 | 1.04 | 1.34 | 1.24 | 8.64E-03 | 0.0321 | (A) NE (B/D) |
| MYLK4 | 1.43 | 0.19 | 1.12 | 0.43 | 1.24 | 0.0102 | 0.0362 | (A) NE (B/D) |
| TMEM173 | 3.2 | 1.96 | 1.09 | 1.3 | 1.24 | 0.011 | 0.0381 | (A) NE (B/D) |
| ZNF318 | 3.15 | 1.91 | 0.32 | 0.63 | 1.23 | 1.28E-08 | 9.47E-07 | (A) NE (B/D) |
| ABCF1 | 2.03 | 0.8 | 0.31 | 0.81 | 1.23 | 2.46E-08 | 1.55E-06 | (A) NE (B/D) |
| SNORD52 | 2.14 | 0.91 | 0.34 | 0.78 | 1.23 | 7.39E-08 | 3.79E-06 | (A) NE (B/D) |
| ABL1 | 3.04 | 1.81 | 0.36 | 0.79 | 1.23 | 1.40E-07 | 6.35E-06 | (A) NE (B/D) |
| SAMD4B | 2.68 | 1.45 | 0.36 | 0.79 | 1.23 | 1.95E-07 | 8.21E-06 | (A) NE (B/D) |
| SLC27A3 | 1.73 | 0.5 | 0.38 | 0.59 | 1.23 | 2.62E-07 | 1.03E-05 | (A) NE (B/D) |
| ARAP1 | 7.16 | 5.94 | 0.27 | 1.05 | 1.23 | 3.41E-07 | 1.26E-05 | (A) NE (B/D) |
| TRAM2-AS1 | 6.13 | 4.9 | 0.28 | 1.07 | 1.23 | 4.98E-07 | 1.72E-05 | (A) NE (B/D) |
| ZNF554 | 4.34 | 3.12 | 0.34 | 0.99 | 1.23 | 5.31E-07 | 1.81E-05 | (A) NE (B/D) |
| TUBB6 | 3.35 | 2.11 | 0.38 | 0.94 | 1.23 | 7.77E-07 | 2.43E-05 | (A) NE (B/D) |
| TP53I13 | 2.19 | 0.96 | 0.42 | 0.58 | 1.23 | 1.55E-06 | 4.23E-05 | (A) NE (B/D) |
| AFG3L1P | 5.77 | 4.54 | 0.44 | 0.8 | 1.23 | 2.68E-06 | 6.42E-05 | (A) NE (B/D) |
| SRSF2 | 9.2 | 7.98 | 0.41 | 1.16 | 1.23 | 1.11E-05 | 1.98E-04 | (A) NE (B/D) |
| CDK5 | 1.94 | 0.71 | 0.48 | 0.93 | 1.23 | 1.15E-05 | 2.05E-04 | (A) NE (B/D) |
| RNF5 | 6.03 | 4.8 | 0.19 | 1.41 | 1.23 | 1.91E-05 | 2.98E-04 | (A) NE (B/D) |
| SMG7 | 2.46 | 1.23 | 0.5 | 1.02 | 1.23 | 2.54E-05 | 3.70E-04 | (A) NE (B/D) |
| INSR | 2.39 | 1.16 | 0.55 | 0.8 | 1.23 | 3.86E-05 | 5.17E-04 | (A) NE (B/D) |
| UBE2MP1 | 8.15 | 6.92 | 0.39 | 1.41 | 1.23 | 6.28E-05 | 7.56E-04 | (A) NE (B/D) |
| ZCCHC8 | 3.04 | 1.81 | 0.58 | 1.04 | 1.23 | 1.03E-04 | 1.11E-03 | (A) NE (B/D) |
| C11ORF71 | 2.83 | 1.6 | 0.57 | 1.12 | 1.23 | 1.08E-04 | 1.15E-03 | (A) NE (B/D) |
| PRSS53 | 4.81 | 3.58 | 0.62 | 0.91 | 1.23 | 1.59E-04 | 1.55E-03 | (A) NE (B/D) |
| ROGDI | 2.43 | 1.19 | 0.64 | 1.01 | 1.23 | 2.34E-04 | 2.08E-03 | (A) NE (B/D) |
| EXOSC9 | 6.85 | 5.62 | 0.66 | 1.16 | 1.23 | 3.80E-04 | 3.03E-03 | (A) NE (B/D) |
| C2ORF81 | 1.35 | 0.11 | 0.65 | 0.29 | 1.23 | 3.92E-04 | 3.11E-03 | (A) NE (B/D) |
| KLHL31 | 2.61 | 1.38 | 0.7 | 0.88 | 1.23 | 4.33E-04 | 3.34E-03 | (A) NE (B/D) |
| CHMP4A | 3.93 | 2.7 | 0.74 | 1.39 | 1.23 | 1.51E-03 | 8.65E-03 | (A) NE (B/D) |
| PCDHGB7 | 1.81 | 0.58 | 1.12 | 0.73 | 1.23 | 0.0109 | 0.038 | (A) NE (B/D) |
| CRIP1P4 | 2.49 | 1.25 | 1.1 | 1.4 | 1.23 | 0.0125 | 0.0419 | (A) NE (B/D) |
| ANKRD49 | 2.02 | 0.78 | 1.17 | 0.96 | 1.23 | 0.0143 | 0.0462 | (A) NE (B/D) |
| TRIM60P19 | 1.47 | 0.24 | 1.19 | 0.57 | 1.23 | 0.0148 | 0.0475 | (A) NE (B/D) |
| POM121B | 7.36 | 6.14 | 0.3 | 0.88 | 1.22 | 5.00E-08 | 2.78E-06 | (A) NE (B/D) |
| DOC2A | 2.49 | 1.27 | 0.33 | 0.85 | 1.22 | 1.14E-07 | 5.37E-06 | (A) NE (B/D) |
| MIR-5010/3P | 5.4 | 4.18 | 0.35 | 0.8 | 1.22 | 1.72E-07 | 7.47E-06 | (A) NE (B/D) |
| MIR-5010/5P | 5.4 | 4.18 | 0.35 | 0.8 | 1.22 | 1.72E-07 | 7.47E-06 | (A) NE (B/D) |
| FCHSD1 | 2.52 | 1.3 | 0.48 | 0.8 | 1.22 | 8.97E-06 | 1.67E-04 | (A) NE (B/D) |
| SLC38A10 | 2.61 | 1.4 | 0.48 | 0.88 | 1.22 | 1.13E-05 | 2.01E-04 | (A) NE (B/D) |
| MLXIP | 1.52 | 0.3 | 0.48 | 0.4 | 1.22 | 2.24E-05 | 3.34E-04 | (A) NE (B/D) |
| CCT6P2 | 3.06 | 1.85 | 0.52 | 0.94 | 1.22 | 3.11E-05 | 4.36E-04 | (A) NE (B/D) |
| GTF3A | 9.41 | 8.19 | 0.41 | 1.34 | 1.22 | 4.82E-05 | 6.13E-04 | (A) NE (B/D) |

| | | | | | | | | |
|---|---|---|---|---|---|---|---|---|
| MTND4P26 | 2.74 | 1.52 | 0.58 | 0.99 | 1.22 | 9.61E-05 | 1.06E-03 | (A) NE (B/D) |
| PGLS | 1.51 | 0.28 | 0.62 | 0.46 | 1.22 | 1.99E-04 | 1.83E-03 | (A) NE (B/D) |
| RFC2 | 2.02 | 0.8 | 0.63 | 0.88 | 1.22 | 2.03E-04 | 1.86E-03 | (A) NE (B/D) |
| BCL3 | 2.34 | 1.11 | 0.64 | 0.86 | 1.22 | 2.07E-04 | 1.89E-03 | (A) NE (B/D) |
| GRIPAP1 | 7.86 | 6.63 | 0.6 | 1.19 | 1.22 | 2.17E-04 | 1.96E-03 | (A) NE (B/D) |
| HNRNPDP1 | 2.74 | 1.52 | 0.63 | 1.02 | 1.22 | 2.42E-04 | 2.13E-03 | (A) NE (B/D) |
| MAP3K6 | 1.77 | 0.55 | 0.66 | 0.65 | 1.22 | 3.26E-04 | 2.68E-03 | (A) NE (B/D) |
| TCEAL3 | 6.03 | 4.81 | 0.55 | 1.64 | 1.22 | 7.83E-04 | 5.27E-03 | (A) NE (B/D) |
| DNTTIP1 | 5.4 | 4.18 | 0.68 | 1.32 | 1.22 | 8.07E-04 | 5.40E-03 | (A) NE (B/D) |
| DSP | 9.48 | 8.25 | 0.56 | 1.71 | 1.22 | 1.15E-03 | 7.04E-03 | (A) NE (B/D) |
| CTSF | 2.13 | 0.91 | 0.73 | 1.3 | 1.22 | 1.15E-03 | 7.04E-03 | (A) NE (B/D) |
| RBBP4 | 5.09 | 3.87 | 0.79 | 0.95 | 1.22 | 1.36E-03 | 7.97E-03 | (A) NE (B/D) |
| SKA2P1 | 4.81 | 3.59 | 0.71 | 1.45 | 1.22 | 1.44E-03 | 8.35E-03 | (A) NE (B/D) |
| OR8B7P | 1.44 | 0.21 | 0.91 | 0.45 | 1.22 | 3.53E-03 | 0.0165 | (A) NE (B/D) |
| KNSTRN | 1.44 | 0.22 | 0.91 | 0.43 | 1.22 | 3.67E-03 | 0.017 | (A) NE (B/D) |
| CACNA2D3-A | 1.41 | 0.19 | 0.92 | 0.34 | 1.22 | 3.79E-03 | 0.0174 | (A) NE (B/D) |
| HTATSF1 | 2.44 | 1.21 | 0.9 | 1.29 | 1.22 | 4.17E-03 | 0.0188 | (A) NE (B/D) |
| NHS | 1.96 | 0.74 | 1.17 | 0.86 | 1.22 | 0.0145 | 0.0467 | (A) NE (B/D) |
| HMGA1P1 | 1.66 | 0.44 | 1.2 | 0.52 | 1.22 | 0.0158 | 0.0499 | (A) NE (B/D) |
| MFSD11 | 11.06 | 9.85 | 0.15 | 0.66 | 1.21 | 4.10E-12 | 2.03E-09 | (A) NE (B/D) |
| FAM168A | 4.5 | 3.29 | 0.24 | 0.65 | 1.21 | 1.95E-10 | 4.08E-08 | (A) NE (B/D) |
| COPG2 | 2.98 | 1.76 | 0.27 | 0.75 | 1.21 | 2.78E-09 | 3.03E-07 | (A) NE (B/D) |
| KDM4A-AS1 | 3.12 | 1.91 | 0.27 | 0.73 | 1.21 | 2.95E-09 | 3.17E-07 | (A) NE (B/D) |
| VPS26C | 4.58 | 3.37 | 0.29 | 0.69 | 1.21 | 5.82E-09 | 5.24E-07 | (A) NE (B/D) |
| KIF3B | 6.39 | 5.18 | 0.34 | 0.82 | 1.21 | 1.15E-07 | 5.40E-06 | (A) NE (B/D) |
| PRKAB1 | 1.91 | 0.7 | 0.42 | 0.82 | 1.21 | 1.98E-06 | 5.11E-05 | (A) NE (B/D) |
| GART | 3.05 | 1.84 | 0.31 | 1.16 | 1.21 | 3.00E-06 | 7.02E-05 | (A) NE (B/D) |
| SLC25A25 | 5.94 | 4.73 | 0.47 | 0.86 | 1.21 | 7.84E-06 | 1.50E-04 | (A) NE (B/D) |
| IGSF8 | 3.73 | 2.52 | 0.39 | 1.2 | 1.21 | 1.46E-05 | 2.46E-04 | (A) NE (B/D) |
| PEF1 | 8.72 | 7.51 | 0.42 | 1.22 | 1.21 | 2.36E-05 | 3.48E-04 | (A) NE (B/D) |
| GPANK1 | 7.28 | 6.07 | 0.49 | 1.12 | 1.21 | 3.47E-05 | 4.74E-04 | (A) NE (B/D) |
| YARS | 7.08 | 5.87 | 0.54 | 0.84 | 1.21 | 4.16E-05 | 5.47E-04 | (A) NE (B/D) |
| YBX1P1 | 7.27 | 6.06 | 0.37 | 1.36 | 1.21 | 4.23E-05 | 5.55E-04 | (A) NE (B/D) |
| TMEM222 | 7.82 | 6.6 | 0.38 | 1.36 | 1.21 | 4.31E-05 | 5.64E-04 | (A) NE (B/D) |
| EIF1 | 3.16 | 1.95 | 0.37 | 1.42 | 1.21 | 6.56E-05 | 7.81E-04 | (A) NE (B/D) |
| ZNF786 | 1.94 | 0.73 | 0.57 | 0.75 | 1.21 | 7.63E-05 | 8.80E-04 | (A) NE (B/D) |
| ADI1 | 2.84 | 1.63 | 0.33 | 1.5 | 1.21 | 9.61E-05 | 1.06E-03 | (A) NE (B/D) |
| ATP9A | 7.51 | 6.3 | 0.47 | 1.35 | 1.21 | 9.84E-05 | 1.07E-03 | (A) NE (B/D) |
| C17ORF49 | 5.56 | 4.35 | 0.4 | 1.45 | 1.21 | 1.06E-04 | 1.14E-03 | (A) NE (B/D) |
| TOR2A | 1.78 | 0.57 | 0.61 | 0.66 | 1.21 | 1.50E-04 | 1.48E-03 | (A) NE (B/D) |
| PTPN23 | 5.07 | 3.86 | 0.58 | 1.17 | 1.21 | 1.74E-04 | 1.65E-03 | (A) NE (B/D) |
| SEPHS1P1 | 2.33 | 1.11 | 0.62 | 0.84 | 1.21 | 1.80E-04 | 1.70E-03 | (A) NE (B/D) |
| TIMM17B | 3.54 | 2.33 | 0.41 | 1.56 | 1.21 | 2.46E-04 | 2.16E-03 | (A) NE (B/D) |
| HSPA5 | 7.23 | 6.01 | 0.6 | 1.29 | 1.21 | 3.25E-04 | 2.68E-03 | (A) NE (B/D) |
| TCEAL8 | 9.35 | 8.14 | 0.59 | 1.44 | 1.21 | 5.39E-04 | 3.96E-03 | (A) NE (B/D) |
| RGS14 | 2.82 | 1.6 | 0.66 | 1.25 | 1.21 | 6.08E-04 | 4.35E-03 | (A) NE (B/D) |
| CSTB | 3.32 | 2.11 | 0.61 | 1.59 | 1.21 | 1.14E-03 | 6.97E-03 | (A) NE (B/D) |
| TPMT | 7.83 | 6.61 | 0.76 | 1.07 | 1.21 | 1.22E-03 | 7.34E-03 | (A) NE (B/D) |
| NLGN2 | 1.53 | 0.33 | 0.81 | 0.63 | 1.21 | 1.71E-03 | 9.50E-03 | (A) NE (B/D) |
| GPR137C | 1.42 | 0.21 | 0.92 | 0.56 | 1.21 | 3.96E-03 | 0.018 | (A) NE (B/D) |
| VWDE | 1.21 | 0.01 | 0.93 | 0.05 | 1.21 | 4.48E-03 | 0.0198 | (A) NE (B/D) |
| SOX2-OT | 1.76 | 0.55 | 0.99 | 0.68 | 1.21 | 5.97E-03 | 0.0245 | (A) NE (B/D) |
| GEM | 2.68 | 1.47 | 0.96 | 1.32 | 1.21 | 6.86E-03 | 0.0272 | (A) NE (B/D) |
| TMEM191C | 1.24 | 0.02 | 1.02 | 0.12 | 1.21 | 7.27E-03 | 0.0283 | (A) NE (B/D) |
| ZNF738 | 1.34 | 0.13 | 1.04 | 0.39 | 1.21 | 8.39E-03 | 0.0314 | (A) NE (B/D) |
| HMGB1P35 | 1.37 | 0.16 | 1.07 | 0.29 | 1.21 | 9.22E-03 | 0.0337 | (A) NE (B/D) |
| MIR-7-1/1* | 5.39 | 4.18 | 1.05 | 1.23 | 1.21 | 9.82E-03 | 0.0352 | (A) NE (B/D) |
| MIR-7-1/3P | 5.39 | 4.18 | 1.05 | 1.23 | 1.21 | 9.82E-03 | 0.0352 | (A) NE (B/D) |
| MIR-7-1/5P | 5.39 | 4.18 | 1.05 | 1.23 | 1.21 | 9.82E-03 | 0.0352 | (A) NE (B/D) |
| MIR-7-1/7 | 5.39 | 4.18 | 1.05 | 1.23 | 1.21 | 9.82E-03 | 0.0352 | (A) NE (B/D) |
| SP5 | 2.57 | 1.36 | 1.04 | 1.46 | 1.21 | 0.0114 | 0.0392 | (A) NE (B/D) |
| RPS2P1 | 1.57 | 0.36 | 1.13 | 0.51 | 1.21 | 0.0125 | 0.042 | (A) NE (B/D) |
| RPL35P2 | 6.72 | 5.51 | 1 | 1.8 | 1.21 | 0.0137 | 0.0449 | (A) NE (B/D) |
| GJC3 | 1.47 | 0.26 | 1.16 | 0.64 | 1.21 | 0.014 | 0.0456 | (A) NE (B/D) |
| TIRAP | 7.78 | 6.58 | 0.29 | 0.82 | 1.2 | 2.47E-08 | 1.56E-06 | (A) NE (B/D) |

| | | | | | | | | |
|---|---|---|---|---|---|---|---|---|
| PSMD8 | 8.45 | 7.25 | 0.23 | 1.16 | 1.2 | 1.68E-06 | 4.49E-05 | (A) NE (B/D) |
| PBX2P1 | 7.56 | 6.37 | 0.4 | 1.13 | 1.2 | 1.11E-05 | 1.98E-04 | (A) NE (B/D) |
| SULT1A4 | 4.41 | 3.21 | 0.28 | 1.29 | 1.2 | 1.15E-05 | 2.04E-04 | (A) NE (B/D) |
| AFG3L2 | 1.89 | 0.69 | 0.48 | 0.77 | 1.2 | 1.19E-05 | 2.09E-04 | (A) NE (B/D) |
| SNAPIN | 2.85 | 1.64 | 0.27 | 1.33 | 1.2 | 1.60E-05 | 2.61E-04 | (A) NE (B/D) |
| EIF2B5 | 7.34 | 6.14 | 0.47 | 1.1 | 1.2 | 3.05E-05 | 4.28E-04 | (A) NE (B/D) |
| ARHGEF25 | 2.68 | 1.47 | 0.38 | 1.32 | 1.2 | 3.83E-05 | 5.14E-04 | (A) NE (B/D) |
| TRAPPC3 | 8.11 | 6.91 | 0.45 | 1.23 | 1.2 | 4.37E-05 | 5.68E-04 | (A) NE (B/D) |
| SF3B5 | 9.17 | 7.97 | 0.41 | 1.4 | 1.2 | 9.56E-05 | 1.05E-03 | (A) NE (B/D) |
| AGBL5 | 7.04 | 5.84 | 0.63 | 0.9 | 1.2 | 2.13E-04 | 1.93E-03 | (A) NE (B/D) |
| YWHAE | 5.83 | 4.63 | 0.5 | 1.41 | 1.2 | 2.21E-04 | 1.98E-03 | (A) NE (B/D) |
| YBX1 | 5.94 | 4.74 | 0.58 | 1.28 | 1.2 | 2.65E-04 | 2.28E-03 | (A) NE (B/D) |
| GSE1 | 6.25 | 5.05 | 0.66 | 0.71 | 1.2 | 3.52E-04 | 2.85E-03 | (A) NE (B/D) |
| ELMO3 | 2.1 | 0.91 | 0.67 | 0.93 | 1.2 | 4.26E-04 | 3.30E-03 | (A) NE (B/D) |
| MSI2 | 2.51 | 1.31 | 0.73 | 0.64 | 1.2 | 8.33E-04 | 5.53E-03 | (A) NE (B/D) |
| TMEM164 | 2.19 | 0.99 | 0.74 | 0.7 | 1.2 | 8.56E-04 | 5.63E-03 | (A) NE (B/D) |
| APEX1 | 10.68 | 9.48 | 0.53 | 1.68 | 1.2 | 1.03E-03 | 6.45E-03 | (A) NE (B/D) |
| RNU4-39P | 2.13 | 0.93 | 0.75 | 0.97 | 1.2 | 1.06E-03 | 6.57E-03 | (A) NE (B/D) |
| MIR-4700/3P | 5.34 | 4.14 | 0.53 | 1.7 | 1.2 | 1.14E-03 | 6.97E-03 | (A) NE (B/D) |
| MIR-4700/5P | 5.34 | 4.14 | 0.53 | 1.7 | 1.2 | 1.14E-03 | 6.97E-03 | (A) NE (B/D) |
| KIF24 | 5.27 | 4.08 | 0.76 | 0.93 | 1.2 | 1.20E-03 | 7.29E-03 | (A) NE (B/D) |
| NEDD8-MDP1 | 5.02 | 3.82 | 0.67 | 1.49 | 1.2 | 1.43E-03 | 8.32E-03 | (A) NE (B/D) |
| FANCA | 2.3 | 1.1 | 0.79 | 0.97 | 1.2 | 1.46E-03 | 8.46E-03 | (A) NE (B/D) |
| FTOP1 | 2.93 | 1.73 | 0.85 | 1.01 | 1.2 | 2.68E-03 | 0.0134 | (A) NE (B/D) |
| C1QTNF1 | 4.21 | 3.01 | 0.74 | 1.63 | 1.2 | 3.16E-03 | 0.0152 | (A) NE (B/D) |
| MIR-5001/3P | 5.25 | 4.05 | 0.81 | 1.51 | 1.2 | 3.61E-03 | 0.0168 | (A) NE (B/D) |
| MIR-5001/5P | 5.25 | 4.05 | 0.81 | 1.51 | 1.2 | 3.61E-03 | 0.0168 | (A) NE (B/D) |
| COX6B1P5 | 3.63 | 2.43 | 0.91 | 1.06 | 1.2 | 4.38E-03 | 0.0195 | (A) NE (B/D) |
| NCAPD3 | 5.46 | 4.27 | 0.83 | 1.6 | 1.2 | 4.91E-03 | 0.0212 | (A) NE (B/D) |
| MPC1 | 2.56 | 1.37 | 0.91 | 1.28 | 1.2 | 4.98E-03 | 0.0214 | (A) NE (B/D) |
| KPNA5 | 1.42 | 0.23 | 1.04 | 0.47 | 1.2 | 8.40E-03 | 0.0314 | (A) NE (B/D) |
| CLASP2 | 1.33 | 0.14 | 1.04 | 0.23 | 1.2 | 8.82E-03 | 0.0326 | (A) NE (B/D) |
| PCDHGA12 | 1.67 | 0.47 | 1.05 | 0.64 | 1.2 | 8.87E-03 | 0.0327 | (A) NE (B/D) |
| OR8G3P | 1.44 | 0.25 | 1.16 | 0.55 | 1.2 | 0.015 | 0.0481 | (A) NE (B/D) |
| TMEM63B | 2.27 | 1.08 | 0.15 | 0.81 | 1.19 | 7.71E-10 | 1.15E-07 | (A) NE (B/D) |
| LINC00511 | 1.46 | 0.26 | 0.27 | 0.57 | 1.19 | 8.43E-10 | 1.23E-07 | (A) NE (B/D) |
| UNC45A | 7.96 | 6.77 | 0.27 | 0.82 | 1.19 | 1.48E-08 | 1.05E-06 | (A) NE (B/D) |
| RBM33 | 1.89 | 0.7 | 0.36 | 0.63 | 1.19 | 2.00E-07 | 8.36E-06 | (A) NE (B/D) |
| CAMTA2 | 2.25 | 1.06 | 0.36 | 0.84 | 1.19 | 4.02E-07 | 1.44E-05 | (A) NE (B/D) |
| EML3 | 6.59 | 5.41 | 0.33 | 0.97 | 1.19 | 6.71E-07 | 2.16E-05 | (A) NE (B/D) |
| TAF8 | 5.81 | 4.63 | 0.4 | 0.58 | 1.19 | 1.10E-06 | 3.22E-05 | (A) NE (B/D) |
| DNMT3A | 2.32 | 1.13 | 0.44 | 0.64 | 1.19 | 4.34E-06 | 9.47E-05 | (A) NE (B/D) |
| GNB1 | 6.73 | 5.54 | 0.33 | 1.19 | 1.19 | 7.09E-06 | 1.39E-04 | (A) NE (B/D) |
| TATDN3 | 5.03 | 3.84 | 0.42 | 1.05 | 1.19 | 1.05E-05 | 1.89E-04 | (A) NE (B/D) |
| USP35 | 4.48 | 3.3 | 0.48 | 0.66 | 1.19 | 1.38E-05 | 2.36E-04 | (A) NE (B/D) |
| DNAJB2 | 8.64 | 7.45 | 0.58 | 0.9 | 1.19 | 1.25E-04 | 1.28E-03 | (A) NE (B/D) |
| PHACTR4 | 4.78 | 3.59 | 0.68 | 0.81 | 1.19 | 4.49E-04 | 3.44E-03 | (A) NE (B/D) |
| BEND3P3 | 5.28 | 4.1 | 0.66 | 1.11 | 1.19 | 5.32E-04 | 3.92E-03 | (A) NE (B/D) |
| BICD1 | 3.57 | 2.38 | 0.7 | 0.67 | 1.19 | 5.88E-04 | 4.24E-03 | (A) NE (B/D) |
| TRAPPC2B | 3.36 | 2.17 | 0.73 | 1.33 | 1.19 | 1.58E-03 | 8.94E-03 | (A) NE (B/D) |
| CROCCP2 | 1.83 | 0.64 | 0.8 | 1.06 | 1.19 | 1.93E-03 | 0.0104 | (A) NE (B/D) |
| MIR-1234/12 | 2.01 | 0.82 | 0.87 | 0.85 | 1.19 | 3.02E-03 | 0.0147 | (A) NE (B/D) |
| MIR-1234/3P | 2.01 | 0.82 | 0.87 | 0.85 | 1.19 | 3.02E-03 | 0.0147 | (A) NE (B/D) |
| DUSP8P5 | 6.54 | 5.34 | 0.91 | 0.75 | 1.19 | 3.94E-03 | 0.0179 | (A) NE (B/D) |
| FTLP14 | 5.51 | 4.33 | 0.83 | 1.47 | 1.19 | 4.27E-03 | 0.0191 | (A) NE (B/D) |
| GGTLC3 | 1.72 | 0.53 | 1.05 | 0.72 | 1.19 | 9.64E-03 | 0.0348 | (A) NE (B/D) |
| LBX2-AS1 | 3.7 | 2.51 | 1.05 | 1.2 | 1.19 | 0.011 | 0.0382 | (A) NE (B/D) |
| FBP1 | 7.86 | 6.67 | 0.81 | 2.11 | 1.19 | 0.0125 | 0.042 | (A) NE (B/D) |
| CRNDE | 8.03 | 6.84 | 1.07 | 1.42 | 1.19 | 0.0138 | 0.0452 | (A) NE (B/D) |
| MFSD12 | 4.35 | 3.17 | 0.24 | 0.93 | 1.18 | 6.06E-08 | 3.21E-06 | (A) NE (B/D) |
| BCKDHA | 1.91 | 0.73 | 0.36 | 0.73 | 1.18 | 2.69E-07 | 1.05E-05 | (A) NE (B/D) |
| CLK2 | 4.26 | 3.07 | 0.3 | 1.04 | 1.18 | 8.80E-07 | 2.69E-05 | (A) NE (B/D) |
| LINC00957 | 1.65 | 0.46 | 0.38 | 0.44 | 1.18 | 1.14E-06 | 3.30E-05 | (A) NE (B/D) |
| GPN2 | 4.7 | 3.52 | 0.42 | 0.91 | 1.18 | 4.79E-06 | 1.02E-04 | (A) NE (B/D) |
| ITPA | 9.96 | 8.79 | 0.35 | 1.12 | 1.18 | 5.69E-06 | 1.17E-04 | (A) NE (B/D) |

| | | | | | | | | |
|---|---|---|---|---|---|---|---|---|
| RPL28 | 3.67 | 2.49 | 0.42 | 1.21 | 1.18 | 3.02E-05 | 4.26E-04 | (A) NE (B/D) |
| C19ORF71 | 1.39 | 0.21 | 0.5 | 0.41 | 1.18 | 4.12E-05 | 5.42E-04 | (A) NE (B/D) |
| GLOD4 | 3.94 | 2.76 | 0.46 | 1.18 | 1.18 | 4.75E-05 | 6.06E-04 | (A) NE (B/D) |
| SOX4 | 2.21 | 1.03 | 0.54 | 0.79 | 1.18 | 4.87E-05 | 6.18E-04 | (A) NE (B/D) |
| PPOX | 6.17 | 4.99 | 0.34 | 1.47 | 1.18 | 1.10E-04 | 1.17E-03 | (A) NE (B/D) |
| CFAP410 | 1.33 | 0.15 | 0.6 | 0.32 | 1.18 | 2.58E-04 | 2.24E-03 | (A) NE (B/D) |
| FAHD2CP | 2.79 | 1.61 | 0.6 | 1.22 | 1.18 | 3.39E-04 | 2.77E-03 | (A) NE (B/D) |
| PPAN-P2RY1 | 5.25 | 4.07 | 0.66 | 0.97 | 1.18 | 4.47E-04 | 3.43E-03 | (A) NE (B/D) |
| SNORD36A | 6.84 | 5.67 | 0.53 | 1.44 | 1.18 | 4.52E-04 | 3.46E-03 | (A) NE (B/D) |
| WASF1 | 1.49 | 0.3 | 0.76 | 0.5 | 1.18 | 1.26E-03 | 7.52E-03 | (A) NE (B/D) |
| TMEM129 | 2.84 | 1.66 | 0.76 | 0.96 | 1.18 | 1.34E-03 | 7.90E-03 | (A) NE (B/D) |
| BANF1 | 2.92 | 1.74 | 0.56 | 1.68 | 1.18 | 1.37E-03 | 8.01E-03 | (A) NE (B/D) |
| TSPYL4 | 5.3 | 4.12 | 0.75 | 1.14 | 1.18 | 1.51E-03 | 8.65E-03 | (A) NE (B/D) |
| MTERF4 | 4.09 | 2.91 | 0.76 | 1.08 | 1.18 | 1.56E-03 | 8.89E-03 | (A) NE (B/D) |
| RTN3P1 | 7.85 | 6.67 | 0.73 | 1.29 | 1.18 | 1.57E-03 | 8.92E-03 | (A) NE (B/D) |
| RPS4X | 6.58 | 5.4 | 0.35 | 1.95 | 1.18 | 1.87E-03 | 0.0102 | (A) NE (B/D) |
| TELO2 | 3.32 | 2.14 | 0.76 | 1.32 | 1.18 | 2.15E-03 | 0.0113 | (A) NE (B/D) |
| PKD1P6 | 6.4 | 5.22 | 0.84 | 0.67 | 1.18 | 2.46E-03 | 0.0126 | (A) NE (B/D) |
| TCTE3 | 2.61 | 1.42 | 0.85 | 0.58 | 1.18 | 2.81E-03 | 0.0139 | (A) NE (B/D) |
| TENM4 | 1.26 | 0.08 | 0.85 | 0.34 | 1.18 | 3.02E-03 | 0.0147 | (A) NE (B/D) |
| SAC3D1 | 2.63 | 1.45 | 0.88 | 1.02 | 1.18 | 3.90E-03 | 0.0178 | (A) NE (B/D) |
| RAD51-AS1 | 3.02 | 1.84 | 0.89 | 0.73 | 1.18 | 3.90E-03 | 0.0178 | (A) NE (B/D) |
| PCDHA13 | 1.43 | 0.25 | 0.94 | 0.45 | 1.18 | 5.20E-03 | 0.0221 | (A) NE (B/D) |
| NAIP | 2.44 | 1.26 | 0.96 | 1.29 | 1.18 | 7.80E-03 | 0.0297 | (A) NE (B/D) |
| PPM1G | 4.3 | 3.12 | 1 | 1.22 | 1.18 | 8.99E-03 | 0.0331 | (A) NE (B/D) |
| NSRP1 | 2.54 | 1.36 | 1.02 | 1.19 | 1.18 | 9.57E-03 | 0.0346 | (A) NE (B/D) |
| PBK | 1.34 | 0.16 | 1.14 | 0.47 | 1.18 | 0.0144 | 0.0466 | (A) NE (B/D) |
| MYO9B | 5.14 | 3.97 | 0.3 | 0.67 | 1.17 | 1.66E-08 | 1.15E-06 | (A) NE (B/D) |
| BAHD1 | 4.42 | 3.25 | 0.32 | 0.84 | 1.17 | 1.35E-07 | 6.17E-06 | (A) NE (B/D) |
| FAF1 | 3.94 | 2.77 | 0.44 | 0.62 | 1.17 | 5.10E-06 | 1.07E-04 | (A) NE (B/D) |
| PELP1 | 1.79 | 0.62 | 0.45 | 0.83 | 1.17 | 7.10E-06 | 1.39E-04 | (A) NE (B/D) |
| ATXN7L2 | 1.26 | 0.09 | 0.42 | 0.27 | 1.17 | 1.50E-05 | 2.50E-04 | (A) NE (B/D) |
| ATP6V0A1 | 1.58 | 0.41 | 0.48 | 0.57 | 1.17 | 1.67E-05 | 2.69E-04 | (A) NE (B/D) |
| KRTAP5-1 | 1.41 | 0.25 | 0.47 | 0.48 | 1.17 | 1.91E-05 | 2.98E-04 | (A) NE (B/D) |
| MSH5 | 5.73 | 4.57 | 0.34 | 1.27 | 1.17 | 2.58E-05 | 3.73E-04 | (A) NE (B/D) |
| POLH | 6.37 | 5.2 | 0.52 | 0.64 | 1.17 | 3.84E-05 | 5.15E-04 | (A) NE (B/D) |
| CCDC167 | 9.21 | 8.04 | 0.31 | 1.35 | 1.17 | 3.96E-05 | 5.26E-04 | (A) NE (B/D) |
| MPDU1 | 2.67 | 1.5 | 0.45 | 1.21 | 1.17 | 4.84E-05 | 6.15E-04 | (A) NE (B/D) |
| IFT46 | 3.69 | 2.53 | 0.51 | 1.04 | 1.17 | 6.31E-05 | 7.59E-04 | (A) NE (B/D) |
| CC2D1B | 2.38 | 1.22 | 0.56 | 0.8 | 1.17 | 8.91E-05 | 9.94E-04 | (A) NE (B/D) |
| ACTL6A | 7.37 | 6.2 | 0.4 | 1.35 | 1.17 | 9.01E-05 | 1.00E-03 | (A) NE (B/D) |
| RPS20P33 | 3.88 | 2.7 | 0.55 | 0.94 | 1.17 | 9.17E-05 | 1.02E-03 | (A) NE (B/D) |
| GORASP2 | 2.24 | 1.06 | 0.51 | 1.17 | 1.17 | 9.70E-05 | 1.06E-03 | (A) NE (B/D) |
| SMPD2 | 3.19 | 2.02 | 0.53 | 1.1 | 1.17 | 1.05E-04 | 1.13E-03 | (A) NE (B/D) |
| RIMS2 | 1.53 | 0.35 | 0.57 | 0.56 | 1.17 | 1.21E-04 | 1.25E-03 | (A) NE (B/D) |
| LAMP1 | 4.09 | 2.93 | 0.56 | 1.03 | 1.17 | 1.42E-04 | 1.42E-03 | (A) NE (B/D) |
| INTS4P2 | 5.39 | 4.21 | 0.6 | 0.66 | 1.17 | 1.66E-04 | 1.59E-03 | (A) NE (B/D) |
| NR2F1-AS1 | 2.28 | 1.11 | 0.61 | 0.98 | 1.17 | 2.32E-04 | 2.06E-03 | (A) NE (B/D) |
| PCK2 | 1.83 | 0.66 | 0.64 | 0.76 | 1.17 | 3.27E-04 | 2.69E-03 | (A) NE (B/D) |
| OCRL | 2.9 | 1.73 | 0.61 | 1.19 | 1.17 | 3.80E-04 | 3.03E-03 | (A) NE (B/D) |
| C6ORF118 | 1.78 | 0.61 | 0.66 | 0.96 | 1.17 | 4.62E-04 | 3.52E-03 | (A) NE (B/D) |
| DNASE1 | 1.7 | 0.53 | 0.67 | 0.67 | 1.17 | 4.63E-04 | 3.53E-03 | (A) NE (B/D) |
| NPIPP1 | 6.54 | 5.37 | 0.67 | 1 | 1.17 | 5.63E-04 | 4.10E-03 | (A) NE (B/D) |
| ZNF627 | 3.33 | 2.16 | 0.68 | 0.87 | 1.17 | 5.68E-04 | 4.13E-03 | (A) NE (B/D) |
| TMEM205 | 2.2 | 1.04 | 0.67 | 1.19 | 1.17 | 8.09E-04 | 5.41E-03 | (A) NE (B/D) |
| DUSP10 | 1.53 | 0.37 | 0.71 | 0.55 | 1.17 | 8.36E-04 | 5.54E-03 | (A) NE (B/D) |
| SLC37A4 | 3.74 | 2.57 | 0.51 | 1.68 | 1.17 | 1.18E-03 | 7.17E-03 | (A) NE (B/D) |
| FIGNL1 | 4.85 | 3.68 | 0.74 | 0.97 | 1.17 | 1.19E-03 | 7.24E-03 | (A) NE (B/D) |
| FBXL6 | 4.7 | 3.52 | 0.76 | 1.06 | 1.17 | 1.51E-03 | 8.68E-03 | (A) NE (B/D) |
| CHD1L | 4.73 | 3.56 | 0.75 | 1.12 | 1.17 | 1.52E-03 | 8.71E-03 | (A) NE (B/D) |
| EEF2 | 7.43 | 6.26 | 0.88 | 0.98 | 1.17 | 3.99E-03 | 0.0181 | (A) NE (B/D) |
| YAP1P1 | 1.56 | 0.39 | 0.97 | 0.71 | 1.17 | 6.58E-03 | 0.0263 | (A) NE (B/D) |
| RPS11P5 | 11.5 | 10.33 | 0.95 | 1.48 | 1.17 | 9.06E-03 | 0.0332 | (A) NE (B/D) |
| RPS3AP5 | 7.66 | 6.49 | 0.92 | 1.59 | 1.17 | 9.44E-03 | 0.0342 | (A) NE (B/D) |
| CLDN4 | 10.24 | 9.07 | 0.88 | 1.92 | 1.17 | 0.013 | 0.0432 | (A) NE (B/D) |

| | | | | | | | | |
|---|---|---|---|---|---|---|---|---|
| ZSWIM1 | 6.86 | 5.7 | 0.26 | 0.79 | 1.16 | 9.75E-09 | 7.79E-07 | (A) NE (B/D) |
| FAHD2A | 1.99 | 0.83 | 0.17 | 0.96 | 1.16 | 5.76E-08 | 3.09E-06 | (A) NE (B/D) |
| TMED4 | 3.1 | 1.94 | 0.13 | 1.05 | 1.16 | 2.80E-07 | 1.09E-05 | (A) NE (B/D) |
| CASKIN2 | 4.45 | 3.29 | 0.27 | 1.01 | 1.16 | 4.66E-07 | 1.63E-05 | (A) NE (B/D) |
| LINC00649 | 2 | 0.84 | 0.44 | 0.67 | 1.16 | 5.04E-06 | 1.06E-04 | (A) NE (B/D) |
| GIPC1 | 2.06 | 0.9 | 0.43 | 0.84 | 1.16 | 5.40E-06 | 1.12E-04 | (A) NE (B/D) |
| SNORD96A | 3.57 | 2.41 | 0.37 | 1.09 | 1.16 | 7.22E-06 | 1.41E-04 | (A) NE (B/D) |
| YEATS2 | 3.2 | 2.04 | 0.43 | 0.94 | 1.16 | 9.12E-06 | 1.69E-04 | (A) NE (B/D) |
| INTS7 | 1.33 | 0.18 | 0.45 | 0.41 | 1.16 | 1.51E-05 | 2.51E-04 | (A) NE (B/D) |
| TSC2 | 3.24 | 2.09 | 0.47 | 0.92 | 1.16 | 1.92E-05 | 2.99E-04 | (A) NE (B/D) |
| EXOSC2 | 6.4 | 5.24 | 0.46 | 1.04 | 1.16 | 2.82E-05 | 4.02E-04 | (A) NE (B/D) |
| CHTF18 | 1.31 | 0.15 | 0.51 | 0.3 | 1.16 | 7.87E-05 | 8.98E-04 | (A) NE (B/D) |
| SNORA70 | 4.29 | 3.13 | 0.53 | 1.2 | 1.16 | 1.63E-04 | 1.57E-03 | (A) NE (B/D) |
| SFXN1 | 4.15 | 2.98 | 0.59 | 0.62 | 1.16 | 1.78E-04 | 1.68E-03 | (A) NE (B/D) |
| CDK11A | 1.42 | 0.26 | 0.71 | 0.52 | 1.16 | 8.90E-04 | 5.79E-03 | (A) NE (B/D) |
| SYCE2 | 1.28 | 0.12 | 0.75 | 0.32 | 1.16 | 1.49E-03 | 8.60E-03 | (A) NE (B/D) |
| COX4I1P2 | 1.18 | 0.03 | 0.82 | 0.13 | 1.16 | 2.86E-03 | 0.0141 | (A) NE (B/D) |
| RALGDS | 2.32 | 1.16 | 0.84 | 0.95 | 1.16 | 2.96E-03 | 0.0144 | (A) NE (B/D) |
| HOXA9 | 1.25 | 0.09 | 0.85 | 0.39 | 1.16 | 3.27E-03 | 0.0156 | (A) NE (B/D) |
| NACAD | 1.19 | 0.03 | 0.87 | 0.12 | 1.16 | 3.99E-03 | 0.0181 | (A) NE (B/D) |
| CCDC173 | 1.32 | 0.17 | 0.9 | 0.27 | 1.16 | 4.59E-03 | 0.0202 | (A) NE (B/D) |
| SKA2 | 2.45 | 1.29 | 0.92 | 1.26 | 1.16 | 6.74E-03 | 0.0268 | (A) NE (B/D) |
| LINC00908 | 1.75 | 0.59 | 0.99 | 0.89 | 1.16 | 7.66E-03 | 0.0293 | (A) NE (B/D) |
| RNU4-78P | 2.99 | 1.83 | 0.98 | 1.32 | 1.16 | 9.92E-03 | 0.0355 | (A) NE (B/D) |
| TRIM60P18 | 4.91 | 3.75 | 1.11 | 0.91 | 1.16 | 0.0146 | 0.047 | (A) NE (B/D) |
| C11ORF49 | 1.65 | 0.51 | 0.28 | 0.5 | 1.15 | 3.69E-09 | 3.69E-07 | (A) NE (B/D) |
| DCPS | 2.54 | 1.39 | 0.28 | 0.77 | 1.15 | 2.31E-08 | 1.47E-06 | (A) NE (B/D) |
| CNOT9 | 8.45 | 7.3 | 0.24 | 0.87 | 1.15 | 2.91E-08 | 1.81E-06 | (A) NE (B/D) |
| CTTN | 6.1 | 4.94 | 0.22 | 0.97 | 1.15 | 1.32E-07 | 6.04E-06 | (A) NE (B/D) |
| COX5B | 4.08 | 2.93 | 0.31 | 0.87 | 1.15 | 2.48E-07 | 9.87E-06 | (A) NE (B/D) |
| GBAP1 | 3.12 | 1.96 | 0.27 | 1.01 | 1.15 | 6.42E-07 | 2.09E-05 | (A) NE (B/D) |
| MIR-943/943 | 7.13 | 5.98 | 0.38 | 0.59 | 1.15 | 7.83E-07 | 2.45E-05 | (A) NE (B/D) |
| CTSA | 5.95 | 4.81 | 0.34 | 1.05 | 1.15 | 3.83E-06 | 8.57E-05 | (A) NE (B/D) |
| ACOT8 | 4.98 | 3.82 | 0.46 | 0.87 | 1.15 | 1.39E-05 | 2.36E-04 | (A) NE (B/D) |
| PMF1-BGLAP | 6.48 | 5.33 | 0.3 | 1.27 | 1.15 | 2.11E-05 | 3.19E-04 | (A) NE (B/D) |
| BCS1L | 3.23 | 2.08 | 0.27 | 1.3 | 1.15 | 2.42E-05 | 3.55E-04 | (A) NE (B/D) |
| C1ORF50 | 4.35 | 3.2 | 0.49 | 0.93 | 1.15 | 3.72E-05 | 5.03E-04 | (A) NE (B/D) |
| ZNF561 | 2.8 | 1.64 | 0.53 | 0.81 | 1.15 | 5.79E-05 | 7.09E-04 | (A) NE (B/D) |
| WBP2 | 5.24 | 4.1 | 0.28 | 1.39 | 1.15 | 6.31E-05 | 7.58E-04 | (A) NE (B/D) |
| FUNDC2 | 5.56 | 4.41 | 0.45 | 1.26 | 1.15 | 9.21E-05 | 1.02E-03 | (A) NE (B/D) |
| TMEM234 | 3.57 | 2.42 | 0.55 | 0.9 | 1.15 | 1.01E-04 | 1.09E-03 | (A) NE (B/D) |
| SF3A3 | 6.87 | 5.73 | 0.54 | 1 | 1.15 | 1.16E-04 | 1.21E-03 | (A) NE (B/D) |
| RPS8 | 9.99 | 8.84 | 0.29 | 1.47 | 1.15 | 1.25E-04 | 1.29E-03 | (A) NE (B/D) |
| LRP11 | 2.41 | 1.26 | 0.58 | 0.96 | 1.15 | 1.84E-04 | 1.73E-03 | (A) NE (B/D) |
| EDC4 | 2.23 | 1.07 | 0.61 | 0.84 | 1.15 | 2.27E-04 | 2.02E-03 | (A) NE (B/D) |
| HNRNPA1P7 | 7.42 | 6.28 | 0.35 | 1.56 | 1.15 | 3.24E-04 | 2.67E-03 | (A) NE (B/D) |
| NDUFB11 | 7.24 | 6.09 | 0.38 | 1.57 | 1.15 | 3.94E-04 | 3.12E-03 | (A) NE (B/D) |
| MRPS12 | 5.99 | 4.84 | 0.64 | 1.22 | 1.15 | 6.75E-04 | 4.71E-03 | (A) NE (B/D) |
| CENPB | 6.58 | 5.44 | 0.69 | 0.7 | 1.15 | 7.15E-04 | 4.92E-03 | (A) NE (B/D) |
| SNAPC2 | 1.48 | 0.33 | 0.69 | 0.53 | 1.15 | 7.87E-04 | 5.30E-03 | (A) NE (B/D) |
| TUBB8 | 1.25 | 0.1 | 0.71 | 0.31 | 1.15 | 1.10E-03 | 6.76E-03 | (A) NE (B/D) |
| SRRM5 | 2.98 | 1.82 | 0.7 | 1.23 | 1.15 | 1.31E-03 | 7.74E-03 | (A) NE (B/D) |
| TP73-AS1 | 1.41 | 0.26 | 0.73 | 0.42 | 1.15 | 1.32E-03 | 7.78E-03 | (A) NE (B/D) |
| MGAT4A | 1.52 | 0.37 | 0.77 | 0.59 | 1.15 | 1.65E-03 | 9.28E-03 | (A) NE (B/D) |
| JUN | 8.5 | 7.34 | 0.7 | 1.39 | 1.15 | 1.92E-03 | 0.0104 | (A) NE (B/D) |
| ATOH7 | 1.96 | 0.81 | 0.79 | 0.91 | 1.15 | 2.03E-03 | 0.0109 | (A) NE (B/D) |
| NUDT15 | 2.4 | 1.25 | 0.78 | 1.1 | 1.15 | 2.38E-03 | 0.0122 | (A) NE (B/D) |
| BANF1P3 | 8.21 | 7.06 | 0.47 | 1.94 | 1.15 | 3.35E-03 | 0.0159 | (A) NE (B/D) |
| RRM1 | 3.71 | 2.56 | 0.79 | 1.41 | 1.15 | 3.65E-03 | 0.0169 | (A) NE (B/D) |
| ZNF331 | 6.22 | 5.06 | 0.88 | 0.73 | 1.15 | 3.99E-03 | 0.0181 | (A) NE (B/D) |
| SUMO2 | 4.25 | 3.1 | 0.86 | 1.63 | 1.15 | 8.23E-03 | 0.0309 | (A) NE (B/D) |
| JAM3 | 1.28 | 0.13 | 1 | 0.54 | 1.15 | 8.33E-03 | 0.0312 | (A) NE (B/D) |
| NEIL1 | 1.56 | 0.41 | 1 | 0.72 | 1.15 | 8.88E-03 | 0.0327 | (A) NE (B/D) |
| ALG1L12P | 1.85 | 0.7 | 1.03 | 0.74 | 1.15 | 0.0101 | 0.0358 | (A) NE (B/D) |
| C1QTNF2 | 1.46 | 0.31 | 1.07 | 0.6 | 1.15 | 0.0117 | 0.04 | (A) NE (B/D) |

| | | | | | | | | |
|---|---|---|---|---|---|---|---|---|
| GAS6-DT | 1.59 | 0.44 | 1.11 | 0.74 | 1.15 | 0.0147 | 0.0471 | (A) NE (B/D) |
| FAM160B2 | 2.29 | 1.14 | 1.08 | 1.15 | 1.15 | 0.0148 | 0.0474 | (A) NE (B/D) |
| NIPSNAP3A | 6.46 | 5.31 | 0.97 | 1.72 | 1.15 | 0.0151 | 0.0482 | (A) NE (B/D) |
| RETREG2 | 2.77 | 1.63 | 0.27 | 0.76 | 1.14 | 1.66E-08 | 1.15E-06 | (A) NE (B/D) |
| MTA3 | 6.26 | 5.12 | 0.31 | 0.95 | 1.14 | 6.50E-07 | 2.11E-05 | (A) NE (B/D) |
| UBE2M | 2.24 | 1.09 | 0.34 | 0.95 | 1.14 | 1.37E-06 | 3.83E-05 | (A) NE (B/D) |
| SNX15 | 6.94 | 5.8 | 0.23 | 1.14 | 1.14 | 2.97E-06 | 6.97E-05 | (A) NE (B/D) |
| ETV6 | 3.97 | 2.83 | 0.52 | 0.63 | 1.14 | 4.94E-05 | 6.25E-04 | (A) NE (B/D) |
| TRIM65 | 3.26 | 2.12 | 0.47 | 1.19 | 1.14 | 8.62E-05 | 9.70E-04 | (A) NE (B/D) |
| SNX30 | 4.71 | 3.58 | 0.55 | 0.76 | 1.14 | 9.33E-05 | 1.03E-03 | (A) NE (B/D) |
| ACOT7 | 1.47 | 0.33 | 0.63 | 0.57 | 1.14 | 3.98E-04 | 3.14E-03 | (A) NE (B/D) |
| RBCK1 | 5.87 | 4.74 | 0.56 | 1.25 | 1.14 | 4.00E-04 | 3.15E-03 | (A) NE (B/D) |
| SPG7 | 4.25 | 3.1 | 0.64 | 0.73 | 1.14 | 4.09E-04 | 3.21E-03 | (A) NE (B/D) |
| RHOT2 | 4.39 | 3.25 | 0.58 | 1.23 | 1.14 | 4.44E-04 | 3.42E-03 | (A) NE (B/D) |
| DYNLRB1 | 2.65 | 1.5 | 0.58 | 1.26 | 1.14 | 4.74E-04 | 3.59E-03 | (A) NE (B/D) |
| ILF2 | 7.35 | 6.21 | 0.57 | 1.33 | 1.14 | 5.53E-04 | 4.05E-03 | (A) NE (B/D) |
| H3F3C | 7.36 | 6.22 | 0.66 | 1.04 | 1.14 | 6.36E-04 | 4.50E-03 | (A) NE (B/D) |
| TFDP1 | 7.94 | 6.79 | 0.55 | 1.42 | 1.14 | 6.65E-04 | 4.66E-03 | (A) NE (B/D) |
| NDUFS6 | 2.52 | 1.39 | 0.62 | 1.28 | 1.14 | 7.33E-04 | 5.00E-03 | (A) NE (B/D) |
| ABHD11 | 3.31 | 2.17 | 0.64 | 1.24 | 1.14 | 7.72E-04 | 5.22E-03 | (A) NE (B/D) |
| DRC3 | 1.79 | 0.64 | 0.72 | 0.69 | 1.14 | 1.04E-03 | 6.50E-03 | (A) NE (B/D) |
| SOX12 | 6.08 | 4.93 | 0.65 | 1.33 | 1.14 | 1.18E-03 | 7.16E-03 | (A) NE (B/D) |
| CCDC183-AS1 | 4.09 | 2.95 | 0.72 | 1.21 | 1.14 | 1.56E-03 | 8.87E-03 | (A) NE (B/D) |
| ZNF428 | 2.85 | 1.72 | 0.78 | 1.08 | 1.14 | 2.45E-03 | 0.0125 | (A) NE (B/D) |
| S100PBP | 2.55 | 1.41 | 0.82 | 0.47 | 1.14 | 2.87E-03 | 0.0141 | (A) NE (B/D) |
| RN7SL589P | 2.36 | 1.21 | 0.84 | 1.08 | 1.14 | 3.56E-03 | 0.0166 | (A) NE (B/D) |
| TCF24 | 1.19 | 0.05 | 0.88 | 0.21 | 1.14 | 4.54E-03 | 0.02 | (A) NE (B/D) |
| PAK3 | 1.15 | 0.01 | 0.9 | 0.04 | 1.14 | 5.02E-03 | 0.0216 | (A) NE (B/D) |
| RPL36 | 5 | 3.87 | 0.85 | 1.4 | 1.14 | 5.97E-03 | 0.0245 | (A) NE (B/D) |
| CCT6P3 | 6.95 | 5.81 | 0.94 | 0.85 | 1.14 | 6.40E-03 | 0.0258 | (A) NE (B/D) |
| AIP | 8.31 | 7.17 | 0.89 | 1.35 | 1.14 | 6.89E-03 | 0.0273 | (A) NE (B/D) |
| CALY | 1.24 | 0.1 | 0.96 | 0.29 | 1.14 | 7.21E-03 | 0.0282 | (A) NE (B/D) |
| DPY19L2P1 | 1.42 | 0.28 | 0.96 | 0.75 | 1.14 | 7.26E-03 | 0.0283 | (A) NE (B/D) |
| PRKAA2 | 3.62 | 2.48 | 0.9 | 1.39 | 1.14 | 7.52E-03 | 0.029 | (A) NE (B/D) |
| DICER1-AS1 | 3.64 | 2.5 | 0.93 | 1.18 | 1.14 | 7.56E-03 | 0.0291 | (A) NE (B/D) |
| ZNF436-AS1 | 2.46 | 1.32 | 1 | 1.08 | 1.14 | 0.0102 | 0.0361 | (A) NE (B/D) |
| RPS7P11 | 10.05 | 8.91 | 0.95 | 1.47 | 1.14 | 0.0104 | 0.0367 | (A) NE (B/D) |
| RXFP4 | 1.21 | 0.07 | 1.05 | 0.24 | 1.14 | 0.0116 | 0.0397 | (A) NE (B/D) |
| C1ORF35 | 1.85 | 0.72 | 0.29 | 0.6 | 1.13 | 1.43E-08 | 1.03E-06 | (A) NE (B/D) |
| STK16 | 4.09 | 2.96 | 0.18 | 0.94 | 1.13 | 6.91E-08 | 3.58E-06 | (A) NE (B/D) |
| WASH5P | 7.18 | 6.05 | 0.17 | 0.95 | 1.13 | 8.48E-08 | 4.21E-06 | (A) NE (B/D) |
| ZNF581 | 3.66 | 2.53 | 0.36 | 0.88 | 1.13 | 1.57E-06 | 4.25E-05 | (A) NE (B/D) |
| SLC39A13 | 2.78 | 1.64 | 0.3 | 1.04 | 1.13 | 1.95E-06 | 5.04E-05 | (A) NE (B/D) |
| DIP2A | 5.49 | 4.36 | 0.41 | 0.58 | 1.13 | 3.02E-06 | 7.04E-05 | (A) NE (B/D) |
| DEDD2 | 7.93 | 6.8 | 0.38 | 1.01 | 1.13 | 6.64E-06 | 1.32E-04 | (A) NE (B/D) |
| MMP24 | 1.37 | 0.24 | 0.42 | 0.41 | 1.13 | 9.88E-06 | 1.80E-04 | (A) NE (B/D) |
| JMJD7-PLA2G | 3.66 | 2.52 | 0.44 | 1 | 1.13 | 2.05E-05 | 3.14E-04 | (A) NE (B/D) |
| DNPEP | 2.09 | 0.96 | 0.39 | 1.13 | 1.13 | 2.14E-05 | 3.23E-04 | (A) NE (B/D) |
| PRMT2 | 2.08 | 0.95 | 0.49 | 0.78 | 1.13 | 3.03E-05 | 4.26E-04 | (A) NE (B/D) |
| NDUFV1 | 3.99 | 2.85 | 0.41 | 1.36 | 1.13 | 1.46E-04 | 1.45E-03 | (A) NE (B/D) |
| CWC15 | 5.72 | 4.59 | 0.47 | 1.28 | 1.13 | 1.78E-04 | 1.68E-03 | (A) NE (B/D) |
| CHN1 | 1.3 | 0.17 | 0.57 | 0.41 | 1.13 | 2.06E-04 | 1.88E-03 | (A) NE (B/D) |
| ANKRA2 | 7.13 | 6 | 0.59 | 0.88 | 1.13 | 2.39E-04 | 2.11E-03 | (A) NE (B/D) |
| URGCP-MRP | 10.12 | 8.99 | 0.55 | 1.43 | 1.13 | 7.13E-04 | 4.91E-03 | (A) NE (B/D) |
| MORN2 | 7.02 | 5.89 | 0.59 | 1.33 | 1.13 | 7.34E-04 | 5.01E-03 | (A) NE (B/D) |
| TPRN | 1.59 | 0.46 | 0.69 | 0.62 | 1.13 | 8.51E-04 | 5.61E-03 | (A) NE (B/D) |
| GABARAP | 2.97 | 1.83 | 0.4 | 1.69 | 1.13 | 1.01E-03 | 6.37E-03 | (A) NE (B/D) |
| PDCD7 | 2.09 | 0.97 | 0.73 | 0.86 | 1.13 | 1.34E-03 | 7.89E-03 | (A) NE (B/D) |
| C1ORF159 | 1.45 | 0.33 | 0.75 | 0.54 | 1.13 | 1.68E-03 | 9.37E-03 | (A) NE (B/D) |
| TCEAL4 | 7.03 | 5.91 | 0.44 | 1.82 | 1.13 | 2.24E-03 | 0.0116 | (A) NE (B/D) |
| GALT | 6.07 | 4.94 | 0.71 | 1.47 | 1.13 | 3.02E-03 | 0.0146 | (A) NE (B/D) |
| PSENEN | 2.8 | 1.67 | 0.75 | 1.36 | 1.13 | 3.21E-03 | 0.0154 | (A) NE (B/D) |
| FOXH1 | 6.57 | 5.44 | 0.43 | 1.96 | 1.13 | 3.66E-03 | 0.0169 | (A) NE (B/D) |
| PCMTD2 | 1.54 | 0.42 | 0.86 | 0.7 | 1.13 | 3.96E-03 | 0.018 | (A) NE (B/D) |
| OXCT2P1 | 2.38 | 1.25 | 0.87 | 0.65 | 1.13 | 4.28E-03 | 0.0192 | (A) NE (B/D) |

| | | | | | | | | |
|---|---|---|---|---|---|---|---|---|
| PAQR4 | 3.05 | 1.92 | 0.85 | 1.1 | 1.13 | 4.37E-03 | 0.0194 | (A) NE (B/D) |
| RNU6-1053P | 3.28 | 2.15 | 0.98 | 0.84 | 1.13 | 8.76E-03 | 0.0324 | (A) NE (B/D) |
| ARL6IP1 | 3.37 | 2.24 | 0.89 | 1.53 | 1.13 | 8.99E-03 | 0.033 | (A) NE (B/D) |
| TDP2 | 8.49 | 7.36 | 1.01 | 1.16 | 1.13 | 0.0121 | 0.041 | (A) NE (B/D) |
| MB21D2 | 2.89 | 1.76 | 1.09 | 0.96 | 1.13 | 0.0153 | 0.0487 | (A) NE (B/D) |
| RNF44 | 8.18 | 7.06 | 0.32 | 0.72 | 1.12 | 9.88E-08 | 4.74E-06 | (A) NE (B/D) |
| SLC5A6 | 1.83 | 0.71 | 0.33 | 0.88 | 1.12 | 6.07E-07 | 2.00E-05 | (A) NE (B/D) |
| LSM14B | 3.5 | 2.38 | 0.24 | 1.05 | 1.12 | 1.14E-06 | 3.30E-05 | (A) NE (B/D) |
| CHKB-CPT1B | 8.34 | 7.23 | 0.33 | 1.05 | 1.12 | 4.61E-06 | 9.92E-05 | (A) NE (B/D) |
| ZER1 | 3.79 | 2.66 | 0.43 | 1.03 | 1.12 | 2.36E-05 | 3.48E-04 | (A) NE (B/D) |
| KAT5 | 6.12 | 5.01 | 0.41 | 1.14 | 1.12 | 3.64E-05 | 4.94E-04 | (A) NE (B/D) |
| ARSA | 4.81 | 3.69 | 0.46 | 1.06 | 1.12 | 4.71E-05 | 6.02E-04 | (A) NE (B/D) |
| FURIN | 4.61 | 3.5 | 0.48 | 0.98 | 1.12 | 4.95E-05 | 6.27E-04 | (A) NE (B/D) |
| TSPAN3 | 6.21 | 5.08 | 0.49 | 1.2 | 1.12 | 1.42E-04 | 1.42E-03 | (A) NE (B/D) |
| PPIB | 4.26 | 3.14 | 0.35 | 1.44 | 1.12 | 1.90E-04 | 1.77E-03 | (A) NE (B/D) |
| LRRC75A | 8.46 | 7.34 | 0.33 | 1.48 | 1.12 | 2.16E-04 | 1.95E-03 | (A) NE (B/D) |
| YBX1P10 | 4.68 | 3.56 | 0.49 | 1.32 | 1.12 | 2.74E-04 | 2.34E-03 | (A) NE (B/D) |
| SCRN1 | 1.44 | 0.32 | 0.6 | 0.62 | 1.12 | 2.81E-04 | 2.39E-03 | (A) NE (B/D) |
| LPIN1 | 1.14 | 0.02 | 0.56 | 0.11 | 1.12 | 3.17E-04 | 2.63E-03 | (A) NE (B/D) |
| NUDT9 | 7.13 | 6.01 | 0.41 | 1.49 | 1.12 | 3.69E-04 | 2.96E-03 | (A) NE (B/D) |
| HNRNPA2B1 | 4.98 | 3.86 | 0.55 | 1.3 | 1.12 | 5.11E-04 | 3.80E-03 | (A) NE (B/D) |
| ANAPC2 | 1.18 | 0.06 | 0.62 | 0.17 | 1.12 | 6.00E-04 | 4.31E-03 | (A) NE (B/D) |
| DELE1 | 2.17 | 1.05 | 0.66 | 0.97 | 1.12 | 6.95E-04 | 4.83E-03 | (A) NE (B/D) |
| ZFP37 | 1.27 | 0.15 | 0.73 | 0.32 | 1.12 | 1.63E-03 | 9.20E-03 | (A) NE (B/D) |
| IGBP1 | 4.02 | 2.9 | 0.36 | 1.82 | 1.12 | 1.80E-03 | 9.89E-03 | (A) NE (B/D) |
| RNF19B | 2.41 | 1.29 | 0.73 | 1.1 | 1.12 | 1.81E-03 | 9.93E-03 | (A) NE (B/D) |
| IFI27L2 | 1.47 | 0.35 | 0.96 | 0.75 | 1.12 | 8.00E-03 | 0.0303 | (A) NE (B/D) |
| MIR-4749/3P | 4.25 | 3.13 | 0.93 | 1.16 | 1.12 | 8.09E-03 | 0.0305 | (A) NE (B/D) |
| MIR-4749/5P | 4.25 | 3.13 | 0.93 | 1.16 | 1.12 | 8.09E-03 | 0.0305 | (A) NE (B/D) |
| CEP57L1P1 | 4.54 | 3.41 | 0.94 | 1.29 | 1.12 | 9.55E-03 | 0.0345 | (A) NE (B/D) |
| HOXA-AS2 | 2.17 | 1.05 | 0.99 | 0.85 | 1.12 | 9.73E-03 | 0.035 | (A) NE (B/D) |
| ZBED3 | 1.23 | 0.11 | 1 | 0.3 | 1.12 | 9.80E-03 | 0.0352 | (A) NE (B/D) |
| MCFD2P1 | 1.33 | 0.21 | 1.05 | 0.38 | 1.12 | 0.0123 | 0.0415 | (A) NE (B/D) |
| USP42 | 3.34 | 2.23 | 0.34 | 0.66 | 1.11 | 2.64E-07 | 1.04E-05 | (A) NE (B/D) |
| NGRN | 3.71 | 2.6 | 0.27 | 0.94 | 1.11 | 4.04E-07 | 1.45E-05 | (A) NE (B/D) |
| SETDB1 | 2.87 | 1.76 | 0.28 | 0.98 | 1.11 | 8.23E-07 | 2.56E-05 | (A) NE (B/D) |
| DTNBP1 | 2.5 | 1.39 | 0.34 | 0.85 | 1.11 | 8.33E-07 | 2.58E-05 | (A) NE (B/D) |
| SNORD84 | 3.54 | 2.43 | 0.35 | 0.99 | 1.11 | 4.01E-06 | 8.89E-05 | (A) NE (B/D) |
| APTR | 2.07 | 0.96 | 0.39 | 0.88 | 1.11 | 4.14E-06 | 9.14E-05 | (A) NE (B/D) |
| PRPF4 | 5.33 | 4.21 | 0.37 | 1.06 | 1.11 | 1.03E-05 | 1.86E-04 | (A) NE (B/D) |
| CEP104 | 6.01 | 4.9 | 0.45 | 0.75 | 1.11 | 1.21E-05 | 2.12E-04 | (A) NE (B/D) |
| SH3GL1 | 5.45 | 4.34 | 0.49 | 0.94 | 1.11 | 5.65E-05 | 6.97E-04 | (A) NE (B/D) |
| FAM219B | 1.89 | 0.79 | 0.54 | 0.78 | 1.11 | 9.79E-05 | 1.07E-03 | (A) NE (B/D) |
| COQ8B | 1.67 | 0.56 | 0.56 | 0.63 | 1.11 | 1.65E-04 | 1.59E-03 | (A) NE (B/D) |
| GPAT4 | 7.73 | 6.62 | 0.55 | 0.96 | 1.11 | 1.81E-04 | 1.70E-03 | (A) NE (B/D) |
| GAK | 1.89 | 0.78 | 0.58 | 0.72 | 1.11 | 2.00E-04 | 1.84E-03 | (A) NE (B/D) |
| RCC1 | 6.62 | 5.51 | 0.52 | 1.23 | 1.11 | 2.75E-04 | 2.35E-03 | (A) NE (B/D) |
| OLMALINC | 6.76 | 5.64 | 0.38 | 1.56 | 1.11 | 5.04E-04 | 3.76E-03 | (A) NE (B/D) |
| ARF1 | 4.67 | 3.56 | 0.56 | 1.28 | 1.11 | 5.33E-04 | 3.92E-03 | (A) NE (B/D) |
| SNRPB | 8.38 | 7.27 | 0.57 | 1.26 | 1.11 | 5.97E-04 | 4.29E-03 | (A) NE (B/D) |
| ZNF789 | 2.71 | 1.6 | 0.65 | 1.02 | 1.11 | 7.25E-04 | 4.97E-03 | (A) NE (B/D) |
| SNORD58B | 6.9 | 5.79 | 0.59 | 1.34 | 1.11 | 9.48E-04 | 6.06E-03 | (A) NE (B/D) |
| RPS3 | 2.5 | 1.38 | 0.62 | 1.48 | 1.11 | 1.81E-03 | 9.92E-03 | (A) NE (B/D) |
| KRT18P38 | 2.43 | 1.32 | 0.71 | 1.25 | 1.11 | 1.98E-03 | 0.0106 | (A) NE (B/D) |
| FKBP3 | 4.44 | 3.32 | 0.71 | 1.33 | 1.11 | 2.41E-03 | 0.0124 | (A) NE (B/D) |
| OSER1-DT | 7.23 | 6.11 | 0.77 | 1.11 | 1.11 | 2.77E-03 | 0.0138 | (A) NE (B/D) |
| ROPN1B | 1.64 | 0.53 | 0.8 | 0.73 | 1.11 | 2.82E-03 | 0.0139 | (A) NE (B/D) |
| KRT8P15 | 3.07 | 1.96 | 0.76 | 1.21 | 1.11 | 2.94E-03 | 0.0144 | (A) NE (B/D) |
| ZDHHC8 | 2.11 | 1.01 | 0.8 | 0.93 | 1.11 | 3.01E-03 | 0.0146 | (A) NE (B/D) |
| COX17P1 | 6.34 | 5.23 | 0.71 | 1.46 | 1.11 | 3.16E-03 | 0.0152 | (A) NE (B/D) |
| RPS4XP6 | 5.33 | 4.22 | 0.68 | 1.53 | 1.11 | 3.23E-03 | 0.0154 | (A) NE (B/D) |
| RBBP4P5 | 2.56 | 1.44 | 0.82 | 0.88 | 1.11 | 3.37E-03 | 0.0159 | (A) NE (B/D) |
| LINC00184 | 1.4 | 0.29 | 0.85 | 0.49 | 1.11 | 4.21E-03 | 0.0189 | (A) NE (B/D) |
| HARS2 | 3.33 | 2.22 | 0.83 | 1.17 | 1.11 | 4.76E-03 | 0.0208 | (A) NE (B/D) |
| CTRC | 1.62 | 0.51 | 0.88 | 0.68 | 1.11 | 5.08E-03 | 0.0217 | (A) NE (B/D) |

| | | | | | | | | |
|---|---|---|---|---|---|---|---|---|
| CRYZL1 | 1.36 | 0.25 | 0.89 | 0.5 | 1.11 | 5.47E-03 | 0.0229 | (A) NE (B/D) |
| FAM131A | 1.4 | 0.29 | 0.9 | 0.64 | 1.11 | 5.93E-03 | 0.0244 | (A) NE (B/D) |
| TMEM106C | 2.67 | 1.56 | 0.8 | 1.49 | 1.11 | 6.42E-03 | 0.0259 | (A) NE (B/D) |
| VPS51 | 2.95 | 1.85 | 0.92 | 0.95 | 1.11 | 7.39E-03 | 0.0287 | (A) NE (B/D) |
| ATP6V1C2 | 2.8 | 1.69 | 0.91 | 1.23 | 1.11 | 7.75E-03 | 0.0296 | (A) NE (B/D) |
| USP9YP1 | 1.29 | 0.18 | 1 | 0.46 | 1.11 | 0.0103 | 0.0364 | (A) NE (B/D) |
| DGCR10 | 1.15 | 0.04 | 1.03 | 0.25 | 1.11 | 0.0117 | 0.0399 | (A) NE (B/D) |
| RPL31P15 | 2.16 | 1.05 | 1 | 1.03 | 1.11 | 0.0119 | 0.0405 | (A) NE (B/D) |
| TMEM132D | 1.29 | 0.19 | 1.04 | 0.52 | 1.11 | 0.0128 | 0.0427 | (A) NE (B/D) |
| RN7SL381P | 3.99 | 2.88 | 1.02 | 1.02 | 1.11 | 0.0131 | 0.0435 | (A) NE (B/D) |
| ZNF785 | 2.4 | 1.3 | 0.3 | 0.75 | 1.1 | 1.05E-07 | 5.02E-06 | (A) NE (B/D) |
| DHX34 | 2.47 | 1.37 | 0.32 | 0.65 | 1.1 | 1.09E-07 | 5.17E-06 | (A) NE (B/D) |
| EGLN2 | 5.2 | 4.1 | 0.23 | 0.91 | 1.1 | 1.54E-07 | 6.80E-06 | (A) NE (B/D) |
| TCP11L1 | 1.93 | 0.83 | 0.36 | 0.68 | 1.1 | 6.29E-07 | 2.06E-05 | (A) NE (B/D) |
| DVL3 | 1.58 | 0.48 | 0.38 | 0.68 | 1.1 | 1.39E-06 | 3.88E-05 | (A) NE (B/D) |
| STARD3 | 7.92 | 6.82 | 0.3 | 1.03 | 1.1 | 3.25E-06 | 7.50E-05 | (A) NE (B/D) |
| MIR-5004/3P | 5.94 | 4.84 | 0.38 | 0.91 | 1.1 | 4.44E-06 | 9.62E-05 | (A) NE (B/D) |
| MIR-5004/5P | 5.94 | 4.84 | 0.38 | 0.91 | 1.1 | 4.44E-06 | 9.62E-05 | (A) NE (B/D) |
| NCOA5 | 1.61 | 0.51 | 0.45 | 0.58 | 1.1 | 1.63E-05 | 2.65E-04 | (A) NE (B/D) |
| MCRS1 | 8.36 | 7.26 | 0.36 | 1.2 | 1.1 | 3.98E-05 | 5.27E-04 | (A) NE (B/D) |
| ACTR1A | 2.81 | 1.71 | 0.3 | 1.27 | 1.1 | 4.12E-05 | 5.43E-04 | (A) NE (B/D) |
| CRY1 | 2.98 | 1.88 | 0.44 | 1.12 | 1.1 | 5.73E-05 | 7.04E-04 | (A) NE (B/D) |
| STK19B | 2.26 | 1.15 | 0.44 | 1.12 | 1.1 | 6.40E-05 | 7.66E-04 | (A) NE (B/D) |
| CD3EAP | 2.02 | 0.92 | 0.53 | 0.91 | 1.1 | 1.25E-04 | 1.29E-03 | (A) NE (B/D) |
| PPDPF | 7.01 | 5.92 | 0.5 | 1.1 | 1.1 | 1.34E-04 | 1.36E-03 | (A) NE (B/D) |
| GTF2IRD1P1 | 6.89 | 5.79 | 0.53 | 1.02 | 1.1 | 1.50E-04 | 1.48E-03 | (A) NE (B/D) |
| CARHSP1 | 2.28 | 1.18 | 0.58 | 0.88 | 1.1 | 2.70E-04 | 2.31E-03 | (A) NE (B/D) |
| PUSL1 | 4.09 | 2.99 | 0.6 | 0.62 | 1.1 | 3.36E-04 | 2.75E-03 | (A) NE (B/D) |
| TOR1A | 2.51 | 1.41 | 0.54 | 1.31 | 1.1 | 5.48E-04 | 4.01E-03 | (A) NE (B/D) |
| DCAF5 | 2.7 | 1.6 | 0.66 | 1.18 | 1.1 | 1.22E-03 | 7.35E-03 | (A) NE (B/D) |
| SMIM13 | 2.98 | 1.88 | 0.71 | 0.71 | 1.1 | 1.25E-03 | 7.49E-03 | (A) NE (B/D) |
| RPL29P3 | 1.38 | 0.28 | 0.72 | 0.69 | 1.1 | 1.40E-03 | 8.17E-03 | (A) NE (B/D) |
| BLMH | 1.42 | 0.32 | 0.72 | 0.42 | 1.1 | 1.49E-03 | 8.58E-03 | (A) NE (B/D) |
| MIR-4426/44 | 7.89 | 6.79 | 0.36 | 1.76 | 1.1 | 1.66E-03 | 9.29E-03 | (A) NE (B/D) |
| TRIM52 | 6.01 | 4.92 | 0.73 | 0.87 | 1.1 | 1.75E-03 | 9.68E-03 | (A) NE (B/D) |
| ACADVL | 5.5 | 4.4 | 0.76 | 1.12 | 1.1 | 2.86E-03 | 0.0141 | (A) NE (B/D) |
| CRYBB2 | 2.87 | 1.77 | 0.75 | 1.32 | 1.1 | 3.51E-03 | 0.0164 | (A) NE (B/D) |
| SNORD38B | 1.11 | 0 | 0.86 | 0.02 | 1.1 | 5.04E-03 | 0.0216 | (A) NE (B/D) |
| CENPJ | 6.33 | 5.23 | 0.87 | 0.93 | 1.1 | 5.12E-03 | 0.0219 | (A) NE (B/D) |
| PSAT1 | 8.57 | 7.47 | 0.65 | 1.74 | 1.1 | 5.19E-03 | 0.0221 | (A) NE (B/D) |
| ZNF582-AS1 | 1.34 | 0.24 | 0.89 | 0.49 | 1.1 | 5.53E-03 | 0.0231 | (A) NE (B/D) |
| THAP9-AS1 | 4.48 | 3.38 | 0.88 | 0.9 | 1.1 | 5.65E-03 | 0.0235 | (A) NE (B/D) |
| MYLK3 | 1.63 | 0.53 | 0.96 | 0.52 | 1.1 | 8.47E-03 | 0.0316 | (A) NE (B/D) |
| SNRPN | 5.74 | 4.64 | 0.88 | 1.41 | 1.1 | 8.65E-03 | 0.0321 | (A) NE (B/D) |
| CMIP | 6.5 | 5.4 | 1 | 0.59 | 1.1 | 0.0106 | 0.0372 | (A) NE (B/D) |
| LINC00881 | 1.42 | 0.32 | 1 | 0.48 | 1.1 | 0.0107 | 0.0374 | (A) NE (B/D) |
| IFT122 | 1.46 | 0.37 | 0.26 | 0.4 | 1.09 | 3.47E-09 | 3.55E-07 | (A) NE (B/D) |
| RRP1 | 3.8 | 2.7 | 0.25 | 0.75 | 1.09 | 1.79E-08 | 1.23E-06 | (A) NE (B/D) |
| NRF1 | 1.95 | 0.86 | 0.25 | 0.81 | 1.09 | 4.89E-08 | 2.73E-06 | (A) NE (B/D) |
| IL10RB | 3.39 | 2.3 | 0.29 | 0.82 | 1.09 | 1.59E-07 | 6.97E-06 | (A) NE (B/D) |
| PWP2 | 1.6 | 0.51 | 0.34 | 0.56 | 1.09 | 2.97E-07 | 1.14E-05 | (A) NE (B/D) |
| ERN1 | 7.4 | 6.31 | 0.38 | 0.67 | 1.09 | 1.86E-06 | 4.88E-05 | (A) NE (B/D) |
| ICMT | 7.94 | 6.86 | 0.26 | 1.05 | 1.09 | 2.51E-06 | 6.12E-05 | (A) NE (B/D) |
| ASB16-AS1 | 6.43 | 5.35 | 0.3 | 1.09 | 1.09 | 7.63E-06 | 1.47E-04 | (A) NE (B/D) |
| AKAP8 | 1.31 | 0.21 | 0.4 | 0.41 | 1.09 | 8.22E-06 | 1.56E-04 | (A) NE (B/D) |
| MSTO2P | 4.27 | 3.18 | 0.3 | 1.11 | 1.09 | 9.12E-06 | 1.69E-04 | (A) NE (B/D) |
| SEMA4F | 2.37 | 1.27 | 0.44 | 0.77 | 1.09 | 1.27E-05 | 2.21E-04 | (A) NE (B/D) |
| GNAS | 6.13 | 5.03 | 0.23 | 1.23 | 1.09 | 1.83E-05 | 2.88E-04 | (A) NE (B/D) |
| PRXL2C | 5.74 | 4.65 | 0.44 | 1 | 1.09 | 3.68E-05 | 4.97E-04 | (A) NE (B/D) |
| SLC66A1 | 5.46 | 4.37 | 0.43 | 1.08 | 1.09 | 4.58E-05 | 5.90E-04 | (A) NE (B/D) |
| CNPY3 | 8.62 | 7.53 | 0.34 | 1.24 | 1.09 | 5.24E-05 | 6.57E-04 | (A) NE (B/D) |
| H3F3B | 3.22 | 2.13 | 0.34 | 1.26 | 1.09 | 6.35E-05 | 7.62E-04 | (A) NE (B/D) |
| TANC2 | 7.58 | 6.49 | 0.5 | 0.9 | 1.09 | 6.62E-05 | 7.87E-04 | (A) NE (B/D) |
| PRPF8 | 5.38 | 4.29 | 0.52 | 0.79 | 1.09 | 9.15E-05 | 1.02E-03 | (A) NE (B/D) |
| DOLPP1 | 2.3 | 1.21 | 0.45 | 1.15 | 1.09 | 9.96E-05 | 1.08E-03 | (A) NE (B/D) |

| | | | | | | | | |
|---|---|---|---|---|---|---|---|---|
| ZNF32 | 5.87 | 4.78 | 0.31 | 1.39 | 1.09 | 1.42E-04 | 1.41E-03 | (A) NE (B/D) |
| NMB | 1.28 | 0.19 | 0.54 | 0.38 | 1.09 | 1.95E-04 | 1.81E-03 | (A) NE (B/D) |
| ZNF579 | 1.17 | 0.08 | 0.57 | 0.2 | 1.09 | 3.55E-04 | 2.87E-03 | (A) NE (B/D) |
| PTBP1P | 4.82 | 3.74 | 0.61 | 0.97 | 1.09 | 5.18E-04 | 3.84E-03 | (A) NE (B/D) |
| PIAS3 | 3.79 | 2.69 | 0.58 | 1.25 | 1.09 | 6.68E-04 | 4.67E-03 | (A) NE (B/D) |
| SPAG7 | 2.57 | 1.47 | 0.51 | 1.4 | 1.09 | 6.98E-04 | 4.84E-03 | (A) NE (B/D) |
| SMIM12 | 6.99 | 5.89 | 0.66 | 0.82 | 1.09 | 8.00E-04 | 5.38E-03 | (A) NE (B/D) |
| PSMG1 | 2.12 | 1.03 | 0.64 | 1.06 | 1.09 | 8.30E-04 | 5.51E-03 | (A) NE (B/D) |
| PARP1 | 2.26 | 1.17 | 0.68 | 1.11 | 1.09 | 1.49E-03 | 8.58E-03 | (A) NE (B/D) |
| NAPG | 1.23 | 0.14 | 0.72 | 0.33 | 1.09 | 1.70E-03 | 9.46E-03 | (A) NE (B/D) |
| B4GAT1 | 7.7 | 6.6 | 0.61 | 1.43 | 1.09 | 1.75E-03 | 9.67E-03 | (A) NE (B/D) |
| SH3GLB2 | 7.77 | 6.67 | 0.6 | 1.48 | 1.09 | 1.86E-03 | 0.0101 | (A) NE (B/D) |
| PPP1R13B | 4.01 | 2.93 | 0.71 | 1.26 | 1.09 | 2.48E-03 | 0.0127 | (A) NE (B/D) |
| RPS7 | 3.28 | 2.19 | 0.45 | 1.77 | 1.09 | 2.50E-03 | 0.0127 | (A) NE (B/D) |
| ANKRD46 | 6.17 | 5.08 | 0.79 | 0.93 | 1.09 | 3.07E-03 | 0.0149 | (A) NE (B/D) |
| SLC39A3 | 1.22 | 0.13 | 0.84 | 0.34 | 1.09 | 4.43E-03 | 0.0196 | (A) NE (B/D) |
| STIM2 | 1.72 | 0.63 | 0.85 | 0.83 | 1.09 | 4.68E-03 | 0.0205 | (A) NE (B/D) |
| SOWAHB | 3.92 | 2.83 | 0.8 | 1.46 | 1.09 | 6.45E-03 | 0.0259 | (A) NE (B/D) |
| PCDHB13 | 1.1 | 0.01 | 0.92 | 0.05 | 1.09 | 7.55E-03 | 0.0291 | (A) NE (B/D) |
| TRIM59 | 1.54 | 0.46 | 0.96 | 0.73 | 1.09 | 9.44E-03 | 0.0342 | (A) NE (B/D) |
| DEPP1 | 1.4 | 0.31 | 1.01 | 0.5 | 1.09 | 0.0118 | 0.0401 | (A) NE (B/D) |
| MIS18A-AS1 | 4.19 | 3.1 | 0.97 | 1.15 | 1.09 | 0.0118 | 0.0403 | (A) NE (B/D) |
| HNRNPLP1 | 3.45 | 2.36 | 0.95 | 1.28 | 1.09 | 0.0121 | 0.041 | (A) NE (B/D) |
| AP5S1 | 3.29 | 2.21 | 0.2 | 0.63 | 1.08 | 1.57E-10 | 3.48E-08 | (A) NE (B/D) |
| MAML1 | 5.78 | 4.7 | 0.26 | 0.54 | 1.08 | 4.11E-09 | 4.00E-07 | (A) NE (B/D) |
| BRD4 | 7.15 | 6.07 | 0.26 | 0.36 | 1.08 | 1.25E-08 | 9.36E-07 | (A) NE (B/D) |
| MED24 | 1.62 | 0.54 | 0.3 | 0.62 | 1.08 | 6.43E-08 | 3.36E-06 | (A) NE (B/D) |
| SENP5 | 5.61 | 4.53 | 0.33 | 0.45 | 1.08 | 3.31E-07 | 1.25E-05 | (A) NE (B/D) |
| INIP | 4.86 | 3.78 | 0.27 | 0.95 | 1.08 | 8.88E-07 | 2.70E-05 | (A) NE (B/D) |
| KLHL26 | 1.32 | 0.24 | 0.35 | 0.38 | 1.08 | 1.78E-06 | 4.70E-05 | (A) NE (B/D) |
| PPP4R1 | 1.76 | 0.69 | 0.38 | 0.64 | 1.08 | 1.83E-06 | 4.82E-05 | (A) NE (B/D) |
| FBLIM1 | 2.68 | 1.61 | 0.27 | 1.05 | 1.08 | 3.51E-06 | 8.01E-05 | (A) NE (B/D) |
| NUDC | 2.01 | 0.94 | 0.33 | 0.99 | 1.08 | 4.27E-06 | 9.37E-05 | (A) NE (B/D) |
| YEATS2-AS1 | 3.52 | 2.45 | 0.41 | 0.66 | 1.08 | 5.97E-06 | 1.21E-04 | (A) NE (B/D) |
| NDUFAF8 | 5.15 | 4.07 | 0.28 | 1.11 | 1.08 | 8.82E-06 | 1.65E-04 | (A) NE (B/D) |
| CHERP | 1.09 | 0.01 | 0.34 | 0.04 | 1.08 | 1.27E-05 | 2.20E-04 | (A) NE (B/D) |
| SNRNP40 | 3.92 | 2.84 | 0.41 | 1.08 | 1.08 | 3.89E-05 | 5.18E-04 | (A) NE (B/D) |
| HYPK | 7.69 | 6.61 | 0.39 | 1.15 | 1.08 | 5.23E-05 | 6.55E-04 | (A) NE (B/D) |
| NBR2 | 5.18 | 4.1 | 0.49 | 0.79 | 1.08 | 5.92E-05 | 7.22E-04 | (A) NE (B/D) |
| CARD19 | 3.13 | 2.05 | 0.47 | 1.05 | 1.08 | 8.21E-05 | 9.30E-04 | (A) NE (B/D) |
| ZNF32-AS2 | 1.58 | 0.5 | 0.52 | 0.73 | 1.08 | 8.76E-05 | 9.82E-04 | (A) NE (B/D) |
| PQBP1 | 2.16 | 1.09 | 0.44 | 1.16 | 1.08 | 1.07E-04 | 1.15E-03 | (A) NE (B/D) |
| TECPR1 | 5.06 | 3.99 | 0.52 | 0.79 | 1.08 | 1.08E-04 | 1.15E-03 | (A) NE (B/D) |
| FIS1 | 5.52 | 4.44 | 0.42 | 1.22 | 1.08 | 1.20E-04 | 1.24E-03 | (A) NE (B/D) |
| DHX9P1 | 4.15 | 3.07 | 0.53 | 0.84 | 1.08 | 1.22E-04 | 1.26E-03 | (A) NE (B/D) |
| FLOT2 | 7.87 | 6.79 | 0.44 | 1.25 | 1.08 | 1.74E-04 | 1.65E-03 | (A) NE (B/D) |
| POGK | 7.77 | 6.69 | 0.55 | 0.89 | 1.08 | 1.87E-04 | 1.75E-03 | (A) NE (B/D) |
| SMC1A | 2.94 | 1.86 | 0.52 | 1.09 | 1.08 | 2.39E-04 | 2.11E-03 | (A) NE (B/D) |
| TBC1D10B | 5.68 | 4.61 | 0.57 | 0.82 | 1.08 | 2.55E-04 | 2.21E-03 | (A) NE (B/D) |
| PGP | 4.38 | 3.3 | 0.58 | 1.03 | 1.08 | 4.19E-04 | 3.27E-03 | (A) NE (B/D) |
| C16ORF86 | 4.11 | 3.03 | 0.6 | 0.94 | 1.08 | 4.42E-04 | 3.41E-03 | (A) NE (B/D) |
| SRGAP1 | 2.24 | 1.17 | 0.66 | 0.72 | 1.08 | 8.36E-04 | 5.54E-03 | (A) NE (B/D) |
| IRF2BP2 | 6.37 | 5.29 | 0.66 | 0.61 | 1.08 | 8.50E-04 | 5.61E-03 | (A) NE (B/D) |
| ZBTB14 | 6.23 | 5.14 | 0.65 | 1.09 | 1.08 | 1.08E-03 | 6.71E-03 | (A) NE (B/D) |
| HSPA1L | 4.83 | 3.75 | 0.73 | 0.93 | 1.08 | 1.96E-03 | 0.0106 | (A) NE (B/D) |
| CCND3 | 5.01 | 3.92 | 0.68 | 1.25 | 1.08 | 2.07E-03 | 0.011 | (A) NE (B/D) |
| TGDS | 1.3 | 0.23 | 0.74 | 0.37 | 1.08 | 2.28E-03 | 0.0118 | (A) NE (B/D) |
| SLC35E2B | 1.35 | 0.27 | 0.78 | 0.43 | 1.08 | 3.00E-03 | 0.0146 | (A) NE (B/D) |
| DDAH2 | 4.01 | 2.93 | 0.73 | 1.36 | 1.08 | 3.63E-03 | 0.0168 | (A) NE (B/D) |
| RNU6-610P | 1.28 | 0.2 | 0.84 | 0.43 | 1.08 | 4.59E-03 | 0.0202 | (A) NE (B/D) |
| ZNF20 | 3.68 | 2.6 | 0.87 | 0.74 | 1.08 | 5.84E-03 | 0.0241 | (A) NE (B/D) |
| THA1P | 2.32 | 1.24 | 0.9 | 0.73 | 1.08 | 6.80E-03 | 0.027 | (A) NE (B/D) |
| PRELID3BP1 | 1.19 | 0.11 | 0.9 | 0.39 | 1.08 | 6.89E-03 | 0.0273 | (A) NE (B/D) |
| CYCSP55 | 1.87 | 0.79 | 0.91 | 0.66 | 1.08 | 7.23E-03 | 0.0282 | (A) NE (B/D) |
| PNPT1 | 1.51 | 0.43 | 0.91 | 0.56 | 1.08 | 7.25E-03 | 0.0283 | (A) NE (B/D) |

| | | | | | | | | |
|---|---|---|---|---|---|---|---|---|
| ZNF235 | 4.51 | 3.43 | 0.97 | 0.67 | 1.08 | 0.0104 | 0.0367 | (A) NE (B/D) |
| PPFIA4 | 1.28 | 0.2 | 1 | 0.52 | 1.08 | 0.0122 | 0.0413 | (A) NE (B/D) |
| SPIN4-AS1 | 1.68 | 0.6 | 1 | 0.88 | 1.08 | 0.0126 | 0.0421 | (A) NE (B/D) |
| MICOS10-NB | 4.06 | 2.99 | 0.91 | 1.5 | 1.08 | 0.0131 | 0.0434 | (A) NE (B/D) |
| MCM10 | 3.7 | 2.62 | 0.92 | 1.53 | 1.08 | 0.0142 | 0.0461 | (A) NE (B/D) |
| DAZAP1 | 8.49 | 7.41 | 0.1 | 0.8 | 1.07 | 5.23E-09 | 4.77E-07 | (A) NE (B/D) |
| ZKSCAN4 | 4.81 | 3.74 | 0.18 | 0.82 | 1.07 | 1.49E-08 | 1.06E-06 | (A) NE (B/D) |
| DHRSX | 4.22 | 3.16 | 0.27 | 0.68 | 1.07 | 2.49E-08 | 1.56E-06 | (A) NE (B/D) |
| PEX14 | 3.6 | 2.53 | 0.26 | 0.84 | 1.07 | 1.38E-07 | 6.28E-06 | (A) NE (B/D) |
| NUDT22 | 4.19 | 3.12 | 0.29 | 0.94 | 1.07 | 1.33E-06 | 3.74E-05 | (A) NE (B/D) |
| E2F3 | 2.47 | 1.4 | 0.28 | 1 | 1.07 | 2.26E-06 | 5.64E-05 | (A) NE (B/D) |
| PARP16 | 2.02 | 0.96 | 0.36 | 0.84 | 1.07 | 3.07E-06 | 7.15E-05 | (A) NE (B/D) |
| JOSD2 | 4.93 | 3.86 | 0.34 | 0.96 | 1.07 | 4.63E-06 | 9.93E-05 | (A) NE (B/D) |
| DYNC1I2P1 | 7.21 | 6.14 | 0.33 | 0.99 | 1.07 | 5.33E-06 | 1.11E-04 | (A) NE (B/D) |
| FAM122C | 2.89 | 1.82 | 0.43 | 0.68 | 1.07 | 9.86E-06 | 1.80E-04 | (A) NE (B/D) |
| ZFP69 | 4.47 | 3.39 | 0.4 | 0.96 | 1.07 | 1.48E-05 | 2.48E-04 | (A) NE (B/D) |
| SDF4 | 4.57 | 3.5 | 0.42 | 0.9 | 1.07 | 1.65E-05 | 2.67E-04 | (A) NE (B/D) |
| PPP6R2 | 3.5 | 2.43 | 0.35 | 1.08 | 1.07 | 1.72E-05 | 2.77E-04 | (A) NE (B/D) |
| WWOX | 5.28 | 4.21 | 0.43 | 0.85 | 1.07 | 1.90E-05 | 2.97E-04 | (A) NE (B/D) |
| SMIM7 | 5.51 | 4.43 | 0.4 | 1.02 | 1.07 | 2.45E-05 | 3.59E-04 | (A) NE (B/D) |
| NDOR1 | 1.19 | 0.12 | 0.41 | 0.25 | 1.07 | 2.53E-05 | 3.68E-04 | (A) NE (B/D) |
| B3GALT4 | 2.18 | 1.12 | 0.46 | 0.76 | 1.07 | 2.64E-05 | 3.80E-04 | (A) NE (B/D) |
| MAPRE1 | 9.15 | 8.08 | 0.44 | 0.89 | 1.07 | 2.88E-05 | 4.08E-04 | (A) NE (B/D) |
| TBC1D3B | 4.57 | 3.51 | 0.27 | 1.23 | 1.07 | 3.42E-05 | 4.69E-04 | (A) NE (B/D) |
| SLC25A11 | 2.41 | 1.34 | 0.41 | 1.21 | 1.07 | 1.10E-04 | 1.17E-03 | (A) NE (B/D) |
| PPP1R12C | 3.18 | 2.11 | 0.54 | 0.69 | 1.07 | 1.57E-04 | 1.53E-03 | (A) NE (B/D) |
| SNORD121B | 3.25 | 2.18 | 0.52 | 1.04 | 1.07 | 2.01E-04 | 1.85E-03 | (A) NE (B/D) |
| TMEM9 | 8.13 | 7.06 | 0.31 | 1.41 | 1.07 | 2.23E-04 | 2.00E-03 | (A) NE (B/D) |
| KDM5D | 1.44 | 0.37 | 0.58 | 0.61 | 1.07 | 3.09E-04 | 2.58E-03 | (A) NE (B/D) |
| NARF-IT1 | 2.36 | 1.29 | 0.61 | 0.86 | 1.07 | 4.89E-04 | 3.68E-03 | (A) NE (B/D) |
| C20ORF204 | 2.12 | 1.05 | 0.65 | 0.67 | 1.07 | 7.73E-04 | 5.22E-03 | (A) NE (B/D) |
| NDUFAF5 | 4.77 | 3.7 | 0.67 | 0.84 | 1.07 | 1.01E-03 | 6.38E-03 | (A) NE (B/D) |
| RPS11 | 13.86 | 12.78 | 0.48 | 1.53 | 1.07 | 1.18E-03 | 7.19E-03 | (A) NE (B/D) |
| SNORD119 | 1.86 | 0.79 | 0.68 | 0.72 | 1.07 | 1.19E-03 | 7.22E-03 | (A) NE (B/D) |
| TIMM9 | 2.37 | 1.29 | 0.62 | 1.23 | 1.07 | 1.19E-03 | 7.24E-03 | (A) NE (B/D) |
| CAD | 2.06 | 0.99 | 0.7 | 0.8 | 1.07 | 1.38E-03 | 8.09E-03 | (A) NE (B/D) |
| WDR45 | 2.74 | 1.67 | 0.58 | 1.41 | 1.07 | 1.47E-03 | 8.47E-03 | (A) NE (B/D) |
| CDC26 | 6.14 | 5.07 | 0.65 | 1.35 | 1.07 | 2.12E-03 | 0.0112 | (A) NE (B/D) |
| CORO7 | 2.06 | 0.99 | 0.75 | 0.99 | 1.07 | 2.62E-03 | 0.0132 | (A) NE (B/D) |
| NIPSNAP2 | 7.74 | 6.66 | 0.74 | 1.21 | 1.07 | 3.22E-03 | 0.0154 | (A) NE (B/D) |
| CYP2D8P | 4.12 | 3.05 | 0.85 | 0.86 | 1.07 | 5.33E-03 | 0.0225 | (A) NE (B/D) |
| APOO | 4.26 | 3.19 | 0.77 | 1.65 | 1.07 | 9.01E-03 | 0.0331 | (A) NE (B/D) |
| EEF1A1P6 | 7.94 | 6.88 | 0.83 | 1.44 | 1.07 | 9.02E-03 | 0.0331 | (A) NE (B/D) |
| GPRIN3 | 1.38 | 0.31 | 1.04 | 0.52 | 1.07 | 0.0152 | 0.0485 | (A) NE (B/D) |
| REXO1 | 1.23 | 0.17 | 0.23 | 0.3 | 1.06 | 4.77E-09 | 4.50E-07 | (A) NE (B/D) |
| PRRC2A | 6.86 | 5.79 | 0.23 | 0.84 | 1.06 | 8.83E-08 | 4.34E-06 | (A) NE (B/D) |
| PABPC4 | 6.09 | 5.03 | 0.41 | 0.86 | 1.06 | 1.31E-05 | 2.26E-04 | (A) NE (B/D) |
| PCNT | 1.68 | 0.62 | 0.41 | 0.41 | 1.06 | 1.46E-05 | 2.46E-04 | (A) NE (B/D) |
| RALA | 6.41 | 5.35 | 0.32 | 1.12 | 1.06 | 1.98E-05 | 3.06E-04 | (A) NE (B/D) |
| RBM4B | 2.49 | 1.42 | 0.35 | 1.09 | 1.06 | 2.18E-05 | 3.28E-04 | (A) NE (B/D) |
| GOLGA2 | 8.98 | 7.93 | 0.42 | 0.97 | 1.06 | 3.33E-05 | 4.58E-04 | (A) NE (B/D) |
| UBL4A | 2.16 | 1.09 | 0.34 | 1.18 | 1.06 | 4.28E-05 | 5.61E-04 | (A) NE (B/D) |
| ZNF865 | 3.34 | 2.28 | 0.48 | 0.54 | 1.06 | 5.05E-05 | 6.37E-04 | (A) NE (B/D) |
| RALY | 5.67 | 4.6 | 0.42 | 1.1 | 1.06 | 6.37E-05 | 7.65E-04 | (A) NE (B/D) |
| AKT2 | 1.43 | 0.37 | 0.47 | 0.35 | 1.06 | 7.75E-05 | 8.89E-04 | (A) NE (B/D) |
| AAMP | 3.93 | 2.87 | 0.2 | 1.35 | 1.06 | 8.69E-05 | 9.76E-04 | (A) NE (B/D) |
| LENG9 | 3.24 | 2.18 | 0.54 | 0.86 | 1.06 | 2.05E-04 | 1.87E-03 | (A) NE (B/D) |
| INTS3 | 3.11 | 2.05 | 0.55 | 0.88 | 1.06 | 2.13E-04 | 1.93E-03 | (A) NE (B/D) |
| UTP11 | 6.15 | 5.09 | 0.54 | 1 | 1.06 | 2.57E-04 | 2.22E-03 | (A) NE (B/D) |
| KCNH6 | 1.08 | 0.02 | 0.57 | 0.05 | 1.06 | 5.24E-04 | 3.87E-03 | (A) NE (B/D) |
| SLX1B-SULT1 | 9.39 | 8.33 | 0.59 | 1.06 | 1.06 | 5.89E-04 | 4.25E-03 | (A) NE (B/D) |
| TMEM203 | 2.48 | 1.42 | 0.52 | 1.28 | 1.06 | 6.15E-04 | 4.38E-03 | (A) NE (B/D) |
| C1ORF52 | 1.84 | 0.78 | 0.63 | 0.84 | 1.06 | 7.11E-04 | 4.91E-03 | (A) NE (B/D) |
| GNPAT | 5.88 | 4.82 | 0.62 | 1.02 | 1.06 | 8.11E-04 | 5.42E-03 | (A) NE (B/D) |
| RPA2 | 2.52 | 1.47 | 0.64 | 1.22 | 1.06 | 1.64E-03 | 9.21E-03 | (A) NE (B/D) |

| | | | | | | | | |
|---|---|---|---|---|---|---|---|---|
| ZNF625-ZNF | 3.86 | 2.81 | 0.71 | 0.75 | 1.06 | 1.66E-03 | 9.29E-03 | (A) NE (B/D) |
| RNA5SP323 | 1.38 | 0.32 | 0.76 | 0.62 | 1.06 | 2.68E-03 | 0.0134 | (A) NE (B/D) |
| NCBP2AS2 | 5.84 | 4.78 | 0.76 | 1.15 | 1.06 | 3.65E-03 | 0.0169 | (A) NE (B/D) |
| SIK3 | 4.24 | 3.18 | 0.82 | 0.66 | 1.06 | 4.20E-03 | 0.0189 | (A) NE (B/D) |
| MTCP1 | 7.52 | 6.46 | 0.79 | 1.3 | 1.06 | 5.83E-03 | 0.0241 | (A) NE (B/D) |
| ADAMTS13 | 3.15 | 2.09 | 0.82 | 1.22 | 1.06 | 6.17E-03 | 0.0251 | (A) NE (B/D) |
| PLPP5 | 7.95 | 6.89 | 0.84 | 1.17 | 1.06 | 6.61E-03 | 0.0264 | (A) NE (B/D) |
| GRIK1-AS1 | 1.22 | 0.17 | 0.89 | 0.35 | 1.06 | 7.07E-03 | 0.0278 | (A) NE (B/D) |
| DOK6 | 1.19 | 0.13 | 0.89 | 0.25 | 1.06 | 7.23E-03 | 0.0282 | (A) NE (B/D) |
| SLC12A4 | 1.64 | 0.58 | 0.9 | 0.69 | 1.06 | 7.52E-03 | 0.029 | (A) NE (B/D) |
| ECT2 | 1.13 | 0.07 | 0.92 | 0.33 | 1.06 | 8.74E-03 | 0.0323 | (A) NE (B/D) |
| CAMK2B | 1.07 | 0.01 | 0.99 | 0.05 | 1.06 | 0.0123 | 0.0415 | (A) NE (B/D) |
| PTTG1IP | 6.84 | 5.79 | 0.19 | 0.9 | 1.05 | 1.76E-07 | 7.60E-06 | (A) NE (B/D) |
| KDM4A | 4.91 | 3.86 | 0.33 | 0.73 | 1.05 | 5.53E-07 | 1.87E-05 | (A) NE (B/D) |
| ACAD9 | 2.23 | 1.18 | 0.28 | 0.94 | 1.05 | 1.40E-06 | 3.88E-05 | (A) NE (B/D) |
| IL10RB-DT | 7.41 | 6.36 | 0.27 | 0.98 | 1.05 | 1.79E-06 | 4.73E-05 | (A) NE (B/D) |
| TRMU | 2.57 | 1.52 | 0.27 | 1.03 | 1.05 | 4.32E-06 | 9.46E-05 | (A) NE (B/D) |
| KANSL3 | 1.56 | 0.51 | 0.4 | 0.49 | 1.05 | 6.79E-06 | 1.34E-04 | (A) NE (B/D) |
| DDX27 | 7.15 | 6.1 | 0.36 | 0.93 | 1.05 | 7.99E-06 | 1.53E-04 | (A) NE (B/D) |
| HMGN4 | 6.47 | 5.42 | 0.35 | 1.04 | 1.05 | 1.78E-05 | 2.82E-04 | (A) NE (B/D) |
| SUMF2 | 5.4 | 4.35 | 0.36 | 1.04 | 1.05 | 2.16E-05 | 3.26E-04 | (A) NE (B/D) |
| RABL2B | 5.84 | 4.79 | 0.35 | 1.09 | 1.05 | 2.85E-05 | 4.05E-04 | (A) NE (B/D) |
| AUP1 | 8.06 | 7.01 | 0.37 | 1.06 | 1.05 | 2.90E-05 | 4.10E-04 | (A) NE (B/D) |
| ZBTB48 | 2.01 | 0.96 | 0.45 | 0.77 | 1.05 | 3.00E-05 | 4.23E-04 | (A) NE (B/D) |
| EED | 3.85 | 2.8 | 0.46 | 0.82 | 1.05 | 4.41E-05 | 5.73E-04 | (A) NE (B/D) |
| TCTN2 | 6.46 | 5.41 | 0.39 | 1.11 | 1.05 | 5.50E-05 | 6.81E-04 | (A) NE (B/D) |
| LYPLA2 | 7.08 | 6.03 | 0.31 | 1.3 | 1.05 | 1.07E-04 | 1.15E-03 | (A) NE (B/D) |
| MIR-2277/3F | 2.5 | 1.45 | 0.58 | 0.63 | 1.05 | 3.25E-04 | 2.68E-03 | (A) NE (B/D) |
| MIR-2277/5F | 2.5 | 1.45 | 0.58 | 0.63 | 1.05 | 3.25E-04 | 2.68E-03 | (A) NE (B/D) |
| LMAN2 | 2.77 | 1.72 | 0.64 | 0.8 | 1.05 | 8.69E-04 | 5.69E-03 | (A) NE (B/D) |
| TMEM250 | 4.63 | 3.58 | 0.64 | 0.99 | 1.05 | 9.81E-04 | 6.22E-03 | (A) NE (B/D) |
| PRXL2B | 7.48 | 6.43 | 0.64 | 1.06 | 1.05 | 1.15E-03 | 7.04E-03 | (A) NE (B/D) |
| ARL16 | 6.09 | 5.04 | 0.53 | 1.4 | 1.05 | 1.16E-03 | 7.08E-03 | (A) NE (B/D) |
| C15ORF65 | 3.11 | 2.05 | 0.64 | 1.13 | 1.05 | 1.31E-03 | 7.78E-03 | (A) NE (B/D) |
| RPLP1 | 3.81 | 2.77 | 0.53 | 1.45 | 1.05 | 1.55E-03 | 8.84E-03 | (A) NE (B/D) |
| ATF5 | 5.6 | 4.55 | 0.6 | 1.38 | 1.05 | 1.96E-03 | 0.0106 | (A) NE (B/D) |
| HSBP1 | 8.89 | 7.84 | 0.7 | 1.14 | 1.05 | 2.33E-03 | 0.012 | (A) NE (B/D) |
| LYRM2 | 2.98 | 1.93 | 0.72 | 1 | 1.05 | 2.37E-03 | 0.0122 | (A) NE (B/D) |
| RPL23P2 | 1.36 | 0.31 | 0.75 | 0.51 | 1.05 | 2.65E-03 | 0.0133 | (A) NE (B/D) |
| TMEM126B | 6.86 | 5.82 | 0.67 | 1.32 | 1.05 | 2.94E-03 | 0.0144 | (A) NE (B/D) |
| CREBZF | 1.5 | 0.45 | 0.77 | 0.55 | 1.05 | 3.11E-03 | 0.015 | (A) NE (B/D) |
| PRIMPOL | 2.84 | 1.79 | 0.79 | 0.82 | 1.05 | 3.72E-03 | 0.0172 | (A) NE (B/D) |
| MPHOSPH9 | 1.76 | 0.71 | 0.79 | 0.95 | 1.05 | 4.11E-03 | 0.0186 | (A) NE (B/D) |
| RHEB | 1.67 | 0.62 | 0.86 | 0.71 | 1.05 | 6.40E-03 | 0.0258 | (A) NE (B/D) |
| RPF1 | 3.66 | 2.61 | 0.8 | 1.29 | 1.05 | 6.79E-03 | 0.027 | (A) NE (B/D) |
| SHC2 | 1.41 | 0.36 | 0.88 | 0.64 | 1.05 | 7.08E-03 | 0.0278 | (A) NE (B/D) |
| CEP97 | 2.82 | 1.77 | 0.89 | 0.65 | 1.05 | 7.56E-03 | 0.0291 | (A) NE (B/D) |
| COX19 | 4.39 | 3.34 | 0.95 | 0.72 | 1.05 | 0.0108 | 0.0376 | (A) NE (B/D) |
| RPL35 | 10.23 | 9.18 | 0.85 | 1.54 | 1.05 | 0.0129 | 0.0429 | (A) NE (B/D) |
| GALNT10 | 1.49 | 0.44 | 1.01 | 0.58 | 1.05 | 0.0149 | 0.0477 | (A) NE (B/D) |
| APBA3 | 1.62 | 0.58 | 0.2 | 0.31 | 1.04 | 1.76E-10 | 3.84E-08 | (A) NE (B/D) |
| MAPK11 | 1.47 | 0.43 | 0.22 | 0.44 | 1.04 | 2.37E-10 | 4.65E-08 | (A) NE (B/D) |
| CIAO1 | 2.22 | 1.18 | 0.15 | 0.72 | 1.04 | 1.13E-09 | 1.48E-07 | (A) NE (B/D) |
| CEP250 | 2.81 | 1.77 | 0.14 | 0.86 | 1.04 | 5.35E-08 | 2.92E-06 | (A) NE (B/D) |
| TCTEX1D2 | 1.86 | 0.82 | 0.2 | 0.97 | 1.04 | 9.16E-07 | 2.77E-05 | (A) NE (B/D) |
| SRGAP2 | 2.99 | 1.95 | 0.35 | 0.75 | 1.04 | 1.95E-06 | 5.04E-05 | (A) NE (B/D) |
| RNF114 | 2.58 | 1.54 | 0.2 | 1.1 | 1.04 | 5.93E-06 | 1.21E-04 | (A) NE (B/D) |
| PGAP2 | 3.64 | 2.6 | 0.34 | 1.07 | 1.04 | 2.39E-05 | 3.52E-04 | (A) NE (B/D) |
| BSG | 4.83 | 3.79 | 0.32 | 1.13 | 1.04 | 3.26E-05 | 4.54E-04 | (A) NE (B/D) |
| GOLGA1 | 5.01 | 3.97 | 0.46 | 0.77 | 1.04 | 3.89E-05 | 5.19E-04 | (A) NE (B/D) |
| KCTD6 | 2.49 | 1.46 | 0.47 | 0.81 | 1.04 | 6.38E-05 | 7.65E-04 | (A) NE (B/D) |
| HARS | 6.71 | 5.67 | 0.47 | 0.9 | 1.04 | 6.86E-05 | 8.10E-04 | (A) NE (B/D) |
| STAG3L1 | 2.46 | 1.42 | 0.53 | 0.47 | 1.04 | 2.01E-04 | 1.85E-03 | (A) NE (B/D) |
| RPS9 | 3.98 | 2.94 | 0.3 | 1.37 | 1.04 | 2.04E-04 | 1.86E-03 | (A) NE (B/D) |
| FAM104A | 8.15 | 7.11 | 0.49 | 1.07 | 1.04 | 2.16E-04 | 1.95E-03 | (A) NE (B/D) |

| | | | | | | | | |
|---|---|---|---|---|---|---|---|---|
| LRRC24 | 2.32 | 1.28 | 0.54 | 0.78 | 1.04 | 2.20E-04 | 1.97E-03 | (A) NE (B/D) |
| SMIM26 | 7.29 | 6.25 | 0.36 | 1.32 | 1.04 | 2.33E-04 | 2.07E-03 | (A) NE (B/D) |
| PHC1 | 7.69 | 6.66 | 0.48 | 1.14 | 1.04 | 2.59E-04 | 2.24E-03 | (A) NE (B/D) |
| SEMA4C | 1.31 | 0.27 | 0.56 | 0.4 | 1.04 | 3.37E-04 | 2.76E-03 | (A) NE (B/D) |
| SLC39A7 | 10.82 | 9.78 | 0.39 | 1.48 | 1.04 | 7.29E-04 | 4.99E-03 | (A) NE (B/D) |
| PGAM1P8 | 2.64 | 1.6 | 0.59 | 1.08 | 1.04 | 7.52E-04 | 5.11E-03 | (A) NE (B/D) |
| LRRC34 | 1.06 | 0.01 | 0.61 | 0.06 | 1.04 | 9.24E-04 | 5.96E-03 | (A) NE (B/D) |
| ATRAID | 5.31 | 4.27 | 0.57 | 1.28 | 1.04 | 1.15E-03 | 7.01E-03 | (A) NE (B/D) |
| CAPNS1 | 3.15 | 2.1 | 0.56 | 1.3 | 1.04 | 1.15E-03 | 7.02E-03 | (A) NE (B/D) |
| RNU6-48P | 2.41 | 1.37 | 0.66 | 0.86 | 1.04 | 1.27E-03 | 7.56E-03 | (A) NE (B/D) |
| FPGS | 6.9 | 5.86 | 0.61 | 1.2 | 1.04 | 1.37E-03 | 8.05E-03 | (A) NE (B/D) |
| DHRS1 | 2.13 | 1.09 | 0.68 | 0.98 | 1.04 | 1.78E-03 | 9.81E-03 | (A) NE (B/D) |
| MRPL52 | 4.11 | 3.07 | 0.68 | 1.11 | 1.04 | 2.00E-03 | 0.0107 | (A) NE (B/D) |
| EXD3 | 1.29 | 0.26 | 0.72 | 0.52 | 1.04 | 2.10E-03 | 0.0111 | (A) NE (B/D) |
| CABP4 | 1.45 | 0.41 | 0.73 | 0.66 | 1.04 | 2.39E-03 | 0.0123 | (A) NE (B/D) |
| CTU1 | 5.34 | 4.3 | 0.75 | 0.76 | 1.04 | 2.91E-03 | 0.0143 | (A) NE (B/D) |
| MIR-106B/10 | 2.83 | 1.78 | 0.66 | 1.4 | 1.04 | 3.35E-03 | 0.0159 | (A) NE (B/D) |
| MIR-106B/10 | 2.83 | 1.78 | 0.66 | 1.4 | 1.04 | 3.35E-03 | 0.0159 | (A) NE (B/D) |
| MIR-106B/3P | 2.83 | 1.78 | 0.66 | 1.4 | 1.04 | 3.35E-03 | 0.0159 | (A) NE (B/D) |
| MIR-106B/5P | 2.83 | 1.78 | 0.66 | 1.4 | 1.04 | 3.35E-03 | 0.0159 | (A) NE (B/D) |
| CD276 | 3.48 | 2.44 | 0.59 | 1.65 | 1.04 | 4.42E-03 | 0.0196 | (A) NE (B/D) |
| ILDR1 | 1.8 | 0.76 | 0.86 | 0.91 | 1.04 | 7.28E-03 | 0.0284 | (A) NE (B/D) |
| PCDHGC5 | 1.41 | 0.36 | 0.89 | 0.51 | 1.04 | 7.71E-03 | 0.0295 | (A) NE (B/D) |
| ZBTB8OS | 5.06 | 4.01 | 0.88 | 0.84 | 1.04 | 7.79E-03 | 0.0297 | (A) NE (B/D) |
| RPS3AP26 | 8.69 | 7.65 | 0.74 | 1.62 | 1.04 | 9.22E-03 | 0.0336 | (A) NE (B/D) |
| TMSB4XP4 | 3.57 | 2.53 | 0.77 | 1.58 | 1.04 | 9.67E-03 | 0.0348 | (A) NE (B/D) |
| HMGN2P15 | 3.92 | 2.88 | 0.85 | 1.4 | 1.04 | 0.0107 | 0.0374 | (A) NE (B/D) |
| FAM220CP | 3 | 1.96 | 0.79 | 1.7 | 1.04 | 0.0125 | 0.042 | (A) NE (B/D) |
| FRMPD1 | 1.68 | 0.64 | 1.01 | 0.66 | 1.04 | 0.0152 | 0.0485 | (A) NE (B/D) |
| SH3BP5L | 2.39 | 1.36 | 0.19 | 0.78 | 1.03 | 1.84E-08 | 1.26E-06 | (A) NE (B/D) |
| AP5Z1 | 1.23 | 0.19 | 0.29 | 0.38 | 1.03 | 1.41E-07 | 6.37E-06 | (A) NE (B/D) |
| IPPK | 6.05 | 5.02 | 0.3 | 0.85 | 1.03 | 1.10E-06 | 3.21E-05 | (A) NE (B/D) |
| GIT1 | 1.47 | 0.45 | 0.35 | 0.57 | 1.03 | 1.46E-06 | 4.02E-05 | (A) NE (B/D) |
| LMBR1L | 1.68 | 0.65 | 0.35 | 0.78 | 1.03 | 2.33E-06 | 5.80E-05 | (A) NE (B/D) |
| DHX16 | 6.33 | 5.3 | 0.26 | 1 | 1.03 | 3.87E-06 | 8.64E-05 | (A) NE (B/D) |
| SRSF3 | 2.95 | 1.93 | 0.3 | 0.98 | 1.03 | 5.72E-06 | 1.17E-04 | (A) NE (B/D) |
| MAFG | 4.58 | 3.55 | 0.32 | 0.97 | 1.03 | 6.71E-06 | 1.33E-04 | (A) NE (B/D) |
| TMEM18 | 3.45 | 2.41 | 0.31 | 1.01 | 1.03 | 7.83E-06 | 1.50E-04 | (A) NE (B/D) |
| IPO9-AS1 | 7.32 | 6.28 | 0.41 | 0.75 | 1.03 | 1.06E-05 | 1.91E-04 | (A) NE (B/D) |
| INO80E | 2.71 | 1.68 | 0.23 | 1.11 | 1.03 | 1.08E-05 | 1.94E-04 | (A) NE (B/D) |
| ZNF2 | 2.08 | 1.05 | 0.32 | 1.05 | 1.03 | 1.56E-05 | 2.57E-04 | (A) NE (B/D) |
| MTCH1 | 6.98 | 5.95 | 0.35 | 1.04 | 1.03 | 2.02E-05 | 3.10E-04 | (A) NE (B/D) |
| LRRC47 | 5.59 | 4.55 | 0.38 | 1 | 1.03 | 2.50E-05 | 3.64E-04 | (A) NE (B/D) |
| DKC1 | 3.55 | 2.52 | 0.35 | 1.12 | 1.03 | 3.97E-05 | 5.27E-04 | (A) NE (B/D) |
| MRFAP1 | 4.95 | 3.92 | 0.26 | 1.2 | 1.03 | 4.05E-05 | 5.35E-04 | (A) NE (B/D) |
| ZNF783 | 3.63 | 2.61 | 0.4 | 1.03 | 1.03 | 5.28E-05 | 6.60E-04 | (A) NE (B/D) |
| PMF1 | 2.38 | 1.35 | 0.43 | 1.16 | 1.03 | 1.72E-04 | 1.64E-03 | (A) NE (B/D) |
| TTYH3 | 6.73 | 5.7 | 0.51 | 1.08 | 1.03 | 3.12E-04 | 2.60E-03 | (A) NE (B/D) |
| CHCHD2 | 11.85 | 10.82 | 0.46 | 1.21 | 1.03 | 3.22E-04 | 2.66E-03 | (A) NE (B/D) |
| INAFM2 | 8.88 | 7.85 | 0.51 | 1.12 | 1.03 | 3.41E-04 | 2.78E-03 | (A) NE (B/D) |
| RPN1 | 3.95 | 2.93 | 0.41 | 1.32 | 1.03 | 3.67E-04 | 2.95E-03 | (A) NE (B/D) |
| BBS4 | 2.75 | 1.72 | 0.53 | 1.1 | 1.03 | 4.24E-04 | 3.29E-03 | (A) NE (B/D) |
| EME1 | 4.52 | 3.49 | 0.45 | 1.3 | 1.03 | 4.72E-04 | 3.58E-03 | (A) NE (B/D) |
| KIAA0895L | 1.15 | 0.12 | 0.58 | 0.31 | 1.03 | 5.73E-04 | 4.16E-03 | (A) NE (B/D) |
| MRPS33 | 4.59 | 3.57 | 0.37 | 1.48 | 1.03 | 7.70E-04 | 5.20E-03 | (A) NE (B/D) |
| HSP90B2P | 4.73 | 3.7 | 0.63 | 0.92 | 1.03 | 9.52E-04 | 6.08E-03 | (A) NE (B/D) |
| COX7B | 7.64 | 6.61 | 0.26 | 1.61 | 1.03 | 1.02E-03 | 6.39E-03 | (A) NE (B/D) |
| GAPDHP40 | 3.17 | 2.14 | 0.55 | 1.29 | 1.03 | 1.12E-03 | 6.90E-03 | (A) NE (B/D) |
| RACK1 | 8.51 | 7.48 | 0.54 | 1.32 | 1.03 | 1.13E-03 | 6.94E-03 | (A) NE (B/D) |
| HMGN1P4 | 6.74 | 5.71 | 0.67 | 1.06 | 1.03 | 1.89E-03 | 0.0103 | (A) NE (B/D) |
| RNF146 | 2.56 | 1.53 | 0.68 | 1.17 | 1.03 | 2.53E-03 | 0.0128 | (A) NE (B/D) |
| POLE3 | 5.55 | 4.52 | 0.5 | 1.6 | 1.03 | 2.62E-03 | 0.0132 | (A) NE (B/D) |
| MASP2 | 1.22 | 0.2 | 0.81 | 0.48 | 1.03 | 4.87E-03 | 0.0211 | (A) NE (B/D) |
| GSTA4 | 7.37 | 6.34 | 0.72 | 1.38 | 1.03 | 5.31E-03 | 0.0224 | (A) NE (B/D) |
| FYN | 1.56 | 0.54 | 0.83 | 0.96 | 1.03 | 6.62E-03 | 0.0264 | (A) NE (B/D) |

| | | | | | | | | |
|---|---|---|---|---|---|---|---|---|
| HPSE | 1.42 | 0.4 | 0.87 | 0.62 | 1.03 | 7.34E-03 | 0.0285 | (A) NE (B/D) |
| COA6 | 7.45 | 6.43 | 0.72 | 1.53 | 1.03 | 7.55E-03 | 0.0291 | (A) NE (B/D) |
| CBX3P2 | 2.09 | 1.07 | 0.85 | 1.13 | 1.03 | 8.40E-03 | 0.0314 | (A) NE (B/D) |
| TCF12 | 3.5 | 2.47 | 0.91 | 0.42 | 1.03 | 9.19E-03 | 0.0336 | (A) NE (B/D) |
| ATP6V0E2 | 4.53 | 3.51 | 0.87 | 1.2 | 1.03 | 0.0102 | 0.0361 | (A) NE (B/D) |
| PSIP1 | 1.78 | 0.75 | 0.92 | 0.7 | 1.03 | 0.0102 | 0.0363 | (A) NE (B/D) |
| ANGPTL6 | 3.75 | 2.72 | 0.86 | 1.31 | 1.03 | 0.0105 | 0.037 | (A) NE (B/D) |
| FOXN4 | 1.19 | 0.17 | 0.96 | 0.52 | 1.03 | 0.0122 | 0.0412 | (A) NE (B/D) |
| PDXK | 7.23 | 6.21 | 0.32 | 0.65 | 1.02 | 3.58E-07 | 1.31E-05 | (A) NE (B/D) |
| PCGF3 | 4.78 | 3.77 | 0.4 | 0.68 | 1.02 | 7.98E-06 | 1.53E-04 | (A) NE (B/D) |
| CCZ1B | 2.64 | 1.61 | 0.37 | 0.9 | 1.02 | 1.16E-05 | 2.06E-04 | (A) NE (B/D) |
| TACO1 | 4.68 | 3.66 | 0.26 | 1.1 | 1.02 | 1.56E-05 | 2.56E-04 | (A) NE (B/D) |
| CENPS-CORT | 2.35 | 1.33 | 0.41 | 0.91 | 1.02 | 2.84E-05 | 4.04E-04 | (A) NE (B/D) |
| HSPBP1 | 1.28 | 0.26 | 0.43 | 0.44 | 1.02 | 3.26E-05 | 4.54E-04 | (A) NE (B/D) |
| MEPCE | 2.04 | 1.02 | 0.41 | 1 | 1.02 | 4.65E-05 | 5.96E-04 | (A) NE (B/D) |
| VPS72 | 5.46 | 4.44 | 0.39 | 1.15 | 1.02 | 1.05E-04 | 1.13E-03 | (A) NE (B/D) |
| ATP5MG | 2.5 | 1.48 | 0.37 | 1.24 | 1.02 | 1.73E-04 | 1.65E-03 | (A) NE (B/D) |
| TTC4 | 3.36 | 2.34 | 0.5 | 0.96 | 1.02 | 1.81E-04 | 1.70E-03 | (A) NE (B/D) |
| NAXD | 1.15 | 0.13 | 0.54 | 0.33 | 1.02 | 3.49E-04 | 2.83E-03 | (A) NE (B/D) |
| SMIM11A | 3.29 | 2.27 | 0.56 | 0.86 | 1.02 | 3.77E-04 | 3.01E-03 | (A) NE (B/D) |
| AKIRIN1 | 7.63 | 6.61 | 0.57 | 0.93 | 1.02 | 5.23E-04 | 3.86E-03 | (A) NE (B/D) |
| BRD9 | 1.35 | 0.33 | 0.59 | 0.42 | 1.02 | 6.38E-04 | 4.51E-03 | (A) NE (B/D) |
| SARM1 | 3.28 | 2.27 | 0.64 | 0.88 | 1.02 | 1.21E-03 | 7.30E-03 | (A) NE (B/D) |
| XKR8 | 6.1 | 5.07 | 0.62 | 1.09 | 1.02 | 1.30E-03 | 7.72E-03 | (A) NE (B/D) |
| TTC13 | 8.13 | 7.11 | 0.63 | 1.08 | 1.02 | 1.44E-03 | 8.33E-03 | (A) NE (B/D) |
| MIR-641/641 | 2.12 | 1.1 | 0.67 | 0.95 | 1.02 | 1.66E-03 | 9.32E-03 | (A) NE (B/D) |
| ZNF202 | 1.62 | 0.6 | 0.68 | 0.81 | 1.02 | 1.72E-03 | 9.55E-03 | (A) NE (B/D) |
| TPRG1L | 3.25 | 2.23 | 0.41 | 1.59 | 1.02 | 1.78E-03 | 9.81E-03 | (A) NE (B/D) |
| GSTO1 | 3.34 | 2.32 | 0.57 | 1.4 | 1.02 | 2.13E-03 | 0.0112 | (A) NE (B/D) |
| SEL1L3 | 1.39 | 0.37 | 0.71 | 0.88 | 1.02 | 2.36E-03 | 0.0121 | (A) NE (B/D) |
| F8A2 | 1.45 | 0.43 | 0.73 | 0.75 | 1.02 | 2.55E-03 | 0.0129 | (A) NE (B/D) |
| MTFR1L | 5.16 | 4.14 | 0.7 | 1.16 | 1.02 | 3.14E-03 | 0.0151 | (A) NE (B/D) |
| SLC29A3 | 6.26 | 5.24 | 0.7 | 1.29 | 1.02 | 4.17E-03 | 0.0188 | (A) NE (B/D) |
| TTLL10-AS1 | 1.89 | 0.87 | 0.79 | 0.83 | 1.02 | 4.61E-03 | 0.0202 | (A) NE (B/D) |
| MIR-4453/44 | 4.64 | 3.63 | 0.8 | 0.76 | 1.02 | 4.91E-03 | 0.0212 | (A) NE (B/D) |
| ERMAP | 2.54 | 1.52 | 0.68 | 1.57 | 1.02 | 6.73E-03 | 0.0268 | (A) NE (B/D) |
| TMEM185AP | 5.45 | 4.44 | 0.82 | 1.12 | 1.02 | 7.27E-03 | 0.0283 | (A) NE (B/D) |
| TMEM35B | 10.3 | 9.28 | 0.79 | 1.36 | 1.02 | 8.58E-03 | 0.0319 | (A) NE (B/D) |
| RNA5SP473 | 1.43 | 0.41 | 0.89 | 0.79 | 1.02 | 9.02E-03 | 0.0331 | (A) NE (B/D) |
| SULF1 | 3.4 | 2.39 | 0.77 | 1.77 | 1.02 | 0.0157 | 0.0496 | (A) NE (B/D) |
| GCSHP3 | 4.29 | 3.28 | 0.36 | 0.68 | 1.01 | 3.01E-06 | 7.02E-05 | (A) NE (B/D) |
| FAAP100 | 5.6 | 4.59 | 0.33 | 0.95 | 1.01 | 9.14E-06 | 1.69E-04 | (A) NE (B/D) |
| MRGBP | 6.93 | 5.92 | 0.23 | 1.08 | 1.01 | 9.96E-06 | 1.81E-04 | (A) NE (B/D) |
| IPO9 | 7.92 | 6.91 | 0.34 | 0.96 | 1.01 | 1.15E-05 | 2.04E-04 | (A) NE (B/D) |
| C19ORF54 | 1.81 | 0.8 | 0.4 | 0.82 | 1.01 | 1.53E-05 | 2.53E-04 | (A) NE (B/D) |
| VEZF1 | 3.26 | 2.25 | 0.4 | 0.84 | 1.01 | 2.09E-05 | 3.18E-04 | (A) NE (B/D) |
| KATNA1 | 7.4 | 6.4 | 0.35 | 1 | 1.01 | 2.13E-05 | 3.22E-04 | (A) NE (B/D) |
| TIMM22 | 3.8 | 2.79 | 0.41 | 1 | 1.01 | 6.05E-05 | 7.33E-04 | (A) NE (B/D) |
| QRICH2 | 1.06 | 0.05 | 0.43 | 0.2 | 1.01 | 7.33E-05 | 8.53E-04 | (A) NE (B/D) |
| PRMT7 | 5.59 | 4.58 | 0.5 | 0.84 | 1.01 | 1.47E-04 | 1.46E-03 | (A) NE (B/D) |
| FAM136A | 8.5 | 7.49 | 0.37 | 1.23 | 1.01 | 1.70E-04 | 1.63E-03 | (A) NE (B/D) |
| MMP23A | 2.39 | 1.37 | 0.47 | 1.05 | 1.01 | 1.79E-04 | 1.69E-03 | (A) NE (B/D) |
| CKAP5 | 2.39 | 1.39 | 0.53 | 1.02 | 1.01 | 4.29E-04 | 3.32E-03 | (A) NE (B/D) |
| ABHD12 | 5.87 | 4.86 | 0.38 | 1.39 | 1.01 | 5.63E-04 | 4.10E-03 | (A) NE (B/D) |
| BGLAP | 1.42 | 0.41 | 0.61 | 0.79 | 1.01 | 8.34E-04 | 5.53E-03 | (A) NE (B/D) |
| IRF7 | 2.03 | 1.03 | 0.59 | 1.06 | 1.01 | 9.80E-04 | 6.22E-03 | (A) NE (B/D) |
| AP2S1 | 5.88 | 4.87 | 0.49 | 1.33 | 1.01 | 1.01E-03 | 6.35E-03 | (A) NE (B/D) |
| MTLN | 3.4 | 2.39 | 0.62 | 1.07 | 1.01 | 1.31E-03 | 7.75E-03 | (A) NE (B/D) |
| PKDCC | 1.15 | 0.14 | 0.67 | 0.35 | 1.01 | 1.69E-03 | 9.42E-03 | (A) NE (B/D) |
| DCAF11 | 2.78 | 1.77 | 0.62 | 1.15 | 1.01 | 1.76E-03 | 9.73E-03 | (A) NE (B/D) |
| LENG8-AS1 | 7.92 | 6.92 | 0.68 | 0.79 | 1.01 | 1.79E-03 | 9.84E-03 | (A) NE (B/D) |
| RPL37A | 8.7 | 7.69 | 0.48 | 1.52 | 1.01 | 1.99E-03 | 0.0107 | (A) NE (B/D) |
| RER1 | 3.45 | 2.44 | 0.48 | 1.52 | 1.01 | 2.09E-03 | 0.0111 | (A) NE (B/D) |
| SEC31B | 4.21 | 3.2 | 0.62 | 1.24 | 1.01 | 2.12E-03 | 0.0112 | (A) NE (B/D) |
| RN7SL663P | 1.11 | 0.09 | 0.71 | 0.21 | 1.01 | 2.60E-03 | 0.0131 | (A) NE (B/D) |

| | | | | | | | | |
|---|---|---|---|---|---|---|---|---|
| ITPK1 | 1.86 | 0.85 | 0.71 | 0.9 | 1.01 | 2.74E-03 | 0.0137 | (A) NE (B/D) |
| FKBPL | 3.91 | 2.9 | 0.62 | 1.33 | 1.01 | 2.80E-03 | 0.0139 | (A) NE (B/D) |
| MYL12B | 2.62 | 1.62 | 0.56 | 1.48 | 1.01 | 2.92E-03 | 0.0143 | (A) NE (B/D) |
| CHCHD2P6 | 8.75 | 7.75 | 0.67 | 1.22 | 1.01 | 3.06E-03 | 0.0148 | (A) NE (B/D) |
| SOCS5P4 | 4.5 | 3.49 | 0.78 | 0.78 | 1.01 | 4.48E-03 | 0.0198 | (A) NE (B/D) |
| RN7SL614P | 2.49 | 1.48 | 0.8 | 1 | 1.01 | 6.01E-03 | 0.0246 | (A) NE (B/D) |
| PJVK | 4.17 | 3.16 | 0.74 | 1.31 | 1.01 | 6.10E-03 | 0.0249 | (A) NE (B/D) |
| HCRTR1 | 3.65 | 2.64 | 0.84 | 0.75 | 1.01 | 6.65E-03 | 0.0265 | (A) NE (B/D) |
| SPR | 3.14 | 2.13 | 0.79 | 1.37 | 1.01 | 9.24E-03 | 0.0337 | (A) NE (B/D) |
| SNX10 | 1.26 | 0.24 | 0.91 | 0.49 | 1.01 | 0.0103 | 0.0365 | (A) NE (B/D) |
| HPS3 | 1.97 | 0.96 | 0.93 | 1 | 1.01 | 0.0138 | 0.0452 | (A) NE (B/D) |
| ELMO2 | 6.14 | 5.14 | 0.32 | 0.92 | 1 | 6.41E-06 | 1.29E-04 | (A) NE (B/D) |
| DAXX | 1.73 | 0.73 | 0.38 | 0.76 | 1 | 8.09E-06 | 1.54E-04 | (A) NE (B/D) |
| PPP2R1A | 2.38 | 1.38 | 0.35 | 0.93 | 1 | 1.37E-05 | 2.34E-04 | (A) NE (B/D) |
| PRPF3 | 3.8 | 2.81 | 0.34 | 1.08 | 1 | 3.97E-05 | 5.27E-04 | (A) NE (B/D) |
| C1ORF174 | 2.19 | 1.19 | 0.43 | 0.94 | 1 | 6.99E-05 | 8.22E-04 | (A) NE (B/D) |
| MAGI1 | 4.97 | 3.97 | 0.39 | 1.13 | 1 | 1.21E-04 | 1.25E-03 | (A) NE (B/D) |
| KHDRBS1 | 8.19 | 7.18 | 0.47 | 0.98 | 1 | 1.55E-04 | 1.52E-03 | (A) NE (B/D) |
| TBC1D3G | 4.89 | 3.9 | 0.3 | 1.29 | 1 | 1.94E-04 | 1.80E-03 | (A) NE (B/D) |
| OTUD3 | 5.13 | 4.14 | 0.51 | 0.92 | 1 | 2.68E-04 | 2.30E-03 | (A) NE (B/D) |
| DYNLL1 | 2.48 | 1.48 | 0.2 | 1.43 | 1 | 3.42E-04 | 2.79E-03 | (A) NE (B/D) |
| ASB1 | 1.01 | 0.01 | 0.51 | 0.05 | 1 | 3.54E-04 | 2.87E-03 | (A) NE (B/D) |
| PKP4P1 | 1.87 | 0.87 | 0.57 | 0.73 | 1 | 4.33E-04 | 3.34E-03 | (A) NE (B/D) |
| EIF3I | 7.2 | 6.21 | 0.36 | 1.46 | 1 | 8.87E-04 | 5.78E-03 | (A) NE (B/D) |
| PTPN18 | 2.99 | 1.99 | 0.44 | 1.37 | 1 | 8.89E-04 | 5.79E-03 | (A) NE (B/D) |
| BCL2L2 | 4.09 | 3.09 | 0.6 | 1.12 | 1 | 1.50E-03 | 8.62E-03 | (A) NE (B/D) |
| EXO1 | 1.96 | 0.96 | 0.66 | 0.97 | 1 | 1.83E-03 | 0.01 | (A) NE (B/D) |
| KPNA6 | 6.13 | 5.13 | 0.7 | 0.53 | 1 | 2.34E-03 | 0.0121 | (A) NE (B/D) |
| EI24 | 3.81 | 2.8 | 0.61 | 1.3 | 1 | 2.51E-03 | 0.0128 | (A) NE (B/D) |
| PASK | 5.96 | 4.96 | 0.56 | 1.44 | 1 | 2.76E-03 | 0.0137 | (A) NE (B/D) |
| IQCH-AS1 | 2.32 | 1.33 | 0.75 | 0.62 | 1 | 3.81E-03 | 0.0175 | (A) NE (B/D) |
| GRM8 | 1.01 | 0.01 | 0.78 | 0.06 | 1 | 4.87E-03 | 0.0211 | (A) NE (B/D) |
| CORO2B | 1.01 | 0.01 | 0.8 | 0.08 | 1 | 5.66E-03 | 0.0236 | (A) NE (B/D) |
| MMACHC | 1.43 | 0.43 | 0.82 | 0.55 | 1 | 5.82E-03 | 0.0241 | (A) NE (B/D) |
| PVRIG | 1.07 | 0.07 | 0.85 | 0.23 | 1 | 7.69E-03 | 0.0294 | (A) NE (B/D) |
| POM121C | 7.7 | 6.72 | 0.2 | 0.63 | 0.99 | 1.41E-09 | 1.76E-07 | (A) NE (B/D) |
| NUDCD3 | 8.52 | 7.53 | 0.15 | 0.88 | 0.99 | 2.61E-07 | 1.03E-05 | (A) NE (B/D) |
| ZNF212 | 4.16 | 3.16 | 0.3 | 0.98 | 0.99 | 9.62E-06 | 1.76E-04 | (A) NE (B/D) |
| SLC41A3 | 3.47 | 2.49 | 0.23 | 1.07 | 0.99 | 1.26E-05 | 2.19E-04 | (A) NE (B/D) |
| ROMO1 | 2.32 | 1.33 | 0.32 | 1 | 0.99 | 1.47E-05 | 2.47E-04 | (A) NE (B/D) |
| ZNF514 | 2.65 | 1.67 | 0.42 | 0.77 | 0.99 | 3.23E-05 | 4.52E-04 | (A) NE (B/D) |
| TRAPPC12-AS | 5.15 | 4.16 | 0.44 | 0.85 | 0.99 | 7.14E-05 | 8.34E-04 | (A) NE (B/D) |
| UQCC1 | 7.08 | 6.09 | 0.48 | 0.79 | 0.99 | 1.19E-04 | 1.23E-03 | (A) NE (B/D) |
| RDH14 | 8.91 | 7.93 | 0.47 | 1.06 | 0.99 | 2.97E-04 | 2.50E-03 | (A) NE (B/D) |
| MTSS2 | 7.05 | 6.06 | 0.42 | 1.23 | 0.99 | 3.45E-04 | 2.80E-03 | (A) NE (B/D) |
| UBL7-AS1 | 3.02 | 2.03 | 0.56 | 0.58 | 0.99 | 4.05E-04 | 3.18E-03 | (A) NE (B/D) |
| HNRNPH3 | 2.26 | 1.26 | 0.48 | 1.26 | 0.99 | 6.97E-04 | 4.84E-03 | (A) NE (B/D) |
| INPP5K | 3.04 | 2.04 | 0.46 | 1.31 | 0.99 | 8.02E-04 | 5.38E-03 | (A) NE (B/D) |
| U2AF2 | 1.42 | 0.43 | 0.6 | 0.54 | 0.99 | 8.17E-04 | 5.45E-03 | (A) NE (B/D) |
| RELT | 2.19 | 1.2 | 0.59 | 0.88 | 0.99 | 8.33E-04 | 5.53E-03 | (A) NE (B/D) |
| RSPH9 | 1.99 | 1 | 0.6 | 0.87 | 0.99 | 8.94E-04 | 5.81E-03 | (A) NE (B/D) |
| IRF3 | 3.72 | 2.74 | 0.58 | 1.01 | 0.99 | 8.99E-04 | 5.84E-03 | (A) NE (B/D) |
| LINC00999 | 3.86 | 2.86 | 0.62 | 0.63 | 0.99 | 9.26E-04 | 5.96E-03 | (A) NE (B/D) |
| MRPL37 | 3.72 | 2.73 | 0.58 | 1.31 | 0.99 | 2.19E-03 | 0.0114 | (A) NE (B/D) |
| FAM229A | 6.8 | 5.81 | 0.69 | 1 | 0.99 | 2.84E-03 | 0.014 | (A) NE (B/D) |
| BSCL2 | 6.57 | 5.58 | 0.66 | 1.19 | 0.99 | 2.89E-03 | 0.0142 | (A) NE (B/D) |
| AHDC1 | 1.09 | 0.1 | 0.71 | 0.23 | 0.99 | 3.04E-03 | 0.0147 | (A) NE (B/D) |
| PNN | 3.02 | 2.03 | 0.66 | 1.2 | 0.99 | 3.15E-03 | 0.0151 | (A) NE (B/D) |
| PXDN | 1.43 | 0.44 | 0.73 | 0.69 | 0.99 | 3.32E-03 | 0.0158 | (A) NE (B/D) |
| CYP27A1 | 1.73 | 0.74 | 0.73 | 1.03 | 0.99 | 4.27E-03 | 0.0191 | (A) NE (B/D) |
| NEK9 | 3.01 | 2.02 | 0.73 | 1.08 | 0.99 | 4.35E-03 | 0.0194 | (A) NE (B/D) |
| TMEM35A | 1.21 | 0.22 | 0.77 | 0.68 | 0.99 | 4.37E-03 | 0.0194 | (A) NE (B/D) |
| EZH2 | 1.08 | 0.09 | 0.78 | 0.33 | 0.99 | 4.77E-03 | 0.0208 | (A) NE (B/D) |
| TDRD7 | 4.71 | 3.72 | 0.76 | 0.92 | 0.99 | 4.84E-03 | 0.021 | (A) NE (B/D) |
| NKX2-8 | 1 | 0 | 0.77 | 0.02 | 0.99 | 4.88E-03 | 0.0211 | (A) NE (B/D) |

| | | | | | | | | |
|---|---|---|---|---|---|---|---|---|
| ATF4P3 | 4.15 | 3.16 | 0.79 | 1.02 | 0.99 | 6.15E-03 | 0.0251 | (A) NE (B/D) |
| CDAN1 | 1.04 | 0.05 | 0.81 | 0.2 | 0.99 | 6.27E-03 | 0.0254 | (A) NE (B/D) |
| ZNF204P | 1 | 0.01 | 0.83 | 0.06 | 0.99 | 7.14E-03 | 0.028 | (A) NE (B/D) |
| GPR137 | 2.25 | 1.26 | 0.78 | 1.21 | 0.99 | 7.49E-03 | 0.0289 | (A) NE (B/D) |
| PDLIM7 | 5.35 | 4.36 | 0.75 | 1.41 | 0.99 | 8.93E-03 | 0.0329 | (A) NE (B/D) |
| CHMP4C | 2.33 | 1.34 | 0.8 | 1.28 | 0.99 | 9.19E-03 | 0.0336 | (A) NE (B/D) |
| SEPHS1P6 | 3.74 | 2.75 | 0.87 | 0.99 | 0.99 | 0.0105 | 0.037 | (A) NE (B/D) |
| GMNN | 4.02 | 3.02 | 0.6 | 1.84 | 0.99 | 0.0113 | 0.0389 | (A) NE (B/D) |
| CDKAL1 | 1.25 | 0.26 | 0.94 | 0.53 | 0.99 | 0.0133 | 0.044 | (A) NE (B/D) |
| PAFAH1B1 | 4.41 | 3.42 | 0.93 | 0.81 | 0.99 | 0.0134 | 0.044 | (A) NE (B/D) |
| ZNF316 | 2.68 | 1.69 | 0.96 | 0.52 | 0.99 | 0.0151 | 0.0483 | (A) NE (B/D) |
| ZNF3 | 1.34 | 0.36 | 0.21 | 0.51 | 0.98 | 3.65E-10 | 6.52E-08 | (A) NE (B/D) |
| LPCAT1 | 1.74 | 0.76 | 0.19 | 0.65 | 0.98 | 2.79E-09 | 3.03E-07 | (A) NE (B/D) |
| CDK5RAP1 | 1.33 | 0.35 | 0.26 | 0.45 | 0.98 | 2.05E-08 | 1.35E-06 | (A) NE (B/D) |
| PRPF31 | 1.57 | 0.59 | 0.29 | 0.47 | 0.98 | 1.71E-07 | 7.44E-06 | (A) NE (B/D) |
| ZNF142 | 7.2 | 6.22 | 0.35 | 0.62 | 0.98 | 2.74E-06 | 6.54E-05 | (A) NE (B/D) |
| MED22 | 3.27 | 2.29 | 0.33 | 0.83 | 0.98 | 4.76E-06 | 1.02E-04 | (A) NE (B/D) |
| ZMYND19 | 1.66 | 0.67 | 0.4 | 0.77 | 0.98 | 1.74E-05 | 2.77E-04 | (A) NE (B/D) |
| NIPA2 | 7.14 | 6.16 | 0.38 | 0.88 | 0.98 | 2.14E-05 | 3.23E-04 | (A) NE (B/D) |
| FEM1A | 5.02 | 4.03 | 0.45 | 0.7 | 0.98 | 5.60E-05 | 6.92E-04 | (A) NE (B/D) |
| XPO5 | 1.45 | 0.47 | 0.49 | 0.67 | 0.98 | 1.34E-04 | 1.35E-03 | (A) NE (B/D) |
| AGO1 | 5.71 | 4.73 | 0.49 | 0.51 | 0.98 | 1.59E-04 | 1.55E-03 | (A) NE (B/D) |
| POLR1D | 2.71 | 1.74 | 0.13 | 1.32 | 0.98 | 1.64E-04 | 1.58E-03 | (A) NE (B/D) |
| PIM1 | 2.1 | 1.13 | 0.34 | 1.21 | 0.98 | 1.81E-04 | 1.70E-03 | (A) NE (B/D) |
| MTIF3 | 5.29 | 4.31 | 0.37 | 1.23 | 0.98 | 2.54E-04 | 2.21E-03 | (A) NE (B/D) |
| NDUFA2 | 3.4 | 2.42 | 0.27 | 1.35 | 0.98 | 3.16E-04 | 2.63E-03 | (A) NE (B/D) |
| PSMD10 | 3.05 | 2.07 | 0.33 | 1.38 | 0.98 | 5.15E-04 | 3.82E-03 | (A) NE (B/D) |
| BLOC1S3 | 4.41 | 3.43 | 0.49 | 1.16 | 0.98 | 6.01E-04 | 4.32E-03 | (A) NE (B/D) |
| SYNJ2 | 2.85 | 1.87 | 0.63 | 0.68 | 0.98 | 1.28E-03 | 7.61E-03 | (A) NE (B/D) |
| SINHCAF | 2.64 | 1.66 | 0.66 | 0.72 | 0.98 | 1.61E-03 | 9.08E-03 | (A) NE (B/D) |
| POLR2K | 7.28 | 6.3 | 0.46 | 1.44 | 0.98 | 1.83E-03 | 0.01 | (A) NE (B/D) |
| FBXL2 | 1.35 | 0.37 | 0.77 | 0.5 | 0.98 | 4.58E-03 | 0.0201 | (A) NE (B/D) |
| CICP22 | 1.21 | 0.23 | 0.8 | 0.4 | 0.98 | 5.84E-03 | 0.0241 | (A) NE (B/D) |
| RAC1P2 | 3.99 | 3 | 0.75 | 1.12 | 0.98 | 5.90E-03 | 0.0243 | (A) NE (B/D) |
| ATPAF1 | 1.9 | 0.92 | 0.79 | 1.02 | 0.98 | 6.61E-03 | 0.0264 | (A) NE (B/D) |
| GUSBP5 | 2.68 | 1.69 | 0.76 | 1.28 | 0.98 | 7.92E-03 | 0.0301 | (A) NE (B/D) |
| FANCI | 1.09 | 0.11 | 0.86 | 0.37 | 0.98 | 9.13E-03 | 0.0334 | (A) NE (B/D) |
| RPL36AL | 3.65 | 2.68 | 0.61 | 1.71 | 0.98 | 9.36E-03 | 0.034 | (A) NE (B/D) |
| ERP29 | 2.93 | 1.96 | 0.68 | 1.63 | 0.98 | 0.0106 | 0.037 | (A) NE (B/D) |
| PCBP4 | 1.74 | 0.76 | 0.86 | 1.01 | 0.98 | 0.0106 | 0.0371 | (A) NE (B/D) |
| TINAGL1 | 2.78 | 1.8 | 0.83 | 1.43 | 0.98 | 0.0143 | 0.0462 | (A) NE (B/D) |
| IQCJ-SCHIP1 | 1.24 | 0.26 | 0.96 | 0.52 | 0.98 | 0.0155 | 0.0491 | (A) NE (B/D) |
| C2ORF68 | 7.91 | 6.93 | 0.3 | 0.77 | 0.97 | 1.10E-06 | 3.22E-05 | (A) NE (B/D) |
| CABLES2 | 1.72 | 0.75 | 0.32 | 0.76 | 0.97 | 2.39E-06 | 5.91E-05 | (A) NE (B/D) |
| LRRC41 | 3.01 | 2.03 | 0.29 | 0.85 | 0.97 | 2.40E-06 | 5.92E-05 | (A) NE (B/D) |
| PDCL | 5.87 | 4.9 | 0.24 | 0.93 | 0.97 | 2.67E-06 | 6.40E-05 | (A) NE (B/D) |
| URGCP | 2.63 | 1.67 | 0.28 | 0.91 | 0.97 | 4.37E-06 | 9.52E-05 | (A) NE (B/D) |
| RNF121 | 1.99 | 1.01 | 0.36 | 0.9 | 0.97 | 1.94E-05 | 3.02E-04 | (A) NE (B/D) |
| XRCC5 | 9.89 | 8.91 | 0.33 | 0.98 | 0.97 | 2.22E-05 | 3.31E-04 | (A) NE (B/D) |
| TSEN54 | 1.12 | 0.14 | 0.41 | 0.31 | 0.97 | 4.25E-05 | 5.57E-04 | (A) NE (B/D) |
| LIFR-AS1 | 1.98 | 1.01 | 0.4 | 0.93 | 0.97 | 5.19E-05 | 6.52E-04 | (A) NE (B/D) |
| HACD3 | 4.18 | 3.21 | 0.43 | 1.06 | 0.97 | 1.92E-04 | 1.79E-03 | (A) NE (B/D) |
| CNPPD1 | 2.6 | 1.63 | 0.19 | 1.34 | 0.97 | 2.38E-04 | 2.11E-03 | (A) NE (B/D) |
| HMGN1 | 3.09 | 2.12 | 0.52 | 0.94 | 0.97 | 4.13E-04 | 3.23E-03 | (A) NE (B/D) |
| SULT1A3 | 4.89 | 3.92 | 0.48 | 1.11 | 0.97 | 5.10E-04 | 3.79E-03 | (A) NE (B/D) |
| TMEM256-PL | 3.69 | 2.72 | 0.38 | 1.35 | 0.97 | 6.99E-04 | 4.85E-03 | (A) NE (B/D) |
| LSS | 3.54 | 2.57 | 0.57 | 0.91 | 0.97 | 7.46E-04 | 5.08E-03 | (A) NE (B/D) |
| MIR-1254-1/ | 3.58 | 2.61 | 0.59 | 0.85 | 0.97 | 9.86E-04 | 6.25E-03 | (A) NE (B/D) |
| PSMB2 | 5.88 | 4.91 | 0.5 | 1.27 | 0.97 | 1.13E-03 | 6.91E-03 | (A) NE (B/D) |
| SUCLA2-AS1 | 2.78 | 1.81 | 0.59 | 1.02 | 0.97 | 1.25E-03 | 7.51E-03 | (A) NE (B/D) |
| ZBED1 | 6.85 | 5.88 | 0.63 | 0.63 | 0.97 | 1.29E-03 | 7.66E-03 | (A) NE (B/D) |
| OXA1L | 4.52 | 3.55 | 0.47 | 1.4 | 0.97 | 1.77E-03 | 9.76E-03 | (A) NE (B/D) |
| PARK7 | 3.83 | 2.86 | 0.33 | 1.62 | 0.97 | 2.37E-03 | 0.0122 | (A) NE (B/D) |
| METTL4 | 1.01 | 0.04 | 0.67 | 0.13 | 0.97 | 2.40E-03 | 0.0123 | (A) NE (B/D) |
| LINC00501 | 1.64 | 0.67 | 0.69 | 0.76 | 0.97 | 2.65E-03 | 0.0133 | (A) NE (B/D) |

| | | | | | | | | |
|---|---|---|---|---|---|---|---|---|
| KIF9-AS1 | 2.23 | 1.27 | 0.72 | 0.74 | 0.97 | 3.68E-03 | 0.017 | (A) NE (B/D) |
| EIF1B | 4.05 | 3.08 | 0.6 | 1.52 | 0.97 | 5.77E-03 | 0.0239 | (A) NE (B/D) |
| TMEM8B | 4.59 | 3.62 | 0.75 | 1.24 | 0.97 | 7.22E-03 | 0.0282 | (A) NE (B/D) |
| SLC25A12 | 1.08 | 0.11 | 0.82 | 0.27 | 0.97 | 7.43E-03 | 0.0288 | (A) NE (B/D) |
| FAM13A | 1.04 | 0.06 | 0.83 | 0.36 | 0.97 | 7.93E-03 | 0.0301 | (A) NE (B/D) |
| MPL | 3.53 | 2.56 | 0.81 | 0.96 | 0.97 | 7.95E-03 | 0.0301 | (A) NE (B/D) |
| UBTFL6 | 1.6 | 0.62 | 0.83 | 0.81 | 0.97 | 8.16E-03 | 0.0307 | (A) NE (B/D) |
| EVL | 7.24 | 6.27 | 0.76 | 1.33 | 0.97 | 9.19E-03 | 0.0336 | (A) NE (B/D) |
| RPS10P3 | 5.02 | 4.05 | 0.47 | 1.87 | 0.97 | 0.01 | 0.0358 | (A) NE (B/D) |
| UMODL1 | 1.67 | 0.69 | 0.87 | 1.08 | 0.97 | 0.0124 | 0.0418 | (A) NE (B/D) |
| COX8C | 1.46 | 0.49 | 0.94 | 0.62 | 0.97 | 0.0153 | 0.0486 | (A) NE (B/D) |
| ARHGEF18 | 1.27 | 0.31 | 0.24 | 0.56 | 0.96 | 8.96E-09 | 7.25E-07 | (A) NE (B/D) |
| MBD1 | 1.47 | 0.52 | 0.28 | 0.65 | 0.96 | 2.29E-07 | 9.32E-06 | (A) NE (B/D) |
| OSBPL2 | 6.72 | 5.76 | 0.25 | 0.81 | 0.96 | 5.42E-07 | 1.84E-05 | (A) NE (B/D) |
| U2AF1 | 1.07 | 0.11 | 0.3 | 0.23 | 0.96 | 2.71E-06 | 6.48E-05 | (A) NE (B/D) |
| FSCN3 | 1.97 | 1.01 | 0.1 | 1.01 | 0.96 | 4.32E-06 | 9.46E-05 | (A) NE (B/D) |
| ATAD3A | 2.07 | 1.11 | 0.37 | 0.55 | 0.96 | 7.94E-06 | 1.52E-04 | (A) NE (B/D) |
| CUL4A | 2.01 | 1.06 | 0.34 | 0.85 | 0.96 | 1.02E-05 | 1.85E-04 | (A) NE (B/D) |
| PI4KA | 1.29 | 0.33 | 0.39 | 0.51 | 0.96 | 1.55E-05 | 2.55E-04 | (A) NE (B/D) |
| DCTD | 2.16 | 1.2 | 0.35 | 0.92 | 0.96 | 1.89E-05 | 2.95E-04 | (A) NE (B/D) |
| KLHDC2 | 1.78 | 0.82 | 0.4 | 0.82 | 0.96 | 3.25E-05 | 4.53E-04 | (A) NE (B/D) |
| SPAG5-AS1 | 8.18 | 7.22 | 0.41 | 0.8 | 0.96 | 3.59E-05 | 4.87E-04 | (A) NE (B/D) |
| RTN3 | 2.69 | 1.73 | 0.36 | 1.02 | 0.96 | 5.91E-05 | 7.21E-04 | (A) NE (B/D) |
| WDTC1 | 2.24 | 1.28 | 0.45 | 0.76 | 0.96 | 9.06E-05 | 1.01E-03 | (A) NE (B/D) |
| SHLD1 | 2.67 | 1.71 | 0.43 | 0.9 | 0.96 | 9.23E-05 | 1.02E-03 | (A) NE (B/D) |
| GEMIN4 | 5.71 | 4.75 | 0.4 | 1.09 | 0.96 | 1.73E-04 | 1.64E-03 | (A) NE (B/D) |
| MOSPD3 | 2.08 | 1.12 | 0.46 | 0.95 | 0.96 | 1.96E-04 | 1.82E-03 | (A) NE (B/D) |
| PEX6 | 4.44 | 3.48 | 0.43 | 1.05 | 0.96 | 2.05E-04 | 1.87E-03 | (A) NE (B/D) |
| DCTN2 | 8.27 | 7.3 | 0.46 | 1.07 | 0.96 | 3.42E-04 | 2.79E-03 | (A) NE (B/D) |
| YKT6 | 4.22 | 3.25 | 0.35 | 1.28 | 0.96 | 3.83E-04 | 3.05E-03 | (A) NE (B/D) |
| DHRS4-AS1 | 1.81 | 0.85 | 0.51 | 0.89 | 0.96 | 3.89E-04 | 3.09E-03 | (A) NE (B/D) |
| DNAJC9 | 5.39 | 4.43 | 0.49 | 1.14 | 0.96 | 6.69E-04 | 4.68E-03 | (A) NE (B/D) |
| LINC00467 | 4.66 | 3.71 | 0.26 | 1.42 | 0.96 | 6.93E-04 | 4.81E-03 | (A) NE (B/D) |
| REV1 | 8.59 | 7.63 | 0.59 | 0.65 | 0.96 | 8.06E-04 | 5.40E-03 | (A) NE (B/D) |
| SSSCA1-AS1 | 4.43 | 3.47 | 0.59 | 0.91 | 0.96 | 1.17E-03 | 7.14E-03 | (A) NE (B/D) |
| CHEK2 | 6.2 | 5.24 | 0.59 | 0.9 | 0.96 | 1.18E-03 | 7.16E-03 | (A) NE (B/D) |
| EPB41L4A-AS | 7.45 | 6.49 | 0.49 | 1.27 | 0.96 | 1.26E-03 | 7.52E-03 | (A) NE (B/D) |
| MORF4L2 | 10.72 | 9.76 | 0.52 | 1.27 | 0.96 | 1.49E-03 | 8.58E-03 | (A) NE (B/D) |
| TBCE | 8.02 | 7.06 | 0.52 | 1.29 | 0.96 | 1.71E-03 | 9.50E-03 | (A) NE (B/D) |
| DPM2 | 2.94 | 1.99 | 0.56 | 1.19 | 0.96 | 1.72E-03 | 9.58E-03 | (A) NE (B/D) |
| GNG5 | 7.43 | 6.46 | 0.49 | 1.37 | 0.96 | 1.79E-03 | 9.87E-03 | (A) NE (B/D) |
| ELOVL1 | 5.39 | 4.44 | 0.46 | 1.43 | 0.96 | 1.95E-03 | 0.0105 | (A) NE (B/D) |
| DPY30 | 3.02 | 2.07 | 0.41 | 1.49 | 0.96 | 2.01E-03 | 0.0108 | (A) NE (B/D) |
| BTBD6 | 3.74 | 2.78 | 0.58 | 1.34 | 0.96 | 3.12E-03 | 0.015 | (A) NE (B/D) |
| UBE2Q2P2 | 2.27 | 1.31 | 0.7 | 0.76 | 0.96 | 3.16E-03 | 0.0152 | (A) NE (B/D) |
| MYO19 | 1.27 | 0.32 | 0.71 | 0.54 | 0.96 | 3.44E-03 | 0.0162 | (A) NE (B/D) |
| AARD | 1.13 | 0.17 | 0.72 | 0.46 | 0.96 | 3.66E-03 | 0.0169 | (A) NE (B/D) |
| NEGR1 | 1.13 | 0.17 | 0.74 | 0.38 | 0.96 | 4.32E-03 | 0.0193 | (A) NE (B/D) |
| RPL29P24 | 1.85 | 0.89 | 0.74 | 0.76 | 0.96 | 4.40E-03 | 0.0195 | (A) NE (B/D) |
| NOP10 | 10.41 | 9.45 | 0.59 | 1.44 | 0.96 | 4.76E-03 | 0.0208 | (A) NE (B/D) |
| FAM167B | 1.41 | 0.46 | 0.77 | 0.65 | 0.96 | 5.37E-03 | 0.0226 | (A) NE (B/D) |
| SURF1 | 5.51 | 4.55 | 0.51 | 1.66 | 0.96 | 5.86E-03 | 0.0242 | (A) NE (B/D) |
| HINT2 | 4.55 | 3.59 | 0.5 | 1.67 | 0.96 | 5.97E-03 | 0.0245 | (A) NE (B/D) |
| NDRG3 | 9.1 | 8.13 | 0.59 | 1.54 | 0.96 | 6.10E-03 | 0.0249 | (A) NE (B/D) |
| DPY19L2P2 | 1.1 | 0.14 | 0.81 | 0.53 | 0.96 | 7.11E-03 | 0.0279 | (A) NE (B/D) |
| POP7 | 2.55 | 1.59 | 0.73 | 1.24 | 0.96 | 7.26E-03 | 0.0283 | (A) NE (B/D) |
| EBP | 7.41 | 6.45 | 0.5 | 1.74 | 0.96 | 7.72E-03 | 0.0295 | (A) NE (B/D) |
| CTBP2P8 | 4.1 | 3.15 | 0.8 | 0.84 | 0.96 | 7.78E-03 | 0.0297 | (A) NE (B/D) |
| WSB1 | 2.85 | 1.89 | 0.77 | 1.08 | 0.96 | 7.80E-03 | 0.0297 | (A) NE (B/D) |
| FAM71F2 | 1.82 | 0.86 | 0.84 | 0.77 | 0.96 | 9.41E-03 | 0.0342 | (A) NE (B/D) |
| RPL35A | 8.57 | 7.61 | 0.69 | 1.56 | 0.96 | 0.0109 | 0.0379 | (A) NE (B/D) |
| SLC6A6 | 1.73 | 0.77 | 0.83 | 1.12 | 0.96 | 0.011 | 0.038 | (A) NE (B/D) |
| CIR1P2 | 2 | 1.04 | 0.88 | 0.67 | 0.96 | 0.0115 | 0.0393 | (A) NE (B/D) |
| HSPD1P8 | 1.14 | 0.18 | 0.91 | 0.42 | 0.96 | 0.0137 | 0.0449 | (A) NE (B/D) |
| ABHD18 | 4.12 | 3.16 | 0.92 | 0.62 | 0.96 | 0.0143 | 0.0462 | (A) NE (B/D) |

| | | | | | | | | |
|---|---|---|---|---|---|---|---|---|
| TRIM39 | 3.43 | 2.48 | 0.11 | 0.78 | 0.95 | 4.29E-08 | 2.45E-06 | (A) NE (B/D) |
| IFNAR2 | 5.42 | 4.46 | 0.23 | 0.89 | 0.95 | 1.71E-06 | 4.56E-05 | (A) NE (B/D) |
| TAF6L | 1.48 | 0.52 | 0.33 | 0.52 | 0.95 | 1.71E-06 | 4.56E-05 | (A) NE (B/D) |
| EDC3 | 3.08 | 2.13 | 0.22 | 0.91 | 0.95 | 2.49E-06 | 6.09E-05 | (A) NE (B/D) |
| KIAA2013 | 5.75 | 4.79 | 0.32 | 0.86 | 0.95 | 7.14E-06 | 1.40E-04 | (A) NE (B/D) |
| JARID2 | 3.6 | 2.65 | 0.39 | 0.68 | 0.95 | 1.90E-05 | 2.97E-04 | (A) NE (B/D) |
| B4GALT5 | 6.84 | 5.88 | 0.38 | 0.77 | 0.95 | 1.96E-05 | 3.03E-04 | (A) NE (B/D) |
| HSPA4L | 2.3 | 1.35 | 0.43 | 0.65 | 0.95 | 4.45E-05 | 5.77E-04 | (A) NE (B/D) |
| MRPS11 | 4.9 | 3.95 | 0.36 | 0.97 | 0.95 | 4.81E-05 | 6.12E-04 | (A) NE (B/D) |
| SLC4A2 | 1.65 | 0.7 | 0.4 | 0.88 | 0.95 | 5.25E-05 | 6.57E-04 | (A) NE (B/D) |
| MBOAT7 | 2.59 | 1.64 | 0.43 | 0.73 | 0.95 | 5.84E-05 | 7.14E-04 | (A) NE (B/D) |
| PRR14 | 1.93 | 0.98 | 0.37 | 1.03 | 0.95 | 8.15E-05 | 9.26E-04 | (A) NE (B/D) |
| NUCB1 | 6.61 | 5.66 | 0.35 | 1.1 | 0.95 | 1.06E-04 | 1.13E-03 | (A) NE (B/D) |
| SEC11A | 3.69 | 2.75 | 0.36 | 1.09 | 0.95 | 1.18E-04 | 1.23E-03 | (A) NE (B/D) |
| ZNF512B | 3.09 | 2.14 | 0.4 | 1.06 | 0.95 | 1.67E-04 | 1.61E-03 | (A) NE (B/D) |
| STX16-NPEPL | 6.38 | 5.43 | 0.45 | 0.91 | 0.95 | 1.68E-04 | 1.61E-03 | (A) NE (B/D) |
| PHKG1 | 1.37 | 0.42 | 0.48 | 0.37 | 0.95 | 2.07E-04 | 1.89E-03 | (A) NE (B/D) |
| MIR-590/3P | 2.51 | 1.56 | 0.5 | 0.73 | 0.95 | 2.43E-04 | 2.14E-03 | (A) NE (B/D) |
| MIR-590/590 | 2.51 | 1.56 | 0.5 | 0.73 | 0.95 | 2.43E-04 | 2.14E-03 | (A) NE (B/D) |
| MIR-590/5P | 2.51 | 1.56 | 0.5 | 0.73 | 0.95 | 2.43E-04 | 2.14E-03 | (A) NE (B/D) |
| VASH1 | 2.81 | 1.86 | 0.42 | 1.14 | 0.95 | 3.52E-04 | 2.85E-03 | (A) NE (B/D) |
| MSH6 | 1.24 | 0.28 | 0.53 | 0.48 | 0.95 | 3.75E-04 | 3.00E-03 | (A) NE (B/D) |
| NT5C1B-RDH | 4.61 | 3.66 | 0.53 | 0.74 | 0.95 | 4.17E-04 | 3.25E-03 | (A) NE (B/D) |
| EWSR1 | 4.28 | 3.33 | 0.25 | 1.36 | 0.95 | 4.51E-04 | 3.45E-03 | (A) NE (B/D) |
| FAM3C2 | 9.07 | 8.12 | 0.46 | 1.17 | 0.95 | 5.94E-04 | 4.28E-03 | (A) NE (B/D) |
| TMC2 | 0.96 | 0.01 | 0.55 | 0.05 | 0.95 | 8.11E-04 | 5.42E-03 | (A) NE (B/D) |
| RPS27A | 4.12 | 3.16 | 0.33 | 1.41 | 0.95 | 8.51E-04 | 5.61E-03 | (A) NE (B/D) |
| SEPTIN9 | 4.97 | 4.01 | 0.56 | 1.02 | 0.95 | 1.01E-03 | 6.35E-03 | (A) NE (B/D) |
| INA | 0.96 | 0.01 | 0.57 | 0.04 | 0.95 | 1.04E-03 | 6.50E-03 | (A) NE (B/D) |
| NSUN5P2 | 1.28 | 0.33 | 0.63 | 0.51 | 0.95 | 1.55E-03 | 8.86E-03 | (A) NE (B/D) |
| SLX1A-SULT1 | 8.1 | 7.15 | 0.59 | 1.07 | 0.95 | 1.63E-03 | 9.17E-03 | (A) NE (B/D) |
| MIR-4658/46 | 7.54 | 6.59 | 0.46 | 1.41 | 0.95 | 1.98E-03 | 0.0106 | (A) NE (B/D) |
| SEM1 | 1.8 | 0.86 | 0.63 | 0.95 | 0.95 | 2.09E-03 | 0.0111 | (A) NE (B/D) |
| PGD | 7.28 | 6.33 | 0.52 | 1.32 | 0.95 | 2.23E-03 | 0.0116 | (A) NE (B/D) |
| LIPT1 | 3.29 | 2.34 | 0.63 | 1.05 | 0.95 | 2.51E-03 | 0.0128 | (A) NE (B/D) |
| TAF1A | 3.39 | 2.44 | 0.7 | 0.77 | 0.95 | 3.31E-03 | 0.0157 | (A) NE (B/D) |
| NDUFA6 | 7.42 | 6.47 | 0.59 | 1.39 | 0.95 | 4.26E-03 | 0.0191 | (A) NE (B/D) |
| TP53BP1 | 1.12 | 0.18 | 0.74 | 0.3 | 0.95 | 4.71E-03 | 0.0206 | (A) NE (B/D) |
| GMPSP1 | 3.36 | 2.42 | 0.72 | 1.23 | 0.95 | 7.28E-03 | 0.0283 | (A) NE (B/D) |
| VEGFD | 1.86 | 0.91 | 0.81 | 0.77 | 0.95 | 8.49E-03 | 0.0316 | (A) NE (B/D) |
| HOXB3 | 0.99 | 0.04 | 0.83 | 0.2 | 0.95 | 9.00E-03 | 0.0331 | (A) NE (B/D) |
| HPCAL1 | 4.42 | 3.47 | 0.83 | 0.83 | 0.95 | 9.86E-03 | 0.0353 | (A) NE (B/D) |
| CNIH1 | 3.27 | 2.32 | 0.73 | 1.43 | 0.95 | 0.0108 | 0.0376 | (A) NE (B/D) |
| TNFRSF6B | 3.04 | 2.09 | 0.83 | 1.09 | 0.95 | 0.0116 | 0.0398 | (A) NE (B/D) |
| PPP5D1 | 1.14 | 0.19 | 0.89 | 0.37 | 0.95 | 0.0128 | 0.0428 | (A) NE (B/D) |
| IDS | 2.21 | 1.27 | 0.88 | 0.99 | 0.95 | 0.0144 | 0.0465 | (A) NE (B/D) |
| FUT4 | 4.14 | 3.19 | 0.87 | 1.15 | 0.95 | 0.0146 | 0.047 | (A) NE (B/D) |
| PIP5K1C | 1.56 | 0.61 | 0.23 | 0.47 | 0.94 | 3.64E-09 | 3.67E-07 | (A) NE (B/D) |
| DDX39B-AS1 | 3.02 | 2.08 | 0.21 | 1.01 | 0.94 | 1.06E-05 | 1.91E-04 | (A) NE (B/D) |
| CENPP | 2.1 | 1.16 | 0.4 | 0.76 | 0.94 | 3.11E-05 | 4.36E-04 | (A) NE (B/D) |
| GMEB2 | 1.88 | 0.94 | 0.42 | 0.76 | 0.94 | 5.77E-05 | 7.08E-04 | (A) NE (B/D) |
| TRMT2B | 3.72 | 2.78 | 0.41 | 0.91 | 0.94 | 8.17E-05 | 9.28E-04 | (A) NE (B/D) |
| SURF6 | 9.36 | 8.42 | 0.39 | 1.07 | 0.94 | 1.53E-04 | 1.50E-03 | (A) NE (B/D) |
| CCNB1IP1 | 1.52 | 0.58 | 0.48 | 0.56 | 0.94 | 1.68E-04 | 1.62E-03 | (A) NE (B/D) |
| ERCC1 | 1.46 | 0.52 | 0.5 | 0.68 | 0.94 | 2.43E-04 | 2.14E-03 | (A) NE (B/D) |
| MRPL48 | 1.71 | 0.78 | 0.47 | 0.9 | 0.94 | 2.46E-04 | 2.16E-03 | (A) NE (B/D) |
| DROSHA | 2.47 | 1.53 | 0.38 | 1.15 | 0.94 | 2.70E-04 | 2.32E-03 | (A) NE (B/D) |
| GLE1 | 4.18 | 3.24 | 0.46 | 1 | 0.94 | 2.94E-04 | 2.48E-03 | (A) NE (B/D) |
| EIF4A2 | 1.96 | 1.03 | 0.49 | 0.88 | 0.94 | 3.06E-04 | 2.56E-03 | (A) NE (B/D) |
| TPGS2 | 7.45 | 6.52 | 0.32 | 1.24 | 0.94 | 3.22E-04 | 2.66E-03 | (A) NE (B/D) |
| LRRFIP1 | 3.23 | 2.29 | 0.51 | 0.5 | 0.94 | 3.35E-04 | 2.74E-03 | (A) NE (B/D) |
| TMEM69 | 6.63 | 5.68 | 0.46 | 1.11 | 0.94 | 5.35E-04 | 3.93E-03 | (A) NE (B/D) |
| SRSF6 | 6.32 | 5.38 | 0.42 | 1.28 | 0.94 | 8.65E-04 | 5.67E-03 | (A) NE (B/D) |
| NOC2L | 2.01 | 1.06 | 0.56 | 0.93 | 0.94 | 9.85E-04 | 6.24E-03 | (A) NE (B/D) |
| ZNF768 | 8.99 | 8.05 | 0.55 | 1.05 | 0.94 | 1.16E-03 | 7.07E-03 | (A) NE (B/D) |

| | | | | | | | | |
|---|---|---|---|---|---|---|---|---|
| FARP1 | 1.46 | 0.51 | 0.6 | 0.59 | 0.94 | 1.17E-03 | 7.14E-03 | (A) NE (B/D) |
| SRSF9P1 | 5.17 | 4.23 | 0.56 | 1.09 | 0.94 | 1.44E-03 | 8.33E-03 | (A) NE (B/D) |
| MIF4GD | 1.78 | 0.85 | 0.59 | 0.95 | 0.94 | 1.47E-03 | 8.51E-03 | (A) NE (B/D) |
| ZBTB9 | 6.3 | 5.36 | 0.52 | 1.23 | 0.94 | 1.72E-03 | 9.55E-03 | (A) NE (B/D) |
| EEF1DP5 | 1.01 | 0.07 | 0.68 | 0.42 | 0.94 | 2.80E-03 | 0.0139 | (A) NE (B/D) |
| KBTBD2 | 3.33 | 2.39 | 0.66 | 0.88 | 0.94 | 2.84E-03 | 0.014 | (A) NE (B/D) |
| UBTD2 | 6.84 | 5.9 | 0.67 | 1.05 | 0.94 | 3.95E-03 | 0.018 | (A) NE (B/D) |
| RN7SL130P | 1.58 | 0.64 | 0.72 | 0.48 | 0.94 | 4.36E-03 | 0.0194 | (A) NE (B/D) |
| FSIP2 | 1.1 | 0.16 | 0.83 | 0.34 | 0.94 | 9.82E-03 | 0.0352 | (A) NE (B/D) |
| ZNF329 | 4.47 | 3.53 | 0.83 | 0.86 | 0.94 | 0.01 | 0.0357 | (A) NE (B/D) |
| CIAO2B | 5.2 | 4.26 | 0.77 | 1.35 | 0.94 | 0.0122 | 0.0412 | (A) NE (B/D) |
| RPL4P5 | 3.73 | 2.79 | 0.84 | 1.17 | 0.94 | 0.0139 | 0.0454 | (A) NE (B/D) |
| MIR-3655/36 | 5.42 | 4.49 | 0.88 | 0.91 | 0.94 | 0.0145 | 0.0468 | (A) NE (B/D) |
| LCTL | 2.14 | 1.2 | 0.9 | 0.81 | 0.94 | 0.015 | 0.048 | (A) NE (B/D) |
| NPTXR | 1.09 | 0.16 | 0.15 | 0.34 | 0.93 | 4.17E-13 | 4.12E-10 | (A) NE (B/D) |
| SIRT6 | 1.15 | 0.22 | 0.18 | 0.32 | 0.93 | 4.65E-11 | 1.25E-08 | (A) NE (B/D) |
| INTS4 | 7.57 | 6.64 | 0.14 | 0.64 | 0.93 | 1.54E-09 | 1.89E-07 | (A) NE (B/D) |
| MAPKAPK5-A | 5.42 | 4.5 | 0.22 | 0.75 | 0.93 | 1.99E-07 | 8.33E-06 | (A) NE (B/D) |
| SMG5 | 4.49 | 3.56 | 0.27 | 0.84 | 0.93 | 2.59E-06 | 6.27E-05 | (A) NE (B/D) |
| TBC1D22B | 4.93 | 4 | 0.22 | 0.96 | 0.93 | 7.15E-06 | 1.40E-04 | (A) NE (B/D) |
| CTDNEP1 | 2.05 | 1.12 | 0.19 | 1.03 | 0.93 | 1.43E-05 | 2.42E-04 | (A) NE (B/D) |
| MMP24OS | 5.81 | 4.88 | 0.36 | 0.91 | 0.93 | 4.33E-05 | 5.65E-04 | (A) NE (B/D) |
| FTSJ3 | 5.65 | 4.72 | 0.14 | 1.17 | 0.93 | 6.16E-05 | 7.44E-04 | (A) NE (B/D) |
| HERC2P9 | 3 | 2.07 | 0.43 | 0.71 | 0.93 | 7.48E-05 | 8.68E-04 | (A) NE (B/D) |
| C6ORF89 | 3.35 | 2.42 | 0.2 | 1.19 | 0.93 | 1.00E-04 | 1.09E-03 | (A) NE (B/D) |
| ITGB1BP1 | 3.35 | 2.42 | 0.39 | 0.98 | 0.93 | 1.11E-04 | 1.17E-03 | (A) NE (B/D) |
| ARAP1-AS2 | 6.81 | 5.88 | 0.44 | 0.82 | 0.93 | 1.15E-04 | 1.20E-03 | (A) NE (B/D) |
| PPIL3 | 6.74 | 5.81 | 0.34 | 1.1 | 0.93 | 1.31E-04 | 1.34E-03 | (A) NE (B/D) |
| AAAS | 2.32 | 1.38 | 0.34 | 1.11 | 0.93 | 1.45E-04 | 1.45E-03 | (A) NE (B/D) |
| OAZ2 | 2.51 | 1.58 | 0.41 | 0.97 | 0.93 | 1.46E-04 | 1.45E-03 | (A) NE (B/D) |
| NDUFS2 | 3.19 | 2.26 | 0.29 | 1.16 | 0.93 | 1.52E-04 | 1.49E-03 | (A) NE (B/D) |
| SPIN1 | 8.21 | 7.29 | 0.42 | 0.98 | 0.93 | 1.99E-04 | 1.83E-03 | (A) NE (B/D) |
| JPT1 | 1.86 | 0.93 | 0.45 | 0.98 | 0.93 | 2.65E-04 | 2.28E-03 | (A) NE (B/D) |
| STMP1 | 6.24 | 5.32 | 0.36 | 1.24 | 0.93 | 4.94E-04 | 3.71E-03 | (A) NE (B/D) |
| RBMXP4 | 3.86 | 2.93 | 0.48 | 1.06 | 0.93 | 6.06E-04 | 4.34E-03 | (A) NE (B/D) |
| RSAD1 | 6.46 | 5.53 | 0.34 | 1.3 | 0.93 | 6.12E-04 | 4.37E-03 | (A) NE (B/D) |
| ARRDC1 | 6.31 | 5.38 | 0.52 | 0.98 | 0.93 | 7.19E-04 | 4.95E-03 | (A) NE (B/D) |
| RRP36 | 7.94 | 7.02 | 0.53 | 0.94 | 0.93 | 7.71E-04 | 5.21E-03 | (A) NE (B/D) |
| MT1P1 | 1.87 | 0.95 | 0.56 | 0.89 | 0.93 | 9.66E-04 | 6.16E-03 | (A) NE (B/D) |
| NUBP2 | 2.08 | 1.15 | 0.57 | 0.84 | 0.93 | 1.08E-03 | 6.71E-03 | (A) NE (B/D) |
| KBTBD4 | 5.93 | 5 | 0.55 | 1.05 | 0.93 | 1.34E-03 | 7.90E-03 | (A) NE (B/D) |
| SLC25A28 | 8.46 | 7.53 | 0.3 | 1.46 | 0.93 | 1.41E-03 | 8.18E-03 | (A) NE (B/D) |
| DGKZP1 | 1.54 | 0.61 | 0.61 | 0.76 | 0.93 | 1.51E-03 | 8.65E-03 | (A) NE (B/D) |
| CFAP36 | 1.39 | 0.46 | 0.64 | 0.69 | 0.93 | 1.95E-03 | 0.0105 | (A) NE (B/D) |
| SNHG29 | 7.23 | 6.3 | 0.31 | 1.54 | 0.93 | 2.25E-03 | 0.0117 | (A) NE (B/D) |
| PLK1 | 1.12 | 0.19 | 0.69 | 0.48 | 0.93 | 3.30E-03 | 0.0157 | (A) NE (B/D) |
| RNF113A | 5.16 | 4.23 | 0.46 | 1.49 | 0.93 | 3.47E-03 | 0.0163 | (A) NE (B/D) |
| HSFX1 | 1.39 | 0.47 | 0.69 | 0.47 | 0.93 | 3.51E-03 | 0.0164 | (A) NE (B/D) |
| DNAJC28 | 1.38 | 0.45 | 0.74 | 0.48 | 0.93 | 5.06E-03 | 0.0217 | (A) NE (B/D) |
| CORO7-PAM | 11.51 | 10.58 | 0.71 | 0.98 | 0.93 | 5.17E-03 | 0.0221 | (A) NE (B/D) |
| TOMM6 | 10.09 | 9.16 | 0.46 | 1.62 | 0.93 | 5.48E-03 | 0.023 | (A) NE (B/D) |
| ATN1 | 5.39 | 4.46 | 0.53 | 1.53 | 0.93 | 5.51E-03 | 0.0231 | (A) NE (B/D) |
| SNORD34 | 4.79 | 3.86 | 0.63 | 1.37 | 0.93 | 5.77E-03 | 0.0239 | (A) NE (B/D) |
| CYCS | 4.35 | 3.42 | 0.71 | 1.27 | 0.93 | 8.22E-03 | 0.0309 | (A) NE (B/D) |
| SERPINB6 | 3.62 | 2.69 | 0.59 | 1.6 | 0.93 | 9.15E-03 | 0.0335 | (A) NE (B/D) |
| C8ORF82 | 2.2 | 1.27 | 0.79 | 1.11 | 0.93 | 0.0106 | 0.0372 | (A) NE (B/D) |
| CALCOCO1 | 8.92 | 7.99 | 0.82 | 1.04 | 0.93 | 0.0116 | 0.0396 | (A) NE (B/D) |
| SNAPC3 | 4.05 | 3.12 | 0.88 | 0.74 | 0.93 | 0.014 | 0.0455 | (A) NE (B/D) |
| CCDC114 | 1.27 | 0.34 | 0.88 | 0.76 | 0.93 | 0.0143 | 0.0464 | (A) NE (B/D) |
| SLC9A6 | 3.45 | 2.51 | 0.87 | 0.93 | 0.93 | 0.0146 | 0.0469 | (A) NE (B/D) |
| RRNAD1 | 1.49 | 0.57 | 0.21 | 0.71 | 0.92 | 6.49E-08 | 3.39E-06 | (A) NE (B/D) |
| STK11 | 2.43 | 1.51 | 0.23 | 0.78 | 0.92 | 4.69E-07 | 1.64E-05 | (A) NE (B/D) |
| TOP3A | 2.05 | 1.13 | 0.34 | 0.64 | 0.92 | 3.94E-06 | 8.76E-05 | (A) NE (B/D) |
| LIME1 | 1.34 | 0.41 | 0.38 | 0.56 | 0.92 | 1.44E-05 | 2.43E-04 | (A) NE (B/D) |
| ASB6 | 1.52 | 0.6 | 0.44 | 0.63 | 0.92 | 7.89E-05 | 9.00E-04 | (A) NE (B/D) |

| | | | | | | | | |
|---|---|---|---|---|---|---|---|---|
| MIR570HG | 4.42 | 3.5 | 0.46 | 0.65 | 0.92 | 1.36E-04 | 1.37E-03 | (A) NE (B/D) |
| RNPS1 | 8.18 | 7.26 | 0.43 | 0.99 | 0.92 | 2.30E-04 | 2.05E-03 | (A) NE (B/D) |
| MRPL43 | 2 | 1.08 | 0.35 | 1.15 | 0.92 | 2.36E-04 | 2.09E-03 | (A) NE (B/D) |
| DDX19B | 2.48 | 1.56 | 0.49 | 0.68 | 0.92 | 2.74E-04 | 2.34E-03 | (A) NE (B/D) |
| NOP16 | 5.58 | 4.67 | 0.27 | 1.26 | 0.92 | 3.46E-04 | 2.81E-03 | (A) NE (B/D) |
| YPEL3 | 1.56 | 0.63 | 0.5 | 0.9 | 0.92 | 4.99E-04 | 3.74E-03 | (A) NE (B/D) |
| ZNF396 | 2.86 | 1.95 | 0.5 | 0.92 | 0.92 | 5.07E-04 | 3.78E-03 | (A) NE (B/D) |
| UBOX5-AS1 | 1.57 | 0.66 | 0.56 | 0.59 | 0.92 | 8.17E-04 | 5.45E-03 | (A) NE (B/D) |
| PIN4 | 2.65 | 1.73 | 0.22 | 1.45 | 0.92 | 1.06E-03 | 6.59E-03 | (A) NE (B/D) |
| MIR497HG | 3.71 | 2.79 | 0.38 | 1.36 | 0.92 | 1.23E-03 | 7.41E-03 | (A) NE (B/D) |
| C6ORF47 | 5.9 | 4.98 | 0.56 | 0.99 | 0.92 | 1.39E-03 | 8.09E-03 | (A) NE (B/D) |
| NAA20 | 3.72 | 2.8 | 0.51 | 1.2 | 0.92 | 1.63E-03 | 9.20E-03 | (A) NE (B/D) |
| LAMTOR1 | 7.17 | 6.25 | 0.45 | 1.33 | 0.92 | 1.69E-03 | 9.44E-03 | (A) NE (B/D) |
| ZNF22 | 2.9 | 1.98 | 0.48 | 1.37 | 0.92 | 2.57E-03 | 0.013 | (A) NE (B/D) |
| MCTS1 | 7.25 | 6.33 | 0.48 | 1.39 | 0.92 | 2.82E-03 | 0.0139 | (A) NE (B/D) |
| DVL1 | 10.06 | 9.15 | 0.66 | 0.84 | 0.92 | 3.12E-03 | 0.015 | (A) NE (B/D) |
| WDR13 | 1.52 | 0.6 | 0.69 | 0.76 | 0.92 | 3.80E-03 | 0.0175 | (A) NE (B/D) |
| AGPAT2 | 6.01 | 5.09 | 0.69 | 1.04 | 0.92 | 5.27E-03 | 0.0223 | (A) NE (B/D) |
| PTPRS | 1.15 | 0.22 | 0.77 | 0.28 | 0.92 | 7.06E-03 | 0.0277 | (A) NE (B/D) |
| RNA5SP162 | 1.49 | 0.57 | 0.75 | 0.94 | 0.92 | 7.28E-03 | 0.0284 | (A) NE (B/D) |
| HPF1 | 4.47 | 3.55 | 0.56 | 1.56 | 0.92 | 7.59E-03 | 0.0292 | (A) NE (B/D) |
| PECR | 4.54 | 3.62 | 0.76 | 0.98 | 0.92 | 7.87E-03 | 0.0299 | (A) NE (B/D) |
| ATP5IF1 | 2.03 | 1.11 | 0.76 | 1.01 | 0.92 | 8.15E-03 | 0.0307 | (A) NE (B/D) |
| OTX2-AS1 | 1.11 | 0.18 | 0.83 | 0.42 | 0.92 | 0.01 | 0.0357 | (A) NE (B/D) |
| OCIAD1-AS1 | 1.17 | 0.25 | 0.83 | 0.31 | 0.92 | 0.0103 | 0.0364 | (A) NE (B/D) |
| RNA5SP82 | 1.21 | 0.29 | 0.83 | 0.52 | 0.92 | 0.0107 | 0.0374 | (A) NE (B/D) |
| LTC4S | 1.34 | 0.42 | 0.84 | 0.48 | 0.92 | 0.011 | 0.0381 | (A) NE (B/D) |
| ZNF404 | 0.95 | 0.03 | 0.85 | 0.12 | 0.92 | 0.0118 | 0.0402 | (A) NE (B/D) |
| SGPP2 | 1 | 0.08 | 0.87 | 0.33 | 0.92 | 0.0137 | 0.0449 | (A) NE (B/D) |
| UBE2A | 7.68 | 6.75 | 0.77 | 1.36 | 0.92 | 0.0142 | 0.0461 | (A) NE (B/D) |
| HSPA13 | 2.89 | 1.97 | 0.87 | 0.9 | 0.92 | 0.0147 | 0.0473 | (A) NE (B/D) |
| SBF1 | 4.31 | 3.39 | 0.19 | 0.63 | 0.91 | 6.00E-09 | 5.37E-07 | (A) NE (B/D) |
| TRAPPC10 | 6.51 | 5.6 | 0.33 | 0.62 | 0.91 | 3.52E-06 | 8.03E-05 | (A) NE (B/D) |
| PACS2 | 1.57 | 0.66 | 0.35 | 0.45 | 0.91 | 8.05E-06 | 1.54E-04 | (A) NE (B/D) |
| ARHGAP35 | 10.09 | 9.17 | 0.37 | 0.65 | 0.91 | 1.52E-05 | 2.52E-04 | (A) NE (B/D) |
| LRPAP1 | 3.09 | 2.17 | 0.38 | 0.63 | 0.91 | 1.71E-05 | 2.76E-04 | (A) NE (B/D) |
| TARS2 | 3.19 | 2.28 | 0.33 | 0.89 | 0.91 | 2.19E-05 | 3.28E-04 | (A) NE (B/D) |
| TGFBRAP1 | 1.67 | 0.77 | 0.36 | 0.85 | 0.91 | 3.42E-05 | 4.69E-04 | (A) NE (B/D) |
| PIK3R2 | 1.66 | 0.75 | 0.4 | 0.44 | 0.91 | 3.80E-05 | 5.11E-04 | (A) NE (B/D) |
| BPNT1 | 5.2 | 4.3 | 0.36 | 1.01 | 0.91 | 1.09E-04 | 1.16E-03 | (A) NE (B/D) |
| MSANTD3 | 3.78 | 2.86 | 0.42 | 0.89 | 0.91 | 1.33E-04 | 1.35E-03 | (A) NE (B/D) |
| HYOU1 | 6.76 | 5.85 | 0.37 | 1.1 | 0.91 | 2.49E-04 | 2.17E-03 | (A) NE (B/D) |
| TPRA1 | 3.44 | 2.53 | 0.35 | 1.17 | 0.91 | 2.98E-04 | 2.51E-03 | (A) NE (B/D) |
| DCDC2B | 3.7 | 2.79 | 0.49 | 0.8 | 0.91 | 3.36E-04 | 2.75E-03 | (A) NE (B/D) |
| UBXN6 | 4.52 | 3.61 | 0.51 | 1.14 | 0.91 | 1.36E-03 | 7.99E-03 | (A) NE (B/D) |
| FAM27E3 | 1.9 | 0.98 | 0.6 | 0.88 | 0.91 | 1.87E-03 | 0.0102 | (A) NE (B/D) |
| HNRNPCP2 | 8.14 | 7.23 | 0.45 | 1.32 | 0.91 | 1.88E-03 | 0.0102 | (A) NE (B/D) |
| PIGY | 5.93 | 5.02 | 0.4 | 1.41 | 0.91 | 2.02E-03 | 0.0108 | (A) NE (B/D) |
| CBX7 | 2.59 | 1.69 | 0.44 | 1.37 | 0.91 | 2.23E-03 | 0.0116 | (A) NE (B/D) |
| IDH3A | 1.24 | 0.33 | 0.64 | 0.42 | 0.91 | 2.46E-03 | 0.0126 | (A) NE (B/D) |
| FAM156A | 3.2 | 2.29 | 0.46 | 1.37 | 0.91 | 2.47E-03 | 0.0126 | (A) NE (B/D) |
| ELOF1 | 1.12 | 0.21 | 0.66 | 0.39 | 0.91 | 2.99E-03 | 0.0146 | (A) NE (B/D) |
| WDSUB1 | 2.86 | 1.95 | 0.7 | 0.78 | 0.91 | 4.51E-03 | 0.0199 | (A) NE (B/D) |
| SMCO2 | 1.04 | 0.13 | 0.74 | 0.34 | 0.91 | 5.91E-03 | 0.0244 | (A) NE (B/D) |
| METTL21A | 5.86 | 4.96 | 0.78 | 0.75 | 0.91 | 8.45E-03 | 0.0315 | (A) NE (B/D) |
| XPR1 | 1.97 | 1.06 | 0.82 | 0.83 | 0.91 | 0.0114 | 0.0392 | (A) NE (B/D) |
| SNORA18 | 4.39 | 3.48 | 0.82 | 0.92 | 0.91 | 0.0118 | 0.0403 | (A) NE (B/D) |
| MIR-4691/3P | 3.13 | 2.22 | 0.75 | 1.27 | 0.91 | 0.0124 | 0.0418 | (A) NE (B/D) |
| MIR-4691/5P | 3.13 | 2.22 | 0.75 | 1.27 | 0.91 | 0.0124 | 0.0418 | (A) NE (B/D) |
| CACYBPP3 | 1.62 | 0.71 | 0.82 | 0.97 | 0.91 | 0.0127 | 0.0424 | (A) NE (B/D) |
| OR8A2P | 1.26 | 0.35 | 0.85 | 0.87 | 0.91 | 0.0137 | 0.0448 | (A) NE (B/D) |
| FRS3 | 1.31 | 0.41 | 0.37 | 0.59 | 0.9 | 1.40E-05 | 2.38E-04 | (A) NE (B/D) |
| POMZP3 | 2.95 | 2.05 | 0.38 | 0.62 | 0.9 | 2.02E-05 | 3.10E-04 | (A) NE (B/D) |
| LMTK2 | 3.61 | 2.71 | 0.4 | 0.56 | 0.9 | 3.67E-05 | 4.96E-04 | (A) NE (B/D) |
| ATP9B | 3.21 | 2.32 | 0.4 | 0.63 | 0.9 | 4.71E-05 | 6.02E-04 | (A) NE (B/D) |

| | | | | | | | | |
|---|---|---|---|---|---|---|---|---|
| NELFA | 2.83 | 1.94 | 0.42 | 0.66 | 0.9 | 7.07E-05 | 8.28E-04 | (A) NE (B/D) |
| PHF13 | 3.56 | 2.66 | 0.36 | 0.95 | 0.9 | 8.72E-05 | 9.79E-04 | (A) NE (B/D) |
| MTMR4 | 2.02 | 1.12 | 0.42 | 0.98 | 0.9 | 2.60E-04 | 2.25E-03 | (A) NE (B/D) |
| PHLDB1 | 1.76 | 0.86 | 0.46 | 0.85 | 0.9 | 2.83E-04 | 2.40E-03 | (A) NE (B/D) |
| NXF1 | 3.41 | 2.51 | 0.27 | 1.31 | 0.9 | 6.09E-04 | 4.35E-03 | (A) NE (B/D) |
| ATP13A2 | 2.67 | 1.77 | 0.49 | 0.95 | 0.9 | 6.33E-04 | 4.48E-03 | (A) NE (B/D) |
| ABHD14A | 2.57 | 1.67 | 0.52 | 1.09 | 0.9 | 1.36E-03 | 7.98E-03 | (A) NE (B/D) |
| UBXN11 | 3.64 | 2.74 | 0.22 | 1.5 | 0.9 | 1.78E-03 | 9.80E-03 | (A) NE (B/D) |
| RYR2 | 1.13 | 0.23 | 0.62 | 0.53 | 0.9 | 2.03E-03 | 0.0109 | (A) NE (B/D) |
| TRA2B | 2.88 | 1.98 | 0.6 | 0.86 | 0.9 | 2.05E-03 | 0.0109 | (A) NE (B/D) |
| CCNQP3 | 2.55 | 1.65 | 0.54 | 1.13 | 0.9 | 2.14E-03 | 0.0113 | (A) NE (B/D) |
| UQCRFS1 | 3.88 | 2.98 | 0.38 | 1.43 | 0.9 | 2.17E-03 | 0.0113 | (A) NE (B/D) |
| EEF1A1P5 | 13.67 | 12.77 | 0.33 | 1.48 | 0.9 | 2.28E-03 | 0.0118 | (A) NE (B/D) |
| AMZ2 | 7.98 | 7.08 | 0.44 | 1.37 | 0.9 | 2.36E-03 | 0.0121 | (A) NE (B/D) |
| NAXE | 2.52 | 1.63 | 0.59 | 0.99 | 0.9 | 2.38E-03 | 0.0122 | (A) NE (B/D) |
| LINC00339 | 7.26 | 6.37 | 0.44 | 1.45 | 0.9 | 3.54E-03 | 0.0166 | (A) NE (B/D) |
| HAX1 | 2.58 | 1.68 | 0.51 | 1.39 | 0.9 | 4.15E-03 | 0.0187 | (A) NE (B/D) |
| MVB12B | 6.33 | 5.44 | 0.7 | 0.89 | 0.9 | 5.73E-03 | 0.0238 | (A) NE (B/D) |
| CPTP | 1.64 | 0.74 | 0.77 | 0.73 | 0.9 | 8.26E-03 | 0.031 | (A) NE (B/D) |
| MROH7-TTC4 | 1.04 | 0.14 | 0.8 | 0.31 | 0.9 | 9.42E-03 | 0.0342 | (A) NE (B/D) |
| HSPA8P11 | 1.32 | 0.41 | 0.83 | 0.54 | 0.9 | 0.0114 | 0.039 | (A) NE (B/D) |
| DDT | 2.73 | 1.83 | 0.8 | 0.95 | 0.9 | 0.0123 | 0.0415 | (A) NE (B/D) |
| NFKBIB | 1.75 | 0.85 | 0.81 | 0.92 | 0.9 | 0.0126 | 0.0422 | (A) NE (B/D) |
| STRIP1 | 1.61 | 0.72 | 0.21 | 0.74 | 0.89 | 3.20E-07 | 1.21E-05 | (A) NE (B/D) |
| PANX1 | 2.28 | 1.38 | 0.28 | 0.62 | 0.89 | 6.22E-07 | 2.04E-05 | (A) NE (B/D) |
| GEMIN7 | 1.46 | 0.56 | 0.33 | 0.56 | 0.89 | 3.26E-06 | 7.52E-05 | (A) NE (B/D) |
| FAM219A | 1.57 | 0.67 | 0.21 | 0.88 | 0.89 | 3.64E-06 | 8.25E-05 | (A) NE (B/D) |
| RNF216 | 4.74 | 3.85 | 0.39 | 0.56 | 0.89 | 3.25E-05 | 4.54E-04 | (A) NE (B/D) |
| DDX23 | 8.66 | 7.77 | 0.4 | 0.9 | 0.89 | 1.26E-04 | 1.30E-03 | (A) NE (B/D) |
| NICN1 | 3.36 | 2.48 | 0.3 | 1.11 | 0.89 | 1.68E-04 | 1.62E-03 | (A) NE (B/D) |
| RBBP4P4 | 1.96 | 1.08 | 0.46 | 0.7 | 0.89 | 2.28E-04 | 2.03E-03 | (A) NE (B/D) |
| NDUFV3 | 4.52 | 3.63 | 0.49 | 0.72 | 0.89 | 3.55E-04 | 2.87E-03 | (A) NE (B/D) |
| NUTF2 | 2.09 | 1.2 | 0.5 | 0.99 | 0.89 | 8.69E-04 | 5.69E-03 | (A) NE (B/D) |
| RHOQP1 | 6.63 | 5.73 | 0.57 | 0.76 | 0.89 | 1.23E-03 | 7.39E-03 | (A) NE (B/D) |
| TNK1 | 2.73 | 1.84 | 0.44 | 1.23 | 0.89 | 1.49E-03 | 8.59E-03 | (A) NE (B/D) |
| SNORA40 | 5.51 | 4.62 | 0.6 | 0.82 | 0.89 | 2.03E-03 | 0.0109 | (A) NE (B/D) |
| C8ORF44 | 2.9 | 2.01 | 0.62 | 0.66 | 0.89 | 2.38E-03 | 0.0122 | (A) NE (B/D) |
| KDM1B | 2.03 | 1.13 | 0.64 | 0.75 | 0.89 | 2.74E-03 | 0.0137 | (A) NE (B/D) |
| MIR-1825/18 | 9.66 | 8.77 | 0.44 | 1.38 | 0.89 | 2.86E-03 | 0.0141 | (A) NE (B/D) |
| RPL26L1 | 2.76 | 1.87 | 0.19 | 1.61 | 0.89 | 3.26E-03 | 0.0156 | (A) NE (B/D) |
| LETM2 | 1 | 0.11 | 0.68 | 0.33 | 0.89 | 4.37E-03 | 0.0194 | (A) NE (B/D) |
| RPL4P4 | 8.33 | 7.44 | 0.57 | 1.29 | 0.89 | 4.51E-03 | 0.0199 | (A) NE (B/D) |
| AHRR | 3.1 | 2.2 | 0.58 | 1.29 | 0.89 | 4.92E-03 | 0.0212 | (A) NE (B/D) |
| MIER1 | 1.18 | 0.29 | 0.7 | 0.44 | 0.89 | 5.03E-03 | 0.0216 | (A) NE (B/D) |
| KLHDC4 | 4.61 | 3.72 | 0.72 | 0.65 | 0.89 | 5.59E-03 | 0.0233 | (A) NE (B/D) |
| ZIK1 | 0.92 | 0.03 | 0.72 | 0.12 | 0.89 | 5.90E-03 | 0.0243 | (A) NE (B/D) |
| MIR-1226/12 | 2.51 | 1.62 | 0.67 | 1.08 | 0.89 | 5.99E-03 | 0.0246 | (A) NE (B/D) |
| MIR-1226/12 | 2.51 | 1.62 | 0.67 | 1.08 | 0.89 | 5.99E-03 | 0.0246 | (A) NE (B/D) |
| MIR-1226/3P | 2.51 | 1.62 | 0.67 | 1.08 | 0.89 | 5.99E-03 | 0.0246 | (A) NE (B/D) |
| MIR-1226/5P | 2.51 | 1.62 | 0.67 | 1.08 | 0.89 | 5.99E-03 | 0.0246 | (A) NE (B/D) |
| THOC6 | 6.22 | 5.33 | 0.65 | 1.22 | 0.89 | 6.79E-03 | 0.027 | (A) NE (B/D) |
| SUPT16H | 1.61 | 0.72 | 0.75 | 0.77 | 0.89 | 7.68E-03 | 0.0294 | (A) NE (B/D) |
| UBXN1 | 4.53 | 3.64 | 0.7 | 1.2 | 0.89 | 8.75E-03 | 0.0323 | (A) NE (B/D) |
| PPP1R21 | 5.12 | 4.23 | 0.8 | 0.74 | 0.89 | 0.0109 | 0.0378 | (A) NE (B/D) |
| MIR-431/3P | 0.97 | 0.09 | 0.82 | 0.32 | 0.89 | 0.0115 | 0.0394 | (A) NE (B/D) |
| MIR-431/431 | 0.97 | 0.09 | 0.82 | 0.32 | 0.89 | 0.0115 | 0.0394 | (A) NE (B/D) |
| MIR-431/431 | 0.97 | 0.09 | 0.82 | 0.32 | 0.89 | 0.0115 | 0.0394 | (A) NE (B/D) |
| MIR-431/5P | 0.97 | 0.09 | 0.82 | 0.32 | 0.89 | 0.0115 | 0.0394 | (A) NE (B/D) |
| YY2 | 3.66 | 2.77 | 0.67 | 1.45 | 0.89 | 0.0121 | 0.041 | (A) NE (B/D) |
| CCDC13-AS1 | 1.73 | 0.85 | 0.83 | 0.62 | 0.89 | 0.0129 | 0.043 | (A) NE (B/D) |
| TRAPPC6A | 2.6 | 1.72 | 0.64 | 1.55 | 0.89 | 0.0136 | 0.0445 | (A) NE (B/D) |
| FAM174A | 1.59 | 0.71 | 0.81 | 0.96 | 0.89 | 0.014 | 0.0456 | (A) NE (B/D) |
| MBD4 | 2.1 | 1.2 | 0.82 | 0.97 | 0.89 | 0.0142 | 0.046 | (A) NE (B/D) |
| TRPC4AP | 5.92 | 5.04 | 0.2 | 0.75 | 0.88 | 3.09E-07 | 1.18E-05 | (A) NE (B/D) |
| ANKMY1 | 3.68 | 2.8 | 0.25 | 0.66 | 0.88 | 4.20E-07 | 1.49E-05 | (A) NE (B/D) |

| | | | | | | | | |
|---|---|---|---|---|---|---|---|---|
| RAB8A | 1.37 | 0.49 | 0.3 | 0.55 | 0.88 | 1.12E-06 | 3.25E-05 | (A) NE (B/D) |
| TTLL4 | 7.33 | 6.46 | 0.32 | 0.65 | 0.88 | 4.40E-06 | 9.56E-05 | (A) NE (B/D) |
| TRABD | 4.22 | 3.34 | 0.31 | 0.86 | 0.88 | 2.23E-05 | 3.32E-04 | (A) NE (B/D) |
| GGA1 | 2.77 | 1.89 | 0.33 | 0.87 | 0.88 | 3.89E-05 | 5.18E-04 | (A) NE (B/D) |
| LAMTOR4 | 2.03 | 1.15 | 0.2 | 1.05 | 0.88 | 5.02E-05 | 6.33E-04 | (A) NE (B/D) |
| UBE2O | 1.73 | 0.85 | 0.36 | 0.92 | 0.88 | 8.89E-05 | 9.93E-04 | (A) NE (B/D) |
| MPV17 | 2.03 | 1.15 | 0.33 | 1 | 0.88 | 1.08E-04 | 1.15E-03 | (A) NE (B/D) |
| OAZ1 | 4.7 | 3.82 | 0.13 | 1.16 | 0.88 | 1.18E-04 | 1.23E-03 | (A) NE (B/D) |
| PAIP1P1 | 4.97 | 4.09 | 0.45 | 0.69 | 0.88 | 1.61E-04 | 1.56E-03 | (A) NE (B/D) |
| ZBTB40 | 3.8 | 2.92 | 0.46 | 0.56 | 0.88 | 2.08E-04 | 1.89E-03 | (A) NE (B/D) |
| CYB5R3 | 3.26 | 2.38 | 0.24 | 1.28 | 0.88 | 5.03E-04 | 3.76E-03 | (A) NE (B/D) |
| OSER1 | 3.22 | 2.34 | 0.17 | 1.31 | 0.88 | 5.22E-04 | 3.86E-03 | (A) NE (B/D) |
| SWSAP1 | 2.01 | 1.13 | 0.46 | 0.96 | 0.88 | 5.52E-04 | 4.04E-03 | (A) NE (B/D) |
| SRA1 | 1.68 | 0.81 | 0.5 | 0.8 | 0.88 | 5.65E-04 | 4.11E-03 | (A) NE (B/D) |
| RGL2 | 6.04 | 5.16 | 0.34 | 1.25 | 0.88 | 8.22E-04 | 5.47E-03 | (A) NE (B/D) |
| CRELD2 | 2.05 | 1.17 | 0.51 | 0.9 | 0.88 | 8.54E-04 | 5.62E-03 | (A) NE (B/D) |
| WSB2 | 2.48 | 1.6 | 0.48 | 1.02 | 0.88 | 9.20E-04 | 5.94E-03 | (A) NE (B/D) |
| TOB2P1 | 1.73 | 0.85 | 0.58 | 0.71 | 0.88 | 1.57E-03 | 8.92E-03 | (A) NE (B/D) |
| SNRPA1 | 1.3 | 0.42 | 0.59 | 0.57 | 0.88 | 1.60E-03 | 9.05E-03 | (A) NE (B/D) |
| RPS19 | 6.04 | 5.15 | 0.37 | 1.36 | 0.88 | 1.79E-03 | 9.88E-03 | (A) NE (B/D) |
| LGMN | 2 | 1.13 | 0.56 | 1.2 | 0.88 | 3.58E-03 | 0.0167 | (A) NE (B/D) |
| MORF4L2-AS | 2.3 | 1.41 | 0.67 | 0.85 | 0.88 | 4.48E-03 | 0.0198 | (A) NE (B/D) |
| TTBK2 | 0.95 | 0.07 | 0.73 | 0.24 | 0.88 | 6.90E-03 | 0.0273 | (A) NE (B/D) |
| TBC1D24 | 1.65 | 0.77 | 0.74 | 0.55 | 0.88 | 6.98E-03 | 0.0275 | (A) NE (B/D) |
| C1ORF134 | 5 | 4.12 | 0.55 | 1.45 | 0.88 | 7.31E-03 | 0.0284 | (A) NE (B/D) |
| COMMD3 | 5.72 | 4.84 | 0.65 | 1.3 | 0.88 | 8.73E-03 | 0.0323 | (A) NE (B/D) |
| TIMM8B | 6.2 | 5.32 | 0.57 | 1.5 | 0.88 | 8.95E-03 | 0.0329 | (A) NE (B/D) |
| RPL9P8 | 10.31 | 9.43 | 0.34 | 1.76 | 0.88 | 9.20E-03 | 0.0336 | (A) NE (B/D) |
| ZNF326 | 5.25 | 4.37 | 0.78 | 0.82 | 0.88 | 0.0104 | 0.0368 | (A) NE (B/D) |
| DCTN4 | 7.24 | 6.36 | 0.83 | 0.73 | 0.88 | 0.0135 | 0.0444 | (A) NE (B/D) |
| PARP10 | 6.41 | 5.53 | 0.58 | 1.7 | 0.88 | 0.0158 | 0.0498 | (A) NE (B/D) |
| PDRG1 | 1.72 | 0.84 | 0.24 | 0.71 | 0.87 | 5.52E-07 | 1.87E-05 | (A) NE (B/D) |
| DDX56 | 1.31 | 0.44 | 0.35 | 0.6 | 0.87 | 1.17E-05 | 2.06E-04 | (A) NE (B/D) |
| CRTC2 | 1.66 | 0.79 | 0.34 | 0.74 | 0.87 | 1.46E-05 | 2.45E-04 | (A) NE (B/D) |
| PIGCP1 | 0.88 | 0.01 | 0.29 | 0.06 | 0.87 | 1.61E-05 | 2.63E-04 | (A) NE (B/D) |
| MED17 | 3.22 | 2.35 | 0.33 | 0.77 | 0.87 | 1.68E-05 | 2.72E-04 | (A) NE (B/D) |
| CDK16 | 1.4 | 0.53 | 0.37 | 0.58 | 0.87 | 2.23E-05 | 3.32E-04 | (A) NE (B/D) |
| FAXC | 1.44 | 0.57 | 0.32 | 0.83 | 0.87 | 2.24E-05 | 3.33E-04 | (A) NE (B/D) |
| HCG4P3 | 5.17 | 4.3 | 0.25 | 0.95 | 0.87 | 2.29E-05 | 3.40E-04 | (A) NE (B/D) |
| CRAMP1 | 1.98 | 1.11 | 0.39 | 0.74 | 0.87 | 6.48E-05 | 7.74E-04 | (A) NE (B/D) |
| YIF1B | 3.39 | 2.52 | 0.37 | 0.91 | 0.87 | 1.24E-04 | 1.28E-03 | (A) NE (B/D) |
| CXORF40A | 3.72 | 2.85 | 0.28 | 1.12 | 0.87 | 1.93E-04 | 1.79E-03 | (A) NE (B/D) |
| ZEB1-AS1 | 3.7 | 2.84 | 0.53 | 0.67 | 0.87 | 8.60E-04 | 5.64E-03 | (A) NE (B/D) |
| SAMD10 | 1.17 | 0.3 | 0.54 | 0.66 | 0.87 | 9.10E-04 | 5.89E-03 | (A) NE (B/D) |
| GNAQP1 | 6.16 | 5.29 | 0.54 | 0.66 | 0.87 | 9.13E-04 | 5.90E-03 | (A) NE (B/D) |
| PCBP2 | 9.53 | 8.66 | 0.44 | 1.15 | 0.87 | 1.12E-03 | 6.88E-03 | (A) NE (B/D) |
| CCDC22 | 6.27 | 5.41 | 0.48 | 1.09 | 0.87 | 1.29E-03 | 7.66E-03 | (A) NE (B/D) |
| TTLL3 | 2.49 | 1.62 | 0.46 | 1.18 | 0.87 | 1.65E-03 | 9.26E-03 | (A) NE (B/D) |
| PNPLA2 | 8.8 | 7.93 | 0.54 | 0.94 | 0.87 | 1.67E-03 | 9.34E-03 | (A) NE (B/D) |
| BCL9 | 1.66 | 0.8 | 0.55 | 0.94 | 0.87 | 1.84E-03 | 0.0101 | (A) NE (B/D) |
| TAZ | 1.11 | 0.24 | 0.6 | 0.4 | 0.87 | 2.16E-03 | 0.0113 | (A) NE (B/D) |
| RNU6-1223P | 3.42 | 2.56 | 0.45 | 1.29 | 0.87 | 2.52E-03 | 0.0128 | (A) NE (B/D) |
| IK | 6.39 | 5.53 | 0.49 | 1.24 | 0.87 | 2.85E-03 | 0.0141 | (A) NE (B/D) |
| EIF2B2 | 6.7 | 5.83 | 0.57 | 1.08 | 0.87 | 3.00E-03 | 0.0146 | (A) NE (B/D) |
| TMEM177 | 6.01 | 5.14 | 0.56 | 1.2 | 0.87 | 3.99E-03 | 0.0181 | (A) NE (B/D) |
| CERS6-AS1 | 5.98 | 5.11 | 0.56 | 1.19 | 0.87 | 4.06E-03 | 0.0184 | (A) NE (B/D) |
| PRPSAP2 | 1.78 | 0.91 | 0.64 | 0.97 | 0.87 | 4.55E-03 | 0.0201 | (A) NE (B/D) |
| EFCAB10 | 3.17 | 2.3 | 0.61 | 1.13 | 0.87 | 4.85E-03 | 0.021 | (A) NE (B/D) |
| ST6GALNAC4 | 2.01 | 1.14 | 0.63 | 1.16 | 0.87 | 5.85E-03 | 0.0241 | (A) NE (B/D) |
| BSDC1 | 4.63 | 3.76 | 0.69 | 0.94 | 0.87 | 6.55E-03 | 0.0262 | (A) NE (B/D) |
| WDR27 | 0.93 | 0.06 | 0.74 | 0.21 | 0.87 | 7.63E-03 | 0.0293 | (A) NE (B/D) |
| RPS7P1 | 11.97 | 11.1 | 0.46 | 1.57 | 0.87 | 7.96E-03 | 0.0302 | (A) NE (B/D) |
| SMIM3 | 7.28 | 6.42 | 0.71 | 1.17 | 0.87 | 0.0107 | 0.0375 | (A) NE (B/D) |
| FAM173A | 6.66 | 5.79 | 0.79 | 0.55 | 0.87 | 0.0109 | 0.0378 | (A) NE (B/D) |
| RNU6-1085P | 1.16 | 0.29 | 0.79 | 0.74 | 0.87 | 0.0117 | 0.0399 | (A) NE (B/D) |

| | | | | | | | | |
|---|---|---|---|---|---|---|---|---|
| OR2L13 | 0.96 | 0.09 | 0.81 | 0.29 | 0.87 | 0.0124 | 0.0418 | (A) NE (B/D) |
| ALKBH3 | 2.38 | 1.51 | 0.71 | 1.32 | 0.87 | 0.0141 | 0.0458 | (A) NE (B/D) |
| PKNOX1 | 6.75 | 5.89 | 0.24 | 0.65 | 0.86 | 3.42E-07 | 1.27E-05 | (A) NE (B/D) |
| CTDSP2 | 3.14 | 2.28 | 0.14 | 0.81 | 0.86 | 7.59E-07 | 2.40E-05 | (A) NE (B/D) |
| USP36 | 3.88 | 3.02 | 0.21 | 0.77 | 0.86 | 1.05E-06 | 3.10E-05 | (A) NE (B/D) |
| RPL9P25 | 1.6 | 0.74 | 0.29 | 0.67 | 0.86 | 2.98E-06 | 6.98E-05 | (A) NE (B/D) |
| NOL7 | 6.56 | 5.71 | 0.19 | 0.95 | 0.86 | 1.64E-05 | 2.66E-04 | (A) NE (B/D) |
| EPN1 | 3.79 | 2.92 | 0.36 | 0.69 | 0.86 | 2.59E-05 | 3.75E-04 | (A) NE (B/D) |
| CNOT3 | 0.98 | 0.12 | 0.35 | 0.21 | 0.86 | 5.01E-05 | 6.32E-04 | (A) NE (B/D) |
| PDXDC2P-NP | 4.42 | 3.56 | 0.35 | 0.88 | 0.86 | 7.94E-05 | 9.05E-04 | (A) NE (B/D) |
| PPP1R11 | 4.68 | 3.81 | 0.25 | 1.15 | 0.86 | 2.26E-04 | 2.02E-03 | (A) NE (B/D) |
| IGLV5-52 | 9.69 | 8.83 | 0.43 | 0.89 | 0.86 | 3.10E-04 | 2.59E-03 | (A) NE (B/D) |
| PCDHB2 | 0.87 | 0.01 | 0.43 | 0.05 | 0.86 | 3.24E-04 | 2.67E-03 | (A) NE (B/D) |
| KTI12 | 5.35 | 4.49 | 0.41 | 0.97 | 0.86 | 3.66E-04 | 2.94E-03 | (A) NE (B/D) |
| REXO4 | 7.55 | 6.69 | 0.4 | 1.09 | 0.86 | 6.38E-04 | 4.51E-03 | (A) NE (B/D) |
| FBXO9 | 3.25 | 2.39 | 0.41 | 1.08 | 0.86 | 6.66E-04 | 4.66E-03 | (A) NE (B/D) |
| CCNQ | 3.95 | 3.09 | 0.27 | 1.29 | 0.86 | 9.46E-04 | 6.06E-03 | (A) NE (B/D) |
| HIPK2 | 2.99 | 2.13 | 0.45 | 1.07 | 0.86 | 9.94E-04 | 6.28E-03 | (A) NE (B/D) |
| RGMB | 1.9 | 1.04 | 0.38 | 1.22 | 0.86 | 1.03E-03 | 6.45E-03 | (A) NE (B/D) |
| YWHAH | 8.11 | 7.25 | 0.52 | 0.93 | 0.86 | 1.33E-03 | 7.84E-03 | (A) NE (B/D) |
| SBDSP1 | 6.37 | 5.51 | 0.48 | 1.07 | 0.86 | 1.35E-03 | 7.92E-03 | (A) NE (B/D) |
| CYTH2 | 0.96 | 0.1 | 0.56 | 0.22 | 0.86 | 1.57E-03 | 8.92E-03 | (A) NE (B/D) |
| SRD5A3-AS1 | 3.32 | 2.46 | 0.52 | 0.98 | 0.86 | 1.59E-03 | 9.00E-03 | (A) NE (B/D) |
| PSMD2 | 3.29 | 2.43 | 0.43 | 1.25 | 0.86 | 1.92E-03 | 0.0104 | (A) NE (B/D) |
| PREB | 2 | 1.15 | 0.54 | 0.99 | 0.86 | 1.97E-03 | 0.0106 | (A) NE (B/D) |
| AHCYP2 | 1.21 | 0.36 | 0.59 | 0.4 | 0.86 | 2.02E-03 | 0.0108 | (A) NE (B/D) |
| GNB2 | 10.65 | 9.8 | 0.58 | 0.94 | 0.86 | 2.81E-03 | 0.0139 | (A) NE (B/D) |
| PGM1 | 7.28 | 6.41 | 0.59 | 0.95 | 0.86 | 2.93E-03 | 0.0143 | (A) NE (B/D) |
| LRRC42 | 1.6 | 0.74 | 0.6 | 0.9 | 0.86 | 2.99E-03 | 0.0146 | (A) NE (B/D) |
| OSTCP4 | 2.3 | 1.44 | 0.57 | 1.09 | 0.86 | 3.27E-03 | 0.0156 | (A) NE (B/D) |
| CDC16 | 0.91 | 0.05 | 0.64 | 0.11 | 0.86 | 3.67E-03 | 0.017 | (A) NE (B/D) |
| ZNF791 | 6.04 | 5.17 | 0.68 | 0.51 | 0.86 | 4.94E-03 | 0.0213 | (A) NE (B/D) |
| GPT2 | 1.9 | 1.03 | 0.6 | 1.15 | 0.86 | 5.17E-03 | 0.0221 | (A) NE (B/D) |
| ILK | 3.06 | 2.19 | 0.49 | 1.41 | 0.86 | 5.20E-03 | 0.0221 | (A) NE (B/D) |
| TSN | 7.5 | 6.64 | 0.58 | 1.25 | 0.86 | 5.56E-03 | 0.0232 | (A) NE (B/D) |
| ANAPC13 | 8.71 | 7.84 | 0.48 | 1.45 | 0.86 | 6.07E-03 | 0.0248 | (A) NE (B/D) |
| AGAP10P | 6.56 | 5.7 | 0.63 | 1.17 | 0.86 | 6.43E-03 | 0.0259 | (A) NE (B/D) |
| KCP | 2.47 | 1.62 | 0.67 | 0.98 | 0.86 | 6.49E-03 | 0.0261 | (A) NE (B/D) |
| TREX2 | 2.51 | 1.65 | 0.48 | 1.51 | 0.86 | 7.60E-03 | 0.0292 | (A) NE (B/D) |
| POLR3F | 4.38 | 3.52 | 0.72 | 0.89 | 0.86 | 8.62E-03 | 0.032 | (A) NE (B/D) |
| ADAM22 | 0.95 | 0.09 | 0.76 | 0.2 | 0.86 | 9.10E-03 | 0.0333 | (A) NE (B/D) |
| HCG14 | 0.99 | 0.13 | 0.76 | 0.26 | 0.86 | 9.77E-03 | 0.0351 | (A) NE (B/D) |
| TEP1 | 1.93 | 1.07 | 0.77 | 0.51 | 0.86 | 9.79E-03 | 0.0351 | (A) NE (B/D) |
| VEGFA | 3.38 | 2.53 | 0.6 | 1.41 | 0.86 | 0.01 | 0.0358 | (A) NE (B/D) |
| ENO3 | 5.4 | 4.54 | 0.63 | 1.36 | 0.86 | 0.0107 | 0.0375 | (A) NE (B/D) |
| MYDGF | 1.55 | 0.69 | 0.75 | 0.9 | 0.86 | 0.011 | 0.0383 | (A) NE (B/D) |
| STX10 | 1.59 | 0.74 | 0.31 | 0.74 | 0.85 | 1.17E-05 | 2.06E-04 | (A) NE (B/D) |
| PMS2CL | 7.1 | 6.25 | 0.35 | 0.75 | 0.85 | 3.92E-05 | 5.22E-04 | (A) NE (B/D) |
| MRPS31P5 | 4.8 | 3.95 | 0.28 | 0.93 | 0.85 | 4.60E-05 | 5.92E-04 | (A) NE (B/D) |
| DENND1A | 3.9 | 3.05 | 0.41 | 0.56 | 0.85 | 9.84E-05 | 1.07E-03 | (A) NE (B/D) |
| ANKRD13D | 4.11 | 3.26 | 0.34 | 0.98 | 0.85 | 1.55E-04 | 1.52E-03 | (A) NE (B/D) |
| CCT6A | 2.97 | 2.12 | 0.25 | 1.15 | 0.85 | 3.08E-04 | 2.57E-03 | (A) NE (B/D) |
| ABCB7 | 7.53 | 6.68 | 0.42 | 0.91 | 0.85 | 3.17E-04 | 2.63E-03 | (A) NE (B/D) |
| DTD1 | 6.76 | 5.91 | 0.42 | 0.97 | 0.85 | 4.05E-04 | 3.18E-03 | (A) NE (B/D) |
| LMF2 | 1.68 | 0.83 | 0.46 | 0.83 | 0.85 | 4.73E-04 | 3.58E-03 | (A) NE (B/D) |
| PKMYT1 | 1.16 | 0.31 | 0.51 | 0.49 | 0.85 | 7.57E-04 | 5.14E-03 | (A) NE (B/D) |
| ING4 | 6.12 | 5.27 | 0.21 | 1.32 | 0.85 | 9.24E-04 | 5.96E-03 | (A) NE (B/D) |
| ZFAS1 | 4.42 | 3.57 | 0.27 | 1.29 | 0.85 | 9.46E-04 | 6.06E-03 | (A) NE (B/D) |
| SERF2 | 6.85 | 6 | 0.33 | 1.25 | 0.85 | 1.03E-03 | 6.46E-03 | (A) NE (B/D) |
| INHA | 1.31 | 0.46 | 0.54 | 0.53 | 0.85 | 1.05E-03 | 6.56E-03 | (A) NE (B/D) |
| ZFPL1 | 2.52 | 1.67 | 0.52 | 0.94 | 0.85 | 1.52E-03 | 8.72E-03 | (A) NE (B/D) |
| LMNTD2 | 1.48 | 0.63 | 0.55 | 0.79 | 0.85 | 1.53E-03 | 8.74E-03 | (A) NE (B/D) |
| USF1 | 2.75 | 1.9 | 0.4 | 1.29 | 0.85 | 2.29E-03 | 0.0118 | (A) NE (B/D) |
| MTHFD2P7 | 2.82 | 1.96 | 0.6 | 0.8 | 0.85 | 2.78E-03 | 0.0138 | (A) NE (B/D) |
| MRPL9 | 8.27 | 7.42 | 0.49 | 1.28 | 0.85 | 3.57E-03 | 0.0166 | (A) NE (B/D) |

| | | | | | | | | |
|---|---|---|---|---|---|---|---|---|
| UBE2Q1 | 2.22 | 1.37 | 0.56 | 1.13 | 0.85 | 3.82E-03 | 0.0175 | (A) NE (B/D) |
| FAM156B | 6.3 | 5.46 | 0.48 | 1.39 | 0.85 | 5.15E-03 | 0.022 | (A) NE (B/D) |
| EML4 | 3.34 | 2.49 | 0.66 | 0.87 | 0.85 | 5.92E-03 | 0.0244 | (A) NE (B/D) |
| HSD17B14 | 1.27 | 0.42 | 0.69 | 0.75 | 0.85 | 6.34E-03 | 0.0256 | (A) NE (B/D) |
| CEP68 | 0.94 | 0.09 | 0.71 | 0.28 | 0.85 | 6.73E-03 | 0.0268 | (A) NE (B/D) |
| GPSM2 | 3.55 | 2.7 | 0.7 | 0.75 | 0.85 | 6.81E-03 | 0.027 | (A) NE (B/D) |
| ZNF664 | 3.21 | 2.36 | 0.6 | 1.25 | 0.85 | 7.48E-03 | 0.0289 | (A) NE (B/D) |
| FAM153CP | 0.89 | 0.03 | 0.72 | 0.14 | 0.85 | 7.57E-03 | 0.0291 | (A) NE (B/D) |
| SCAPER | 4.52 | 3.68 | 0.73 | 0.66 | 0.85 | 8.17E-03 | 0.0307 | (A) NE (B/D) |
| ZSCAN29 | 4.44 | 3.59 | 0.75 | 0.75 | 0.85 | 9.72E-03 | 0.035 | (A) NE (B/D) |
| SHF | 4.31 | 3.46 | 0.73 | 0.94 | 0.85 | 0.0106 | 0.0371 | (A) NE (B/D) |
| BRCA1 | 5.91 | 5.06 | 0.72 | 1.15 | 0.85 | 0.0129 | 0.0429 | (A) NE (B/D) |
| NPPA-AS1 | 6.18 | 5.33 | 0.45 | 1.7 | 0.85 | 0.0137 | 0.0449 | (A) NE (B/D) |
| FAM27C | 2.9 | 2.05 | 0.62 | 1.48 | 0.85 | 0.0145 | 0.0468 | (A) NE (B/D) |
| RPRD2 | 1.72 | 0.89 | 0.2 | 0.59 | 0.84 | 3.45E-08 | 2.07E-06 | (A) NE (B/D) |
| NELFCD | 2.17 | 1.33 | 0.16 | 0.83 | 0.84 | 2.26E-06 | 5.65E-05 | (A) NE (B/D) |
| KDELR1 | 1.33 | 0.49 | 0.32 | 0.63 | 0.84 | 6.69E-06 | 1.33E-04 | (A) NE (B/D) |
| MADD | 0.85 | 0.01 | 0.27 | 0.03 | 0.84 | 1.50E-05 | 2.49E-04 | (A) NE (B/D) |
| BMS1P1 | 1.71 | 0.87 | 0.33 | 0.72 | 0.84 | 1.89E-05 | 2.95E-04 | (A) NE (B/D) |
| CHKB | 4.9 | 4.05 | 0.33 | 0.92 | 0.84 | 9.28E-05 | 1.03E-03 | (A) NE (B/D) |
| SMARCC2 | 6.62 | 5.79 | 0.37 | 0.84 | 0.84 | 1.12E-04 | 1.18E-03 | (A) NE (B/D) |
| MYO1C | 3.33 | 2.49 | 0.34 | 1.01 | 0.84 | 2.21E-04 | 1.98E-03 | (A) NE (B/D) |
| EHMT1 | 2.23 | 1.39 | 0.45 | 0.54 | 0.84 | 2.72E-04 | 2.33E-03 | (A) NE (B/D) |
| YWHAQ | 1.83 | 0.99 | 0.4 | 1.05 | 0.84 | 6.26E-04 | 4.44E-03 | (A) NE (B/D) |
| MYO18A | 2.09 | 1.25 | 0.53 | 0.56 | 0.84 | 1.02E-03 | 6.41E-03 | (A) NE (B/D) |
| MPRIP | 3.54 | 2.69 | 0.53 | 0.76 | 0.84 | 1.19E-03 | 7.21E-03 | (A) NE (B/D) |
| SMAP1 | 2.44 | 1.59 | 0.56 | 0.66 | 0.84 | 1.60E-03 | 9.04E-03 | (A) NE (B/D) |
| PSMA7 | 6.71 | 5.87 | 0.37 | 1.26 | 0.84 | 1.63E-03 | 9.17E-03 | (A) NE (B/D) |
| MASTL | 5.2 | 4.36 | 0.57 | 0.75 | 0.84 | 1.96E-03 | 0.0105 | (A) NE (B/D) |
| CACNA1B | 0.89 | 0.05 | 0.59 | 0.13 | 0.84 | 2.69E-03 | 0.0135 | (A) NE (B/D) |
| ILRUN | 8.35 | 7.52 | 0.57 | 0.88 | 0.84 | 2.71E-03 | 0.0135 | (A) NE (B/D) |
| PPCS | 4.83 | 3.99 | 0.45 | 1.25 | 0.84 | 2.71E-03 | 0.0135 | (A) NE (B/D) |
| LMO4 | 2.96 | 2.12 | 0.56 | 0.96 | 0.84 | 2.78E-03 | 0.0138 | (A) NE (B/D) |
| NACC1 | 4.66 | 3.81 | 0.6 | 0.71 | 0.84 | 2.84E-03 | 0.014 | (A) NE (B/D) |
| IRAK1 | 3.18 | 2.33 | 0.43 | 1.31 | 0.84 | 3.10E-03 | 0.015 | (A) NE (B/D) |
| SELENOS | 7.12 | 6.28 | 0.57 | 1.06 | 0.84 | 3.61E-03 | 0.0168 | (A) NE (B/D) |
| KPTN | 1.02 | 0.18 | 0.63 | 0.31 | 0.84 | 3.72E-03 | 0.0172 | (A) NE (B/D) |
| WDR45B | 7.94 | 7.1 | 0.63 | 0.88 | 0.84 | 4.57E-03 | 0.0201 | (A) NE (B/D) |
| VAPA | 7.75 | 6.91 | 0.65 | 0.76 | 0.84 | 5.09E-03 | 0.0218 | (A) NE (B/D) |
| KIT | 0.84 | 0 | 0.67 | 0.01 | 0.84 | 5.30E-03 | 0.0224 | (A) NE (B/D) |
| RNU7-1 | 3.38 | 2.55 | 0.45 | 1.41 | 0.84 | 5.45E-03 | 0.0229 | (A) NE (B/D) |
| MTHFD2 | 4.19 | 3.36 | 0.66 | 0.85 | 0.84 | 5.83E-03 | 0.0241 | (A) NE (B/D) |
| SRPK3 | 1.18 | 0.35 | 0.73 | 0.47 | 0.84 | 8.52E-03 | 0.0317 | (A) NE (B/D) |
| SYVN1 | 3.29 | 2.45 | 0.67 | 1.13 | 0.84 | 9.15E-03 | 0.0334 | (A) NE (B/D) |
| DNAJC10 | 4.68 | 3.84 | 0.72 | 0.88 | 0.84 | 0.0101 | 0.0359 | (A) NE (B/D) |
| TTC19 | 3.42 | 2.58 | 0.72 | 0.97 | 0.84 | 0.0105 | 0.037 | (A) NE (B/D) |
| RAB2A | 3.63 | 2.79 | 0.66 | 1.2 | 0.84 | 0.0106 | 0.0371 | (A) NE (B/D) |
| HLF | 1.99 | 1.14 | 0.66 | 1.25 | 0.84 | 0.0111 | 0.0383 | (A) NE (B/D) |
| RPL26 | 11.24 | 10.39 | 0.43 | 1.65 | 0.84 | 0.0111 | 0.0385 | (A) NE (B/D) |
| SPTSSA | 1.6 | 0.76 | 0.76 | 0.8 | 0.84 | 0.0117 | 0.04 | (A) NE (B/D) |
| PDK4 | 0.88 | 0.04 | 0.78 | 0.12 | 0.84 | 0.0119 | 0.0405 | (A) NE (B/D) |
| ZNF474 | 0.98 | 0.15 | 0.78 | 0.43 | 0.84 | 0.0124 | 0.0417 | (A) NE (B/D) |
| RFK | 1.51 | 0.67 | 0.77 | 0.74 | 0.84 | 0.0128 | 0.0427 | (A) NE (B/D) |
| ZNF17 | 1.44 | 0.6 | 0.82 | 0.41 | 0.84 | 0.0152 | 0.0485 | (A) NE (B/D) |
| PDCD5 | 2.84 | 2.01 | 0.77 | 1 | 0.84 | 0.0153 | 0.0486 | (A) NE (B/D) |
| TRIM39-RPP2 | 1.32 | 0.49 | 0.3 | 0.6 | 0.83 | 4.68E-06 | 1.00E-04 | (A) NE (B/D) |
| ZFAND3 | 2.82 | 2 | 0.3 | 0.68 | 0.83 | 8.53E-06 | 1.61E-04 | (A) NE (B/D) |
| LINC02210 | 1.71 | 0.88 | 0.34 | 0.75 | 0.83 | 3.57E-05 | 4.85E-04 | (A) NE (B/D) |
| MORC2 | 2.82 | 1.99 | 0.24 | 1.02 | 0.83 | 9.87E-05 | 1.07E-03 | (A) NE (B/D) |
| CDS2 | 6.88 | 6.06 | 0.43 | 0.43 | 0.83 | 2.03E-04 | 1.86E-03 | (A) NE (B/D) |
| PDAP1 | 3.97 | 3.14 | 0.31 | 1.05 | 0.83 | 2.83E-04 | 2.40E-03 | (A) NE (B/D) |
| NCOR2 | 3.11 | 2.29 | 0.44 | 0.64 | 0.83 | 2.89E-04 | 2.45E-03 | (A) NE (B/D) |
| UBE2D4 | 2.2 | 1.37 | 0.37 | 0.98 | 0.83 | 3.38E-04 | 2.76E-03 | (A) NE (B/D) |
| SENP3-EIF4A | 9.59 | 8.76 | 0.31 | 1.14 | 0.83 | 5.01E-04 | 3.75E-03 | (A) NE (B/D) |
| SEPTIN7P3 | 4.24 | 3.41 | 0.37 | 1.05 | 0.83 | 5.07E-04 | 3.78E-03 | (A) NE (B/D) |

| | | | | | | | | |
|---|---|---|---|---|---|---|---|---|
| INO80B | 4.07 | 3.24 | 0.35 | 1.17 | 0.83 | 9.46E-04 | 6.06E-03 | (A) NE (B/D) |
| BAK1 | 5.93 | 5.1 | 0.37 | 1.15 | 0.83 | 9.93E-04 | 6.28E-03 | (A) NE (B/D) |
| ECI1 | 2.14 | 1.31 | 0.49 | 0.96 | 0.83 | 1.45E-03 | 8.41E-03 | (A) NE (B/D) |
| ATP5MPL | 3.53 | 2.7 | 0.36 | 1.23 | 0.83 | 1.50E-03 | 8.64E-03 | (A) NE (B/D) |
| ERVK3-1 | 8.23 | 7.4 | 0.25 | 1.34 | 0.83 | 1.51E-03 | 8.65E-03 | (A) NE (B/D) |
| NAPA | 5.02 | 4.18 | 0.26 | 1.34 | 0.83 | 1.55E-03 | 8.85E-03 | (A) NE (B/D) |
| PSPH | 8.24 | 7.42 | 0.48 | 1.05 | 0.83 | 1.74E-03 | 9.63E-03 | (A) NE (B/D) |
| TAOK2 | 2.12 | 1.28 | 0.51 | 0.99 | 0.83 | 1.83E-03 | 0.01 | (A) NE (B/D) |
| TMEM260 | 1.13 | 0.29 | 0.56 | 0.28 | 0.83 | 1.93E-03 | 0.0104 | (A) NE (B/D) |
| CCDC97 | 1.49 | 0.66 | 0.56 | 0.7 | 0.83 | 1.98E-03 | 0.0106 | (A) NE (B/D) |
| HNRNPUL2 | 2.19 | 1.36 | 0.48 | 1.15 | 0.83 | 2.53E-03 | 0.0128 | (A) NE (B/D) |
| LENG1 | 0.87 | 0.04 | 0.61 | 0.15 | 0.83 | 3.38E-03 | 0.016 | (A) NE (B/D) |
| NRDC | 7.54 | 6.71 | 0.59 | 0.94 | 0.83 | 3.87E-03 | 0.0177 | (A) NE (B/D) |
| PAN2 | 0.88 | 0.05 | 0.66 | 0.13 | 0.83 | 5.21E-03 | 0.0222 | (A) NE (B/D) |
| FAM27E2 | 2.44 | 1.61 | 0.58 | 1.13 | 0.83 | 5.40E-03 | 0.0227 | (A) NE (B/D) |
| NDUFC2-KCT | 7.81 | 6.98 | 0.29 | 1.55 | 0.83 | 5.54E-03 | 0.0232 | (A) NE (B/D) |
| TMEM63A | 1.88 | 1.06 | 0.66 | 0.84 | 0.83 | 6.13E-03 | 0.025 | (A) NE (B/D) |
| PFN1 | 4.95 | 4.12 | 0.37 | 1.51 | 0.83 | 6.36E-03 | 0.0257 | (A) NE (B/D) |
| SMC5-AS1 | 4.27 | 3.44 | 0.68 | 0.88 | 0.83 | 7.77E-03 | 0.0296 | (A) NE (B/D) |
| CDH12P4 | 4.9 | 4.07 | 0.47 | 1.53 | 0.83 | 9.46E-03 | 0.0342 | (A) NE (B/D) |
| HNRNPCP4 | 2.04 | 1.21 | 0.68 | 1.1 | 0.83 | 0.011 | 0.038 | (A) NE (B/D) |
| SNORD118 | 1.4 | 0.57 | 0.76 | 0.58 | 0.83 | 0.0111 | 0.0385 | (A) NE (B/D) |
| ZNF180 | 1.28 | 0.45 | 0.78 | 0.49 | 0.83 | 0.0128 | 0.0428 | (A) NE (B/D) |
| CLPP | 1.09 | 0.26 | 0.81 | 0.41 | 0.83 | 0.0153 | 0.0487 | (A) NE (B/D) |
| PFDN5 | 3.36 | 2.53 | 0.43 | 1.74 | 0.83 | 0.0157 | 0.0496 | (A) NE (B/D) |
| APOM | 4.43 | 3.6 | 0.18 | 0.72 | 0.82 | 4.50E-07 | 1.59E-05 | (A) NE (B/D) |
| OLA1P1 | 1.6 | 0.78 | 0.22 | 0.73 | 0.82 | 1.54E-06 | 4.21E-05 | (A) NE (B/D) |
| RAE1 | 6.48 | 5.66 | 0.18 | 0.94 | 0.82 | 2.29E-05 | 3.41E-04 | (A) NE (B/D) |
| GUSBP11 | 8.46 | 7.64 | 0.29 | 0.82 | 0.82 | 2.59E-05 | 3.75E-04 | (A) NE (B/D) |
| COX10 | 2.06 | 1.25 | 0.32 | 0.74 | 0.82 | 2.78E-05 | 3.97E-04 | (A) NE (B/D) |
| STX18 | 1.9 | 1.08 | 0.36 | 0.59 | 0.82 | 3.37E-05 | 4.63E-04 | (A) NE (B/D) |
| KCNK3 | 0.96 | 0.14 | 0.36 | 0.39 | 0.82 | 3.83E-05 | 5.14E-04 | (A) NE (B/D) |
| IKBKGP1 | 4.27 | 3.45 | 0.3 | 1.01 | 0.82 | 1.96E-04 | 1.81E-03 | (A) NE (B/D) |
| UBE2G2 | 3.6 | 2.78 | 0.41 | 0.8 | 0.82 | 2.35E-04 | 2.08E-03 | (A) NE (B/D) |
| C5ORF38 | 1.16 | 0.34 | 0.49 | 0.62 | 0.82 | 7.17E-04 | 4.93E-03 | (A) NE (B/D) |
| FMNL2 | 1.26 | 0.45 | 0.49 | 0.7 | 0.82 | 7.67E-04 | 5.19E-03 | (A) NE (B/D) |
| NPRL2 | 2.07 | 1.25 | 0.4 | 1.08 | 0.82 | 8.98E-04 | 5.83E-03 | (A) NE (B/D) |
| CASP3 | 3.04 | 2.21 | 0.51 | 1.02 | 0.82 | 2.22E-03 | 0.0116 | (A) NE (B/D) |
| DNAL4 | 4.14 | 3.33 | 0.49 | 1.05 | 0.82 | 2.25E-03 | 0.0117 | (A) NE (B/D) |
| SNORD57 | 1 | 0.19 | 0.57 | 0.3 | 0.82 | 2.25E-03 | 0.0117 | (A) NE (B/D) |
| HERC2P2 | 7.16 | 6.34 | 0.47 | 1.18 | 0.82 | 2.78E-03 | 0.0138 | (A) NE (B/D) |
| DLG5 | 6.5 | 5.68 | 0.43 | 1.33 | 0.82 | 4.33E-03 | 0.0193 | (A) NE (B/D) |
| MYBL2 | 1.21 | 0.4 | 0.6 | 0.92 | 0.82 | 4.72E-03 | 0.0206 | (A) NE (B/D) |
| ZRSR2 | 3.1 | 2.28 | 0.58 | 1.05 | 0.82 | 4.97E-03 | 0.0214 | (A) NE (B/D) |
| NOL4 | 0.84 | 0.01 | 0.65 | 0.08 | 0.82 | 5.26E-03 | 0.0223 | (A) NE (B/D) |
| OR8D2 | 1.12 | 0.31 | 0.65 | 0.77 | 0.82 | 6.08E-03 | 0.0248 | (A) NE (B/D) |
| MTIF2 | 1.3 | 0.48 | 0.67 | 0.6 | 0.82 | 6.34E-03 | 0.0257 | (A) NE (B/D) |
| MLF2 | 4.91 | 4.09 | 0.46 | 1.42 | 0.82 | 6.95E-03 | 0.0275 | (A) NE (B/D) |
| SYNCRIP | 2.2 | 1.38 | 0.62 | 1.08 | 0.82 | 7.39E-03 | 0.0287 | (A) NE (B/D) |
| VDAC3 | 2.28 | 1.46 | 0.59 | 1.2 | 0.82 | 7.95E-03 | 0.0301 | (A) NE (B/D) |
| SET | 4.74 | 3.92 | 0.61 | 1.18 | 0.82 | 8.31E-03 | 0.0311 | (A) NE (B/D) |
| ZNF93 | 0.83 | 0.01 | 0.71 | 0.04 | 0.82 | 8.47E-03 | 0.0316 | (A) NE (B/D) |
| SPIN3 | 2.23 | 1.41 | 0.66 | 1.03 | 0.82 | 8.81E-03 | 0.0325 | (A) NE (B/D) |
| TMEM108 | 0.83 | 0.01 | 0.77 | 0.07 | 0.82 | 0.0128 | 0.0426 | (A) NE (B/D) |
| RPS20 | 11.45 | 10.63 | 0.36 | 1.7 | 0.82 | 0.0131 | 0.0434 | (A) NE (B/D) |
| GTF2IRD2P1 | 1.02 | 0.2 | 0.78 | 0.38 | 0.82 | 0.0135 | 0.0444 | (A) NE (B/D) |
| USP18 | 1.24 | 0.42 | 0.76 | 0.72 | 0.82 | 0.0139 | 0.0452 | (A) NE (B/D) |
| HIC2 | 2.34 | 1.54 | 0.2 | 0.41 | 0.81 | 6.62E-09 | 5.77E-07 | (A) NE (B/D) |
| BUD31 | 1.5 | 0.68 | 0.21 | 0.79 | 0.81 | 3.87E-06 | 8.64E-05 | (A) NE (B/D) |
| MAP7D1 | 1.39 | 0.57 | 0.32 | 0.61 | 0.81 | 1.17E-05 | 2.06E-04 | (A) NE (B/D) |
| TM9SF4 | 2.02 | 1.21 | 0.24 | 0.85 | 0.81 | 1.74E-05 | 2.77E-04 | (A) NE (B/D) |
| MED25 | 0.88 | 0.07 | 0.38 | 0.21 | 0.81 | 1.34E-04 | 1.36E-03 | (A) NE (B/D) |
| CACNB1 | 1.37 | 0.56 | 0.41 | 0.67 | 0.81 | 1.91E-04 | 1.78E-03 | (A) NE (B/D) |
| C19ORF53 | 1.36 | 0.55 | 0.43 | 0.66 | 0.81 | 3.05E-04 | 2.56E-03 | (A) NE (B/D) |
| C1ORF131 | 2.72 | 1.9 | 0.44 | 0.72 | 0.81 | 3.40E-04 | 2.78E-03 | (A) NE (B/D) |

| | | | | | | | | |
|---|---|---|---|---|---|---|---|---|
| LINC00866 | 0.85 | 0.04 | 0.43 | 0.15 | 0.81 | 4.55E-04 | 3.47E-03 | (A) NE (B/D) |
| HSD17B11 | 1.34 | 0.53 | 0.47 | 0.65 | 0.81 | 5.89E-04 | 4.25E-03 | (A) NE (B/D) |
| QDPR | 1.08 | 0.27 | 0.49 | 0.45 | 0.81 | 8.37E-04 | 5.54E-03 | (A) NE (B/D) |
| KLF10 | 1.94 | 1.12 | 0.48 | 0.84 | 0.81 | 9.36E-04 | 6.01E-03 | (A) NE (B/D) |
| CDCP2 | 1.45 | 0.64 | 0.51 | 0.5 | 0.81 | 9.92E-04 | 6.28E-03 | (A) NE (B/D) |
| KCTD7 | 2.92 | 2.11 | 0.51 | 0.69 | 0.81 | 1.24E-03 | 7.41E-03 | (A) NE (B/D) |
| HADH | 8.85 | 8.04 | 0.33 | 1.22 | 0.81 | 1.35E-03 | 7.96E-03 | (A) NE (B/D) |
| DFFB | 1.12 | 0.31 | 0.55 | 0.45 | 0.81 | 1.80E-03 | 9.91E-03 | (A) NE (B/D) |
| GNL2 | 8.6 | 7.78 | 0.45 | 1.1 | 0.81 | 1.90E-03 | 0.0103 | (A) NE (B/D) |
| STT3A | 4.88 | 4.06 | 0.42 | 1.16 | 0.81 | 1.98E-03 | 0.0106 | (A) NE (B/D) |
| OLA1 | 1.83 | 1.03 | 0.51 | 0.93 | 0.81 | 2.10E-03 | 0.0111 | (A) NE (B/D) |
| MRPS24 | 2.11 | 1.3 | 0.52 | 1 | 0.81 | 2.63E-03 | 0.0132 | (A) NE (B/D) |
| TRAPPC12 | 1.99 | 1.18 | 0.59 | 0.54 | 0.81 | 2.97E-03 | 0.0145 | (A) NE (B/D) |
| TINF2 | 1.66 | 0.85 | 0.58 | 0.77 | 0.81 | 3.13E-03 | 0.0151 | (A) NE (B/D) |
| NPEPL1 | 1.26 | 0.45 | 0.6 | 0.58 | 0.81 | 3.41E-03 | 0.0161 | (A) NE (B/D) |
| SLC20A1 | 4.75 | 3.94 | 0.62 | 0.76 | 0.81 | 4.50E-03 | 0.0199 | (A) NE (B/D) |
| PAIP1 | 2.07 | 1.26 | 0.58 | 0.99 | 0.81 | 4.51E-03 | 0.0199 | (A) NE (B/D) |
| PAXIP1 | 3.14 | 2.33 | 0.62 | 0.72 | 0.81 | 4.55E-03 | 0.0201 | (A) NE (B/D) |
| PTCH2 | 1.31 | 0.5 | 0.59 | 0.92 | 0.81 | 4.77E-03 | 0.0208 | (A) NE (B/D) |
| LRCH4 | 0.97 | 0.16 | 0.64 | 0.39 | 0.81 | 5.02E-03 | 0.0216 | (A) NE (B/D) |
| IRGQ | 6.12 | 5.31 | 0.64 | 0.68 | 0.81 | 5.16E-03 | 0.022 | (A) NE (B/D) |
| RIPK1 | 1.41 | 0.6 | 0.64 | 0.59 | 0.81 | 5.38E-03 | 0.0226 | (A) NE (B/D) |
| ZNF45 | 0.94 | 0.13 | 0.67 | 0.21 | 0.81 | 6.32E-03 | 0.0256 | (A) NE (B/D) |
| VPS54 | 4.93 | 4.12 | 0.68 | 0.66 | 0.81 | 7.42E-03 | 0.0288 | (A) NE (B/D) |
| MBIP | 1.17 | 0.36 | 0.68 | 0.54 | 0.81 | 7.56E-03 | 0.0291 | (A) NE (B/D) |
| MIR-93-1/93 | 1.68 | 0.87 | 0.62 | 1.09 | 0.81 | 7.64E-03 | 0.0293 | (A) NE (B/D) |
| MIR-93-7.1/9 | 1.68 | 0.87 | 0.62 | 1.09 | 0.81 | 7.64E-03 | 0.0293 | (A) NE (B/D) |
| MIR-93/3P | 1.68 | 0.87 | 0.62 | 1.09 | 0.81 | 7.64E-03 | 0.0293 | (A) NE (B/D) |
| MIR-93/5P | 1.68 | 0.87 | 0.62 | 1.09 | 0.81 | 7.64E-03 | 0.0293 | (A) NE (B/D) |
| MIR-93/93 | 1.68 | 0.87 | 0.62 | 1.09 | 0.81 | 7.64E-03 | 0.0293 | (A) NE (B/D) |
| MIR-93/93* | 1.68 | 0.87 | 0.62 | 1.09 | 0.81 | 7.64E-03 | 0.0293 | (A) NE (B/D) |
| AMIGO3 | 3.68 | 2.87 | 0.62 | 1.06 | 0.81 | 7.71E-03 | 0.0295 | (A) NE (B/D) |
| NUP210P3 | 0.82 | 0.02 | 0.71 | 0.09 | 0.81 | 8.89E-03 | 0.0328 | (A) NE (B/D) |
| ARHGDIA | 5.5 | 4.69 | 0.67 | 1.13 | 0.81 | 0.0117 | 0.0401 | (A) NE (B/D) |
| PDIA6 | 2.11 | 1.3 | 0.73 | 0.98 | 0.81 | 0.0141 | 0.0458 | (A) NE (B/D) |
| ARFRP1 | 6.33 | 5.53 | 0.23 | 0.68 | 0.8 | 1.14E-06 | 3.30E-05 | (A) NE (B/D) |
| JSRP1 | 1.73 | 0.93 | 0.12 | 1 | 0.8 | 5.70E-05 | 7.01E-04 | (A) NE (B/D) |
| UBA6-AS1 | 4.41 | 3.61 | 0.38 | 0.78 | 0.8 | 1.78E-04 | 1.68E-03 | (A) NE (B/D) |
| RAI1 | 0.85 | 0.04 | 0.39 | 0.11 | 0.8 | 2.29E-04 | 2.04E-03 | (A) NE (B/D) |
| UPP1 | 3.65 | 2.85 | 0.36 | 0.9 | 0.8 | 2.52E-04 | 2.19E-03 | (A) NE (B/D) |
| HDAC3 | 1.67 | 0.87 | 0.43 | 0.67 | 0.8 | 3.04E-04 | 2.55E-03 | (A) NE (B/D) |
| SH3PXD2A | 1.14 | 0.34 | 0.47 | 0.65 | 0.8 | 6.63E-04 | 4.65E-03 | (A) NE (B/D) |
| RPS19BP1 | 2.78 | 1.98 | 0.47 | 0.72 | 0.8 | 6.83E-04 | 4.75E-03 | (A) NE (B/D) |
| PIGFP1 | 3.71 | 2.92 | 0.42 | 0.93 | 0.8 | 6.95E-04 | 4.83E-03 | (A) NE (B/D) |
| DDOST | 7.37 | 6.57 | 0.23 | 1.19 | 0.8 | 7.05E-04 | 4.88E-03 | (A) NE (B/D) |
| MRPL20 | 2.1 | 1.3 | 0.36 | 1.07 | 0.8 | 7.98E-04 | 5.36E-03 | (A) NE (B/D) |
| EIF5 | 5.08 | 4.27 | 0.26 | 1.2 | 0.8 | 8.35E-04 | 5.54E-03 | (A) NE (B/D) |
| ATG13 | 7.37 | 6.57 | 0.44 | 0.9 | 0.8 | 8.57E-04 | 5.63E-03 | (A) NE (B/D) |
| PLA2G12AP1 | 3.62 | 2.83 | 0.4 | 1.01 | 0.8 | 9.08E-04 | 5.88E-03 | (A) NE (B/D) |
| SLC25A13 | 3.51 | 2.7 | 0.48 | 0.82 | 0.8 | 1.05E-03 | 6.56E-03 | (A) NE (B/D) |
| RPA1 | 2.06 | 1.26 | 0.37 | 1.2 | 0.8 | 1.84E-03 | 0.0101 | (A) NE (B/D) |
| UBA1 | 8.06 | 7.25 | 0.39 | 1.21 | 0.8 | 2.11E-03 | 0.0112 | (A) NE (B/D) |
| KCTD3 | 3.35 | 2.55 | 0.44 | 1.12 | 0.8 | 2.14E-03 | 0.0113 | (A) NE (B/D) |
| POP4 | 1.53 | 0.73 | 0.55 | 0.74 | 0.8 | 2.38E-03 | 0.0122 | (A) NE (B/D) |
| FXR2 | 0.9 | 0.11 | 0.56 | 0.21 | 0.8 | 2.48E-03 | 0.0127 | (A) NE (B/D) |
| HAGH | 7.4 | 6.6 | 0.34 | 1.32 | 0.8 | 2.79E-03 | 0.0138 | (A) NE (B/D) |
| ECH1 | 1.75 | 0.94 | 0.48 | 1.12 | 0.8 | 3.04E-03 | 0.0147 | (A) NE (B/D) |
| PRKCZ | 5.86 | 5.05 | 0.53 | 0.97 | 0.8 | 3.10E-03 | 0.015 | (A) NE (B/D) |
| PSEN2 | 2.83 | 2.02 | 0.46 | 1.18 | 0.8 | 3.23E-03 | 0.0155 | (A) NE (B/D) |
| PCDHB16 | 0.83 | 0.03 | 0.59 | 0.12 | 0.8 | 3.40E-03 | 0.0161 | (A) NE (B/D) |
| BIVM-ERCC5 | 6.58 | 5.78 | 0.59 | 0.71 | 0.8 | 3.69E-03 | 0.0171 | (A) NE (B/D) |
| PSMD14 | 4.77 | 3.97 | 0.53 | 1.04 | 0.8 | 3.75E-03 | 0.0173 | (A) NE (B/D) |
| SPATA16 | 0.9 | 0.1 | 0.62 | 0.26 | 0.8 | 4.61E-03 | 0.0202 | (A) NE (B/D) |
| MED11 | 5.96 | 5.16 | 0.44 | 1.32 | 0.8 | 5.26E-03 | 0.0223 | (A) NE (B/D) |
| NT5C3B | 1.69 | 0.9 | 0.57 | 1.11 | 0.8 | 6.37E-03 | 0.0257 | (A) NE (B/D) |

| | | | | | | | | |
|---|---|---|---|---|---|---|---|---|
| PDCD6 | 1.73 | 0.93 | 0.63 | 0.91 | 0.8 | 6.45E-03 | 0.0259 | (A) NE (B/D) |
| OSGEPL1 | 1.63 | 0.83 | 0.66 | 0.72 | 0.8 | 6.84E-03 | 0.0271 | (A) NE (B/D) |
| PPP1R13L | 0.93 | 0.13 | 0.68 | 0.31 | 0.8 | 7.54E-03 | 0.0291 | (A) NE (B/D) |
| ZMAT2 | 3.33 | 2.53 | 0.4 | 1.47 | 0.8 | 8.12E-03 | 0.0306 | (A) NE (B/D) |
| ZNF559 | 1.19 | 0.39 | 0.7 | 0.68 | 0.8 | 9.26E-03 | 0.0337 | (A) NE (B/D) |
| SEC24B-AS1 | 2.67 | 1.87 | 0.71 | 0.74 | 0.8 | 0.0104 | 0.0368 | (A) NE (B/D) |
| KIAA1586 | 5.52 | 4.72 | 0.7 | 0.94 | 0.8 | 0.0118 | 0.0402 | (A) NE (B/D) |
| DGKQ | 1.65 | 0.84 | 0.77 | 0.65 | 0.8 | 0.0152 | 0.0485 | (A) NE (B/D) |
| ZNF787 | 3.05 | 2.25 | 0.2 | 0.49 | 0.79 | 2.26E-08 | 1.44E-06 | (A) NE (B/D) |
| TTI1 | 6.18 | 5.4 | 0.29 | 0.59 | 0.79 | 5.27E-06 | 1.10E-04 | (A) NE (B/D) |
| ZC3H4 | 1.88 | 1.09 | 0.34 | 0.49 | 0.79 | 2.43E-05 | 3.56E-04 | (A) NE (B/D) |
| POM121 | 1 | 0.21 | 0.37 | 0.34 | 0.79 | 9.34E-05 | 1.03E-03 | (A) NE (B/D) |
| MIR-3620/36 | 1.26 | 0.47 | 0.38 | 0.38 | 0.79 | 1.15E-04 | 1.21E-03 | (A) NE (B/D) |
| MIR-3620/3P | 1.26 | 0.47 | 0.38 | 0.38 | 0.79 | 1.15E-04 | 1.21E-03 | (A) NE (B/D) |
| MIR-3620/5P | 1.26 | 0.47 | 0.38 | 0.38 | 0.79 | 1.15E-04 | 1.21E-03 | (A) NE (B/D) |
| SNUPN | 1.68 | 0.89 | 0.43 | 0.87 | 0.79 | 7.08E-04 | 4.89E-03 | (A) NE (B/D) |
| STX16 | 3.86 | 3.07 | 0.24 | 1.17 | 0.79 | 7.63E-04 | 5.18E-03 | (A) NE (B/D) |
| INE1 | 4.54 | 3.76 | 0.5 | 0.7 | 0.79 | 1.24E-03 | 7.42E-03 | (A) NE (B/D) |
| GPR107 | 7.27 | 6.48 | 0.42 | 1.05 | 0.79 | 1.53E-03 | 8.74E-03 | (A) NE (B/D) |
| NDUFC1 | 2.93 | 2.14 | 0.35 | 1.24 | 0.79 | 2.21E-03 | 0.0115 | (A) NE (B/D) |
| CLTB | 2.51 | 1.72 | 0.55 | 0.75 | 0.79 | 2.45E-03 | 0.0125 | (A) NE (B/D) |
| CCT4 | 6.4 | 5.61 | 0.49 | 1.04 | 0.79 | 2.96E-03 | 0.0144 | (A) NE (B/D) |
| SNX22 | 4.81 | 4.01 | 0.36 | 1.32 | 0.79 | 3.36E-03 | 0.0159 | (A) NE (B/D) |
| PPP4R3A | 3.38 | 2.59 | 0.53 | 0.96 | 0.79 | 3.48E-03 | 0.0163 | (A) NE (B/D) |
| ANKS3 | 3.7 | 2.9 | 0.56 | 0.87 | 0.79 | 3.51E-03 | 0.0164 | (A) NE (B/D) |
| PAXBP1 | 1.77 | 0.98 | 0.6 | 0.55 | 0.79 | 3.65E-03 | 0.0169 | (A) NE (B/D) |
| RPN2 | 4.7 | 3.91 | 0.41 | 1.27 | 0.79 | 4.08E-03 | 0.0184 | (A) NE (B/D) |
| ZNF140 | 1.34 | 0.55 | 0.58 | 0.87 | 0.79 | 4.28E-03 | 0.0192 | (A) NE (B/D) |
| SIK2 | 1.92 | 1.13 | 0.61 | 0.52 | 0.79 | 4.28E-03 | 0.0192 | (A) NE (B/D) |
| MYL6B | 2.75 | 1.96 | 0.29 | 1.43 | 0.79 | 4.70E-03 | 0.0205 | (A) NE (B/D) |
| SPSB3 | 4.28 | 3.5 | 0.58 | 0.97 | 0.79 | 5.39E-03 | 0.0227 | (A) NE (B/D) |
| TGIF2-RAB5I | 5.14 | 4.36 | 0.46 | 1.31 | 0.79 | 6.65E-03 | 0.0265 | (A) NE (B/D) |
| PGBD2 | 6.67 | 5.88 | 0.62 | 0.93 | 0.79 | 7.16E-03 | 0.028 | (A) NE (B/D) |
| LINC01001 | 6.92 | 6.13 | 0.58 | 1.14 | 0.79 | 7.86E-03 | 0.0299 | (A) NE (B/D) |
| EXOC3 | 0.93 | 0.14 | 0.68 | 0.25 | 0.79 | 8.37E-03 | 0.0313 | (A) NE (B/D) |
| MED28 | 1.48 | 0.69 | 0.69 | 0.58 | 0.79 | 9.35E-03 | 0.034 | (A) NE (B/D) |
| MIR-4656/46 | 3.4 | 2.61 | 0.7 | 0.78 | 0.79 | 0.0112 | 0.0386 | (A) NE (B/D) |
| ANKIB1 | 1.96 | 1.16 | 0.71 | 0.82 | 0.79 | 0.0119 | 0.0405 | (A) NE (B/D) |
| LYSMD4 | 6.54 | 5.75 | 0.66 | 1.12 | 0.79 | 0.0127 | 0.0426 | (A) NE (B/D) |
| HMGB1P10 | 2.66 | 1.87 | 0.52 | 1.44 | 0.79 | 0.0133 | 0.0438 | (A) NE (B/D) |
| PFN4 | 3.11 | 2.32 | 0.69 | 1.01 | 0.79 | 0.0133 | 0.044 | (A) NE (B/D) |
| GDPD5 | 1.93 | 1.13 | 0.64 | 1.2 | 0.79 | 0.0134 | 0.0442 | (A) NE (B/D) |
| TLE1 | 4.66 | 3.87 | 0.68 | 1.13 | 0.79 | 0.0152 | 0.0485 | (A) NE (B/D) |
| FTH1P23 | 7.79 | 7.01 | 0.47 | 1.58 | 0.79 | 0.0158 | 0.0498 | (A) NE (B/D) |
| YTHDF1 | 6.09 | 5.32 | 0.22 | 0.89 | 0.78 | 3.76E-05 | 5.06E-04 | (A) NE (B/D) |
| ZFYVE9P1 | 1.44 | 0.66 | 0.35 | 0.64 | 0.78 | 5.11E-05 | 6.43E-04 | (A) NE (B/D) |
| FAM32A | 1.55 | 0.77 | 0.34 | 0.79 | 0.78 | 9.70E-05 | 1.06E-03 | (A) NE (B/D) |
| ABCB8 | 0.94 | 0.16 | 0.37 | 0.31 | 0.78 | 1.26E-04 | 1.29E-03 | (A) NE (B/D) |
| EIF4H | 5.08 | 4.31 | 0.29 | 0.92 | 0.78 | 1.57E-04 | 1.53E-03 | (A) NE (B/D) |
| KCTD18 | 9.45 | 8.67 | 0.39 | 0.7 | 0.78 | 1.87E-04 | 1.75E-03 | (A) NE (B/D) |
| ANKRD52 | 3.02 | 2.24 | 0.37 | 0.84 | 0.78 | 2.65E-04 | 2.28E-03 | (A) NE (B/D) |
| RABL2A | 5.91 | 5.12 | 0.24 | 1.07 | 0.78 | 3.44E-04 | 2.80E-03 | (A) NE (B/D) |
| ZKSCAN5 | 1.54 | 0.76 | 0.45 | 0.59 | 0.78 | 4.99E-04 | 3.74E-03 | (A) NE (B/D) |
| RBM14 | 6.38 | 5.59 | 0.47 | 0.56 | 0.78 | 7.31E-04 | 5.00E-03 | (A) NE (B/D) |
| SNORA67 | 4.08 | 3.3 | 0.29 | 1.27 | 0.78 | 2.17E-03 | 0.0113 | (A) NE (B/D) |
| TBC1D20 | 1.64 | 0.86 | 0.57 | 0.75 | 0.78 | 3.52E-03 | 0.0165 | (A) NE (B/D) |
| CDK5RAP3 | 7.89 | 7.11 | 0.41 | 1.24 | 0.78 | 3.70E-03 | 0.0171 | (A) NE (B/D) |
| WSCD1 | 0.79 | 0.01 | 0.6 | 0.04 | 0.78 | 4.29E-03 | 0.0192 | (A) NE (B/D) |
| PRDX3P2 | 3.47 | 2.69 | 0.56 | 0.94 | 0.78 | 4.92E-03 | 0.0212 | (A) NE (B/D) |
| DDX17 | 4.3 | 3.52 | 0.59 | 0.85 | 0.78 | 4.94E-03 | 0.0213 | (A) NE (B/D) |
| RPS6KB2 | 3.82 | 3.04 | 0.5 | 1.15 | 0.78 | 5.19E-03 | 0.0221 | (A) NE (B/D) |
| ENSA | 2.6 | 1.82 | 0.48 | 1.2 | 0.78 | 5.34E-03 | 0.0225 | (A) NE (B/D) |
| MAMDC2-AS | 1.07 | 0.29 | 0.63 | 0.3 | 0.78 | 5.78E-03 | 0.0239 | (A) NE (B/D) |
| CBX6 | 3.42 | 2.64 | 0.59 | 1.08 | 0.78 | 7.70E-03 | 0.0294 | (A) NE (B/D) |
| TBC1D2B | 2.13 | 1.35 | 0.66 | 0.71 | 0.78 | 8.10E-03 | 0.0305 | (A) NE (B/D) |

| | | | | | | | | |
|---|---|---|---|---|---|---|---|---|
| DNAJC19 | 2.53 | 1.75 | 0.64 | 0.92 | 0.78 | 8.65E-03 | 0.0321 | (A) NE (B/D) |
| C9ORF85 | 5.48 | 4.7 | 0.65 | 0.89 | 0.78 | 9.05E-03 | 0.0332 | (A) NE (B/D) |
| SGK3 | 3.02 | 2.25 | 0.64 | 0.93 | 0.78 | 9.13E-03 | 0.0334 | (A) NE (B/D) |
| RPL13AP5 | 8.59 | 7.81 | 0.47 | 1.4 | 0.78 | 9.42E-03 | 0.0342 | (A) NE (B/D) |
| IGBP1-AS2 | 6.99 | 6.2 | 0.31 | 1.58 | 0.78 | 0.0102 | 0.0363 | (A) NE (B/D) |
| TRAM2 | 1.78 | 1 | 0.59 | 1.2 | 0.78 | 0.0104 | 0.0368 | (A) NE (B/D) |
| RUSC1-AS1 | 7.84 | 7.05 | 0.64 | 1.07 | 0.78 | 0.0107 | 0.0375 | (A) NE (B/D) |
| KLF12 | 1.02 | 0.24 | 0.71 | 0.49 | 0.78 | 0.0115 | 0.0393 | (A) NE (B/D) |
| TRIM41 | 1.37 | 0.6 | 0.24 | 0.62 | 0.77 | 1.33E-06 | 3.73E-05 | (A) NE (B/D) |
| CYBC1 | 3.88 | 3.11 | 0.2 | 0.8 | 0.77 | 1.01E-05 | 1.82E-04 | (A) NE (B/D) |
| TRPT1 | 1.29 | 0.52 | 0.31 | 0.7 | 0.77 | 2.99E-05 | 4.22E-04 | (A) NE (B/D) |
| SDHAP1 | 7.89 | 7.13 | 0.36 | 0.54 | 0.77 | 7.55E-05 | 8.74E-04 | (A) NE (B/D) |
| TPT1-AS1 | 5.32 | 4.55 | 0.41 | 0.71 | 0.77 | 3.40E-04 | 2.78E-03 | (A) NE (B/D) |
| BICDL1 | 0.79 | 0.02 | 0.41 | 0.1 | 0.77 | 5.17E-04 | 3.83E-03 | (A) NE (B/D) |
| SRPRA | 5.06 | 4.3 | 0.28 | 1.09 | 0.77 | 7.08E-04 | 4.89E-03 | (A) NE (B/D) |
| KIAA0319L | 3.55 | 2.78 | 0.41 | 0.9 | 0.77 | 7.68E-04 | 5.20E-03 | (A) NE (B/D) |
| NUP214 | 2.46 | 1.7 | 0.43 | 0.91 | 0.77 | 1.18E-03 | 7.16E-03 | (A) NE (B/D) |
| TMEM199 | 1.29 | 0.53 | 0.5 | 0.6 | 0.77 | 1.45E-03 | 8.41E-03 | (A) NE (B/D) |
| TLE2 | 0.81 | 0.04 | 0.49 | 0.12 | 0.77 | 1.58E-03 | 8.94E-03 | (A) NE (B/D) |
| CLUH | 1.93 | 1.16 | 0.49 | 0.81 | 0.77 | 1.68E-03 | 9.40E-03 | (A) NE (B/D) |
| CCDC189 | 5.44 | 4.67 | 0.49 | 0.79 | 0.77 | 1.74E-03 | 9.64E-03 | (A) NE (B/D) |
| B4GALT3 | 5.93 | 5.16 | 0.36 | 1.14 | 0.77 | 1.74E-03 | 9.66E-03 | (A) NE (B/D) |
| ESPNP | 0.79 | 0.01 | 0.51 | 0.08 | 0.77 | 1.86E-03 | 0.0102 | (A) NE (B/D) |
| HRAS | 1.64 | 0.87 | 0.52 | 0.73 | 0.77 | 2.13E-03 | 0.0112 | (A) NE (B/D) |
| SURF4 | 1.62 | 0.85 | 0.51 | 0.9 | 0.77 | 2.71E-03 | 0.0135 | (A) NE (B/D) |
| ING3 | 0.9 | 0.14 | 0.55 | 0.31 | 0.77 | 2.79E-03 | 0.0138 | (A) NE (B/D) |
| PYURF | 5.82 | 5.06 | 0.38 | 1.29 | 0.77 | 4.44E-03 | 0.0197 | (A) NE (B/D) |
| GOLGA8B | 1.39 | 0.62 | 0.61 | 0.63 | 0.77 | 5.29E-03 | 0.0224 | (A) NE (B/D) |
| C12ORF73 | 3.17 | 2.4 | 0.46 | 1.24 | 0.77 | 5.92E-03 | 0.0244 | (A) NE (B/D) |
| COA4 | 2.7 | 1.93 | 0.47 | 1.29 | 0.77 | 6.85E-03 | 0.0271 | (A) NE (B/D) |
| COQ3 | 1.93 | 1.16 | 0.51 | 1.22 | 0.77 | 7.39E-03 | 0.0287 | (A) NE (B/D) |
| PDSS2 | 1.34 | 0.57 | 0.63 | 0.76 | 0.77 | 7.40E-03 | 0.0287 | (A) NE (B/D) |
| ZNF222 | 0.8 | 0.03 | 0.66 | 0.09 | 0.77 | 7.70E-03 | 0.0294 | (A) NE (B/D) |
| EGLN1 | 2.79 | 2.02 | 0.66 | 0.65 | 0.77 | 7.92E-03 | 0.0301 | (A) NE (B/D) |
| BCAP31 | 4.75 | 3.98 | 0.4 | 1.41 | 0.77 | 8.52E-03 | 0.0317 | (A) NE (B/D) |
| SNORD20 | 3.11 | 2.33 | 0.59 | 1.09 | 0.77 | 8.77E-03 | 0.0324 | (A) NE (B/D) |
| PRMT5-AS1 | 7.65 | 6.88 | 0.44 | 1.43 | 0.77 | 0.0105 | 0.037 | (A) NE (B/D) |
| MPP3 | 4.62 | 3.85 | 0.54 | 1.3 | 0.77 | 0.0116 | 0.0397 | (A) NE (B/D) |
| SNORD2 | 7.79 | 7.02 | 0.69 | 0.81 | 0.77 | 0.0124 | 0.0416 | (A) NE (B/D) |
| REX1BD | 1.55 | 0.79 | 0.31 | 0.58 | 0.76 | 1.77E-05 | 2.81E-04 | (A) NE (B/D) |
| SH3GL1P1 | 3.89 | 3.13 | 0.31 | 0.6 | 0.76 | 1.83E-05 | 2.88E-04 | (A) NE (B/D) |
| ARHGEF11 | 3.3 | 2.54 | 0.31 | 0.67 | 0.76 | 3.26E-05 | 4.54E-04 | (A) NE (B/D) |
| PTPN9 | 2.92 | 2.16 | 0.34 | 0.69 | 0.76 | 7.41E-05 | 8.61E-04 | (A) NE (B/D) |
| EZH1 | 5.5 | 4.73 | 0.26 | 0.87 | 0.76 | 7.64E-05 | 8.81E-04 | (A) NE (B/D) |
| KANSL2 | 5.15 | 4.39 | 0.22 | 0.91 | 0.76 | 7.74E-05 | 8.89E-04 | (A) NE (B/D) |
| LPCAT4 | 0.79 | 0.02 | 0.34 | 0.07 | 0.76 | 1.54E-04 | 1.51E-03 | (A) NE (B/D) |
| PRR12 | 1.43 | 0.67 | 0.39 | 0.57 | 0.76 | 1.61E-04 | 1.56E-03 | (A) NE (B/D) |
| COPS8P2 | 2.27 | 1.51 | 0.26 | 1.05 | 0.76 | 4.60E-04 | 3.51E-03 | (A) NE (B/D) |
| WIPI2 | 1.25 | 0.49 | 0.45 | 0.56 | 0.76 | 5.94E-04 | 4.28E-03 | (A) NE (B/D) |
| KDM2B | 5.5 | 4.74 | 0.38 | 0.9 | 0.76 | 5.97E-04 | 4.30E-03 | (A) NE (B/D) |
| FOXK2 | 1.26 | 0.5 | 0.45 | 0.59 | 0.76 | 7.06E-04 | 4.88E-03 | (A) NE (B/D) |
| MAPK9 | 1.59 | 0.82 | 0.44 | 0.78 | 0.76 | 8.05E-04 | 5.40E-03 | (A) NE (B/D) |
| AKT1 | 8.48 | 7.71 | 0.43 | 0.85 | 0.76 | 9.23E-04 | 5.96E-03 | (A) NE (B/D) |
| ATF4 | 7.26 | 6.5 | 0.39 | 1.06 | 0.76 | 1.63E-03 | 9.18E-03 | (A) NE (B/D) |
| ASS1P12 | 3.51 | 2.74 | 0.47 | 0.91 | 0.76 | 1.84E-03 | 0.01 | (A) NE (B/D) |
| SERF1B | 1.58 | 0.82 | 0.41 | 1.04 | 0.76 | 1.85E-03 | 0.0101 | (A) NE (B/D) |
| INTS4P1 | 1.23 | 0.48 | 0.51 | 0.39 | 0.76 | 1.91E-03 | 0.0104 | (A) NE (B/D) |
| RNF34 | 2.14 | 1.38 | 0.49 | 0.96 | 0.76 | 2.98E-03 | 0.0145 | (A) NE (B/D) |
| MIR-3124/31 | 1.49 | 0.73 | 0.56 | 0.67 | 0.76 | 3.61E-03 | 0.0168 | (A) NE (B/D) |
| MIR-3124/3P | 1.49 | 0.73 | 0.56 | 0.67 | 0.76 | 3.61E-03 | 0.0168 | (A) NE (B/D) |
| MIR-3124/5P | 1.49 | 0.73 | 0.56 | 0.67 | 0.76 | 3.61E-03 | 0.0168 | (A) NE (B/D) |
| STEAP3 | 1.14 | 0.37 | 0.59 | 0.74 | 0.76 | 4.95E-03 | 0.0213 | (A) NE (B/D) |
| WDR59 | 2.55 | 1.79 | 0.58 | 0.78 | 0.76 | 5.07E-03 | 0.0217 | (A) NE (B/D) |
| NDUFC2 | 4.65 | 3.9 | 0.24 | 1.44 | 0.76 | 5.74E-03 | 0.0238 | (A) NE (B/D) |
| CADM1 | 0.98 | 0.22 | 0.62 | 0.56 | 0.76 | 6.34E-03 | 0.0257 | (A) NE (B/D) |

| | | | | | | | | |
|---|---|---|---|---|---|---|---|---|
| MMP23B | 3.93 | 3.17 | 0.54 | 1.1 | 0.76 | 7.12E-03 | 0.0279 | (A) NE (B/D) |
| NDUFA3 | 0.99 | 0.23 | 0.64 | 0.36 | 0.76 | 7.33E-03 | 0.0285 | (A) NE (B/D) |
| ADH5P4 | 3.22 | 2.45 | 0.44 | 1.34 | 0.76 | 7.82E-03 | 0.0297 | (A) NE (B/D) |
| YDJC | 1.38 | 0.62 | 0.65 | 0.58 | 0.76 | 8.21E-03 | 0.0308 | (A) NE (B/D) |
| DAB2IP | 3.16 | 2.4 | 0.52 | 1.2 | 0.76 | 8.25E-03 | 0.031 | (A) NE (B/D) |
| IFNAR1 | 3.07 | 2.31 | 0.65 | 0.64 | 0.76 | 8.59E-03 | 0.0319 | (A) NE (B/D) |
| APH1B | 0.81 | 0.05 | 0.68 | 0.23 | 0.76 | 9.86E-03 | 0.0353 | (A) NE (B/D) |
| PFKFB2 | 1.52 | 0.76 | 0.68 | 0.58 | 0.76 | 0.0105 | 0.037 | (A) NE (B/D) |
| HSD17B7P2 | 1.16 | 0.4 | 0.69 | 0.53 | 0.76 | 0.0105 | 0.037 | (A) NE (B/D) |
| TBRG1 | 2.61 | 1.86 | 0.64 | 0.97 | 0.76 | 0.012 | 0.0406 | (A) NE (B/D) |
| EIF4A1P10 | 10.78 | 10.02 | 0.59 | 1.24 | 0.76 | 0.0146 | 0.0469 | (A) NE (B/D) |
| RPL9P7 | 9.39 | 8.62 | 0.41 | 1.57 | 0.76 | 0.0157 | 0.0496 | (A) NE (B/D) |
| ISL1 | 0.81 | 0.04 | 0.75 | 0.19 | 0.76 | 0.0158 | 0.0499 | (A) NE (B/D) |
| MPLKIP | 2.78 | 2.03 | 0.25 | 0.8 | 0.75 | 3.61E-05 | 4.89E-04 | (A) NE (B/D) |
| VPS39 | 7.91 | 7.16 | 0.3 | 0.87 | 0.75 | 1.68E-04 | 1.62E-03 | (A) NE (B/D) |
| IFT43 | 2.52 | 1.77 | 0.31 | 0.92 | 0.75 | 3.18E-04 | 2.64E-03 | (A) NE (B/D) |
| OGDH | 2.37 | 1.62 | 0.37 | 0.8 | 0.75 | 3.41E-04 | 2.78E-03 | (A) NE (B/D) |
| DFFA | 6.37 | 5.63 | 0.4 | 0.84 | 0.75 | 7.20E-04 | 4.95E-03 | (A) NE (B/D) |
| DLGAP4 | 0.76 | 0.02 | 0.43 | 0.05 | 0.75 | 7.73E-04 | 5.22E-03 | (A) NE (B/D) |
| TCF3 | 2.53 | 1.78 | 0.42 | 0.86 | 0.75 | 1.03E-03 | 6.45E-03 | (A) NE (B/D) |
| POLR2J2 | 8.19 | 7.44 | 0.32 | 1.06 | 0.75 | 1.06E-03 | 6.60E-03 | (A) NE (B/D) |
| HTRA2 | 3.83 | 3.08 | 0.27 | 1.15 | 0.75 | 1.28E-03 | 7.63E-03 | (A) NE (B/D) |
| SEC61A1 | 5.71 | 4.96 | 0.33 | 1.1 | 0.75 | 1.38E-03 | 8.09E-03 | (A) NE (B/D) |
| MAP3K4 | 4.36 | 3.62 | 0.49 | 0.59 | 0.75 | 1.40E-03 | 8.15E-03 | (A) NE (B/D) |
| YY1AP1 | 3.09 | 2.34 | 0.43 | 0.92 | 0.75 | 1.56E-03 | 8.90E-03 | (A) NE (B/D) |
| NIF3L1 | 1.77 | 1.02 | 0.42 | 0.99 | 0.75 | 1.79E-03 | 9.85E-03 | (A) NE (B/D) |
| LUC7L2 | 6.73 | 5.98 | 0.32 | 1.18 | 0.75 | 1.96E-03 | 0.0106 | (A) NE (B/D) |
| STX2 | 1.07 | 0.32 | 0.53 | 0.41 | 0.75 | 2.63E-03 | 0.0132 | (A) NE (B/D) |
| SIVA1 | 1.24 | 0.49 | 0.55 | 0.59 | 0.75 | 3.01E-03 | 0.0146 | (A) NE (B/D) |
| RBM5 | 6.16 | 5.41 | 0.5 | 0.9 | 0.75 | 3.06E-03 | 0.0148 | (A) NE (B/D) |
| NR2F6 | 1.27 | 0.52 | 0.59 | 0.35 | 0.75 | 4.83E-03 | 0.021 | (A) NE (B/D) |
| LONP1 | 0.84 | 0.09 | 0.59 | 0.15 | 0.75 | 4.89E-03 | 0.0211 | (A) NE (B/D) |
| RPL4 | 5.36 | 4.61 | 0.34 | 1.33 | 0.75 | 5.33E-03 | 0.0225 | (A) NE (B/D) |
| GLS | 0.81 | 0.06 | 0.64 | 0.19 | 0.75 | 7.41E-03 | 0.0287 | (A) NE (B/D) |
| MIR-3176/31 | 6.22 | 5.48 | 0.58 | 0.97 | 0.75 | 7.44E-03 | 0.0288 | (A) NE (B/D) |
| PHACTR1 | 1.27 | 0.52 | 0.64 | 0.44 | 0.75 | 7.79E-03 | 0.0297 | (A) NE (B/D) |
| COPS8 | 6.97 | 6.21 | 0.58 | 1.1 | 0.75 | 9.63E-03 | 0.0347 | (A) NE (B/D) |
| APBB3 | 0.87 | 0.12 | 0.67 | 0.27 | 0.75 | 0.01 | 0.0358 | (A) NE (B/D) |
| TRAF3IP1 | 11.13 | 10.39 | 0.62 | 0.92 | 0.75 | 0.0101 | 0.0358 | (A) NE (B/D) |
| GMPPB | 5.94 | 5.19 | 0.61 | 0.98 | 0.75 | 0.0101 | 0.0359 | (A) NE (B/D) |
| C3ORF14 | 0.86 | 0.12 | 0.68 | 0.28 | 0.75 | 0.0107 | 0.0374 | (A) NE (B/D) |
| TRUB2 | 5.86 | 5.11 | 0.56 | 1.23 | 0.75 | 0.0125 | 0.0419 | (A) NE (B/D) |
| PINK1-AS | 6.95 | 6.19 | 0.64 | 1.05 | 0.75 | 0.0131 | 0.0433 | (A) NE (B/D) |
| CAMTA1 | 0.83 | 0.08 | 0.72 | 0.2 | 0.75 | 0.0143 | 0.0463 | (A) NE (B/D) |
| PPP1R14A | 1.45 | 0.7 | 0.68 | 0.97 | 0.75 | 0.0153 | 0.0487 | (A) NE (B/D) |
| KMT2E-AS1 | 4.7 | 3.96 | 0.19 | 0.63 | 0.74 | 6.97E-07 | 2.23E-05 | (A) NE (B/D) |
| CDC5L | 3.75 | 3.01 | 0.29 | 0.68 | 0.74 | 2.89E-05 | 4.10E-04 | (A) NE (B/D) |
| IAH1 | 6.23 | 5.49 | 0.23 | 0.81 | 0.74 | 3.29E-05 | 4.56E-04 | (A) NE (B/D) |
| SNX21 | 1.38 | 0.64 | 0.34 | 0.74 | 0.74 | 1.58E-04 | 1.54E-03 | (A) NE (B/D) |
| SCAMP2 | 2.54 | 1.8 | 0.3 | 0.96 | 0.74 | 4.27E-04 | 3.31E-03 | (A) NE (B/D) |
| SUMO3 | 2.83 | 2.09 | 0.2 | 1.07 | 0.74 | 4.79E-04 | 3.62E-03 | (A) NE (B/D) |
| GPATCH3 | 6.84 | 6.1 | 0.27 | 1.17 | 0.74 | 1.66E-03 | 9.29E-03 | (A) NE (B/D) |
| PDE6D | 1.55 | 0.81 | 0.49 | 0.75 | 0.74 | 1.92E-03 | 0.0104 | (A) NE (B/D) |
| PAAF1 | 7.23 | 6.49 | 0.39 | 1.06 | 0.74 | 2.08E-03 | 0.011 | (A) NE (B/D) |
| SHPK | 1.72 | 0.98 | 0.49 | 0.85 | 0.74 | 2.58E-03 | 0.0131 | (A) NE (B/D) |
| USP24 | 0.93 | 0.19 | 0.55 | 0.34 | 0.74 | 3.35E-03 | 0.0159 | (A) NE (B/D) |
| VN1R81P | 0.79 | 0.05 | 0.55 | 0.13 | 0.74 | 3.72E-03 | 0.0172 | (A) NE (B/D) |
| IPO5 | 0.9 | 0.16 | 0.56 | 0.32 | 0.74 | 3.75E-03 | 0.0173 | (A) NE (B/D) |
| SLC10A3 | 1.94 | 1.19 | 0.42 | 1.14 | 0.74 | 3.80E-03 | 0.0175 | (A) NE (B/D) |
| METTL23 | 7.49 | 6.75 | 0.48 | 1.03 | 0.74 | 4.08E-03 | 0.0185 | (A) NE (B/D) |
| ZNF232 | 0.9 | 0.16 | 0.58 | 0.32 | 0.74 | 4.98E-03 | 0.0214 | (A) NE (B/D) |
| DCLK2 | 0.77 | 0.03 | 0.59 | 0.1 | 0.74 | 5.37E-03 | 0.0226 | (A) NE (B/D) |
| CDIPT | 6.51 | 5.76 | 0.52 | 1.03 | 0.74 | 5.79E-03 | 0.024 | (A) NE (B/D) |
| COG4 | 1.92 | 1.18 | 0.53 | 0.99 | 0.74 | 5.92E-03 | 0.0244 | (A) NE (B/D) |
| EEF1A1 | 7.62 | 6.88 | 0.31 | 1.39 | 0.74 | 6.94E-03 | 0.0274 | (A) NE (B/D) |

| | | | | | | | | |
|---|---|---|---|---|---|---|---|---|
| MMADHC | 3.63 | 2.89 | 0.51 | 1.12 | 0.74 | 7.09E-03 | 0.0278 | (A) NE (B/D) |
| HDAC2 | 1.5 | 0.76 | 0.6 | 0.75 | 0.74 | 7.13E-03 | 0.0279 | (A) NE (B/D) |
| CTNNBIP1 | 6.06 | 5.32 | 0.51 | 1.16 | 0.74 | 7.93E-03 | 0.0301 | (A) NE (B/D) |
| PLXNB2 | 4.58 | 3.84 | 0.6 | 0.92 | 0.74 | 8.98E-03 | 0.033 | (A) NE (B/D) |
| HNRNPUL2-E | 7.37 | 6.63 | 0.56 | 1.07 | 0.74 | 9.41E-03 | 0.0341 | (A) NE (B/D) |
| TAF10 | 3.76 | 3.02 | 0.48 | 1.27 | 0.74 | 9.89E-03 | 0.0354 | (A) NE (B/D) |
| RRAGC | 2.6 | 1.86 | 0.56 | 1.09 | 0.74 | 0.0101 | 0.0361 | (A) NE (B/D) |
| RABL6 | 0.89 | 0.14 | 0.67 | 0.34 | 0.74 | 0.0105 | 0.0369 | (A) NE (B/D) |
| ATG5 | 3.23 | 2.49 | 0.62 | 1.01 | 0.74 | 0.012 | 0.0407 | (A) NE (B/D) |
| DHRS13 | 3.38 | 2.64 | 0.27 | 1.58 | 0.74 | 0.0134 | 0.0442 | (A) NE (B/D) |
| RNU6-850P | 6.16 | 5.42 | 0.57 | 1.22 | 0.74 | 0.0135 | 0.0445 | (A) NE (B/D) |
| DIDO1 | 1.37 | 0.64 | 0.25 | 0.45 | 0.73 | 1.25E-06 | 3.53E-05 | (A) NE (B/D) |
| COBL | 1.54 | 0.81 | 0.3 | 0.73 | 0.73 | 6.83E-05 | 8.07E-04 | (A) NE (B/D) |
| DGKZ | 1.5 | 0.78 | 0.34 | 0.58 | 0.73 | 9.37E-05 | 1.03E-03 | (A) NE (B/D) |
| GTF3C3 | 1 | 0.27 | 0.38 | 0.37 | 0.73 | 2.03E-04 | 1.86E-03 | (A) NE (B/D) |
| IP6K1 | 1.74 | 1.01 | 0.36 | 0.77 | 0.73 | 2.82E-04 | 2.40E-03 | (A) NE (B/D) |
| DPF2 | 2.52 | 1.79 | 0.36 | 1.05 | 0.73 | 1.66E-03 | 9.29E-03 | (A) NE (B/D) |
| MTMR1 | 6.62 | 5.89 | 0.4 | 1.01 | 0.73 | 2.13E-03 | 0.0112 | (A) NE (B/D) |
| COX7A2L | 5.14 | 4.41 | 0.31 | 1.17 | 0.73 | 2.28E-03 | 0.0118 | (A) NE (B/D) |
| ZBED5 | 6.2 | 5.48 | 0.45 | 0.98 | 0.73 | 3.14E-03 | 0.0151 | (A) NE (B/D) |
| PYY2 | 0.87 | 0.14 | 0.54 | 0.32 | 0.73 | 3.49E-03 | 0.0164 | (A) NE (B/D) |
| RECQL4 | 1.1 | 0.37 | 0.56 | 0.54 | 0.73 | 4.14E-03 | 0.0187 | (A) NE (B/D) |
| RBM8A | 2.46 | 1.73 | 0.39 | 1.19 | 0.73 | 4.55E-03 | 0.0201 | (A) NE (B/D) |
| PRPF38AP2 | 3.91 | 3.18 | 0.42 | 1.23 | 0.73 | 6.39E-03 | 0.0258 | (A) NE (B/D) |
| CICP14 | 4.53 | 3.8 | 0.6 | 0.62 | 0.73 | 6.50E-03 | 0.0261 | (A) NE (B/D) |
| WDR5 | 1.52 | 0.79 | 0.59 | 0.8 | 0.73 | 7.54E-03 | 0.029 | (A) NE (B/D) |
| CBX1 | 8.13 | 7.4 | 0.34 | 1.36 | 0.73 | 7.68E-03 | 0.0294 | (A) NE (B/D) |
| SNORD102 | 6.2 | 5.47 | 0.6 | 0.77 | 0.73 | 7.73E-03 | 0.0295 | (A) NE (B/D) |
| VTI1BP2 | 1.75 | 1.02 | 0.61 | 0.74 | 0.73 | 8.20E-03 | 0.0308 | (A) NE (B/D) |
| CA5BP1 | 3.31 | 2.59 | 0.46 | 1.25 | 0.73 | 8.63E-03 | 0.032 | (A) NE (B/D) |
| MED18 | 5.84 | 5.12 | 0.56 | 1.02 | 0.73 | 8.72E-03 | 0.0323 | (A) NE (B/D) |
| AIFM1 | 6.42 | 5.69 | 0.44 | 1.3 | 0.73 | 9.28E-03 | 0.0338 | (A) NE (B/D) |
| H2AFY | 2.18 | 1.46 | 0.61 | 0.84 | 0.73 | 9.78E-03 | 0.0351 | (A) NE (B/D) |
| ZNF528 | 0.85 | 0.11 | 0.66 | 0.28 | 0.73 | 9.99E-03 | 0.0357 | (A) NE (B/D) |
| C9ORF78 | 5.87 | 5.13 | 0.29 | 1.5 | 0.73 | 0.0109 | 0.0379 | (A) NE (B/D) |
| RABGGTB | 2.99 | 2.26 | 0.64 | 0.88 | 0.73 | 0.0119 | 0.0405 | (A) NE (B/D) |
| PLRG1 | 3.06 | 2.33 | 0.5 | 1.26 | 0.73 | 0.0119 | 0.0406 | (A) NE (B/D) |
| UBB | 6.04 | 5.3 | 0.21 | 1.59 | 0.73 | 0.0131 | 0.0435 | (A) NE (B/D) |
| LINC00363 | 0.96 | 0.23 | 0.69 | 0.54 | 0.73 | 0.0136 | 0.0447 | (A) NE (B/D) |
| PPIAP19 | 1.31 | 0.58 | 0.69 | 0.63 | 0.73 | 0.0145 | 0.0468 | (A) NE (B/D) |
| CRCP | 3.3 | 2.58 | 0.12 | 0.58 | 0.72 | 3.99E-08 | 2.31E-06 | (A) NE (B/D) |
| DYNLT1 | 1.71 | 0.99 | 0.23 | 0.72 | 0.72 | 1.47E-05 | 2.46E-04 | (A) NE (B/D) |
| WDR31 | 1.3 | 0.58 | 0.16 | 0.81 | 0.72 | 1.98E-05 | 3.06E-04 | (A) NE (B/D) |
| LRRC37A16P | 6.3 | 5.58 | 0.1 | 0.86 | 0.72 | 2.96E-05 | 4.18E-04 | (A) NE (B/D) |
| CSNK1G2 | 0.77 | 0.05 | 0.3 | 0.15 | 0.72 | 7.09E-05 | 8.30E-04 | (A) NE (B/D) |
| NCBP1 | 6.36 | 5.65 | 0.29 | 0.81 | 0.72 | 1.35E-04 | 1.36E-03 | (A) NE (B/D) |
| RABIF | 2.31 | 1.59 | 0.33 | 0.84 | 0.72 | 3.51E-04 | 2.84E-03 | (A) NE (B/D) |
| COA1 | 0.84 | 0.12 | 0.43 | 0.21 | 0.72 | 9.00E-04 | 5.84E-03 | (A) NE (B/D) |
| CSRNP2 | 5.34 | 4.62 | 0.33 | 0.98 | 0.72 | 1.02E-03 | 6.39E-03 | (A) NE (B/D) |
| EIPR1 | 6.99 | 6.27 | 0.43 | 0.75 | 0.72 | 1.13E-03 | 6.94E-03 | (A) NE (B/D) |
| NPLOC4 | 1.63 | 0.92 | 0.41 | 0.83 | 0.72 | 1.22E-03 | 7.34E-03 | (A) NE (B/D) |
| RASA4DP | 0.82 | 0.1 | 0.45 | 0.22 | 0.72 | 1.28E-03 | 7.61E-03 | (A) NE (B/D) |
| CLK3 | 1.93 | 1.22 | 0.43 | 0.8 | 0.72 | 1.35E-03 | 7.95E-03 | (A) NE (B/D) |
| C12ORF76 | 6.19 | 5.47 | 0.45 | 0.83 | 0.72 | 1.77E-03 | 9.76E-03 | (A) NE (B/D) |
| TTC3-AS1 | 5.72 | 5 | 0.49 | 0.78 | 0.72 | 2.50E-03 | 0.0127 | (A) NE (B/D) |
| RCE1 | 1.24 | 0.53 | 0.5 | 0.62 | 0.72 | 2.58E-03 | 0.013 | (A) NE (B/D) |
| ANKEF1 | 6.27 | 5.55 | 0.47 | 0.98 | 0.72 | 4.29E-03 | 0.0192 | (A) NE (B/D) |
| PHYKPL | 2.53 | 1.81 | 0.55 | 0.6 | 0.72 | 4.44E-03 | 0.0197 | (A) NE (B/D) |
| SAXO2 | 0.75 | 0.03 | 0.59 | 0.12 | 0.72 | 6.40E-03 | 0.0258 | (A) NE (B/D) |
| FBXW4P1 | 2 | 1.28 | 0.58 | 0.65 | 0.72 | 6.48E-03 | 0.026 | (A) NE (B/D) |
| CUEDC2 | 5.45 | 4.73 | 0.35 | 1.35 | 0.72 | 8.41E-03 | 0.0314 | (A) NE (B/D) |
| WIZ | 0.83 | 0.11 | 0.62 | 0.29 | 0.72 | 8.42E-03 | 0.0314 | (A) NE (B/D) |
| GPBP1L1 | 1.52 | 0.8 | 0.64 | 0.55 | 0.72 | 9.45E-03 | 0.0342 | (A) NE (B/D) |
| SGSM3 | 3.1 | 2.38 | 0.53 | 1.1 | 0.72 | 9.67E-03 | 0.0348 | (A) NE (B/D) |
| RBM18 | 3.8 | 3.08 | 0.57 | 1.04 | 0.72 | 0.011 | 0.038 | (A) NE (B/D) |

| | | | | | | | | |
|---|---|---|---|---|---|---|---|---|
| POLR2F | 4.95 | 4.23 | 0.51 | 1.18 | 0.72 | 0.011 | 0.0381 | (A) NE (B/D) |
| MIR-138-1/1 | 0.77 | 0.05 | 0.67 | 0.19 | 0.72 | 0.0123 | 0.0416 | (A) NE (B/D) |
| MIR-138-1/1 | 0.77 | 0.05 | 0.67 | 0.19 | 0.72 | 0.0123 | 0.0416 | (A) NE (B/D) |
| MIR-138-1/3 | 0.77 | 0.05 | 0.67 | 0.19 | 0.72 | 0.0123 | 0.0416 | (A) NE (B/D) |
| MIR-138-1/5 | 0.77 | 0.05 | 0.67 | 0.19 | 0.72 | 0.0123 | 0.0416 | (A) NE (B/D) |
| TP53I3 | 0.8 | 0.08 | 0.68 | 0.22 | 0.72 | 0.0129 | 0.0429 | (A) NE (B/D) |
| PHIP | 1.2 | 0.48 | 0.69 | 0.4 | 0.72 | 0.0144 | 0.0464 | (A) NE (B/D) |
| CSE1L | 2.38 | 1.66 | 0.61 | 1.1 | 0.72 | 0.0153 | 0.0486 | (A) NE (B/D) |
| ANKS1A | 0.94 | 0.23 | 0.18 | 0.34 | 0.71 | 1.04E-08 | 8.23E-07 | (A) NE (B/D) |
| TBC1D9B | 1.22 | 0.51 | 0.14 | 0.62 | 0.71 | 3.05E-07 | 1.17E-05 | (A) NE (B/D) |
| MTHFS | 1.03 | 0.32 | 0.3 | 0.46 | 0.71 | 2.20E-05 | 3.29E-04 | (A) NE (B/D) |
| ST6GALNAC2 | 4.58 | 3.87 | 0.24 | 0.85 | 0.71 | 1.14E-04 | 1.20E-03 | (A) NE (B/D) |
| PTPRA | 1.74 | 1.03 | 0.34 | 0.73 | 0.71 | 2.39E-04 | 2.11E-03 | (A) NE (B/D) |
| ARL8A | 5.82 | 5.11 | 0.16 | 0.98 | 0.71 | 2.61E-04 | 2.25E-03 | (A) NE (B/D) |
| POR | 7.55 | 6.84 | 0.25 | 0.94 | 0.71 | 3.43E-04 | 2.79E-03 | (A) NE (B/D) |
| RNF168 | 2.63 | 1.91 | 0.39 | 0.54 | 0.71 | 3.57E-04 | 2.88E-03 | (A) NE (B/D) |
| KDM5C | 6.41 | 5.7 | 0.33 | 0.92 | 0.71 | 7.32E-04 | 5.00E-03 | (A) NE (B/D) |
| MRPL30 | 2.67 | 1.96 | 0.33 | 0.99 | 0.71 | 1.16E-03 | 7.07E-03 | (A) NE (B/D) |
| PSMC3IP | 1.28 | 0.57 | 0.46 | 0.71 | 0.71 | 1.88E-03 | 0.0102 | (A) NE (B/D) |
| TOMM20L | 2.99 | 2.28 | 0.39 | 1 | 0.71 | 2.25E-03 | 0.0117 | (A) NE (B/D) |
| ALG9 | 0.77 | 0.06 | 0.5 | 0.15 | 0.71 | 2.45E-03 | 0.0125 | (A) NE (B/D) |
| UPK3BL1 | 1.86 | 1.15 | 0.41 | 1.03 | 0.71 | 3.27E-03 | 0.0156 | (A) NE (B/D) |
| TSC1 | 1.17 | 0.46 | 0.52 | 0.62 | 0.71 | 3.59E-03 | 0.0167 | (A) NE (B/D) |
| CDC42BPA | 5.39 | 4.68 | 0.52 | 0.75 | 0.71 | 4.10E-03 | 0.0185 | (A) NE (B/D) |
| EIF4A1P2 | 7.83 | 7.12 | 0.37 | 1.17 | 0.71 | 4.43E-03 | 0.0196 | (A) NE (B/D) |
| ACP1 | 2.1 | 1.39 | 0.55 | 0.82 | 0.71 | 6.50E-03 | 0.0261 | (A) NE (B/D) |
| CNTNAP3 | 0.95 | 0.24 | 0.6 | 0.51 | 0.71 | 7.48E-03 | 0.0289 | (A) NE (B/D) |
| NDUFV2P1 | 5.19 | 4.48 | 0.29 | 1.39 | 0.71 | 8.53E-03 | 0.0317 | (A) NE (B/D) |
| ARMH4 | 0.9 | 0.19 | 0.62 | 0.4 | 0.71 | 9.18E-03 | 0.0335 | (A) NE (B/D) |
| PCDHA10 | 0.85 | 0.14 | 0.63 | 0.29 | 0.71 | 9.47E-03 | 0.0343 | (A) NE (B/D) |
| SCNM1 | 5.44 | 4.72 | 0.49 | 1.18 | 0.71 | 9.68E-03 | 0.0349 | (A) NE (B/D) |
| CBX3 | 3.12 | 2.41 | 0.45 | 1.33 | 0.71 | 0.0127 | 0.0425 | (A) NE (B/D) |
| ZSCAN31 | 0.71 | 0.01 | 0.69 | 0.03 | 0.71 | 0.0153 | 0.0488 | (A) NE (B/D) |
| MIDN | 1.2 | 0.5 | 0.25 | 0.32 | 0.7 | 2.87E-06 | 6.77E-05 | (A) NE (B/D) |
| BAZ1B | 1.77 | 1.07 | 0.21 | 0.68 | 0.7 | 6.94E-06 | 1.36E-04 | (A) NE (B/D) |
| ZSCAN30 | 3.66 | 2.95 | 0.3 | 0.75 | 0.7 | 1.32E-04 | 1.34E-03 | (A) NE (B/D) |
| COPS7B | 1.53 | 0.83 | 0.25 | 0.84 | 0.7 | 1.50E-04 | 1.48E-03 | (A) NE (B/D) |
| RBMX | 1.56 | 0.86 | 0.4 | 0.63 | 0.7 | 5.28E-04 | 3.89E-03 | (A) NE (B/D) |
| STARD7 | 5.16 | 4.45 | 0.25 | 0.99 | 0.7 | 6.22E-04 | 4.42E-03 | (A) NE (B/D) |
| BAG5 | 2.01 | 1.31 | 0.33 | 0.89 | 0.7 | 7.06E-04 | 4.88E-03 | (A) NE (B/D) |
| TMEM189-U | 7.91 | 7.21 | 0.19 | 1.05 | 0.7 | 7.93E-04 | 5.34E-03 | (A) NE (B/D) |
| ANKRD20A2 | 1.59 | 0.9 | 0.41 | 0.83 | 0.7 | 1.43E-03 | 8.31E-03 | (A) NE (B/D) |
| OGFR | 3.18 | 2.48 | 0.41 | 0.87 | 0.7 | 1.67E-03 | 9.34E-03 | (A) NE (B/D) |
| SRSF1 | 2.39 | 1.69 | 0.33 | 1.03 | 0.7 | 1.79E-03 | 9.87E-03 | (A) NE (B/D) |
| DCAF16 | 2.74 | 2.04 | 0.47 | 0.68 | 0.7 | 2.10E-03 | 0.0111 | (A) NE (B/D) |
| ZSWIM8 | 5.97 | 5.27 | 0.44 | 0.84 | 0.7 | 2.22E-03 | 0.0116 | (A) NE (B/D) |
| UBA2 | 1.04 | 0.34 | 0.49 | 0.56 | 0.7 | 2.44E-03 | 0.0125 | (A) NE (B/D) |
| TBCCD1 | 1.47 | 0.78 | 0.48 | 0.79 | 0.7 | 3.42E-03 | 0.0161 | (A) NE (B/D) |
| LRRC37BP1 | 6.13 | 5.43 | 0.44 | 0.97 | 0.7 | 3.47E-03 | 0.0163 | (A) NE (B/D) |
| UNC50 | 1.6 | 0.9 | 0.47 | 0.93 | 0.7 | 3.94E-03 | 0.018 | (A) NE (B/D) |
| MTA1 | 0.98 | 0.28 | 0.54 | 0.41 | 0.7 | 4.10E-03 | 0.0185 | (A) NE (B/D) |
| SPTBN1 | 3.26 | 2.55 | 0.41 | 1.12 | 0.7 | 5.07E-03 | 0.0217 | (A) NE (B/D) |
| PRICKLE2 | 1.03 | 0.34 | 0.55 | 0.54 | 0.7 | 5.14E-03 | 0.022 | (A) NE (B/D) |
| SNORD16 | 4.45 | 3.75 | 0.49 | 1 | 0.7 | 6.37E-03 | 0.0257 | (A) NE (B/D) |
| NPTN | 1.15 | 0.45 | 0.56 | 0.76 | 0.7 | 6.70E-03 | 0.0267 | (A) NE (B/D) |
| C22ORF46 | 6.97 | 6.28 | 0.43 | 1.15 | 0.7 | 6.98E-03 | 0.0275 | (A) NE (B/D) |
| MSANTD2 | 0.82 | 0.12 | 0.59 | 0.33 | 0.7 | 7.36E-03 | 0.0286 | (A) NE (B/D) |
| TBC1D26 | 0.73 | 0.03 | 0.6 | 0.15 | 0.7 | 7.86E-03 | 0.0299 | (A) NE (B/D) |
| GGN | 0.91 | 0.21 | 0.63 | 0.34 | 0.7 | 0.0101 | 0.0361 | (A) NE (B/D) |
| EIF3EP1 | 7.35 | 6.65 | 0.55 | 1.03 | 0.7 | 0.0106 | 0.0373 | (A) NE (B/D) |
| KIAA0753 | 1.8 | 1.1 | 0.63 | 0.52 | 0.7 | 0.0109 | 0.0378 | (A) NE (B/D) |
| MRFAP1L1 | 4.11 | 3.41 | 0.5 | 1.18 | 0.7 | 0.0126 | 0.0423 | (A) NE (B/D) |
| WRNIP1 | 4.3 | 3.6 | 0.61 | 0.9 | 0.7 | 0.013 | 0.043 | (A) NE (B/D) |
| TOMM7 | 2.7 | 2 | 0.35 | 1.45 | 0.7 | 0.0145 | 0.0468 | (A) NE (B/D) |
| PTAR1 | 1.07 | 0.37 | 0.68 | 0.45 | 0.7 | 0.0152 | 0.0485 | (A) NE (B/D) |

| | | | | | | | | |
|---|---|---|---|---|---|---|---|---|
| CPSF4 | 1.35 | 0.67 | 0.13 | 0.44 | 0.69 | 9.52E-10 | 1.32E-07 | (A) NE (B/D) |
| ZRSR2P1 | 2.76 | 2.07 | 0.31 | 0.76 | 0.69 | 2.18E-04 | 1.96E-03 | (A) NE (B/D) |
| TBC1D3 | 1.59 | 0.9 | 0.27 | 0.92 | 0.69 | 4.64E-04 | 3.53E-03 | (A) NE (B/D) |
| MAP3K3 | 1.54 | 0.86 | 0.4 | 0.53 | 0.69 | 6.33E-04 | 4.49E-03 | (A) NE (B/D) |
| CHURC1-FNT | 6.68 | 5.99 | 0.27 | 0.96 | 0.69 | 6.66E-04 | 4.66E-03 | (A) NE (B/D) |
| UBE2V1 | 2.19 | 1.49 | 0.23 | 1.05 | 0.69 | 9.76E-04 | 6.21E-03 | (A) NE (B/D) |
| SENP3 | 1.27 | 0.58 | 0.43 | 0.63 | 0.69 | 1.07E-03 | 6.62E-03 | (A) NE (B/D) |
| SCGB2B2 | 0.76 | 0.06 | 0.48 | 0.15 | 0.69 | 2.37E-03 | 0.0122 | (A) NE (B/D) |
| SERINC3 | 1.73 | 1.04 | 0.45 | 0.81 | 0.69 | 2.66E-03 | 0.0133 | (A) NE (B/D) |
| KYAT1 | 1.24 | 0.55 | 0.49 | 0.76 | 0.69 | 3.38E-03 | 0.016 | (A) NE (B/D) |
| LYPLA2P1 | 2.88 | 2.19 | 0.39 | 1.1 | 0.69 | 4.58E-03 | 0.0202 | (A) NE (B/D) |
| CSTF2 | 2.6 | 1.92 | 0.3 | 1.21 | 0.69 | 4.77E-03 | 0.0208 | (A) NE (B/D) |
| AUTS2 | 1.1 | 0.41 | 0.54 | 0.47 | 0.69 | 4.86E-03 | 0.021 | (A) NE (B/D) |
| SF1 | 4.64 | 3.94 | 0.39 | 1.13 | 0.69 | 5.17E-03 | 0.022 | (A) NE (B/D) |
| RAB1A | 2.53 | 1.84 | 0.39 | 1.16 | 0.69 | 5.94E-03 | 0.0244 | (A) NE (B/D) |
| PDZD11 | 1.79 | 1.1 | 0.26 | 1.31 | 0.69 | 6.31E-03 | 0.0256 | (A) NE (B/D) |
| TATDN2P2 | 1.74 | 1.04 | 0.56 | 0.67 | 0.69 | 6.47E-03 | 0.026 | (A) NE (B/D) |
| IRX2 | 0.87 | 0.18 | 0.59 | 0.35 | 0.69 | 7.93E-03 | 0.0301 | (A) NE (B/D) |
| SIPA1L2 | 2.38 | 1.69 | 0.51 | 1.01 | 0.69 | 8.56E-03 | 0.0318 | (A) NE (B/D) |
| USE1 | 0.91 | 0.22 | 0.61 | 0.42 | 0.69 | 9.56E-03 | 0.0345 | (A) NE (B/D) |
| DGUOK-AS1 | 6.74 | 6.05 | 0.52 | 1.06 | 0.69 | 0.0101 | 0.036 | (A) NE (B/D) |
| CBY1 | 2.01 | 1.32 | 0.39 | 1.28 | 0.69 | 0.0102 | 0.0362 | (A) NE (B/D) |
| PTPA | 6.32 | 5.62 | 0.47 | 1.2 | 0.69 | 0.0113 | 0.0388 | (A) NE (B/D) |
| BAIAP2-DT | 5.46 | 4.77 | 0.55 | 1.02 | 0.69 | 0.0119 | 0.0404 | (A) NE (B/D) |
| COX6C | 2.1 | 1.41 | 0.42 | 1.32 | 0.69 | 0.0129 | 0.043 | (A) NE (B/D) |
| RPS7P4 | 5.51 | 4.82 | 0.4 | 1.36 | 0.69 | 0.0138 | 0.045 | (A) NE (B/D) |
| ADAM1A | 2.51 | 1.82 | 0.63 | 0.81 | 0.69 | 0.0151 | 0.0483 | (A) NE (B/D) |
| TBRG4 | 1.03 | 0.35 | 0.08 | 0.51 | 0.68 | 6.73E-09 | 5.81E-07 | (A) NE (B/D) |
| GNA11 | 0.74 | 0.06 | 0.21 | 0.14 | 0.68 | 2.60E-06 | 6.29E-05 | (A) NE (B/D) |
| YBEY | 2.45 | 1.77 | 0.12 | 0.77 | 0.68 | 1.82E-05 | 2.87E-04 | (A) NE (B/D) |
| ABI2 | 0.92 | 0.23 | 0.29 | 0.45 | 0.68 | 2.14E-05 | 3.23E-04 | (A) NE (B/D) |
| JMJD7 | 2.25 | 1.56 | 0.22 | 0.84 | 0.68 | 1.25E-04 | 1.29E-03 | (A) NE (B/D) |
| ZNF335 | 2.12 | 1.44 | 0.32 | 0.71 | 0.68 | 2.02E-04 | 1.85E-03 | (A) NE (B/D) |
| COQ9 | 1.82 | 1.14 | 0.34 | 0.66 | 0.68 | 2.71E-04 | 2.32E-03 | (A) NE (B/D) |
| MTG2 | 6.19 | 5.51 | 0.28 | 0.87 | 0.68 | 3.80E-04 | 3.03E-03 | (A) NE (B/D) |
| RNASEH2C | 1.29 | 0.61 | 0.37 | 0.58 | 0.68 | 3.91E-04 | 3.10E-03 | (A) NE (B/D) |
| AP1M1 | 0.7 | 0.02 | 0.4 | 0.08 | 0.68 | 8.68E-04 | 5.68E-03 | (A) NE (B/D) |
| PRUNE1 | 1.71 | 1.03 | 0.31 | 0.95 | 0.68 | 9.73E-04 | 6.19E-03 | (A) NE (B/D) |
| MED23 | 1.9 | 1.22 | 0.4 | 0.78 | 0.68 | 1.27E-03 | 7.59E-03 | (A) NE (B/D) |
| TOR1B | 2.93 | 2.25 | 0.22 | 1.09 | 0.68 | 1.53E-03 | 8.74E-03 | (A) NE (B/D) |
| GNPTG | 6.78 | 6.1 | 0.3 | 1.01 | 0.68 | 1.61E-03 | 9.08E-03 | (A) NE (B/D) |
| RANBP9 | 2.53 | 1.85 | 0.37 | 0.91 | 0.68 | 1.67E-03 | 9.34E-03 | (A) NE (B/D) |
| RNPEP | 2.44 | 1.76 | 0.39 | 0.93 | 0.68 | 2.27E-03 | 0.0118 | (A) NE (B/D) |
| ABCB6 | 0.87 | 0.2 | 0.47 | 0.48 | 0.68 | 2.32E-03 | 0.012 | (A) NE (B/D) |
| FAM220A | 2.27 | 1.59 | 0.36 | 1.06 | 0.68 | 3.78E-03 | 0.0174 | (A) NE (B/D) |
| NOP58 | 2.26 | 1.58 | 0.36 | 1.12 | 0.68 | 4.39E-03 | 0.0195 | (A) NE (B/D) |
| SNORA66 | 6.87 | 6.19 | 0.48 | 0.89 | 0.68 | 4.77E-03 | 0.0208 | (A) NE (B/D) |
| UBA6 | 0.9 | 0.22 | 0.53 | 0.41 | 0.68 | 4.86E-03 | 0.021 | (A) NE (B/D) |
| LCMT1 | 3.52 | 2.84 | 0.41 | 1.08 | 0.68 | 5.23E-03 | 0.0222 | (A) NE (B/D) |
| E2F4 | 3.21 | 2.53 | 0.4 | 1.14 | 0.68 | 6.61E-03 | 0.0264 | (A) NE (B/D) |
| TUBA1C | 1.3 | 0.62 | 0.53 | 0.8 | 0.68 | 6.63E-03 | 0.0265 | (A) NE (B/D) |
| TOMM34 | 6.51 | 5.84 | 0.42 | 1.11 | 0.68 | 6.97E-03 | 0.0275 | (A) NE (B/D) |
| BBS5 | 0.79 | 0.1 | 0.57 | 0.19 | 0.68 | 7.13E-03 | 0.0279 | (A) NE (B/D) |
| HLTF-AS1 | 3.78 | 3.09 | 0.47 | 1.03 | 0.68 | 7.25E-03 | 0.0283 | (A) NE (B/D) |
| NAA38 | 2 | 1.32 | 0.4 | 1.22 | 0.68 | 9.65E-03 | 0.0348 | (A) NE (B/D) |
| SEPTIN2 | 4.71 | 4.03 | 0.55 | 0.97 | 0.68 | 0.0109 | 0.0378 | (A) NE (B/D) |
| ATG4A | 5.41 | 4.73 | 0.42 | 1.24 | 0.68 | 0.0111 | 0.0385 | (A) NE (B/D) |
| USP9YP14 | 0.72 | 0.04 | 0.62 | 0.16 | 0.68 | 0.0112 | 0.0386 | (A) NE (B/D) |
| ZC3H8 | 1.11 | 0.43 | 0.63 | 0.58 | 0.68 | 0.0129 | 0.0429 | (A) NE (B/D) |
| RAB24 | 1.33 | 0.65 | 0.61 | 0.77 | 0.68 | 0.0133 | 0.0438 | (A) NE (B/D) |
| THBS3 | 6.32 | 5.64 | 0.45 | 1.29 | 0.68 | 0.0158 | 0.0498 | (A) NE (B/D) |
| USP21 | 1.19 | 0.52 | 0.3 | 0.65 | 0.67 | 1.29E-04 | 1.31E-03 | (A) NE (B/D) |
| VIPAS39 | 4.65 | 3.99 | 0.31 | 0.86 | 0.67 | 6.87E-04 | 4.78E-03 | (A) NE (B/D) |
| DOP1B | 5.77 | 5.11 | 0.4 | 0.88 | 0.67 | 2.48E-03 | 0.0126 | (A) NE (B/D) |
| AGAP5 | 5.42 | 4.75 | 0.42 | 0.87 | 0.67 | 2.75E-03 | 0.0137 | (A) NE (B/D) |

| | | | | | | | | |
|---|---|---|---|---|---|---|---|---|
| GEMIN8 | 1.41 | 0.74 | 0.47 | 0.69 | 0.67 | 2.98E-03 | 0.0145 | (A) NE (B/D) |
| RECQL5 | 1.55 | 0.88 | 0.48 | 0.54 | 0.67 | 3.00E-03 | 0.0146 | (A) NE (B/D) |
| NKAP | 4.99 | 4.32 | 0.35 | 1.05 | 0.67 | 3.40E-03 | 0.0161 | (A) NE (B/D) |
| TBC1D25 | 3.26 | 2.58 | 0.38 | 1.05 | 0.67 | 4.32E-03 | 0.0193 | (A) NE (B/D) |
| ZNF672 | 2.03 | 1.37 | 0.51 | 0.61 | 0.67 | 4.50E-03 | 0.0199 | (A) NE (B/D) |
| ORC2 | 0.67 | 0 | 0.52 | 0.01 | 0.67 | 4.67E-03 | 0.0204 | (A) NE (B/D) |
| CCDC9 | 1.02 | 0.35 | 0.53 | 0.43 | 0.67 | 4.86E-03 | 0.021 | (A) NE (B/D) |
| VOPP1 | 1.03 | 0.36 | 0.52 | 0.63 | 0.67 | 5.26E-03 | 0.0223 | (A) NE (B/D) |
| MORC2-AS1 | 6.37 | 5.71 | 0.43 | 1.03 | 0.67 | 6.16E-03 | 0.0251 | (A) NE (B/D) |
| SNHG17 | 1.77 | 1.11 | 0.54 | 0.7 | 0.67 | 6.89E-03 | 0.0273 | (A) NE (B/D) |
| DPH3P1 | 5.65 | 4.98 | 0.52 | 0.88 | 0.67 | 7.56E-03 | 0.0291 | (A) NE (B/D) |
| CHAMP1 | 5.59 | 4.92 | 0.45 | 1.08 | 0.67 | 8.82E-03 | 0.0326 | (A) NE (B/D) |
| YTHDF2 | 7.87 | 7.2 | 0.52 | 0.98 | 0.67 | 0.0112 | 0.0386 | (A) NE (B/D) |
| MOB3C | 4.13 | 3.45 | 0.5 | 1.08 | 0.67 | 0.0113 | 0.039 | (A) NE (B/D) |
| BAIAP2L1 | 3.16 | 2.48 | 0.35 | 1.35 | 0.67 | 0.0129 | 0.0428 | (A) NE (B/D) |
| SOD1 | 4.44 | 3.76 | 0.25 | 1.43 | 0.67 | 0.0135 | 0.0444 | (A) NE (B/D) |
| PDIA3P1 | 9.4 | 8.73 | 0.49 | 1.18 | 0.67 | 0.0142 | 0.046 | (A) NE (B/D) |
| COASY | 7.31 | 6.64 | 0.54 | 1.06 | 0.67 | 0.0143 | 0.0463 | (A) NE (B/D) |
| GSTK1 | 9.25 | 8.58 | 0.49 | 1.19 | 0.67 | 0.0152 | 0.0485 | (A) NE (B/D) |
| EIF2B3 | 2.93 | 2.26 | 0.52 | 1.15 | 0.67 | 0.0155 | 0.0492 | (A) NE (B/D) |
| AKIRIN2 | 1.05 | 0.4 | 0.24 | 0.48 | 0.66 | 5.39E-06 | 1.12E-04 | (A) NE (B/D) |
| NONO | 1.01 | 0.34 | 0.21 | 0.63 | 0.66 | 6.53E-06 | 1.31E-04 | (A) NE (B/D) |
| ZNF445 | 4.72 | 4.05 | 0.32 | 0.59 | 0.66 | 1.50E-04 | 1.48E-03 | (A) NE (B/D) |
| ZFP62 | 5.51 | 4.85 | 0.29 | 0.74 | 0.66 | 2.48E-04 | 2.17E-03 | (A) NE (B/D) |
| LMAN2L | 1.15 | 0.49 | 0.37 | 0.57 | 0.66 | 4.67E-04 | 3.54E-03 | (A) NE (B/D) |
| NUB1 | 7.29 | 6.63 | 0.28 | 0.9 | 0.66 | 7.43E-04 | 5.06E-03 | (A) NE (B/D) |
| BICRA | 0.8 | 0.14 | 0.39 | 0.22 | 0.66 | 7.70E-04 | 5.21E-03 | (A) NE (B/D) |
| ERCC3 | 1.28 | 0.62 | 0.36 | 0.71 | 0.66 | 7.86E-04 | 5.30E-03 | (A) NE (B/D) |
| ZBTB17 | 5.9 | 5.25 | 0.37 | 0.78 | 0.66 | 1.13E-03 | 6.93E-03 | (A) NE (B/D) |
| RTL6 | 1 | 0.34 | 0.42 | 0.51 | 0.66 | 1.14E-03 | 6.98E-03 | (A) NE (B/D) |
| DHX9 | 3.59 | 2.92 | 0.29 | 0.98 | 0.66 | 1.38E-03 | 8.06E-03 | (A) NE (B/D) |
| SUN2 | 2.48 | 1.82 | 0.31 | 0.96 | 0.66 | 1.61E-03 | 9.08E-03 | (A) NE (B/D) |
| CROCC | 3.75 | 3.08 | 0.39 | 0.89 | 0.66 | 2.43E-03 | 0.0124 | (A) NE (B/D) |
| CCT8 | 2.87 | 2.21 | 0.3 | 1.08 | 0.66 | 3.20E-03 | 0.0153 | (A) NE (B/D) |
| GLB1 | 1.65 | 1 | 0.4 | 0.98 | 0.66 | 4.44E-03 | 0.0197 | (A) NE (B/D) |
| MAT2A | 2.81 | 2.15 | 0.34 | 1.13 | 0.66 | 5.44E-03 | 0.0228 | (A) NE (B/D) |
| ACTBP11 | 3.8 | 3.14 | 0.44 | 0.96 | 0.66 | 5.99E-03 | 0.0246 | (A) NE (B/D) |
| GPLD1 | 0.72 | 0.06 | 0.54 | 0.15 | 0.66 | 6.22E-03 | 0.0253 | (A) NE (B/D) |
| MYLIP | 3.35 | 2.68 | 0.36 | 1.14 | 0.66 | 6.25E-03 | 0.0254 | (A) NE (B/D) |
| RXYLT1-AS1 | 2.03 | 1.37 | 0.38 | 1.12 | 0.66 | 6.51E-03 | 0.0261 | (A) NE (B/D) |
| ARHGEF7-AS | 0.86 | 0.2 | 0.55 | 0.49 | 0.66 | 7.16E-03 | 0.028 | (A) NE (B/D) |
| UTP14A | 4.77 | 4.12 | 0.48 | 0.91 | 0.66 | 7.17E-03 | 0.028 | (A) NE (B/D) |
| PGM2L1 | 0.74 | 0.08 | 0.56 | 0.18 | 0.66 | 7.79E-03 | 0.0297 | (A) NE (B/D) |
| FBXO46 | 0.82 | 0.16 | 0.57 | 0.36 | 0.66 | 8.05E-03 | 0.0304 | (A) NE (B/D) |
| RANBP1 | 1.35 | 0.69 | 0.55 | 0.67 | 0.66 | 8.29E-03 | 0.0311 | (A) NE (B/D) |
| CHMP1B | 4.1 | 3.44 | 0.47 | 1.02 | 0.66 | 9.49E-03 | 0.0343 | (A) NE (B/D) |
| KRIT1 | 0.84 | 0.18 | 0.59 | 0.31 | 0.66 | 9.81E-03 | 0.0352 | (A) NE (B/D) |
| TXNDC15 | 1.76 | 1.11 | 0.52 | 1.03 | 0.66 | 0.0133 | 0.0439 | (A) NE (B/D) |
| NFATC2IP | 2.83 | 2.17 | 0.54 | 1.01 | 0.66 | 0.0135 | 0.0444 | (A) NE (B/D) |
| MAPK3 | 4.42 | 3.76 | 0.5 | 1.1 | 0.66 | 0.0137 | 0.045 | (A) NE (B/D) |
| KSR1 | 1.28 | 0.62 | 0.62 | 0.59 | 0.66 | 0.0144 | 0.0465 | (A) NE (B/D) |
| CFAP300 | 0.97 | 0.3 | 0.64 | 0.5 | 0.66 | 0.0148 | 0.0474 | (A) NE (B/D) |
| ENTR1 | 0.99 | 0.35 | 0.26 | 0.38 | 0.65 | 1.06E-05 | 1.91E-04 | (A) NE (B/D) |
| TIMM21 | 1.51 | 0.86 | 0.17 | 0.85 | 0.65 | 1.64E-04 | 1.58E-03 | (A) NE (B/D) |
| EIF4A1 | 3.06 | 2.41 | 0.26 | 0.84 | 0.65 | 3.96E-04 | 3.13E-03 | (A) NE (B/D) |
| INTS14 | 2.68 | 2.03 | 0.26 | 0.86 | 0.65 | 4.85E-04 | 3.65E-03 | (A) NE (B/D) |
| CHMP4BP1 | 4.36 | 3.71 | 0.31 | 0.9 | 0.65 | 1.30E-03 | 7.72E-03 | (A) NE (B/D) |
| ZNF746 | 6.58 | 5.93 | 0.42 | 0.53 | 0.65 | 1.41E-03 | 8.18E-03 | (A) NE (B/D) |
| C15ORF62 | 0.69 | 0.05 | 0.44 | 0.14 | 0.65 | 2.19E-03 | 0.0114 | (A) NE (B/D) |
| PFDN1 | 4.46 | 3.81 | 0.3 | 1 | 0.65 | 2.23E-03 | 0.0116 | (A) NE (B/D) |
| C1ORF115 | 1.6 | 0.95 | 0.26 | 1.12 | 0.65 | 3.72E-03 | 0.0172 | (A) NE (B/D) |
| GATD1 | 8.4 | 7.74 | 0.43 | 0.89 | 0.65 | 4.33E-03 | 0.0193 | (A) NE (B/D) |
| PFDN6 | 4.23 | 3.57 | 0.22 | 1.22 | 0.65 | 5.56E-03 | 0.0232 | (A) NE (B/D) |
| RNF126 | 0.71 | 0.06 | 0.52 | 0.25 | 0.65 | 5.67E-03 | 0.0236 | (A) NE (B/D) |
| MAP2 | 0.67 | 0.02 | 0.53 | 0.09 | 0.65 | 6.68E-03 | 0.0266 | (A) NE (B/D) |

| | | | | | | | | |
|---|---|---|---|---|---|---|---|---|
| TMEM60 | 2.7 | 2.05 | 0.42 | 1.04 | 0.65 | 7.39E-03 | 0.0287 | (A) NE (B/D) |
| SPDL1 | 0.74 | 0.09 | 0.55 | 0.22 | 0.65 | 7.42E-03 | 0.0288 | (A) NE (B/D) |
| NDUFAF4 | 1.22 | 0.57 | 0.54 | 0.65 | 0.65 | 7.65E-03 | 0.0293 | (A) NE (B/D) |
| PTER | 1.19 | 0.54 | 0.56 | 0.48 | 0.65 | 8.13E-03 | 0.0306 | (A) NE (B/D) |
| ZNF200 | 3.08 | 2.43 | 0.54 | 0.71 | 0.65 | 8.84E-03 | 0.0326 | (A) NE (B/D) |
| RBM8B | 8.29 | 7.64 | 0.28 | 1.27 | 0.65 | 9.06E-03 | 0.0332 | (A) NE (B/D) |
| ATP5MF-PTC | 8.42 | 7.77 | 0.36 | 1.22 | 0.65 | 9.72E-03 | 0.035 | (A) NE (B/D) |
| SDCBPP3 | 3.98 | 3.33 | 0.47 | 1.02 | 0.65 | 9.81E-03 | 0.0352 | (A) NE (B/D) |
| SDHAP2 | 8.12 | 7.48 | 0.57 | 0.66 | 0.65 | 0.0108 | 0.0377 | (A) NE (B/D) |
| CNBP | 11.52 | 10.87 | 0.26 | 1.33 | 0.65 | 0.0111 | 0.0384 | (A) NE (B/D) |
| THAP2 | 4.49 | 3.84 | 0.6 | 0.42 | 0.65 | 0.0121 | 0.0409 | (A) NE (B/D) |
| GOLGA8K | 1.22 | 0.57 | 0.62 | 0.59 | 0.65 | 0.0147 | 0.0472 | (A) NE (B/D) |
| ZNF250 | 3.91 | 3.27 | 0.53 | 1.03 | 0.65 | 0.0157 | 0.0498 | (A) NE (B/D) |
| RFC5 | 1.56 | 0.92 | 0.23 | 0.81 | 0.64 | 2.22E-04 | 1.99E-03 | (A) NE (B/D) |
| TNIP1 | 2.95 | 2.31 | 0.35 | 0.73 | 0.64 | 7.22E-04 | 4.96E-03 | (A) NE (B/D) |
| C2ORF15 | 7.83 | 7.18 | 0.33 | 0.83 | 0.64 | 1.04E-03 | 6.51E-03 | (A) NE (B/D) |
| TMEM189 | 4.93 | 4.29 | 0.2 | 0.98 | 0.64 | 1.09E-03 | 6.72E-03 | (A) NE (B/D) |
| KLF7 | 1.72 | 1.07 | 0.4 | 0.62 | 0.64 | 1.35E-03 | 7.92E-03 | (A) NE (B/D) |
| FAM122A | 4.42 | 3.78 | 0.27 | 0.96 | 0.64 | 1.47E-03 | 8.50E-03 | (A) NE (B/D) |
| SUSD1 | 3.28 | 2.64 | 0.41 | 0.73 | 0.64 | 2.10E-03 | 0.0111 | (A) NE (B/D) |
| ZNF749 | 0.68 | 0.03 | 0.44 | 0.14 | 0.64 | 2.14E-03 | 0.0113 | (A) NE (B/D) |
| MIR-4775/47 | 5.11 | 4.47 | 0.26 | 1.04 | 0.64 | 2.35E-03 | 0.0121 | (A) NE (B/D) |
| ZNF496 | 0.66 | 0.03 | 0.45 | 0.11 | 0.64 | 2.87E-03 | 0.0141 | (A) NE (B/D) |
| STK32C | 0.88 | 0.24 | 0.51 | 0.32 | 0.64 | 5.44E-03 | 0.0229 | (A) NE (B/D) |
| RMND1 | 9.83 | 9.18 | 0.39 | 1.01 | 0.64 | 5.55E-03 | 0.0232 | (A) NE (B/D) |
| ZNF491 | 0.66 | 0.02 | 0.55 | 0.07 | 0.64 | 8.17E-03 | 0.0307 | (A) NE (B/D) |
| MARCH6 | 1.61 | 0.97 | 0.55 | 0.62 | 0.64 | 9.32E-03 | 0.0339 | (A) NE (B/D) |
| SBDS | 2.22 | 1.58 | 0.32 | 1.22 | 0.64 | 9.84E-03 | 0.0353 | (A) NE (B/D) |
| RPTOR | 0.65 | 0.01 | 0.58 | 0.05 | 0.64 | 0.0106 | 0.0372 | (A) NE (B/D) |
| ANKHD1-EIF4 | 8.27 | 7.63 | 0.58 | 0.56 | 0.64 | 0.0114 | 0.0392 | (A) NE (B/D) |
| VPS4A | 2.43 | 1.79 | 0.5 | 0.96 | 0.64 | 0.012 | 0.0408 | (A) NE (B/D) |
| NDUFB3 | 7.24 | 6.6 | 0.37 | 1.27 | 0.64 | 0.0143 | 0.0463 | (A) NE (B/D) |
| RBM17 | 7.15 | 6.51 | 0.49 | 1.09 | 0.64 | 0.0146 | 0.047 | (A) NE (B/D) |
| KAT8 | 1.24 | 0.61 | 0.27 | 0.61 | 0.63 | 6.70E-05 | 7.94E-04 | (A) NE (B/D) |
| HM13-AS1 | 1.96 | 1.32 | 0.22 | 0.83 | 0.63 | 3.11E-04 | 2.59E-03 | (A) NE (B/D) |
| SLC23A3 | 7.29 | 6.65 | 0.2 | 0.89 | 0.63 | 4.99E-04 | 3.74E-03 | (A) NE (B/D) |
| GPN1 | 2.39 | 1.76 | 0.27 | 0.89 | 0.63 | 1.02E-03 | 6.40E-03 | (A) NE (B/D) |
| PET117 | 6.79 | 6.16 | 0.28 | 0.88 | 0.63 | 1.02E-03 | 6.43E-03 | (A) NE (B/D) |
| ZSWIM8-AS1 | 8.62 | 7.99 | 0.38 | 0.9 | 0.63 | 3.33E-03 | 0.0158 | (A) NE (B/D) |
| ORC3 | 1.55 | 0.93 | 0.46 | 0.64 | 0.63 | 4.14E-03 | 0.0187 | (A) NE (B/D) |
| FMC1-LUC7L | 2.24 | 1.6 | 0.36 | 1.08 | 0.63 | 6.83E-03 | 0.0271 | (A) NE (B/D) |
| TSGA10 | 0.67 | 0.04 | 0.55 | 0.25 | 0.63 | 8.41E-03 | 0.0314 | (A) NE (B/D) |
| ZNF862 | 1.83 | 1.2 | 0.51 | 0.87 | 0.63 | 0.0105 | 0.0369 | (A) NE (B/D) |
| PGAM2 | 1.17 | 0.54 | 0.56 | 0.65 | 0.63 | 0.0118 | 0.0403 | (A) NE (B/D) |
| CMTM1 | 2.52 | 1.89 | 0.5 | 0.95 | 0.63 | 0.0119 | 0.0405 | (A) NE (B/D) |
| MAPK8 | 6.14 | 5.52 | 0.55 | 0.7 | 0.63 | 0.012 | 0.0406 | (A) NE (B/D) |
| C7ORF50 | 1.63 | 1 | 0.51 | 0.92 | 0.63 | 0.0131 | 0.0435 | (A) NE (B/D) |
| TRRAP | 5.29 | 4.68 | 0.18 | 0.57 | 0.62 | 3.08E-06 | 7.16E-05 | (A) NE (B/D) |
| GATAD2B | 2.23 | 1.61 | 0.24 | 0.52 | 0.62 | 1.52E-05 | 2.52E-04 | (A) NE (B/D) |
| ZDHHC16 | 2.92 | 2.31 | 0.21 | 0.89 | 0.62 | 7.13E-04 | 4.91E-03 | (A) NE (B/D) |
| R3HDM2 | 2.06 | 1.44 | 0.37 | 0.68 | 0.62 | 1.35E-03 | 7.93E-03 | (A) NE (B/D) |
| TAF1C | 1.42 | 0.8 | 0.41 | 0.53 | 0.62 | 1.76E-03 | 9.73E-03 | (A) NE (B/D) |
| MARK4 | 0.71 | 0.09 | 0.42 | 0.25 | 0.62 | 2.05E-03 | 0.0109 | (A) NE (B/D) |
| MTF1 | 2.78 | 2.15 | 0.44 | 0.62 | 0.62 | 3.03E-03 | 0.0147 | (A) NE (B/D) |
| CXORF58 | 1.02 | 0.4 | 0.41 | 0.76 | 0.62 | 3.37E-03 | 0.016 | (A) NE (B/D) |
| CPT2 | 7.85 | 7.23 | 0.25 | 1.05 | 0.62 | 3.45E-03 | 0.0163 | (A) NE (B/D) |
| FTLP12 | 0.73 | 0.11 | 0.48 | 0.47 | 0.62 | 4.53E-03 | 0.02 | (A) NE (B/D) |
| MGME1 | 5.7 | 5.08 | 0.3 | 1.08 | 0.62 | 5.32E-03 | 0.0225 | (A) NE (B/D) |
| SDCBPP2 | 3.22 | 2.6 | 0.46 | 0.94 | 0.62 | 9.27E-03 | 0.0338 | (A) NE (B/D) |
| ZNF23 | 0.65 | 0.03 | 0.56 | 0.15 | 0.62 | 0.0102 | 0.0362 | (A) NE (B/D) |
| ZBTB7A | 1.42 | 0.79 | 0.56 | 0.6 | 0.62 | 0.0117 | 0.0399 | (A) NE (B/D) |
| MVD | 0.84 | 0.22 | 0.58 | 0.4 | 0.62 | 0.0131 | 0.0435 | (A) NE (B/D) |
| GTPBP3 | 0.82 | 0.21 | 0.24 | 0.31 | 0.61 | 7.61E-06 | 1.47E-04 | (A) NE (B/D) |
| DDX42 | 1.33 | 0.72 | 0.22 | 0.78 | 0.61 | 2.78E-04 | 2.37E-03 | (A) NE (B/D) |
| STAT2 | 3.08 | 2.47 | 0.27 | 0.85 | 0.61 | 1.04E-03 | 6.48E-03 | (A) NE (B/D) |

| | | | | | | | | |
|---|---|---|---|---|---|---|---|---|
| TMEM43 | 1.99 | 1.38 | 0.25 | 0.92 | 0.61 | 1.47E-03 | 8.47E-03 | (A) NE (B/D) |
| NCKAP5L | 5.15 | 4.53 | 0.37 | 0.77 | 0.61 | 2.11E-03 | 0.0112 | (A) NE (B/D) |
| MOV10 | 5.7 | 5.09 | 0.2 | 1.06 | 0.61 | 2.95E-03 | 0.0144 | (A) NE (B/D) |
| WASF3 | 1.86 | 1.25 | 0.43 | 0.68 | 0.61 | 3.34E-03 | 0.0159 | (A) NE (B/D) |
| GPX7 | 0.65 | 0.04 | 0.47 | 0.15 | 0.61 | 4.23E-03 | 0.019 | (A) NE (B/D) |
| SUDS3P1 | 4.27 | 3.67 | 0.45 | 0.6 | 0.61 | 4.35E-03 | 0.0194 | (A) NE (B/D) |
| KIAA1191 | 1.99 | 1.38 | 0.43 | 0.85 | 0.61 | 6.19E-03 | 0.0252 | (A) NE (B/D) |
| FLCN | 0.88 | 0.27 | 0.51 | 0.49 | 0.61 | 7.20E-03 | 0.0281 | (A) NE (B/D) |
| TCHP | 2.99 | 2.38 | 0.37 | 1.04 | 0.61 | 7.70E-03 | 0.0294 | (A) NE (B/D) |
| CAMTA1-IT1 | 0.93 | 0.33 | 0.52 | 0.44 | 0.61 | 7.97E-03 | 0.0302 | (A) NE (B/D) |
| BARD1 | 0.64 | 0.02 | 0.53 | 0.14 | 0.61 | 8.79E-03 | 0.0325 | (A) NE (B/D) |
| RGP1 | 8.16 | 7.56 | 0.36 | 1.09 | 0.61 | 9.27E-03 | 0.0338 | (A) NE (B/D) |
| PLA2G6 | 0.83 | 0.22 | 0.54 | 0.42 | 0.61 | 9.46E-03 | 0.0342 | (A) NE (B/D) |
| FAM207A | 4.12 | 3.51 | 0.53 | 0.54 | 0.61 | 9.61E-03 | 0.0347 | (A) NE (B/D) |
| TTK | 0.66 | 0.05 | 0.55 | 0.19 | 0.61 | 0.0109 | 0.038 | (A) NE (B/D) |
| RWDD2B | 2.28 | 1.67 | 0.3 | 1.25 | 0.61 | 0.0128 | 0.0428 | (A) NE (B/D) |
| IPP | 3.92 | 3.31 | 0.56 | 0.69 | 0.61 | 0.0143 | 0.0463 | (A) NE (B/D) |
| RALGPS1 | 0.71 | 0.1 | 0.32 | 0.23 | 0.6 | 3.41E-04 | 2.78E-03 | (A) NE (B/D) |
| RAB26 | 1.22 | 0.62 | 0.32 | 0.58 | 0.6 | 4.03E-04 | 3.17E-03 | (A) NE (B/D) |
| SMARCD1 | 1.15 | 0.56 | 0.32 | 0.6 | 0.6 | 4.89E-04 | 3.68E-03 | (A) NE (B/D) |
| BMERB1 | 0.83 | 0.23 | 0.35 | 0.42 | 0.6 | 5.53E-04 | 4.05E-03 | (A) NE (B/D) |
| SNORD51 | 0.72 | 0.12 | 0.36 | 0.16 | 0.6 | 8.83E-04 | 5.76E-03 | (A) NE (B/D) |
| TAPBP | 10.11 | 9.51 | 0.26 | 0.84 | 0.6 | 9.78E-04 | 6.21E-03 | (A) NE (B/D) |
| PRELID1 | 3.51 | 2.91 | 0.2 | 0.93 | 0.6 | 1.15E-03 | 7.04E-03 | (A) NE (B/D) |
| ANKRD42 | 2.48 | 1.88 | 0.33 | 0.73 | 0.6 | 1.18E-03 | 7.19E-03 | (A) NE (B/D) |
| CTNND1 | 6.43 | 5.83 | 0.23 | 0.96 | 0.6 | 1.90E-03 | 0.0103 | (A) NE (B/D) |
| RBAK | 6.83 | 6.23 | 0.39 | 0.63 | 0.6 | 1.97E-03 | 0.0106 | (A) NE (B/D) |
| ARHGAP19-S | 0.72 | 0.12 | 0.42 | 0.43 | 0.6 | 2.40E-03 | 0.0123 | (A) NE (B/D) |
| ZNF513 | 1.12 | 0.52 | 0.42 | 0.61 | 0.6 | 2.89E-03 | 0.0142 | (A) NE (B/D) |
| UBE2F-SCLY | 7.21 | 6.61 | 0.3 | 0.96 | 0.6 | 3.49E-03 | 0.0164 | (A) NE (B/D) |
| EPC2 | 0.68 | 0.08 | 0.46 | 0.19 | 0.6 | 4.45E-03 | 0.0197 | (A) NE (B/D) |
| AHCYL2 | 3.13 | 2.53 | 0.43 | 0.72 | 0.6 | 5.01E-03 | 0.0215 | (A) NE (B/D) |
| POLR1C | 0.91 | 0.3 | 0.48 | 0.52 | 0.6 | 5.37E-03 | 0.0226 | (A) NE (B/D) |
| FAM209A | 6.31 | 5.71 | 0.26 | 1.13 | 0.6 | 7.54E-03 | 0.029 | (A) NE (B/D) |
| ZFP14 | 0.73 | 0.12 | 0.51 | 0.18 | 0.6 | 7.70E-03 | 0.0294 | (A) NE (B/D) |
| WTAP | 2.14 | 1.55 | 0.45 | 0.91 | 0.6 | 0.0103 | 0.0364 | (A) NE (B/D) |
| SENP1 | 5.65 | 5.05 | 0.52 | 0.76 | 0.6 | 0.013 | 0.0432 | (A) NE (B/D) |
| NCBP2 | 1.59 | 0.99 | 0.46 | 1 | 0.6 | 0.0138 | 0.0451 | (A) NE (B/D) |
| NOL12 | 3.11 | 2.51 | 0.52 | 0.8 | 0.6 | 0.014 | 0.0455 | (A) NE (B/D) |
| BCDIN3D-AS1 | 5.69 | 5.09 | 0.43 | 1.11 | 0.6 | 0.0157 | 0.0496 | (A) NE (B/D) |
| CLASP1 | 6.9 | 6.31 | 0.19 | 0.46 | 0.59 | 1.19E-06 | 3.40E-05 | (A) NE (B/D) |
| FAM27E4 | 0.81 | 0.22 | 0.32 | 0.47 | 0.59 | 3.19E-04 | 2.64E-03 | (A) NE (B/D) |
| POLR2J3 | 3.67 | 3.08 | 0.18 | 0.82 | 0.59 | 4.48E-04 | 3.44E-03 | (A) NE (B/D) |
| COX5A | 1.23 | 0.64 | 0.3 | 0.73 | 0.59 | 7.57E-04 | 5.14E-03 | (A) NE (B/D) |
| SEC31A | 4.04 | 3.45 | 0.32 | 0.68 | 0.59 | 7.69E-04 | 5.20E-03 | (A) NE (B/D) |
| TRIM32 | 2.06 | 1.47 | 0.34 | 0.73 | 0.59 | 1.50E-03 | 8.64E-03 | (A) NE (B/D) |
| SEH1L | 0.75 | 0.16 | 0.4 | 0.35 | 0.59 | 1.73E-03 | 9.59E-03 | (A) NE (B/D) |
| EIF4ENIF1 | 5.88 | 5.29 | 0.34 | 0.74 | 0.59 | 1.73E-03 | 9.59E-03 | (A) NE (B/D) |
| ZNF561-AS1 | 0.78 | 0.19 | 0.42 | 0.43 | 0.59 | 2.48E-03 | 0.0126 | (A) NE (B/D) |
| SSX2IP | 0.65 | 0.06 | 0.42 | 0.27 | 0.59 | 2.82E-03 | 0.0139 | (A) NE (B/D) |
| LETMD1 | 1.66 | 1.07 | 0.41 | 0.87 | 0.59 | 7.05E-03 | 0.0277 | (A) NE (B/D) |
| SLC27A1 | 0.66 | 0.07 | 0.54 | 0.26 | 0.59 | 0.0103 | 0.0364 | (A) NE (B/D) |
| C2CD2L | 3.45 | 2.87 | 0.42 | 0.96 | 0.59 | 0.0109 | 0.0379 | (A) NE (B/D) |
| ERLIN1 | 7.01 | 6.42 | 0.47 | 0.85 | 0.59 | 0.0109 | 0.038 | (A) NE (B/D) |
| UBAP2L | 1.85 | 1.25 | 0.47 | 0.97 | 0.59 | 0.0141 | 0.0457 | (A) NE (B/D) |
| EXOSC1 | 5.04 | 4.45 | 0.41 | 1.07 | 0.59 | 0.0149 | 0.0477 | (A) NE (B/D) |
| POM121L4P | 0.04 | 0.63 | 0.11 | 0.68 | -0.59 | 2.61E-05 | 3.77E-04 | (B) Pathology_classA-RNA |
| MIR-5009/3P | 0.03 | 0.62 | 0.1 | 0.79 | -0.59 | 1.58E-04 | 1.54E-03 | (B) Pathology_classA-RNA |
| MIR-5009/5P | 0.03 | 0.62 | 0.1 | 0.79 | -0.59 | 1.58E-04 | 1.54E-03 | (B) Pathology_classA-RNA |
| TRDMT1 | 0.19 | 0.78 | 0.3 | 0.66 | -0.59 | 5.26E-04 | 3.88E-03 | (B) Pathology_classA-RNA |
| RN7SL306P | 0.14 | 0.73 | 0.21 | 0.85 | -0.59 | 7.52E-04 | 5.11E-03 | (B) Pathology_classA-RNA |
| MIPOL1 | 0.32 | 0.91 | 0.21 | 0.88 | -0.59 | 9.28E-04 | 5.96E-03 | (B) Pathology_classA-RNA |
| ANKRD33B-A | 0.02 | 0.61 | 0.06 | 0.94 | -0.59 | 9.35E-04 | 6.00E-03 | (B) Pathology_classA-RNA |
| RPS15AP34 | 0.04 | 0.63 | 0.12 | 0.93 | -0.59 | 1.00E-03 | 6.31E-03 | (B) Pathology_classA-RNA |
| GREB1 | 0.02 | 0.6 | 0.05 | 0.99 | -0.59 | 1.54E-03 | 8.77E-03 | (B) Pathology_classA-RNA |

| | | | | | | | | |
|---|---|---|---|---|---|---|---|---|
| USP17L7 | 0.02 | 0.61 | 0.05 | 1.02 | -0.59 | 1.87E-03 | 0.0102 | (B) Pathology_classA-RNA |
| IFI44L | 0 | 0.6 | 0.01 | 1.04 | -0.59 | 2.04E-03 | 0.0109 | (B) Pathology_classA-RNA |
| HNRNPDLP4 | 0.09 | 0.68 | 0.28 | 0.96 | -0.59 | 3.42E-03 | 0.0161 | (B) Pathology_classA-RNA |
| IZUMO3 | 0.01 | 0.6 | 0.03 | 1.14 | -0.59 | 5.01E-03 | 0.0215 | (B) Pathology_classA-RNA |
| RN7SL449P | 0.24 | 0.83 | 0.35 | 0.96 | -0.59 | 5.78E-03 | 0.024 | (B) Pathology_classA-RNA |
| ZNF385A | 0.15 | 0.74 | 0.37 | 0.98 | -0.59 | 7.04E-03 | 0.0277 | (B) Pathology_classA-RNA |
| MIR-569/569 | 0.15 | 0.74 | 0.24 | 1.18 | -0.59 | 9.22E-03 | 0.0336 | (B) Pathology_classA-RNA |
| PARVB | 0.83 | 1.42 | 0.49 | 0.71 | -0.59 | 9.27E-03 | 0.0338 | (B) Pathology_classA-RNA |
| PRRG3 | 0.17 | 0.76 | 0.5 | 0.68 | -0.59 | 0.0103 | 0.0363 | (B) Pathology_classA-RNA |
| QKI | 0.07 | 0.67 | 0.14 | 0.39 | -0.6 | 5.23E-09 | 4.77E-07 | (B) Pathology_classA-RNA |
| HLA-DRB9 | 0.05 | 0.65 | 0.11 | 0.81 | -0.6 | 1.51E-04 | 1.49E-03 | (B) Pathology_classA-RNA |
| HMGN1P10 | 0.01 | 0.61 | 0.03 | 0.88 | -0.6 | 4.14E-04 | 3.24E-03 | (B) Pathology_classA-RNA |
| SIGLEC22P | 0.01 | 0.6 | 0.02 | 0.93 | -0.6 | 6.97E-04 | 4.84E-03 | (B) Pathology_classA-RNA |
| FAM118A | 0.06 | 0.66 | 0.17 | 0.92 | -0.6 | 8.61E-04 | 5.64E-03 | (B) Pathology_classA-RNA |
| MRPL45P2 | 4.72 | 5.32 | 0.36 | 0.82 | -0.6 | 3.00E-03 | 0.0146 | (B) Pathology_classA-RNA |
| PPFIBP1 | 0.25 | 0.86 | 0.37 | 0.87 | -0.6 | 3.90E-03 | 0.0178 | (B) Pathology_classA-RNA |
| BMS1P4 | 2.44 | 3.04 | 0.46 | 0.54 | -0.6 | 4.80E-03 | 0.0209 | (B) Pathology_classA-RNA |
| MUC2 | 0.01 | 0.61 | 0.02 | 1.23 | -0.6 | 7.26E-03 | 0.0283 | (B) Pathology_classA-RNA |
| MBNL1 | 0.49 | 1.09 | 0.44 | 0.85 | -0.6 | 7.56E-03 | 0.0291 | (B) Pathology_classA-RNA |
| RNU7-4P | 0.08 | 0.68 | 0.24 | 1.3 | -0.6 | 0.015 | 0.048 | (B) Pathology_classA-RNA |
| RYKP1 | 3.85 | 4.45 | 0.55 | 0.74 | -0.6 | 0.0156 | 0.0494 | (B) Pathology_classA-RNA |
| RN7SL172P | 0.07 | 0.68 | 0.14 | 0.9 | -0.61 | 5.20E-04 | 3.84E-03 | (B) Pathology_classA-RNA |
| GAPDHP36 | 0.01 | 0.62 | 0.03 | 0.97 | -0.61 | 8.46E-04 | 5.59E-03 | (B) Pathology_classA-RNA |
| RPL34P34 | 0.18 | 0.79 | 0.28 | 0.83 | -0.61 | 8.62E-04 | 5.65E-03 | (B) Pathology_classA-RNA |
| CD4 | 0.19 | 0.8 | 0.41 | 0.96 | -0.61 | 7.20E-03 | 0.0281 | (B) Pathology_classA-RNA |
| TRBV6-9 | 0.12 | 0.73 | 0.37 | 1.07 | -0.61 | 8.91E-03 | 0.0328 | (B) Pathology_classA-RNA |
| SLC6A21P | 0.01 | 0.63 | 0.04 | 1.3 | -0.61 | 9.81E-03 | 0.0352 | (B) Pathology_classA-RNA |
| RNU6ATAC10 | 0.1 | 0.71 | 0.3 | 1.19 | -0.61 | 0.0101 | 0.036 | (B) Pathology_classA-RNA |
| RAB21 | 3.47 | 4.07 | 0.54 | 0.66 | -0.61 | 0.012 | 0.0406 | (B) Pathology_classA-RNA |
| HNRNPMP1 | 0.17 | 0.78 | 0.51 | 0.91 | -0.61 | 0.0149 | 0.0478 | (B) Pathology_classA-RNA |
| PHKB | 3.98 | 4.61 | 0.29 | 0.6 | -0.62 | 1.50E-04 | 1.48E-03 | (B) Pathology_classA-RNA |
| RNU6-878P | 0 | 0.63 | 0.01 | 0.99 | -0.62 | 8.91E-04 | 5.80E-03 | (B) Pathology_classA-RNA |
| SFMBT2 | 0.52 | 1.14 | 0.4 | 0.62 | -0.62 | 1.58E-03 | 8.95E-03 | (B) Pathology_classA-RNA |
| NIPA2P2 | 0.04 | 0.67 | 0.13 | 1.07 | -0.62 | 2.13E-03 | 0.0113 | (B) Pathology_classA-RNA |
| LINC00508 | 0.02 | 0.64 | 0.05 | 1.14 | -0.62 | 3.30E-03 | 0.0157 | (B) Pathology_classA-RNA |
| HSPA8P4 | 0.35 | 0.97 | 0.39 | 0.84 | -0.62 | 3.62E-03 | 0.0168 | (B) Pathology_classA-RNA |
| RNFT1P2 | 0.06 | 0.68 | 0.19 | 1.1 | -0.62 | 3.73E-03 | 0.0172 | (B) Pathology_classA-RNA |
| MLANA | 1.44 | 2.06 | 0.36 | 0.93 | -0.62 | 4.08E-03 | 0.0184 | (B) Pathology_classA-RNA |
| TRAV34 | 0.09 | 0.71 | 0.28 | 1.06 | -0.62 | 4.23E-03 | 0.019 | (B) Pathology_classA-RNA |
| CDC42SE2 | 0.15 | 0.77 | 0.38 | 0.92 | -0.62 | 4.35E-03 | 0.0194 | (B) Pathology_classA-RNA |
| RPL15P18 | 2.37 | 2.99 | 0.35 | 1.02 | -0.62 | 5.80E-03 | 0.024 | (B) Pathology_classA-RNA |
| CFLAR-AS1 | 1.51 | 2.13 | 0.5 | 0.76 | -0.62 | 8.72E-03 | 0.0323 | (B) Pathology_classA-RNA |
| MIR-1250/12 | 0.23 | 0.85 | 0.43 | 1.04 | -0.62 | 0.0109 | 0.0378 | (B) Pathology_classA-RNA |
| MIR-1250/3P | 0.23 | 0.85 | 0.43 | 1.04 | -0.62 | 0.0109 | 0.0378 | (B) Pathology_classA-RNA |
| MIR-1250/5P | 0.23 | 0.85 | 0.43 | 1.04 | -0.62 | 0.0109 | 0.0378 | (B) Pathology_classA-RNA |
| WWC2-AS2 | 0 | 0.63 | 0.01 | 0.62 | -0.63 | 1.39E-06 | 3.88E-05 | (B) Pathology_classA-RNA |
| RAD50 | 0.4 | 1.03 | 0.26 | 0.65 | -0.63 | 8.22E-05 | 9.31E-04 | (B) Pathology_classA-RNA |
| RNU6-79P | 0.02 | 0.65 | 0.06 | 0.85 | -0.63 | 1.53E-04 | 1.51E-03 | (B) Pathology_classA-RNA |
| SRFBP1 | 1.13 | 1.76 | 0.34 | 0.51 | -0.63 | 2.99E-04 | 2.52E-03 | (B) Pathology_classA-RNA |
| NOC3L | 2.69 | 3.31 | 0.38 | 0.66 | -0.63 | 1.32E-03 | 7.81E-03 | (B) Pathology_classA-RNA |
| ENTPD5 | 0.25 | 0.89 | 0.16 | 1.06 | -0.63 | 1.83E-03 | 0.01 | (B) Pathology_classA-RNA |
| RNU6-504P | 0.2 | 0.83 | 0.3 | 0.98 | -0.63 | 2.76E-03 | 0.0137 | (B) Pathology_classA-RNA |
| MED13L | 6.03 | 6.66 | 0.47 | 0.63 | -0.63 | 4.26E-03 | 0.0191 | (B) Pathology_classA-RNA |
| TRANK1 | 2.55 | 3.18 | 0.45 | 0.91 | -0.63 | 7.14E-03 | 0.0279 | (B) Pathology_classA-RNA |
| TMA16 | 4.24 | 4.87 | 0.5 | 0.76 | -0.63 | 7.50E-03 | 0.029 | (B) Pathology_classA-RNA |
| THUMPD1 | 5.77 | 6.4 | 0.45 | 0.95 | -0.63 | 8.16E-03 | 0.0307 | (B) Pathology_classA-RNA |
| GPRASP1 | 2.32 | 2.94 | 0.41 | 1.08 | -0.63 | 0.0103 | 0.0364 | (B) Pathology_classA-RNA |
| RHOT1P2 | 1.26 | 1.89 | 0.56 | 0.7 | -0.63 | 0.0122 | 0.0412 | (B) Pathology_classA-RNA |
| RNU6-150P | 0.09 | 0.72 | 0.28 | 1.32 | -0.63 | 0.0146 | 0.0469 | (B) Pathology_classA-RNA |
| EMC3 | 0.06 | 0.7 | 0.13 | 0.81 | -0.64 | 9.43E-05 | 1.04E-03 | (B) Pathology_classA-RNA |
| ARHGAP23P1 | 0.06 | 0.7 | 0.12 | 0.83 | -0.64 | 9.70E-05 | 1.06E-03 | (B) Pathology_classA-RNA |
| FAM205A | 0.18 | 0.82 | 0.17 | 0.86 | -0.64 | 2.25E-04 | 2.01E-03 | (B) Pathology_classA-RNA |
| VPS13B | 3.97 | 4.61 | 0.41 | 0.62 | -0.64 | 1.58E-03 | 8.98E-03 | (B) Pathology_classA-RNA |
| PGBD4P7 | 0.08 | 0.72 | 0.25 | 0.99 | -0.64 | 1.63E-03 | 9.19E-03 | (B) Pathology_classA-RNA |
| P2RY13 | 0.05 | 0.68 | 0.14 | 1.22 | -0.64 | 5.11E-03 | 0.0219 | (B) Pathology_classA-RNA |

| | | | | | | | | |
|---|---|---|---|---|---|---|---|---|
| NFAM1 | 0.03 | 0.67 | 0.09 | 1.24 | -0.64 | 5.25E-03 | 0.0223 | (B) Pathology_classA-RNA |
| PIEZO1 | 0.22 | 0.86 | 0.47 | 1 | -0.64 | 0.0101 | 0.0359 | (B) Pathology_classA-RNA |
| PMS2P9 | 2.04 | 2.68 | 0.4 | 1.14 | -0.64 | 0.0101 | 0.036 | (B) Pathology_classA-RNA |
| SCYL2 | 0.41 | 1.05 | 0.53 | 0.96 | -0.64 | 0.0137 | 0.0449 | (B) Pathology_classA-RNA |
| MS4A7 | 0.2 | 0.85 | 0.21 | 0.57 | -0.65 | 4.74E-06 | 1.02E-04 | (B) Pathology_classA-RNA |
| CASP10 | 0.08 | 0.73 | 0.19 | 0.62 | -0.65 | 5.80E-06 | 1.18E-04 | (B) Pathology_classA-RNA |
| KDELC1P1 | 0.08 | 0.73 | 0.15 | 0.88 | -0.65 | 2.09E-04 | 1.90E-03 | (B) Pathology_classA-RNA |
| ZNF859P | 0.06 | 0.71 | 0.17 | 0.91 | -0.65 | 3.62E-04 | 2.92E-03 | (B) Pathology_classA-RNA |
| FDPSP2 | 0.07 | 0.72 | 0.21 | 0.92 | -0.65 | 5.17E-04 | 3.83E-03 | (B) Pathology_classA-RNA |
| GPD1L | 0.27 | 0.92 | 0.28 | 0.95 | -0.65 | 1.31E-03 | 7.76E-03 | (B) Pathology_classA-RNA |
| KAT2B | 0.04 | 0.69 | 0.1 | 1.12 | -0.65 | 2.12E-03 | 0.0112 | (B) Pathology_classA-RNA |
| DNMBP | 0.4 | 1.05 | 0.48 | 0.61 | -0.65 | 3.82E-03 | 0.0175 | (B) Pathology_classA-RNA |
| FARSBP1 | 0.07 | 0.72 | 0.22 | 1.23 | -0.65 | 6.08E-03 | 0.0249 | (B) Pathology_classA-RNA |
| HMGB3P18 | 0.1 | 0.75 | 0.2 | 1.28 | -0.65 | 6.84E-03 | 0.0271 | (B) Pathology_classA-RNA |
| HNRNPRP1 | 4.67 | 5.32 | 0.54 | 0.59 | -0.65 | 6.99E-03 | 0.0275 | (B) Pathology_classA-RNA |
| VTI1A | 3.62 | 4.27 | 0.59 | 0.37 | -0.65 | 0.0107 | 0.0374 | (B) Pathology_classA-RNA |
| PLEKHO2 | 5.64 | 6.29 | 0.48 | 1.12 | -0.65 | 0.0136 | 0.0447 | (B) Pathology_classA-RNA |
| RPS20P31 | 0.04 | 0.7 | 0.1 | 0.85 | -0.66 | 7.85E-05 | 8.97E-04 | (B) Pathology_classA-RNA |
| TMPPE | 0.2 | 0.86 | 0.35 | 0.59 | -0.66 | 3.37E-04 | 2.75E-03 | (B) Pathology_classA-RNA |
| RNU6-260P | 0.02 | 0.68 | 0.06 | 1.04 | -0.66 | 8.44E-04 | 5.58E-03 | (B) Pathology_classA-RNA |
| ADGRE1 | 0.03 | 0.69 | 0.1 | 1.12 | -0.66 | 1.82E-03 | 9.97E-03 | (B) Pathology_classA-RNA |
| OR4K2 | 0.04 | 0.71 | 0.13 | 1.19 | -0.66 | 3.10E-03 | 0.015 | (B) Pathology_classA-RNA |
| LRRIQ1 | 0.16 | 0.82 | 0.33 | 1.08 | -0.66 | 4.34E-03 | 0.0194 | (B) Pathology_classA-RNA |
| DPPA4 | 0.02 | 0.68 | 0.06 | 1.27 | -0.66 | 4.83E-03 | 0.021 | (B) Pathology_classA-RNA |
| GPR1 | 0.22 | 0.88 | 0.32 | 1.27 | -0.66 | 9.52E-03 | 0.0344 | (B) Pathology_classA-RNA |
| MRPL19 | 5.75 | 6.41 | 0.58 | 0.65 | -0.66 | 0.0109 | 0.0379 | (B) Pathology_classA-RNA |
| LRP1B | 0.36 | 1.02 | 0.4 | 1.24 | -0.66 | 0.0119 | 0.0404 | (B) Pathology_classA-RNA |
| ETF1P2 | 0.06 | 0.73 | 0.14 | 0.69 | -0.67 | 5.02E-06 | 1.06E-04 | (B) Pathology_classA-RNA |
| VSIG10L | 0.11 | 0.77 | 0.22 | 0.91 | -0.67 | 3.83E-04 | 3.05E-03 | (B) Pathology_classA-RNA |
| RNU6-141P | 0.08 | 0.75 | 0.24 | 0.93 | -0.67 | 5.69E-04 | 4.14E-03 | (B) Pathology_classA-RNA |
| ANKRD26P3 | 0.13 | 0.8 | 0.23 | 1.04 | -0.67 | 1.32E-03 | 7.80E-03 | (B) Pathology_classA-RNA |
| RNU6-176P | 0.05 | 0.72 | 0.15 | 1.1 | -0.67 | 1.50E-03 | 8.60E-03 | (B) Pathology_classA-RNA |
| PDE12 | 3.05 | 3.72 | 0.43 | 0.58 | -0.67 | 1.55E-03 | 8.84E-03 | (B) Pathology_classA-RNA |
| RNU6-319P | 0.07 | 0.74 | 0.22 | 1.07 | -0.67 | 1.60E-03 | 9.06E-03 | (B) Pathology_classA-RNA |
| PRRC1 | 0.52 | 1.19 | 0.43 | 0.77 | -0.67 | 2.25E-03 | 0.0117 | (B) Pathology_classA-RNA |
| PHBP12 | 0.13 | 0.8 | 0.39 | 0.98 | -0.67 | 3.51E-03 | 0.0164 | (B) Pathology_classA-RNA |
| GAPDHP49 | 0.06 | 0.73 | 0.19 | 1.25 | -0.67 | 4.61E-03 | 0.0202 | (B) Pathology_classA-RNA |
| PRORY | 0.02 | 0.68 | 0.05 | 1.33 | -0.67 | 6.32E-03 | 0.0256 | (B) Pathology_classA-RNA |
| RB1-DT | 1.77 | 2.45 | 0.51 | 0.98 | -0.67 | 9.44E-03 | 0.0342 | (B) Pathology_classA-RNA |
| USP32 | 0.29 | 0.96 | 0.57 | 0.89 | -0.67 | 0.0126 | 0.0422 | (B) Pathology_classA-RNA |
| HIST1H2AE | 0.32 | 0.99 | 0.52 | 1.15 | -0.67 | 0.0154 | 0.049 | (B) Pathology_classA-RNA |
| MYCBP2 | 0.11 | 0.79 | 0.14 | 0.71 | -0.68 | 5.01E-06 | 1.06E-04 | (B) Pathology_classA-RNA |
| GLIPR1 | 0.18 | 0.85 | 0.28 | 0.84 | -0.68 | 3.26E-04 | 2.68E-03 | (B) Pathology_classA-RNA |
| RN7SL563P | 0.01 | 0.69 | 0.03 | 1 | -0.68 | 3.91E-04 | 3.10E-03 | (B) Pathology_classA-RNA |
| SCARA3 | 0 | 0.68 | 0.01 | 1 | -0.68 | 3.99E-04 | 3.15E-03 | (B) Pathology_classA-RNA |
| ZNF185 | 0.09 | 0.77 | 0.27 | 0.95 | -0.68 | 7.58E-04 | 5.14E-03 | (B) Pathology_classA-RNA |
| MIR-4419A/4 | 0.11 | 0.79 | 0.26 | 1 | -0.68 | 1.01E-03 | 6.36E-03 | (B) Pathology_classA-RNA |
| TRBV6-8 | 0.06 | 0.74 | 0.15 | 1.09 | -0.68 | 1.12E-03 | 6.88E-03 | (B) Pathology_classA-RNA |
| NFIL3 | 1.93 | 2.62 | 0.35 | 0.93 | -0.68 | 1.42E-03 | 8.25E-03 | (B) Pathology_classA-RNA |
| MTCYBP4 | 0 | 0.69 | 0.01 | 1.18 | -0.68 | 1.97E-03 | 0.0106 | (B) Pathology_classA-RNA |
| MTND2P18 | 0.16 | 0.83 | 0.24 | 1.11 | -0.68 | 2.22E-03 | 0.0116 | (B) Pathology_classA-RNA |
| NPM1P11 | 0.25 | 0.93 | 0.29 | 1.13 | -0.68 | 2.94E-03 | 0.0144 | (B) Pathology_classA-RNA |
| CSNK1A1L | 2.01 | 2.69 | 0.44 | 0.97 | -0.68 | 4.45E-03 | 0.0197 | (B) Pathology_classA-RNA |
| DTX2P1 | 0.64 | 1.32 | 0.24 | 1.22 | -0.68 | 4.47E-03 | 0.0198 | (B) Pathology_classA-RNA |
| RNU6-1217P | 0.17 | 0.84 | 0.33 | 1.19 | -0.68 | 5.58E-03 | 0.0233 | (B) Pathology_classA-RNA |
| RNU6-199P | 0.27 | 0.95 | 0.43 | 1.06 | -0.68 | 6.05E-03 | 0.0248 | (B) Pathology_classA-RNA |
| PAICSP6 | 0.27 | 0.95 | 0.35 | 1.21 | -0.68 | 7.26E-03 | 0.0283 | (B) Pathology_classA-RNA |
| IGHV3-30 | 0.08 | 0.76 | 0.24 | 1.45 | -0.68 | 0.0132 | 0.0436 | (B) Pathology_classA-RNA |
| RNU6-498P | 0.27 | 0.95 | 0.39 | 1.36 | -0.68 | 0.0149 | 0.0476 | (B) Pathology_classA-RNA |
| FGF13-AS1 | 0.01 | 0.71 | 0.04 | 0.71 | -0.69 | 2.52E-06 | 6.14E-05 | (B) Pathology_classA-RNA |
| NADK2 | 0.02 | 0.71 | 0.07 | 0.76 | -0.69 | 8.43E-06 | 1.59E-04 | (B) Pathology_classA-RNA |
| RNU6-879P | 0.01 | 0.7 | 0.02 | 0.84 | -0.69 | 3.39E-05 | 4.65E-04 | (B) Pathology_classA-RNA |
| ELOCP33 | 0.06 | 0.75 | 0.18 | 0.88 | -0.69 | 1.34E-04 | 1.36E-03 | (B) Pathology_classA-RNA |
| FGD2 | 0.11 | 0.8 | 0.24 | 0.94 | -0.69 | 4.15E-04 | 3.24E-03 | (B) Pathology_classA-RNA |
| RN7SL434P | 0.07 | 0.76 | 0.2 | 1.01 | -0.69 | 5.93E-04 | 4.27E-03 | (B) Pathology_classA-RNA |

| | | | | | | | | |
|---|---|---|---|---|---|---|---|---|
| LINC00582 | 0.06 | 0.75 | 0.13 | 1.05 | -0.69 | 6.68E-04 | 4.67E-03 | (B) Pathology_classA-RNA |
| FZD10-AS1 | 0.1 | 0.8 | 0.26 | 1.04 | -0.69 | 1.20E-03 | 7.25E-03 | (B) Pathology_classA-RNA |
| MIR-554/554 | 0.07 | 0.76 | 0.22 | 1.11 | -0.69 | 1.64E-03 | 9.21E-03 | (B) Pathology_classA-RNA |
| TTLL11-IT1 | 0.29 | 0.97 | 0.45 | 0.87 | -0.69 | 3.14E-03 | 0.0151 | (B) Pathology_classA-RNA |
| DDX11L10 | 0.09 | 0.78 | 0.27 | 1.21 | -0.69 | 3.90E-03 | 0.0178 | (B) Pathology_classA-RNA |
| LILRB4 | 0.22 | 0.91 | 0.51 | 0.86 | -0.69 | 5.79E-03 | 0.024 | (B) Pathology_classA-RNA |
| ENO1P3 | 0.19 | 0.89 | 0.46 | 1.03 | -0.69 | 6.14E-03 | 0.025 | (B) Pathology_classA-RNA |
| MS4A4A | 0.21 | 0.9 | 0.5 | 0.97 | -0.69 | 7.14E-03 | 0.0279 | (B) Pathology_classA-RNA |
| IGKV2-40 | 0.11 | 0.8 | 0.33 | 1.31 | -0.69 | 8.69E-03 | 0.0322 | (B) Pathology_classA-RNA |
| RNU6-626P | 0.07 | 0.76 | 0.2 | 1.41 | -0.69 | 9.02E-03 | 0.0331 | (B) Pathology_classA-RNA |
| HIST1H4A | 0.08 | 0.77 | 0.25 | 1.43 | -0.69 | 0.0116 | 0.0398 | (B) Pathology_classA-RNA |
| ENTPD7 | 3.61 | 4.31 | 0.26 | 0.68 | -0.7 | 2.56E-05 | 3.72E-04 | (B) Pathology_classA-RNA |
| KCNRG | 4.9 | 5.6 | 0.43 | 0.72 | -0.7 | 1.22E-03 | 7.35E-03 | (B) Pathology_classA-RNA |
| BCRP6 | 0.06 | 0.76 | 0.17 | 1.12 | -0.7 | 1.25E-03 | 7.51E-03 | (B) Pathology_classA-RNA |
| CENPVL3 | 0.05 | 0.74 | 0.14 | 1.13 | -0.7 | 1.28E-03 | 7.61E-03 | (B) Pathology_classA-RNA |
| RN7SL242P | 0.09 | 0.79 | 0.22 | 1.12 | -0.7 | 1.41E-03 | 8.21E-03 | (B) Pathology_classA-RNA |
| NPIPB8 | 4.17 | 4.86 | 0.47 | 0.57 | -0.7 | 1.82E-03 | 9.96E-03 | (B) Pathology_classA-RNA |
| TRNT1 | 3.29 | 3.99 | 0.45 | 1.11 | -0.7 | 6.46E-03 | 0.026 | (B) Pathology_classA-RNA |
| USP32P2 | 0.42 | 1.11 | 0.49 | 1.03 | -0.7 | 7.32E-03 | 0.0285 | (B) Pathology_classA-RNA |
| RNU6-517P | 0.06 | 0.75 | 0.14 | 1.43 | -0.7 | 8.63E-03 | 0.032 | (B) Pathology_classA-RNA |
| RNU6-1111P | 0.22 | 0.92 | 0.47 | 1.19 | -0.7 | 0.0107 | 0.0374 | (B) Pathology_classA-RNA |
| RNU2-13P | 0.26 | 0.96 | 0.38 | 1.39 | -0.7 | 0.0132 | 0.0436 | (B) Pathology_classA-RNA |
| UQCRBP2 | 0.12 | 0.81 | 0.35 | 1.46 | -0.7 | 0.0156 | 0.0495 | (B) Pathology_classA-RNA |
| PPP1R3C | 0 | 0.71 | 0.01 | 0.95 | -0.71 | 1.29E-04 | 1.31E-03 | (B) Pathology_classA-RNA |
| MTND4P4 | 0.03 | 0.73 | 0.05 | 0.98 | -0.71 | 1.94E-04 | 1.80E-03 | (B) Pathology_classA-RNA |
| DCAF13P2 | 0 | 0.72 | 0 | 1.04 | -0.71 | 3.11E-04 | 2.60E-03 | (B) Pathology_classA-RNA |
| CDK12 | 4.52 | 5.23 | 0.42 | 0.51 | -0.71 | 6.68E-04 | 4.67E-03 | (B) Pathology_classA-RNA |
| RN7SKP86 | 0.03 | 0.75 | 0.1 | 1.1 | -0.71 | 6.89E-04 | 4.79E-03 | (B) Pathology_classA-RNA |
| TMED10P2 | 0.13 | 0.83 | 0.28 | 0.99 | -0.71 | 8.05E-04 | 5.40E-03 | (B) Pathology_classA-RNA |
| RN7SL37P | 0.1 | 0.81 | 0.24 | 1.07 | -0.71 | 9.80E-04 | 6.22E-03 | (B) Pathology_classA-RNA |
| MIR-4518/45 | 0.02 | 0.73 | 0.06 | 1.18 | -0.71 | 1.32E-03 | 7.79E-03 | (B) Pathology_classA-RNA |
| GRHL1 | 0.14 | 0.85 | 0.41 | 0.93 | -0.71 | 1.94E-03 | 0.0105 | (B) Pathology_classA-RNA |
| GCNT1P5 | 0.16 | 0.87 | 0.31 | 1.11 | -0.71 | 2.16E-03 | 0.0113 | (B) Pathology_classA-RNA |
| HIST1H2AJ | 0.02 | 0.73 | 0.05 | 1.25 | -0.71 | 2.36E-03 | 0.0122 | (B) Pathology_classA-RNA |
| DPRXP2 | 0.16 | 0.87 | 0.21 | 1.22 | -0.71 | 2.67E-03 | 0.0134 | (B) Pathology_classA-RNA |
| PSMC1P2 | 0.26 | 0.97 | 0.33 | 1.19 | -0.71 | 3.56E-03 | 0.0166 | (B) Pathology_classA-RNA |
| IL6STP1 | 1.54 | 2.25 | 0.3 | 1.28 | -0.71 | 5.12E-03 | 0.0219 | (B) Pathology_classA-RNA |
| NPM1P26 | 6.42 | 7.13 | 0.64 | 0.64 | -0.71 | 0.0112 | 0.0386 | (B) Pathology_classA-RNA |
| LPP-AS2 | 2.04 | 2.75 | 0.68 | 0.56 | -0.71 | 0.0152 | 0.0485 | (B) Pathology_classA-RNA |
| NCOA2 | 1.81 | 2.52 | 0.65 | 0.88 | -0.71 | 0.0153 | 0.0487 | (B) Pathology_classA-RNA |
| MIR-4720/3P | 0.19 | 0.92 | 0.18 | 0.98 | -0.72 | 2.25E-04 | 2.01E-03 | (B) Pathology_classA-RNA |
| MIR-4720/5P | 0.19 | 0.92 | 0.18 | 0.98 | -0.72 | 2.25E-04 | 2.01E-03 | (B) Pathology_classA-RNA |
| RNU6-1062P | 0.08 | 0.81 | 0.25 | 0.98 | -0.72 | 4.25E-04 | 3.30E-03 | (B) Pathology_classA-RNA |
| SAMD8 | 7.07 | 7.79 | 0.41 | 0.48 | -0.72 | 4.47E-04 | 3.43E-03 | (B) Pathology_classA-RNA |
| UTP20 | 2.01 | 2.73 | 0.43 | 0.6 | -0.72 | 7.06E-04 | 4.88E-03 | (B) Pathology_classA-RNA |
| NBPF8 | 0.88 | 1.6 | 0.4 | 0.77 | -0.72 | 7.43E-04 | 5.06E-03 | (B) Pathology_classA-RNA |
| ATP6V1A | 0.42 | 1.15 | 0.4 | 0.82 | -0.72 | 8.56E-04 | 5.63E-03 | (B) Pathology_classA-RNA |
| SLC25A15P4 | 0.06 | 0.78 | 0.19 | 1.38 | -0.72 | 5.59E-03 | 0.0233 | (B) Pathology_classA-RNA |
| FUT10 | 3.02 | 3.75 | 0.55 | 0.82 | -0.72 | 5.69E-03 | 0.0237 | (B) Pathology_classA-RNA |
| NXNL2 | 0.88 | 1.59 | 0.47 | 1.06 | -0.72 | 5.71E-03 | 0.0237 | (B) Pathology_classA-RNA |
| NLGN1-AS1 | 0 | 0.72 | 0 | 1.42 | -0.72 | 5.79E-03 | 0.024 | (B) Pathology_classA-RNA |
| DDI2 | 0.47 | 1.2 | 0.17 | 0.58 | -0.73 | 1.27E-07 | 5.89E-06 | (B) Pathology_classA-RNA |
| OR7E36P | 0.31 | 1.04 | 0.3 | 0.85 | -0.73 | 1.96E-04 | 1.81E-03 | (B) Pathology_classA-RNA |
| RNU6-597P | 0.06 | 0.8 | 0.14 | 1.01 | -0.73 | 2.09E-04 | 1.90E-03 | (B) Pathology_classA-RNA |
| MTND4P8 | 0.1 | 0.83 | 0.3 | 1.11 | -0.73 | 1.45E-03 | 8.38E-03 | (B) Pathology_classA-RNA |
| GNG5P3 | 0.12 | 0.85 | 0.35 | 1.13 | -0.73 | 2.58E-03 | 0.013 | (B) Pathology_classA-RNA |
| GM2A | 0.42 | 1.16 | 0.49 | 0.96 | -0.73 | 3.95E-03 | 0.018 | (B) Pathology_classA-RNA |
| ERVFRD-1 | 0.25 | 0.98 | 0.6 | 1.11 | -0.73 | 0.0144 | 0.0464 | (B) Pathology_classA-RNA |
| OR52H2P | 0.03 | 0.76 | 0.07 | 0.94 | -0.74 | 6.83E-05 | 8.07E-04 | (B) Pathology_classA-RNA |
| ATP5MC1P3 | 0.22 | 0.97 | 0.27 | 0.94 | -0.74 | 2.25E-04 | 2.01E-03 | (B) Pathology_classA-RNA |
| PGPEP1L | 0.13 | 0.87 | 0.22 | 1.01 | -0.74 | 3.16E-04 | 2.63E-03 | (B) Pathology_classA-RNA |
| TMPRSS11CP | 0.5 | 1.24 | 0.35 | 0.9 | -0.74 | 5.33E-04 | 3.92E-03 | (B) Pathology_classA-RNA |
| MTND5P25 | 0.18 | 0.92 | 0.28 | 1.07 | -0.74 | 8.51E-04 | 5.61E-03 | (B) Pathology_classA-RNA |
| RNU4ATAC1[illegible] | 0.12 | 0.86 | 0.37 | 1.03 | -0.74 | 1.36E-03 | 8.00E-03 | (B) Pathology_classA-RNA |
| ERCC4 | 0.85 | 1.59 | 0.43 | 0.91 | -0.74 | 1.41E-03 | 8.20E-03 | (B) Pathology_classA-RNA |

| | | | | | | | | |
|---|---|---|---|---|---|---|---|---|
| PGAM1P6 | 0.12 | 0.86 | 0.34 | 1.09 | -0.74 | 1.68E-03 | 9.40E-03 | (B) Pathology_classA-RNA |
| TRAJ1 | 0.11 | 0.84 | 0.25 | 1.19 | -0.74 | 1.82E-03 | 1.00E-02 | (B) Pathology_classA-RNA |
| TMEM254-AS | 0.17 | 0.9 | 0.33 | 1.17 | -0.74 | 2.61E-03 | 0.0131 | (B) Pathology_classA-RNA |
| RN7SKP58 | 0.15 | 0.89 | 0.3 | 1.27 | -0.74 | 3.59E-03 | 0.0167 | (B) Pathology_classA-RNA |
| NRBF2 | 0.42 | 1.16 | 0.48 | 1.1 | -0.74 | 5.27E-03 | 0.0223 | (B) Pathology_classA-RNA |
| OR7E2P | 0.14 | 0.88 | 0.41 | 1.22 | -0.74 | 5.44E-03 | 0.0228 | (B) Pathology_classA-RNA |
| ZNRF3 | 0.2 | 0.93 | 0.42 | 1.27 | -0.74 | 7.11E-03 | 0.0279 | (B) Pathology_classA-RNA |
| C1ORF195 | 0.18 | 0.93 | 0.55 | 1.09 | -0.74 | 8.31E-03 | 0.0311 | (B) Pathology_classA-RNA |
| STEAP2 | 0.19 | 0.93 | 0.56 | 1.07 | -0.74 | 8.82E-03 | 0.0326 | (B) Pathology_classA-RNA |
| RNU6-805P | 0.36 | 1.1 | 0.53 | 1.16 | -0.74 | 9.54E-03 | 0.0345 | (B) Pathology_classA-RNA |
| RNU6-733P | 0.22 | 0.96 | 0.47 | 1.32 | -0.74 | 0.0107 | 0.0374 | (B) Pathology_classA-RNA |
| MIR-2110/21 | 3.46 | 4.19 | 0.54 | 1.19 | -0.74 | 0.0109 | 0.0378 | (B) Pathology_classA-RNA |
| WDFY3-AS1 | 2.02 | 2.76 | 0.38 | 1.49 | -0.74 | 0.0128 | 0.0428 | (B) Pathology_classA-RNA |
| BFSP2-AS1 | 0.02 | 0.77 | 0.06 | 0.78 | -0.75 | 3.40E-06 | 7.79E-05 | (B) Pathology_classA-RNA |
| RPL24P7 | 0.01 | 0.76 | 0.03 | 0.98 | -0.75 | 9.61E-05 | 1.06E-03 | (B) Pathology_classA-RNA |
| RNU6-942P | 0.04 | 0.79 | 0.11 | 1 | -0.75 | 1.20E-04 | 1.25E-03 | (B) Pathology_classA-RNA |
| LINC00457 | 0.12 | 0.87 | 0.26 | 0.98 | -0.75 | 2.74E-04 | 2.35E-03 | (B) Pathology_classA-RNA |
| PTPN7 | 0.03 | 0.78 | 0.06 | 1.09 | -0.75 | 3.08E-04 | 2.57E-03 | (B) Pathology_classA-RNA |
| NPM1P43 | 0.15 | 0.9 | 0.25 | 1.01 | -0.75 | 3.55E-04 | 2.87E-03 | (B) Pathology_classA-RNA |
| MTND5P17 | 0.04 | 0.79 | 0.11 | 1.24 | -0.75 | 1.28E-03 | 7.62E-03 | (B) Pathology_classA-RNA |
| TPM4 | 0.28 | 1.04 | 0.39 | 1.05 | -0.75 | 1.65E-03 | 9.26E-03 | (B) Pathology_classA-RNA |
| PLXNC1 | 0.08 | 0.83 | 0.25 | 1.2 | -0.75 | 1.73E-03 | 9.59E-03 | (B) Pathology_classA-RNA |
| TAB3 | 0.55 | 1.3 | 0.35 | 1.12 | -0.75 | 1.80E-03 | 9.89E-03 | (B) Pathology_classA-RNA |
| MIR-744/3P | 0.19 | 0.93 | 0.47 | 1.08 | -0.75 | 4.33E-03 | 0.0193 | (B) Pathology_classA-RNA |
| MIR-744/5P | 0.19 | 0.93 | 0.47 | 1.08 | -0.75 | 4.33E-03 | 0.0193 | (B) Pathology_classA-RNA |
| MIR-744/744 | 0.19 | 0.93 | 0.47 | 1.08 | -0.75 | 4.33E-03 | 0.0193 | (B) Pathology_classA-RNA |
| MIR-744/744 | 0.19 | 0.93 | 0.47 | 1.08 | -0.75 | 4.33E-03 | 0.0193 | (B) Pathology_classA-RNA |
| ICAM3 | 0.18 | 0.94 | 0.36 | 1.36 | -0.75 | 6.27E-03 | 0.0254 | (B) Pathology_classA-RNA |
| RILPL2 | 1.02 | 1.77 | 0.6 | 1.05 | -0.75 | 0.0107 | 0.0374 | (B) Pathology_classA-RNA |
| CLTC | 1.02 | 1.77 | 0.69 | 0.76 | -0.75 | 0.0139 | 0.0452 | (B) Pathology_classA-RNA |
| STAB2 | 0.4 | 1.16 | 0.26 | 0.76 | -0.76 | 2.46E-05 | 3.60E-04 | (B) Pathology_classA-RNA |
| RN7SL706P | 0.02 | 0.78 | 0.07 | 0.92 | -0.76 | 3.55E-05 | 4.83E-04 | (B) Pathology_classA-RNA |
| CCDC125 | 4.54 | 5.29 | 0.2 | 0.99 | -0.76 | 1.73E-04 | 1.64E-03 | (B) Pathology_classA-RNA |
| MIR-3611/36 | 0.03 | 0.79 | 0.09 | 1.08 | -0.76 | 2.81E-04 | 2.39E-03 | (B) Pathology_classA-RNA |
| RNU6-162P | 0.02 | 0.78 | 0.06 | 1.14 | -0.76 | 4.45E-04 | 3.42E-03 | (B) Pathology_classA-RNA |
| RN7SKP163 | 0.01 | 0.77 | 0.02 | 1.13 | -0.76 | 4.48E-04 | 3.44E-03 | (B) Pathology_classA-RNA |
| CRISPLD2 | 0.25 | 1.01 | 0.49 | 0.99 | -0.76 | 3.48E-03 | 0.0163 | (B) Pathology_classA-RNA |
| PGGT1B | 0.94 | 1.7 | 0.58 | 0.72 | -0.76 | 4.51E-03 | 0.0199 | (B) Pathology_classA-RNA |
| ADAP2 | 1.26 | 2.02 | 0.62 | 0.67 | -0.76 | 6.21E-03 | 0.0253 | (B) Pathology_classA-RNA |
| RDH11 | 1.22 | 1.98 | 0.26 | 1.47 | -0.76 | 6.60E-03 | 0.0264 | (B) Pathology_classA-RNA |
| SQOR | 0.27 | 1.03 | 0.59 | 0.95 | -0.76 | 7.58E-03 | 0.0291 | (B) Pathology_classA-RNA |
| PTCD2 | 2.42 | 3.18 | 0.4 | 0.69 | -0.77 | 2.73E-04 | 2.34E-03 | (B) Pathology_classA-RNA |
| NGLY1 | 5.84 | 6.61 | 0.43 | 0.82 | -0.77 | 7.51E-04 | 5.11E-03 | (B) Pathology_classA-RNA |
| FHL1 | 0.26 | 1.02 | 0.22 | 1.17 | -0.77 | 9.06E-04 | 5.87E-03 | (B) Pathology_classA-RNA |
| PPP1R12BP1 | 0.02 | 0.79 | 0.05 | 1.3 | -0.77 | 1.56E-03 | 8.90E-03 | (B) Pathology_classA-RNA |
| UBE2V2P2 | 0.45 | 1.21 | 0.35 | 1.29 | -0.77 | 3.88E-03 | 0.0177 | (B) Pathology_classA-RNA |
| AHR | 0.04 | 0.81 | 0.12 | 1.45 | -0.77 | 4.28E-03 | 0.0192 | (B) Pathology_classA-RNA |
| RAC1P6 | 0.07 | 0.84 | 0.22 | 1.45 | -0.77 | 5.30E-03 | 0.0224 | (B) Pathology_classA-RNA |
| MIR-566/566 | 0.08 | 0.85 | 0.24 | 1.45 | -0.77 | 5.61E-03 | 0.0234 | (B) Pathology_classA-RNA |
| TASOR2 | 1.27 | 2.05 | 0.66 | 0.63 | -0.77 | 8.52E-03 | 0.0317 | (B) Pathology_classA-RNA |
| MIR-126/126 | 0.3 | 1.06 | 0.53 | 1.27 | -0.77 | 9.73E-03 | 0.035 | (B) Pathology_classA-RNA |
| MIR-126/126 | 0.3 | 1.06 | 0.53 | 1.27 | -0.77 | 9.73E-03 | 0.035 | (B) Pathology_classA-RNA |
| MIR-126/3P | 0.3 | 1.06 | 0.53 | 1.27 | -0.77 | 9.73E-03 | 0.035 | (B) Pathology_classA-RNA |
| MIR-126/5P | 0.3 | 1.06 | 0.53 | 1.27 | -0.77 | 9.73E-03 | 0.035 | (B) Pathology_classA-RNA |
| NUP93 | 2.23 | 2.99 | 0.6 | 1.19 | -0.77 | 0.0121 | 0.041 | (B) Pathology_classA-RNA |
| RPL35P3 | 0.44 | 1.21 | 0.56 | 1.37 | -0.77 | 0.0148 | 0.0475 | (B) Pathology_classA-RNA |
| MTND4P11 | 0.08 | 0.86 | 0.24 | 1.04 | -0.78 | 2.54E-04 | 2.20E-03 | (B) Pathology_classA-RNA |
| TRBV6-5 | 0.02 | 0.79 | 0.06 | 1.21 | -0.78 | 7.07E-04 | 4.89E-03 | (B) Pathology_classA-RNA |
| NSUN3 | 1.35 | 2.13 | 0.5 | 0.91 | -0.78 | 2.09E-03 | 0.0111 | (B) Pathology_classA-RNA |
| ADAM9 | 7.94 | 8.73 | 0.36 | 1.2 | -0.78 | 2.13E-03 | 0.0112 | (B) Pathology_classA-RNA |
| RNU5B-1 | 0.15 | 0.93 | 0.32 | 1.28 | -0.78 | 2.77E-03 | 0.0138 | (B) Pathology_classA-RNA |
| WWP1 | 4.34 | 5.12 | 0.6 | 0.86 | -0.78 | 5.65E-03 | 0.0235 | (B) Pathology_classA-RNA |
| TP53INP1 | 1.31 | 2.09 | 0.56 | 1.19 | -0.78 | 8.91E-03 | 0.0328 | (B) Pathology_classA-RNA |
| MIR-608/608 | 0.33 | 1.11 | 0.52 | 1.44 | -0.78 | 0.0139 | 0.0454 | (B) Pathology_classA-RNA |
| SMARCA2 | 0.09 | 0.88 | 0.24 | 0.82 | -0.79 | 1.64E-05 | 2.66E-04 | (B) Pathology_classA-RNA |

| | | | | | | | | |
|---|---|---|---|---|---|---|---|---|
| SUCLG2P2 | 0.17 | 0.97 | 0.33 | 0.76 | -0.79 | 6.21E-05 | 7.48E-04 | (B) Pathology_classA-RNA |
| ALDH7A1P1 | 0.22 | 1.01 | 0.33 | 0.77 | -0.79 | 6.75E-05 | 8.00E-04 | (B) Pathology_classA-RNA |
| RNU6-1005P | 0.07 | 0.86 | 0.22 | 1.01 | -0.79 | 1.37E-04 | 1.38E-03 | (B) Pathology_classA-RNA |
| CACYBPP1 | 0.03 | 0.82 | 0.08 | 1.11 | -0.79 | 2.25E-04 | 2.01E-03 | (B) Pathology_classA-RNA |
| ACTG1P1 | 2.38 | 3.18 | 0.36 | 0.94 | -0.79 | 3.57E-04 | 2.88E-03 | (B) Pathology_classA-RNA |
| SNORA84 | 0.05 | 0.84 | 0.16 | 1.19 | -0.79 | 6.54E-04 | 4.60E-03 | (B) Pathology_classA-RNA |
| HPGDS | 0.09 | 0.88 | 0.26 | 1.18 | -0.79 | 8.82E-04 | 5.75E-03 | (B) Pathology_classA-RNA |
| LCP2 | 0.12 | 0.91 | 0.37 | 1.09 | -0.79 | 1.14E-03 | 7.00E-03 | (B) Pathology_classA-RNA |
| GCNT1P3 | 0.22 | 1 | 0.37 | 1.11 | -0.79 | 1.30E-03 | 7.72E-03 | (B) Pathology_classA-RNA |
| SLC25A39P1 | 0.13 | 0.92 | 0.39 | 1.14 | -0.79 | 1.84E-03 | 0.0101 | (B) Pathology_classA-RNA |
| KDM5A | 7.92 | 8.71 | 0.61 | 0.39 | -0.79 | 4.23E-03 | 0.019 | (B) Pathology_classA-RNA |
| CEACAM3 | 0.03 | 0.82 | 0.1 | 1.68 | -0.79 | 0.0103 | 0.0364 | (B) Pathology_classA-RNA |
| RNA5SP33 | 0.17 | 0.96 | 0.51 | 1.37 | -0.79 | 0.0104 | 0.0368 | (B) Pathology_classA-RNA |
| PTP4A1P2 | 0.45 | 1.25 | 0.46 | 1.51 | -0.79 | 0.012 | 0.0408 | (B) Pathology_classA-RNA |
| WDR3 | 5.05 | 5.84 | 0.69 | 0.93 | -0.79 | 0.0121 | 0.0409 | (B) Pathology_classA-RNA |
| RTP5 | 0.04 | 0.85 | 0.13 | 0.96 | -0.8 | 3.31E-05 | 4.58E-04 | (B) Pathology_classA-RNA |
| MTND1P9 | 0.07 | 0.88 | 0.12 | 1.01 | -0.8 | 6.19E-05 | 7.46E-04 | (B) Pathology_classA-RNA |
| RN7SL735P | 0.04 | 0.84 | 0.09 | 1.09 | -0.8 | 1.55E-04 | 1.52E-03 | (B) Pathology_classA-RNA |
| PRR16 | 0.13 | 0.94 | 0.35 | 1.05 | -0.8 | 5.62E-04 | 4.10E-03 | (B) Pathology_classA-RNA |
| BRD7P4 | 0.33 | 1.13 | 0.48 | 0.7 | -0.8 | 8.75E-04 | 5.71E-03 | (B) Pathology_classA-RNA |
| PACSIN3 | 0.14 | 0.94 | 0.41 | 1.02 | -0.8 | 9.52E-04 | 6.08E-03 | (B) Pathology_classA-RNA |
| SETP11 | 0.09 | 0.89 | 0.27 | 1.22 | -0.8 | 1.11E-03 | 6.84E-03 | (B) Pathology_classA-RNA |
| CCDC144CP | 0.42 | 1.23 | 0.44 | 1.04 | -0.8 | 1.39E-03 | 8.12E-03 | (B) Pathology_classA-RNA |
| CLEC4GP1 | 0.16 | 0.97 | 0.27 | 1.28 | -0.8 | 1.59E-03 | 8.99E-03 | (B) Pathology_classA-RNA |
| CDY9P | 0.08 | 0.88 | 0.23 | 1.45 | -0.8 | 3.92E-03 | 0.0179 | (B) Pathology_classA-RNA |
| NFIA | 0.4 | 1.2 | 0.61 | 0.75 | -0.8 | 4.56E-03 | 0.0201 | (B) Pathology_classA-RNA |
| INPP5A | 3.74 | 4.54 | 0.62 | 0.91 | -0.8 | 6.11E-03 | 0.0249 | (B) Pathology_classA-RNA |
| PABPC1P11 | 0.07 | 0.88 | 0.21 | 1.15 | -0.81 | 3.73E-04 | 2.99E-03 | (B) Pathology_classA-RNA |
| ZFAT-AS1 | 1.62 | 2.43 | 0.46 | 0.6 | -0.81 | 4.64E-04 | 3.53E-03 | (B) Pathology_classA-RNA |
| LMAN1L | 0.02 | 0.83 | 0.07 | 1.25 | -0.81 | 6.40E-04 | 4.52E-03 | (B) Pathology_classA-RNA |
| PTK2B | 2.01 | 2.81 | 0.43 | 0.98 | -0.81 | 9.24E-04 | 5.96E-03 | (B) Pathology_classA-RNA |
| DPRXP4 | 2.53 | 3.35 | 0.49 | 0.9 | -0.81 | 1.36E-03 | 7.98E-03 | (B) Pathology_classA-RNA |
| ZHX1 | 0.73 | 1.53 | 0.5 | 1.01 | -0.81 | 2.20E-03 | 0.0115 | (B) Pathology_classA-RNA |
| RNU1-100P | 0.2 | 1.01 | 0.45 | 1.13 | -0.81 | 2.29E-03 | 0.0119 | (B) Pathology_classA-RNA |
| MIR-3121/31 | 0.12 | 0.93 | 0.35 | 1.44 | -0.81 | 4.85E-03 | 0.021 | (B) Pathology_classA-RNA |
| MIR-3121/3P | 0.12 | 0.93 | 0.35 | 1.44 | -0.81 | 4.85E-03 | 0.021 | (B) Pathology_classA-RNA |
| MIR-3121/5P | 0.12 | 0.93 | 0.35 | 1.44 | -0.81 | 4.85E-03 | 0.021 | (B) Pathology_classA-RNA |
| HIF1A-AS1 | 4.63 | 5.44 | 0.58 | 1.34 | -0.81 | 0.0117 | 0.0399 | (B) Pathology_classA-RNA |
| SNORA7A | 5.99 | 6.8 | 0.73 | 1.01 | -0.81 | 0.015 | 0.048 | (B) Pathology_classA-RNA |
| ACLY | 5.33 | 6.14 | 0.75 | 0.95 | -0.81 | 0.0158 | 0.05 | (B) Pathology_classA-RNA |
| TMLHE | 0.11 | 0.93 | 0.21 | 0.71 | -0.82 | 6.34E-07 | 2.07E-05 | (B) Pathology_classA-RNA |
| SULT1C2P1 | 0.1 | 0.92 | 0.22 | 0.92 | -0.82 | 2.73E-05 | 3.91E-04 | (B) Pathology_classA-RNA |
| NXPE4 | 0.07 | 0.89 | 0.14 | 1.15 | -0.82 | 2.35E-04 | 2.08E-03 | (B) Pathology_classA-RNA |
| THNSL1 | 0.26 | 1.07 | 0.46 | 0.79 | -0.82 | 6.07E-04 | 4.35E-03 | (B) Pathology_classA-RNA |
| MTND1P17 | 0.16 | 0.98 | 0.32 | 1.15 | -0.82 | 6.75E-04 | 4.71E-03 | (B) Pathology_classA-RNA |
| RNU6-669P | 0.13 | 0.95 | 0.27 | 1.22 | -0.82 | 8.74E-04 | 5.71E-03 | (B) Pathology_classA-RNA |
| IRF8 | 0.17 | 0.99 | 0.37 | 1.2 | -0.82 | 1.47E-03 | 8.50E-03 | (B) Pathology_classA-RNA |
| NEK6 | 1.08 | 1.9 | 0.58 | 0.66 | -0.82 | 2.59E-03 | 0.0131 | (B) Pathology_classA-RNA |
| LINC00972 | 0.08 | 0.89 | 0.17 | 1.51 | -0.82 | 3.87E-03 | 0.0177 | (B) Pathology_classA-RNA |
| RN7SL125P | 0.22 | 1.03 | 0.46 | 1.36 | -0.82 | 5.76E-03 | 0.0239 | (B) Pathology_classA-RNA |
| ARL8B | 0.78 | 1.6 | 0.61 | 1.11 | -0.82 | 7.02E-03 | 0.0276 | (B) Pathology_classA-RNA |
| UBE2E2 | 0.88 | 1.7 | 0.68 | 0.81 | -0.82 | 7.43E-03 | 0.0288 | (B) Pathology_classA-RNA |
| RN7SL733P | 0.17 | 1 | 0.41 | 0.96 | -0.83 | 4.78E-04 | 3.61E-03 | (B) Pathology_classA-RNA |
| HIGD1AP11 | 0.12 | 0.95 | 0.15 | 1.26 | -0.83 | 6.46E-04 | 4.56E-03 | (B) Pathology_classA-RNA |
| MRPS33P2 | 0.22 | 1.05 | 0.44 | 1.31 | -0.83 | 3.58E-03 | 0.0167 | (B) Pathology_classA-RNA |
| NBPF15 | 5.18 | 6 | 0.62 | 0.72 | -0.83 | 3.87E-03 | 0.0177 | (B) Pathology_classA-RNA |
| RN7SL815P | 0.18 | 1.01 | 0.36 | 1.45 | -0.83 | 4.60E-03 | 0.0202 | (B) Pathology_classA-RNA |
| EMX2 | 0.43 | 1.26 | 0.62 | 1.06 | -0.83 | 6.57E-03 | 0.0263 | (B) Pathology_classA-RNA |
| SIRPAP1 | 0.75 | 1.58 | 0.72 | 1.01 | -0.83 | 0.0119 | 0.0404 | (B) Pathology_classA-RNA |
| TCF4 | 0.52 | 1.36 | 0.73 | 0.97 | -0.83 | 0.0121 | 0.0409 | (B) Pathology_classA-RNA |
| BNIP3L | 6.05 | 6.88 | 0.64 | 1.36 | -0.83 | 0.0144 | 0.0464 | (B) Pathology_classA-RNA |
| ZNF469 | 1.09 | 1.92 | 0.77 | 0.88 | -0.83 | 0.0145 | 0.0468 | (B) Pathology_classA-RNA |
| RNU6-853P | 0.69 | 1.51 | 0.67 | 1.31 | -0.83 | 0.015 | 0.0479 | (B) Pathology_classA-RNA |
| RNA5SP498 | 0 | 0.85 | 0.01 | 1 | -0.84 | 2.43E-05 | 3.56E-04 | (B) Pathology_classA-RNA |
| TMEM64 | 0.64 | 1.48 | 0.31 | 0.87 | -0.84 | 4.29E-05 | 5.62E-04 | (B) Pathology_classA-RNA |

| | | | | | | | | |
|---|---|---|---|---|---|---|---|---|
| OR4F8P | 0.08 | 0.92 | 0.21 | 1.1 | -0.84 | 1.71E-04 | 1.63E-03 | (B) Pathology_classA-RNA |
| MRPS28 | 0.04 | 0.88 | 0.11 | 1.18 | -0.84 | 2.20E-04 | 1.98E-03 | (B) Pathology_classA-RNA |
| MTND2P4 | 0.06 | 0.9 | 0.19 | 1.3 | -0.84 | 9.23E-04 | 5.96E-03 | (B) Pathology_classA-RNA |
| REEP6 | 0.14 | 0.98 | 0.41 | 1.17 | -0.84 | 1.40E-03 | 8.16E-03 | (B) Pathology_classA-RNA |
| DISC1-IT1 | 0.16 | 1 | 0.4 | 1.19 | -0.84 | 1.45E-03 | 8.41E-03 | (B) Pathology_classA-RNA |
| LEF1-AS1 | 0.92 | 1.76 | 0.5 | 1.07 | -0.84 | 1.98E-03 | 0.0106 | (B) Pathology_classA-RNA |
| GIMAP8 | 0.42 | 1.26 | 0.56 | 0.92 | -0.84 | 2.51E-03 | 0.0127 | (B) Pathology_classA-RNA |
| RAB18 | 0.46 | 1.3 | 0.56 | 1.09 | -0.84 | 3.85E-03 | 0.0177 | (B) Pathology_classA-RNA |
| SNORA70G | 0.18 | 1.03 | 0.55 | 1.17 | -0.84 | 4.14E-03 | 0.0187 | (B) Pathology_classA-RNA |
| CEBPB | 1.8 | 2.64 | 0.53 | 1.29 | -0.84 | 5.34E-03 | 0.0225 | (B) Pathology_classA-RNA |
| EPN2-AS1 | 0.31 | 1.16 | 0.59 | 1.13 | -0.84 | 5.37E-03 | 0.0226 | (B) Pathology_classA-RNA |
| MIR-4767/47 | 0.3 | 1.14 | 0.5 | 1.45 | -0.84 | 7.71E-03 | 0.0295 | (B) Pathology_classA-RNA |
| RNU6-757P | 0.84 | 1.68 | 0.6 | 1.3 | -0.84 | 8.91E-03 | 0.0328 | (B) Pathology_classA-RNA |
| LMOD3 | 0.05 | 0.9 | 0.14 | 0.61 | -0.85 | 3.86E-09 | 3.79E-07 | (B) Pathology_classA-RNA |
| SAR1B | 8.13 | 8.98 | 0.2 | 0.82 | -0.85 | 2.41E-06 | 5.94E-05 | (B) Pathology_classA-RNA |
| PIAS2 | 0.17 | 1.02 | 0.33 | 0.89 | -0.85 | 6.91E-05 | 8.15E-04 | (B) Pathology_classA-RNA |
| MIR-4762/3P | 0.07 | 0.92 | 0.1 | 1.11 | -0.85 | 9.29E-05 | 1.03E-03 | (B) Pathology_classA-RNA |
| MIR-4762/5P | 0.07 | 0.92 | 0.1 | 1.11 | -0.85 | 9.29E-05 | 1.03E-03 | (B) Pathology_classA-RNA |
| EDEM1 | 3.01 | 3.86 | 0.43 | 0.66 | -0.85 | 1.50E-04 | 1.48E-03 | (B) Pathology_classA-RNA |
| FRMD4B | 0.24 | 1.08 | 0.42 | 0.78 | -0.85 | 1.99E-04 | 1.83E-03 | (B) Pathology_classA-RNA |
| DPEP2 | 0.02 | 0.86 | 0.03 | 1.27 | -0.85 | 4.65E-04 | 3.54E-03 | (B) Pathology_classA-RNA |
| ULK4P1 | 0.32 | 1.17 | 0.58 | 0.91 | -0.85 | 2.55E-03 | 0.0129 | (B) Pathology_classA-RNA |
| TPSAB1 | 0.23 | 1.08 | 0.47 | 1.49 | -0.85 | 7.52E-03 | 0.029 | (B) Pathology_classA-RNA |
| SLC2A4 | 0.68 | 1.53 | 0.6 | 1.33 | -0.85 | 8.74E-03 | 0.0323 | (B) Pathology_classA-RNA |
| MYL6P2 | 0.54 | 1.39 | 0.69 | 1.12 | -0.85 | 9.77E-03 | 0.0351 | (B) Pathology_classA-RNA |
| ATP5MC2P3 | 2.58 | 3.43 | 0.57 | 1.46 | -0.85 | 0.0105 | 0.0369 | (B) Pathology_classA-RNA |
| RSC1A1 | 0.02 | 0.88 | 0.06 | 0.6 | -0.86 | 9.72E-10 | 1.34E-07 | (B) Pathology_classA-RNA |
| LINC02603 | 2.97 | 3.83 | 0.32 | 0.71 | -0.86 | 8.61E-06 | 1.62E-04 | (B) Pathology_classA-RNA |
| AKR1D1P1 | 0.02 | 0.87 | 0.05 | 1 | -0.86 | 2.08E-05 | 3.17E-04 | (B) Pathology_classA-RNA |
| GAPDHP33 | 0.2 | 1.06 | 0.28 | 1.15 | -0.86 | 3.02E-04 | 2.54E-03 | (B) Pathology_classA-RNA |
| OR7K1P | 0.04 | 0.9 | 0.13 | 1.23 | -0.86 | 3.14E-04 | 2.62E-03 | (B) Pathology_classA-RNA |
| CACUL1 | 5.39 | 6.25 | 0.49 | 0.45 | -0.86 | 4.89E-04 | 3.68E-03 | (B) Pathology_classA-RNA |
| SLC24A4 | 0.09 | 0.96 | 0.28 | 1.22 | -0.86 | 5.15E-04 | 3.82E-03 | (B) Pathology_classA-RNA |
| RN7SL693P | 0.59 | 1.45 | 0.53 | 1.16 | -0.86 | 2.86E-03 | 0.0141 | (B) Pathology_classA-RNA |
| MIR-505/3P | 0.13 | 0.99 | 0.35 | 1.45 | -0.86 | 3.44E-03 | 0.0162 | (B) Pathology_classA-RNA |
| MIR-505/505 | 0.13 | 0.99 | 0.35 | 1.45 | -0.86 | 3.44E-03 | 0.0162 | (B) Pathology_classA-RNA |
| MIR-505/505 | 0.13 | 0.99 | 0.35 | 1.45 | -0.86 | 3.44E-03 | 0.0162 | (B) Pathology_classA-RNA |
| MIR-505/5P | 0.13 | 0.99 | 0.35 | 1.45 | -0.86 | 3.44E-03 | 0.0162 | (B) Pathology_classA-RNA |
| MIR-4731/3P | 0.18 | 1.05 | 0.39 | 1.43 | -0.86 | 3.53E-03 | 0.0165 | (B) Pathology_classA-RNA |
| MIR-4731/5P | 0.18 | 1.05 | 0.39 | 1.43 | -0.86 | 3.53E-03 | 0.0165 | (B) Pathology_classA-RNA |
| SLC9B1P3 | 0.08 | 0.94 | 0.25 | 1.54 | -0.86 | 3.71E-03 | 0.0171 | (B) Pathology_classA-RNA |
| MS4A14 | 0.94 | 1.8 | 0.62 | 1.04 | -0.86 | 4.74E-03 | 0.0207 | (B) Pathology_classA-RNA |
| TAS2R20 | 3.78 | 4.63 | 0.66 | 0.82 | -0.86 | 5.14E-03 | 0.022 | (B) Pathology_classA-RNA |
| MED15P7 | 0.32 | 1.19 | 0.5 | 1.49 | -0.86 | 7.34E-03 | 0.0285 | (B) Pathology_classA-RNA |
| LINC00160 | 0.24 | 1.1 | 0.4 | 1.78 | -0.86 | 0.0134 | 0.0442 | (B) Pathology_classA-RNA |
| YAP1 | 0 | 0.87 | 0.01 | 0.7 | -0.87 | 3.07E-08 | 1.88E-06 | (B) Pathology_classA-RNA |
| MAP2K4P1 | 0.32 | 1.18 | 0.34 | 0.85 | -0.87 | 4.06E-05 | 5.35E-04 | (B) Pathology_classA-RNA |
| VPS13D | 1.41 | 2.28 | 0.42 | 0.43 | -0.87 | 1.08E-04 | 1.16E-03 | (B) Pathology_classA-RNA |
| GAPDHP16 | 0.04 | 0.91 | 0.09 | 1.16 | -0.87 | 1.28E-04 | 1.31E-03 | (B) Pathology_classA-RNA |
| RN7SL672P | 0.08 | 0.95 | 0.2 | 1.14 | -0.87 | 1.62E-04 | 1.57E-03 | (B) Pathology_classA-RNA |
| RN7SL757P | 0.02 | 0.9 | 0.06 | 1.2 | -0.87 | 1.72E-04 | 1.64E-03 | (B) Pathology_classA-RNA |
| ACTN1-AS1 | 0.26 | 1.12 | 0.41 | 0.95 | -0.87 | 2.70E-04 | 2.31E-03 | (B) Pathology_classA-RNA |
| UBTFL5 | 0.22 | 1.09 | 0.34 | 1.24 | -0.87 | 8.27E-04 | 5.49E-03 | (B) Pathology_classA-RNA |
| RNU4ATAC11 | 0.24 | 1.11 | 0.43 | 1.19 | -0.87 | 1.33E-03 | 7.85E-03 | (B) Pathology_classA-RNA |
| PRKX-AS1 | 0.19 | 1.06 | 0.4 | 1.29 | -0.87 | 1.69E-03 | 9.44E-03 | (B) Pathology_classA-RNA |
| PAOX | 5.01 | 5.88 | 0.53 | 1.04 | -0.87 | 1.77E-03 | 9.76E-03 | (B) Pathology_classA-RNA |
| C10ORF131 | 2.71 | 3.58 | 0.51 | 1.11 | -0.87 | 1.85E-03 | 0.0101 | (B) Pathology_classA-RNA |
| SHOC2 | 1.67 | 2.54 | 0.59 | 0.48 | -0.87 | 1.98E-03 | 0.0106 | (B) Pathology_classA-RNA |
| CDC27P1 | 4.36 | 5.23 | 0.73 | 0.92 | -0.87 | 8.43E-03 | 0.0315 | (B) Pathology_classA-RNA |
| ADIRF-AS1 | 0.3 | 1.18 | 0.51 | 1.63 | -0.87 | 0.0112 | 0.0387 | (B) Pathology_classA-RNA |
| IFFO2 | 3.85 | 4.73 | 0.82 | 0.83 | -0.87 | 0.0144 | 0.0465 | (B) Pathology_classA-RNA |
| LINC00968 | 0.03 | 0.9 | 0.07 | 0.97 | -0.88 | 8.98E-06 | 1.67E-04 | (B) Pathology_classA-RNA |
| RNU7-28P | 0.03 | 0.91 | 0.1 | 1.09 | -0.88 | 5.08E-05 | 6.39E-04 | (B) Pathology_classA-RNA |
| HNRNPA1P62 | 0.1 | 0.98 | 0.22 | 1.14 | -0.88 | 1.49E-04 | 1.48E-03 | (B) Pathology_classA-RNA |
| HIST2H2AC | 0.14 | 1.02 | 0.2 | 1.2 | -0.88 | 2.39E-04 | 2.11E-03 | (B) Pathology_classA-RNA |

| | | | | | | | | |
|---|---|---|---|---|---|---|---|---|
| C9ORF170 | 0.31 | 1.2 | 0.42 | 1.04 | -0.88 | 4.44E-04 | 3.42E-03 | (B) Pathology_classA-RNA |
| RHEBP3 | 0.2 | 1.08 | 0.41 | 1.1 | -0.88 | 5.83E-04 | 4.21E-03 | (B) Pathology_classA-RNA |
| API5P2 | 0.28 | 1.16 | 0.25 | 1.45 | -0.88 | 1.76E-03 | 9.71E-03 | (B) Pathology_classA-RNA |
| GAN | 4.96 | 5.84 | 0.55 | 1.09 | -0.88 | 2.24E-03 | 0.0116 | (B) Pathology_classA-RNA |
| RPS15AP16 | 3.13 | 4 | 0.57 | 1.52 | -0.88 | 9.93E-03 | 0.0355 | (B) Pathology_classA-RNA |
| BTF3P12 | 2.72 | 3.59 | 0.78 | 0.83 | -0.88 | 0.011 | 0.0382 | (B) Pathology_classA-RNA |
| RBL2 | 1.17 | 2.04 | 0.79 | 1.07 | -0.88 | 0.0139 | 0.0454 | (B) Pathology_classA-RNA |
| MIR-30E/30E | 0.16 | 1.03 | 0.48 | 1.76 | -0.88 | 0.0141 | 0.0458 | (B) Pathology_classA-RNA |
| MIR-30E/30E | 0.16 | 1.03 | 0.48 | 1.76 | -0.88 | 0.0141 | 0.0458 | (B) Pathology_classA-RNA |
| MIR-30E/3P | 0.16 | 1.03 | 0.48 | 1.76 | -0.88 | 0.0141 | 0.0458 | (B) Pathology_classA-RNA |
| MIR-30E/5P | 0.16 | 1.03 | 0.48 | 1.76 | -0.88 | 0.0141 | 0.0458 | (B) Pathology_classA-RNA |
| POM121L8P | 0.04 | 0.93 | 0.09 | 1.17 | -0.89 | 1.04E-04 | 1.12E-03 | (B) Pathology_classA-RNA |
| RPL36AP41 | 0.04 | 0.93 | 0.12 | 1.22 | -0.89 | 1.82E-04 | 1.71E-03 | (B) Pathology_classA-RNA |
| RN7SL748P | 0.07 | 0.95 | 0.2 | 1.19 | -0.89 | 1.85E-04 | 1.73E-03 | (B) Pathology_classA-RNA |
| TMEM266 | 1.2 | 2.09 | 0.47 | 0.97 | -0.89 | 5.59E-04 | 4.08E-03 | (B) Pathology_classA-RNA |
| TRBV25-1 | 0.21 | 1.1 | 0.45 | 1.19 | -0.89 | 1.21E-03 | 7.30E-03 | (B) Pathology_classA-RNA |
| NADK2-AS1 | 3.69 | 4.58 | 0.4 | 1.42 | -0.89 | 2.51E-03 | 0.0128 | (B) Pathology_classA-RNA |
| TRBV10-2 | 0.13 | 1.02 | 0.35 | 1.56 | -0.89 | 3.97E-03 | 0.0181 | (B) Pathology_classA-RNA |
| NPM1P25 | 3.04 | 3.92 | 0.66 | 1.26 | -0.89 | 8.06E-03 | 0.0305 | (B) Pathology_classA-RNA |
| MIR-4511/45 | 0.21 | 1.1 | 0.5 | 1.62 | -0.89 | 9.27E-03 | 0.0338 | (B) Pathology_classA-RNA |
| MIR-151B/15 | 0.44 | 1.33 | 0.68 | 1.44 | -0.89 | 0.0128 | 0.0428 | (B) Pathology_classA-RNA |
| PTPRK | 0.66 | 1.56 | 0.82 | 0.88 | -0.89 | 0.013 | 0.0432 | (B) Pathology_classA-RNA |
| TGIF2 | 2.82 | 3.71 | 0.86 | 0.82 | -0.89 | 0.0156 | 0.0494 | (B) Pathology_classA-RNA |
| RNU7-175P | 0.05 | 0.94 | 0.07 | 1.09 | -0.9 | 3.59E-05 | 4.88E-04 | (B) Pathology_classA-RNA |
| DEFB109B | 0.15 | 1.04 | 0.44 | 1.17 | -0.9 | 9.09E-04 | 5.89E-03 | (B) Pathology_classA-RNA |
| RNF148 | 1.1 | 1.99 | 0.52 | 1.31 | -0.9 | 3.48E-03 | 0.0163 | (B) Pathology_classA-RNA |
| SMIM10L2B | 1.35 | 2.25 | 0.64 | 1.29 | -0.9 | 6.64E-03 | 0.0265 | (B) Pathology_classA-RNA |
| SLC25A3P2 | 0.65 | 1.55 | 0.75 | 0.69 | -0.9 | 6.97E-03 | 0.0275 | (B) Pathology_classA-RNA |
| TSPOAP1-AS | 1.64 | 2.54 | 0.71 | 1.25 | -0.9 | 9.97E-03 | 0.0356 | (B) Pathology_classA-RNA |
| TRAJ9 | 0.01 | 0.92 | 0.03 | 1.07 | -0.91 | 2.10E-05 | 3.19E-04 | (B) Pathology_classA-RNA |
| SYT15 | 0.71 | 1.61 | 0.36 | 0.9 | -0.91 | 5.74E-05 | 7.05E-04 | (B) Pathology_classA-RNA |
| FMR1-AS1 | 0.14 | 1.05 | 0.26 | 1.26 | -0.91 | 3.46E-04 | 2.81E-03 | (B) Pathology_classA-RNA |
| DUX4L9 | 0.13 | 1.05 | 0.4 | 1.18 | -0.91 | 5.79E-04 | 4.19E-03 | (B) Pathology_classA-RNA |
| RNU6-194P | 0.06 | 0.97 | 0.19 | 1.4 | -0.91 | 7.95E-04 | 5.35E-03 | (B) Pathology_classA-RNA |
| RPL23AP50 | 0.14 | 1.05 | 0.42 | 1.24 | -0.91 | 1.00E-03 | 6.33E-03 | (B) Pathology_classA-RNA |
| LRRN1 | 0.05 | 0.96 | 0.15 | 1.46 | -0.91 | 1.07E-03 | 6.65E-03 | (B) Pathology_classA-RNA |
| GLB1L3 | 0.16 | 1.07 | 0.32 | 1.44 | -0.91 | 1.57E-03 | 8.93E-03 | (B) Pathology_classA-RNA |
| OR2A3P | 0.29 | 1.2 | 0.55 | 1.34 | -0.91 | 3.92E-03 | 0.0179 | (B) Pathology_classA-RNA |
| FAM45BP | 3.63 | 4.53 | 0.69 | 0.98 | -0.91 | 5.17E-03 | 0.022 | (B) Pathology_classA-RNA |
| HMGB1P21 | 0.23 | 1.13 | 0.57 | 1.37 | -0.91 | 5.34E-03 | 0.0225 | (B) Pathology_classA-RNA |
| RNU1-19P | 0.18 | 1.09 | 0.55 | 1.53 | -0.91 | 7.36E-03 | 0.0286 | (B) Pathology_classA-RNA |
| RN7SL92P | 0.58 | 1.49 | 0.68 | 1.32 | -0.91 | 8.35E-03 | 0.0312 | (B) Pathology_classA-RNA |
| H2BFS | 1.76 | 2.67 | 0.73 | 1.18 | -0.91 | 9.07E-03 | 0.0332 | (B) Pathology_classA-RNA |
| AKR1B10P1 | 0.03 | 0.95 | 0.09 | 1.05 | -0.92 | 1.62E-05 | 2.63E-04 | (B) Pathology_classA-RNA |
| TRAV39 | 0.07 | 1 | 0.22 | 1.16 | -0.92 | 8.82E-05 | 9.88E-04 | (B) Pathology_classA-RNA |
| UBL5P4 | 0.32 | 1.23 | 0.39 | 1.29 | -0.92 | 9.18E-04 | 5.93E-03 | (B) Pathology_classA-RNA |
| BBOF1 | 3.71 | 4.64 | 0.58 | 0.92 | -0.92 | 1.32E-03 | 7.81E-03 | (B) Pathology_classA-RNA |
| ZNF697 | 0.35 | 1.27 | 0.56 | 0.99 | -0.92 | 1.38E-03 | 8.06E-03 | (B) Pathology_classA-RNA |
| ZNF652 | 1.59 | 2.5 | 0.62 | 0.72 | -0.92 | 1.78E-03 | 9.81E-03 | (B) Pathology_classA-RNA |
| CTSS | 0.14 | 1.06 | 0.41 | 1.47 | -0.92 | 2.58E-03 | 0.0131 | (B) Pathology_classA-RNA |
| PCDH9-AS3 | 0.13 | 1.05 | 0.29 | 1.65 | -0.92 | 3.80E-03 | 0.0175 | (B) Pathology_classA-RNA |
| ADAMTS9 | 0.24 | 1.16 | 0.51 | 1.5 | -0.92 | 4.97E-03 | 0.0214 | (B) Pathology_classA-RNA |
| LILRA5 | 0.05 | 0.97 | 0.15 | 1.81 | -0.92 | 6.10E-03 | 0.0249 | (B) Pathology_classA-RNA |
| SHMT1P1 | 0.62 | 1.54 | 0.83 | 1.18 | -0.92 | 0.0155 | 0.0491 | (B) Pathology_classA-RNA |
| COQ10BP2 | 0.29 | 1.22 | 0.26 | 1.12 | -0.93 | 7.47E-05 | 8.68E-04 | (B) Pathology_classA-RNA |
| HTR1F | 0 | 0.93 | 0 | 1.2 | -0.93 | 7.77E-05 | 8.91E-04 | (B) Pathology_classA-RNA |
| IFNG-AS1 | 0.07 | 1 | 0.12 | 1.22 | -0.93 | 1.08E-04 | 1.15E-03 | (B) Pathology_classA-RNA |
| C1QTNF9-AS | 0.02 | 0.96 | 0.07 | 1.28 | -0.93 | 1.76E-04 | 1.67E-03 | (B) Pathology_classA-RNA |
| SNORD116-1 | 0.22 | 1.15 | 0.3 | 1.3 | -0.93 | 4.79E-04 | 3.62E-03 | (B) Pathology_classA-RNA |
| THAP12P3 | 1 | 1.92 | 0.44 | 1.41 | -0.93 | 2.16E-03 | 0.0113 | (B) Pathology_classA-RNA |
| HERPUD1 | 0.69 | 1.62 | 0.54 | 1.24 | -0.93 | 2.19E-03 | 0.0114 | (B) Pathology_classA-RNA |
| CASC11 | 0.21 | 1.15 | 0.55 | 1.23 | -0.93 | 2.20E-03 | 0.0115 | (B) Pathology_classA-RNA |
| PRSS48 | 0.28 | 1.21 | 0.54 | 1.3 | -0.93 | 2.73E-03 | 0.0136 | (B) Pathology_classA-RNA |
| CDK17 | 1.07 | 2 | 0.71 | 0.74 | -0.93 | 4.10E-03 | 0.0185 | (B) Pathology_classA-RNA |
| CT45A6 | 0.02 | 0.95 | 0.06 | 1.9 | -0.93 | 7.62E-03 | 0.0293 | (B) Pathology_classA-RNA |

| | | | | | | | | |
|---|---|---|---|---|---|---|---|---|
| ZNF652P1 | 0.02 | 0.96 | 0.07 | 1.27 | -0.94 | 1.31E-04 | 1.33E-03 | (B) Pathology_classA-RNA |
| TIFAB | 0.06 | 1 | 0.14 | 1.3 | -0.94 | 2.06E-04 | 1.88E-03 | (B) Pathology_classA-RNA |
| TRBJ2-1 | 0.03 | 0.97 | 0.09 | 1.38 | -0.94 | 4.22E-04 | 3.29E-03 | (B) Pathology_classA-RNA |
| IQGAP2 | 0.17 | 1.12 | 0.52 | 0.95 | -0.94 | 5.79E-04 | 4.20E-03 | (B) Pathology_classA-RNA |
| CYP4Z2P | 0.12 | 1.06 | 0.36 | 1.32 | -0.94 | 7.28E-04 | 4.99E-03 | (B) Pathology_classA-RNA |
| RN7SKP54 | 0.05 | 0.99 | 0.1 | 1.52 | -0.94 | 1.07E-03 | 6.62E-03 | (B) Pathology_classA-RNA |
| CSKMT | 3.12 | 4.06 | 0.66 | 0.75 | -0.94 | 2.32E-03 | 0.012 | (B) Pathology_classA-RNA |
| HEG1 | 4.86 | 5.8 | 0.7 | 1.22 | -0.94 | 6.35E-03 | 0.0257 | (B) Pathology_classA-RNA |
| RPL7AP60 | 0.61 | 1.56 | 0.78 | 0.76 | -0.94 | 6.72E-03 | 0.0267 | (B) Pathology_classA-RNA |
| RNU6-1016P | 3.8 | 4.74 | 0.79 | 1.19 | -0.94 | 0.0105 | 0.037 | (B) Pathology_classA-RNA |
| DUBR | 2.26 | 3.21 | 0.83 | 1.2 | -0.94 | 0.0133 | 0.044 | (B) Pathology_classA-RNA |
| DICER1 | 3.42 | 4.37 | 0.17 | 0.55 | -0.95 | 1.01E-10 | 2.41E-08 | (B) Pathology_classA-RNA |
| PNCK | 0.04 | 0.99 | 0.11 | 1.24 | -0.95 | 9.14E-05 | 1.02E-03 | (B) Pathology_classA-RNA |
| RNU2-65P | 0.08 | 1.03 | 0.14 | 1.26 | -0.95 | 1.36E-04 | 1.37E-03 | (B) Pathology_classA-RNA |
| RN7SL204P | 0.12 | 1.06 | 0.25 | 1.22 | -0.95 | 1.53E-04 | 1.50E-03 | (B) Pathology_classA-RNA |
| RPL7L1P2 | 0.08 | 1.03 | 0.24 | 1.36 | -0.95 | 4.30E-04 | 3.33E-03 | (B) Pathology_classA-RNA |
| TAS2R40 | 0.24 | 1.2 | 0.49 | 1.21 | -0.95 | 9.46E-04 | 6.06E-03 | (B) Pathology_classA-RNA |
| OR5P4P | 0.1 | 1.05 | 0.19 | 1.59 | -0.95 | 1.63E-03 | 9.16E-03 | (B) Pathology_classA-RNA |
| RNF149 | 3.61 | 4.55 | 0.68 | 0.81 | -0.95 | 2.87E-03 | 0.0141 | (B) Pathology_classA-RNA |
| GHRLOS | 0.83 | 1.78 | 0.71 | 0.74 | -0.95 | 3.59E-03 | 0.0167 | (B) Pathology_classA-RNA |
| MIR-603/603 | 0.2 | 1.15 | 0.5 | 1.53 | -0.95 | 3.93E-03 | 0.0179 | (B) Pathology_classA-RNA |
| DKK1 | 0.2 | 1.15 | 0.61 | 1.4 | -0.95 | 5.13E-03 | 0.0219 | (B) Pathology_classA-RNA |
| CSF1R | 0.37 | 1.33 | 0.81 | 1.31 | -0.95 | 0.0127 | 0.0424 | (B) Pathology_classA-RNA |
| GPR160 | 5.75 | 6.69 | 0.75 | 1.55 | -0.95 | 0.0154 | 0.0488 | (B) Pathology_classA-RNA |
| SLC2A3P4 | 0.03 | 0.99 | 0.08 | 1.06 | -0.96 | 8.12E-06 | 1.55E-04 | (B) Pathology_classA-RNA |
| VAV3 | 0.28 | 1.24 | 0.33 | 0.95 | -0.96 | 2.09E-05 | 3.17E-04 | (B) Pathology_classA-RNA |
| LARGE-AS1 | 0.04 | 0.99 | 0.11 | 1.12 | -0.96 | 2.32E-05 | 3.44E-04 | (B) Pathology_classA-RNA |
| DUXAP7 | 0.22 | 1.18 | 0.32 | 1.1 | -0.96 | 6.90E-05 | 8.14E-04 | (B) Pathology_classA-RNA |
| DNMBP-AS1 | 0.88 | 1.84 | 0.52 | 0.68 | -0.96 | 2.77E-04 | 2.36E-03 | (B) Pathology_classA-RNA |
| RNU6-513P | 1.97 | 2.93 | 0.38 | 1.27 | -0.96 | 4.88E-04 | 3.68E-03 | (B) Pathology_classA-RNA |
| FCRL2 | 0.01 | 0.97 | 0.03 | 1.46 | -0.96 | 5.44E-04 | 4.00E-03 | (B) Pathology_classA-RNA |
| RNU6-132P | 0.14 | 1.09 | 0.28 | 1.39 | -0.96 | 5.83E-04 | 4.21E-03 | (B) Pathology_classA-RNA |
| RESF1 | 0.21 | 1.18 | 0.41 | 1.32 | -0.96 | 7.69E-04 | 5.20E-03 | (B) Pathology_classA-RNA |
| UHRF2P1 | 1.07 | 2.03 | 0.51 | 1.39 | -0.96 | 2.27E-03 | 0.0118 | (B) Pathology_classA-RNA |
| CCDC83 | 0.19 | 1.16 | 0.58 | 1.46 | -0.96 | 4.51E-03 | 0.0199 | (B) Pathology_classA-RNA |
| LSM3P5 | 0.27 | 1.23 | 0.6 | 1.45 | -0.96 | 5.11E-03 | 0.0219 | (B) Pathology_classA-RNA |
| TBX3 | 0.27 | 1.23 | 0.82 | 1.04 | -0.96 | 9.80E-03 | 0.0352 | (B) Pathology_classA-RNA |
| FDFT1 | 1.22 | 2.18 | 0.85 | 1.19 | -0.96 | 0.0133 | 0.0439 | (B) Pathology_classA-RNA |
| RNU6-118P | 2.7 | 3.66 | 0.85 | 1.29 | -0.96 | 0.0153 | 0.0488 | (B) Pathology_classA-RNA |
| MARCH1 | 0.07 | 1.03 | 0.2 | 0.69 | -0.97 | 1.10E-08 | 8.60E-07 | (B) Pathology_classA-RNA |
| HIST1H2BN | 0.24 | 1.21 | 0.28 | 0.81 | -0.97 | 1.06E-06 | 3.13E-05 | (B) Pathology_classA-RNA |
| SLMAP | 2.47 | 3.44 | 0.34 | 0.47 | -0.97 | 2.59E-06 | 6.28E-05 | (B) Pathology_classA-RNA |
| GLI1 | 0.06 | 1.04 | 0.1 | 1.12 | -0.97 | 1.72E-05 | 2.77E-04 | (B) Pathology_classA-RNA |
| USP12-AS1 | 0.04 | 1.01 | 0.08 | 1.13 | -0.97 | 1.84E-05 | 2.89E-04 | (B) Pathology_classA-RNA |
| MRPS10P2 | 0.08 | 1.04 | 0.22 | 1.2 | -0.97 | 7.81E-05 | 8.94E-04 | (B) Pathology_classA-RNA |
| TENT4B | 0.92 | 1.89 | 0.52 | 0.55 | -0.97 | 2.57E-04 | 2.22E-03 | (B) Pathology_classA-RNA |
| GLUD1 | 0.24 | 1.2 | 0.49 | 1.27 | -0.97 | 1.13E-03 | 6.92E-03 | (B) Pathology_classA-RNA |
| OACYLP | 1 | 1.97 | 0.33 | 1.6 | -0.97 | 2.03E-03 | 0.0109 | (B) Pathology_classA-RNA |
| PDCD4-AS1 | 2.67 | 3.64 | 0.67 | 0.81 | -0.97 | 2.12E-03 | 0.0112 | (B) Pathology_classA-RNA |
| DDX58 | 4.72 | 5.69 | 0.78 | 0.96 | -0.97 | 6.44E-03 | 0.0259 | (B) Pathology_classA-RNA |
| RN7SL382P | 0.58 | 1.55 | 0.59 | 1.6 | -0.97 | 6.90E-03 | 0.0273 | (B) Pathology_classA-RNA |
| EIF1AX-AS1 | 0.76 | 1.73 | 0.61 | 1.59 | -0.97 | 7.45E-03 | 0.0288 | (B) Pathology_classA-RNA |
| GPR171 | 0.01 | 1 | 0.04 | 0.99 | -0.98 | 1.78E-06 | 4.70E-05 | (B) Pathology_classA-RNA |
| STARD13-IT1 | 0.03 | 1.01 | 0.08 | 1.18 | -0.98 | 2.94E-05 | 4.16E-04 | (B) Pathology_classA-RNA |
| GCSHP5 | 3.21 | 4.19 | 0.4 | 1.19 | -0.98 | 2.51E-04 | 2.19E-03 | (B) Pathology_classA-RNA |
| RNU6-1208P | 0.15 | 1.13 | 0.34 | 1.41 | -0.98 | 7.12E-04 | 4.91E-03 | (B) Pathology_classA-RNA |
| RPS3P1 | 0.24 | 1.22 | 0.38 | 1.43 | -0.98 | 9.52E-04 | 6.08E-03 | (B) Pathology_classA-RNA |
| IQCJ-SCHIP1- | 0.47 | 1.46 | 0.48 | 1.62 | -0.98 | 3.92E-03 | 0.0179 | (B) Pathology_classA-RNA |
| CASP16P | 0.73 | 1.72 | 0.8 | 0.83 | -0.98 | 6.36E-03 | 0.0257 | (B) Pathology_classA-RNA |
| RN7SKP292 | 0.6 | 1.59 | 0.69 | 1.5 | -0.98 | 7.75E-03 | 0.0296 | (B) Pathology_classA-RNA |
| RNU6-969P | 0.91 | 1.89 | 0.72 | 1.7 | -0.98 | 0.0143 | 0.0463 | (B) Pathology_classA-RNA |
| RPS7P7 | 0.01 | 1.01 | 0.03 | 1.08 | -0.99 | 6.79E-06 | 1.34E-04 | (B) Pathology_classA-RNA |
| MRPL15P1 | 0.17 | 1.16 | 0.23 | 1.18 | -0.99 | 4.35E-05 | 5.67E-04 | (B) Pathology_classA-RNA |
| RN7SL846P | 0.09 | 1.09 | 0.19 | 1.44 | -0.99 | 3.91E-04 | 3.10E-03 | (B) Pathology_classA-RNA |
| TRBV6-1 | 0.09 | 1.09 | 0.28 | 1.47 | -0.99 | 6.41E-04 | 4.53E-03 | (B) Pathology_classA-RNA |

| | | | | | | | | |
|---|---|---|---|---|---|---|---|---|
| EIF4A1P11 | 0.36 | 1.35 | 0.58 | 1.5 | -0.99 | 4.02E-03 | 0.0182 | (B) Pathology_classA-RNA |
| HIST1H4H | 0.1 | 1.1 | 0.3 | 1.06 | -1 | 1.92E-05 | 2.99E-04 | (B) Pathology_classA-RNA |
| ANTXRL | 0.04 | 1.04 | 0.12 | 1.26 | -1 | 5.99E-05 | 7.28E-04 | (B) Pathology_classA-RNA |
| NEIL2 | 4.51 | 5.51 | 0.48 | 1.05 | -1 | 2.54E-04 | 2.21E-03 | (B) Pathology_classA-RNA |
| TRAJ38 | 0.06 | 1.06 | 0.18 | 1.43 | -1 | 3.60E-04 | 2.91E-03 | (B) Pathology_classA-RNA |
| CCBE1 | 0.33 | 1.32 | 0.56 | 1.09 | -1 | 8.72E-04 | 5.70E-03 | (B) Pathology_classA-RNA |
| RAB27B | 0.2 | 1.2 | 0.6 | 1.1 | -1 | 1.28E-03 | 7.64E-03 | (B) Pathology_classA-RNA |
| SDC2 | 0.68 | 1.68 | 0.46 | 1.49 | -1 | 1.76E-03 | 9.71E-03 | (B) Pathology_classA-RNA |
| AK4 | 0.81 | 1.81 | 0.64 | 1.41 | -1 | 4.37E-03 | 0.0195 | (B) Pathology_classA-RNA |
| MIR-3150A/3 | 0.76 | 1.77 | 0.72 | 1.27 | -1 | 4.99E-03 | 0.0215 | (B) Pathology_classA-RNA |
| MIR-3150A/3 | 0.76 | 1.77 | 0.72 | 1.27 | -1 | 4.99E-03 | 0.0215 | (B) Pathology_classA-RNA |
| MIR-3150A/5 | 0.76 | 1.77 | 0.72 | 1.27 | -1 | 4.99E-03 | 0.0215 | (B) Pathology_classA-RNA |
| TMEM150B | 0.85 | 1.85 | 0.88 | 1.31 | -1 | 0.0138 | 0.0452 | (B) Pathology_classA-RNA |
| MIR-2276/22 | 0.01 | 1.03 | 0.03 | 1.18 | -1.01 | 1.87E-05 | 2.93E-04 | (B) Pathology_classA-RNA |
| MTND5P16 | 0.02 | 1.03 | 0.06 | 1.25 | -1.01 | 4.57E-05 | 5.89E-04 | (B) Pathology_classA-RNA |
| RNU2-6P | 0.06 | 1.07 | 0.17 | 1.27 | -1.01 | 6.15E-05 | 7.43E-04 | (B) Pathology_classA-RNA |
| LSM3P4 | 0.03 | 1.04 | 0.1 | 1.31 | -1.01 | 9.57E-05 | 1.05E-03 | (B) Pathology_classA-RNA |
| TRBV7-4 | 0.04 | 1.05 | 0.1 | 1.37 | -1.01 | 1.53E-04 | 1.50E-03 | (B) Pathology_classA-RNA |
| ELK4 | 6.24 | 7.25 | 0.47 | 1.09 | -1.01 | 2.65E-04 | 2.28E-03 | (B) Pathology_classA-RNA |
| KRT223P | 0.16 | 1.17 | 0.33 | 1.33 | -1.01 | 2.82E-04 | 2.40E-03 | (B) Pathology_classA-RNA |
| VTRNA1-1 | 0.15 | 1.16 | 0.31 | 1.38 | -1.01 | 3.49E-04 | 2.84E-03 | (B) Pathology_classA-RNA |
| RNU6-135P | 0.23 | 1.24 | 0.37 | 1.33 | -1.01 | 3.76E-04 | 3.01E-03 | (B) Pathology_classA-RNA |
| PLEK | 0.03 | 1.04 | 0.1 | 1.5 | -1.01 | 4.31E-04 | 3.33E-03 | (B) Pathology_classA-RNA |
| BAZ1A | 0.46 | 1.47 | 0.58 | 0.77 | -1.01 | 5.09E-04 | 3.79E-03 | (B) Pathology_classA-RNA |
| RPS26P11 | 4.68 | 5.7 | 0.44 | 1.38 | -1.01 | 7.66E-04 | 5.19E-03 | (B) Pathology_classA-RNA |
| SETP9 | 0.3 | 1.31 | 0.52 | 1.25 | -1.01 | 8.56E-04 | 5.63E-03 | (B) Pathology_classA-RNA |
| MIR-4665/3P | 0.25 | 1.26 | 0.45 | 1.4 | -1.01 | 9.31E-04 | 5.98E-03 | (B) Pathology_classA-RNA |
| MIR-4665/5P | 0.25 | 1.26 | 0.45 | 1.4 | -1.01 | 9.31E-04 | 5.98E-03 | (B) Pathology_classA-RNA |
| HBBP1 | 0.01 | 1.02 | 0.03 | 1.66 | -1.01 | 1.22E-03 | 7.33E-03 | (B) Pathology_classA-RNA |
| FERMT3 | 1.29 | 2.3 | 0.53 | 1.43 | -1.01 | 1.93E-03 | 0.0104 | (B) Pathology_classA-RNA |
| WWC2 | 2.88 | 3.89 | 0.66 | 1.24 | -1.01 | 2.92E-03 | 0.0143 | (B) Pathology_classA-RNA |
| USP15 | 0.96 | 1.97 | 0.65 | 1.27 | -1.01 | 3.08E-03 | 0.0149 | (B) Pathology_classA-RNA |
| RSF1-IT1 | 0.65 | 1.66 | 0.64 | 1.34 | -1.01 | 3.33E-03 | 0.0158 | (B) Pathology_classA-RNA |
| PDCD4 | 0.46 | 1.48 | 0.31 | 0.91 | -1.02 | 3.74E-06 | 8.42E-05 | (B) Pathology_classA-RNA |
| DUTP7 | 0.05 | 1.07 | 0.16 | 1.24 | -1.02 | 4.43E-05 | 5.74E-04 | (B) Pathology_classA-RNA |
| LAX1 | 0.04 | 1.06 | 0.12 | 1.34 | -1.02 | 1.02E-04 | 1.10E-03 | (B) Pathology_classA-RNA |
| RNU6-123P | 0.09 | 1.11 | 0.24 | 1.31 | -1.02 | 1.29E-04 | 1.32E-03 | (B) Pathology_classA-RNA |
| MIR-2117/21 | 0.07 | 1.09 | 0.21 | 1.35 | -1.02 | 1.51E-04 | 1.49E-03 | (B) Pathology_classA-RNA |
| ADARB1 | 0.21 | 1.23 | 0.47 | 1.02 | -1.02 | 1.63E-04 | 1.58E-03 | (B) Pathology_classA-RNA |
| RPL23AP51 | 0.21 | 1.23 | 0.29 | 1.35 | -1.02 | 2.20E-04 | 1.97E-03 | (B) Pathology_classA-RNA |
| RNU6-1143P | 0.34 | 1.36 | 0.45 | 1.33 | -1.02 | 6.02E-04 | 4.32E-03 | (B) Pathology_classA-RNA |
| RNU6-541P | 0.24 | 1.26 | 0.37 | 1.43 | -1.02 | 6.40E-04 | 4.53E-03 | (B) Pathology_classA-RNA |
| ACADSB | 0.71 | 1.73 | 0.45 | 1.39 | -1.02 | 8.54E-04 | 5.62E-03 | (B) Pathology_classA-RNA |
| RN7SL656P | 0.56 | 1.58 | 0.67 | 0.9 | -1.02 | 1.59E-03 | 8.99E-03 | (B) Pathology_classA-RNA |
| RNU6-720P | 0.64 | 1.66 | 0.6 | 1.29 | -1.02 | 1.85E-03 | 0.0101 | (B) Pathology_classA-RNA |
| IGKV2D-40 | 0.16 | 1.18 | 0.47 | 1.61 | -1.02 | 2.49E-03 | 0.0127 | (B) Pathology_classA-RNA |
| NHSL2 | 2.31 | 3.34 | 0.94 | 1.23 | -1.02 | 0.0154 | 0.049 | (B) Pathology_classA-RNA |
| GPR132 | 0.08 | 1.11 | 0.2 | 1.15 | -1.03 | 1.60E-05 | 2.61E-04 | (B) Pathology_classA-RNA |
| ENTPD1-AS1 | 0.63 | 1.66 | 0.5 | 0.85 | -1.03 | 1.26E-04 | 1.30E-03 | (B) Pathology_classA-RNA |
| IFT57 | 0.06 | 1.09 | 0.18 | 1.4 | -1.03 | 1.78E-04 | 1.68E-03 | (B) Pathology_classA-RNA |
| LINC00621 | 0.16 | 1.19 | 0.22 | 1.6 | -1.03 | 8.83E-04 | 5.76E-03 | (B) Pathology_classA-RNA |
| LINC00676 | 0.06 | 1.1 | 0.13 | 1.64 | -1.03 | 9.44E-04 | 6.05E-03 | (B) Pathology_classA-RNA |
| KRR1P1 | 2.19 | 3.21 | 0.54 | 1.38 | -1.03 | 1.47E-03 | 8.47E-03 | (B) Pathology_classA-RNA |
| MTND2P29 | 0.22 | 1.24 | 0.65 | 1.58 | -1.03 | 5.77E-03 | 0.0239 | (B) Pathology_classA-RNA |
| RNU6-539P | 0.59 | 1.62 | 0.7 | 1.61 | -1.03 | 7.92E-03 | 0.0301 | (B) Pathology_classA-RNA |
| SHOC1 | 0 | 1.05 | 0.01 | 1.14 | -1.04 | 6.88E-06 | 1.36E-04 | (B) Pathology_classA-RNA |
| AKR1B15 | 0.1 | 1.14 | 0.15 | 1.39 | -1.04 | 1.36E-04 | 1.37E-03 | (B) Pathology_classA-RNA |
| LINC01549 | 0.35 | 1.39 | 0.42 | 1.51 | -1.04 | 1.05E-03 | 6.56E-03 | (B) Pathology_classA-RNA |
| TRAJ57 | 0.15 | 1.19 | 0.34 | 1.62 | -1.04 | 1.31E-03 | 7.77E-03 | (B) Pathology_classA-RNA |
| DNM3-IT1 | 0.43 | 1.47 | 0.59 | 1.37 | -1.04 | 1.88E-03 | 0.0102 | (B) Pathology_classA-RNA |
| RNA5SP92 | 0.32 | 1.36 | 0.65 | 1.44 | -1.04 | 3.50E-03 | 0.0164 | (B) Pathology_classA-RNA |
| JTBP1 | 0.74 | 1.78 | 0.72 | 1.47 | -1.04 | 5.74E-03 | 0.0238 | (B) Pathology_classA-RNA |
| RNVU1-14 | 0.96 | 2 | 0.8 | 1.45 | -1.04 | 8.71E-03 | 0.0322 | (B) Pathology_classA-RNA |
| RNU6-105P | 3.14 | 4.18 | 0.88 | 1.51 | -1.04 | 0.0146 | 0.047 | (B) Pathology_classA-RNA |
| ZBTB20-AS3 | 0 | 1.05 | 0.01 | 1.1 | -1.05 | 3.53E-06 | 8.03E-05 | (B) Pathology_classA-RNA |

| | | | | | | | | |
|---|---|---|---|---|---|---|---|---|
| RBPMSLP | 0.06 | 1.11 | 0.18 | 1.26 | -1.05 | 3.85E-05 | 5.15E-04 | (B) Pathology_classA-RNA |
| RNA5SP221 | 0.03 | 1.07 | 0.08 | 1.44 | -1.05 | 1.71E-04 | 1.63E-03 | (B) Pathology_classA-RNA |
| RPS20P15 | 0.18 | 1.22 | 0.4 | 1.27 | -1.05 | 2.00E-04 | 1.84E-03 | (B) Pathology_classA-RNA |
| RNU2-72P | 0.13 | 1.17 | 0.26 | 1.65 | -1.05 | 1.03E-03 | 6.45E-03 | (B) Pathology_classA-RNA |
| CAPN11 | 0.48 | 1.53 | 0.69 | 1.01 | -1.05 | 1.87E-03 | 0.0102 | (B) Pathology_classA-RNA |
| RN7SL749P | 0.27 | 1.33 | 0.6 | 1.57 | -1.05 | 3.48E-03 | 0.0163 | (B) Pathology_classA-RNA |
| SERPINB4 | 0.36 | 1.41 | 0.54 | 1.88 | -1.05 | 6.66E-03 | 0.0265 | (B) Pathology_classA-RNA |
| MZB1 | 0.35 | 1.4 | 0.7 | 1.81 | -1.05 | 0.0105 | 0.0368 | (B) Pathology_classA-RNA |
| CCDC196 | 0.05 | 1.11 | 0.15 | 0.97 | -1.06 | 4.52E-07 | 1.59E-05 | (B) Pathology_classA-RNA |
| CEBPA-DT | 0.09 | 1.15 | 0.27 | 1.2 | -1.06 | 2.70E-05 | 3.88E-04 | (B) Pathology_classA-RNA |
| SEL1L | 5.46 | 6.52 | 0.43 | 0.95 | -1.06 | 3.14E-05 | 4.40E-04 | (B) Pathology_classA-RNA |
| AIMP1P1 | 0.11 | 1.17 | 0.28 | 1.31 | -1.06 | 7.87E-05 | 8.98E-04 | (B) Pathology_classA-RNA |
| RNU6-906P | 0.06 | 1.12 | 0.19 | 1.41 | -1.06 | 1.39E-04 | 1.39E-03 | (B) Pathology_classA-RNA |
| CLDN24 | 0.37 | 1.43 | 0.51 | 1.19 | -1.06 | 3.61E-04 | 2.91E-03 | (B) Pathology_classA-RNA |
| NBAT1 | 0.34 | 1.4 | 0.31 | 1.5 | -1.06 | 4.63E-04 | 3.52E-03 | (B) Pathology_classA-RNA |
| CD53 | 0.16 | 1.22 | 0.48 | 1.5 | -1.06 | 1.20E-03 | 7.25E-03 | (B) Pathology_classA-RNA |
| CYBRD1 | 0.96 | 2.01 | 0.64 | 1.13 | -1.06 | 1.31E-03 | 7.78E-03 | (B) Pathology_classA-RNA |
| LINC00694 | 0.05 | 1.11 | 0.14 | 1.77 | -1.06 | 1.43E-03 | 8.32E-03 | (B) Pathology_classA-RNA |
| LST1 | 0.13 | 1.2 | 0.4 | 1.65 | -1.06 | 1.54E-03 | 8.80E-03 | (B) Pathology_classA-RNA |
| NAP1L3 | 0.33 | 1.38 | 0.62 | 1.35 | -1.06 | 1.97E-03 | 0.0106 | (B) Pathology_classA-RNA |
| MAST4-IT1 | 0.24 | 1.31 | 0.47 | 1.64 | -1.06 | 2.00E-03 | 0.0107 | (B) Pathology_classA-RNA |
| PCMTD1P2 | 0.32 | 1.38 | 0.46 | 1.71 | -1.06 | 2.61E-03 | 0.0132 | (B) Pathology_classA-RNA |
| MYO1F | 0.17 | 1.23 | 0.44 | 1.74 | -1.06 | 2.69E-03 | 0.0135 | (B) Pathology_classA-RNA |
| RNU6-155P | 0.34 | 1.39 | 0.68 | 1.64 | -1.06 | 6.27E-03 | 0.0254 | (B) Pathology_classA-RNA |
| KLHDC7B | 0.56 | 1.62 | 0.81 | 1.38 | -1.06 | 7.15E-03 | 0.028 | (B) Pathology_classA-RNA |
| TBCAP3 | 1.53 | 2.59 | 0.75 | 1.59 | -1.06 | 7.62E-03 | 0.0293 | (B) Pathology_classA-RNA |
| RNA5SP256 | 0.22 | 1.28 | 0.66 | 1.83 | -1.06 | 8.47E-03 | 0.0316 | (B) Pathology_classA-RNA |
| MSRB3 | 0.94 | 2 | 0.83 | 1.42 | -1.06 | 8.72E-03 | 0.0323 | (B) Pathology_classA-RNA |
| SNORA37 | 0.03 | 1.1 | 0.1 | 1.42 | -1.07 | 1.19E-04 | 1.23E-03 | (B) Pathology_classA-RNA |
| TRBV24-1 | 0.07 | 1.14 | 0.14 | 1.56 | -1.07 | 3.74E-04 | 3.00E-03 | (B) Pathology_classA-RNA |
| RPL6P8 | 0.13 | 1.2 | 0.28 | 1.51 | -1.07 | 3.91E-04 | 3.10E-03 | (B) Pathology_classA-RNA |
| ANPEP | 0.16 | 1.23 | 0.31 | 1.62 | -1.07 | 8.43E-04 | 5.58E-03 | (B) Pathology_classA-RNA |
| RNU6-1316P | 0.2 | 1.26 | 0.39 | 1.56 | -1.07 | 8.88E-04 | 5.78E-03 | (B) Pathology_classA-RNA |
| CMTM6 | 0.77 | 1.84 | 0.7 | 0.91 | -1.07 | 1.49E-03 | 8.59E-03 | (B) Pathology_classA-RNA |
| ADAM8 | 5.18 | 6.24 | 0.89 | 1.4 | -1.07 | 0.0109 | 0.038 | (B) Pathology_classA-RNA |
| CHST15 | 5.02 | 6.08 | 0.99 | 1.12 | -1.07 | 0.0139 | 0.0454 | (B) Pathology_classA-RNA |
| DDX18P1 | 0.18 | 1.26 | 0.32 | 1.19 | -1.08 | 3.27E-05 | 4.54E-04 | (B) Pathology_classA-RNA |
| DPYD-AS1 | 1.68 | 2.76 | 0.44 | 1.23 | -1.08 | 1.71E-04 | 1.63E-03 | (B) Pathology_classA-RNA |
| RNU6-523P | 0.15 | 1.23 | 0.44 | 1.43 | -1.08 | 5.08E-04 | 3.78E-03 | (B) Pathology_classA-RNA |
| DDX21 | 1.68 | 2.76 | 0.64 | 0.76 | -1.08 | 6.45E-04 | 4.55E-03 | (B) Pathology_classA-RNA |
| HMGN2P9 | 0.16 | 1.24 | 0.49 | 1.5 | -1.08 | 1.07E-03 | 6.65E-03 | (B) Pathology_classA-RNA |
| PRRX1 | 0.05 | 1.13 | 0.11 | 1.75 | -1.08 | 1.08E-03 | 6.70E-03 | (B) Pathology_classA-RNA |
| BLZF2P | 0.27 | 1.35 | 0.59 | 1.48 | -1.08 | 1.90E-03 | 0.0103 | (B) Pathology_classA-RNA |
| CCDC88B | 0.38 | 1.46 | 0.68 | 1.61 | -1.08 | 4.86E-03 | 0.021 | (B) Pathology_classA-RNA |
| RNF43 | 4.96 | 6.04 | 0.81 | 1.7 | -1.08 | 0.0109 | 0.0379 | (B) Pathology_classA-RNA |
| CAB39L | 3.1 | 4.18 | 0.88 | 1.64 | -1.08 | 0.0137 | 0.0449 | (B) Pathology_classA-RNA |
| RN7SKP64 | 0.34 | 1.42 | 0.89 | 1.62 | -1.08 | 0.0138 | 0.0451 | (B) Pathology_classA-RNA |
| UBE2CP3 | 0.06 | 1.15 | 0.14 | 1.36 | -1.09 | 5.67E-05 | 6.99E-04 | (B) Pathology_classA-RNA |
| RPS2P44 | 0.14 | 1.23 | 0.29 | 1.34 | -1.09 | 8.62E-05 | 9.70E-04 | (B) Pathology_classA-RNA |
| HMGB1P27 | 0.25 | 1.34 | 0.43 | 1.22 | -1.09 | 1.16E-04 | 1.21E-03 | (B) Pathology_classA-RNA |
| CSF2RB | 0.05 | 1.14 | 0.14 | 1.52 | -1.09 | 2.22E-04 | 1.99E-03 | (B) Pathology_classA-RNA |
| RNU1-51P | 0.26 | 1.35 | 0.55 | 1.37 | -1.09 | 8.69E-04 | 5.69E-03 | (B) Pathology_classA-RNA |
| POTEE | 0.57 | 1.66 | 0.58 | 1.39 | -1.09 | 1.21E-03 | 7.32E-03 | (B) Pathology_classA-RNA |
| RN7SL258P | 0.93 | 2.02 | 0.78 | 1.19 | -1.09 | 3.69E-03 | 0.017 | (B) Pathology_classA-RNA |
| SULT1C4 | 1.12 | 2.21 | 0.71 | 1.51 | -1.09 | 4.13E-03 | 0.0186 | (B) Pathology_classA-RNA |
| C5AR2 | 0.57 | 1.66 | 0.72 | 1.65 | -1.09 | 6.02E-03 | 0.0247 | (B) Pathology_classA-RNA |
| HCG23 | 0.49 | 1.58 | 0.96 | 1.46 | -1.09 | 0.0149 | 0.0477 | (B) Pathology_classA-RNA |
| ZNF552 | 2.51 | 3.62 | 0.41 | 1.04 | -1.1 | 2.16E-05 | 3.25E-04 | (B) Pathology_classA-RNA |
| RN7SL350P | 0.03 | 1.14 | 0.1 | 1.36 | -1.1 | 4.43E-05 | 5.74E-04 | (B) Pathology_classA-RNA |
| GULOP | 0.25 | 1.35 | 0.52 | 1.34 | -1.1 | 5.05E-04 | 3.77E-03 | (B) Pathology_classA-RNA |
| RPL34P21 | 0.23 | 1.33 | 0.52 | 1.4 | -1.1 | 6.82E-04 | 4.75E-03 | (B) Pathology_classA-RNA |
| CACNA1C-IT3 | 0.13 | 1.23 | 0.27 | 1.66 | -1.1 | 7.41E-04 | 5.06E-03 | (B) Pathology_classA-RNA |
| TMCC3 | 0.72 | 1.81 | 0.91 | 0.88 | -1.1 | 6.81E-03 | 0.027 | (B) Pathology_classA-RNA |
| LSM3P2 | 0.32 | 1.43 | 0.77 | 1.71 | -1.1 | 8.17E-03 | 0.0307 | (B) Pathology_classA-RNA |
| ABHD11-AS1 | 0.48 | 1.58 | 0.93 | 1.38 | -1.1 | 0.0107 | 0.0374 | (B) Pathology_classA-RNA |

| | | | | | | | | |
|---|---|---|---|---|---|---|---|---|
| CT45A4 | 0.22 | 1.32 | 0.66 | 2.12 | -1.1 | 0.014 | 0.0455 | (B) Pathology_classA-RNA |
| TRAV8-4 | 0.01 | 1.13 | 0.04 | 1.28 | -1.11 | 1.43E-05 | 2.42E-04 | (B) Pathology_classA-RNA |
| SLC6A8 | 0.24 | 1.36 | 0.59 | 0.98 | -1.11 | 2.94E-04 | 2.48E-03 | (B) Pathology_classA-RNA |
| HNRNPA1P53 | 0.36 | 1.46 | 0.66 | 1.27 | -1.11 | 1.43E-03 | 8.30E-03 | (B) Pathology_classA-RNA |
| GK-IT1 | 0.22 | 1.33 | 0.62 | 1.45 | -1.11 | 1.74E-03 | 9.65E-03 | (B) Pathology_classA-RNA |
| AQP3 | 0.3 | 1.42 | 0.64 | 1.46 | -1.11 | 1.96E-03 | 0.0105 | (B) Pathology_classA-RNA |
| SNORA14B | 0.81 | 1.91 | 0.71 | 1.39 | -1.11 | 2.93E-03 | 0.0143 | (B) Pathology_classA-RNA |
| TDO2 | 0.07 | 1.18 | 0.21 | 1.99 | -1.11 | 2.96E-03 | 0.0145 | (B) Pathology_classA-RNA |
| FAM166B | 2.44 | 3.55 | 0.8 | 1.28 | -1.11 | 4.24E-03 | 0.019 | (B) Pathology_classA-RNA |
| CD82 | 1.38 | 2.49 | 0.95 | 1.17 | -1.11 | 9.86E-03 | 0.0353 | (B) Pathology_classA-RNA |
| ARHGAP10 | 4.73 | 5.84 | 1.03 | 1.19 | -1.11 | 0.0147 | 0.0472 | (B) Pathology_classA-RNA |
| GBP2 | 2.67 | 3.78 | 0.99 | 1.45 | -1.11 | 0.0153 | 0.0486 | (B) Pathology_classA-RNA |
| ASH1L-IT1 | 0.26 | 1.38 | 0.29 | 1.31 | -1.12 | 4.32E-05 | 5.65E-04 | (B) Pathology_classA-RNA |
| DCAF6 | 2.36 | 3.48 | 0.54 | 1.03 | -1.12 | 1.53E-04 | 1.50E-03 | (B) Pathology_classA-RNA |
| RNU7-41P | 0.17 | 1.28 | 0.35 | 1.5 | -1.12 | 3.05E-04 | 2.56E-03 | (B) Pathology_classA-RNA |
| MIR-3136/3P | 0.33 | 1.45 | 0.55 | 1.21 | -1.12 | 3.43E-04 | 2.79E-03 | (B) Pathology_classA-RNA |
| MIR-3136/5P | 0.33 | 1.45 | 0.55 | 1.21 | -1.12 | 3.43E-04 | 2.79E-03 | (B) Pathology_classA-RNA |
| IGKV1OR10-1 | 0.24 | 1.36 | 0.48 | 1.52 | -1.12 | 6.77E-04 | 4.72E-03 | (B) Pathology_classA-RNA |
| MTRNR2L2 | 0.15 | 1.27 | 0.44 | 1.58 | -1.12 | 7.27E-04 | 4.99E-03 | (B) Pathology_classA-RNA |
| MIR-711/711 | 1.7 | 2.81 | 0.68 | 1.49 | -1.12 | 2.74E-03 | 0.0137 | (B) Pathology_classA-RNA |
| RFLNB | 1.34 | 2.45 | 0.72 | 1.62 | -1.12 | 4.79E-03 | 0.0208 | (B) Pathology_classA-RNA |
| RNU6-101P | 0.33 | 1.44 | 0.65 | 1.79 | -1.12 | 5.19E-03 | 0.0221 | (B) Pathology_classA-RNA |
| LINC00327 | 1.56 | 2.68 | 0.65 | 1.85 | -1.12 | 5.97E-03 | 0.0245 | (B) Pathology_classA-RNA |
| APOBR | 0.86 | 1.99 | 0.81 | 1.7 | -1.12 | 8.60E-03 | 0.0319 | (B) Pathology_classA-RNA |
| FCF1P7 | 0.63 | 1.75 | 0.9 | 1.5 | -1.12 | 9.79E-03 | 0.0351 | (B) Pathology_classA-RNA |
| MED28P3 | 0.05 | 1.18 | 0.06 | 1.1 | -1.13 | 9.12E-07 | 2.76E-05 | (B) Pathology_classA-RNA |
| HSPA8P19 | 0 | 1.13 | 0.01 | 1.21 | -1.13 | 4.93E-06 | 1.04E-04 | (B) Pathology_classA-RNA |
| MTND2P33 | 0.1 | 1.23 | 0.17 | 1.46 | -1.13 | 9.52E-05 | 1.05E-03 | (B) Pathology_classA-RNA |
| ATP6V0E1P2 | 1.06 | 2.19 | 0.53 | 1.2 | -1.13 | 2.38E-04 | 2.10E-03 | (B) Pathology_classA-RNA |
| CTSO | 6.27 | 7.4 | 0.59 | 1.46 | -1.13 | 1.24E-03 | 7.43E-03 | (B) Pathology_classA-RNA |
| RPS3P7 | 0.5 | 1.63 | 0.64 | 1.66 | -1.13 | 3.17E-03 | 0.0152 | (B) Pathology_classA-RNA |
| HNRNPA1P15 | 0.44 | 1.57 | 0.87 | 1.61 | -1.13 | 9.45E-03 | 0.0342 | (B) Pathology_classA-RNA |
| RPL7L1P9 | 1.33 | 2.46 | 0.92 | 1.62 | -1.13 | 0.0117 | 0.04 | (B) Pathology_classA-RNA |
| SSR4P1 | 2.42 | 3.55 | 1.04 | 0.99 | -1.13 | 0.0126 | 0.0423 | (B) Pathology_classA-RNA |
| MACROD2-IT | 0.26 | 1.4 | 0.38 | 1.41 | -1.14 | 1.60E-04 | 1.55E-03 | (B) Pathology_classA-RNA |
| ARID5B | 4.86 | 6 | 0.65 | 0.87 | -1.14 | 4.78E-04 | 3.61E-03 | (B) Pathology_classA-RNA |
| RWDD4P1 | 2.76 | 3.9 | 0.68 | 0.7 | -1.14 | 6.47E-04 | 4.57E-03 | (B) Pathology_classA-RNA |
| RPL9P30 | 0.24 | 1.38 | 0.49 | 1.51 | -1.14 | 6.48E-04 | 4.57E-03 | (B) Pathology_classA-RNA |
| USP46-AS1 | 3.47 | 4.61 | 0.71 | 1.17 | -1.14 | 1.40E-03 | 8.16E-03 | (B) Pathology_classA-RNA |
| LRRC37A15P | 3.69 | 4.82 | 0.76 | 1.67 | -1.14 | 5.58E-03 | 0.0233 | (B) Pathology_classA-RNA |
| RABGAP1L-IT | 0.88 | 2.02 | 0.81 | 1.69 | -1.14 | 7.54E-03 | 0.0291 | (B) Pathology_classA-RNA |
| GAPDHP42 | 1.2 | 2.33 | 0.84 | 1.91 | -1.14 | 0.013 | 0.0432 | (B) Pathology_classA-RNA |
| WWC3-AS1 | 0.46 | 1.61 | 0.45 | 1.5 | -1.15 | 3.74E-04 | 2.99E-03 | (B) Pathology_classA-RNA |
| OSMR | 0.41 | 1.56 | 0.61 | 1.24 | -1.15 | 5.47E-04 | 4.01E-03 | (B) Pathology_classA-RNA |
| RNU6-1294P | 0.19 | 1.34 | 0.57 | 1.6 | -1.15 | 1.47E-03 | 8.50E-03 | (B) Pathology_classA-RNA |
| PPP1R12BP2 | 0.16 | 1.31 | 0.38 | 1.91 | -1.15 | 2.22E-03 | 0.0115 | (B) Pathology_classA-RNA |
| RNU7-48P | 0.54 | 1.69 | 0.71 | 1.5 | -1.15 | 2.68E-03 | 0.0134 | (B) Pathology_classA-RNA |
| EMBP1 | 1.72 | 2.88 | 0.89 | 0.97 | -1.15 | 4.64E-03 | 0.0203 | (B) Pathology_classA-RNA |
| RN7SL767P | 1.16 | 2.31 | 0.85 | 1.74 | -1.15 | 9.12E-03 | 0.0334 | (B) Pathology_classA-RNA |
| PLAAT4 | 1.69 | 2.84 | 0.92 | 1.69 | -1.15 | 0.0119 | 0.0405 | (B) Pathology_classA-RNA |
| CHSY1 | 0.2 | 1.36 | 0.31 | 0.81 | -1.16 | 8.23E-08 | 4.12E-06 | (B) Pathology_classA-RNA |
| PLA2G4A | 0.01 | 1.17 | 0.04 | 1.04 | -1.16 | 2.32E-07 | 9.34E-06 | (B) Pathology_classA-RNA |
| RNU6-314P | 0.12 | 1.28 | 0.32 | 1.45 | -1.16 | 1.10E-04 | 1.17E-03 | (B) Pathology_classA-RNA |
| TUSC7 | 0.01 | 1.18 | 0.04 | 1.7 | -1.16 | 3.50E-04 | 2.84E-03 | (B) Pathology_classA-RNA |
| CAT | 1.71 | 2.86 | 0.51 | 1.73 | -1.16 | 1.61E-03 | 9.06E-03 | (B) Pathology_classA-RNA |
| ZNF727 | 0.23 | 1.39 | 0.68 | 1.59 | -1.16 | 2.52E-03 | 0.0128 | (B) Pathology_classA-RNA |
| RN7SL284P | 0.27 | 1.43 | 0.8 | 1.75 | -1.16 | 6.97E-03 | 0.0275 | (B) Pathology_classA-RNA |
| TNFAIP8L2 | 4.4 | 5.56 | 0.95 | 1.85 | -1.16 | 0.0155 | 0.0491 | (B) Pathology_classA-RNA |
| MIR-1273C/1 | 0.41 | 1.58 | 0.51 | 1.53 | -1.17 | 5.81E-04 | 4.21E-03 | (B) Pathology_classA-RNA |
| ZFP36L2 | 1.72 | 2.88 | 0.67 | 1.12 | -1.17 | 6.52E-04 | 4.59E-03 | (B) Pathology_classA-RNA |
| TRAV20 | 0.23 | 1.4 | 0.39 | 1.78 | -1.17 | 1.04E-03 | 6.48E-03 | (B) Pathology_classA-RNA |
| FAM238A | 1.03 | 2.21 | 0.52 | 1.74 | -1.17 | 1.54E-03 | 8.79E-03 | (B) Pathology_classA-RNA |
| SNORA11B | 0.35 | 1.52 | 0.73 | 1.78 | -1.17 | 5.18E-03 | 0.0221 | (B) Pathology_classA-RNA |
| MTND5P14 | 0.02 | 1.19 | 0.04 | 1.26 | -1.18 | 4.93E-06 | 1.04E-04 | (B) Pathology_classA-RNA |
| HMGB3P14 | 0.1 | 1.28 | 0.17 | 1.34 | -1.18 | 1.58E-05 | 2.59E-04 | (B) Pathology_classA-RNA |

| | | | | | | | | |
|---|---|---|---|---|---|---|---|---|
| TNFSF10 | 0.21 | 1.38 | 0.34 | 1.32 | -1.18 | 3.39E-05 | 4.65E-04 | (B) Pathology_classA-RNA |
| SUCLG2 | 0.21 | 1.39 | 0.56 | 1.13 | -1.18 | 1.57E-04 | 1.53E-03 | (B) Pathology_classA-RNA |
| RN7SL20P | 0.11 | 1.29 | 0.28 | 1.61 | -1.18 | 2.45E-04 | 2.14E-03 | (B) Pathology_classA-RNA |
| TSEN15P2 | 0.34 | 1.52 | 0.5 | 1.46 | -1.18 | 3.56E-04 | 2.87E-03 | (B) Pathology_classA-RNA |
| RAD21-AS1 | 3.17 | 4.35 | 0.68 | 1.01 | -1.18 | 6.09E-04 | 4.35E-03 | (B) Pathology_classA-RNA |
| IGHV4OR15-: | 0.15 | 1.33 | 0.24 | 1.96 | -1.18 | 1.58E-03 | 8.98E-03 | (B) Pathology_classA-RNA |
| SNORA14A | 0.36 | 1.54 | 0.55 | 1.84 | -1.18 | 2.47E-03 | 0.0126 | (B) Pathology_classA-RNA |
| DCSTAMP | 0.46 | 1.65 | 1.03 | 1.29 | -1.18 | 0.0108 | 0.0377 | (B) Pathology_classA-RNA |
| MIR-1296/12 | 0.05 | 1.23 | 0.14 | 1.46 | -1.19 | 4.33E-05 | 5.65E-04 | (B) Pathology_classA-RNA |
| MIR-1296/3P | 0.05 | 1.23 | 0.14 | 1.46 | -1.19 | 4.33E-05 | 5.65E-04 | (B) Pathology_classA-RNA |
| MIR-1296/5P | 0.05 | 1.23 | 0.14 | 1.46 | -1.19 | 4.33E-05 | 5.65E-04 | (B) Pathology_classA-RNA |
| NDUFA5P2 | 0.16 | 1.35 | 0.34 | 1.53 | -1.19 | 1.65E-04 | 1.59E-03 | (B) Pathology_classA-RNA |
| DCANP1 | 0.16 | 1.34 | 0.39 | 1.55 | -1.19 | 2.45E-04 | 2.15E-03 | (B) Pathology_classA-RNA |
| GIMAP7 | 0.52 | 1.71 | 0.87 | 1.84 | -1.19 | 9.85E-03 | 0.0353 | (B) Pathology_classA-RNA |
| THRB-AS1 | 0.08 | 1.28 | 0.22 | 0.79 | -1.2 | 1.18E-09 | 1.51E-07 | (B) Pathology_classA-RNA |
| RGS10 | 0.09 | 1.3 | 0.28 | 1.16 | -1.2 | 2.38E-06 | 5.88E-05 | (B) Pathology_classA-RNA |
| TCF7L1-IT1 | 0.05 | 1.25 | 0.16 | 1.37 | -1.2 | 1.50E-05 | 2.49E-04 | (B) Pathology_classA-RNA |
| MIR-4755/3P | 0.12 | 1.31 | 0.28 | 1.53 | -1.2 | 1.11E-04 | 1.17E-03 | (B) Pathology_classA-RNA |
| MIR-4755/5P | 0.12 | 1.31 | 0.28 | 1.53 | -1.2 | 1.11E-04 | 1.17E-03 | (B) Pathology_classA-RNA |
| KIRREL1-IT1 | 0.15 | 1.35 | 0.34 | 1.51 | -1.2 | 1.21E-04 | 1.26E-03 | (B) Pathology_classA-RNA |
| FAM30A | 0.11 | 1.31 | 0.33 | 1.65 | -1.2 | 2.96E-04 | 2.50E-03 | (B) Pathology_classA-RNA |
| RNU6-1327P | 0.16 | 1.36 | 0.39 | 1.64 | -1.2 | 3.80E-04 | 3.03E-03 | (B) Pathology_classA-RNA |
| COL18A1-AS1 | 0.22 | 1.42 | 0.43 | 1.67 | -1.2 | 5.32E-04 | 3.92E-03 | (B) Pathology_classA-RNA |
| GIMAP4 | 0.26 | 1.46 | 0.52 | 1.6 | -1.2 | 6.34E-04 | 4.49E-03 | (B) Pathology_classA-RNA |
| GRHL2 | 0.26 | 1.46 | 0.77 | 1.07 | -1.2 | 1.42E-03 | 8.25E-03 | (B) Pathology_classA-RNA |
| POC1B-GALN | 2.23 | 3.43 | 0.85 | 0.62 | -1.2 | 2.37E-03 | 0.0122 | (B) Pathology_classA-RNA |
| C15ORF48 | 0.36 | 1.56 | 0.8 | 1.49 | -1.2 | 3.45E-03 | 0.0163 | (B) Pathology_classA-RNA |
| RN7SL472P | 0.65 | 1.85 | 0.84 | 1.52 | -1.2 | 4.61E-03 | 0.0202 | (B) Pathology_classA-RNA |
| EAF2 | 3.69 | 4.9 | 1 | 1.15 | -1.2 | 7.60E-03 | 0.0292 | (B) Pathology_classA-RNA |
| LATS2 | 0.18 | 1.39 | 0.3 | 0.68 | -1.21 | 1.01E-08 | 8.01E-07 | (B) Pathology_classA-RNA |
| DPF3 | 0.2 | 1.41 | 0.28 | 1.52 | -1.21 | 8.92E-05 | 9.95E-04 | (B) Pathology_classA-RNA |
| IFI16 | 0.21 | 1.42 | 0.47 | 1.34 | -1.21 | 9.27E-05 | 1.03E-03 | (B) Pathology_classA-RNA |
| POTEF | 0.46 | 1.68 | 0.37 | 1.67 | -1.21 | 3.77E-04 | 3.01E-03 | (B) Pathology_classA-RNA |
| RNU6-608P | 0.11 | 1.32 | 0.32 | 1.76 | -1.21 | 5.19E-04 | 3.84E-03 | (B) Pathology_classA-RNA |
| IGHD5-12 | 0.14 | 1.35 | 0.41 | 1.96 | -1.21 | 1.81E-03 | 9.94E-03 | (B) Pathology_classA-RNA |
| SNORD71 | 0.33 | 1.54 | 0.74 | 1.64 | -1.21 | 3.00E-03 | 0.0146 | (B) Pathology_classA-RNA |
| ITGA8 | 1.68 | 2.89 | 0.65 | 1.87 | -1.21 | 3.44E-03 | 0.0162 | (B) Pathology_classA-RNA |
| TAF9BP1 | 1.28 | 2.48 | 0.92 | 1.45 | -1.21 | 6.29E-03 | 0.0255 | (B) Pathology_classA-RNA |
| TPTEP1 | 0.98 | 2.18 | 0.98 | 1.4 | -1.21 | 8.03E-03 | 0.0304 | (B) Pathology_classA-RNA |
| RNU4-38P | 0.75 | 1.96 | 0.98 | 1.56 | -1.21 | 9.63E-03 | 0.0347 | (B) Pathology_classA-RNA |
| TRBV4-2 | 0.15 | 1.37 | 0.24 | 1.62 | -1.22 | 1.46E-04 | 1.45E-03 | (B) Pathology_classA-RNA |
| CD1C | 0.18 | 1.41 | 0.55 | 1.49 | -1.22 | 4.07E-04 | 3.19E-03 | (B) Pathology_classA-RNA |
| STK17B | 0.75 | 1.96 | 0.63 | 1.26 | -1.22 | 4.23E-04 | 3.29E-03 | (B) Pathology_classA-RNA |
| MIR-581/581 | 0.41 | 1.63 | 0.59 | 1.45 | -1.22 | 5.11E-04 | 3.80E-03 | (B) Pathology_classA-RNA |
| MFAP4 | 0.25 | 1.47 | 0.5 | 2.03 | -1.22 | 3.04E-03 | 0.0147 | (B) Pathology_classA-RNA |
| GP1BA | 1.98 | 3.2 | 0.87 | 1.54 | -1.22 | 4.97E-03 | 0.0214 | (B) Pathology_classA-RNA |
| STON1 | 0.59 | 1.82 | 0.42 | 1.01 | -1.23 | 3.93E-06 | 8.74E-05 | (B) Pathology_classA-RNA |
| EIF2S2P5 | 0.03 | 1.26 | 0.1 | 1.35 | -1.23 | 8.02E-06 | 1.53E-04 | (B) Pathology_classA-RNA |
| ATP6V1G1P6 | 0.11 | 1.34 | 0.21 | 1.48 | -1.23 | 3.95E-05 | 5.24E-04 | (B) Pathology_classA-RNA |
| SOD2 | 0.27 | 1.5 | 0.47 | 1.33 | -1.23 | 7.64E-05 | 8.81E-04 | (B) Pathology_classA-RNA |
| KRT8P4 | 0.06 | 1.29 | 0.13 | 1.78 | -1.23 | 3.37E-04 | 2.75E-03 | (B) Pathology_classA-RNA |
| PNOC | 0.26 | 1.49 | 0.51 | 1.76 | -1.23 | 9.50E-04 | 6.08E-03 | (B) Pathology_classA-RNA |
| HMGN1P8 | 0.94 | 2.17 | 0.69 | 1.45 | -1.23 | 1.09E-03 | 6.72E-03 | (B) Pathology_classA-RNA |
| TRAV8-6 | 0.28 | 1.51 | 0.57 | 1.7 | -1.23 | 1.10E-03 | 6.76E-03 | (B) Pathology_classA-RNA |
| ABCA1 | 3.2 | 4.44 | 0.85 | 0.9 | -1.23 | 2.05E-03 | 0.0109 | (B) Pathology_classA-RNA |
| MTND5P1 | 1.21 | 2.44 | 0.82 | 1.26 | -1.23 | 2.16E-03 | 0.0113 | (B) Pathology_classA-RNA |
| PCED1B-AS1 | 1.32 | 2.55 | 0.69 | 1.88 | -1.23 | 3.80E-03 | 0.0175 | (B) Pathology_classA-RNA |
| DNTT | 0.21 | 1.44 | 0.22 | 2.42 | -1.23 | 5.94E-03 | 0.0244 | (B) Pathology_classA-RNA |
| ERAP2 | 0.88 | 2.1 | 0.98 | 1.49 | -1.23 | 8.15E-03 | 0.0307 | (B) Pathology_classA-RNA |
| TSC22D3 | 1.58 | 2.8 | 1.06 | 1.16 | -1.23 | 9.56E-03 | 0.0345 | (B) Pathology_classA-RNA |
| PI16 | 0.93 | 2.17 | 0.77 | 2.21 | -1.23 | 0.0103 | 0.0364 | (B) Pathology_classA-RNA |
| RPS3AP39 | 0.66 | 1.89 | 0.9 | 2.05 | -1.23 | 0.0126 | 0.0423 | (B) Pathology_classA-RNA |
| HMGB1P23 | 0.5 | 1.72 | 1.04 | 1.79 | -1.23 | 0.0142 | 0.0461 | (B) Pathology_classA-RNA |
| MYO1G | 1.87 | 3.11 | 0.75 | 1.52 | -1.24 | 1.84E-03 | 0.0101 | (B) Pathology_classA-RNA |
| TAB3-AS2 | 6.08 | 7.32 | 0.94 | 1.02 | -1.24 | 4.13E-03 | 0.0186 | (B) Pathology_classA-RNA |

| | | | | | | | | |
|---|---|---|---|---|---|---|---|---|
| RN7SL751P | 0.54 | 1.78 | 0.7 | 2.08 | -1.24 | 6.23E-03 | 0.0253 | (B) Pathology_classA-RNA |
| NPL | 3.04 | 4.28 | 0.99 | 1.47 | -1.24 | 8.06E-03 | 0.0304 | (B) Pathology_classA-RNA |
| MIR-4644/46 | 1.15 | 2.39 | 0.75 | 2.26 | -1.24 | 0.0104 | 0.0367 | (B) Pathology_classA-RNA |
| AIF1 | 0.55 | 1.79 | 0.94 | 1.96 | -1.24 | 0.0119 | 0.0405 | (B) Pathology_classA-RNA |
| MIR-1284/12 | 0.03 | 1.28 | 0.06 | 1.21 | -1.25 | 8.75E-07 | 2.67E-05 | (B) Pathology_classA-RNA |
| GOLGA4 | 1.58 | 2.83 | 0.43 | 0.67 | -1.25 | 1.62E-06 | 4.36E-05 | (B) Pathology_classA-RNA |
| CHPT1 | 0.79 | 2.04 | 0.55 | 1.09 | -1.25 | 5.67E-05 | 6.99E-04 | (B) Pathology_classA-RNA |
| MALT1 | 2.98 | 4.23 | 0.56 | 1.12 | -1.25 | 7.28E-05 | 8.49E-04 | (B) Pathology_classA-RNA |
| SNORA54 | 0.1 | 1.35 | 0.3 | 1.62 | -1.25 | 1.32E-04 | 1.34E-03 | (B) Pathology_classA-RNA |
| TRAJ53 | 0.06 | 1.31 | 0.19 | 1.66 | -1.25 | 1.35E-04 | 1.36E-03 | (B) Pathology_classA-RNA |
| RPL29P14 | 0.12 | 1.37 | 0.36 | 1.6 | -1.25 | 1.61E-04 | 1.56E-03 | (B) Pathology_classA-RNA |
| LINC00161 | 0.32 | 1.57 | 0.41 | 1.62 | -1.25 | 2.26E-04 | 2.02E-03 | (B) Pathology_classA-RNA |
| ST7-OT4 | 1.2 | 2.44 | 0.54 | 1.58 | -1.25 | 4.52E-04 | 3.46E-03 | (B) Pathology_classA-RNA |
| RNU7-19P | 0.14 | 1.4 | 0.43 | 1.93 | -1.25 | 1.23E-03 | 7.38E-03 | (B) Pathology_classA-RNA |
| RN7SL370P | 0.31 | 1.56 | 0.68 | 1.61 | -1.25 | 1.36E-03 | 8.00E-03 | (B) Pathology_classA-RNA |
| GALNT4 | 4.18 | 5.44 | 0.86 | 0.62 | -1.25 | 2.06E-03 | 0.011 | (B) Pathology_classA-RNA |
| RPL4P6 | 1.47 | 2.72 | 0.9 | 1.19 | -1.25 | 3.22E-03 | 0.0154 | (B) Pathology_classA-RNA |
| CYCSP41 | 0.16 | 1.41 | 0.48 | 2.5 | -1.25 | 9.25E-03 | 0.0337 | (B) Pathology_classA-RNA |
| RNU4-49P | 0.02 | 1.28 | 0.07 | 1.46 | -1.26 | 1.73E-05 | 2.77E-04 | (B) Pathology_classA-RNA |
| HIST2H3D | 0.25 | 1.52 | 0.47 | 1.68 | -1.26 | 4.11E-04 | 3.22E-03 | (B) Pathology_classA-RNA |
| EEA1 | 0.3 | 1.56 | 0.74 | 0.76 | -1.26 | 6.05E-04 | 4.34E-03 | (B) Pathology_classA-RNA |
| PREX1 | 1.81 | 3.07 | 0.74 | 1.21 | -1.26 | 7.82E-04 | 5.27E-03 | (B) Pathology_classA-RNA |
| TRAJ52 | 0.26 | 1.52 | 0.53 | 1.84 | -1.26 | 1.26E-03 | 7.53E-03 | (B) Pathology_classA-RNA |
| ZMYM4-AS1 | 4.62 | 5.88 | 0.83 | 0.95 | -1.26 | 1.53E-03 | 8.75E-03 | (B) Pathology_classA-RNA |
| KLF3P1 | 0.63 | 1.89 | 0.68 | 1.86 | -1.26 | 2.54E-03 | 0.0129 | (B) Pathology_classA-RNA |
| TRIM34 | 0.3 | 1.56 | 0.89 | 1.05 | -1.26 | 2.63E-03 | 0.0132 | (B) Pathology_classA-RNA |
| HIST2H2AA3 | 7.72 | 8.98 | 0.87 | 1.37 | -1.26 | 2.96E-03 | 0.0144 | (B) Pathology_classA-RNA |
| KRTAP13-2 | 0.1 | 1.36 | 0.2 | 2.33 | -1.26 | 3.68E-03 | 0.017 | (B) Pathology_classA-RNA |
| BMS1P17 | 2.36 | 3.62 | 1.05 | 1.58 | -1.26 | 0.0108 | 0.0376 | (B) Pathology_classA-RNA |
| HELB | 0.56 | 1.83 | 0.53 | 1.07 | -1.27 | 3.26E-05 | 4.54E-04 | (B) Pathology_classA-RNA |
| P3H2-AS1 | 0.1 | 1.38 | 0.31 | 1.71 | -1.27 | 2.10E-04 | 1.91E-03 | (B) Pathology_classA-RNA |
| GSR | 1.45 | 2.71 | 0.66 | 0.87 | -1.27 | 2.16E-04 | 1.95E-03 | (B) Pathology_classA-RNA |
| AAK1 | 2.47 | 3.73 | 0.71 | 0.81 | -1.27 | 4.25E-04 | 3.30E-03 | (B) Pathology_classA-RNA |
| GAPDHP51 | 0.48 | 1.75 | 0.49 | 1.71 | -1.27 | 5.18E-04 | 3.84E-03 | (B) Pathology_classA-RNA |
| JARID2-AS1 | 1.81 | 3.09 | 0.98 | 1.41 | -1.27 | 5.89E-03 | 0.0243 | (B) Pathology_classA-RNA |
| IL10 | 0.97 | 2.24 | 1.08 | 1.29 | -1.27 | 8.90E-03 | 0.0328 | (B) Pathology_classA-RNA |
| TNS4 | 0.37 | 1.63 | 0.77 | 2.24 | -1.27 | 9.26E-03 | 0.0337 | (B) Pathology_classA-RNA |
| RNU6-701P | 0.01 | 1.29 | 0.02 | 1.37 | -1.28 | 4.86E-06 | 1.03E-04 | (B) Pathology_classA-RNA |
| METTL21AP1 | 0.1 | 1.38 | 0.3 | 1.5 | -1.28 | 3.65E-05 | 4.94E-04 | (B) Pathology_classA-RNA |
| TNRC18P1 | 0.47 | 1.74 | 0.42 | 1.5 | -1.28 | 8.38E-05 | 9.47E-04 | (B) Pathology_classA-RNA |
| MYLK | 4.23 | 5.51 | 0.48 | 1.56 | -1.28 | 2.02E-04 | 1.85E-03 | (B) Pathology_classA-RNA |
| GNAT2 | 1.23 | 2.51 | 0.79 | 1.25 | -1.28 | 1.22E-03 | 7.35E-03 | (B) Pathology_classA-RNA |
| MIR-4728/3P | 2.65 | 3.93 | 0.84 | 1.52 | -1.28 | 2.66E-03 | 0.0133 | (B) Pathology_classA-RNA |
| MIR-4728/5P | 2.65 | 3.93 | 0.84 | 1.52 | -1.28 | 2.66E-03 | 0.0133 | (B) Pathology_classA-RNA |
| GPR34 | 2.7 | 3.98 | 0.93 | 1.04 | -1.28 | 3.06E-03 | 0.0148 | (B) Pathology_classA-RNA |
| MIR-1343/13 | 0.35 | 1.63 | 0.71 | 1.99 | -1.28 | 3.91E-03 | 0.0178 | (B) Pathology_classA-RNA |
| MIR-1343/3P | 0.35 | 1.63 | 0.71 | 1.99 | -1.28 | 3.91E-03 | 0.0178 | (B) Pathology_classA-RNA |
| MIR-1343/5P | 0.35 | 1.63 | 0.71 | 1.99 | -1.28 | 3.91E-03 | 0.0178 | (B) Pathology_classA-RNA |
| RNU105B | 0.37 | 1.65 | 0.81 | 1.85 | -1.28 | 4.38E-03 | 0.0195 | (B) Pathology_classA-RNA |
| GLTPP1 | 3.42 | 4.69 | 1.07 | 1.11 | -1.28 | 7.52E-03 | 0.029 | (B) Pathology_classA-RNA |
| VN1R40P | 0.42 | 1.7 | 0.5 | 2.53 | -1.28 | 8.80E-03 | 0.0325 | (B) Pathology_classA-RNA |
| BIRC6-AS1 | 0.13 | 1.42 | 0.27 | 1.31 | -1.29 | 3.89E-06 | 8.69E-05 | (B) Pathology_classA-RNA |
| M1AP | 0.15 | 1.44 | 0.3 | 1.34 | -1.29 | 6.60E-06 | 1.32E-04 | (B) Pathology_classA-RNA |
| RN7SKP26 | 0.11 | 1.4 | 0.32 | 1.36 | -1.29 | 9.93E-06 | 1.81E-04 | (B) Pathology_classA-RNA |
| MTCYBP3 | 0.19 | 1.48 | 0.44 | 1.55 | -1.29 | 1.22E-04 | 1.26E-03 | (B) Pathology_classA-RNA |
| APOL3 | 4.83 | 6.11 | 0.68 | 1.31 | -1.29 | 4.60E-04 | 3.51E-03 | (B) Pathology_classA-RNA |
| CCL19 | 0.03 | 1.31 | 0.06 | 1.93 | -1.29 | 4.69E-04 | 3.56E-03 | (B) Pathology_classA-RNA |
| EI24P4 | 0.05 | 1.34 | 0.14 | 2.29 | -1.29 | 2.50E-03 | 0.0127 | (B) Pathology_classA-RNA |
| PRRX2 | 1.26 | 2.55 | 0.93 | 1.84 | -1.29 | 7.11E-03 | 0.0279 | (B) Pathology_classA-RNA |
| VMP1 | 0.93 | 2.22 | 1.09 | 1.04 | -1.29 | 7.80E-03 | 0.0297 | (B) Pathology_classA-RNA |
| OR2B2 | 0.15 | 1.45 | 0.3 | 1.47 | -1.3 | 2.28E-05 | 3.38E-04 | (B) Pathology_classA-RNA |
| RNU5A-3P | 0.21 | 1.51 | 0.44 | 1.96 | -1.3 | 1.03E-03 | 6.47E-03 | (B) Pathology_classA-RNA |
| LINC00937 | 0.38 | 1.68 | 0.8 | 1.3 | -1.3 | 1.25E-03 | 7.48E-03 | (B) Pathology_classA-RNA |
| CT45A3 | 0.16 | 1.46 | 0.48 | 2.16 | -1.3 | 2.45E-03 | 0.0125 | (B) Pathology_classA-RNA |
| MIR-648/648 | 0.43 | 1.73 | 0.89 | 1.78 | -1.3 | 5.18E-03 | 0.0221 | (B) Pathology_classA-RNA |

| | | | | | | | | |
|---|---|---|---|---|---|---|---|---|
| LAPTM5 | 0.64 | 1.94 | 1.05 | 1.74 | -1.3 | 0.0103 | 0.0365 | (B) Pathology_classA-RNA |
| CLN8 | 0.84 | 2.14 | 1.24 | 0.66 | -1.3 | 0.0133 | 0.0439 | (B) Pathology_classA-RNA |
| CCND1 | 1.09 | 2.39 | 1.21 | 1.25 | -1.3 | 0.0135 | 0.0444 | (B) Pathology_classA-RNA |
| EVI2B | 0.08 | 1.4 | 0.24 | 1.56 | -1.31 | 3.26E-05 | 4.54E-04 | (B) Pathology_classA-RNA |
| SNORA46 | 0.36 | 1.67 | 0.39 | 1.5 | -1.31 | 4.60E-05 | 5.92E-04 | (B) Pathology_classA-RNA |
| LINC00211 | 0.03 | 1.34 | 0.06 | 1.74 | -1.31 | 1.09E-04 | 1.16E-03 | (B) Pathology_classA-RNA |
| TRIM29 | 0.09 | 1.4 | 0.26 | 1.94 | -1.31 | 4.96E-04 | 3.72E-03 | (B) Pathology_classA-RNA |
| ODCP | 0.79 | 2.1 | 0.85 | 1.34 | -1.31 | 1.87E-03 | 0.0102 | (B) Pathology_classA-RNA |
| LINC01010 | 0.32 | 1.63 | 0.79 | 1.64 | -1.31 | 1.97E-03 | 0.0106 | (B) Pathology_classA-RNA |
| TGFBR2 | 5.61 | 6.92 | 0.93 | 0.98 | -1.31 | 2.54E-03 | 0.0129 | (B) Pathology_classA-RNA |
| TMEM71 | 0.27 | 1.59 | 0.44 | 1.8 | -1.32 | 4.05E-04 | 3.18E-03 | (B) Pathology_classA-RNA |
| PRKN | 0.49 | 1.8 | 0.86 | 0.81 | -1.32 | 1.40E-03 | 8.15E-03 | (B) Pathology_classA-RNA |
| MILR1 | 1.67 | 2.99 | 0.87 | 1.39 | -1.32 | 2.28E-03 | 0.0118 | (B) Pathology_classA-RNA |
| FOXN3-AS2 | 0.28 | 1.59 | 0.74 | 1.89 | -1.32 | 2.68E-03 | 0.0134 | (B) Pathology_classA-RNA |
| DDIT4L | 0.39 | 1.71 | 0.45 | 2.44 | -1.32 | 5.13E-03 | 0.0219 | (B) Pathology_classA-RNA |
| CACNA1C-AS | 0.42 | 1.74 | 0.85 | 1.99 | -1.32 | 5.54E-03 | 0.0232 | (B) Pathology_classA-RNA |
| TPI1P4 | 0.2 | 1.53 | 0.42 | 1.61 | -1.33 | 1.06E-04 | 1.14E-03 | (B) Pathology_classA-RNA |
| RNU6-1003P | 0.18 | 1.51 | 0.37 | 1.8 | -1.33 | 2.73E-04 | 2.34E-03 | (B) Pathology_classA-RNA |
| RNU7-172P | 0.13 | 1.45 | 0.38 | 1.81 | -1.33 | 3.07E-04 | 2.57E-03 | (B) Pathology_classA-RNA |
| RNU6-789P | 0.42 | 1.75 | 0.84 | 1.86 | -1.33 | 3.98E-03 | 0.0181 | (B) Pathology_classA-RNA |
| MRVI1 | 3.04 | 4.36 | 0.99 | 1.5 | -1.33 | 5.09E-03 | 0.0218 | (B) Pathology_classA-RNA |
| XAF1 | 2.39 | 3.72 | 1.08 | 1.48 | -1.33 | 7.57E-03 | 0.0291 | (B) Pathology_classA-RNA |
| RN7SL674P | 2.63 | 3.97 | 0.96 | 2.08 | -1.33 | 9.46E-03 | 0.0342 | (B) Pathology_classA-RNA |
| IGHV6-1 | 0.37 | 1.7 | 0.98 | 2.27 | -1.33 | 0.0137 | 0.0449 | (B) Pathology_classA-RNA |
| BCAS2P2 | 0.43 | 1.77 | 0.34 | 0.99 | -1.34 | 9.75E-08 | 4.69E-06 | (B) Pathology_classA-RNA |
| TRAJ21 | 0.01 | 1.34 | 0.02 | 1.19 | -1.34 | 1.94E-07 | 8.17E-06 | (B) Pathology_classA-RNA |
| PTPRVP | 0.54 | 1.88 | 0.54 | 0.99 | -1.34 | 1.50E-05 | 2.49E-04 | (B) Pathology_classA-RNA |
| ATP8A2P2 | 0.25 | 1.59 | 0.36 | 1.48 | -1.34 | 2.17E-05 | 3.26E-04 | (B) Pathology_classA-RNA |
| ZBTB8OSP1 | 0.51 | 1.85 | 0.41 | 1.53 | -1.34 | 4.72E-05 | 6.03E-04 | (B) Pathology_classA-RNA |
| RN7SL254P | 0.09 | 1.42 | 0.26 | 1.64 | -1.34 | 5.23E-05 | 6.56E-04 | (B) Pathology_classA-RNA |
| TRBV6-6 | 0.16 | 1.5 | 0.49 | 1.58 | -1.34 | 1.24E-04 | 1.28E-03 | (B) Pathology_classA-RNA |
| GPR21 | 0.79 | 2.12 | 0.66 | 1.46 | -1.34 | 3.65E-04 | 2.94E-03 | (B) Pathology_classA-RNA |
| MIR-3619/36 | 0.52 | 1.86 | 0.73 | 1.52 | -1.34 | 8.07E-04 | 5.40E-03 | (B) Pathology_classA-RNA |
| MIR-3619/3P | 0.52 | 1.86 | 0.73 | 1.52 | -1.34 | 8.07E-04 | 5.40E-03 | (B) Pathology_classA-RNA |
| MIR-3619/5P | 0.52 | 1.86 | 0.73 | 1.52 | -1.34 | 8.07E-04 | 5.40E-03 | (B) Pathology_classA-RNA |
| NFIB | 0.62 | 1.96 | 0.91 | 0.99 | -1.34 | 1.91E-03 | 0.0103 | (B) Pathology_classA-RNA |
| BIRC3 | 0.98 | 2.32 | 1.06 | 1.04 | -1.34 | 5.23E-03 | 0.0222 | (B) Pathology_classA-RNA |
| GTF2A1L | 1.43 | 2.76 | 1.05 | 1.62 | -1.34 | 7.34E-03 | 0.0285 | (B) Pathology_classA-RNA |
| ACACA | 3.54 | 4.89 | 1.18 | 1.2 | -1.34 | 9.82E-03 | 0.0352 | (B) Pathology_classA-RNA |
| RNU6-198P | 0.08 | 1.44 | 0.16 | 1.56 | -1.35 | 1.73E-05 | 2.77E-04 | (B) Pathology_classA-RNA |
| HMGN2P20 | 0.22 | 1.56 | 0.55 | 1.5 | -1.35 | 1.31E-04 | 1.33E-03 | (B) Pathology_classA-RNA |
| SGMS1 | 2.76 | 4.12 | 0.69 | 0.84 | -1.35 | 1.77E-04 | 1.68E-03 | (B) Pathology_classA-RNA |
| UGT1A1 | 0.11 | 1.47 | 0.34 | 2.05 | -1.35 | 7.45E-04 | 5.07E-03 | (B) Pathology_classA-RNA |
| FCGR2A | 1.47 | 2.82 | 0.92 | 1.3 | -1.35 | 2.33E-03 | 0.012 | (B) Pathology_classA-RNA |
| SYTL2 | 0.37 | 1.72 | 1.11 | 1.72 | -1.35 | 9.98E-03 | 0.0356 | (B) Pathology_classA-RNA |
| RNU6-1100P | 1.98 | 3.33 | 0.96 | 2.42 | -1.35 | 0.0148 | 0.0475 | (B) Pathology_classA-RNA |
| DUSP9 | 0.14 | 1.49 | 0.34 | 1.06 | -1.36 | 1.52E-07 | 6.76E-06 | (B) Pathology_classA-RNA |
| RNU11-2P | 0.04 | 1.4 | 0.12 | 1.6 | -1.36 | 2.15E-05 | 3.24E-04 | (B) Pathology_classA-RNA |
| TRAJ41 | 0.11 | 1.47 | 0.34 | 1.57 | -1.36 | 3.39E-05 | 4.66E-04 | (B) Pathology_classA-RNA |
| IL18R1 | 0.3 | 1.66 | 0.69 | 1.03 | -1.36 | 1.79E-04 | 1.69E-03 | (B) Pathology_classA-RNA |
| NR2F2-AS1 | 1.12 | 2.48 | 0.61 | 1.45 | -1.36 | 1.80E-04 | 1.70E-03 | (B) Pathology_classA-RNA |
| RAD17P2 | 0.04 | 1.41 | 0.08 | 1.51 | -1.37 | 8.27E-06 | 1.57E-04 | (B) Pathology_classA-RNA |
| RNU6-378P | 0.1 | 1.47 | 0.3 | 1.65 | -1.37 | 4.75E-05 | 6.06E-04 | (B) Pathology_classA-RNA |
| TRAJ40 | 0.08 | 1.45 | 0.24 | 1.7 | -1.37 | 5.79E-05 | 7.09E-04 | (B) Pathology_classA-RNA |
| MIR-4725/3P | 0.15 | 1.51 | 0.44 | 1.7 | -1.37 | 1.39E-04 | 1.40E-03 | (B) Pathology_classA-RNA |
| MIR-4725/5P | 0.15 | 1.51 | 0.44 | 1.7 | -1.37 | 1.39E-04 | 1.40E-03 | (B) Pathology_classA-RNA |
| APOBEC3G | 2.32 | 3.69 | 0.53 | 1.7 | -1.37 | 2.47E-04 | 2.16E-03 | (B) Pathology_classA-RNA |
| SCARNA12 | 5.5 | 6.86 | 0.51 | 1.74 | -1.37 | 2.85E-04 | 2.42E-03 | (B) Pathology_classA-RNA |
| MIR-4523/45 | 0.28 | 1.66 | 0.48 | 1.82 | -1.37 | 3.42E-04 | 2.79E-03 | (B) Pathology_classA-RNA |
| MTND4P23 | 0.55 | 1.92 | 0.75 | 1.58 | -1.37 | 8.35E-04 | 5.54E-03 | (B) Pathology_classA-RNA |
| GSTM3P2 | 0.26 | 1.63 | 0.62 | 1.88 | -1.37 | 9.38E-04 | 6.01E-03 | (B) Pathology_classA-RNA |
| GBP4 | 2.52 | 3.89 | 1.03 | 1.3 | -1.37 | 4.15E-03 | 0.0187 | (B) Pathology_classA-RNA |
| TRAV26-2 | 0.29 | 1.66 | 0.88 | 2.07 | -1.37 | 5.64E-03 | 0.0235 | (B) Pathology_classA-RNA |
| ZFP36L1 | 2.42 | 3.79 | 1.13 | 1.15 | -1.37 | 6.93E-03 | 0.0274 | (B) Pathology_classA-RNA |
| P2RY8 | 1.19 | 2.56 | 1.06 | 1.72 | -1.37 | 7.27E-03 | 0.0283 | (B) Pathology_classA-RNA |

| | | | | | | | | |
|---|---|---|---|---|---|---|---|---|
| ARF1P2 | 0.64 | 2.01 | 1.12 | 1.79 | -1.37 | 0.0103 | 0.0365 | (B) Pathology_classA-RNA |
| RPS2P39 | 0.1 | 1.48 | 0.3 | 1.65 | -1.38 | 4.34E-05 | 5.65E-04 | (B) Pathology_classA-RNA |
| ANKRD44-IT1 | 0.15 | 1.53 | 0.36 | 1.72 | -1.38 | 9.21E-05 | 1.02E-03 | (B) Pathology_classA-RNA |
| ITGB2 | 0.18 | 1.56 | 0.37 | 1.88 | -1.38 | 2.65E-04 | 2.28E-03 | (B) Pathology_classA-RNA |
| SNORD13 | 0.22 | 1.6 | 0.49 | 1.78 | -1.38 | 2.69E-04 | 2.31E-03 | (B) Pathology_classA-RNA |
| ALDH7A1 | 5.66 | 7.05 | 0.8 | 1.7 | -1.38 | 1.51E-03 | 8.65E-03 | (B) Pathology_classA-RNA |
| AGGF1P3 | 0.81 | 2.18 | 0.84 | 1.74 | -1.38 | 2.31E-03 | 0.0119 | (B) Pathology_classA-RNA |
| TRBJ2-2 | 0.56 | 1.94 | 1.03 | 1.94 | -1.38 | 8.04E-03 | 0.0304 | (B) Pathology_classA-RNA |
| LRMDA | 1.15 | 2.54 | 1.27 | 0.96 | -1.38 | 0.0114 | 0.0393 | (B) Pathology_classA-RNA |
| ARHGEF38-IT | 0.23 | 1.62 | 0.36 | 1.4 | -1.39 | 6.56E-06 | 1.31E-04 | (B) Pathology_classA-RNA |
| RNU6-255P | 0.25 | 1.64 | 0.42 | 1.87 | -1.39 | 2.83E-04 | 2.40E-03 | (B) Pathology_classA-RNA |
| NSFP1 | 0.39 | 1.78 | 0.78 | 0.95 | -1.39 | 3.92E-04 | 3.11E-03 | (B) Pathology_classA-RNA |
| CELF2-AS2 | 0.3 | 1.69 | 0.5 | 1.98 | -1.39 | 6.68E-04 | 4.67E-03 | (B) Pathology_classA-RNA |
| HIST1H2BL | 0.47 | 1.86 | 0.57 | 1.91 | -1.39 | 7.20E-04 | 4.95E-03 | (B) Pathology_classA-RNA |
| NFE4 | 0.94 | 2.32 | 0.6 | 1.87 | -1.39 | 7.25E-04 | 4.98E-03 | (B) Pathology_classA-RNA |
| HIST1H2BE | 1.73 | 3.12 | 0.84 | 1.88 | -1.39 | 2.83E-03 | 0.0139 | (B) Pathology_classA-RNA |
| FDPSP7 | 1.12 | 2.51 | 1.17 | 1.4 | -1.39 | 8.23E-03 | 0.0309 | (B) Pathology_classA-RNA |
| RNU6-761P | 0.63 | 2.02 | 1.23 | 1.8 | -1.39 | 0.0146 | 0.047 | (B) Pathology_classA-RNA |
| CXCR2P1 | 0.01 | 1.41 | 0.03 | 1.44 | -1.4 | 2.65E-06 | 6.36E-05 | (B) Pathology_classA-RNA |
| CD244 | 0.05 | 1.45 | 0.11 | 1.74 | -1.4 | 4.85E-05 | 6.15E-04 | (B) Pathology_classA-RNA |
| FTH1P24 | 0.32 | 1.72 | 0.62 | 1.63 | -1.4 | 2.63E-04 | 2.27E-03 | (B) Pathology_classA-RNA |
| FOSL2 | 0.54 | 1.95 | 0.8 | 1.11 | -1.4 | 4.83E-04 | 3.64E-03 | (B) Pathology_classA-RNA |
| EPHA6 | 1.12 | 2.52 | 0.8 | 1.58 | -1.4 | 1.07E-03 | 6.62E-03 | (B) Pathology_classA-RNA |
| RNU6-1322P | 0.47 | 1.87 | 0.86 | 2.01 | -1.4 | 3.76E-03 | 0.0173 | (B) Pathology_classA-RNA |
| MAN2A1 | 3.42 | 4.82 | 1.1 | 0.72 | -1.4 | 4.76E-03 | 0.0208 | (B) Pathology_classA-RNA |
| MIR-548O-2/ | 0.97 | 2.37 | 1.03 | 1.92 | -1.4 | 7.02E-03 | 0.0276 | (B) Pathology_classA-RNA |
| MIR-548O/54 | 0.97 | 2.37 | 1.03 | 1.92 | -1.4 | 7.02E-03 | 0.0276 | (B) Pathology_classA-RNA |
| NCOA7-AS1 | 0 | 1.42 | 0.01 | 1.61 | -1.41 | 1.29E-05 | 2.24E-04 | (B) Pathology_classA-RNA |
| MPPED2 | 0.11 | 1.52 | 0.34 | 1.57 | -1.41 | 1.92E-05 | 2.99E-04 | (B) Pathology_classA-RNA |
| TRBV5-6 | 0.08 | 1.49 | 0.16 | 1.69 | -1.41 | 3.04E-05 | 4.27E-04 | (B) Pathology_classA-RNA |
| KY | 0.4 | 1.8 | 0.42 | 1.56 | -1.41 | 3.30E-05 | 4.57E-04 | (B) Pathology_classA-RNA |
| RN7SKP248 | 0.46 | 1.86 | 0.73 | 1.66 | -1.41 | 7.17E-04 | 4.94E-03 | (B) Pathology_classA-RNA |
| RNU4-47P | 2.71 | 4.11 | 0.9 | 1.4 | -1.41 | 1.60E-03 | 9.06E-03 | (B) Pathology_classA-RNA |
| IGHJ3P | 0.42 | 1.83 | 0.7 | 2.04 | -1.41 | 1.83E-03 | 0.01 | (B) Pathology_classA-RNA |
| KCNG1 | 0.29 | 1.71 | 0.56 | 1 | -1.42 | 9.57E-06 | 1.76E-04 | (B) Pathology_classA-RNA |
| CDR1 | 0.11 | 1.53 | 0.22 | 1.72 | -1.42 | 4.04E-05 | 5.34E-04 | (B) Pathology_classA-RNA |
| HMGB1P31 | 2.67 | 4.09 | 0.73 | 1.22 | -1.42 | 2.15E-04 | 1.95E-03 | (B) Pathology_classA-RNA |
| RPL21P123 | 0.63 | 2.05 | 0.72 | 2.02 | -1.42 | 1.84E-03 | 0.01 | (B) Pathology_classA-RNA |
| NEAT1 | 8.47 | 9.89 | 1.03 | 1.16 | -1.42 | 2.95E-03 | 0.0144 | (B) Pathology_classA-RNA |
| COLEC12 | 0.53 | 1.95 | 1.1 | 1.61 | -1.42 | 6.28E-03 | 0.0255 | (B) Pathology_classA-RNA |
| HMGN1P33 | 0.09 | 1.51 | 0.26 | 1.53 | -1.43 | 7.59E-06 | 1.47E-04 | (B) Pathology_classA-RNA |
| BRWD1-IT1 | 0.55 | 1.99 | 0.49 | 2 | -1.43 | 5.14E-04 | 3.81E-03 | (B) Pathology_classA-RNA |
| LDHD | 0.66 | 2.09 | 0.84 | 1.07 | -1.43 | 5.97E-04 | 4.29E-03 | (B) Pathology_classA-RNA |
| B4GALT1 | 2.3 | 3.73 | 1.02 | 0.89 | -1.43 | 2.52E-03 | 0.0128 | (B) Pathology_classA-RNA |
| SNORD111B | 0.32 | 1.75 | 0.96 | 2.25 | -1.43 | 7.65E-03 | 0.0293 | (B) Pathology_classA-RNA |
| PRELID1P3 | 0.58 | 2.01 | 1.18 | 2.09 | -1.43 | 0.0131 | 0.0434 | (B) Pathology_classA-RNA |
| ZNF831 | 0.04 | 1.48 | 0.12 | 1.32 | -1.44 | 3.32E-07 | 1.25E-05 | (B) Pathology_classA-RNA |
| LINC00595 | 0.18 | 1.62 | 0.41 | 1.22 | -1.44 | 1.09E-06 | 3.21E-05 | (B) Pathology_classA-RNA |
| RNU6-998P | 0.02 | 1.46 | 0.06 | 1.59 | -1.44 | 7.43E-06 | 1.44E-04 | (B) Pathology_classA-RNA |
| LINC00310 | 3.32 | 4.76 | 0.57 | 1.12 | -1.44 | 1.43E-05 | 2.42E-04 | (B) Pathology_classA-RNA |
| MIR-3175/31 | 0.02 | 1.46 | 0.05 | 1.66 | -1.44 | 1.51E-05 | 2.51E-04 | (B) Pathology_classA-RNA |
| SLFN11 | 0.61 | 2.05 | 0.64 | 1.17 | -1.44 | 5.32E-05 | 6.64E-04 | (B) Pathology_classA-RNA |
| SRGN | 1.02 | 2.47 | 0.93 | 1.77 | -1.44 | 2.69E-03 | 0.0135 | (B) Pathology_classA-RNA |
| CT45A1 | 0.25 | 1.69 | 0.74 | 2.27 | -1.44 | 3.31E-03 | 0.0158 | (B) Pathology_classA-RNA |
| MIR-619/3P | 0.61 | 2.04 | 0.96 | 1.88 | -1.44 | 4.00E-03 | 0.0181 | (B) Pathology_classA-RNA |
| MIR-619/5P | 0.61 | 2.04 | 0.96 | 1.88 | -1.44 | 4.00E-03 | 0.0181 | (B) Pathology_classA-RNA |
| MIR-619/619 | 0.61 | 2.04 | 0.96 | 1.88 | -1.44 | 4.00E-03 | 0.0181 | (B) Pathology_classA-RNA |
| MIR-4787/3P | 0.26 | 1.7 | 0.77 | 2.35 | -1.44 | 4.56E-03 | 0.0201 | (B) Pathology_classA-RNA |
| MIR-4787/5P | 0.26 | 1.7 | 0.77 | 2.35 | -1.44 | 4.56E-03 | 0.0201 | (B) Pathology_classA-RNA |
| CD177 | 0.5 | 1.95 | 1.22 | 2.12 | -1.44 | 0.0147 | 0.0472 | (B) Pathology_classA-RNA |
| RPS24P16 | 0.66 | 2.12 | 0.51 | 0.77 | -1.45 | 1.73E-06 | 4.61E-05 | (B) Pathology_classA-RNA |
| RNA5SP88 | 0.06 | 1.51 | 0.17 | 1.67 | -1.45 | 1.66E-05 | 2.68E-04 | (B) Pathology_classA-RNA |
| RNU6-659P | 0.18 | 1.63 | 0.37 | 1.88 | -1.45 | 1.56E-04 | 1.52E-03 | (B) Pathology_classA-RNA |
| CYP1B1-AS1 | 1.15 | 2.6 | 0.77 | 1.27 | -1.45 | 3.15E-04 | 2.62E-03 | (B) Pathology_classA-RNA |
| RNU6-979P | 0.23 | 1.69 | 0.7 | 1.85 | -1.45 | 7.62E-04 | 5.17E-03 | (B) Pathology_classA-RNA |

| | | | | | | | | |
|---|---|---|---|---|---|---|---|---|
| GLI3 | 0.47 | 1.92 | 0.96 | 1.49 | -1.45 | 2.08E-03 | 0.0111 | (B) Pathology_classA-RNA |
| NCBP2L | 0.73 | 2.19 | 0.91 | 1.71 | -1.45 | 2.09E-03 | 0.0111 | (B) Pathology_classA-RNA |
| RN7SL330P | 0.59 | 2.04 | 1.02 | 1.99 | -1.45 | 5.78E-03 | 0.0239 | (B) Pathology_classA-RNA |
| RNA5SP278 | 0.58 | 2.03 | 1.26 | 1.79 | -1.45 | 0.0124 | 0.0417 | (B) Pathology_classA-RNA |
| PRF1 | 2.53 | 3.98 | 1.21 | 2.04 | -1.45 | 0.0128 | 0.0426 | (B) Pathology_classA-RNA |
| RBMS3-AS1 | 0.06 | 1.52 | 0.14 | 1.77 | -1.46 | 3.62E-05 | 4.91E-04 | (B) Pathology_classA-RNA |
| MIR-4729/47 | 0.23 | 1.68 | 0.31 | 1.84 | -1.46 | 8.60E-05 | 9.69E-04 | (B) Pathology_classA-RNA |
| TAOK3 | 1.48 | 2.93 | 0.78 | 0.96 | -1.46 | 2.69E-04 | 2.31E-03 | (B) Pathology_classA-RNA |
| RNU6-848P | 0.42 | 1.88 | 0.53 | 2 | -1.46 | 4.97E-04 | 3.72E-03 | (B) Pathology_classA-RNA |
| ARL6IP5 | 0.78 | 2.24 | 0.86 | 1.43 | -1.46 | 8.73E-04 | 5.70E-03 | (B) Pathology_classA-RNA |
| LIPN | 0.36 | 1.82 | 0.69 | 1.94 | -1.46 | 9.00E-04 | 5.84E-03 | (B) Pathology_classA-RNA |
| DIO3OS | 1.35 | 2.81 | 1.3 | 1.97 | -1.46 | 0.0155 | 0.0491 | (B) Pathology_classA-RNA |
| SLC9A9-AS2 | 0.27 | 1.74 | 0.37 | 1.26 | -1.47 | 5.97E-07 | 1.98E-05 | (B) Pathology_classA-RNA |
| NKD1 | 0.14 | 1.61 | 0.22 | 1.44 | -1.47 | 1.45E-06 | 4.00E-05 | (B) Pathology_classA-RNA |
| TRAJ44 | 0.13 | 1.6 | 0.33 | 1.43 | -1.47 | 2.40E-06 | 5.92E-05 | (B) Pathology_classA-RNA |
| PBOV1 | 0.49 | 1.97 | 0.37 | 1.71 | -1.47 | 3.73E-05 | 5.03E-04 | (B) Pathology_classA-RNA |
| DOCK9-AS1 | 0.21 | 1.68 | 0.38 | 1.75 | -1.47 | 4.99E-05 | 6.31E-04 | (B) Pathology_classA-RNA |
| HIST1H2APS3 | 0.53 | 2 | 0.71 | 1.88 | -1.47 | 7.46E-04 | 5.08E-03 | (B) Pathology_classA-RNA |
| FCER1A | 0.26 | 1.73 | 0.79 | 1.79 | -1.47 | 1.02E-03 | 6.41E-03 | (B) Pathology_classA-RNA |
| HIST1H2BG | 2.67 | 4.15 | 0.89 | 1.56 | -1.47 | 1.23E-03 | 7.38E-03 | (B) Pathology_classA-RNA |
| LPAR1 | 3.05 | 4.52 | 1.01 | 1.36 | -1.47 | 2.31E-03 | 0.0119 | (B) Pathology_classA-RNA |
| SNORA19 | 1.55 | 3.01 | 1.14 | 1.82 | -1.47 | 7.27E-03 | 0.0283 | (B) Pathology_classA-RNA |
| NCOA4 | 4.42 | 5.91 | 0.68 | 1.2 | -1.48 | 7.14E-05 | 8.34E-04 | (B) Pathology_classA-RNA |
| SNORA12 | 0.48 | 1.96 | 0.41 | 1.89 | -1.48 | 1.40E-04 | 1.40E-03 | (B) Pathology_classA-RNA |
| SPATC1 | 0.74 | 2.22 | 0.82 | 1 | -1.48 | 3.46E-04 | 2.81E-03 | (B) Pathology_classA-RNA |
| RNU6-490P | 0.24 | 1.72 | 0.51 | 2.08 | -1.48 | 5.75E-04 | 4.17E-03 | (B) Pathology_classA-RNA |
| ZCCHC24 | 3.65 | 5.14 | 0.98 | 1.12 | -1.48 | 1.50E-03 | 8.61E-03 | (B) Pathology_classA-RNA |
| IGHJ2P | 0.28 | 1.76 | 0.7 | 2.28 | -1.48 | 2.41E-03 | 0.0124 | (B) Pathology_classA-RNA |
| CYP4X1 | 2.38 | 3.86 | 1.08 | 2.01 | -1.48 | 6.85E-03 | 0.0271 | (B) Pathology_classA-RNA |
| JAK3 | 4.36 | 5.84 | 1.31 | 1.54 | -1.48 | 0.0113 | 0.0388 | (B) Pathology_classA-RNA |
| RNU6-1026P | 0.11 | 1.6 | 0.25 | 1.84 | -1.49 | 4.98E-05 | 6.30E-04 | (B) Pathology_classA-RNA |
| SNORA74B | 0.23 | 1.71 | 0.27 | 1.82 | -1.49 | 5.10E-05 | 6.42E-04 | (B) Pathology_classA-RNA |
| RASSF3 | 0.7 | 2.18 | 0.94 | 1.59 | -1.49 | 1.75E-03 | 9.67E-03 | (B) Pathology_classA-RNA |
| MIR-4435-1/ | 0.53 | 2.02 | 0.92 | 1.95 | -1.49 | 2.62E-03 | 0.0132 | (B) Pathology_classA-RNA |
| TWIST2 | 1.08 | 2.57 | 1.1 | 1.93 | -1.49 | 6.14E-03 | 0.025 | (B) Pathology_classA-RNA |
| HLA-DOB | 0.21 | 1.7 | 0.26 | 1.6 | -1.5 | 6.69E-06 | 1.33E-04 | (B) Pathology_classA-RNA |
| SLC9A9-AS1 | 0.48 | 1.98 | 0.6 | 1.92 | -1.5 | 3.67E-04 | 2.95E-03 | (B) Pathology_classA-RNA |
| BANF1P1 | 0.31 | 1.81 | 0.69 | 1.78 | -1.5 | 4.04E-04 | 3.18E-03 | (B) Pathology_classA-RNA |
| RN7SL577P | 0.43 | 1.93 | 0.88 | 1.88 | -1.5 | 1.81E-03 | 9.93E-03 | (B) Pathology_classA-RNA |
| TRAJ42 | 0.41 | 1.9 | 0.86 | 1.92 | -1.5 | 1.84E-03 | 0.0101 | (B) Pathology_classA-RNA |
| RPL36AP33 | 0.92 | 2.42 | 0.84 | 2.01 | -1.5 | 1.91E-03 | 0.0103 | (B) Pathology_classA-RNA |
| RHOU | 0.42 | 1.92 | 0.97 | 1.65 | -1.5 | 2.12E-03 | 0.0112 | (B) Pathology_classA-RNA |
| MKX | 0.45 | 1.94 | 1.01 | 1.81 | -1.5 | 3.30E-03 | 0.0157 | (B) Pathology_classA-RNA |
| SRGAP3-AS3 | 0.09 | 1.59 | 0.24 | 1.78 | -1.51 | 2.76E-05 | 3.94E-04 | (B) Pathology_classA-RNA |
| QRSL1P3 | 0.36 | 1.87 | 0.45 | 1.79 | -1.51 | 7.16E-05 | 8.36E-04 | (B) Pathology_classA-RNA |
| RSPO1 | 0.59 | 2.1 | 0.62 | 2.03 | -1.51 | 5.84E-04 | 4.22E-03 | (B) Pathology_classA-RNA |
| TNXA | 0.87 | 2.38 | 0.75 | 1.83 | -1.51 | 6.30E-04 | 4.46E-03 | (B) Pathology_classA-RNA |
| RNU7-79P | 0.36 | 1.87 | 0.67 | 2.13 | -1.51 | 1.08E-03 | 6.71E-03 | (B) Pathology_classA-RNA |
| TEAD2 | 1.99 | 3.5 | 0.97 | 1.24 | -1.51 | 1.36E-03 | 8.00E-03 | (B) Pathology_classA-RNA |
| TRPM4 | 0.52 | 2.02 | 1.07 | 1.43 | -1.51 | 2.94E-03 | 0.0144 | (B) Pathology_classA-RNA |
| ITCH-IT1 | 0.18 | 1.7 | 0.38 | 1.69 | -1.52 | 2.03E-05 | 3.12E-04 | (B) Pathology_classA-RNA |
| LAIR1 | 2.07 | 3.59 | 0.83 | 1.44 | -1.52 | 4.73E-04 | 3.58E-03 | (B) Pathology_classA-RNA |
| AZGP1P2 | 0.42 | 1.94 | 0.94 | 1.91 | -1.52 | 2.29E-03 | 0.0118 | (B) Pathology_classA-RNA |
| MARCKSL1P1 | 1.53 | 3.05 | 1.1 | 1.57 | -1.52 | 3.74E-03 | 0.0172 | (B) Pathology_classA-RNA |
| CARMN | 1.58 | 3.1 | 1.25 | 2.4 | -1.52 | 0.0157 | 0.0498 | (B) Pathology_classA-RNA |
| ITGAD | 0.11 | 1.63 | 0.26 | 1.35 | -1.53 | 2.90E-07 | 1.12E-05 | (B) Pathology_classA-RNA |
| SPATA13-AS1 | 0.02 | 1.54 | 0.05 | 1.76 | -1.53 | 1.50E-05 | 2.49E-04 | (B) Pathology_classA-RNA |
| RNU6-402P | 0.91 | 2.44 | 1 | 2.29 | -1.53 | 5.75E-03 | 0.0239 | (B) Pathology_classA-RNA |
| CCL8 | 0.89 | 2.43 | 1.23 | 1.85 | -1.53 | 8.22E-03 | 0.0309 | (B) Pathology_classA-RNA |
| KBTBD11 | 0.9 | 2.42 | 1.37 | 1.65 | -1.53 | 0.0127 | 0.0423 | (B) Pathology_classA-RNA |
| TRAV6 | 0.21 | 1.76 | 0.37 | 1.56 | -1.54 | 5.20E-06 | 1.09E-04 | (B) Pathology_classA-RNA |
| RNU6-977P | 0.11 | 1.65 | 0.34 | 1.93 | -1.54 | 7.84E-05 | 8.96E-04 | (B) Pathology_classA-RNA |
| RNU6-1045P | 0.24 | 1.78 | 0.36 | 2.08 | -1.54 | 2.12E-04 | 1.92E-03 | (B) Pathology_classA-RNA |
| NLRP3 | 1.48 | 3.03 | 0.86 | 1.34 | -1.54 | 4.74E-04 | 3.59E-03 | (B) Pathology_classA-RNA |
| SNORD56B | 0.23 | 1.77 | 0.69 | 2.02 | -1.54 | 6.91E-04 | 4.80E-03 | (B) Pathology_classA-RNA |

| | | | | | | | | |
|---|---|---|---|---|---|---|---|---|
| INPP5D | 2.98 | 4.51 | 1.12 | 1.56 | -1.54 | 3.82E-03 | 0.0175 | (B) Pathology_classA-RNA |
| MRPL53P1 | 0.69 | 2.23 | 1.15 | 1.55 | -1.54 | 4.25E-03 | 0.0191 | (B) Pathology_classA-RNA |
| DAB2 | 4.39 | 5.93 | 1.28 | 1.01 | -1.54 | 6.84E-03 | 0.0271 | (B) Pathology_classA-RNA |
| DUXAP8 | 4.52 | 6.06 | 1.29 | 1.71 | -1.54 | 9.25E-03 | 0.0337 | (B) Pathology_classA-RNA |
| TRAV4 | 0.22 | 1.77 | 0.43 | 1.51 | -1.55 | 5.26E-06 | 1.10E-04 | (B) Pathology_classA-RNA |
| MIR-224/224 | 0.08 | 1.63 | 0.23 | 2.06 | -1.55 | 1.29E-04 | 1.32E-03 | (B) Pathology_classA-RNA |
| MIR-224/224 | 0.08 | 1.63 | 0.23 | 2.06 | -1.55 | 1.29E-04 | 1.32E-03 | (B) Pathology_classA-RNA |
| MIR-224/3P | 0.08 | 1.63 | 0.23 | 2.06 | -1.55 | 1.29E-04 | 1.32E-03 | (B) Pathology_classA-RNA |
| MIR-224/5P | 0.08 | 1.63 | 0.23 | 2.06 | -1.55 | 1.29E-04 | 1.32E-03 | (B) Pathology_classA-RNA |
| RNU7-96P | 0.19 | 1.74 | 0.43 | 2.07 | -1.55 | 2.28E-04 | 2.03E-03 | (B) Pathology_classA-RNA |
| DUXAP10 | 3.47 | 5.02 | 0.86 | 1.75 | -1.55 | 8.64E-04 | 5.66E-03 | (B) Pathology_classA-RNA |
| TBC1D8B | 0.64 | 2.2 | 1.01 | 1.17 | -1.55 | 1.42E-03 | 8.23E-03 | (B) Pathology_classA-RNA |
| RN7SL683P | 0.3 | 1.86 | 0.91 | 1.98 | -1.55 | 1.92E-03 | 0.0104 | (B) Pathology_classA-RNA |
| PAICSP5 | 0.51 | 2.07 | 0.82 | 2.29 | -1.55 | 2.46E-03 | 0.0126 | (B) Pathology_classA-RNA |
| SUSD3 | 1.5 | 3.05 | 1.02 | 1.96 | -1.55 | 3.33E-03 | 0.0158 | (B) Pathology_classA-RNA |
| IGKV2D-29 | 0.33 | 1.88 | 0.65 | 2.81 | -1.55 | 5.49E-03 | 0.023 | (B) Pathology_classA-RNA |
| PPEF1 | 1.47 | 3.02 | 1.23 | 1.73 | -1.55 | 6.83E-03 | 0.0271 | (B) Pathology_classA-RNA |
| TRAJ37 | 0.04 | 1.6 | 0.13 | 1.94 | -1.56 | 4.87E-05 | 6.18E-04 | (B) Pathology_classA-RNA |
| RC3H1-IT1 | 0.34 | 1.9 | 0.54 | 1.8 | -1.56 | 8.61E-05 | 9.70E-04 | (B) Pathology_classA-RNA |
| RNA5SP321 | 0.02 | 1.58 | 0.07 | 2.04 | -1.56 | 9.00E-05 | 1.00E-03 | (B) Pathology_classA-RNA |
| RNU7-38P | 0.17 | 1.73 | 0.5 | 2.05 | -1.56 | 2.54E-04 | 2.21E-03 | (B) Pathology_classA-RNA |
| PLXDC2 | 2.13 | 3.69 | 0.99 | 0.91 | -1.56 | 1.09E-03 | 6.74E-03 | (B) Pathology_classA-RNA |
| SLC35A3 | 1.41 | 2.98 | 1.02 | 0.97 | -1.56 | 1.34E-03 | 7.90E-03 | (B) Pathology_classA-RNA |
| SNRPGP9 | 0.59 | 2.15 | 0.9 | 2.09 | -1.56 | 2.26E-03 | 0.0117 | (B) Pathology_classA-RNA |
| ZBTB20-AS1 | 0.11 | 1.68 | 0.27 | 1.4 | -1.57 | 3.30E-07 | 1.24E-05 | (B) Pathology_classA-RNA |
| EEF1A1P32 | 0.27 | 1.84 | 0.32 | 1.88 | -1.57 | 4.39E-05 | 5.71E-04 | (B) Pathology_classA-RNA |
| HIST1H1PS1 | 4.54 | 6.12 | 0.71 | 1.18 | -1.57 | 5.41E-05 | 6.72E-04 | (B) Pathology_classA-RNA |
| NUDT19P5 | 0.23 | 1.79 | 0.67 | 1.51 | -1.57 | 7.52E-05 | 8.70E-04 | (B) Pathology_classA-RNA |
| MIR-644/644 | 0.3 | 1.86 | 0.52 | 1.86 | -1.57 | 9.84E-05 | 1.07E-03 | (B) Pathology_classA-RNA |
| MIR-644A/64 | 0.3 | 1.86 | 0.52 | 1.86 | -1.57 | 9.84E-05 | 1.07E-03 | (B) Pathology_classA-RNA |
| MIR-644A/64 | 0.3 | 1.86 | 0.52 | 1.86 | -1.57 | 9.84E-05 | 1.07E-03 | (B) Pathology_classA-RNA |
| RN7SKP9 | 0.3 | 1.87 | 0.73 | 2.01 | -1.57 | 6.64E-04 | 4.65E-03 | (B) Pathology_classA-RNA |
| SLC4A7 | 0.92 | 2.49 | 0.94 | 0.87 | -1.57 | 7.53E-04 | 5.12E-03 | (B) Pathology_classA-RNA |
| RXRA | 4.58 | 6.15 | 1.09 | 0.79 | -1.57 | 2.16E-03 | 0.0113 | (B) Pathology_classA-RNA |
| ALOX15B | 1.31 | 2.88 | 1.03 | 2.79 | -1.57 | 0.0115 | 0.0395 | (B) Pathology_classA-RNA |
| TRAV29DV5 | 0.05 | 1.62 | 0.14 | 1.61 | -1.58 | 2.39E-06 | 5.91E-05 | (B) Pathology_classA-RNA |
| CXCL12 | 0.36 | 1.94 | 0.5 | 1.6 | -1.58 | 1.61E-05 | 2.63E-04 | (B) Pathology_classA-RNA |
| TRAV26-1 | 0.09 | 1.67 | 0.22 | 1.84 | -1.58 | 2.10E-05 | 3.18E-04 | (B) Pathology_classA-RNA |
| HIST2H2BE | 2.98 | 4.56 | 0.64 | 1.42 | -1.58 | 3.50E-05 | 4.78E-04 | (B) Pathology_classA-RNA |
| MIR-4459/44 | 0.12 | 1.7 | 0.35 | 1.9 | -1.58 | 4.96E-05 | 6.27E-04 | (B) Pathology_classA-RNA |
| CLMP | 2.1 | 3.68 | 0.83 | 1.43 | -1.58 | 3.16E-04 | 2.63E-03 | (B) Pathology_classA-RNA |
| RNU6-915P | 0.55 | 2.13 | 0.87 | 1.7 | -1.58 | 7.22E-04 | 4.96E-03 | (B) Pathology_classA-RNA |
| SLC4A4 | 0.87 | 2.46 | 1.13 | 2.58 | -1.58 | 0.0104 | 0.0367 | (B) Pathology_classA-RNA |
| PRRG1 | 0.23 | 1.82 | 0.48 | 1.17 | -1.59 | 5.07E-07 | 1.74E-05 | (B) Pathology_classA-RNA |
| NBPF10 | 3.08 | 4.67 | 0.68 | 0.53 | -1.59 | 5.13E-05 | 6.45E-04 | (B) Pathology_classA-RNA |
| STX18-IT1 | 0.32 | 1.92 | 0.54 | 1.83 | -1.59 | 7.66E-05 | 8.82E-04 | (B) Pathology_classA-RNA |
| ABCA13 | 0.05 | 1.64 | 0.15 | 2.17 | -1.59 | 1.64E-04 | 1.58E-03 | (B) Pathology_classA-RNA |
| TBL1XR1-AS1 | 0.47 | 2.06 | 0.59 | 1.99 | -1.59 | 2.42E-04 | 2.13E-03 | (B) Pathology_classA-RNA |
| HMGN2P38 | 0.32 | 1.91 | 0.77 | 1.69 | -1.59 | 2.89E-04 | 2.45E-03 | (B) Pathology_classA-RNA |
| MIR-4774/3P | 0.35 | 1.93 | 0.71 | 2.16 | -1.59 | 8.55E-04 | 5.62E-03 | (B) Pathology_classA-RNA |
| MIR-4774/5P | 0.35 | 1.93 | 0.71 | 2.16 | -1.59 | 8.55E-04 | 5.62E-03 | (B) Pathology_classA-RNA |
| MIR-4789/3P | 0.33 | 1.92 | 0.99 | 2.43 | -1.59 | 5.19E-03 | 0.0221 | (B) Pathology_classA-RNA |
| MIR-4789/5P | 0.33 | 1.92 | 0.99 | 2.43 | -1.59 | 5.19E-03 | 0.0221 | (B) Pathology_classA-RNA |
| CYP7B1 | 1.95 | 3.55 | 1.27 | 1.37 | -1.59 | 5.76E-03 | 0.0239 | (B) Pathology_classA-RNA |
| RN7SKP158 | 0.06 | 1.65 | 0.12 | 1.72 | -1.6 | 5.59E-06 | 1.15E-04 | (B) Pathology_classA-RNA |
| RNU6-146P | 3.23 | 4.82 | 1.03 | 0.82 | -1.6 | 1.32E-03 | 7.81E-03 | (B) Pathology_classA-RNA |
| TRBV7-3 | 0.81 | 2.41 | 0.73 | 2.34 | -1.6 | 1.51E-03 | 8.67E-03 | (B) Pathology_classA-RNA |
| HLA-DQA2 | 0.69 | 2.29 | 0.99 | 2.4 | -1.6 | 4.77E-03 | 0.0208 | (B) Pathology_classA-RNA |
| DMXL1 | 0.02 | 1.62 | 0.05 | 1.36 | -1.61 | 7.58E-08 | 3.87E-06 | (B) Pathology_classA-RNA |
| MRVI1-AS1 | 0.96 | 2.57 | 0.68 | 1.07 | -1.61 | 2.09E-05 | 3.17E-04 | (B) Pathology_classA-RNA |
| JCHAIN | 0.03 | 1.64 | 0.09 | 2.13 | -1.61 | 1.05E-04 | 1.13E-03 | (B) Pathology_classA-RNA |
| CLIC2 | 2.08 | 3.69 | 0.99 | 1.46 | -1.61 | 1.08E-03 | 6.68E-03 | (B) Pathology_classA-RNA |
| MIR-3610/36 | 2.8 | 4.41 | 0.9 | 2.09 | -1.61 | 1.68E-03 | 9.39E-03 | (B) Pathology_classA-RNA |
| RN7SL328P | 0.66 | 2.27 | 1.24 | 2.04 | -1.61 | 7.38E-03 | 0.0287 | (B) Pathology_classA-RNA |
| STAT4 | 1.04 | 2.66 | 0.67 | 1.34 | -1.62 | 2.48E-05 | 3.61E-04 | (B) Pathology_classA-RNA |

| | | | | | | | | |
|---|---|---|---|---|---|---|---|---|
| RNU6-1282P | 0.23 | 1.85 | 0.57 | 2.01 | -1.62 | 1.85E-04 | 1.73E-03 | (B) Pathology_classA-RNA |
| NHS-AS1 | 0.32 | 1.94 | 0.49 | 2.18 | -1.62 | 2.77E-04 | 2.36E-03 | (B) Pathology_classA-RNA |
| HIST2H3PS2 | 0.92 | 2.54 | 0.79 | 1.71 | -1.62 | 3.07E-04 | 2.57E-03 | (B) Pathology_classA-RNA |
| ST8SIA6 | 0.22 | 1.85 | 0.39 | 1.32 | -1.63 | 2.21E-07 | 9.04E-06 | (B) Pathology_classA-RNA |
| TRAV2 | 0.21 | 1.84 | 0.48 | 1.6 | -1.63 | 7.86E-06 | 1.51E-04 | (B) Pathology_classA-RNA |
| SH3BGRL | 5.24 | 6.86 | 1.29 | 1.55 | -1.63 | 5.86E-03 | 0.0242 | (B) Pathology_classA-RNA |
| ARHGAP18 | 0.38 | 2.02 | 0.61 | 0.73 | -1.64 | 5.57E-06 | 1.14E-04 | (B) Pathology_classA-RNA |
| RNU6-1305P | 0.1 | 1.74 | 0.23 | 1.98 | -1.64 | 3.67E-05 | 4.96E-04 | (B) Pathology_classA-RNA |
| TRBV11-2 | 0.14 | 1.78 | 0.23 | 2 | -1.64 | 4.09E-05 | 5.40E-04 | (B) Pathology_classA-RNA |
| RPL31P20 | 0.09 | 1.73 | 0.25 | 2.08 | -1.64 | 7.04E-05 | 8.26E-04 | (B) Pathology_classA-RNA |
| CHAD | 0.65 | 2.28 | 0.85 | 1.62 | -1.64 | 3.45E-04 | 2.81E-03 | (B) Pathology_classA-RNA |
| ADGRE4P | 1.36 | 3 | 0.99 | 1.19 | -1.64 | 7.42E-04 | 5.06E-03 | (B) Pathology_classA-RNA |
| RPS16P5 | 0.9 | 2.54 | 1.11 | 2.29 | -1.64 | 5.16E-03 | 0.022 | (B) Pathology_classA-RNA |
| CLEC4E | 0.49 | 2.13 | 1.24 | 1.97 | -1.64 | 5.96E-03 | 0.0245 | (B) Pathology_classA-RNA |
| BCAS1 | 0.45 | 2.09 | 1.35 | 1.62 | -1.64 | 7.44E-03 | 0.0288 | (B) Pathology_classA-RNA |
| C19ORF33 | 3.32 | 4.96 | 1.48 | 1.48 | -1.64 | 0.0112 | 0.0386 | (B) Pathology_classA-RNA |
| TAS2R63P | 0.22 | 1.86 | 0.44 | 1.47 | -1.65 | 1.58E-06 | 4.29E-05 | (B) Pathology_classA-RNA |
| TMEFF2 | 0.05 | 1.7 | 0.16 | 2.24 | -1.65 | 1.54E-04 | 1.51E-03 | (B) Pathology_classA-RNA |
| FOLR3 | 0.01 | 1.66 | 0.03 | 2.72 | -1.65 | 1.21E-03 | 7.29E-03 | (B) Pathology_classA-RNA |
| MT1XP1 | 5.67 | 7.32 | 1.03 | 1.98 | -1.65 | 2.14E-03 | 0.0113 | (B) Pathology_classA-RNA |
| EID3 | 2.6 | 4.25 | 0.7 | 1.1 | -1.66 | 2.07E-05 | 3.16E-04 | (B) Pathology_classA-RNA |
| TRDJ1 | 0.24 | 1.9 | 0.53 | 2.04 | -1.66 | 1.26E-04 | 1.29E-03 | (B) Pathology_classA-RNA |
| WAS | 3.49 | 5.15 | 0.69 | 1.91 | -1.66 | 1.94E-04 | 1.80E-03 | (B) Pathology_classA-RNA |
| RNA5SP282 | 0.55 | 2.21 | 0.68 | 1.97 | -1.66 | 2.22E-04 | 1.99E-03 | (B) Pathology_classA-RNA |
| LINC01547 | 0.71 | 2.37 | 0.94 | 1.18 | -1.66 | 4.24E-04 | 3.30E-03 | (B) Pathology_classA-RNA |
| HIST1H3J | 4.21 | 5.87 | 0.76 | 2.13 | -1.66 | 6.02E-04 | 4.32E-03 | (B) Pathology_classA-RNA |
| TRBJ2-2P | 0.61 | 2.27 | 0.92 | 2.13 | -1.66 | 1.48E-03 | 8.54E-03 | (B) Pathology_classA-RNA |
| CARD16 | 3.76 | 5.43 | 0.9 | 1.88 | -1.67 | 7.20E-04 | 4.95E-03 | (B) Pathology_classA-RNA |
| HLA-DPA1 | 5.24 | 6.91 | 1.15 | 2.09 | -1.67 | 4.21E-03 | 0.0189 | (B) Pathology_classA-RNA |
| HBM | 0.26 | 1.93 | 0.77 | 3.46 | -1.67 | 0.0132 | 0.0437 | (B) Pathology_classA-RNA |
| CYP4F29P | 0.91 | 2.59 | 0.83 | 1.62 | -1.68 | 2.22E-04 | 1.99E-03 | (B) Pathology_classA-RNA |
| TRBJ2-4 | 0.44 | 2.12 | 0.64 | 2.07 | -1.68 | 2.27E-04 | 2.02E-03 | (B) Pathology_classA-RNA |
| ARHGAP26-A | 0.56 | 2.23 | 0.5 | 2.22 | -1.68 | 2.39E-04 | 2.11E-03 | (B) Pathology_classA-RNA |
| SHISA9 | 0.17 | 1.85 | 0.35 | 2.43 | -1.68 | 4.25E-04 | 3.30E-03 | (B) Pathology_classA-RNA |
| SNORA7B | 3.42 | 5.1 | 1.27 | 2.5 | -1.68 | 9.95E-03 | 0.0356 | (B) Pathology_classA-RNA |
| HLA-DQB1-A | 5.01 | 6.69 | 1.47 | 1.84 | -1.68 | 0.0111 | 0.0383 | (B) Pathology_classA-RNA |
| EGOT | 0.18 | 1.87 | 0.32 | 1.63 | -1.69 | 1.49E-06 | 4.10E-05 | (B) Pathology_classA-RNA |
| ZNF90P1 | 0.6 | 2.29 | 0.57 | 1.96 | -1.69 | 8.38E-05 | 9.47E-04 | (B) Pathology_classA-RNA |
| RNY4P4 | 0.26 | 1.95 | 0.52 | 2.07 | -1.69 | 1.12E-04 | 1.18E-03 | (B) Pathology_classA-RNA |
| RAD51B | 0.86 | 2.56 | 0.54 | 0.9 | -1.7 | 3.74E-07 | 1.36E-05 | (B) Pathology_classA-RNA |
| ZBTB20-AS2 | 0.08 | 1.77 | 0.23 | 1.74 | -1.7 | 2.85E-06 | 6.74E-05 | (B) Pathology_classA-RNA |
| RNU6-331P | 0.23 | 1.93 | 0.69 | 2.11 | -1.7 | 2.98E-04 | 2.51E-03 | (B) Pathology_classA-RNA |
| ATG12P2 | 0.75 | 2.45 | 0.62 | 2.3 | -1.7 | 4.50E-04 | 3.45E-03 | (B) Pathology_classA-RNA |
| NCF1C | 3.77 | 5.47 | 1.08 | 2.55 | -1.7 | 5.14E-03 | 0.022 | (B) Pathology_classA-RNA |
| HIST1H2BH | 0.58 | 2.28 | 1.17 | 2.33 | -1.7 | 5.29E-03 | 0.0224 | (B) Pathology_classA-RNA |
| RN7SL398P | 0.34 | 2.05 | 0.51 | 2.06 | -1.71 | 8.20E-05 | 9.30E-04 | (B) Pathology_classA-RNA |
| MIR-4265/42 | 0.2 | 1.91 | 0.46 | 2.23 | -1.71 | 1.67E-04 | 1.61E-03 | (B) Pathology_classA-RNA |
| RNU6-886P | 0.41 | 2.12 | 0.66 | 2.18 | -1.71 | 3.19E-04 | 2.64E-03 | (B) Pathology_classA-RNA |
| MIR-4296/42 | 0.57 | 2.29 | 0.86 | 1.91 | -1.71 | 4.62E-04 | 3.52E-03 | (B) Pathology_classA-RNA |
| BTBD9-AS1 | 0.58 | 2.29 | 0.88 | 1.94 | -1.71 | 6.03E-04 | 4.32E-03 | (B) Pathology_classA-RNA |
| SPTLC3 | 2.47 | 4.18 | 1.04 | 1.37 | -1.71 | 8.27E-04 | 5.49E-03 | (B) Pathology_classA-RNA |
| WDR74 | 0.7 | 2.41 | 1.02 | 1.61 | -1.71 | 8.86E-04 | 5.77E-03 | (B) Pathology_classA-RNA |
| RASGRP2 | 5.35 | 7.06 | 1.06 | 2.15 | -1.71 | 2.30E-03 | 0.0119 | (B) Pathology_classA-RNA |
| CBX3P3 | 0.08 | 1.8 | 0.18 | 1.63 | -1.72 | 7.15E-07 | 2.27E-05 | (B) Pathology_classA-RNA |
| ARHGAP9 | 0.67 | 2.38 | 0.54 | 2.01 | -1.72 | 7.53E-05 | 8.71E-04 | (B) Pathology_classA-RNA |
| VSIR | 4.69 | 6.4 | 1.12 | 1.78 | -1.72 | 1.91E-03 | 0.0103 | (B) Pathology_classA-RNA |
| TRAV38-1 | 0.11 | 1.85 | 0.34 | 1.93 | -1.73 | 1.44E-05 | 2.43E-04 | (B) Pathology_classA-RNA |
| TRAJ31 | 0.14 | 1.86 | 0.41 | 1.98 | -1.73 | 2.88E-05 | 4.09E-04 | (B) Pathology_classA-RNA |
| CCND2 | 2.02 | 3.75 | 0.82 | 1.51 | -1.73 | 1.17E-04 | 1.23E-03 | (B) Pathology_classA-RNA |
| TRAJ23 | 0.37 | 2.1 | 0.71 | 2.34 | -1.73 | 6.36E-04 | 4.50E-03 | (B) Pathology_classA-RNA |
| FCGR2C | 0.82 | 2.54 | 1.06 | 1.27 | -1.73 | 8.40E-04 | 5.56E-03 | (B) Pathology_classA-RNA |
| MIR-4803/48 | 0.16 | 1.89 | 0.47 | 2.65 | -1.73 | 8.92E-04 | 5.80E-03 | (B) Pathology_classA-RNA |
| NUDT4 | 2.27 | 3.99 | 1.15 | 1.29 | -1.73 | 1.67E-03 | 9.33E-03 | (B) Pathology_classA-RNA |
| IGHD | 0.71 | 2.43 | 1.04 | 2.31 | -1.73 | 2.63E-03 | 0.0132 | (B) Pathology_classA-RNA |
| RPL7AP64 | 0.59 | 2.33 | 0.63 | 1.64 | -1.74 | 1.76E-05 | 2.81E-04 | (B) Pathology_classA-RNA |

| | | | | | | | | |
|---|---|---|---|---|---|---|---|---|
| RUFY4 | 0.76 | 2.5 | 0.87 | 1.86 | -1.74 | 3.78E-04 | 3.02E-03 | (B) Pathology_classA-RNA |
| MMP25 | 6.3 | 8.05 | 0.98 | 1.84 | -1.74 | 7.55E-04 | 5.13E-03 | (B) Pathology_classA-RNA |
| TRIM5 | 3.4 | 5.14 | 1.25 | 1.2 | -1.74 | 2.71E-03 | 0.0136 | (B) Pathology_classA-RNA |
| CD5 | 0.88 | 2.62 | 1.31 | 1.76 | -1.74 | 4.54E-03 | 0.02 | (B) Pathology_classA-RNA |
| TRAV19 | 0.1 | 1.85 | 0.21 | 1.32 | -1.75 | 6.81E-09 | 5.86E-07 | (B) Pathology_classA-RNA |
| ENPP7P10 | 0.17 | 1.92 | 0.34 | 1.26 | -1.75 | 1.02E-08 | 8.14E-07 | (B) Pathology_classA-RNA |
| LARP1P1 | 0.27 | 2.01 | 0.46 | 1.47 | -1.75 | 6.05E-07 | 2.00E-05 | (B) Pathology_classA-RNA |
| MIR-5690/56 | 0.02 | 1.77 | 0.05 | 1.93 | -1.75 | 8.19E-06 | 1.56E-04 | (B) Pathology_classA-RNA |
| GGTA1P | 2.47 | 4.22 | 0.97 | 1.74 | -1.75 | 5.70E-04 | 4.14E-03 | (B) Pathology_classA-RNA |
| IGHV3-22 | 0.28 | 2.03 | 0.56 | 2.51 | -1.75 | 5.98E-04 | 4.30E-03 | (B) Pathology_classA-RNA |
| RPS7P15 | 0.56 | 2.31 | 0.98 | 2.24 | -1.75 | 1.51E-03 | 8.67E-03 | (B) Pathology_classA-RNA |
| RNU7-84P | 0.59 | 2.34 | 1.26 | 2.12 | -1.75 | 4.82E-03 | 0.0209 | (B) Pathology_classA-RNA |
| TRGV3 | 1.1 | 2.86 | 0.65 | 1.7 | -1.76 | 2.66E-05 | 3.83E-04 | (B) Pathology_classA-RNA |
| RPL12P11 | 1.08 | 2.84 | 1.06 | 2.2 | -1.76 | 2.05E-03 | 0.0109 | (B) Pathology_classA-RNA |
| SNORD15B | 1.3 | 3.06 | 1.01 | 2.46 | -1.76 | 2.57E-03 | 0.013 | (B) Pathology_classA-RNA |
| HIST2H2BA | 1.3 | 3.07 | 0.65 | 1.45 | -1.77 | 7.67E-06 | 1.48E-04 | (B) Pathology_classA-RNA |
| ZNRF3-IT1 | 0.43 | 2.2 | 0.9 | 2.27 | -1.77 | 1.00E-03 | 6.31E-03 | (B) Pathology_classA-RNA |
| WWTR1-IT1 | 0.79 | 2.56 | 1.05 | 2.37 | -1.77 | 2.42E-03 | 0.0124 | (B) Pathology_classA-RNA |
| TDRP | 0.87 | 2.64 | 1.27 | 1.67 | -1.77 | 3.14E-03 | 0.0151 | (B) Pathology_classA-RNA |
| LINC00612 | 0.35 | 2.13 | 0.41 | 1.2 | -1.78 | 1.43E-08 | 1.02E-06 | (B) Pathology_classA-RNA |
| RNVU1-19 | 0.62 | 2.4 | 0.55 | 1.73 | -1.78 | 8.63E-06 | 1.62E-04 | (B) Pathology_classA-RNA |
| HIST1H2BF | 2.78 | 4.56 | 0.74 | 1.9 | -1.78 | 1.07E-04 | 1.15E-03 | (B) Pathology_classA-RNA |
| SCD | 5 | 6.77 | 1.36 | 1.7 | -1.78 | 4.77E-03 | 0.0208 | (B) Pathology_classA-RNA |
| OPHN1 | 0.24 | 2.03 | 0.46 | 1.7 | -1.79 | 2.77E-06 | 6.59E-05 | (B) Pathology_classA-RNA |
| TRAV14DV4 | 0.31 | 2.1 | 0.54 | 1.62 | -1.79 | 3.77E-06 | 8.47E-05 | (B) Pathology_classA-RNA |
| TRBV29-1 | 0.49 | 2.28 | 0.73 | 2.11 | -1.79 | 2.03E-04 | 1.86E-03 | (B) Pathology_classA-RNA |
| ATP6V0E1P1 | 0.71 | 2.5 | 1.32 | 2.33 | -1.79 | 6.20E-03 | 0.0252 | (B) Pathology_classA-RNA |
| NUPR1 | 0.92 | 2.72 | 1.59 | 1.68 | -1.79 | 0.0105 | 0.037 | (B) Pathology_classA-RNA |
| MTND2P2 | 0.02 | 1.82 | 0.05 | 1.84 | -1.8 | 2.22E-06 | 5.55E-05 | (B) Pathology_classA-RNA |
| RNU6-883P | 0.37 | 2.18 | 0.45 | 1.71 | -1.8 | 2.51E-06 | 6.13E-05 | (B) Pathology_classA-RNA |
| PTGER4 | 0.24 | 2.05 | 0.66 | 1.13 | -1.8 | 3.17E-06 | 7.35E-05 | (B) Pathology_classA-RNA |
| TRAV38-2DV | 0.23 | 2.03 | 0.47 | 1.91 | -1.8 | 1.27E-05 | 2.21E-04 | (B) Pathology_classA-RNA |
| CAP2P1 | 0.74 | 2.54 | 0.92 | 1.8 | -1.8 | 3.16E-04 | 2.63E-03 | (B) Pathology_classA-RNA |
| LTF | 0.22 | 2.02 | 0.67 | 2.97 | -1.8 | 2.41E-03 | 0.0124 | (B) Pathology_classA-RNA |
| ZNF492 | 0.36 | 2.17 | 0.4 | 1.77 | -1.81 | 2.58E-06 | 6.25E-05 | (B) Pathology_classA-RNA |
| YBX1P3 | 0.32 | 2.13 | 0.65 | 1.96 | -1.81 | 5.48E-05 | 6.79E-04 | (B) Pathology_classA-RNA |
| MIR-920/920 | 0.05 | 1.86 | 0.15 | 2.58 | -1.81 | 2.65E-04 | 2.28E-03 | (B) Pathology_classA-RNA |
| GASK1B | 0.55 | 2.36 | 1.24 | 1.23 | -1.81 | 1.92E-03 | 0.0104 | (B) Pathology_classA-RNA |
| LYN | 4.19 | 6 | 1.5 | 1.2 | -1.81 | 6.78E-03 | 0.0269 | (B) Pathology_classA-RNA |
| CCDC200 | 2.51 | 4.32 | 1.52 | 1.1 | -1.81 | 7.10E-03 | 0.0279 | (B) Pathology_classA-RNA |
| PLEKHS1 | 0.96 | 2.77 | 1.49 | 2.1 | -1.81 | 8.77E-03 | 0.0324 | (B) Pathology_classA-RNA |
| TRBJ2-5 | 0.48 | 2.3 | 0.63 | 2.12 | -1.82 | 9.19E-05 | 1.02E-03 | (B) Pathology_classA-RNA |
| USP51 | 1.31 | 3.13 | 0.81 | 1.74 | -1.82 | 9.31E-05 | 1.03E-03 | (B) Pathology_classA-RNA |
| RNU6-106P | 0.92 | 2.74 | 1.39 | 2.61 | -1.82 | 9.29E-03 | 0.0338 | (B) Pathology_classA-RNA |
| IMMP2L | 0.51 | 2.34 | 0.69 | 1.76 | -1.83 | 2.83E-05 | 4.04E-04 | (B) Pathology_classA-RNA |
| RGS2 | 2.89 | 4.72 | 0.85 | 2.06 | -1.83 | 3.02E-04 | 2.54E-03 | (B) Pathology_classA-RNA |
| UGT1A3 | 0.52 | 2.35 | 0.97 | 2.62 | -1.83 | 2.07E-03 | 0.011 | (B) Pathology_classA-RNA |
| MIR-1280/12 | 0.73 | 2.56 | 1.49 | 2.2 | -1.83 | 8.63E-03 | 0.032 | (B) Pathology_classA-RNA |
| LINC00885 | 0.14 | 1.98 | 0.27 | 1.62 | -1.84 | 1.97E-07 | 8.25E-06 | (B) Pathology_classA-RNA |
| PPP1R2P4 | 1.52 | 3.38 | 0.47 | 1.77 | -1.85 | 2.90E-06 | 6.83E-05 | (B) Pathology_classA-RNA |
| RNU11 | 0.27 | 2.13 | 0.47 | 1.89 | -1.85 | 7.01E-06 | 1.38E-04 | (B) Pathology_classA-RNA |
| RN7SL648P | 0.3 | 2.14 | 0.58 | 1.94 | -1.85 | 2.14E-05 | 3.23E-04 | (B) Pathology_classA-RNA |
| RNY4P7 | 0.34 | 2.19 | 0.64 | 2.24 | -1.85 | 1.46E-04 | 1.46E-03 | (B) Pathology_classA-RNA |
| RNU6-171P | 0.55 | 2.4 | 0.84 | 2.04 | -1.85 | 2.23E-04 | 2.00E-03 | (B) Pathology_classA-RNA |
| ABCC11 | 2.65 | 4.5 | 0.78 | 2.22 | -1.85 | 2.79E-04 | 2.38E-03 | (B) Pathology_classA-RNA |
| RNU1-3 | 0.95 | 2.8 | 0.88 | 2.88 | -1.85 | 2.55E-03 | 0.0129 | (B) Pathology_classA-RNA |
| RNU6-1190P | 0.27 | 2.13 | 0.45 | 2.15 | -1.86 | 3.34E-05 | 4.60E-04 | (B) Pathology_classA-RNA |
| HMGN2P27 | 0.19 | 2.05 | 0.4 | 2.28 | -1.86 | 6.01E-05 | 7.30E-04 | (B) Pathology_classA-RNA |
| ADAMTS9-AS | 1.67 | 3.53 | 0.61 | 2.18 | -1.86 | 8.67E-05 | 9.74E-04 | (B) Pathology_classA-RNA |
| FASLG | 0.55 | 2.41 | 1.07 | 1.8 | -1.86 | 7.18E-04 | 4.94E-03 | (B) Pathology_classA-RNA |
| HIST1H2AM | 1.19 | 3.05 | 1.03 | 2.14 | -1.86 | 9.25E-04 | 5.96E-03 | (B) Pathology_classA-RNA |
| RNU6-711P | 0.83 | 2.69 | 1.04 | 2.64 | -1.86 | 2.48E-03 | 0.0126 | (B) Pathology_classA-RNA |
| SPATA20P1 | 0.6 | 2.46 | 1.27 | 2.04 | -1.86 | 2.86E-03 | 0.0141 | (B) Pathology_classA-RNA |
| NAP1L2 | 0.31 | 2.18 | 0.93 | 1.77 | -1.87 | 2.30E-04 | 2.04E-03 | (B) Pathology_classA-RNA |
| NAP1L6 | 0 | 1.88 | 0 | 2.25 | -1.88 | 2.56E-05 | 3.72E-04 | (B) Pathology_classA-RNA |

| | | | | | | | | |
|---|---|---|---|---|---|---|---|---|
| RNA5SP237 | 0.8 | 2.68 | 1.6 | 2.7 | -1.88 | 0.0142 | 0.046 | (B) Pathology_classA-RNA |
| RASSF2 | 3.42 | 5.3 | 0.82 | 1.46 | -1.89 | 3.53E-05 | 4.81E-04 | (B) Pathology_classA-RNA |
| EIF4BP5 | 0.75 | 2.64 | 0.83 | 1.32 | -1.89 | 3.54E-05 | 4.83E-04 | (B) Pathology_classA-RNA |
| CPED1 | 2.44 | 4.32 | 0.97 | 1.66 | -1.89 | 2.43E-04 | 2.13E-03 | (B) Pathology_classA-RNA |
| PLPP1 | 3.16 | 5.05 | 1.28 | 1.99 | -1.89 | 2.55E-03 | 0.0129 | (B) Pathology_classA-RNA |
| HLA-DQA1 | 0.76 | 2.65 | 1.32 | 1.87 | -1.89 | 2.86E-03 | 0.0141 | (B) Pathology_classA-RNA |
| ADAMTS17 | 0.13 | 2.04 | 0.4 | 1.07 | -1.9 | 5.37E-10 | 9.09E-08 | (B) Pathology_classA-RNA |
| HIST1H2AC | 6.37 | 8.27 | 0.79 | 1.08 | -1.9 | 1.88E-05 | 2.94E-04 | (B) Pathology_classA-RNA |
| NAMPTP1 | 2.48 | 4.38 | 0.79 | 1.21 | -1.9 | 1.92E-05 | 2.99E-04 | (B) Pathology_classA-RNA |
| FGR | 1.16 | 3.06 | 1.23 | 2.32 | -1.9 | 2.82E-03 | 0.0139 | (B) Pathology_classA-RNA |
| RPL23AP53 | 2.97 | 4.86 | 1.57 | 0.81 | -1.9 | 6.52E-03 | 0.0261 | (B) Pathology_classA-RNA |
| NDNF | 1.2 | 3.1 | 1.71 | 2.08 | -1.9 | 0.0127 | 0.0425 | (B) Pathology_classA-RNA |
| IGHEP2 | 0.22 | 2.13 | 0.43 | 1.74 | -1.91 | 9.43E-07 | 2.84E-05 | (B) Pathology_classA-RNA |
| HIST1H4E | 2.49 | 4.4 | 0.83 | 1.73 | -1.91 | 6.17E-05 | 7.44E-04 | (B) Pathology_classA-RNA |
| RNU6-469P | 0.82 | 2.74 | 1.29 | 2.34 | -1.91 | 3.43E-03 | 0.0162 | (B) Pathology_classA-RNA |
| HMGN2P35 | 0.72 | 2.63 | 1.34 | 2.23 | -1.91 | 3.80E-03 | 0.0175 | (B) Pathology_classA-RNA |
| TRAJ13 | 0.16 | 2.08 | 0.48 | 1.98 | -1.92 | 7.67E-06 | 1.48E-04 | (B) Pathology_classA-RNA |
| TRAV1-2 | 0.89 | 2.81 | 1.03 | 2.09 | -1.92 | 6.14E-04 | 4.38E-03 | (B) Pathology_classA-RNA |
| LYST-AS1 | 1.08 | 3 | 1.05 | 2 | -1.92 | 6.18E-04 | 4.40E-03 | (B) Pathology_classA-RNA |
| TRGV2 | 0.27 | 2.2 | 0.45 | 1.81 | -1.93 | 1.63E-06 | 4.39E-05 | (B) Pathology_classA-RNA |
| POU5F1P5 | 0.38 | 2.3 | 0.76 | 2.06 | -1.93 | 8.20E-05 | 9.30E-04 | (B) Pathology_classA-RNA |
| PCAT4 | 0.14 | 2.07 | 0.25 | 2.56 | -1.93 | 1.20E-04 | 1.24E-03 | (B) Pathology_classA-RNA |
| SOCS2-AS1 | 0.86 | 2.79 | 1 | 1.7 | -1.93 | 2.44E-04 | 2.14E-03 | (B) Pathology_classA-RNA |
| KNDC1 | 0.79 | 2.72 | 1.01 | 2.3 | -1.93 | 8.01E-04 | 5.38E-03 | (B) Pathology_classA-RNA |
| FCRL5 | 0.38 | 2.32 | 0.92 | 2.52 | -1.93 | 8.48E-04 | 5.60E-03 | (B) Pathology_classA-RNA |
| GZMA | 0.68 | 2.61 | 1.05 | 2.3 | -1.93 | 1.01E-03 | 6.34E-03 | (B) Pathology_classA-RNA |
| HLA-DRA | 5.84 | 7.77 | 1.34 | 2.23 | -1.93 | 3.57E-03 | 0.0166 | (B) Pathology_classA-RNA |
| OR52K3P | 0.21 | 2.15 | 0.57 | 1.97 | -1.94 | 1.09E-05 | 1.95E-04 | (B) Pathology_classA-RNA |
| TPD52L1 | 3.77 | 5.7 | 1.24 | 1.38 | -1.94 | 1.21E-03 | 7.31E-03 | (B) Pathology_classA-RNA |
| TDRD1 | 0.4 | 2.35 | 1.2 | 2.01 | -1.94 | 1.37E-03 | 8.05E-03 | (B) Pathology_classA-RNA |
| CXCR3 | 1.08 | 3.02 | 1.33 | 1.89 | -1.94 | 2.47E-03 | 0.0126 | (B) Pathology_classA-RNA |
| HLA-DRB1 | 5.59 | 7.53 | 1.6 | 2.1 | -1.94 | 8.15E-03 | 0.0307 | (B) Pathology_classA-RNA |
| MIR-214/214 | 1.2 | 3.15 | 0.78 | 2.6 | -1.95 | 4.96E-04 | 3.72E-03 | (B) Pathology_classA-RNA |
| MIR-214/214 | 1.2 | 3.15 | 0.78 | 2.6 | -1.95 | 4.96E-04 | 3.72E-03 | (B) Pathology_classA-RNA |
| MIR-214/3P | 1.2 | 3.15 | 0.78 | 2.6 | -1.95 | 4.96E-04 | 3.72E-03 | (B) Pathology_classA-RNA |
| MIR-214/5P | 1.2 | 3.15 | 0.78 | 2.6 | -1.95 | 4.96E-04 | 3.72E-03 | (B) Pathology_classA-RNA |
| MIR-620/620 | 0.07 | 2.03 | 0.21 | 2.27 | -1.96 | 1.81E-05 | 2.86E-04 | (B) Pathology_classA-RNA |
| CDKL5 | 3.49 | 5.46 | 0.98 | 1.18 | -1.96 | 1.40E-04 | 1.40E-03 | (B) Pathology_classA-RNA |
| GOLGA5P1 | 0.63 | 2.59 | 0.87 | 2.17 | -1.96 | 2.13E-04 | 1.93E-03 | (B) Pathology_classA-RNA |
| IGKV2-30 | 0.93 | 2.9 | 1.8 | 2.46 | -1.96 | 0.0157 | 0.0498 | (B) Pathology_classA-RNA |
| NDUFB4P2 | 0.02 | 1.99 | 0.04 | 2.12 | -1.97 | 5.56E-06 | 1.14E-04 | (B) Pathology_classA-RNA |
| CASP1 | 0.7 | 2.67 | 0.67 | 1.87 | -1.97 | 1.34E-05 | 2.30E-04 | (B) Pathology_classA-RNA |
| SLC9A9 | 2.4 | 4.38 | 0.83 | 1.32 | -1.97 | 2.18E-05 | 3.28E-04 | (B) Pathology_classA-RNA |
| TIGIT | 0.29 | 2.26 | 0.8 | 1.78 | -1.97 | 3.55E-05 | 4.83E-04 | (B) Pathology_classA-RNA |
| XXYLT1-AS2 | 0.6 | 2.56 | 0.96 | 1.38 | -1.97 | 1.09E-04 | 1.16E-03 | (B) Pathology_classA-RNA |
| RPA4 | 0.44 | 2.41 | 0.54 | 1.31 | -1.98 | 7.53E-08 | 3.85E-06 | (B) Pathology_classA-RNA |
| CR1 | 0.08 | 2.05 | 0.23 | 1.75 | -1.98 | 1.84E-07 | 7.90E-06 | (B) Pathology_classA-RNA |
| HIST1H4B | 0.01 | 1.98 | 0.02 | 2.21 | -1.98 | 9.43E-06 | 1.74E-04 | (B) Pathology_classA-RNA |
| HIST1H1C | 6.48 | 8.47 | 0.78 | 1.42 | -1.98 | 1.05E-05 | 1.89E-04 | (B) Pathology_classA-RNA |
| ARHGAP26-IT | 0.46 | 2.44 | 0.59 | 2.56 | -1.98 | 1.77E-04 | 1.67E-03 | (B) Pathology_classA-RNA |
| BLVRB | 0.81 | 2.79 | 1.39 | 2.1 | -1.98 | 3.16E-03 | 0.0152 | (B) Pathology_classA-RNA |
| GATA1 | 0.36 | 2.35 | 0.38 | 2.71 | -1.99 | 1.96E-04 | 1.82E-03 | (B) Pathology_classA-RNA |
| SNORA57 | 4.35 | 6.36 | 0.57 | 1.62 | -2 | 6.91E-07 | 2.21E-05 | (B) Pathology_classA-RNA |
| MTND5P5 | 0.08 | 2.08 | 0.25 | 2.53 | -2 | 6.59E-05 | 7.84E-04 | (B) Pathology_classA-RNA |
| ATP1B1P1 | 0.33 | 2.33 | 0.96 | 1.64 | -2 | 1.08E-04 | 1.15E-03 | (B) Pathology_classA-RNA |
| SCARNA13 | 7.5 | 9.5 | 0.49 | 2.78 | -2 | 3.02E-04 | 2.54E-03 | (B) Pathology_classA-RNA |
| ALAS2 | 0.06 | 2.06 | 0.18 | 3.6 | -2 | 2.83E-03 | 0.014 | (B) Pathology_classA-RNA |
| TAS2R50 | 0.24 | 2.25 | 0.38 | 1.88 | -2.01 | 8.97E-07 | 2.73E-05 | (B) Pathology_classA-RNA |
| MPEG1 | 0.28 | 2.29 | 0.65 | 1.74 | -2.01 | 3.99E-06 | 8.84E-05 | (B) Pathology_classA-RNA |
| NANOGP5 | 0.62 | 2.63 | 0.83 | 2.33 | -2.01 | 1.91E-04 | 1.78E-03 | (B) Pathology_classA-RNA |
| RNU1-106P | 0.82 | 2.83 | 1.43 | 1.38 | -2.01 | 2.60E-03 | 0.0131 | (B) Pathology_classA-RNA |
| HFE | 3.79 | 5.8 | 1.45 | 0.86 | -2.01 | 2.93E-03 | 0.0143 | (B) Pathology_classA-RNA |
| TRGC1 | 0.51 | 2.52 | 1.54 | 2.68 | -2.01 | 7.96E-03 | 0.0302 | (B) Pathology_classA-RNA |
| RNU4-1 | 0.71 | 2.72 | 1.62 | 2.64 | -2.01 | 9.37E-03 | 0.034 | (B) Pathology_classA-RNA |
| RNU6-957P | 0.13 | 2.15 | 0.39 | 2.43 | -2.02 | 4.14E-05 | 5.45E-04 | (B) Pathology_classA-RNA |

| | | | | | | | | |
|---|---|---|---|---|---|---|---|---|
| IGKV1D-12 | 1.13 | 3.14 | 1.51 | 3.08 | -2.02 | 0.0101 | 0.0359 | (B) Pathology_classA-RNA |
| LTB | 1.21 | 3.23 | 1.81 | 2.54 | -2.02 | 0.0143 | 0.0463 | (B) Pathology_classA-RNA |
| RNU6ATAC23 | 0.33 | 2.36 | 0.98 | 2.79 | -2.03 | 1.22E-03 | 7.36E-03 | (B) Pathology_classA-RNA |
| TAS2R18P | 0.37 | 2.41 | 0.47 | 2.06 | -2.04 | 4.76E-06 | 1.02E-04 | (B) Pathology_classA-RNA |
| ANGPTL3 | 0.37 | 2.41 | 0.69 | 2.02 | -2.04 | 1.81E-05 | 2.86E-04 | (B) Pathology_classA-RNA |
| TRGV4 | 2.21 | 4.26 | 1.16 | 2.1 | -2.05 | 7.49E-04 | 5.10E-03 | (B) Pathology_classA-RNA |
| TMPRSS13 | 2.5 | 4.55 | 1.76 | 1.64 | -2.05 | 8.51E-03 | 0.0317 | (B) Pathology_classA-RNA |
| CFL1P7 | 0.05 | 2.11 | 0.14 | 2.38 | -2.06 | 1.60E-05 | 2.61E-04 | (B) Pathology_classA-RNA |
| CFAP69 | 0.15 | 2.21 | 0.39 | 2.35 | -2.06 | 1.96E-05 | 3.03E-04 | (B) Pathology_classA-RNA |
| PADI2 | 0.49 | 2.56 | 0.86 | 2.11 | -2.06 | 8.53E-05 | 9.62E-04 | (B) Pathology_classA-RNA |
| CYBB | 4.05 | 6.11 | 0.93 | 1.89 | -2.06 | 8.95E-05 | 9.98E-04 | (B) Pathology_classA-RNA |
| PCGEM1 | 0.26 | 2.32 | 0.54 | 2.94 | -2.06 | 4.26E-04 | 3.31E-03 | (B) Pathology_classA-RNA |
| MIR4458HG | 1.69 | 3.76 | 1.51 | 1.49 | -2.06 | 3.11E-03 | 0.015 | (B) Pathology_classA-RNA |
| HMGB3P9 | 0.97 | 3.04 | 1 | 2.07 | -2.07 | 2.12E-04 | 1.92E-03 | (B) Pathology_classA-RNA |
| SIGLEC14 | 1.83 | 3.9 | 1.15 | 2.06 | -2.07 | 5.86E-04 | 4.24E-03 | (B) Pathology_classA-RNA |
| ERBB2 | 0.78 | 2.87 | 1.18 | 1.46 | -2.09 | 4.17E-04 | 3.25E-03 | (B) Pathology_classA-RNA |
| SNORD3B-1 | 0.13 | 2.24 | 0.19 | 2.13 | -2.11 | 1.97E-06 | 5.08E-05 | (B) Pathology_classA-RNA |
| HIST1H1T | 0.5 | 2.6 | 0.63 | 1.8 | -2.11 | 2.02E-06 | 5.18E-05 | (B) Pathology_classA-RNA |
| RN7SL471P | 0.03 | 2.13 | 0.08 | 2.48 | -2.11 | 2.07E-05 | 3.15E-04 | (B) Pathology_classA-RNA |
| IGKV1D-27 | 0.28 | 2.39 | 0.56 | 2.54 | -2.11 | 6.46E-05 | 7.71E-04 | (B) Pathology_classA-RNA |
| TRIM22 | 4.73 | 6.84 | 1.16 | 1.48 | -2.11 | 3.27E-04 | 2.69E-03 | (B) Pathology_classA-RNA |
| CDC14A | 3.84 | 5.95 | 1.26 | 1.38 | -2.11 | 6.82E-04 | 4.75E-03 | (B) Pathology_classA-RNA |
| GSTA1 | 0.18 | 2.3 | 0.39 | 2.14 | -2.12 | 3.11E-06 | 7.24E-05 | (B) Pathology_classA-RNA |
| FTO-IT1 | 1.45 | 3.57 | 0.93 | 1.6 | -2.12 | 4.03E-05 | 5.33E-04 | (B) Pathology_classA-RNA |
| CSF2RA | 2.08 | 4.21 | 1.01 | 1.59 | -2.12 | 9.06E-05 | 1.01E-03 | (B) Pathology_classA-RNA |
| IGHV1OR15- | 0.87 | 2.98 | 0.96 | 2.37 | -2.12 | 2.50E-04 | 2.17E-03 | (B) Pathology_classA-RNA |
| PCDH7 | 1.81 | 3.93 | 1.08 | 2.07 | -2.12 | 2.99E-04 | 2.52E-03 | (B) Pathology_classA-RNA |
| AKT3-IT1 | 1 | 3.12 | 1.39 | 2.09 | -2.12 | 1.89E-03 | 0.0103 | (B) Pathology_classA-RNA |
| RUBCNL | 2.04 | 4.18 | 0.69 | 1.86 | -2.14 | 4.11E-06 | 9.09E-05 | (B) Pathology_classA-RNA |
| RAP2B | 2.04 | 4.18 | 1.59 | 0.91 | -2.14 | 3.49E-03 | 0.0164 | (B) Pathology_classA-RNA |
| MTND1P11 | 0.2 | 2.35 | 0.22 | 1.89 | -2.15 | 1.70E-07 | 7.42E-06 | (B) Pathology_classA-RNA |
| MALAT1 | 11.47 | 13.62 | 0.65 | 1.84 | -2.15 | 2.07E-06 | 5.26E-05 | (B) Pathology_classA-RNA |
| HIST1H4F | 0.14 | 2.29 | 0.31 | 2.72 | -2.15 | 7.11E-05 | 8.32E-04 | (B) Pathology_classA-RNA |
| ZNF812P | 1.06 | 3.22 | 0.82 | 2.32 | -2.15 | 7.14E-05 | 8.34E-04 | (B) Pathology_classA-RNA |
| RNU4ATAC1( | 0.41 | 2.55 | 0.84 | 2.31 | -2.15 | 8.19E-05 | 9.30E-04 | (B) Pathology_classA-RNA |
| AHNAK | 1.09 | 3.24 | 1.15 | 0.78 | -2.15 | 3.53E-04 | 2.86E-03 | (B) Pathology_classA-RNA |
| CST7 | 2.79 | 4.94 | 0.98 | 2.63 | -2.15 | 4.56E-04 | 3.48E-03 | (B) Pathology_classA-RNA |
| CD96 | 1.42 | 3.58 | 0.88 | 1.94 | -2.16 | 3.45E-05 | 4.72E-04 | (B) Pathology_classA-RNA |
| SAMD9L | 1.16 | 3.33 | 0.98 | 1.46 | -2.16 | 5.06E-05 | 6.37E-04 | (B) Pathology_classA-RNA |
| MIR-4722/3F | 2.51 | 4.67 | 1.38 | 0.91 | -2.16 | 1.29E-03 | 7.67E-03 | (B) Pathology_classA-RNA |
| MIR-4722/5F | 2.51 | 4.67 | 1.38 | 0.91 | -2.16 | 1.29E-03 | 7.67E-03 | (B) Pathology_classA-RNA |
| RNU6-107P | 0.33 | 2.5 | 0.46 | 2.2 | -2.17 | 3.97E-06 | 8.82E-05 | (B) Pathology_classA-RNA |
| APBB1IP | 0.42 | 2.59 | 0.72 | 1.86 | -2.17 | 4.17E-06 | 9.18E-05 | (B) Pathology_classA-RNA |
| RNU2-61P | 0.17 | 2.34 | 0.5 | 2.7 | -2.17 | 7.78E-05 | 8.91E-04 | (B) Pathology_classA-RNA |
| CLIP4 | 0.49 | 2.67 | 0.76 | 1.22 | -2.18 | 1.76E-06 | 4.68E-05 | (B) Pathology_classA-RNA |
| TRDJ2 | 0.27 | 2.45 | 0.81 | 2.47 | -2.18 | 1.00E-04 | 1.08E-03 | (B) Pathology_classA-RNA |
| UGT1A6 | 0.83 | 3.02 | 1.33 | 2.73 | -2.18 | 2.15E-03 | 0.0113 | (B) Pathology_classA-RNA |
| RNU6-355P | 0.03 | 2.21 | 0.09 | 1.91 | -2.19 | 1.34E-07 | 6.13E-06 | (B) Pathology_classA-RNA |
| S100P | 0.08 | 2.28 | 0.23 | 1.82 | -2.2 | 4.99E-08 | 2.78E-06 | (B) Pathology_classA-RNA |
| GLIPR2 | 2.93 | 5.12 | 0.97 | 1.73 | -2.2 | 4.58E-05 | 5.89E-04 | (B) Pathology_classA-RNA |
| TRAJ32 | 0.31 | 2.53 | 0.94 | 2.05 | -2.21 | 5.79E-05 | 7.09E-04 | (B) Pathology_classA-RNA |
| TNXB | 2.48 | 4.69 | 1 | 1.89 | -2.21 | 7.30E-05 | 8.51E-04 | (B) Pathology_classA-RNA |
| HLA-DQB1 | 2.04 | 4.25 | 1.68 | 1.78 | -2.21 | 4.10E-03 | 0.0185 | (B) Pathology_classA-RNA |
| FOLH1B | 0.98 | 3.19 | 1.97 | 2.4 | -2.21 | 0.0121 | 0.0409 | (B) Pathology_classA-RNA |
| MIR-3139/31 | 0.2 | 2.42 | 0.6 | 2.64 | -2.22 | 6.02E-05 | 7.30E-04 | (B) Pathology_classA-RNA |
| HIST1H3F | 0.77 | 2.99 | 1.44 | 3.05 | -2.22 | 4.12E-03 | 0.0186 | (B) Pathology_classA-RNA |
| HIST2H2BB | 0.67 | 2.9 | 0.64 | 1.32 | -2.23 | 8.42E-08 | 4.19E-06 | (B) Pathology_classA-RNA |
| CD8B | 1.9 | 4.14 | 1.35 | 2.13 | -2.23 | 9.74E-04 | 6.19E-03 | (B) Pathology_classA-RNA |
| MIR-4802/3F | 0.4 | 2.64 | 0.81 | 2.46 | -2.24 | 6.77E-05 | 8.02E-04 | (B) Pathology_classA-RNA |
| MIR-4802/5F | 0.4 | 2.64 | 0.81 | 2.46 | -2.24 | 6.77E-05 | 8.02E-04 | (B) Pathology_classA-RNA |
| CCL5 | 3.84 | 6.1 | 1 | 2.33 | -2.25 | 1.44E-04 | 1.43E-03 | (B) Pathology_classA-RNA |
| CCDC69 | 2.67 | 4.93 | 1.05 | 1.43 | -2.26 | 6.32E-05 | 7.59E-04 | (B) Pathology_classA-RNA |
| PRR33 | 4.63 | 6.9 | 0.97 | 2.23 | -2.26 | 8.61E-05 | 9.70E-04 | (B) Pathology_classA-RNA |
| SUMO2P8 | 0.39 | 2.66 | 0.81 | 2.4 | -2.27 | 4.53E-05 | 5.84E-04 | (B) Pathology_classA-RNA |
| ELMO1-AS1 | 1.31 | 3.58 | 1.17 | 1.03 | -2.27 | 2.10E-04 | 1.91E-03 | (B) Pathology_classA-RNA |

| | | | | | | | | |
|---|---|---|---|---|---|---|---|---|
| IGKV1-27 | 0.78 | 3.05 | 1.62 | 2.87 | -2.27 | 5.00E-03 | 0.0215 | (B) Pathology_classA-RNA |
| DAPP1 | 0.29 | 2.57 | 0.86 | 2.17 | -2.28 | 2.62E-05 | 3.79E-04 | (B) Pathology_classA-RNA |
| SETP21 | 2.09 | 4.37 | 1.34 | 3.02 | -2.28 | 2.25E-03 | 0.0117 | (B) Pathology_classA-RNA |
| VCAM1 | 0.5 | 2.79 | 0.52 | 1.56 | -2.29 | 1.21E-08 | 9.09E-07 | (B) Pathology_classA-RNA |
| HMGB1P41 | 1.21 | 3.5 | 0.73 | 1.72 | -2.29 | 1.02E-06 | 3.03E-05 | (B) Pathology_classA-RNA |
| DENND2C | 1.27 | 3.57 | 0.41 | 0.87 | -2.3 | 4.36E-12 | 2.12E-09 | (B) Pathology_classA-RNA |
| PART1 | 0.74 | 3.04 | 1.3 | 2.02 | -2.3 | 5.27E-04 | 3.88E-03 | (B) Pathology_classA-RNA |
| CCDC160 | 1.41 | 3.71 | 1.77 | 2.14 | -2.3 | 4.78E-03 | 0.0208 | (B) Pathology_classA-RNA |
| MTRNR2L9 | 0.45 | 2.75 | 0.61 | 2.43 | -2.31 | 1.23E-05 | 2.15E-04 | (B) Pathology_classA-RNA |
| DEFA1B | 0.1 | 2.41 | 0.28 | 4.02 | -2.31 | 2.14E-03 | 0.0113 | (B) Pathology_classA-RNA |
| S100A9 | 6.7 | 9.02 | 1.93 | 3.53 | -2.32 | 0.0147 | 0.0473 | (B) Pathology_classA-RNA |
| SLC25A37 | 1.73 | 4.06 | 0.63 | 1.52 | -2.33 | 5.66E-08 | 3.05E-06 | (B) Pathology_classA-RNA |
| FOXP1-IT1 | 0.13 | 2.48 | 0.23 | 2.16 | -2.35 | 4.01E-07 | 1.44E-05 | (B) Pathology_classA-RNA |
| C19ORF81 | 0.59 | 2.93 | 1.01 | 2.02 | -2.35 | 4.71E-05 | 6.02E-04 | (B) Pathology_classA-RNA |
| RNU6-177P | 0.47 | 2.82 | 1.4 | 2.75 | -2.35 | 1.52E-03 | 8.70E-03 | (B) Pathology_classA-RNA |
| PCAT1 | 0.26 | 2.62 | 0.7 | 2.08 | -2.36 | 2.50E-06 | 6.11E-05 | (B) Pathology_classA-RNA |
| NRIP1 | 1.73 | 4.09 | 1.22 | 1.08 | -2.36 | 2.14E-04 | 1.94E-03 | (B) Pathology_classA-RNA |
| ABCC4 | 3.08 | 5.46 | 1.75 | 1.7 | -2.37 | 3.24E-03 | 0.0155 | (B) Pathology_classA-RNA |
| DIAPH2-AS1 | 1.2 | 3.59 | 0.6 | 1.2 | -2.39 | 6.72E-09 | 5.81E-07 | (B) Pathology_classA-RNA |
| RPS29P22 | 0.8 | 3.19 | 1.16 | 2.23 | -2.39 | 1.68E-04 | 1.62E-03 | (B) Pathology_classA-RNA |
| TREML1 | 1.7 | 4.1 | 1.32 | 2.24 | -2.4 | 4.72E-04 | 3.58E-03 | (B) Pathology_classA-RNA |
| RNF144B | 0.14 | 2.56 | 0.42 | 1.4 | -2.42 | 7.46E-11 | 1.85E-08 | (B) Pathology_classA-RNA |
| TRBJ2-7 | 0.16 | 2.58 | 0.32 | 2.42 | -2.42 | 2.01E-06 | 5.16E-05 | (B) Pathology_classA-RNA |
| PADI4 | 0.49 | 2.91 | 0.84 | 3.52 | -2.42 | 7.76E-04 | 5.23E-03 | (B) Pathology_classA-RNA |
| IGLJ6 | 1.17 | 3.59 | 1.92 | 3.1 | -2.42 | 8.42E-03 | 0.0314 | (B) Pathology_classA-RNA |
| ITK | 0.46 | 2.89 | 0.43 | 1.9 | -2.43 | 2.92E-08 | 1.81E-06 | (B) Pathology_classA-RNA |
| GVINP1 | 0.47 | 2.91 | 0.61 | 1.58 | -2.44 | 2.03E-08 | 1.34E-06 | (B) Pathology_classA-RNA |
| UGT1A7 | 1.19 | 3.65 | 1.7 | 2.84 | -2.46 | 3.35E-03 | 0.0159 | (B) Pathology_classA-RNA |
| RNU1-1 | 0.21 | 2.68 | 0.62 | 2.96 | -2.47 | 5.56E-05 | 6.87E-04 | (B) Pathology_classA-RNA |
| MIR-1205/12 | 0.3 | 2.79 | 0.64 | 2.65 | -2.49 | 1.24E-05 | 2.17E-04 | (B) Pathology_classA-RNA |
| SNORA74A | 2.5 | 4.99 | 1.57 | 3.53 | -2.49 | 3.99E-03 | 0.0181 | (B) Pathology_classA-RNA |
| TRDC | 0.73 | 3.23 | 0.7 | 2.3 | -2.5 | 2.83E-06 | 6.71E-05 | (B) Pathology_classA-RNA |
| OAS2 | 1.31 | 3.81 | 1.4 | 1.86 | -2.5 | 4.12E-04 | 3.23E-03 | (B) Pathology_classA-RNA |
| TRBC2 | 2.68 | 5.17 | 1.25 | 2.98 | -2.5 | 6.17E-04 | 4.40E-03 | (B) Pathology_classA-RNA |
| GPR183 | 1.49 | 4 | 1.28 | 1.46 | -2.51 | 1.62E-04 | 1.57E-03 | (B) Pathology_classA-RNA |
| HIST1H4C | 1.15 | 3.65 | 1.23 | 2.92 | -2.51 | 4.91E-04 | 3.69E-03 | (B) Pathology_classA-RNA |
| TMED11P | 0.62 | 3.14 | 1.22 | 2.1 | -2.52 | 1.29E-04 | 1.32E-03 | (B) Pathology_classA-RNA |
| NOTCH2 | 0.53 | 3.06 | 0.97 | 0.58 | -2.53 | 2.79E-05 | 3.98E-04 | (B) Pathology_classA-RNA |
| PDLIM5 | 0.74 | 3.28 | 0.92 | 2.22 | -2.54 | 1.06E-05 | 1.91E-04 | (B) Pathology_classA-RNA |
| PTGER2 | 1.05 | 3.59 | 1.13 | 1.62 | -2.54 | 3.82E-05 | 5.14E-04 | (B) Pathology_classA-RNA |
| FGL2 | 0.55 | 3.1 | 0.79 | 1.71 | -2.55 | 4.58E-07 | 1.60E-05 | (B) Pathology_classA-RNA |
| HIST2H3C | 3.21 | 5.75 | 1.24 | 1.85 | -2.55 | 1.10E-04 | 1.16E-03 | (B) Pathology_classA-RNA |
| SNORD3D | 1.72 | 4.28 | 0.88 | 3.01 | -2.56 | 9.65E-05 | 1.06E-03 | (B) Pathology_classA-RNA |
| PSTPIP1 | 0.19 | 2.76 | 0.39 | 2.19 | -2.57 | 1.13E-07 | 5.33E-06 | (B) Pathology_classA-RNA |
| RNF138P1 | 4.98 | 7.55 | 0.72 | 2.33 | -2.57 | 2.43E-06 | 5.97E-05 | (B) Pathology_classA-RNA |
| CHRM3-AS2 | 0.16 | 2.74 | 0.3 | 1.66 | -2.58 | 1.56E-10 | 3.48E-08 | (B) Pathology_classA-RNA |
| TRAV12-3 | 0.2 | 2.78 | 0.43 | 2.34 | -2.58 | 4.15E-07 | 1.48E-05 | (B) Pathology_classA-RNA |
| TRAC | 0.59 | 3.17 | 0.66 | 2.56 | -2.58 | 5.07E-06 | 1.07E-04 | (B) Pathology_classA-RNA |
| TERC | 0.11 | 2.69 | 0.21 | 2.99 | -2.58 | 1.80E-05 | 2.85E-04 | (B) Pathology_classA-RNA |
| SNORD3C | 1.96 | 4.55 | 0.94 | 3.88 | -2.58 | 1.09E-03 | 6.73E-03 | (B) Pathology_classA-RNA |
| SCARNA6 | 0.4 | 2.99 | 0.39 | 2.93 | -2.59 | 1.47E-05 | 2.47E-04 | (B) Pathology_classA-RNA |
| LPAR3 | 2.15 | 4.74 | 1.65 | 2.67 | -2.59 | 1.62E-03 | 9.14E-03 | (B) Pathology_classA-RNA |
| HIST2H2AB | 0.56 | 3.16 | 0.39 | 2.29 | -2.6 | 2.07E-07 | 8.60E-06 | (B) Pathology_classA-RNA |
| LINC00861 | 0.89 | 3.51 | 0.82 | 2.24 | -2.61 | 2.84E-06 | 6.73E-05 | (B) Pathology_classA-RNA |
| GBP5 | 0.37 | 2.99 | 0.88 | 2.22 | -2.62 | 4.46E-06 | 9.64E-05 | (B) Pathology_classA-RNA |
| MYBPC1 | 0.46 | 3.1 | 1.12 | 2.17 | -2.64 | 3.44E-05 | 4.71E-04 | (B) Pathology_classA-RNA |
| SERPINB11 | 0.68 | 3.32 | 1.46 | 2.85 | -2.64 | 7.32E-04 | 5.00E-03 | (B) Pathology_classA-RNA |
| SH2D1A | 0.68 | 3.33 | 0.7 | 2.38 | -2.65 | 1.52E-06 | 4.15E-05 | (B) Pathology_classA-RNA |
| RNVU1-7 | 0.4 | 3.05 | 1.18 | 3.26 | -2.65 | 4.36E-04 | 3.36E-03 | (B) Pathology_classA-RNA |
| DES | 0.58 | 3.24 | 1.07 | 3.63 | -2.66 | 6.11E-04 | 4.37E-03 | (B) Pathology_classA-RNA |
| UGT1A5 | 1.54 | 4.21 | 2.01 | 3.02 | -2.67 | 5.34E-03 | 0.0225 | (B) Pathology_classA-RNA |
| HIST1H4D | 5.06 | 7.73 | 2.18 | 1.94 | -2.67 | 6.18E-03 | 0.0252 | (B) Pathology_classA-RNA |
| RN7SKP71 | 0.01 | 2.69 | 0.02 | 2.93 | -2.69 | 6.76E-06 | 1.34E-04 | (B) Pathology_classA-RNA |
| ITGB2-AS1 | 1.94 | 4.63 | 1.11 | 2.44 | -2.69 | 4.04E-05 | 5.34E-04 | (B) Pathology_classA-RNA |
| ARAP2 | 1.59 | 4.3 | 1.68 | 0.97 | -2.71 | 1.07E-03 | 6.63E-03 | (B) Pathology_classA-RNA |

| | | | | | | | | |
|---|---|---|---|---|---|---|---|---|
| ALDH1A3 | 0.97 | 3.69 | 1.26 | 2.04 | -2.72 | 6.69E-05 | 7.94E-04 | (B) Pathology_classA-RNA |
| SCARNA17 | 2.02 | 4.76 | 1.04 | 3 | -2.74 | 8.19E-05 | 9.30E-04 | (B) Pathology_classA-RNA |
| TRBJ2-6 | 1.4 | 4.14 | 1.67 | 2.99 | -2.75 | 1.39E-03 | 8.12E-03 | (B) Pathology_classA-RNA |
| UGT1A10 | 1.56 | 4.32 | 2.11 | 3.14 | -2.75 | 5.95E-03 | 0.0244 | (B) Pathology_classA-RNA |
| STARD13-AS | 0.22 | 2.98 | 0.43 | 2.6 | -2.76 | 7.75E-07 | 2.43E-05 | (B) Pathology_classA-RNA |
| LINC00668 | 0.48 | 3.26 | 0.39 | 3.64 | -2.77 | 1.10E-04 | 1.16E-03 | (B) Pathology_classA-RNA |
| RNU4-25P | 0.81 | 3.58 | 1.56 | 1.8 | -2.77 | 4.12E-04 | 3.23E-03 | (B) Pathology_classA-RNA |
| TRAV8-3 | 0.5 | 3.29 | 0.6 | 2.12 | -2.79 | 4.39E-08 | 2.49E-06 | (B) Pathology_classA-RNA |
| SNORD3A | 5.91 | 8.69 | 1.85 | 4.06 | -2.79 | 5.49E-03 | 0.023 | (B) Pathology_classA-RNA |
| SPATA13 | 2.25 | 5.05 | 1.47 | 0.94 | -2.8 | 3.20E-04 | 2.65E-03 | (B) Pathology_classA-RNA |
| RNU2-59P | 1.29 | 4.09 | 1.34 | 3.49 | -2.8 | 6.31E-04 | 4.47E-03 | (B) Pathology_classA-RNA |
| HLA-DRB5 | 3.64 | 6.44 | 1.49 | 2.11 | -2.81 | 2.60E-04 | 2.24E-03 | (B) Pathology_classA-RNA |
| CYSLTR1 | 2.66 | 5.48 | 1.08 | 1.57 | -2.82 | 6.46E-06 | 1.29E-04 | (B) Pathology_classA-RNA |
| IGKV1-17 | 1.39 | 4.21 | 2.55 | 3.51 | -2.82 | 0.0151 | 0.0481 | (B) Pathology_classA-RNA |
| SERINC5 | 0.55 | 3.38 | 0.82 | 1.25 | -2.83 | 1.19E-07 | 5.51E-06 | (B) Pathology_classA-RNA |
| SIDT1-AS1 | 0.78 | 3.61 | 0.9 | 2.3 | -2.83 | 1.92E-06 | 4.97E-05 | (B) Pathology_classA-RNA |
| MIR-3648-1/ | 0.31 | 3.14 | 0.93 | 3.55 | -2.83 | 1.71E-04 | 1.63E-03 | (B) Pathology_classA-RNA |
| MIR-3648-2/ | 0.31 | 3.14 | 0.93 | 3.55 | -2.83 | 1.71E-04 | 1.63E-03 | (B) Pathology_classA-RNA |
| MIR-3648/36 | 0.31 | 3.14 | 0.93 | 3.55 | -2.83 | 1.71E-04 | 1.63E-03 | (B) Pathology_classA-RNA |
| DEFA1 | 1.34 | 4.2 | 1.7 | 5.19 | -2.86 | 9.82E-03 | 0.0352 | (B) Pathology_classA-RNA |
| SNORA49 | 0.11 | 2.98 | 0.23 | 3.26 | -2.87 | 1.35E-05 | 2.31E-04 | (B) Pathology_classA-RNA |
| TRAV18 | 0.5 | 3.37 | 1.33 | 2.3 | -2.87 | 7.56E-05 | 8.74E-04 | (B) Pathology_classA-RNA |
| MIR-142/3P | 0.95 | 3.83 | 1.29 | 2.33 | -2.88 | 5.84E-05 | 7.14E-04 | (B) Pathology_classA-RNA |
| MIR-142/5P | 0.95 | 3.83 | 1.29 | 2.33 | -2.88 | 5.84E-05 | 7.14E-04 | (B) Pathology_classA-RNA |
| CEBPA | 1.79 | 4.69 | 1.52 | 1.42 | -2.89 | 2.48E-04 | 2.17E-03 | (B) Pathology_classA-RNA |
| RNU2-2P | 6.25 | 9.14 | 1.75 | 4.95 | -2.89 | 8.04E-03 | 0.0304 | (B) Pathology_classA-RNA |
| MNDA | 4.37 | 7.29 | 1.37 | 2.98 | -2.92 | 1.98E-04 | 1.83E-03 | (B) Pathology_classA-RNA |
| FOLH1 | 1.18 | 4.13 | 2.42 | 2.84 | -2.96 | 7.11E-03 | 0.0279 | (B) Pathology_classA-RNA |
| PIGR | 2.01 | 4.98 | 2.04 | 3.45 | -2.97 | 3.36E-03 | 0.0159 | (B) Pathology_classA-RNA |
| NAALADL2 | 4.48 | 7.46 | 1.87 | 1.79 | -2.98 | 1.01E-03 | 6.34E-03 | (B) Pathology_classA-RNA |
| TRAJ6 | 0.25 | 3.24 | 0.76 | 3.12 | -2.99 | 9.90E-06 | 1.81E-04 | (B) Pathology_classA-RNA |
| MS4A1 | 0.44 | 3.44 | 1.23 | 3.07 | -3 | 7.74E-05 | 8.89E-04 | (B) Pathology_classA-RNA |
| DSC2 | 2.51 | 5.52 | 2.73 | 1.68 | -3.01 | 0.0107 | 0.0374 | (B) Pathology_classA-RNA |
| MIR-5704/57 | 0.46 | 3.48 | 1.33 | 2.27 | -3.02 | 3.92E-05 | 5.22E-04 | (B) Pathology_classA-RNA |
| HBD | 1.21 | 4.24 | 1.25 | 5.09 | -3.03 | 3.19E-03 | 0.0153 | (B) Pathology_classA-RNA |
| PYHIN1 | 0.63 | 3.72 | 0.76 | 1.91 | -3.09 | 1.20E-08 | 9.09E-07 | (B) Pathology_classA-RNA |
| TRGV10 | 0.74 | 3.84 | 1.85 | 2.17 | -3.1 | 6.68E-04 | 4.67E-03 | (B) Pathology_classA-RNA |
| RMRP | 0.13 | 3.24 | 0.38 | 3.53 | -3.11 | 1.37E-05 | 2.34E-04 | (B) Pathology_classA-RNA |
| LYZ | 6.24 | 9.34 | 1.42 | 2.95 | -3.11 | 1.17E-04 | 1.22E-03 | (B) Pathology_classA-RNA |
| TRGJ2 | 0.45 | 3.57 | 1.14 | 2.97 | -3.12 | 2.04E-05 | 3.13E-04 | (B) Pathology_classA-RNA |
| TRBV4-1 | 0.25 | 3.38 | 0.49 | 2.82 | -3.14 | 3.37E-07 | 1.26E-05 | (B) Pathology_classA-RNA |
| CPNE4 | 1.15 | 4.3 | 1.57 | 2.12 | -3.14 | 1.34E-04 | 1.35E-03 | (B) Pathology_classA-RNA |
| SCARNA5 | 0.22 | 3.37 | 0.26 | 3.55 | -3.15 | 1.16E-05 | 2.05E-04 | (B) Pathology_classA-RNA |
| TAF4B | 1.22 | 4.41 | 1.75 | 0.71 | -3.18 | 5.32E-04 | 3.91E-03 | (B) Pathology_classA-RNA |
| RN7SKP203 | 0.17 | 3.39 | 0.52 | 3.66 | -3.21 | 1.73E-05 | 2.77E-04 | (B) Pathology_classA-RNA |
| DOCK8 | 2.69 | 5.9 | 1.4 | 1.61 | -3.21 | 3.62E-05 | 4.91E-04 | (B) Pathology_classA-RNA |
| PVT1 | 3.09 | 6.3 | 2.33 | 1.23 | -3.22 | 2.97E-03 | 0.0145 | (B) Pathology_classA-RNA |
| PSTPIP2 | 5.93 | 9.17 | 1.02 | 1.32 | -3.24 | 6.62E-07 | 2.13E-05 | (B) Pathology_classA-RNA |
| PTPRC | 0.7 | 3.95 | 0.99 | 2 | -3.26 | 2.23E-07 | 9.11E-06 | (B) Pathology_classA-RNA |
| MIR-663/663 | 1.95 | 5.22 | 1.69 | 5.18 | -3.26 | 3.55E-03 | 0.0166 | (B) Pathology_classA-RNA |
| MIR-663A/66 | 1.95 | 5.22 | 1.69 | 5.18 | -3.26 | 3.55E-03 | 0.0166 | (B) Pathology_classA-RNA |
| GALNT17 | 0.6 | 3.89 | 0.97 | 2.35 | -3.28 | 3.57E-07 | 1.30E-05 | (B) Pathology_classA-RNA |
| FCN1 | 0.75 | 4.1 | 1.08 | 3.52 | -3.35 | 2.43E-05 | 3.56E-04 | (B) Pathology_classA-RNA |
| NOTCH2NLA | 3.88 | 7.24 | 1.75 | 0.99 | -3.36 | 3.17E-04 | 2.63E-03 | (B) Pathology_classA-RNA |
| SNORA73B | 3.45 | 6.9 | 0.83 | 4.16 | -3.45 | 5.65E-05 | 6.97E-04 | (B) Pathology_classA-RNA |
| CPLX3 | 2.33 | 5.81 | 2.09 | 3.1 | -3.48 | 8.48E-04 | 5.60E-03 | (B) Pathology_classA-RNA |
| TRGJP | 0.95 | 4.56 | 2.85 | 3.78 | -3.61 | 6.17E-03 | 0.0251 | (B) Pathology_classA-RNA |
| SCARNA2 | 0.52 | 4.22 | 0.86 | 4.38 | -3.7 | 4.18E-05 | 5.50E-04 | (B) Pathology_classA-RNA |
| TRAV3 | 0.51 | 4.29 | 1.02 | 3.21 | -3.78 | 8.31E-07 | 2.58E-05 | (B) Pathology_classA-RNA |
| PLA2G2A | 3.63 | 7.45 | 2.03 | 3.82 | -3.81 | 4.48E-04 | 3.44E-03 | (B) Pathology_classA-RNA |
| KLK4 | 0.76 | 4.59 | 2.27 | 3.01 | -3.83 | 6.82E-04 | 4.75E-03 | (B) Pathology_classA-RNA |
| TC2N | 3.36 | 7.24 | 1.89 | 1.11 | -3.88 | 1.87E-04 | 1.75E-03 | (B) Pathology_classA-RNA |
| MIR-3609/36 | 0.63 | 4.55 | 1.02 | 4.14 | -3.92 | 1.19E-05 | 2.09E-04 | (B) Pathology_classA-RNA |
| KLK2 | 1.01 | 4.93 | 2.84 | 3.18 | -3.92 | 3.02E-03 | 0.0147 | (B) Pathology_classA-RNA |
| TFAP2C | 1.26 | 5.26 | 1.79 | 1.72 | -4.01 | 5.64E-05 | 6.97E-04 | (B) Pathology_classA-RNA |

| | | | | | | | | |
|---|---|---|---|---|---|---|---|---|
| MIR-5687/56 | 0.77 | 4.81 | 1.05 | 3.67 | -4.04 | 1.57E-06 | 4.25E-05 | (B) Pathology_classA-RNA |
| RN7SL3 | 2.23 | 6.3 | 1.88 | 4.23 | -4.06 | 1.94E-04 | 1.81E-03 | (B) Pathology_classA-RNA |
| HMGCS2 | 0.77 | 4.85 | 1.15 | 3.14 | -4.08 | 3.82E-07 | 1.39E-05 | (B) Pathology_classA-RNA |
| SCARNA21 | 2.27 | 6.39 | 1.07 | 3.61 | -4.12 | 1.03E-06 | 3.06E-05 | (B) Pathology_classA-RNA |
| TRGJ1 | 0.75 | 5.02 | 2.23 | 3.15 | -4.27 | 2.11E-04 | 1.91E-03 | (B) Pathology_classA-RNA |
| TRGC2 | 1.01 | 5.34 | 2.68 | 3.38 | -4.32 | 9.78E-04 | 6.21E-03 | (B) Pathology_classA-RNA |
| RPPH1 | 0.42 | 4.76 | 0.66 | 5.01 | -4.33 | 2.05E-05 | 3.14E-04 | (B) Pathology_classA-RNA |
| RN7SL4P | 10.48 | 14.91 | 1.68 | 3.84 | -4.44 | 1.51E-05 | 2.51E-04 | (B) Pathology_classA-RNA |
| RN7SL2 | 9.96 | 14.65 | 1.62 | 4.11 | -4.69 | 7.35E-06 | 1.43E-04 | (B) Pathology_classA-RNA |
| AR | 0.57 | 5.28 | 1.72 | 2.57 | -4.7 | 3.42E-06 | 7.83E-05 | (B) Pathology_classA-RNA |
| SCARNA7 | 0.77 | 5.48 | 0.83 | 4.52 | -4.72 | 1.20E-06 | 3.43E-05 | (B) Pathology_classA-RNA |
| PRAC2 | 2.72 | 7.62 | 3.92 | 2.33 | -4.9 | 5.33E-03 | 0.0225 | (B) Pathology_classA-RNA |
| RN7SL5P | 6.06 | 10.97 | 1.73 | 3.94 | -4.91 | 4.85E-06 | 1.03E-04 | (B) Pathology_classA-RNA |
| RN7SK | 2.51 | 7.42 | 1.59 | 5.12 | -4.91 | 2.36E-05 | 3.48E-04 | (B) Pathology_classA-RNA |
| RN7SL1 | 9.18 | 14.14 | 1.7 | 4.17 | -4.96 | 4.97E-06 | 1.05E-04 | (B) Pathology_classA-RNA |
| MIR-3687-1/ | 2.19 | 7.24 | 3.58 | 6.6 | -5.04 | 5.25E-03 | 0.0223 | (B) Pathology_classA-RNA |
| MIR-3687-2/ | 2.19 | 7.24 | 3.58 | 6.6 | -5.04 | 5.25E-03 | 0.0223 | (B) Pathology_classA-RNA |
| MIR-3687/36 | 2.19 | 7.24 | 3.58 | 6.6 | -5.04 | 5.25E-03 | 0.0223 | (B) Pathology_classA-RNA |
| NKX3-1 | 0.79 | 5.99 | 2.36 | 2.25 | -5.21 | 6.55E-05 | 7.80E-04 | (B) Pathology_classA-RNA |
| SPDEF | 2.07 | 7.34 | 3.37 | 2.18 | -5.27 | 1.28E-03 | 7.63E-03 | (B) Pathology_classA-RNA |
| KLK3 | 1.44 | 7.65 | 3.45 | 3.43 | -6.21 | 3.82E-04 | 3.04E-03 | (B) Pathology_classA-RNA |
