## Supplementary material for "KDM4A promotes the progression of neuroendocrine prostate cancer": Suppl. Table 2

| Suppl.Table 2_DE_KDM4A_class B-D_vs_C-E_mRNA expression z-Scores relative to diploid samples | | | | | | | | |
|---|---|---|---|---|---|---|---|---|
| Gene | (A) NE (B/D) | (B) Small cel | (A) NE (B/D) | (B) Small cel | Log Ratio | p-Value | q-Value | Higher expression in |
| CRISP3 | 7.99 | 0.52 | 3.26 | 0.92 | 7.47 | 7.64E-05 | 7.35E-03 | (A) NE (B/D) |
| GPC5-AS1 | 8.95 | 1.83 | 2.12 | 2.38 | 7.12 | 1.48E-04 | 9.80E-03 | (A) NE (B/D) |
| SLC26A9 | 7.66 | 0.99 | 3.59 | 2.04 | 6.67 | 5.38E-04 | 0.0186 | (A) NE (B/D) |
| CSAG2 | 7.11 | 1.17 | 4.05 | 2.04 | 5.94 | 2.66E-03 | 0.0401 | (A) NE (B/D) |
| UCN3 | 6.29 | 0.38 | 3.08 | 0.58 | 5.91 | 3.48E-04 | 0.0151 | (A) NE (B/D) |
| CPHL1P | 7.79 | 1.98 | 1.72 | 1.01 | 5.81 | 1.74E-06 | 8.29E-04 | (A) NE (B/D) |
| TSPAN1 | 6.46 | 1.04 | 1.49 | 1.23 | 5.42 | 5.24E-06 | 1.65E-03 | (A) NE (B/D) |
| CHGB | 16.11 | 11.14 | 2.03 | 2.14 | 4.98 | 1.02E-03 | 0.0256 | (A) NE (B/D) |
| FABP5P7 | 13.5 | 8.64 | 1.96 | 2.07 | 4.86 | 9.50E-04 | 0.0249 | (A) NE (B/D) |
| BCAM | 4.97 | 0.15 | 0.8 | 0.38 | 4.81 | 2.15E-09 | 1.68E-05 | (A) NE (B/D) |
| CALCA | 5.44 | 0.64 | 2.63 | 1.57 | 4.8 | 7.07E-04 | 0.0215 | (A) NE (B/D) |
| SCG2 | 5.99 | 1.22 | 2.35 | 1.96 | 4.77 | 1.07E-03 | 0.026 | (A) NE (B/D) |
| MARCH11 | 6.25 | 1.58 | 1.37 | 2.31 | 4.67 | 2.58E-03 | 0.0397 | (A) NE (B/D) |
| FABP5P1 | 6.15 | 1.49 | 1.74 | 1.24 | 4.66 | 4.36E-05 | 5.44E-03 | (A) NE (B/D) |
| FAM3B | 5.8 | 1.36 | 0.61 | 1.94 | 4.44 | 1.95E-03 | 0.0351 | (A) NE (B/D) |
| LINC00404 | 5.46 | 1.05 | 2.22 | 1.78 | 4.4 | 1.05E-03 | 0.0259 | (A) NE (B/D) |
| MIR-642/642 | 5.3 | 0.91 | 2.56 | 1.43 | 4.39 | 1.00E-03 | 0.0254 | (A) NE (B/D) |
| MIR-642A/3F | 5.3 | 0.91 | 2.56 | 1.43 | 4.39 | 1.00E-03 | 0.0254 | (A) NE (B/D) |
| MIR-642A/5F | 5.3 | 0.91 | 2.56 | 1.43 | 4.39 | 1.00E-03 | 0.0254 | (A) NE (B/D) |
| MIR-642A/64 | 5.3 | 0.91 | 2.56 | 1.43 | 4.39 | 1.00E-03 | 0.0254 | (A) NE (B/D) |
| MIR-642A/64 | 5.3 | 0.91 | 2.56 | 1.43 | 4.39 | 1.00E-03 | 0.0254 | (A) NE (B/D) |
| CEACAM5 | 5.13 | 0.76 | 2.12 | 1.37 | 4.38 | 3.20E-04 | 0.0146 | (A) NE (B/D) |
| RN7SKP106 | 4.99 | 0.62 | 3.2 | 1.52 | 4.37 | 4.03E-03 | 0.0486 | (A) NE (B/D) |
| CAPN13 | 5.64 | 1.32 | 1.28 | 1.83 | 4.33 | 9.32E-04 | 0.0247 | (A) NE (B/D) |
| SLC4A11 | 5.89 | 1.61 | 1.67 | 1.2 | 4.28 | 6.63E-05 | 6.86E-03 | (A) NE (B/D) |
| FABP6 | 5.53 | 1.26 | 2.28 | 1.53 | 4.26 | 8.16E-04 | 0.0231 | (A) NE (B/D) |
| MAGEA2 | 4.94 | 0.68 | 2.81 | 1.66 | 4.26 | 2.79E-03 | 0.041 | (A) NE (B/D) |
| SOX1-OT | 4.87 | 0.74 | 1.97 | 1.49 | 4.13 | 5.08E-04 | 0.0184 | (A) NE (B/D) |
| TMEM125 | 9.27 | 5.18 | 0.75 | 1.75 | 4.09 | 1.46E-03 | 0.0307 | (A) NE (B/D) |
| BPIFB6 | 4.22 | 0.15 | 2.19 | 0.37 | 4.07 | 4.51E-04 | 0.0175 | (A) NE (B/D) |
| LINC00958 | 4.72 | 0.67 | 2.56 | 0.78 | 4.06 | 1.20E-03 | 0.0274 | (A) NE (B/D) |
| SPRY2 | 7.41 | 3.38 | 1.41 | 1.97 | 4.02 | 2.27E-03 | 0.0376 | (A) NE (B/D) |
| BAALC-AS2 | 4.5 | 0.52 | 2.42 | 0.98 | 3.98 | 9.50E-04 | 0.0249 | (A) NE (B/D) |
| FABP5P2 | 7.45 | 3.53 | 1.85 | 0.97 | 3.92 | 1.52E-04 | 9.92E-03 | (A) NE (B/D) |
| SLC1A5 | 9.88 | 5.98 | 0.77 | 1.2 | 3.9 | 1.29E-04 | 9.17E-03 | (A) NE (B/D) |
| EPOR | 5.08 | 1.2 | 1.74 | 1.62 | 3.88 | 9.46E-04 | 0.0249 | (A) NE (B/D) |
| MIR-200B/20 | 5.89 | 2.02 | 1.38 | 2.08 | 3.87 | 4.00E-03 | 0.0484 | (A) NE (B/D) |
| MIR-200B/20 | 5.89 | 2.02 | 1.38 | 2.08 | 3.87 | 4.00E-03 | 0.0484 | (A) NE (B/D) |
| MIR-200B/3F | 5.89 | 2.02 | 1.38 | 2.08 | 3.87 | 4.00E-03 | 0.0484 | (A) NE (B/D) |
| MIR-200B/5F | 5.89 | 2.02 | 1.38 | 2.08 | 3.87 | 4.00E-03 | 0.0484 | (A) NE (B/D) |
| GIPR | 4.12 | 0.25 | 2.12 | 0.39 | 3.86 | 5.15E-04 | 0.0185 | (A) NE (B/D) |
| SLC44A5 | 3.82 | 0.04 | 2.2 | 0.1 | 3.78 | 8.50E-04 | 0.0236 | (A) NE (B/D) |
| NDRG1 | 5.33 | 1.55 | 0.89 | 1.33 | 3.77 | 2.88E-04 | 0.0142 | (A) NE (B/D) |
| HSPA1B | 12.45 | 8.68 | 0.66 | 0.99 | 3.76 | 3.80E-05 | 5.17E-03 | (A) NE (B/D) |
| C6ORF141 | 4.81 | 1.05 | 1.01 | 1.07 | 3.76 | 3.94E-05 | 5.17E-03 | (A) NE (B/D) |
| PHGR1 | 4.47 | 0.71 | 0.9 | 1.74 | 3.76 | 1.96E-03 | 0.0351 | (A) NE (B/D) |
| TGM3 | 4.41 | 0.66 | 1.22 | 1.24 | 3.75 | 1.38E-04 | 9.45E-03 | (A) NE (B/D) |
| MIR-4648/46 | 4.86 | 1.11 | 0.87 | 1.51 | 3.75 | 8.23E-04 | 0.0231 | (A) NE (B/D) |
| ATP1A3 | 4.84 | 1.09 | 1.35 | 1.01 | 3.74 | 3.89E-05 | 5.17E-03 | (A) NE (B/D) |
| FOXD3 | 5.15 | 1.42 | 2.35 | 0.96 | 3.73 | 1.24E-03 | 0.0278 | (A) NE (B/D) |
| MAGEA12 | 4.07 | 0.34 | 2.32 | 0.84 | 3.72 | 1.10E-03 | 0.0263 | (A) NE (B/D) |
| GPC4 | 4.84 | 1.12 | 1.88 | 1.23 | 3.71 | 4.73E-04 | 0.0176 | (A) NE (B/D) |
| RPS14P3 | 6.05 | 2.37 | 1.34 | 0.81 | 3.68 | 1.72E-05 | 3.38E-03 | (A) NE (B/D) |
| SEZ6L2 | 8.19 | 4.51 | 0.71 | 1.77 | 3.68 | 2.76E-03 | 0.0407 | (A) NE (B/D) |
| FAAH | 5.48 | 1.83 | 0.27 | 1.64 | 3.65 | 2.64E-03 | 0.04 | (A) NE (B/D) |
| PHF19 | 4.46 | 0.82 | 0.77 | 1.22 | 3.63 | 2.29E-04 | 0.0123 | (A) NE (B/D) |
| PI4KAP1 | 7 | 3.37 | 1.36 | 1.32 | 3.63 | 3.10E-04 | 0.0145 | (A) NE (B/D) |
| FBLL1 | 6.45 | 2.84 | 0.6 | 1.64 | 3.6 | 2.20E-03 | 0.0372 | (A) NE (B/D) |
| HSPA1A | 10.98 | 7.39 | 0.66 | 1.06 | 3.59 | 1.03E-04 | 8.25E-03 | (A) NE (B/D) |
| CRACR2B | 9 | 5.45 | 0.65 | 1.72 | 3.55 | 3.00E-03 | 0.0424 | (A) NE (B/D) |
| FAM226A | 6.21 | 2.67 | 1.02 | 1.62 | 3.54 | 1.59E-03 | 0.0317 | (A) NE (B/D) |
| MLPH | 4.6 | 1.09 | 0.59 | 1 | 3.51 | 8.91E-05 | 7.77E-03 | (A) NE (B/D) |
| LINC00951 | 3.92 | 0.41 | 2.49 | 0.75 | 3.51 | 2.64E-03 | 0.04 | (A) NE (B/D) |

| | | | | | | | | |
|---|---|---|---|---|---|---|---|---|
| RASD1 | 4.02 | 0.53 | 1.72 | 1.29 | 3.5 | 6.53E-04 | 0.0204 | (A) NE (B/D) |
| TMC4 | 4.43 | 0.93 | 0.68 | 1.21 | 3.49 | 3.27E-04 | 0.0148 | (A) NE (B/D) |
| TEX101 | 4.46 | 1 | 2.4 | 1.31 | 3.46 | 3.34E-03 | 0.0446 | (A) NE (B/D) |
| LRP4 | 5.76 | 2.31 | 2.09 | 0.95 | 3.45 | 9.87E-04 | 0.0253 | (A) NE (B/D) |
| RNF208 | 7.58 | 4.13 | 1.34 | 1.74 | 3.45 | 2.71E-03 | 0.0404 | (A) NE (B/D) |
| VSTM2L | 5.89 | 2.45 | 1.47 | 1.32 | 3.44 | 5.29E-04 | 0.0185 | (A) NE (B/D) |
| MAPK13 | 4.45 | 1.02 | 0.26 | 1.47 | 3.44 | 2.08E-03 | 0.0362 | (A) NE (B/D) |
| NR1D1 | 3.89 | 0.45 | 0.81 | 1 | 3.43 | 5.67E-05 | 6.31E-03 | (A) NE (B/D) |
| FOXP4 | 7.45 | 4.01 | 0.62 | 1.65 | 3.43 | 2.88E-03 | 0.0417 | (A) NE (B/D) |
| SLC25A6 | 12.3 | 8.87 | 0.76 | 0.91 | 3.42 | 2.58E-05 | 4.22E-03 | (A) NE (B/D) |
| QTRT1 | 4.08 | 0.66 | 0.44 | 1.03 | 3.41 | 2.19E-04 | 0.012 | (A) NE (B/D) |
| AP1M2 | 4 | 0.59 | 0.27 | 1 | 3.41 | 2.90E-04 | 0.0142 | (A) NE (B/D) |
| ZNF414 | 4.47 | 1.09 | 0.91 | 0.97 | 3.38 | 3.96E-05 | 5.17E-03 | (A) NE (B/D) |
| SLC18A1 | 3.72 | 0.34 | 1.86 | 0.61 | 3.38 | 4.45E-04 | 0.0174 | (A) NE (B/D) |
| GAPDHP72 | 6.39 | 3.02 | 1.96 | 1.39 | 3.37 | 1.87E-03 | 0.0347 | (A) NE (B/D) |
| TSPYL2 | 8.63 | 5.28 | 1.28 | 1.19 | 3.35 | 2.63E-04 | 0.0136 | (A) NE (B/D) |
| GUCY1B2 | 4.35 | 1 | 1.3 | 1.38 | 3.35 | 7.39E-04 | 0.022 | (A) NE (B/D) |
| ARHGEF16 | 4.71 | 1.37 | 0.64 | 1 | 3.34 | 1.01E-04 | 8.21E-03 | (A) NE (B/D) |
| PARM1 | 4.11 | 0.78 | 1.26 | 1 | 3.33 | 8.85E-05 | 7.77E-03 | (A) NE (B/D) |
| LINC00342 | 7.33 | 4 | 1.14 | 1.39 | 3.32 | 8.04E-04 | 0.0229 | (A) NE (B/D) |
| USP27X | 7.59 | 4.27 | 1 | 1.47 | 3.32 | 1.28E-03 | 0.0283 | (A) NE (B/D) |
| DMPK | 8.79 | 5.49 | 1.07 | 1.14 | 3.31 | 1.91E-04 | 0.0112 | (A) NE (B/D) |
| SNORA56 | 8.93 | 5.65 | 0.82 | 1.06 | 3.28 | 1.33E-04 | 9.26E-03 | (A) NE (B/D) |
| SCGB1B2P | 5.33 | 2.05 | 1.5 | 1.46 | 3.28 | 1.40E-03 | 0.0299 | (A) NE (B/D) |
| CPLX2 | 4.32 | 1.05 | 0.72 | 1.37 | 3.27 | 1.11E-03 | 0.0263 | (A) NE (B/D) |
| MIR-200A/20 | 4.22 | 0.97 | 1.07 | 1.19 | 3.26 | 3.00E-04 | 0.0142 | (A) NE (B/D) |
| MIR-200A/20 | 4.22 | 0.97 | 1.07 | 1.19 | 3.26 | 3.00E-04 | 0.0142 | (A) NE (B/D) |
| MIR-200A/3P | 4.22 | 0.97 | 1.07 | 1.19 | 3.26 | 3.00E-04 | 0.0142 | (A) NE (B/D) |
| MIR-200A/5P | 4.22 | 0.97 | 1.07 | 1.19 | 3.26 | 3.00E-04 | 0.0142 | (A) NE (B/D) |
| CCDC24 | 6.31 | 3.07 | 0.8 | 1.47 | 3.25 | 1.64E-03 | 0.0323 | (A) NE (B/D) |
| VGF | 3.35 | 0.12 | 1.37 | 0.23 | 3.23 | 8.08E-05 | 7.64E-03 | (A) NE (B/D) |
| LINC00460 | 3.82 | 0.59 | 1.83 | 1.32 | 3.23 | 1.66E-03 | 0.0324 | (A) NE (B/D) |
| WLS | 4.63 | 1.4 | 1.84 | 1.48 | 3.23 | 2.56E-03 | 0.0396 | (A) NE (B/D) |
| ANKZF1 | 3.62 | 0.4 | 0.89 | 0.63 | 3.22 | 1.88E-06 | 8.29E-04 | (A) NE (B/D) |
| ZDHHC8P1 | 4 | 0.78 | 1.38 | 0.93 | 3.22 | 1.22E-04 | 9.03E-03 | (A) NE (B/D) |
| SEMA4B | 8.68 | 5.49 | 0.65 | 1.27 | 3.19 | 8.51E-04 | 0.0236 | (A) NE (B/D) |
| TNFRSF25 | 4.01 | 0.83 | 2.06 | 1.09 | 3.18 | 1.97E-03 | 0.0353 | (A) NE (B/D) |
| RASSF6 | 4.35 | 1.18 | 1.8 | 1.26 | 3.17 | 1.50E-03 | 0.031 | (A) NE (B/D) |
| HOOK2 | 3.44 | 0.29 | 0.52 | 0.71 | 3.15 | 9.04E-06 | 2.28E-03 | (A) NE (B/D) |
| LSM7 | 6.63 | 3.48 | 0.33 | 0.81 | 3.15 | 9.63E-05 | 8.02E-03 | (A) NE (B/D) |
| C1ORF210 | 4.22 | 1.07 | 1.16 | 0.98 | 3.15 | 1.02E-04 | 8.21E-03 | (A) NE (B/D) |
| PRC1-AS1 | 8.12 | 4.97 | 0.98 | 1.58 | 3.15 | 2.81E-03 | 0.0412 | (A) NE (B/D) |
| ATP13A1 | 3.75 | 0.62 | 0.6 | 0.78 | 3.13 | 1.76E-05 | 3.41E-03 | (A) NE (B/D) |
| SLC12A9 | 4.43 | 1.3 | 0.53 | 0.89 | 3.13 | 8.74E-05 | 7.77E-03 | (A) NE (B/D) |
| PKM | 5.11 | 1.98 | 0.57 | 1.51 | 3.13 | 2.92E-03 | 0.0419 | (A) NE (B/D) |
| ARHGEF1 | 6.48 | 3.37 | 1.11 | 0.94 | 3.12 | 7.64E-05 | 7.35E-03 | (A) NE (B/D) |
| SSBP4 | 3.85 | 0.73 | 0.51 | 0.93 | 3.12 | 1.36E-04 | 9.37E-03 | (A) NE (B/D) |
| DHPS | 5.46 | 2.34 | 0.71 | 1.33 | 3.12 | 1.22E-03 | 0.0275 | (A) NE (B/D) |
| FLOT1 | 5.35 | 2.23 | 0.31 | 1.52 | 3.12 | 3.62E-03 | 0.0463 | (A) NE (B/D) |
| SPINT2 | 3.85 | 0.74 | 0.78 | 1.15 | 3.11 | 3.98E-04 | 0.0164 | (A) NE (B/D) |
| MAST2 | 7.43 | 4.31 | 0.71 | 1.17 | 3.11 | 5.06E-04 | 0.0184 | (A) NE (B/D) |
| TM4SF1 | 3.84 | 0.73 | 0.66 | 1.33 | 3.11 | 1.30E-03 | 0.0286 | (A) NE (B/D) |
| RPS4XP11 | 7.67 | 4.56 | 0.93 | 1.53 | 3.11 | 2.46E-03 | 0.0388 | (A) NE (B/D) |
| RPS19P3 | 8.23 | 5.12 | 1.18 | 1.65 | 3.11 | 3.69E-03 | 0.0466 | (A) NE (B/D) |
| BAIAP3 | 3.51 | 0.42 | 0.86 | 1.02 | 3.1 | 1.35E-04 | 9.37E-03 | (A) NE (B/D) |
| MCF2L-AS1 | 5.26 | 2.16 | 0.68 | 1.12 | 3.1 | 3.84E-04 | 0.016 | (A) NE (B/D) |
| ETV1 | 3.88 | 0.8 | 1.65 | 1.06 | 3.08 | 7.29E-04 | 0.0218 | (A) NE (B/D) |
| HES1 | 4.35 | 1.28 | 0.44 | 1.43 | 3.08 | 2.68E-03 | 0.0402 | (A) NE (B/D) |
| SELENOW | 9.27 | 6.2 | 0.92 | 1.4 | 3.07 | 1.53E-03 | 0.0311 | (A) NE (B/D) |
| DOCK9-DT | 5.03 | 1.97 | 1.27 | 1.08 | 3.06 | 3.05E-04 | 0.0144 | (A) NE (B/D) |
| CNKSR1 | 3.78 | 0.72 | 0.73 | 1.18 | 3.06 | 5.60E-04 | 0.019 | (A) NE (B/D) |
| RNA5SP244 | 6.26 | 3.2 | 0.38 | 1.55 | 3.06 | 4.16E-03 | 0.049 | (A) NE (B/D) |
| REC8 | 7.3 | 4.27 | 1.26 | 0.84 | 3.04 | 8.56E-05 | 7.74E-03 | (A) NE (B/D) |
| NDUFA13 | 4.44 | 1.4 | 0.52 | 1.41 | 3.04 | 2.45E-03 | 0.0388 | (A) NE (B/D) |
| CDC20 | 4.22 | 1.2 | 0.97 | 1.01 | 3.03 | 1.49E-04 | 9.80E-03 | (A) NE (B/D) |

| | | | | | | | | |
|---|---|---|---|---|---|---|---|---|
| RAB11B | 7.98 | 4.95 | 0.44 | 1.19 | 3.03 | 1.03E-03 | 0.0257 | (A) NE (B/D) |
| FAM83E | 3.36 | 0.34 | 1.12 | 0.83 | 3.02 | 4.86E-05 | 5.69E-03 | (A) NE (B/D) |
| UNC13A | 3.84 | 0.82 | 0.42 | 0.83 | 3.02 | 9.82E-05 | 8.11E-03 | (A) NE (B/D) |
| TYRP1 | 3.34 | 0.32 | 1.92 | 0.64 | 3.02 | 1.28E-03 | 0.0283 | (A) NE (B/D) |
| PLEKHB1 | 3.52 | 0.5 | 2.15 | 0.8 | 3.02 | 2.81E-03 | 0.0412 | (A) NE (B/D) |
| DCAF15 | 3.61 | 0.6 | 0.37 | 0.89 | 3.01 | 1.95E-04 | 0.0113 | (A) NE (B/D) |
| CDC20P1 | 6.05 | 3.04 | 1.11 | 1.13 | 3.01 | 3.81E-04 | 0.0159 | (A) NE (B/D) |
| HOXD4 | 3.66 | 0.66 | 1.53 | 0.98 | 3.01 | 4.71E-04 | 0.0176 | (A) NE (B/D) |
| SPOCK1 | 7.01 | 4 | 1.03 | 1.31 | 3.01 | 1.06E-03 | 0.026 | (A) NE (B/D) |
| TMED1 | 4.68 | 1.67 | 0.53 | 1.52 | 3.01 | 3.78E-03 | 0.0473 | (A) NE (B/D) |
| CDCA8 | 4.45 | 1.45 | 1.24 | 1.19 | 3 | 6.02E-04 | 0.0195 | (A) NE (B/D) |
| HOXD13 | 4.09 | 1.09 | 1.87 | 1.15 | 3 | 2.05E-03 | 0.036 | (A) NE (B/D) |
| DDX39A | 4.37 | 1.37 | 0.38 | 1.42 | 3 | 3.04E-03 | 0.0426 | (A) NE (B/D) |
| CEACAM19 | 4.84 | 1.84 | 1.05 | 1.56 | 3 | 3.27E-03 | 0.0441 | (A) NE (B/D) |
| ITPR3 | 8.22 | 5.22 | 0.39 | 1.5 | 3 | 3.83E-03 | 0.0476 | (A) NE (B/D) |
| SNRPD2 | 6.13 | 3.14 | 0.36 | 1.02 | 2.99 | 5.29E-04 | 0.0185 | (A) NE (B/D) |
| SMIM3 | 7.28 | 4.3 | 0.71 | 1.44 | 2.99 | 2.55E-03 | 0.0396 | (A) NE (B/D) |
| SLC35D3 | 3.11 | 0.13 | 1.42 | 0.31 | 2.98 | 1.75E-04 | 0.0106 | (A) NE (B/D) |
| BEST4 | 4.55 | 1.57 | 1.23 | 1.01 | 2.98 | 2.27E-04 | 0.0123 | (A) NE (B/D) |
| LINC00346 | 3.22 | 0.25 | 1.38 | 0.39 | 2.97 | 1.27E-04 | 9.14E-03 | (A) NE (B/D) |
| PVRIG2P | 5.91 | 2.94 | 1.18 | 1.12 | 2.97 | 4.24E-04 | 0.017 | (A) NE (B/D) |
| FOXD2-AS1 | 4.73 | 1.79 | 1.62 | 0.82 | 2.94 | 5.32E-04 | 0.0185 | (A) NE (B/D) |
| PPM1N | 6.01 | 3.06 | 0.27 | 1.02 | 2.94 | 6.57E-04 | 0.0205 | (A) NE (B/D) |
| PPFIA3 | 2.95 | 0.02 | 0.62 | 0.06 | 2.93 | 4.77E-07 | 4.22E-04 | (A) NE (B/D) |
| BAALC | 3.07 | 0.14 | 1.93 | 0.34 | 2.93 | 1.72E-03 | 0.0329 | (A) NE (B/D) |
| RPL13AP20 | 4.24 | 1.32 | 1.85 | 1.29 | 2.92 | 3.26E-03 | 0.0441 | (A) NE (B/D) |
| ADPRHL1 | 6.03 | 3.12 | 0.95 | 0.74 | 2.91 | 1.95E-05 | 3.50E-03 | (A) NE (B/D) |
| KLHL35 | 4.3 | 1.39 | 1.09 | 0.88 | 2.91 | 8.58E-05 | 7.74E-03 | (A) NE (B/D) |
| LINC01023 | 4.16 | 1.25 | 1.32 | 0.71 | 2.91 | 1.07E-04 | 8.42E-03 | (A) NE (B/D) |
| SHC3 | 3.62 | 0.72 | 1.78 | 0.67 | 2.9 | 1.01E-03 | 0.0255 | (A) NE (B/D) |
| ATP5F1D | 4.72 | 1.82 | 0.69 | 1.24 | 2.9 | 1.20E-03 | 0.0274 | (A) NE (B/D) |
| CRIPAK | 6.73 | 3.83 | 0.88 | 1.5 | 2.9 | 3.26E-03 | 0.0441 | (A) NE (B/D) |
| GYG1 | 3.97 | 1.08 | 1.3 | 1.21 | 2.89 | 9.58E-04 | 0.025 | (A) NE (B/D) |
| NR4A2 | 6.96 | 4.06 | 1.55 | 1.12 | 2.89 | 1.05E-03 | 0.0259 | (A) NE (B/D) |
| TUBB7P | 3.91 | 1.02 | 0.46 | 1.31 | 2.89 | 2.19E-03 | 0.0372 | (A) NE (B/D) |
| NCDN | 3.46 | 0.59 | 0.65 | 0.49 | 2.88 | 2.89E-07 | 4.22E-04 | (A) NE (B/D) |
| GPR108 | 3.49 | 0.61 | 0.45 | 0.95 | 2.88 | 3.14E-04 | 0.0145 | (A) NE (B/D) |
| YJEFN3 | 9.91 | 7.03 | 0.62 | 1.52 | 2.88 | 4.29E-03 | 0.0497 | (A) NE (B/D) |
| NOP53 | 6.82 | 3.95 | 0.31 | 0.66 | 2.87 | 3.83E-05 | 5.17E-03 | (A) NE (B/D) |
| FKBP8 | 4.24 | 1.37 | 0.45 | 1.33 | 2.87 | 2.49E-03 | 0.0391 | (A) NE (B/D) |
| NFKBID | 4.67 | 1.81 | 0.89 | 0.86 | 2.86 | 6.18E-05 | 6.72E-03 | (A) NE (B/D) |
| NRGN | 7 | 4.14 | 1.63 | 1.12 | 2.86 | 1.47E-03 | 0.0307 | (A) NE (B/D) |
| NQO1 | 4.31 | 1.45 | 1.2 | 1.36 | 2.86 | 1.94E-03 | 0.0351 | (A) NE (B/D) |
| KIF2C | 6.69 | 3.83 | 1.2 | 1.39 | 2.86 | 2.24E-03 | 0.0375 | (A) NE (B/D) |
| ULK3 | 8.75 | 5.89 | 0.76 | 1.51 | 2.86 | 3.89E-03 | 0.048 | (A) NE (B/D) |
| WDR83OS | 4.39 | 1.53 | 0.5 | 1.47 | 2.86 | 4.04E-03 | 0.0486 | (A) NE (B/D) |
| CD320 | 3.69 | 0.84 | 0.57 | 1.06 | 2.85 | 5.27E-04 | 0.0185 | (A) NE (B/D) |
| CTBP2P8 | 4.1 | 1.26 | 0.8 | 1.08 | 2.84 | 4.47E-04 | 0.0174 | (A) NE (B/D) |
| ZGLP1 | 3.89 | 1.05 | 0.95 | 1.13 | 2.84 | 5.80E-04 | 0.0193 | (A) NE (B/D) |
| RPL18AP3 | 13.68 | 10.85 | 0.78 | 1.03 | 2.83 | 3.21E-04 | 0.0146 | (A) NE (B/D) |
| EMC9 | 6.34 | 3.5 | 0.75 | 1.25 | 2.83 | 1.38E-03 | 0.0296 | (A) NE (B/D) |
| ACOT11 | 5.13 | 2.31 | 1.8 | 0.89 | 2.83 | 1.61E-03 | 0.0319 | (A) NE (B/D) |
| GNA14 | 6.55 | 3.72 | 0.77 | 0.94 | 2.82 | 1.57E-04 | 0.0101 | (A) NE (B/D) |
| HSD11B1L | 5.24 | 2.43 | 0.51 | 0.95 | 2.82 | 2.96E-04 | 0.0142 | (A) NE (B/D) |
| SDR39U1 | 3.94 | 1.12 | 0.81 | 1.52 | 2.82 | 4.28E-03 | 0.0496 | (A) NE (B/D) |
| PCDHB14 | 3.4 | 0.59 | 1.4 | 0.82 | 2.81 | 2.97E-04 | 0.0142 | (A) NE (B/D) |
| SNAI1P1 | 4.25 | 1.45 | 0.98 | 1.18 | 2.81 | 8.52E-04 | 0.0236 | (A) NE (B/D) |
| KLC3 | 6.54 | 3.73 | 1.58 | 1.41 | 2.81 | 3.78E-03 | 0.0473 | (A) NE (B/D) |
| MIR-637/637 | 3.27 | 0.47 | 0.46 | 0.47 | 2.8 | 2.60E-07 | 4.05E-04 | (A) NE (B/D) |
| RPL37AP1 | 8 | 5.2 | 1.53 | 1.04 | 2.8 | 1.00E-03 | 0.0254 | (A) NE (B/D) |
| PLEKHJ1 | 3.1 | 0.31 | 0.36 | 0.64 | 2.79 | 2.27E-05 | 3.95E-03 | (A) NE (B/D) |
| SMG9 | 3.53 | 0.74 | 0.87 | 0.8 | 2.79 | 4.02E-05 | 5.17E-03 | (A) NE (B/D) |
| RNF31 | 3.45 | 0.67 | 0.8 | 1.09 | 2.79 | 5.22E-04 | 0.0185 | (A) NE (B/D) |
| ATP6AP1 | 4.18 | 1.39 | 0.63 | 1.07 | 2.79 | 5.96E-04 | 0.0194 | (A) NE (B/D) |
| PILRB | 6.3 | 3.51 | 1.07 | 1.19 | 2.79 | 9.15E-04 | 0.0245 | (A) NE (B/D) |

| | | | | | | | | |
|---|---|---|---|---|---|---|---|---|
| MIR-1914/19 | 3.51 | 0.73 | 0.53 | 1.16 | 2.79 | 1.21E-03 | 0.0274 | (A) NE (B/D) |
| MIR-1914/19 | 3.51 | 0.73 | 0.53 | 1.16 | 2.79 | 1.21E-03 | 0.0274 | (A) NE (B/D) |
| MIR-1914/3F | 3.51 | 0.73 | 0.53 | 1.16 | 2.79 | 1.21E-03 | 0.0274 | (A) NE (B/D) |
| MIR-1914/5F | 3.51 | 0.73 | 0.53 | 1.16 | 2.79 | 1.21E-03 | 0.0274 | (A) NE (B/D) |
| RGL2 | 6.04 | 3.25 | 0.34 | 1.11 | 2.79 | 1.26E-03 | 0.0281 | (A) NE (B/D) |
| RPS4XP7 | 4.11 | 1.33 | 1.42 | 1.15 | 2.78 | 1.20E-03 | 0.0274 | (A) NE (B/D) |
| VAMP2 | 6.66 | 3.88 | 0.83 | 1.22 | 2.78 | 1.22E-03 | 0.0275 | (A) NE (B/D) |
| MAP6 | 3.34 | 0.56 | 1.22 | 1.37 | 2.78 | 2.44E-03 | 0.0387 | (A) NE (B/D) |
| TBC1D8 | 5.67 | 2.89 | 0.39 | 1.31 | 2.78 | 2.87E-03 | 0.0417 | (A) NE (B/D) |
| BRAT1 | 3.55 | 0.78 | 0.34 | 0.57 | 2.77 | 1.04E-05 | 2.43E-03 | (A) NE (B/D) |
| OBSL1 | 3.19 | 0.41 | 0.46 | 0.72 | 2.77 | 4.02E-05 | 5.17E-03 | (A) NE (B/D) |
| WFDC2 | 2.8 | 0.03 | 1.21 | 0.08 | 2.77 | 1.26E-04 | 9.14E-03 | (A) NE (B/D) |
| HNRNPA1P27 | 4.41 | 1.63 | 0.71 | 1.3 | 2.77 | 2.03E-03 | 0.0358 | (A) NE (B/D) |
| DPP7 | 9.06 | 6.3 | 0.62 | 0.81 | 2.76 | 6.43E-05 | 6.81E-03 | (A) NE (B/D) |
| ATG4D | 5.67 | 2.91 | 0.56 | 0.99 | 2.76 | 4.22E-04 | 0.0169 | (A) NE (B/D) |
| COPE | 3.37 | 0.61 | 0.45 | 0.99 | 2.76 | 5.29E-04 | 0.0185 | (A) NE (B/D) |
| MAP2K7 | 7.64 | 4.88 | 0.6 | 1.08 | 2.76 | 7.19E-04 | 0.0217 | (A) NE (B/D) |
| CC2D1A | 3.13 | 0.38 | 0.94 | 0.68 | 2.75 | 1.84E-05 | 3.49E-03 | (A) NE (B/D) |
| SLC22A23 | 5.79 | 3.04 | 1.11 | 1.13 | 2.75 | 7.46E-04 | 0.022 | (A) NE (B/D) |
| STRN4 | 4.6 | 1.86 | 0.37 | 0.83 | 2.74 | 2.00E-04 | 0.0113 | (A) NE (B/D) |
| TLE5 | 3.49 | 0.75 | 0.18 | 0.94 | 2.74 | 7.42E-04 | 0.022 | (A) NE (B/D) |
| LPAR2 | 8.04 | 5.29 | 0.31 | 1.27 | 2.74 | 2.81E-03 | 0.0412 | (A) NE (B/D) |
| RPL18A | 3.68 | 0.95 | 0.76 | 0.93 | 2.73 | 1.84E-04 | 0.0109 | (A) NE (B/D) |
| GYG1P3 | 2.94 | 0.22 | 0.64 | 0.34 | 2.72 | 1.02E-07 | 2.56E-04 | (A) NE (B/D) |
| SNORD38A | 3.54 | 0.83 | 0.88 | 0.89 | 2.72 | 1.21E-04 | 9.03E-03 | (A) NE (B/D) |
| SPAG4 | 6.73 | 4.01 | 0.57 | 1.08 | 2.72 | 8.21E-04 | 0.0231 | (A) NE (B/D) |
| FAM183A | 3.34 | 0.62 | 1.45 | 1.15 | 2.72 | 1.53E-03 | 0.031 | (A) NE (B/D) |
| BACE2 | 3.05 | 0.35 | 0.63 | 0.47 | 2.7 | 4.05E-07 | 4.22E-04 | (A) NE (B/D) |
| ZNF165 | 6.75 | 4.05 | 1.09 | 1.09 | 2.7 | 6.81E-04 | 0.0209 | (A) NE (B/D) |
| SP9 | 4.22 | 1.51 | 1.66 | 1.03 | 2.7 | 1.90E-03 | 0.0348 | (A) NE (B/D) |
| STK11IP | 2.95 | 0.26 | 0.72 | 0.45 | 2.69 | 6.52E-07 | 4.92E-04 | (A) NE (B/D) |
| CCDC124 | 4.28 | 1.59 | 0.43 | 0.95 | 2.69 | 4.88E-04 | 0.018 | (A) NE (B/D) |
| MIR-1909/19 | 4.85 | 2.16 | 0.31 | 1 | 2.69 | 8.85E-04 | 0.0241 | (A) NE (B/D) |
| MIR-1909/19 | 4.85 | 2.16 | 0.31 | 1 | 2.69 | 8.85E-04 | 0.0241 | (A) NE (B/D) |
| MIR-1909/3F | 4.85 | 2.16 | 0.31 | 1 | 2.69 | 8.85E-04 | 0.0241 | (A) NE (B/D) |
| MIR-1909/5F | 4.85 | 2.16 | 0.31 | 1 | 2.69 | 8.85E-04 | 0.0241 | (A) NE (B/D) |
| HJURP | 5.38 | 2.69 | 0.83 | 1.27 | 2.69 | 1.86E-03 | 0.0345 | (A) NE (B/D) |
| MAFF | 8.23 | 5.55 | 0.47 | 0.76 | 2.68 | 7.43E-05 | 7.23E-03 | (A) NE (B/D) |
| PRKD2 | 7.7 | 5.02 | 0.32 | 0.75 | 2.68 | 1.41E-04 | 9.54E-03 | (A) NE (B/D) |
| COLGALT1 | 4.62 | 1.94 | 0.25 | 1.08 | 2.68 | 1.52E-03 | 0.031 | (A) NE (B/D) |
| PTOV1 | 5.44 | 2.76 | 0.77 | 1.35 | 2.68 | 2.88E-03 | 0.0417 | (A) NE (B/D) |
| BORCS8-MEF | 4.41 | 1.73 | 0.35 | 1.28 | 2.68 | 3.11E-03 | 0.0429 | (A) NE (B/D) |
| PCSK1N | 2.85 | 0.18 | 1.5 | 0.43 | 2.67 | 5.29E-04 | 0.0185 | (A) NE (B/D) |
| BABAM1 | 9.58 | 6.9 | 0.44 | 1.1 | 2.67 | 1.25E-03 | 0.0281 | (A) NE (B/D) |
| DDX41 | 3.73 | 1.06 | 0.28 | 1.07 | 2.67 | 1.39E-03 | 0.0298 | (A) NE (B/D) |
| AP1G2 | 3.11 | 0.45 | 0.98 | 0.88 | 2.66 | 1.59E-04 | 0.0102 | (A) NE (B/D) |
| FBXW9 | 5.51 | 2.85 | 0.65 | 1 | 2.66 | 4.88E-04 | 0.018 | (A) NE (B/D) |
| MIR-1181/11 | 9.53 | 6.86 | 0.48 | 1.08 | 2.66 | 1.06E-03 | 0.026 | (A) NE (B/D) |
| PRAF2 | 8.64 | 5.98 | 0.51 | 1.32 | 2.66 | 3.34E-03 | 0.0446 | (A) NE (B/D) |
| ABHD17AP1 | 3.59 | 0.94 | 0.5 | 0.7 | 2.65 | 3.48E-05 | 4.95E-03 | (A) NE (B/D) |
| PITX1 | 3 | 0.35 | 1.11 | 0.38 | 2.65 | 4.66E-05 | 5.63E-03 | (A) NE (B/D) |
| GRAMD1A | 5.29 | 2.64 | 0.3 | 0.69 | 2.65 | 8.50E-05 | 7.73E-03 | (A) NE (B/D) |
| CCDC151 | 3.15 | 0.5 | 0.7 | 0.84 | 2.65 | 9.97E-05 | 8.15E-03 | (A) NE (B/D) |
| BPIFB4 | 3.47 | 0.82 | 1.13 | 0.94 | 2.65 | 3.30E-04 | 0.0148 | (A) NE (B/D) |
| TUBB8P8 | 3.31 | 0.65 | 0.48 | 1 | 2.65 | 6.59E-04 | 0.0205 | (A) NE (B/D) |
| FLII | 8.61 | 5.96 | 0.48 | 1.11 | 2.65 | 1.28E-03 | 0.0283 | (A) NE (B/D) |
| ICA1 | 3.62 | 0.97 | 0.6 | 1.25 | 2.65 | 2.28E-03 | 0.0376 | (A) NE (B/D) |
| RPL26P37 | 3.11 | 0.47 | 1.11 | 0.42 | 2.64 | 4.54E-05 | 5.58E-03 | (A) NE (B/D) |
| L3MBTL1 | 4.88 | 2.25 | 1.04 | 0.73 | 2.64 | 6.79E-05 | 6.91E-03 | (A) NE (B/D) |
| IFNGR2 | 4.27 | 1.63 | 0.45 | 1.19 | 2.64 | 2.07E-03 | 0.0361 | (A) NE (B/D) |
| EEF1A1P11 | 5.36 | 2.72 | 1.44 | 1.24 | 2.64 | 2.63E-03 | 0.04 | (A) NE (B/D) |
| HMG20B | 4.1 | 1.47 | 0.34 | 1.33 | 2.64 | 4.14E-03 | 0.049 | (A) NE (B/D) |
| ABHD8 | 3.2 | 0.57 | 0.48 | 0.65 | 2.63 | 1.93E-05 | 3.50E-03 | (A) NE (B/D) |
| SIX2 | 3.29 | 0.66 | 0.74 | 1.19 | 2.63 | 1.55E-03 | 0.0313 | (A) NE (B/D) |
| POMGNT1 | 7.11 | 4.48 | 0.6 | 1.26 | 2.63 | 2.42E-03 | 0.0385 | (A) NE (B/D) |

| | | | | | | | | |
|---|---|---|---|---|---|---|---|---|
| FN3K | 3.05 | 0.43 | 0.84 | 0.74 | 2.62 | 3.95E-05 | 5.17E-03 | (A) NE (B/D) |
| IRS2 | 7.83 | 5.21 | 0.62 | 0.98 | 2.62 | 4.70E-04 | 0.0176 | (A) NE (B/D) |
| HSPA1L | 4.83 | 2.22 | 0.73 | 0.67 | 2.61 | 1.44E-05 | 2.98E-03 | (A) NE (B/D) |
| MAU2 | 4.65 | 2.04 | 0.8 | 0.7 | 2.61 | 2.50E-05 | 4.17E-03 | (A) NE (B/D) |
| RPS15 | 6.92 | 4.31 | 0.49 | 0.67 | 2.61 | 2.64E-05 | 4.26E-03 | (A) NE (B/D) |
| C6ORF136 | 2.98 | 0.37 | 0.27 | 0.64 | 2.61 | 6.66E-05 | 6.86E-03 | (A) NE (B/D) |
| STAG3L5P-P\ | 4.42 | 1.81 | 1.1 | 1.04 | 2.61 | 6.42E-04 | 0.0202 | (A) NE (B/D) |
| ATG4B | 7.49 | 4.89 | 0.54 | 1 | 2.6 | 6.53E-04 | 0.0204 | (A) NE (B/D) |
| TRAPPC5 | 5.08 | 2.48 | 1.21 | 1.08 | 2.6 | 1.01E-03 | 0.0254 | (A) NE (B/D) |
| SERPINB6 | 3.62 | 1.02 | 0.59 | 1.36 | 2.6 | 4.03E-03 | 0.0486 | (A) NE (B/D) |
| CAPS | 3.03 | 0.44 | 0.92 | 0.64 | 2.59 | 2.45E-05 | 4.14E-03 | (A) NE (B/D) |
| SNRNP70 | 6.16 | 3.57 | 1.06 | 1.07 | 2.59 | 8.08E-04 | 0.0229 | (A) NE (B/D) |
| ARTN | 2.83 | 0.25 | 0.73 | 0.28 | 2.58 | 1.04E-06 | 6.56E-04 | (A) NE (B/D) |
| TTC22 | 3.01 | 0.43 | 0.89 | 0.46 | 2.58 | 7.09E-06 | 2.02E-03 | (A) NE (B/D) |
| UBE2C | 3.3 | 0.72 | 0.93 | 0.78 | 2.58 | 8.26E-05 | 7.64E-03 | (A) NE (B/D) |
| SOBP | 2.72 | 0.14 | 1.3 | 0.35 | 2.58 | 2.48E-04 | 0.0131 | (A) NE (B/D) |
| GPR153 | 2.71 | 0.13 | 1.38 | 0.15 | 2.58 | 4.80E-04 | 0.0178 | (A) NE (B/D) |
| RPL18 | 4.83 | 2.25 | 0.54 | 0.98 | 2.58 | 6.03E-04 | 0.0195 | (A) NE (B/D) |
| LINC00265 | 5.63 | 3.05 | 0.62 | 1.12 | 2.58 | 1.31E-03 | 0.0287 | (A) NE (B/D) |
| ATP5MC1P4 | 4.21 | 1.64 | 1.08 | 1.2 | 2.58 | 1.70E-03 | 0.0328 | (A) NE (B/D) |
| SNHG5 | 4.45 | 1.88 | 0.42 | 0.81 | 2.57 | 1.96E-04 | 0.0113 | (A) NE (B/D) |
| PLEKHG5 | 3.19 | 0.62 | 1.6 | 0.78 | 2.57 | 1.32E-03 | 0.0289 | (A) NE (B/D) |
| SUGP1 | 2.82 | 0.26 | 0.93 | 0.63 | 2.56 | 2.60E-05 | 4.22E-03 | (A) NE (B/D) |
| RFXANK | 4.69 | 2.14 | 0.31 | 0.92 | 2.56 | 7.07E-04 | 0.0215 | (A) NE (B/D) |
| CSNK1D | 3 | 0.44 | 0.37 | 0.96 | 2.56 | 7.68E-04 | 0.0223 | (A) NE (B/D) |
| DNAJB2 | 8.64 | 6.08 | 0.58 | 1.02 | 2.56 | 7.81E-04 | 0.0225 | (A) NE (B/D) |
| ENTPD6 | 3.74 | 1.17 | 0.42 | 1 | 2.56 | 8.98E-04 | 0.0243 | (A) NE (B/D) |
| GUCA1B | 3.31 | 0.75 | 1.23 | 1.13 | 2.56 | 1.49E-03 | 0.0309 | (A) NE (B/D) |
| PKP3 | 3.78 | 1.22 | 1.42 | 1.19 | 2.56 | 2.59E-03 | 0.0398 | (A) NE (B/D) |
| PPP2R3B | 3.3 | 0.75 | 0.72 | 0.52 | 2.55 | 2.71E-06 | 1.04E-03 | (A) NE (B/D) |
| EIF3G | 4.19 | 1.63 | 0.49 | 0.87 | 2.55 | 2.94E-04 | 0.0142 | (A) NE (B/D) |
| PHF1 | 6.98 | 4.43 | 0.66 | 0.93 | 2.55 | 3.59E-04 | 0.0154 | (A) NE (B/D) |
| TMEM63C | 4.21 | 1.66 | 1 | 1.13 | 2.55 | 1.22E-03 | 0.0275 | (A) NE (B/D) |
| DMAP1 | 7.88 | 5.33 | 0.82 | 1.33 | 2.55 | 3.45E-03 | 0.0453 | (A) NE (B/D) |
| RPL36AP26 | 3.23 | 0.68 | 1.57 | 1.08 | 2.54 | 2.55E-03 | 0.0396 | (A) NE (B/D) |
| LARGE2 | 3.19 | 0.65 | 0.77 | 1.26 | 2.54 | 2.56E-03 | 0.0396 | (A) NE (B/D) |
| HNRNPA1P4 | 4.47 | 1.93 | 0.44 | 1.24 | 2.54 | 3.10E-03 | 0.0429 | (A) NE (B/D) |
| WDR83 | 3.05 | 0.52 | 0.63 | 0.85 | 2.53 | 1.88E-04 | 0.0111 | (A) NE (B/D) |
| GPS2 | 3.33 | 0.8 | 0.8 | 0.89 | 2.53 | 2.24E-04 | 0.0122 | (A) NE (B/D) |
| HID1 | 2.85 | 0.33 | 0.35 | 0.8 | 2.53 | 2.69E-04 | 0.0137 | (A) NE (B/D) |
| NDN | 7.67 | 5.14 | 1.28 | 0.74 | 2.53 | 3.28E-04 | 0.0148 | (A) NE (B/D) |
| ATP6V1F | 3.29 | 0.75 | 0.45 | 1.02 | 2.53 | 1.03E-03 | 0.0256 | (A) NE (B/D) |
| SLC37A1 | 6.93 | 4.4 | 0.3 | 0.97 | 2.53 | 1.03E-03 | 0.0256 | (A) NE (B/D) |
| UBE2S | 3.85 | 1.32 | 0.75 | 1.15 | 2.53 | 1.49E-03 | 0.0309 | (A) NE (B/D) |
| SCAMP5 | 2.97 | 0.45 | 0.44 | 1.09 | 2.53 | 1.61E-03 | 0.0319 | (A) NE (B/D) |
| SH3BP4 | 3.42 | 0.88 | 0.98 | 1.19 | 2.53 | 1.71E-03 | 0.0329 | (A) NE (B/D) |
| PLP2 | 8.22 | 5.69 | 0.59 | 1.16 | 2.53 | 1.90E-03 | 0.0348 | (A) NE (B/D) |
| MFSD10 | 3.03 | 0.5 | 0.78 | 1.22 | 2.53 | 2.20E-03 | 0.0372 | (A) NE (B/D) |
| RSPH1 | 3 | 0.48 | 0.85 | 0.69 | 2.52 | 3.51E-05 | 4.97E-03 | (A) NE (B/D) |
| ALG1L3P | 2.57 | 0.05 | 1.16 | 0.12 | 2.52 | 1.72E-04 | 0.0106 | (A) NE (B/D) |
| RNF207 | 4.14 | 1.62 | 0.47 | 1.23 | 2.52 | 3.09E-03 | 0.0429 | (A) NE (B/D) |
| SOX21 | 2.57 | 0.06 | 1.55 | 0.14 | 2.51 | 1.20E-03 | 0.0274 | (A) NE (B/D) |
| TMEM198 | 4.3 | 1.79 | 0.34 | 1 | 2.51 | 1.21E-03 | 0.0274 | (A) NE (B/D) |
| POLR2I | 3.87 | 1.35 | 0.85 | 1.14 | 2.51 | 1.38E-03 | 0.0296 | (A) NE (B/D) |
| ASNA1 | 8.83 | 6.32 | 0.37 | 1.19 | 2.51 | 2.94E-03 | 0.042 | (A) NE (B/D) |
| ATF6B | 5.36 | 2.85 | 0.3 | 1.21 | 2.51 | 3.33E-03 | 0.0445 | (A) NE (B/D) |
| C9ORF152 | 2.74 | 0.25 | 1.01 | 0.56 | 2.5 | 3.93E-05 | 5.17E-03 | (A) NE (B/D) |
| EIF4HP2 | 5.53 | 3.03 | 0.8 | 0.76 | 2.5 | 6.85E-05 | 6.91E-03 | (A) NE (B/D) |
| APOD | 3.09 | 0.59 | 0.91 | 0.9 | 2.5 | 2.86E-04 | 0.0142 | (A) NE (B/D) |
| DLL1 | 3.09 | 0.59 | 1.39 | 0.63 | 2.5 | 5.12E-04 | 0.0185 | (A) NE (B/D) |
| ZNF358 | 7.19 | 4.69 | 0.62 | 1.01 | 2.5 | 7.48E-04 | 0.022 | (A) NE (B/D) |
| HDAC10 | 6.46 | 3.96 | 0.34 | 0.93 | 2.5 | 7.85E-04 | 0.0225 | (A) NE (B/D) |
| HOXC5 | 3.9 | 1.39 | 0.87 | 1.1 | 2.5 | 1.13E-03 | 0.0267 | (A) NE (B/D) |
| TECRP1 | 8.17 | 5.67 | 0.42 | 1.03 | 2.5 | 1.22E-03 | 0.0275 | (A) NE (B/D) |
| FAM50A | 10.05 | 7.55 | 0.55 | 1.17 | 2.5 | 2.22E-03 | 0.0374 | (A) NE (B/D) |

| | | | | | | | | |
|---|---|---|---|---|---|---|---|---|
| RPL34P31 | 6.72 | 4.23 | 1.13 | 1.34 | 2.5 | 4.06E-03 | 0.0487 | (A) NE (B/D) |
| CHST2 | 4.15 | 1.66 | 1.33 | 1.11 | 2.49 | 1.98E-03 | 0.0354 | (A) NE (B/D) |
| UBA52 | 7.35 | 4.86 | 0.41 | 1.18 | 2.49 | 2.72E-03 | 0.0404 | (A) NE (B/D) |
| TRMT1 | 7.76 | 5.28 | 0.52 | 1.24 | 2.49 | 3.22E-03 | 0.0438 | (A) NE (B/D) |
| GMPPA | 3.49 | 1 | 0.23 | 1.19 | 2.49 | 3.43E-03 | 0.0452 | (A) NE (B/D) |
| VWA7 | 4.64 | 2.16 | 0.59 | 0.79 | 2.48 | 1.21E-04 | 9.03E-03 | (A) NE (B/D) |
| TEKT4P2 | 8.21 | 5.72 | 0.66 | 0.94 | 2.48 | 4.37E-04 | 0.0173 | (A) NE (B/D) |
| CDK18 | 3.69 | 1.21 | 0.99 | 1.06 | 2.48 | 9.86E-04 | 0.0253 | (A) NE (B/D) |
| S100A13 | 3.15 | 0.67 | 1.18 | 1.03 | 2.48 | 1.04E-03 | 0.0258 | (A) NE (B/D) |
| SLC38A11 | 2.73 | 0.24 | 1.6 | 0.41 | 2.48 | 1.42E-03 | 0.03 | (A) NE (B/D) |
| MIR-939/939 | 9.11 | 6.62 | 0.37 | 1.04 | 2.48 | 1.51E-03 | 0.031 | (A) NE (B/D) |
| IGBP1 | 4.02 | 1.54 | 0.36 | 1.1 | 2.48 | 2.05E-03 | 0.036 | (A) NE (B/D) |
| NAT9 | 3.77 | 1.29 | 0.4 | 1.14 | 2.48 | 2.32E-03 | 0.038 | (A) NE (B/D) |
| FAM13A-AS1 | 7.25 | 4.77 | 0.75 | 1.33 | 2.48 | 4.05E-03 | 0.0487 | (A) NE (B/D) |
| TECR | 3.27 | 0.79 | 0.4 | 0.87 | 2.47 | 4.63E-04 | 0.0175 | (A) NE (B/D) |
| MINK1 | 3.37 | 0.9 | 0.75 | 0.96 | 2.47 | 4.81E-04 | 0.0178 | (A) NE (B/D) |
| LFNG | 2.74 | 0.27 | 0.57 | 0.67 | 2.46 | 2.92E-05 | 4.49E-03 | (A) NE (B/D) |
| USP11 | 3.06 | 0.6 | 1 | 0.93 | 2.46 | 4.54E-04 | 0.0175 | (A) NE (B/D) |
| ZNF618 | 3.41 | 0.95 | 0.54 | 0.91 | 2.46 | 4.56E-04 | 0.0175 | (A) NE (B/D) |
| RPL28 | 3.67 | 1.21 | 0.42 | 0.93 | 2.46 | 7.26E-04 | 0.0218 | (A) NE (B/D) |
| BRD8 | 3.24 | 0.78 | 1.14 | 1.17 | 2.46 | 2.16E-03 | 0.0369 | (A) NE (B/D) |
| SERGEF | 9 | 6.53 | 0.62 | 1.23 | 2.46 | 2.94E-03 | 0.042 | (A) NE (B/D) |
| TMEM240 | 4.12 | 1.66 | 1.26 | 1.24 | 2.46 | 3.24E-03 | 0.044 | (A) NE (B/D) |
| RABAC1 | 2.84 | 0.39 | 0.47 | 0.44 | 2.45 | 3.76E-07 | 4.22E-04 | (A) NE (B/D) |
| GATAD1 | 3.91 | 1.46 | 0.98 | 0.87 | 2.45 | 2.85E-04 | 0.0142 | (A) NE (B/D) |
| PCDHB10 | 2.63 | 0.18 | 1.36 | 0.45 | 2.45 | 4.65E-04 | 0.0175 | (A) NE (B/D) |
| RPS19P1 | 5.89 | 3.44 | 1.11 | 0.93 | 2.45 | 5.69E-04 | 0.0191 | (A) NE (B/D) |
| CPSF1 | 8.74 | 6.29 | 0.56 | 1.03 | 2.45 | 1.09E-03 | 0.0262 | (A) NE (B/D) |
| GIGYF1 | 4.33 | 1.88 | 0.54 | 1.27 | 2.45 | 3.86E-03 | 0.0478 | (A) NE (B/D) |
| EEF1A1P13 | 10.06 | 7.62 | 1.42 | 1.24 | 2.45 | 4.24E-03 | 0.0494 | (A) NE (B/D) |
| CDC37 | 5.86 | 3.42 | 0.16 | 1.05 | 2.44 | 2.17E-03 | 0.037 | (A) NE (B/D) |
| RCCD1 | 7.68 | 5.24 | 0.56 | 1.14 | 2.44 | 2.21E-03 | 0.0372 | (A) NE (B/D) |
| SNHG6 | 9.82 | 7.38 | 0.27 | 1.1 | 2.44 | 2.45E-03 | 0.0388 | (A) NE (B/D) |
| RNPEPL1 | 3.71 | 1.27 | 0.12 | 1.09 | 2.44 | 2.70E-03 | 0.0404 | (A) NE (B/D) |
| C6ORF226 | 5.63 | 3.19 | 0.69 | 1.25 | 2.44 | 3.21E-03 | 0.0438 | (A) NE (B/D) |
| AMIGO2 | 3.35 | 0.91 | 1.45 | 1.21 | 2.44 | 3.99E-03 | 0.0484 | (A) NE (B/D) |
| RIPK4 | 2.71 | 0.27 | 0.47 | 0.44 | 2.43 | 5.09E-07 | 4.22E-04 | (A) NE (B/D) |
| SYMPK | 5.89 | 3.45 | 0.47 | 0.55 | 2.43 | 6.47E-06 | 1.89E-03 | (A) NE (B/D) |
| MIR-10B/10E | 3.9 | 1.47 | 0.99 | 0.79 | 2.43 | 1.70E-04 | 0.0106 | (A) NE (B/D) |
| MIR-10B/10E | 3.9 | 1.47 | 0.99 | 0.79 | 2.43 | 1.70E-04 | 0.0106 | (A) NE (B/D) |
| MIR-10B/3P | 3.9 | 1.47 | 0.99 | 0.79 | 2.43 | 1.70E-04 | 0.0106 | (A) NE (B/D) |
| MIR-10B/5P | 3.9 | 1.47 | 0.99 | 0.79 | 2.43 | 1.70E-04 | 0.0106 | (A) NE (B/D) |
| DUS3L | 3.38 | 0.95 | 0.57 | 0.94 | 2.43 | 5.93E-04 | 0.0194 | (A) NE (B/D) |
| RTBDN | 3.16 | 0.72 | 1.51 | 1.06 | 2.43 | 2.81E-03 | 0.0412 | (A) NE (B/D) |
| MTMR9LP | 7.2 | 4.77 | 1.26 | 1.22 | 2.43 | 3.20E-03 | 0.0437 | (A) NE (B/D) |
| DMWD | 2.91 | 0.49 | 0.66 | 0.49 | 2.42 | 2.04E-06 | 8.29E-04 | (A) NE (B/D) |
| KCNH2 | 2.76 | 0.34 | 0.4 | 0.63 | 2.42 | 3.96E-05 | 5.17E-03 | (A) NE (B/D) |
| EPHB3 | 2.58 | 0.17 | 1.06 | 0.41 | 2.42 | 6.43E-05 | 6.81E-03 | (A) NE (B/D) |
| INAFM1 | 5.46 | 3.04 | 0.53 | 0.77 | 2.42 | 1.34E-04 | 9.33E-03 | (A) NE (B/D) |
| INPP5E | 6.16 | 3.74 | 0.65 | 0.89 | 2.42 | 3.65E-04 | 0.0156 | (A) NE (B/D) |
| EHMT2-AS1 | 7.81 | 5.39 | 0.46 | 1.06 | 2.42 | 1.59E-03 | 0.0318 | (A) NE (B/D) |
| TFPT | 5.01 | 2.6 | 0.47 | 0.81 | 2.41 | 2.69E-04 | 0.0137 | (A) NE (B/D) |
| MIR-639/639 | 3.34 | 0.93 | 0.58 | 0.98 | 2.41 | 8.62E-04 | 0.0237 | (A) NE (B/D) |
| SMTN | 4.99 | 2.58 | 0.63 | 1.01 | 2.41 | 9.68E-04 | 0.0251 | (A) NE (B/D) |
| NXPH1 | 2.56 | 0.15 | 1.49 | 0.38 | 2.41 | 1.05E-03 | 0.0259 | (A) NE (B/D) |
| TNPO2 | 7.32 | 4.91 | 0.41 | 1.07 | 2.41 | 1.94E-03 | 0.0351 | (A) NE (B/D) |
| ARX | 3.18 | 0.77 | 0.79 | 1.17 | 2.41 | 2.16E-03 | 0.0369 | (A) NE (B/D) |
| TK1 | 3.26 | 0.85 | 0.98 | 1.21 | 2.41 | 2.65E-03 | 0.04 | (A) NE (B/D) |
| ELOVL1 | 5.39 | 2.99 | 0.46 | 1.17 | 2.41 | 2.90E-03 | 0.0417 | (A) NE (B/D) |
| UCN | 2.78 | 0.37 | 0.84 | 0.46 | 2.4 | 9.18E-06 | 2.28E-03 | (A) NE (B/D) |
| MAGEA3 | 2.7 | 0.31 | 1.56 | 0.75 | 2.4 | 1.81E-03 | 0.0341 | (A) NE (B/D) |
| CDK11B | 4.85 | 2.45 | 0.33 | 1.17 | 2.4 | 3.40E-03 | 0.0451 | (A) NE (B/D) |
| CLASRP | 2.47 | 0.08 | 0.95 | 0.13 | 2.39 | 5.26E-05 | 5.98E-03 | (A) NE (B/D) |
| CPNE1 | 5.45 | 3.07 | 0.55 | 0.77 | 2.39 | 1.40E-04 | 9.53E-03 | (A) NE (B/D) |
| SNHG20 | 4.66 | 2.27 | 0.77 | 0.8 | 2.39 | 1.51E-04 | 9.92E-03 | (A) NE (B/D) |

| | | | | | | | | |
|---|---|---|---|---|---|---|---|---|
| ZNF266 | 7.36 | 4.97 | 1.04 | 0.99 | 2.39 | 8.55E-04 | 0.0236 | (A) NE (B/D) |
| YTHDF3-AS1 | 6.48 | 4.09 | 0.33 | 0.92 | 2.39 | 9.41E-04 | 0.0248 | (A) NE (B/D) |
| MANEAL | 2.88 | 0.5 | 0.51 | 1.21 | 2.39 | 3.55E-03 | 0.0459 | (A) NE (B/D) |
| RPL32P29 | 3.09 | 0.7 | 1.28 | 1.1 | 2.38 | 2.28E-03 | 0.0376 | (A) NE (B/D) |
| SPPL2B | 6.86 | 4.47 | 0.45 | 1.11 | 2.38 | 2.33E-03 | 0.0382 | (A) NE (B/D) |
| ILF3 | 9.86 | 7.48 | 0.51 | 1.23 | 2.38 | 3.94E-03 | 0.0483 | (A) NE (B/D) |
| ABHD17A | 2.49 | 0.12 | 0.29 | 0.29 | 2.37 | 9.47E-09 | 4.43E-05 | (A) NE (B/D) |
| PRXL2B | 7.48 | 5.11 | 0.64 | 1.01 | 2.37 | 1.08E-03 | 0.0261 | (A) NE (B/D) |
| ZNF853 | 5.02 | 2.65 | 1.42 | 0.74 | 2.37 | 1.08E-03 | 0.0261 | (A) NE (B/D) |
| SNORD62B | 6.95 | 4.58 | 0.71 | 1.04 | 2.37 | 1.18E-03 | 0.0272 | (A) NE (B/D) |
| ARFGAP1 | 3.57 | 1.21 | 0.34 | 1 | 2.37 | 1.59E-03 | 0.0317 | (A) NE (B/D) |
| PRICKLE3 | 4.49 | 2.12 | 0.41 | 1.04 | 2.37 | 1.76E-03 | 0.0336 | (A) NE (B/D) |
| GPI | 4.64 | 2.27 | 0.35 | 1.17 | 2.37 | 3.47E-03 | 0.0454 | (A) NE (B/D) |
| LRP10 | 9.19 | 6.82 | 0.83 | 1.25 | 2.37 | 3.56E-03 | 0.0459 | (A) NE (B/D) |
| TMEM191A | 3.27 | 0.91 | 0.5 | 0.78 | 2.36 | 1.98E-04 | 0.0113 | (A) NE (B/D) |
| CLPTM1 | 4.12 | 1.76 | 0.21 | 0.75 | 2.36 | 4.37E-04 | 0.0173 | (A) NE (B/D) |
| EMC10 | 10.17 | 7.82 | 0.6 | 1.17 | 2.36 | 2.78E-03 | 0.041 | (A) NE (B/D) |
| MKNK2 | 4.14 | 1.78 | 0.48 | 1.16 | 2.36 | 3.05E-03 | 0.0427 | (A) NE (B/D) |
| TRIR | 3.8 | 1.44 | 0.21 | 1.1 | 2.36 | 3.09E-03 | 0.0429 | (A) NE (B/D) |
| SCMH1 | 2.5 | 0.15 | 0.38 | 0.37 | 2.35 | 1.21E-07 | 2.56E-04 | (A) NE (B/D) |
| STXBP2 | 2.63 | 0.28 | 0.3 | 0.55 | 2.35 | 2.89E-05 | 4.49E-03 | (A) NE (B/D) |
| MAFK | 3.36 | 1.01 | 0.69 | 0.71 | 2.35 | 5.97E-05 | 6.53E-03 | (A) NE (B/D) |
| TYK2 | 4.85 | 2.51 | 0.33 | 0.7 | 2.35 | 1.84E-04 | 0.0109 | (A) NE (B/D) |
| SMIM29 | 8.11 | 5.76 | 0.47 | 0.77 | 2.35 | 2.06E-04 | 0.0115 | (A) NE (B/D) |
| OGT | 11.07 | 8.71 | 1.18 | 0.77 | 2.35 | 4.28E-04 | 0.0171 | (A) NE (B/D) |
| POLR2A | 4.01 | 1.66 | 0.38 | 0.87 | 2.35 | 6.36E-04 | 0.0201 | (A) NE (B/D) |
| SGTA | 3.74 | 1.39 | 0.15 | 0.88 | 2.35 | 1.12E-03 | 0.0265 | (A) NE (B/D) |
| MDP1 | 5.33 | 2.99 | 0.81 | 1.13 | 2.35 | 2.08E-03 | 0.0362 | (A) NE (B/D) |
| UNC45A | 7.96 | 5.61 | 0.27 | 1.05 | 2.35 | 2.36E-03 | 0.0383 | (A) NE (B/D) |
| SNORD41 | 3.45 | 1.12 | 0.75 | 0.9 | 2.34 | 4.43E-04 | 0.0174 | (A) NE (B/D) |
| MIF | 5.53 | 3.19 | 1.32 | 0.88 | 2.34 | 1.22E-03 | 0.0275 | (A) NE (B/D) |
| IRAK1 | 3.18 | 0.84 | 0.43 | 1.02 | 2.34 | 1.64E-03 | 0.0323 | (A) NE (B/D) |
| FAM71E1 | 7.43 | 5.09 | 0.7 | 1.12 | 2.34 | 2.06E-03 | 0.0361 | (A) NE (B/D) |
| PYCR2 | 5.06 | 2.72 | 0.31 | 1.12 | 2.34 | 3.13E-03 | 0.043 | (A) NE (B/D) |
| PPIAP6 | 5.9 | 3.56 | 1.42 | 1.09 | 2.34 | 3.33E-03 | 0.0445 | (A) NE (B/D) |
| SLC2A1 | 7.92 | 5.57 | 1.16 | 1.21 | 2.34 | 3.51E-03 | 0.0457 | (A) NE (B/D) |
| RPS8 | 9.99 | 7.65 | 0.29 | 1.17 | 2.34 | 3.93E-03 | 0.0482 | (A) NE (B/D) |
| BMS1P10 | 6.24 | 3.91 | 0.74 | 0.6 | 2.33 | 1.88E-05 | 3.50E-03 | (A) NE (B/D) |
| MBD3 | 3.46 | 1.13 | 0.44 | 0.9 | 2.33 | 7.80E-04 | 0.0225 | (A) NE (B/D) |
| SPINT1 | 2.74 | 0.41 | 0.51 | 1 | 2.33 | 1.28E-03 | 0.0283 | (A) NE (B/D) |
| PRMT1 | 8.2 | 5.87 | 0.39 | 1.03 | 2.33 | 1.90E-03 | 0.0348 | (A) NE (B/D) |
| RPL35P2 | 6.72 | 4.39 | 1 | 1.15 | 2.33 | 2.50E-03 | 0.0391 | (A) NE (B/D) |
| RNF44 | 8.18 | 5.86 | 0.32 | 0.77 | 2.32 | 3.79E-04 | 0.0159 | (A) NE (B/D) |
| HAUS5 | 3.29 | 0.97 | 0.92 | 0.95 | 2.32 | 7.23E-04 | 0.0217 | (A) NE (B/D) |
| PPIE | 2.91 | 0.59 | 0.69 | 1.01 | 2.32 | 1.14E-03 | 0.0268 | (A) NE (B/D) |
| ZNF224 | 2.87 | 0.56 | 1.56 | 0.5 | 2.32 | 1.92E-03 | 0.0349 | (A) NE (B/D) |
| LAMTOR1 | 7.17 | 4.85 | 0.45 | 1.05 | 2.32 | 1.92E-03 | 0.035 | (A) NE (B/D) |
| CHPF | 5.18 | 2.86 | 0.59 | 1.13 | 2.32 | 2.48E-03 | 0.0391 | (A) NE (B/D) |
| HNRNPA1L2 | 2.99 | 0.67 | 0.54 | 1.15 | 2.32 | 2.94E-03 | 0.042 | (A) NE (B/D) |
| MSL3P1 | 6.93 | 4.61 | 0.34 | 1.15 | 2.32 | 3.63E-03 | 0.0463 | (A) NE (B/D) |
| UXT | 4.47 | 2.14 | 0.59 | 1.23 | 2.32 | 4.14E-03 | 0.049 | (A) NE (B/D) |
| PTTG3P | 4.68 | 2.37 | 0.94 | 0.66 | 2.31 | 8.81E-05 | 7.77E-03 | (A) NE (B/D) |
| LMNB2 | 3.04 | 0.73 | 0.5 | 0.7 | 2.31 | 9.38E-05 | 7.97E-03 | (A) NE (B/D) |
| PCDHB8 | 2.47 | 0.15 | 1.08 | 0.38 | 2.31 | 1.12E-04 | 8.69E-03 | (A) NE (B/D) |
| PSMG4 | 3.22 | 0.91 | 0.95 | 0.8 | 2.31 | 2.68E-04 | 0.0137 | (A) NE (B/D) |
| STK36 | 5.16 | 2.85 | 0.78 | 1 | 2.31 | 1.04E-03 | 0.0258 | (A) NE (B/D) |
| NFKBIL1 | 7.75 | 5.44 | 0.38 | 0.98 | 2.31 | 1.48E-03 | 0.0308 | (A) NE (B/D) |
| GAD1 | 2.77 | 0.47 | 1.12 | 0.73 | 2.3 | 3.33E-04 | 0.0149 | (A) NE (B/D) |
| HOTAIRM1 | 2.75 | 0.45 | 1.17 | 0.74 | 2.3 | 4.51E-04 | 0.0175 | (A) NE (B/D) |
| AGAP3 | 5.33 | 3.03 | 0.41 | 0.88 | 2.3 | 7.59E-04 | 0.0222 | (A) NE (B/D) |
| TRIM65 | 3.26 | 0.95 | 0.47 | 1 | 2.3 | 1.52E-03 | 0.031 | (A) NE (B/D) |
| LINC01011 | 6.17 | 3.87 | 1.31 | 0.96 | 2.3 | 1.88E-03 | 0.0347 | (A) NE (B/D) |
| BAIAP2-DT | 5.46 | 3.16 | 0.55 | 1.06 | 2.3 | 1.94E-03 | 0.0351 | (A) NE (B/D) |
| SMPDL3B | 3.36 | 1.06 | 1.17 | 1.1 | 2.3 | 2.49E-03 | 0.0391 | (A) NE (B/D) |
| FZD1 | 3.37 | 1.07 | 0.52 | 1.2 | 2.3 | 4.04E-03 | 0.0486 | (A) NE (B/D) |

| | | | | | | | | |
|---|---|---|---|---|---|---|---|---|
| EPHA10 | 2.81 | 0.51 | 0.4 | 0.53 | 2.29 | 9.61E-06 | 2.34E-03 | (A) NE (B/D) |
| NDUFS7 | 2.92 | 0.63 | 0.4 | 0.61 | 2.29 | 4.58E-05 | 5.58E-03 | (A) NE (B/D) |
| YBX1P2 | 6.29 | 4 | 0.65 | 0.95 | 2.29 | 8.54E-04 | 0.0236 | (A) NE (B/D) |
| ZNF692 | 4.54 | 2.25 | 0.65 | 0.99 | 2.29 | 1.08E-03 | 0.0261 | (A) NE (B/D) |
| CCNL2 | 8.66 | 6.37 | 0.69 | 1.2 | 2.29 | 3.65E-03 | 0.0464 | (A) NE (B/D) |
| NPAS2 | 2.42 | 0.14 | 1.14 | 0.33 | 2.28 | 2.21E-04 | 0.0121 | (A) NE (B/D) |
| CCND3 | 5.01 | 2.73 | 0.68 | 0.81 | 2.28 | 2.51E-04 | 0.0132 | (A) NE (B/D) |
| ZNF133 | 2.86 | 0.57 | 0.78 | 0.86 | 2.28 | 3.68E-04 | 0.0156 | (A) NE (B/D) |
| ODF2 | 8.47 | 6.19 | 0.68 | 0.97 | 2.28 | 9.34E-04 | 0.0247 | (A) NE (B/D) |
| JUN | 8.5 | 6.22 | 0.7 | 1.07 | 2.28 | 1.89E-03 | 0.0347 | (A) NE (B/D) |
| HIGD2A | 3.11 | 0.83 | 0.32 | 1.02 | 2.28 | 2.20E-03 | 0.0372 | (A) NE (B/D) |
| SAFB2 | 7.36 | 5.08 | 0.69 | 1.17 | 2.28 | 3.26E-03 | 0.0441 | (A) NE (B/D) |
| NUMA1 | 5.92 | 3.65 | 0.52 | 0.53 | 2.27 | 5.82E-06 | 1.74E-03 | (A) NE (B/D) |
| PPAN-P2RY1 | 5.25 | 2.98 | 0.66 | 0.57 | 2.27 | 1.31E-05 | 2.81E-03 | (A) NE (B/D) |
| HAPLN2 | 2.55 | 0.28 | 0.61 | 0.69 | 2.27 | 7.29E-05 | 7.15E-03 | (A) NE (B/D) |
| CLSTN1 | 2.53 | 0.26 | 0.98 | 0.64 | 2.27 | 1.14E-04 | 8.73E-03 | (A) NE (B/D) |
| TNK1 | 2.73 | 0.46 | 0.44 | 0.7 | 2.27 | 1.23E-04 | 9.09E-03 | (A) NE (B/D) |
| XCL1 | 2.45 | 0.18 | 1.16 | 0.44 | 2.27 | 2.40E-04 | 0.0128 | (A) NE (B/D) |
| PAQR6 | 2.73 | 0.46 | 0.91 | 0.8 | 2.27 | 2.66E-04 | 0.0137 | (A) NE (B/D) |
| ARGLU1 | 2.76 | 0.48 | 1.2 | 0.54 | 2.27 | 3.35E-04 | 0.0149 | (A) NE (B/D) |
| IDH3G | 2.91 | 0.65 | 0.24 | 0.89 | 2.27 | 1.23E-03 | 0.0276 | (A) NE (B/D) |
| DDX49 | 6.01 | 3.74 | 0.41 | 0.99 | 2.27 | 1.65E-03 | 0.0323 | (A) NE (B/D) |
| RPS4XP1 | 7.81 | 5.55 | 0.8 | 1.09 | 2.27 | 2.00E-03 | 0.0354 | (A) NE (B/D) |
| CA8 | 2.53 | 0.26 | 1.72 | 0.43 | 2.27 | 4.00E-03 | 0.0484 | (A) NE (B/D) |
| MBOAT7 | 2.59 | 0.33 | 0.43 | 0.44 | 2.26 | 1.08E-06 | 6.65E-04 | (A) NE (B/D) |
| FAAP20 | 4.44 | 2.18 | 0.46 | 0.86 | 2.26 | 6.11E-04 | 0.0197 | (A) NE (B/D) |
| CARM1 | 5.47 | 3.21 | 0.46 | 0.86 | 2.26 | 6.14E-04 | 0.0197 | (A) NE (B/D) |
| RUVBL2 | 4.55 | 2.29 | 0.47 | 0.9 | 2.26 | 8.14E-04 | 0.0231 | (A) NE (B/D) |
| ING5 | 7.51 | 5.25 | 0.74 | 0.97 | 2.26 | 9.56E-04 | 0.025 | (A) NE (B/D) |
| HDAC1 | 6.53 | 4.28 | 0.91 | 1.03 | 2.26 | 1.51E-03 | 0.031 | (A) NE (B/D) |
| NAA10 | 4.24 | 1.98 | 0.29 | 0.94 | 2.26 | 1.52E-03 | 0.031 | (A) NE (B/D) |
| ILF3-DT | 3.53 | 1.27 | 0.81 | 1.19 | 2.26 | 3.46E-03 | 0.0454 | (A) NE (B/D) |
| EIF3LP2 | 5.02 | 2.76 | 1.38 | 1.08 | 2.26 | 3.80E-03 | 0.0475 | (A) NE (B/D) |
| P4HTM | 7.37 | 5.12 | 0.47 | 0.78 | 2.25 | 2.82E-04 | 0.0141 | (A) NE (B/D) |
| MIR-1302-8/ | 2.59 | 0.34 | 1.25 | 0.46 | 2.25 | 4.79E-04 | 0.0178 | (A) NE (B/D) |
| PABPC1P3 | 8.04 | 5.79 | 1.26 | 0.85 | 2.25 | 1.18E-03 | 0.0272 | (A) NE (B/D) |
| TIMM13 | 3.5 | 1.25 | 0.55 | 1 | 2.25 | 1.50E-03 | 0.031 | (A) NE (B/D) |
| RPL36 | 5 | 2.75 | 0.85 | 1.09 | 2.25 | 2.17E-03 | 0.0369 | (A) NE (B/D) |
| LOXL1-AS1 | 5.13 | 2.89 | 0.49 | 1.09 | 2.25 | 2.68E-03 | 0.0402 | (A) NE (B/D) |
| NPFF | 2.96 | 0.71 | 0.75 | 1.14 | 2.25 | 2.89E-03 | 0.0417 | (A) NE (B/D) |
| CBLC | 3.34 | 1.1 | 1.45 | 0.94 | 2.25 | 2.96E-03 | 0.0421 | (A) NE (B/D) |
| PLPP5 | 7.95 | 5.71 | 0.84 | 0.82 | 2.24 | 3.19E-04 | 0.0146 | (A) NE (B/D) |
| ZSWIM4 | 3.4 | 1.16 | 0.3 | 0.73 | 2.24 | 3.35E-04 | 0.0149 | (A) NE (B/D) |
| PICK1 | 2.7 | 0.46 | 0.23 | 0.89 | 2.24 | 1.33E-03 | 0.029 | (A) NE (B/D) |
| HNRNPM | 4.96 | 2.72 | 0.26 | 1.08 | 2.24 | 3.43E-03 | 0.0452 | (A) NE (B/D) |
| RN7SKP239 | 3.96 | 1.72 | 0.81 | 1.21 | 2.24 | 4.04E-03 | 0.0486 | (A) NE (B/D) |
| RFNG | 4.76 | 2.52 | 0.65 | 1.2 | 2.24 | 4.10E-03 | 0.0489 | (A) NE (B/D) |
| ENDOG | 8.85 | 6.61 | 0.64 | 1.2 | 2.24 | 4.26E-03 | 0.0495 | (A) NE (B/D) |
| ATG16L2 | 2.66 | 0.44 | 1.15 | 0.46 | 2.23 | 2.58E-04 | 0.0134 | (A) NE (B/D) |
| YIPF2 | 3.71 | 1.47 | 0.59 | 0.81 | 2.23 | 3.24E-04 | 0.0147 | (A) NE (B/D) |
| ANAPC11 | 2.97 | 0.75 | 0.42 | 0.97 | 2.23 | 1.64E-03 | 0.0323 | (A) NE (B/D) |
| RPS19 | 6.04 | 3.81 | 0.37 | 0.97 | 2.23 | 1.73E-03 | 0.0331 | (A) NE (B/D) |
| ACIN1 | 4.02 | 1.79 | 0.79 | 1.07 | 2.23 | 2.02E-03 | 0.0357 | (A) NE (B/D) |
| TSPAN8 | 3.23 | 1 | 1.04 | 1.09 | 2.23 | 2.42E-03 | 0.0385 | (A) NE (B/D) |
| MIR-4746/3P | 6.17 | 3.94 | 0.62 | 1.09 | 2.23 | 2.49E-03 | 0.0391 | (A) NE (B/D) |
| MIR-4746/5P | 6.17 | 3.94 | 0.62 | 1.09 | 2.23 | 2.49E-03 | 0.0391 | (A) NE (B/D) |
| RPL23AP60 | 3.94 | 1.73 | 0.93 | 0.66 | 2.22 | 1.27E-04 | 9.14E-03 | (A) NE (B/D) |
| SLC26A11 | 6.14 | 3.92 | 0.96 | 0.72 | 2.22 | 2.01E-04 | 0.0114 | (A) NE (B/D) |
| PSRC1 | 2.56 | 0.34 | 0.95 | 0.83 | 2.22 | 4.73E-04 | 0.0176 | (A) NE (B/D) |
| CDKN2D | 2.89 | 0.68 | 0.94 | 0.87 | 2.22 | 5.73E-04 | 0.0192 | (A) NE (B/D) |
| C19ORF73 | 5.67 | 3.45 | 0.38 | 0.81 | 2.22 | 5.78E-04 | 0.0192 | (A) NE (B/D) |
| RPS11P5 | 11.5 | 9.29 | 0.95 | 1.11 | 2.22 | 2.64E-03 | 0.04 | (A) NE (B/D) |
| RPP21 | 4.26 | 2.04 | 0.38 | 1.12 | 2.22 | 3.73E-03 | 0.0468 | (A) NE (B/D) |
| DNM2 | 3.51 | 1.3 | 0.31 | 0.52 | 2.21 | 2.54E-05 | 4.21E-03 | (A) NE (B/D) |
| C7ORF26 | 9.53 | 7.32 | 0.47 | 0.74 | 2.21 | 2.36E-04 | 0.0127 | (A) NE (B/D) |

| | | | | | | | | |
|---|---|---|---|---|---|---|---|---|
| SZT2-AS1 | 2.69 | 0.48 | 0.48 | 0.88 | 2.21 | 7.68E-04 | 0.0223 | (A) NE (B/D) |
| RN7SL262P | 3.53 | 1.32 | 1.19 | 0.85 | 2.21 | 1.10E-03 | 0.0263 | (A) NE (B/D) |
| UAP1L1 | 8.08 | 5.88 | 1.07 | 0.96 | 2.21 | 1.40E-03 | 0.0299 | (A) NE (B/D) |
| NELFB | 9.77 | 7.56 | 0.4 | 0.98 | 2.21 | 1.78E-03 | 0.0337 | (A) NE (B/D) |
| HSPB1 | 3.52 | 1.31 | 0.94 | 1.04 | 2.21 | 1.80E-03 | 0.034 | (A) NE (B/D) |
| YIPF3 | 10.18 | 7.97 | 0.31 | 1.01 | 2.21 | 2.37E-03 | 0.0384 | (A) NE (B/D) |
| MAN2B1 | 3.85 | 1.64 | 0.22 | 1.01 | 2.21 | 2.68E-03 | 0.0402 | (A) NE (B/D) |
| SNRNP200 | 3.33 | 1.13 | 0.41 | 1.11 | 2.21 | 3.53E-03 | 0.0457 | (A) NE (B/D) |
| ZFAND2B | 5.44 | 3.23 | 0.56 | 1.16 | 2.21 | 3.96E-03 | 0.0483 | (A) NE (B/D) |
| USP27X-AS1 | 4.06 | 1.85 | 1.08 | 1.18 | 2.21 | 4.15E-03 | 0.049 | (A) NE (B/D) |
| MAP2K2 | 4.02 | 1.82 | 0.38 | 0.44 | 2.2 | 2.28E-06 | 9.05E-04 | (A) NE (B/D) |
| RN7SL521P | 3.06 | 0.86 | 0.92 | 0.7 | 2.2 | 1.70E-04 | 0.0106 | (A) NE (B/D) |
| BSG | 4.83 | 2.62 | 0.32 | 0.74 | 2.2 | 3.75E-04 | 0.0158 | (A) NE (B/D) |
| UCKL1 | 3.91 | 1.71 | 0.44 | 0.85 | 2.2 | 7.27E-04 | 0.0218 | (A) NE (B/D) |
| RPS11P7 | 2.33 | 0.12 | 1.37 | 0.3 | 2.2 | 1.14E-03 | 0.0268 | (A) NE (B/D) |
| SOX21-AS1 | 2.23 | 0.04 | 1.44 | 0.09 | 2.2 | 1.81E-03 | 0.0341 | (A) NE (B/D) |
| LRP4-AS1 | 2.43 | 0.23 | 1.48 | 0.27 | 2.2 | 1.90E-03 | 0.0348 | (A) NE (B/D) |
| FAM166A | 4.5 | 2.3 | 0.94 | 1.05 | 2.2 | 1.98E-03 | 0.0353 | (A) NE (B/D) |
| CCDC22 | 6.27 | 4.08 | 0.48 | 1.02 | 2.2 | 2.09E-03 | 0.0362 | (A) NE (B/D) |
| MIR-4658/46 | 7.54 | 5.34 | 0.46 | 1.14 | 2.2 | 3.95E-03 | 0.0483 | (A) NE (B/D) |
| FDX2 | 2.39 | 0.2 | 0.65 | 0.38 | 2.19 | 1.73E-06 | 8.29E-04 | (A) NE (B/D) |
| RPS3 | 2.5 | 0.31 | 0.62 | 0.5 | 2.19 | 5.85E-06 | 1.74E-03 | (A) NE (B/D) |
| INTS1 | 5.43 | 3.23 | 0.42 | 0.5 | 2.19 | 7.16E-06 | 2.02E-03 | (A) NE (B/D) |
| TCEA2 | 2.59 | 0.4 | 0.56 | 0.65 | 2.19 | 5.73E-05 | 6.35E-03 | (A) NE (B/D) |
| ANKRD10 | 4.61 | 2.42 | 0.78 | 0.65 | 2.19 | 6.84E-05 | 6.91E-03 | (A) NE (B/D) |
| TMEM94 | 3.17 | 0.98 | 0.62 | 0.83 | 2.19 | 4.37E-04 | 0.0173 | (A) NE (B/D) |
| AURKB | 3.76 | 1.56 | 0.8 | 0.86 | 2.19 | 5.29E-04 | 0.0185 | (A) NE (B/D) |
| CCDC120 | 6.67 | 4.48 | 0.31 | 0.9 | 2.19 | 1.36E-03 | 0.0295 | (A) NE (B/D) |
| NECTIN2 | 2.97 | 0.78 | 0.25 | 0.91 | 2.19 | 1.65E-03 | 0.0324 | (A) NE (B/D) |
| LRRC56 | 3.19 | 1 | 0.38 | 0.97 | 2.19 | 1.79E-03 | 0.0339 | (A) NE (B/D) |
| EGFL8 | 7.62 | 5.44 | 0.61 | 1.02 | 2.19 | 1.88E-03 | 0.0347 | (A) NE (B/D) |
| MIB2 | 3.31 | 1.13 | 0.76 | 0.57 | 2.18 | 2.89E-05 | 4.49E-03 | (A) NE (B/D) |
| PPME1 | 2.54 | 0.36 | 0.29 | 0.57 | 2.18 | 7.05E-05 | 7.02E-03 | (A) NE (B/D) |
| SH3GL1P3 | 2.88 | 0.7 | 0.47 | 0.69 | 2.18 | 1.38E-04 | 9.44E-03 | (A) NE (B/D) |
| NT5C | 4.42 | 2.25 | 0.6 | 0.9 | 2.18 | 8.24E-04 | 0.0232 | (A) NE (B/D) |
| STAG3L5P | 5.49 | 3.31 | 1.07 | 0.9 | 2.18 | 1.10E-03 | 0.0263 | (A) NE (B/D) |
| PLK3 | 5.92 | 3.74 | 0.85 | 1.01 | 2.18 | 1.60E-03 | 0.0318 | (A) NE (B/D) |
| UPF3AP2 | 3.21 | 1.03 | 0.66 | 1 | 2.18 | 1.62E-03 | 0.0319 | (A) NE (B/D) |
| VARS2 | 9.14 | 6.96 | 0.48 | 0.99 | 2.18 | 1.83E-03 | 0.0343 | (A) NE (B/D) |
| TRABD | 4.22 | 2.04 | 0.31 | 0.95 | 2.18 | 1.86E-03 | 0.0345 | (A) NE (B/D) |
| CSPG4P10 | 3.98 | 1.8 | 0.33 | 0.95 | 2.18 | 1.90E-03 | 0.0348 | (A) NE (B/D) |
| CCDC130 | 8.97 | 6.79 | 0.35 | 0.97 | 2.18 | 1.98E-03 | 0.0353 | (A) NE (B/D) |
| RUNX2 | 2.67 | 0.49 | 0.93 | 1.07 | 2.18 | 2.40E-03 | 0.0385 | (A) NE (B/D) |
| MIR-196B/19 | 2.35 | 0.17 | 1.56 | 0.22 | 2.18 | 2.90E-03 | 0.0417 | (A) NE (B/D) |
| MIR-196B/19 | 2.35 | 0.17 | 1.56 | 0.22 | 2.18 | 2.90E-03 | 0.0417 | (A) NE (B/D) |
| MIR-196B/3P | 2.35 | 0.17 | 1.56 | 0.22 | 2.18 | 2.90E-03 | 0.0417 | (A) NE (B/D) |
| MIR-196B/5P | 2.35 | 0.17 | 1.56 | 0.22 | 2.18 | 2.90E-03 | 0.0417 | (A) NE (B/D) |
| KLC2 | 3.43 | 1.25 | 0.6 | 1.12 | 2.18 | 3.40E-03 | 0.0451 | (A) NE (B/D) |
| MXD3 | 8.74 | 6.56 | 0.55 | 1.12 | 2.18 | 3.51E-03 | 0.0457 | (A) NE (B/D) |
| MRPL41 | 5.95 | 3.78 | 0.67 | 1.14 | 2.18 | 3.63E-03 | 0.0463 | (A) NE (B/D) |
| MED10 | 2.84 | 0.66 | 0.55 | 1.14 | 2.18 | 3.88E-03 | 0.048 | (A) NE (B/D) |
| KMT2B | 4.02 | 1.86 | 0.6 | 0.51 | 2.17 | 6.77E-06 | 1.95E-03 | (A) NE (B/D) |
| TMCO3 | 5.72 | 3.55 | 0.58 | 0.73 | 2.17 | 1.67E-04 | 0.0106 | (A) NE (B/D) |
| UQCRBP1 | 6.2 | 4.03 | 0.55 | 0.82 | 2.17 | 4.52E-04 | 0.0175 | (A) NE (B/D) |
| RAB11FIP1 | 2.54 | 0.37 | 1.13 | 0.78 | 2.17 | 7.48E-04 | 0.022 | (A) NE (B/D) |
| TSEN34 | 4.3 | 2.13 | 0.45 | 0.87 | 2.17 | 8.57E-04 | 0.0236 | (A) NE (B/D) |
| SNORA11 | 3.17 | 1 | 1.39 | 0.53 | 2.17 | 1.35E-03 | 0.0292 | (A) NE (B/D) |
| LOXL4 | 2.31 | 0.14 | 1.63 | 0.34 | 2.17 | 3.75E-03 | 0.0471 | (A) NE (B/D) |
| CIRBP | 5.19 | 3.03 | 0.65 | 0.79 | 2.16 | 3.23E-04 | 0.0147 | (A) NE (B/D) |
| ACD | 2.49 | 0.32 | 0.49 | 0.8 | 2.16 | 4.25E-04 | 0.017 | (A) NE (B/D) |
| CEMIP | 2.51 | 0.35 | 1.19 | 0.55 | 2.16 | 4.91E-04 | 0.018 | (A) NE (B/D) |
| FGFR1 | 4.28 | 2.12 | 1.17 | 0.75 | 2.16 | 7.70E-04 | 0.0223 | (A) NE (B/D) |
| CCDC183 | 4.12 | 1.96 | 0.97 | 0.92 | 2.16 | 1.12E-03 | 0.0264 | (A) NE (B/D) |
| KLHL17 | 4.09 | 1.93 | 0.5 | 1.03 | 2.16 | 2.32E-03 | 0.038 | (A) NE (B/D) |
| KLF5 | 5.57 | 3.41 | 1.25 | 1.08 | 2.16 | 3.95E-03 | 0.0483 | (A) NE (B/D) |

| | | | | | | | | |
|---|---|---|---|---|---|---|---|---|
| CAPN10-DT | 6.94 | 4.78 | 0.89 | 1.17 | 2.16 | 4.10E-03 | 0.0489 | (A) NE (B/D) |
| DDX39B | 8.07 | 5.92 | 0.28 | 1.09 | 2.16 | 4.16E-03 | 0.049 | (A) NE (B/D) |
| SIN3B | 2.31 | 0.16 | 0.75 | 0.4 | 2.15 | 8.18E-06 | 2.20E-03 | (A) NE (B/D) |
| CACTIN-AS1 | 4.07 | 1.92 | 0.35 | 0.54 | 2.15 | 2.74E-05 | 4.36E-03 | (A) NE (B/D) |
| BFSP1 | 2.59 | 0.44 | 0.9 | 0.49 | 2.15 | 5.23E-05 | 5.97E-03 | (A) NE (B/D) |
| SF3A2 | 2.45 | 0.3 | 0.17 | 0.51 | 2.15 | 6.94E-05 | 6.94E-03 | (A) NE (B/D) |
| TIMM29 | 8.1 | 5.95 | 0.65 | 0.71 | 2.15 | 1.32E-04 | 9.26E-03 | (A) NE (B/D) |
| NEURL4 | 7.09 | 4.94 | 0.88 | 0.67 | 2.15 | 1.38E-04 | 9.44E-03 | (A) NE (B/D) |
| PPP5C | 3.04 | 0.89 | 0.51 | 0.78 | 2.15 | 3.56E-04 | 0.0153 | (A) NE (B/D) |
| GCOM2 | 6.04 | 3.89 | 1.07 | 0.78 | 2.15 | 6.12E-04 | 0.0197 | (A) NE (B/D) |
| ENO1 | 3.83 | 1.68 | 1.05 | 0.92 | 2.15 | 1.28E-03 | 0.0284 | (A) NE (B/D) |
| BUD23 | 3.52 | 1.38 | 0.12 | 0.94 | 2.15 | 2.39E-03 | 0.0384 | (A) NE (B/D) |
| PCDH9-AS1 | 2.26 | 0.11 | 1.52 | 0.26 | 2.15 | 2.68E-03 | 0.0403 | (A) NE (B/D) |
| PRC1 | 3.04 | 0.89 | 1.14 | 1.06 | 2.15 | 3.02E-03 | 0.0425 | (A) NE (B/D) |
| TMEM184A | 4.16 | 2.01 | 0.77 | 1.1 | 2.15 | 3.07E-03 | 0.0427 | (A) NE (B/D) |
| UPF3A | 8.6 | 6.45 | 0.88 | 1.1 | 2.15 | 3.07E-03 | 0.0427 | (A) NE (B/D) |
| OVOL2 | 3.03 | 0.89 | 0.82 | 0.82 | 2.14 | 4.39E-04 | 0.0173 | (A) NE (B/D) |
| GOLGA6L10 | 4.26 | 2.12 | 0.41 | 0.8 | 2.14 | 5.63E-04 | 0.019 | (A) NE (B/D) |
| CTTN | 6.1 | 3.96 | 0.22 | 0.76 | 2.14 | 7.13E-04 | 0.0216 | (A) NE (B/D) |
| GNB2 | 10.65 | 8.51 | 0.58 | 1 | 2.14 | 1.90E-03 | 0.0348 | (A) NE (B/D) |
| ZNF32-AS1 | 6.99 | 4.84 | 0.29 | 0.95 | 2.14 | 2.09E-03 | 0.0362 | (A) NE (B/D) |
| HCN3 | 3.13 | 1 | 0.37 | 0.98 | 2.14 | 2.24E-03 | 0.0374 | (A) NE (B/D) |
| PHLDA2 | 2.3 | 0.16 | 1.51 | 0.39 | 2.14 | 2.56E-03 | 0.0396 | (A) NE (B/D) |
| RANGRF | 2.81 | 0.67 | 0.59 | 1.05 | 2.14 | 2.63E-03 | 0.04 | (A) NE (B/D) |
| ENOSF1 | 3.08 | 0.94 | 1.08 | 1.04 | 2.14 | 2.74E-03 | 0.0405 | (A) NE (B/D) |
| MC1R | 5.08 | 2.94 | 0.86 | 1.1 | 2.14 | 3.01E-03 | 0.0424 | (A) NE (B/D) |
| ABCA7 | 2.67 | 0.54 | 0.99 | 0.33 | 2.13 | 1.21E-04 | 9.03E-03 | (A) NE (B/D) |
| VPS37D | 3.08 | 0.95 | 0.84 | 0.76 | 2.13 | 3.05E-04 | 0.0144 | (A) NE (B/D) |
| DNAJC3-DT | 3.64 | 1.51 | 1.2 | 0.69 | 2.13 | 8.34E-04 | 0.0234 | (A) NE (B/D) |
| REPIN1 | 3.01 | 0.88 | 0.33 | 0.87 | 2.13 | 1.26E-03 | 0.0282 | (A) NE (B/D) |
| TRIB2 | 2.54 | 0.4 | 1.49 | 0.59 | 2.13 | 2.55E-03 | 0.0396 | (A) NE (B/D) |
| ARHGAP33 | 3.48 | 1.35 | 0.92 | 1.09 | 2.13 | 3.12E-03 | 0.0429 | (A) NE (B/D) |
| CDRT4 | 2.46 | 0.34 | 1.15 | 0.48 | 2.12 | 3.91E-04 | 0.0162 | (A) NE (B/D) |
| THAP3 | 4.46 | 2.34 | 0.53 | 0.82 | 2.12 | 5.58E-04 | 0.019 | (A) NE (B/D) |
| ZNF581 | 3.66 | 1.54 | 0.36 | 0.79 | 2.12 | 6.31E-04 | 0.02 | (A) NE (B/D) |
| INTS11 | 2.82 | 0.7 | 0.44 | 0.85 | 2.12 | 8.89E-04 | 0.0242 | (A) NE (B/D) |
| POU5F1P6 | 3.3 | 1.17 | 1.11 | 0.94 | 2.12 | 1.82E-03 | 0.0342 | (A) NE (B/D) |
| TPD52L2 | 5.61 | 3.49 | 0.37 | 0.99 | 2.12 | 2.52E-03 | 0.0393 | (A) NE (B/D) |
| EEF1A1P12 | 4.13 | 2 | 1.28 | 1.03 | 2.12 | 3.92E-03 | 0.0481 | (A) NE (B/D) |
| ZNF331 | 6.22 | 4.1 | 0.88 | 0.45 | 2.11 | 4.57E-05 | 5.58E-03 | (A) NE (B/D) |
| XAB2 | 2.84 | 0.73 | 0.38 | 0.6 | 2.11 | 7.27E-05 | 7.15E-03 | (A) NE (B/D) |
| CRLF1 | 6.14 | 4.04 | 1.06 | 0.36 | 2.11 | 2.15E-04 | 0.0119 | (A) NE (B/D) |
| COG1 | 2.77 | 0.66 | 0.58 | 0.77 | 2.11 | 3.27E-04 | 0.0148 | (A) NE (B/D) |
| MXD4 | 8.07 | 5.96 | 0.82 | 0.79 | 2.11 | 3.72E-04 | 0.0157 | (A) NE (B/D) |
| PTPN23 | 5.07 | 2.96 | 0.58 | 0.84 | 2.11 | 6.60E-04 | 0.0205 | (A) NE (B/D) |
| PSMG3 | 3.34 | 1.22 | 1.02 | 0.8 | 2.11 | 6.66E-04 | 0.0206 | (A) NE (B/D) |
| MIR-1238/12 | 3.33 | 1.23 | 0.66 | 0.88 | 2.11 | 8.02E-04 | 0.0228 | (A) NE (B/D) |
| MIR-1238/3P | 3.33 | 1.23 | 0.66 | 0.88 | 2.11 | 8.02E-04 | 0.0228 | (A) NE (B/D) |
| MIR-1238/5P | 3.33 | 1.23 | 0.66 | 0.88 | 2.11 | 8.02E-04 | 0.0228 | (A) NE (B/D) |
| CARD19 | 3.13 | 1.03 | 0.47 | 0.92 | 2.11 | 1.41E-03 | 0.03 | (A) NE (B/D) |
| ARRDC1 | 6.31 | 4.21 | 0.52 | 0.98 | 2.11 | 2.01E-03 | 0.0356 | (A) NE (B/D) |
| OVCA2 | 3.07 | 0.97 | 0.65 | 1.05 | 2.11 | 2.74E-03 | 0.0406 | (A) NE (B/D) |
| C7ORF13 | 4.58 | 2.47 | 0.87 | 1.08 | 2.11 | 3.11E-03 | 0.0429 | (A) NE (B/D) |
| RPL35 | 10.23 | 8.12 | 0.85 | 1.11 | 2.11 | 3.47E-03 | 0.0454 | (A) NE (B/D) |
| WDR6 | 9.34 | 7.24 | 0.56 | 1.12 | 2.11 | 4.12E-03 | 0.0489 | (A) NE (B/D) |
| CAMSAP3 | 2.62 | 0.52 | 0.43 | 0.56 | 2.1 | 3.34E-05 | 4.88E-03 | (A) NE (B/D) |
| SPRED2 | 8.31 | 6.21 | 0.73 | 0.61 | 2.1 | 5.18E-05 | 5.94E-03 | (A) NE (B/D) |
| TENT5A | 2.32 | 0.23 | 0.93 | 0.41 | 2.1 | 7.60E-05 | 7.35E-03 | (A) NE (B/D) |
| GTF2F1 | 6.79 | 4.7 | 0.38 | 0.62 | 2.1 | 1.05E-04 | 8.38E-03 | (A) NE (B/D) |
| FCHSD1 | 2.52 | 0.42 | 0.48 | 0.68 | 2.1 | 1.49E-04 | 9.80E-03 | (A) NE (B/D) |
| ASIC2 | 2.11 | 0.01 | 0.96 | 0.02 | 2.1 | 1.80E-04 | 0.0108 | (A) NE (B/D) |
| PCAT6 | 2.34 | 0.24 | 1.14 | 0.59 | 2.1 | 5.07E-04 | 0.0184 | (A) NE (B/D) |
| POMT1 | 4.21 | 2.11 | 0.81 | 0.84 | 2.1 | 5.78E-04 | 0.0192 | (A) NE (B/D) |
| FABP5 | 2.18 | 0.08 | 1.26 | 0.2 | 2.1 | 9.93E-04 | 0.0254 | (A) NE (B/D) |
| LAT2 | 7.66 | 5.56 | 0.95 | 0.89 | 2.1 | 1.07E-03 | 0.026 | (A) NE (B/D) |

| | | | | | | | | |
|---|---|---|---|---|---|---|---|---|
| NUDT1 | 2.81 | 0.72 | 0.99 | 0.91 | 2.1 | 1.32E-03 | 0.0289 | (A) NE (B/D) |
| RAC1 | 3.68 | 1.58 | 0.57 | 1.04 | 2.1 | 2.80E-03 | 0.0411 | (A) NE (B/D) |
| CSNK2B | 10.16 | 8.06 | 0.27 | 1.01 | 2.1 | 3.26E-03 | 0.0441 | (A) NE (B/D) |
| GRK2 | 6.18 | 4.1 | 0.67 | 0.77 | 2.09 | 3.15E-04 | 0.0145 | (A) NE (B/D) |
| PPT2 | 7.98 | 5.89 | 0.51 | 0.78 | 2.09 | 4.40E-04 | 0.0173 | (A) NE (B/D) |
| SLC26A2 | 2.34 | 0.25 | 1.38 | 0.38 | 2.09 | 1.66E-03 | 0.0324 | (A) NE (B/D) |
| ZFHX2 | 5.88 | 3.79 | 1.1 | 0.91 | 2.09 | 1.67E-03 | 0.0325 | (A) NE (B/D) |
| SNORD34 | 4.79 | 2.71 | 0.63 | 0.97 | 2.09 | 1.77E-03 | 0.0337 | (A) NE (B/D) |
| WASH7P | 9.28 | 7.19 | 0.25 | 0.95 | 2.09 | 2.55E-03 | 0.0396 | (A) NE (B/D) |
| LINC00926 | 2.42 | 0.32 | 1.47 | 0.79 | 2.09 | 3.60E-03 | 0.0462 | (A) NE (B/D) |
| LLGL2 | 2.51 | 0.43 | 0.49 | 0.67 | 2.08 | 1.44E-04 | 9.66E-03 | (A) NE (B/D) |
| AHSA2P | 4.72 | 2.64 | 0.51 | 0.8 | 2.08 | 5.61E-04 | 0.019 | (A) NE (B/D) |
| RAET1E-AS1 | 2.58 | 0.49 | 1.2 | 0.32 | 2.08 | 6.60E-04 | 0.0205 | (A) NE (B/D) |
| WASH4P | 2.61 | 0.53 | 0.35 | 0.85 | 2.08 | 1.15E-03 | 0.0268 | (A) NE (B/D) |
| FAM193B | 6.79 | 4.71 | 0.46 | 1.03 | 2.08 | 3.05E-03 | 0.0426 | (A) NE (B/D) |
| CD9 | 2.31 | 0.24 | 0.56 | 0.41 | 2.07 | 1.67E-06 | 8.29E-04 | (A) NE (B/D) |
| PRKCSH | 2.37 | 0.3 | 0.27 | 0.5 | 2.07 | 3.53E-05 | 4.97E-03 | (A) NE (B/D) |
| CCDC149 | 5.39 | 3.31 | 0.7 | 0.82 | 2.07 | 5.33E-04 | 0.0186 | (A) NE (B/D) |
| IRF9 | 2.31 | 0.24 | 1.22 | 0.37 | 2.07 | 7.38E-04 | 0.022 | (A) NE (B/D) |
| PIAS3 | 3.79 | 1.72 | 0.58 | 0.85 | 2.07 | 7.84E-04 | 0.0225 | (A) NE (B/D) |
| NSUN5P1 | 7.6 | 5.53 | 0.68 | 0.9 | 2.07 | 1.07E-03 | 0.026 | (A) NE (B/D) |
| HNRNPL | 4.7 | 2.63 | 0.57 | 0.91 | 2.07 | 1.27E-03 | 0.0282 | (A) NE (B/D) |
| ERGIC3 | 5.59 | 3.52 | 0.4 | 0.96 | 2.07 | 2.24E-03 | 0.0375 | (A) NE (B/D) |
| AKAP17A | 2.45 | 0.38 | 0.34 | 0.62 | 2.06 | 1.41E-04 | 9.54E-03 | (A) NE (B/D) |
| MIR-4800/3P | 7.07 | 5.02 | 0.84 | 0.75 | 2.06 | 3.43E-04 | 0.015 | (A) NE (B/D) |
| MIR-4800/5P | 7.07 | 5.02 | 0.84 | 0.75 | 2.06 | 3.43E-04 | 0.015 | (A) NE (B/D) |
| THOP1 | 4.6 | 2.54 | 0.18 | 0.78 | 2.06 | 1.07E-03 | 0.026 | (A) NE (B/D) |
| TBC1D13 | 4.82 | 2.76 | 0.44 | 0.93 | 2.06 | 1.79E-03 | 0.0339 | (A) NE (B/D) |
| RBM23 | 3.16 | 1.09 | 0.85 | 0.98 | 2.06 | 1.91E-03 | 0.0348 | (A) NE (B/D) |
| CENPW | 4.05 | 1.98 | 0.89 | 1.12 | 2.06 | 4.22E-03 | 0.0493 | (A) NE (B/D) |
| TRIM73 | 5.06 | 3.01 | 0.83 | 0.71 | 2.05 | 2.63E-04 | 0.0136 | (A) NE (B/D) |
| TOMM40 | 5.13 | 3.08 | 0.29 | 0.66 | 2.05 | 3.00E-04 | 0.0142 | (A) NE (B/D) |
| SAFB | 7.51 | 5.46 | 0.36 | 0.87 | 2.05 | 1.43E-03 | 0.0303 | (A) NE (B/D) |
| JUND | 7.09 | 5.04 | 0.44 | 0.91 | 2.05 | 1.63E-03 | 0.0322 | (A) NE (B/D) |
| KLHDC3 | 3.48 | 1.44 | 0.31 | 1.05 | 2.05 | 4.15E-03 | 0.049 | (A) NE (B/D) |
| RPS12P28 | 2.64 | 0.6 | 0.88 | 0.47 | 2.04 | 6.62E-05 | 6.86E-03 | (A) NE (B/D) |
| RPS4XP16 | 3.28 | 1.25 | 0.78 | 0.73 | 2.04 | 2.81E-04 | 0.0141 | (A) NE (B/D) |
| PRKCZ | 5.86 | 3.82 | 0.53 | 0.75 | 2.04 | 3.70E-04 | 0.0157 | (A) NE (B/D) |
| PLCG1 | 9.25 | 7.21 | 0.94 | 0.83 | 2.04 | 8.41E-04 | 0.0235 | (A) NE (B/D) |
| PI4KAP2 | 3.02 | 0.98 | 1 | 0.85 | 2.04 | 1.15E-03 | 0.0268 | (A) NE (B/D) |
| CTSV | 2.35 | 0.3 | 1.17 | 0.75 | 2.04 | 1.17E-03 | 0.0272 | (A) NE (B/D) |
| GOLGA2 | 8.98 | 6.94 | 0.42 | 0.87 | 2.04 | 1.27E-03 | 0.0282 | (A) NE (B/D) |
| MPV17L2 | 5.65 | 3.62 | 0.38 | 0.88 | 2.04 | 1.52E-03 | 0.031 | (A) NE (B/D) |
| RACK1 | 8.51 | 6.47 | 0.54 | 0.97 | 2.04 | 2.21E-03 | 0.0372 | (A) NE (B/D) |
| ZNF524 | 3.07 | 1.03 | 0.31 | 0.95 | 2.04 | 2.58E-03 | 0.0397 | (A) NE (B/D) |
| REEP4 | 5.27 | 3.23 | 0.91 | 1.05 | 2.04 | 3.28E-03 | 0.0442 | (A) NE (B/D) |
| TEKT4 | 2.8 | 0.76 | 1.5 | 0.58 | 2.04 | 3.65E-03 | 0.0464 | (A) NE (B/D) |
| BLOC1S3 | 4.41 | 2.38 | 0.49 | 0.67 | 2.03 | 1.58E-04 | 0.0102 | (A) NE (B/D) |
| ATF3 | 6.54 | 4.51 | 0.92 | 1.04 | 2.03 | 3.11E-03 | 0.0429 | (A) NE (B/D) |
| RCC1 | 6.62 | 4.59 | 0.52 | 1.03 | 2.03 | 3.25E-03 | 0.0441 | (A) NE (B/D) |
| SLC26A6 | 2.05 | 0.03 | 0.65 | 0.08 | 2.02 | 1.18E-05 | 2.62E-03 | (A) NE (B/D) |
| SP5 | 2.57 | 0.54 | 1.04 | 0.6 | 2.02 | 3.87E-04 | 0.0161 | (A) NE (B/D) |
| PROB1 | 2.7 | 0.68 | 1.2 | 0.57 | 2.02 | 8.97E-04 | 0.0243 | (A) NE (B/D) |
| THAP4 | 5.63 | 3.61 | 0.41 | 0.88 | 2.02 | 1.46E-03 | 0.0307 | (A) NE (B/D) |
| WRAP73 | 5.62 | 3.6 | 0.27 | 0.91 | 2.02 | 2.26E-03 | 0.0376 | (A) NE (B/D) |
| EML3 | 6.59 | 4.57 | 0.33 | 0.97 | 2.02 | 2.99E-03 | 0.0423 | (A) NE (B/D) |
| EHMT2 | 5.97 | 3.95 | 0.31 | 0.98 | 2.02 | 3.27E-03 | 0.0441 | (A) NE (B/D) |
| RPL7A | 7.04 | 5.01 | 1.11 | 1.04 | 2.02 | 4.02E-03 | 0.0486 | (A) NE (B/D) |
| ZNF783 | 3.63 | 1.63 | 0.4 | 0.32 | 2.01 | 1.14E-07 | 2.56E-04 | (A) NE (B/D) |
| TIMM44 | 2.08 | 0.08 | 0.66 | 0.19 | 2.01 | 7.30E-06 | 2.03E-03 | (A) NE (B/D) |
| GAS2L1 | 2.71 | 0.69 | 0.84 | 0.72 | 2.01 | 3.44E-04 | 0.015 | (A) NE (B/D) |
| TRIM27 | 13.89 | 11.88 | 0.39 | 0.72 | 2.01 | 4.15E-04 | 0.0169 | (A) NE (B/D) |
| PFKL | 4.97 | 2.96 | 0.49 | 0.79 | 2.01 | 6.58E-04 | 0.0205 | (A) NE (B/D) |
| PIN1 | 5.18 | 3.17 | 0.32 | 0.8 | 2.01 | 1.05E-03 | 0.0259 | (A) NE (B/D) |
| SDHAF1 | 3.03 | 1.02 | 0.36 | 0.82 | 2.01 | 1.14E-03 | 0.0268 | (A) NE (B/D) |

| | | | | | | | | |
|---|---|---|---|---|---|---|---|---|
| CNPPD1 | 2.6 | 0.59 | 0.19 | 0.82 | 2.01 | 1.58E-03 | 0.0316 | (A) NE (B/D) |
| MAP3K9 | 7.93 | 5.91 | 0.85 | 0.99 | 2.01 | 2.44E-03 | 0.0387 | (A) NE (B/D) |
| RAI2 | 3.48 | 1.47 | 1.19 | 0.88 | 2.01 | 2.50E-03 | 0.0391 | (A) NE (B/D) |
| CDC34 | 3.69 | 1.69 | 0.37 | 0.96 | 2.01 | 2.74E-03 | 0.0405 | (A) NE (B/D) |
| RBM10 | 3.46 | 1.45 | 0.46 | 1.01 | 2.01 | 3.31E-03 | 0.0444 | (A) NE (B/D) |
| ARF5 | 3.08 | 1.07 | 0.29 | 1.03 | 2.01 | 4.21E-03 | 0.0493 | (A) NE (B/D) |
| PANK4 | 5.66 | 3.67 | 0.31 | 0.45 | 2 | 9.74E-06 | 2.34E-03 | (A) NE (B/D) |
| TBCB | 2.4 | 0.4 | 0.51 | 0.62 | 2 | 9.09E-05 | 7.90E-03 | (A) NE (B/D) |
| XRCC1 | 6.4 | 4.4 | 0.82 | 0.7 | 2 | 2.89E-04 | 0.0142 | (A) NE (B/D) |
| ZNF436-AS1 | 2.46 | 0.46 | 1 | 0.67 | 2 | 4.64E-04 | 0.0175 | (A) NE (B/D) |
| RPS4XP6 | 5.33 | 3.33 | 0.68 | 0.81 | 2 | 6.20E-04 | 0.0198 | (A) NE (B/D) |
| AKAP8L | 2.41 | 0.42 | 0.48 | 0.81 | 2 | 8.48E-04 | 0.0236 | (A) NE (B/D) |
| FARSA | 2.73 | 0.73 | 0.51 | 0.89 | 2 | 1.46E-03 | 0.0307 | (A) NE (B/D) |
| MTERF4 | 4.09 | 2.09 | 0.76 | 0.91 | 2 | 1.47E-03 | 0.0308 | (A) NE (B/D) |
| DNPEP | 2.09 | 0.1 | 0.39 | 0.25 | 1.99 | 2.01E-08 | 6.77E-05 | (A) NE (B/D) |
| MED16 | 2.03 | 0.04 | 0.46 | 0.11 | 1.99 | 3.97E-07 | 4.22E-04 | (A) NE (B/D) |
| DDA1 | 2.13 | 0.15 | 0.46 | 0.36 | 1.99 | 5.08E-07 | 4.22E-04 | (A) NE (B/D) |
| FAHD2CP | 2.79 | 0.8 | 0.6 | 0.62 | 1.99 | 7.81E-05 | 7.49E-03 | (A) NE (B/D) |
| EEF2 | 7.43 | 5.44 | 0.88 | 0.67 | 1.99 | 2.97E-04 | 0.0142 | (A) NE (B/D) |
| HSFX2 | 2.9 | 0.92 | 1.04 | 0.6 | 1.99 | 4.53E-04 | 0.0175 | (A) NE (B/D) |
| EIF3K | 2.84 | 0.85 | 0.22 | 0.76 | 1.99 | 1.00E-03 | 0.0254 | (A) NE (B/D) |
| DPH7 | 8.37 | 6.38 | 0.44 | 0.83 | 1.99 | 1.09E-03 | 0.0262 | (A) NE (B/D) |
| ZNF311 | 2.98 | 0.99 | 1.23 | 0.58 | 1.99 | 1.20E-03 | 0.0274 | (A) NE (B/D) |
| KHSRP | 6.06 | 4.07 | 0.15 | 0.88 | 1.99 | 2.45E-03 | 0.0388 | (A) NE (B/D) |
| TNNC2 | 2.55 | 0.56 | 1.3 | 0.78 | 1.99 | 2.65E-03 | 0.04 | (A) NE (B/D) |
| FAM95B1 | 3.61 | 1.62 | 0.46 | 1.06 | 1.99 | 4.31E-03 | 0.0497 | (A) NE (B/D) |
| BTBD2 | 2.36 | 0.38 | 0.44 | 0.45 | 1.98 | 4.37E-06 | 1.50E-03 | (A) NE (B/D) |
| AHRR | 3.1 | 1.12 | 0.58 | 0.62 | 1.98 | 8.70E-05 | 7.77E-03 | (A) NE (B/D) |
| POLRMT | 7.83 | 5.85 | 0.3 | 0.67 | 1.98 | 4.02E-04 | 0.0165 | (A) NE (B/D) |
| ALDH16A1 | 2.2 | 0.22 | 1.11 | 0.52 | 1.98 | 5.64E-04 | 0.019 | (A) NE (B/D) |
| SNORD14E | 3.9 | 1.92 | 1.11 | 0.59 | 1.98 | 6.69E-04 | 0.0206 | (A) NE (B/D) |
| FAM27C | 2.9 | 0.92 | 0.62 | 0.82 | 1.98 | 7.81E-04 | 0.0225 | (A) NE (B/D) |
| TVP23C-CDRT | 5.34 | 3.36 | 0.89 | 0.88 | 1.98 | 1.40E-03 | 0.0299 | (A) NE (B/D) |
| SRRM3 | 2.05 | 0.07 | 1.29 | 0.17 | 1.98 | 1.70E-03 | 0.0328 | (A) NE (B/D) |
| CD3EAP | 2.02 | 0.05 | 0.53 | 0.13 | 1.97 | 1.57E-06 | 8.16E-04 | (A) NE (B/D) |
| ALKBH6 | 2.67 | 0.7 | 0.49 | 0.59 | 1.97 | 6.29E-05 | 6.77E-03 | (A) NE (B/D) |
| UVSSA | 4.71 | 2.74 | 0.69 | 0.6 | 1.97 | 7.91E-05 | 7.53E-03 | (A) NE (B/D) |
| CCT6P2 | 3.06 | 1.09 | 0.52 | 0.7 | 1.97 | 2.57E-04 | 0.0134 | (A) NE (B/D) |
| IQSEC2 | 4.3 | 2.33 | 0.78 | 0.85 | 1.97 | 1.01E-03 | 0.0254 | (A) NE (B/D) |
| PGPEP1 | 2.55 | 0.58 | 1.09 | 0.75 | 1.97 | 1.17E-03 | 0.0271 | (A) NE (B/D) |
| ZNF350-AS1 | 3.04 | 1.07 | 1.25 | 0.6 | 1.97 | 1.49E-03 | 0.0309 | (A) NE (B/D) |
| DDRGK1 | 6.22 | 4.25 | 0.32 | 0.93 | 1.97 | 2.67E-03 | 0.0401 | (A) NE (B/D) |
| CRYBB2 | 2.87 | 0.9 | 0.75 | 1.01 | 1.97 | 2.98E-03 | 0.0423 | (A) NE (B/D) |
| COX8A | 9.31 | 7.34 | 0.8 | 1.03 | 1.97 | 3.29E-03 | 0.0442 | (A) NE (B/D) |
| MIR-4640/3P | 2.47 | 0.51 | 0.53 | 0.48 | 1.96 | 9.05E-06 | 2.28E-03 | (A) NE (B/D) |
| MIR-4640/5P | 2.47 | 0.51 | 0.53 | 0.48 | 1.96 | 9.05E-06 | 2.28E-03 | (A) NE (B/D) |
| ISYNA1 | 2.15 | 0.19 | 0.89 | 0.46 | 1.96 | 1.04E-04 | 8.30E-03 | (A) NE (B/D) |
| MICALL2 | 2.14 | 0.18 | 1.1 | 0.45 | 1.96 | 5.29E-04 | 0.0185 | (A) NE (B/D) |
| CLIC1P1 | 2.83 | 0.87 | 0.75 | 0.91 | 1.96 | 1.59E-03 | 0.0317 | (A) NE (B/D) |
| HOXC-AS1 | 2.21 | 0.25 | 1.32 | 0.41 | 1.96 | 1.92E-03 | 0.0349 | (A) NE (B/D) |
| ZNF226 | 5.21 | 3.26 | 1.32 | 0.29 | 1.96 | 1.97E-03 | 0.0353 | (A) NE (B/D) |
| MAGEA6 | 2.21 | 0.25 | 1.3 | 0.61 | 1.96 | 2.02E-03 | 0.0356 | (A) NE (B/D) |
| ARRDC1-AS1 | 5.02 | 3.06 | 0.4 | 0.92 | 1.96 | 2.26E-03 | 0.0376 | (A) NE (B/D) |
| CCDC183-AS1 | 4.09 | 2.13 | 0.72 | 0.97 | 1.96 | 2.41E-03 | 0.0385 | (A) NE (B/D) |
| MYADM | 6.81 | 4.85 | 0.54 | 0.97 | 1.96 | 2.62E-03 | 0.04 | (A) NE (B/D) |
| PTRHD1 | 6.18 | 4.22 | 0.79 | 1.07 | 1.96 | 4.18E-03 | 0.049 | (A) NE (B/D) |
| DHX34 | 2.47 | 0.51 | 0.32 | 0.43 | 1.95 | 7.82E-06 | 2.13E-03 | (A) NE (B/D) |
| AP3D1 | 6.6 | 4.65 | 0.16 | 0.4 | 1.95 | 2.31E-05 | 3.97E-03 | (A) NE (B/D) |
| SH3GL1 | 5.45 | 3.5 | 0.49 | 0.63 | 1.95 | 1.32E-04 | 9.26E-03 | (A) NE (B/D) |
| HNRNPCP7 | 4.48 | 2.53 | 1.01 | 0.58 | 1.95 | 3.99E-04 | 0.0164 | (A) NE (B/D) |
| ZBTB22 | 6.99 | 5.05 | 0.55 | 0.76 | 1.95 | 5.54E-04 | 0.019 | (A) NE (B/D) |
| MVB12A | 3.42 | 1.47 | 0.23 | 0.67 | 1.95 | 5.58E-04 | 0.019 | (A) NE (B/D) |
| CAPNS1 | 3.15 | 1.19 | 0.56 | 0.8 | 1.95 | 7.82E-04 | 0.0225 | (A) NE (B/D) |
| TRIM52 | 6.01 | 4.07 | 0.73 | 0.84 | 1.95 | 9.72E-04 | 0.0252 | (A) NE (B/D) |
| FASTK | 2.49 | 0.53 | 0.49 | 0.84 | 1.95 | 1.17E-03 | 0.0271 | (A) NE (B/D) |

| | | | | | | | | |
|---|---|---|---|---|---|---|---|---|
| EIF2B5 | 7.34 | 5.39 | 0.47 | 0.9 | 1.95 | 1.85E-03 | 0.0345 | (A) NE (B/D) |
| MAN2C1 | 3.12 | 1.18 | 0.51 | 0.91 | 1.95 | 1.94E-03 | 0.0351 | (A) NE (B/D) |
| GOLGA6L3 | 2.55 | 0.6 | 0.72 | 0.94 | 1.95 | 2.13E-03 | 0.0366 | (A) NE (B/D) |
| C9ORF16 | 7.85 | 5.9 | 0.69 | 0.99 | 1.95 | 2.89E-03 | 0.0417 | (A) NE (B/D) |
| ABHD17AP3 | 3.83 | 1.88 | 0.78 | 1 | 1.95 | 2.94E-03 | 0.042 | (A) NE (B/D) |
| NELFE | 4.21 | 2.26 | 0.23 | 0.92 | 1.95 | 3.03E-03 | 0.0425 | (A) NE (B/D) |
| IER5L | 6.83 | 4.89 | 0.53 | 1 | 1.95 | 3.35E-03 | 0.0447 | (A) NE (B/D) |
| CBR3-AS1 | 2.8 | 0.86 | 0.58 | 1.03 | 1.95 | 3.81E-03 | 0.0475 | (A) NE (B/D) |
| TCEAL9 | 6.76 | 4.81 | 1.32 | 0.87 | 1.95 | 4.18E-03 | 0.0491 | (A) NE (B/D) |
| FOXK1 | 2.27 | 0.33 | 0.51 | 0.19 | 1.94 | 4.85E-07 | 4.22E-04 | (A) NE (B/D) |
| HNRNPUL1 | 3.91 | 1.97 | 0.46 | 0.57 | 1.94 | 5.61E-05 | 6.27E-03 | (A) NE (B/D) |
| ZNF223 | 5.5 | 3.57 | 0.9 | 0.39 | 1.94 | 1.13E-04 | 8.69E-03 | (A) NE (B/D) |
| STX16-NPEPL | 6.38 | 4.44 | 0.45 | 0.65 | 1.94 | 2.05E-04 | 0.0115 | (A) NE (B/D) |
| TLNRD1 | 4.68 | 2.74 | 0.4 | 0.65 | 1.94 | 2.40E-04 | 0.0128 | (A) NE (B/D) |
| RABEP2 | 5.43 | 3.48 | 0.42 | 0.68 | 1.94 | 3.14E-04 | 0.0145 | (A) NE (B/D) |
| SLC16A6P1 | 3.42 | 1.47 | 0.64 | 0.79 | 1.94 | 6.77E-04 | 0.0209 | (A) NE (B/D) |
| ZNF700 | 2.87 | 0.92 | 1.13 | 0.78 | 1.94 | 1.71E-03 | 0.0329 | (A) NE (B/D) |
| BCL2L2 | 4.09 | 2.16 | 0.6 | 0.93 | 1.94 | 2.03E-03 | 0.0357 | (A) NE (B/D) |
| ROGDI | 2.43 | 0.48 | 0.64 | 0.94 | 1.94 | 2.18E-03 | 0.037 | (A) NE (B/D) |
| FBXW5 | 8.24 | 6.3 | 0.58 | 0.97 | 1.94 | 2.73E-03 | 0.0404 | (A) NE (B/D) |
| ADM5 | 5.21 | 3.27 | 0.73 | 1.05 | 1.94 | 4.11E-03 | 0.0489 | (A) NE (B/D) |
| NPHP4 | 3.11 | 1.17 | 0.75 | 0.68 | 1.93 | 2.71E-04 | 0.0137 | (A) NE (B/D) |
| WDR45B | 7.94 | 6 | 0.63 | 0.76 | 1.93 | 5.42E-04 | 0.0186 | (A) NE (B/D) |
| ANKRD23 | 2.38 | 0.45 | 0.37 | 0.77 | 1.93 | 9.24E-04 | 0.0247 | (A) NE (B/D) |
| SNORD33 | 7.5 | 5.57 | 0.53 | 0.87 | 1.93 | 1.48E-03 | 0.0308 | (A) NE (B/D) |
| ANKRD13D | 4.11 | 2.18 | 0.34 | 0.83 | 1.93 | 1.51E-03 | 0.031 | (A) NE (B/D) |
| BAG6 | 2.59 | 0.67 | 0.23 | 0.81 | 1.93 | 1.72E-03 | 0.0329 | (A) NE (B/D) |
| TRA2A | 2.63 | 0.7 | 0.84 | 0.9 | 1.93 | 1.83E-03 | 0.0342 | (A) NE (B/D) |
| UBAC1 | 6.15 | 4.22 | 0.44 | 0.92 | 1.93 | 2.37E-03 | 0.0384 | (A) NE (B/D) |
| SIPA1L3 | 3.58 | 1.65 | 0.3 | 0.89 | 1.93 | 2.42E-03 | 0.0385 | (A) NE (B/D) |
| KHDC1 | 2.59 | 0.66 | 0.88 | 1.02 | 1.93 | 3.81E-03 | 0.0475 | (A) NE (B/D) |
| COPS6 | 3.3 | 1.37 | 0.15 | 0.95 | 1.93 | 3.95E-03 | 0.0483 | (A) NE (B/D) |
| TOE1 | 6.14 | 4.22 | 0.64 | 0.76 | 1.92 | 5.21E-04 | 0.0185 | (A) NE (B/D) |
| PMS2P1 | 6.8 | 4.88 | 0.86 | 0.74 | 1.92 | 6.01E-04 | 0.0195 | (A) NE (B/D) |
| ARAP1 | 7.16 | 5.24 | 0.27 | 0.7 | 1.92 | 6.64E-04 | 0.0206 | (A) NE (B/D) |
| ALDOA | 3.37 | 1.45 | 1.12 | 0.51 | 1.92 | 7.42E-04 | 0.022 | (A) NE (B/D) |
| MIR-1233-2/ | 2.78 | 0.86 | 0.47 | 0.8 | 1.92 | 9.59E-04 | 0.025 | (A) NE (B/D) |
| MIR-1233-2/ | 2.78 | 0.86 | 0.47 | 0.8 | 1.92 | 9.59E-04 | 0.025 | (A) NE (B/D) |
| MIR-1233-2/ | 2.78 | 0.86 | 0.47 | 0.8 | 1.92 | 9.59E-04 | 0.025 | (A) NE (B/D) |
| POM121B | 7.36 | 5.44 | 0.3 | 0.9 | 1.92 | 2.65E-03 | 0.04 | (A) NE (B/D) |
| RGS14 | 2.82 | 0.89 | 0.66 | 1.01 | 1.92 | 3.59E-03 | 0.0461 | (A) NE (B/D) |
| ZNF230 | 5.71 | 3.8 | 1 | 0.39 | 1.91 | 2.97E-04 | 0.0142 | (A) NE (B/D) |
| DOT1L | 4.73 | 2.82 | 0.29 | 0.72 | 1.91 | 7.52E-04 | 0.0221 | (A) NE (B/D) |
| ZNF816 | 6.52 | 4.61 | 0.73 | 0.96 | 1.91 | 2.62E-03 | 0.0399 | (A) NE (B/D) |
| ZSCAN12P1 | 4.17 | 2.26 | 1.24 | 0.83 | 1.91 | 3.30E-03 | 0.0443 | (A) NE (B/D) |
| LINC00888 | 6.04 | 4.13 | 1.18 | 0.88 | 1.91 | 3.55E-03 | 0.0459 | (A) NE (B/D) |
| RHPN2 | 2.12 | 0.22 | 0.96 | 0.37 | 1.9 | 2.13E-04 | 0.0118 | (A) NE (B/D) |
| PLEKHM1P1 | 7.53 | 5.62 | 0.59 | 0.69 | 1.9 | 2.79E-04 | 0.014 | (A) NE (B/D) |
| ZNF227 | 6.59 | 4.68 | 0.99 | 0.38 | 1.9 | 2.91E-04 | 0.0142 | (A) NE (B/D) |
| UBXN6 | 4.52 | 2.62 | 0.51 | 0.77 | 1.9 | 7.30E-04 | 0.0218 | (A) NE (B/D) |
| MIR-4292/42 | 4.16 | 2.27 | 0.35 | 0.79 | 1.9 | 1.19E-03 | 0.0273 | (A) NE (B/D) |
| LEMD2 | 5.15 | 3.25 | 0.25 | 0.79 | 1.9 | 1.56E-03 | 0.0313 | (A) NE (B/D) |
| HSP90AB1 | 4.26 | 2.35 | 0.63 | 0.93 | 1.9 | 2.30E-03 | 0.0379 | (A) NE (B/D) |
| CUL9 | 2.85 | 0.95 | 0.38 | 0.89 | 1.9 | 2.34E-03 | 0.0382 | (A) NE (B/D) |
| DENND4B | 2.57 | 0.66 | 0.35 | 0.89 | 1.9 | 2.39E-03 | 0.0384 | (A) NE (B/D) |
| DPH2 | 2.87 | 0.97 | 0.23 | 0.87 | 1.9 | 2.61E-03 | 0.0399 | (A) NE (B/D) |
| MAGIX | 4.29 | 2.39 | 1 | 0.95 | 1.9 | 3.22E-03 | 0.0438 | (A) NE (B/D) |
| DNAJA4 | 2.31 | 0.42 | 0.67 | 0.54 | 1.89 | 5.46E-05 | 6.16E-03 | (A) NE (B/D) |
| GGA3 | 2.16 | 0.26 | 0.37 | 0.61 | 1.89 | 1.78E-04 | 0.0107 | (A) NE (B/D) |
| DNLZ | 3.78 | 1.89 | 0.52 | 0.67 | 1.89 | 2.45E-04 | 0.013 | (A) NE (B/D) |
| SCAND1 | 2.49 | 0.6 | 0.78 | 0.7 | 1.89 | 4.06E-04 | 0.0166 | (A) NE (B/D) |
| RNF220 | 2.71 | 0.82 | 0.34 | 0.7 | 1.89 | 5.96E-04 | 0.0194 | (A) NE (B/D) |
| PRRC2A | 6.86 | 4.97 | 0.23 | 0.73 | 1.89 | 1.04E-03 | 0.0258 | (A) NE (B/D) |
| FAM53C | 2.47 | 0.58 | 0.53 | 0.81 | 1.89 | 1.06E-03 | 0.026 | (A) NE (B/D) |
| WRB | 2.99 | 1.1 | 0.66 | 0.86 | 1.89 | 1.43E-03 | 0.0303 | (A) NE (B/D) |

| | | | | | | | | |
|---|---|---|---|---|---|---|---|---|
| PGD | 7.28 | 5.39 | 0.52 | 0.89 | 1.89 | 1.96E-03 | 0.0351 | (A) NE (B/D) |
| LINC01002 | 2.26 | 0.38 | 0.41 | 0.93 | 1.89 | 2.93E-03 | 0.042 | (A) NE (B/D) |
| DYNLRB1 | 2.65 | 0.76 | 0.58 | 0.96 | 1.89 | 2.97E-03 | 0.0422 | (A) NE (B/D) |
| PSMD8 | 8.45 | 6.56 | 0.23 | 0.9 | 1.89 | 3.13E-03 | 0.043 | (A) NE (B/D) |
| SLC25A35 | 5.92 | 4.03 | 0.35 | 0.94 | 1.89 | 3.43E-03 | 0.0452 | (A) NE (B/D) |
| LGALS3BP | 2.62 | 0.73 | 1.07 | 0.94 | 1.89 | 3.68E-03 | 0.0466 | (A) NE (B/D) |
| ACTN4P1 | 3.67 | 1.78 | 0.6 | 1.02 | 1.89 | 4.17E-03 | 0.049 | (A) NE (B/D) |
| GBA2 | 7.46 | 5.58 | 0.29 | 0.74 | 1.88 | 9.72E-04 | 0.0252 | (A) NE (B/D) |
| RAC1P2 | 3.99 | 2.11 | 0.75 | 0.84 | 1.88 | 1.27E-03 | 0.0282 | (A) NE (B/D) |
| ZNF875 | 5.36 | 3.48 | 1.27 | 0.55 | 1.88 | 2.11E-03 | 0.0365 | (A) NE (B/D) |
| NSRP1 | 2.54 | 0.66 | 1.02 | 0.9 | 1.88 | 2.79E-03 | 0.041 | (A) NE (B/D) |
| CBX8 | 2.48 | 0.6 | 0.68 | 0.96 | 1.88 | 2.89E-03 | 0.0417 | (A) NE (B/D) |
| DNTTIP1 | 5.4 | 3.52 | 0.68 | 0.97 | 1.88 | 3.20E-03 | 0.0437 | (A) NE (B/D) |
| ZNF461 | 4.59 | 2.72 | 1.28 | 0.76 | 1.88 | 3.52E-03 | 0.0457 | (A) NE (B/D) |
| SURF2 | 5.59 | 3.71 | 0.64 | 1 | 1.88 | 3.70E-03 | 0.0466 | (A) NE (B/D) |
| PAK6 | 2.3 | 0.41 | 0.91 | 1.01 | 1.88 | 4.15E-03 | 0.049 | (A) NE (B/D) |
| CIZ1 | 2.12 | 0.25 | 0.37 | 0.61 | 1.87 | 1.95E-04 | 0.0113 | (A) NE (B/D) |
| ZNF767P | 7.68 | 5.81 | 0.43 | 0.7 | 1.87 | 4.81E-04 | 0.0178 | (A) NE (B/D) |
| IVNS1ABP | 2.33 | 0.46 | 1.09 | 0.62 | 1.87 | 1.04E-03 | 0.0258 | (A) NE (B/D) |
| PTGES2 | 4.96 | 3.09 | 0.85 | 0.84 | 1.87 | 1.48E-03 | 0.0308 | (A) NE (B/D) |
| ARMC9 | 5.63 | 3.76 | 0.79 | 0.86 | 1.87 | 1.56E-03 | 0.0314 | (A) NE (B/D) |
| ZER1 | 3.79 | 1.92 | 0.43 | 0.86 | 1.87 | 1.99E-03 | 0.0354 | (A) NE (B/D) |
| PLXNB1 | 2.55 | 0.67 | 0.45 | 0.89 | 1.87 | 2.27E-03 | 0.0376 | (A) NE (B/D) |
| SEC14L1 | 5.49 | 3.62 | 0.64 | 0.94 | 1.87 | 2.59E-03 | 0.0397 | (A) NE (B/D) |
| SOX12 | 6.08 | 4.2 | 0.65 | 0.94 | 1.87 | 2.72E-03 | 0.0404 | (A) NE (B/D) |
| EXOSC10 | 2.56 | 0.69 | 0.89 | 0.95 | 1.87 | 3.13E-03 | 0.043 | (A) NE (B/D) |
| PCED1A | 5.31 | 3.44 | 0.39 | 0.96 | 1.87 | 3.85E-03 | 0.0478 | (A) NE (B/D) |
| CALM3 | 2.41 | 0.54 | 0.5 | 0.45 | 1.86 | 8.71E-06 | 2.28E-03 | (A) NE (B/D) |
| ELMO3 | 2.1 | 0.24 | 0.67 | 0.59 | 1.86 | 1.11E-04 | 8.64E-03 | (A) NE (B/D) |
| STXBP5 | 2.12 | 0.27 | 0.89 | 0.29 | 1.86 | 1.46E-04 | 9.72E-03 | (A) NE (B/D) |
| LIN37 | 7.1 | 5.24 | 0.64 | 0.74 | 1.86 | 5.57E-04 | 0.019 | (A) NE (B/D) |
| ZNF284 | 2.82 | 0.96 | 1.06 | 0.25 | 1.86 | 6.28E-04 | 0.02 | (A) NE (B/D) |
| PCP2 | 1.88 | 0.02 | 1.05 | 0.04 | 1.86 | 6.96E-04 | 0.0213 | (A) NE (B/D) |
| NFYC | 5.1 | 3.24 | 0.47 | 0.79 | 1.86 | 1.11E-03 | 0.0263 | (A) NE (B/D) |
| RINL | 5.82 | 3.97 | 1.15 | 0.49 | 1.86 | 1.11E-03 | 0.0264 | (A) NE (B/D) |
| SYCE1L | 6.05 | 4.18 | 0.88 | 0.85 | 1.86 | 1.65E-03 | 0.0324 | (A) NE (B/D) |
| PPOX | 6.17 | 4.31 | 0.34 | 0.85 | 1.86 | 2.10E-03 | 0.0363 | (A) NE (B/D) |
| LINC00623 | 6.66 | 4.79 | 0.32 | 0.86 | 1.86 | 2.38E-03 | 0.0384 | (A) NE (B/D) |
| IRF3 | 3.72 | 1.86 | 0.58 | 0.98 | 1.86 | 3.68E-03 | 0.0466 | (A) NE (B/D) |
| ZMYM3 | 2.62 | 0.77 | 0.39 | 0.7 | 1.85 | 5.82E-04 | 0.0193 | (A) NE (B/D) |
| MAGED2 | 3.4 | 1.56 | 0.54 | 0.99 | 1.85 | 4.11E-03 | 0.0489 | (A) NE (B/D) |
| RELL2 | 2.02 | 0.18 | 0.43 | 0.45 | 1.84 | 1.00E-05 | 2.37E-03 | (A) NE (B/D) |
| KCP | 2.47 | 0.63 | 0.67 | 0.61 | 1.84 | 1.49E-04 | 9.80E-03 | (A) NE (B/D) |
| CAMTA2 | 2.25 | 0.41 | 0.36 | 0.59 | 1.84 | 1.66E-04 | 0.0106 | (A) NE (B/D) |
| C9ORF40 | 2.76 | 0.91 | 0.66 | 0.62 | 1.84 | 1.70E-04 | 0.0106 | (A) NE (B/D) |
| PDCD7 | 2.09 | 0.25 | 0.73 | 0.62 | 1.84 | 1.98E-04 | 0.0113 | (A) NE (B/D) |
| TXLNA | 5.85 | 4.01 | 0.61 | 0.78 | 1.84 | 8.95E-04 | 0.0242 | (A) NE (B/D) |
| SKIV2L | 6.96 | 5.13 | 0.36 | 0.86 | 1.84 | 2.38E-03 | 0.0384 | (A) NE (B/D) |
| MAPK3 | 4.42 | 2.58 | 0.5 | 0.92 | 1.84 | 2.91E-03 | 0.0419 | (A) NE (B/D) |
| CCNQP3 | 2.55 | 0.72 | 0.54 | 0.68 | 1.83 | 3.66E-04 | 0.0156 | (A) NE (B/D) |
| CYP4F27P | 2.31 | 0.49 | 0.82 | 0.76 | 1.83 | 9.71E-04 | 0.0252 | (A) NE (B/D) |
| HERC2P3 | 7.49 | 5.67 | 1.06 | 0.67 | 1.83 | 1.31E-03 | 0.0287 | (A) NE (B/D) |
| PLXND1 | 2.43 | 0.6 | 1.01 | 0.79 | 1.83 | 1.88E-03 | 0.0347 | (A) NE (B/D) |
| ELAVL1 | 5.75 | 3.92 | 0.38 | 0.85 | 1.83 | 2.09E-03 | 0.0362 | (A) NE (B/D) |
| TCEAL3 | 6.03 | 4.2 | 0.55 | 0.93 | 1.83 | 3.07E-03 | 0.0427 | (A) NE (B/D) |
| PPT2-EGFL8 | 8.59 | 6.76 | 0.26 | 0.91 | 1.83 | 3.62E-03 | 0.0463 | (A) NE (B/D) |
| B3GALT2 | 2.08 | 0.25 | 1.37 | 0.57 | 1.83 | 4.09E-03 | 0.0488 | (A) NE (B/D) |
| ELP5 | 7.95 | 6.13 | 0.55 | 0.98 | 1.83 | 4.21E-03 | 0.0493 | (A) NE (B/D) |
| COA8 | 4.7 | 2.87 | 0.45 | 0.98 | 1.83 | 4.31E-03 | 0.0497 | (A) NE (B/D) |
| ARHGAP5-AS | 2.55 | 0.73 | 0.96 | 0.7 | 1.82 | 9.78E-04 | 0.0253 | (A) NE (B/D) |
| CCDC180 | 3.85 | 2.03 | 1.11 | 0.7 | 1.82 | 1.82E-03 | 0.0342 | (A) NE (B/D) |
| MFSD12 | 4.35 | 2.53 | 0.24 | 0.8 | 1.82 | 2.06E-03 | 0.0361 | (A) NE (B/D) |
| SAE1 | 4.7 | 2.88 | 0.57 | 0.87 | 1.82 | 2.06E-03 | 0.0361 | (A) NE (B/D) |
| OAZ1 | 4.7 | 2.88 | 0.13 | 0.84 | 1.82 | 2.98E-03 | 0.0423 | (A) NE (B/D) |
| SMPD2 | 3.19 | 1.37 | 0.53 | 0.93 | 1.82 | 3.05E-03 | 0.0427 | (A) NE (B/D) |

| | | | | | | | | |
|---|---|---|---|---|---|---|---|---|
| YBX1P10 | 4.68 | 2.86 | 0.49 | 0.93 | 1.82 | 3.17E-03 | 0.0435 | (A) NE (B/D) |
| JOSD2 | 4.93 | 3.11 | 0.34 | 0.92 | 1.82 | 3.79E-03 | 0.0474 | (A) NE (B/D) |
| TTC14 | 1.83 | 0.01 | 1.39 | 0.03 | 1.82 | 4.27E-03 | 0.0495 | (A) NE (B/D) |
| DEDD2 | 7.93 | 6.12 | 0.38 | 0.57 | 1.81 | 1.45E-04 | 9.72E-03 | (A) NE (B/D) |
| TTYH3 | 6.73 | 4.91 | 0.51 | 0.64 | 1.81 | 2.54E-04 | 0.0133 | (A) NE (B/D) |
| IGBP1-AS2 | 6.99 | 5.18 | 0.31 | 0.66 | 1.81 | 5.81E-04 | 0.0193 | (A) NE (B/D) |
| ASXL1 | 2.44 | 0.63 | 0.31 | 0.72 | 1.81 | 9.84E-04 | 0.0253 | (A) NE (B/D) |
| P4HA2-AS1 | 3.43 | 1.61 | 0.75 | 0.78 | 1.81 | 1.02E-03 | 0.0255 | (A) NE (B/D) |
| DAB2IP | 3.16 | 1.35 | 0.52 | 0.86 | 1.81 | 2.07E-03 | 0.0361 | (A) NE (B/D) |
| PIGS | 4.92 | 3.11 | 0.22 | 0.9 | 1.81 | 3.86E-03 | 0.0478 | (A) NE (B/D) |
| PCDHB18P | 2.97 | 1.16 | 1.27 | 0.72 | 1.81 | 3.95E-03 | 0.0483 | (A) NE (B/D) |
| IQCN | 5.57 | 3.76 | 1.06 | 0.92 | 1.81 | 4.19E-03 | 0.0492 | (A) NE (B/D) |
| PIAS4 | 1.93 | 0.13 | 0.27 | 0.25 | 1.8 | 2.03E-08 | 6.77E-05 | (A) NE (B/D) |
| HOXC-AS3 | 2.27 | 0.47 | 0.94 | 0.41 | 1.8 | 3.06E-04 | 0.0144 | (A) NE (B/D) |
| RPAIN | 2.26 | 0.45 | 0.98 | 0.61 | 1.8 | 7.30E-04 | 0.0218 | (A) NE (B/D) |
| MCM3AP-AS | 8.56 | 6.76 | 0.73 | 0.77 | 1.8 | 9.35E-04 | 0.0247 | (A) NE (B/D) |
| PPP2R1A | 2.38 | 0.58 | 0.35 | 0.75 | 1.8 | 1.18E-03 | 0.0272 | (A) NE (B/D) |
| RETREG2 | 2.77 | 0.97 | 0.27 | 0.75 | 1.8 | 1.43E-03 | 0.0303 | (A) NE (B/D) |
| PPP1R12C | 3.18 | 1.39 | 0.54 | 0.83 | 1.8 | 1.61E-03 | 0.0319 | (A) NE (B/D) |
| PPP6R1 | 6.64 | 4.83 | 0.49 | 0.82 | 1.8 | 1.66E-03 | 0.0324 | (A) NE (B/D) |
| LINC00847 | 5.43 | 3.64 | 0.82 | 0.9 | 1.8 | 2.72E-03 | 0.0404 | (A) NE (B/D) |
| MBOAT1 | 2.16 | 0.35 | 0.99 | 0.86 | 1.8 | 2.83E-03 | 0.0413 | (A) NE (B/D) |
| WASH2P | 2.28 | 0.49 | 0.49 | 0.9 | 1.8 | 2.89E-03 | 0.0417 | (A) NE (B/D) |
| PANX3 | 2.03 | 0.23 | 1.32 | 0.57 | 1.8 | 3.64E-03 | 0.0464 | (A) NE (B/D) |
| TMEM63A | 1.88 | 0.09 | 0.66 | 0.19 | 1.79 | 1.72E-05 | 3.38E-03 | (A) NE (B/D) |
| MAML1 | 5.78 | 3.99 | 0.26 | 0.57 | 1.79 | 2.57E-04 | 0.0134 | (A) NE (B/D) |
| ZNF865 | 3.34 | 1.55 | 0.48 | 0.66 | 1.79 | 3.65E-04 | 0.0156 | (A) NE (B/D) |
| SBNO2 | 5.63 | 3.83 | 0.56 | 0.72 | 1.79 | 5.92E-04 | 0.0194 | (A) NE (B/D) |
| CNNM3 | 2.21 | 0.42 | 0.41 | 0.74 | 1.79 | 9.75E-04 | 0.0252 | (A) NE (B/D) |
| AMZ2P1 | 4.29 | 2.5 | 0.82 | 0.76 | 1.79 | 1.11E-03 | 0.0263 | (A) NE (B/D) |
| CBARP | 2.56 | 0.77 | 0.99 | 0.74 | 1.79 | 1.54E-03 | 0.0311 | (A) NE (B/D) |
| IPO13 | 3.56 | 1.77 | 0.36 | 0.79 | 1.79 | 1.65E-03 | 0.0324 | (A) NE (B/D) |
| TBC1D22B | 4.93 | 3.14 | 0.22 | 0.84 | 1.79 | 2.96E-03 | 0.0421 | (A) NE (B/D) |
| CNPY3 | 8.62 | 6.84 | 0.34 | 0.89 | 1.79 | 3.37E-03 | 0.0448 | (A) NE (B/D) |
| TMED4 | 3.1 | 1.31 | 0.13 | 0.89 | 1.79 | 4.25E-03 | 0.0494 | (A) NE (B/D) |
| CTU1 | 5.34 | 3.55 | 0.75 | 0.47 | 1.78 | 7.92E-05 | 7.53E-03 | (A) NE (B/D) |
| SHLD1 | 2.67 | 0.89 | 0.43 | 0.57 | 1.78 | 1.26E-04 | 9.14E-03 | (A) NE (B/D) |
| ZBTB4 | 3.7 | 1.93 | 0.73 | 0.57 | 1.78 | 1.72E-04 | 0.0106 | (A) NE (B/D) |
| DOCK9 | 3.93 | 2.15 | 1.12 | 0.62 | 1.78 | 1.80E-03 | 0.034 | (A) NE (B/D) |
| PPP2R5D | 3.08 | 1.31 | 0.33 | 0.82 | 1.78 | 2.27E-03 | 0.0376 | (A) NE (B/D) |
| STAG3 | 2.48 | 0.71 | 1.29 | 0.62 | 1.78 | 3.76E-03 | 0.0471 | (A) NE (B/D) |
| GNL1 | 7.2 | 5.42 | 0.34 | 0.93 | 1.78 | 4.28E-03 | 0.0496 | (A) NE (B/D) |
| ACTN4 | 10.25 | 8.47 | 0.58 | 0.53 | 1.77 | 6.35E-05 | 6.81E-03 | (A) NE (B/D) |
| ZNF317 | 4.98 | 3.21 | 0.42 | 0.6 | 1.77 | 2.03E-04 | 0.0114 | (A) NE (B/D) |
| RNA5SP434 | 2.03 | 0.26 | 0.95 | 0.46 | 1.77 | 4.08E-04 | 0.0166 | (A) NE (B/D) |
| CRTC1 | 1.84 | 0.07 | 0.94 | 0.17 | 1.77 | 4.20E-04 | 0.0169 | (A) NE (B/D) |
| SFPQ | 6.3 | 4.54 | 0.78 | 0.76 | 1.77 | 1.11E-03 | 0.0263 | (A) NE (B/D) |
| PKP4 | 6.65 | 4.89 | 0.51 | 0.78 | 1.77 | 1.31E-03 | 0.0287 | (A) NE (B/D) |
| TCEAL1 | 4.75 | 2.98 | 0.73 | 0.8 | 1.77 | 1.44E-03 | 0.0304 | (A) NE (B/D) |
| TDP2 | 8.49 | 6.72 | 1.01 | 0.74 | 1.77 | 1.90E-03 | 0.0348 | (A) NE (B/D) |
| AURKA | 2.16 | 0.39 | 1.14 | 0.61 | 1.77 | 1.92E-03 | 0.035 | (A) NE (B/D) |
| VPS16 | 9.39 | 7.62 | 0.47 | 0.85 | 1.77 | 2.24E-03 | 0.0374 | (A) NE (B/D) |
| TTLL4 | 7.33 | 5.58 | 0.32 | 0.43 | 1.76 | 1.66E-05 | 3.33E-03 | (A) NE (B/D) |
| COQ4 | 1.93 | 0.17 | 0.68 | 0.41 | 1.76 | 3.17E-05 | 4.73E-03 | (A) NE (B/D) |
| APTR | 2.07 | 0.3 | 0.39 | 0.54 | 1.76 | 1.03E-04 | 8.25E-03 | (A) NE (B/D) |
| TNRC18 | 1.98 | 0.22 | 0.86 | 0.35 | 1.76 | 1.75E-04 | 0.0106 | (A) NE (B/D) |
| ZNF337 | 2.49 | 0.73 | 0.59 | 0.64 | 1.76 | 2.89E-04 | 0.0142 | (A) NE (B/D) |
| TRAPPC2B | 3.36 | 1.6 | 0.73 | 0.83 | 1.76 | 1.76E-03 | 0.0335 | (A) NE (B/D) |
| RPS10P7 | 3.31 | 1.56 | 0.38 | 0.83 | 1.76 | 2.45E-03 | 0.0388 | (A) NE (B/D) |
| TMEM81 | 3.98 | 2.21 | 0.87 | 0.85 | 1.76 | 2.48E-03 | 0.0391 | (A) NE (B/D) |
| HLA-T | 2.82 | 1.06 | 0.62 | 0.91 | 1.76 | 3.21E-03 | 0.0438 | (A) NE (B/D) |
| FAM156A | 3.2 | 1.45 | 0.46 | 0.9 | 1.76 | 3.37E-03 | 0.0448 | (A) NE (B/D) |
| FZR1 | 1.78 | 0.03 | 0.32 | 0.07 | 1.75 | 4.83E-08 | 1.41E-04 | (A) NE (B/D) |
| PELP1 | 1.79 | 0.04 | 0.45 | 0.09 | 1.75 | 1.23E-06 | 7.17E-04 | (A) NE (B/D) |
| NPRL2 | 2.07 | 0.32 | 0.4 | 0.5 | 1.75 | 4.46E-05 | 5.52E-03 | (A) NE (B/D) |

| | | | | | | | | |
|---|---|---|---|---|---|---|---|---|
| GIPC1 | 2.06 | 0.31 | 0.43 | 0.51 | 1.75 | 4.80E-05 | 5.67E-03 | (A) NE (B/D) |
| TREX1 | 2.22 | 0.47 | 0.7 | 0.6 | 1.75 | 2.24E-04 | 0.0122 | (A) NE (B/D) |
| TGFB1 | 2.26 | 0.51 | 0.6 | 0.63 | 1.75 | 2.60E-04 | 0.0135 | (A) NE (B/D) |
| BTF3P2 | 2.26 | 0.51 | 0.68 | 0.62 | 1.75 | 2.62E-04 | 0.0136 | (A) NE (B/D) |
| THTPA | 1.99 | 0.24 | 0.85 | 0.6 | 1.75 | 4.43E-04 | 0.0174 | (A) NE (B/D) |
| SMPD4 | 4.66 | 2.91 | 0.66 | 0.69 | 1.75 | 5.18E-04 | 0.0185 | (A) NE (B/D) |
| RNU4-39P | 2.13 | 0.38 | 0.75 | 0.76 | 1.75 | 1.14E-03 | 0.0268 | (A) NE (B/D) |
| NAIF1 | 4.54 | 2.79 | 0.71 | 0.87 | 1.75 | 2.56E-03 | 0.0396 | (A) NE (B/D) |
| OSBP2 | 3.6 | 1.85 | 0.8 | 0.92 | 1.75 | 3.62E-03 | 0.0463 | (A) NE (B/D) |
| OGFR | 3.18 | 1.43 | 0.41 | 0.9 | 1.75 | 3.65E-03 | 0.0464 | (A) NE (B/D) |
| PABPC4 | 6.09 | 4.34 | 0.41 | 0.92 | 1.75 | 4.05E-03 | 0.0486 | (A) NE (B/D) |
| CHMP4C | 2.33 | 0.58 | 0.8 | 0.94 | 1.75 | 4.10E-03 | 0.0489 | (A) NE (B/D) |
| ROM1 | 4.01 | 2.26 | 0.47 | 0.48 | 1.74 | 2.60E-05 | 4.22E-03 | (A) NE (B/D) |
| INTS3 | 3.11 | 1.37 | 0.55 | 0.62 | 1.74 | 2.35E-04 | 0.0127 | (A) NE (B/D) |
| EXO5 | 6.78 | 5.04 | 0.76 | 0.7 | 1.74 | 7.55E-04 | 0.0222 | (A) NE (B/D) |
| CYP4A22-AS1 | 2.16 | 0.42 | 1.22 | 0.36 | 1.74 | 2.52E-03 | 0.0393 | (A) NE (B/D) |
| ANKRD49 | 2.02 | 0.28 | 1.17 | 0.68 | 1.74 | 3.06E-03 | 0.0427 | (A) NE (B/D) |
| FOXJ3 | 9.23 | 7.49 | 0.71 | 0.91 | 1.74 | 3.49E-03 | 0.0456 | (A) NE (B/D) |
| MCOLN1 | 2.68 | 0.95 | 0.2 | 0.85 | 1.74 | 3.63E-03 | 0.0463 | (A) NE (B/D) |
| NIPSNAP3B | 6.61 | 4.88 | 1.14 | 0.8 | 1.74 | 4.30E-03 | 0.0497 | (A) NE (B/D) |
| RTN2 | 1.87 | 0.14 | 0.37 | 0.35 | 1.73 | 1.44E-06 | 7.75E-04 | (A) NE (B/D) |
| RN7SL573P | 5.36 | 3.63 | 0.9 | 0.57 | 1.73 | 5.38E-04 | 0.0186 | (A) NE (B/D) |
| LRPAP1 | 3.09 | 1.36 | 0.38 | 0.68 | 1.73 | 7.05E-04 | 0.0215 | (A) NE (B/D) |
| HSP90AB3P | 7.19 | 5.47 | 0.98 | 0.69 | 1.73 | 1.53E-03 | 0.0311 | (A) NE (B/D) |
| RNY5P8 | 6.78 | 5.05 | 1.12 | 0.54 | 1.73 | 1.76E-03 | 0.0335 | (A) NE (B/D) |
| TMEM44 | 2.24 | 0.51 | 0.5 | 0.82 | 1.73 | 2.07E-03 | 0.0361 | (A) NE (B/D) |
| EIF3LP3 | 1.98 | 0.26 | 1.19 | 0.55 | 1.73 | 2.65E-03 | 0.04 | (A) NE (B/D) |
| CNTROB | 6.62 | 4.89 | 0.51 | 0.9 | 1.73 | 3.44E-03 | 0.0452 | (A) NE (B/D) |
| TIMM9 | 2.37 | 0.64 | 0.62 | 0.92 | 1.73 | 3.80E-03 | 0.0474 | (A) NE (B/D) |
| VASP | 3.24 | 1.51 | 0.58 | 0.93 | 1.73 | 4.12E-03 | 0.0489 | (A) NE (B/D) |
| HCG4P3 | 5.17 | 3.45 | 0.25 | 0.58 | 1.72 | 4.13E-04 | 0.0168 | (A) NE (B/D) |
| BBS12 | 2.75 | 1.02 | 0.88 | 0.59 | 1.72 | 5.79E-04 | 0.0192 | (A) NE (B/D) |
| SUGP2 | 1.94 | 0.22 | 1.15 | 0.54 | 1.72 | 2.10E-03 | 0.0363 | (A) NE (B/D) |
| SNORD48 | 6.54 | 4.83 | 0.67 | 0.85 | 1.72 | 2.37E-03 | 0.0384 | (A) NE (B/D) |
| RALY | 5.67 | 3.95 | 0.42 | 0.83 | 1.72 | 2.54E-03 | 0.0396 | (A) NE (B/D) |
| TMEM8B | 4.59 | 2.87 | 0.75 | 0.86 | 1.72 | 2.61E-03 | 0.0399 | (A) NE (B/D) |
| TUBGCP6 | 6.58 | 4.85 | 0.25 | 0.81 | 1.72 | 2.84E-03 | 0.0413 | (A) NE (B/D) |
| UNK | 8.55 | 6.83 | 0.45 | 0.86 | 1.72 | 2.87E-03 | 0.0417 | (A) NE (B/D) |
| PABPC1 | 4.2 | 2.48 | 0.81 | 0.87 | 1.72 | 3.03E-03 | 0.0425 | (A) NE (B/D) |
| CLN6 | 2.62 | 0.9 | 0.6 | 0.89 | 1.72 | 3.29E-03 | 0.0442 | (A) NE (B/D) |
| FOXO4 | 4.23 | 2.5 | 0.75 | 0.91 | 1.72 | 3.56E-03 | 0.0459 | (A) NE (B/D) |
| LRRFIP1 | 3.23 | 1.51 | 0.51 | 0.92 | 1.72 | 4.03E-03 | 0.0486 | (A) NE (B/D) |
| MIR-943/943 | 7.13 | 5.41 | 0.38 | 0.91 | 1.72 | 4.24E-03 | 0.0494 | (A) NE (B/D) |
| KCNMB3 | 2.92 | 1.2 | 1.16 | 0.77 | 1.72 | 4.26E-03 | 0.0495 | (A) NE (B/D) |
| MCM3AP | 5.86 | 4.15 | 0.57 | 0.58 | 1.71 | 1.73E-04 | 0.0106 | (A) NE (B/D) |
| BRD4 | 7.15 | 5.44 | 0.26 | 0.58 | 1.71 | 3.94E-04 | 0.0163 | (A) NE (B/D) |
| RTEL1-TNFRS | 3.02 | 1.31 | 0.36 | 0.62 | 1.71 | 4.46E-04 | 0.0174 | (A) NE (B/D) |
| NRF1 | 1.95 | 0.25 | 0.25 | 0.6 | 1.71 | 5.19E-04 | 0.0185 | (A) NE (B/D) |
| RBMX2 | 2.19 | 0.48 | 0.46 | 0.67 | 1.71 | 5.72E-04 | 0.0191 | (A) NE (B/D) |
| ARNT2 | 2.02 | 0.31 | 0.68 | 0.7 | 1.71 | 7.14E-04 | 0.0216 | (A) NE (B/D) |
| ANKRD10-IT1 | 7.44 | 5.73 | 1.01 | 0.49 | 1.71 | 8.76E-04 | 0.024 | (A) NE (B/D) |
| MTA3 | 6.26 | 4.55 | 0.31 | 0.79 | 1.71 | 2.20E-03 | 0.0372 | (A) NE (B/D) |
| PSENEN | 2.8 | 1.09 | 0.75 | 0.88 | 1.71 | 3.04E-03 | 0.0426 | (A) NE (B/D) |
| PLEKHH2 | 1.83 | 0.12 | 1.25 | 0.28 | 1.71 | 3.13E-03 | 0.043 | (A) NE (B/D) |
| DDX27 | 7.15 | 5.45 | 0.36 | 0.88 | 1.71 | 3.90E-03 | 0.0481 | (A) NE (B/D) |
| LSM4 | 3.35 | 1.65 | 0.5 | 0.91 | 1.71 | 3.98E-03 | 0.0484 | (A) NE (B/D) |
| LINC00111 | 2.12 | 0.41 | 1.15 | 0.75 | 1.71 | 4.02E-03 | 0.0486 | (A) NE (B/D) |
| KIF9-AS1 | 2.23 | 0.53 | 0.72 | 0.43 | 1.7 | 7.28E-05 | 7.15E-03 | (A) NE (B/D) |
| DAZAP1 | 8.49 | 6.78 | 0.1 | 0.53 | 1.7 | 4.68E-04 | 0.0176 | (A) NE (B/D) |
| RTEL1 | 4.84 | 3.13 | 0.4 | 0.64 | 1.7 | 4.71E-04 | 0.0176 | (A) NE (B/D) |
| PQBP1 | 2.16 | 0.46 | 0.44 | 0.65 | 1.7 | 4.82E-04 | 0.0178 | (A) NE (B/D) |
| SRRM1 | 2.03 | 0.32 | 0.83 | 0.64 | 1.7 | 6.69E-04 | 0.0206 | (A) NE (B/D) |
| MRPL34 | 2.32 | 0.61 | 0.55 | 0.79 | 1.7 | 1.72E-03 | 0.0329 | (A) NE (B/D) |
| C17ORF100 | 5.41 | 3.71 | 0.85 | 0.8 | 1.7 | 2.23E-03 | 0.0374 | (A) NE (B/D) |
| NIPSNAP2 | 7.74 | 6.03 | 0.74 | 0.83 | 1.7 | 2.28E-03 | 0.0376 | (A) NE (B/D) |

| | | | | | | | | |
|---|---|---|---|---|---|---|---|---|
| MSTO2P | 4.27 | 2.57 | 0.3 | 0.85 | 1.7 | 3.47E-03 | 0.0454 | (A) NE (B/D) |
| OSBPL2 | 6.72 | 5.02 | 0.25 | 0.85 | 1.7 | 3.73E-03 | 0.0468 | (A) NE (B/D) |
| RAVER1 | 1.76 | 0.07 | 0.24 | 0.17 | 1.69 | 8.36E-10 | 1.04E-05 | (A) NE (B/D) |
| ZSCAN21 | 1.88 | 0.19 | 0.46 | 0.3 | 1.69 | 1.01E-06 | 6.56E-04 | (A) NE (B/D) |
| DUSP4 | 1.89 | 0.19 | 0.59 | 0.41 | 1.69 | 1.95E-05 | 3.50E-03 | (A) NE (B/D) |
| TMUB1 | 2 | 0.31 | 0.43 | 0.53 | 1.69 | 9.48E-05 | 8.00E-03 | (A) NE (B/D) |
| LAMTOR4 | 2.03 | 0.34 | 0.2 | 0.55 | 1.69 | 3.73E-04 | 0.0157 | (A) NE (B/D) |
| MPV17 | 2.03 | 0.34 | 0.33 | 0.59 | 1.69 | 3.79E-04 | 0.0159 | (A) NE (B/D) |
| AGBL5 | 7.04 | 5.35 | 0.63 | 0.76 | 1.69 | 1.29E-03 | 0.0285 | (A) NE (B/D) |
| LINC00312 | 2.42 | 0.73 | 1.04 | 0.6 | 1.69 | 1.56E-03 | 0.0313 | (A) NE (B/D) |
| ZNF789 | 2.71 | 1.03 | 0.65 | 0.78 | 1.69 | 1.61E-03 | 0.0319 | (A) NE (B/D) |
| LSR | 3.08 | 1.39 | 0.77 | 0.81 | 1.69 | 2.17E-03 | 0.0369 | (A) NE (B/D) |
| UBA1 | 8.06 | 6.37 | 0.39 | 0.84 | 1.69 | 3.10E-03 | 0.0429 | (A) NE (B/D) |
| PCIF1 | 8.08 | 6.39 | 0.24 | 0.84 | 1.69 | 3.58E-03 | 0.046 | (A) NE (B/D) |
| WDR73 | 8.31 | 6.62 | 0.43 | 0.89 | 1.69 | 3.91E-03 | 0.0481 | (A) NE (B/D) |
| MOSPD3 | 2.08 | 0.4 | 0.46 | 0.48 | 1.68 | 3.61E-05 | 5.05E-03 | (A) NE (B/D) |
| IRF2BP1 | 4.84 | 3.16 | 0.48 | 0.53 | 1.68 | 9.93E-05 | 8.15E-03 | (A) NE (B/D) |
| CDK5 | 1.94 | 0.26 | 0.48 | 0.64 | 1.68 | 4.33E-04 | 0.0173 | (A) NE (B/D) |
| SEPTIN9 | 4.97 | 3.29 | 0.56 | 0.66 | 1.68 | 5.24E-04 | 0.0185 | (A) NE (B/D) |
| COX19 | 4.39 | 2.71 | 0.95 | 0.49 | 1.68 | 6.86E-04 | 0.021 | (A) NE (B/D) |
| MIR-3655/36 | 5.42 | 3.75 | 0.88 | 0.63 | 1.68 | 9.09E-04 | 0.0245 | (A) NE (B/D) |
| POM121C | 7.7 | 6.02 | 0.2 | 0.64 | 1.68 | 9.66E-04 | 0.0251 | (A) NE (B/D) |
| PDCD2L | 2.77 | 1.08 | 0.33 | 0.7 | 1.68 | 1.13E-03 | 0.0268 | (A) NE (B/D) |
| GP1BB | 2.04 | 0.37 | 0.94 | 0.71 | 1.68 | 1.83E-03 | 0.0343 | (A) NE (B/D) |
| GPR160 | 5.75 | 4.06 | 0.75 | 0.79 | 1.68 | 1.93E-03 | 0.035 | (A) NE (B/D) |
| ZNF787 | 3.05 | 1.37 | 0.2 | 0.73 | 1.68 | 2.01E-03 | 0.0355 | (A) NE (B/D) |
| BIRC5 | 2.03 | 0.35 | 0.93 | 0.75 | 1.68 | 2.13E-03 | 0.0366 | (A) NE (B/D) |
| POLR2J4 | 7.78 | 6.1 | 0.46 | 0.82 | 1.68 | 2.42E-03 | 0.0385 | (A) NE (B/D) |
| CYTH3 | 7.6 | 5.91 | 0.92 | 0.79 | 1.68 | 2.63E-03 | 0.04 | (A) NE (B/D) |
| LINC00680 | 4.72 | 3.04 | 1 | 0.8 | 1.68 | 3.44E-03 | 0.0452 | (A) NE (B/D) |
| HNRNPA1P5( | 5.15 | 3.47 | 0.8 | 0.87 | 1.68 | 3.57E-03 | 0.0459 | (A) NE (B/D) |
| ZNF880 | 4.37 | 2.7 | 0.67 | 0.91 | 1.68 | 4.23E-03 | 0.0494 | (A) NE (B/D) |
| STK11 | 2.43 | 0.76 | 0.23 | 0.4 | 1.67 | 3.11E-05 | 4.66E-03 | (A) NE (B/D) |
| SSSCA1-AS1 | 4.43 | 2.76 | 0.59 | 0.51 | 1.67 | 8.13E-05 | 7.64E-03 | (A) NE (B/D) |
| KEAP1 | 1.98 | 0.31 | 0.35 | 0.5 | 1.67 | 8.21E-05 | 7.64E-03 | (A) NE (B/D) |
| PDXK | 7.23 | 5.55 | 0.32 | 0.57 | 1.67 | 3.12E-04 | 0.0145 | (A) NE (B/D) |
| EVI5L | 3.08 | 1.41 | 0.33 | 0.58 | 1.67 | 3.16E-04 | 0.0145 | (A) NE (B/D) |
| ZNF322 | 7.43 | 5.76 | 0.54 | 0.66 | 1.67 | 5.19E-04 | 0.0185 | (A) NE (B/D) |
| NR4A1 | 2.05 | 0.38 | 0.37 | 0.7 | 1.67 | 1.12E-03 | 0.0265 | (A) NE (B/D) |
| IKBKB | 6.56 | 4.9 | 0.55 | 0.74 | 1.67 | 1.25E-03 | 0.0279 | (A) NE (B/D) |
| LTBP3 | 1.87 | 0.21 | 1.04 | 0.5 | 1.67 | 1.35E-03 | 0.0292 | (A) NE (B/D) |
| SURF6 | 9.36 | 7.69 | 0.39 | 0.77 | 1.67 | 1.94E-03 | 0.0351 | (A) NE (B/D) |
| C19ORF57 | 2.52 | 0.85 | 0.55 | 0.82 | 1.67 | 2.30E-03 | 0.0379 | (A) NE (B/D) |
| STARD3 | 7.92 | 6.25 | 0.3 | 0.83 | 1.67 | 3.32E-03 | 0.0444 | (A) NE (B/D) |
| TAOK2 | 2.12 | 0.45 | 0.51 | 0.87 | 1.67 | 3.41E-03 | 0.0452 | (A) NE (B/D) |
| TMEM69 | 6.63 | 4.95 | 0.46 | 0.89 | 1.67 | 3.97E-03 | 0.0484 | (A) NE (B/D) |
| CRB3 | 2.34 | 0.68 | 0.44 | 0.54 | 1.66 | 1.33E-04 | 9.26E-03 | (A) NE (B/D) |
| ALKBH7 | 1.86 | 0.19 | 0.84 | 0.31 | 1.66 | 2.17E-04 | 0.012 | (A) NE (B/D) |
| NUDCD3 | 8.52 | 6.86 | 0.15 | 0.58 | 1.66 | 7.40E-04 | 0.022 | (A) NE (B/D) |
| ABCF1 | 2.03 | 0.37 | 0.31 | 0.66 | 1.66 | 9.12E-04 | 0.0245 | (A) NE (B/D) |
| MIR-7-1/1* | 5.39 | 3.74 | 1.05 | 0.59 | 1.66 | 1.86E-03 | 0.0345 | (A) NE (B/D) |
| MIR-7-1/3P | 5.39 | 3.74 | 1.05 | 0.59 | 1.66 | 1.86E-03 | 0.0345 | (A) NE (B/D) |
| MIR-7-1/5P | 5.39 | 3.74 | 1.05 | 0.59 | 1.66 | 1.86E-03 | 0.0345 | (A) NE (B/D) |
| MIR-7-1/7 | 5.39 | 3.74 | 1.05 | 0.59 | 1.66 | 1.86E-03 | 0.0345 | (A) NE (B/D) |
| MRI1 | 2.31 | 0.65 | 0.37 | 0.76 | 1.66 | 1.94E-03 | 0.0351 | (A) NE (B/D) |
| VPS51 | 2.95 | 1.3 | 0.92 | 0.75 | 1.66 | 2.36E-03 | 0.0383 | (A) NE (B/D) |
| PPP1R26-AS1 | 4.03 | 2.37 | 0.69 | 0.84 | 1.66 | 2.87E-03 | 0.0417 | (A) NE (B/D) |
| OXLD1 | 7.13 | 5.47 | 0.5 | 0.87 | 1.66 | 3.73E-03 | 0.0469 | (A) NE (B/D) |
| LRRFIP1P1 | 6.86 | 5.2 | 0.84 | 0.88 | 1.66 | 4.10E-03 | 0.0489 | (A) NE (B/D) |
| GAK | 1.89 | 0.24 | 0.58 | 0.6 | 1.65 | 2.92E-04 | 0.0142 | (A) NE (B/D) |
| NFRKB | 4.69 | 3.03 | 0.65 | 0.63 | 1.65 | 4.40E-04 | 0.0173 | (A) NE (B/D) |
| KDM4A-AS1 | 3.12 | 1.48 | 0.27 | 0.6 | 1.65 | 6.05E-04 | 0.0196 | (A) NE (B/D) |
| TENT4A | 1.95 | 0.3 | 0.54 | 0.67 | 1.65 | 6.24E-04 | 0.0199 | (A) NE (B/D) |
| LRRC41 | 3.01 | 1.36 | 0.29 | 0.63 | 1.65 | 7.31E-04 | 0.0218 | (A) NE (B/D) |
| CTDNEP1 | 2.05 | 0.4 | 0.19 | 0.61 | 1.65 | 8.84E-04 | 0.0241 | (A) NE (B/D) |

| | | | | | | | | |
|---|---|---|---|---|---|---|---|---|
| PIF1 | 1.88 | 0.24 | 1 | 0.42 | 1.65 | 9.63E-04 | 0.0251 | (A) NE (B/D) |
| CA5BP1 | 3.31 | 1.66 | 0.46 | 0.7 | 1.65 | 1.02E-03 | 0.0255 | (A) NE (B/D) |
| SIRT2 | 1.76 | 0.11 | 1.03 | 0.19 | 1.65 | 1.21E-03 | 0.0275 | (A) NE (B/D) |
| ZSWIM1 | 6.86 | 5.22 | 0.26 | 0.72 | 1.65 | 1.84E-03 | 0.0344 | (A) NE (B/D) |
| TBKBP1 | 3.35 | 1.7 | 1.07 | 0.56 | 1.65 | 1.97E-03 | 0.0353 | (A) NE (B/D) |
| CENPS | 3.12 | 1.47 | 0.39 | 0.78 | 1.65 | 2.33E-03 | 0.0381 | (A) NE (B/D) |
| HEXD | 2.04 | 0.4 | 0.69 | 0.81 | 1.65 | 2.33E-03 | 0.0382 | (A) NE (B/D) |
| ZNF688 | 2.59 | 0.94 | 0.32 | 0.79 | 1.65 | 2.58E-03 | 0.0397 | (A) NE (B/D) |
| SENP3-EIF4A | 9.59 | 7.94 | 0.31 | 0.82 | 1.65 | 3.45E-03 | 0.0453 | (A) NE (B/D) |
| HNRNPLP2 | 6.61 | 4.96 | 0.32 | 0.84 | 1.65 | 3.77E-03 | 0.0472 | (A) NE (B/D) |
| TRIM60P18 | 4.91 | 3.26 | 1.11 | 0.73 | 1.65 | 4.08E-03 | 0.0488 | (A) NE (B/D) |
| DUSP7 | 3.73 | 2.08 | 0.76 | 0.44 | 1.64 | 1.54E-04 | 0.01 | (A) NE (B/D) |
| XRCC3 | 5.34 | 3.7 | 0.49 | 0.56 | 1.64 | 1.70E-04 | 0.0106 | (A) NE (B/D) |
| NEURL2 | 5.29 | 3.65 | 0.36 | 0.54 | 1.64 | 1.97E-04 | 0.0113 | (A) NE (B/D) |
| B4GALT3 | 5.93 | 4.29 | 0.36 | 0.65 | 1.64 | 7.94E-04 | 0.0227 | (A) NE (B/D) |
| LGALSL | 1.87 | 0.23 | 1.1 | 0.25 | 1.64 | 1.82E-03 | 0.0342 | (A) NE (B/D) |
| NGDN | 6.6 | 4.97 | 0.74 | 0.84 | 1.64 | 3.10E-03 | 0.0429 | (A) NE (B/D) |
| ERN1 | 7.4 | 5.76 | 0.38 | 0.82 | 1.64 | 3.25E-03 | 0.0441 | (A) NE (B/D) |
| MAP3K6 | 1.77 | 0.14 | 0.66 | 0.29 | 1.63 | 3.41E-05 | 4.93E-03 | (A) NE (B/D) |
| OSGEPL1-AS | 4.68 | 3.05 | 0.92 | 0.52 | 1.63 | 7.67E-04 | 0.0223 | (A) NE (B/D) |
| ZFPL1 | 2.52 | 0.89 | 0.52 | 0.71 | 1.63 | 1.09E-03 | 0.0262 | (A) NE (B/D) |
| KDM4A | 4.91 | 3.28 | 0.33 | 0.68 | 1.63 | 1.18E-03 | 0.0272 | (A) NE (B/D) |
| MYO9B | 5.14 | 3.51 | 0.3 | 0.73 | 1.63 | 1.97E-03 | 0.0353 | (A) NE (B/D) |
| LRRC47 | 5.59 | 3.96 | 0.38 | 0.77 | 1.63 | 2.17E-03 | 0.0369 | (A) NE (B/D) |
| HCG25 | 2.23 | 0.6 | 0.27 | 0.78 | 1.63 | 2.82E-03 | 0.0413 | (A) NE (B/D) |
| TMEM175 | 2.58 | 0.95 | 0.88 | 0.8 | 1.63 | 3.09E-03 | 0.0429 | (A) NE (B/D) |
| CTBP2P7 | 1.8 | 0.16 | 1.21 | 0.34 | 1.63 | 3.52E-03 | 0.0457 | (A) NE (B/D) |
| SNHG10 | 5.7 | 4.07 | 0.73 | 0.88 | 1.63 | 4.15E-03 | 0.049 | (A) NE (B/D) |
| DNASE1 | 1.7 | 0.08 | 0.67 | 0.19 | 1.62 | 4.67E-05 | 5.63E-03 | (A) NE (B/D) |
| THAP8 | 2.23 | 0.61 | 0.44 | 0.53 | 1.62 | 1.28E-04 | 9.17E-03 | (A) NE (B/D) |
| NACC1 | 4.66 | 3.03 | 0.6 | 0.65 | 1.62 | 6.12E-04 | 0.0197 | (A) NE (B/D) |
| KDM5C | 6.41 | 4.78 | 0.33 | 0.63 | 1.62 | 7.20E-04 | 0.0217 | (A) NE (B/D) |
| GMEB2 | 1.88 | 0.27 | 0.42 | 0.65 | 1.62 | 7.48E-04 | 0.022 | (A) NE (B/D) |
| SMAGP | 3.2 | 1.58 | 0.63 | 0.71 | 1.62 | 1.17E-03 | 0.0271 | (A) NE (B/D) |
| ARHGDIA | 5.5 | 3.88 | 0.67 | 0.8 | 1.62 | 2.39E-03 | 0.0384 | (A) NE (B/D) |
| ABL1 | 3.04 | 1.42 | 0.36 | 0.78 | 1.62 | 2.54E-03 | 0.0395 | (A) NE (B/D) |
| MATN1-AS1 | 4.4 | 2.78 | 0.86 | 0.8 | 1.62 | 3.12E-03 | 0.043 | (A) NE (B/D) |
| NLRP1 | 1.84 | 0.21 | 1.22 | 0.37 | 1.62 | 3.86E-03 | 0.0478 | (A) NE (B/D) |
| LYPD5 | 2.37 | 0.75 | 0.83 | 0.86 | 1.62 | 4.33E-03 | 0.0498 | (A) NE (B/D) |
| SLC10A3 | 1.94 | 0.32 | 0.42 | 0.55 | 1.61 | 1.98E-04 | 0.0113 | (A) NE (B/D) |
| SPATA24 | 2.36 | 0.75 | 0.81 | 0.54 | 1.61 | 4.65E-04 | 0.0175 | (A) NE (B/D) |
| NOC2L | 2.01 | 0.4 | 0.56 | 0.63 | 1.61 | 5.08E-04 | 0.0184 | (A) NE (B/D) |
| MPP3 | 4.62 | 3.01 | 0.54 | 0.68 | 1.61 | 8.33E-04 | 0.0234 | (A) NE (B/D) |
| RAD51-AS1 | 3.02 | 1.4 | 0.89 | 0.73 | 1.61 | 2.34E-03 | 0.0382 | (A) NE (B/D) |
| RBM5 | 6.16 | 4.55 | 0.5 | 0.41 | 1.6 | 1.73E-05 | 3.38E-03 | (A) NE (B/D) |
| MAPK12 | 1.81 | 0.21 | 0.4 | 0.44 | 1.6 | 3.02E-05 | 4.57E-03 | (A) NE (B/D) |
| PIGL | 2.13 | 0.53 | 0.51 | 0.47 | 1.6 | 5.07E-05 | 5.87E-03 | (A) NE (B/D) |
| MIR-3605/3F | 6.92 | 5.32 | 0.66 | 0.57 | 1.6 | 3.31E-04 | 0.0148 | (A) NE (B/D) |
| MIR-3605/5F | 6.92 | 5.32 | 0.66 | 0.57 | 1.6 | 3.31E-04 | 0.0148 | (A) NE (B/D) |
| FNBP1 | 7.6 | 6.01 | 0.5 | 0.63 | 1.6 | 5.57E-04 | 0.019 | (A) NE (B/D) |
| CIRBP-AS1 | 2.36 | 0.76 | 0.67 | 0.66 | 1.6 | 8.19E-04 | 0.0231 | (A) NE (B/D) |
| IRF2BP2 | 6.37 | 4.77 | 0.66 | 0.72 | 1.6 | 1.37E-03 | 0.0296 | (A) NE (B/D) |
| TMEM164 | 2.19 | 0.59 | 0.74 | 0.72 | 1.6 | 1.49E-03 | 0.0308 | (A) NE (B/D) |
| RRP1 | 3.8 | 2.2 | 0.25 | 0.68 | 1.6 | 1.51E-03 | 0.031 | (A) NE (B/D) |
| HARS | 6.71 | 5.11 | 0.47 | 0.78 | 1.6 | 2.26E-03 | 0.0376 | (A) NE (B/D) |
| DPM2 | 2.94 | 1.34 | 0.56 | 0.79 | 1.6 | 2.35E-03 | 0.0383 | (A) NE (B/D) |
| HERC2P7 | 6.03 | 4.43 | 0.84 | 0.76 | 1.6 | 2.53E-03 | 0.0394 | (A) NE (B/D) |
| ZNF846 | 4.82 | 3.21 | 1.14 | 0.32 | 1.6 | 2.70E-03 | 0.0404 | (A) NE (B/D) |
| UBE2M | 2.24 | 0.64 | 0.34 | 0.79 | 1.6 | 3.04E-03 | 0.0426 | (A) NE (B/D) |
| ELP1 | 8.77 | 7.17 | 0.79 | 0.8 | 1.6 | 3.07E-03 | 0.0427 | (A) NE (B/D) |
| RPS18P9 | 2.1 | 0.49 | 1.07 | 0.68 | 1.6 | 3.57E-03 | 0.0459 | (A) NE (B/D) |
| YARS | 7.08 | 5.48 | 0.54 | 0.84 | 1.6 | 3.59E-03 | 0.0461 | (A) NE (B/D) |
| UPP1 | 3.65 | 2.05 | 0.36 | 0.83 | 1.6 | 3.83E-03 | 0.0476 | (A) NE (B/D) |
| MED12 | 1.63 | 0.04 | 0.38 | 0.09 | 1.59 | 5.26E-07 | 4.22E-04 | (A) NE (B/D) |
| STAG3L2 | 2.61 | 1.02 | 0.5 | 0.37 | 1.59 | 9.81E-06 | 2.34E-03 | (A) NE (B/D) |

| | | | | | | | | |
|---|---|---|---|---|---|---|---|---|
| ZNF514 | 2.65 | 1.06 | 0.42 | 0.64 | 1.59 | 7.09E-04 | 0.0215 | (A) NE (B/D) |
| DTNBP1 | 2.5 | 0.91 | 0.34 | 0.64 | 1.59 | 8.66E-04 | 0.0238 | (A) NE (B/D) |
| SLX1B | 2.9 | 1.3 | 0.73 | 0.68 | 1.59 | 1.14E-03 | 0.0268 | (A) NE (B/D) |
| YY1AP1 | 3.09 | 1.49 | 0.43 | 0.71 | 1.59 | 1.43E-03 | 0.0303 | (A) NE (B/D) |
| LINC00999 | 3.86 | 2.27 | 0.62 | 0.76 | 1.59 | 1.97E-03 | 0.0353 | (A) NE (B/D) |
| CDKN2C | 1.82 | 0.23 | 1.03 | 0.57 | 1.59 | 2.12E-03 | 0.0365 | (A) NE (B/D) |
| TBC1D10B | 5.68 | 4.1 | 0.57 | 0.79 | 1.59 | 2.42E-03 | 0.0385 | (A) NE (B/D) |
| USF2 | 1.81 | 0.24 | 0.43 | 0.43 | 1.58 | 2.39E-05 | 4.09E-03 | (A) NE (B/D) |
| CLK3 | 1.93 | 0.36 | 0.43 | 0.5 | 1.58 | 9.57E-05 | 8.02E-03 | (A) NE (B/D) |
| RAB17 | 1.78 | 0.2 | 0.63 | 0.49 | 1.58 | 1.29E-04 | 9.17E-03 | (A) NE (B/D) |
| SPIN3 | 2.23 | 0.65 | 0.66 | 0.54 | 1.58 | 2.46E-04 | 0.013 | (A) NE (B/D) |
| SCARNA15 | 2.89 | 1.3 | 1.05 | 0.5 | 1.58 | 2.09E-03 | 0.0362 | (A) NE (B/D) |
| FAM104A | 8.15 | 6.57 | 0.49 | 0.76 | 1.58 | 2.21E-03 | 0.0372 | (A) NE (B/D) |
| CAB39L | 3.1 | 1.52 | 0.88 | 0.75 | 1.58 | 2.98E-03 | 0.0423 | (A) NE (B/D) |
| PTBP1 | 10.09 | 8.51 | 0.23 | 0.78 | 1.58 | 3.70E-03 | 0.0466 | (A) NE (B/D) |
| PRR14 | 1.93 | 0.35 | 0.37 | 0.83 | 1.58 | 4.03E-03 | 0.0486 | (A) NE (B/D) |
| ZBED6CL | 3.17 | 1.6 | 0.5 | 0.3 | 1.57 | 4.37E-06 | 1.50E-03 | (A) NE (B/D) |
| BCL9 | 1.66 | 0.09 | 0.55 | 0.17 | 1.57 | 1.11E-05 | 2.53E-03 | (A) NE (B/D) |
| NCKIPSD | 1.89 | 0.33 | 0.68 | 0.34 | 1.57 | 6.94E-05 | 6.94E-03 | (A) NE (B/D) |
| MAP1S | 1.75 | 0.18 | 0.64 | 0.44 | 1.57 | 8.22E-05 | 7.64E-03 | (A) NE (B/D) |
| KLHL31 | 2.61 | 1.04 | 0.7 | 0.39 | 1.57 | 9.33E-05 | 7.97E-03 | (A) NE (B/D) |
| DHDH | 1.86 | 0.29 | 0.73 | 0.36 | 1.57 | 1.21E-04 | 9.03E-03 | (A) NE (B/D) |
| CRAMP1 | 1.98 | 0.41 | 0.39 | 0.6 | 1.57 | 5.14E-04 | 0.0185 | (A) NE (B/D) |
| FBXL19 | 2.31 | 0.75 | 0.6 | 0.63 | 1.57 | 5.96E-04 | 0.0194 | (A) NE (B/D) |
| ZNF786 | 1.94 | 0.37 | 0.57 | 0.63 | 1.57 | 6.25E-04 | 0.0199 | (A) NE (B/D) |
| MIR-4517/45 | 4.38 | 2.81 | 0.5 | 0.66 | 1.57 | 8.54E-04 | 0.0236 | (A) NE (B/D) |
| ZNF234 | 6.05 | 4.48 | 1.01 | 0.34 | 1.57 | 1.34E-03 | 0.0292 | (A) NE (B/D) |
| CCNB1 | 1.72 | 0.15 | 1.02 | 0.28 | 1.57 | 1.52E-03 | 0.031 | (A) NE (B/D) |
| FAM219B | 1.89 | 0.33 | 0.54 | 0.77 | 1.57 | 2.38E-03 | 0.0384 | (A) NE (B/D) |
| LRIG1 | 2.3 | 0.73 | 0.77 | 0.77 | 1.57 | 2.61E-03 | 0.0399 | (A) NE (B/D) |
| ZNF440 | 6.44 | 4.87 | 1.13 | 0.44 | 1.57 | 3.06E-03 | 0.0427 | (A) NE (B/D) |
| SRRM5 | 2.98 | 1.41 | 0.7 | 0.82 | 1.57 | 3.47E-03 | 0.0454 | (A) NE (B/D) |
| ILVBL | 1.6 | 0.04 | 0.51 | 0.11 | 1.56 | 9.79E-06 | 2.34E-03 | (A) NE (B/D) |
| NT5C3B | 1.69 | 0.13 | 0.57 | 0.2 | 1.56 | 1.31E-05 | 2.81E-03 | (A) NE (B/D) |
| TM9SF4 | 2.02 | 0.46 | 0.24 | 0.58 | 1.56 | 6.97E-04 | 0.0213 | (A) NE (B/D) |
| SNORD32A | 5.34 | 3.78 | 0.8 | 0.58 | 1.56 | 7.84E-04 | 0.0225 | (A) NE (B/D) |
| NR6A1 | 2.18 | 0.62 | 0.74 | 0.64 | 1.56 | 1.02E-03 | 0.0255 | (A) NE (B/D) |
| TARBP2 | 1.99 | 0.43 | 0.36 | 0.76 | 1.56 | 2.71E-03 | 0.0404 | (A) NE (B/D) |
| TMEM234 | 3.57 | 2.01 | 0.55 | 0.83 | 1.56 | 3.74E-03 | 0.047 | (A) NE (B/D) |
| DKC1 | 3.55 | 1.99 | 0.35 | 0.82 | 1.56 | 4.17E-03 | 0.049 | (A) NE (B/D) |
| LINC00649 | 2 | 0.44 | 0.44 | 0.43 | 1.55 | 2.72E-05 | 4.36E-03 | (A) NE (B/D) |
| RABGGTA | 1.73 | 0.18 | 0.61 | 0.34 | 1.55 | 3.03E-05 | 4.57E-03 | (A) NE (B/D) |
| GYS1 | 2.48 | 0.92 | 0.37 | 0.48 | 1.55 | 9.64E-05 | 8.02E-03 | (A) NE (B/D) |
| CRB2 | 3.16 | 1.61 | 0.49 | 0.52 | 1.55 | 1.53E-04 | 1.00E-02 | (A) NE (B/D) |
| PGBD2 | 6.67 | 5.12 | 0.62 | 0.52 | 1.55 | 1.99E-04 | 0.0113 | (A) NE (B/D) |
| ACOT8 | 4.98 | 3.43 | 0.46 | 0.56 | 1.55 | 2.98E-04 | 0.0142 | (A) NE (B/D) |
| IMPDH1 | 1.77 | 0.22 | 0.74 | 0.53 | 1.55 | 4.08E-04 | 0.0166 | (A) NE (B/D) |
| LINC01006 | 3.01 | 1.46 | 0.69 | 0.68 | 1.55 | 1.23E-03 | 0.0276 | (A) NE (B/D) |
| SNORD58B | 6.9 | 5.35 | 0.59 | 0.71 | 1.55 | 1.48E-03 | 0.0308 | (A) NE (B/D) |
| DLGAP1-AS1 | 1.8 | 0.26 | 1 | 0.41 | 1.55 | 1.50E-03 | 0.031 | (A) NE (B/D) |
| PPIL6 | 5.14 | 3.59 | 0.64 | 0.76 | 1.55 | 2.24E-03 | 0.0375 | (A) NE (B/D) |
| CNPY4 | 3.87 | 2.31 | 0.45 | 0.78 | 1.55 | 2.78E-03 | 0.041 | (A) NE (B/D) |
| SH3BP5L | 2.39 | 0.85 | 0.19 | 0.78 | 1.55 | 4.22E-03 | 0.0493 | (A) NE (B/D) |
| UBALD2 | 1.86 | 0.32 | 0.43 | 0.34 | 1.54 | 4.53E-06 | 1.51E-03 | (A) NE (B/D) |
| STAG3L3 | 3.43 | 1.89 | 0.65 | 0.48 | 1.54 | 1.67E-04 | 0.0106 | (A) NE (B/D) |
| EBF4 | 1.65 | 0.11 | 0.9 | 0.26 | 1.54 | 7.00E-04 | 0.0214 | (A) NE (B/D) |
| DIP2A | 5.49 | 3.96 | 0.41 | 0.66 | 1.54 | 1.15E-03 | 0.0268 | (A) NE (B/D) |
| ZNF316 | 2.68 | 1.15 | 0.96 | 0.37 | 1.54 | 1.18E-03 | 0.0272 | (A) NE (B/D) |
| ANKRD39 | 5.85 | 4.32 | 0.25 | 0.64 | 1.54 | 1.37E-03 | 0.0296 | (A) NE (B/D) |
| SH3KBP1 | 2.56 | 1.02 | 0.89 | 0.68 | 1.54 | 2.40E-03 | 0.0385 | (A) NE (B/D) |
| DELE1 | 2.17 | 0.63 | 0.66 | 0.77 | 1.54 | 2.65E-03 | 0.04 | (A) NE (B/D) |
| KRT8P11 | 1.8 | 0.26 | 1.09 | 0.53 | 1.54 | 3.20E-03 | 0.0437 | (A) NE (B/D) |
| PRMT2 | 2.08 | 0.54 | 0.49 | 0.8 | 1.54 | 3.30E-03 | 0.0442 | (A) NE (B/D) |
| TTC28-AS1 | 2.47 | 0.93 | 0.83 | 0.79 | 1.54 | 3.86E-03 | 0.0478 | (A) NE (B/D) |
| MAPK8IP3 | 1.88 | 0.34 | 0.47 | 0.83 | 1.54 | 4.12E-03 | 0.0489 | (A) NE (B/D) |

| | | | | | | | | |
|---|---|---|---|---|---|---|---|---|
| PSMG2 | 7.43 | 5.89 | 0.56 | 0.84 | 1.54 | 4.18E-03 | 0.049 | (A) NE (B/D) |
| GTPBP2 | 1.73 | 0.2 | 0.33 | 0.49 | 1.53 | 1.55E-04 | 0.0101 | (A) NE (B/D) |
| DGKQ | 1.65 | 0.12 | 0.77 | 0.29 | 1.53 | 2.27E-04 | 0.0123 | (A) NE (B/D) |
| RPS6KB2 | 3.82 | 2.3 | 0.5 | 0.71 | 1.53 | 1.63E-03 | 0.0322 | (A) NE (B/D) |
| C16ORF46 | 1.94 | 0.42 | 0.68 | 0.71 | 1.53 | 1.80E-03 | 0.034 | (A) NE (B/D) |
| CNNM4 | 5.42 | 3.88 | 0.31 | 0.7 | 1.53 | 2.00E-03 | 0.0354 | (A) NE (B/D) |
| ROMO1 | 2.32 | 0.79 | 0.32 | 0.71 | 1.53 | 2.12E-03 | 0.0365 | (A) NE (B/D) |
| DHX16 | 6.33 | 4.8 | 0.26 | 0.72 | 1.53 | 2.62E-03 | 0.04 | (A) NE (B/D) |
| AMIGO3 | 3.68 | 2.15 | 0.62 | 0.77 | 1.53 | 2.70E-03 | 0.0404 | (A) NE (B/D) |
| QARS | 2.55 | 1.01 | 0.62 | 0.8 | 1.53 | 3.29E-03 | 0.0442 | (A) NE (B/D) |
| UBE2V1 | 2.19 | 0.66 | 0.23 | 0.75 | 1.53 | 3.44E-03 | 0.0452 | (A) NE (B/D) |
| OCRL | 2.9 | 1.37 | 0.61 | 0.82 | 1.53 | 3.85E-03 | 0.0478 | (A) NE (B/D) |
| MYO1C | 3.33 | 1.8 | 0.34 | 0.79 | 1.53 | 3.92E-03 | 0.0481 | (A) NE (B/D) |
| ZSWIM5 | 1.91 | 0.39 | 0.57 | 0.47 | 1.52 | 9.87E-05 | 8.13E-03 | (A) NE (B/D) |
| ANKS3 | 3.7 | 2.17 | 0.56 | 0.56 | 1.52 | 3.23E-04 | 0.0147 | (A) NE (B/D) |
| C1ORF174 | 2.19 | 0.67 | 0.43 | 0.58 | 1.52 | 4.61E-04 | 0.0175 | (A) NE (B/D) |
| CAPN10 | 1.78 | 0.26 | 0.42 | 0.64 | 1.52 | 9.33E-04 | 0.0247 | (A) NE (B/D) |
| ZNF350 | 3.52 | 2 | 0.97 | 0.36 | 1.52 | 1.38E-03 | 0.0296 | (A) NE (B/D) |
| AMH | 1.75 | 0.23 | 0.93 | 0.52 | 1.52 | 1.46E-03 | 0.0307 | (A) NE (B/D) |
| GPR107 | 7.27 | 5.75 | 0.42 | 0.72 | 1.52 | 2.11E-03 | 0.0365 | (A) NE (B/D) |
| CCZ1B | 2.64 | 1.12 | 0.37 | 0.71 | 1.52 | 2.12E-03 | 0.0365 | (A) NE (B/D) |
| BTF3P7 | 2.16 | 0.64 | 0.73 | 0.75 | 1.52 | 2.70E-03 | 0.0404 | (A) NE (B/D) |
| AARS2 | 1.55 | 0.04 | 0.39 | 0.1 | 1.51 | 8.59E-07 | 5.91E-04 | (A) NE (B/D) |
| ZNF746 | 6.58 | 5.07 | 0.42 | 0.29 | 1.51 | 1.72E-06 | 8.29E-04 | (A) NE (B/D) |
| B9D2 | 1.54 | 0.03 | 0.4 | 0.07 | 1.51 | 2.06E-06 | 8.29E-04 | (A) NE (B/D) |
| UBE2O | 1.73 | 0.22 | 0.36 | 0.54 | 1.51 | 3.41E-04 | 0.015 | (A) NE (B/D) |
| KDM6B | 1.98 | 0.46 | 0.55 | 0.65 | 1.51 | 9.86E-04 | 0.0253 | (A) NE (B/D) |
| ZNF707 | 3.57 | 2.06 | 0.63 | 0.7 | 1.51 | 1.66E-03 | 0.0324 | (A) NE (B/D) |
| TMEM160 | 1.77 | 0.26 | 1 | 0.41 | 1.51 | 1.79E-03 | 0.0339 | (A) NE (B/D) |
| NEIL1 | 1.56 | 0.06 | 1 | 0.14 | 1.51 | 1.91E-03 | 0.0348 | (A) NE (B/D) |
| IPO4 | 1.86 | 0.35 | 0.83 | 0.68 | 1.51 | 2.23E-03 | 0.0374 | (A) NE (B/D) |
| GNL2 | 8.6 | 7.08 | 0.45 | 0.81 | 1.51 | 4.16E-03 | 0.049 | (A) NE (B/D) |
| PER2 | 1.63 | 0.13 | 0.49 | 0.27 | 1.5 | 4.29E-06 | 1.50E-03 | (A) NE (B/D) |
| SDCBP | 1.66 | 0.16 | 0.98 | 0.39 | 1.5 | 1.57E-03 | 0.0314 | (A) NE (B/D) |
| ZNF419 | 1.59 | 0.09 | 1.01 | 0.22 | 1.5 | 1.96E-03 | 0.0351 | (A) NE (B/D) |
| DUSP8P5 | 6.54 | 5.03 | 0.91 | 0.63 | 1.5 | 2.23E-03 | 0.0374 | (A) NE (B/D) |
| LENG8-AS1 | 7.92 | 6.43 | 0.68 | 0.73 | 1.5 | 2.46E-03 | 0.0388 | (A) NE (B/D) |
| CANT1 | 1.94 | 0.44 | 0.56 | 0.75 | 1.5 | 2.61E-03 | 0.0399 | (A) NE (B/D) |
| PGLS | 1.51 | 0.01 | 0.62 | 0.04 | 1.49 | 8.25E-05 | 7.64E-03 | (A) NE (B/D) |
| RFX2 | 1.59 | 0.1 | 0.71 | 0.24 | 1.49 | 1.40E-04 | 9.53E-03 | (A) NE (B/D) |
| PPP1R35 | 1.59 | 0.11 | 0.31 | 0.27 | 1.48 | 4.06E-07 | 4.22E-04 | (A) NE (B/D) |
| PNKP | 1.63 | 0.15 | 0.42 | 0.32 | 1.48 | 3.95E-06 | 1.42E-03 | (A) NE (B/D) |
| SLC35E1 | 8.21 | 6.73 | 0.5 | 0.49 | 1.48 | 1.42E-04 | 9.54E-03 | (A) NE (B/D) |
| PPP1R37 | 1.64 | 0.16 | 0.72 | 0.26 | 1.48 | 1.60E-04 | 0.0103 | (A) NE (B/D) |
| ZNF322P1 | 5.35 | 3.88 | 0.4 | 0.52 | 1.48 | 2.43E-04 | 0.0129 | (A) NE (B/D) |
| TOR2A | 1.78 | 0.3 | 0.61 | 0.57 | 1.48 | 4.83E-04 | 0.0178 | (A) NE (B/D) |
| USP36 | 3.88 | 2.4 | 0.21 | 0.67 | 1.48 | 2.27E-03 | 0.0376 | (A) NE (B/D) |
| KIF3B | 6.39 | 4.91 | 0.34 | 0.73 | 1.48 | 2.98E-03 | 0.0423 | (A) NE (B/D) |
| PWP2 | 1.6 | 0.13 | 0.34 | 0.3 | 1.47 | 1.37E-06 | 7.73E-04 | (A) NE (B/D) |
| C19ORF44 | 1.7 | 0.23 | 0.32 | 0.38 | 1.47 | 1.91E-05 | 3.50E-03 | (A) NE (B/D) |
| FAM27E3 | 1.9 | 0.42 | 0.6 | 0.24 | 1.47 | 3.43E-05 | 4.93E-03 | (A) NE (B/D) |
| MMEL1 | 1.72 | 0.25 | 0.67 | 0.48 | 1.47 | 2.56E-04 | 0.0134 | (A) NE (B/D) |
| HTR5BP | 4.25 | 2.78 | 0.48 | 0.55 | 1.47 | 3.50E-04 | 0.0152 | (A) NE (B/D) |
| JSRP1 | 1.73 | 0.26 | 0.12 | 0.46 | 1.47 | 4.01E-04 | 0.0165 | (A) NE (B/D) |
| NAPA-AS1 | 6.97 | 5.5 | 0.17 | 0.55 | 1.47 | 9.45E-04 | 0.0249 | (A) NE (B/D) |
| ABHD4 | 1.82 | 0.35 | 0.8 | 0.57 | 1.47 | 1.15E-03 | 0.0268 | (A) NE (B/D) |
| GALNT10 | 1.49 | 0.02 | 1.01 | 0.04 | 1.47 | 2.42E-03 | 0.0385 | (A) NE (B/D) |
| PUSL1 | 4.09 | 2.61 | 0.6 | 0.77 | 1.47 | 3.41E-03 | 0.0452 | (A) NE (B/D) |
| CCDC12 | 2.17 | 0.7 | 0.5 | 0.78 | 1.47 | 3.84E-03 | 0.0477 | (A) NE (B/D) |
| KPNA6 | 6.13 | 4.67 | 0.7 | 0.37 | 1.46 | 1.76E-04 | 0.0107 | (A) NE (B/D) |
| NBPF2P | 4.98 | 3.51 | 0.75 | 0.51 | 1.46 | 6.03E-04 | 0.0195 | (A) NE (B/D) |
| ZNF142 | 7.2 | 5.73 | 0.35 | 0.6 | 1.46 | 8.43E-04 | 0.0236 | (A) NE (B/D) |
| PLD2 | 1.63 | 0.17 | 0.92 | 0.29 | 1.46 | 1.19E-03 | 0.0273 | (A) NE (B/D) |
| URGCP | 2.63 | 1.17 | 0.28 | 0.67 | 1.46 | 2.14E-03 | 0.0367 | (A) NE (B/D) |
| ZNF558 | 1.56 | 0.1 | 1.02 | 0.24 | 1.46 | 2.42E-03 | 0.0385 | (A) NE (B/D) |

| | | | | | | | | |
|---|---|---|---|---|---|---|---|---|
| RGS12 | 1.73 | 0.27 | 0.84 | 0.66 | 1.46 | 2.45E-03 | 0.0388 | (A) NE (B/D) |
| MIF4GD | 1.78 | 0.33 | 0.59 | 0.72 | 1.46 | 2.49E-03 | 0.0391 | (A) NE (B/D) |
| DPF2 | 2.52 | 1.06 | 0.36 | 0.77 | 1.46 | 4.03E-03 | 0.0486 | (A) NE (B/D) |
| NELFA | 2.83 | 1.38 | 0.42 | 0.78 | 1.46 | 4.08E-03 | 0.0488 | (A) NE (B/D) |
| WASH5P | 7.18 | 5.72 | 0.17 | 0.46 | 1.45 | 3.55E-04 | 0.0153 | (A) NE (B/D) |
| TP53I13 | 2.19 | 0.74 | 0.42 | 0.62 | 1.45 | 1.00E-03 | 0.0254 | (A) NE (B/D) |
| WBP1LP2 | 4.46 | 3 | 0.91 | 0.35 | 1.45 | 1.15E-03 | 0.0268 | (A) NE (B/D) |
| MIR-4467/44 | 1.62 | 0.17 | 0.91 | 0.41 | 1.45 | 1.33E-03 | 0.0289 | (A) NE (B/D) |
| TAF6 | 1.69 | 0.24 | 0.19 | 0.6 | 1.45 | 1.48E-03 | 0.0308 | (A) NE (B/D) |
| NCOR2 | 3.11 | 1.66 | 0.44 | 0.73 | 1.45 | 2.92E-03 | 0.0419 | (A) NE (B/D) |
| IPPK | 6.05 | 4.61 | 0.3 | 0.71 | 1.45 | 3.06E-03 | 0.0427 | (A) NE (B/D) |
| ARMC6 | 1.73 | 0.29 | 0.64 | 0.35 | 1.44 | 9.22E-05 | 7.97E-03 | (A) NE (B/D) |
| ZNF862 | 1.83 | 0.39 | 0.51 | 0.6 | 1.44 | 8.22E-04 | 0.0231 | (A) NE (B/D) |
| HERC3 | 3.35 | 1.91 | 0.46 | 0.61 | 1.44 | 8.50E-04 | 0.0236 | (A) NE (B/D) |
| SMIM7 | 5.51 | 4.07 | 0.4 | 0.6 | 1.44 | 9.04E-04 | 0.0244 | (A) NE (B/D) |
| PAAF1 | 7.23 | 5.79 | 0.39 | 0.64 | 1.44 | 1.32E-03 | 0.0288 | (A) NE (B/D) |
| RASSF1 | 4.75 | 3.31 | 0.27 | 0.68 | 1.44 | 2.48E-03 | 0.0391 | (A) NE (B/D) |
| USP19 | 3.62 | 2.18 | 0.51 | 0.73 | 1.44 | 2.85E-03 | 0.0416 | (A) NE (B/D) |
| CEP95 | 7.9 | 6.46 | 0.8 | 0.71 | 1.44 | 3.44E-03 | 0.0452 | (A) NE (B/D) |
| PCDHB19P | 1.51 | 0.07 | 1.09 | 0.17 | 1.44 | 3.99E-03 | 0.0484 | (A) NE (B/D) |
| RBM33 | 1.89 | 0.45 | 0.36 | 0.76 | 1.44 | 3.99E-03 | 0.0484 | (A) NE (B/D) |
| PPP4R1 | 1.76 | 0.33 | 0.38 | 0.45 | 1.43 | 9.58E-05 | 8.02E-03 | (A) NE (B/D) |
| ACOT7 | 1.47 | 0.04 | 0.63 | 0.09 | 1.43 | 1.16E-04 | 8.81E-03 | (A) NE (B/D) |
| RNF121 | 1.99 | 0.56 | 0.36 | 0.46 | 1.43 | 1.20E-04 | 9.03E-03 | (A) NE (B/D) |
| POMZP3 | 2.95 | 1.52 | 0.38 | 0.48 | 1.43 | 1.80E-04 | 0.0108 | (A) NE (B/D) |
| GPBP1 | 1.81 | 0.38 | 0.79 | 0.47 | 1.43 | 7.61E-04 | 0.0223 | (A) NE (B/D) |
| ZNF329 | 4.47 | 3.05 | 0.83 | 0.49 | 1.43 | 1.09E-03 | 0.0262 | (A) NE (B/D) |
| C4ORF47 | 4.11 | 2.68 | 0.51 | 0.66 | 1.43 | 1.57E-03 | 0.0314 | (A) NE (B/D) |
| GMPPB | 5.94 | 4.51 | 0.61 | 0.69 | 1.43 | 2.15E-03 | 0.0368 | (A) NE (B/D) |
| SNX8 | 1.77 | 0.35 | 0.98 | 0.38 | 1.43 | 2.27E-03 | 0.0376 | (A) NE (B/D) |
| ZNF615 | 3.8 | 2.37 | 1.04 | 0.2 | 1.43 | 3.11E-03 | 0.0429 | (A) NE (B/D) |
| IL10RB | 3.39 | 1.96 | 0.29 | 0.71 | 1.43 | 3.37E-03 | 0.0448 | (A) NE (B/D) |
| C1ORF52 | 1.84 | 0.42 | 0.63 | 0.77 | 1.43 | 4.05E-03 | 0.0487 | (A) NE (B/D) |
| ABCC10 | 1.42 | 0 | 0.61 | 0 | 1.42 | 1.15E-04 | 8.73E-03 | (A) NE (B/D) |
| KHSRPP1 | 4.81 | 3.39 | 0.4 | 0.51 | 1.42 | 2.87E-04 | 0.0142 | (A) NE (B/D) |
| KMT2E-AS1 | 4.7 | 3.28 | 0.19 | 0.59 | 1.42 | 1.46E-03 | 0.0307 | (A) NE (B/D) |
| PHF13 | 3.56 | 2.14 | 0.36 | 0.66 | 1.42 | 1.88E-03 | 0.0347 | (A) NE (B/D) |
| CCT6P3 | 6.95 | 5.53 | 0.94 | 0.62 | 1.42 | 3.70E-03 | 0.0466 | (A) NE (B/D) |
| UQCRB | 1.9 | 0.48 | 0.77 | 0.74 | 1.42 | 4.19E-03 | 0.0491 | (A) NE (B/D) |
| YIF1B | 3.39 | 1.97 | 0.37 | 0.76 | 1.42 | 4.32E-03 | 0.0498 | (A) NE (B/D) |
| ZNF202 | 1.62 | 0.21 | 0.68 | 0.52 | 1.41 | 5.90E-04 | 0.0194 | (A) NE (B/D) |
| RNA5-8SP2 | 1.42 | 0 | 0.97 | 0.01 | 1.41 | 2.35E-03 | 0.0383 | (A) NE (B/D) |
| STIM2 | 1.72 | 0.31 | 0.85 | 0.61 | 1.41 | 2.42E-03 | 0.0385 | (A) NE (B/D) |
| TFAP2E | 3.61 | 2.2 | 0.81 | 0.67 | 1.41 | 3.01E-03 | 0.0424 | (A) NE (B/D) |
| TSTD2 | 2.87 | 1.46 | 0.58 | 0.72 | 1.41 | 3.03E-03 | 0.0426 | (A) NE (B/D) |
| CWC22 | 2.8 | 1.38 | 1.02 | 0.49 | 1.41 | 3.72E-03 | 0.0468 | (A) NE (B/D) |
| LINC00663 | 4.51 | 3.1 | 1.05 | 0.42 | 1.41 | 3.95E-03 | 0.0483 | (A) NE (B/D) |
| ASB6 | 1.52 | 0.12 | 0.44 | 0.29 | 1.4 | 4.86E-06 | 1.58E-03 | (A) NE (B/D) |
| TAF8 | 5.81 | 4.41 | 0.4 | 0.32 | 1.4 | 5.89E-06 | 1.74E-03 | (A) NE (B/D) |
| UNC119 | 1.61 | 0.21 | 0.65 | 0.35 | 1.4 | 1.27E-04 | 9.14E-03 | (A) NE (B/D) |
| TRAF3IP1 | 11.13 | 9.73 | 0.62 | 0.47 | 1.4 | 2.91E-04 | 0.0142 | (A) NE (B/D) |
| SNORD36C | 7.51 | 6.11 | 0.52 | 0.72 | 1.4 | 2.92E-03 | 0.0419 | (A) NE (B/D) |
| DPP9 | 1.85 | 0.45 | 0.19 | 0.68 | 1.4 | 3.46E-03 | 0.0454 | (A) NE (B/D) |
| GPAT2 | 5.11 | 3.72 | 0.7 | 0.25 | 1.39 | 2.09E-04 | 0.0116 | (A) NE (B/D) |
| PKN1 | 1.59 | 0.2 | 0.66 | 0.49 | 1.39 | 4.48E-04 | 0.0174 | (A) NE (B/D) |
| FAM122C | 2.89 | 1.5 | 0.43 | 0.55 | 1.39 | 5.27E-04 | 0.0185 | (A) NE (B/D) |
| PTPRH | 1.46 | 0.07 | 0.79 | 0.17 | 1.39 | 6.46E-04 | 0.0203 | (A) NE (B/D) |
| B3GALT4 | 2.18 | 0.8 | 0.46 | 0.59 | 1.39 | 9.34E-04 | 0.0247 | (A) NE (B/D) |
| MTF1 | 2.78 | 1.39 | 0.44 | 0.61 | 1.39 | 1.19E-03 | 0.0273 | (A) NE (B/D) |
| NIF3L1 | 1.77 | 0.38 | 0.42 | 0.62 | 1.39 | 1.34E-03 | 0.0292 | (A) NE (B/D) |
| EIF4A1 | 3.06 | 1.66 | 0.26 | 0.6 | 1.39 | 1.51E-03 | 0.031 | (A) NE (B/D) |
| DAXX | 1.73 | 0.34 | 0.38 | 0.68 | 1.39 | 2.55E-03 | 0.0396 | (A) NE (B/D) |
| RNF216 | 4.74 | 3.36 | 0.39 | 0.32 | 1.38 | 5.74E-06 | 1.74E-03 | (A) NE (B/D) |
| ZBTB7A | 1.42 | 0.04 | 0.56 | 0.08 | 1.38 | 6.55E-05 | 6.86E-03 | (A) NE (B/D) |
| KDM5D | 1.44 | 0.05 | 0.58 | 0.12 | 1.38 | 6.75E-05 | 6.91E-03 | (A) NE (B/D) |

| | | | | | | | | |
|---|---|---|---|---|---|---|---|---|
| NCOA5 | 1.61 | 0.22 | 0.45 | 0.55 | 1.38 | 5.60E-04 | 0.019 | (A) NE (B/D) |
| UBE2SP2 | 1.56 | 0.18 | 0.78 | 0.44 | 1.38 | 8.01E-04 | 0.0228 | (A) NE (B/D) |
| ZNF425 | 3.27 | 1.89 | 0.57 | 0.58 | 1.38 | 9.17E-04 | 0.0245 | (A) NE (B/D) |
| STRIP1 | 1.61 | 0.23 | 0.21 | 0.55 | 1.38 | 1.09E-03 | 0.0262 | (A) NE (B/D) |
| PRPF8 | 5.38 | 4 | 0.52 | 0.62 | 1.38 | 1.36E-03 | 0.0294 | (A) NE (B/D) |
| PNPT1 | 1.51 | 0.13 | 0.91 | 0.31 | 1.38 | 1.63E-03 | 0.0321 | (A) NE (B/D) |
| HOXD-AS2 | 1.7 | 0.32 | 0.9 | 0.52 | 1.38 | 2.49E-03 | 0.0391 | (A) NE (B/D) |
| BMS1P9 | 3.31 | 1.92 | 0.58 | 0.73 | 1.38 | 3.51E-03 | 0.0457 | (A) NE (B/D) |
| PDCD6 | 1.73 | 0.35 | 0.63 | 0.72 | 1.38 | 3.61E-03 | 0.0463 | (A) NE (B/D) |
| PIP5K1C | 1.56 | 0.19 | 0.23 | 0.24 | 1.37 | 4.77E-07 | 4.22E-04 | (A) NE (B/D) |
| HELZ2 | 1.67 | 0.3 | 0.53 | 0.42 | 1.37 | 1.07E-04 | 8.41E-03 | (A) NE (B/D) |
| C8ORF44 | 2.9 | 1.53 | 0.62 | 0.37 | 1.37 | 1.48E-04 | 9.80E-03 | (A) NE (B/D) |
| YPEL3 | 1.56 | 0.19 | 0.5 | 0.46 | 1.37 | 1.73E-04 | 0.0106 | (A) NE (B/D) |
| ZNF561 | 2.8 | 1.43 | 0.53 | 0.54 | 1.37 | 5.85E-04 | 0.0193 | (A) NE (B/D) |
| ZNF841 | 2.52 | 1.14 | 0.83 | 0.31 | 1.37 | 9.19E-04 | 0.0246 | (A) NE (B/D) |
| WDR75 | 7.74 | 6.37 | 0.78 | 0.48 | 1.37 | 1.01E-03 | 0.0254 | (A) NE (B/D) |
| ZNF785 | 2.4 | 1.03 | 0.3 | 0.58 | 1.37 | 1.22E-03 | 0.0275 | (A) NE (B/D) |
| CYCSP55 | 1.87 | 0.49 | 0.91 | 0.46 | 1.37 | 2.21E-03 | 0.0372 | (A) NE (B/D) |
| CACNA2D2 | 1.63 | 0.26 | 0.94 | 0.41 | 1.37 | 2.27E-03 | 0.0376 | (A) NE (B/D) |
| CUL4A | 2.01 | 0.64 | 0.34 | 0.67 | 1.37 | 2.54E-03 | 0.0395 | (A) NE (B/D) |
| LIPT1 | 3.29 | 1.92 | 0.63 | 0.7 | 1.37 | 3.02E-03 | 0.0424 | (A) NE (B/D) |
| BCRP2 | 3.28 | 1.91 | 0.4 | 0.73 | 1.37 | 3.99E-03 | 0.0484 | (A) NE (B/D) |
| ZFYVE9P1 | 1.44 | 0.08 | 0.35 | 0.09 | 1.36 | 8.30E-07 | 5.88E-04 | (A) NE (B/D) |
| SNHG17 | 1.77 | 0.41 | 0.54 | 0.44 | 1.36 | 1.64E-04 | 0.0105 | (A) NE (B/D) |
| IP6K1 | 1.74 | 0.38 | 0.36 | 0.46 | 1.36 | 1.77E-04 | 0.0107 | (A) NE (B/D) |
| HMGXB3 | 6.79 | 5.44 | 0.6 | 0.53 | 1.36 | 6.09E-04 | 0.0197 | (A) NE (B/D) |
| MRPL48 | 1.71 | 0.35 | 0.47 | 0.55 | 1.36 | 6.26E-04 | 0.0199 | (A) NE (B/D) |
| MED22 | 3.27 | 1.9 | 0.33 | 0.59 | 1.36 | 1.31E-03 | 0.0287 | (A) NE (B/D) |
| HDAC5 | 2.25 | 0.89 | 0.33 | 0.69 | 1.36 | 3.25E-03 | 0.0441 | (A) NE (B/D) |
| ZSWIM7 | 1.46 | 0.09 | 1.01 | 0.22 | 1.36 | 3.49E-03 | 0.0456 | (A) NE (B/D) |
| MYLIP | 3.35 | 1.99 | 0.36 | 0.44 | 1.35 | 1.27E-04 | 9.14E-03 | (A) NE (B/D) |
| VAPA | 7.75 | 6.4 | 0.65 | 0.41 | 1.35 | 2.71E-04 | 0.0137 | (A) NE (B/D) |
| ARAP1-AS2 | 6.81 | 5.46 | 0.44 | 0.53 | 1.35 | 4.72E-04 | 0.0176 | (A) NE (B/D) |
| PKP4P1 | 1.87 | 0.52 | 0.57 | 0.53 | 1.35 | 5.76E-04 | 0.0192 | (A) NE (B/D) |
| CCNJL | 1.57 | 0.22 | 0.62 | 0.53 | 1.35 | 7.21E-04 | 0.0217 | (A) NE (B/D) |
| UBAP2L | 1.85 | 0.49 | 0.47 | 0.6 | 1.35 | 1.16E-03 | 0.027 | (A) NE (B/D) |
| NELFCD | 2.17 | 0.82 | 0.16 | 0.58 | 1.35 | 1.97E-03 | 0.0353 | (A) NE (B/D) |
| ARL2 | 2.05 | 0.7 | 0.75 | 0.61 | 1.35 | 2.28E-03 | 0.0376 | (A) NE (B/D) |
| FERP1 | 6.34 | 4.99 | 0.86 | 0.58 | 1.35 | 3.19E-03 | 0.0437 | (A) NE (B/D) |
| ARMCX6 | 1.82 | 0.47 | 0.23 | 0.67 | 1.35 | 3.33E-03 | 0.0445 | (A) NE (B/D) |
| C19ORF71 | 1.39 | 0.04 | 0.5 | 0.11 | 1.34 | 2.48E-05 | 4.17E-03 | (A) NE (B/D) |
| STAG3L1 | 2.46 | 1.12 | 0.53 | 0.3 | 1.34 | 3.32E-05 | 4.88E-03 | (A) NE (B/D) |
| FRG1BP | 1.7 | 0.36 | 0.6 | 0.41 | 1.34 | 1.84E-04 | 0.0109 | (A) NE (B/D) |
| MED28 | 1.48 | 0.14 | 0.69 | 0.33 | 1.34 | 2.82E-04 | 0.0141 | (A) NE (B/D) |
| CRTC2 | 1.66 | 0.32 | 0.34 | 0.51 | 1.34 | 4.70E-04 | 0.0176 | (A) NE (B/D) |
| CHST12 | 1.49 | 0.15 | 0.88 | 0.34 | 1.34 | 1.67E-03 | 0.0325 | (A) NE (B/D) |
| CIAO1 | 2.22 | 0.88 | 0.15 | 0.58 | 1.34 | 1.90E-03 | 0.0348 | (A) NE (B/D) |
| UBE2G2 | 3.6 | 2.26 | 0.41 | 0.64 | 1.34 | 2.10E-03 | 0.0363 | (A) NE (B/D) |
| KMT5A | 4.19 | 2.85 | 0.38 | 0.69 | 1.34 | 3.40E-03 | 0.0451 | (A) NE (B/D) |
| STK24 | 5.84 | 4.5 | 0.86 | 0.61 | 1.34 | 3.62E-03 | 0.0463 | (A) NE (B/D) |
| MED24 | 1.62 | 0.29 | 0.3 | 0.71 | 1.34 | 4.26E-03 | 0.0495 | (A) NE (B/D) |
| DENND1A | 3.9 | 2.57 | 0.41 | 0.24 | 1.33 | 2.94E-06 | 1.11E-03 | (A) NE (B/D) |
| GIT1 | 1.47 | 0.14 | 0.35 | 0.35 | 1.33 | 1.65E-05 | 3.33E-03 | (A) NE (B/D) |
| NARF | 1.49 | 0.15 | 0.37 | 0.37 | 1.33 | 3.21E-05 | 4.76E-03 | (A) NE (B/D) |
| WDR13 | 1.52 | 0.19 | 0.69 | 0.32 | 1.33 | 3.04E-04 | 0.0144 | (A) NE (B/D) |
| PM20D2 | 1.63 | 0.3 | 0.71 | 0.55 | 1.33 | 1.44E-03 | 0.0304 | (A) NE (B/D) |
| ZNF30 | 1.83 | 0.5 | 0.83 | 0.45 | 1.33 | 1.68E-03 | 0.0325 | (A) NE (B/D) |
| AFG3L2 | 1.89 | 0.56 | 0.48 | 0.62 | 1.33 | 1.69E-03 | 0.0328 | (A) NE (B/D) |
| DVL3 | 1.58 | 0.25 | 0.38 | 0.61 | 1.33 | 1.71E-03 | 0.0328 | (A) NE (B/D) |
| ARHGEF7 | 2.5 | 1.17 | 0.66 | 0.6 | 1.33 | 1.85E-03 | 0.0345 | (A) NE (B/D) |
| ZNF20 | 3.68 | 2.34 | 0.87 | 0.51 | 1.33 | 2.58E-03 | 0.0397 | (A) NE (B/D) |
| GYG1P1 | 1.79 | 0.45 | 0.71 | 0.7 | 1.33 | 4.08E-03 | 0.0488 | (A) NE (B/D) |
| C19ORF53 | 1.36 | 0.04 | 0.43 | 0.06 | 1.32 | 1.32E-05 | 2.81E-03 | (A) NE (B/D) |
| SLFNL1 | 4.21 | 2.89 | 0.73 | 0.46 | 1.32 | 9.09E-04 | 0.0245 | (A) NE (B/D) |
| SDCBPP3 | 3.98 | 2.66 | 0.47 | 0.61 | 1.32 | 1.54E-03 | 0.0311 | (A) NE (B/D) |

| | | | | | | | | |
|---|---|---|---|---|---|---|---|---|
| KDM3B | 2.33 | 1.01 | 0.57 | 0.61 | 1.32 | 1.66E-03 | 0.0324 | (A) NE (B/D) |
| TJP1 | 2 | 0.68 | 0.51 | 0.67 | 1.32 | 2.81E-03 | 0.0412 | (A) NE (B/D) |
| FAM71F2 | 1.82 | 0.5 | 0.84 | 0.6 | 1.32 | 3.68E-03 | 0.0466 | (A) NE (B/D) |
| APBA3 | 1.62 | 0.31 | 0.2 | 0.13 | 1.31 | 8.87E-10 | 1.04E-05 | (A) NE (B/D) |
| SNX21 | 1.38 | 0.08 | 0.34 | 0.13 | 1.31 | 5.41E-07 | 4.22E-04 | (A) NE (B/D) |
| DFFA | 6.37 | 5.06 | 0.4 | 0.37 | 1.31 | 3.99E-05 | 5.17E-03 | (A) NE (B/D) |
| CDK11A | 1.42 | 0.11 | 0.71 | 0.28 | 1.31 | 3.93E-04 | 0.0163 | (A) NE (B/D) |
| C1ORF35 | 1.85 | 0.54 | 0.29 | 0.49 | 1.31 | 4.50E-04 | 0.0175 | (A) NE (B/D) |
| RBAKDN | 1.48 | 0.17 | 0.76 | 0.43 | 1.31 | 9.48E-04 | 0.0249 | (A) NE (B/D) |
| METTL3 | 1.52 | 0.21 | 0.7 | 0.51 | 1.31 | 1.07E-03 | 0.0261 | (A) NE (B/D) |
| FAM131A | 1.4 | 0.08 | 0.9 | 0.2 | 1.31 | 2.21E-03 | 0.0372 | (A) NE (B/D) |
| JRKL | 2.95 | 1.64 | 0.87 | 0.46 | 1.31 | 2.44E-03 | 0.0387 | (A) NE (B/D) |
| MRPS24 | 2.11 | 0.8 | 0.52 | 0.68 | 1.31 | 3.22E-03 | 0.0439 | (A) NE (B/D) |
| PRCC | 4.31 | 3 | 0.64 | 0.68 | 1.31 | 3.66E-03 | 0.0464 | (A) NE (B/D) |
| CICP13 | 2.44 | 1.13 | 0.97 | 0.42 | 1.31 | 3.91E-03 | 0.0481 | (A) NE (B/D) |
| SGSH | 1.7 | 0.4 | 0.42 | 0.32 | 1.3 | 1.41E-05 | 2.93E-03 | (A) NE (B/D) |
| OAZ3 | 3.33 | 2.03 | 0.47 | 0.4 | 1.3 | 9.28E-05 | 7.97E-03 | (A) NE (B/D) |
| STX10 | 1.59 | 0.29 | 0.31 | 0.46 | 1.3 | 2.99E-04 | 0.0142 | (A) NE (B/D) |
| ARHGAP4 | 1.39 | 0.09 | 0.73 | 0.16 | 1.3 | 5.58E-04 | 0.019 | (A) NE (B/D) |
| NBR2 | 5.18 | 3.88 | 0.49 | 0.53 | 1.3 | 6.78E-04 | 0.0209 | (A) NE (B/D) |
| ATRIP | 2.01 | 0.71 | 0.74 | 0.43 | 1.3 | 9.25E-04 | 0.0247 | (A) NE (B/D) |
| HSF1 | 2.62 | 1.32 | 0.7 | 0.54 | 1.3 | 1.41E-03 | 0.0299 | (A) NE (B/D) |
| ETV2 | 2.19 | 0.89 | 0.46 | 0.6 | 1.3 | 1.60E-03 | 0.0318 | (A) NE (B/D) |
| BAP1 | 1.79 | 0.48 | 0.55 | 0.67 | 1.3 | 3.01E-03 | 0.0424 | (A) NE (B/D) |
| NPLOC4 | 1.63 | 0.34 | 0.41 | 0.68 | 1.3 | 3.52E-03 | 0.0457 | (A) NE (B/D) |
| GDPD5 | 1.93 | 0.63 | 0.64 | 0.67 | 1.3 | 3.53E-03 | 0.0457 | (A) NE (B/D) |
| MIR-1224/3P | 1.73 | 0.43 | 0.76 | 0.63 | 1.3 | 3.65E-03 | 0.0464 | (A) NE (B/D) |
| MIR-1224/5P | 1.73 | 0.43 | 0.76 | 0.63 | 1.3 | 3.65E-03 | 0.0464 | (A) NE (B/D) |
| PCMTD2 | 1.54 | 0.24 | 0.86 | 0.59 | 1.3 | 4.17E-03 | 0.049 | (A) NE (B/D) |
| PILRA | 1.38 | 0.07 | 1 | 0.18 | 1.3 | 4.26E-03 | 0.0495 | (A) NE (B/D) |
| AHCYL2 | 3.13 | 1.83 | 0.43 | 0.42 | 1.29 | 1.23E-04 | 9.09E-03 | (A) NE (B/D) |
| BGLAP | 1.42 | 0.13 | 0.61 | 0.26 | 1.29 | 1.36E-04 | 9.40E-03 | (A) NE (B/D) |
| ZNF200 | 3.08 | 1.79 | 0.54 | 0.41 | 1.29 | 1.81E-04 | 0.0108 | (A) NE (B/D) |
| SPIDR | 3.17 | 1.87 | 0.58 | 0.48 | 1.29 | 5.17E-04 | 0.0185 | (A) NE (B/D) |
| VWA5A | 1.41 | 0.12 | 0.74 | 0.19 | 1.29 | 6.36E-04 | 0.0201 | (A) NE (B/D) |
| EHMT1 | 2.23 | 0.94 | 0.45 | 0.54 | 1.29 | 8.17E-04 | 0.0231 | (A) NE (B/D) |
| VN2R19P | 1.36 | 0.07 | 0.77 | 0.08 | 1.29 | 1.01E-03 | 0.0254 | (A) NE (B/D) |
| WDTC1 | 2.24 | 0.95 | 0.45 | 0.57 | 1.29 | 1.15E-03 | 0.0268 | (A) NE (B/D) |
| TRIM39 | 3.43 | 2.14 | 0.11 | 0.52 | 1.29 | 1.54E-03 | 0.0311 | (A) NE (B/D) |
| NOL7 | 6.56 | 5.27 | 0.19 | 0.56 | 1.29 | 1.85E-03 | 0.0345 | (A) NE (B/D) |
| NUBP2 | 2.08 | 0.79 | 0.57 | 0.69 | 1.29 | 3.97E-03 | 0.0484 | (A) NE (B/D) |
| KRTAP5-1 | 1.41 | 0.14 | 0.47 | 0.33 | 1.28 | 3.43E-05 | 4.93E-03 | (A) NE (B/D) |
| CEP104 | 6.01 | 4.72 | 0.45 | 0.36 | 1.28 | 4.04E-05 | 5.17E-03 | (A) NE (B/D) |
| INHA | 1.31 | 0.03 | 0.54 | 0.05 | 1.28 | 8.67E-05 | 7.77E-03 | (A) NE (B/D) |
| PRPF31 | 1.57 | 0.29 | 0.29 | 0.43 | 1.28 | 1.99E-04 | 0.0113 | (A) NE (B/D) |
| ABCF2 | 1.29 | 0.01 | 0.68 | 0.03 | 1.28 | 4.61E-04 | 0.0175 | (A) NE (B/D) |
| ZNF761 | 1.29 | 0.01 | 0.7 | 0.03 | 1.28 | 5.84E-04 | 0.0193 | (A) NE (B/D) |
| BUD31 | 1.5 | 0.22 | 0.21 | 0.53 | 1.28 | 1.30E-03 | 0.0287 | (A) NE (B/D) |
| CRYZL1 | 1.36 | 0.08 | 0.89 | 0.21 | 1.28 | 2.36E-03 | 0.0383 | (A) NE (B/D) |
| SEM1 | 1.8 | 0.53 | 0.63 | 0.63 | 1.28 | 2.73E-03 | 0.0405 | (A) NE (B/D) |
| RRP12 | 4.34 | 3.07 | 0.7 | 0.62 | 1.28 | 2.97E-03 | 0.0422 | (A) NE (B/D) |
| KTI12 | 5.35 | 4.06 | 0.41 | 0.66 | 1.28 | 3.18E-03 | 0.0435 | (A) NE (B/D) |
| MRRFP1 | 1.73 | 0.44 | 0.92 | 0.39 | 1.28 | 3.20E-03 | 0.0438 | (A) NE (B/D) |
| WASF2 | 4.83 | 3.55 | 0.28 | 0.66 | 1.28 | 3.92E-03 | 0.0481 | (A) NE (B/D) |
| GEMIN7 | 1.46 | 0.19 | 0.33 | 0.24 | 1.27 | 1.01E-06 | 6.56E-04 | (A) NE (B/D) |
| TCF3 | 2.53 | 1.27 | 0.42 | 0.32 | 1.27 | 1.83E-05 | 3.49E-03 | (A) NE (B/D) |
| CHTF18 | 1.31 | 0.04 | 0.51 | 0.1 | 1.27 | 4.87E-05 | 5.69E-03 | (A) NE (B/D) |
| INE1 | 4.54 | 3.27 | 0.5 | 0.47 | 1.27 | 3.49E-04 | 0.0152 | (A) NE (B/D) |
| HSFX1 | 1.39 | 0.12 | 0.69 | 0.2 | 1.27 | 4.21E-04 | 0.0169 | (A) NE (B/D) |
| FCF1P2 | 5.07 | 3.8 | 0.38 | 0.49 | 1.27 | 4.59E-04 | 0.0175 | (A) NE (B/D) |
| HDAC3 | 1.67 | 0.41 | 0.43 | 0.56 | 1.27 | 1.19E-03 | 0.0273 | (A) NE (B/D) |
| ARAP3 | 1.43 | 0.16 | 0.82 | 0.27 | 1.27 | 1.38E-03 | 0.0296 | (A) NE (B/D) |
| CENPS-CORT | 2.35 | 1.08 | 0.41 | 0.61 | 1.27 | 1.99E-03 | 0.0354 | (A) NE (B/D) |
| CCDC114 | 1.27 | 0 | 0.88 | 0.01 | 1.27 | 2.60E-03 | 0.0399 | (A) NE (B/D) |
| UQCC3 | 6.06 | 4.79 | 0.67 | 0.65 | 1.27 | 3.62E-03 | 0.0463 | (A) NE (B/D) |

| | | | | | | | | |
|---|---|---|---|---|---|---|---|---|
| C9ORF85 | 5.48 | 4.22 | 0.65 | 0.38 | 1.26 | 3.94E-04 | 0.0163 | (A) NE (B/D) |
| YY2 | 3.66 | 2.4 | 0.67 | 0.41 | 1.26 | 5.70E-04 | 0.0191 | (A) NE (B/D) |
| DALRD3 | 1.45 | 0.19 | 0.67 | 0.42 | 1.26 | 6.21E-04 | 0.0198 | (A) NE (B/D) |
| ANKMY1 | 3.68 | 2.42 | 0.25 | 0.5 | 1.26 | 7.92E-04 | 0.0227 | (A) NE (B/D) |
| SDCBPP2 | 3.22 | 1.95 | 0.46 | 0.61 | 1.26 | 2.16E-03 | 0.0369 | (A) NE (B/D) |
| FAM219A | 1.57 | 0.31 | 0.21 | 0.58 | 1.26 | 2.30E-03 | 0.0379 | (A) NE (B/D) |
| EIF3EP1 | 7.35 | 6.09 | 0.55 | 0.65 | 1.26 | 3.14E-03 | 0.043 | (A) NE (B/D) |
| EZH1 | 5.5 | 4.24 | 0.26 | 0.64 | 1.26 | 3.52E-03 | 0.0457 | (A) NE (B/D) |
| TBC1D20 | 1.64 | 0.38 | 0.57 | 0.67 | 1.26 | 3.82E-03 | 0.0476 | (A) NE (B/D) |
| POLD2 | 2.22 | 0.95 | 0.6 | 0.68 | 1.26 | 4.07E-03 | 0.0488 | (A) NE (B/D) |
| DDX56 | 1.31 | 0.06 | 0.35 | 0.15 | 1.25 | 7.19E-07 | 5.25E-04 | (A) NE (B/D) |
| FRS3 | 1.31 | 0.06 | 0.37 | 0.15 | 1.25 | 1.46E-06 | 7.75E-04 | (A) NE (B/D) |
| NMB | 1.28 | 0.03 | 0.54 | 0.06 | 1.25 | 1.08E-04 | 8.50E-03 | (A) NE (B/D) |
| TINF2 | 1.66 | 0.41 | 0.58 | 0.48 | 1.25 | 6.50E-04 | 0.0204 | (A) NE (B/D) |
| ZKSCAN4 | 4.81 | 3.56 | 0.18 | 0.52 | 1.25 | 1.52E-03 | 0.031 | (A) NE (B/D) |
| SRSF9P1 | 5.17 | 3.92 | 0.56 | 0.59 | 1.25 | 1.85E-03 | 0.0345 | (A) NE (B/D) |
| PTBP1P | 4.82 | 3.57 | 0.61 | 0.59 | 1.25 | 2.15E-03 | 0.0368 | (A) NE (B/D) |
| RN7SL657P | 4.53 | 3.29 | 0.44 | 0.63 | 1.25 | 2.74E-03 | 0.0406 | (A) NE (B/D) |
| FAM173A | 6.66 | 5.41 | 0.79 | 0.58 | 1.25 | 3.89E-03 | 0.048 | (A) NE (B/D) |
| SDHAP1 | 7.89 | 6.65 | 0.36 | 0.34 | 1.24 | 2.90E-05 | 4.49E-03 | (A) NE (B/D) |
| SLC4A2 | 1.65 | 0.41 | 0.4 | 0.54 | 1.24 | 1.01E-03 | 0.0255 | (A) NE (B/D) |
| GMPR2 | 1.6 | 0.36 | 0.71 | 0.56 | 1.24 | 2.58E-03 | 0.0397 | (A) NE (B/D) |
| NATD1 | 3.32 | 2.08 | 0.4 | 0.64 | 1.24 | 3.42E-03 | 0.0452 | (A) NE (B/D) |
| REXO1 | 1.23 | 0 | 0.23 | 0.01 | 1.23 | 2.51E-07 | 4.05E-04 | (A) NE (B/D) |
| KLHL26 | 1.32 | 0.09 | 0.35 | 0.22 | 1.23 | 1.39E-06 | 7.73E-04 | (A) NE (B/D) |
| UTP14A | 4.77 | 3.54 | 0.48 | 0.45 | 1.23 | 3.41E-04 | 0.015 | (A) NE (B/D) |
| SRGAP2 | 2.99 | 1.77 | 0.35 | 0.54 | 1.23 | 1.27E-03 | 0.0282 | (A) NE (B/D) |
| ELL | 6.37 | 5.14 | 0.44 | 0.63 | 1.23 | 2.96E-03 | 0.0421 | (A) NE (B/D) |
| INTS4 | 7.57 | 6.35 | 0.14 | 0.25 | 1.22 | 1.38E-05 | 2.90E-03 | (A) NE (B/D) |
| CFAP410 | 1.33 | 0.11 | 0.6 | 0.27 | 1.22 | 1.74E-04 | 0.0106 | (A) NE (B/D) |
| RELT | 2.19 | 0.97 | 0.59 | 0.38 | 1.22 | 3.11E-04 | 0.0145 | (A) NE (B/D) |
| PMS2CL | 7.1 | 5.88 | 0.35 | 0.52 | 1.22 | 1.04E-03 | 0.0258 | (A) NE (B/D) |
| MAGEA2B | 1.29 | 0.06 | 0.75 | 0.16 | 1.22 | 1.07E-03 | 0.026 | (A) NE (B/D) |
| ARFRP1 | 6.33 | 5.11 | 0.23 | 0.54 | 1.22 | 1.71E-03 | 0.0329 | (A) NE (B/D) |
| KRBA2 | 3.86 | 2.63 | 0.74 | 0.5 | 1.22 | 2.08E-03 | 0.0362 | (A) NE (B/D) |
| LTC4S | 1.34 | 0.12 | 0.84 | 0.25 | 1.22 | 2.16E-03 | 0.0369 | (A) NE (B/D) |
| ARIH2P1 | 1.76 | 0.54 | 0.33 | 0.58 | 1.22 | 2.23E-03 | 0.0374 | (A) NE (B/D) |
| NKAP | 4.99 | 3.77 | 0.35 | 0.64 | 1.22 | 3.70E-03 | 0.0466 | (A) NE (B/D) |
| KDELR1 | 1.33 | 0.12 | 0.32 | 0.19 | 1.21 | 4.70E-07 | 4.22E-04 | (A) NE (B/D) |
| ERCC3 | 1.28 | 0.07 | 0.36 | 0.16 | 1.21 | 1.79E-06 | 8.29E-04 | (A) NE (B/D) |
| PIK3R2 | 1.66 | 0.45 | 0.4 | 0.39 | 1.21 | 1.13E-04 | 8.69E-03 | (A) NE (B/D) |
| DBNL | 4.8 | 3.59 | 0.23 | 0.42 | 1.21 | 3.52E-04 | 0.0152 | (A) NE (B/D) |
| FAM168A | 4.5 | 3.29 | 0.24 | 0.47 | 1.21 | 7.21E-04 | 0.0217 | (A) NE (B/D) |
| KRTCAP3 | 1.34 | 0.13 | 0.74 | 0.22 | 1.21 | 9.81E-04 | 0.0253 | (A) NE (B/D) |
| ANKHD1-EIF4 | 8.27 | 7.06 | 0.58 | 0.56 | 1.21 | 1.79E-03 | 0.0339 | (A) NE (B/D) |
| NUB1 | 7.29 | 6.09 | 0.28 | 0.57 | 1.21 | 2.29E-03 | 0.0377 | (A) NE (B/D) |
| MFSD11 | 11.06 | 9.84 | 0.15 | 0.58 | 1.21 | 3.11E-03 | 0.0429 | (A) NE (B/D) |
| CDK5RAP1 | 1.33 | 0.13 | 0.26 | 0.31 | 1.2 | 1.89E-05 | 3.50E-03 | (A) NE (B/D) |
| TNFSF12 | 1.58 | 0.38 | 0.36 | 0.44 | 1.2 | 3.45E-04 | 0.015 | (A) NE (B/D) |
| YDJC | 1.38 | 0.18 | 0.65 | 0.32 | 1.2 | 4.64E-04 | 0.0175 | (A) NE (B/D) |
| RRNAD1 | 1.49 | 0.29 | 0.21 | 0.5 | 1.2 | 1.29E-03 | 0.0284 | (A) NE (B/D) |
| SPTAN1 | 6.5 | 5.29 | 0.24 | 0.58 | 1.2 | 2.60E-03 | 0.0399 | (A) NE (B/D) |
| MAF1 | 1.71 | 0.51 | 0.63 | 0.6 | 1.2 | 3.24E-03 | 0.0441 | (A) NE (B/D) |
| PRMT9 | 1.77 | 0.57 | 0.82 | 0.49 | 1.2 | 3.92E-03 | 0.0481 | (A) NE (B/D) |
| TUBB8 | 1.25 | 0.06 | 0.71 | 0.15 | 1.19 | 8.60E-04 | 0.0237 | (A) NE (B/D) |
| VPS26C | 4.58 | 3.39 | 0.29 | 0.58 | 1.19 | 2.73E-03 | 0.0404 | (A) NE (B/D) |
| EEF1A1P38 | 1.83 | 0.64 | 0.43 | 0.61 | 1.19 | 2.99E-03 | 0.0423 | (A) NE (B/D) |
| AP5Z1 | 1.23 | 0.05 | 0.29 | 0.12 | 1.18 | 2.45E-07 | 4.05E-04 | (A) NE (B/D) |
| ARHGEF18 | 1.27 | 0.09 | 0.24 | 0.22 | 1.18 | 5.21E-07 | 4.22E-04 | (A) NE (B/D) |
| ZNF554 | 4.34 | 3.16 | 0.34 | 0.26 | 1.18 | 4.52E-06 | 1.51E-03 | (A) NE (B/D) |
| RAB24 | 1.33 | 0.15 | 0.61 | 0.37 | 1.18 | 4.20E-04 | 0.0169 | (A) NE (B/D) |
| EXD3 | 1.29 | 0.12 | 0.72 | 0.28 | 1.18 | 9.55E-04 | 0.025 | (A) NE (B/D) |
| ZFP37 | 1.27 | 0.09 | 0.73 | 0.22 | 1.18 | 1.08E-03 | 0.0261 | (A) NE (B/D) |
| PIM1 | 2.1 | 0.92 | 0.34 | 0.53 | 1.18 | 1.39E-03 | 0.0297 | (A) NE (B/D) |
| CFAP36 | 1.39 | 0.21 | 0.64 | 0.52 | 1.18 | 1.95E-03 | 0.0351 | (A) NE (B/D) |

| | | | | | | | | |
|---|---|---|---|---|---|---|---|---|
| MPL | 3.53 | 2.35 | 0.81 | 0.42 | 1.18 | 2.89E-03 | 0.0417 | (A) NE (B/D) |
| SENP3 | 1.27 | 0.1 | 0.43 | 0.25 | 1.17 | 1.56E-05 | 3.20E-03 | (A) NE (B/D) |
| PRR12 | 1.43 | 0.26 | 0.39 | 0.47 | 1.17 | 6.38E-04 | 0.0201 | (A) NE (B/D) |
| RPL9P25 | 1.6 | 0.43 | 0.29 | 0.52 | 1.17 | 1.52E-03 | 0.031 | (A) NE (B/D) |
| RPL23P2 | 1.36 | 0.19 | 0.75 | 0.36 | 1.17 | 1.58E-03 | 0.0315 | (A) NE (B/D) |
| ACTBP11 | 3.8 | 2.62 | 0.44 | 0.61 | 1.17 | 3.28E-03 | 0.0442 | (A) NE (B/D) |
| COPS7B | 1.53 | 0.36 | 0.25 | 0.59 | 1.17 | 3.44E-03 | 0.0452 | (A) NE (B/D) |
| AGO4 | 5.58 | 4.41 | 0.74 | 0.56 | 1.17 | 4.29E-03 | 0.0497 | (A) NE (B/D) |
| WIPI2 | 1.25 | 0.1 | 0.45 | 0.24 | 1.16 | 2.28E-05 | 3.95E-03 | (A) NE (B/D) |
| CDK16 | 1.4 | 0.23 | 0.37 | 0.32 | 1.16 | 2.95E-05 | 4.51E-03 | (A) NE (B/D) |
| TRIM39-RPP1 | 1.32 | 0.16 | 0.3 | 0.39 | 1.16 | 1.91E-04 | 0.0112 | (A) NE (B/D) |
| DIDO1 | 1.37 | 0.21 | 0.25 | 0.38 | 1.16 | 2.00E-04 | 0.0113 | (A) NE (B/D) |
| TOMM20L | 2.99 | 1.83 | 0.39 | 0.44 | 1.16 | 3.75E-04 | 0.0158 | (A) NE (B/D) |
| MSMP | 4.8 | 3.64 | 0.48 | 0.45 | 1.16 | 4.99E-04 | 0.0182 | (A) NE (B/D) |
| ZNF559 | 1.19 | 0.03 | 0.7 | 0.08 | 1.16 | 1.05E-03 | 0.0259 | (A) NE (B/D) |
| MAP7D1 | 1.39 | 0.22 | 0.32 | 0.55 | 1.16 | 2.00E-03 | 0.0354 | (A) NE (B/D) |
| ATAD3A | 2.07 | 0.91 | 0.37 | 0.58 | 1.16 | 2.73E-03 | 0.0405 | (A) NE (B/D) |
| PLEKHM1 | 3.37 | 2.21 | 0.43 | 0.63 | 1.16 | 4.07E-03 | 0.0488 | (A) NE (B/D) |
| FEM1A | 5.02 | 3.86 | 0.45 | 0.63 | 1.16 | 4.15E-03 | 0.049 | (A) NE (B/D) |
| UBA6-AS1 | 4.41 | 3.25 | 0.38 | 0.63 | 1.16 | 4.34E-03 | 0.05 | (A) NE (B/D) |
| ZNF335 | 2.12 | 0.98 | 0.32 | 0.41 | 1.15 | 2.96E-04 | 0.0142 | (A) NE (B/D) |
| KANSL3 | 1.56 | 0.41 | 0.4 | 0.53 | 1.15 | 1.57E-03 | 0.0314 | (A) NE (B/D) |
| UIMC1 | 2.85 | 1.7 | 0.48 | 0.58 | 1.15 | 2.70E-03 | 0.0404 | (A) NE (B/D) |
| SNTA1 | 1.16 | 0.01 | 0.81 | 0.02 | 1.15 | 2.72E-03 | 0.0404 | (A) NE (B/D) |
| TYW3 | 5.4 | 4.25 | 0.8 | 0.43 | 1.15 | 3.34E-03 | 0.0446 | (A) NE (B/D) |
| ZNF701 | 3.26 | 2.11 | 0.53 | 0.62 | 1.15 | 4.22E-03 | 0.0493 | (A) NE (B/D) |
| TRMT2B | 3.72 | 2.57 | 0.41 | 0.37 | 1.14 | 1.21E-04 | 9.03E-03 | (A) NE (B/D) |
| REX1BD | 1.55 | 0.41 | 0.31 | 0.37 | 1.14 | 1.29E-04 | 9.17E-03 | (A) NE (B/D) |
| NSUN5P2 | 1.28 | 0.14 | 0.63 | 0.33 | 1.14 | 5.56E-04 | 0.019 | (A) NE (B/D) |
| PDCL3 | 1.31 | 0.16 | 0.59 | 0.4 | 1.14 | 6.25E-04 | 0.0199 | (A) NE (B/D) |
| CD55 | 1.26 | 0.13 | 0.65 | 0.18 | 1.14 | 6.38E-04 | 0.0201 | (A) NE (B/D) |
| OSGEPL1 | 1.63 | 0.5 | 0.66 | 0.33 | 1.14 | 7.63E-04 | 0.0223 | (A) NE (B/D) |
| NDUFV3 | 4.52 | 3.38 | 0.49 | 0.47 | 1.14 | 8.68E-04 | 0.0238 | (A) NE (B/D) |
| PKDCC | 1.15 | 0.01 | 0.67 | 0.02 | 1.14 | 8.83E-04 | 0.0241 | (A) NE (B/D) |
| SNORD4A | 5.59 | 4.45 | 0.25 | 0.51 | 1.14 | 1.67E-03 | 0.0324 | (A) NE (B/D) |
| BRD9 | 1.35 | 0.21 | 0.59 | 0.51 | 1.14 | 1.84E-03 | 0.0344 | (A) NE (B/D) |
| PTPN12 | 1.4 | 0.26 | 0.65 | 0.5 | 1.14 | 2.12E-03 | 0.0365 | (A) NE (B/D) |
| STX4 | 1.49 | 0.36 | 0.6 | 0.55 | 1.14 | 2.78E-03 | 0.0409 | (A) NE (B/D) |
| OGDH | 2.37 | 1.23 | 0.37 | 0.58 | 1.14 | 2.99E-03 | 0.0423 | (A) NE (B/D) |
| CABP4 | 1.45 | 0.31 | 0.73 | 0.54 | 1.14 | 4.24E-03 | 0.0494 | (A) NE (B/D) |
| SIRT6 | 1.15 | 0.02 | 0.18 | 0.05 | 1.13 | 8.72E-09 | 4.43E-05 | (A) NE (B/D) |
| GMIP | 1.21 | 0.08 | 0.49 | 0.18 | 1.13 | 5.89E-05 | 6.47E-03 | (A) NE (B/D) |
| HSPBP1 | 1.28 | 0.15 | 0.43 | 0.37 | 1.13 | 1.65E-04 | 0.0106 | (A) NE (B/D) |
| ZNF579 | 1.17 | 0.04 | 0.57 | 0.1 | 1.13 | 2.74E-04 | 0.0138 | (A) NE (B/D) |
| PTCH2 | 1.31 | 0.17 | 0.59 | 0.25 | 1.13 | 2.99E-04 | 0.0142 | (A) NE (B/D) |
| DDX42 | 1.33 | 0.2 | 0.22 | 0.48 | 1.13 | 1.31E-03 | 0.0287 | (A) NE (B/D) |
| ARIH2 | 4.02 | 2.89 | 0.35 | 0.55 | 1.13 | 2.21E-03 | 0.0372 | (A) NE (B/D) |
| XPO5 | 1.45 | 0.33 | 0.49 | 0.56 | 1.13 | 2.60E-03 | 0.0398 | (A) NE (B/D) |
| HIPK2 | 2.99 | 1.87 | 0.45 | 0.46 | 1.12 | 7.46E-04 | 0.022 | (A) NE (B/D) |
| STX18 | 1.9 | 0.77 | 0.36 | 0.48 | 1.12 | 9.79E-04 | 0.0253 | (A) NE (B/D) |
| SLC35A2 | 1.53 | 0.41 | 0.58 | 0.53 | 1.12 | 2.34E-03 | 0.0382 | (A) NE (B/D) |
| ZNF3 | 1.34 | 0.22 | 0.21 | 0.41 | 1.11 | 5.90E-04 | 0.0194 | (A) NE (B/D) |
| NDE1 | 1.71 | 0.6 | 0.45 | 0.5 | 1.11 | 1.42E-03 | 0.03 | (A) NE (B/D) |
| VPS54 | 4.93 | 3.82 | 0.68 | 0.46 | 1.11 | 2.24E-03 | 0.0374 | (A) NE (B/D) |
| MSANTD1 | 2.7 | 1.59 | 0.58 | 0.56 | 1.11 | 3.53E-03 | 0.0457 | (A) NE (B/D) |
| MIDN | 1.2 | 0.1 | 0.25 | 0.17 | 1.1 | 1.52E-07 | 2.95E-04 | (A) NE (B/D) |
| ZFP69 | 4.47 | 3.36 | 0.4 | 0.15 | 1.1 | 1.13E-05 | 2.53E-03 | (A) NE (B/D) |
| ANKRD42 | 2.48 | 1.38 | 0.33 | 0.34 | 1.1 | 8.49E-05 | 7.73E-03 | (A) NE (B/D) |
| MMP24 | 1.37 | 0.27 | 0.42 | 0.43 | 1.1 | 5.36E-04 | 0.0186 | (A) NE (B/D) |
| TAZ | 1.11 | 0.01 | 0.6 | 0.02 | 1.1 | 5.88E-04 | 0.0194 | (A) NE (B/D) |
| ATP6V1H | 1.49 | 0.39 | 0.38 | 0.46 | 1.1 | 7.70E-04 | 0.0223 | (A) NE (B/D) |
| PPIAL4D | 1.25 | 0.15 | 0.69 | 0.12 | 1.1 | 1.26E-03 | 0.0281 | (A) NE (B/D) |
| ZNF791 | 6.04 | 4.94 | 0.68 | 0.4 | 1.1 | 1.80E-03 | 0.034 | (A) NE (B/D) |
| TBRG1 | 2.61 | 1.51 | 0.64 | 0.53 | 1.1 | 3.52E-03 | 0.0457 | (A) NE (B/D) |
| PACS2 | 1.57 | 0.47 | 0.35 | 0.59 | 1.1 | 4.07E-03 | 0.0488 | (A) NE (B/D) |

| | | | | | | | | |
|---|---|---|---|---|---|---|---|---|
| HCG15 | 2.77 | 1.67 | 0.66 | 0.55 | 1.1 | 4.14E-03 | 0.049 | (A) NE (B/D) |
| NDOR1 | 1.19 | 0.1 | 0.41 | 0.24 | 1.09 | 1.94E-05 | 3.50E-03 | (A) NE (B/D) |
| ZC3H4 | 1.88 | 0.79 | 0.34 | 0.34 | 1.09 | 8.35E-05 | 7.68E-03 | (A) NE (B/D) |
| IBSP | 1.21 | 0.12 | 0.59 | 0.3 | 1.09 | 5.00E-04 | 0.0182 | (A) NE (B/D) |
| GEMIN8 | 1.41 | 0.32 | 0.47 | 0.48 | 1.09 | 1.27E-03 | 0.0282 | (A) NE (B/D) |
| RN7SL663P | 1.11 | 0.02 | 0.71 | 0.05 | 1.09 | 1.77E-03 | 0.0337 | (A) NE (B/D) |
| BLMH | 1.42 | 0.33 | 0.72 | 0.45 | 1.09 | 3.09E-03 | 0.0429 | (A) NE (B/D) |
| RN7SKP23 | 1.51 | 0.42 | 0.42 | 0.57 | 1.09 | 3.25E-03 | 0.0441 | (A) NE (B/D) |
| SNHG16 | 1.78 | 0.69 | 0.52 | 0.56 | 1.09 | 3.45E-03 | 0.0453 | (A) NE (B/D) |
| SMYD5 | 1.9 | 0.8 | 0.64 | 0.53 | 1.09 | 3.55E-03 | 0.0459 | (A) NE (B/D) |
| DHX9P1 | 4.15 | 3.06 | 0.53 | 0.57 | 1.09 | 3.78E-03 | 0.0473 | (A) NE (B/D) |
| TMEM87B | 1.3 | 0.21 | 0.72 | 0.51 | 1.09 | 4.33E-03 | 0.0498 | (A) NE (B/D) |
| RNASET2 | 1.16 | 0.08 | 0.37 | 0.12 | 1.08 | 8.75E-06 | 2.28E-03 | (A) NE (B/D) |
| EMD | 1.18 | 0.1 | 0.36 | 0.24 | 1.08 | 9.09E-06 | 2.28E-03 | (A) NE (B/D) |
| PHKG1 | 1.37 | 0.29 | 0.48 | 0.25 | 1.08 | 8.20E-05 | 7.64E-03 | (A) NE (B/D) |
| IDH3A | 1.24 | 0.16 | 0.64 | 0.35 | 1.08 | 1.09E-03 | 0.0262 | (A) NE (B/D) |
| KCTD7 | 2.92 | 1.84 | 0.51 | 0.49 | 1.08 | 1.63E-03 | 0.0322 | (A) NE (B/D) |
| ARHGEF11 | 3.3 | 2.22 | 0.31 | 0.53 | 1.08 | 2.51E-03 | 0.0391 | (A) NE (B/D) |
| CHPF2 | 1.94 | 0.86 | 0.59 | 0.53 | 1.08 | 3.14E-03 | 0.0431 | (A) NE (B/D) |
| FBXW7 | 1.23 | 0.16 | 0.79 | 0.18 | 1.08 | 3.25E-03 | 0.0441 | (A) NE (B/D) |
| SNRPA1 | 1.3 | 0.22 | 0.59 | 0.54 | 1.08 | 3.66E-03 | 0.0465 | (A) NE (B/D) |
| CACNB1 | 1.37 | 0.3 | 0.41 | 0.58 | 1.08 | 4.10E-03 | 0.0489 | (A) NE (B/D) |
| SERTAD2 | 4.35 | 3.29 | 0.25 | 0.36 | 1.07 | 2.16E-04 | 0.0119 | (A) NE (B/D) |
| NAXD | 1.15 | 0.08 | 0.54 | 0.21 | 1.07 | 2.25E-04 | 0.0122 | (A) NE (B/D) |
| RPL22 | 1.31 | 0.24 | 0.51 | 0.33 | 1.07 | 2.73E-04 | 0.0138 | (A) NE (B/D) |
| NFYA | 1.23 | 0.16 | 0.24 | 0.37 | 1.07 | 2.96E-04 | 0.0142 | (A) NE (B/D) |
| TSC22D1 | 1.11 | 0.05 | 0.57 | 0.11 | 1.07 | 4.45E-04 | 0.0174 | (A) NE (B/D) |
| ENOX2 | 2.82 | 1.75 | 0.75 | 0.25 | 1.07 | 2.47E-03 | 0.0389 | (A) NE (B/D) |
| FTSJ1 | 2.13 | 1.06 | 0.36 | 0.54 | 1.07 | 2.71E-03 | 0.0404 | (A) NE (B/D) |
| TEP1 | 1.93 | 0.86 | 0.77 | 0.42 | 1.07 | 4.21E-03 | 0.0493 | (A) NE (B/D) |
| MIR-3620/36 | 1.26 | 0.2 | 0.38 | 0.29 | 1.06 | 4.02E-05 | 5.17E-03 | (A) NE (B/D) |
| MIR-3620/3P | 1.26 | 0.2 | 0.38 | 0.29 | 1.06 | 4.02E-05 | 5.17E-03 | (A) NE (B/D) |
| MIR-3620/5P | 1.26 | 0.2 | 0.38 | 0.29 | 1.06 | 4.02E-05 | 5.17E-03 | (A) NE (B/D) |
| ZBTB49 | 1.2 | 0.14 | 0.48 | 0.23 | 1.06 | 1.03E-04 | 8.25E-03 | (A) NE (B/D) |
| TBC1D9B | 1.22 | 0.16 | 0.14 | 0.36 | 1.06 | 5.00E-04 | 0.0182 | (A) NE (B/D) |
| ANKHD1 | 2.41 | 1.36 | 0.52 | 0.45 | 1.06 | 1.28E-03 | 0.0284 | (A) NE (B/D) |
| ANKFY1 | 5.95 | 4.89 | 0.64 | 0.41 | 1.06 | 1.76E-03 | 0.0335 | (A) NE (B/D) |
| CLPP | 1.09 | 0.03 | 0.81 | 0.08 | 1.06 | 4.29E-03 | 0.0497 | (A) NE (B/D) |
| USP21 | 1.19 | 0.13 | 0.3 | 0.33 | 1.05 | 8.70E-05 | 7.77E-03 | (A) NE (B/D) |
| SPAG1 | 1.12 | 0.07 | 0.63 | 0.11 | 1.05 | 9.30E-04 | 0.0247 | (A) NE (B/D) |
| MIR-4520A/3 | 1.34 | 0.28 | 0.64 | 0.49 | 1.05 | 3.41E-03 | 0.0452 | (A) NE (B/D) |
| MIR-4520A/5 | 1.34 | 0.28 | 0.64 | 0.49 | 1.05 | 3.41E-03 | 0.0452 | (A) NE (B/D) |
| APPL2 | 2.41 | 1.37 | 0.62 | 0.5 | 1.05 | 3.44E-03 | 0.0453 | (A) NE (B/D) |
| ZNF799 | 4.29 | 3.24 | 0.8 | 0.27 | 1.05 | 4.23E-03 | 0.0494 | (A) NE (B/D) |
| CBX4 | 1.04 | 0.01 | 0.45 | 0.02 | 1.04 | 1.14E-04 | 8.73E-03 | (A) NE (B/D) |
| ZNF394 | 2.96 | 1.92 | 0.13 | 0.33 | 1.04 | 3.07E-04 | 0.0144 | (A) NE (B/D) |
| TSC1 | 1.17 | 0.13 | 0.52 | 0.31 | 1.04 | 3.16E-04 | 0.0145 | (A) NE (B/D) |
| RGP1 | 8.16 | 7.13 | 0.36 | 0.45 | 1.04 | 1.10E-03 | 0.0262 | (A) NE (B/D) |
| SDHAP2 | 8.12 | 7.08 | 0.57 | 0.44 | 1.04 | 1.63E-03 | 0.0321 | (A) NE (B/D) |
| DNMT1 | 1.35 | 0.31 | 0.44 | 0.53 | 1.04 | 2.99E-03 | 0.0423 | (A) NE (B/D) |
| C1ORF131 | 2.72 | 1.68 | 0.44 | 0.55 | 1.04 | 3.69E-03 | 0.0466 | (A) NE (B/D) |
| CITED2 | 1.61 | 0.57 | 0.59 | 0.52 | 1.04 | 3.75E-03 | 0.047 | (A) NE (B/D) |
| AP5S1 | 3.29 | 2.25 | 0.2 | 0.54 | 1.04 | 4.32E-03 | 0.0498 | (A) NE (B/D) |
| ZNF888 | 1.14 | 0.11 | 0.33 | 0.17 | 1.03 | 3.50E-06 | 1.28E-03 | (A) NE (B/D) |
| UBE2HP1 | 1.07 | 0.03 | 0.45 | 0.08 | 1.03 | 9.40E-05 | 7.97E-03 | (A) NE (B/D) |
| RAB8A | 1.37 | 0.34 | 0.3 | 0.39 | 1.03 | 4.61E-04 | 0.0175 | (A) NE (B/D) |
| MFF | 5.1 | 4.07 | 0.51 | 0.47 | 1.03 | 1.88E-03 | 0.0347 | (A) NE (B/D) |
| CICP14 | 4.53 | 3.51 | 0.6 | 0.44 | 1.03 | 2.16E-03 | 0.0369 | (A) NE (B/D) |
| COX10 | 2.06 | 1.04 | 0.32 | 0.42 | 1.02 | 7.69E-04 | 0.0223 | (A) NE (B/D) |
| TIRAP | 7.78 | 6.76 | 0.29 | 0.51 | 1.02 | 2.83E-03 | 0.0413 | (A) NE (B/D) |
| VAMP3 | 1.2 | 0.19 | 0.41 | 0.3 | 1.01 | 1.06E-04 | 8.39E-03 | (A) NE (B/D) |
| PLK1 | 1.12 | 0.11 | 0.69 | 0.26 | 1.01 | 2.00E-03 | 0.0354 | (A) NE (B/D) |
| ZBTB43 | 5.71 | 4.71 | 0.26 | 0.25 | 1 | 1.25E-05 | 2.74E-03 | (A) NE (B/D) |
| ZNF703 | 1.11 | 0.11 | 0.37 | 0.27 | 1 | 3.89E-05 | 5.17E-03 | (A) NE (B/D) |
| KAT8 | 1.24 | 0.24 | 0.27 | 0.37 | 1 | 3.41E-04 | 0.015 | (A) NE (B/D) |

| | | | | | | | | |
|---|---|---|---|---|---|---|---|---|
| RPS20P33 | 3.88 | 2.87 | 0.55 | 0.41 | 1 | 1.48E-03 | 0.0308 | (A) NE (B/D) |
| ZKSCAN5 | 1.54 | 0.54 | 0.45 | 0.48 | 1 | 1.99E-03 | 0.0354 | (A) NE (B/D) |
| FKTN | 1.37 | 0.37 | 0.6 | 0.42 | 1 | 2.34E-03 | 0.0382 | (A) NE (B/D) |
| TOM1L2 | 1.36 | 0.36 | 0.44 | 0.51 | 1 | 2.89E-03 | 0.0417 | (A) NE (B/D) |
| ARHGAP35 | 10.09 | 9.09 | 0.37 | 0.53 | 1 | 3.49E-03 | 0.0456 | (A) NE (B/D) |
| U2AF1 | 1.07 | 0.09 | 0.3 | 0.21 | 0.99 | 4.98E-06 | 1.60E-03 | (A) NE (B/D) |
| AP2M1 | 1.15 | 0.16 | 0.41 | 0.38 | 0.99 | 5.24E-04 | 0.0185 | (A) NE (B/D) |
| RIPPLY2 | 1.15 | 0.16 | 0.68 | 0.24 | 0.99 | 2.21E-03 | 0.0372 | (A) NE (B/D) |
| DFFB | 1.12 | 0.14 | 0.55 | 0.32 | 0.98 | 7.96E-04 | 0.0228 | (A) NE (B/D) |
| KPTN | 1.02 | 0.04 | 0.63 | 0.07 | 0.98 | 1.56E-03 | 0.0313 | (A) NE (B/D) |
| KIAA0753 | 1.8 | 0.82 | 0.63 | 0.3 | 0.98 | 1.67E-03 | 0.0324 | (A) NE (B/D) |
| UGDH-AS1 | 2.87 | 1.89 | 0.66 | 0.41 | 0.98 | 3.64E-03 | 0.0464 | (A) NE (B/D) |
| MAP3K3 | 1.54 | 0.57 | 0.4 | 0.34 | 0.97 | 2.70E-04 | 0.0137 | (A) NE (B/D) |
| TSEN54 | 1.12 | 0.15 | 0.41 | 0.36 | 0.97 | 4.51E-04 | 0.0175 | (A) NE (B/D) |
| TAF6L | 1.48 | 0.5 | 0.33 | 0.38 | 0.97 | 5.18E-04 | 0.0185 | (A) NE (B/D) |
| NDUFA3 | 0.99 | 0.03 | 0.64 | 0.06 | 0.97 | 1.95E-03 | 0.0351 | (A) NE (B/D) |
| SLC25A26 | 1.1 | 0.13 | 0.69 | 0.29 | 0.97 | 2.94E-03 | 0.042 | (A) NE (B/D) |
| PDSS2 | 1.34 | 0.37 | 0.63 | 0.44 | 0.97 | 3.87E-03 | 0.0479 | (A) NE (B/D) |
| RNASEH2C | 1.29 | 0.33 | 0.37 | 0.42 | 0.96 | 1.02E-03 | 0.0256 | (A) NE (B/D) |
| LRCH4 | 0.97 | 0.01 | 0.64 | 0.02 | 0.96 | 1.99E-03 | 0.0354 | (A) NE (B/D) |
| IRGQ | 6.12 | 5.15 | 0.64 | 0.37 | 0.96 | 2.66E-03 | 0.0401 | (A) NE (B/D) |
| TBRG4 | 1.03 | 0.08 | 0.08 | 0.19 | 0.95 | 1.99E-05 | 3.53E-03 | (A) NE (B/D) |
| BCL10 | 5.56 | 4.61 | 0.49 | 0.41 | 0.95 | 1.48E-03 | 0.0308 | (A) NE (B/D) |
| FAM207A | 4.12 | 3.17 | 0.53 | 0.42 | 0.95 | 2.08E-03 | 0.0362 | (A) NE (B/D) |
| POLH | 6.37 | 5.42 | 0.52 | 0.49 | 0.95 | 4.09E-03 | 0.0488 | (A) NE (B/D) |
| NOL8 | 2.35 | 1.42 | 0.48 | 0.34 | 0.94 | 7.06E-04 | 0.0215 | (A) NE (B/D) |
| DND1 | 1.16 | 0.21 | 0.34 | 0.42 | 0.94 | 1.15E-03 | 0.0268 | (A) NE (B/D) |
| GID8 | 1.65 | 0.71 | 0.13 | 0.42 | 0.94 | 2.25E-03 | 0.0375 | (A) NE (B/D) |
| RINT1 | 2.18 | 1.24 | 0.33 | 0.46 | 0.94 | 2.50E-03 | 0.0391 | (A) NE (B/D) |
| ESYT2 | 2.89 | 1.96 | 0.31 | 0.27 | 0.93 | 5.50E-05 | 6.19E-03 | (A) NE (B/D) |
| MIR-555/555 | 6.62 | 5.69 | 0.5 | 0.41 | 0.93 | 1.96E-03 | 0.0351 | (A) NE (B/D) |
| SRD5A3-AS1 | 3.32 | 2.39 | 0.52 | 0.44 | 0.93 | 2.96E-03 | 0.0421 | (A) NE (B/D) |
| CNOT3 | 0.98 | 0.06 | 0.35 | 0.14 | 0.92 | 1.96E-05 | 3.50E-03 | (A) NE (B/D) |
| FAM104B | 1.05 | 0.13 | 0.48 | 0.25 | 0.92 | 3.66E-04 | 0.0156 | (A) NE (B/D) |
| RECQL5 | 1.55 | 0.62 | 0.48 | 0.42 | 0.92 | 2.14E-03 | 0.0367 | (A) NE (B/D) |
| TOX4 | 1.27 | 0.35 | 0.4 | 0.48 | 0.92 | 3.32E-03 | 0.0444 | (A) NE (B/D) |
| POM121 | 1 | 0.09 | 0.37 | 0.21 | 0.91 | 4.20E-05 | 5.30E-03 | (A) NE (B/D) |
| ABCB8 | 0.94 | 0.03 | 0.37 | 0.08 | 0.91 | 5.87E-05 | 6.47E-03 | (A) NE (B/D) |
| SRP68 | 1.03 | 0.13 | 0.38 | 0.31 | 0.91 | 2.40E-04 | 0.0128 | (A) NE (B/D) |
| CREB3 | 1.14 | 0.24 | 0.5 | 0.27 | 0.91 | 5.69E-04 | 0.0191 | (A) NE (B/D) |
| TAX1BP1 | 7.56 | 6.65 | 0.33 | 0.49 | 0.91 | 3.90E-03 | 0.048 | (A) NE (B/D) |
| ACBD3 | 8.48 | 7.57 | 0.32 | 0.49 | 0.91 | 3.96E-03 | 0.0484 | (A) NE (B/D) |
| RN7SL244P | 1.23 | 0.33 | 0.39 | 0.21 | 0.9 | 6.52E-05 | 6.86E-03 | (A) NE (B/D) |
| B4GALT7 | 1.04 | 0.14 | 0.23 | 0.35 | 0.9 | 5.37E-04 | 0.0186 | (A) NE (B/D) |
| MTA1 | 0.98 | 0.08 | 0.54 | 0.2 | 0.9 | 8.06E-04 | 0.0229 | (A) NE (B/D) |
| ASB1 | 1.01 | 0.12 | 0.51 | 0.3 | 0.89 | 8.94E-04 | 0.0242 | (A) NE (B/D) |
| IGBP1P1 | 4.73 | 3.84 | 0.4 | 0.38 | 0.89 | 1.11E-03 | 0.0263 | (A) NE (B/D) |
| DOCK6 | 1.01 | 0.12 | 0.6 | 0.11 | 0.89 | 1.90E-03 | 0.0348 | (A) NE (B/D) |
| RABGEF1 | 1.51 | 0.62 | 0.29 | 0.48 | 0.89 | 4.01E-03 | 0.0486 | (A) NE (B/D) |
| ALG1L6P | 1.63 | 0.75 | 0.38 | 0.22 | 0.88 | 8.43E-05 | 7.73E-03 | (A) NE (B/D) |
| KAZN | 1.31 | 0.43 | 0.39 | 0.39 | 0.88 | 1.33E-03 | 0.0289 | (A) NE (B/D) |
| FXR2 | 0.9 | 0.02 | 0.56 | 0.05 | 0.88 | 1.40E-03 | 0.0299 | (A) NE (B/D) |
| SEPTIN8 | 2.64 | 1.76 | 0.38 | 0.41 | 0.88 | 1.75E-03 | 0.0334 | (A) NE (B/D) |
| LARP1 | 0.89 | 0.01 | 0.37 | 0.03 | 0.87 | 8.83E-05 | 7.77E-03 | (A) NE (B/D) |
| LMAN2L | 1.15 | 0.28 | 0.37 | 0.41 | 0.87 | 2.00E-03 | 0.0354 | (A) NE (B/D) |
| ZBTB2 | 4.71 | 3.85 | 0.31 | 0.27 | 0.86 | 1.13E-04 | 8.69E-03 | (A) NE (B/D) |
| PPP2R5C | 0.96 | 0.1 | 0.55 | 0.26 | 0.86 | 1.53E-03 | 0.0311 | (A) NE (B/D) |
| CNOT6LP1 | 4.97 | 4.12 | 0.41 | 0.41 | 0.86 | 2.35E-03 | 0.0383 | (A) NE (B/D) |
| POLR2E | 0.93 | 0.08 | 0.36 | 0.19 | 0.85 | 4.93E-05 | 5.73E-03 | (A) NE (B/D) |
| HDAC7 | 0.93 | 0.08 | 0.39 | 0.12 | 0.85 | 1.01E-04 | 8.21E-03 | (A) NE (B/D) |
| E2F5 | 0.94 | 0.08 | 0.44 | 0.19 | 0.85 | 2.77E-04 | 0.0139 | (A) NE (B/D) |
| PCDHB2 | 0.87 | 0.03 | 0.43 | 0.06 | 0.85 | 3.15E-04 | 0.0145 | (A) NE (B/D) |
| ZNF473 | 1.29 | 0.44 | 0.4 | 0.37 | 0.85 | 1.40E-03 | 0.0299 | (A) NE (B/D) |
| IPO5 | 0.9 | 0.04 | 0.56 | 0.07 | 0.85 | 1.70E-03 | 0.0328 | (A) NE (B/D) |
| TMEM260 | 1.13 | 0.28 | 0.56 | 0.25 | 0.85 | 1.88E-03 | 0.0347 | (A) NE (B/D) |

| | | | | | | | | |
|---|---|---|---|---|---|---|---|---|
| MED30 | 1.57 | 0.73 | 0.59 | 0.27 | 0.85 | 2.76E-03 | 0.0407 | (A) NE (B/D) |
| ENTR1 | 0.99 | 0.15 | 0.26 | 0.24 | 0.84 | 4.10E-05 | 5.21E-03 | (A) NE (B/D) |
| HOXC6 | 0.95 | 0.11 | 0.37 | 0.22 | 0.84 | 9.77E-05 | 8.10E-03 | (A) NE (B/D) |
| POLR1C | 0.91 | 0.07 | 0.48 | 0.17 | 0.84 | 5.66E-04 | 0.0191 | (A) NE (B/D) |
| RNF24 | 1.9 | 1.06 | 0.38 | 0.41 | 0.84 | 2.61E-03 | 0.0399 | (A) NE (B/D) |
| CYTH2 | 0.96 | 0.13 | 0.56 | 0.31 | 0.84 | 2.67E-03 | 0.0401 | (A) NE (B/D) |
| RPL32P3 | 0.95 | 0.12 | 0.41 | 0.3 | 0.83 | 6.41E-04 | 0.0202 | (A) NE (B/D) |
| CSK | 1.03 | 0.2 | 0.46 | 0.36 | 0.83 | 1.74E-03 | 0.0333 | (A) NE (B/D) |
| TRAPPC10 | 6.51 | 5.68 | 0.33 | 0.45 | 0.83 | 4.15E-03 | 0.049 | (A) NE (B/D) |
| ANKS1A | 0.94 | 0.13 | 0.18 | 0.31 | 0.82 | 5.41E-04 | 0.0186 | (A) NE (B/D) |
| CHD2 | 0.89 | 0.08 | 0.5 | 0.19 | 0.82 | 9.37E-04 | 0.0248 | (A) NE (B/D) |
| SRSF8 | 1.11 | 0.28 | 0.31 | 0.42 | 0.82 | 2.92E-03 | 0.0419 | (A) NE (B/D) |
| MAVS | 2.42 | 1.61 | 0.26 | 0.24 | 0.81 | 5.15E-05 | 5.93E-03 | (A) NE (B/D) |
| FAM27E4 | 0.81 | 0.01 | 0.32 | 0.01 | 0.81 | 6.27E-05 | 6.77E-03 | (A) NE (B/D) |
| QRICH1 | 0.92 | 0.12 | 0.41 | 0.24 | 0.81 | 3.51E-04 | 0.0152 | (A) NE (B/D) |
| PLEKHH1 | 0.94 | 0.12 | 0.48 | 0.3 | 0.81 | 1.50E-03 | 0.031 | (A) NE (B/D) |
| UBA6 | 0.9 | 0.09 | 0.53 | 0.21 | 0.81 | 1.69E-03 | 0.0327 | (A) NE (B/D) |
| USP24 | 0.93 | 0.12 | 0.55 | 0.21 | 0.81 | 2.06E-03 | 0.0361 | (A) NE (B/D) |
| NAAA | 1.04 | 0.23 | 0.43 | 0.38 | 0.81 | 2.39E-03 | 0.0384 | (A) NE (B/D) |
| MSANTD2 | 0.82 | 0.01 | 0.59 | 0.02 | 0.81 | 3.51E-03 | 0.0457 | (A) NE (B/D) |
| RBMX | 1.56 | 0.75 | 0.4 | 0.43 | 0.81 | 4.15E-03 | 0.049 | (A) NE (B/D) |
| WDR60 | 0.91 | 0.11 | 0.32 | 0.17 | 0.8 | 2.85E-05 | 4.49E-03 | (A) NE (B/D) |
| RAI1 | 0.85 | 0.05 | 0.39 | 0.13 | 0.8 | 1.71E-04 | 0.0106 | (A) NE (B/D) |
| ZNF763 | 0.86 | 0.05 | 0.53 | 0.13 | 0.8 | 1.74E-03 | 0.0332 | (A) NE (B/D) |
| INTS4P1 | 1.23 | 0.43 | 0.51 | 0.36 | 0.8 | 3.44E-03 | 0.0452 | (A) NE (B/D) |
| ALG13 | 1.58 | 0.8 | 0.48 | 0.21 | 0.79 | 1.00E-03 | 0.0254 | (A) NE (B/D) |
| GTF3C3 | 1 | 0.21 | 0.38 | 0.37 | 0.79 | 1.90E-03 | 0.0348 | (A) NE (B/D) |
| LARP6 | 0.89 | 0.11 | 0.4 | 0.28 | 0.78 | 6.58E-04 | 0.0205 | (A) NE (B/D) |
| FNTA | 1.57 | 0.78 | 0.47 | 0.34 | 0.78 | 2.57E-03 | 0.0396 | (A) NE (B/D) |
| MAP7 | 1.04 | 0.26 | 0.44 | 0.38 | 0.78 | 3.56E-03 | 0.0459 | (A) NE (B/D) |
| LONP1 | 0.84 | 0.06 | 0.59 | 0.15 | 0.78 | 3.85E-03 | 0.0478 | (A) NE (B/D) |
| D2HGDH | 0.93 | 0.16 | 0.37 | 0.25 | 0.77 | 3.39E-04 | 0.015 | (A) NE (B/D) |
| MED25 | 0.88 | 0.12 | 0.38 | 0.28 | 0.77 | 6.16E-04 | 0.0198 | (A) NE (B/D) |
| SSU72 | 0.89 | 0.12 | 0.45 | 0.29 | 0.77 | 1.41E-03 | 0.0299 | (A) NE (B/D) |
| CBLL1 | 2.8 | 2.03 | 0.31 | 0.35 | 0.77 | 1.49E-03 | 0.0309 | (A) NE (B/D) |
| ADAM17 | 1.02 | 0.25 | 0.42 | 0.35 | 0.77 | 2.28E-03 | 0.0376 | (A) NE (B/D) |
| TRAPPC12-AS | 5.15 | 4.39 | 0.44 | 0.37 | 0.76 | 3.58E-03 | 0.0461 | (A) NE (B/D) |
| COA1 | 0.84 | 0.08 | 0.43 | 0.17 | 0.75 | 5.88E-04 | 0.0194 | (A) NE (B/D) |
| ZNF587 | 1.95 | 1.19 | 0.47 | 0.25 | 0.75 | 1.60E-03 | 0.0318 | (A) NE (B/D) |
| ALG9 | 0.77 | 0.02 | 0.5 | 0.04 | 0.75 | 1.83E-03 | 0.0343 | (A) NE (B/D) |
| GTPBP3 | 0.82 | 0.09 | 0.24 | 0.21 | 0.74 | 4.73E-05 | 5.64E-03 | (A) NE (B/D) |
| COG7 | 0.78 | 0.05 | 0.46 | 0.11 | 0.74 | 1.15E-03 | 0.0268 | (A) NE (B/D) |
| CCDC150 | 0.77 | 0.03 | 0.56 | 0.08 | 0.74 | 4.13E-03 | 0.049 | (A) NE (B/D) |
| METTL26 | 0.85 | 0.12 | 0.51 | 0.21 | 0.73 | 2.56E-03 | 0.0396 | (A) NE (B/D) |
| ZNHIT6 | 4.22 | 3.49 | 0.29 | 0.38 | 0.73 | 3.40E-03 | 0.0451 | (A) NE (B/D) |
| ZNF282 | 0.81 | 0.1 | 0.25 | 0.15 | 0.72 | 1.12E-05 | 2.53E-03 | (A) NE (B/D) |
| FAM122A | 4.42 | 3.7 | 0.27 | 0.23 | 0.72 | 1.25E-04 | 9.14E-03 | (A) NE (B/D) |
| POC1A | 0.89 | 0.17 | 0.34 | 0.15 | 0.72 | 1.27E-04 | 9.14E-03 | (A) NE (B/D) |
| WHAMM | 0.78 | 0.05 | 0.39 | 0.13 | 0.72 | 4.06E-04 | 0.0166 | (A) NE (B/D) |
| NRBP1 | 0.92 | 0.2 | 0.13 | 0.33 | 0.72 | 2.09E-03 | 0.0362 | (A) NE (B/D) |
| SPIN2A | 0.92 | 0.21 | 0.26 | 0.23 | 0.71 | 1.28E-04 | 9.17E-03 | (A) NE (B/D) |
| LPCAT4 | 0.79 | 0.08 | 0.34 | 0.19 | 0.71 | 2.09E-04 | 0.0116 | (A) NE (B/D) |
| UBE4B | 0.75 | 0.04 | 0.43 | 0.08 | 0.71 | 1.02E-03 | 0.0255 | (A) NE (B/D) |
| HTT | 3.97 | 3.26 | 0.38 | 0.3 | 0.71 | 1.51E-03 | 0.031 | (A) NE (B/D) |
| OTUD5 | 0.94 | 0.22 | 0.23 | 0.33 | 0.71 | 1.56E-03 | 0.0313 | (A) NE (B/D) |
| AFDN | 0.76 | 0.05 | 0.49 | 0.12 | 0.71 | 2.25E-03 | 0.0375 | (A) NE (B/D) |
| EIF2B4 | 0.7 | 0 | 0.3 | 0 | 0.7 | 1.11E-04 | 8.64E-03 | (A) NE (B/D) |
| COG5 | 2.3 | 1.6 | 0.38 | 0.32 | 0.7 | 2.40E-03 | 0.0385 | (A) NE (B/D) |
| PIK3R3 | 0.69 | 0.01 | 0.44 | 0.02 | 0.69 | 1.54E-03 | 0.0312 | (A) NE (B/D) |
| FAM86DP | 1.06 | 0.38 | 0.21 | 0.24 | 0.68 | 1.96E-04 | 0.0113 | (A) NE (B/D) |
| DSTYK | 0.77 | 0.09 | 0.41 | 0.22 | 0.68 | 1.19E-03 | 0.0273 | (A) NE (B/D) |
| SCAND2P | 0.7 | 0.02 | 0.43 | 0.06 | 0.68 | 1.52E-03 | 0.031 | (A) NE (B/D) |
| ANKRD17 | 0.77 | 0.09 | 0.49 | 0.22 | 0.68 | 3.53E-03 | 0.0457 | (A) NE (B/D) |
| RICTOR | 0.83 | 0.16 | 0.49 | 0.21 | 0.67 | 3.63E-03 | 0.0463 | (A) NE (B/D) |
| ZZEF1 | 1.71 | 1.06 | 0.45 | 0.2 | 0.66 | 2.25E-03 | 0.0375 | (A) NE (B/D) |

| | | | | | | | | |
|---|---|---|---|---|---|---|---|---|
| SNIP1 | 0.76 | 0.11 | 0.35 | 0.26 | 0.65 | 1.18E-03 | 0.0272 | (A) NE (B/D) |
| C15ORF62 | 0.69 | 0.05 | 0.44 | 0.1 | 0.65 | 2.07E-03 | 0.0361 | (A) NE (B/D) |
| C8ORF37 | 0.68 | 0.02 | 0.45 | 0.06 | 0.65 | 2.37E-03 | 0.0384 | (A) NE (B/D) |
| SNORD51 | 0.72 | 0.08 | 0.36 | 0.15 | 0.64 | 5.41E-04 | 0.0186 | (A) NE (B/D) |
| CAMK2D | 0.81 | 0.17 | 0.32 | 0.28 | 0.64 | 1.56E-03 | 0.0313 | (A) NE (B/D) |
| HSPB1P1 | 0.82 | 0.17 | 0.41 | 0.27 | 0.64 | 2.64E-03 | 0.04 | (A) NE (B/D) |
| NUP42 | 0.66 | 0.02 | 0.44 | 0.06 | 0.63 | 2.38E-03 | 0.0384 | (A) NE (B/D) |
| SLC35A4 | 0.81 | 0.18 | 0.4 | 0.3 | 0.63 | 4.11E-03 | 0.0489 | (A) NE (B/D) |
| FARSB | 1.09 | 0.47 | 0.21 | 0.33 | 0.62 | 3.68E-03 | 0.0466 | (A) NE (B/D) |
| TMSB4XP2 | 0.67 | 0.06 | 0.45 | 0.15 | 0.61 | 3.29E-03 | 0.0442 | (A) NE (B/D) |
| SH2B1 | 0.7 | 0.1 | 0.14 | 0.23 | 0.6 | 5.93E-04 | 0.0194 | (A) NE (B/D) |
| ABHD17B | 0.74 | 0.14 | 0.37 | 0.17 | 0.6 | 1.27E-03 | 0.0282 | (A) NE (B/D) |
| ATP5MF | 0.79 | 0.19 | 0.16 | 0.3 | 0.6 | 2.72E-03 | 0.0404 | (A) NE (B/D) |
| MIGA2 | 0.65 | 0.05 | 0.43 | 0.13 | 0.6 | 2.94E-03 | 0.042 | (A) NE (B/D) |
| PGM3 | 0.67 | 0.07 | 0.4 | 0.17 | 0.59 | 1.94E-03 | 0.0351 | (A) NE (B/D) |
| ZNF749 | 0.68 | 0.09 | 0.44 | 0.21 | 0.59 | 4.30E-03 | 0.0497 | (A) NE (B/D) |
| GOLGA4 | 1.58 | 2.19 | 0.43 | 0.16 | -0.6 | 2.95E-03 | 0.042 | (B) Small cell-mixed small cell (C/E) |
| OXNAD1 | 0.66 | 1.3 | 0.22 | 0.32 | -0.64 | 2.37E-03 | 0.0384 | (B) Small cell-mixed small cell (C/E) |
| BCAS2P2 | 0.43 | 1.2 | 0.34 | 0.39 | -0.78 | 2.80E-03 | 0.0411 | (B) Small cell-mixed small cell (C/E) |
| BMS1P2 | 2.1 | 3.03 | 0.32 | 0.39 | -0.92 | 9.03E-04 | 0.0244 | (B) Small cell-mixed small cell (C/E) |
| EEA1 | 0.3 | 1.32 | 0.74 | 0.38 | -1.02 | 4.14E-03 | 0.049 | (B) Small cell-mixed small cell (C/E) |
| SLC25A3P2 | 0.65 | 1.75 | 0.75 | 0.35 | -1.1 | 2.39E-03 | 0.0384 | (B) Small cell-mixed small cell (C/E) |
| MTND2P12 | 0.07 | 1.21 | 0.17 | 0.54 | -1.13 | 2.88E-03 | 0.0417 | (B) Small cell-mixed small cell (C/E) |
| RARB | 1.21 | 2.39 | 0.82 | 0.5 | -1.19 | 4.17E-03 | 0.049 | (B) Small cell-mixed small cell (C/E) |
| RPS6P16 | 1.13 | 2.55 | 0.69 | 0.37 | -1.42 | 2.05E-04 | 0.0115 | (B) Small cell-mixed small cell (C/E) |
| RNU6-892P | 1.35 | 2.88 | 0.93 | 0.73 | -1.54 | 3.49E-03 | 0.0456 | (B) Small cell-mixed small cell (C/E) |
| LINC02603 | 2.97 | 4.55 | 0.32 | 0.51 | -1.58 | 1.85E-04 | 0.011 | (B) Small cell-mixed small cell (C/E) |
| GK3P | 1.88 | 3.52 | 0.76 | 0.46 | -1.64 | 1.75E-04 | 0.0106 | (B) Small cell-mixed small cell (C/E) |
| BMS1P3 | 1.47 | 3.16 | 1.15 | 0.56 | -1.69 | 2.50E-03 | 0.0391 | (B) Small cell-mixed small cell (C/E) |
| ZNRF3-IT1 | 0.43 | 2.24 | 0.9 | 0.96 | -1.81 | 4.04E-03 | 0.0486 | (B) Small cell-mixed small cell (C/E) |
| RNU6-146P | 3.23 | 5.08 | 1.03 | 0.89 | -1.86 | 3.03E-03 | 0.0426 | (B) Small cell-mixed small cell (C/E) |
| BANF1P4 | 2.54 | 4.44 | 1.22 | 0.39 | -1.9 | 1.37E-03 | 0.0296 | (B) Small cell-mixed small cell (C/E) |
| TRDC | 0.73 | 2.67 | 0.7 | 1.01 | -1.94 | 3.31E-03 | 0.0444 | (B) Small cell-mixed small cell (C/E) |
| LINC00515 | 1.61 | 3.7 | 1.11 | 1.06 | -2.1 | 3.56E-03 | 0.0459 | (B) Small cell-mixed small cell (C/E) |
| ITCH-IT1 | 0.18 | 2.41 | 0.38 | 1.13 | -2.24 | 3.77E-03 | 0.0472 | (B) Small cell-mixed small cell (C/E) |
| SHMT1P1 | 0.62 | 2.87 | 0.83 | 0.99 | -2.25 | 1.10E-03 | 0.0263 | (B) Small cell-mixed small cell (C/E) |
| LINC00310 | 3.32 | 5.69 | 0.57 | 0.47 | -2.37 | 1.14E-06 | 6.83E-04 | (B) Small cell-mixed small cell (C/E) |
| PTGES3P2 | 0.91 | 3.31 | 0.99 | 0.62 | -2.4 | 6.61E-05 | 6.86E-03 | (B) Small cell-mixed small cell (C/E) |
| PDZK1P1 | 2.13 | 4.58 | 1.02 | 1.27 | -2.45 | 3.28E-03 | 0.0442 | (B) Small cell-mixed small cell (C/E) |
| ARAP2 | 1.59 | 4.11 | 1.68 | 1 | -2.52 | 3.03E-03 | 0.0425 | (B) Small cell-mixed small cell (C/E) |
| SNORD56B | 0.23 | 3.07 | 0.69 | 0.88 | -2.84 | 9.30E-05 | 7.97E-03 | (B) Small cell-mixed small cell (C/E) |
| MIR-214/214 | 1.2 | 4.57 | 0.78 | 0.71 | -3.37 | 2.03E-06 | 8.29E-04 | (B) Small cell-mixed small cell (C/E) |
| MIR-214/214 | 1.2 | 4.57 | 0.78 | 0.71 | -3.37 | 2.03E-06 | 8.29E-04 | (B) Small cell-mixed small cell (C/E) |
| MIR-214/3P | 1.2 | 4.57 | 0.78 | 0.71 | -3.37 | 2.03E-06 | 8.29E-04 | (B) Small cell-mixed small cell (C/E) |
| MIR-214/5P | 1.2 | 4.57 | 0.78 | 0.71 | -3.37 | 2.03E-06 | 8.29E-04 | (B) Small cell-mixed small cell (C/E) |
| AKR1B10P1 | 0.03 | 3.61 | 0.09 | 1.3 | -3.58 | 1.06E-03 | 0.0259 | (B) Small cell-mixed small cell (C/E) |
| RN7SL5P | 6.06 | 11.79 | 1.73 | 2.91 | -5.73 | 3.01E-03 | 0.0424 | (B) Small cell-mixed small cell (C/E) |
| RN7SL2 | 9.96 | 15.72 | 1.62 | 3.04 | -5.76 | 3.90E-03 | 0.048 | (B) Small cell-mixed small cell (C/E) |
| RN7SL1 | 9.18 | 14.98 | 1.7 | 3.09 | -5.8 | 3.97E-03 | 0.0484 | (B) Small cell-mixed small cell (C/E) |
