## Supplementary material for "KDM4A promotes the progression of neuroendocrine prostate cancer": Suppl. Table 3

| Suppl. Table 3_DEGs_shKdm4a_vs_shControl | | | | | | | |
|---|---|---|---|---|---|---|---|
| Ensembl Gene ID | Gene Symbo | Mean_Contro | Mean_shKdm | Mean_Chang | LogFC | Pvalue | FDR |
| ENSMUSG00000030087 | Klf15 | 16.1321959 | 284.738457 | 268.606261 | 3.66607867 | 1.1147E-62 | 3.2299E-60 |
| ENSMUSG00000039158 | Akna | 29.338233 | 357.501622 | 328.163389 | 3.2765742 | 5.8933E-60 | 1.5745E-57 |
| ENSMUSG00000030088 | Aldh1l1 | 37.2099581 | 407.156637 | 369.946679 | 3.26520311 | 1.3212E-91 | 1.0067E-88 |
| ENSMUSG00000030827 | Fgf21 | 9.5102673 | 149.260816 | 139.750548 | 3.20546557 | 1.4009E-32 | 8.5613E-31 |
| ENSMUSG00000021536 | Adcy2 | 4.37293722 | 87.1757293 | 82.802792 | 3.06770028 | 4.4701E-23 | 1.4482E-21 |
| ENSMUSG00000000386 | - | 62.9504922 | 678.796187 | 615.845695 | 3.06100614 | 5.5149E-44 | 6.3381E-42 |
| ENSMUSG00000032715 | Trib3 | 1593.69161 | 13561.7233 | 11968.0317 | 2.96300683 | 3.0423E-95 | 2.7211E-92 |
| ENSMUSG00000074570 | Cass4 | 9.48413267 | 110.742375 | 101.258242 | 2.94282599 | 1.7164E-30 | 9.4412E-29 |
| ENSMUSG00000035948 | Acss3 | 20.414082 | 211.374797 | 190.960715 | 2.9305191 | 7.111E-36 | 5.3981E-34 |
| ENSMUSG00000084807 | - | 3.72094269 | 64.1137256 | 60.392783 | 2.92803704 | 9.4472E-22 | 2.7526E-20 |
| ENSMUSG00000031202 | Rab39b | 146.49897 | 1213.46389 | 1066.96492 | 2.92303124 | 1.9761E-93 | 1.6939E-90 |
| ENSMUSG00000027016 | Zfp385b | 29.6130666 | 271.739084 | 242.126017 | 2.85260198 | 9.693E-40 | 9.4954E-38 |
| ENSMUSG00000031616 | Ednra | 127.528251 | 979.608339 | 852.080088 | 2.84086542 | 1.061E-110 | 1.818E-107 |
| ENSMUSG00000030077 | Chl1 | 12.4854008 | 174.861488 | 162.376087 | 2.82585036 | 1.0222E-20 | 2.7451E-19 |
| ENSMUSG00000038496 | Slc19a3 | 1.87099048 | 39.8880425 | 38.017052 | 2.80517437 | 9.9719E-18 | 2.0576E-16 |
| ENSMUSG00000003420 | Fcgrt | 94.082683 | 739.109516 | 645.026833 | 2.78607215 | 6.7755E-58 | 1.6021E-55 |
| ENSMUSG00000019906 | Lin7a | 12.274522 | 121.86409 | 109.589568 | 2.7644063 | 2.5717E-26 | 1.0415E-24 |
| ENSMUSG00000041132 | N4bp2l1 | 83.5821638 | 607.838008 | 524.255844 | 2.744657 | 1.4419E-81 | 8.4754E-79 |
| ENSMUSG00000090084 | Srpx | 16.4173439 | 191.610066 | 175.192722 | 2.7127992 | 4.1908E-20 | 1.0657E-18 |
| ENSMUSG00000022548 | Apod | 6.13003651 | 78.949156 | 72.8191195 | 2.70579348 | 1.0857E-18 | 2.4789E-17 |
| ENSMUSG00000023045 | Bijeta Pradha | 0.40743019 | 33.9061445 | 33.4987143 | 2.69195459 | 2.1087E-14 | 3.1232E-13 |
| ENSMUSG00000097348 | - | 4.19838578 | 58.7497032 | 54.5513174 | 2.68435794 | 9.1421E-18 | 1.8997E-16 |
| ENSMUSG00000086264 | Gm15850 | 1.73096465 | 47.5459906 | 45.815026 | 2.67853074 | 4.5302E-15 | 7.2526E-14 |
| ENSMUSG00000085429 | - | 1.65256076 | 36.5455659 | 34.8930051 | 2.64680439 | 2.2842E-15 | 3.7835E-14 |
| ENSMUSG00000057074 | Ces1g | 29.1599564 | 215.621756 | 186.4618 | 2.63266682 | 2.9268E-39 | 2.7619E-37 |
| ENSMUSG00000056133 | Unc93a2 | 2.01611266 | 36.4824905 | 34.4663779 | 2.6108826 | 2.0973E-15 | 3.4823E-14 |
| ENSMUSG00000054136 | Adm2 | 234.633532 | 1556.33414 | 1321.70061 | 2.60063142 | 5.1887E-64 | 1.6679E-61 |
| ENSMUSG00000030222 | Rerg | 3.30841614 | 40.5601087 | 37.2516926 | 2.59294299 | 8.2558E-17 | 1.577E-15 |
| ENSMUSG00000049723 | Mmp12 | 31.6493775 | 221.503223 | 189.853846 | 2.58633871 | 6.6561E-40 | 6.6794E-38 |
| ENSMUSG00000071047 | Ces1a | 13.0338185 | 143.228463 | 130.194645 | 2.58466567 | 1.3643E-17 | 2.7815E-16 |
| ENSMUSG00000020340 | Cyfip2 | 7.97159773 | 74.7079013 | 66.7363035 | 2.57763255 | 1.7743E-20 | 4.6205E-19 |
| ENSMUSG00000073418 | C4b | 17.4320503 | 136.932633 | 119.500583 | 2.54341851 | 1.5048E-23 | 5.0011E-22 |
| ENSMUSG00000008461 | Fut1 | 3.28737787 | 41.6565579 | 38.3691801 | 2.52699981 | 7.0796E-16 | 1.2311E-14 |
| ENSMUSG00000092021 | Gbp11 | 0.71871283 | 36.9340805 | 36.2153676 | 2.52566499 | 9.9503E-13 | 1.2272E-11 |
| ENSMUSG00000028838 | Extl1 | 12.1648872 | 123.484487 | 111.319599 | 2.49722931 | 2.6432E-16 | 4.7866E-15 |
| ENSMUSG00000020010 | Vnn3 | 22.6925026 | 166.900872 | 144.208369 | 2.49691688 | 1.0785E-25 | 4.218E-24 |
| ENSMUSG00000054123 | - | 0.74484746 | 23.2213699 | 22.4765224 | 2.47862814 | 1.0575E-12 | 1.3011E-11 |
| ENSMUSG00000027358 | Bmp2 | 9.15510638 | 101.79228 | 92.637174 | 2.47039939 | 4.4759E-15 | 7.1768E-14 |
| ENSMUSG00000004609 | Cd33 | 8.09897627 | 68.7596245 | 60.6606482 | 2.46636853 | 2.7371E-19 | 6.5459E-18 |
| ENSMUSG00000064294 | Aox3 | 1.98997803 | 35.5033449 | 33.5133669 | 2.46373117 | 2.8629E-13 | 3.7562E-12 |
| ENSMUSG00000022330 | Osr2 | 1.32353447 | 54.8340299 | 53.5104954 | 2.45170884 | 5.5094E-12 | 6.2653E-11 |
| ENSMUSG00000022661 | Cd200 | 179.525113 | 985.519374 | 805.99426 | 2.38932091 | 3.6963E-93 | 3.0416E-90 |
| ENSMUSG00000052631 | Sh2d6 | 6.05672898 | 59.6106439 | 53.5539149 | 2.38344997 | 1.2073E-14 | 1.8425E-13 |
| ENSMUSG00000043664 | Tmem221 | 26.0549898 | 167.128895 | 141.073905 | 2.38252344 | 5.5155E-28 | 2.5103E-26 |
| ENSMUSG00000036862 | Dchs1 | 7.13060273 | 73.7397389 | 66.6091362 | 2.37413916 | 6.9121E-14 | 9.6558E-13 |
| ENSMUSG00000034463 | Scara3 | 28.4767309 | 205.877813 | 177.401082 | 2.35510048 | 1.9231E-18 | 4.2631E-17 |
| ENSMUSG00000041642 | Kif21b | 394.007211 | 2102.2765 | 1708.26929 | 2.3427942 | 2.9107E-85 | 1.8145E-82 |
| ENSMUSG00000053835 | H2-T24 | 15.4302651 | 150.389161 | 134.958896 | 2.34252737 | 1.4455E-13 | 1.959E-12 |
| ENSMUSG00000041481 | Serpina3g | 371.244335 | 2262.29201 | 1891.04767 | 2.32928364 | 7.313E-26 | 2.8876E-24 |
| ENSMUSG00000053552 | Ebf4 | 6.53417208 | 51.0951517 | 44.5609796 | 2.32737294 | 2.9216E-16 | 5.2722E-15 |
| ENSMUSG00000030717 | Nupr1 | 2414.40061 | 12755.4455 | 10341.0449 | 2.32102624 | 8.2712E-71 | 3.5449E-68 |
| ENSMUSG00000027460 | Angpt4 | 32.7923025 | 206.606071 | 173.813769 | 2.3193711 | 2.5218E-22 | 7.6405E-21 |
| ENSMUSG00000054728 | Phactr1 | 54.9416054 | 309.793632 | 254.852027 | 2.31476289 | 6.9094E-35 | 4.9873E-33 |
| ENSMUSG00000007653 | Gabrb2 | 6.93830764 | 64.9602648 | 58.0219571 | 2.31074614 | 1.2103E-13 | 1.6537E-12 |
| ENSMUSG00000028339 | Col15a1 | 150.16028 | 825.94064 | 675.78036 | 2.27776308 | 9.3211E-35 | 6.6351E-33 |
| ENSMUSG00000038319 | Kcnh2 | 8.57641937 | 59.7257402 | 51.1493209 | 2.27447671 | 5.031E-17 | 9.7732E-16 |
| ENSMUSG00000097758 | - | 1.80097757 | 25.8152651 | 24.0142875 | 2.26972121 | 2.8091E-11 | 2.9499E-10 |
| ENSMUSG00000060924 | Csmd1 | 31.0682359 | 184.689295 | 153.621059 | 2.26832171 | 1.4867E-23 | 4.9561E-22 |
| ENSMUSG00000036390 | Gadd45a | 1689.31154 | 8446.47604 | 6757.1645 | 2.26635767 | 2.0457E-92 | 1.6186E-89 |
| ENSMUSG00000098979 | - | 0.31128264 | 18.2954425 | 17.9841598 | 2.25442943 | 2.8815E-10 | 2.6823E-09 |
| ENSMUSG00000059810 | Rgs3 | 73.3657094 | 378.960056 | 305.594347 | 2.25263828 | 1.7904E-47 | 2.4887E-45 |
| ENSMUSG00000056296 | Synpr | 3.98325069 | 36.0493373 | 32.0660866 | 2.24820521 | 1.4816E-12 | 1.7961E-11 |
| ENSMUSG00000028211 | Trp53inp1 | 492.712871 | 2443.03557 | 1950.3227 | 2.24778207 | 9.0773E-82 | 5.4923E-79 |
| ENSMUSG00000001741 | Il16 | 12.5596701 | 88.0948934 | 75.5352233 | 2.24584066 | 1.7146E-15 | 2.8654E-14 |
| ENSMUSG00000028179 | Cth | 659.191001 | 3337.63913 | 2678.44813 | 2.23411253 | 9.9651E-50 | 1.6532E-47 |
| ENSMUSG00000082483 | - | 10.1454799 | 66.7845754 | 56.6390955 | 2.20918145 | 9.3534E-16 | 1.6115E-14 |
| ENSMUSG00000026638 | Irf6 | 10.6919742 | 68.2486627 | 57.5566885 | 2.20587888 | 4.2614E-16 | 7.5638E-15 |
| ENSMUSG00000046618 | Olfml2a | 9.68152412 | 65.6767215 | 55.9951973 | 2.20341483 | 1.4826E-15 | 2.5E-14 |
| ENSMUSG00000021986 | Amer2 | 82.3822828 | 409.776651 | 327.394368 | 2.20254875 | 4.0074E-44 | 4.6576E-42 |
| ENSMUSG00000025408 | Ddit3 | 2029.32614 | 9497.59502 | 7468.26888 | 2.19949897 | 1.674E-163 | 1.148E-159 |
| ENSMUSG00000006369 | Fbln1 | 440.692738 | 2192.85774 | 1752.16501 | 2.19877586 | 3.9159E-44 | 4.5771E-42 |
| ENSMUSG00000039883 | Lrrc17 | 2.92712058 | 30.4290123 | 27.5018918 | 2.19836484 | 3.5668E-11 | 3.6966E-10 |
| ENSMUSG00000045573 | Penk | 171.864712 | 890.426447 | 718.561736 | 2.19175621 | 8.2029E-30 | 4.2399E-28 |

| | | | | | | | |
|---|---|---|---|---|---|---|---|
| ENSMUSG00000020524 | Gria1 | 8.2077711 | 55.7552781 | 47.547507 | 2.18050608 | 4.4867E-14 | 6.4455E-13 |
| ENSMUSG00000037685 | Atp8a1 | 13.2885756 | 104.267445 | 90.978869 | 2.17737366 | 1.9367E-12 | 2.311E-11 |
| ENSMUSG00000086451 | - | 1.0561301 | 19.6116659 | 18.5555358 | 2.16743503 | 6.4159E-10 | 5.6599E-09 |
| ENSMUSG00000055976 | Cldn23 | 77.8180122 | 365.444044 | 287.626032 | 2.16384064 | 1.4655E-65 | 4.9424E-63 |
| ENSMUSG00000033022 | Cdo1 | 167.697644 | 781.776547 | 614.078903 | 2.15722238 | 3.7236E-75 | 1.741E-72 |
| ENSMUSG00000096221 | - | 69.5341701 | 339.62871 | 270.09454 | 2.15346934 | 1.6942E-36 | 1.3405E-34 |
| ENSMUSG00000048503 | Tlcd5 | 6.21959481 | 81.8573455 | 75.6377507 | 2.14665493 | 6.6683E-10 | 5.8574E-09 |
| ENSMUSG00000087222 | - | 11.2445266 | 66.8875791 | 55.6430525 | 2.14503938 | 4.0227E-16 | 7.1712E-15 |
| ENSMUSG00000049176 | Frmpd4 | 1.60868248 | 22.2048675 | 20.596185 | 2.14082964 | 4.4877E-10 | 4.0527E-09 |
| ENSMUSG00000054409 | Tmem74 | 146.63474 | 682.16873 | 535.53399 | 2.13760047 | 5.4631E-59 | 1.3541E-56 |
| ENSMUSG00000030701 | Plekhb1 | 23.7174018 | 119.118837 | 95.4014356 | 2.13180189 | 1.9918E-26 | 8.1141E-25 |
| ENSMUSG00000085912 | Trp53cor1 | 13.0548568 | 87.2959588 | 74.241102 | 2.1238636 | 6.4514E-13 | 8.1124E-12 |
| ENSMUSG00000021388 | Aspn | 134.000801 | 704.510116 | 570.509315 | 2.12298673 | 7.3843E-21 | 2.0067E-19 |
| ENSMUSG00000048826 | Dact2 | 106.752641 | 488.647145 | 381.894504 | 2.12160729 | 8.2187E-56 | 1.7612E-53 |
| ENSMUSG00000035829 | Ppp1r26 | 26.8226773 | 193.425491 | 166.602814 | 2.11726908 | 6.1851E-12 | 6.9989E-11 |
| ENSMUSG00000037005 | Xpnpep2 | 5.39963807 | 51.0414217 | 45.6417836 | 2.1001836 | 3.0567E-10 | 2.8325E-09 |
| ENSMUSG00000028893 | Sesn2 | 1393.96817 | 6139.54725 | 4745.57909 | 2.09685576 | 2.3084E-96 | 2.1586E-93 |
| ENSMUSG00000013367 | Iglon5 | 8.01547601 | 72.8205821 | 64.8051061 | 2.09515149 | 3.2222E-10 | 2.9765E-09 |
| ENSMUSG00000022758 | P2rx6 | 4.22122579 | 43.1898224 | 38.9685966 | 2.09279877 | 5.4045E-10 | 4.8151E-09 |
| ENSMUSG00000091649 | Phf11b | 57.7236038 | 272.647421 | 214.923817 | 2.09060718 | 2.7766E-30 | 1.4837E-28 |
| ENSMUSG00000073491 | Ifi213 | 0.95998256 | 17.2548881 | 16.2949055 | 2.09001105 | 2.6553E-09 | 2.1677E-08 |
| ENSMUSG00000048004 | Tmem196 | 0.40743019 | 18.1970646 | 17.7896344 | 2.08647557 | 7.3034E-09 | 5.6589E-08 |
| ENSMUSG00000032776 | Mctp2 | 1.98997803 | 24.7470637 | 22.7570856 | 2.08240597 | 1.497E-09 | 1.2611E-08 |
| ENSMUSG00000066363 | Serpina3f | 128.953251 | 609.837689 | 480.884438 | 2.08076166 | 8.0239E-29 | 3.9023E-27 |
| ENSMUSG00000031554 | Adam5 | 3.07224277 | 27.2739176 | 24.2016749 | 2.07695634 | 2.567E-10 | 2.407E-09 |
| ENSMUSG00000043953 | Ccrl2 | 18.9301362 | 103.384049 | 84.4539126 | 2.07016887 | 3.2711E-15 | 5.3154E-14 |
| ENSMUSG00000049690 | Nckap5 | 5.04447715 | 38.5600546 | 33.5155774 | 2.06585787 | 6.4609E-11 | 6.4709E-10 |
| ENSMUSG00000020303 | Stc2 | 661.162805 | 2840.04546 | 2178.88266 | 2.0653313 | 8.6164E-98 | 8.4408E-95 |
| ENSMUSG00000027359 | Slc27a2 | 70.2000825 | 332.55136 | 262.351278 | 2.06345832 | 1.6956E-26 | 6.9902E-25 |
| ENSMUSG00000079363 | Gbp4 | 99.7590781 | 451.180764 | 351.421686 | 2.05734481 | 1.055E-36 | 8.412E-35 |
| ENSMUSG00000005045 | Chd5 | 301.166343 | 1328.21799 | 1027.05164 | 2.05586465 | 6.1438E-51 | 1.0621E-48 |
| ENSMUSG00000039485 | Tspyl4 | 702.6945 | 3046.66319 | 2343.96869 | 2.05368182 | 6.0902E-63 | 1.8158E-60 |
| ENSMUSG00000078144 | Capns2 | 0.31128264 | 15.5954966 | 15.284214 | 2.05070146 | 1.3006E-08 | 9.7044E-08 |
| ENSMUSG00000033152 | Podxl2 | 16.1136122 | 83.4100868 | 67.2964746 | 2.04969589 | 2.5229E-16 | 4.5728E-15 |
| ENSMUSG00000053886 | Sh2d4a | 33.0554506 | 162.304117 | 129.248667 | 2.04849489 | 2.3642E-20 | 6.0948E-19 |
| ENSMUSG00000025776 | Crispld1 | 32.8557262 | 164.349033 | 131.493307 | 2.04747794 | 6.0746E-19 | 1.4073E-17 |
| ENSMUSG00000039364 | Sectm1b | 7.22675027 | 58.4143206 | 51.1875704 | 2.04226966 | 4.6299E-10 | 4.172E-09 |
| ENSMUSG00000020388 | Pdlim4 | 20.645159 | 110.572572 | 89.9274129 | 2.04207703 | 3.2404E-15 | 5.2697E-14 |
| ENSMUSG00000060459 | Kng2 | 38.3388645 | 218.452441 | 180.113577 | 2.04151411 | 1.4642E-13 | 1.9829E-12 |
| ENSMUSG00000029333 | - | 5.22838126 | 36.1309776 | 30.9025964 | 2.04131374 | 4.0083E-11 | 4.125E-10 |
| ENSMUSG00000028031 | Dkk2 | 41.4811202 | 590.562046 | 549.080926 | 2.04017924 | 9.7618E-09 | 7.4268E-08 |
| ENSMUSG00000021226 | Acot2 | 405.898771 | 1771.46662 | 1365.56785 | 2.03761351 | 2.1733E-45 | 2.7096E-43 |
| ENSMUSG00000063142 | Kcnma1 | 2.76096013 | 31.0191629 | 28.2582028 | 2.03202644 | 4.2107E-09 | 3.3471E-08 |
| ENSMUSG00000052572 | Dlg2 | 58.7141646 | 262.662933 | 203.948768 | 2.02693574 | 1.0501E-35 | 7.8843E-34 |
| ENSMUSG00000027562 | Car2 | 22.2226105 | 122.362898 | 100.140288 | 2.02320212 | 1.2106E-13 | 1.6537E-12 |
| ENSMUSG00000025911 | Adhfe1 | 32.7737188 | 160.142589 | 127.36887 | 2.02314525 | 6.8504E-18 | 1.4424E-16 |
| ENSMUSG00000078616 | Trim30c | 14.2645001 | 77.3661805 | 63.1016804 | 2.02224773 | 3.293E-14 | 4.8044E-13 |
| ENSMUSG00000023945 | Slc5a7 | 18.5524442 | 91.3586377 | 72.8061936 | 2.02041778 | 1.2193E-19 | 2.9826E-18 |
| ENSMUSG00000039529 | Atp8b1 | 15.5535089 | 105.144854 | 89.5913455 | 2.01991522 | 9.8678E-11 | 9.6898E-10 |
| ENSMUSG00000042258 | Isl1 | 81.6602753 | 347.356093 | 265.695818 | 2.01929065 | 1.4131E-52 | 2.6427E-50 |
| ENSMUSG00000034610 | Tut4 | 418.674316 | 1756.25217 | 1337.57786 | 2.01644189 | 1.3299E-72 | 5.8211E-70 |
| ENSMUSG00000022510 | Trp63 | 203.909899 | 852.056297 | 648.146398 | 2.01231138 | 4.1336E-69 | 1.6045E-66 |
| ENSMUSG00000071984 | Fndc1 | 2.73482549 | 59.1834347 | 56.4486092 | 2.01078119 | 2.4291E-08 | 1.7473E-07 |
| ENSMUSG00000006777 | Krt23 | 3.86426313 | 31.0924533 | 27.2281902 | 2.01029261 | 1.6569E-09 | 1.3867E-08 |
| ENSMUSG00000078588 | Ccdc24 | 3.91323777 | 41.2660127 | 37.3527749 | 2.01009854 | 6.3806E-09 | 4.9702E-08 |
| ENSMUSG00000032940 | Rbm11 | 2.30126067 | 27.197471 | 24.8962103 | 2.00567808 | 8.0983E-09 | 6.228E-08 |
| ENSMUSG00000085316 | D330050G23 | 1.9638434 | 21.2146397 | 19.2507963 | 2.00344183 | 4.9856E-09 | 3.9327E-08 |
| ENSMUSG00000042500 | Ago4 | 17.6919038 | 89.5471717 | 71.8552679 | 2.00294269 | 8.3335E-16 | 1.4407E-14 |
| ENSMUSG00000045658 | Pid1 | 41.5315877 | 186.959625 | 145.428037 | 2.0025787 | 3.5342E-25 | 1.334E-23 |
| ENSMUSG00000007682 | Dio2 | 72.8936715 | 358.626647 | 285.732976 | 2.00100406 | 4.1584E-17 | 8.1163E-16 |
| ENSMUSG00000000290 | Itgb2 | 9.36843973 | 55.4110424 | 46.0426027 | 1.99835645 | 1.4397E-11 | 1.5622E-10 |
| ENSMUSG00000028542 | Slc6a9 | 1487.44762 | 6144.37602 | 4656.9284 | 1.99271626 | 4.9925E-67 | 1.8019E-64 |
| ENSMUSG00000027313 | Chac1 | 1935.06091 | 7759.56675 | 5824.50584 | 1.99092237 | 1.626E-243 | 3.345E-239 |
| ENSMUSG00000049687 | Pheta2 | 3.00222986 | 28.8304055 | 25.8281756 | 1.98564971 | 5.3306E-09 | 4.1951E-08 |
| ENSMUSG00000000416 | Cttnbp2 | 14.7158085 | 74.745389 | 60.0295805 | 1.9841165 | 1.1088E-14 | 1.7023E-13 |
| ENSMUSG00000093661 | Eif4e3 | 66.316274 | 281.182278 | 214.866004 | 1.98241579 | 1.9748E-39 | 1.8984E-37 |
| ENSMUSG00000052295 | 8030423F21I | 0 | 16.3986259 | 16.3986259 | 1.98048225 | 5.0653E-08 | 3.4642E-07 |
| ENSMUSG00000094910 | D430019H16 | 9.83419723 | 59.6474844 | 49.8132871 | 1.97246472 | 7.0814E-11 | 7.0615E-10 |
| ENSMUSG00000037440 | Vnn1 | 204.647196 | 839.31923 | 634.672034 | 1.96781122 | 1.5355E-51 | 2.6998E-49 |
| ENSMUSG00000042604 | Kcna4 | 38.5456086 | 201.688901 | 163.143292 | 1.96649017 | 3.5456E-13 | 4.6078E-12 |
| ENSMUSG00000074115 | Saa1 | 93.8486553 | 402.288611 | 308.439955 | 1.96494048 | 2.486E-27 | 1.0812E-25 |
| ENSMUSG00000033192 | Lpcat2 | 74.5950196 | 315.584987 | 240.989968 | 1.96318976 | 4.9013E-32 | 2.9311E-30 |
| ENSMUSG00000069733 | Ube2u | 3.3135125 | 32.717766 | 29.4042535 | 1.9607119 | 1.2133E-08 | 9.0999E-08 |
| ENSMUSG00000021390 | Ogn | 424.906902 | 1812.03292 | 1387.12602 | 1.95639237 | 4.1204E-28 | 1.9005E-26 |
| ENSMUSG00000041301 | Cftr | 117.408596 | 724.868179 | 607.459583 | 1.95631807 | 2.6378E-10 | 2.4666E-09 |

| | | | | | | | |
|---|---|---|---|---|---|---|---|
| ENSMUSG00000053141 | Ptprt | 10.6364103 | 59.4386868 | 48.8022764 | 1.95050948 | 1.2041E-11 | 1.3191E-10 |
| ENSMUSG00000054404 | Slfn5 | 2.15613849 | 26.1505071 | 23.9943686 | 1.94776353 | 3.3496E-08 | 2.3607E-07 |
| ENSMUSG00000030930 | Chst15 | 73.3618835 | 331.582309 | 258.220425 | 1.944651 | 7.181E-19 | 1.6543E-17 |
| ENSMUSG00000018427 | Ypel2 | 103.034462 | 414.214237 | 311.179775 | 1.94404003 | 1.6232E-50 | 2.7827E-48 |
| ENSMUSG00000024331 | Dsc2 | 28.5079619 | 163.968557 | 135.460595 | 1.94329223 | 7.1572E-11 | 7.1267E-10 |
| ENSMUSG00000037490 | Slc2a12 | 3.0983774 | 31.5463593 | 28.4479819 | 1.94081031 | 2.2841E-08 | 1.6499E-07 |
| ENSMUSG00000020053 | Igf1 | 14.5775845 | 79.957869 | 65.3802845 | 1.93808887 | 8.2547E-12 | 9.2391E-11 |
| ENSMUSG00000036699 | Zcchc12 | 5.01834252 | 58.2161897 | 53.1978472 | 1.93710048 | 4.1138E-08 | 2.8617E-07 |
| ENSMUSG00000021223 | Papln | 32.8440406 | 160.594381 | 127.75034 | 1.93213949 | 6.3869E-14 | 8.9686E-13 |
| ENSMUSG00000043496 | Tril | 8.57641937 | 49.2164468 | 40.6400275 | 1.92821063 | 8.0963E-11 | 8.0114E-10 |
| ENSMUSG00000027408 | Cpxm1 | 80.7941074 | 332.089244 | 251.295137 | 1.92804396 | 1.4043E-31 | 8.2071E-30 |
| ENSMUSG00000037849 | Ifi206 | 129.765556 | 526.737721 | 396.972165 | 1.92655481 | 1.1641E-38 | 1.0549E-36 |
| ENSMUSG00000043099 | Hic1 | 18.0013847 | 109.534306 | 91.5329212 | 1.92615276 | 6.6259E-10 | 5.8252E-09 |
| ENSMUSG00000029375 | Cxcl15 | 56.9435779 | 276.066036 | 219.122458 | 1.92609478 | 6.2051E-14 | 8.7313E-13 |
| ENSMUSG00000086020 | - | 1.91996512 | 20.0006369 | 18.0806718 | 1.92503219 | 2.4064E-08 | 1.732E-07 |
| ENSMUSG00000020642 | Rnf144a | 42.7361346 | 198.794553 | 156.058418 | 1.92337056 | 2.7567E-15 | 4.5224E-14 |
| ENSMUSG00000028269 | - | 124.44995 | 524.073955 | 399.624006 | 1.91927494 | 1.8999E-24 | 6.7387E-23 |
| ENSMUSG00000025997 | Ikzf2 | 52.8731019 | 225.001944 | 172.128842 | 1.9125938 | 6.5993E-22 | 1.9534E-20 |
| ENSMUSG00000097727 | F630040K05 | 16.0266957 | 85.0205163 | 68.9938206 | 1.9121706 | 1.8401E-11 | 1.9768E-10 |
| ENSMUSG00000040093 | Bmf | 109.151851 | 435.712762 | 326.560911 | 1.91012926 | 1.2586E-37 | 1.0568E-35 |
| ENSMUSG00000097324 | - | 10.3479677 | 55.9010087 | 45.553041 | 1.90540624 | 2.5231E-11 | 2.6659E-10 |
| ENSMUSG00000012819 | Cdh23 | 5.01834252 | 36.7393996 | 31.7210571 | 1.90428489 | 8.27E-09 | 6.3505E-08 |
| ENSMUSG00000049313 | Sorl1 | 64.1846556 | 283.142672 | 218.958016 | 1.90407804 | 5.9929E-18 | 1.2684E-16 |
| ENSMUSG00000046027 | Stard5 | 1465.90215 | 5583.85602 | 4117.95388 | 1.9033987 | 1.296E-110 | 2.051E-107 |
| ENSMUSG00000028751 | Pla2g2e | 2.61254332 | 21.8814795 | 19.2689361 | 1.89903962 | 1.7602E-08 | 1.291E-07 |
| ENSMUSG00000015085 | Entpd2 | 7.69250777 | 57.2427599 | 49.5502522 | 1.89712965 | 1.3578E-08 | 1.0091E-07 |
| ENSMUSG00000025491 | Ifitm1 | 7.86525746 | 66.1905272 | 58.3252697 | 1.89654475 | 3.6862E-08 | 2.5811E-07 |
| ENSMUSG00000043556 | Fbxl7 | 1.7487083 | 20.4013168 | 18.6526085 | 1.89496462 | 7.7644E-08 | 5.1709E-07 |
| ENSMUSG00000028972 | Car6 | 6.889333 | 56.6846856 | 49.7953526 | 1.89349172 | 3.038E-08 | 2.1514E-07 |
| ENSMUSG00000021091 | Serpina3n | 343.030939 | 1527.20697 | 1184.17603 | 1.89207252 | 7.9781E-16 | 1.3815E-14 |
| ENSMUSG00000072653 | Zfp783 | 67.0816285 | 282.590667 | 215.509038 | 1.89146849 | 5.4157E-21 | 1.4915E-19 |
| ENSMUSG00000040569 | Slc26a7 | 16.5918954 | 85.3871946 | 68.7952992 | 1.89017467 | 2.3889E-11 | 2.5306E-10 |
| ENSMUSG00000053199 | Arhgap20 | 7.12550637 | 45.2383289 | 38.1128225 | 1.88494302 | 4.6954E-09 | 3.7151E-08 |
| ENSMUSG00000025656 | Arhgef9 | 52.6454411 | 216.171487 | 163.526046 | 1.88396327 | 8.032E-23 | 2.5578E-21 |
| ENSMUSG00000006386 | Tek | 45.5385185 | 194.876619 | 149.338101 | 1.88212662 | 1.8267E-18 | 4.0581E-17 |
| ENSMUSG00000060487 | Samd5 | 121.376214 | 472.020161 | 350.643947 | 1.88123815 | 6.8754E-46 | 8.7851E-44 |
| ENSMUSG00000003341 | Atp8b3 | 6.38904989 | 34.4482399 | 28.05919 | 1.88084157 | 2.2967E-10 | 2.1654E-09 |
| ENSMUSG00000086283 | - | 8.03321966 | 50.993316 | 42.9600963 | 1.87804946 | 5.4587E-09 | 4.2927E-08 |
| ENSMUSG00000037660 | Gdf7 | 2.13839484 | 21.0838833 | 18.9454885 | 1.87784842 | 7.4602E-08 | 4.9844E-07 |
| ENSMUSG00000028957 | Per3 | 434.446442 | 1648.76047 | 1214.31403 | 1.87722909 | 2.6112E-67 | 9.7667E-65 |
| ENSMUSG00000027070 | Lrp2 | 14.5202188 | 72.3110564 | 57.7908376 | 1.87496179 | 7.539E-12 | 8.4704E-11 |
| ENSMUSG00000059588 | Calcrl | 4.39577723 | 37.5332437 | 33.1374665 | 1.87397324 | 5.0918E-08 | 3.48E-07 |
| ENSMUSG00000043289 | Mei4 | 274.40467 | 1064.47787 | 790.073196 | 1.87297169 | 9.0802E-39 | 8.3021E-37 |
| ENSMUSG00000055368 | Slc6a2 | 3.0983774 | 89.0969485 | 85.9985711 | 1.87101473 | 2.6267E-07 | 1.6261E-06 |
| ENSMUSG00000038151 | Prdm1 | 9.79541531 | 52.6654837 | 42.8700684 | 1.86907074 | 2.168E-10 | 2.0487E-09 |
| ENSMUSG00000039601 | Rcan2 | 13.7492367 | 65.498456 | 51.7492193 | 1.86885729 | 6.8275E-13 | 8.5435E-12 |
| ENSMUSG00000071604 | Fam189a2 | 242.403669 | 948.925886 | 706.522217 | 1.86272267 | 2.3393E-30 | 1.2565E-28 |
| ENSMUSG00000039252 | Lgi2 | 314.492364 | 1568.82747 | 1254.33511 | 1.86133128 | 5.6488E-11 | 5.6908E-10 |
| ENSMUSG00000031099 | Smarca1 | 123.771512 | 484.662076 | 360.890564 | 1.86041193 | 2.6281E-29 | 1.3028E-27 |
| ENSMUSG00000027765 | P2ry1 | 18.6620791 | 84.2355096 | 65.5734306 | 1.85493548 | 1.8107E-14 | 2.7071E-13 |
| ENSMUSG00000030670 | Cyp2r1 | 5.90321581 | 39.5962422 | 33.6930264 | 1.85461173 | 1.7848E-08 | 1.3076E-07 |
| ENSMUSG00000039814 | Xkr5 | 27.2568949 | 118.065271 | 90.8083763 | 1.85439499 | 1.6193E-14 | 2.4369E-13 |
| ENSMUSG00000024924 | Vldlr | 1909.123 | 7027.2851 | 5118.1621 | 1.84880679 | 5.673E-89 | 3.8901E-86 |
| ENSMUSG00000027792 | Bche | 24.4216656 | 131.004596 | 106.582931 | 1.8460988 | 1.1192E-09 | 9.5576E-09 |
| ENSMUSG00000020810 | Cygb | 1.01225182 | 18.9104808 | 17.898229 | 1.84478601 | 3.1379E-07 | 1.9281E-06 |
| ENSMUSG00000032528 | Vipr1 | 57.3636554 | 228.296751 | 170.933096 | 1.83900679 | 2.5312E-22 | 7.6577E-21 |
| ENSMUSG00000031965 | Tbx20 | 6.38904989 | 35.5574811 | 29.1684312 | 1.83370376 | 3.3899E-09 | 2.7359E-08 |
| ENSMUSG00000049422 | Chchd10 | 15.8764772 | 81.9350623 | 66.0585851 | 1.82993119 | 8.2793E-10 | 7.1866E-09 |
| ENSMUSG00000034457 | Eda2r | 55.2538497 | 217.638835 | 162.384986 | 1.82864098 | 2.097E-21 | 5.9502E-20 |
| ENSMUSG00000085443 | - | 2.16452947 | 17.859025 | 15.6944955 | 1.82858295 | 1.0936E-07 | 7.1262E-07 |
| ENSMUSG00000040017 | Saa4 | 1.41968201 | 16.8551366 | 15.4354546 | 1.82816617 | 2.6879E-07 | 1.6625E-06 |
| ENSMUSG00000034438 | Gbp8 | 13.18553 | 64.9315498 | 51.7460199 | 1.82763781 | 2.5568E-10 | 2.3985E-09 |
| ENSMUSG00000030218 | Mgp | 3.93937241 | 29.815117 | 25.8757446 | 1.82296476 | 9.6146E-08 | 6.323E-07 |
| ENSMUSG00000040557 | Mettl27 | 161.888874 | 588.987045 | 427.098171 | 1.81806746 | 1.4329E-52 | 2.6556E-50 |
| ENSMUSG00000039552 | Rsph4a | 4.22122579 | 26.7401988 | 22.518973 | 1.81434246 | 1.5906E-08 | 1.172E-07 |
| ENSMUSG00000026921 | Egfl7 | 50.3180458 | 192.331676 | 142.01363 | 1.81354104 | 7.6698E-25 | 2.8025E-23 |
| ENSMUSG00000060183 | - | 34.7916332 | 289.275958 | 254.484325 | 1.81044565 | 2.2275E-07 | 1.395E-06 |
| ENSMUSG00000041261 | Car8 | 178.095218 | 692.389996 | 514.294778 | 1.80918046 | 5.1369E-21 | 1.4166E-19 |
| ENSMUSG00000021806 | Nid2 | 37.5693753 | 162.160552 | 124.591177 | 1.80662977 | 8.4568E-13 | 1.0499E-11 |
| ENSMUSG00000072568 | Lratd2 | 322.364024 | 1151.52938 | 829.165356 | 1.80600891 | 2.2547E-77 | 1.1313E-74 |
| ENSMUSG00000019890 | Nts | 3.35739078 | 30.075942 | 26.7185512 | 1.79755724 | 2.8269E-07 | 1.7443E-06 |
| ENSMUSG00000042918 | Mamstr | 11.8064316 | 51.5515744 | 39.7451428 | 1.79436324 | 3.0807E-12 | 3.605E-11 |
| ENSMUSG00000052372 | Il1rapl1 | 0.64869992 | 14.1740814 | 13.5253815 | 1.79408028 | 7.069E-07 | 4.108E-06 |
| ENSMUSG00000034949 | Zfr2 | 21.9206805 | 101.432967 | 79.5122863 | 1.79402606 | 1.5059E-10 | 1.4524E-09 |
| ENSMUSG00000098132 | Rassf10 | 89.5199053 | 361.105309 | 271.585404 | 1.79229457 | 1.4509E-15 | 2.4506E-14 |

| | | | | | | | |
|---|---|---|---|---|---|---|---|
| ENSMUSG00000015243 | Abca1 | 42.3113913 | 322.416591 | 280.1052 | 1.79052013 | 2.3829E-07 | 1.4869E-06 |
| ENSMUSG00000026923 | Notch1 | 43.9074264 | 211.042348 | 167.134922 | 1.78987821 | 6.1196E-10 | 5.4077E-09 |
| ENSMUSG00000085862 | Gm13483 | 65.3898553 | 249.505425 | 184.11557 | 1.78947172 | 8.7869E-22 | 2.5787E-20 |
| ENSMUSG00000062960 | Kdr | 45.101659 | 174.949286 | 129.847627 | 1.78798373 | 1.3856E-20 | 3.659E-19 |
| ENSMUSG00000042359 | Osbpl6 | 939.107537 | 3292.7389 | 2353.63136 | 1.78485237 | 9.391E-110 | 1.38E-106 |
| ENSMUSG00000032011 | Thy1 | 31.2679603 | 160.232458 | 128.964498 | 1.78344492 | 4.8284E-09 | 3.8131E-08 |
| ENSMUSG00000027849 | Syt6 | 106.306206 | 387.707238 | 281.401032 | 1.77658327 | 2.3129E-34 | 1.5808E-32 |
| ENSMUSG00000011154 | Cfap161 | 58.3545601 | 240.61358 | 182.25902 | 1.77409129 | 3.3701E-13 | 4.3963E-12 |
| ENSMUSG00000027716 | Trpc3 | 8.87421467 | 40.7926375 | 31.9184228 | 1.76994557 | 4.9691E-10 | 4.4523E-09 |
| ENSMUSG00000038508 | Gdf15 | 41.8573194 | 161.722775 | 119.865455 | 1.76934175 | 9.1927E-18 | 1.9083E-16 |
| ENSMUSG00000051065 | Mb21d2 | 44.5533629 | 167.374396 | 122.821033 | 1.75800114 | 8.7908E-21 | 2.3733E-19 |
| ENSMUSG00000079012 | Serpina3m | 10.7764361 | 50.6271612 | 39.850725 | 1.75783227 | 1.3167E-09 | 1.1157E-08 |
| ENSMUSG00000042675 | Ypel3 | 754.245717 | 2658.73596 | 1904.49025 | 1.7576067 | 2.3026E-41 | 2.4168E-39 |
| ENSMUSG00000020620 | Abca8b | 209.821836 | 761.512234 | 551.690398 | 1.75552254 | 1.498E-26 | 6.2381E-25 |
| ENSMUSG00000079179 | Rab10os | 73.9284545 | 265.404329 | 191.475875 | 1.75408594 | 4.4979E-32 | 2.7056E-30 |
| ENSMUSG00000073568 | Arl14epl | 1.15227765 | 14.7963326 | 13.6440549 | 1.75267758 | 1.0911E-06 | 6.1445E-06 |
| ENSMUSG00000036446 | Lum | 13.1500427 | 66.4750378 | 53.3249951 | 1.75017007 | 1.3882E-08 | 1.0306E-07 |
| ENSMUSG00000031740 | Mmp2 | 597.703746 | 2573.28392 | 1975.58017 | 1.75009044 | 7.0993E-11 | 7.0725E-10 |
| ENSMUSG00000031337 | Mtm1 | 2003.14292 | 6837.9253 | 4834.78238 | 1.74810648 | 1.331E-100 | 1.4414E-97 |
| ENSMUSG00000043259 | Fam13c | 125.54306 | 437.013752 | 311.470692 | 1.7475879 | 5.5745E-47 | 7.5946E-45 |
| ENSMUSG00000031700 | Gpt2 | 6462.80482 | 22176.6207 | 15713.8159 | 1.73942736 | 1.0777E-59 | 2.8064E-57 |
| ENSMUSG00000021416 | Eci3 | 18.1609559 | 72.8975539 | 54.7365981 | 1.73809023 | 1.0666E-12 | 1.3116E-11 |
| ENSMUSG00000016024 | Lbp | 186.363361 | 672.074406 | 485.711045 | 1.73684561 | 3.6559E-23 | 1.1938E-21 |
| ENSMUSG00000062369 | Usp17lb | 19.919548 | 164.299861 | 144.380313 | 1.73531955 | 8.9706E-07 | 5.1434E-06 |
| ENSMUSG00000057465 | Saa2 | 200.98908 | 768.182128 | 567.193048 | 1.73467721 | 2.6929E-15 | 4.4319E-14 |
| ENSMUSG00000062593 | - | 74.203941 | 309.710807 | 235.506866 | 1.73212162 | 4.3935E-11 | 4.4788E-10 |
| ENSMUSG00000021799 | Opn4 | 1.0561301 | 13.5966743 | 12.5405442 | 1.73090586 | 1.452E-06 | 8.036E-06 |
| ENSMUSG00000026051 | Ecrg4 | 18.5506424 | 77.7078746 | 59.1572322 | 1.72897394 | 6.0136E-11 | 6.0524E-10 |
| ENSMUSG00000037211 | Spry1 | 518.565803 | 2445.53546 | 1926.96966 | 1.72876878 | 7.5558E-09 | 5.8413E-08 |
| ENSMUSG00000055639 | Dach1 | 10.1944545 | 78.7228911 | 68.5284366 | 1.72441165 | 8.6298E-07 | 4.9576E-06 |
| ENSMUSG00000034829 | Nxnl1 | 5.47294561 | 30.6321791 | 25.1592335 | 1.7212496 | 1.5346E-07 | 9.8287E-07 |
| ENSMUSG00000029134 | Plb1 | 15.0144439 | 62.440486 | 47.4260421 | 1.72066991 | 2.5573E-11 | 2.6993E-10 |
| ENSMUSG00000068748 | Ptprz1 | 656.201105 | 2858.39087 | 2202.18977 | 1.71533876 | 1.149E-09 | 9.7955E-09 |
| ENSMUSG00000042807 | Hecw2 | 219.998526 | 755.877336 | 535.87881 | 1.71349475 | 6.098E-36 | 4.6635E-34 |
| ENSMUSG00000071317 | Bves | 3.47967295 | 23.7831366 | 20.3034637 | 1.70924897 | 6.2888E-07 | 3.6848E-06 |
| ENSMUSG00000086997 | - | 1.22229056 | 15.8868528 | 14.6645623 | 1.70892343 | 2.2256E-06 | 1.1948E-05 |
| ENSMUSG00000038742 | Angptl6 | 121.278078 | 461.898052 | 340.619974 | 1.70177886 | 2.4104E-13 | 3.1848E-12 |
| ENSMUSG00000034614 | Pik3ip1 | 452.761498 | 1492.38064 | 1039.61914 | 1.70148213 | 7.597E-113 | 1.563E-109 |
| ENSMUSG00000048779 | P2ry6 | 6.56636475 | 46.283682 | 39.7173173 | 1.70055726 | 9.0108E-07 | 5.1606E-06 |
| ENSMUSG00000052920 | Prkg1 | 48.9509419 | 201.029338 | 152.078396 | 1.70002583 | 1.2038E-10 | 1.1776E-09 |
| ENSMUSG00000006931 | P3h4 | 336.660752 | 1157.14452 | 820.48377 | 1.6995185 | 2.9529E-28 | 1.3744E-26 |
| ENSMUSG00000039621 | Prex1 | 20.1298956 | 91.9941275 | 71.8642319 | 1.69926904 | 9.221E-09 | 7.044E-08 |
| ENSMUSG00000040618 | Pck2 | 6866.88209 | 22880.7482 | 16013.8661 | 1.69814725 | 4.7892E-57 | 1.0947E-54 |
| ENSMUSG00000024986 | Hhex | 79.0250136 | 269.279897 | 190.254884 | 1.69703482 | 1.6026E-28 | 7.6496E-27 |
| ENSMUSG00000036356 | Csgalnact1 | 362.233698 | 1187.3369 | 825.103205 | 1.69653159 | 8.125E-107 | 1.045E-103 |
| ENSMUSG00000037625 | Cldn11 | 40.2696752 | 377.056867 | 336.787192 | 1.69586943 | 2.2198E-06 | 1.192E-05 |
| ENSMUSG00000047875 | Gpr157 | 30.7924406 | 122.628338 | 91.8358977 | 1.69485877 | 2.1997E-11 | 2.3386E-10 |
| ENSMUSG00000021994 | Wnt5a | 250.916543 | 890.107016 | 639.190473 | 1.69367217 | 1.4514E-19 | 3.5211E-18 |
| ENSMUSG00000044365 | Cxxc4 | 3.93607778 | 25.5514554 | 21.6153776 | 1.69173788 | 6.022E-07 | 3.5375E-06 |
| ENSMUSG00000049848 | Ceacam19 | 1.02999547 | 20.5259509 | 19.4959554 | 1.68942559 | 3.3237E-06 | 1.7421E-05 |
| ENSMUSG00000036928 | Stag3 | 85.0852245 | 306.18384 | 221.098616 | 1.68937284 | 2.9821E-17 | 5.8988E-16 |
| ENSMUSG00000048852 | Gm12185 | 41.9179796 | 261.142654 | 219.224674 | 1.68643377 | 8.0179E-07 | 4.6281E-06 |
| ENSMUSG00000009185 | Ccl8 | 104.02897 | 375.800928 | 271.771958 | 1.68224199 | 7.1827E-16 | 1.2469E-14 |
| ENSMUSG00000014776 | Nol3 | 107.855944 | 368.708673 | 260.852729 | 1.68132829 | 1.7593E-24 | 6.2615E-23 |
| ENSMUSG00000045620 | Odf3l1 | 1.15227765 | 13.2819156 | 12.129638 | 1.68107162 | 3.0147E-06 | 1.5923E-05 |
| ENSMUSG00000086587 | Gm11837 | 2.54582503 | 18.366121 | 15.820296 | 1.67726827 | 1.5252E-06 | 8.421E-06 |
| ENSMUSG00000054931 | Zkscan4 | 27.7715054 | 101.952047 | 74.1805415 | 1.67484812 | 1.1204E-14 | 1.7175E-13 |
| ENSMUSG00000017943 | Gdap1l1 | 7.9454631 | 35.6896345 | 27.7441714 | 1.67078124 | 1.6352E-08 | 1.2031E-07 |
| ENSMUSG00000048249 | Crebrf | 830.12133 | 2703.94434 | 1873.82301 | 1.67032779 | 7.1396E-64 | 2.2254E-61 |
| ENSMUSG00000086813 | - | 15.8807335 | 83.2392507 | 67.3585172 | 1.6686935 | 3.3483E-07 | 2.0452E-06 |
| ENSMUSG00000025867 | Cplx2 | 41.9324286 | 149.960906 | 108.028477 | 1.66849516 | 4.3042E-16 | 7.6332E-15 |
| ENSMUSG00000085918 | Gm13032 | 4.36454624 | 24.5728688 | 20.2083225 | 1.66824909 | 6.0355E-07 | 3.5436E-06 |
| ENSMUSG00000026489 | Coq8a | 42.2577298 | 149.535764 | 107.278034 | 1.66796783 | 1.5732E-16 | 2.9103E-15 |
| ENSMUSG00000097227 | - | 19.4540984 | 74.4824886 | 55.0283892 | 1.66756305 | 5.4883E-12 | 6.2447E-11 |
| ENSMUSG00000015568 | Lpl | 130.71193 | 579.908837 | 449.196907 | 1.66632725 | 2.2337E-08 | 1.6157E-07 |
| ENSMUSG00000049871 | Nlrc3 | 32.1429498 | 125.1146 | 92.9716505 | 1.66510019 | 3.0547E-11 | 3.1948E-10 |
| ENSMUSG00000028369 | Svep1 | 151.020411 | 534.116371 | 383.09596 | 1.66494563 | 6.5534E-17 | 1.2635E-15 |
| ENSMUSG00000023935 | Spats1 | 19.0677075 | 76.2076132 | 57.1399057 | 1.65921084 | 5.6616E-10 | 5.0312E-09 |
| ENSMUSG00000072620 | Slfn2 | 776.043376 | 2506.11588 | 1730.0725 | 1.65880615 | 1.7358E-58 | 4.2511E-56 |
| ENSMUSG00000085517 | - | 6.12344727 | 31.9669675 | 25.8435203 | 1.65850401 | 1.9839E-07 | 1.2519E-06 |
| ENSMUSG00000039891 | Txlnb | 14.2626983 | 71.9451407 | 57.6824424 | 1.65791417 | 3.117E-07 | 1.9164E-06 |
| ENSMUSG00000053646 | Plxnb1 | 140.370736 | 468.501493 | 328.130757 | 1.65565203 | 4.2472E-28 | 1.9547E-26 |
| ENSMUSG00000035852 | Misp | 6.595794 | 34.5578211 | 27.9620271 | 1.65460779 | 3.9491E-07 | 2.3894E-06 |
| ENSMUSG00000047793 | Sned1 | 250.384276 | 866.775533 | 616.391258 | 1.65416789 | 4.3715E-18 | 9.3482E-17 |
| ENSMUSG00000041992 | Rapgef5 | 5.18450298 | 26.5288203 | 21.3443174 | 1.65231691 | 2.8805E-07 | 1.7747E-06 |

| | | | | | | | |
|---|---|---|---|---|---|---|---|
| ENSMUSG00000023999 | Kif6 | 9.22511929 | 40.6277715 | 31.4026522 | 1.64940978 | 2.4367E-08 | 1.7515E-07 |
| ENSMUSG00000019256 | Ahr | 541.623108 | 1795.42727 | 1253.80417 | 1.64829852 | 3.152E-27 | 1.3623E-25 |
| ENSMUSG00000046447 | Camk2n1 | 137.0553 | 448.066914 | 311.011614 | 1.6479885 | 1.0321E-38 | 9.395E-37 |
| ENSMUSG00000046807 | Lrrc75b | 96.8502324 | 342.519707 | 245.669475 | 1.64349557 | 1.8151E-14 | 2.7117E-13 |
| ENSMUSG00000019647 | Sema6a | 74.6727707 | 353.415086 | 278.742315 | 1.6420426 | 1.9332E-07 | 1.2225E-06 |
| ENSMUSG00000025272 | Tro | 5.1406247 | 25.8981377 | 20.7575131 | 1.6402748 | 3.205E-07 | 1.9658E-06 |
| ENSMUSG00000021792 | Prxl2a | 18.1609559 | 76.5519904 | 58.3910345 | 1.63935619 | 1.6576E-08 | 1.2188E-07 |
| ENSMUSG00000074796 | Slc4a11 | 18.7726756 | 83.2320646 | 64.459389 | 1.63770047 | 4.5177E-08 | 3.1198E-07 |
| ENSMUSG00000056821 | 1700028I16R | 15.0676748 | 61.5220174 | 46.4543425 | 1.63675569 | 4.132E-09 | 3.2896E-08 |
| ENSMUSG00000000794 | Kcnn3 | 4.42191187 | 24.7867819 | 20.36487 | 1.63508267 | 9.4121E-07 | 5.3621E-06 |
| ENSMUSG00000079484 | Phyhd1 | 1.77484293 | 15.8036374 | 14.0287944 | 1.63411433 | 4.7529E-06 | 2.4274E-05 |
| ENSMUSG00000032291 | Crabp1 | 1165.96151 | 4972.83781 | 3806.8763 | 1.63388843 | 3.3331E-08 | 2.3499E-07 |
| ENSMUSG00000032363 | Adamts7 | 338.927704 | 1103.04881 | 764.121104 | 1.63072221 | 4.4814E-30 | 2.3579E-28 |
| ENSMUSG00000051225 | Fam83a | 5.30349053 | 26.9555273 | 21.6520368 | 1.63045016 | 3.8172E-07 | 2.3158E-06 |
| ENSMUSG00000025938 | Slco5a1 | 64.9314264 | 225.240448 | 160.309021 | 1.63015858 | 4.0842E-16 | 7.2681E-15 |
| ENSMUSG00000021203 | Otub2 | 1542.81555 | 4911.571 | 3368.75545 | 1.62947753 | 1.0767E-47 | 1.5171E-45 |
| ENSMUSG00000001986 | Gria3 | 16.61803 | 136.214483 | 119.596453 | 1.62565375 | 5.461E-06 | 2.7657E-05 |
| ENSMUSG00000017754 | Pltp | 488.350966 | 1545.38119 | 1057.03022 | 1.62551373 | 5.2948E-54 | 1.0575E-51 |
| ENSMUSG00000024600 | Slc27a6 | 0.31128264 | 8.72896955 | 8.4176869 | 1.62463876 | 7.8438E-06 | 3.8798E-05 |
| ENSMUSG00000037681 | Esyt3 | 5.57418952 | 30.0816673 | 24.5074778 | 1.62393738 | 1.0867E-06 | 6.124E-06 |
| ENSMUSG00000029298 | Gbp9 | 1455.90957 | 4740.13191 | 3284.22234 | 1.6237221 | 1.645E-26 | 6.8089E-25 |
| ENSMUSG00000046607 | Hrk | 0 | 8.35047864 | 8.35047864 | 1.62232303 | 7.9269E-06 | 3.9181E-05 |
| ENSMUSG00000084915 | - | 2.08612558 | 15.5842722 | 13.4981466 | 1.62204605 | 4.0479E-06 | 2.0928E-05 |
| ENSMUSG00000033595 | Lgi3 | 7.94216848 | 54.1018394 | 46.1596709 | 1.61991728 | 3.7743E-06 | 1.9627E-05 |
| ENSMUSG00000008489 | Elavl2 | 98.0776193 | 317.038341 | 218.960722 | 1.61754323 | 3.1024E-27 | 1.3437E-25 |
| ENSMUSG00000030411 | Nova2 | 1.12614302 | 14.0924682 | 12.9663251 | 1.61611473 | 8.1155E-06 | 4.0027E-05 |
| ENSMUSG00000097812 | - | 6.58314672 | 43.9251546 | 37.3420079 | 1.61584963 | 3.6562E-06 | 1.9052E-05 |
| ENSMUSG00000002012 | Pnck | 9.49677995 | 57.8157818 | 48.3190018 | 1.61581854 | 2.8666E-06 | 1.5195E-05 |
| ENSMUSG00000004113 | Cacna1b | 5.90321581 | 31.5084564 | 25.6052406 | 1.61574824 | 1.2411E-06 | 6.9265E-06 |
| ENSMUSG00000097084 | Foxl1 | 3.13896106 | 20.8196251 | 17.6806641 | 1.61408371 | 2.8337E-06 | 1.5024E-05 |
| ENSMUSG00000032359 | Ctsh | 26.7322789 | 108.13875 | 81.4064713 | 1.61386024 | 1.7173E-08 | 1.26E-07 |
| ENSMUSG00000040829 | Zmynd15 | 45.6533713 | 156.195845 | 110.542473 | 1.61347387 | 2.4347E-15 | 4.0231E-14 |
| ENSMUSG00000043498 | - | 62.574293 | 204.003588 | 141.429295 | 1.61335985 | 9.9941E-25 | 3.6261E-23 |
| ENSMUSG00000055737 | Ghr | 691.174876 | 2207.25312 | 1516.07825 | 1.60984638 | 1.831E-30 | 1.0018E-28 |
| ENSMUSG00000019848 | Popdc3 | 1.36741275 | 12.9608656 | 11.5934529 | 1.60835031 | 7.1946E-06 | 3.5751E-05 |
| ENSMUSG00000067586 | S1pr3 | 192.483701 | 613.617492 | 421.133791 | 1.60772179 | 1.8507E-31 | 1.0725E-29 |
| ENSMUSG00000038456 | Dennd2a | 271.639884 | 895.475453 | 623.835569 | 1.60451959 | 5.3022E-19 | 1.2325E-17 |
| ENSMUSG00000097286 | - | 38.4468193 | 132.694269 | 94.2474502 | 1.60225423 | 6.6006E-13 | 8.2797E-12 |
| ENSMUSG00000008206 | Cers4 | 10.8692891 | 114.084255 | 103.214966 | 1.601973 | 9.6693E-06 | 4.7081E-05 |
| ENSMUSG00000049791 | Fzd4 | 275.549705 | 876.238162 | 600.688456 | 1.59818927 | 2.0673E-29 | 1.0347E-27 |
| ENSMUSG00000040717 | Il17rd | 226.769388 | 748.75926 | 521.989871 | 1.59589374 | 9.7848E-18 | 2.0231E-16 |
| ENSMUSG00000045912 | C2cd4c | 0.64869992 | 11.5584437 | 10.9097438 | 1.59550736 | 1.1242E-05 | 5.4212E-05 |
| ENSMUSG00000021314 | Amph | 11.5912965 | 56.4574935 | 44.866197 | 1.5942097 | 1.0448E-06 | 5.9114E-06 |
| ENSMUSG00000005125 | Ndrg1 | 2648.29726 | 8287.2601 | 5638.96284 | 1.59346468 | 6.5507E-36 | 4.9912E-34 |
| ENSMUSG00000028150 | Rorc | 8.96706759 | 39.6353084 | 30.6682408 | 1.59213871 | 2.6171E-07 | 1.6207E-06 |
| ENSMUSG00000026398 | Nr5a2 | 7.40130171 | 37.5689075 | 30.1676058 | 1.58870453 | 1.693E-06 | 9.2705E-06 |
| ENSMUSG00000067780 | Pi15 | 319.136122 | 1529.54356 | 1210.40744 | 1.58451525 | 1.2268E-06 | 6.8506E-06 |
| ENSMUSG00000059108 | Ifitm6 | 149.031718 | 461.728199 | 312.696482 | 1.582071 | 3.3644E-38 | 2.9203E-36 |
| ENSMUSG00000031389 | Arhgap4 | 6.62192864 | 45.8045901 | 39.1826615 | 1.58168715 | 7.7389E-06 | 3.8316E-05 |
| ENSMUSG00000092274 | Neat1 | 11838.718 | 40072.678 | 28233.96 | 1.58110264 | 3.5791E-13 | 4.6454E-12 |
| ENSMUSG00000006014 | Prg4 | 151.922597 | 669.521372 | 517.598774 | 1.5790611 | 4.8473E-07 | 2.8854E-06 |
| ENSMUSG00000051029 | Serpinb1b | 11.6047839 | 46.4607737 | 34.8559898 | 1.57893305 | 4.1608E-08 | 2.8923E-07 |
| ENSMUSG00000040264 | Gbp2b | 910.068547 | 3294.0409 | 2383.97235 | 1.57889059 | 6.9122E-10 | 6.0639E-09 |
| ENSMUSG00000040613 | Apobec1 | 887.648164 | 2735.73643 | 1848.08826 | 1.57603576 | 2.5828E-37 | 2.1253E-35 |
| ENSMUSG00000071890 | Mroh9 | 2.8832423 | 21.8369195 | 18.9536772 | 1.57600939 | 9.6743E-06 | 4.7094E-05 |
| ENSMUSG00000039084 | Chad | 3.12451203 | 20.1063291 | 16.9818171 | 1.57558681 | 6.4773E-06 | 3.2421E-05 |
| ENSMUSG00000040957 | Cables1 | 18.8829633 | 79.4703717 | 60.5874084 | 1.57273854 | 2.7902E-07 | 1.7227E-06 |
| ENSMUSG00000048368 | Omd | 312.504101 | 1207.09322 | 894.589116 | 1.56550683 | 3.9185E-08 | 2.7345E-07 |
| ENSMUSG00000001663 | Gstt1 | 225.005527 | 679.47721 | 454.471683 | 1.56527411 | 1.4257E-51 | 2.5284E-49 |
| ENSMUSG00000074264 | Amy1 | 15.9086699 | 79.5987886 | 63.6901187 | 1.56441259 | 2.7327E-06 | 1.4526E-05 |
| ENSMUSG00000034059 | Ypel4 | 157.578486 | 494.532307 | 336.953822 | 1.56331698 | 9.5489E-24 | 3.215E-22 |
| ENSMUSG00000054568 | Usp17la | 53.3920903 | 303.24452 | 249.85243 | 1.5628108 | 6.1179E-06 | 3.0772E-05 |
| ENSMUSG00000021306 | Gpr137b | 374.568349 | 1173.77704 | 799.208694 | 1.56067409 | 7.1767E-22 | 2.1182E-20 |
| ENSMUSG00000097325 | Gm16897 | 7.64257144 | 33.6899341 | 26.0473626 | 1.56035295 | 7.2561E-07 | 4.2049E-06 |
| ENSMUSG00000020627 | Klhl29 | 13.6876148 | 51.9467266 | 38.2591118 | 1.55946972 | 8.5298E-09 | 6.5329E-08 |
| ENSMUSG00000050370 | Ch25h | 898.870118 | 2699.73021 | 1800.86009 | 1.55900371 | 4.1472E-58 | 1.0013E-55 |
| ENSMUSG00000045790 | Ccdc149 | 45.1735953 | 168.403711 | 123.230115 | 1.55895672 | 1.1273E-08 | 8.4884E-08 |
| ENSMUSG00000023473 | Celsr3 | 161.121891 | 529.441965 | 368.320074 | 1.5573635 | 3.9536E-14 | 5.7237E-13 |
| ENSMUSG00000049999 | Ppp1r3d | 10.6102757 | 48.9137712 | 38.3034955 | 1.55483184 | 1.4928E-06 | 8.251E-06 |
| ENSMUSG00000033308 | Dpyd | 9.43515802 | 48.2319537 | 38.7967957 | 1.55400799 | 3.9423E-06 | 2.0434E-05 |
| ENSMUSG00000026447 | Pik3c2b | 38.1381784 | 129.712673 | 91.5744943 | 1.55281371 | 1.2417E-11 | 1.358E-10 |
| ENSMUSG00000029061 | Mmp23 | 270.515613 | 831.471054 | 560.955441 | 1.55181659 | 5.2343E-25 | 1.9437E-23 |
| ENSMUSG00000089669 | Tnfsf13 | 9.55414558 | 38.6760852 | 29.1219397 | 1.55169649 | 1.8271E-07 | 1.159E-06 |
| ENSMUSG00000022875 | Kng1 | 8.81439448 | 117.95708 | 109.142685 | 1.54828725 | 2.0665E-05 | 9.5681E-05 |
| ENSMUSG00000052688 | Rab7b | 117.312449 | 394.414537 | 277.102088 | 1.54808638 | 1.6461E-11 | 1.7758E-10 |

| | | | | | | | |
|---|---|---|---|---|---|---|---|
| ENSMUSG00000034839 | Larp6 | 68.7421497 | 212.952654 | 144.210504 | 1.54718227 | 2.2883E-22 | 7.0051E-21 |
| ENSMUSG00000039611 | Pgap4 | 2.82767842 | 18.1079626 | 15.2802842 | 1.54687099 | 8.5939E-06 | 4.2228E-05 |
| ENSMUSG00000078485 | Plekhn1 | 406.990045 | 1210.37946 | 803.389419 | 1.5465855 | 1.0225E-74 | 4.5728E-72 |
| ENSMUSG00000034258 | Flvcr2 | 16.2511835 | 57.3614733 | 41.1102898 | 1.54532485 | 4.3542E-10 | 3.9408E-09 |
| ENSMUSG00000045672 | Col27a1 | 41.1343502 | 164.449544 | 123.315194 | 1.54174046 | 2.6839E-07 | 1.6605E-06 |
| ENSMUSG00000031486 | Adgra2 | 1090.34698 | 3284.96633 | 2194.61935 | 1.54171406 | 1.0504E-34 | 7.4516E-33 |
| ENSMUSG00000022562 | Oplah | 858.478956 | 2513.72518 | 1655.24622 | 1.53762177 | 1.243E-100 | 1.4204E-97 |
| ENSMUSG00000097749 | - | 45.5899477 | 147.166254 | 101.576306 | 1.53556941 | 1.8682E-13 | 2.4908E-12 |
| ENSMUSG00000086782 | E130102H24 | 14.56559 | 51.0296777 | 36.4640877 | 1.53326392 | 1.1128E-08 | 8.3887E-08 |
| ENSMUSG00000026315 | Serpinb8 | 71.9854621 | 221.702899 | 149.717437 | 1.53284477 | 9.0243E-18 | 1.8771E-16 |
| ENSMUSG00000031343 | Gabra3 | 319.775778 | 986.520808 | 666.74503 | 1.53131463 | 7.8257E-20 | 1.942E-18 |
| ENSMUSG00000096940 | - | 7.71768072 | 33.4131208 | 25.6954401 | 1.53092574 | 1.0252E-06 | 5.8087E-06 |
| ENSMUSG00000021281 | Tnfaip2 | 4900.59631 | 14234.7985 | 9334.20221 | 1.52943044 | 2.89E-165 | 2.973E-161 |
| ENSMUSG00000024610 | Cd74 | 37.3374582 | 195.36056 | 158.023102 | 1.52889798 | 7.9727E-06 | 3.9389E-05 |
| ENSMUSG00000055301 | Adh7 | 74.2221151 | 267.845114 | 193.622999 | 1.52811682 | 1.9073E-08 | 1.3914E-07 |
| ENSMUSG00000064202 | Spata6l | 1.39354738 | 14.327581 | 12.9340336 | 1.5276284 | 2.428E-05 | 0.00011097 |
| ENSMUSG00000027977 | Ndst3 | 118.965319 | 356.859422 | 237.894104 | 1.52672806 | 2.9952E-34 | 2.0336E-32 |
| ENSMUSG00000006651 | Aplp1 | 1.67869539 | 18.9825055 | 17.3038101 | 1.52635957 | 2.5917E-05 | 0.0001178 |
| ENSMUSG00000042851 | Zc3h6 | 293.175853 | 867.73085 | 574.554997 | 1.52349068 | 1.771E-44 | 2.1182E-42 |
| ENSMUSG00000017412 | Cacnb4 | 23.0113362 | 77.6898956 | 54.6785594 | 1.52320046 | 7.7386E-10 | 6.74E-09 |
| ENSMUSG00000026678 | Rgs5 | 2466.99988 | 8567.81626 | 6100.81638 | 1.52184854 | 3.9542E-09 | 3.159E-08 |
| ENSMUSG00000097960 | A330074K22 | 106.504129 | 320.072586 | 213.568457 | 1.51899389 | 2.7298E-27 | 1.1848E-25 |
| ENSMUSG00000068606 | Gm4841 | 43.9340923 | 222.39243 | 178.458338 | 1.51869667 | 8.1444E-06 | 4.016E-05 |
| ENSMUSG00000048450 | Msx1 | 6.96264053 | 42.8163522 | 35.8537117 | 1.5181635 | 1.6889E-05 | 7.92E-05 |
| ENSMUSG00000037363 | Letm2 | 88.6914359 | 265.534141 | 176.842705 | 1.51679167 | 1.2706E-26 | 5.3564E-25 |
| ENSMUSG00000036334 | Igsf10 | 4942.73266 | 14702.0772 | 9759.34455 | 1.51357823 | 3.7552E-28 | 1.736E-26 |
| ENSMUSG00000032010 | Usp2 | 62.4180164 | 199.77442 | 137.356403 | 1.51269125 | 2.2632E-12 | 2.6897E-11 |
| ENSMUSG00000033871 | Ppargc1b | 7.31450683 | 32.0455666 | 24.7310597 | 1.5098455 | 2.8757E-06 | 1.5235E-05 |
| ENSMUSG00000028328 | Tmod1 | 126.569761 | 400.123223 | 273.553463 | 1.50846063 | 3.7694E-13 | 4.877E-12 |
| ENSMUSG00000097323 | - | 1.02999547 | 12.8531137 | 11.8231182 | 1.50740443 | 3.3313E-05 | 0.00014882 |
| ENSMUSG00000084094 | - | 3.02836449 | 26.5763002 | 23.5479357 | 1.50739282 | 3.0101E-05 | 0.00013553 |
| ENSMUSG00000043531 | Sorcs1 | 130.418801 | 426.484407 | 296.065607 | 1.50735377 | 7.0019E-11 | 6.989E-10 |
| ENSMUSG00000049086 | Bmyc | 34.3090937 | 120.729112 | 86.4200187 | 1.50503963 | 2.8897E-08 | 2.0532E-07 |
| ENSMUSG00000017969 | Ptgis | 5.09165005 | 23.6755919 | 18.5839418 | 1.5039065 | 6.3978E-06 | 3.2062E-05 |
| ENSMUSG00000038576 | Susd4 | 2.56356867 | 16.4268474 | 13.8632787 | 1.50228489 | 2.259E-05 | 0.00010378 |
| ENSMUSG00000030785 | Cox6a2 | 17.1165114 | 67.0435276 | 49.9270163 | 1.5021653 | 8.3117E-07 | 4.7851E-06 |
| ENSMUSG00000055373 | Fut9 | 467.141944 | 1371.86147 | 904.719523 | 1.50130118 | 1.9816E-31 | 1.1419E-29 |
| ENSMUSG00000038725 | Pkhd1l1 | 22.4722712 | 77.056556 | 54.5842848 | 1.49975405 | 1.0441E-08 | 7.9087E-08 |
| ENSMUSG00000062866 | Phactr2 | 16.2740235 | 68.9292867 | 52.6552632 | 1.49866776 | 2.38E-06 | 1.2724E-05 |
| ENSMUSG00000063488 | Zkscan7 | 36.4795596 | 114.109832 | 77.6302719 | 1.49834867 | 1.103E-13 | 1.5167E-12 |
| ENSMUSG00000086589 | Gm12915 | 0.74484746 | 9.8381175 | 9.09327004 | 1.49832416 | 3.7444E-05 | 0.00016562 |
| ENSMUSG00000027004 | Frzb | 10.685385 | 40.7842801 | 30.0988951 | 1.49819829 | 2.4785E-07 | 1.5418E-06 |
| ENSMUSG00000000782 | Tcf7 | 118.843568 | 359.725072 | 240.881504 | 1.49696198 | 9.2321E-18 | 1.9145E-16 |
| ENSMUSG00000030465 | Psd3 | 998.180386 | 2909.64602 | 1911.46563 | 1.49691925 | 1.6442E-34 | 1.1427E-32 |
| ENSMUSG00000087522 | - | 0.40743019 | 11.1949885 | 10.7875583 | 1.49593034 | 3.7894E-05 | 0.0001675 |
| ENSMUSG00000054640 | Slc8a1 | 20.5979861 | 72.8608487 | 52.2628626 | 1.49232572 | 4.9604E-08 | 3.3981E-07 |
| ENSMUSG00000017446 | C1qtnf1 | 268.129942 | 772.574408 | 504.444467 | 1.4904029 | 1.2868E-48 | 1.9182E-46 |
| ENSMUSG00000048388 | Fam171b | 147.817596 | 439.18398 | 291.366384 | 1.48814366 | 9.1188E-22 | 2.6646E-20 |
| ENSMUSG00000029771 | Irf5 | 68.6619441 | 204.300751 | 135.638807 | 1.48413969 | 3.1255E-18 | 6.8112E-17 |
| ENSMUSG00000041658 | Rragb | 107.46063 | 308.417792 | 200.957163 | 1.48113436 | 5.2696E-33 | 3.285E-31 |
| ENSMUSG00000058254 | Tspan7 | 378.162409 | 1421.38133 | 1043.21892 | 1.48035412 | 7.3761E-07 | 4.2732E-06 |
| ENSMUSG00000039706 | Ldb2 | 148.350603 | 433.949806 | 285.599203 | 1.48006836 | 1.5558E-25 | 5.9937E-24 |
| ENSMUSG00000019823 | Mical1 | 475.350934 | 1352.03698 | 876.686049 | 1.47925618 | 6.3887E-49 | 9.8819E-47 |
| ENSMUSG00000032066 | Bco2 | 5.7977156 | 28.7930362 | 22.9953206 | 1.47767178 | 1.4053E-05 | 6.6859E-05 |
| ENSMUSG00000097194 | - | 40.9183751 | 129.709349 | 88.7909741 | 1.4758755 | 1.2425E-10 | 1.2137E-09 |
| ENSMUSG00000038349 | Plcl1 | 24.5508458 | 91.2558281 | 66.7049822 | 1.47569636 | 5.6266E-07 | 3.32E-06 |
| ENSMUSG00000087535 | - | 21.9113278 | 68.9274762 | 47.0161484 | 1.47185332 | 1.7567E-10 | 1.6785E-09 |
| ENSMUSG00000087095 | Emx2os | 54.8911379 | 165.052979 | 110.161841 | 1.47118864 | 1.7603E-15 | 2.937E-14 |
| ENSMUSG00000090248 | - | 4.86992571 | 21.7750712 | 16.9051455 | 1.47052008 | 5.599E-06 | 2.8307E-05 |
| ENSMUSG00000037813 | D630003M21 | 13.0809914 | 56.3361598 | 43.2551683 | 1.46959675 | 7.3799E-06 | 3.6601E-05 |
| ENSMUSG00000079017 | Ifi27l2a | 274.674442 | 820.998707 | 546.324265 | 1.46614762 | 5.328E-15 | 8.4314E-14 |
| ENSMUSG00000062901 | Klhl24 | 1505.5836 | 4255.88882 | 2750.30522 | 1.46518782 | 2.2644E-42 | 2.5045E-40 |
| ENSMUSG00000033794 | Lpcat2b | 1.0561301 | 9.76675284 | 8.71062273 | 1.46471312 | 5.1602E-05 | 0.00022339 |
| ENSMUSG00000052117 | D630039A03 | 8.37902792 | 30.1254648 | 21.7464369 | 1.4646352 | 4.0109E-07 | 2.424E-06 |
| ENSMUSG00000053218 | - | 25.7773927 | 184.629112 | 158.851719 | 1.46441134 | 4.793E-05 | 0.00020859 |
| ENSMUSG00000089672 | - | 38.2241332 | 195.418374 | 157.194241 | 1.46418492 | 2.3717E-05 | 0.00010853 |
| ENSMUSG00000020102 | Slc16a7 | 5.61477317 | 27.0945634 | 21.4797902 | 1.46339103 | 1.4455E-05 | 6.8599E-05 |
| ENSMUSG00000045467 | Ttll13 | 2.68585085 | 14.4913842 | 11.8055334 | 1.46337472 | 2.98E-05 | 0.00013424 |
| ENSMUSG00000056498 | Tmem154 | 0 | 7.45649552 | 7.45649552 | 1.46331371 | 5.0953E-05 | 0.00022095 |
| ENSMUSG00000099065 | - | 112.501999 | 334.615852 | 222.113853 | 1.46182555 | 5.5266E-16 | 9.6924E-15 |
| ENSMUSG00000054612 | Mgmt | 59.1128949 | 202.158164 | 143.045269 | 1.46166195 | 8.9951E-08 | 5.9405E-07 |
| ENSMUSG00000090231 | Cfb | 38.7198512 | 122.423447 | 83.7035955 | 1.46095878 | 1.0649E-09 | 9.1168E-09 |
| ENSMUSG00000025175 | Fn3k | 0.74484746 | 9.44037036 | 8.6955229 | 1.460091 | 5.9241E-05 | 0.00025411 |
| ENSMUSG00000008932 | Slc1a7 | 9.06321513 | 34.7919842 | 25.7287691 | 1.45894495 | 2.0328E-06 | 1.0982E-05 |
| ENSMUSG00000053414 | Hunk | 16.3404329 | 58.1939382 | 41.8535052 | 1.45768393 | 6.1642E-07 | 3.6159E-06 |

| | | | | | | | |
|---|---|---|---|---|---|---|---|
| ENSMUSG00000018819 | Lsp1 | 178.108296 | 553.979943 | 375.871648 | 1.45520433 | 1.4205E-10 | 1.3765E-09 |
| ENSMUSG00000019539 | Rcn3 | 537.17043 | 1784.86875 | 1247.69832 | 1.45482006 | 3.4401E-08 | 2.4228E-07 |
| ENSMUSG00000023913 | Pla2g7 | 652.824052 | 1840.671 | 1187.84695 | 1.45446763 | 1.887E-33 | 1.2284E-31 |
| ENSMUSG00000097768 | 2310043M15 | 3.57252588 | 20.2477103 | 16.6751844 | 1.45343881 | 3.1981E-05 | 0.00014315 |
| ENSMUSG00000070737 | Tmem35b | 71.6143593 | 205.411613 | 133.797254 | 1.45318934 | 1.5927E-19 | 3.8592E-18 |
| ENSMUSG00000034613 | Ppm1h | 38.0841074 | 113.692217 | 75.6081094 | 1.45281476 | 4.8612E-14 | 6.9544E-13 |
| ENSMUSG00000032243 | Itga11 | 10.3116403 | 37.5804019 | 27.2687616 | 1.45183251 | 1.2177E-06 | 6.8052E-06 |
| ENSMUSG00000056004 | Elapor2 | 272.138056 | 782.770993 | 510.632936 | 1.44913153 | 1.64E-20 | 4.2868E-19 |
| ENSMUSG00000021998 | Lcp1 | 129.7219 | 374.458583 | 244.736683 | 1.44867213 | 8.9938E-19 | 2.0696E-17 |
| ENSMUSG00000046714 | Foxc2 | 32.9527138 | 101.695221 | 68.7425069 | 1.44830772 | 1.7022E-11 | 1.8334E-10 |
| ENSMUSG00000032496 | Ltf | 4.56703405 | 25.9316798 | 21.3646457 | 1.44679103 | 3.7985E-05 | 0.00016783 |
| ENSMUSG00000039196 | Orm1 | 2.74321648 | 18.0368688 | 15.2936523 | 1.44356278 | 5.3325E-05 | 0.00023027 |
| ENSMUSG00000060548 | Tnfrsf19 | 123.855321 | 398.80436 | 274.949039 | 1.44296391 | 1.0739E-08 | 8.1281E-08 |
| ENSMUSG00000026748 | Plxdc2 | 34.911992 | 123.292149 | 88.3801574 | 1.4424126 | 7.077E-07 | 4.1104E-06 |
| ENSMUSG00000046157 | Tmem229b | 17.7096474 | 55.6050748 | 37.8954274 | 1.44148261 | 1.5089E-09 | 1.2696E-08 |
| ENSMUSG00000021596 | Mctp1 | 14.0199357 | 48.9369497 | 34.917014 | 1.44047066 | 3.3164E-07 | 2.027E-06 |
| ENSMUSG00000031431 | Tsc22d3 | 2037.13968 | 6665.03111 | 4627.89143 | 1.44028221 | 3.897E-08 | 2.7204E-07 |
| ENSMUSG00000033538 | Casp4 | 610.977169 | 1748.16252 | 1137.18535 | 1.43985446 | 2.0523E-19 | 4.9379E-18 |
| ENSMUSG00000097993 | Ptprv | 12.1438489 | 44.5226437 | 32.3787948 | 1.43976917 | 1.6095E-06 | 8.853E-06 |
| ENSMUSG00000024397 | Aif1 | 19.1377204 | 65.2017994 | 46.064079 | 1.43942117 | 2.309E-07 | 1.4434E-06 |
| ENSMUSG00000070644 | Etnk2 | 3.39191639 | 19.4085823 | 16.0166659 | 1.43871435 | 4.4124E-05 | 0.00019317 |
| ENSMUSG00000053541 | - | 0.33741727 | 10.3381427 | 10.0007254 | 1.43864474 | 7.2151E-05 | 0.00030478 |
| ENSMUSG00000097804 | Gm16685 | 29.4054825 | 92.1352051 | 62.7297226 | 1.43861905 | 2.888E-09 | 2.3474E-08 |
| ENSMUSG00000034127 | Tspan8 | 1.0561301 | 9.03773896 | 7.98160886 | 1.43611857 | 7.1037E-05 | 0.00030045 |
| ENSMUSG00000026765 | Lypd6b | 109.267012 | 315.566407 | 206.299395 | 1.43546232 | 3.5761E-17 | 7.0265E-16 |
| ENSMUSG00000027009 | Itga4 | 122.222427 | 365.199564 | 242.977137 | 1.43542028 | 3.59E-12 | 4.1538E-11 |
| ENSMUSG00000028524 | Sgip1 | 9.96996675 | 37.4443676 | 27.4744008 | 1.43374849 | 2.9572E-06 | 1.5635E-05 |
| ENSMUSG00000064373 | Selenop | 1198.19022 | 3725.44164 | 2527.25142 | 1.4324284 | 1.7903E-09 | 1.4947E-08 |
| ENSMUSG00000074971 | Fibin | 48.3323241 | 417.050107 | 368.717783 | 1.43121448 | 8.1118E-05 | 0.00033897 |
| ENSMUSG00000025105 | Bnc1 | 121.31854 | 355.133029 | 233.81449 | 1.43074579 | 1.0381E-13 | 1.4313E-12 |
| ENSMUSG00000060969 | Irx1 | 1392.29213 | 3852.97928 | 2460.68715 | 1.4280251 | 1.1942E-32 | 7.3776E-31 |
| ENSMUSG00000040276 | Pacsin1 | 2.06838193 | 12.5931404 | 10.5247584 | 1.42788208 | 5.6358E-05 | 0.0002425 |
| ENSMUSG00000029121 | Crmp1 | 33.5407534 | 126.168118 | 92.6273642 | 1.42774029 | 4.0274E-06 | 2.0827E-05 |
| ENSMUSG00000042249 | Grk3 | 7.03445518 | 30.4737296 | 23.4392744 | 1.42616501 | 1.6489E-05 | 7.7481E-05 |
| ENSMUSG00000051811 | Cox6b2 | 1.48969492 | 13.276809 | 11.7871141 | 1.42531423 | 8.5441E-05 | 0.00035573 |
| ENSMUSG00000087006 | Gm13889 | 25.0129998 | 78.5429926 | 53.5299928 | 1.42502805 | 1.7923E-08 | 1.3116E-07 |
| ENSMUSG00000020865 | Abcc3 | 39.7577064 | 117.709053 | 77.9513464 | 1.42493387 | 5.6008E-12 | 6.3587E-11 |
| ENSMUSG00000021846 | Gm6498; Pel | 398.763999 | 1092.04248 | 693.278476 | 1.42470632 | 5.9745E-54 | 1.1705E-51 |
| ENSMUSG00000040026 | Saa3 | 74015.8488 | 204339.932 | 130324.083 | 1.42270381 | 1.3135E-31 | 7.6987E-30 |
| ENSMUSG00000059824 | Dbp | 407.674267 | 1140.58044 | 732.906173 | 1.42200254 | 1.9717E-22 | 6.0722E-21 |
| ENSMUSG00000072115 | Ang | 16.9993256 | 52.9550855 | 35.9557599 | 1.42183753 | 3.9774E-09 | 3.1764E-08 |
| ENSMUSG00000000753 | Serpinf1 | 1648.0343 | 4873.29998 | 3225.26568 | 1.42161986 | 8.0055E-12 | 8.9846E-11 |
| ENSMUSG00000037362 | Ccn3 | 9141.09028 | 25135.2178 | 15994.1275 | 1.42114101 | 1.3715E-34 | 9.6623E-33 |
| ENSMUSG00000010660 | Plcd1 | 192.405828 | 553.227915 | 360.822087 | 1.42073004 | 4.895E-15 | 7.8062E-14 |
| ENSMUSG00000036766 | Dner | 1.98997803 | 11.7449948 | 9.75501678 | 1.42046563 | 5.6233E-05 | 0.00024206 |
| ENSMUSG00000027221 | Chst1 | 508.421917 | 1430.32799 | 921.906068 | 1.41996191 | 8.8711E-20 | 2.1961E-18 |
| ENSMUSG00000019932 | Kera | 1.01225182 | 11.9158683 | 10.9036165 | 1.41817091 | 9.6183E-05 | 0.00039733 |
| ENSMUSG00000031997 | Trpc6 | 154.232406 | 442.535106 | 288.3027 | 1.41766089 | 1.3575E-14 | 2.0625E-13 |
| ENSMUSG00000074818 | Pdzd7 | 28.5383528 | 95.1683833 | 66.6300305 | 1.41636659 | 3.5532E-07 | 2.1626E-06 |
| ENSMUSG00000047370 | - | 5.60967681 | 24.2634479 | 18.6537711 | 1.41559884 | 2.254E-05 | 0.00010359 |
| ENSMUSG00000028766 | Alpl | 112.887121 | 314.120631 | 201.23351 | 1.41442003 | 7.227E-24 | 2.4493E-22 |
| ENSMUSG00000079657 | Rab26 | 4.94323325 | 28.0482524 | 23.1050191 | 1.41408794 | 6.7172E-05 | 0.00028592 |
| ENSMUSG00000002565 | Scin | 12.2805801 | 45.8854701 | 33.60489 | 1.41315388 | 5.0514E-06 | 2.5729E-05 |
| ENSMUSG00000029009 | Mthfr | 1560.69815 | 4212.15878 | 2651.46062 | 1.40850457 | 2.7581E-52 | 5.0661E-50 |
| ENSMUSG00000022883 | Robo1 | 0.98611719 | 9.21515095 | 8.22903376 | 1.40820364 | 0.00010273 | 0.00042217 |
| ENSMUSG00000042686 | Jph1 | 242.310285 | 671.014824 | 428.704539 | 1.4074138 | 1.6317E-22 | 5.0706E-21 |
| ENSMUSG00000012428 | Steap4 | 1920.51646 | 5136.47587 | 3215.95941 | 1.40519892 | 6.7942E-86 | 4.5087E-83 |
| ENSMUSG00000050035 | Fhl4 | 97.8282674 | 272.496717 | 174.66845 | 1.40320439 | 5.0297E-21 | 1.3907E-19 |
| ENSMUSG00000076441 | Ass1 | 126.911004 | 367.963308 | 241.052304 | 1.40012524 | 1.2698E-11 | 1.3858E-10 |
| ENSMUSG00000021260 | Hhipl1 | 239.593491 | 650.377367 | 410.783875 | 1.39964983 | 1.9567E-33 | 1.2698E-31 |
| ENSMUSG00000044667 | Plppr4 | 119.452114 | 376.988443 | 257.536328 | 1.3978866 | 8.5243E-08 | 5.6459E-07 |
| ENSMUSG00000025783 | - | 2.30126067 | 13.1732114 | 10.8719507 | 1.39633002 | 7.6762E-05 | 0.00032195 |
| ENSMUSG00000000409 | Lck | 149.264409 | 408.582563 | 259.318155 | 1.39619563 | 1.7298E-21 | 4.9425E-20 |
| ENSMUSG00000022408 | Fam83f | 1.0561301 | 8.83564665 | 7.77951655 | 1.39527951 | 0.00011698 | 0.00047563 |
| ENSMUSG00000023952 | Gtpbp2 | 3099.81761 | 8230.31407 | 5130.49646 | 1.39332156 | 4.2739E-77 | 2.0934E-74 |
| ENSMUSG00000035873 | Pawr | 409.088631 | 1114.4264 | 705.337765 | 1.39302344 | 5.7468E-25 | 2.1263E-23 |
| ENSMUSG00000052921 | Arhgef15 | 54.8793307 | 171.26301 | 116.383679 | 1.39186826 | 4.9978E-08 | 3.4226E-07 |
| ENSMUSG00000049555 | Tmie | 8.24229672 | 38.5948717 | 30.352575 | 1.39175621 | 5.7606E-05 | 0.00024761 |
| ENSMUSG00000072949 | Acot1 | 281.356865 | 789.693001 | 508.336136 | 1.39118516 | 1.4569E-14 | 2.2022E-13 |
| ENSMUSG00000061740 | Cyp2d22 | 213.191013 | 620.020006 | 406.828993 | 1.39104626 | 7.3258E-11 | 7.2841E-10 |
| ENSMUSG00000074207 | Adh1 | 0.64869992 | 11.1880833 | 10.5393834 | 1.39035712 | 0.00012906 | 0.0005207 |
| ENSMUSG00000037902 | Sirpa | 226.090555 | 828.858139 | 602.767584 | 1.38997915 | 9.2104E-06 | 4.5017E-05 |
| ENSMUSG00000045180 | Shroom2 | 264.308373 | 833.123658 | 568.815285 | 1.38981938 | 1.3703E-07 | 8.8146E-07 |
| ENSMUSG00000078952 | - | 8.6380413 | 32.0386616 | 23.4006203 | 1.38884579 | 9.061E-06 | 4.436E-05 |
| ENSMUSG00000038370 | Pcp4l1 | 2.37127359 | 15.3214943 | 12.9502208 | 1.38852814 | 0.00010966 | 0.00044806 |

| | | | | | | | |
|---|---|---|---|---|---|---|---|
| ENSMUSG00000032839 | Trpc1 | 102.968052 | 299.25355 | 196.285498 | 1.38820513 | 7.609E-11 | 7.5546E-10 |
| ENSMUSG00000001493 | Meox1 | 55.7032349 | 157.161655 | 101.458421 | 1.38737314 | 3.1002E-13 | 4.0597E-12 |
| ENSMUSG00000049929 | Lpar4 | 2625.43309 | 6939.86492 | 4314.43183 | 1.38579966 | 1.8725E-69 | 7.5531E-67 |
| ENSMUSG00000027583 | Zbtb46 | 33.5164205 | 111.459781 | 77.9433606 | 1.38562127 | 9.8558E-07 | 5.5932E-06 |
| ENSMUSG00000053049 | Gm15413 | 26.707946 | 83.7145077 | 57.0065617 | 1.38552109 | 8.9801E-08 | 5.9325E-07 |
| ENSMUSG00000074577 | Ripor3 | 4.6319506 | 19.1879896 | 14.556039 | 1.38506147 | 3.4551E-05 | 0.00015392 |
| ENSMUSG00000044703 | Phf11a | 4.22452042 | 18.0653351 | 13.8408146 | 1.38479929 | 4.0816E-05 | 0.00017961 |
| ENSMUSG00000023947 | Nfkbie | 499.248105 | 1335.32678 | 836.07867 | 1.38438995 | 4.1043E-34 | 2.7593E-32 |
| ENSMUSG00000097715 | - | 308.985955 | 821.410542 | 512.424587 | 1.38423045 | 2.7339E-45 | 3.388E-43 |
| ENSMUSG00000068740 | Celsr2 | 3.64583341 | 17.4065825 | 13.7607491 | 1.38165385 | 7.3547E-05 | 0.00030998 |
| ENSMUSG00000049307 | Fut4 | 66.9896156 | 187.525078 | 120.535463 | 1.38134443 | 2.7868E-15 | 4.5681E-14 |
| ENSMUSG00000021944 | Gata4 | 2.54582503 | 13.0490244 | 10.5031994 | 1.38109443 | 8.5981E-05 | 0.00035777 |
| ENSMUSG00000019320 | Noxo1 | 23.3766899 | 69.491119 | 46.1144292 | 1.38029209 | 1.8307E-09 | 1.526E-08 |
| ENSMUSG00000026649 | Cfap126 | 26.3314379 | 77.3099779 | 50.9785399 | 1.37791092 | 7.6993E-09 | 5.9411E-08 |
| ENSMUSG00000036306 | Lzts1 | 11.2081992 | 37.6745463 | 26.4663471 | 1.37644226 | 5.2241E-06 | 2.6569E-05 |
| ENSMUSG00000058325 | Dock1 | 626.663326 | 1663.05798 | 1036.39465 | 1.37630817 | 1.3771E-37 | 1.1469E-35 |
| ENSMUSG00000086539 | - | 3.55478223 | 17.8731966 | 14.3184144 | 1.37626776 | 7.5504E-05 | 0.00031719 |
| ENSMUSG00000038550 | Ciart | 513.360464 | 1369.82866 | 856.468199 | 1.37602014 | 4.9847E-30 | 2.6159E-28 |
| ENSMUSG00000073514 | Dok6 | 23.1582601 | 68.7452535 | 45.5869933 | 1.37457547 | 2.4168E-09 | 1.9856E-08 |
| ENSMUSG00000036611 | Eepd1 | 69.9963241 | 189.380917 | 119.384593 | 1.37343189 | 3.5143E-17 | 6.9116E-16 |
| ENSMUSG00000097021 | 4933433G15 | 8.20447648 | 28.6775188 | 20.4730423 | 1.37329437 | 5.3124E-06 | 2.6951E-05 |
| ENSMUSG00000022788 | Fgd4 | 30.3360358 | 100.614055 | 70.2780193 | 1.37198991 | 2.2052E-06 | 1.1851E-05 |
| ENSMUSG00000097164 | - | 182.666817 | 484.361633 | 301.694816 | 1.37173091 | 5.9404E-34 | 3.9807E-32 |
| ENSMUSG00000024990 | Rbp4 | 433.281056 | 1169.89768 | 736.616625 | 1.37152711 | 1.8971E-20 | 4.934E-19 |
| ENSMUSG00000025255 | Zfhx4 | 76.6132429 | 231.806005 | 155.192762 | 1.37057961 | 3.4783E-08 | 2.4447E-07 |
| ENSMUSG00000019856 | Fam184a | 25.8380529 | 80.0965018 | 54.2584489 | 1.36844018 | 2.4647E-07 | 1.5341E-06 |
| ENSMUSG00000096210 | H1f0 | 5730.18951 | 15074.992 | 9344.80248 | 1.36815895 | 5.5851E-42 | 6.0472E-40 |
| ENSMUSG00000069170 | Adgrv1 | 59.1533569 | 169.438533 | 110.285176 | 1.36760592 | 1.0916E-10 | 1.0683E-09 |
| ENSMUSG00000035211 | Xrra1 | 13.1281643 | 43.1412464 | 30.013082 | 1.36747406 | 3.5058E-06 | 1.8305E-05 |
| ENSMUSG00000086401 | Gm15559 | 19.527407 | 58.5988307 | 39.0714238 | 1.36543495 | 4.4689E-08 | 3.0892E-07 |
| ENSMUSG00000029757 | Dync1i1 | 19.8970169 | 59.3528128 | 39.4557959 | 1.36528315 | 1.4036E-08 | 1.0409E-07 |
| ENSMUSG00000060429 | Sntb1 | 25.5775467 | 76.7692717 | 51.191725 | 1.36387025 | 1.227E-08 | 9.1954E-08 |
| ENSMUSG00000069873 | 4930438A08 | 0.98611719 | 8.56204603 | 7.57592884 | 1.36077567 | 0.00017536 | 0.00069029 |
| ENSMUSG00000071656 | Lrrn4cl | 32.4422379 | 98.9832886 | 66.5410507 | 1.36038135 | 1.6939E-07 | 1.0792E-06 |
| ENSMUSG00000036067 | Slc2a6 | 197.284267 | 526.544034 | 329.259767 | 1.36024777 | 6.278E-22 | 1.8636E-20 |
| ENSMUSG00000031297 | Slc7a3 | 1915.25019 | 5115.27779 | 3200.0276 | 1.35967753 | 2.0391E-21 | 5.794E-20 |
| ENSMUSG00000040428 | Plekha4 | 13.5196526 | 47.6620275 | 34.1423749 | 1.35904905 | 1.1636E-05 | 5.6021E-05 |
| ENSMUSG00000054588 | - | 850.591507 | 2406.38892 | 1555.79742 | 1.35765554 | 1.4415E-10 | 1.3949E-09 |
| ENSMUSG00000032333 | Stoml1 | 676.297751 | 1759.82985 | 1083.5321 | 1.35755984 | 8.7803E-47 | 1.1806E-44 |
| ENSMUSG00000057706 | Mex3b | 1436.1548 | 3766.41877 | 2330.26397 | 1.35553668 | 1.6435E-32 | 9.9733E-31 |
| ENSMUSG00000035773 | Kiss1r | 44.8664473 | 130.183444 | 85.3169964 | 1.3552896 | 1.6849E-09 | 1.4096E-08 |
| ENSMUSG00000047842 | Diras2 | 153.337183 | 416.114142 | 262.776959 | 1.35459551 | 1.8253E-16 | 3.3407E-15 |
| ENSMUSG00000097184 | 4632428C04 | 2.99713349 | 14.612008 | 11.6148745 | 1.35390895 | 0.00012039 | 0.00048819 |
| ENSMUSG00000034780 | B3galt1 | 36.9901571 | 106.731526 | 69.7413686 | 1.35191614 | 2.5464E-09 | 2.0862E-08 |
| ENSMUSG00000053111 | Fank1 | 3.55807685 | 15.5676354 | 12.0095586 | 1.35019719 | 8.4888E-05 | 0.00035365 |
| ENSMUSG00000018925 | Heatr9 | 25.9747841 | 77.422155 | 51.4473708 | 1.34873376 | 7.6177E-08 | 5.0814E-07 |
| ENSMUSG00000036587 | Fut7 | 3.33125615 | 17.2661101 | 13.934854 | 1.34790226 | 0.00013441 | 0.00054038 |
| ENSMUSG00000038486 | Sv2a | 46.3188532 | 127.85664 | 81.5377866 | 1.34377027 | 3.346E-12 | 3.8889E-11 |
| ENSMUSG00000015305 | Sash1 | 607.334768 | 1581.89068 | 974.555912 | 1.34330081 | 5.4293E-31 | 3.06E-29 |
| ENSMUSG00000032596 | Uba7 | 789.086356 | 2101.07385 | 1311.9875 | 1.34282993 | 1.2679E-17 | 2.5903E-16 |
| ENSMUSG00000031391 | L1cam | 80.0513705 | 224.605385 | 144.554015 | 1.34141923 | 1.5663E-10 | 1.5064E-09 |
| ENSMUSG00000062609 | Kcnj15 | 5.30349053 | 22.3648042 | 17.0613137 | 1.34122169 | 7.3162E-05 | 0.00030855 |
| ENSMUSG00000030747 | Dgat2 | 954.536824 | 2508.37619 | 1553.83937 | 1.34070951 | 5.7625E-22 | 1.7156E-20 |
| ENSMUSG00000017002 | Slpi | 1547.21912 | 4515.33628 | 2968.11716 | 1.34054847 | 4.0825E-08 | 2.8412E-07 |
| ENSMUSG00000023982 | Guca1a | 7.73871899 | 31.8039139 | 24.0651949 | 1.33905012 | 7.7121E-05 | 0.0003231 |
| ENSMUSG00000030562 | Nox4 | 124.70747 | 338.925011 | 214.217541 | 1.33872729 | 2.2328E-13 | 2.9634E-12 |
| ENSMUSG00000028664 | Ephb2 | 418.994951 | 1112.79523 | 693.800278 | 1.33835061 | 1.8904E-17 | 3.7793E-16 |
| ENSMUSG00000037820 | Tgm2 | 2916.59687 | 7482.36179 | 4565.76492 | 1.33769533 | 6.619E-49 | 1.0162E-46 |
| ENSMUSG00000031548 | Sfrp1 | 329.479934 | 1176.5096 | 847.029668 | 1.33569036 | 2.8129E-05 | 0.00012718 |
| ENSMUSG00000084713 | - | 2.99713349 | 14.125949 | 11.1288155 | 1.33566522 | 0.00014521 | 0.00057926 |
| ENSMUSG00000025777 | Gdap1 | 21.3344426 | 70.1786746 | 48.844232 | 1.33485595 | 6.9316E-06 | 3.4502E-05 |
| ENSMUSG00000052726 | Kcnt2 | 64.843361 | 181.194468 | 116.351107 | 1.33374823 | 5.103E-10 | 4.5683E-09 |
| ENSMUSG00000030653 | Pde2a | 23.3454589 | 67.4085393 | 44.0630804 | 1.33357837 | 3.1279E-08 | 2.2097E-07 |
| ENSMUSG00000020108 | Ddit4 | 7110.73087 | 18347.5108 | 11236.7799 | 1.33306031 | 1.0637E-31 | 6.2702E-30 |
| ENSMUSG00000075707 | Dio3 | 48.8764855 | 137.327831 | 88.4513453 | 1.33055154 | 4.3456E-09 | 3.4503E-08 |
| ENSMUSG00000024867 | Pip5k1b | 40.9767024 | 130.946299 | 89.9695971 | 1.33038095 | 4.5162E-06 | 2.318E-05 |
| ENSMUSG00000090622 | A930033H14 | 56.9426162 | 162.881174 | 105.938557 | 1.32956864 | 1.1827E-08 | 8.893E-08 |
| ENSMUSG00000063564 | Col23a1 | 9.20737564 | 38.9972738 | 29.7898981 | 1.3294745 | 0.00010348 | 0.00042509 |
| ENSMUSG00000079450 | Cldn34c2; Cld | 17.2961592 | 53.2621683 | 35.9660091 | 1.32921931 | 1.9324E-06 | 1.0489E-05 |
| ENSMUSG00000038608 | Dock10 | 33.6300029 | 94.0593765 | 60.4293736 | 1.32758359 | 3.436E-09 | 2.7704E-08 |
| ENSMUSG00000073405 | H2-T-ps | 331.663478 | 848.600439 | 516.936961 | 1.32675304 | 1.5519E-39 | 1.4988E-37 |
| ENSMUSG00000050822 | Slc29a4 | 0.40743019 | 6.51243596 | 6.10500577 | 1.32613842 | 0.00025687 | 0.00097968 |
| ENSMUSG00000021559 | Dapk1 | 543.529141 | 1532.51535 | 988.986207 | 1.32605157 | 6.4254E-09 | 5.0031E-08 |
| ENSMUSG00000079362 | - | 418.792751 | 1135.79561 | 717.00286 | 1.32588594 | 9.2384E-12 | 1.027E-10 |
| ENSMUSG00000029664 | Tfpi2 | 11.8899319 | 39.8086853 | 27.9187534 | 1.32452655 | 1.1129E-05 | 5.3694E-05 |

| | | | | | | | |
|---|---|---|---|---|---|---|---|
| ENSMUSG00000025466 | Fuom | 63.3906463 | 169.679798 | 106.289152 | 1.32259605 | 8.9835E-13 | 1.1106E-11 |
| ENSMUSG00000022824 | Muc13 | 8.04161065 | 40.1310656 | 32.0894549 | 1.32245577 | 0.0001942 | 0.00075778 |
| ENSMUSG00000096999 | - | 0.40743019 | 9.1715946 | 8.76416441 | 1.32156609 | 0.00025302 | 0.00096643 |
| ENSMUSG00000097292 | A230107N01 | 8.67766328 | 42.6534505 | 33.9757873 | 1.32003724 | 0.00019028 | 0.00074377 |
| ENSMUSG00000070424 | Art5 | 1.12614302 | 9.03204832 | 7.9059053 | 1.31989339 | 0.00028039 | 0.00106052 |
| ENSMUSG00000040747 | Cd53 | 4.4168155 | 26.3648726 | 21.9480571 | 1.31938623 | 0.00025495 | 0.00097317 |
| ENSMUSG00000068735 | Trp53i11 | 9.40392703 | 37.7345983 | 28.3306713 | 1.3187551 | 0.00010372 | 0.00042597 |
| ENSMUSG00000016494 | Cd34 | 1924.22758 | 6298.98804 | 4374.76046 | 1.31872488 | 1.2235E-05 | 5.8671E-05 |
| ENSMUSG00000023885 | Thbs2 | 3146.04684 | 11908.3638 | 8762.31697 | 1.3180448 | 7.2639E-05 | 0.00030659 |
| ENSMUSG00000001506 | Col1a1 | 485.64842 | 1238.34016 | 752.691735 | 1.31804358 | 1.6203E-35 | 1.1947E-33 |
| ENSMUSG00000097666 | - | 0.33741727 | 6.17825089 | 5.84083362 | 1.31784449 | 0.00028093 | 0.00106218 |
| ENSMUSG00000058897 | Col25a1 | 1.46356029 | 18.5300797 | 17.0665194 | 1.31659769 | 0.00028426 | 0.00107378 |
| ENSMUSG00000015533 | Itga2 | 6.70542889 | 28.6462344 | 21.9408055 | 1.31649551 | 0.00012623 | 0.00051008 |
| ENSMUSG00000020676 | Ccl11 | 19.7948113 | 60.5399068 | 40.7450955 | 1.31638341 | 1.8925E-06 | 1.0299E-05 |
| ENSMUSG00000078607 | 1810010H24 | 32.93329 | 90.7526951 | 57.8194051 | 1.31579623 | 4.1891E-09 | 3.3325E-08 |
| ENSMUSG00000015340 | Cybb | 1.66095174 | 22.7047229 | 21.0437712 | 1.31547101 | 0.00028398 | 0.0010729 |
| ENSMUSG00000090799 | Klhl33 | 15.2861046 | 45.6292144 | 30.3431098 | 1.31500172 | 8.4302E-07 | 4.847E-06 |
| ENSMUSG00000096971 | 4930556M19 | 246.725709 | 626.808056 | 380.082348 | 1.3148775 | 1.1875E-34 | 8.3948E-33 |
| ENSMUSG00000036158 | Prickle1 | 481.738734 | 1444.94801 | 963.209273 | 1.31420312 | 1.1995E-06 | 6.7185E-06 |
| ENSMUSG00000026821 | Ralgds | 430.54331 | 1126.57462 | 696.031308 | 1.31391518 | 5.5435E-16 | 9.7138E-15 |
| ENSMUSG00000031870 | Pgr | 571.656748 | 1477.14724 | 905.490493 | 1.31288708 | 4.404E-20 | 1.1157E-18 |
| ENSMUSG00000043969 | Emx2 | 308.872308 | 778.780697 | 469.90839 | 1.31248398 | 8.0927E-43 | 9.048E-41 |
| ENSMUSG00000085781 | - | 12.5251444 | 37.6375338 | 25.1123894 | 1.31184933 | 1.2246E-06 | 6.8401E-06 |
| ENSMUSG00000028289 | Epha7 | 278.194355 | 713.244483 | 435.050128 | 1.31096502 | 3.7079E-23 | 1.2088E-21 |
| ENSMUSG00000065341 | Gm25579 | 12.7831961 | 42.5205158 | 29.7373196 | 1.31092845 | 1.6887E-05 | 7.92E-05 |
| ENSMUSG00000039496 | Cdnf | 67.8219108 | 178.190941 | 110.36903 | 1.31092086 | 3.4534E-14 | 5.0278E-13 |
| ENSMUSG00000004035 | Gstm7 | 78.025718 | 208.782886 | 130.757168 | 1.31074208 | 9.6251E-12 | 1.0674E-10 |
| ENSMUSG00000048126 | Col6a3 | 2277.49776 | 5898.17795 | 3620.68019 | 1.31005065 | 3.6742E-18 | 7.9397E-17 |
| ENSMUSG00000027071 | P2rx3 | 18.9809126 | 55.4847929 | 36.5038802 | 1.30800787 | 4.2137E-07 | 2.5339E-06 |
| ENSMUSG00000070421 | Olfr658 | 3.79934658 | 16.8082815 | 13.0089349 | 1.30745517 | 0.00016462 | 0.00065089 |
| ENSMUSG00000028435 | Aqp3 | 21.9924951 | 63.4314924 | 41.4389973 | 1.30738288 | 1.6825E-07 | 1.0726E-06 |
| ENSMUSG00000050050 | Ccdc158 | 13.1248697 | 61.7037537 | 48.578884 | 1.30700607 | 0.00021616 | 0.00083793 |
| ENSMUSG00000021876 | Rnase4 | 1199.52901 | 3044.89829 | 1845.36929 | 1.30543253 | 8.6646E-27 | 3.6601E-25 |
| ENSMUSG00000031722 | Hp | 2162.45546 | 10155.5737 | 7993.11827 | 1.30143104 | 0.00023591 | 0.00090798 |
| ENSMUSG00000056755 | Grm7 | 2.43193383 | 12.6612233 | 10.2292895 | 1.30110782 | 0.00025991 | 0.0009896 |
| ENSMUSG00000030849 | Fgfr2 | 341.006314 | 1000.59706 | 659.590746 | 1.30108905 | 9.3232E-07 | 5.3144E-06 |
| ENSMUSG00000003273 | Car11 | 32.1048207 | 93.8043171 | 61.6994964 | 1.30008926 | 7.8514E-07 | 4.5396E-06 |
| ENSMUSG00000079481 | Nhsl2 | 4.29123871 | 18.720091 | 14.4288523 | 1.2997645 | 0.0001688 | 0.00066663 |
| ENSMUSG00000027931 | Npr1 | 27.1499018 | 77.0553933 | 49.9054915 | 1.29971477 | 8.3856E-08 | 5.5612E-07 |
| ENSMUSG00000053263 | Gm12592 | 8.91989469 | 31.3394429 | 22.4195482 | 1.29824654 | 4.2328E-05 | 0.00018574 |
| ENSMUSG00000009097 | Tbx1 | 20.9065053 | 77.9490905 | 57.0425852 | 1.29749106 | 9.7603E-05 | 0.00040263 |
| ENSMUSG00000026602 | Nphs2 | 20.052023 | 61.6168493 | 41.5648263 | 1.29686818 | 7.2743E-06 | 3.6112E-05 |
| ENSMUSG00000091089 | - | 29.7606434 | 90.3501362 | 60.5894928 | 1.29503901 | 4.5975E-06 | 2.3533E-05 |
| ENSMUSG00000031189 | Aff2 | 9.32456145 | 33.83732 | 24.5127585 | 1.29487378 | 8.7427E-05 | 0.00036342 |
| ENSMUSG00000022512 | Cldn1 | 441.838613 | 1130.05449 | 688.215872 | 1.29467305 | 2.0467E-18 | 4.5128E-17 |
| ENSMUSG00000037166 | Ppp1r14a | 1.65256076 | 11.0656913 | 9.41313053 | 1.29387375 | 0.00034803 | 0.00129143 |
| ENSMUSG00000031808 | Slc27a1 | 906.326459 | 2244.55389 | 1338.22743 | 1.29319088 | 5.8456E-64 | 1.8501E-61 |
| ENSMUSG00000028539 | Artn | 1059.91872 | 2629.86232 | 1569.9436 | 1.29305819 | 2.0816E-54 | 4.2399E-52 |
| ENSMUSG00000010051 | Hyal1 | 36.5142069 | 104.994847 | 68.4806404 | 1.29162908 | 6.8177E-07 | 3.9676E-06 |
| ENSMUSG00000029797 | Sspo | 6.03994701 | 27.3785304 | 21.3385834 | 1.29111082 | 0.00023156 | 0.00089255 |
| ENSMUSG00000074465 | - | 2.20511313 | 12.7516969 | 10.5465838 | 1.29104229 | 0.00031907 | 0.00119259 |
| ENSMUSG00000002083 | Bbc3 | 329.534901 | 828.217317 | 498.682416 | 1.29085779 | 8.4632E-25 | 3.076E-23 |
| ENSMUSG00000006395 | - | 331.7251 | 835.779119 | 504.054019 | 1.29037082 | 1.2221E-24 | 4.4106E-23 |
| ENSMUSG00000034156 | Tspoap1 | 101.975037 | 259.684106 | 157.709069 | 1.28913475 | 1.8354E-17 | 3.6766E-16 |
| ENSMUSG00000027737 | Slc7a11 | 5526.56315 | 15143.4667 | 9616.90357 | 1.28870064 | 1.8871E-08 | 1.3781E-07 |
| ENSMUSG00000074896 | Ifit3 | 16124.7785 | 40505.8893 | 24381.1108 | 1.28829605 | 6.1069E-25 | 2.2474E-23 |
| ENSMUSG00000041313 | Slc7a1 | 7330.21122 | 18335.4866 | 11005.2753 | 1.28763908 | 1.8568E-28 | 8.7811E-27 |
| ENSMUSG00000019982 | Myb | 66.465222 | 172.363439 | 105.898217 | 1.28737764 | 2.397E-12 | 2.8438E-11 |
| ENSMUSG00000085419 | - | 1.0561301 | 9.40848759 | 8.35235748 | 1.28582974 | 0.000407 | 0.00148877 |
| ENSMUSG00000029661 | Col1a2 | 5994.80237 | 15746.8974 | 9752.09507 | 1.28533866 | 3.002E-11 | 3.1412E-10 |
| ENSMUSG00000074277 | Phldb3 | 39.2504036 | 115.825901 | 76.5754978 | 1.28500862 | 3.1033E-06 | 1.6349E-05 |
| ENSMUSG00000038793 | Lefty1 | 83.0048481 | 218.876935 | 135.872087 | 1.28436111 | 1.3992E-10 | 1.3578E-09 |
| ENSMUSG00000032579 | Hemk1 | 363.629356 | 914.451141 | 550.821785 | 1.28427639 | 2.2364E-22 | 6.8668E-21 |
| ENSMUSG00000040605 | Bace2 | 2.18227312 | 15.0867736 | 12.9045005 | 1.28406359 | 0.00040584 | 0.00148586 |
| ENSMUSG00000027068 | Dhrs9 | 2002.25471 | 5026.76235 | 3024.50763 | 1.28363581 | 5.8258E-23 | 1.8639E-21 |
| ENSMUSG00000039699 | Batf2 | 2.0422473 | 12.4554166 | 10.4131693 | 1.28349793 | 0.00037992 | 0.00139714 |
| ENSMUSG00000001552 | Jup | 266.419463 | 699.12813 | 432.708667 | 1.28290997 | 3.2427E-11 | 3.3776E-10 |
| ENSMUSG00000038884 | Shfl | 428.47505 | 1060.79449 | 632.319437 | 1.28199804 | 2.8721E-37 | 2.3446E-35 |
| ENSMUSG00000028518 | Prkaa2 | 130.62927 | 346.777834 | 216.148565 | 1.28084562 | 3.5865E-10 | 3.2869E-09 |
| ENSMUSG00000048652 | Samd13 | 1.8271122 | 13.1523156 | 11.3252034 | 1.27693159 | 0.00044177 | 0.00160681 |
| ENSMUSG00000025083 | Afap1l2 | 77.5304096 | 203.051947 | 125.521538 | 1.27681873 | 4.4562E-11 | 4.5383E-10 |
| ENSMUSG00000069893 | 9930111J21F | 34.9216535 | 102.715927 | 67.7942737 | 1.27594365 | 2.8047E-06 | 1.4878E-05 |
| ENSMUSG00000029417 | Cxcl9 | 65.3400406 | 271.422167 | 206.082127 | 1.27465993 | 0.00024778 | 0.00094905 |
| ENSMUSG00000015222 | Map2 | 455.711115 | 1179.64085 | 723.929738 | 1.27354336 | 8.5423E-12 | 9.5403E-11 |
| ENSMUSG00000046058 | Eid2 | 151.943549 | 380.281361 | 228.337812 | 1.27267805 | 4.4279E-19 | 1.0375E-17 |

| | | | | | | | |
|---|---|---|---|---|---|---|---|
| ENSMUSG00000029228 | Lnx1 | 67.0710053 | 170.955785 | 103.88478 | 1.27178215 | 4.4905E-13 | 5.7593E-12 |
| ENSMUSG00000054200 | Ffar4 | 13.9607684 | 42.8104624 | 28.849694 | 1.27175971 | 2.2183E-05 | 0.00010216 |
| ENSMUSG00000074676 | Foxs1 | 10.4996791 | 32.1396849 | 21.6400058 | 1.27155681 | 1.4426E-05 | 6.8475E-05 |
| ENSMUSG00000055963 | Triqk | 379.395702 | 938.595837 | 559.200135 | 1.27088136 | 3.1717E-25 | 1.2016E-23 |
| ENSMUSG00000042857 | Gm9776 | 126.70426 | 314.294658 | 187.590398 | 1.26958061 | 4.4566E-21 | 1.2373E-19 |
| ENSMUSG00000060176 | Kif27 | 12.2577401 | 41.065959 | 28.8082189 | 1.26878328 | 7.0377E-05 | 0.00029818 |
| ENSMUSG00000085794 | Vax2os | 3.35739078 | 13.7526266 | 10.3952358 | 1.26855786 | 0.00021162 | 0.00082187 |
| ENSMUSG00000020363 | Gfpt2 | 76.507125 | 190.758341 | 114.251216 | 1.26796428 | 1.4127E-17 | 2.8717E-16 |
| ENSMUSG00000040488 | Ltbp4 | 739.792577 | 1857.49486 | 1117.70229 | 1.2665127 | 8.6242E-17 | 1.6412E-15 |
| ENSMUSG00000001435 | Col18a1 | 561.539791 | 1376.88366 | 815.343874 | 1.26626982 | 3.3579E-34 | 2.2723E-32 |
| ENSMUSG00000020256 | Aldh1l2 | 7250.526 | 18008.1553 | 10757.6292 | 1.26471261 | 1.184E-20 | 3.1509E-19 |
| ENSMUSG00000035778 | Ggta1 | 1164.90205 | 2924.16398 | 1759.26193 | 1.26455852 | 2.6892E-16 | 4.8571E-15 |
| ENSMUSG00000075044 | Slc22a29 | 29.2131874 | 78.0853165 | 48.8721292 | 1.26404855 | 2.3447E-08 | 1.6919E-07 |
| ENSMUSG00000025584 | Pde8a | 1028.52855 | 2517.05743 | 1488.52889 | 1.26385405 | 3.6906E-34 | 2.4892E-32 |
| ENSMUSG00000022150 | Dab2 | 1650.57488 | 4035.97096 | 2385.39608 | 1.26327866 | 1.666E-34 | 1.154E-32 |
| ENSMUSG00000053846 | Lipg | 11.0690135 | 54.5190046 | 43.4499912 | 1.26184672 | 0.00042186 | 0.00153984 |
| ENSMUSG00000097493 | - | 24.7894738 | 65.6296173 | 40.8401435 | 1.2611293 | 5.9919E-08 | 4.0588E-07 |
| ENSMUSG00000092075 | - | 3.70319904 | 17.9200614 | 14.2168624 | 1.26015234 | 0.00039876 | 0.00146175 |
| ENSMUSG00000085071 | - | 51.2858881 | 131.487795 | 80.2019064 | 1.25839851 | 3.0687E-12 | 3.5931E-11 |
| ENSMUSG00000005583 | Mef2c | 369.457959 | 912.186103 | 542.728144 | 1.25827433 | 3.9682E-22 | 1.1917E-20 |
| ENSMUSG00000040794 | C1qtnf4 | 1.02999547 | 8.79798807 | 7.76799259 | 1.25786263 | 0.00054298 | 0.00193592 |
| ENSMUSG00000036298 | Slc2a13 | 3.46192931 | 14.1457594 | 10.6838301 | 1.25696714 | 0.00027293 | 0.00103441 |
| ENSMUSG00000028017 | Egf | 113.613506 | 278.71165 | 165.098144 | 1.25591193 | 1.2324E-22 | 3.8647E-21 |
| ENSMUSG00000022103 | Gfra2 | 2048.36179 | 4935.64625 | 2887.28446 | 1.25571342 | 8.224E-70 | 3.3837E-67 |
| ENSMUSG00000004267 | Eno2 | 209.909471 | 545.182498 | 335.273027 | 1.25408895 | 9.6051E-10 | 8.2641E-09 |
| ENSMUSG00000052914 | Cyp2j6 | 68.9723867 | 184.260185 | 115.287799 | 1.25373053 | 7.7548E-08 | 5.1662E-07 |
| ENSMUSG00000095457 | - | 6.21959481 | 32.5067322 | 26.2871374 | 1.2525769 | 0.00049523 | 0.00178391 |
| ENSMUSG00000020312 | Shc2 | 109.731191 | 283.650119 | 173.918929 | 1.25125097 | 1.438E-09 | 1.2149E-08 |
| ENSMUSG00000032252 | Glce | 2112.85478 | 5099.77852 | 2986.92375 | 1.25097412 | 5.0874E-44 | 5.8797E-42 |
| ENSMUSG00000039013 | Siglecf | 4.20677677 | 20.5345203 | 16.3277436 | 1.25071564 | 0.0004693 | 0.0016968 |
| ENSMUSG00000037664 | Cdkn1c | 10.4339225 | 37.0256083 | 26.5916858 | 1.25013255 | 0.00018124 | 0.00071181 |
| ENSMUSG00000088217 | Gm22079 | 1.32353447 | 8.72306949 | 7.39953503 | 1.24708736 | 0.00057998 | 0.00205747 |
| ENSMUSG00000046709 | Mapk10 | 35.4388402 | 94.8517366 | 59.4128964 | 1.24707072 | 6.5531E-08 | 4.4113E-07 |
| ENSMUSG00000017195 | Zpbp2 | 3.29067249 | 13.2204355 | 9.92976306 | 1.24592786 | 0.00034597 | 0.00128471 |
| ENSMUSG00000067212 | H2-T23 | 769.169433 | 1853.55077 | 1084.38134 | 1.24550816 | 1.0488E-33 | 6.9153E-32 |
| ENSMUSG00000097125 | - | 11.1348917 | 34.2957965 | 23.1609049 | 1.24419142 | 5.121E-05 | 0.00022193 |
| ENSMUSG00000042099 | Kank3 | 118.992946 | 331.547189 | 212.554242 | 1.24372991 | 2.217E-06 | 1.1908E-05 |
| ENSMUSG00000016028 | Celsr1 | 104.522011 | 390.76164 | 286.239629 | 1.24363321 | 0.00026419 | 0.00100459 |
| ENSMUSG00000044629 | Cnrip1 | 3.53194222 | 17.3674627 | 13.8355204 | 1.2434085 | 0.00051842 | 0.00185769 |
| ENSMUSG00000097069 | Gm16998 | 5.00059887 | 18.243586 | 13.2429871 | 1.24149911 | 0.00018787 | 0.00073517 |
| ENSMUSG00000025946 | Pth2r | 22.564284 | 66.7221609 | 44.1578769 | 1.24070413 | 2.4065E-05 | 0.00011006 |
| ENSMUSG00000024481 | Lvrn | 207.241364 | 508.768506 | 301.527142 | 1.23944821 | 3.6546E-18 | 7.9056E-17 |
| ENSMUSG00000042010 | Acacb | 232.402128 | 580.091401 | 347.689273 | 1.23917049 | 3.6804E-13 | 4.7678E-12 |
| ENSMUSG00000025017 | Pik3ap1 | 5.06551542 | 21.4514727 | 16.3859573 | 1.23820706 | 0.00044089 | 0.00160417 |
| ENSMUSG00000025586 | Cpeb1 | 68.7184697 | 272.358304 | 203.639835 | 1.23795025 | 0.00036805 | 0.00135788 |
| ENSMUSG00000044231 | Nhlrc1 | 37.2869907 | 102.185426 | 64.8984355 | 1.23791726 | 1.2026E-06 | 6.7321E-06 |
| ENSMUSG00000029544 | Cabp1 | 4.04061631 | 15.2056442 | 11.1650279 | 1.23748427 | 0.00024429 | 0.00093707 |
| ENSMUSG00000064329 | Scn1a | 5.15507372 | 19.7293444 | 14.5742707 | 1.23731881 | 0.00024949 | 0.00095489 |
| ENSMUSG00000022237 | Ankrd33b | 3.40966004 | 19.6330726 | 16.2234126 | 1.23698806 | 0.0006331 | 0.00222406 |
| ENSMUSG00000037341 | Slc9a7 | 216.917174 | 552.120511 | 335.203338 | 1.23662435 | 5.6604E-10 | 5.0312E-09 |
| ENSMUSG00000010803 | Gabra1 | 9.0799971 | 29.4863248 | 20.4063277 | 1.23537494 | 0.00010179 | 0.00041872 |
| ENSMUSG00000052560 | Cpne8 | 432.906436 | 1041.13134 | 608.224902 | 1.23464123 | 3.6675E-28 | 1.6993E-26 |
| ENSMUSG00000079018 | Ly6c1 | 5249.51709 | 12588.973 | 7339.4559 | 1.2342617 | 5.3228E-31 | 3.0083E-29 |
| ENSMUSG00000035459 | Stab2 | 82.7503999 | 208.119755 | 125.369355 | 1.2333644 | 3.111E-11 | 3.2504E-10 |
| ENSMUSG00000069518 | - | 1.7487083 | 9.38115277 | 7.63244447 | 1.23219366 | 0.00062515 | 0.00220028 |
| ENSMUSG00000021451 | Sema4d | 13.3838831 | 42.3204421 | 28.936559 | 1.23177393 | 9.1414E-05 | 0.00037907 |
| ENSMUSG00000042873 | Lhfpl4 | 131.013673 | 316.45196 | 185.438287 | 1.23175921 | 1.3635E-21 | 3.9121E-20 |
| ENSMUSG00000030022 | Adamts9 | 96.516297 | 261.754505 | 165.238208 | 1.23167054 | 9.9312E-07 | 5.6329E-06 |
| ENSMUSG00000050505 | Pcdh20 | 3.26453786 | 13.56846 | 10.3039221 | 1.23166486 | 0.00045233 | 0.00164232 |
| ENSMUSG00000051817 | Sox12 | 453.68407 | 1081.05633 | 627.372255 | 1.23119365 | 1.0132E-40 | 1.0422E-38 |
| ENSMUSG00000073485 | - | 27.1867604 | 76.1058176 | 48.9190572 | 1.23088316 | 5.2232E-06 | 2.6569E-05 |
| ENSMUSG00000029206 | Nsun7 | 21.0441982 | 59.8448352 | 38.800637 | 1.22609806 | 1.347E-05 | 6.4218E-05 |
| ENSMUSG00000035566 | Pcdh17 | 29.24538 | 78.5750808 | 49.3297008 | 1.22580741 | 8.763E-07 | 5.0299E-06 |
| ENSMUSG00000020950 | Foxg1 | 1182.01999 | 2816.4514 | 1634.43141 | 1.2253233 | 2.2954E-30 | 1.2362E-28 |
| ENSMUSG00000010529 | Gm266 | 4.78036741 | 20.7967737 | 16.0164063 | 1.22484494 | 0.0005589 | 0.00198717 |
| ENSMUSG00000020142 | Slc1a4 | 4949.92666 | 11651.3268 | 6701.40009 | 1.22299133 | 1.4221E-67 | 5.4178E-65 |
| ENSMUSG00000039934 | Gsap | 497.015056 | 1198.26143 | 701.246377 | 1.22297748 | 7.0008E-19 | 1.6146E-17 |
| ENSMUSG00000087153 | - | 235.33733 | 559.976223 | 324.638893 | 1.22148705 | 1.9559E-34 | 1.3412E-32 |
| ENSMUSG00000026768 | Itga8 | 3.88200678 | 21.0355314 | 17.1535246 | 1.22102066 | 0.00074752 | 0.00259195 |
| ENSMUSG00000017737 | Mmp9 | 221.63185 | 530.946923 | 309.315073 | 1.22023654 | 6.1406E-24 | 2.088E-22 |
| ENSMUSG00000046442 | Ppm1e | 38.0681655 | 101.19786 | 63.1296946 | 1.21921838 | 4.7027E-07 | 2.8066E-06 |
| ENSMUSG00000026890 | Lhx6 | 96.2385126 | 239.533931 | 143.295419 | 1.21838552 | 3.0696E-10 | 2.8432E-09 |
| ENSMUSG00000083763 | - | 0.33741727 | 6.40653234 | 6.06911506 | 1.2183124 | 0.00072645 | 0.00252315 |
| ENSMUSG00000028782 | Adgrb2 | 6.92385861 | 34.0851408 | 27.1612822 | 1.21757551 | 0.00071146 | 0.00247524 |
| ENSMUSG00000085873 | - | 9.06650976 | 29.0265241 | 19.9600144 | 1.21732884 | 0.00016927 | 0.00066814 |

| | | | | | | | |
|---|---|---|---|---|---|---|---|
| ENSMUSG00000070866 | Zfp804a | 6.33678062 | 24.3135565 | 17.9767759 | 1.21628205 | 0.00044209 | 0.0016074 |
| ENSMUSG00000029322 | Plac8 | 40.042424 | 108.034003 | 67.991579 | 1.21575437 | 3.1177E-06 | 1.642E-05 |
| ENSMUSG00000043487 | Acot6 | 40.2785973 | 108.300974 | 68.022377 | 1.21413794 | 3.3632E-06 | 1.7623E-05 |
| ENSMUSG00000027217 | Tspan18 | 11.3863541 | 38.9719455 | 27.5855914 | 1.21411212 | 0.00023814 | 0.00091501 |
| ENSMUSG00000025993 | Slc40a1 | 2.71198548 | 13.8561954 | 11.1442099 | 1.21349878 | 0.00078411 | 0.0027083 |
| ENSMUSG00000017652 | Cd40 | 36.8320787 | 106.297111 | 69.465032 | 1.21334486 | 3.3507E-05 | 0.00014962 |
| ENSMUSG00000031119 | Gpc4 | 2262.28057 | 5676.25898 | 3413.97841 | 1.21255426 | 2.4027E-09 | 1.9755E-08 |
| ENSMUSG00000041794 | Myrip | 10.5174228 | 36.7817885 | 26.2643657 | 1.21145807 | 0.0003168 | 0.0011858 |
| ENSMUSG00000084845 | Tmem240 | 14.357353 | 40.0352674 | 25.6779144 | 1.21120898 | 9.0726E-06 | 4.4407E-05 |
| ENSMUSG00000028654 | Mycl | 24.1636139 | 67.5948533 | 43.4312395 | 1.20979811 | 1.8716E-05 | 8.7108E-05 |
| ENSMUSG00000086610 | Gm15408 | 84.5048574 | 203.974843 | 119.469986 | 1.20940191 | 4.8908E-14 | 6.9918E-13 |
| ENSMUSG00000021903 | Galnt15 | 7.48905827 | 30.7299304 | 23.2408722 | 1.20784609 | 0.00061398 | 0.0021654 |
| ENSMUSG00000059173 | Pde1a | 255.017223 | 613.926482 | 358.909259 | 1.20695066 | 8.188E-15 | 1.2684E-13 |
| ENSMUSG00000002791 | Pdxk-ps | 1.32353447 | 8.66491086 | 7.3413764 | 1.20660887 | 0.00088999 | 0.00303934 |
| ENSMUSG00000040003 | Magi2 | 42.6610253 | 104.673817 | 62.0127916 | 1.20615411 | 4.0776E-10 | 3.7101E-09 |
| ENSMUSG00000023043 | Krt18 | 92.3210186 | 225.654043 | 133.333024 | 1.2060229 | 2.6045E-11 | 2.7463E-10 |
| ENSMUSG00000074682 | Zcchc3 | 19.5517398 | 53.7170997 | 34.1653599 | 1.20469308 | 1.5162E-05 | 7.1621E-05 |
| ENSMUSG00000021803 | Cdhr1 | 2.54582503 | 13.3158479 | 10.7700229 | 1.20443655 | 0.00086306 | 0.00295294 |
| ENSMUSG00000032198 | Dock6 | 340.073428 | 799.410519 | 459.337091 | 1.20259285 | 2.7817E-26 | 1.1243E-24 |
| ENSMUSG00000024593 | Megf10 | 310.433372 | 902.577099 | 592.143727 | 1.20220949 | 5.2064E-05 | 0.0002252 |
| ENSMUSG00000000732 | Icosl | 20.5928897 | 55.5840607 | 34.991171 | 1.20091294 | 6.902E-06 | 3.4371E-05 |
| ENSMUSG00000041477 | Dcp1b | 145.320659 | 342.730211 | 197.409553 | 1.20029131 | 4.0153E-20 | 1.0223E-18 |
| ENSMUSG00000029314 | Gpat3 | 884.463675 | 2106.46397 | 1222.0003 | 1.19980173 | 1.0279E-16 | 1.9435E-15 |
| ENSMUSG00000041134 | Cyyr1 | 3.04610814 | 13.1984457 | 10.1523375 | 1.19959203 | 0.00070506 | 0.00245588 |
| ENSMUSG00000032766 | Gng11 | 441.77317 | 1068.18924 | 626.416071 | 1.19937095 | 3.2807E-12 | 3.8195E-11 |
| ENSMUSG00000044921 | Rassf9 | 233.550837 | 560.904305 | 327.353469 | 1.19909562 | 1.8498E-13 | 2.4727E-12 |
| ENSMUSG00000074039 | - | 11.5296746 | 33.0855262 | 21.5558517 | 1.19907723 | 4.1793E-05 | 0.00018356 |
| ENSMUSG00000085984 | 1700001G11 | 38.1471006 | 96.7625898 | 58.6154891 | 1.19869715 | 8.7484E-08 | 5.7883E-07 |
| ENSMUSG00000010476 | Ebf3 | 13.7035567 | 48.8582437 | 35.154687 | 1.19849887 | 0.00043651 | 0.00158991 |
| ENSMUSG00000017417 | Plxdc1 | 20.628377 | 53.7376676 | 33.1092905 | 1.19842164 | 1.2147E-06 | 6.7923E-06 |
| ENSMUSG00000021990 | Spata13 | 329.34789 | 788.862392 | 459.514502 | 1.19758864 | 3.1626E-14 | 4.6208E-13 |
| ENSMUSG00000035878 | Hykk | 91.6240833 | 228.820485 | 137.196401 | 1.19613237 | 9.3188E-09 | 7.1108E-08 |
| ENSMUSG00000032118 | Fez1 | 47.9543231 | 135.021683 | 87.0673595 | 1.19539723 | 3.4191E-05 | 0.00015243 |
| ENSMUSG00000038523 | 1700003F12 | 13.479069 | 39.1574662 | 25.6783973 | 1.19478982 | 7.0972E-05 | 0.0003003 |
| ENSMUSG00000031758 | Cdyl2 | 107.550927 | 264.77081 | 157.219883 | 1.19306616 | 6.4315E-10 | 5.6712E-09 |
| ENSMUSG00000006784 | Odad4 | 22.3350088 | 60.0241456 | 37.6891368 | 1.1930526 | 4.0536E-06 | 2.0953E-05 |
| ENSMUSG00000043557 | Mdga1 | 459.548347 | 1113.81256 | 654.264212 | 1.19263166 | 2.5351E-11 | 2.6773E-10 |
| ENSMUSG00000042246 | Tmc7 | 29.9243493 | 155.62667 | 125.702321 | 1.19253938 | 0.00099029 | 0.00335122 |
| ENSMUSG00000029084 | Cd38 | 4.75093816 | 20.4676407 | 15.7167025 | 1.19248358 | 0.00078484 | 0.00270993 |
| ENSMUSG00000014361 | Mertk | 11.7541623 | 34.9769242 | 23.2227619 | 1.19080009 | 0.00010461 | 0.0004293 |
| ENSMUSG00000030671 | Pde3b | 118.723087 | 297.109014 | 178.385927 | 1.19064051 | 3.1784E-08 | 2.2438E-07 |
| ENSMUSG00000095649 | - | 22.2751886 | 71.6028913 | 49.3277027 | 1.19030362 | 0.00023801 | 0.0009147 |
| ENSMUSG00000025140 | Pycr1 | 1843.51739 | 4262.29164 | 2418.77425 | 1.189876 | 1.3778E-38 | 1.2432E-36 |
| ENSMUSG00000003282 | Plag1 | 11.4657197 | 35.0815173 | 23.6157976 | 1.18887417 | 0.00014484 | 0.00057823 |
| ENSMUSG00000040280 | Ndufa4l2 | 2.16452947 | 10.9822466 | 8.81771715 | 1.18837982 | 0.00099767 | 0.003374 |
| ENSMUSG00000048402 | Gli2 | 46.4952064 | 138.887699 | 92.3924923 | 1.18837676 | 0.0001177 | 0.00047823 |
| ENSMUSG00000078349 | AW011738 | 330.623411 | 787.599991 | 456.976579 | 1.18792694 | 1.102E-13 | 1.5165E-12 |
| ENSMUSG00000035121 | Neil2 | 129.01254 | 302.215044 | 173.202504 | 1.18758254 | 2.7011E-18 | 5.9177E-17 |
| ENSMUSG00000035864 | Syt1 | 4.92219498 | 18.7522482 | 13.8300532 | 1.18753126 | 0.00058334 | 0.00206692 |
| ENSMUSG00000058420 | Syt17 | 57.7561054 | 158.827691 | 101.071586 | 1.18532944 | 2.2324E-05 | 0.00010269 |
| ENSMUSG00000020875 | Hoxb9 | 1004.14637 | 2395.07799 | 1390.93162 | 1.18529569 | 7.2295E-13 | 9.0246E-12 |
| ENSMUSG00000046215 | Rprml | 5.92935044 | 19.0466944 | 13.1173439 | 1.18500654 | 0.00029145 | 0.00109911 |
| ENSMUSG00000074052 | - | 11.1577317 | 34.3038876 | 23.146156 | 1.18455665 | 0.00019483 | 0.00075997 |
| ENSMUSG00000028024 | Enpep | 102.976974 | 328.738117 | 225.761143 | 1.18388883 | 0.00028683 | 0.0010831 |
| ENSMUSG00000022037 | Clu | 856.355132 | 2209.80214 | 1353.44701 | 1.18333456 | 1.5725E-06 | 8.6691E-06 |
| ENSMUSG00000074398 | Gm15441 | 13.6825185 | 39.0346089 | 25.3520904 | 1.18248538 | 5.1946E-05 | 0.00022476 |
| ENSMUSG00000055003 | Lrtm2 | 17.5780126 | 49.2972601 | 31.7192475 | 1.18122349 | 4.4957E-05 | 0.00019649 |
| ENSMUSG00000029217 | Tec | 142.097666 | 350.153604 | 208.055937 | 1.18036463 | 2.1016E-08 | 1.5234E-07 |
| ENSMUSG00000021069 | Pygl | 76.7314911 | 207.17873 | 130.447239 | 1.17939935 | 1.7451E-05 | 8.1608E-05 |
| ENSMUSG00000022425 | Enpp2 | 78733.7888 | 179935.024 | 101201.235 | 1.17868988 | 3.5386E-52 | 6.4421E-50 |
| ENSMUSG00000035914 | Cd276 | 128.135096 | 300.832791 | 172.697695 | 1.17790858 | 5.6138E-15 | 8.8496E-14 |
| ENSMUSG00000051022 | Hs3st1 | 233.483157 | 639.727259 | 406.244102 | 1.17699481 | 3.1509E-05 | 0.00014128 |
| ENSMUSG00000027684 | Mecom | 101.788678 | 240.553066 | 138.764388 | 1.17671721 | 3.0269E-14 | 4.432E-13 |
| ENSMUSG00000054072 | Iigp1 | 2138.35477 | 5914.56964 | 3776.21487 | 1.17643828 | 4.1301E-05 | 0.00018155 |
| ENSMUSG00000026073 | Il1r2 | 24.9201469 | 62.4723682 | 37.5522213 | 1.17578559 | 1.376E-06 | 7.6404E-06 |
| ENSMUSG00000029490 | Mfsd7a | 30.2114206 | 85.0524559 | 54.8410352 | 1.1745321 | 5.9055E-05 | 0.00025347 |
| ENSMUSG00000036908 | Unc93b1 | 748.828401 | 1749.03849 | 1000.21008 | 1.17427107 | 3.0489E-16 | 5.4875E-15 |
| ENSMUSG00000097558 | - | 148.620153 | 343.222571 | 194.602419 | 1.17380139 | 2.2815E-22 | 6.9947E-21 |
| ENSMUSG00000020866 | Cacna1g | 128.694668 | 303.041363 | 174.346695 | 1.17338592 | 2.5038E-13 | 3.2976E-12 |
| ENSMUSG00000026616 | Cr2 | 3.45353832 | 13.3886197 | 9.93508139 | 1.1733591 | 0.00080574 | 0.00277743 |
| ENSMUSG00000067613 | Krt83 | 0.74484746 | 7.9387802 | 7.19393274 | 1.17290318 | 0.00118686 | 0.00395146 |
| ENSMUSG00000073490 | Ifi207 | 560.825436 | 1288.89377 | 728.068331 | 1.17192566 | 1.2659E-25 | 4.9137E-24 |
| ENSMUSG00000007021 | Syngr3 | 54.3479382 | 143.76711 | 89.4191718 | 1.17146092 | 1.136E-05 | 5.4743E-05 |
| ENSMUSG00000097002 | - | 0.67483455 | 6.57343123 | 5.89859668 | 1.17083097 | 0.00125314 | 0.00415263 |
| ENSMUSG00000056476 | Med12l | 158.705813 | 378.360752 | 219.654939 | 1.17051136 | 7.2928E-11 | 7.2547E-10 |

| | | | | | | | |
|---|---|---|---|---|---|---|---|
| ENSMUSG00000048806 | Ifnb1 | 19.8023622 | 68.6486444 | 48.8462821 | 1.16933949 | 0.00062504 | 0.00220025 |
| ENSMUSG00000047238 | Mageh1 | 438.860033 | 1007.84382 | 568.98379 | 1.16918302 | 5.2719E-23 | 1.692E-21 |
| ENSMUSG00000040387 | Klhl32 | 31.3711276 | 83.7612596 | 52.390132 | 1.16916567 | 1.4942E-05 | 7.0681E-05 |
| ENSMUSG00000031364 | Grpr | 3.86096851 | 14.0021573 | 10.1411888 | 1.1684301 | 0.00071041 | 0.00247203 |
| ENSMUSG00000006930 | Hap1 | 158.697422 | 363.355935 | 204.658514 | 1.16689621 | 1.7177E-27 | 7.5831E-26 |
| ENSMUSG00000034634 | Ly6d | 73.4820551 | 174.266997 | 100.784942 | 1.16686283 | 1.5198E-10 | 1.4644E-09 |
| ENSMUSG00000085936 | - | 83.7077406 | 198.908609 | 115.200869 | 1.16674526 | 1.6712E-10 | 1.6028E-09 |
| ENSMUSG00000025500 | Lmntd2 | 45.4433326 | 111.529505 | 66.0861721 | 1.16672238 | 2.4046E-08 | 1.7315E-07 |
| ENSMUSG00000042726 | Trafd1 | 2536.22161 | 5733.13785 | 3196.91623 | 1.16652479 | 2.6513E-64 | 8.6574E-62 |
| ENSMUSG00000087179 | - | 14.0133465 | 46.13809 | 32.1247435 | 1.16636454 | 0.00044192 | 0.00160707 |
| ENSMUSG00000040270 | Bach2 | 21.3260516 | 54.9990363 | 33.6729847 | 1.165648 | 3.8461E-06 | 1.995E-05 |
| ENSMUSG00000043993 | - | 67.7890653 | 163.998314 | 96.2092488 | 1.16502433 | 3.5223E-09 | 2.8349E-08 |
| ENSMUSG00000045725 | Prr15 | 0.74484746 | 7.27172292 | 6.52687546 | 1.16478657 | 0.00130208 | 0.00429613 |
| ENSMUSG00000097471 | 5830432E09 | 7.48576365 | 26.999316 | 19.5135523 | 1.16407036 | 0.00074527 | 0.00258456 |
| ENSMUSG00000074093 | Svip | 50.4626368 | 126.408243 | 75.9456064 | 1.16396877 | 4.9244E-07 | 2.9287E-06 |
| ENSMUSG00000047992 | Dipk1c | 8.78136174 | 26.9230065 | 18.1416448 | 1.16372942 | 0.00034624 | 0.00128549 |
| ENSMUSG00000091157 | - | 1.3496691 | 8.55003564 | 7.20036654 | 1.1633396 | 0.00137311 | 0.00450808 |
| ENSMUSG00000046961 | Gpr156 | 5.45190734 | 18.5208855 | 13.0689782 | 1.16305668 | 0.00056497 | 0.00200803 |
| ENSMUSG00000042659 | Arrdc4 | 970.633245 | 2195.19822 | 1224.56497 | 1.16273827 | 1.9027E-45 | 2.4013E-43 |
| ENSMUSG00000031425 | Plp1 | 41.589915 | 102.353871 | 60.7639564 | 1.16263594 | 1.0995E-07 | 7.1578E-07 |
| ENSMUSG00000078307 | AI593442 | 1.0561301 | 9.90549718 | 8.84936708 | 1.162626 | 0.00133376 | 0.0043887 |
| ENSMUSG00000036030 | Prtg | 44.5323247 | 108.643835 | 64.1115101 | 1.16237109 | 1.4661E-08 | 1.0857E-07 |
| ENSMUSG00000037447 | Arid5a | 1123.24373 | 2580.51138 | 1457.26765 | 1.16129405 | 1.3213E-19 | 3.2283E-18 |
| ENSMUSG00000032036 | Kirrel3 | 33.1366179 | 81.7802195 | 48.6436016 | 1.16121615 | 1.0187E-07 | 6.668E-07 |
| ENSMUSG00000029636 | Wasf3 | 66.4431213 | 183.85487 | 117.411749 | 1.1611097 | 6.7871E-05 | 0.00028842 |
| ENSMUSG00000090942 | F830016B08 | 71.3003997 | 200.092777 | 128.792377 | 1.16059106 | 9.2095E-05 | 0.00038167 |
| ENSMUSG00000027346 | Gpcpd1 | 1867.09946 | 4232.05955 | 2364.9601 | 1.16048909 | 2.0657E-35 | 1.5123E-33 |
| ENSMUSG00000038393 | Txnip | 9162.93282 | 20996.6252 | 11833.6924 | 1.16014096 | 1.0159E-20 | 2.7318E-19 |
| ENSMUSG00000044407 | - | 1.0561301 | 7.26794098 | 6.21181087 | 1.1596231 | 0.00143017 | 0.00467231 |
| ENSMUSG00000057315 | Arhgap24 | 440.548268 | 998.732014 | 558.183746 | 1.15890047 | 1.4728E-29 | 7.4813E-28 |
| ENSMUSG00000098055 | - | 1.36741275 | 8.35040468 | 6.98299194 | 1.15874857 | 0.00142937 | 0.00467044 |
| ENSMUSG00000024620 | Pdgfrb | 2887.79516 | 6532.83454 | 3645.03938 | 1.15814809 | 3.7559E-36 | 2.9048E-34 |
| ENSMUSG00000063388 | - | 123.177536 | 409.765047 | 286.587511 | 1.15624826 | 0.00062943 | 0.00221345 |
| ENSMUSG00000048772 | Tmem53 | 174.470544 | 402.760608 | 228.290064 | 1.15619153 | 1.798E-14 | 2.6901E-13 |
| ENSMUSG00000085982 | 9530051G07 | 3.55478223 | 15.0127153 | 11.4579331 | 1.15594808 | 0.00112327 | 0.003758 |
| ENSMUSG00000074604 | Mgst2 | 488.735767 | 1121.72568 | 632.990056 | 1.15495246 | 1.2232E-16 | 2.298E-15 |
| ENSMUSG00000055322 | Tns1 | 478.945203 | 1103.41884 | 624.47364 | 1.15321265 | 1.0928E-15 | 1.8672E-14 |
| ENSMUSG00000086477 | - | 6.80157644 | 21.0502871 | 14.2487107 | 1.1530654 | 0.00032844 | 0.00122447 |
| ENSMUSG00000022747 | St3gal6 | 329.614089 | 751.763081 | 422.148991 | 1.15262321 | 6.3881E-21 | 1.7499E-19 |
| ENSMUSG00000049653 | Spatc1 | 1.32353447 | 9.81329005 | 8.48975559 | 1.15198474 | 0.00153098 | 0.00496994 |
| ENSMUSG00000023809 | Rps6ka2 | 1191.6436 | 2680.25377 | 1488.61017 | 1.14998805 | 1.8779E-36 | 1.4802E-34 |
| ENSMUSG00000033213 | AA467197 | 1160.88335 | 2643.59906 | 1482.71571 | 1.14692939 | 6.4978E-18 | 1.3724E-16 |
| ENSMUSG00000020684 | Rasl10b | 19.795773 | 71.8798478 | 52.0840748 | 1.14646931 | 0.0010173 | 0.00343305 |
| ENSMUSG00000020068 | - | 85.8086241 | 200.018841 | 114.210217 | 1.14578554 | 3.0295E-12 | 3.5512E-11 |
| ENSMUSG00000017740 | Slc12a5 | 71.2267833 | 174.336735 | 103.109952 | 1.14434745 | 4.626E-07 | 2.7657E-06 |
| ENSMUSG00000085118 | - | 2.42354285 | 11.0231527 | 8.59960989 | 1.14422904 | 0.00146534 | 0.00477961 |
| ENSMUSG00000086841 | 2410006H16 | 4120.21644 | 9287.47964 | 5167.2632 | 1.14380581 | 3.5252E-24 | 1.2271E-22 |
| ENSMUSG00000026822 | Lcn2 | 14699.1272 | 33409.1693 | 18710.0421 | 1.14327628 | 8.8496E-18 | 1.8445E-16 |
| ENSMUSG00000074056 | - | 15.5568036 | 59.7347882 | 44.1779847 | 1.14296775 | 0.00122858 | 0.00407914 |
| ENSMUSG00000093682 | - | 5.16675933 | 21.8801885 | 16.7134291 | 1.14276319 | 0.00140908 | 0.00461218 |
| ENSMUSG00000054263 | Lifr | 1313.9721 | 3025.79984 | 1711.82774 | 1.14268753 | 5.5292E-13 | 7.0041E-12 |
| ENSMUSG00000045215 | Asxl3 | 85.8975296 | 219.69883 | 133.8013 | 1.14244211 | 1.0139E-05 | 4.923E-05 |
| ENSMUSG00000025887 | Casp12 | 1872.13088 | 4190.89825 | 2318.76738 | 1.1418605 | 1.5349E-31 | 8.9195E-30 |
| ENSMUSG00000022453 | Naga | 1319.71834 | 2938.248 | 1618.52966 | 1.1418185 | 4.8176E-49 | 7.6237E-47 |
| ENSMUSG00000040729 | Cep126 | 22.6369388 | 69.9662343 | 47.3292956 | 1.14085866 | 0.00048486 | 0.0017487 |
| ENSMUSG00000091400 | - | 31.0757869 | 79.77908 | 48.7032931 | 1.14010363 | 1.7729E-05 | 8.2833E-05 |
| ENSMUSG00000000392 | Fap | 14.5067315 | 48.8822097 | 34.3754782 | 1.13995171 | 0.00084853 | 0.00290981 |
| ENSMUSG00000029321 | Slc10a6 | 53.3455702 | 130.86074 | 77.5151694 | 1.13986504 | 1.6018E-06 | 8.8177E-06 |
| ENSMUSG00000085583 | - | 33.1913417 | 77.9145418 | 44.7232001 | 1.1395783 | 1.3083E-08 | 9.7517E-08 |
| ENSMUSG00000085586 | - | 50.2471928 | 120.267939 | 70.0207462 | 1.1395685 | 1.6186E-07 | 1.0344E-06 |
| ENSMUSG00000037393 | Nmur2 | 0.31128264 | 5.38041714 | 5.0691345 | 1.13935003 | 0.00151304 | 0.00492116 |
| ENSMUSG00000086006 | Gm13293 | 4.06675094 | 16.0180802 | 11.9513292 | 1.13885726 | 0.00128382 | 0.00424132 |
| ENSMUSG00000028871 | Rspo1 | 4.77197643 | 16.0972869 | 11.3253105 | 1.13877164 | 0.00096694 | 0.0032787 |
| ENSMUSG00000089984 | Fbxo24 | 59.2058694 | 138.106508 | 78.9006383 | 1.13842048 | 1.8468E-09 | 1.5388E-08 |
| ENSMUSG00000024232 | Bambi | 218.986087 | 492.809846 | 273.823759 | 1.13762376 | 1.9377E-20 | 5.0268E-19 |
| ENSMUSG00000043085 | Tmem82 | 2.61583794 | 12.9571863 | 10.3413484 | 1.13761451 | 0.00170313 | 0.00546767 |
| ENSMUSG00000005951 | Shpk | 374.496191 | 853.957928 | 479.461738 | 1.13555641 | 2.9894E-14 | 4.3802E-13 |
| ENSMUSG00000026170 | Cyp27a1 | 20.9488907 | 53.0356494 | 32.0867587 | 1.13183758 | 6.1732E-06 | 3.1027E-05 |
| ENSMUSG00000005973 | Rcn1 | 2341.3896 | 5209.46939 | 2868.07979 | 1.1308096 | 1.8159E-28 | 8.6075E-27 |
| ENSMUSG00000027692 | Tnik | 68.1722842 | 179.761828 | 111.589543 | 1.13032089 | 6.3363E-05 | 0.00027094 |
| ENSMUSG00000029695 | Aass | 1.5335732 | 14.5184014 | 12.9848282 | 1.13025589 | 0.00173855 | 0.00557441 |
| ENSMUSG00000028362 | Tnfsf8 | 1.41968201 | 8.98684446 | 7.56716245 | 1.12987907 | 0.00189186 | 0.00601906 |
| ENSMUSG00000090069 | Gm1604a | 1.29739983 | 7.47375203 | 6.17635219 | 1.12971591 | 0.0018614 | 0.00592859 |
| ENSMUSG00000052392 | Acot4 | 2.78709476 | 19.6263038 | 16.839209 | 1.12877242 | 0.00188408 | 0.00599524 |
| ENSMUSG00000057726 | Serpinb9g | 3.76811559 | 14.6261691 | 10.8580535 | 1.128586 | 0.00153695 | 0.00498709 |

| | | | | | | | |
|---|---|---|---|---|---|---|---|
| ENSMUSG00000097458 | - | 1.01225182 | 9.63433838 | 8.62208656 | 1.1280073 | 0.00180948 | 0.00578023 |
| ENSMUSG00000054889 | Dsp | 12.7174395 | 33.9089092 | 21.1914697 | 1.12756504 | 0.00011519 | 0.00046897 |
| ENSMUSG00000050138 | Kcnk12 | 54.9284269 | 143.572995 | 88.6445677 | 1.12735446 | 5.3576E-05 | 0.00023121 |
| ENSMUSG00000040329 | Il7 | 126.520786 | 296.54451 | 170.023724 | 1.12727126 | 1.4899E-08 | 1.1017E-07 |
| ENSMUSG00000040435 | Ppp1r15a | 2855.1602 | 6431.87742 | 3576.71722 | 1.12724973 | 4.3963E-16 | 7.7899E-15 |
| ENSMUSG00000051243 | Islr2 | 4.00428895 | 18.4318089 | 14.4275199 | 1.12695003 | 0.00185902 | 0.00592232 |
| ENSMUSG00000085829 | - | 51.9342792 | 119.475497 | 67.5412177 | 1.12645653 | 9.8407E-10 | 8.4598E-09 |
| ENSMUSG00000011382 | Dhdh | 11.5912965 | 31.3839175 | 19.792621 | 1.12638061 | 0.00014265 | 0.00057051 |
| ENSMUSG00000050022 | Amz1 | 135.776828 | 310.485458 | 174.70863 | 1.12567089 | 5.2055E-11 | 5.2622E-10 |
| ENSMUSG00000087018 | - | 5.63581144 | 17.6322819 | 11.9964705 | 1.12525779 | 0.00076294 | 0.00264184 |
| ENSMUSG00000041608 | Entpd3 | 10.7815325 | 38.7646757 | 27.9831432 | 1.12521951 | 0.00128331 | 0.0042411 |
| ENSMUSG00000017390 | Aldoc | 128.540194 | 290.564463 | 162.02427 | 1.12483286 | 4.3348E-14 | 6.2448E-13 |
| ENSMUSG00000023883 | Phf10 | 6407.76084 | 14061.6941 | 7653.93323 | 1.12392555 | 6.7703E-61 | 1.8821E-58 |
| ENSMUSG00000006221 | Hspb7 | 1.46356029 | 8.4921519 | 7.02859161 | 1.1237398 | 0.00200141 | 0.00633236 |
| ENSMUSG00000095403 | - | 647.915174 | 1429.86913 | 781.953954 | 1.12370152 | 2.7256E-39 | 2.5959E-37 |
| ENSMUSG00000047822 | Angptl8 | 1.3496691 | 9.35944173 | 8.00977263 | 1.12357243 | 0.00199689 | 0.00632098 |
| ENSMUSG00000086043 | - | 6.47680645 | 19.2149885 | 12.7381821 | 1.12304547 | 0.00059528 | 0.00210524 |
| ENSMUSG00000047963 | Stbd1 | 1454.42013 | 3266.93272 | 1812.51259 | 1.12294804 | 8.3542E-16 | 1.443E-14 |
| ENSMUSG00000008035 | Mid1ip1 | 1491.99762 | 3302.57223 | 1810.57461 | 1.12290852 | 1.8946E-27 | 8.3459E-26 |
| ENSMUSG00000050447 | Lypd6 | 131.92593 | 303.942014 | 172.016083 | 1.12289848 | 5E-10 | 4.478E-09 |
| ENSMUSG00000037868 | Egr2 | 28.8556935 | 70.8382326 | 41.9825391 | 1.12248103 | 3.3053E-06 | 1.7329E-05 |
| ENSMUSG00000021662 | Arhgef28 | 182.192668 | 422.128845 | 239.936177 | 1.12125857 | 4.1509E-09 | 3.3034E-08 |
| ENSMUSG00000040181 | Fmo1 | 2314.91707 | 5709.91109 | 3394.99403 | 1.12109627 | 7.2262E-06 | 3.5882E-05 |
| ENSMUSG00000000197 | Nalcn | 1.43742566 | 7.84464619 | 6.40722053 | 1.12087187 | 0.00204016 | 0.00644208 |
| ENSMUSG00000034997 | Htr2a | 431.591014 | 977.137224 | 545.54621 | 1.11999894 | 6.5854E-13 | 8.2657E-12 |
| ENSMUSG00000006958 | Chrd | 62.5184203 | 154.433835 | 91.9154146 | 1.11977714 | 9.4336E-06 | 4.602E-05 |
| ENSMUSG00000020427 | Igfbp3 | 433.922104 | 1479.37858 | 1045.45648 | 1.11884428 | 0.00123572 | 0.00410088 |
| ENSMUSG00000026424 | Gpr37l1 | 6.28960772 | 20.3990673 | 14.1094596 | 1.11854723 | 0.00084417 | 0.00289728 |
| ENSMUSG00000018166 | Erbb3 | 1.91996512 | 9.46862044 | 7.54865532 | 1.11850952 | 0.00197862 | 0.00626991 |
| ENSMUSG00000097504 | - | 4.72809815 | 15.0337747 | 10.3056765 | 1.11781305 | 0.00099647 | 0.00337051 |
| ENSMUSG00000032327 | Stra6 | 1.08226474 | 13.227602 | 12.1453373 | 1.11778977 | 0.00182403 | 0.005824 |
| ENSMUSG00000050821 | Fam131a | 287.176337 | 648.232826 | 361.056488 | 1.1177827 | 1.6363E-12 | 1.9743E-11 |
| ENSMUSG00000097134 | - | 8.66417593 | 23.7585134 | 15.0943374 | 1.11736975 | 0.00019808 | 0.00077205 |
| ENSMUSG00000042279 | H1f8 | 0.31128264 | 4.85338884 | 4.54210619 | 1.11732927 | 0.00188299 | 0.00599269 |
| ENSMUSG00000089817 | - | 4.68931623 | 15.6815798 | 10.9922636 | 1.11685511 | 0.00101017 | 0.00341181 |
| ENSMUSG00000059326 | Csf2ra | 110.017487 | 250.889922 | 140.872435 | 1.11677665 | 4.9872E-11 | 5.0466E-10 |
| ENSMUSG00000039853 | Trim14 | 127.504327 | 301.406666 | 173.902339 | 1.11628186 | 2.1313E-07 | 1.338E-06 |
| ENSMUSG00000040183 | Ankrd6 | 101.092496 | 245.132978 | 144.040482 | 1.11595046 | 3.5013E-06 | 1.8286E-05 |
| ENSMUSG00000032507 | Fbxl2 | 160.268506 | 357.464763 | 197.196257 | 1.11549401 | 3.1405E-15 | 5.1276E-14 |
| ENSMUSG00000050538 | - | 2.0422473 | 10.0382631 | 7.99601585 | 1.11520862 | 0.00208103 | 0.00655828 |
| ENSMUSG00000029060 | Mib2 | 1561.62778 | 3436.26421 | 1874.63643 | 1.11507025 | 5.5317E-28 | 2.5121E-26 |
| ENSMUSG00000022871 | Fetub | 61.8555802 | 144.23312 | 82.3775401 | 1.1146512 | 1.9956E-08 | 1.4511E-07 |
| ENSMUSG00000039633 | Lonrf1 | 826.521253 | 1813.13333 | 986.612079 | 1.11302303 | 6.7714E-32 | 4.0144E-30 |
| ENSMUSG00000002771 | Grin2d | 34.4358194 | 86.0162386 | 51.5804191 | 1.11254394 | 1.2248E-05 | 5.8719E-05 |
| ENSMUSG00000032482 | Cspg5 | 8.95771493 | 31.938934 | 22.9812191 | 1.1122879 | 0.00148848 | 0.00484587 |
| ENSMUSG00000032271 | Nnmt | 401.795197 | 982.045123 | 580.249926 | 1.11152586 | 8.0703E-06 | 3.9833E-05 |
| ENSMUSG00000023055 | Calcoco1 | 1445.08367 | 3142.42416 | 1697.34049 | 1.11024816 | 3.171E-55 | 6.6565E-53 |
| ENSMUSG00000054679 | Srsf12 | 75.4620277 | 175.534515 | 100.072487 | 1.10913106 | 5.0472E-08 | 3.4529E-07 |
| ENSMUSG00000028059 | Arhgef2 | 14664.749 | 31850.4939 | 17185.7449 | 1.10899768 | 9.2402E-60 | 2.437E-57 |
| ENSMUSG00000040936 | Ulk4 | 108.645287 | 243.228933 | 134.583645 | 1.1088471 | 4.0321E-16 | 7.1816E-15 |
| ENSMUSG00000028801 | Stpg1 | 0.74484746 | 5.90206517 | 5.15721771 | 1.10810736 | 0.00224126 | 0.0070157 |
| ENSMUSG00000028864 | Hgf | 157.929612 | 525.614064 | 367.684451 | 1.10793473 | 0.00132089 | 0.00435115 |
| ENSMUSG00000021913 | Ogdhl | 10.5039354 | 28.5717303 | 18.0677948 | 1.10744728 | 0.0003572 | 0.00132353 |
| ENSMUSG00000025427 | Rnf165 | 167.73227 | 372.28228 | 204.55001 | 1.10741989 | 8.6359E-15 | 1.3348E-13 |
| ENSMUSG00000059022 | Kcp | 7.41065438 | 23.9337834 | 16.523129 | 1.10649918 | 0.00119119 | 0.00396525 |
| ENSMUSG00000025170 | Rab40b | 35.7701995 | 88.2239717 | 52.4537723 | 1.106318 | 1.493E-05 | 7.0639E-05 |
| ENSMUSG00000036820 | Amdhd2 | 780.653084 | 1748.83665 | 968.183564 | 1.10593862 | 2.654E-12 | 3.1288E-11 |
| ENSMUSG00000075256 | Cerkl | 0.33741727 | 5.70703779 | 5.36962051 | 1.1057636 | 0.00198922 | 0.0062996 |
| ENSMUSG00000021125 | Arg2 | 26.386349 | 69.4155807 | 43.0292316 | 1.1046412 | 0.00014194 | 0.00056841 |
| ENSMUSG00000031770 | Herpud1 | 6452.9584 | 14109.5238 | 7656.56541 | 1.10459776 | 7.8518E-26 | 3.0885E-24 |
| ENSMUSG00000097574 | C920006O11 | 67.2287748 | 150.753797 | 83.5250222 | 1.10392754 | 7.3941E-11 | 7.3484E-10 |
| ENSMUSG00000028133 | Rwdd3 | 254.348547 | 554.862838 | 300.514291 | 1.10379872 | 4.7194E-24 | 1.6208E-22 |
| ENSMUSG00000026875 | Traf1 | 123.386491 | 279.115539 | 155.729048 | 1.10359046 | 5.113E-10 | 4.5733E-09 |
| ENSMUSG00000071796 | 6820431F20I | 337.338689 | 738.425647 | 401.086958 | 1.10306113 | 2.8284E-28 | 1.3194E-26 |
| ENSMUSG00000020227 | Irak3 | 313.645964 | 686.74807 | 373.102106 | 1.10285198 | 8.3722E-23 | 2.6579E-21 |
| ENSMUSG00000074024 | 4632427E13 | 115.657211 | 273.604691 | 157.94748 | 1.10255817 | 9.0446E-07 | 5.1757E-06 |
| ENSMUSG00000097885 | - | 26.5838621 | 62.5985137 | 36.0146517 | 1.10240939 | 1.4344E-06 | 7.9474E-06 |
| ENSMUSG00000093452 | - | 48.9897238 | 115.17692 | 66.1871966 | 1.10156158 | 3.2992E-07 | 2.0176E-06 |
| ENSMUSG00000041935 | AW549877 | 4162.24098 | 9034.54349 | 4872.30251 | 1.10114864 | 1.1913E-35 | 8.9122E-34 |
| ENSMUSG00000033576 | Apol6 | 464.269582 | 1059.81061 | 595.541032 | 1.1010782 | 8.1664E-09 | 6.2733E-08 |
| ENSMUSG00000069792 | Wfdc17 | 0 | 8.21707055 | 8.21707055 | 1.09953336 | 0.0015695 | 0.00507749 |
| ENSMUSG00000027995 | Tlr2 | 3699.76003 | 8474.45431 | 4774.69428 | 1.09947453 | 2.3537E-08 | 1.6977E-07 |
| ENSMUSG00000027315 | Spint1 | 36.4241174 | 89.527499 | 53.1033817 | 1.09929869 | 2.1999E-05 | 0.0001014 |
| ENSMUSG00000032006 | Pdgfd | 74.1465753 | 309.522955 | 235.37638 | 1.09909135 | 0.00228826 | 0.00714542 |
| ENSMUSG00000097016 | - | 4.01448168 | 17.603951 | 13.5894694 | 1.09899274 | 0.00231906 | 0.00723065 |

| | | | | | | | |
|---|---|---|---|---|---|---|---|
| ENSMUSG00000043613 | Mmp3 | 41281.3532 | 91375.3548 | 50094.0016 | 1.09812277 | 4.6578E-14 | 6.6774E-13 |
| ENSMUSG00000026764 | Kif5c | 411.102188 | 893.37246 | 482.270272 | 1.09691619 | 1.2608E-30 | 6.9909E-29 |
| ENSMUSG00000058656 | Samd12 | 143.192478 | 317.064956 | 173.872478 | 1.09649232 | 2.8263E-14 | 4.1501E-13 |
| ENSMUSG00000020681 | Ace | 3.30841614 | 11.9364075 | 8.62799137 | 1.09625303 | 0.00198583 | 0.00629079 |
| ENSMUSG00000057176 | Ccdc189 | 80.8590239 | 180.759048 | 99.9000243 | 1.09622105 | 1.2633E-10 | 1.2328E-09 |
| ENSMUSG00000032565 | Nudt16 | 398.009647 | 866.173898 | 468.164251 | 1.09609124 | 2.5159E-22 | 7.6339E-21 |
| ENSMUSG00000018920 | Cxcl16 | 663.486719 | 1484.6991 | 821.212378 | 1.09538218 | 1.0277E-10 | 1.0082E-09 |
| ENSMUSG00000044197 | Gpr146 | 108.049919 | 241.789495 | 133.739576 | 1.09404024 | 1.6915E-10 | 1.6207E-09 |
| ENSMUSG00000050103 | Agmo | 1.0561301 | 7.65648901 | 6.6003589 | 1.09382551 | 0.00257605 | 0.00794879 |
| ENSMUSG00000097339 | - | 15.5390599 | 38.7891502 | 23.2500903 | 1.09307856 | 6.5254E-05 | 0.00027851 |
| ENSMUSG00000041809 | Efhc1 | 126.885953 | 281.979083 | 155.09313 | 1.0920684 | 6.6921E-11 | 6.696E-10 |
| ENSMUSG00000090523 | Gypc | 3089.75458 | 6637.40937 | 3547.65479 | 1.09202655 | 5.0942E-49 | 7.9392E-47 |
| ENSMUSG00000007039 | Ddah2 | 1183.00866 | 2556.1231 | 1373.11445 | 1.08938598 | 2.8374E-26 | 1.1445E-24 |
| ENSMUSG00000097150 | - | 12.6558176 | 32.1440516 | 19.488234 | 1.08924617 | 0.00012141 | 0.00049185 |
| ENSMUSG00000030443 | Zfp583 | 40.7624073 | 94.3857852 | 53.6233779 | 1.08913634 | 3.249E-07 | 1.991E-06 |
| ENSMUSG00000097731 | - | 1.0561301 | 6.60233199 | 5.54620189 | 1.08846823 | 0.00275751 | 0.00843531 |
| ENSMUSG00000097657 | - | 32.488758 | 88.3074527 | 55.8186946 | 1.08758561 | 0.00039501 | 0.00144825 |
| ENSMUSG00000026456 | Cyb5r1 | 4503.70763 | 9704.86861 | 5201.16098 | 1.08739043 | 7.8702E-29 | 3.8366E-27 |
| ENSMUSG00000085085 | - | 17.1020624 | 45.5780959 | 28.4760335 | 1.08738059 | 0.0004021 | 0.00147266 |
| ENSMUSG00000028600 | Podn | 2.11226021 | 8.79296008 | 6.68069987 | 1.08728782 | 0.00251687 | 0.0077825 |
| ENSMUSG00000091764 | Zfp964 | 147.366631 | 321.137444 | 173.770812 | 1.08697685 | 1.7282E-16 | 3.1772E-15 |
| ENSMUSG00000068105 | Tnfrsf13c | 0.33741727 | 4.69946955 | 4.36205227 | 1.08695568 | 0.00245402 | 0.00760532 |
| ENSMUSG00000027463 | Slc52a3 | 73.9387688 | 171.047381 | 97.1086118 | 1.08691585 | 5.0819E-07 | 3.0155E-06 |
| ENSMUSG00000067215 | Usp51 | 21.0609802 | 56.4027509 | 35.3417708 | 1.08689649 | 0.00036285 | 0.0013411 |
| ENSMUSG00000041836 | Ptpre | 50.6278356 | 121.954711 | 71.3268756 | 1.0851574 | 1.544E-05 | 7.2866E-05 |
| ENSMUSG00000071553 | Cpa2 | 18.5902644 | 46.8620361 | 28.2717717 | 1.08503055 | 8.5056E-05 | 0.00035428 |
| ENSMUSG00000020930 | Ccdc103 | 31.4345513 | 78.3143069 | 46.8797557 | 1.0843929 | 5.3011E-05 | 0.00022906 |
| ENSMUSG00000025355 | Mmp19 | 1029.86908 | 2205.90358 | 1176.03449 | 1.0842562 | 2.7384E-39 | 2.596E-37 |
| ENSMUSG00000034842 | Art3 | 6.43292817 | 17.7841558 | 11.3512276 | 1.08403352 | 0.00066906 | 0.0023404 |
| ENSMUSG00000000489 | Pdgfb | 24.3348707 | 64.2406561 | 39.9057854 | 1.08323362 | 0.00033835 | 0.00125777 |
| ENSMUSG00000063652 | Slc22a21 | 85.0820515 | 191.356167 | 106.274116 | 1.08272542 | 9.824E-09 | 7.47E-08 |
| ENSMUSG00000032207 | Lipc | 8.65907957 | 28.0781717 | 19.4190921 | 1.08247497 | 0.00173958 | 0.00557685 |
| ENSMUSG00000047686 | Rtl3 | 560.080158 | 1196.6696 | 636.589445 | 1.08224554 | 1.9229E-40 | 1.9391E-38 |
| ENSMUSG00000052160 | Pld4 | 1.80097757 | 8.75119798 | 6.95022042 | 1.08166325 | 0.00289788 | 0.00882272 |
| ENSMUSG00000035513 | Ntng2 | 4.79811106 | 26.6620396 | 21.8639285 | 1.08163441 | 0.00293574 | 0.00893007 |
| ENSMUSG00000014773 | Dll1 | 4.04996898 | 14.0695308 | 10.0195618 | 1.08121197 | 0.00196418 | 0.00622701 |
| ENSMUSG00000068747 | Sort1 | 105.948282 | 258.466437 | 152.518155 | 1.08097276 | 4.0405E-05 | 0.00017795 |
| ENSMUSG00000031506 | Ptpn7 | 8.30572039 | 23.0781121 | 14.7723917 | 1.0807518 | 0.0004822 | 0.00173941 |
| ENSMUSG00000021464 | Ror2 | 30.2515738 | 73.6268703 | 43.3752965 | 1.07969953 | 4.0181E-05 | 0.00017704 |
| ENSMUSG00000038530 | Rgs4 | 3277.34963 | 7745.21658 | 4467.86695 | 1.07960842 | 8.7147E-06 | 4.2757E-05 |
| ENSMUSG00000052131 | Akr1b7 | 5.89482482 | 19.0852477 | 13.1904229 | 1.07879722 | 0.0017314 | 0.00555321 |
| ENSMUSG00000033361 | Prrg3 | 345.622875 | 976.787284 | 631.164409 | 1.07841367 | 0.00083506 | 0.00286886 |
| ENSMUSG00000030433 | Sbk2 | 3.69810267 | 19.822338 | 16.1242353 | 1.07815692 | 0.00303179 | 0.00919776 |
| ENSMUSG00000028789 | Azin2 | 461.299646 | 1001.2234 | 539.923759 | 1.07737716 | 2.903E-15 | 4.751E-14 |
| ENSMUSG00000085749 | - | 1.70483002 | 8.24300476 | 6.53817473 | 1.0772053 | 0.00297344 | 0.00903405 |
| ENSMUSG00000030699 | Tbx6 | 3.67706441 | 14.2302148 | 10.5531504 | 1.07678374 | 0.00254468 | 0.00786024 |
| ENSMUSG00000040987 | Mill2 | 11.8782463 | 31.3955711 | 19.5173248 | 1.07654402 | 0.00023133 | 0.00089184 |
| ENSMUSG00000004631 | Sgce | 1772.7504 | 3792.78418 | 2020.03378 | 1.07651108 | 5.6854E-27 | 2.4265E-25 |
| ENSMUSG00000047638 | Nr1h4 | 70.1174223 | 155.844064 | 85.7266417 | 1.07645947 | 1.4888E-10 | 1.4372E-09 |
| ENSMUSG00000022584 | Ly6c2 | 16.7057866 | 40.2813709 | 23.5755844 | 1.07479626 | 3.1623E-05 | 0.00014173 |
| ENSMUSG00000005705 | Agrp | 14.0899486 | 34.8428413 | 20.7528927 | 1.07422603 | 6.8452E-05 | 0.00029059 |
| ENSMUSG00000031513 | Leprotl1 | 5345.58571 | 11375.1063 | 6029.52057 | 1.07227651 | 3.639E-32 | 2.1953E-30 |
| ENSMUSG00000087624 | - | 2.01611266 | 10.9387415 | 8.92262879 | 1.07186783 | 0.00320503 | 0.00966914 |
| ENSMUSG00000085596 | - | 16.7874851 | 45.68694 | 28.8994549 | 1.07153368 | 0.00069531 | 0.00242561 |
| ENSMUSG00000022012 | Enox1 | 51.4871054 | 116.55022 | 65.0631149 | 1.070926 | 3.3913E-07 | 2.0702E-06 |
| ENSMUSG00000043972 | Opn5 | 1.12614302 | 8.17489883 | 7.04875582 | 1.07090676 | 0.00310762 | 0.00940563 |
| ENSMUSG00000034271 | Jdp2 | 2591.64979 | 5460.64823 | 2868.99845 | 1.07070359 | 1.7686E-90 | 1.2546E-87 |
| ENSMUSG00000001930 | Vwf | 13.5229472 | 37.5150086 | 23.9920613 | 1.07020074 | 0.0008671 | 0.0029656 |
| ENSMUSG00000028354 | Fmn2 | 3.72094269 | 12.789593 | 9.06865033 | 1.06988663 | 0.00219266 | 0.00687613 |
| ENSMUSG00000031434 | Morc4 | 620.290936 | 1343.19415 | 722.903216 | 1.06942476 | 6.4459E-14 | 9.0454E-13 |
| ENSMUSG00000009628 | Tex15 | 52.3555056 | 171.470139 | 119.114633 | 1.06938648 | 0.00210931 | 0.00664004 |
| ENSMUSG00000039763 | Dnajc28 | 242.614548 | 520.089161 | 277.474613 | 1.06880387 | 6.9273E-21 | 1.8926E-19 |
| ENSMUSG00000053687 | Dpep2 | 28.5510001 | 68.8394287 | 40.2884286 | 1.06855674 | 5.8417E-05 | 0.00025084 |
| ENSMUSG00000029126 | Nsg1 | 1888.21167 | 3974.35535 | 2086.14367 | 1.06807371 | 1.2802E-77 | 6.5841E-75 |
| ENSMUSG00000085054 | - | 9.98261404 | 26.6461523 | 16.6635383 | 1.06705423 | 0.00063163 | 0.00221966 |
| ENSMUSG00000004508 | Gab2 | 506.918204 | 1127.69675 | 620.778545 | 1.06662078 | 2.5514E-08 | 1.8269E-07 |
| ENSMUSG00000087014 | - | 32.1478589 | 74.7171918 | 42.569333 | 1.06553224 | 1.5392E-05 | 7.2659E-05 |
| ENSMUSG00000032826 | Ank2 | 666.119994 | 1428.83503 | 762.715034 | 1.06532017 | 1.0355E-17 | 2.1323E-16 |
| ENSMUSG00000021751 | Acox2 | 1.02999547 | 6.27349188 | 5.24349641 | 1.06511613 | 0.00339494 | 0.01018827 |
| ENSMUSG00000028266 | Lmo4 | 8636.426 | 18192.39 | 9555.964 | 1.06508184 | 7.7364E-53 | 1.4601E-50 |
| ENSMUSG00000046413 | - | 8.6380413 | 22.7269642 | 14.0889229 | 1.06422984 | 0.00045729 | 0.00165943 |
| ENSMUSG00000097633 | Gm16617 | 58.08417 | 132.248331 | 74.1641612 | 1.06341379 | 6.6643E-07 | 3.8849E-06 |
| ENSMUSG00000045917 | Tmem268 | 442.893716 | 953.2699 | 510.376184 | 1.06283728 | 3.4321E-14 | 5.0004E-13 |
| ENSMUSG00000019359 | Gdpd2 | 2052.05243 | 4311.82453 | 2259.7721 | 1.06082724 | 4.0344E-57 | 9.3254E-55 |
| ENSMUSG00000031490 | Eif4ebp1 | 8385.35205 | 17995.6593 | 9610.30729 | 1.06050403 | 1.4263E-14 | 2.1623E-13 |

| | | | | | | | |
|---|---|---|---|---|---|---|---|
| ENSMUSG00000032717 | Mdfi | 15.7500603 | 47.2876415 | 31.5375812 | 1.06041501 | 0.00174565 | 0.00559368 |
| ENSMUSG00000035314 | Gdpd5 | 302.822168 | 649.888536 | 347.066368 | 1.05959329 | 1.0053E-13 | 1.3871E-12 |
| ENSMUSG00000033540 | Idua | 382.400969 | 818.976708 | 436.575739 | 1.05938227 | 9.9765E-16 | 1.7089E-14 |
| ENSMUSG00000026220 | Slc16a14 | 1.70483002 | 8.69037053 | 6.98554051 | 1.05913282 | 0.00357453 | 0.01067584 |
| ENSMUSG00000026303 | Mlph | 93.9375608 | 209.646113 | 115.708552 | 1.05889098 | 1.671E-07 | 1.0666E-06 |
| ENSMUSG00000022773 | Ypel1 | 105.641068 | 228.034084 | 122.393016 | 1.05791202 | 4.3319E-11 | 4.4354E-10 |
| ENSMUSG00000079057 | Cyp4v3 | 1184.1445 | 2506.46187 | 1322.31738 | 1.0566484 | 1.8557E-21 | 5.2874E-20 |
| ENSMUSG00000098332 | Pigbos1 | 8.55028474 | 24.5526431 | 16.0023584 | 1.05608958 | 0.00130461 | 0.00430241 |
| ENSMUSG00000030877 | Mfsd13b | 4.51805941 | 14.6706051 | 10.1525457 | 1.05587102 | 0.00236251 | 0.00735163 |
| ENSMUSG00000093390 | - | 4.88766936 | 16.2153646 | 11.3276953 | 1.05556474 | 0.00231965 | 0.00723137 |
| ENSMUSG00000019832 | Rab32 | 440.084778 | 926.250044 | 486.165266 | 1.05523273 | 3.5391E-24 | 1.2299E-22 |
| ENSMUSG00000081490 | - | 2.59479967 | 10.4695635 | 7.87476383 | 1.0545625 | 0.00326911 | 0.00984227 |
| ENSMUSG00000075284 | Wipf1 | 709.157455 | 1493.6877 | 784.530244 | 1.0536549 | 1.3002E-25 | 5.0372E-24 |
| ENSMUSG00000020814 | Mxra7 | 464.465394 | 1024.30799 | 559.842595 | 1.05343412 | 4.9581E-08 | 3.3976E-07 |
| ENSMUSG00000018570 | 2810408A11 | 150.17063 | 320.461195 | 170.290565 | 1.05341263 | 8.0196E-14 | 1.117E-12 |
| ENSMUSG00000023972 | Ptk7 | 980.311217 | 2047.66832 | 1067.3571 | 1.05274446 | 3.2747E-59 | 8.317E-57 |
| ENSMUSG00000021720 | Rnf180 | 0.81486037 | 6.40371693 | 5.58885655 | 1.05267325 | 0.00353066 | 0.01055709 |
| ENSMUSG00000023951 | Vegfa | 7722.03412 | 16153.7598 | 8431.72563 | 1.05219514 | 1.0677E-40 | 1.0928E-38 |
| ENSMUSG00000055945 | Prr18 | 21.7535584 | 50.1644908 | 28.4109325 | 1.05154766 | 1.9636E-05 | 9.1226E-05 |
| ENSMUSG00000028463 | Car9 | 452.571414 | 1141.84847 | 689.277055 | 1.05059503 | 0.00036548 | 0.00134936 |
| ENSMUSG00000026730 | Pter | 371.0247 | 788.964629 | 417.939928 | 1.05049865 | 5.0365E-15 | 8.0132E-14 |
| ENSMUSG00000045312 | Lhfpl2 | 5218.934 | 10855.6855 | 5636.75151 | 1.05043693 | 1.5281E-85 | 9.8237E-83 |
| ENSMUSG00000029092 | D5Ertd615e | 0.33741727 | 4.33308912 | 3.99567185 | 1.04954169 | 0.00338666 | 0.01016788 |
| ENSMUSG00000001076 | C1ql4 | 6.44131915 | 19.0208988 | 12.5795796 | 1.04868391 | 0.00198526 | 0.00628997 |
| ENSMUSG00000015829 | Tnr | 19.2119896 | 46.4098413 | 27.1978517 | 1.04864906 | 0.00017921 | 0.00070439 |
| ENSMUSG00000032623 | Oas1d | 5.01834252 | 16.3935231 | 11.3751806 | 1.04813152 | 0.00253784 | 0.00784263 |
| ENSMUSG00000020848 | Doc2b | 7.16938465 | 21.1491344 | 13.9797497 | 1.04811764 | 0.00196339 | 0.00622643 |
| ENSMUSG00000038522 | Mfsd4b1 | 2.64706893 | 12.4858475 | 9.83877861 | 1.04784076 | 0.00390519 | 0.0115577 |
| ENSMUSG00000053161 | Daw1 | 17.7913459 | 46.2523979 | 28.461052 | 1.04749977 | 0.00070064 | 0.00244175 |
| ENSMUSG00000097322 | - | 2.78709476 | 12.2267864 | 9.43969168 | 1.04736094 | 0.00387677 | 0.01148847 |
| ENSMUSG00000042190 | Cmklr1 | 2.18227312 | 9.8065539 | 7.62428078 | 1.04710421 | 0.00392658 | 0.01161599 |
| ENSMUSG00000017713 | Tha1 | 187.348517 | 406.937278 | 219.588761 | 1.04695029 | 5.4745E-09 | 4.3034E-08 |
| ENSMUSG00000086290 | Snhg12 | 1114.00999 | 2346.37641 | 1232.36641 | 1.04669113 | 2.2824E-19 | 5.4724E-18 |
| ENSMUSG00000005958 | Ephb3 | 559.26796 | 1170.41409 | 611.146126 | 1.04629814 | 7.6693E-28 | 3.4675E-26 |
| ENSMUSG00000027669 | Gnb4 | 1554.96829 | 3228.14833 | 1673.18004 | 1.04626229 | 4.429E-56 | 9.7972E-54 |
| ENSMUSG00000022126 | Acod1 | 98.6007424 | 269.459928 | 170.859186 | 1.04544697 | 0.00117967 | 0.00393007 |
| ENSMUSG00000026413 | Pkp1 | 726.528379 | 1654.07305 | 927.544669 | 1.04531553 | 8.0407E-06 | 3.9706E-05 |
| ENSMUSG00000015709 | Arnt2 | 410.637027 | 881.528287 | 470.89126 | 1.04516891 | 4.1932E-11 | 4.3016E-10 |
| ENSMUSG00000082100 | - | 18.522706 | 52.3211866 | 33.7984805 | 1.04508878 | 0.00161282 | 0.00520128 |
| ENSMUSG00000073409 | H2-Q6 | 183.629907 | 475.959395 | 292.329488 | 1.04507447 | 0.00066711 | 0.00233476 |
| ENSMUSG00000046167 | Gldn | 11.3238922 | 39.3586688 | 28.0347766 | 1.04505394 | 0.00325881 | 0.009817 |
| ENSMUSG00000058715 | Fcer1g | 1.70483002 | 8.05851348 | 6.35368346 | 1.0448758 | 0.00401161 | 0.01185218 |
| ENSMUSG00000034330 | Plcg2 | 56.3014672 | 122.961698 | 66.6602305 | 1.04466774 | 1.9221E-08 | 1.4012E-07 |
| ENSMUSG00000038390 | Gpr162 | 37.1487666 | 86.7249263 | 49.5761597 | 1.04431181 | 2.6579E-05 | 0.00012062 |
| ENSMUSG00000041920 | Slc16a6 | 97.2109201 | 208.07033 | 110.85941 | 1.04374906 | 2.0356E-12 | 2.4276E-11 |
| ENSMUSG00000060441 | Trim5 | 154.985953 | 332.212358 | 177.226405 | 1.04325678 | 1.0409E-10 | 1.0207E-09 |
| ENSMUSG00000092219 | - | 27.2973569 | 67.1943231 | 39.8969661 | 1.04289632 | 0.0002512 | 0.00096034 |
| ENSMUSG00000001604 | Tcea3 | 34.076215 | 77.967176 | 43.890961 | 1.04289241 | 2.3683E-05 | 0.00010841 |
| ENSMUSG00000005667 | Mthfd2 | 11529.4454 | 24048.4471 | 12519.0016 | 1.04266469 | 1.0931E-28 | 5.2789E-27 |
| ENSMUSG00000081611 | - | 9.50187632 | 24.2256644 | 14.7237881 | 1.04189865 | 0.0004103 | 0.00149975 |
| ENSMUSG00000097212 | - | 28.073557 | 67.2544552 | 39.1808982 | 1.04104131 | 0.00015168 | 0.00060332 |
| ENSMUSG00000028255 | Clca1 | 0.33741727 | 4.3977747 | 4.06035743 | 1.04019251 | 0.00363617 | 0.01083791 |
| ENSMUSG00000048281 | Dleu7 | 0 | 6.09461735 | 6.09461735 | 1.04007284 | 0.00261728 | 0.00805184 |
| ENSMUSG00000044948 | Cfap43 | 119.440342 | 255.525435 | 136.085092 | 1.03930016 | 1.3927E-09 | 1.1786E-08 |
| ENSMUSG00000085269 | - | 39.4388729 | 87.9140002 | 48.4751273 | 1.03890644 | 1.3056E-06 | 7.2691E-06 |
| ENSMUSG00000037580 | Gch1 | 3660.38982 | 7637.03848 | 3976.64866 | 1.03862715 | 4.6617E-23 | 1.5079E-21 |
| ENSMUSG00000027230 | Creb3l1 | 559.62302 | 1178.37175 | 618.748732 | 1.03853183 | 5.0573E-16 | 8.8999E-15 |
| ENSMUSG00000040010 | Slc7a5 | 14900.9932 | 31097.0182 | 16196.025 | 1.03830832 | 9.7074E-23 | 3.0723E-21 |
| ENSMUSG00000024905 | Tesmin | 7.56926391 | 21.8116276 | 14.2423636 | 1.03702138 | 0.00161533 | 0.00520774 |
| ENSMUSG00000034570 | Inpp5j | 6.66910154 | 18.7044733 | 12.0353718 | 1.03698957 | 0.00195323 | 0.00619805 |
| ENSMUSG00000035509 | Fbxl21 | 71.3318531 | 166.594564 | 95.2627105 | 1.03696395 | 6.3708E-05 | 0.00027236 |
| ENSMUSG00000086328 | - | 1.55970783 | 7.89994156 | 6.34023372 | 1.03682233 | 0.00435681 | 0.01275123 |
| ENSMUSG00000086873 | - | 9.86033186 | 24.4027921 | 14.5424603 | 1.03619292 | 0.00043003 | 0.00156826 |
| ENSMUSG00000052613 | Pcdh15 | 2.54253041 | 10.9843191 | 8.44178868 | 1.03611181 | 0.00418159 | 0.01228735 |
| ENSMUSG00000020034 | Tcp11l2 | 1367.51604 | 2836.32757 | 1468.81153 | 1.03565909 | 2.2511E-30 | 1.2228E-28 |
| ENSMUSG00000044254 | Pcsk9 | 19.1732077 | 102.156235 | 82.9830273 | 1.03520263 | 0.00439683 | 0.01284825 |
| ENSMUSG00000091514 | Gm17484 | 38.5452997 | 84.3011181 | 45.7558184 | 1.03334324 | 2.0205E-06 | 1.0921E-05 |
| ENSMUSG00000053007 | Creb5 | 381.1916 | 821.75644 | 440.56484 | 1.0329988 | 8.5153E-09 | 6.5242E-08 |
| ENSMUSG00000097386 | - | 19.6422598 | 46.1286359 | 26.4863761 | 1.03293216 | 0.00011944 | 0.00048454 |
| ENSMUSG00000021619 | Atg10 | 501.903565 | 1041.84801 | 539.944444 | 1.03285212 | 5.3062E-24 | 1.8133E-22 |
| ENSMUSG00000064325 | Hhip | 0.71871283 | 5.8434823 | 5.12476947 | 1.03272548 | 0.00417638 | 0.01227729 |
| ENSMUSG00000026981 | Il1rn | 160.151629 | 510.357327 | 350.205698 | 1.03204894 | 0.00315903 | 0.00954857 |
| ENSMUSG00000022847 | Thpo | 8.51575912 | 23.2147792 | 14.6990201 | 1.03176596 | 0.00131408 | 0.00433155 |
| ENSMUSG00000073988 | Ttpa | 66.0349518 | 142.392781 | 76.3578289 | 1.03166953 | 7.7937E-08 | 5.1887E-07 |
| ENSMUSG00000063506 | Arhgap22 | 92.3313329 | 210.851822 | 118.520489 | 1.03161441 | 3.0576E-05 | 0.00013745 |

| | | | | | | | |
|---|---|---|---|---|---|---|---|
| ENSMUSG00000091472 | Gm3739 | 58.7012084 | 127.12623 | 68.4250214 | 1.03158561 | 5.6635E-08 | 3.8466E-07 |
| ENSMUSG00000086924 | - | 6.95934591 | 18.7682607 | 11.8089148 | 1.03115232 | 0.0013224 | 0.0043548 |
| ENSMUSG00000021747 | Cfap20dc | 158.072933 | 351.552815 | 193.479882 | 1.03072753 | 3.6373E-06 | 1.8963E-05 |
| ENSMUSG00000037386 | Rims2 | 18.4806295 | 50.4068055 | 31.926176 | 1.02994571 | 0.00157331 | 0.0050874 |
| ENSMUSG00000051920 | Rspo2 | 5210.14573 | 10687.1414 | 5476.99566 | 1.02978205 | 1.9335E-69 | 7.6494E-67 |
| ENSMUSG00000086224 | - | 0 | 5.10999891 | 5.10999891 | 1.02940744 | 0.0029112 | 0.00886066 |
| ENSMUSG00000091418 | - | 0 | 3.62574287 | 3.62574287 | 1.0291412 | 0.00307445 | 0.00931069 |
| ENSMUSG00000029660 | Tex26 | 23.3664971 | 52.5475285 | 29.1810314 | 1.0281662 | 2.3395E-05 | 0.00010717 |
| ENSMUSG00000056427 | Slit3 | 785.439769 | 2090.73337 | 1305.2936 | 1.02788719 | 0.00132108 | 0.00435115 |
| ENSMUSG00000050737 | Ptges | 145.652879 | 314.111063 | 168.458184 | 1.02713227 | 5.1857E-08 | 3.5395E-07 |
| ENSMUSG00000066800 | Rnasel | 104.70135 | 224.300004 | 119.598654 | 1.02710484 | 6.876E-09 | 5.3399E-08 |
| ENSMUSG00000022676 | Snai2 | 7753.81491 | 16180.227 | 8426.41209 | 1.02688624 | 1.3795E-15 | 2.3358E-14 |
| ENSMUSG00000039328 | Rnf122 | 388.912215 | 798.292062 | 409.379847 | 1.02677896 | 1.5312E-39 | 1.4859E-37 |
| ENSMUSG00000029307 | Dmp1 | 6.6741979 | 63.3817851 | 56.7075872 | 1.02660023 | 0.0040035 | 0.01183164 |
| ENSMUSG00000024897 | Apba1 | 9.54904922 | 25.6347655 | 16.0857163 | 1.02615519 | 0.00156295 | 0.00505947 |
| ENSMUSG00000085971 | - | 27.4990047 | 61.4401257 | 33.941121 | 1.0258896 | 6.9399E-06 | 3.4535E-05 |
| ENSMUSG00000023912 | Slc25a27 | 140.875376 | 294.407746 | 153.53237 | 1.02541001 | 8.7978E-14 | 1.2188E-12 |
| ENSMUSG00000094417 | - | 1.08226474 | 6.30156 | 5.21929526 | 1.025238 | 0.00477277 | 0.01382114 |
| ENSMUSG00000071335 | Mfsd4b3-ps | 57.1748773 | 125.709048 | 68.5341709 | 1.02479988 | 5.9489E-06 | 2.9951E-05 |
| ENSMUSG00000045555 | Mettl24 | 14.9393346 | 36.4136795 | 21.4743449 | 1.02476974 | 0.00040667 | 0.00148826 |
| ENSMUSG00000023781 | Hes7 | 17.8766479 | 44.1140056 | 26.2373577 | 1.02441553 | 0.00036196 | 0.00133876 |
| ENSMUSG00000062345 | Serpinb2 | 49.0473118 | 109.267207 | 60.2198948 | 1.02406186 | 1.439E-05 | 6.8319E-05 |
| ENSMUSG00000074500 | Zfp558 | 55.6251398 | 123.041101 | 67.4159616 | 1.0239447 | 1.605E-06 | 8.8327E-06 |
| ENSMUSG00000034035 | Ccdc17 | 33.8727655 | 75.5716162 | 41.6988508 | 1.02333047 | 2.5537E-05 | 0.0001162 |
| ENSMUSG00000028943 | Espn | 1.41968201 | 7.43969082 | 6.02000881 | 1.0225112 | 0.00492597 | 0.01421078 |
| ENSMUSG00000044288 | Cnr1 | 20.5812041 | 48.0003117 | 27.4191076 | 1.02131384 | 0.0001041 | 0.00042746 |
| ENSMUSG00000031778 | Cx3cl1 | 1461.63953 | 3007.2242 | 1545.58467 | 1.02091574 | 2.5528E-24 | 8.977E-23 |
| ENSMUSG00000034245 | Hdac11 | 537.892969 | 1121.23339 | 583.340419 | 1.02075738 | 1.8549E-13 | 2.4762E-12 |
| ENSMUSG00000086719 | - | 1.46356029 | 8.47197275 | 7.00841246 | 1.02074936 | 0.00492136 | 0.01420146 |
| ENSMUSG00000020806 | Rhbdf2 | 1096.29066 | 2239.00102 | 1142.71037 | 1.0206347 | 5.5457E-56 | 1.2137E-53 |
| ENSMUSG00000064179 | Tnnt1 | 5.70156805 | 18.6682764 | 12.9667083 | 1.02036652 | 0.00368955 | 0.01097634 |
| ENSMUSG00000039316 | Rftn1 | 553.846479 | 1152.93282 | 599.086337 | 1.02005827 | 8.2719E-14 | 1.1506E-12 |
| ENSMUSG00000015619 | Gata3 | 256.747945 | 528.782587 | 272.034642 | 1.01997292 | 3.3283E-22 | 1.0054E-20 |
| ENSMUSG00000001240 | Ramp2 | 310.325366 | 803.639717 | 493.314351 | 1.01960304 | 0.00118531 | 0.00394693 |
| ENSMUSG00000033207 | Mamdc2 | 22.3466944 | 53.264061 | 30.9173666 | 1.01886886 | 0.00026818 | 0.0010179 |
| ENSMUSG00000018500 | Adora2b | 423.347948 | 873.436681 | 450.088734 | 1.01876442 | 7.3954E-18 | 1.5524E-16 |
| ENSMUSG00000087651 | 1500009L16I | 1.94609975 | 10.488822 | 8.54272229 | 1.01871096 | 0.00508333 | 0.01460535 |
| ENSMUSG00000048794 | Cfap100 | 79.4416749 | 237.237468 | 157.795794 | 1.01870436 | 0.00306229 | 0.00927935 |
| ENSMUSG00000030344 | Akap3 | 32.2433536 | 74.1794355 | 41.9360819 | 1.01865247 | 5.5092E-05 | 0.00023735 |
| ENSMUSG00000039375 | Wdr17 | 8.94422758 | 28.0314374 | 19.0872099 | 1.01814716 | 0.00361986 | 0.01079399 |
| ENSMUSG00000031112 | Stk26 | 3.62479514 | 12.3380161 | 8.71322091 | 1.01692866 | 0.00397738 | 0.01175614 |
| ENSMUSG00000040537 | Adam22 | 574.954353 | 1188.81283 | 613.858481 | 1.01680429 | 1.6056E-16 | 2.9676E-15 |
| ENSMUSG00000074771 | Ankef1 | 10.106698 | 25.4323648 | 15.3256668 | 1.01632065 | 0.00065094 | 0.00228167 |
| ENSMUSG00000085584 | Rtl9 | 5.00995154 | 19.2176652 | 14.2077136 | 1.01600728 | 0.00484618 | 0.01401178 |
| ENSMUSG00000070632 | - | 51.9868573 | 111.344666 | 59.3578085 | 1.01531799 | 2.6936E-08 | 1.9254E-07 |
| ENSMUSG00000062488 | Ifit3b | 5038.49496 | 10510.5012 | 5472.0062 | 1.01498524 | 6.0486E-12 | 6.8482E-11 |
| ENSMUSG00000018008 | Cyth4 | 11.1577317 | 35.3302909 | 24.1725593 | 1.0149688 | 0.00379662 | 0.01127369 |
| ENSMUSG00000075042 | - | 129.21065 | 298.316671 | 169.106021 | 1.01484908 | 0.00011755 | 0.00047772 |
| ENSMUSG00000075254 | Heg1 | 1189.27546 | 2445.87271 | 1256.59725 | 1.01464659 | 8.3215E-20 | 2.0625E-18 |
| ENSMUSG00000070691 | Runx3 | 512.807258 | 1073.141 | 560.333739 | 1.01400804 | 7.0754E-11 | 7.0589E-10 |
| ENSMUSG00000032572 | Col6a4 | 1.41968201 | 8.56759108 | 7.14790907 | 1.01385541 | 0.00517555 | 0.01484336 |
| ENSMUSG00000041552 | Ptchd1 | 183.044717 | 391.118907 | 208.07419 | 1.01354566 | 6.9765E-08 | 4.6749E-07 |
| ENSMUSG00000021185 | Dglucy | 78.9234258 | 168.237008 | 89.313582 | 1.01336413 | 8.2746E-09 | 6.3517E-08 |
| ENSMUSG00000036617 | Etl4 | 189.73846 | 404.777001 | 215.038541 | 1.01242862 | 6.2012E-08 | 4.1909E-07 |
| ENSMUSG00000086150 | Bach2os | 1.15227765 | 6.66609853 | 5.51382089 | 1.01175972 | 0.00528723 | 0.01511939 |
| ENSMUSG00000031596 | Slc7a2 | 13486.1511 | 27733.9081 | 14247.7569 | 1.01169673 | 1.2411E-17 | 2.5379E-16 |
| ENSMUSG00000066361 | Serpina3c | 0.33741727 | 4.31398574 | 3.97656847 | 1.01097383 | 0.00456149 | 0.01326908 |
| ENSMUSG00000028373 | Astn2 | 26.7397082 | 60.2794067 | 33.5396985 | 1.01080614 | 6.8146E-05 | 0.00028935 |
| ENSMUSG00000063594 | Gng8 | 6.55521034 | 18.2267816 | 11.6715713 | 1.00995045 | 0.00276747 | 0.00846453 |
| ENSMUSG00000038756 | Ttll6 | 11.4900526 | 27.2180982 | 15.7280456 | 1.00930745 | 0.00052177 | 0.00186806 |
| ENSMUSG00000042759 | Apobr | 392.215965 | 834.008754 | 441.792789 | 1.00879717 | 9.5788E-08 | 6.3017E-07 |
| ENSMUSG00000054901 | Arhgef33 | 1.73096465 | 7.82472547 | 6.09376081 | 1.00879716 | 0.00549674 | 0.01565216 |
| ENSMUSG00000053615 | - | 23.4289591 | 52.1483927 | 28.7194336 | 1.00717818 | 1.9898E-05 | 9.2299E-05 |
| ENSMUSG00000097313 | - | 12.3412403 | 32.4914963 | 20.1502559 | 1.00684114 | 0.00167198 | 0.00537521 |
| ENSMUSG00000047531 | Rtp2 | 0 | 3.51170483 | 3.51170483 | 1.00678555 | 0.00365976 | 0.01089559 |
| ENSMUSG00000019880 | Rspo3 | 300.221067 | 623.923223 | 323.702157 | 1.0057069 | 6.953E-11 | 6.9469E-10 |
| ENSMUSG00000074151 | Nlrc5 | 756.736657 | 1560.14502 | 803.408358 | 1.00565262 | 4.1805E-14 | 6.0348E-13 |
| ENSMUSG00000039217 | Il18 | 88.8763017 | 189.936256 | 101.059955 | 1.00534099 | 4.5332E-07 | 2.7149E-06 |
| ENSMUSG00000097462 | 9530026P05 | 16.2628691 | 42.9394287 | 26.6765596 | 1.00517555 | 0.00184779 | 0.00589528 |
| ENSMUSG00000050921 | P2ry10 | 6.75865984 | 21.5165792 | 14.7579194 | 1.00506712 | 0.00414644 | 0.01219798 |
| ENSMUSG00000053626 | Tll1 | 3.16839031 | 10.8547196 | 7.68632931 | 1.00482537 | 0.00480164 | 0.01390084 |
| ENSMUSG00000063659 | Zbtb18 | 1829.33235 | 3690.21163 | 1860.87929 | 1.00470623 | 1.1336E-62 | 3.239E-60 |
| ENSMUSG00000006920 | Ezh1 | 2046.91388 | 4134.35301 | 2087.43912 | 1.0043305 | 4.7611E-49 | 7.6163E-47 |
| ENSMUSG00000031129 | Slc9a9 | 185.012386 | 382.099614 | 197.087228 | 1.00376459 | 5.8261E-13 | 7.3531E-12 |
| ENSMUSG00000070509 | Rgma | 490.81686 | 1003.26924 | 512.452384 | 1.00328071 | 1.3561E-19 | 3.3053E-18 |

| | | | | | | | |
|---|---|---|---|---|---|---|---|
| ENSMUSG00000000791 | Il12rb1 | 5.14901568 | 17.1725747 | 12.023559 | 1.00308801 | 0.004673 | 0.01355513 |
| ENSMUSG00000075410 | Prcd | 11.4857963 | 28.2056356 | 16.7198393 | 1.00305164 | 0.00086631 | 0.00296336 |
| ENSMUSG00000010142 | Tnfrsf13b | 3.97995606 | 12.708112 | 8.72815589 | 1.00237915 | 0.0041163 | 0.01212319 |
| ENSMUSG00000064165 | Krt39 | 4.89096398 | 16.3811683 | 11.4902043 | 1.00237333 | 0.00482817 | 0.01397371 |
| ENSMUSG00000032925 | Itgbl1 | 3.43579467 | 15.6490474 | 12.2132527 | 1.00231842 | 0.00584337 | 0.01653505 |
| ENSMUSG00000030020 | Prickle2 | 100.211227 | 270.759581 | 170.548354 | 1.0021016 | 0.00237488 | 0.00738454 |
| ENSMUSG00000030707 | Coro1a | 17.6066018 | 43.4270594 | 25.8204576 | 1.00183599 | 0.00114236 | 0.00381564 |
| ENSMUSG00000097013 | - | 21.1444804 | 49.9169103 | 28.7724299 | 1.00178547 | 0.00041828 | 0.00152786 |
| ENSMUSG00000038515 | Grtp1 | 98.6377226 | 210.910295 | 112.272572 | 1.00174512 | 1.0401E-06 | 5.8866E-06 |
| ENSMUSG00000073437 | D330041H03 | 253.150985 | 518.700919 | 265.549934 | 1.00138796 | 7.0337E-17 | 1.3536E-15 |
| ENSMUSG00000044060 | - | 325.560624 | 672.047204 | 346.486579 | 1.00098153 | 1.286E-12 | 1.5691E-11 |
| ENSMUSG00000072244 | Trim6 | 13.8225443 | 32.9963757 | 19.1738314 | 1.00048423 | 0.00047987 | 0.0017316 |
| ENSMUSG00000036995 | Asap3 | 139.55004 | 286.239774 | 146.689734 | 1.00006081 | 1.8086E-12 | 2.1682E-11 |
| ENSMUSG00000040811 | Eml2 | 849.599509 | 1707.75567 | 858.156162 | 0.9995086 | 6.8622E-58 | 1.6042E-55 |
| ENSMUSG00000040328 | Olfr56 | 36.2443479 | 78.5420006 | 42.2976527 | 0.99890621 | 3.3818E-06 | 1.7716E-05 |
| ENSMUSG00000035234 | Abraxas1 | 1137.75261 | 2287.18844 | 1149.43583 | 0.99863152 | 3.9912E-46 | 5.1316E-44 |
| ENSMUSG00000074766 | Ism1 | 1557.67877 | 3129.19659 | 1571.51782 | 0.99850338 | 2.1749E-51 | 3.7917E-49 |
| ENSMUSG00000022537 | Tmem44 | 209.406855 | 430.143162 | 220.736307 | 0.99828493 | 6.1654E-14 | 8.6814E-13 |
| ENSMUSG00000030800 | Prss8 | 1.46356029 | 9.10713117 | 7.64357088 | 0.99766441 | 0.00582675 | 0.01649711 |
| ENSMUSG00000063683 | Glyat | 3.46192931 | 17.1046847 | 13.6427554 | 0.99722059 | 0.00609487 | 0.01716409 |
| ENSMUSG00000014030 | Pax5 | 19.0070473 | 45.1671569 | 26.1601096 | 0.99712051 | 0.00056732 | 0.00201605 |
| ENSMUSG00000044229 | Nxpe4 | 146.140616 | 298.997612 | 152.856996 | 0.99613762 | 1.2892E-12 | 1.5712E-11 |
| ENSMUSG00000032281 | Acsbg1 | 470.321868 | 1050.82028 | 580.49841 | 0.99613533 | 7.4516E-05 | 0.00031361 |
| ENSMUSG00000042035 | Igsf3 | 1028.42796 | 2071.7861 | 1043.35814 | 0.99607102 | 3.5871E-35 | 2.6168E-33 |
| ENSMUSG00000030532 | Hddc3 | 282.919093 | 574.013314 | 291.094221 | 0.99583226 | 3.7806E-17 | 7.4071E-16 |
| ENSMUSG00000034910 | Pygo1 | 11.1610263 | 31.2283834 | 20.0673571 | 0.99575731 | 0.00327944 | 0.00986903 |
| ENSMUSG00000078306 | - | 4.49192478 | 13.2681806 | 8.7762558 | 0.99488316 | 0.00381814 | 0.01132942 |
| ENSMUSG00000002459 | Rgs20 | 33.591221 | 76.5257386 | 42.9345177 | 0.99486109 | 0.00017854 | 0.00070214 |
| ENSMUSG00000097393 | D030068K23 | 41.1175683 | 88.0397798 | 46.9222116 | 0.99471517 | 1.0238E-06 | 5.802E-06 |
| ENSMUSG00000037138 | Aff3 | 17.4767687 | 40.9323264 | 23.4555577 | 0.99416727 | 0.0004583 | 0.00166192 |
| ENSMUSG00000097177 | 9330159M07 | 288.346202 | 596.176179 | 307.829977 | 0.99380525 | 3.3516E-10 | 3.0892E-09 |
| ENSMUSG00000045114 | Prrt2 | 0.64869992 | 6.35460382 | 5.7059039 | 0.99337951 | 0.00543215 | 0.01548429 |
| ENSMUSG00000052062 | Pard3b | 44.6383561 | 96.0241636 | 51.3858075 | 0.99328415 | 5.8236E-06 | 2.9371E-05 |
| ENSMUSG00000085395 | Gm13056 | 215.849272 | 443.022143 | 227.172871 | 0.99282704 | 1.4471E-10 | 1.3996E-09 |
| ENSMUSG00000030123 | Plxnd1 | 2180.263 | 4410.15746 | 2229.89446 | 0.99260132 | 7.0428E-20 | 1.754E-18 |
| ENSMUSG00000050473 | Slc35d3 | 1.39354738 | 7.34897295 | 5.95542557 | 0.99237878 | 0.00631643 | 0.01771287 |
| ENSMUSG00000067158 | Col4a4 | 10.5724554 | 83.9238415 | 73.3513861 | 0.99223423 | 0.00553244 | 0.01574183 |
| ENSMUSG00000095930 | Nim1k | 542.332711 | 1094.64535 | 552.312641 | 0.99209657 | 7.7987E-22 | 2.2985E-20 |
| ENSMUSG00000030074 | Gxylt2 | 543.114883 | 1106.46072 | 563.345841 | 0.99069331 | 2.3041E-13 | 3.0502E-12 |
| ENSMUSG00000051113 | Fam71e1 | 10.5823393 | 25.1211963 | 14.538857 | 0.9903721 | 0.00105204 | 0.00354101 |
| ENSMUSG00000068758 | Il3ra | 748.959519 | 1506.59564 | 757.636119 | 0.99006507 | 3.883E-22 | 1.1679E-20 |
| ENSMUSG00000097921 | - | 10.5318718 | 26.2548003 | 15.7229285 | 0.98973237 | 0.00124354 | 0.00412414 |
| ENSMUSG00000072707 | Olfr31 | 0.71871283 | 6.86400741 | 6.14529459 | 0.98961723 | 0.00555021 | 0.01578582 |
| ENSMUSG00000087480 | Gm15910 | 2.01611266 | 9.3347881 | 7.31867544 | 0.98860816 | 0.00655736 | 0.01830401 |
| ENSMUSG00000051224 | Tceanc | 326.054044 | 658.362602 | 332.308557 | 0.98858361 | 1.7289E-18 | 3.845E-17 |
| ENSMUSG00000024901 | Peli3 | 74.1679225 | 153.426961 | 79.2590381 | 0.98838624 | 3.0098E-09 | 2.4425E-08 |
| ENSMUSG00000032177 | Pde4a | 136.372196 | 285.565671 | 149.193475 | 0.98803021 | 1.4397E-07 | 9.2471E-07 |
| ENSMUSG00000067199 | Frat1 | 11.9464574 | 28.3285602 | 16.3821028 | 0.98772448 | 0.00076612 | 0.00265195 |
| ENSMUSG00000073647 | - | 162.473088 | 330.278986 | 167.805898 | 0.98766777 | 4.3863E-13 | 5.6362E-12 |
| ENSMUSG00000004791 | Pgf | 146.580791 | 327.869125 | 181.288334 | 0.98622428 | 0.00013577 | 0.00054563 |
| ENSMUSG00000027955 | Gask1b | 18.0209301 | 50.919516 | 32.8985859 | 0.98533726 | 0.0038673 | 0.01146205 |
| ENSMUSG00000028773 | Fabp3 | 16.1667216 | 37.674566 | 21.5078445 | 0.98504953 | 0.00047754 | 0.00172411 |
| ENSMUSG00000047419 | Cmya5 | 16.6745556 | 39.1852692 | 22.5107136 | 0.98495581 | 0.00080743 | 0.00278245 |
| ENSMUSG00000033174 | Mgll | 30.9684849 | 66.0850046 | 35.1165197 | 0.98403771 | 1.5068E-05 | 7.1209E-05 |
| ENSMUSG00000042804 | Gpr153 | 30.4261253 | 65.7999645 | 35.3738393 | 0.98294311 | 3.0122E-05 | 0.0001356 |
| ENSMUSG00000097336 | Fendrr | 16.8356197 | 54.749012 | 37.9133924 | 0.98221491 | 0.005859 | 0.01657244 |
| ENSMUSG00000058022 | Adtrp | 53.2469681 | 118.654582 | 65.4076136 | 0.9822014 | 0.00012736 | 0.00051435 |
| ENSMUSG00000031748 | Gnao1 | 756.561488 | 1519.50949 | 762.948003 | 0.98194503 | 2.0045E-20 | 5.1936E-19 |
| ENSMUSG00000041482 | Piezo2 | 64.3930142 | 140.474952 | 76.0819378 | 0.98168856 | 4.2951E-05 | 0.00018836 |
| ENSMUSG00000016756 | Cmah | 4.0532636 | 14.1049039 | 10.0516403 | 0.98068051 | 0.00645948 | 0.01805985 |
| ENSMUSG00000089989 | - | 106.772186 | 218.380449 | 111.608262 | 0.98018067 | 9.1596E-10 | 7.8941E-09 |
| ENSMUSG00000085614 | - | 26.4563619 | 58.5211656 | 32.0648036 | 0.97928987 | 8.4859E-05 | 0.0003536 |
| ENSMUSG00000029108 | Pcdh7 | 1450.85447 | 2912.35338 | 1461.4989 | 0.9789951 | 1.3996E-17 | 2.848E-16 |
| ENSMUSG00000033063 | Cntnap3 | 0 | 3.24827312 | 3.24827312 | 0.97847109 | 0.00457308 | 0.01329815 |
| ENSMUSG00000020081 | Tacr2 | 6.33678062 | 18.2543771 | 11.9175964 | 0.97842359 | 0.00451542 | 0.01314995 |
| ENSMUSG00000022219 | Cideb | 18.8492778 | 41.7411323 | 22.8918546 | 0.9782252 | 8.6026E-05 | 0.00035788 |
| ENSMUSG00000026601 | Axdnd1 | 15.1199441 | 34.7998361 | 19.679892 | 0.97772996 | 0.00046037 | 0.00166876 |
| ENSMUSG00000097536 | - | 16.8998834 | 46.0273454 | 29.127462 | 0.97730195 | 0.00354963 | 0.01060919 |
| ENSMUSG00000011148 | Adssl1 | 388.370574 | 783.170093 | 394.799519 | 0.97711027 | 9.3942E-14 | 1.2996E-12 |
| ENSMUSG00000038775 | Vill | 1187.70625 | 2359.75713 | 1172.05088 | 0.97703207 | 3.3235E-32 | 2.0109E-30 |
| ENSMUSG00000079505 | - | 26.4065472 | 57.6203406 | 31.2137933 | 0.97660035 | 0.00013132 | 0.00052878 |
| ENSMUSG00000085125 | - | 0 | 3.84053933 | 3.84053933 | 0.97659659 | 0.00451369 | 0.01314676 |
| ENSMUSG00000031551 | Ido1 | 31.0519852 | 73.7426257 | 42.6906405 | 0.97594691 | 0.00107431 | 0.00360768 |
| ENSMUSG00000087113 | Gm11714 | 4.01448168 | 12.186954 | 8.17247236 | 0.97535321 | 0.00496587 | 0.01431787 |
| ENSMUSG00000033717 | Adra2a | 216.718699 | 440.971666 | 224.252967 | 0.97517975 | 2.7265E-10 | 2.5414E-09 |

| | | | | | | | |
|---|---|---|---|---|---|---|---|
| ENSMUSG00000086638 | 4930405A21 | 33.8240997 | 70.8366529 | 37.0125532 | 0.97516325 | 8.7266E-07 | 5.0104E-06 |
| ENSMUSG00000020961 | Ston2 | 113.135289 | 239.987778 | 126.852489 | 0.97459414 | 6.1918E-06 | 3.1113E-05 |
| ENSMUSG00000038910 | Plcl2 | 1041.42206 | 2060.56542 | 1019.14335 | 0.97449045 | 1.6833E-38 | 1.4926E-36 |
| ENSMUSG00000030228 | Pik3c2g | 0.64869992 | 5.0795403 | 4.43084039 | 0.97331777 | 0.00685416 | 0.01903142 |
| ENSMUSG00000070371 | Prss36 | 187.56269 | 388.810729 | 201.248039 | 0.9732429 | 3.9209E-07 | 2.3738E-06 |
| ENSMUSG00000038615 | Nfe2l1 | 25648.8535 | 50556.5364 | 24907.6829 | 0.9729217 | 5.1496E-66 | 1.7956E-63 |
| ENSMUSG00000039410 | Prdm16 | 188.041282 | 383.616588 | 195.575306 | 0.97268421 | 3.3926E-09 | 2.7369E-08 |
| ENSMUSG00000085049 | Parvaos | 22.643528 | 49.6935256 | 27.0499976 | 0.97174253 | 0.00011306 | 0.00046139 |
| ENSMUSG00000038175 | Mylip | 319.922393 | 645.844858 | 325.922465 | 0.9715311 | 1.0004E-11 | 1.1059E-10 |
| ENSMUSG00000021702 | Thbs4 | 0.31128264 | 5.14502743 | 4.83374479 | 0.9712247 | 0.00576803 | 0.01634659 |
| ENSMUSG00000097787 | 2700046G09 | 66.8871012 | 139.343646 | 72.4565449 | 0.96972796 | 1.6645E-06 | 9.1286E-06 |
| ENSMUSG00000039676 | Capsl | 26.586195 | 58.8428694 | 32.2566743 | 0.9686234 | 0.00024808 | 0.00095001 |
| ENSMUSG00000041757 | Plekha6 | 93.8763694 | 257.391319 | 163.514949 | 0.96857922 | 0.00430573 | 0.01261608 |
| ENSMUSG00000027200 | Sema6d | 7.34573783 | 19.9594125 | 12.6136746 | 0.96846161 | 0.00365911 | 0.01089524 |
| ENSMUSG00000035064 | Eef2k | 1101.51337 | 2176.62349 | 1075.11012 | 0.96827563 | 3.2187E-26 | 1.2882E-24 |
| ENSMUSG00000097090 | - | 31.9311093 | 71.9234913 | 39.992382 | 0.96825121 | 0.00032327 | 0.0012074 |
| ENSMUSG00000024866 | Acy3 | 57.8172968 | 119.831496 | 62.0141988 | 0.96813331 | 5.1823E-07 | 3.0688E-06 |
| ENSMUSG00000040520 | Manea | 3332.68161 | 6550.06311 | 3217.3815 | 0.96800348 | 8.8268E-64 | 2.6704E-61 |
| ENSMUSG00000096401 | - | 298.8749 | 592.622564 | 293.747663 | 0.96785463 | 2.1867E-28 | 1.0247E-26 |
| ENSMUSG00000080783 | - | 11.7018931 | 28.9228316 | 17.2209385 | 0.96755509 | 0.0019348 | 0.0061424 |
| ENSMUSG00000073555 | Gm4951 | 189.098187 | 453.86709 | 264.768903 | 0.96740875 | 0.00151773 | 0.00493249 |
| ENSMUSG00000085112 | A530072M11 | 101.440106 | 203.706734 | 102.266628 | 0.96733634 | 8.5437E-13 | 1.0594E-11 |
| ENSMUSG00000031428 | Zcchc18 | 13.7609223 | 37.5420623 | 23.7811399 | 0.96710873 | 0.00417437 | 0.01227314 |
| ENSMUSG00000052749 | Gm38525; Tr | 12.8686198 | 33.0614466 | 20.1928268 | 0.9668532 | 0.0027713 | 0.00847499 |
| ENSMUSG00000040722 | Scamp5 | 227.116259 | 465.462763 | 238.346503 | 0.96615908 | 4.5109E-08 | 3.1162E-07 |
| ENSMUSG00000072774 | Zfp951 | 57.0827637 | 117.236507 | 60.1537438 | 0.96590138 | 1.011E-07 | 6.6237E-07 |
| ENSMUSG00000087371 | - | 8.08548893 | 19.9727579 | 11.887269 | 0.96519234 | 0.00225409 | 0.00705157 |
| ENSMUSG00000062432 | Cyp26c1 | 2.56866504 | 11.1846511 | 8.61598611 | 0.96506464 | 0.00793723 | 0.0216357 |
| ENSMUSG00000029470 | P2rx4 | 1957.59548 | 3894.3868 | 1936.79132 | 0.96498041 | 6.1696E-16 | 1.0774E-14 |
| ENSMUSG00000052276 | Ostn | 6.06937626 | 20.315723 | 14.2463467 | 0.96497311 | 0.00717015 | 0.01979923 |
| ENSMUSG00000098615 | - | 1.15227765 | 5.9846432 | 4.83236555 | 0.9648617 | 0.00781749 | 0.02134634 |
| ENSMUSG00000029468 | P2rx7 | 551.816191 | 1133.57068 | 581.754487 | 0.96396428 | 2.5311E-07 | 1.5712E-06 |
| ENSMUSG00000024164 | C3 | 15928.6888 | 31734.294 | 15805.6052 | 0.96362736 | 1.7636E-14 | 2.6405E-13 |
| ENSMUSG00000041351 | Rap1gap | 113.221896 | 237.87336 | 124.651463 | 0.96344423 | 1.0247E-05 | 4.9695E-05 |
| ENSMUSG00000030406 | Gipr | 0.74484746 | 5.0937594 | 4.34891194 | 0.9633516 | 0.00747389 | 0.02052502 |
| ENSMUSG00000097598 | - | 1.46356029 | 7.49210496 | 6.02854467 | 0.96331494 | 0.00795057 | 0.02166634 |
| ENSMUSG00000087424 | 5730405O15 | 15.8031696 | 35.9610913 | 20.1579216 | 0.96327364 | 0.00058334 | 0.00206692 |
| ENSMUSG00000047495 | Dlgap2 | 53.8601807 | 111.779783 | 57.9196023 | 0.96324066 | 1.6721E-06 | 9.1657E-06 |
| ENSMUSG00000082229 | Nap1l2 | 41.1731322 | 86.1140965 | 44.9409643 | 0.96264941 | 5.124E-06 | 2.6086E-05 |
| ENSMUSG00000044734 | Serpinb1a | 334.374967 | 669.327651 | 334.952684 | 0.96255822 | 1.4646E-11 | 1.5874E-10 |
| ENSMUSG00000020467 | Efemp1 | 6070.71457 | 12103.0341 | 6032.31957 | 0.96142138 | 3.3737E-13 | 4.3983E-12 |
| ENSMUSG00000036568 | Bicral | 1260.97083 | 2475.95935 | 1214.98852 | 0.96128134 | 1.4528E-34 | 1.0201E-32 |
| ENSMUSG00000086534 | - | 0.64869992 | 4.98936723 | 4.34066731 | 0.96027866 | 0.00757333 | 0.02075653 |
| ENSMUSG00000045251 | Zfp688 | 435.829401 | 858.366587 | 422.537186 | 0.95995767 | 8.9489E-22 | 2.6225E-20 |
| ENSMUSG00000070336 | Fbxo47 | 105.992691 | 214.498626 | 108.505934 | 0.95984999 | 8.3981E-09 | 6.4392E-08 |
| ENSMUSG00000022286 | Grhl2 | 0.71871283 | 5.81067052 | 5.09195769 | 0.95976858 | 0.00730397 | 0.02010669 |
| ENSMUSG00000097055 | - | 177.548723 | 352.130837 | 174.582114 | 0.95964293 | 4.1832E-14 | 6.0348E-13 |
| ENSMUSG00000097934 | 6720483E21 | 1.60868248 | 8.96768839 | 7.35900591 | 0.95943052 | 0.00812957 | 0.02209559 |
| ENSMUSG00000020604 | Arsg | 94.9903963 | 191.842856 | 96.8524596 | 0.95924551 | 1.3099E-09 | 1.1103E-08 |
| ENSMUSG00000031558 | Slit2 | 1037.40846 | 2278.74262 | 1241.33416 | 0.95921189 | 0.00024125 | 0.00092595 |
| ENSMUSG00000052675 | Zfp112 | 92.9673855 | 186.602446 | 93.6350603 | 0.95885946 | 1.5997E-09 | 1.341E-08 |
| ENSMUSG00000034855 | Cxcl10 | 3111.51221 | 7709.59212 | 4598.07991 | 0.95871706 | 0.00262452 | 0.00806928 |
| ENSMUSG00000022996 | Wnt10b | 3.0199735 | 10.5250306 | 7.50505714 | 0.95816468 | 0.00767215 | 0.0209978 |
| ENSMUSG00000047497 | Adamts12 | 189.681404 | 385.194929 | 195.513525 | 0.95773043 | 1.5771E-08 | 1.1625E-07 |
| ENSMUSG00000029348 | Asphd2 | 10.6970706 | 27.5602602 | 16.8631897 | 0.95771117 | 0.00350124 | 0.01047614 |
| ENSMUSG00000022311 | Csmd3 | 0.95998256 | 5.71370518 | 4.75372262 | 0.95729782 | 0.00826356 | 0.02242717 |
| ENSMUSG00000032246 | Calml4 | 48.5563814 | 105.002981 | 56.4466 | 0.95726101 | 0.00018389 | 0.00072138 |
| ENSMUSG00000040258 | Nxph4 | 39.1284303 | 88.3075046 | 49.1790743 | 0.95711892 | 0.0005005 | 0.00179974 |
| ENSMUSG00000020357 | Flt4 | 103.90149 | 309.607945 | 205.706455 | 0.95632472 | 0.00655142 | 0.01829453 |
| ENSMUSG00000035697 | Arhgap45 | 52.0219141 | 109.469833 | 57.447919 | 0.95546239 | 2.9578E-05 | 0.00013335 |
| ENSMUSG00000029816 | Gpnmb | 701.40621 | 1400.58321 | 699.177002 | 0.95507893 | 7.6446E-11 | 7.5863E-10 |
| ENSMUSG00000097815 | - | 844.886234 | 1694.61103 | 849.724798 | 0.95503559 | 8.9852E-10 | 7.7535E-09 |
| ENSMUSG00000070476 | Fam217b | 139.87524 | 291.240644 | 151.365404 | 0.95439573 | 9.0406E-06 | 4.4271E-05 |
| ENSMUSG00000092216 | Gm19345 | 7.95895045 | 24.4796955 | 16.5207451 | 0.95434288 | 0.00683861 | 0.01899337 |
| ENSMUSG00000041649 | Klf8 | 65.0735979 | 144.86409 | 79.7904917 | 0.95420889 | 0.00050322 | 0.00180792 |
| ENSMUSG00000064115 | Cadm2 | 644.809468 | 1299.88982 | 655.080348 | 0.95378378 | 1.3128E-08 | 9.7812E-08 |
| ENSMUSG00000019843 | Fyn | 5158.15757 | 10099.7849 | 4941.62733 | 0.95354858 | 4.1009E-25 | 1.5367E-23 |
| ENSMUSG00000046402 | Rbp1 | 700.350581 | 1412.75969 | 712.409107 | 0.95251917 | 2.8863E-08 | 2.0517E-07 |
| ENSMUSG00000021373 | Cap2 | 271.759833 | 570.173062 | 298.413229 | 0.9524568 | 2.2216E-05 | 0.00010229 |
| ENSMUSG00000038305 | Spats2l | 7.32619244 | 20.3943778 | 13.0681854 | 0.95216216 | 0.00598531 | 0.01688561 |
| ENSMUSG00000032841 | Prr5l | 58.8061774 | 127.154591 | 68.348414 | 0.95199699 | 0.00014853 | 0.00059184 |
| ENSMUSG00000028389 | Zfp37 | 322.397997 | 629.577758 | 307.17976 | 0.95144833 | 1.4889E-23 | 4.9561E-22 |
| ENSMUSG00000056216 | Cebpg | 5871.07396 | 11533.6787 | 5662.60478 | 0.95120151 | 5.5433E-18 | 1.1756E-16 |
| ENSMUSG00000087365 | - | 271.287865 | 536.981959 | 265.694094 | 0.95109105 | 1.0384E-11 | 1.145E-10 |
| ENSMUSG00000081189 | - | 1.70483002 | 6.66581456 | 4.96098454 | 0.95007921 | 0.00879471 | 0.02366887 |

| | | | | | | | |
|---|---|---|---|---|---|---|---|
| ENSMUSG00000084440 | - | 0.33741727 | 4.69966072 | 4.36224344 | 0.95003612 | 0.00694697 | 0.01924237 |
| ENSMUSG00000037762 | Slc16a9 | 34.7821589 | 78.2740749 | 43.491916 | 0.94987102 | 0.00071175 | 0.00247541 |
| ENSMUSG00000022357 | Klhl38 | 22.5212458 | 51.6511994 | 29.1299536 | 0.94980169 | 0.00112198 | 0.00375551 |
| ENSMUSG00000084809 | - | 1.80097757 | 7.08377249 | 5.28279493 | 0.9494939 | 0.00893914 | 0.02401358 |
| ENSMUSG00000019872 | Smpdl3a | 160.168224 | 323.129566 | 162.961342 | 0.94903908 | 1.1663E-07 | 7.5663E-07 |
| ENSMUSG00000072966 | Gprasp2 | 2.69094721 | 8.71647174 | 6.02552452 | 0.94893818 | 0.00764568 | 0.02093178 |
| ENSMUSG00000026207 | Speg | 245.778991 | 511.130427 | 265.351436 | 0.9487993 | 1.3123E-05 | 6.2679E-05 |
| ENSMUSG00000095868 | - | 2.47581211 | 8.73182383 | 6.25601171 | 0.94860543 | 0.00862617 | 0.02329757 |
| ENSMUSG00000097855 | A930007I19R | 20.8224738 | 45.5829271 | 24.7604533 | 0.94773285 | 0.00024954 | 0.0009549 |
| ENSMUSG00000049556 | Lingo1 | 3.77321195 | 14.7280039 | 10.9547919 | 0.9471986 | 0.00909938 | 0.02440261 |
| ENSMUSG00000046897 | Zfp740 | 4043.268 | 7834.47176 | 3791.20376 | 0.94650976 | 6.9541E-50 | 1.1726E-47 |
| ENSMUSG00000018930 | Ccl4 | 20.1686776 | 50.5185568 | 30.3498792 | 0.94634283 | 0.00354328 | 0.01059173 |
| ENSMUSG00000024302 | Dtna | 598.086909 | 1199.87863 | 601.791719 | 0.94580259 | 2.4928E-08 | 1.7887E-07 |
| ENSMUSG00000027661 | Slc2a10 | 32.5528345 | 73.1956026 | 40.6427681 | 0.94576987 | 0.00085267 | 0.00292206 |
| ENSMUSG00000020435 | Osbp2 | 59.4251392 | 121.956572 | 62.5314326 | 0.94565549 | 4.6546E-06 | 2.3784E-05 |
| ENSMUSG00000046186 | Cd109 | 2863.64298 | 5689.51618 | 2825.87319 | 0.94496207 | 4.7581E-10 | 4.2772E-09 |
| ENSMUSG00000037735 | 2810032G03 | 2.71198548 | 15.8636366 | 13.1516512 | 0.94484352 | 0.00875808 | 0.02359497 |
| ENSMUSG00000002032 | Tmem25 | 26.2142521 | 72.4653747 | 46.2511225 | 0.94477753 | 0.00615101 | 0.0172985 |
| ENSMUSG00000075316 | Scn9a | 5.56579853 | 16.7861472 | 11.2203486 | 0.94475657 | 0.0075007 | 0.02058491 |
| ENSMUSG00000056919 | Cep162 | 2507.79218 | 4913.44063 | 2405.64844 | 0.94473027 | 2.3916E-16 | 4.3501E-15 |
| ENSMUSG00000029605 | Oas1b | 1672.71718 | 3283.06892 | 1610.35174 | 0.94467148 | 7.0456E-15 | 1.1014E-13 |
| ENSMUSG00000094497 | - | 153.838751 | 307.847586 | 154.008835 | 0.94400813 | 2.7335E-08 | 1.9519E-07 |
| ENSMUSG00000023911 | Flywch2 | 212.271958 | 428.847546 | 216.575587 | 0.94398672 | 4.5403E-07 | 2.7184E-06 |
| ENSMUSG00000031586 | Rbpms | 8.30062402 | 26.7193905 | 18.4187665 | 0.94395993 | 0.00837052 | 0.02269949 |
| ENSMUSG00000086827 | - | 0.74484746 | 5.72355589 | 4.97870843 | 0.94369294 | 0.00827599 | 0.02245794 |
| ENSMUSG00000029370 | Rassf6 | 147.347926 | 290.531127 | 143.183201 | 0.94347594 | 6.3196E-12 | 7.1394E-11 |
| ENSMUSG00000021499 | Catsper3 | 1.0561301 | 7.38285689 | 6.32672679 | 0.94156654 | 0.00864362 | 0.02333857 |
| ENSMUSG00000022146 | Osmr | 5723.43895 | 11056.3497 | 5332.91072 | 0.94122782 | 1.3781E-43 | 1.5663E-41 |
| ENSMUSG00000088015 | Mir1932 | 2.23454238 | 8.44361119 | 6.20906881 | 0.94087492 | 0.00959977 | 0.0255679 |
| ENSMUSG00000079069 | 8430423G03 | 6.79318545 | 16.7961871 | 10.0030016 | 0.9408125 | 0.00314729 | 0.00951729 |
| ENSMUSG00000020682 | Mmp28 | 424.882878 | 830.999706 | 406.116828 | 0.94076683 | 5.9339E-16 | 1.038E-14 |
| ENSMUSG00000044678 | Ly6k | 1.97223438 | 8.39466717 | 6.42243279 | 0.93988112 | 0.00973521 | 0.02589511 |
| ENSMUSG00000036545 | Adamts2 | 2072.74755 | 3990.95896 | 1918.21141 | 0.93954682 | 4.7226E-70 | 1.9827E-67 |
| ENSMUSG00000034858 | Fam214a | 519.543474 | 1010.66279 | 491.119315 | 0.93941234 | 2.8718E-21 | 8.093E-20 |
| ENSMUSG00000007480 | Mc5r | 149.174816 | 316.670914 | 167.496098 | 0.93934789 | 0.00013092 | 0.00052746 |
| ENSMUSG00000027122 | Arl14ep | 9383.38557 | 18126.3622 | 8742.97666 | 0.93830188 | 5.8642E-32 | 3.4867E-30 |
| ENSMUSG00000037708 | Spag6 | 0.64869992 | 5.05019849 | 4.40149857 | 0.93764198 | 0.00889909 | 0.02391849 |
| ENSMUSG00000034731 | Dgkh | 78.2948024 | 158.015118 | 79.7203157 | 0.93747491 | 3.2578E-07 | 1.994E-06 |
| ENSMUSG00000085891 | Gm14634 | 4.31227698 | 12.0030422 | 7.69076523 | 0.93746338 | 0.00701264 | 0.01939815 |
| ENSMUSG00000026692 | Fmo4 | 86.0593472 | 169.712067 | 83.6527194 | 0.93506636 | 2.1022E-09 | 1.741E-08 |
| ENSMUSG00000010755 | Cars | 7470.51164 | 14380.0226 | 6909.51092 | 0.93478338 | 9.2052E-36 | 6.9366E-34 |
| ENSMUSG00000031934 | Panx1 | 293.548483 | 570.832052 | 277.283568 | 0.93475019 | 4.1972E-15 | 6.7509E-14 |
| ENSMUSG00000031294 | D630029K05 | 5.69317707 | 15.458552 | 9.7653749 | 0.93457087 | 0.0059836 | 0.0168831 |
| ENSMUSG00000085639 | - | 26.6049004 | 55.6726834 | 29.067783 | 0.93446662 | 0.00012251 | 0.00049613 |
| ENSMUSG00000083945 | - | 14.30934 | 34.044407 | 19.7350669 | 0.93436804 | 0.00284435 | 0.0086829 |
| ENSMUSG00000055137 | Sugct | 31.55407 | 68.4818703 | 36.9278003 | 0.93409782 | 0.00039089 | 0.00143468 |
| ENSMUSG00000020057 | Dram1 | 2428.7717 | 4988.96103 | 2560.18934 | 0.93400095 | 1.7899E-05 | 8.3574E-05 |
| ENSMUSG00000046991 | Wdr27 | 27.0047796 | 59.2249205 | 32.2201409 | 0.93369102 | 0.00065128 | 0.00228249 |
| ENSMUSG00000055421 | Pcdh9 | 45.0858387 | 156.315052 | 111.229214 | 0.9336455 | 0.00985642 | 0.02619043 |
| ENSMUSG00000027962 | Vcam1 | 6440.97807 | 12507.1107 | 6066.13259 | 0.93352719 | 1.9274E-16 | 3.5213E-15 |
| ENSMUSG00000027188 | Pamr1 | 2.70869086 | 9.91645804 | 7.20776718 | 0.93224983 | 0.01003777 | 0.02660358 |
| ENSMUSG00000040596 | Pogk | 1113.97632 | 2153.57787 | 1039.60155 | 0.93146565 | 1.7688E-20 | 4.6118E-19 |
| ENSMUSG00000032860 | P2ry2 | 10.4567625 | 31.0366072 | 20.5798447 | 0.93111898 | 0.00849993 | 0.02299586 |
| ENSMUSG00000087231 | E230016M11 | 64.247892 | 130.054314 | 65.8064221 | 0.93087867 | 2.924E-06 | 1.5471E-05 |
| ENSMUSG00000002409 | Dyrk1b | 458.365936 | 889.238336 | 430.872399 | 0.93071609 | 1.92E-15 | 3.1956E-14 |
| ENSMUSG00000037306 | Man1c1 | 340.983905 | 671.097911 | 330.114007 | 0.93037 | 1.5847E-09 | 1.3301E-08 |
| ENSMUSG00000041750 | Cd1d2 | 71.5251098 | 144.25436 | 72.7292505 | 0.93013477 | 2.1674E-06 | 1.1672E-05 |
| ENSMUSG00000040310 | Alx4 | 14.1449813 | 34.3132907 | 20.1683094 | 0.92981768 | 0.00309466 | 0.00937051 |
| ENSMUSG00000058152 | Chsy3 | 18.8525724 | 42.5127912 | 23.6602188 | 0.92923723 | 0.00157183 | 0.00508421 |
| ENSMUSG00000056413 | Adap1 | 8.9544203 | 22.1669734 | 13.2125531 | 0.92838582 | 0.00358726 | 0.01070921 |
| ENSMUSG00000035085 | 1700020L24R | 58.4600603 | 117.025823 | 58.5657626 | 0.92830472 | 1.9557E-06 | 1.0599E-05 |
| ENSMUSG00000053797 | Krt16 | 145.815623 | 289.934605 | 144.118981 | 0.928043 | 8.2519E-08 | 5.4786E-07 |
| ENSMUSG00000026786 | Apbb1ip | 37.7592158 | 125.055318 | 87.2961027 | 0.92789224 | 0.01006701 | 0.02666732 |
| ENSMUSG00000043190 | Rfesd | 933.510957 | 1792.43415 | 858.923192 | 0.92739593 | 4.8732E-24 | 1.6709E-22 |
| ENSMUSG00000011589 | Fsd1 | 267.812823 | 518.8619 | 251.049077 | 0.92710049 | 4.6437E-15 | 7.4284E-14 |
| ENSMUSG00000073008 | Gpr174 | 56.5216987 | 113.648888 | 57.1271898 | 0.92622313 | 9.5609E-07 | 5.4424E-06 |
| ENSMUSG00000074824 | Rslcan18 | 141.145235 | 272.347146 | 131.201911 | 0.92566207 | 6.8123E-14 | 9.5271E-13 |
| ENSMUSG00000068686 | Cd59b | 40.8973368 | 86.236418 | 45.3390812 | 0.9255805 | 0.00020553 | 0.00079971 |
| ENSMUSG00000014837 | 4931428F04R | 101.875473 | 201.126884 | 99.2514109 | 0.92320908 | 2.1455E-08 | 1.5541E-07 |
| ENSMUSG00000087477 | - | 6.05672898 | 15.9092454 | 9.85251644 | 0.9223057 | 0.00585847 | 0.01657244 |
| ENSMUSG00000059714 | Flot1 | 3940.4926 | 7673.55853 | 3733.06593 | 0.92203061 | 1.6216E-10 | 1.5567E-09 |
| ENSMUSG00000035967 | Ints6l | 978.195145 | 1903.40609 | 925.210941 | 0.92188595 | 9.8539E-11 | 9.6807E-10 |
| ENSMUSG00000012396 | Nanog | 73.9006397 | 149.097278 | 75.1966387 | 0.9205828 | 1.2273E-05 | 5.8812E-05 |
| ENSMUSG00000026463 | Atp2b4 | 1507.24189 | 2924.65392 | 1417.41203 | 0.92041008 | 1.6703E-11 | 1.8E-10 |
| ENSMUSG00000038292 | Kash5 | 6.72646716 | 16.7930986 | 10.0666314 | 0.92002192 | 0.00566628 | 0.01608705 |

| | | | | | | | |
|---|---|---|---|---|---|---|---|
| ENSMUSG00000085375 | - | 0.33741727 | 4.17616847 | 3.83875119 | 0.91947651 | 0.00892408 | 0.02397939 |
| ENSMUSG00000070604 | Vsig10l | 69.3143691 | 136.807229 | 67.4928603 | 0.91898009 | 1.9643E-07 | 1.2399E-06 |
| ENSMUSG00000084843 | - | 0.64869992 | 5.01827912 | 4.3695792 | 0.91892546 | 0.01017091 | 0.02690792 |
| ENSMUSG00000041729 | Coro2b | 35.3152875 | 83.1802286 | 47.8649411 | 0.91874023 | 0.00360127 | 0.01074792 |
| ENSMUSG00000062044 | Lmtk3 | 81.1739101 | 165.876076 | 84.7021661 | 0.91825958 | 4.3725E-05 | 0.00019155 |
| ENSMUSG00000090459 | - | 3.65092977 | 11.5289259 | 7.87799615 | 0.91780616 | 0.01018241 | 0.02693141 |
| ENSMUSG00000040231 | Syngr4 | 12.096676 | 28.8531581 | 16.7564821 | 0.917616 | 0.00344892 | 0.01033371 |
| ENSMUSG00000027438 | Napb | 559.127489 | 1081.30107 | 522.17358 | 0.91741581 | 2.7659E-12 | 3.2551E-11 |
| ENSMUSG00000097847 | - | 6.3806589 | 19.2681394 | 12.8874805 | 0.91695603 | 0.01016601 | 0.02690479 |
| ENSMUSG00000027360 | Hdc | 34.5638508 | 69.9724644 | 35.4086136 | 0.91673461 | 2.144E-05 | 9.8984E-05 |
| ENSMUSG00000019838 | Slc16a10 | 184.964561 | 359.442943 | 174.478382 | 0.91651528 | 1.4912E-11 | 1.6146E-10 |
| ENSMUSG00000019817 | Plagl1 | 15.5418233 | 35.0112078 | 19.4693845 | 0.91616543 | 0.00156731 | 0.00507199 |
| ENSMUSG00000021483 | Cdk20 | 373.636733 | 711.924381 | 338.287648 | 0.91610912 | 1.2995E-26 | 5.4669E-25 |
| ENSMUSG00000053783 | 1700016K19 | 0.31128264 | 4.08876738 | 3.77748474 | 0.91572059 | 0.00912375 | 0.02444885 |
| ENSMUSG00000074715 | Ccl28 | 8.70805421 | 20.8040508 | 12.0959966 | 0.91526429 | 0.00411783 | 0.01212598 |
| ENSMUSG00000061603 | Akap6 | 390.839539 | 763.754537 | 372.914998 | 0.91503311 | 7.4007E-09 | 5.7279E-08 |
| ENSMUSG00000098489 | Mir7678 | 0 | 3.085108 | 3.085108 | 0.91451111 | 0.00723096 | 0.01993691 |
| ENSMUSG00000041567 | Serpina12 | 0 | 3.09489979 | 3.09489979 | 0.91410072 | 0.00724968 | 0.01998397 |
| ENSMUSG00000069892 | 9930111J21R | 271.196845 | 542.904084 | 271.707239 | 0.91375305 | 6.9971E-06 | 3.4803E-05 |
| ENSMUSG00000086076 | - | 0.31128264 | 4.28398183 | 3.97269919 | 0.91367677 | 0.00912001 | 0.02444456 |
| ENSMUSG00000037375 | Hhat | 226.237701 | 432.043973 | 205.806272 | 0.91341894 | 2.5731E-14 | 3.7919E-13 |
| ENSMUSG00000031891 | Hsd11b2 | 8.04670701 | 27.6145597 | 19.5678527 | 0.91288106 | 0.01166253 | 0.03037751 |
| ENSMUSG00000000303 | Cdh1 | 12.1243035 | 27.9845722 | 15.8602687 | 0.9124644 | 0.00402213 | 0.0118786 |
| ENSMUSG00000040554 | Aipl1 | 10.036685 | 23.4344532 | 13.3977682 | 0.91242174 | 0.00271802 | 0.00832939 |
| ENSMUSG00000020486 | Septin4 | 4.42191187 | 13.2805702 | 8.85865829 | 0.91233595 | 0.01023865 | 0.02706276 |
| ENSMUSG00000025504 | Eps8l2 | 21.1505385 | 49.2918927 | 28.1413542 | 0.91226256 | 0.00342478 | 0.01026734 |
| ENSMUSG00000050106 | Tmc8 | 17.6876475 | 38.579044 | 20.8913965 | 0.91223642 | 0.00095408 | 0.00323882 |
| ENSMUSG00000031385 | Plxnb3 | 1.0561301 | 5.12597744 | 4.06984734 | 0.91158519 | 0.01197719 | 0.03109868 |
| ENSMUSG00000022324 | Matn2 | 3103.75512 | 5856.06192 | 2752.3068 | 0.91085838 | 2.1734E-59 | 5.5889E-57 |
| ENSMUSG00000037801 | Iqch | 2.76096013 | 9.18172768 | 6.42076756 | 0.91058097 | 0.01165118 | 0.03035178 |
| ENSMUSG00000027210 | Meis2 | 185.891202 | 357.19836 | 171.307158 | 0.9099789 | 7.2166E-13 | 9.014E-12 |
| ENSMUSG00000067203 | - | 399.466496 | 761.000391 | 361.533895 | 0.90958958 | 2.2385E-17 | 4.4579E-16 |
| ENSMUSG00000011008 | Mcoln2 | 2.08612558 | 7.68720132 | 5.60107574 | 0.90925576 | 0.01228807 | 0.03177753 |
| ENSMUSG00000059895 | Ptp4a3 | 620.335811 | 1178.94544 | 558.609633 | 0.9076435 | 3.7539E-18 | 8.0864E-17 |
| ENSMUSG00000081282 | - | 0.31128264 | 3.68381883 | 3.37253619 | 0.90706319 | 0.01006491 | 0.0266652 |
| ENSMUSG00000086077 | - | 1.41968201 | 7.11095922 | 5.69127721 | 0.90692456 | 0.01227829 | 0.03176021 |
| ENSMUSG00000053297 | - | 2.58970331 | 11.1456495 | 8.55594624 | 0.90654319 | 0.01257954 | 0.03241311 |
| ENSMUSG00000038894 | Irs2 | 17.99809 | 39.6681902 | 21.6701002 | 0.9064128 | 0.00215459 | 0.00677118 |
| ENSMUSG00000028145 | Them4 | 607.754429 | 1166.584 | 558.829575 | 0.906221 | 3.0797E-11 | 3.2192E-10 |
| ENSMUSG00000085282 | Gm15663 | 0.71871283 | 5.75218272 | 5.03346989 | 0.90588064 | 0.01076354 | 0.02831555 |
| ENSMUSG00000076435 | Acsf2 | 914.927885 | 1730.77019 | 815.842302 | 0.90584793 | 5.2687E-24 | 1.8035E-22 |
| ENSMUSG00000031980 | Agt | 4146.69831 | 8778.67213 | 4631.97383 | 0.90519557 | 0.00069856 | 0.00243573 |
| ENSMUSG00000047441 | Antxrl | 0.95998256 | 6.07191182 | 5.11192926 | 0.90509716 | 0.01182621 | 0.03072992 |
| ENSMUSG00000073617 | - | 273.564981 | 519.439792 | 245.874811 | 0.90462042 | 5.2826E-16 | 9.2804E-15 |
| ENSMUSG00000038665 | Dgki | 2.80483841 | 8.49115407 | 5.68631567 | 0.90446944 | 0.01137024 | 0.02971022 |
| ENSMUSG00000038020 | Rapgefl1 | 96.1158865 | 191.515813 | 95.3999261 | 0.90437028 | 6.6939E-06 | 3.344E-05 |
| ENSMUSG00000085408 | C530005A16 | 12.2922657 | 27.8054598 | 15.5131942 | 0.90432955 | 0.00306641 | 0.00929035 |
| ENSMUSG00000062380 | Tubb3 | 778.978099 | 1475.54704 | 696.568939 | 0.90404247 | 5.0433E-20 | 1.273E-18 |
| ENSMUSG00000043644 | 0610009L18R | 33.7177594 | 67.1584027 | 33.4406433 | 0.90324777 | 2.1886E-05 | 0.00010095 |
| ENSMUSG00000023224 | Serping1 | 2.58970331 | 9.15069459 | 6.56099128 | 0.90310125 | 0.01287984 | 0.03308329 |
| ENSMUSG00000084932 | - | 0.31128264 | 4.66027464 | 4.34899199 | 0.90290604 | 0.0096031 | 0.02557016 |
| ENSMUSG00000045608 | Dbx2 | 68.7136822 | 138.222237 | 69.508555 | 0.90290251 | 3.501E-05 | 0.00015579 |
| ENSMUSG00000041550 | Serpina5 | 0.64869992 | 4.79561348 | 4.14691357 | 0.90280743 | 0.01160802 | 0.03025853 |
| ENSMUSG00000036534 | Slc38a7 | 3059.52715 | 5814.32672 | 2754.79957 | 0.9022308 | 5.3356E-15 | 8.4368E-14 |
| ENSMUSG00000035041 | Creb3l3 | 1.36741275 | 6.16408131 | 4.79666856 | 0.90186804 | 0.01303637 | 0.03343942 |
| ENSMUSG00000006586 | Runx1t1 | 800.773664 | 1534.65805 | 733.884385 | 0.90173695 | 8.8573E-11 | 8.7434E-10 |
| ENSMUSG00000081123 | - | 0.62256529 | 5.47492239 | 4.85235711 | 0.90160762 | 0.01101872 | 0.0288908 |
| ENSMUSG00000034777 | Vax2 | 136.086309 | 267.548855 | 131.462546 | 0.90105626 | 4.0264E-06 | 2.0827E-05 |
| ENSMUSG00000028675 | Pnrc2 | 5985.36112 | 11280.2 | 5294.83889 | 0.90105369 | 4.326E-25 | 1.6181E-23 |
| ENSMUSG00000053821 | - | 8.77381082 | 23.4169605 | 14.6431497 | 0.90089869 | 0.00895928 | 0.02405512 |
| ENSMUSG00000047747 | Rnf150 | 6943.07804 | 13715.0254 | 6771.94736 | 0.90062306 | 6.5442E-06 | 3.2732E-05 |
| ENSMUSG00000053641 | Dennd4a | 1600.91101 | 3011.74285 | 1410.83184 | 0.90047223 | 7.2228E-35 | 5.1953E-33 |
| ENSMUSG00000087507 | - | 1.15227765 | 8.08292489 | 6.93064724 | 0.90040059 | 0.01143443 | 0.02985518 |
| ENSMUSG00000087366 | - | 35.7651031 | 76.1546951 | 40.3895921 | 0.90029038 | 0.00101132 | 0.00341456 |
| ENSMUSG00000032492 | Pth1r | 7.75136627 | 18.7606845 | 11.0093183 | 0.90023182 | 0.00689461 | 0.01912309 |
| ENSMUSG00000029999 | Tgfa | 27.6012103 | 56.5826663 | 28.981456 | 0.89960461 | 0.000211 | 0.00081961 |
| ENSMUSG00000029156 | Sgcb | 387.764603 | 730.943831 | 343.179228 | 0.89940596 | 6.6656E-20 | 1.6641E-18 |
| ENSMUSG00000051375 | Pcdh1 | 89.7008236 | 254.922608 | 165.221784 | 0.89928989 | 0.01099297 | 0.02884621 |
| ENSMUSG00000059401 | Mamld1 | 13.1644917 | 34.7828829 | 21.6183912 | 0.8986775 | 0.00924433 | 0.02473008 |
| ENSMUSG00000025235 | Bbs4 | 383.812583 | 724.826582 | 341.013999 | 0.89864552 | 3.0127E-18 | 6.5794E-17 |
| ENSMUSG00000031595 | Pdgfrl | 824.763408 | 1551.97869 | 727.21528 | 0.89859731 | 1.2412E-22 | 3.8865E-21 |
| ENSMUSG00000029330 | Cds1 | 3.97995606 | 13.5799738 | 9.60001774 | 0.89849876 | 0.01302578 | 0.03342476 |
| ENSMUSG00000098488 | Pla2g4b | 9.56253656 | 23.3333077 | 13.7707712 | 0.89801758 | 0.00619338 | 0.01740339 |
| ENSMUSG00000079671 | 2610203C22 | 3.5758205 | 10.3394523 | 6.76363177 | 0.89782257 | 0.01164622 | 0.03034271 |
| ENSMUSG00000050628 | Ubald2 | 1367.92723 | 2593.61909 | 1225.69185 | 0.89782106 | 8.5046E-14 | 1.179E-12 |

| | | | | | | | |
|---|---|---|---|---|---|---|---|
| ENSMUSG00000043518 | Rai2 | 29.3294115 | 61.4511106 | 32.1216991 | 0.89729246 | 0.0007648 | 0.00264783 |
| ENSMUSG00000097474 | - | 3.26963422 | 11.7398151 | 8.47018086 | 0.89727627 | 0.01340143 | 0.03427329 |
| ENSMUSG00000034009 | Rxfp1 | 2918.99962 | 6153.86875 | 3234.86913 | 0.89677406 | 0.00085759 | 0.00293744 |
| ENSMUSG00000018986 | Slfn3 | 80.8634675 | 162.235455 | 81.3719873 | 0.89632868 | 4.58E-05 | 0.00019987 |
| ENSMUSG00000049092 | Gpr137c | 85.8549569 | 171.599272 | 85.7443148 | 0.89600808 | 4.9295E-05 | 0.00021408 |
| ENSMUSG00000044364 | Tmem74b | 6.78808909 | 22.9439821 | 16.155893 | 0.8955537 | 0.01348842 | 0.03446575 |
| ENSMUSG00000061451 | Tmem151a | 154.958426 | 348.818242 | 193.859816 | 0.89531286 | 0.00341221 | 0.01023563 |
| ENSMUSG00000036278 | Dennd2d | 62.8433119 | 127.768871 | 64.9255595 | 0.89486649 | 0.00024212 | 0.0009291 |
| ENSMUSG00000015890 | Amdhd1 | 4.1545075 | 14.7870165 | 10.6325089 | 0.89472742 | 0.01375831 | 0.03506391 |
| ENSMUSG00000005950 | P2rx5 | 116.473377 | 223.596651 | 107.123274 | 0.89443584 | 1.3943E-08 | 1.0348E-07 |
| ENSMUSG00000002983 | Relb | 1969.33286 | 3708.56763 | 1739.23476 | 0.89431887 | 7.8121E-18 | 1.6349E-16 |
| ENSMUSG00000070337 | Gpr179 | 21.3479299 | 44.8812072 | 23.5332773 | 0.8943165 | 0.00054228 | 0.00193376 |
| ENSMUSG00000062480 | Acat3 | 43.7478552 | 88.9094922 | 45.161637 | 0.89388626 | 0.00015012 | 0.0005978 |
| ENSMUSG00000097123 | - | 10.0036523 | 23.3931315 | 13.3894792 | 0.89384658 | 0.00628834 | 0.01764131 |
| ENSMUSG00000072915 | Gm12258 | 387.858074 | 744.950507 | 357.092434 | 0.89367064 | 1.5255E-08 | 1.126E-07 |
| ENSMUSG00000036278 | Macrod1 | 923.543753 | 1746.08675 | 822.542999 | 0.89210653 | 5.3035E-13 | 6.7265E-12 |
| ENSMUSG00000058163 | Gm5431 | 4.78546377 | 14.86715 | 10.0816862 | 0.89195371 | 0.01300858 | 0.03338897 |
| ENSMUSG00000078435 | AU041133 | 143.854356 | 273.144957 | 129.2906 | 0.89181921 | 1.4821E-10 | 1.4321E-09 |
| ENSMUSG00000026725 | Tnn | 996.548225 | 1902.05003 | 905.501808 | 0.89162448 | 1.4489E-09 | 1.2231E-08 |
| ENSMUSG00000083736 | - | 13.1966844 | 28.6352056 | 15.4385213 | 0.891597 | 0.00159966 | 0.0051632 |
| ENSMUSG00000068874 | Selenbp1 | 113.814192 | 237.015708 | 123.201516 | 0.89137517 | 0.00072812 | 0.00252808 |
| ENSMUSG00000036295 | Lrrn3 | 86.4215279 | 173.580172 | 87.158644 | 0.89111274 | 0.00011028 | 0.00045033 |
| ENSMUSG00000060131 | Atp8b4 | 820.224382 | 1549.58715 | 729.362764 | 0.89058748 | 3.5823E-13 | 4.6466E-12 |
| ENSMUSG00000032724 | Abtb2 | 1199.77193 | 2286.36914 | 1086.59721 | 0.89046411 | 1.2392E-09 | 1.0526E-08 |
| ENSMUSG00000055538 | Zcchc24 | 12254.1581 | 23147.3779 | 10893.2198 | 0.89017834 | 5.7801E-13 | 7.2995E-12 |
| ENSMUSG00000052852 | Reep1 | 49.6759351 | 97.4669453 | 47.7910101 | 0.88987395 | 2.3749E-05 | 0.00010864 |
| ENSMUSG00000007097 | Atp1a2 | 32.572071 | 82.1639265 | 49.5918554 | 0.88983357 | 0.00886194 | 0.02383027 |
| ENSMUSG00000036832 | Lpar3 | 61.4103297 | 127.056121 | 65.6457909 | 0.88953721 | 0.00051706 | 0.00185376 |
| ENSMUSG00000026620 | Mark1 | 2293.81246 | 4265.99652 | 1972.18406 | 0.88949619 | 2.822E-54 | 5.6916E-52 |
| ENSMUSG00000024044 | Epb41l3 | 3686.39218 | 6956.88206 | 3270.48987 | 0.88948514 | 2.7989E-13 | 3.6768E-12 |
| ENSMUSG00000022864 | D16Ertd472e | 901.994192 | 1710.29332 | 808.299131 | 0.88915664 | 4.4996E-11 | 4.578E-10 |
| ENSMUSG00000032174 | Icam5 | 35.2657817 | 75.780578 | 40.5147963 | 0.88913452 | 0.00186481 | 0.00593761 |
| ENSMUSG00000055782 | Abcd2 | 84.7908454 | 165.28389 | 80.4930443 | 0.88853396 | 4.0812E-06 | 2.1079E-05 |
| ENSMUSG00000022763 | Aifm3 | 2.76096013 | 8.08223634 | 5.32127622 | 0.88836135 | 0.01242645 | 0.03207893 |
| ENSMUSG00000035279 | Ssc5d | 910.557842 | 1712.00007 | 801.442233 | 0.88819656 | 1.657E-15 | 2.7714E-14 |
| ENSMUSG00000038893 | Fam117a | 168.837805 | 376.426318 | 207.588513 | 0.88768409 | 0.00355552 | 0.0106237 |
| ENSMUSG00000087252 | Gm14379 | 22.5660858 | 48.8323055 | 26.2662197 | 0.88760612 | 0.00215701 | 0.00677732 |
| ENSMUSG00000026956 | Uap1l1 | 1970.17064 | 3679.83024 | 1709.6596 | 0.88743926 | 1.5217E-22 | 4.743E-21 |
| ENSMUSG00000039058 | Ak5 | 283.383545 | 553.970062 | 270.586518 | 0.88737222 | 7.8779E-06 | 3.8958E-05 |
| ENSMUSG00000063765 | Chadl | 29.3430205 | 60.2592746 | 30.9162542 | 0.88686109 | 0.00070545 | 0.00245643 |
| ENSMUSG00000025007 | Aldh18a1 | 15062.6397 | 27998.2434 | 12935.6037 | 0.88574977 | 5.8419E-36 | 4.4843E-34 |
| ENSMUSG00000098434 | - | 3.91323777 | 10.3890577 | 6.4758199 | 0.8857048 | 0.01120014 | 0.0293067 |
| ENSMUSG00000063568 | Jazf1 | 22.1997705 | 46.3448612 | 24.1450908 | 0.88515462 | 0.00133274 | 0.00438605 |
| ENSMUSG00000031289 | Il13ra2 | 454.043831 | 878.671965 | 424.628134 | 0.885109 | 2.1576E-06 | 1.1626E-05 |
| ENSMUSG00000026347 | Tmem163 | 48.4268571 | 97.3155128 | 48.8886556 | 0.88491098 | 0.00011453 | 0.00046646 |
| ENSMUSG00000095648 | Gm2004 | 0 | 4.12739212 | 4.12739212 | 0.88442902 | 0.00849371 | 0.02298509 |
| ENSMUSG00000028469 | Npr2 | 234.218207 | 441.876108 | 207.657901 | 0.88388545 | 1.7843E-12 | 2.1416E-11 |
| ENSMUSG00000095600 | - | 0.98611719 | 5.8853534 | 4.8992362 | 0.88303321 | 0.01397955 | 0.03555288 |
| ENSMUSG00000029648 | Flt1 | 491.890324 | 934.379267 | 442.488944 | 0.88212523 | 5.3746E-09 | 4.2281E-08 |
| ENSMUSG00000029053 | Prkcz | 1.29739983 | 5.92161471 | 4.62421488 | 0.88186605 | 0.01516529 | 0.03820012 |
| ENSMUSG00000059857 | Ntng1 | 1.60868248 | 8.34833649 | 6.73965401 | 0.88080788 | 0.01473594 | 0.03727379 |
| ENSMUSG00000087351 | - | 0.40743019 | 3.99376209 | 3.5863319 | 0.88050219 | 0.01198768 | 0.03111021 |
| ENSMUSG00000038065 | Mturn | 178.639931 | 334.92307 | 156.283139 | 0.88038463 | 4.2788E-12 | 4.9258E-11 |
| ENSMUSG00000066258 | Trim12a | 1164.27269 | 2190.45052 | 1026.17783 | 0.88027495 | 2.6807E-11 | 2.8209E-10 |
| ENSMUSG00000063234 | Gpr84 | 1.97223438 | 9.22984676 | 7.25761237 | 0.88013917 | 0.01518333 | 0.03824088 |
| ENSMUSG00000024084 | Qpct | 965.590483 | 1947.08263 | 981.492144 | 0.87920676 | 0.00031695 | 0.00118616 |
| ENSMUSG00000002980 | Bcam | 5.97322872 | 23.016191 | 17.0429623 | 0.87904651 | 0.01563795 | 0.0392274 |
| ENSMUSG00000068245 | Phf11d | 4287.17285 | 7941.57874 | 3654.40588 | 0.87892297 | 3.4824E-28 | 1.6172E-26 |
| ENSMUSG00000000982 | Ccl3 | 19.5096633 | 48.2457236 | 28.7360603 | 0.8784418 | 0.00926398 | 0.0247762 |
| ENSMUSG00000020653 | Klf11 | 1468.90258 | 2711.6149 | 1242.71232 | 0.87787575 | 2.1517E-41 | 2.27E-39 |
| ENSMUSG00000025069 | Gsto2 | 66.0613953 | 148.974839 | 82.9134434 | 0.87763934 | 0.00497803 | 0.01434286 |
| ENSMUSG00000029446 | Psph | 5920.97829 | 11098.9751 | 5177.99684 | 0.87740552 | 8.0155E-12 | 8.991E-11 |
| ENSMUSG00000045532 | C1ql1 | 19.5890289 | 46.3747844 | 26.7857555 | 0.87735948 | 0.00756818 | 0.02074519 |
| ENSMUSG00000084917 | - | 10.9215583 | 27.8742156 | 16.9526573 | 0.87714958 | 0.01047237 | 0.02761514 |
| ENSMUSG00000033685 | Ucp2 | 334.245321 | 641.469192 | 307.223871 | 0.87663963 | 1.5409E-06 | 8.5029E-06 |
| ENSMUSG00000028456 | Unc13b | 22.943125 | 48.7264753 | 25.7833503 | 0.87662498 | 0.00196034 | 0.00621921 |
| ENSMUSG00000097085 | - | 4.54419404 | 14.2600137 | 9.71581961 | 0.87648765 | 0.01513329 | 0.03814753 |
| ENSMUSG00000085197 | 1700125H03 | 0 | 3.28387243 | 3.28387243 | 0.87642746 | 0.00923271 | 0.02470544 |
| ENSMUSG00000027750 | Postn | 5751.07797 | 11080.2641 | 5329.1861 | 0.87633688 | 3.6879E-06 | 1.9207E-05 |
| ENSMUSG00000037747 | D630013N20 | 8.80749638 | 24.2567217 | 15.4492254 | 0.87565328 | 0.01343409 | 0.03434826 |
| ENSMUSG00000029469 | Ift81 | 1706.22444 | 3144.22561 | 1438.00116 | 0.87520544 | 6.4533E-42 | 6.8432E-40 |
| ENSMUSG00000040907 | Atp1a3 | 3.64253879 | 13.3036962 | 9.66115739 | 0.87492517 | 0.01610009 | 0.04022969 |
| ENSMUSG00000010080 | Epn3 | 22.8054322 | 46.8625014 | 24.0570692 | 0.87488059 | 0.00077512 | 0.0026804 |
| ENSMUSG00000040855 | Reps2 | 14.6939302 | 32.2217193 | 17.5277891 | 0.87417589 | 0.00450313 | 0.01311788 |
| ENSMUSG00000022805 | Cfap91 | 42.2685754 | 84.3474419 | 42.0788666 | 0.87416629 | 0.00036545 | 0.00134936 |

| | | | | | | | |
|---|---|---|---|---|---|---|---|
| ENSMUSG00000025854 | Fam20c | 23.9997863 | 54.5206558 | 30.5208694 | 0.87350205 | 0.00612549 | 0.01723853 |
| ENSMUSG00000022206 | Npr3 | 12283.0072 | 23069.6413 | 10786.6341 | 0.87283467 | 8.9366E-10 | 7.7148E-09 |
| ENSMUSG00000021313 | Ryr2 | 1.89712511 | 8.02388494 | 6.12675983 | 0.87241454 | 0.0161891 | 0.0404016 |
| ENSMUSG00000056481 | Cd248 | 550.021903 | 1352.64242 | 802.620514 | 0.87216264 | 0.0099171 | 0.02632447 |
| ENSMUSG00000030220 | Arhgdib | 6.24063308 | 21.1741727 | 14.9335396 | 0.87162925 | 0.01636827 | 0.04080068 |
| ENSMUSG00000012350 | Ehf | 794.646304 | 1472.51042 | 677.864119 | 0.87153894 | 4.0312E-18 | 8.6374E-17 |
| ENSMUSG00000018339 | Gpx3 | 30.3225484 | 61.2363566 | 30.9138082 | 0.87119504 | 0.00049782 | 0.00179198 |
| ENSMUSG00000047787 | Flrt1 | 55.1864786 | 109.88499 | 54.698511 | 0.87106339 | 0.00018657 | 0.00073051 |
| ENSMUSG00000025658 | Cnksr2 | 12.7141449 | 29.0775016 | 16.3633566 | 0.87081982 | 0.00608773 | 0.01714634 |
| ENSMUSG00000071724 | Smpd5 | 54.776285 | 108.502096 | 53.7258113 | 0.87069274 | 0.0001409 | 0.00056459 |
| ENSMUSG00000068742 | Cry2 | 782.176516 | 1440.16467 | 657.988151 | 0.87066362 | 5.2406E-29 | 2.5608E-27 |
| ENSMUSG00000050164 | Mchr1 | 1.77484293 | 7.00448454 | 5.2296416 | 0.87013192 | 0.016703 | 0.04154444 |
| ENSMUSG00000034664 | Itga2b | 197.843099 | 369.217836 | 171.374737 | 0.86991918 | 6.2055E-10 | 5.4813E-09 |
| ENSMUSG00000049526 | Tmem202 | 13.4968126 | 28.2602487 | 14.7634361 | 0.86989041 | 0.0022592 | 0.00706433 |
| ENSMUSG00000004098 | Col5a3 | 901.821952 | 1710.92776 | 809.105807 | 0.86956559 | 3.3923E-07 | 2.0702E-06 |
| ENSMUSG00000029320 | 1700016H13 | 48.3071511 | 95.8862824 | 47.5791313 | 0.86955726 | 0.00028122 | 0.00106308 |
| ENSMUSG00000020953 | Coch | 2.0422473 | 8.08724886 | 6.04500157 | 0.86932336 | 0.01680939 | 0.04179896 |
| ENSMUSG00000029641 | Rasl11a | 836.359082 | 1721.70922 | 885.350136 | 0.86930128 | 0.00120231 | 0.00399836 |
| ENSMUSG00000081142 | - | 4.91709861 | 12.5977542 | 7.68065564 | 0.86898849 | 0.01209462 | 0.03136794 |
| ENSMUSG00000035498 | Cdcp1 | 65.9011056 | 128.854807 | 62.9537017 | 0.86878638 | 0.0001282 | 0.00051755 |
| ENSMUSG00000028020 | Glrb | 520.677977 | 1080.37064 | 559.692662 | 0.8673568 | 0.00162753 | 0.00524542 |
| ENSMUSG00000066842 | Hmcn1 | 1.9638434 | 10.1976915 | 8.23384814 | 0.86711227 | 0.01607156 | 0.04018274 |
| ENSMUSG00000031290 | Lrch2 | 1000.81623 | 1882.01225 | 881.196018 | 0.86702704 | 2.5037E-08 | 1.7952E-07 |
| ENSMUSG00000087750 | - | 3.21226859 | 9.69948626 | 6.48721767 | 0.86702381 | 0.01614641 | 0.0403224 |
| ENSMUSG00000097745 | AI115009 | 2.09451656 | 7.97270668 | 5.87819012 | 0.86599253 | 0.01725567 | 0.04273308 |
| ENSMUSG00000027667 | Zfp639 | 2097.28156 | 3865.28361 | 1768.00205 | 0.86596334 | 5.2401E-19 | 1.2195E-17 |
| ENSMUSG00000010461 | Eya4 | 1121.92715 | 2079.42527 | 957.498125 | 0.86551889 | 1.9274E-13 | 2.5664E-12 |
| ENSMUSG00000005611 | Irag1 | 10.7731415 | 28.7488586 | 17.9757171 | 0.8655025 | 0.0135158 | 0.03451857 |
| ENSMUSG00000049939 | Lrrc4 | 22.0447644 | 47.7416605 | 25.6968961 | 0.86522898 | 0.00386418 | 0.01145611 |
| ENSMUSG00000083353 | - | 4.69866889 | 13.9594152 | 9.26074628 | 0.86510706 | 0.01569741 | 0.03934776 |
| ENSMUSG00000055923 | Aasdh | 636.522165 | 1167.77633 | 531.25417 | 0.86476779 | 2.1574E-30 | 1.1772E-28 |
| ENSMUSG00000066867 | Oas1e | 8.15124554 | 20.9296629 | 12.7784173 | 0.86447939 | 0.01211942 | 0.03142829 |
| ENSMUSG00000079580 | Tmem217 | 11.8670919 | 27.6519223 | 15.7848305 | 0.86436472 | 0.00834865 | 0.02264317 |
| ENSMUSG00000039431 | Mtmr7 | 1972.58394 | 3623.11156 | 1650.52763 | 0.86429 | 5.5967E-24 | 1.9094E-22 |
| ENSMUSG00000097120 | - | 0.40743019 | 4.13529994 | 3.72786975 | 0.86379317 | 0.01330376 | 0.03405736 |
| ENSMUSG00000042501 | Cpa6 | 31.4847099 | 78.9809835 | 47.4962736 | 0.86360762 | 0.01206536 | 0.03129599 |
| ENSMUSG00000024379 | Tslp | 379.915444 | 865.851272 | 485.935828 | 0.86343966 | 0.00719188 | 0.01985391 |
| ENSMUSG00000004633 | Chn2 | 17.0464985 | 36.4349597 | 19.3884612 | 0.86324335 | 0.00379599 | 0.01127344 |
| ENSMUSG00000085675 | Gm12022 | 2.85710767 | 10.166029 | 7.30892131 | 0.86305223 | 0.01761596 | 0.0435153 |
| ENSMUSG00000073802 | Cdkn2b | 2339.25429 | 4288.04004 | 1948.78574 | 0.86291837 | 1.2289E-24 | 4.4276E-23 |
| ENSMUSG00000087692 | - | 5.64090781 | 14.0435005 | 8.40259273 | 0.86260032 | 0.01048149 | 0.02763363 |
| ENSMUSG00000023044 | Csad | 833.485057 | 1521.29934 | 687.814288 | 0.86251593 | 3.2362E-49 | 5.2422E-47 |
| ENSMUSG00000020432 | Tcn2 | 1463.13197 | 2722.88347 | 1259.75151 | 0.86241581 | 4.623E-10 | 4.1694E-09 |
| ENSMUSG00000048721 | Fndc9 | 0.31128264 | 4.78231884 | 4.4710362 | 0.8620148 | 0.0124351 | 0.03209723 |
| ENSMUSG00000082361 | Btc | 229.136972 | 432.862526 | 203.725554 | 0.86135184 | 6.0205E-07 | 3.5375E-06 |
| ENSMUSG00000027824 | Vmn2r1 | 3.04610814 | 9.40779632 | 6.36168819 | 0.86129271 | 0.01694045 | 0.04206892 |
| ENSMUSG00000097714 | - | 0.71871283 | 5.47377162 | 4.75505879 | 0.86097267 | 0.0148325 | 0.03749959 |
| ENSMUSG00000032348 | Gsta4 | 237.191574 | 456.583271 | 219.391697 | 0.86062354 | 3.4671E-05 | 0.00015438 |
| ENSMUSG00000073679 | - | 2.68255623 | 11.2911382 | 8.60858197 | 0.86054788 | 0.01772907 | 0.0437842 |
| ENSMUSG00000096956 | Snhg18 | 71.5185206 | 186.689976 | 115.171456 | 0.86053968 | 0.01381711 | 0.03519634 |
| ENSMUSG00000034173 | Zbed5 | 206.547206 | 382.342899 | 175.795693 | 0.86046607 | 1.5675E-11 | 1.6936E-10 |
| ENSMUSG00000059554 | Ccdc28a | 636.417519 | 1161.47329 | 525.055775 | 0.86037846 | 1.4179E-30 | 7.8411E-29 |
| ENSMUSG00000024295 | Cyp4f41-ps | 7.42414172 | 17.4234302 | 9.99928852 | 0.8603712 | 0.00786322 | 0.02145955 |
| ENSMUSG00000021775 | Nr1d2 | 7689.02494 | 14001.2364 | 6312.21142 | 0.86011613 | 2.1976E-60 | 6.0278E-58 |
| ENSMUSG00000067724 | Gbx1 | 1.02999547 | 4.85288729 | 3.82289181 | 0.86005955 | 0.01750807 | 0.04327478 |
| ENSMUSG00000004837 | Grap | 1.0561301 | 4.87964639 | 3.82351629 | 0.85992437 | 0.01747938 | 0.04321946 |
| ENSMUSG00000038916 | Soga3 | 233.332903 | 429.998102 | 196.665198 | 0.8595014 | 1.327E-15 | 2.2523E-14 |
| ENSMUSG00000022803 | Popdc2 | 10.3233259 | 23.6197319 | 13.2964059 | 0.85918299 | 0.00916696 | 0.02455183 |
| ENSMUSG00000090936 | - | 4.85218206 | 17.2509984 | 12.3988163 | 0.85913711 | 0.01811311 | 0.04460413 |
| ENSMUSG00000066233 | Tmem42 | 187.582767 | 346.879341 | 159.296574 | 0.85904001 | 2.0205E-10 | 1.9181E-09 |
| ENSMUSG00000054619 | Mettl7a1 | 336.361185 | 617.986913 | 281.625728 | 0.8587035 | 3.7105E-18 | 8.0013E-17 |
| ENSMUSG00000035835 | Plppr3 | 96.0021169 | 180.523165 | 84.5210477 | 0.85842391 | 4.3007E-08 | 2.983E-07 |
| ENSMUSG00000006218 | Fam131c | 32.0684933 | 68.5924787 | 36.5239854 | 0.85736937 | 0.00435883 | 0.01275533 |
| ENSMUSG00000091119 | Ccdc152 | 4.86812847 | 12.7049242 | 7.83680028 | 0.85701588 | 0.01525551 | 0.03840855 |
| ENSMUSG00000071793 | 2610005L07I | 135.96974 | 252.909167 | 116.939426 | 0.85623353 | 9.3345E-11 | 9.2012E-10 |
| ENSMUSG00000006204 | Cdcp3 | 7.16512834 | 19.3653266 | 12.2001982 | 0.85615539 | 0.01523465 | 0.03836075 |
| ENSMUSG00000079419 | Ms4a6c | 0.40743019 | 4.41169927 | 4.00426909 | 0.85609756 | 0.01369183 | 0.03491179 |
| ENSMUSG00000053559 | Smagp | 8.29223304 | 21.3734437 | 13.0812107 | 0.85604144 | 0.01373608 | 0.0350116 |
| ENSMUSG00000039481 | Nrtn | 10.9593786 | 23.8979434 | 12.9385649 | 0.85509625 | 0.00582877 | 0.01650054 |
| ENSMUSG00000007035 | Msh5 | 97.8770548 | 181.373296 | 83.4962413 | 0.85499515 | 1.0991E-08 | 8.2973E-08 |
| ENSMUSG00000096847 | Tmem151b | 14.6719303 | 31.8361317 | 17.1642015 | 0.85446599 | 0.0049837 | 0.01435195 |
| ENSMUSG00000090558 | - | 2.11226021 | 7.91981182 | 5.80755162 | 0.85436033 | 0.0188 | 0.046075 |
| ENSMUSG00000035900 | Gramd4 | 1338.08029 | 2474.75327 | 1136.67298 | 0.85416328 | 5.3418E-10 | 4.7675E-09 |
| ENSMUSG00000049128 | Ivl | 2.49026114 | 11.2573249 | 8.76706371 | 0.85343116 | 0.01847293 | 0.04539257 |
| ENSMUSG00000086765 | Gm11827 | 28.7895064 | 57.3422218 | 28.5527154 | 0.8524157 | 0.00113291 | 0.00378715 |

| | | | | | | | |
|---|---|---|---|---|---|---|---|
| ENSMUSG00000030256 | Bhlhe41 | 269.947392 | 523.779915 | 253.832522 | 0.85210131 | 0.00015588 | 0.00061849 |
| ENSMUSG00000068252 | Apol7b | 1.15227765 | 5.75032553 | 4.59804788 | 0.85196841 | 0.0178537 | 0.04404969 |
| ENSMUSG00000061111 | Mcrip1 | 908.422403 | 1658.12923 | 749.706829 | 0.85164029 | 1.2031E-17 | 2.4652E-16 |
| ENSMUSG00000049191 | Rtl5 | 7.19126298 | 25.5831954 | 18.3919324 | 0.85151114 | 0.01921241 | 0.04690692 |
| ENSMUSG00000022018 | Rgcc | 71.2348654 | 181.225891 | 109.991026 | 0.85146833 | 0.01429595 | 0.03630371 |
| ENSMUSG00000041592 | Sdk2 | 1.36741275 | 6.47719519 | 5.10978245 | 0.85093826 | 0.01852758 | 0.04548876 |
| ENSMUSG00000097576 | - | 23.3412026 | 46.3781685 | 23.0369659 | 0.85039461 | 0.00052147 | 0.00186763 |
| ENSMUSG00000061524 | Zic2 | 4.09714188 | 11.5492228 | 7.4520809 | 0.84969475 | 0.01785216 | 0.04404969 |
| ENSMUSG00000000359 | Rem1 | 68.2050081 | 127.722868 | 59.5178596 | 0.84966392 | 7.1911E-07 | 4.1721E-06 |
| ENSMUSG00000042671 | Rgs8 | 3.53194222 | 9.8470635 | 6.31512128 | 0.84958772 | 0.01724651 | 0.04271554 |
| ENSMUSG00000081670 | - | 9.55414558 | 21.5656848 | 12.0115392 | 0.84912874 | 0.00744993 | 0.02046195 |
| ENSMUSG00000040272 | Accs | 496.224737 | 901.078376 | 404.853639 | 0.8489006 | 1.1467E-22 | 3.618E-21 |
| ENSMUSG00000036944 | Tmem71 | 12.6702666 | 32.2663453 | 19.5960786 | 0.84886347 | 0.0145921 | 0.03695086 |
| ENSMUSG00000054545 | Ugt1a6a | 378.525774 | 689.045372 | 310.519598 | 0.84848522 | 2.7397E-19 | 6.5459E-18 |
| ENSMUSG00000000204 | Slfn4 | 83.6461187 | 173.086101 | 89.4399819 | 0.84791746 | 0.00286303 | 0.00872827 |
| ENSMUSG00000029378 | Areg | 540.046688 | 1182.85634 | 642.809655 | 0.84790574 | 0.00664015 | 0.01850843 |
| ENSMUSG00000087187 | - | 28.4716345 | 57.0807948 | 28.6091603 | 0.84770138 | 0.00131252 | 0.00432711 |
| ENSMUSG00000086568 | - | 2.08612558 | 9.24913509 | 7.16300951 | 0.84694016 | 0.01921073 | 0.04690692 |
| ENSMUSG00000055629 | B4galnt4 | 396.903136 | 723.340256 | 326.43712 | 0.84666617 | 9.8988E-18 | 2.0446E-16 |
| ENSMUSG00000090659 | Zfp493 | 556.697716 | 1014.88629 | 458.188575 | 0.84649233 | 1.6549E-14 | 2.4868E-13 |
| ENSMUSG00000085582 | 3110099E03 | 0.31128264 | 4.46403142 | 4.15274878 | 0.84648146 | 0.01404388 | 0.03570766 |
| ENSMUSG00000078763 | Slfn1 | 0.40743019 | 5.26502191 | 4.85759172 | 0.84627691 | 0.01386795 | 0.0353101 |
| ENSMUSG00000026610 | Esrrg | 0.71871283 | 4.62028465 | 3.90157182 | 0.84592483 | 0.0173187 | 0.04286335 |
| ENSMUSG00000031255 | Sytl4 | 4.96607326 | 13.1257413 | 8.15966807 | 0.84591053 | 0.01537384 | 0.03865445 |
| ENSMUSG00000056313 | Tcim | 257.776569 | 473.978757 | 216.202189 | 0.84573602 | 4.7366E-10 | 4.2606E-09 |
| ENSMUSG00000026313 | Hdac4 | 455.789109 | 829.660494 | 373.871385 | 0.84531114 | 2.6024E-18 | 5.7076E-17 |
| ENSMUSG00000096932 | - | 25.4273281 | 49.7182172 | 24.290889 | 0.84501826 | 0.00034326 | 0.00127532 |
| ENSMUSG00000042599 | Kdm7a | 791.795341 | 1475.38356 | 683.588223 | 0.84495944 | 6.6733E-07 | 3.889E-06 |
| ENSMUSG00000037405 | Icam1 | 17.0675367 | 35.0362328 | 17.968696 | 0.84491614 | 0.00371881 | 0.01105379 |
| ENSMUSG00000039899 | Fgl2 | 28.782386 | 63.716415 | 34.934029 | 0.84481547 | 0.00750997 | 0.02060485 |
| ENSMUSG00000050463 | Krt78 | 2.20840775 | 6.74706116 | 4.53865341 | 0.8445097 | 0.01974015 | 0.04804144 |
| ENSMUSG00000084844 | - | 190.250221 | 348.158974 | 157.908753 | 0.84428385 | 2.0325E-10 | 1.9286E-09 |
| ENSMUSG00000096858 | Olfr805 | 5.92605582 | 14.7497827 | 8.82372686 | 0.84382978 | 0.01291508 | 0.03316553 |
| ENSMUSG00000035640 | Cbarp | 417.2506 | 780.295297 | 363.044697 | 0.84365325 | 2.7241E-06 | 1.4484E-05 |
| ENSMUSG00000032595 | Cdhr4 | 0.40743019 | 3.98691678 | 3.57948659 | 0.84353036 | 0.01527803 | 0.03845585 |
| ENSMUSG00000022364 | Tbc1d31 | 4315.51369 | 7770.62792 | 3455.11423 | 0.84302875 | 4.7759E-49 | 7.6163E-47 |
| ENSMUSG00000030553 | Pgpep1l | 20.7894411 | 47.5363272 | 26.7468861 | 0.84297754 | 0.01055396 | 0.02780332 |
| ENSMUSG00000007812 | Zfp655 | 3452.21516 | 6207.73751 | 2755.52235 | 0.84292963 | 8.8547E-78 | 4.6707E-75 |
| ENSMUSG00000023067 | Cdkn1a | 691.332868 | 1257.03021 | 565.697342 | 0.84289807 | 1.1296E-14 | 1.729E-13 |
| ENSMUSG00000018012 | Rac3 | 504.666262 | 937.958988 | 433.292727 | 0.84255923 | 6.9677E-07 | 4.0514E-06 |
| ENSMUSG00000044276 | 4933427E11 | 9.87711383 | 21.7078238 | 11.8307099 | 0.84253622 | 0.00764639 | 0.02093178 |
| ENSMUSG00000037922 | Bank1 | 2.61254332 | 10.2348718 | 7.62232845 | 0.84232623 | 0.02048345 | 0.04956893 |
| ENSMUSG00000002228 | Ppm1j | 444.300751 | 970.556762 | 526.256011 | 0.84198787 | 0.00722393 | 0.01992664 |
| ENSMUSG00000097820 | - | 11.8223735 | 28.4784849 | 16.6561114 | 0.84178985 | 0.01434782 | 0.03641294 |
| ENSMUSG00000086447 | - | 1.15227765 | 8.15500894 | 7.00273129 | 0.84175609 | 0.01684929 | 0.041883 |
| ENSMUSG00000030748 | Il4ra | 1486.48861 | 2845.92743 | 1359.43883 | 0.8413973 | 0.00012309 | 0.00049829 |
| ENSMUSG00000059327 | Eda | 213.845649 | 463.633461 | 249.787811 | 0.841346 | 0.00676298 | 0.01880616 |
| ENSMUSG00000058952 | Cfi | 4.57638672 | 12.987268 | 8.41088133 | 0.84130965 | 0.0183188 | 0.04505671 |
| ENSMUSG00000009633 | G0s2 | 589.993409 | 1105.21508 | 515.221673 | 0.84015782 | 7.8087E-06 | 3.8643E-05 |
| ENSMUSG00000097645 | - | 7.77420628 | 20.3953664 | 12.6211601 | 0.84011521 | 0.01717764 | 0.04257058 |
| ENSMUSG00000022203 | Efs | 0.31128264 | 3.85590921 | 3.54462656 | 0.83988267 | 0.01538729 | 0.03867881 |
| ENSMUSG00000017723 | Wfdc2 | 4553.07362 | 8271.83177 | 3718.75815 | 0.83933858 | 3.3885E-13 | 4.4147E-12 |
| ENSMUSG00000006724 | Cyp27b1 | 5.80706826 | 14.9361187 | 9.12905039 | 0.83909896 | 0.0165128 | 0.04112608 |
| ENSMUSG00000086296 | - | 3.26963422 | 9.54838655 | 6.27875233 | 0.83877738 | 0.01903515 | 0.04654042 |
| ENSMUSG00000005397 | Nid1 | 2094.99927 | 4376.99676 | 2281.99749 | 0.83830267 | 0.00443068 | 0.01293245 |
| ENSMUSG00000061816 | Myl1 | 1.94100339 | 8.01498258 | 6.07397919 | 0.83819479 | 0.02063173 | 0.04990428 |
| ENSMUSG00000062175 | Tgif2 | 371.474294 | 672.274284 | 300.79999 | 0.83807179 | 1.4557E-18 | 3.2729E-17 |
| ENSMUSG00000044103 | Il36g | 47.6336878 | 103.098442 | 55.4647542 | 0.83765862 | 0.00729316 | 0.0200823 |
| ENSMUSG00000043614 | Vps37d | 155.081361 | 284.377586 | 129.296224 | 0.83720229 | 1.1998E-07 | 7.7665E-07 |
| ENSMUSG00000036186 | Dipk1b | 138.977411 | 261.740495 | 122.763085 | 0.83652041 | 3.721E-05 | 0.00016466 |
| ENSMUSG00000022620 | Arsa | 1975.12178 | 3539.33895 | 1564.21716 | 0.8361236 | 1.0618E-45 | 1.3484E-43 |
| ENSMUSG00000040327 | Cul9 | 700.38457 | 1264.71468 | 564.330112 | 0.83597271 | 1.3424E-18 | 3.0314E-17 |
| ENSMUSG00000078598 | Skint5 | 29.4914373 | 61.6253839 | 32.1339466 | 0.83574244 | 0.00528491 | 0.01511487 |
| ENSMUSG00000040710 | St8sia4 | 92.0110065 | 169.773978 | 77.7629715 | 0.83382787 | 2.5183E-07 | 1.5647E-06 |
| ENSMUSG00000020395 | Itk | 0 | 2.95840866 | 2.95840866 | 0.83326263 | 0.01246302 | 0.03216122 |
| ENSMUSG00000092564 | - | 146.940739 | 268.572171 | 121.631432 | 0.8332602 | 4.9544E-08 | 3.3963E-07 |
| ENSMUSG00000040812 | Agbl2 | 39.075321 | 76.2216281 | 37.1463071 | 0.83299219 | 0.00070615 | 0.00245843 |
| ENSMUSG00000097373 | - | 9.12387538 | 21.4192419 | 12.2953666 | 0.8326579 | 0.0133762 | 0.03422151 |
| ENSMUSG00000054708 | Ankrd24 | 397.657249 | 720.646045 | 322.988796 | 0.83235797 | 3.5484E-11 | 3.6793E-10 |
| ENSMUSG00000041390 | Mdfic | 6094.66673 | 10919.2173 | 4824.55054 | 0.83191191 | 8.7287E-28 | 3.9293E-26 |
| ENSMUSG00000040795 | Iqcc | 678.206369 | 1223.15588 | 544.949506 | 0.83188176 | 1.7773E-16 | 3.2587E-15 |
| ENSMUSG00000097017 | - | 4.28943697 | 11.2981162 | 7.00867925 | 0.83178344 | 0.02009344 | 0.0487628 |
| ENSMUSG00000074934 | Grem1 | 6577.99775 | 14293.4583 | 7715.46057 | 0.831235 | 0.00837737 | 0.02270909 |
| ENSMUSG00000066026 | Dhrs3 | 159.392677 | 291.65822 | 132.265543 | 0.83100022 | 3.1018E-07 | 1.9077E-06 |
| ENSMUSG00000051037 | Zfp455 | 216.256767 | 392.112889 | 175.856121 | 0.83072653 | 4.9301E-10 | 4.4212E-09 |

| | | | | | | | |
|---|---|---|---|---|---|---|---|
| ENSMUSG00000047759 | Hs3st3a1 | 26.7845482 | 51.2109504 | 24.4264022 | 0.83057263 | 0.00037613 | 0.00138519 |
| ENSMUSG00000028300 | C9orf72 | 1636.22417 | 2923.50651 | 1287.28234 | 0.83028604 | 7.3644E-37 | 5.9412E-35 |
| ENSMUSG00000042200 | Cdrt4 | 10.5747884 | 22.4023185 | 11.8275301 | 0.82980453 | 0.00531487 | 0.01518787 |
| ENSMUSG00000032456 | Nmnat3 | 283.179786 | 508.68434 | 225.504554 | 0.82916345 | 4.9795E-16 | 8.7704E-15 |
| ENSMUSG00000024521 | Pmaip1 | 394.681584 | 798.944919 | 404.263334 | 0.82894402 | 0.00313396 | 0.00948116 |
| ENSMUSG00000086881 | - | 11.3145395 | 24.5430398 | 13.2285003 | 0.82894225 | 0.00886279 | 0.02383027 |
| ENSMUSG00000032577 | Mapkapk3 | 248.831071 | 451.384382 | 202.553312 | 0.82835111 | 1.2557E-09 | 1.0661E-08 |
| ENSMUSG00000028661 | Epha8 | 0.33741727 | 3.233766 | 2.89634872 | 0.82829852 | 0.01777199 | 0.04387439 |
| ENSMUSG00000032578 | Cish | 153.018141 | 278.980179 | 125.962038 | 0.82823235 | 2.4323E-08 | 1.749E-07 |
| ENSMUSG00000006720 | Zfp184 | 361.608901 | 647.705116 | 286.096216 | 0.82792142 | 1.4101E-16 | 2.6181E-15 |
| ENSMUSG00000020333 | Acsl6 | 207.080522 | 385.727188 | 178.646666 | 0.82768281 | 1.8743E-05 | 8.7216E-05 |
| ENSMUSG00000055692 | Tmem191c | 12.946062 | 27.8076845 | 14.8616225 | 0.82766789 | 0.00709004 | 0.01959381 |
| ENSMUSG00000028278 | Rragd | 56.8909998 | 124.563233 | 67.6722328 | 0.82764669 | 0.00963143 | 0.02563562 |
| ENSMUSG00000015377 | Dennd6b | 278.215601 | 502.969436 | 224.753835 | 0.82743194 | 2.4901E-13 | 3.2832E-12 |
| ENSMUSG00000078653 | Cntd1 | 104.751286 | 191.848559 | 87.0972732 | 0.82733118 | 1.1103E-07 | 7.2193E-07 |
| ENSMUSG00000006418 | Rnf114 | 4807.93536 | 8564.04257 | 3756.10721 | 0.8271875 | 8.832E-40 | 8.6934E-38 |
| ENSMUSG00000060402 | Chst8 | 13.0464658 | 27.356295 | 14.3098292 | 0.82677232 | 0.00626525 | 0.01758373 |
| ENSMUSG00000078117 | - | 65.8943947 | 124.446235 | 58.5518407 | 0.82663442 | 0.00015532 | 0.0006166 |
| ENSMUSG00000060791 | Gmfg | 8.73748346 | 19.7103987 | 10.9729153 | 0.82608081 | 0.01294792 | 0.03324155 |
| ENSMUSG00000013353 | 4931406B18 | 0 | 3.52919403 | 3.52919403 | 0.82561704 | 0.01276539 | 0.0328221 |
| ENSMUSG00000079722 | Ttll2 | 0 | 2.7863089 | 2.7863089 | 0.82523308 | 0.01322821 | 0.0338766 |
| ENSMUSG00000056124 | B4galt6 | 1174.40353 | 2094.15669 | 919.753154 | 0.82464689 | 1.8545E-26 | 7.6008E-25 |
| ENSMUSG00000015837 | Sqstm1 | 29788.4962 | 53436.7206 | 23648.2243 | 0.82458958 | 9.6103E-15 | 1.4798E-13 |
| ENSMUSG00000048058 | Ldlrad3 | 833.666916 | 1507.99807 | 674.331158 | 0.82454585 | 8.6748E-10 | 7.4951E-09 |
| ENSMUSG00000025592 | Dach2 | 0.33741727 | 5.93294197 | 5.5955247 | 0.82422578 | 0.01508728 | 0.03805021 |
| ENSMUSG00000054822 | - | 0.31128264 | 4.3217458 | 4.01046316 | 0.82417073 | 0.0164224 | 0.04092569 |
| ENSMUSG00000032373 | Car12 | 419.430971 | 751.284029 | 331.853059 | 0.82410163 | 6.7179E-18 | 1.416E-16 |
| ENSMUSG00000056144 | Trim34a | 1023.71602 | 1865.26593 | 841.549913 | 0.82389917 | 1.1766E-07 | 7.6287E-07 |
| ENSMUSG00000054793 | Cadm4 | 20.5075877 | 49.2652947 | 28.757707 | 0.82354531 | 0.0169566 | 0.04209083 |
| ENSMUSG00000030247 | Kcnj8 | 5.38189443 | 13.4302754 | 8.04838099 | 0.82288752 | 0.01735022 | 0.0429362 |
| ENSMUSG00000044167 | Foxo1 | 3636.18997 | 6457.40921 | 2821.21924 | 0.82211525 | 1.3126E-40 | 1.3368E-38 |
| ENSMUSG00000034912 | Mdga2 | 0 | 2.60814548 | 2.60814548 | 0.82198625 | 0.01363242 | 0.03478618 |
| ENSMUSG00000035295 | Wdr38 | 5.01324616 | 12.8124959 | 7.79924978 | 0.82189885 | 0.01989732 | 0.04836674 |
| ENSMUSG00000055471 | Alk | 7.59636022 | 18.3827475 | 10.7863872 | 0.82186193 | 0.01570636 | 0.0393654 |
| ENSMUSG00000040584 | Abcb1a | 630.295515 | 1141.90529 | 511.609779 | 0.82181119 | 8.3232E-09 | 6.3842E-08 |
| ENSMUSG00000071181 | 3830408C21 | 14.4745388 | 29.3606476 | 14.8861088 | 0.82124358 | 0.00439295 | 0.01284418 |
| ENSMUSG00000053040 | Aph1c | 69.1765546 | 133.516433 | 64.3398786 | 0.82117842 | 0.00078933 | 0.00272387 |
| ENSMUSG00000064307 | Lrrc51 | 142.61209 | 261.921831 | 119.309742 | 0.82094425 | 6.8522E-06 | 3.414E-05 |
| ENSMUSG00000046005 | - | 0 | 3.04666794 | 3.04666794 | 0.81971684 | 0.01350625 | 0.03450546 |
| ENSMUSG00000032311 | Nrg4 | 37.3891963 | 71.945069 | 34.5558727 | 0.81936186 | 0.00061076 | 0.0021548 |
| ENSMUSG00000036452 | Arhgap26 | 480.187914 | 857.293437 | 377.105523 | 0.81901272 | 3.1303E-16 | 5.6242E-15 |
| ENSMUSG00000025991 | Cps1 | 0 | 2.89700603 | 2.89700603 | 0.81854378 | 0.0137039 | 0.03493823 |
| ENSMUSG00000032420 | Nt5e | 1602.51319 | 2985.9773 | 1383.46411 | 0.81773588 | 7.6542E-05 | 0.00032109 |
| ENSMUSG00000031937 | Vstm5 | 31.9119944 | 68.120275 | 36.2082806 | 0.81764401 | 0.008433 | 0.02283581 |
| ENSMUSG00000032417 | Rwdd2a | 86.8662471 | 165.786572 | 78.9203245 | 0.8175499 | 0.00055427 | 0.00197171 |
| ENSMUSG00000032514 | Ttc21a | 9.12387538 | 19.2616761 | 10.1378007 | 0.81711215 | 0.00837438 | 0.02270397 |
| ENSMUSG00000066607 | Insyn1 | 431.848109 | 900.131503 | 468.283394 | 0.81652923 | 0.00712344 | 0.01967819 |
| ENSMUSG00000021872 | Rnase10 | 13.732986 | 28.6075473 | 14.8745614 | 0.81604128 | 0.00887443 | 0.02385845 |
| ENSMUSG00000047407 | Tgif1 | 4657.56791 | 8226.33327 | 3568.76537 | 0.81575562 | 4.1549E-45 | 5.0577E-43 |
| ENSMUSG00000066892 | Fbxl12 | 681.852891 | 1204.64913 | 522.796239 | 0.81525513 | 1.2514E-33 | 8.2251E-32 |
| ENSMUSG00000039099 | Wdr93 | 25.9705278 | 52.5573186 | 26.5867908 | 0.81500727 | 0.00404957 | 0.0119472 |
| ENSMUSG00000084805 | Gm12273 | 0 | 3.11124641 | 3.11124641 | 0.81491637 | 0.01390136 | 0.03537589 |
| ENSMUSG00000032012 | Nectin1 | 130.218889 | 242.572401 | 112.353512 | 0.81480624 | 0.00010253 | 0.0004215 |
| ENSMUSG00000057596 | Trim30d | 3462.40094 | 6496.20492 | 3033.80398 | 0.81440194 | 0.00021433 | 0.00083144 |
| ENSMUSG00000029859 | Epha1 | 140.587895 | 268.708937 | 128.121043 | 0.81435368 | 0.00078369 | 0.00270733 |
| ENSMUSG00000031398 | Plxna3 | 784.796762 | 1418.16936 | 633.372595 | 0.81393873 | 8.5601E-08 | 5.6659E-07 |
| ENSMUSG00000040640 | Erc2 | 289.068401 | 521.852554 | 232.784153 | 0.81310592 | 5.0792E-07 | 3.0147E-06 |
| ENSMUSG00000042293 | Gm5617 | 124.342461 | 224.50458 | 100.162119 | 0.81294999 | 5.1006E-07 | 3.0239E-06 |
| ENSMUSG00000020175 | Rab36 | 73.2559529 | 134.51988 | 61.2639267 | 0.81220739 | 1.4663E-05 | 6.9439E-05 |
| ENSMUSG00000053475 | Tnfaip6 | 652.978017 | 1173.08435 | 520.106331 | 0.81194341 | 1.0906E-08 | 8.2393E-08 |
| ENSMUSG00000024055 | Cyp4f13 | 468.663766 | 839.766931 | 371.103165 | 0.81164353 | 7.3751E-10 | 6.4398E-09 |
| ENSMUSG00000042406 | Atf4 | 27822.0587 | 49094.438 | 21272.3793 | 0.81161194 | 1.1532E-30 | 6.4116E-29 |
| ENSMUSG00000034255 | Arhgap27 | 216.640073 | 396.361995 | 179.721922 | 0.81131145 | 7.3045E-06 | 3.6253E-05 |
| ENSMUSG00000030510 | Cers3 | 43.0786482 | 79.5742736 | 36.4956254 | 0.81115845 | 4.1685E-05 | 0.00018315 |
| ENSMUSG00000044499 | Hs3st5 | 10.360615 | 22.3789502 | 12.0183353 | 0.81074118 | 0.01039095 | 0.02744065 |
| ENSMUSG00000039109 | F13a1 | 1055.06268 | 1877.03122 | 821.96854 | 0.80993672 | 3.3551E-12 | 3.8973E-11 |
| ENSMUSG00000074354 | Arhgap20os | 0.31128264 | 3.45041082 | 3.13912818 | 0.80957442 | 0.01940199 | 0.04733045 |
| ENSMUSG00000023467 | Tulp2 | 20.35162 | 42.6179928 | 22.2663728 | 0.80926643 | 0.00878453 | 0.02364766 |
| ENSMUSG00000090891 | D6Ertd527e | 19.0956439 | 39.1968529 | 20.101209 | 0.80830221 | 0.00561648 | 0.01595887 |
| ENSMUSG00000047084 | Ngrn | 1848.7917 | 3265.51508 | 1416.72338 | 0.80797677 | 1.5967E-18 | 3.5665E-17 |
| ENSMUSG00000036377 | Cracd | 38.118633 | 77.6677226 | 39.5490896 | 0.80794413 | 0.00632967 | 0.01774273 |
| ENSMUSG00000024899 | Papss2 | 141.528763 | 264.057823 | 122.52906 | 0.80751544 | 0.00024709 | 0.00094691 |
| ENSMUSG00000037031 | Tspan15 | 298.69175 | 543.279998 | 244.588248 | 0.80710133 | 6.9016E-06 | 3.4371E-05 |
| ENSMUSG00000042826 | Fgf11 | 15.5433162 | 31.6851878 | 16.1418716 | 0.80688744 | 0.00812115 | 0.02207853 |
| ENSMUSG00000097257 | - | 5.25451589 | 12.5953116 | 7.34079574 | 0.8067158 | 0.01998713 | 0.04854489 |

| | | | | | | | |
|---|---|---|---|---|---|---|---|
| ENSMUSG00000041797 | Abca9 | 23.9385949 | 48.5872898 | 24.6486949 | 0.80565966 | 0.00590773 | 0.01668503 |
| ENSMUSG00000023755 | Rhebl1 | 360.583161 | 636.861372 | 276.278211 | 0.80557406 | 3.5985E-14 | 5.2243E-13 |
| ENSMUSG00000028807 | Zbtb8a | 34.8051205 | 67.0140448 | 32.2089242 | 0.80523508 | 0.00181229 | 0.00578831 |
| ENSMUSG00000024778 | Fas | 2098.44533 | 3694.5273 | 1596.08196 | 0.80480336 | 1.5354E-20 | 4.034E-19 |
| ENSMUSG00000063286 | Gm8995 | 255.300722 | 579.853787 | 324.553065 | 0.80346653 | 0.01695746 | 0.04209083 |
| ENSMUSG00000000253 | Gmpr | 181.103936 | 327.885131 | 146.781195 | 0.80336482 | 4.0984E-06 | 2.1157E-05 |
| ENSMUSG00000050578 | Mmp13 | 39627.3007 | 71451.5899 | 31824.2892 | 0.80269718 | 1.5524E-06 | 8.5644E-06 |
| ENSMUSG00000040147 | Maob | 376.992201 | 741.430768 | 364.438567 | 0.80167387 | 0.00374564 | 0.01113034 |
| ENSMUSG00000098206 | - | 0.31128264 | 4.4847058 | 4.17342316 | 0.80140913 | 0.01856649 | 0.04557331 |
| ENSMUSG00000044628 | Rnf208 | 0.40743019 | 3.79926297 | 3.39183278 | 0.80131859 | 0.0203286 | 0.04924055 |
| ENSMUSG00000048078 | Tenm4 | 165.380628 | 301.754389 | 136.37376 | 0.80109216 | 1.8685E-05 | 8.7006E-05 |
| ENSMUSG00000025780 | Itih5 | 17.3806211 | 35.8525843 | 18.4719632 | 0.80075055 | 0.00868451 | 0.02342746 |
| ENSMUSG00000097448 | - | 24.4048836 | 48.3869623 | 23.9820787 | 0.80065489 | 0.00552988 | 0.01573672 |
| ENSMUSG00000038700 | Hoxb5 | 122.461465 | 223.971322 | 101.509858 | 0.80056887 | 6.4283E-05 | 0.00027465 |
| ENSMUSG00000036155 | Mgat5 | 474.846811 | 849.373705 | 374.526894 | 0.80041096 | 7.7476E-08 | 5.1631E-07 |
| ENSMUSG00000040703 | Cyp2s1 | 16.5031771 | 36.0782366 | 19.5750594 | 0.80017166 | 0.01415246 | 0.03597039 |
| ENSMUSG00000099094 | - | 46.9843351 | 87.4847027 | 40.5003676 | 0.79908555 | 0.00024739 | 0.00094775 |
| ENSMUSG00000005836 | Gata6 | 261.948081 | 460.543406 | 198.595326 | 0.79898222 | 1.1991E-13 | 1.6413E-12 |
| ENSMUSG00000087519 | AV039307 | 10.5453591 | 23.2999322 | 12.754573 | 0.79898046 | 0.01338404 | 0.03423305 |
| ENSMUSG00000097944 | - | 99.7656673 | 179.017959 | 79.2522919 | 0.79874063 | 6.7678E-07 | 3.9407E-06 |
| ENSMUSG00000028392 | Bspry | 10.2855057 | 22.2988206 | 12.0133149 | 0.79867638 | 0.01471112 | 0.03722713 |
| ENSMUSG00000032679 | Cd59a | 0.33741727 | 4.08226261 | 3.74484533 | 0.79860389 | 0.01965849 | 0.0478767 |
| ENSMUSG00000038540 | Tmc3 | 42.7845781 | 80.0700285 | 37.2854504 | 0.79831353 | 0.0003864 | 0.00141896 |
| ENSMUSG00000068457 | Uty | 4168.48959 | 7262.12371 | 3093.63412 | 0.79699548 | 4.186E-58 | 1.0013E-55 |
| ENSMUSG00000092097 | - | 19.5806379 | 38.3192693 | 18.7386315 | 0.79675713 | 0.00407306 | 0.01201131 |
| ENSMUSG00000059839 | Zfp874b | 821.761386 | 1435.18823 | 613.426846 | 0.79663056 | 2.9727E-26 | 1.1944E-24 |
| ENSMUSG00000026017 | Carf | 196.438363 | 348.152176 | 151.713814 | 0.7965532 | 1.311E-10 | 1.2758E-09 |
| ENSMUSG00000091586 | Cyp4f17 | 245.382219 | 431.630003 | 186.247784 | 0.79641509 | 2.5836E-13 | 3.4005E-12 |
| ENSMUSG00000044250 | Pced1b | 0 | 2.80701853 | 2.80701853 | 0.79553758 | 0.01592194 | 0.03985717 |
| ENSMUSG00000006154 | Eps8l1 | 15.9795228 | 34.1219666 | 18.1424438 | 0.79518365 | 0.01169862 | 0.0304549 |
| ENSMUSG00000029436 | Mmp17 | 75.9731563 | 141.838814 | 65.8656578 | 0.79490735 | 0.00067057 | 0.0023449 |
| ENSMUSG00000026825 | Dnm1 | 1312.77589 | 2297.03722 | 984.261334 | 0.79477552 | 1.3838E-20 | 3.659E-19 |
| ENSMUSG00000072592 | Gm10373 | 26.5694131 | 49.7907077 | 23.2212946 | 0.79362028 | 0.00128354 | 0.0042411 |
| ENSMUSG00000000532 | Acvr1b | 944.423057 | 1665.79154 | 721.368487 | 0.79314351 | 2.2676E-10 | 2.1389E-09 |
| ENSMUSG00000020101 | Vsir | 621.287403 | 1087.32467 | 466.037271 | 0.79289444 | 8.1102E-15 | 1.2601E-13 |
| ENSMUSG00000022265 | Ank | 4642.49824 | 8082.94404 | 3440.4458 | 0.79261311 | 2.2527E-30 | 1.2228E-28 |
| ENSMUSG00000075514 | - | 147.758772 | 261.221457 | 113.462685 | 0.79225942 | 4.1148E-08 | 2.8617E-07 |
| ENSMUSG00000039646 | Vasn | 1915.73882 | 3368.08674 | 1452.34792 | 0.79208531 | 1.9749E-11 | 2.1127E-10 |
| ENSMUSG00000006764 | Tph2 | 183.749735 | 327.890939 | 144.141205 | 0.79177846 | 1.6604E-06 | 9.1109E-06 |
| ENSMUSG00000028487 | Bnc2 | 462.240069 | 810.306864 | 348.066795 | 0.79144724 | 1.6231E-15 | 2.7212E-14 |
| ENSMUSG00000039512 | Uhrf1bp1 | 3554.30824 | 6219.19173 | 2664.88349 | 0.79087297 | 4.8091E-15 | 7.6811E-14 |
| ENSMUSG00000029096 | Htra3 | 20.5675951 | 41.8965528 | 21.3289576 | 0.78889808 | 0.00803034 | 0.02185187 |
| ENSMUSG00000067928 | Zfp760 | 1102.24072 | 1934.43934 | 832.198628 | 0.78872112 | 4.0636E-11 | 4.1757E-10 |
| ENSMUSG00000031534 | Smim19 | 840.603599 | 1468.83157 | 628.227971 | 0.78820805 | 1.1281E-13 | 1.5472E-12 |
| ENSMUSG00000038425 | Poli | 711.530666 | 1240.63928 | 529.108617 | 0.7879844 | 4.7624E-15 | 7.6124E-14 |
| ENSMUSG00000051043 | Gprc5c | 0 | 4.26118139 | 4.26118139 | 0.78792848 | 0.01588485 | 0.03977883 |
| ENSMUSG00000030089 | Slc41a3 | 712.601352 | 1240.14588 | 527.544529 | 0.78728862 | 1.9259E-19 | 4.6448E-18 |
| ENSMUSG00000075389 | - | 86.5465734 | 160.045206 | 73.4986328 | 0.78665597 | 0.00054515 | 0.00194296 |
| ENSMUSG00000045231 | BC106179 | 8.1605982 | 18.3211294 | 10.1605312 | 0.7862347 | 0.01857333 | 0.04558464 |
| ENSMUSG00000055632 | Hmcn2 | 19.4195738 | 38.0729517 | 18.6533779 | 0.78529949 | 0.00483787 | 0.0139939 |
| ENSMUSG00000039508 | Calhm4 | 31.3861078 | 59.7526538 | 28.3665459 | 0.78520044 | 0.00286215 | 0.00872687 |
| ENSMUSG00000025176 | Hoga1 | 123.102735 | 219.175737 | 96.0730018 | 0.78455812 | 3.7886E-06 | 1.9692E-05 |
| ENSMUSG00000070524 | Fcrlb | 32.1725006 | 63.5558789 | 31.3833783 | 0.78451424 | 0.00643652 | 0.01800545 |
| ENSMUSG00000068551 | Zfp467 | 609.619792 | 1075.37246 | 465.752663 | 0.78397007 | 1.0117E-07 | 6.6263E-07 |
| ENSMUSG00000025875 | Tspan17 | 522.095513 | 979.983673 | 457.88816 | 0.78388257 | 0.00163477 | 0.00526463 |
| ENSMUSG00000000983 | Wfdc18 | 0 | 2.53284451 | 2.53284451 | 0.78380059 | 0.01730447 | 0.04283329 |
| ENSMUSG00000037977 | 6430571L13I | 21.6490198 | 40.3007444 | 18.6517246 | 0.78343802 | 0.00130272 | 0.00429688 |
| ENSMUSG00000030255 | Sspn | 919.301096 | 1694.70271 | 775.401615 | 0.78300227 | 0.00061888 | 0.0021808 |
| ENSMUSG00000022610 | Mapk12 | 394.337613 | 684.868829 | 290.531216 | 0.78277194 | 6.0612E-13 | 7.6357E-12 |
| ENSMUSG00000033949 | Trim36 | 91.2122096 | 164.188914 | 72.9767046 | 0.78265107 | 6.089E-05 | 0.0002608 |
| ENSMUSG00000033318 | Gstt2 | 32.3620323 | 67.347612 | 34.9855797 | 0.78234202 | 0.01377555 | 0.03510351 |
| ENSMUSG00000060380 | - | 124.516137 | 222.225919 | 97.7097821 | 0.78230911 | 1.5365E-06 | 8.4812E-06 |
| ENSMUSG00000036885 | Arhgef26 | 281.760191 | 495.298862 | 213.538671 | 0.78216982 | 4.3707E-09 | 3.4689E-08 |
| ENSMUSG00000026249 | Serpine2 | 19017.5896 | 33317.7401 | 14300.1505 | 0.78208813 | 1.8793E-09 | 1.5639E-08 |
| ENSMUSG00000043251 | Exoc3l | 106.977316 | 196.158911 | 89.181595 | 0.78169696 | 0.00040077 | 0.00146833 |
| ENSMUSG00000039770 | Ypel5 | 3752.86659 | 6506.96469 | 2754.0981 | 0.7812379 | 1.623E-17 | 3.2606E-16 |
| ENSMUSG00000049811 | Fam161a | 397.226857 | 689.849968 | 292.623111 | 0.78119037 | 5.826E-14 | 8.243E-13 |
| ENSMUSG00000086513 | - | 107.771138 | 237.02352 | 129.252382 | 0.78071436 | 0.01954446 | 0.04763635 |
| ENSMUSG00000070323 | Mmp27 | 0.40743019 | 5.80216507 | 5.39473489 | 0.77908971 | 0.02063488 | 0.04990605 |
| ENSMUSG00000050108 | Bpifc | 21.2905643 | 40.6077897 | 19.3172254 | 0.77819766 | 0.00288359 | 0.00878296 |
| ENSMUSG00000037321 | Tap1 | 5066.46218 | 8822.49558 | 3756.0334 | 0.77784916 | 8.3462E-11 | 8.2547E-10 |
| ENSMUSG00000054939 | Zfp174 | 213.484021 | 372.67554 | 159.191519 | 0.77761671 | 1.0939E-09 | 9.3532E-09 |
| ENSMUSG00000031066 | Usp11 | 1572.66894 | 2741.44153 | 1168.77258 | 0.77727293 | 4.3717E-10 | 3.9549E-09 |
| ENSMUSG00000025486 | Sirt3 | 425.422266 | 740.153236 | 314.73097 | 0.77645018 | 3.2877E-10 | 3.0356E-09 |
| ENSMUSG00000027827 | Kcnab1 | 62.3068886 | 112.129963 | 49.8230741 | 0.77595515 | 7.1837E-05 | 0.00030364 |

| | | | | | | | |
|---|---|---|---|---|---|---|---|
| ENSMUSG00000022687 | Boc | 169.686351 | 316.775952 | 147.089601 | 0.77476452 | 0.00207289 | 0.00653642 |
| ENSMUSG00000087368 | BC065397 | 92.9363418 | 168.682479 | 75.7461377 | 0.77473982 | 0.00023304 | 0.00089761 |
| ENSMUSG00000025665 | Rps6ka6 | 484.433629 | 842.998846 | 358.565217 | 0.77441795 | 2.133E-09 | 1.7658E-08 |
| ENSMUSG00000020400 | Tnip1 | 2296.3216 | 3934.7988 | 1638.47719 | 0.77416517 | 7.7865E-64 | 2.3908E-61 |
| ENSMUSG00000027938 | Creb3l4 | 78.3546226 | 139.896174 | 61.5415516 | 0.77370972 | 3.3652E-05 | 0.0001502 |
| ENSMUSG00000037003 | Tns2 | 2680.68617 | 4645.1784 | 1964.49223 | 0.77369597 | 4.5508E-12 | 5.2156E-11 |
| ENSMUSG00000032308 | Ulk3 | 788.946796 | 1355.72712 | 566.780328 | 0.77356911 | 2.9172E-23 | 9.556E-22 |
| ENSMUSG00000032262 | Elovl4 | 385.774403 | 664.921916 | 279.147513 | 0.77338379 | 1.2481E-20 | 3.3131E-19 |
| ENSMUSG00000036552 | Ermard | 258.949454 | 451.140103 | 192.190649 | 0.77307226 | 7.9088E-09 | 6.0914E-08 |
| ENSMUSG00000049420 | Tmem200a | 191.532519 | 382.572169 | 191.03965 | 0.77285587 | 0.01041457 | 0.02749949 |
| ENSMUSG00000066113 | Adamtsl1 | 58.0763102 | 104.620906 | 46.5445958 | 0.77283348 | 0.00016378 | 0.00064782 |
| ENSMUSG00000025006 | Sorbs1 | 185.949407 | 328.781725 | 142.832317 | 0.77274735 | 5.3158E-06 | 2.6962E-05 |
| ENSMUSG00000089940 | - | 134.497789 | 240.932037 | 106.434248 | 0.77262107 | 5.7707E-05 | 0.00024796 |
| ENSMUSG00000042797 | Aqp11 | 18.8652197 | 36.3696662 | 17.5044465 | 0.77190593 | 0.00675951 | 0.0188005 |
| ENSMUSG00000043073 | Usp17le | 4.30802068 | 134.632586 | 130.324565 | 0.77177753 | 0.01809 | 0.04456322 |
| ENSMUSG00000074782 | 4833422C13 | 37.7490231 | 69.4100923 | 31.6610692 | 0.77166732 | 0.00140769 | 0.00460837 |
| ENSMUSG00000026544 | Dusp23 | 44.6281634 | 80.9781983 | 36.3500349 | 0.77145023 | 0.00060323 | 0.0021304 |
| ENSMUSG00000027514 | Zbp1 | 8218.73667 | 14223.1312 | 6004.39448 | 0.77086613 | 2.2721E-11 | 2.4131E-10 |
| ENSMUSG00000033763 | Mtss2 | 2285.92035 | 4288.00935 | 2002.089 | 0.77081679 | 0.00288396 | 0.00878296 |
| ENSMUSG00000033730 | Egr3 | 9.08934977 | 18.4775646 | 9.38821482 | 0.77058357 | 0.01384184 | 0.03525061 |
| ENSMUSG00000030782 | Tgfb1i1 | 536.46326 | 943.930327 | 407.467066 | 0.77056585 | 3.5848E-06 | 1.8694E-05 |
| ENSMUSG00000027386 | Fbln7 | 33.6924649 | 68.8744696 | 35.1820047 | 0.77026658 | 0.01386864 | 0.0353101 |
| ENSMUSG00000047562 | Mmp10 | 2305.00727 | 3995.24373 | 1690.23646 | 0.76944504 | 8.4539E-10 | 7.3227E-09 |
| ENSMUSG00000059142 | Zfp945 | 1039.2625 | 1800.87964 | 761.617145 | 0.76938719 | 3.625E-10 | 3.3203E-09 |
| ENSMUSG00000045506 | - | 44.8212985 | 83.2273817 | 38.4060832 | 0.76918557 | 0.00199155 | 0.00630601 |
| ENSMUSG00000079465 | Col4a3 | 22.7228936 | 43.4168236 | 20.69393 | 0.76883949 | 0.00567613 | 0.01611057 |
| ENSMUSG00000033577 | Myo6 | 1595.92868 | 2734.88303 | 1138.95436 | 0.76880837 | 3.3226E-27 | 1.433E-25 |
| ENSMUSG00000021066 | Atl1 | 167.350752 | 298.38192 | 131.031168 | 0.76847158 | 7.0447E-05 | 0.00029826 |
| ENSMUSG00000024066 | Xdh | 7156.37156 | 12235.7219 | 5079.35036 | 0.76822652 | 2.0851E-36 | 1.631E-34 |
| ENSMUSG00000057098 | Ebf1 | 51.4029523 | 94.2001021 | 42.7971498 | 0.76781047 | 0.00081476 | 0.00280521 |
| ENSMUSG00000035735 | Dagla | 1805.20336 | 3104.44985 | 1299.24649 | 0.76760309 | 1.0567E-15 | 1.8071E-14 |
| ENSMUSG00000029752 | Asns | 21711.5778 | 37496.7896 | 15785.2118 | 0.76755598 | 4.1622E-11 | 4.2749E-10 |
| ENSMUSG00000027544 | Nfatc2 | 12.8448181 | 24.843551 | 11.9987329 | 0.76752872 | 0.00734555 | 0.02020493 |
| ENSMUSG00000043015 | Nemp2 | 440.766972 | 759.520259 | 318.753288 | 0.76735316 | 3.161E-15 | 5.1569E-14 |
| ENSMUSG00000033863 | Klf9 | 3502.37213 | 5989.46503 | 2487.0929 | 0.76724757 | 2.2588E-30 | 1.2228E-28 |
| ENSMUSG00000026317 | Cln8 | 1858.33421 | 3212.03997 | 1353.70576 | 0.76722229 | 2.2675E-10 | 2.1389E-09 |
| ENSMUSG00000082292 | - | 57.0328274 | 112.924103 | 55.8912759 | 0.76719922 | 0.01046045 | 0.02759231 |
| ENSMUSG00000042608 | Stk40 | 863.637542 | 1480.42424 | 616.786698 | 0.76630287 | 2.1939E-20 | 5.6629E-19 |
| ENSMUSG00000036890 | Gtdc1 | 262.327352 | 452.972663 | 190.645311 | 0.76630157 | 2.7479E-11 | 2.8871E-10 |
| ENSMUSG00000043165 | Lor | 8.49801548 | 18.0577209 | 9.55970545 | 0.76629397 | 0.01903807 | 0.04654204 |
| ENSMUSG00000073096 | Lrrc61 | 292.5664 | 504.933546 | 212.367146 | 0.76592679 | 6.3885E-11 | 6.4078E-10 |
| ENSMUSG00000023960 | Enpp5 | 1400.08509 | 2627.0625 | 1226.97741 | 0.7654967 | 0.00360384 | 0.01075277 |
| ENSMUSG00000054146 | Krt15 | 71.7090139 | 130.363482 | 58.6544679 | 0.76495766 | 0.0009455 | 0.00321183 |
| ENSMUSG00000040570 | Rundc3b | 96.3479252 | 170.650386 | 74.3024606 | 0.76449597 | 1.7423E-05 | 8.1534E-05 |
| ENSMUSG00000052949 | Rnf157 | 1295.66144 | 2227.45612 | 931.794687 | 0.76432094 | 7.6545E-13 | 9.5377E-12 |
| ENSMUSG00000028088 | Fmo5 | 126.951279 | 219.483083 | 92.5318039 | 0.76417341 | 4.1959E-07 | 2.5239E-06 |
| ENSMUSG00000022865 | Cxadr | 2188.59076 | 3750.75823 | 1562.16748 | 0.76406993 | 1.1647E-16 | 2.1922E-15 |
| ENSMUSG00000005800 | Mmp8 | 0 | 2.49854274 | 2.49854274 | 0.7638187 | 0.01954822 | 0.04763635 |
| ENSMUSG00000085996 | - | 0 | 2.63806434 | 2.63806434 | 0.76370336 | 0.01949147 | 0.04752056 |
| ENSMUSG00000044566 | Cage1 | 65.153682 | 115.210924 | 50.057242 | 0.76332915 | 1.6811E-05 | 7.8869E-05 |
| ENSMUSG00000072812 | - | 2201.17432 | 3819.36099 | 1618.18668 | 0.76290677 | 8.253E-08 | 5.4786E-07 |
| ENSMUSG00000031351 | Zfp185 | 16.2965546 | 32.5817553 | 16.2852006 | 0.76256144 | 0.01521312 | 0.0383112 |
| ENSMUSG00000054256 | Msi1 | 63.7638597 | 113.883586 | 50.119726 | 0.76242227 | 0.00011813 | 0.0004798 |
| ENSMUSG00000028634 | Hivep3 | 126.970602 | 221.912162 | 94.9415604 | 0.76242062 | 2.2547E-06 | 1.2089E-05 |
| ENSMUSG00000043029 | Trpv3 | 13.0860878 | 25.2608608 | 12.174773 | 0.76219399 | 0.00683749 | 0.01899281 |
| ENSMUSG00000053519 | Kcnip1 | 0 | 2.77137331 | 2.77137331 | 0.76207003 | 0.01954657 | 0.04763635 |
| ENSMUSG00000056501 | Cebpb | 7759.3359 | 13329.3493 | 5570.01338 | 0.76186123 | 8.7134E-12 | 9.7262E-11 |
| ENSMUSG00000031565 | Fgfr1 | 4497.24558 | 7713.59154 | 3216.34596 | 0.76101078 | 1.2127E-12 | 1.4815E-11 |
| ENSMUSG00000032182 | Yipf2 | 496.420205 | 848.541581 | 352.121377 | 0.7604943 | 1.3136E-16 | 2.4499E-15 |
| ENSMUSG00000039304 | Tnfsf10 | 107.757772 | 200.111362 | 92.3535901 | 0.7599968 | 0.00309608 | 0.00937345 |
| ENSMUSG00000003344 | Btbd2 | 2143.96761 | 3647.4806 | 1503.51299 | 0.75994285 | 2.3965E-27 | 1.0445E-25 |
| ENSMUSG00000038058 | Nod1 | 290.060298 | 509.89194 | 219.831642 | 0.75990171 | 3.0519E-05 | 0.00013723 |
| ENSMUSG00000035818 | Plekhs1 | 10.8287054 | 22.3728772 | 11.5441718 | 0.75965635 | 0.01791801 | 0.04418716 |
| ENSMUSG00000054203 | Ifi205 | 859.362336 | 1464.12473 | 604.762394 | 0.75954366 | 5.6822E-19 | 1.3193E-17 |
| ENSMUSG00000097573 | - | 17.3983648 | 32.2783265 | 14.8799618 | 0.75892248 | 0.00351202 | 0.01050441 |
| ENSMUSG00000018800 | Abca5 | 1479.50095 | 2524.04339 | 1044.54245 | 0.75848973 | 1.9998E-18 | 4.4189E-17 |
| ENSMUSG00000061666 | Gdpd1 | 1721.43786 | 2950.19439 | 1228.75652 | 0.75813186 | 2.5983E-11 | 2.7411E-10 |
| ENSMUSG00000055866 | Per2 | 292.050241 | 508.498188 | 216.447947 | 0.75776521 | 1.5731E-06 | 8.6691E-06 |
| ENSMUSG00000032911 | Cspg4 | 7.89648846 | 16.2009442 | 8.30445571 | 0.75761999 | 0.02023585 | 0.0490546 |
| ENSMUSG00000056367 | Actr3b | 51.383407 | 94.1634086 | 42.7800017 | 0.75743038 | 0.00190448 | 0.00605453 |
| ENSMUSG00000061654 | - | 656.904033 | 1117.55361 | 460.649579 | 0.75729891 | 6.6942E-22 | 1.9786E-20 |
| ENSMUSG00000027276 | Jag1 | 1589.80873 | 2726.75479 | 1136.94605 | 0.7555046 | 5.395E-10 | 4.8087E-09 |
| ENSMUSG00000097892 | - | 17.0155763 | 34.3419088 | 17.3263325 | 0.7554929 | 0.01248834 | 0.03221444 |
| ENSMUSG00000086608 | - | 0 | 2.70303574 | 2.70303574 | 0.75524606 | 0.02039952 | 0.04939488 |
| ENSMUSG00000027357 | Crls1 | 2623.94927 | 4525.9311 | 1901.98184 | 0.75455194 | 1.4789E-07 | 9.4839E-07 |

| | | | | | | | |
|---|---|---|---|---|---|---|---|
| ENSMUSG00000021534 | 1700001L19I | 33.0176303 | 60.6135092 | 27.5958789 | 0.75400113 | 0.00222743 | 0.00697453 |
| ENSMUSG00000074578 | Zfas1 | 3841.6261 | 6618.01954 | 2776.39344 | 0.75387226 | 9.5167E-08 | 6.2649E-07 |
| ENSMUSG00000046329 | Slc25a23 | 559.114856 | 969.778043 | 410.663187 | 0.75310099 | 3.0626E-06 | 1.6151E-05 |
| ENSMUSG00000023266 | Frs3 | 192.890357 | 336.258682 | 143.368325 | 0.75210557 | 6.2832E-06 | 3.1534E-05 |
| ENSMUSG00000097578 | - | 26.924574 | 48.3611396 | 21.4365657 | 0.75201286 | 0.00176291 | 0.00564286 |
| ENSMUSG00000075012 | Fjx1 | 106.596142 | 198.837858 | 92.2417158 | 0.75174705 | 0.0049142 | 0.01418279 |
| ENSMUSG00000073412 | Lst1 | 35.4235511 | 67.1271201 | 31.703569 | 0.75168221 | 0.00759759 | 0.02081569 |
| ENSMUSG00000045193 | Cirbp | 935.677065 | 1584.0832 | 648.406136 | 0.75159356 | 2.3374E-21 | 6.614E-20 |
| ENSMUSG00000027985 | Lef1 | 0 | 3.30313676 | 3.30313676 | 0.75146197 | 0.02043017 | 0.04944583 |
| ENSMUSG00000073910 | Mob3b | 46.9474765 | 84.060111 | 37.1126344 | 0.75134893 | 0.00031003 | 0.00116258 |
| ENSMUSG00000071369 | Map3k5 | 484.834557 | 822.635065 | 337.800509 | 0.75127633 | 3.165E-18 | 6.89E-17 |
| ENSMUSG00000028572 | Hook1 | 9.2834466 | 19.4850213 | 10.2015748 | 0.75118408 | 0.0195918 | 0.04773688 |
| ENSMUSG00000027224 | Duoxa1 | 12.2087654 | 23.8399418 | 11.6311764 | 0.75077024 | 0.01483474 | 0.03750064 |
| ENSMUSG00000037379 | Spon2 | 30.5502092 | 56.9616421 | 26.4114329 | 0.75064991 | 0.00441303 | 0.01288932 |
| ENSMUSG00000010095 | Slc3a2 | 23017.8606 | 39083.8306 | 16065.9699 | 0.75043988 | 3.5899E-15 | 5.815E-14 |
| ENSMUSG00000097121 | D130020L05 | 29.3919951 | 53.1828655 | 23.7908704 | 0.7503895 | 0.00258225 | 0.00796551 |
| ENSMUSG00000024059 | Clip4 | 96.7671626 | 169.063963 | 72.2968006 | 0.75026388 | 3.6831E-05 | 0.00016312 |
| ENSMUSG00000003070 | Efna2 | 88.342333 | 154.229679 | 65.8873457 | 0.74950611 | 1.4034E-05 | 6.6785E-05 |
| ENSMUSG00000039450 | Dcxr | 767.778205 | 1311.29705 | 543.518846 | 0.74949603 | 1.4269E-09 | 1.206E-08 |
| ENSMUSG00000028860 | Sytl1 | 13.3534922 | 25.6792298 | 12.3257376 | 0.74947054 | 0.00867766 | 0.0234151 |
| ENSMUSG00000021895 | Arhgef3 | 2221.46749 | 3753.93972 | 1532.47223 | 0.7492517 | 5.8778E-25 | 2.1709E-23 |
| ENSMUSG00000091898 | Tnnc1 | 2235.07464 | 4202.40772 | 1967.33308 | 0.74891554 | 0.006735 | 0.01874356 |
| ENSMUSG00000053469 | Tg | 30.0424968 | 57.1521287 | 27.109632 | 0.74601732 | 0.00867065 | 0.02340473 |
| ENSMUSG00000038517 | Tbkbp1 | 76.520734 | 147.668971 | 71.1482366 | 0.74568824 | 0.01198452 | 0.03110593 |
| ENSMUSG00000036912 | Piwil4 | 212.54123 | 363.881342 | 151.340112 | 0.74533122 | 6.7792E-08 | 4.5472E-07 |
| ENSMUSG00000072621 | Slfn10-ps | 749.869689 | 1284.14214 | 534.272451 | 0.74488069 | 1.091E-07 | 7.1113E-07 |
| ENSMUSG00000097208 | - | 10.4684481 | 20.3349561 | 9.86650799 | 0.744871 | 0.01622301 | 0.04047293 |
| ENSMUSG00000055305 | Zfp93 | 428.967352 | 726.168835 | 297.201483 | 0.74482108 | 1.1608E-13 | 1.591E-12 |
| ENSMUSG00000072941 | Sod3 | 8404.85283 | 14714.5432 | 6309.69035 | 0.74454489 | 0.00013439 | 0.00054038 |
| ENSMUSG00000058297 | Spock2 | 12.3538876 | 24.1728679 | 11.8189803 | 0.74335387 | 0.01563762 | 0.0392274 |
| ENSMUSG00000028444 | Cntfr | 81.4216474 | 151.615626 | 70.1939788 | 0.74291009 | 0.00618726 | 0.01738857 |
| ENSMUSG00000029049 | Morn1 | 26.9431577 | 48.1058607 | 21.162703 | 0.74229012 | 0.00089897 | 0.00306593 |
| ENSMUSG00000025357 | Dgka | 1807.96781 | 3036.97809 | 1229.01028 | 0.74210601 | 1.0198E-27 | 4.5705E-26 |
| ENSMUSG00000041272 | Tox | 255.612039 | 432.387686 | 176.775646 | 0.74172009 | 2.5256E-12 | 2.9878E-11 |
| ENSMUSG00000032076 | Cadm1 | 76.0211693 | 141.865351 | 65.844182 | 0.74154865 | 0.00693773 | 0.01922455 |
| ENSMUSG00000053477 | Tcf4 | 6892.24013 | 11690.1404 | 4797.90024 | 0.74153426 | 3.1195E-10 | 2.8856E-09 |
| ENSMUSG00000023330 | Dtwd1 | 3427.94527 | 5765.4231 | 2337.47783 | 0.74107719 | 9.1516E-20 | 2.2601E-18 |
| ENSMUSG00000081824 | - | 257.166995 | 442.293386 | 185.126392 | 0.74059235 | 9.2912E-06 | 4.539E-05 |
| ENSMUSG00000023367 | Tmem176a | 6741.22059 | 11370.2927 | 4629.07214 | 0.74046638 | 3.6951E-14 | 5.3607E-13 |
| ENSMUSG00000022382 | Wnt7b | 24.1215373 | 43.7032812 | 19.5817439 | 0.73987014 | 0.00245132 | 0.00759811 |
| ENSMUSG00000028015 | Ctso | 855.387168 | 1441.81524 | 586.428073 | 0.73920688 | 5.9108E-13 | 7.4508E-12 |
| ENSMUSG00000049001 | Ndnf | 712.565278 | 1199.58955 | 487.024272 | 0.73910092 | 1.3942E-18 | 3.1415E-17 |
| ENSMUSG00000039908 | Slc26a11 | 407.455185 | 687.173167 | 279.717982 | 0.73807146 | 3.1982E-12 | 3.7277E-11 |
| ENSMUSG00000025485 | Ric8a | 3345.33584 | 5608.46841 | 2263.13257 | 0.737799 | 2.8594E-23 | 9.3817E-22 |
| ENSMUSG00000009905 | Kdsr | 1822.30772 | 3080.72174 | 1258.41401 | 0.73709388 | 4.6521E-10 | 4.1883E-09 |
| ENSMUSG00000037921 | Ddx60 | 6748.96596 | 12681.6022 | 5932.63619 | 0.73671444 | 0.00941654 | 0.02513847 |
| ENSMUSG00000054181 | - | 24.023588 | 42.563555 | 18.539967 | 0.73629854 | 0.00164437 | 0.00529306 |
| ENSMUSG00000036775 | Decr2 | 972.980408 | 1627.59237 | 654.611963 | 0.73623072 | 8.9461E-24 | 3.022E-22 |
| ENSMUSG00000020883 | Fbxl20 | 2406.71968 | 4027.29657 | 1620.57689 | 0.73425247 | 1.9641E-22 | 6.0576E-21 |
| ENSMUSG00000056900 | Usp13 | 66.3289212 | 125.18191 | 58.8529887 | 0.73344771 | 0.01108761 | 0.02904374 |
| ENSMUSG00000049404 | Rarres1 | 834.114878 | 1402.86036 | 568.745484 | 0.73337793 | 4.3735E-11 | 4.4654E-10 |
| ENSMUSG00000095440 | Fignl2 | 72.9468159 | 131.48182 | 58.5350036 | 0.733207 | 0.00403063 | 0.01189985 |
| ENSMUSG00000027204 | Fbn1 | 20070.6894 | 34014.3233 | 13943.6339 | 0.73307869 | 7.8504E-08 | 5.2214E-07 |
| ENSMUSG00000038886 | Man2a2 | 2653.7628 | 4441.33478 | 1787.57198 | 0.73295134 | 1.4483E-19 | 3.5177E-18 |
| ENSMUSG00000036257 | Pnpla8 | 4549.79132 | 7584.42358 | 3034.63226 | 0.73283464 | 2.582E-37 | 2.1253E-35 |
| ENSMUSG00000030424 | Zfp939 | 108.585158 | 188.734291 | 80.1491333 | 0.73270943 | 0.00018433 | 0.00072282 |
| ENSMUSG00000003279 | Dlgap1 | 15.9431955 | 31.4932926 | 15.5500971 | 0.73264817 | 0.01972523 | 0.04801651 |
| ENSMUSG00000085412 | Halr1 | 29.7461944 | 52.5821061 | 22.8359118 | 0.73246893 | 0.0022966 | 0.00716822 |
| ENSMUSG00000097360 | 9430065F17I | 22.8898941 | 41.9422586 | 19.0523645 | 0.73224923 | 0.00478523 | 0.01385527 |
| ENSMUSG00000049742 | - | 35.6732119 | 64.3678588 | 28.694647 | 0.73220833 | 0.00382798 | 0.01135533 |
| ENSMUSG00000085385 | - | 1678.0882 | 2808.97991 | 1130.89171 | 0.73210995 | 3.7743E-18 | 8.1135E-17 |
| ENSMUSG00000022090 | Pdlim2 | 473.757906 | 801.690763 | 327.932858 | 0.73172749 | 7.6293E-08 | 5.0875E-07 |
| ENSMUSG00000087331 | 1810021B22 | 40.2225023 | 70.9856163 | 30.763114 | 0.7309976 | 0.00114833 | 0.00383373 |
| ENSMUSG00000020105 | Lrig3 | 643.618874 | 1081.05121 | 437.43234 | 0.73091936 | 3.8756E-11 | 4.0004E-10 |
| ENSMUSG00000021884 | Hacl1 | 420.203668 | 706.007647 | 285.803979 | 0.7308051 | 1.0426E-10 | 1.0218E-09 |
| ENSMUSG00000039787 | Cercam | 132.767477 | 226.773993 | 94.0065156 | 0.73077683 | 4.9916E-06 | 2.5449E-05 |
| ENSMUSG00000021133 | Susd6 | 8961.72394 | 14959.2091 | 5997.48519 | 0.73031768 | 1.5572E-20 | 4.086E-19 |
| ENSMUSG00000097027 | - | 110.021313 | 186.08471 | 76.0633967 | 0.72974591 | 8.3133E-07 | 4.7851E-06 |
| ENSMUSG00000001911 | Nfix | 4189.44063 | 7007.86429 | 2818.42365 | 0.72950994 | 4.1578E-15 | 6.6928E-14 |
| ENSMUSG00000078247 | Airn | 41.3915619 | 78.2611256 | 36.8695638 | 0.72948749 | 0.01262493 | 0.03249751 |
| ENSMUSG00000032454 | Rbp2 | 186.917507 | 326.389985 | 139.472478 | 0.72929275 | 0.00069507 | 0.00242521 |
| ENSMUSG00000087177 | E130307A14 | 329.230704 | 551.461551 | 222.230847 | 0.72837395 | 2.3891E-12 | 2.8361E-11 |
| ENSMUSG00000087159 | Gm15246 | 75.2686702 | 133.660823 | 58.3921528 | 0.72827322 | 0.0012559 | 0.00416113 |
| ENSMUSG00000025464 | Paox | 217.415655 | 365.162198 | 147.746543 | 0.72826874 | 2.3547E-09 | 1.9384E-08 |
| ENSMUSG00000047153 | Khnyn | 1878.06477 | 3130.79779 | 1252.73302 | 0.72794613 | 9.053E-22 | 2.6492E-20 |

| | | | | | | | |
|---|---|---|---|---|---|---|---|
| ENSMUSG00000038067 | Csf3 | 525.583182 | 908.066295 | 382.483113 | 0.72776711 | 0.00019242 | 0.00075126 |
| ENSMUSG00000071202 | Ccdc78 | 12.9671002 | 24.6285248 | 11.6614245 | 0.72762472 | 0.01482795 | 0.03749271 |
| ENSMUSG00000089818 | - | 10.4474098 | 20.3549586 | 9.90754874 | 0.72713273 | 0.01899007 | 0.04646334 |
| ENSMUSG00000017830 | Dhx58 | 3836.25083 | 6449.69238 | 2613.44155 | 0.72703865 | 4.9623E-09 | 3.9158E-08 |
| ENSMUSG00000056656 | Apol8 | 61.9979389 | 108.059055 | 46.0611163 | 0.72702649 | 0.00046266 | 0.00167509 |
| ENSMUSG00000046191 | Pcdhb20 | 151.806287 | 262.616726 | 110.810439 | 0.7267444 | 0.00024981 | 0.00095577 |
| ENSMUSG00000033256 | Shf | 547.007061 | 915.296228 | 368.289167 | 0.72656217 | 5.2449E-11 | 5.2995E-10 |
| ENSMUSG00000078671 | Chd2 | 3632.29026 | 6111.57087 | 2479.28061 | 0.72619429 | 1.5488E-08 | 1.1424E-07 |
| ENSMUSG00000007989 | Fzd3 | 42.7201927 | 76.3365405 | 33.6163478 | 0.72597472 | 0.00275094 | 0.00841774 |
| ENSMUSG00000041624 | Gucy1a2 | 189.991107 | 319.882107 | 129.891 | 0.72548582 | 1.5766E-08 | 1.1625E-07 |
| ENSMUSG00000047013 | Fbxo41 | 177.818016 | 302.834965 | 125.016948 | 0.72531644 | 2.0453E-06 | 1.1041E-05 |
| ENSMUSG00000036273 | Lrrk2 | 1753.73324 | 3039.61234 | 1285.8791 | 0.72504592 | 0.00036122 | 0.0013365 |
| ENSMUSG00000040350 | Trim7 | 106.033928 | 181.079401 | 75.0454728 | 0.72464085 | 6.176E-05 | 0.00026436 |
| ENSMUSG00000021684 | Pde8b | 76.6810236 | 131.030884 | 54.3498608 | 0.72420415 | 9.2019E-06 | 4.4997E-05 |
| ENSMUSG00000040187 | Arntl2 | 2347.65487 | 3919.64278 | 1571.98791 | 0.72386107 | 4.0846E-12 | 4.7049E-11 |
| ENSMUSG00000066553 | - | 17.5906599 | 32.2528276 | 14.6621677 | 0.72368154 | 0.00898206 | 0.02411314 |
| ENSMUSG00000029769 | Ccdc136 | 979.386513 | 1624.04535 | 644.658836 | 0.72343138 | 1.3691E-26 | 5.7364E-25 |
| ENSMUSG00000026166 | Ccl20 | 3986.78478 | 6651.85601 | 2665.07123 | 0.72324981 | 4.2901E-12 | 4.936E-11 |
| ENSMUSG00000083240 | - | 93.379072 | 158.576617 | 65.1975448 | 0.7225157 | 2.1917E-05 | 0.00010107 |
| ENSMUSG00000048756 | Foxo3 | 4823.40312 | 8023.69852 | 3200.2954 | 0.722483 | 1.2113E-15 | 2.0628E-14 |
| ENSMUSG00000055862 | Izumo4 | 101.067202 | 172.647081 | 71.5798795 | 0.72243466 | 4.0432E-05 | 0.00017803 |
| ENSMUSG00000026839 | Upp2 | 117.908571 | 197.813996 | 79.9054254 | 0.72162538 | 7.6982E-07 | 4.4561E-06 |
| ENSMUSG00000026158 | Ogfrl1 | 389.866858 | 661.217662 | 271.350804 | 0.71993235 | 1.4243E-05 | 6.7702E-05 |
| ENSMUSG00000083282 | Ctsf | 616.879853 | 1023.63048 | 406.750629 | 0.71990531 | 1.7674E-13 | 2.3702E-12 |
| ENSMUSG00000033249 | Hsf4 | 135.638416 | 228.030795 | 92.3923782 | 0.71975853 | 5.0909E-06 | 2.5923E-05 |
| ENSMUSG00000032589 | Bsn | 205.662333 | 343.386135 | 137.723802 | 0.71970199 | 4.6345E-09 | 3.6712E-08 |
| ENSMUSG00000071506 | Tmem139 | 54.7688557 | 94.5039857 | 39.73513 | 0.71944594 | 0.00033503 | 0.00124656 |
| ENSMUSG00000059288 | Cdyl | 1882.68281 | 3117.68666 | 1235.00386 | 0.71914623 | 8.4123E-21 | 2.2801E-19 |
| ENSMUSG00000039745 | Htatip2 | 833.849276 | 1412.68431 | 578.835035 | 0.7185337 | 1.7E-05 | 7.9665E-05 |
| ENSMUSG00000040441 | Slc26a10 | 33.5936755 | 62.0993846 | 28.5057091 | 0.71843355 | 0.01286735 | 0.03305945 |
| ENSMUSG00000053574 | 4930563E22 | 90.9188579 | 162.374652 | 71.4557943 | 0.71825175 | 0.00388001 | 0.01149641 |
| ENSMUSG00000026873 | Phf19 | 271.776306 | 453.60078 | 181.824474 | 0.71744357 | 8.3364E-10 | 7.23E-09 |
| ENSMUSG00000071757 | Zhx2 | 1303.57651 | 2195.37351 | 891.797 | 0.71668255 | 2.8999E-06 | 1.5356E-05 |
| ENSMUSG00000049907 | Rasl11b | 118.15854 | 198.011666 | 79.853126 | 0.71618921 | 2.3943E-07 | 1.4935E-06 |
| ENSMUSG00000042256 | Ptchd4 | 991.725077 | 1641.31995 | 649.594873 | 0.7160489 | 9.9588E-17 | 1.8847E-15 |
| ENSMUSG00000093880 | - | 9.83419723 | 18.5534006 | 8.71920342 | 0.71500479 | 0.01806082 | 0.044518 |
| ENSMUSG00000034872 | Gipc3 | 25.7116361 | 47.0097099 | 21.2980738 | 0.71478535 | 0.01115886 | 0.02920982 |
| ENSMUSG00000024975 | Pdcd4 | 2503.59896 | 4128.07293 | 1624.47398 | 0.71378584 | 9.4665E-20 | 2.3323E-18 |
| ENSMUSG00000046598 | Bdh1 | 28.2676538 | 52.080704 | 23.8130502 | 0.71371636 | 0.01122471 | 0.02936725 |
| ENSMUSG00000027858 | Tspan2 | 43.9635215 | 76.3367174 | 32.3731959 | 0.71333002 | 0.00153735 | 0.0049876 |
| ENSMUSG00000040265 | Dnm3 | 25.804489 | 46.1699713 | 20.3654823 | 0.71317889 | 0.00595831 | 0.01681634 |
| ENSMUSG00000042284 | Itga1 | 542.358092 | 901.551534 | 359.193442 | 0.71278772 | 9.4257E-10 | 8.12E-09 |
| ENSMUSG00000084616 | - | 18.0587503 | 32.6237695 | 14.5650192 | 0.71239664 | 0.01007107 | 0.02667465 |
| ENSMUSG00000090124 | Ugt1a7c | 85.7955463 | 151.462331 | 65.6667845 | 0.71175098 | 0.00338873 | 0.01017261 |
| ENSMUSG00000031709 | Tbc1d9 | 1298.34894 | 2142.40963 | 844.060682 | 0.71141944 | 1.4649E-15 | 2.4721E-14 |
| ENSMUSG00000040254 | Sema3d | 245.42843 | 405.451407 | 160.022977 | 0.71057093 | 9.9102E-12 | 1.0967E-10 |
| ENSMUSG00000022105 | Rb1 | 247.17978 | 411.394562 | 164.214782 | 0.71054912 | 2.8595E-08 | 2.0341E-07 |
| ENSMUSG00000030720 | Cln3 | 553.86568 | 913.549204 | 359.683524 | 0.71007419 | 4.4616E-14 | 6.4139E-13 |
| ENSMUSG00000009828 | Cilk1 | 783.074411 | 1292.19287 | 509.118463 | 0.70922393 | 1.5366E-14 | 2.3158E-13 |
| ENSMUSG00000043300 | B3galnt1 | 909.63491 | 1524.30679 | 614.671882 | 0.70912574 | 5.2585E-06 | 2.6724E-05 |
| ENSMUSG00000040410 | Fbxl4 | 912.905349 | 1500.87096 | 587.965612 | 0.70821385 | 7.5418E-20 | 1.8738E-18 |
| ENSMUSG00000087403 | - | 56.069019 | 96.6185493 | 40.5495303 | 0.70791796 | 0.00053888 | 0.00192261 |
| ENSMUSG00000063894 | Zkscan8 | 1726.75125 | 2893.19853 | 1166.44728 | 0.70766655 | 6.097E-06 | 3.0682E-05 |
| ENSMUSG00000050957 | Insl6 | 546.245123 | 903.994445 | 357.749323 | 0.70748387 | 9.3417E-09 | 7.1256E-08 |
| ENSMUSG00000028799 | Zfp362 | 327.961772 | 545.493403 | 217.531631 | 0.70743321 | 1.456E-07 | 9.3428E-07 |
| ENSMUSG00000034557 | Zfyve9 | 2241.54577 | 3680.02284 | 1438.47707 | 0.70724218 | 2.6691E-24 | 9.3542E-23 |
| ENSMUSG00000024789 | Jak2 | 7914.16456 | 13002.486 | 5088.32142 | 0.70699901 | 3.5922E-18 | 7.7788E-17 |
| ENSMUSG00000000223 | Drp2 | 65.2535546 | 111.502513 | 46.2489586 | 0.70688056 | 0.00026522 | 0.00100797 |
| ENSMUSG00000010651 | Acaa1b | 34.0356313 | 60.26515 | 26.2295186 | 0.7059536 | 0.00741769 | 0.02037885 |
| ENSMUSG00000084925 | 1810062O18 | 10.4945828 | 19.3386534 | 8.84407063 | 0.70550674 | 0.02005594 | 0.04868328 |
| ENSMUSG00000060988 | Galnt13 | 395.576098 | 653.960454 | 258.384356 | 0.70479859 | 1.0357E-08 | 7.8534E-08 |
| ENSMUSG00000038260 | Trpm4 | 813.541141 | 1335.85155 | 522.310406 | 0.70468735 | 2.9189E-19 | 6.9339E-18 |
| ENSMUSG00000024885 | Aldh3b1 | 466.626994 | 780.750434 | 314.12344 | 0.70436404 | 1.5525E-05 | 7.3253E-05 |
| ENSMUSG00000079173 | Zan | 81.2481793 | 136.604263 | 55.3560832 | 0.70348806 | 3.4791E-05 | 0.00015485 |
| ENSMUSG00000037846 | Rtkn2 | 442.977525 | 728.381799 | 285.404274 | 0.70218986 | 7.474E-12 | 8.4065E-11 |
| ENSMUSG00000039840 | Epg5 | 3899.83971 | 6411.96056 | 2512.12086 | 0.70197925 | 1.0958E-11 | 1.2055E-10 |
| ENSMUSG00000025227 | Mfsd13a | 178.260007 | 294.945426 | 116.685419 | 0.70149858 | 5.5878E-07 | 3.2994E-06 |
| ENSMUSG00000002578 | Ikzf4 | 169.374724 | 282.579184 | 113.20446 | 0.70141154 | 1.1216E-06 | 6.3062E-06 |
| ENSMUSG00000031785 | Adgrg1 | 202.627157 | 340.131135 | 137.503979 | 0.70126635 | 3.0975E-05 | 0.0001391 |
| ENSMUSG00000042213 | Zfand4 | 617.643881 | 1017.00641 | 399.362527 | 0.70089268 | 4.2735E-09 | 3.3957E-08 |
| ENSMUSG00000034748 | Sirt6 | 531.910984 | 870.792219 | 338.881234 | 0.70065271 | 4.1528E-14 | 5.9994E-13 |
| ENSMUSG00000060594 | Layn | 437.05691 | 730.940735 | 293.883825 | 0.70054093 | 2.9668E-05 | 0.0001337 |
| ENSMUSG00000036432 | Siah2 | 1745.46282 | 2854.98198 | 1109.51916 | 0.69956262 | 8.9725E-15 | 1.3847E-13 |
| ENSMUSG00000048763 | Hoxb3 | 67.7555014 | 114.291992 | 46.536491 | 0.69954331 | 7.3926E-05 | 0.00031145 |
| ENSMUSG00000040209 | Zfp704 | 1523.98512 | 2525.66388 | 1001.67876 | 0.69845538 | 1.2026E-06 | 6.7321E-06 |

| | | | | | | | |
|---|---|---|---|---|---|---|---|
| ENSMUSG00000037032 | Apbb1 | 1613.60535 | 2633.79532 | 1020.18997 | 0.69792959 | 2.0029E-18 | 4.4209E-17 |
| ENSMUSG00000024501 | Dpysl3 | 5356.72077 | 8769.06134 | 3412.34057 | 0.69786256 | 8.7125E-13 | 1.0797E-11 |
| ENSMUSG00000097331 | F420014N23 | 30.1843243 | 52.5785073 | 22.394183 | 0.69775519 | 0.00222359 | 0.00696462 |
| ENSMUSG00000032902 | Slc16a1 | 17922.505 | 29142.239 | 11219.734 | 0.69774234 | 7.1354E-40 | 7.0913E-38 |
| ENSMUSG00000019194 | Scn1b | 149.834892 | 250.482817 | 100.647925 | 0.69762012 | 3.6341E-05 | 0.00016119 |
| ENSMUSG00000075590 | Nrbp2 | 566.287192 | 929.454301 | 363.167109 | 0.69736963 | 4.6996E-11 | 4.7744E-10 |
| ENSMUSG00000062054 | Iah1 | 1280.17244 | 2106.77302 | 826.600577 | 0.69695095 | 2.7885E-08 | 1.9875E-07 |
| ENSMUSG00000085207 | - | 13.6951657 | 25.2644456 | 11.5692798 | 0.69679649 | 0.01962682 | 0.04781089 |
| ENSMUSG00000028680 | Plk3 | 699.410584 | 1151.08775 | 451.67717 | 0.69659489 | 2.8063E-08 | 1.999E-07 |
| ENSMUSG00000026471 | Mr1 | 650.891061 | 1072.24903 | 421.357973 | 0.69639669 | 5.7946E-08 | 3.9316E-07 |
| ENSMUSG00000030083 | Abtb1 | 658.37973 | 1075.41133 | 417.031602 | 0.69547657 | 2.0888E-15 | 3.471E-14 |
| ENSMUSG00000068015 | Lrch1 | 1437.14859 | 2337.69504 | 900.546455 | 0.69512569 | 1.0486E-21 | 3.0426E-20 |
| ENSMUSG00000056673 | Kdm5d | 3642.94877 | 5923.41785 | 2280.46907 | 0.69505697 | 2.9678E-25 | 1.1264E-23 |
| ENSMUSG00000031539 | Ap3m2 | 1052.65971 | 1726.96984 | 674.310124 | 0.69476194 | 6.1746E-09 | 4.8188E-08 |
| ENSMUSG00000024065 | Ehd3 | 130.306833 | 216.753141 | 86.4463086 | 0.69464062 | 3.0257E-06 | 1.5972E-05 |
| ENSMUSG00000027309 | Dnaaf9 | 2198.43719 | 3596.42559 | 1397.9884 | 0.69445752 | 3.5722E-11 | 3.7003E-10 |
| ENSMUSG00000001751 | Naglu | 2224.72269 | 3621.15344 | 1396.43075 | 0.69379794 | 6.8762E-17 | 1.3245E-15 |
| ENSMUSG00000026473 | Glul | 553.53053 | 900.209823 | 346.679293 | 0.69376064 | 1.1901E-15 | 2.0283E-14 |
| ENSMUSG00000034708 | Grn | 16937.0994 | 27463.2307 | 10526.1312 | 0.69352105 | 2.5138E-38 | 2.1913E-36 |
| ENSMUSG00000087593 | Gm16174 | 19.6184581 | 34.1706098 | 14.5521517 | 0.69297086 | 0.00817719 | 0.02221335 |
| ENSMUSG00000018476 | Kdm6b | 1654.22194 | 2728.06469 | 1073.84275 | 0.69247177 | 9.09E-07 | 5.1944E-06 |
| ENSMUSG00000095105 | Edaradd | 21.1342877 | 37.360083 | 16.2257953 | 0.69211793 | 0.01113924 | 0.0291696 |
| ENSMUSG00000015542 | Nat9 | 891.678178 | 1448.38753 | 556.709352 | 0.69181977 | 1.2801E-16 | 2.3963E-15 |
| ENSMUSG00000032014 | Oaf | 708.475039 | 1241.08945 | 532.614415 | 0.69109257 | 0.00682129 | 0.01895038 |
| ENSMUSG00000046311 | Zfp62 | 2965.43975 | 4823.81263 | 1858.37288 | 0.69108834 | 3.6159E-15 | 5.8526E-14 |
| ENSMUSG00000071324 | Armc2 | 260.547878 | 426.202434 | 165.654555 | 0.69099003 | 1.4498E-07 | 9.3061E-07 |
| ENSMUSG00000027470 | Mylk2 | 54.3274311 | 91.4924515 | 37.1650204 | 0.69097062 | 0.00079549 | 0.00274441 |
| ENSMUSG00000092626 | - | 22.5904187 | 41.246443 | 18.6560243 | 0.69085977 | 0.02012126 | 0.0488188 |
| ENSMUSG00000097404 | Gm10814 | 28.4776926 | 49.9605476 | 21.482855 | 0.6908272 | 0.00686223 | 0.01904799 |
| ENSMUSG00000090641 | Zfp712 | 166.84062 | 273.489874 | 106.649254 | 0.69010136 | 1.9653E-06 | 1.0647E-05 |
| ENSMUSG00000006462 | - | 30.2913174 | 52.6748875 | 22.3835701 | 0.69007333 | 0.00636831 | 0.01783647 |
| ENSMUSG00000049164 | Zfp518a | 754.927855 | 1256.27654 | 501.348681 | 0.6896629 | 6.792E-05 | 0.00028851 |
| ENSMUSG00000021366 | Hivep1 | 2827.43372 | 4621.44347 | 1794.00974 | 0.68927602 | 4.4847E-09 | 3.5567E-08 |
| ENSMUSG00000012126 | Ubxn11 | 29.809618 | 52.3706405 | 22.5610224 | 0.68926508 | 0.00713239 | 0.01970025 |
| ENSMUSG00000022108 | Itm2b | 28315.9961 | 46486.0951 | 18170.099 | 0.68920693 | 3.9811E-07 | 2.4074E-06 |
| ENSMUSG00000039850 | Endov | 1087.36846 | 1758.33514 | 670.966672 | 0.68909238 | 2.2873E-33 | 1.4659E-31 |
| ENSMUSG00000063458 | Lrmda | 396.760569 | 645.257523 | 248.496955 | 0.68888389 | 8.8666E-13 | 1.0968E-11 |
| ENSMUSG00000032000 | Birc3 | 1365.87453 | 2225.88279 | 860.008262 | 0.68875608 | 1.4275E-10 | 1.3826E-09 |
| ENSMUSG00000039191 | Rbpj | 8705.37473 | 14109.344 | 5403.9693 | 0.68874861 | 1.4193E-19 | 3.4554E-18 |
| ENSMUSG00000034007 | Scaper | 1276.30537 | 2068.11752 | 791.812144 | 0.68809031 | 2.4185E-22 | 7.36E-21 |
| ENSMUSG00000026979 | Psd4 | 118.745087 | 200.750585 | 82.0054977 | 0.68769693 | 0.0009414 | 0.00319991 |
| ENSMUSG00000023191 | P3h3 | 901.638327 | 1478.96111 | 577.322787 | 0.68761904 | 5.6275E-07 | 3.32E-06 |
| ENSMUSG00000082419 | - | 61.7656123 | 108.144135 | 46.3785228 | 0.68710968 | 0.00771374 | 0.02110481 |
| ENSMUSG00000021319 | Sfrp4 | 2254.89454 | 3927.69141 | 1672.79687 | 0.68647401 | 0.006683 | 0.01861395 |
| ENSMUSG00000029699 | Ssc4d | 125.366398 | 210.555708 | 85.1893099 | 0.68645189 | 0.00079793 | 0.00275235 |
| ENSMUSG00000041609 | Bicdl1 | 233.62169 | 378.825474 | 145.203785 | 0.68611195 | 2.8945E-10 | 2.6919E-09 |
| ENSMUSG00000020605 | Hs1bp3 | 1774.35054 | 2868.1895 | 1093.83896 | 0.6859547 | 4.2524E-20 | 1.08E-18 |
| ENSMUSG00000097729 | - | 330.75247 | 541.918834 | 211.166364 | 0.6859541 | 3.9523E-07 | 2.3907E-06 |
| ENSMUSG00000028337 | Coro2a | 109.123383 | 184.331039 | 75.2076562 | 0.68570122 | 0.00112495 | 0.00376299 |
| ENSMUSG00000098050 | - | 788.138368 | 1275.86965 | 487.731283 | 0.68561172 | 2.5355E-17 | 5.0318E-16 |
| ENSMUSG00000020889 | Nr1d1 | 2698.612 | 4799.13802 | 2100.52602 | 0.68495797 | 0.0127402 | 0.03276552 |
| ENSMUSG00000019944 | Rhobtb1 | 860.747078 | 1421.1306 | 560.383518 | 0.6848975 | 2.3384E-05 | 0.00010716 |
| ENSMUSG00000051499 | Zfp786 | 69.7955164 | 116.988207 | 47.1926903 | 0.6844314 | 0.00076244 | 0.00264057 |
| ENSMUSG00000020641 | Rsad2 | 4460.63645 | 7871.34543 | 3410.70898 | 0.68421945 | 0.01081729 | 0.02842061 |
| ENSMUSG00000028104 | Polr3gl | 746.428129 | 1203.2631 | 456.834973 | 0.68362618 | 6.6441E-23 | 2.1191E-21 |
| ENSMUSG00000020831 | 0610010K14 | 26.8031319 | 46.4672581 | 19.6641262 | 0.68326196 | 0.00539843 | 0.01540312 |
| ENSMUSG00000031502 | Col4a1 | 9637.23004 | 16975.8027 | 7338.57266 | 0.68313487 | 0.01069481 | 0.02815633 |
| ENSMUSG00000044576 | Garem2 | 133.911895 | 224.26856 | 90.3566652 | 0.68260627 | 0.00070385 | 0.0024521 |
| ENSMUSG00000031214 | Ophn1 | 638.806688 | 1032.90116 | 394.094472 | 0.68201663 | 1.64E-16 | 3.0205E-15 |
| ENSMUSG00000002885 | Adgre5 | 500.411811 | 822.017252 | 321.605441 | 0.68149355 | 8.6501E-06 | 4.246E-05 |
| ENSMUSG00000046341 | - | 661.253812 | 1075.96209 | 414.708279 | 0.68136589 | 9.1774E-08 | 6.0473E-07 |
| ENSMUSG00000030780 | Rusf1 | 521.240069 | 838.610032 | 317.369962 | 0.68049667 | 2.2909E-21 | 6.4915E-20 |
| ENSMUSG00000024678 | Ms4a4d | 135.326481 | 223.942262 | 88.6157814 | 0.68038915 | 0.00016978 | 0.00066981 |
| ENSMUSG00000060519 | Tor3a | 7156.48515 | 11717.9366 | 4561.45147 | 0.68031928 | 4.61E-06 | 2.3591E-05 |
| ENSMUSG00000073968 | Trim68 | 174.623182 | 285.153798 | 110.530616 | 0.67996813 | 5.9223E-08 | 4.0143E-07 |
| ENSMUSG00000032733 | Snx33 | 2076.60795 | 3334.27222 | 1257.66427 | 0.67975692 | 5.4736E-45 | 6.6237E-43 |
| ENSMUSG00000025920 | Stau2 | 1634.96301 | 2651.31705 | 1016.35404 | 0.67963388 | 3.8268E-09 | 3.0632E-08 |
| ENSMUSG00000028555 | Ttc39a | 67.5740518 | 114.034345 | 46.4602935 | 0.67962369 | 0.00151444 | 0.00492492 |
| ENSMUSG00000000567 | Sox9 | 782.947807 | 1266.58854 | 483.640731 | 0.67907136 | 5.4967E-11 | 5.5404E-10 |
| ENSMUSG00000021550 | 2210016F16I | 1199.5958 | 1959.59909 | 760.003286 | 0.67888414 | 2.9344E-06 | 1.5523E-05 |
| ENSMUSG00000035692 | Isg15 | 14848.3078 | 24002.8947 | 9154.58695 | 0.67839334 | 8.0809E-11 | 8E-10 |
| ENSMUSG00000051285 | Pcmtd1 | 2604.36112 | 4183.80258 | 1579.44146 | 0.67800694 | 1.2337E-23 | 4.1269E-22 |
| ENSMUSG00000026142 | Rhbdd1 | 3599.68711 | 5783.90445 | 2184.21734 | 0.67532112 | 1.6326E-16 | 3.0095E-15 |
| ENSMUSG00000061353 | Cxcl12 | 5316.84284 | 8538.72789 | 3221.88506 | 0.67418438 | 2.2292E-15 | 3.6954E-14 |
| ENSMUSG00000068250 | Amn1 | 1233.07522 | 1989.67581 | 756.600585 | 0.67384472 | 1.9363E-09 | 1.6088E-08 |

| | | | | | | | |
|---|---|---|---|---|---|---|---|
| ENSMUSG00000029510 | Gpc2 | 367.224722 | 599.430173 | 232.205451 | 0.67268088 | 1.6738E-05 | 7.8561E-05 |
| ENSMUSG00000071691 | Gm960 | 24.4817946 | 43.5266522 | 19.0448576 | 0.67233345 | 0.01561847 | 0.03919288 |
| ENSMUSG00000020251 | Glt8d2 | 264.094335 | 430.856784 | 166.762448 | 0.67204814 | 2.044E-05 | 9.4726E-05 |
| ENSMUSG00000063558 | Aox1 | 912.42902 | 1473.25105 | 560.822032 | 0.67185785 | 6.9158E-09 | 5.3687E-08 |
| ENSMUSG00000057054 | Inca1 | 99.7774746 | 162.922785 | 63.1453102 | 0.67156784 | 4.8051E-05 | 0.00020903 |
| ENSMUSG00000072763 | 5430403G16 | 219.269742 | 354.173754 | 134.904012 | 0.67133807 | 9.1357E-08 | 6.0237E-07 |
| ENSMUSG00000039233 | Tbce | 1545.01227 | 2491.52471 | 946.51244 | 0.67086357 | 1.0275E-08 | 7.7943E-08 |
| ENSMUSG00000063698 | Sfxn4 | 340.18533 | 550.211952 | 210.026622 | 0.67070416 | 6.3476E-07 | 3.7168E-06 |
| ENSMUSG00000097928 | - | 511.818176 | 825.937237 | 314.119061 | 0.67013302 | 2.3798E-08 | 1.7148E-07 |
| ENSMUSG00000087543 | Gm16576 | 93.2474372 | 152.774398 | 59.5269605 | 0.67005548 | 0.00025075 | 0.00095892 |
| ENSMUSG00000016239 | Lonrf3 | 351.419994 | 565.470295 | 214.050301 | 0.66993115 | 6.0074E-09 | 4.6937E-08 |
| ENSMUSG00000089736 | Tgfbr3l | 27.5928193 | 48.1848931 | 20.5920738 | 0.66978962 | 0.0126219 | 0.03249377 |
| ENSMUSG00000008429 | Herpud2 | 1788.25037 | 2878.04587 | 1089.7955 | 0.66977945 | 2.2013E-09 | 1.8194E-08 |
| ENSMUSG00000024424 | Ttc39c | 3745.7711 | 5994.36256 | 2248.59147 | 0.66947757 | 1.2425E-15 | 2.1142E-14 |
| ENSMUSG00000040841 | Six5 | 479.717834 | 776.523778 | 296.805944 | 0.6694164 | 8.8356E-07 | 5.0702E-06 |
| ENSMUSG00000079055 | Slc8a3 | 24.4655438 | 42.1140126 | 17.6484687 | 0.66892457 | 0.01173689 | 0.03052868 |
| ENSMUSG00000038797 | Zscan2 | 290.633257 | 469.722171 | 179.088914 | 0.66885383 | 2.8076E-07 | 1.7329E-06 |
| ENSMUSG00000028028 | Alpk1 | 1169.1265 | 1891.17782 | 722.051313 | 0.66875799 | 4.9375E-07 | 2.9357E-06 |
| ENSMUSG00000066829 | Zfp810 | 366.758534 | 589.619399 | 222.860865 | 0.66849162 | 2.4615E-09 | 2.0191E-08 |
| ENSMUSG00000060301 | 2610008E11 | 978.657132 | 1564.41897 | 585.761842 | 0.66813661 | 7.6214E-15 | 1.1878E-13 |
| ENSMUSG00000021469 | Msx2 | 200.309989 | 322.198848 | 121.888858 | 0.66765822 | 6.375E-08 | 4.2985E-07 |
| ENSMUSG00000037336 | Mfsd2b | 19.7486001 | 34.3390834 | 14.5904833 | 0.66682574 | 0.01021143 | 0.02699775 |
| ENSMUSG00000052139 | Babam2 | 1882.69618 | 3007.77601 | 1125.07983 | 0.6668118 | 6.8307E-15 | 1.071E-13 |
| ENSMUSG00000028167 | Bdh2 | 395.377867 | 637.954674 | 242.576807 | 0.66664989 | 6.3792E-07 | 3.7335E-06 |
| ENSMUSG00000046572 | Zfp518b | 1016.25706 | 1630.81761 | 614.560548 | 0.66565617 | 8.461E-10 | 7.3257E-09 |
| ENSMUSG00000004558 | Ndrg2 | 1235.78153 | 2015.34858 | 779.567052 | 0.6656258 | 7.2434E-05 | 0.00030592 |
| ENSMUSG00000024143 | Rhoq | 5459.37611 | 8690.16491 | 3230.7888 | 0.66531397 | 2.338E-25 | 8.9399E-24 |
| ENSMUSG00000038521 | C1s1 | 14850.1055 | 23590.3584 | 8740.25287 | 0.66507136 | 7.9465E-47 | 1.0755E-44 |
| ENSMUSG00000024727 | Trpm6 | 173.383801 | 278.532544 | 105.148743 | 0.66481114 | 1.1011E-09 | 9.4112E-09 |
| ENSMUSG00000086554 | 9530034E10 | 17.7652113 | 31.2316075 | 13.4663962 | 0.66481052 | 0.02041746 | 0.04942716 |
| ENSMUSG00000036561 | Ppp6r2 | 750.041592 | 1197.72831 | 447.686719 | 0.66456203 | 1.0486E-13 | 1.4448E-12 |
| ENSMUSG00000064080 | Fbln2 | 24301.1915 | 40818.9928 | 16517.8012 | 0.66412698 | 0.00370681 | 0.01102291 |
| ENSMUSG00000020173 | Cobl | 221.811741 | 375.85824 | 154.046499 | 0.66407702 | 0.00589002 | 0.01664417 |
| ENSMUSG00000028051 | Hcn3 | 20.0225937 | 34.8431579 | 14.8205642 | 0.66400774 | 0.01548681 | 0.03890995 |
| ENSMUSG00000028811 | Yars | 10443.437 | 16641.578 | 6198.14091 | 0.66371542 | 3.0033E-16 | 5.4148E-15 |
| ENSMUSG00000000901 | Mmp11 | 1150.51947 | 1831.76632 | 681.246849 | 0.66362983 | 9.227E-16 | 1.5911E-14 |
| ENSMUSG00000050910 | Cdr2l | 1239.46379 | 1975.05714 | 735.593346 | 0.66322702 | 9.8105E-16 | 1.6832E-14 |
| ENSMUSG00000070639 | Lrrc8b | 271.100632 | 447.135704 | 176.035072 | 0.66320094 | 0.00081961 | 0.00282059 |
| ENSMUSG00000027420 | Bfsp1 | 70.3802615 | 123.643566 | 53.2633044 | 0.66202355 | 0.01864089 | 0.04573409 |
| ENSMUSG00000022999 | Lmbr1l | 648.260895 | 1031.96496 | 383.704067 | 0.66192764 | 1.2035E-16 | 2.2631E-15 |
| ENSMUSG00000002489 | Tiam1 | 745.747662 | 1196.40564 | 450.657981 | 0.66160493 | 1.7867E-08 | 1.3085E-07 |
| ENSMUSG00000002688 | Prkd1 | 1198.77291 | 1906.33704 | 707.564131 | 0.6615559 | 7.1531E-18 | 1.5031E-16 |
| ENSMUSG00000008734 | Gprc5b | 954.809907 | 1527.92322 | 573.113312 | 0.66031245 | 9.8258E-09 | 7.47E-08 |
| ENSMUSG00000026942 | Traf2 | 4329.28739 | 6885.96068 | 2556.67329 | 0.66025893 | 1.5498E-14 | 2.3339E-13 |
| ENSMUSG00000028197 | Col24a1 | 2539.81579 | 4078.30473 | 1538.48894 | 0.65999842 | 4.8632E-07 | 2.8932E-06 |
| ENSMUSG00000062031 | Pgghg | 1626.37549 | 2583.18275 | 956.807263 | 0.65991055 | 4.2468E-19 | 1.0018E-17 |
| ENSMUSG00000052934 | Fbxo31 | 1815.90568 | 2888.53723 | 1072.63155 | 0.65985459 | 1.9715E-14 | 2.9284E-13 |
| ENSMUSG00000022419 | Deptor | 1449.23525 | 2400.49581 | 951.260564 | 0.65976852 | 0.00178459 | 0.00570425 |
| ENSMUSG00000018417 | Myo1b | 4424.83664 | 7059.08323 | 2634.24659 | 0.65964595 | 2.3014E-10 | 2.1688E-09 |
| ENSMUSG00000032402 | Smad3 | 2982.52708 | 4802.92851 | 1820.40144 | 0.65958514 | 3.8289E-06 | 1.9876E-05 |
| ENSMUSG00000055809 | Dnaaf3 | 72.9769845 | 119.489596 | 46.5126113 | 0.65925245 | 0.00062033 | 0.0021848 |
| ENSMUSG00000025241 | Fyco1 | 3429.28301 | 5471.39049 | 2042.10748 | 0.65908612 | 4.0036E-10 | 3.6492E-09 |
| ENSMUSG00000024883 | Rin1 | 3870.71818 | 6144.83907 | 2274.1209 | 0.65907358 | 3.1692E-18 | 6.8918E-17 |
| ENSMUSG00000050270 | Tmem220 | 104.016444 | 167.866702 | 63.8502577 | 0.65887377 | 3.7388E-05 | 0.00016541 |
| ENSMUSG00000061393 | Acvr2b | 31.0319086 | 52.7755652 | 21.7436567 | 0.65863887 | 0.00951334 | 0.02535783 |
| ENSMUSG00000097058 | - | 25.1058528 | 41.9378388 | 16.8319861 | 0.6577318 | 0.006443 | 0.01802114 |
| ENSMUSG00000030486 | Zfp108 | 94.8190179 | 155.621665 | 60.8026466 | 0.65727389 | 0.00068172 | 0.00238225 |
| ENSMUSG00000062980 | Cped1 | 171.623285 | 277.972406 | 106.349121 | 0.65658364 | 4.208E-05 | 0.0001847 |
| ENSMUSG00000042506 | Usp22 | 9863.72073 | 15583.3899 | 5719.66916 | 0.65638718 | 1.9173E-38 | 1.6856E-36 |
| ENSMUSG00000029136 | Rbks | 69.101567 | 112.786015 | 43.684448 | 0.6563858 | 0.00054881 | 0.00195466 |
| ENSMUSG00000034771 | Tle2 | 406.281094 | 650.585249 | 244.304155 | 0.65617472 | 2.2879E-07 | 1.4311E-06 |
| ENSMUSG00000070003 | Ssbp4 | 1718.14533 | 2731.42117 | 1013.27584 | 0.65586599 | 8.7094E-11 | 8.6015E-10 |
| ENSMUSG00000059772 | Slx1b | 409.276548 | 652.262128 | 242.985581 | 0.65579727 | 7.3442E-10 | 6.4155E-09 |
| ENSMUSG00000032905 | Atg12 | 5810.94417 | 9201.24847 | 3390.30429 | 0.65573243 | 5.9105E-18 | 1.2522E-16 |
| ENSMUSG00000022092 | Ppp3cc | 586.877475 | 933.321745 | 346.44427 | 0.65563198 | 4.0366E-10 | 3.676E-09 |
| ENSMUSG00000030872 | Gga2 | 1865.06196 | 2950.28886 | 1085.2269 | 0.65559843 | 2.3232E-22 | 7.0909E-21 |
| ENSMUSG00000062470 | Fbxl12os | 112.601563 | 182.052122 | 69.4505592 | 0.6549872 | 2.4087E-05 | 0.00011014 |
| ENSMUSG00000054582 | Pabpc1l | 73.083856 | 121.164384 | 48.0805276 | 0.65461734 | 0.00316389 | 0.00956045 |
| ENSMUSG00000087022 | 9130024F11 | 24.7996665 | 41.8439591 | 17.0442927 | 0.65364232 | 0.00829308 | 0.02250135 |
| ENSMUSG00000042404 | Dennd4b | 457.64405 | 724.647433 | 267.003383 | 0.65328446 | 5.3837E-12 | 6.1291E-11 |
| ENSMUSG00000071359 | Tbpl1 | 3419.83909 | 5403.30346 | 1983.46437 | 0.65316334 | 1.3303E-18 | 3.0106E-17 |
| ENSMUSG00000021756 | Il6st | 22725.103 | 36017.5746 | 13292.4716 | 0.65222493 | 2.0165E-11 | 2.1561E-10 |
| ENSMUSG00000053411 | Cbx7 | 587.390714 | 930.239475 | 342.848761 | 0.65219732 | 1.2568E-11 | 1.3738E-10 |
| ENSMUSG00000030041 | M1ap | 54.5486243 | 90.4986791 | 35.9500549 | 0.65173727 | 0.00356677 | 0.01065576 |
| ENSMUSG00000057554 | Lgals8 | 6734.46081 | 10709.1107 | 3974.64992 | 0.65163513 | 2.0658E-08 | 1.4996E-07 |

| | | | | | | | |
|---|---|---|---|---|---|---|---|
| ENSMUSG00000073374 | - | 49.2045501 | 84.9584425 | 35.7538924 | 0.65138314 | 0.0177509 | 0.04383285 |
| ENSMUSG00000026958 | Dpp7 | 1721.23195 | 2715.10953 | 993.877581 | 0.6513765 | 1.146E-18 | 2.6051E-17 |
| ENSMUSG00000053091 | Lins1 | 810.476341 | 1281.16251 | 470.686167 | 0.65108677 | 2.7357E-14 | 4.0199E-13 |
| ENSMUSG00000040420 | Cdh18 | 52.8647109 | 90.3201779 | 37.455467 | 0.65099489 | 0.0128491 | 0.0330167 |
| ENSMUSG00000090576 | - | 46.6545904 | 77.3445648 | 30.6899744 | 0.65059614 | 0.00592963 | 0.01674229 |
| ENSMUSG00000028100 | Nudt17 | 110.519263 | 179.218348 | 68.6990843 | 0.65031246 | 0.00036522 | 0.0013489 |
| ENSMUSG00000025403 | Shmt2 | 16438.6961 | 26044.1304 | 9605.43432 | 0.64978564 | 5.4519E-10 | 4.8532E-09 |
| ENSMUSG00000040415 | Dtx3 | 1824.64101 | 2874.49071 | 1049.8497 | 0.64910707 | 1.4843E-18 | 3.3335E-17 |
| ENSMUSG00000029173 | Sepsecs | 1768.31694 | 2807.08449 | 1038.76755 | 0.64857825 | 3.4551E-08 | 2.4308E-07 |
| ENSMUSG00000002325 | Irf9 | 9355.73527 | 14749.2078 | 5393.47249 | 0.64849122 | 1.4007E-15 | 2.3696E-14 |
| ENSMUSG00000034462 | Pkd2 | 6237.9216 | 9893.6053 | 3655.6837 | 0.64848429 | 1.3049E-08 | 9.73E-08 |
| ENSMUSG00000063849 | Ppcdc | 845.834889 | 1333.59105 | 487.756162 | 0.64829416 | 8.3487E-14 | 1.1597E-12 |
| ENSMUSG00000057137 | Tmem140 | 726.237927 | 1145.68177 | 419.443847 | 0.64788064 | 1.7133E-13 | 2.3006E-12 |
| ENSMUSG00000064141 | Zfp69 | 56.7038011 | 93.6531798 | 36.9493787 | 0.64696655 | 0.00448804 | 0.01308317 |
| ENSMUSG00000097347 | - | 101.813133 | 170.257531 | 68.4443982 | 0.64695315 | 0.00768148 | 0.02101934 |
| ENSMUSG00000020905 | Usp43 | 91.753486 | 147.17512 | 55.4216338 | 0.64680151 | 9.6436E-05 | 0.00039813 |
| ENSMUSG00000038387 | Rras | 2545.89977 | 4013.60941 | 1467.70964 | 0.64664064 | 1.7311E-12 | 2.0801E-11 |
| ENSMUSG00000000296 | Tpd52l1 | 2278.99891 | 3590.31312 | 1311.31421 | 0.64656154 | 1.2475E-13 | 1.7007E-12 |
| ENSMUSG00000044719 | E230025N22 | 96.1576542 | 155.887342 | 59.7296883 | 0.64650839 | 0.00052483 | 0.00187772 |
| ENSMUSG00000027478 | Dnmt3b | 76.7664472 | 128.986961 | 52.2205134 | 0.64642465 | 0.0086004 | 0.02323712 |
| ENSMUSG00000023411 | Nfatc4 | 1315.99772 | 2069.20544 | 753.207712 | 0.6463926 | 1.5955E-18 | 3.5665E-17 |
| ENSMUSG00000053216 | Btn2a2 | 108.875094 | 185.356762 | 76.4816681 | 0.64616795 | 0.01451264 | 0.03678133 |
| ENSMUSG00000032449 | Slc25a36 | 2354.90816 | 3702.26153 | 1347.35337 | 0.6457864 | 5.4795E-20 | 1.3797E-18 |
| ENSMUSG00000026004 | Kansl1l | 960.160915 | 1528.86582 | 568.704908 | 0.64567389 | 3.1426E-06 | 1.6543E-05 |
| ENSMUSG00000097039 | - | 838.540015 | 1336.69179 | 498.151773 | 0.64555994 | 5.8906E-06 | 2.968E-05 |
| ENSMUSG00000037992 | Rara | 1144.2146 | 1794.86074 | 650.646141 | 0.64552808 | 1.8684E-26 | 7.6414E-25 |
| ENSMUSG00000048264 | Dip2c | 1577.94115 | 2484.13208 | 906.190925 | 0.64539972 | 2.1578E-15 | 3.5798E-14 |
| ENSMUSG00000059669 | Taf1b | 1451.92895 | 2289.51103 | 837.58208 | 0.64493978 | 9.875E-11 | 9.6922E-10 |
| ENSMUSG00000028226 | Mmp16 | 408.375323 | 646.584991 | 238.209668 | 0.64450547 | 1.2906E-08 | 9.6375E-08 |
| ENSMUSG00000075122 | Cd80 | 26.2403868 | 44.9065608 | 18.666174 | 0.64384427 | 0.02023753 | 0.0490546 |
| ENSMUSG00000049225 | Pdp1 | 1301.19131 | 2040.22305 | 739.031742 | 0.64372583 | 2.8433E-20 | 7.2934E-19 |
| ENSMUSG00000020656 | Grhl1 | 116.653025 | 189.185442 | 72.5324169 | 0.64340086 | 0.0009481 | 0.00321959 |
| ENSMUSG00000070427 | Il18bp | 128.907349 | 205.897131 | 76.9897826 | 0.64311751 | 9.5887E-06 | 4.6722E-05 |
| ENSMUSG00000034334 | Fam151b | 126.404784 | 201.141097 | 74.7363121 | 0.64269755 | 5.4061E-05 | 0.0002332 |
| ENSMUSG00000056271 | Lman1l | 190.123804 | 306.339898 | 116.216094 | 0.64205055 | 0.00046248 | 0.00167472 |
| ENSMUSG00000030527 | Crtc3 | 981.315761 | 1543.67162 | 562.355864 | 0.64200399 | 1.0849E-13 | 1.4938E-12 |
| ENSMUSG00000074141 | Il4i1; Il4i1b | 54.6843938 | 90.1295769 | 35.4451831 | 0.6414823 | 0.00402228 | 0.0118786 |
| ENSMUSG00000034636 | Zyg11b | 4879.41533 | 7643.38499 | 2763.96966 | 0.64077298 | 1.7468E-19 | 4.2226E-18 |
| ENSMUSG00000056267 | Cep70 | 536.459521 | 841.550361 | 305.09084 | 0.64061221 | 2.9497E-18 | 6.4485E-17 |
| ENSMUSG00000032712 | Resf1 | 5470.73338 | 8671.28173 | 3200.54835 | 0.64054334 | 2.1576E-06 | 1.1626E-05 |
| ENSMUSG00000021180 | Rps6ka5 | 551.529241 | 864.975301 | 313.44606 | 0.64048823 | 2.7352E-15 | 4.4942E-14 |
| ENSMUSG00000056648 | Hoxb8 | 442.900427 | 699.881013 | 256.980586 | 0.6404057 | 3.6398E-07 | 2.2127E-06 |
| ENSMUSG00000037938 | Chchd5 | 717.466591 | 1128.48732 | 411.020734 | 0.63975576 | 2.9843E-10 | 2.7705E-09 |
| ENSMUSG00000028412 | Slc44a1 | 3081.97946 | 4818.8009 | 1736.82144 | 0.63964586 | 2.5063E-23 | 8.2365E-22 |
| ENSMUSG00000023341 | Mx2 | 2470.67413 | 3919.51504 | 1448.84091 | 0.63949443 | 6.156E-06 | 3.0948E-05 |
| ENSMUSG00000001995 | Sipa1l2 | 1652.94712 | 2632.76886 | 979.821739 | 0.63908732 | 3.842E-05 | 0.00016965 |
| ENSMUSG00000027173 | Depdc7 | 48.3671585 | 82.7203293 | 34.3531707 | 0.638267 | 0.01867114 | 0.0457974 |
| ENSMUSG00000097188 | - | 8310.83203 | 13131.2725 | 4820.44051 | 0.63813644 | 9.2831E-07 | 5.2945E-06 |
| ENSMUSG00000030854 | Ptpn5 | 2852.70259 | 4472.79557 | 1620.09298 | 0.63794199 | 9.2406E-12 | 1.027E-10 |
| ENSMUSG00000009654 | Oit3 | 71.672565 | 119.133482 | 47.4609169 | 0.63715227 | 0.00968164 | 0.02576105 |
| ENSMUSG00000040848 | Sft2d2 | 2618.70524 | 4086.98512 | 1468.27988 | 0.63670471 | 5.862E-23 | 1.8726E-21 |
| ENSMUSG00000015980 | Lrrc27 | 173.037031 | 279.095378 | 106.058347 | 0.63639417 | 0.00087777 | 0.00300107 |
| ENSMUSG00000013495 | Tmem175 | 1040.74613 | 1624.81791 | 584.071782 | 0.63601146 | 1.7951E-21 | 5.122E-20 |
| ENSMUSG00000021460 | Auh | 1749.29529 | 2728.44287 | 979.147584 | 0.63599256 | 4.2656E-19 | 1.0033E-17 |
| ENSMUSG00000086725 | - | 73.6516974 | 119.629965 | 45.9782675 | 0.63597368 | 0.00161272 | 0.00520128 |
| ENSMUSG00000029082 | Bst1 | 63.9001604 | 107.317392 | 43.4172315 | 0.63535402 | 0.01393541 | 0.0354494 |
| ENSMUSG00000023186 | Vwa5a | 3823.20929 | 6024.65495 | 2201.44566 | 0.63517617 | 6.3124E-07 | 3.6976E-06 |
| ENSMUSG00000002006 | Pdzd4 | 193.331594 | 304.893386 | 111.561791 | 0.6342178 | 6.764E-08 | 4.5384E-07 |
| ENSMUSG00000067297 | Ifit1bl2 | 953.127831 | 1518.03335 | 564.905518 | 0.63404899 | 0.00013609 | 0.0005468 |
| ENSMUSG00000039126 | Prune2 | 5874.9117 | 9228.29982 | 3353.38812 | 0.63359141 | 7.6117E-08 | 5.0799E-07 |
| ENSMUSG00000026938 | Fcna | 71.51725 | 120.652593 | 49.1353433 | 0.63315654 | 0.01834819 | 0.0451236 |
| ENSMUSG00000034647 | Ankrd12 | 1781.20612 | 2793.14899 | 1011.94286 | 0.63278896 | 1.6815E-08 | 1.235E-07 |
| ENSMUSG00000061288 | Taok3 | 2136.28961 | 3328.7378 | 1192.44819 | 0.63238637 | 9.9168E-17 | 1.8785E-15 |
| ENSMUSG00000001558 | Klhl10 | 36.0030782 | 59.452821 | 23.4497428 | 0.6323764 | 0.00701235 | 0.01939815 |
| ENSMUSG00000071855 | Ccdc112 | 1027.1646 | 1595.74282 | 568.578216 | 0.63189927 | 2.5093E-24 | 8.8391E-23 |
| ENSMUSG00000087142 | - | 50.9692003 | 83.2322236 | 32.2630233 | 0.63116166 | 0.007611 | 0.0208459 |
| ENSMUSG00000032298 | Neil1 | 404.446401 | 634.128293 | 229.681892 | 0.63004414 | 9.2992E-07 | 5.3022E-06 |
| ENSMUSG00000079608 | Stard6 | 150.862003 | 240.177006 | 89.3150028 | 0.62933916 | 0.0003323 | 0.00123729 |
| ENSMUSG00000043004 | Gng2 | 2669.23219 | 4164.64456 | 1495.41237 | 0.62922825 | 7.1762E-10 | 6.274E-09 |
| ENSMUSG00000083649 | Rasl2-9 | 25.6028413 | 42.3853304 | 16.7824891 | 0.62878604 | 0.01170489 | 0.03046086 |
| ENSMUSG00000054752 | Fsd1l | 528.851618 | 826.393783 | 297.542165 | 0.62875516 | 8.1359E-09 | 6.2522E-08 |
| ENSMUSG00000001323 | Srr | 954.033333 | 1480.07158 | 526.038248 | 0.62792423 | 3.1316E-21 | 8.789E-20 |
| ENSMUSG00000024014 | Pim1 | 1309.60602 | 2057.3664 | 747.760375 | 0.62786227 | 3.7157E-06 | 1.9342E-05 |
| ENSMUSG00000052676 | Zmat1 | 933.372597 | 1448.93739 | 515.564789 | 0.627853 | 4.9539E-16 | 8.7329E-15 |
| ENSMUSG00000042444 | Mindy2 | 1334.70686 | 2082.49829 | 747.791431 | 0.62776358 | 2.2659E-09 | 1.869E-08 |

| | | | | | | | |
|---|---|---|---|---|---|---|---|
| ENSMUSG00000028549 | Itgb3bp | 465.275437 | 732.70104 | 267.425603 | 0.62769316 | 3.263E-05 | 0.0001459 |
| ENSMUSG00000002020 | Ltbp2 | 23370.917 | 36360.986 | 12990.069 | 0.62746581 | 5.2024E-12 | 5.9358E-11 |
| ENSMUSG00000057143 | Trim12c | 3820.85589 | 5965.94462 | 2145.08873 | 0.62723762 | 1.7928E-08 | 1.3116E-07 |
| ENSMUSG00000072872 | Rybp; Rybp-p | 1451.28226 | 2252.34902 | 801.066765 | 0.62605955 | 1.8393E-15 | 3.0663E-14 |
| ENSMUSG00000061143 | Maml3 | 730.552644 | 1196.86764 | 466.314996 | 0.62587948 | 0.00805378 | 0.02191276 |
| ENSMUSG00000052525 | Spdya | 48.5753955 | 78.1266755 | 29.55128 | 0.62541607 | 0.00163353 | 0.00526146 |
| ENSMUSG00000006403 | Adamts4 | 1432.52212 | 2245.06068 | 812.538565 | 0.6252567 | 2.6425E-06 | 1.4072E-05 |
| ENSMUSG00000039349 | C130074G19 | 27.3584477 | 46.3607454 | 19.0022978 | 0.62511764 | 0.01630141 | 0.04065374 |
| ENSMUSG00000023961 | Enpp4 | 3043.4468 | 4708.85821 | 1665.41141 | 0.62425925 | 5.634E-21 | 1.5495E-19 |
| ENSMUSG00000042320 | Prox2 | 97.4275481 | 154.064581 | 56.6370329 | 0.62418189 | 0.00017487 | 0.00068864 |
| ENSMUSG00000070315 | 4930581F22I | 117.726777 | 186.852808 | 69.1260304 | 0.62407107 | 0.00036432 | 0.0013462 |
| ENSMUSG00000073591 | Pcdhb22 | 156.439817 | 253.226125 | 96.7863083 | 0.62381408 | 0.00386131 | 0.01144926 |
| ENSMUSG00000065954 | Tacc1 | 5851.08733 | 9055.02733 | 3203.94 | 0.62320071 | 1.548E-17 | 3.1251E-16 |
| ENSMUSG00000030287 | Itpr2 | 3431.28215 | 5340.57064 | 1909.28849 | 0.62227847 | 2.4925E-08 | 1.7887E-07 |
| ENSMUSG00000056938 | Acbd4 | 1408.11881 | 2174.48293 | 766.364119 | 0.62188837 | 1.1659E-19 | 2.8588E-18 |
| ENSMUSG00000029456 | Acad10 | 375.917918 | 583.065542 | 207.147624 | 0.62159756 | 2.1746E-11 | 2.3143E-10 |
| ENSMUSG00000098041 | - | 48.7228507 | 79.006628 | 30.2837774 | 0.62149586 | 0.00545221 | 0.01553502 |
| ENSMUSG00000044164 | Rnf182 | 167.894604 | 265.8611 | 97.9664959 | 0.62135887 | 0.00044996 | 0.00163428 |
| ENSMUSG00000025232 | Hexa | 5202.13423 | 8043.17637 | 2841.04214 | 0.62121802 | 1.1733E-14 | 1.7919E-13 |
| ENSMUSG00000048490 | Nrip1 | 1435.9283 | 2244.21244 | 808.284133 | 0.62120349 | 3.0186E-06 | 1.5939E-05 |
| ENSMUSG00000005534 | Insr | 1970.87116 | 3062.73853 | 1091.86736 | 0.62099947 | 7.7675E-09 | 5.9893E-08 |
| ENSMUSG00000031109 | Enox2 | 1472.52549 | 2276.30429 | 803.778799 | 0.62037556 | 2.1742E-13 | 2.8875E-12 |
| ENSMUSG00000085024 | C230035I16R | 33.4166695 | 54.796562 | 21.3798925 | 0.61991031 | 0.00861117 | 0.02326317 |
| ENSMUSG00000045854 | Lyrm2 | 516.083826 | 806.962632 | 290.878806 | 0.61964222 | 1.4348E-05 | 6.8151E-05 |
| ENSMUSG00000027894 | Slc6a17 | 1189.88129 | 1862.38022 | 672.498925 | 0.61958479 | 1.6339E-05 | 7.6848E-05 |
| ENSMUSG00000054967 | Zfp647 | 486.188031 | 751.427908 | 265.239878 | 0.61957558 | 3.1754E-12 | 3.7074E-11 |
| ENSMUSG00000030231 | Plekha5 | 1959.24743 | 3024.6062 | 1065.35878 | 0.61949894 | 1.4302E-17 | 2.9017E-16 |
| ENSMUSG00000097785 | - | 81.7970066 | 129.542281 | 47.7452749 | 0.61940275 | 0.00054729 | 0.00194991 |
| ENSMUSG00000029403 | Cdkl2 | 817.562111 | 1264.49314 | 446.931028 | 0.61896966 | 1.164E-14 | 1.779E-13 |
| ENSMUSG00000038539 | Atf5 | 3112.17607 | 4816.61853 | 1704.44246 | 0.61855146 | 2.1135E-10 | 2E-09 |
| ENSMUSG00000074863 | - | 37.1781959 | 60.5830594 | 23.4048635 | 0.61740097 | 0.00588021 | 0.01661927 |
| ENSMUSG00000026880 | Stom | 12712.2671 | 19763.9627 | 7051.69559 | 0.61703103 | 9.0028E-07 | 5.1575E-06 |
| ENSMUSG00000040624 | Plekhg1 | 1338.69671 | 2073.32554 | 734.62883 | 0.6167042 | 3.748E-09 | 3.0025E-08 |
| ENSMUSG00000029163 | Emilin1 | 670.915578 | 1056.14034 | 385.224764 | 0.61648629 | 0.00028302 | 0.00106948 |
| ENSMUSG00000028476 | Reck | 210.840864 | 337.584351 | 126.743487 | 0.61638721 | 0.0028824 | 0.0087808 |
| ENSMUSG00000041775 | Mapk1ip1 | 271.323505 | 421.858404 | 150.534899 | 0.61620818 | 3.7634E-08 | 2.6316E-07 |
| ENSMUSG00000033368 | Trim69 | 53.1059806 | 87.3472194 | 34.2412387 | 0.61616341 | 0.01387915 | 0.03532376 |
| ENSMUSG00000097267 | 1700109K24 | 63.9369182 | 105.815474 | 41.8785559 | 0.61583486 | 0.01555458 | 0.03905636 |
| ENSMUSG00000040883 | Tmem205 | 481.983743 | 754.27527 | 272.291527 | 0.61574808 | 9.2029E-05 | 0.00038147 |
| ENSMUSG00000047414 | Flrt2 | 485.478757 | 760.038673 | 274.559916 | 0.6155535 | 7.0316E-05 | 0.00029807 |
| ENSMUSG00000066191 | Anks6 | 425.686806 | 655.657535 | 229.970728 | 0.61555169 | 3.1863E-12 | 3.7159E-11 |
| ENSMUSG00000078580 | E430018J23I | 271.121239 | 420.142817 | 149.021578 | 0.6147179 | 1.0812E-07 | 7.0525E-07 |
| ENSMUSG00000032376 | Usp3 | 2557.94722 | 3931.38379 | 1373.43656 | 0.61398882 | 4.5218E-17 | 8.8006E-16 |
| ENSMUSG00000020886 | Dlg4 | 942.889498 | 1447.40001 | 504.510516 | 0.61332679 | 8.7763E-22 | 2.5787E-20 |
| ENSMUSG00000028480 | Glipr2 | 3700.05883 | 5702.6443 | 2002.58547 | 0.61308715 | 2.8892E-10 | 2.6882E-09 |
| ENSMUSG00000097193 | - | 20.9681272 | 34.3072044 | 13.3390772 | 0.61251596 | 0.02003463 | 0.04864681 |
| ENSMUSG00000062101 | Zfp119b | 500.976894 | 773.683337 | 272.706443 | 0.6117502 | 4.2561E-07 | 2.5586E-06 |
| ENSMUSG00000057367 | Birc2 | 1937.50692 | 2984.10914 | 1046.60222 | 0.61072404 | 1.7676E-09 | 1.4764E-08 |
| ENSMUSG00000064246 | Chil1 | 395.764238 | 626.116048 | 230.35181 | 0.61061708 | 0.0017411 | 0.00558085 |
| ENSMUSG00000021904 | Sema3g | 372.02752 | 575.907146 | 203.879626 | 0.6103987 | 1.4172E-06 | 7.8585E-06 |
| ENSMUSG00000061461 | Smim20 | 1498.14957 | 2302.27117 | 804.121597 | 0.60937524 | 3.7784E-10 | 3.4516E-09 |
| ENSMUSG00000003617 | Cp | 22714.532 | 34752.6947 | 12038.1627 | 0.60917352 | 1.1137E-23 | 3.7316E-22 |
| ENSMUSG00000044348 | Slc25a53 | 375.251575 | 579.506118 | 204.254544 | 0.60881425 | 1.08E-06 | 6.0956E-06 |
| ENSMUSG00000097757 | - | 34.3616718 | 56.0580782 | 21.6964063 | 0.60838127 | 0.00867268 | 0.02340473 |
| ENSMUSG00000027215 | Cd82 | 2030.06361 | 3098.44261 | 1068.379 | 0.6080551 | 1.9721E-36 | 1.5485E-34 |
| ENSMUSG00000051735 | Rinl | 123.739441 | 191.002069 | 67.262628 | 0.60796877 | 1.1979E-05 | 5.7562E-05 |
| ENSMUSG00000024426 | Atat1 | 395.822343 | 606.674061 | 210.851718 | 0.60761385 | 3.3055E-12 | 3.8463E-11 |
| ENSMUSG00000032411 | Tfdp2 | 2124.08818 | 3239.61724 | 1115.52907 | 0.60689809 | 3.1831E-41 | 3.324E-39 |
| ENSMUSG00000028830 | AU040320 | 2783.16891 | 4266.84626 | 1483.67735 | 0.60685165 | 3.5936E-12 | 4.1556E-11 |
| ENSMUSG00000020528 | Prpsap2 | 1868.8402 | 2864.40185 | 995.561647 | 0.6066335 | 1.4395E-11 | 1.5622E-10 |
| ENSMUSG00000032489 | Kif9 | 52.4144857 | 82.8703609 | 30.4558751 | 0.60546863 | 0.00330398 | 0.00993706 |
| ENSMUSG00000038301 | Snx10 | 4350.607 | 6640.71447 | 2290.10747 | 0.60523956 | 1.4323E-19 | 3.4828E-18 |
| ENSMUSG00000032186 | Tmod2 | 90.0583175 | 142.675084 | 52.6167669 | 0.60495837 | 0.00275305 | 0.00842292 |
| ENSMUSG00000021830 | Txndc16 | 3117.39365 | 4770.14379 | 1652.75014 | 0.60491298 | 2.6293E-12 | 3.1033E-11 |
| ENSMUSG00000025359 | Pmel | 67.0971399 | 104.565066 | 37.4679259 | 0.60427405 | 0.00078941 | 0.00272387 |
| ENSMUSG00000022091 | Sorbs3 | 952.73276 | 1454.75981 | 502.027052 | 0.60410449 | 8.2376E-15 | 1.2751E-13 |
| ENSMUSG00000042684 | Npl | 68.3295016 | 108.715011 | 40.385509 | 0.60283638 | 0.00465034 | 0.01349701 |
| ENSMUSG00000097207 | 6030443J06R | 78.8034765 | 124.098847 | 45.2953705 | 0.6028215 | 0.00090171 | 0.00307424 |
| ENSMUSG00000029313 | Aff1 | 7906.04601 | 12096.3965 | 4190.3505 | 0.6024649 | 3.4712E-10 | 3.1865E-09 |
| ENSMUSG00000022228 | Zscan26 | 2410.91074 | 3680.81408 | 1269.90334 | 0.60210797 | 4.1131E-13 | 5.3016E-12 |
| ENSMUSG00000081137 | - | 301.702492 | 463.514768 | 161.812276 | 0.60182428 | 2.4658E-06 | 1.3159E-05 |
| ENSMUSG00000029279 | Brdt | 335.946138 | 516.452807 | 180.506668 | 0.60169547 | 1.9658E-06 | 1.0647E-05 |
| ENSMUSG00000034959 | Rubcnl | 374.497805 | 573.679793 | 199.181988 | 0.60153652 | 7.5914E-09 | 5.8644E-08 |
| ENSMUSG00000008855 | Hdac5 | 4599.18755 | 6989.17276 | 2389.98522 | 0.60152002 | 1.4532E-40 | 1.4727E-38 |
| ENSMUSG00000022440 | C1qtnf6 | 130.747984 | 203.561723 | 72.8137396 | 0.6011237 | 0.00131244 | 0.00432711 |

| | | | | | | | |
|---|---|---|---|---|---|---|---|
| ENSMUSG00000051457 | Spn | 152.697749 | 243.922234 | 91.2244843 | 0.60096175 | 0.00874252 | 0.02356232 |
| ENSMUSG00000022292 | Rrm2b | 2183.8619 | 3322.23434 | 1138.37245 | 0.60076417 | 3.9687E-24 | 1.3699E-22 |
| ENSMUSG00000028347 | Tmeff1 | 2014.39717 | 3081.30283 | 1066.90566 | 0.6007634 | 5.5993E-09 | 4.3898E-08 |
| ENSMUSG00000056749 | Nfil3 | 3990.99549 | 6092.78899 | 2101.7935 | 0.59998974 | 1.9133E-10 | 1.8206E-09 |
| ENSMUSG00000047617 | Paxx | 428.08845 | 653.396847 | 225.308397 | 0.59971987 | 5.5721E-09 | 4.3702E-08 |
| ENSMUSG00000087385 | Frg2f1 | 90.727094 | 142.283175 | 51.5560809 | 0.5992912 | 0.00124041 | 0.00411511 |
| ENSMUSG00000026259 | Ngef | 117.730072 | 182.437163 | 64.707091 | 0.59918047 | 0.00012506 | 0.00050555 |
| ENSMUSG00000038593 | Tctn1 | 272.889615 | 416.394555 | 143.50494 | 0.59899943 | 3.5838E-09 | 2.8793E-08 |
| ENSMUSG00000020882 | Cacnb1 | 316.712149 | 486.737055 | 170.024906 | 0.59859675 | 5.1548E-06 | 2.6236E-05 |
| ENSMUSG00000097162 | 2310010J17R | 76.1423025 | 120.217903 | 44.0756007 | 0.59854716 | 0.00671472 | 0.01868965 |
| ENSMUSG00000038564 | Ift172 | 2241.97087 | 3412.08201 | 1170.11114 | 0.59825797 | 1.9124E-14 | 2.8508E-13 |
| ENSMUSG00000005897 | Nr2c1 | 1094.27253 | 1661.86728 | 567.594741 | 0.5981807 | 1.2937E-21 | 3.7274E-20 |
| ENSMUSG00000003746 | Man1a | 3105.84232 | 4918.21987 | 1812.37755 | 0.5978479 | 0.00564864 | 0.01604138 |
| ENSMUSG00000097418 | Mir155hg | 78.5921673 | 124.002333 | 45.4101654 | 0.59781969 | 0.00410417 | 0.01209268 |
| ENSMUSG00000026628 | Atf3 | 3947.30512 | 6010.76853 | 2063.4634 | 0.59697026 | 6.9084E-11 | 6.9057E-10 |
| ENSMUSG00000019467 | Arhgef25 | 2061.88604 | 3121.91578 | 1060.02973 | 0.59575291 | 2.6366E-24 | 9.2559E-23 |
| ENSMUSG00000086158 | Pierce2 | 198.682889 | 306.426887 | 107.743997 | 0.59571437 | 0.00011494 | 0.00046802 |
| ENSMUSG00000086560 | - | 74.3665844 | 118.090595 | 43.7240101 | 0.59502404 | 0.00549713 | 0.01565216 |
| ENSMUSG00000022369 | Mtbp | 4029.52164 | 6106.12632 | 2076.60467 | 0.59499856 | 1.65E-22 | 5.1119E-21 |
| ENSMUSG00000054021 | Sirt5 | 514.446575 | 783.050637 | 268.604062 | 0.59449314 | 1.9936E-08 | 1.4502E-07 |
| ENSMUSG00000032387 | Rbpms2 | 388.572996 | 600.583832 | 212.010836 | 0.59429712 | 0.00032764 | 0.00122195 |
| ENSMUSG00000095990 | Zfp97 | 140.981838 | 215.117657 | 74.1358195 | 0.59421299 | 8.2243E-07 | 4.7392E-06 |
| ENSMUSG00000041594 | Tmtc4 | 3553.55596 | 5650.92541 | 2097.36945 | 0.59393191 | 0.0095088 | 0.02535186 |
| ENSMUSG00000031015 | Swap70 | 5192.68662 | 7964.07493 | 2771.38831 | 0.59372647 | 1.8101E-05 | 8.4479E-05 |
| ENSMUSG00000021280 | Exoc3l4 | 81.6425317 | 127.255366 | 45.6128345 | 0.59330833 | 0.00165288 | 0.00531713 |
| ENSMUSG00000067629 | Syngap1 | 523.430542 | 807.303141 | 283.8726 | 0.59298838 | 0.00014605 | 0.00058226 |
| ENSMUSG00000004661 | Arid3b | 118.696644 | 181.234927 | 62.5382838 | 0.5916239 | 6.7019E-06 | 3.3472E-05 |
| ENSMUSG00000037509 | Arhgef4 | 188.190129 | 288.56665 | 100.376521 | 0.59154109 | 8.5946E-06 | 4.2228E-05 |
| ENSMUSG00000039759 | Thap3 | 613.810971 | 929.661022 | 315.850051 | 0.59109408 | 1.846E-10 | 1.7582E-09 |
| ENSMUSG00000085218 | BB218582 | 94.946862 | 149.293867 | 54.3470051 | 0.59106649 | 0.00820348 | 0.02227508 |
| ENSMUSG00000038042 | Ptpdc1 | 1038.54531 | 1570.1878 | 531.642494 | 0.59102931 | 4.7877E-16 | 8.4544E-15 |
| ENSMUSG00000032109 | Nlrx1 | 442.147497 | 670.170454 | 228.022957 | 0.59048739 | 4.7592E-10 | 4.2772E-09 |
| ENSMUSG00000032637 | Atxn2l | 7932.29906 | 12017.5642 | 4085.26519 | 0.58969386 | 6.4568E-11 | 6.47E-10 |
| ENSMUSG00000050379 | Septin6 | 522.479795 | 788.201037 | 265.721242 | 0.58912653 | 3.3096E-15 | 5.3737E-14 |
| ENSMUSG00000005949 | Ctns | 1220.50723 | 1846.62508 | 626.117849 | 0.58881666 | 1.3893E-11 | 1.5105E-10 |
| ENSMUSG00000026791 | Slc2a8 | 716.811302 | 1086.97306 | 370.161759 | 0.58806798 | 1.0077E-08 | 7.6551E-08 |
| ENSMUSG00000020135 | Apc2 | 42.3305062 | 66.701607 | 24.3711009 | 0.58784079 | 0.00717533 | 0.01981088 |
| ENSMUSG00000034684 | Sema3f | 1000.24351 | 1509.95046 | 509.706948 | 0.58767745 | 6.8454E-15 | 1.0725E-13 |
| ENSMUSG00000039461 | Tcta | 954.738107 | 1442.10084 | 487.362733 | 0.58724776 | 1.0681E-12 | 1.3127E-11 |
| ENSMUSG00000046314 | Stxbp6 | 424.901705 | 644.427323 | 219.525618 | 0.58723072 | 1.8993E-08 | 1.386E-07 |
| ENSMUSG00000078302 | Foxd1 | 181.153872 | 276.357245 | 95.2033722 | 0.58720079 | 1.0919E-05 | 5.274E-05 |
| ENSMUSG00000043501 | Lgals2 | 29.5463483 | 47.3111154 | 17.764767 | 0.58637729 | 0.01485911 | 0.03754841 |
| ENSMUSG00000025815 | Dhtkd1 | 325.205721 | 495.338866 | 170.133145 | 0.58624372 | 2.2702E-05 | 0.00010426 |
| ENSMUSG00000031577 | Tti2 | 1259.35004 | 1897.54139 | 638.191354 | 0.58616912 | 4.2673E-19 | 1.0033E-17 |
| ENSMUSG00000025855 | Prkar1b | 723.431408 | 1091.77444 | 368.343031 | 0.5859434 | 1.0715E-12 | 1.3152E-11 |
| ENSMUSG00000035713 | Usp35 | 531.948051 | 803.298565 | 271.350514 | 0.58584678 | 7.9392E-12 | 8.9151E-11 |
| ENSMUSG00000066894 | Vsig10 | 770.283942 | 1169.27975 | 398.995811 | 0.58539284 | 1.1708E-06 | 6.5682E-06 |
| ENSMUSG00000085741 | 5430405H02 | 62.8035683 | 97.0706846 | 34.2671163 | 0.58537966 | 0.00221662 | 0.00694491 |
| ENSMUSG00000039145 | Camk1d | 1385.87866 | 2094.78946 | 708.910797 | 0.58527094 | 1.7521E-10 | 1.6749E-09 |
| ENSMUSG00000038594 | Cep85l | 470.544727 | 717.659156 | 247.114429 | 0.58501778 | 1.845E-05 | 8.6007E-05 |
| ENSMUSG00000034437 | - | 117.964752 | 75.9466927 | -42.01806 | -0.585051 | 0.00461144 | 0.01339735 |
| ENSMUSG00000023272 | Creld2 | 4461.48305 | 2945.78636 | -1515.6967 | -0.5856823 | 3.5648E-08 | 2.5046E-07 |
| ENSMUSG00000001175 | Calm1 | 57264.4312 | 38005.2054 | -19259.226 | -0.585969 | 3.4446E-17 | 6.781E-16 |
| ENSMUSG00000028124 | Gclm | 10365.6364 | 6857.1483 | -3508.4881 | -0.5862169 | 2.6305E-10 | 2.4609E-09 |
| ENSMUSG00000037946 | Fgd3 | 5124.16563 | 3377.72262 | -1746.443 | -0.5864675 | 2.1788E-07 | 1.3665E-06 |
| ENSMUSG00000031924 | Cyb5b | 33923.6973 | 22540.0221 | -11383.675 | -0.5865347 | 4.0573E-27 | 1.7425E-25 |
| ENSMUSG00000031971 | Ccsap | 988.70409 | 649.184149 | -339.51994 | -0.5866623 | 3.7758E-06 | 1.963E-05 |
| ENSMUSG00000021062 | Rab15 | 3957.87152 | 2625.34613 | -1332.5254 | -0.5870142 | 1.4806E-17 | 2.998E-16 |
| ENSMUSG00000071054 | Safb | 7487.58653 | 4955.32636 | -2532.2602 | -0.5870151 | 1.1931E-11 | 1.3084E-10 |
| ENSMUSG00000008036 | Ap2s1 | 6180.15913 | 4083.92946 | -2096.2297 | -0.5871662 | 5.7388E-10 | 5.0953E-09 |
| ENSMUSG00000037221 | Mospd3 | 1510.49839 | 1001.23076 | -509.26763 | -0.5871761 | 3.5533E-16 | 6.3619E-15 |
| ENSMUSG00000052337 | Immt | 29325.1447 | 19468.9304 | -9856.2143 | -0.5872611 | 1.2878E-24 | 4.6317E-23 |
| ENSMUSG00000071716 | Apol7e | 64.0924555 | 40.8925634 | -23.199892 | -0.5882402 | 0.00608575 | 0.0171431 |
| ENSMUSG00000007655 | Cav1 | 23962.1218 | 15710.1792 | -8251.9426 | -0.5883389 | 8.1924E-06 | 4.0368E-05 |
| ENSMUSG00000021276 | Cinp | 3399.43365 | 2240.46599 | -1158.9677 | -0.5889499 | 9.2534E-09 | 7.0635E-08 |
| ENSMUSG00000027108 | Ola1 | 14280.0851 | 9469.07784 | -4811.0073 | -0.5889899 | 5.131E-25 | 1.9088E-23 |
| ENSMUSG00000046623 | Gjb4 | 1583.22021 | 1045.75291 | -537.4673 | -0.5890025 | 2.6061E-11 | 2.7466E-10 |
| ENSMUSG00000073131 | Vma21 | 20219.0354 | 12833.6783 | -7385.3571 | -0.5891302 | 0.00753344 | 0.02065822 |
| ENSMUSG00000036599 | Chst12 | 1938.76015 | 1281.72182 | -657.03833 | -0.5893428 | 2.2388E-13 | 2.9695E-12 |
| ENSMUSG00000016921 | Srsf6 | 20077.2543 | 13289.3096 | -6787.9447 | -0.5893879 | 3.0081E-16 | 5.4188E-15 |
| ENSMUSG00000057113 | Npm1 | 127791.836 | 84130.9924 | -43660.844 | -0.589535 | 4.6226E-08 | 3.1847E-07 |
| ENSMUSG00000031381 | Piga | 2963.97775 | 1964.31454 | -999.66322 | -0.5906287 | 1.5962E-26 | 6.6205E-25 |
| ENSMUSG00000022178 | Ajuba | 8651.61798 | 5721.94486 | -2929.6731 | -0.5916132 | 5.2019E-19 | 1.2133E-17 |
| ENSMUSG00000090674 | - | 137.199704 | 88.3962998 | -48.803404 | -0.591621 | 0.00091657 | 0.00312129 |
| ENSMUSG00000067189 | - | 87.1130228 | 55.789823 | -31.3232 | -0.5916471 | 0.00379952 | 0.0112774 |

| | | | | | | | |
|---|---|---|---|---|---|---|---|
| ENSMUSG00000022205 | Sub1 | 18076.2488 | 11901.1033 | -6175.1454 | -0.5918161 | 1.4791E-09 | 1.2476E-08 |
| ENSMUSG00000032330 | Cox7a2 | 7562.36661 | 4992.259 | -2570.1076 | -0.5919396 | 2.3473E-14 | 3.4715E-13 |
| ENSMUSG00000045160 | Bola3 | 2829.85464 | 1857.26975 | -972.58488 | -0.5919562 | 1.9618E-07 | 1.2391E-06 |
| ENSMUSG00000002820 | Atg4d | 1100.50727 | 726.308804 | -374.19846 | -0.5920248 | 1.6293E-16 | 3.0062E-15 |
| ENSMUSG00000025156 | Gps1 | 10291.7039 | 6796.3209 | -3495.383 | -0.5921443 | 2.754E-15 | 4.5216E-14 |
| ENSMUSG00000036309 | Skp1 | 25128.9092 | 16590.0732 | -8538.836 | -0.5923968 | 5.2799E-15 | 8.3682E-14 |
| ENSMUSG00000027933 | Ints3 | 8733.65406 | 5770.23544 | -2963.4186 | -0.5924208 | 3.0338E-17 | 5.9954E-16 |
| ENSMUSG00000079311 | - | 96.3313656 | 60.9165259 | -35.41484 | -0.5929838 | 0.00490098 | 0.01415059 |
| ENSMUSG00000027752 | Exosc8 | 5250.14927 | 3463.3887 | -1786.7606 | -0.5931396 | 1.4791E-14 | 2.234E-13 |
| ENSMUSG00000085436 | Zfp335os | 140.31965 | 87.735481 | -52.584169 | -0.5931806 | 0.01409689 | 0.03583359 |
| ENSMUSG00000020549 | Elac2 | 4273.36882 | 2823.8682 | -1449.5006 | -0.5934533 | 4.7556E-21 | 1.3185E-19 |
| ENSMUSG00000020782 | Llgl2 | 177.634678 | 113.332364 | -64.302314 | -0.5935552 | 0.00252805 | 0.00781591 |
| ENSMUSG00000002129 | Sf3a1 | 6069.94673 | 4008.70547 | -2061.2413 | -0.593737 | 6.6807E-19 | 1.5459E-17 |
| ENSMUSG00000030603 | Psmc4 | 10955.2681 | 7213.08962 | -3742.1785 | -0.5937489 | 2.0681E-11 | 2.2067E-10 |
| ENSMUSG00000076928 | - | 94.8998763 | 60.725498 | -34.174378 | -0.5938724 | 0.00184976 | 0.00589972 |
| ENSMUSG00000001525 | Tubb5 | 126829.66 | 83624.2079 | -43205.452 | -0.5942052 | 6.9443E-15 | 1.0864E-13 |
| ENSMUSG00000037466 | Tedc1 | 2511.30861 | 1652.13895 | -859.16967 | -0.5943855 | 9.6137E-11 | 9.4493E-10 |
| ENSMUSG00000004929 | Thop1 | 7209.34837 | 4760.47298 | -2448.8754 | -0.5944874 | 1.9037E-23 | 6.2963E-22 |
| ENSMUSG00000079111 | Kdelr2 | 13941.8567 | 9199.8121 | -4742.0446 | -0.594601 | 2.1729E-19 | 5.2159E-18 |
| ENSMUSG00000063018 | - | 646.730895 | 422.152974 | -224.57792 | -0.5948615 | 3.4038E-06 | 1.7822E-05 |
| ENSMUSG00000031896 | Ctrl | 40.4982976 | 25.2866214 | -15.211676 | -0.5951878 | 0.0116927 | 0.03044452 |
| ENSMUSG00000090841 | Myl6 | 50919.5707 | 33580.331 | -17339.24 | -0.5953578 | 1.0161E-18 | 2.3278E-17 |
| ENSMUSG00000026028 | Trak2 | 6205.42317 | 4084.37834 | -2121.0448 | -0.5955904 | 7.1029E-13 | 8.8773E-12 |
| ENSMUSG00000034765 | Dusp5 | 923.983442 | 587.30708 | -336.67636 | -0.5956947 | 0.00406041 | 0.01197574 |
| ENSMUSG00000054408 | Spcs3 | 20014.1534 | 13173.3585 | -6840.795 | -0.5957736 | 1.9499E-13 | 2.5946E-12 |
| ENSMUSG00000041360 | Pum3 | 11661.56 | 7681.10555 | -3980.4545 | -0.5961857 | 4.6934E-16 | 8.2949E-15 |
| ENSMUSG00000026479 | Lamc2 | 10893.8028 | 7164.19292 | -3729.6099 | -0.5965783 | 7.5994E-13 | 9.4748E-12 |
| ENSMUSG00000053317 | Sec61b | 10107.8877 | 6590.83009 | -3517.0576 | -0.5966042 | 4.3741E-06 | 2.2502E-05 |
| ENSMUSG00000030423 | Pop4 | 5628.48094 | 3701.1219 | -1927.359 | -0.5972 | 6.3822E-14 | 8.9682E-13 |
| ENSMUSG00000034165 | Ccnd3 | 9633.81823 | 6341.3302 | -3292.488 | -0.5980866 | 3.156E-19 | 7.4712E-18 |
| ENSMUSG00000005505 | Kbtbd4 | 901.744059 | 592.144392 | -309.59967 | -0.5982046 | 5.7E-11 | 5.7396E-10 |
| ENSMUSG00000059323 | Tonsl | 3068.40067 | 2014.88636 | -1053.5143 | -0.5984221 | 1.7297E-12 | 2.0797E-11 |
| ENSMUSG00000030204 | Ddx47 | 10741.7117 | 7071.37807 | -3670.3336 | -0.5985051 | 2.9327E-21 | 8.2532E-20 |
| ENSMUSG00000032366 | Tpm1 | 76897.0971 | 50265.803 | -26631.294 | -0.5986981 | 7.1303E-08 | 4.7733E-07 |
| ENSMUSG00000098154 | - | 50.7894309 | 31.3886987 | -19.400732 | -0.5987442 | 0.017268 | 0.04275845 |
| ENSMUSG00000043889 | - | 181.948944 | 116.610923 | -65.338021 | -0.598895 | 0.00095219 | 0.00323296 |
| ENSMUSG00000031787 | Katnb1 | 3309.18841 | 2175.51901 | -1133.6694 | -0.5990652 | 7.5726E-17 | 1.4546E-15 |
| ENSMUSG00000024312 | Wdr46 | 5013.5737 | 3297.27148 | -1716.3022 | -0.5997157 | 5.5536E-20 | 1.3967E-18 |
| ENSMUSG00000020185 | E2f7 | 2491.30476 | 1607.5022 | -883.80256 | -0.5999415 | 0.00018552 | 0.00072697 |
| ENSMUSG00000079480 | Pin4 | 1373.16356 | 895.098248 | -478.06531 | -0.6007366 | 1.714E-07 | 1.0906E-06 |
| ENSMUSG00000020863 | Luc7l3 | 13233.2135 | 8629.61084 | -4603.6026 | -0.6010404 | 1.6445E-07 | 1.0507E-06 |
| ENSMUSG00000026854 | Usp20 | 4538.28913 | 2983.03804 | -1555.2511 | -0.6011432 | 2.3675E-23 | 7.8051E-22 |
| ENSMUSG00000039234 | Sec24d | 5742.63631 | 3763.6974 | -1978.9389 | -0.6014346 | 5.2025E-13 | 6.6066E-12 |
| ENSMUSG00000063316 | Rpl27 | 15796.9923 | 10329.7415 | -5467.2508 | -0.6015924 | 6.507E-10 | 5.7304E-09 |
| ENSMUSG00000063754 | - | 76.577978 | 48.1331896 | -28.444788 | -0.6018544 | 0.00650851 | 0.01818459 |
| ENSMUSG00000022297 | Fzd6 | 1771.4521 | 1155.55806 | -615.89404 | -0.6019592 | 5.3757E-08 | 3.657E-07 |
| ENSMUSG00000098749 | - | 88.8377421 | 55.2639333 | -33.573809 | -0.6024165 | 0.00883529 | 0.02376252 |
| ENSMUSG00000000399 | Ndufa9 | 12057.5489 | 7900.4912 | -4157.0577 | -0.6024308 | 4.3943E-14 | 6.3217E-13 |
| ENSMUSG00000017781 | Pitpna | 20644.1406 | 13558.868 | -7085.2726 | -0.6026255 | 5.8578E-26 | 2.3309E-24 |
| ENSMUSG00000029992 | Gfpt1 | 15221.23 | 9926.69559 | -5294.5344 | -0.602924 | 2.089E-08 | 1.5159E-07 |
| ENSMUSG00000041012 | Cmtm8 | 57.1620076 | 35.7024411 | -21.459567 | -0.6031277 | 0.01105275 | 0.02896892 |
| ENSMUSG00000030058 | Copg1 | 29814.3083 | 19389.9479 | -10424.36 | -0.6032278 | 5.4967E-07 | 3.2484E-06 |
| ENSMUSG00000093565 | Rab26os | 278.650285 | 173.204329 | -105.44596 | -0.6036877 | 0.01126902 | 0.02947422 |
| ENSMUSG00000081684 | - | 59.3368515 | 37.3432274 | -21.993624 | -0.6037378 | 0.00687401 | 0.01907367 |
| ENSMUSG00000021340 | Gpld1 | 615.060475 | 397.021529 | -218.03895 | -0.6042549 | 4.6982E-05 | 0.00020477 |
| ENSMUSG00000020899 | Pfas | 3685.75793 | 2411.84005 | -1273.9179 | -0.60436 | 1.2124E-13 | 1.6551E-12 |
| ENSMUSG00000022972 | Cfap298 | 1965.23751 | 1284.72497 | -680.51255 | -0.6043862 | 1.0999E-12 | 1.3485E-11 |
| ENSMUSG00000007050 | Lsm2 | 4057.84412 | 2652.46098 | -1405.3831 | -0.6044521 | 2.7218E-12 | 3.2069E-11 |
| ENSMUSG00000081485 | - | 240.84058 | 154.336882 | -86.503698 | -0.6048283 | 0.0003365 | 0.00125136 |
| ENSMUSG00000081721 | - | 107.963777 | 68.4618098 | -39.501967 | -0.6050673 | 0.0008467 | 0.00290498 |
| ENSMUSG00000024190 | Dusp1 | 5780.64163 | 3586.65775 | -2193.9839 | -0.6051184 | 0.01108001 | 0.02902929 |
| ENSMUSG00000029246 | Ppat | 8805.72542 | 5774.64501 | -3031.0804 | -0.6052843 | 4.3637E-29 | 2.1374E-27 |
| ENSMUSG00000033031 | Cip2a | 14156.913 | 9278.84449 | -4878.0685 | -0.6054285 | 5.6152E-25 | 2.0814E-23 |
| ENSMUSG00000096006 | - | 94.6900808 | 59.3968496 | -35.293231 | -0.6056874 | 0.00422712 | 0.01240228 |
| ENSMUSG00000029840 | Mtpn | 73159.7648 | 47808.3004 | -25351.464 | -0.6061396 | 5.7029E-14 | 8.0911E-13 |
| ENSMUSG00000022034 | Esco2 | 2959.83534 | 1933.75128 | -1026.0841 | -0.6064348 | 1.0936E-14 | 1.6802E-13 |
| ENSMUSG00000061414 | Cracr2a | 74.1422183 | 46.0113595 | -28.130859 | -0.6067815 | 0.01082984 | 0.02844631 |
| ENSMUSG00000027342 | Pcna | 27589.2879 | 18031.6439 | -9557.644 | -0.6069192 | 5.931E-16 | 1.038E-14 |
| ENSMUSG00000031952 | Chst5 | 412.487136 | 265.160079 | -147.32706 | -0.6076827 | 9.1156E-05 | 0.00037808 |
| ENSMUSG00000062110 | Scfd2 | 1447.84367 | 945.467274 | -502.3764 | -0.608136 | 5.1784E-17 | 1.005E-15 |
| ENSMUSG00000026087 | Mrpl30 | 5526.98533 | 3587.06124 | -1939.9241 | -0.6090024 | 2.8904E-08 | 2.0532E-07 |
| ENSMUSG00000022711 | Pmm2 | 4312.76316 | 2806.0942 | -1506.669 | -0.6090591 | 2.0405E-10 | 1.9354E-09 |
| ENSMUSG00000083179 | - | 91.6361129 | 57.1015026 | -34.53461 | -0.6095435 | 0.00412266 | 0.01213844 |
| ENSMUSG00000056665 | Them6 | 2067.21395 | 1340.98248 | -726.23147 | -0.609546 | 4.4918E-08 | 3.104E-07 |
| ENSMUSG00000046694 | Tent5b | 427.011094 | 275.618473 | -151.39262 | -0.6096712 | 1.1054E-06 | 6.22E-06 |

| | | | | | | | |
|---|---|---|---|---|---|---|---|
| ENSMUSG00000021737 | Psmd6 | 14102.5284 | 9180.49364 | -4922.0347 | -0.6098952 | 6.8379E-12 | 7.7122E-11 |
| ENSMUSG00000063275 | Hacd1 | 2068.06625 | 1330.68405 | -737.3822 | -0.610275 | 2.4592E-05 | 0.0001123 |
| ENSMUSG00000006494 | Pdk1 | 7365.20874 | 4800.24375 | -2564.965 | -0.6103945 | 3.1757E-15 | 5.1726E-14 |
| ENSMUSG00000082465 | - | 58.0592194 | 35.3349332 | -22.724286 | -0.6105434 | 0.01803899 | 0.04447484 |
| ENSMUSG00000025791 | Pgm1 | 8733.21709 | 5694.02418 | -3039.1929 | -0.6112247 | 1.4895E-18 | 3.3416E-17 |
| ENSMUSG00000015932 | Dstn | 37309.0966 | 24195.3187 | -13113.778 | -0.6115267 | 5.5162E-09 | 4.3346E-08 |
| ENSMUSG00000024833 | Pola2 | 5022.79527 | 3278.83482 | -1743.9605 | -0.6119905 | 3.5884E-33 | 2.2714E-31 |
| ENSMUSG00000029250 | Polr2b | 14514.6972 | 9476.52069 | -5038.1765 | -0.6124451 | 2.0759E-37 | 1.722E-35 |
| ENSMUSG00000031668 | Eif2ak3 | 3006.5037 | 1955.60027 | -1050.9034 | -0.6127094 | 5.8244E-14 | 8.243E-13 |
| ENSMUSG00000006998 | Psmd2 | 54725.6546 | 35718.9777 | -19006.677 | -0.6129977 | 6.8116E-40 | 6.8023E-38 |
| ENSMUSG00000019297 | Nop9 | 5967.68917 | 3884.20212 | -2083.487 | -0.6135265 | 5.1703E-18 | 1.0977E-16 |
| ENSMUSG00000027671 | Actl6a | 6013.24672 | 3910.85731 | -2102.3894 | -0.6135424 | 1.6312E-15 | 2.7327E-14 |
| ENSMUSG00000035890 | Rnf126 | 6063.46891 | 3946.35039 | -2117.1185 | -0.6136818 | 1.989E-18 | 4.3999E-17 |
| ENSMUSG00000029642 | Polr1d | 8113.68728 | 5269.86424 | -2843.823 | -0.6140548 | 2.6913E-13 | 3.5399E-12 |
| ENSMUSG00000024378 | Stard4 | 5036.08646 | 3235.51161 | -1800.5749 | -0.6142181 | 9.4333E-06 | 4.602E-05 |
| ENSMUSG00000031523 | Dlc1 | 8314.09347 | 5373.89312 | -2940.2003 | -0.6146104 | 3.4487E-08 | 2.4272E-07 |
| ENSMUSG00000021359 | Tfap2a | 2100.0075 | 1361.88084 | -738.12666 | -0.6146953 | 2.9378E-11 | 3.0788E-10 |
| ENSMUSG00000014867 | Surf4 | 39632.5309 | 25846.822 | -13785.709 | -0.6148194 | 2.5691E-53 | 4.8937E-51 |
| ENSMUSG00000048164 | - | 105.834082 | 66.5978539 | -39.236228 | -0.6152181 | 0.00192112 | 0.00610461 |
| ENSMUSG00000001424 | Snd1 | 22280.7645 | 14457.6588 | -7823.1057 | -0.615347 | 4.6999E-13 | 6.0017E-12 |
| ENSMUSG00000047539 | Fbxo28 | 3525.62365 | 2294.25145 | -1231.3722 | -0.6160145 | 1.8563E-25 | 7.1379E-24 |
| ENSMUSG00000035337 | Uchl4 | 74.6237961 | 46.1933943 | -28.430402 | -0.6170903 | 0.0087453 | 0.02356671 |
| ENSMUSG00000020988 | L2hgdh | 2137.76563 | 1389.67563 | -748.09 | -0.6175085 | 3.5195E-24 | 1.2271E-22 |
| ENSMUSG00000025224 | Gbf1 | 8299.64933 | 5371.06437 | -2928.585 | -0.6176565 | 2.625E-11 | 2.765E-10 |
| ENSMUSG00000060152 | Pop5 | 2668.50824 | 1723.97548 | -944.53276 | -0.6179873 | 6.9587E-10 | 6.0968E-09 |
| ENSMUSG00000030718 | Ppme1 | 10723.5829 | 6968.79199 | -3754.7909 | -0.6181446 | 1.1763E-28 | 5.6672E-27 |
| ENSMUSG00000063802 | Hspbp1 | 2530.84074 | 1638.64922 | -892.19152 | -0.618709 | 7.1042E-14 | 9.915E-13 |
| ENSMUSG00000028639 | Ybx1 | 60783.5631 | 39409.9957 | -21373.567 | -0.6194267 | 2.632E-19 | 6.3034E-18 |
| ENSMUSG00000032802 | Srxn1 | 5068.22969 | 3282.79728 | -1785.4324 | -0.6194352 | 3.0707E-16 | 5.5219E-15 |
| ENSMUSG00000070287 | Slc35g2 | 1122.96034 | 711.593726 | -411.36661 | -0.6198733 | 0.0002613 | 0.00099473 |
| ENSMUSG00000069744 | Psmb3 | 11843.5307 | 7653.56436 | -4189.9663 | -0.6204025 | 1.9284E-12 | 2.3025E-11 |
| ENSMUSG00000004356 | Utp20 | 7638.83875 | 4928.90474 | -2709.934 | -0.6207379 | 1.9957E-10 | 1.8954E-09 |
| ENSMUSG00000023944 | Hsp90ab1 | 281345.606 | 182335.469 | -99010.137 | -0.6209193 | 1.7574E-22 | 5.4283E-21 |
| ENSMUSG00000071653 | 1810009A15 | 4455.19044 | 2877.2726 | -1577.9178 | -0.6210507 | 2.9179E-12 | 3.4281E-11 |
| ENSMUSG00000036751 | Cox6b1 | 9090.25684 | 5865.97595 | -3224.2809 | -0.6211779 | 3.1293E-11 | 3.2645E-10 |
| ENSMUSG00000004054 | Map3k11 | 4671.85414 | 3028.39096 | -1643.4632 | -0.6224695 | 4.1211E-42 | 4.4857E-40 |
| ENSMUSG00000034875 | Nudt19 | 1953.33663 | 1255.77184 | -697.56478 | -0.6227669 | 7.298E-09 | 5.6569E-08 |
| ENSMUSG00000042111 | Ccdc115 | 1990.81384 | 1285.36534 | -705.4485 | -0.6227675 | 8.3743E-14 | 1.1625E-12 |
| ENSMUSG00000065265 | - | 84.4417425 | 51.5354495 | -32.906293 | -0.6228168 | 0.01074145 | 0.02826428 |
| ENSMUSG00000052738 | Suclg1 | 11152.3607 | 7208.3416 | -3944.0191 | -0.6231095 | 1.4146E-17 | 2.8727E-16 |
| ENSMUSG00000038028 | Tigar | 2998.42334 | 1942.63133 | -1055.792 | -0.6232648 | 1.0236E-36 | 8.1935E-35 |
| ENSMUSG00000067608 | - | 311.936978 | 198.456226 | -113.48075 | -0.6235593 | 2.0917E-05 | 9.6802E-05 |
| ENSMUSG00000004044 | Cavin1 | 7394.21138 | 4784.29969 | -2609.9117 | -0.6240605 | 7.229E-30 | 3.7649E-28 |
| ENSMUSG00000078348 | Sf3b5 | 3977.40567 | 2558.29889 | -1419.1068 | -0.6243567 | 3.0871E-10 | 2.8568E-09 |
| ENSMUSG00000084111 | - | 141.52653 | 88.9029599 | -52.62357 | -0.6253053 | 0.00056874 | 0.00202073 |
| ENSMUSG00000020994 | Pnn | 11537.8299 | 7417.45683 | -4120.373 | -0.6256853 | 1.5352E-10 | 1.4772E-09 |
| ENSMUSG00000070372 | Capza1 | 7731.21986 | 4981.99718 | -2749.2227 | -0.6258622 | 1.0751E-14 | 1.6529E-13 |
| ENSMUSG00000028729 | Ebna1bp2 | 8448.34473 | 5444.52625 | -3003.8185 | -0.6260441 | 5.163E-15 | 8.1954E-14 |
| ENSMUSG00000020553 | Pctp | 839.372466 | 539.067014 | -300.30545 | -0.6261918 | 2.6056E-12 | 3.0789E-11 |
| ENSMUSG00000075266 | Cenpw | 2124.73549 | 1360.39 | -764.34549 | -0.6264689 | 5.2085E-08 | 3.5527E-07 |
| ENSMUSG00000025962 | Fastkd2 | 2939.08102 | 1893.04254 | -1046.0385 | -0.6265028 | 5.3265E-14 | 7.5832E-13 |
| ENSMUSG00000029701 | Rbm28 | 13818.0628 | 8934.35995 | -4883.7029 | -0.6265471 | 2.0028E-42 | 2.2271E-40 |
| ENSMUSG00000043998 | Mgat2 | 11171.2731 | 7218.86828 | -3952.4048 | -0.6267118 | 8.4758E-35 | 6.0543E-33 |
| ENSMUSG00000074781 | Ube2n | 7450.11527 | 4811.82055 | -2638.2947 | -0.6268089 | 3.6546E-30 | 1.9427E-28 |
| ENSMUSG00000009687 | Fxyd5 | 6862.66682 | 4416.17877 | -2446.4881 | -0.6269736 | 1.1058E-13 | 1.5197E-12 |
| ENSMUSG00000047757 | Fancb | 2054.362 | 1326.76685 | -727.59515 | -0.6273397 | 1.5555E-30 | 8.5792E-29 |
| ENSMUSG00000082896 | - | 171.633944 | 108.07695 | -63.556994 | -0.6275521 | 0.00017667 | 0.0006952 |
| ENSMUSG00000039231 | Suv39h1 | 5374.42865 | 3470.95453 | -1903.4741 | -0.6277902 | 3.4865E-38 | 3.0136E-36 |
| ENSMUSG00000019837 | Gtf3c6 | 3048.09917 | 1949.74817 | -1098.351 | -0.6280273 | 4.9422E-08 | 3.3891E-07 |
| ENSMUSG00000031843 | Mphosph6 | 4420.85186 | 2847.10178 | -1573.7501 | -0.6281819 | 1.8494E-18 | 4.1041E-17 |
| ENSMUSG00000004268 | Emg1 | 8989.24149 | 5771.01683 | -3218.2247 | -0.6288553 | 1.0372E-11 | 1.1447E-10 |
| ENSMUSG00000026177 | Slc11a1 | 140.49492 | 88.4907837 | -52.004137 | -0.6289547 | 0.00021572 | 0.00083651 |
| ENSMUSG00000020496 | Rnf187 | 13775.2518 | 8871.58723 | -4903.6646 | -0.6291901 | 1.5796E-20 | 4.1395E-19 |
| ENSMUSG00000070871 | Ccnyl1 | 1890.60446 | 1205.07461 | -685.52985 | -0.6292445 | 6.5942E-07 | 3.8495E-06 |
| ENSMUSG00000035021 | Baz1a | 8027.90644 | 5140.65392 | -2887.2525 | -0.6292501 | 2.8009E-09 | 2.282E-08 |
| ENSMUSG00000092072 | - | 135.090651 | 82.9621852 | -52.128466 | -0.6292927 | 0.00366119 | 0.01089827 |
| ENSMUSG00000096965 | 3300005D01 | 893.656349 | 573.58127 | -320.07508 | -0.6293772 | 2.0013E-13 | 2.6613E-12 |
| ENSMUSG00000049230 | - | 1168.49184 | 743.207797 | -425.28404 | -0.6297263 | 1.1816E-06 | 6.6234E-06 |
| ENSMUSG00000071662 | Polr2g | 10185.7171 | 6542.37702 | -3643.3401 | -0.6298942 | 6.3396E-14 | 8.9144E-13 |
| ENSMUSG00000017221 | Psmd3 | 18875.4171 | 12168.1982 | -6707.2189 | -0.630294 | 1.3821E-35 | 1.0265E-33 |
| ENSMUSG00000026712 | Mrc1 | 963.831046 | 612.74445 | -351.0866 | -0.6303379 | 3.8204E-06 | 1.9837E-05 |
| ENSMUSG00000024772 | Ehd1 | 24709.7865 | 15898.8367 | -8810.9498 | -0.63035 | 7.0262E-20 | 1.752E-18 |
| ENSMUSG00000045302 | Preb | 8503.81551 | 5484.03579 | -3019.7797 | -0.630549 | 6.5149E-48 | 9.2431E-46 |
| ENSMUSG00000024732 | Ccdc86 | 2785.07784 | 1788.63283 | -996.445 | -0.6309578 | 5.1168E-15 | 8.1346E-14 |
| ENSMUSG00000006057 | Atp5g1 | 6726.74513 | 4307.04509 | -2419.7 | -0.631535 | 6.4153E-11 | 6.4316E-10 |

| | | | | | | | |
|---|---|---|---|---|---|---|---|
| ENSMUSG00000041763 | Tpp2 | 13031.8726 | 8378.92526 | -4652.9474 | -0.6319576 | 9.46E-22 | 2.7526E-20 |
| ENSMUSG00000029447 | Cct6a | 31219.7387 | 20091.8061 | -11127.933 | -0.6321615 | 7.311E-31 | 4.1093E-29 |
| ENSMUSG00000020929 | Eftud2 | 16175.5133 | 10415.5301 | -5759.9832 | -0.6326527 | 1.6183E-46 | 2.1341E-44 |
| ENSMUSG00000064068 | Mtx1 | 3663.06749 | 2350.24618 | -1312.8213 | -0.6326789 | 1.1163E-16 | 2.103E-15 |
| ENSMUSG00000030189 | Ybx3 | 50729.1271 | 32474.6132 | -18254.514 | -0.6327423 | 1.1842E-11 | 1.2993E-10 |
| ENSMUSG00000054074 | Skida1 | 490.825508 | 313.428042 | -177.39747 | -0.6329073 | 1.5416E-09 | 1.2955E-08 |
| ENSMUSG00000046841 | Ckap4 | 22869.8866 | 14721.7297 | -8148.1569 | -0.632945 | 2.0098E-45 | 2.5211E-43 |
| ENSMUSG00000021763 | Cspg4b | 119.215819 | 73.9691649 | -45.246654 | -0.6339138 | 0.00185637 | 0.00591601 |
| ENSMUSG00000034880 | Mrpl34 | 2815.83495 | 1800.65411 | -1015.1808 | -0.6340334 | 8.8333E-12 | 9.8492E-11 |
| ENSMUSG00000025036 | Sfxn2 | 1222.08122 | 775.742284 | -446.33894 | -0.6340468 | 9.6084E-07 | 5.4679E-06 |
| ENSMUSG00000002718 | Cse1l | 18447.8724 | 11869.633 | -6578.2394 | -0.6342081 | 6.0298E-55 | 1.253E-52 |
| ENSMUSG00000020288 | Ahsa2 | 3261.25617 | 2089.77467 | -1171.4815 | -0.6343551 | 7.011E-16 | 1.2202E-14 |
| ENSMUSG00000005370 | Msh6 | 10702.6337 | 6873.107 | -3829.5267 | -0.6347216 | 2.1557E-27 | 9.4355E-26 |
| ENSMUSG00000095098 | Ccdc85b | 2889.49077 | 1834.40186 | -1055.0889 | -0.6350176 | 4.642E-07 | 2.7738E-06 |
| ENSMUSG00000017774 | Myo1c | 26295.5461 | 16886.6143 | -9408.9317 | -0.6350207 | 2.3602E-29 | 1.1728E-27 |
| ENSMUSG00000027940 | Tpm3 | 35579.8003 | 22834.9487 | -12744.852 | -0.6350616 | 1.5189E-24 | 5.4249E-23 |
| ENSMUSG00000048922 | Cdca2 | 4998.48788 | 3205.71973 | -1792.7682 | -0.6351143 | 7.9742E-22 | 2.3469E-20 |
| ENSMUSG00000025144 | Cenpx | 1532.41834 | 970.340502 | -562.07784 | -0.6363191 | 1.3082E-06 | 7.2813E-06 |
| ENSMUSG00000020873 | Slc35b1 | 7667.03031 | 4906.57477 | -2760.4555 | -0.6367742 | 4.1526E-17 | 8.1128E-16 |
| ENSMUSG00000024683 | Mrpl16 | 3816.73184 | 2433.97373 | -1382.7581 | -0.6368427 | 1.2724E-10 | 1.2411E-09 |
| ENSMUSG00000024871 | Doc2g | 114.572527 | 71.8056237 | -42.766903 | -0.6369978 | 0.00011798 | 0.00047927 |
| ENSMUSG00000082209 | - | 1593.88787 | 1014.6125 | -579.27537 | -0.6373939 | 1.4919E-09 | 1.2573E-08 |
| ENSMUSG00000081272 | - | 2894.52105 | 1831.8023 | -1062.7188 | -0.63746 | 1.3884E-06 | 7.7071E-06 |
| ENSMUSG00000084830 | - | 171.88115 | 107.883183 | -63.997967 | -0.6379258 | 9.8412E-05 | 0.00040572 |
| ENSMUSG00000015944 | Castor2 | 2171.36014 | 1372.21576 | -799.14437 | -0.6386249 | 2.1181E-06 | 1.143E-05 |
| ENSMUSG00000030007 | Cct7 | 50666.012 | 32433.4123 | -18232.6 | -0.6386412 | 3.785E-24 | 1.3109E-22 |
| ENSMUSG00000004500 | Zfp324 | 845.498586 | 539.119288 | -306.3793 | -0.6386441 | 2.7049E-11 | 2.8448E-10 |
| ENSMUSG00000021102 | Glrx5 | 5909.4161 | 3760.93209 | -2148.484 | -0.638837 | 2.9495E-10 | 2.7419E-09 |
| ENSMUSG00000026385 | Dbi | 5943.8571 | 3787.68642 | -2156.1707 | -0.6388923 | 1.0386E-11 | 1.145E-10 |
| ENSMUSG00000054115 | Skp2 | 4394.6343 | 2803.81803 | -1590.8163 | -0.6391172 | 1.4245E-13 | 1.933E-12 |
| ENSMUSG00000059518 | Znhit1 | 2970.18954 | 1893.14273 | -1077.0468 | -0.6391985 | 8.3788E-13 | 1.0421E-11 |
| ENSMUSG00000040550 | Otud6b | 7360.39807 | 4712.61625 | -2647.7818 | -0.639279 | 9.8629E-28 | 4.4301E-26 |
| ENSMUSG00000022905 | Kpna1 | 8499.57598 | 5448.51368 | -3051.0623 | -0.6393272 | 1.2579E-46 | 1.6804E-44 |
| ENSMUSG00000020180 | Snrpd3 | 10123.8556 | 6476.91143 | -3646.9442 | -0.6395851 | 2.3045E-26 | 9.3508E-25 |
| ENSMUSG00000035364 | - | 174.986956 | 109.770082 | -65.216874 | -0.6397104 | 5.0495E-05 | 0.00021901 |
| ENSMUSG00000017801 | Mlx | 2996.85612 | 1914.2394 | -1082.6167 | -0.6397299 | 1.6976E-20 | 4.4318E-19 |
| ENSMUSG00000027808 | Serp1 | 33957.2949 | 21751.8143 | -12205.481 | -0.6397349 | 4.8651E-41 | 5.0548E-39 |
| ENSMUSG00000060214 | - | 708.197077 | 446.527831 | -261.66925 | -0.6397799 | 4.5729E-06 | 2.3431E-05 |
| ENSMUSG00000037351 | Actr1b | 9509.28329 | 6079.30559 | -3429.9777 | -0.6400957 | 2.1078E-22 | 6.4816E-21 |
| ENSMUSG00000062203 | Gspt1 | 23632.1295 | 15137.2835 | -8494.846 | -0.6404022 | 8.8906E-50 | 1.487E-47 |
| ENSMUSG00000026639 | Lamb3 | 534.278088 | 335.846872 | -198.43122 | -0.6407945 | 2.1312E-05 | 9.8458E-05 |
| ENSMUSG00000068039 | Tcp1 | 43451.8687 | 27738.1998 | -15713.669 | -0.6410389 | 9.8824E-19 | 2.2665E-17 |
| ENSMUSG00000031824 | 6430548M08 | 1568.48581 | 994.920264 | -573.56555 | -0.6415018 | 5.9479E-09 | 4.6507E-08 |
| ENSMUSG00000097954 | - | 31.1364471 | 18.3297349 | -12.806712 | -0.6420219 | 0.01790309 | 0.04416097 |
| ENSMUSG00000041164 | Zmiz2 | 3849.77707 | 2444.00633 | -1405.7707 | -0.6422585 | 3.6768E-10 | 3.3647E-09 |
| ENSMUSG00000038352 | Arl5c | 551.593863 | 337.93377 | -213.66009 | -0.6425502 | 0.0020927 | 0.00659282 |
| ENSMUSG00000039680 | Mrps6 | 4120.23363 | 2618.85725 | -1501.3764 | -0.6427082 | 8.5391E-12 | 9.5403E-11 |
| ENSMUSG00000001305 | Rrp15 | 2748.09264 | 1742.90982 | -1005.1828 | -0.643311 | 5.9537E-10 | 5.268E-09 |
| ENSMUSG00000024646 | Cyb5a | 9738.52911 | 6190.17581 | -3548.3533 | -0.6440233 | 1.4366E-13 | 1.9481E-12 |
| ENSMUSG00000026127 | Imp4 | 3955.67951 | 2512.34494 | -1443.3346 | -0.6444683 | 2.0904E-12 | 2.4887E-11 |
| ENSMUSG00000031232 | Magt1 | 14907.2675 | 9491.62031 | -5415.6472 | -0.6444998 | 1.5358E-18 | 3.4416E-17 |
| ENSMUSG00000074886 | Grk6 | 8209.40354 | 5235.82401 | -2973.5795 | -0.6448819 | 1.2407E-31 | 7.2927E-30 |
| ENSMUSG00000058050 | - | 70.7648517 | 42.9471031 | -27.817749 | -0.6454094 | 0.00383691 | 0.01138017 |
| ENSMUSG00000053038 | - | 481.064188 | 300.866752 | -180.19744 | -0.6454117 | 2.7856E-05 | 0.00012609 |
| ENSMUSG00000060467 | - | 147.616292 | 92.6132304 | -55.003062 | -0.6454967 | 1.1044E-05 | 5.3318E-05 |
| ENSMUSG00000025041 | Nt5c2 | 5190.88821 | 3258.93324 | -1931.955 | -0.6455248 | 4.3563E-06 | 2.2416E-05 |
| ENSMUSG00000026974 | Zmynd19 | 5374.69209 | 3418.72413 | -1955.968 | -0.645716 | 8.5165E-18 | 1.7787E-16 |
| ENSMUSG00000062070 | Pgk1 | 83628.3386 | 53273.2981 | -30355.04 | -0.6458176 | 1.5442E-25 | 5.9599E-24 |
| ENSMUSG00000025487 | Psmd13 | 9821.61319 | 6247.75269 | -3573.8605 | -0.6459683 | 2.902E-19 | 6.9017E-18 |
| ENSMUSG00000020917 | Acly | 17539.4182 | 11151.4592 | -6387.9589 | -0.6463455 | 7.4562E-18 | 1.5636E-16 |
| ENSMUSG00000075229 | Ccdc58 | 2965.26445 | 1878.72732 | -1086.5371 | -0.646504 | 8.1959E-12 | 9.1834E-11 |
| ENSMUSG00000036372 | Tmem258 | 4648.03239 | 2953.92801 | -1694.1044 | -0.6465352 | 1.5198E-17 | 3.0713E-16 |
| ENSMUSG00000060860 | Ube2s | 6912.47762 | 4375.28925 | -2537.1884 | -0.6466795 | 1.6724E-10 | 1.6032E-09 |
| ENSMUSG00000046722 | Cdc42se1 | 13539.187 | 8616.43748 | -4922.7495 | -0.6478361 | 2.2106E-29 | 1.1011E-27 |
| ENSMUSG00000027843 | Ptpn22 | 6725.00114 | 4240.57138 | -2484.4298 | -0.6479179 | 2.0988E-08 | 1.5225E-07 |
| ENSMUSG00000081974 | - | 54.6429701 | 32.4382655 | -22.204705 | -0.6480463 | 0.00845458 | 0.02288822 |
| ENSMUSG00000059743 | Fdps | 8024.71012 | 4768.99061 | -3255.7195 | -0.6481192 | 0.00980192 | 0.02606167 |
| ENSMUSG00000022962 | Gart | 7669.0213 | 4872.51043 | -2796.5109 | -0.6484356 | 8.592E-21 | 2.3227E-19 |
| ENSMUSG00000026078 | Pdcl3 | 6569.94157 | 4164.5426 | -2405.399 | -0.6484908 | 3.1595E-14 | 4.6195E-13 |
| ENSMUSG00000071172 | Srsf3 | 28097.9478 | 17738.0552 | -10359.893 | -0.6486787 | 1.8606E-09 | 1.5496E-08 |
| ENSMUSG00000067547 | - | 153.562116 | 95.2854858 | -58.27663 | -0.6492857 | 4.995E-05 | 0.00021679 |
| ENSMUSG00000079523 | Tmsb10 | 35888.2597 | 22735.8356 | -13152.424 | -0.6494089 | 2.0409E-14 | 3.0249E-13 |
| ENSMUSG00000044595 | Dnd1 | 165.849594 | 103.133836 | -62.715757 | -0.6497141 | 1.9818E-05 | 9.2012E-05 |
| ENSMUSG00000028063 | Lmna | 56234.4575 | 35753.5075 | -20480.95 | -0.6497737 | 7.3922E-34 | 4.9214E-32 |
| ENSMUSG00000025027 | Xpnpep1 | 17462.9424 | 11099.5495 | -6363.3929 | -0.6500361 | 4.0336E-33 | 2.5376E-31 |

| | | | | | | | |
|---|---|---|---|---|---|---|---|
| ENSMUSG00000001761 | Smo | 10103.1354 | 6402.64224 | -3700.4932 | -0.6503962 | 9.8703E-17 | 1.8715E-15 |
| ENSMUSG00000090761 | - | 344.808424 | 216.495519 | -128.31291 | -0.6504511 | 4.2963E-07 | 2.5798E-06 |
| ENSMUSG00000034926 | Dhcr24 | 2962.37163 | 1865.84337 | -1096.5283 | -0.650546 | 7.1824E-09 | 5.5715E-08 |
| ENSMUSG00000030056 | Isy1 | 6018.74247 | 3792.98929 | -2225.7532 | -0.6509981 | 1.8208E-09 | 1.5184E-08 |
| ENSMUSG00000003051 | Elf3 | 1077.10505 | 671.20368 | -405.90137 | -0.6511565 | 1.7771E-05 | 8.301E-05 |
| ENSMUSG00000021707 | Dhfr | 6382.58928 | 4023.56789 | -2359.0214 | -0.6513136 | 5.9391E-10 | 5.2587E-09 |
| ENSMUSG00000030216 | Wbp11 | 10366.0396 | 6577.63254 | -3788.4071 | -0.6517402 | 1.2422E-27 | 5.5196E-26 |
| ENSMUSG00000090210 | Itga10 | 191.962689 | 119.165478 | -72.797211 | -0.6523874 | 0.00010005 | 0.00041197 |
| ENSMUSG00000097751 | - | 57.8149639 | 34.6901042 | -23.12486 | -0.6524047 | 0.00519235 | 0.01488323 |
| ENSMUSG00000085183 | - | 1225.57539 | 759.643754 | -465.93163 | -0.6525516 | 6.7677E-05 | 0.00028771 |
| ENSMUSG00000026869 | Psmd5 | 12897.108 | 8159.25797 | -4737.8501 | -0.6527252 | 8.4466E-17 | 1.6104E-15 |
| ENSMUSG00000075570 | Krt26 | 1.98997803 | 0 | -1.989978 | -0.6529698 | 0.0115849 | 0.03020207 |
| ENSMUSG00000091243 | Vgll3 | 4319.97942 | 2715.72761 | -1604.2518 | -0.6531315 | 7.4613E-09 | 5.7726E-08 |
| ENSMUSG00000024534 | Sncaip | 2364.29039 | 1366.03673 | -998.25366 | -0.6533667 | 0.01851561 | 0.0454701 |
| ENSMUSG00000040850 | Psme4 | 13734.5094 | 8638.8687 | -5095.6407 | -0.6535294 | 2.1445E-09 | 1.7739E-08 |
| ENSMUSG00000059316 | Slc27a4 | 4588.4047 | 2898.13394 | -1690.2708 | -0.6541312 | 4.4343E-15 | 7.1185E-14 |
| ENSMUSG00000015217 | Hmgb3 | 2252.3864 | 1405.80628 | -846.58011 | -0.65426 | 2.4337E-06 | 1.2997E-05 |
| ENSMUSG00000026525 | Opn3 | 72.334221 | 43.1073313 | -29.22689 | -0.6548411 | 0.00686291 | 0.01904799 |
| ENSMUSG00000023089 | Ndufa5 | 5719.1554 | 3603.91559 | -2115.2398 | -0.6551667 | 1.312E-12 | 1.5961E-11 |
| ENSMUSG00000090137 | Uba52 | 1726.36945 | 1082.68245 | -643.687 | -0.6557169 | 1.3135E-08 | 9.7829E-08 |
| ENSMUSG00000095454 | - | 154.542074 | 95.015829 | -59.526245 | -0.6558539 | 0.00040672 | 0.00148826 |
| ENSMUSG00000023988 | Bysl | 4484.21885 | 2833.07419 | -1651.1447 | -0.6559354 | 1.0192E-19 | 2.505E-18 |
| ENSMUSG00000026980 | Ly75 | 6899.14629 | 4350.97822 | -2548.1681 | -0.6561812 | 8.1352E-15 | 1.2628E-13 |
| ENSMUSG00000024269 | Tpgs2 | 3618.03079 | 2288.53625 | -1329.4945 | -0.6567763 | 9.9988E-37 | 8.035E-35 |
| ENSMUSG00000073542 | Cep76 | 3663.52479 | 2316.74979 | -1346.775 | -0.6576961 | 4.1433E-38 | 3.5515E-36 |
| ENSMUSG00000024844 | Banf1 | 13346.7496 | 8403.54405 | -4943.2055 | -0.6583106 | 5.1718E-15 | 8.2031E-14 |
| ENSMUSG00000029371 | Cxcl5 | 345943.823 | 216703.308 | -129240.52 | -0.6586639 | 3.2292E-09 | 2.6144E-08 |
| ENSMUSG00000092358 | - | 30.2413811 | 17.1341475 | -13.107234 | -0.6587573 | 0.01849516 | 0.04542532 |
| ENSMUSG00000030823 | 9130019O22 | 134.878776 | 81.8996993 | -52.979077 | -0.6594246 | 0.00126837 | 0.00419768 |
| ENSMUSG00000022792 | Yars2 | 3160.84934 | 1981.60502 | -1179.2443 | -0.6599093 | 1.0453E-10 | 1.024E-09 |
| ENSMUSG00000082878 | - | 50.9593164 | 30.0309203 | -20.928396 | -0.6600502 | 0.00687157 | 0.01906948 |
| ENSMUSG00000046756 | Mrps7 | 6567.17063 | 4127.23744 | -2439.9332 | -0.6603804 | 1.5114E-14 | 2.2812E-13 |
| ENSMUSG00000096536 | - | 36.1085784 | 21.0829925 | -15.025586 | -0.6606238 | 0.01170407 | 0.03046086 |
| ENSMUSG00000030512 | Snrpa1 | 5041.76397 | 3169.23137 | -1872.5326 | -0.6611593 | 2.6568E-16 | 4.8071E-15 |
| ENSMUSG00000015671 | Psma2 | 22991.5637 | 14456.996 | -8534.5677 | -0.6613716 | 3.7633E-17 | 7.3803E-16 |
| ENSMUSG00000056076 | Eif3b | 41055.1843 | 25871.7959 | -15183.388 | -0.6614606 | 2.3386E-27 | 1.0214E-25 |
| ENSMUSG00000031231 | Cox7b | 14497.5809 | 9103.51823 | -5394.0627 | -0.661844 | 7.644E-15 | 1.1904E-13 |
| ENSMUSG00000028961 | Pgd | 20161.9192 | 12713.0613 | -7448.8579 | -0.6619428 | 4.8452E-38 | 4.1359E-36 |
| ENSMUSG00000090610 | - | 74.4377814 | 44.2173157 | -30.220466 | -0.6623286 | 0.00512244 | 0.01470743 |
| ENSMUSG00000081999 | - | 859.935805 | 534.727225 | -325.20858 | -0.6625531 | 5.4209E-07 | 3.2046E-06 |
| ENSMUSG00000030335 | Mrpl51 | 9078.26752 | 5676.42602 | -3401.8415 | -0.6640669 | 4.8534E-11 | 4.9208E-10 |
| ENSMUSG00000057228 | Aadat | 46.0210579 | 26.3670281 | -19.65403 | -0.6642797 | 0.0187326 | 0.04593718 |
| ENSMUSG00000095203 | - | 38.0906967 | 22.0639427 | -16.026754 | -0.6643932 | 0.01118599 | 0.02927341 |
| ENSMUSG00000036867 | Smad6 | 1410.31469 | 874.888808 | -535.42588 | -0.6644906 | 7.9663E-07 | 4.6021E-06 |
| ENSMUSG00000051246 | Msantd1 | 40.3682773 | 22.7952679 | -17.573009 | -0.6646025 | 0.01884603 | 0.04615483 |
| ENSMUSG00000069310 | H3c3 | 46.0976601 | 26.8857839 | -19.211876 | -0.6647416 | 0.0098029 | 0.02606167 |
| ENSMUSG00000070461 | - | 147.974869 | 89.6124757 | -58.362394 | -0.6648717 | 0.00069975 | 0.00243945 |
| ENSMUSG00000039356 | Exosc2 | 4712.6512 | 2956.80858 | -1755.8426 | -0.6654677 | 2.0829E-20 | 5.3898E-19 |
| ENSMUSG00000041773 | Enc1 | 5881.24266 | 3675.93155 | -2205.3111 | -0.665478 | 1.2976E-11 | 1.4146E-10 |
| ENSMUSG00000097537 | - | 174.619392 | 107.139119 | -67.480273 | -0.6658922 | 3.2747E-05 | 0.00014639 |
| ENSMUSG00000051579 | Tceal8 | 11147.3112 | 6985.99733 | -4161.3138 | -0.6662938 | 7.8267E-18 | 1.6363E-16 |
| ENSMUSG00000002017 | Fam98a | 8249.41422 | 5159.46772 | -3089.9465 | -0.6666267 | 9.4053E-14 | 1.3003E-12 |
| ENSMUSG00000070972 | Dnajc25 | 1639.95727 | 1028.22761 | -611.72965 | -0.6667623 | 3.8043E-21 | 1.0648E-19 |
| ENSMUSG00000096917 | 2500002B13 | 697.408733 | 434.162862 | -263.24587 | -0.6673928 | 2.5824E-09 | 2.1115E-08 |
| ENSMUSG00000091460 | - | 58.1725793 | 33.9586817 | -24.213898 | -0.6674447 | 0.00879953 | 0.02367565 |
| ENSMUSG00000038965 | Ube2l3 | 6942.15707 | 4360.36783 | -2581.7892 | -0.6677687 | 5.0349E-43 | 5.6911E-41 |
| ENSMUSG00000063889 | Crem | 5510.46706 | 3436.14442 | -2074.3226 | -0.6684143 | 1.5194E-11 | 1.6425E-10 |
| ENSMUSG00000005615 | Pcyt1a | 6825.58015 | 4273.19224 | -2552.3879 | -0.6686106 | 1.1579E-19 | 2.8425E-18 |
| ENSMUSG00000046312 | Myorg | 997.452592 | 625.087667 | -372.36492 | -0.6686223 | 9.3986E-21 | 2.5307E-19 |
| ENSMUSG00000071866 | Ppia | 92462.6594 | 57659.448 | -34803.211 | -0.668993 | 5.218E-12 | 5.9504E-11 |
| ENSMUSG00000051339 | 2900026A02 | 2033.224 | 1264.01157 | -769.21243 | -0.6691271 | 2.1601E-09 | 1.7861E-08 |
| ENSMUSG00000023236 | Scg5 | 74.1661207 | 41.8670211 | -32.2991 | -0.6692716 | 0.01863157 | 0.04571668 |
| ENSMUSG00000028068 | Iqgap3 | 5781.63334 | 3607.02041 | -2174.6129 | -0.6696636 | 3.4147E-13 | 4.4432E-12 |
| ENSMUSG00000020163 | Uqcr11 | 6484.10961 | 4042.12775 | -2441.9819 | -0.670062 | 8.446E-13 | 1.0492E-11 |
| ENSMUSG00000091478 | - | 88.6253704 | 53.1072215 | -35.518149 | -0.6701184 | 0.00094244 | 0.00320197 |
| ENSMUSG00000062737 | Prl3d2 | 2.11226021 | 0.14394265 | -1.9683176 | -0.6701438 | 0.02004457 | 0.04866142 |
| ENSMUSG00000029430 | Ran | 57420.0342 | 35849.5777 | -21570.456 | -0.6703303 | 1.9073E-15 | 3.1771E-14 |
| ENSMUSG00000098123 | - | 83.0917646 | 48.9454419 | -34.146323 | -0.6707293 | 0.00444526 | 0.01297317 |
| ENSMUSG00000009376 | Met | 8641.89357 | 5382.60653 | -3259.287 | -0.6711008 | 2.2962E-12 | 2.7273E-11 |
| ENSMUSG00000022519 | Srl | 919.09551 | 567.52631 | -351.5692 | -0.6711558 | 9.2023E-07 | 5.2542E-06 |
| ENSMUSG00000040269 | Mrps28 | 1775.1048 | 1100.055 | -675.0498 | -0.6715158 | 1.0225E-08 | 7.7621E-08 |
| ENSMUSG00000033735 | Spr | 5575.0023 | 3467.62555 | -2107.3768 | -0.671519 | 3.1166E-11 | 3.2529E-10 |
| ENSMUSG00000030276 | Ttll3 | 340.812304 | 207.431022 | -133.38128 | -0.6721062 | 0.00012095 | 0.00049018 |
| ENSMUSG00000024810 | Il33 | 29041.4713 | 16810.121 | -12231.35 | -0.6723091 | 0.00950158 | 0.0253359 |
| ENSMUSG00000054312 | Mrps21 | 4285.20645 | 2655.9042 | -1629.3023 | -0.6723825 | 5.0196E-09 | 3.958E-08 |

| | | | | | | | |
|---|---|---|---|---|---|---|---|
| ENSMUSG00000093483 | AA465934 | 925.172016 | 573.05006 | -352.12196 | -0.6727767 | 3.3052E-09 | 2.6706E-08 |
| ENSMUSG00000072594 | - | 71.2156289 | 41.8521509 | -29.363478 | -0.672875 | 0.00498506 | 0.01435305 |
| ENSMUSG00000022814 | Umps | 8859.05372 | 5535.33022 | -3323.7235 | -0.6729543 | 4.9169E-25 | 1.8324E-23 |
| ENSMUSG00000065226 | Gm25791 | 97.6558617 | 59.1358113 | -38.52005 | -0.673078 | 0.00018615 | 0.0007291 |
| ENSMUSG00000037572 | Wdhd1 | 7527.74169 | 4705.4607 | -2822.281 | -0.6733308 | 2.5401E-30 | 1.3608E-28 |
| ENSMUSG00000035834 | Polr3g | 1284.41597 | 800.391605 | -484.02437 | -0.6738608 | 6.9472E-19 | 1.604E-17 |
| ENSMUSG00000049124 | - | 1987.94387 | 1232.55261 | -755.39126 | -0.6738948 | 4.5173E-10 | 4.0759E-09 |
| ENSMUSG00000039942 | Ptger4 | 367.823521 | 216.576215 | -151.24731 | -0.6738979 | 0.00394873 | 0.01167985 |
| ENSMUSG00000053801 | Grwd1 | 3500.75427 | 2179.31674 | -1321.4375 | -0.6744129 | 9.506E-16 | 1.6351E-14 |
| ENSMUSG00000091639 | - | 38.6651274 | 22.3114587 | -16.353669 | -0.6750591 | 0.00929831 | 0.02484542 |
| ENSMUSG00000041220 | Elovl6 | 3351.46673 | 2001.68903 | -1349.7777 | -0.6753716 | 0.00134315 | 0.00441605 |
| ENSMUSG00000024965 | Fermt3 | 89.7700972 | 53.7480717 | -36.022025 | -0.6760547 | 0.00051006 | 0.00183064 |
| ENSMUSG00000031633 | Slc25a4 | 45680.8048 | 28458.2514 | -17222.553 | -0.6761918 | 1.1754E-21 | 3.4009E-20 |
| ENSMUSG00000083044 | - | 130.579146 | 79.9149695 | -50.664177 | -0.6763259 | 1.5189E-05 | 7.1718E-05 |
| ENSMUSG00000043155 | Hpdl | 139.103949 | 83.7718007 | -55.332148 | -0.6764933 | 0.00031715 | 0.00118646 |
| ENSMUSG00000038759 | Nup205 | 20291.9035 | 12649.143 | -7642.7605 | -0.6768086 | 6.5754E-27 | 2.7948E-25 |
| ENSMUSG00000002102 | Psmc3 | 23544.2905 | 14640.9306 | -8903.3599 | -0.6770557 | 1.5542E-17 | 3.1346E-16 |
| ENSMUSG00000059796 | Eif4a1 | 76443.8752 | 47362.269 | -29081.606 | -0.6770689 | 2.1285E-11 | 2.2676E-10 |
| ENSMUSG00000031613 | Hpgd | 160.523385 | 93.43829 | -67.085095 | -0.6790335 | 0.00558432 | 0.01587627 |
| ENSMUSG00000034424 | Gcsh | 6505.35699 | 4037.74568 | -2467.6113 | -0.6792874 | 7.8164E-17 | 1.4958E-15 |
| ENSMUSG00000068749 | Psma5 | 17234.7774 | 10671.7253 | -6563.0521 | -0.6800293 | 2.4874E-13 | 3.2823E-12 |
| ENSMUSG00000056952 | Tatdn2 | 8347.59427 | 5201.45115 | -3146.1431 | -0.6802877 | 1.0567E-65 | 3.623E-63 |
| ENSMUSG00000001105 | Ift20 | 3689.98467 | 2285.15441 | -1404.8303 | -0.6807987 | 9.3637E-15 | 1.4429E-13 |
| ENSMUSG00000036850 | Mrpl41 | 2309.9388 | 1423.77209 | -886.16671 | -0.6808185 | 1.007E-09 | 8.6319E-09 |
| ENSMUSG00000062580 | Timm17a | 6119.68202 | 3785.15153 | -2334.5305 | -0.6809137 | 7.3858E-13 | 9.2141E-12 |
| ENSMUSG00000077575 | Gm24888 | 35.9396545 | 20.440814 | -15.498841 | -0.6809237 | 0.0118033 | 0.03067427 |
| ENSMUSG00000029394 | Cdk2ap1 | 3589.14643 | 2220.36621 | -1368.7802 | -0.6810504 | 4.9021E-13 | 6.2405E-12 |
| ENSMUSG00000006641 | Slc5a6 | 1158.02182 | 719.02256 | -438.99926 | -0.6812888 | 4.1979E-21 | 1.167E-19 |
| ENSMUSG00000001517 | Foxm1 | 10194.1461 | 6317.23746 | -3876.9086 | -0.6815544 | 7.6865E-17 | 1.4751E-15 |
| ENSMUSG00000018293 | Pfn1 | 41761.8558 | 25865.5113 | -15896.345 | -0.6820727 | 1.7298E-16 | 3.1772E-15 |
| ENSMUSG00000030494 | Rhpn2 | 776.297345 | 468.725166 | -307.57218 | -0.6832162 | 8.8731E-05 | 0.00036854 |
| ENSMUSG00000092074 | Dynlt1a | 466.059441 | 286.243609 | -179.81583 | -0.6832381 | 2.6051E-09 | 2.1292E-08 |
| ENSMUSG00000090136 | - | 440.686715 | 269.267581 | -171.41913 | -0.6833223 | 4.3926E-07 | 2.6361E-06 |
| ENSMUSG00000003444 | Med29 | 1032.21356 | 633.366852 | -398.84671 | -0.6834521 | 4.0723E-08 | 2.836E-07 |
| ENSMUSG00000036299 | BC031181 | 6728.77156 | 4168.63897 | -2560.1326 | -0.684386 | 1.4061E-23 | 4.6958E-22 |
| ENSMUSG00000020439 | Smtn | 3036.9718 | 1877.67805 | -1159.2937 | -0.6844917 | 2.0314E-16 | 3.7049E-15 |
| ENSMUSG00000026192 | Atic | 16995.6384 | 10543.6209 | -6452.0175 | -0.6845719 | 2.1947E-33 | 1.4198E-31 |
| ENSMUSG00000029547 | Ints1 | 8616.39855 | 5305.46319 | -3310.9354 | -0.684721 | 7.674E-11 | 7.6119E-10 |
| ENSMUSG00000064341 | ND1 | 269349.602 | 164526.002 | -104823.6 | -0.6848126 | 6.4487E-07 | 3.7709E-06 |
| ENSMUSG00000038236 | Hoxa7 | 2159.87293 | 1335.31853 | -824.5544 | -0.6851085 | 1.1299E-17 | 2.3198E-16 |
| ENSMUSG00000024268 | Celf4 | 6762.74039 | 4000.8516 | -2761.8888 | -0.6851997 | 0.00137987 | 0.00452623 |
| ENSMUSG00000032279 | Idh3a | 10328.3579 | 6360.97864 | -3967.3792 | -0.685999 | 4.5445E-12 | 5.2112E-11 |
| ENSMUSG00000042541 | Sem1 | 22992.8316 | 14027.3908 | -8965.4408 | -0.6863995 | 6.4074E-07 | 3.7479E-06 |
| ENSMUSG00000024952 | - | 8028.48043 | 4969.32462 | -3059.1558 | -0.6870877 | 3.7226E-30 | 1.9737E-28 |
| ENSMUSG00000004665 | Cnn2 | 29091.8957 | 17954.0348 | -11137.861 | -0.6875421 | 1.6773E-17 | 3.3664E-16 |
| ENSMUSG00000050856 | Atp5k | 5203.00011 | 3205.23296 | -1997.7672 | -0.6881377 | 7.1574E-15 | 1.1172E-13 |
| ENSMUSG00000026864 | Hspa5 | 136866.221 | 83788.0093 | -53078.212 | -0.6882595 | 8.9924E-09 | 6.877E-08 |
| ENSMUSG00000030357 | Fkbp4 | 36864.2721 | 22759.4463 | -14104.826 | -0.6884813 | 1.2896E-20 | 3.4187E-19 |
| ENSMUSG00000022978 | Mis18a | 3048.92581 | 1879.79862 | -1169.1272 | -0.6885527 | 2.0815E-17 | 4.1493E-16 |
| ENSMUSG00000097769 | - | 1353.737 | 827.416924 | -526.32007 | -0.6888569 | 3.6602E-08 | 2.5638E-07 |
| ENSMUSG00000028318 | Polr1e | 1758.97577 | 1082.63716 | -676.33861 | -0.6893407 | 3.5953E-14 | 5.2234E-13 |
| ENSMUSG00000024764 | Naa40 | 4051.5645 | 2500.92354 | -1550.641 | -0.6900828 | 1.3703E-25 | 5.2987E-24 |
| ENSMUSG00000068303 | - | 98.9692034 | 58.0566301 | -40.912573 | -0.6902392 | 0.00151881 | 0.00493447 |
| ENSMUSG00000082816 | - | 74.2641916 | 43.3723165 | -30.891875 | -0.6910627 | 0.00174699 | 0.00559712 |
| ENSMUSG00000031137 | Fgf13 | 8099.72843 | 4773.46472 | -3326.2637 | -0.6912181 | 0.0011648 | 0.00388555 |
| ENSMUSG00000070063 | - | 97.26161 | 55.0261459 | -42.235464 | -0.6918884 | 0.00852731 | 0.02306387 |
| ENSMUSG00000044937 | Ttc41 | 186.78692 | 112.241072 | -74.545849 | -0.6920684 | 3.8948E-05 | 0.00017183 |
| ENSMUSG00000098238 | - | 67.5064935 | 38.6233438 | -28.88315 | -0.6921657 | 0.00637811 | 0.0178598 |
| ENSMUSG00000096171 | - | 48.0741508 | 27.054813 | -21.019338 | -0.6926802 | 0.00855796 | 0.02313765 |
| ENSMUSG00000037070 | Rbmxl1 | 7438.4912 | 4584.57408 | -2853.9171 | -0.6929189 | 2.5315E-28 | 1.1836E-26 |
| ENSMUSG00000029147 | Ppm1g | 19546.7825 | 12053.7378 | -7493.0446 | -0.6929993 | 3.675E-33 | 2.3191E-31 |
| ENSMUSG00000048482 | Bdnf | 7501.90442 | 4595.44742 | -2906.457 | -0.6932585 | 5.5133E-12 | 6.2663E-11 |
| ENSMUSG00000021131 | Erh | 8681.36611 | 5322.21877 | -3359.1473 | -0.6937399 | 2.2915E-13 | 3.0354E-12 |
| ENSMUSG00000085030 | - | 147.614612 | 87.4983331 | -60.116279 | -0.6945308 | 0.00022663 | 0.00087503 |
| ENSMUSG00000081669 | - | 584.413879 | 357.105106 | -227.30877 | -0.6945697 | 1.5027E-11 | 1.6262E-10 |
| ENSMUSG00000030554 | Synm | 68.9685609 | 38.3854115 | -30.583149 | -0.6947663 | 0.01221391 | 0.03163744 |
| ENSMUSG00000019731 | Slc35e1 | 12708.105 | 7826.46251 | -4881.6425 | -0.6951353 | 1.2088E-35 | 9.01E-34 |
| ENSMUSG00000040721 | Zfhx2 | 700.52729 | 416.725943 | -283.80135 | -0.6955726 | 0.00023646 | 0.00090977 |
| ENSMUSG00000030008 | Pradc1 | 894.467807 | 545.538368 | -348.92944 | -0.6957723 | 3.5652E-10 | 3.2713E-09 |
| ENSMUSG00000078945 | Naip2 | 32.0339677 | 17.8533878 | -14.18058 | -0.6964801 | 0.00877841 | 0.02363427 |
| ENSMUSG00000022204 | Ngdn | 4772.2345 | 2912.53354 | -1859.701 | -0.6972187 | 1.8939E-11 | 2.0302E-10 |
| ENSMUSG00000019970 | Sgk1 | 13033.4193 | 7918.78462 | -5114.6347 | -0.6972894 | 1.9324E-08 | 1.4077E-07 |
| ENSMUSG00000078974 | Sec61g | 5292.46329 | 3236.52899 | -2055.9343 | -0.6983781 | 3.8419E-15 | 6.1989E-14 |
| ENSMUSG00000001729 | Akt1 | 29674.3074 | 18218.5518 | -11455.756 | -0.6984274 | 1.9105E-28 | 9.0146E-27 |
| ENSMUSG00000094151 | - | 47.0702899 | 25.3518494 | -21.718441 | -0.6987266 | 0.0199385 | 0.04844394 |

| | | | | | | | |
|---|---|---|---|---|---|---|---|
| ENSMUSG00000001403 | Ube2c | 15179.5136 | 9292.79027 | -5886.7233 | -0.6987463 | 1.5692E-17 | 3.1619E-16 |
| ENSMUSG00000063800 | Prpf38a | 4853.99142 | 2963.59889 | -1890.3925 | -0.6989202 | 4.4462E-13 | 5.706E-12 |
| ENSMUSG00000028194 | Ddah1 | 5941.24082 | 3586.38848 | -2354.8523 | -0.6992343 | 9.1088E-07 | 5.2037E-06 |
| ENSMUSG00000057666 | Gapdh | 206706.753 | 126694.192 | -80012.561 | -0.6992567 | 2.4464E-22 | 7.4338E-21 |
| ENSMUSG00000021290 | Atp5mpl | 6270.35531 | 3819.97796 | -2450.3774 | -0.6995536 | 2.983E-11 | 3.1246E-10 |
| ENSMUSG00000028655 | Mfsd2a | 214.886369 | 129.277313 | -85.609056 | -0.6998945 | 4.6165E-06 | 2.3619E-05 |
| ENSMUSG00000029484 | Anxa3 | 62747.6294 | 38273.7302 | -24473.899 | -0.6999026 | 1.1654E-12 | 1.4254E-11 |
| ENSMUSG00000055675 | Kbtbd11 | 27.4771264 | 14.8925705 | -12.584556 | -0.7006071 | 0.01562595 | 0.03920685 |
| ENSMUSG00000098187 | - | 35.5550644 | 19.0422207 | -16.512844 | -0.7009384 | 0.02064054 | 0.04991386 |
| ENSMUSG00000094873 | - | 38.0906967 | 21.4056045 | -16.685092 | -0.700993 | 0.00749203 | 0.0205666 |
| ENSMUSG00000045282 | Tmem86b | 66.7699154 | 37.8116667 | -28.958249 | -0.7015564 | 0.00526635 | 0.01507225 |
| ENSMUSG00000091021 | - | 46.805649 | 27.1615698 | -19.644079 | -0.7025398 | 0.00265253 | 0.00814324 |
| ENSMUSG00000021951 | Eef1akmt1 | 2903.40779 | 1763.5251 | -1139.8827 | -0.7025553 | 1.3067E-10 | 1.2721E-09 |
| ENSMUSG00000033938 | Ndufb7 | 8147.27737 | 4957.50227 | -3189.7751 | -0.7027094 | 2.2358E-12 | 2.6602E-11 |
| ENSMUSG00000006442 | Srm | 8897.14045 | 5398.2162 | -3498.9242 | -0.7036022 | 2.5778E-10 | 2.416E-09 |
| ENSMUSG00000078440 | Dohh | 4771.24407 | 2909.47717 | -1861.7669 | -0.7038367 | 5.5136E-17 | 1.068E-15 |
| ENSMUSG00000003948 | Mmd | 5266.3913 | 3192.49044 | -2073.9009 | -0.704444 | 3.4413E-10 | 3.1633E-09 |
| ENSMUSG00000024258 | Polr2d | 3692.39223 | 2251.25972 | -1441.1325 | -0.7054535 | 5.0005E-20 | 1.2638E-18 |
| ENSMUSG00000089838 | - | 68.6806494 | 40.1649802 | -28.515669 | -0.7055635 | 0.00076834 | 0.00265829 |
| ENSMUSG00000030431 | Tmem238 | 294.016352 | 174.391751 | -119.6246 | -0.7055881 | 7.8758E-05 | 0.00032971 |
| ENSMUSG00000038429 | Usp5 | 18326.0926 | 11215.2284 | -7110.8642 | -0.7056835 | 9.1775E-56 | 1.9464E-53 |
| ENSMUSG00000082809 | - | 1856.74096 | 1119.8132 | -736.92776 | -0.7058662 | 3.4666E-08 | 2.4381E-07 |
| ENSMUSG00000060639 | H4c9 | 596.279944 | 359.374842 | -236.9051 | -0.7059988 | 4.9048E-08 | 3.3667E-07 |
| ENSMUSG00000035754 | Wdr18 | 5178.93194 | 3153.61789 | -2025.3141 | -0.7064331 | 3.107E-18 | 6.778E-17 |
| ENSMUSG00000055762 | Eef1d | 25657.8989 | 15622.7421 | -10035.157 | -0.7067593 | 2.4131E-18 | 5.3037E-17 |
| ENSMUSG00000028937 | Acot7 | 8664.96305 | 5242.8715 | -3422.0916 | -0.707789 | 1.2526E-10 | 1.223E-09 |
| ENSMUSG00000001380 | Hars | 18225.0178 | 11094.058 | -7130.9598 | -0.7088556 | 2.2999E-22 | 7.0302E-21 |
| ENSMUSG00000070493 | Chchd2 | 23953.1999 | 14524.6837 | -9428.5162 | -0.7090083 | 1.1664E-13 | 1.5975E-12 |
| ENSMUSG00000028851 | Nudc | 12388.499 | 7517.61431 | -4870.8847 | -0.7092644 | 3.9739E-15 | 6.4068E-14 |
| ENSMUSG00000049751 | Rpl36al | 10335.913 | 6268.56417 | -4067.3488 | -0.709326 | 2.7016E-14 | 3.9755E-13 |
| ENSMUSG00000058126 | - | 443.678616 | 267.434711 | -176.24391 | -0.7097775 | 3.5807E-09 | 2.8786E-08 |
| ENSMUSG00000029632 | Ndufa4 | 18538.816 | 11180.7293 | -7358.0867 | -0.7102106 | 1.218E-09 | 1.0363E-08 |
| ENSMUSG00000027531 | Impa1 | 11181.1271 | 6791.79935 | -4389.3278 | -0.7103155 | 1.1055E-18 | 2.5186E-17 |
| ENSMUSG00000036285 | Noa1 | 5924.72435 | 3599.1171 | -2325.6072 | -0.7104517 | 1.9685E-19 | 4.7418E-18 |
| ENSMUSG00000007721 | Ccdc124 | 8365.25904 | 5064.59789 | -3300.6612 | -0.7105851 | 3.362E-13 | 4.3885E-12 |
| ENSMUSG00000028367 | Txn1 | 37216.7605 | 22537.8705 | -14678.89 | -0.710682 | 1.6038E-13 | 2.1564E-12 |
| ENSMUSG00000004460 | Dnajb11 | 17761.2798 | 10802.7244 | -6958.5554 | -0.7108735 | 2.4504E-25 | 9.3523E-24 |
| ENSMUSG00000058351 | Smim4 | 700.497549 | 420.994621 | -279.50293 | -0.7112358 | 2.8393E-08 | 2.0204E-07 |
| ENSMUSG00000026201 | Stk16 | 5263.01094 | 3196.11352 | -2066.8974 | -0.7117564 | 5.1122E-21 | 1.4116E-19 |
| ENSMUSG00000025935 | Tram1 | 11496.9163 | 6996.09704 | -4500.8192 | -0.7123149 | 5.1245E-39 | 4.7919E-37 |
| ENSMUSG00000024870 | Rab1b | 16853.3537 | 10219.3377 | -6634.0159 | -0.7132965 | 5.9403E-20 | 1.4885E-18 |
| ENSMUSG00000026068 | Il18rap | 1815.19368 | 1078.10476 | -737.08892 | -0.7141766 | 6.7093E-06 | 3.3501E-05 |
| ENSMUSG00000002835 | Chaf1a | 8010.53953 | 4832.76899 | -3177.7705 | -0.7143413 | 2.944E-12 | 3.4549E-11 |
| ENSMUSG00000039220 | Ppp1r10 | 4546.12048 | 2754.36196 | -1791.7585 | -0.7146243 | 6.3989E-20 | 1.6014E-18 |
| ENSMUSG00000047821 | Trim16 | 2344.11984 | 1410.62686 | -933.49297 | -0.7148971 | 1.3219E-10 | 1.2857E-09 |
| ENSMUSG00000082884 | - | 57.7530331 | 32.687672 | -25.065361 | -0.7161803 | 0.00253719 | 0.00784181 |
| ENSMUSG00000039329 | Tex19.1 | 2.39740822 | 0.17533831 | -2.2220699 | -0.7167005 | 0.01453137 | 0.03682427 |
| ENSMUSG00000031974 | Abcb10 | 4311.61603 | 2611.14379 | -1700.4722 | -0.7182119 | 4.8763E-31 | 2.7711E-29 |
| ENSMUSG00000083669 | - | 231.508391 | 130.683453 | -100.82494 | -0.7192607 | 0.0033436 | 0.01005033 |
| ENSMUSG00000005378 | Bud23 | 3329.66965 | 2013.18665 | -1316.483 | -0.7197436 | 1.7342E-29 | 8.7654E-28 |
| ENSMUSG00000054091 | 1810037I17R | 5981.19301 | 3595.62701 | -2385.566 | -0.7202281 | 3.3523E-13 | 4.3787E-12 |
| ENSMUSG00000025134 | Alyref | 8305.85975 | 5017.76516 | -3288.0946 | -0.7206966 | 1.477E-26 | 6.1633E-25 |
| ENSMUSG00000083579 | - | 3.30512151 | 0.30640226 | -2.9987193 | -0.7207699 | 0.01636422 | 0.04079553 |
| ENSMUSG00000014077 | Chp1 | 18545.444 | 11207.71 | -7337.734 | -0.7209488 | 8.7168E-30 | 4.4943E-28 |
| ENSMUSG00000097610 | A930012L18 | 44.9277604 | 25.1257427 | -19.802018 | -0.7209936 | 0.00361379 | 0.01077745 |
| ENSMUSG00000075701 | Selenos | 4838.44004 | 2918.8192 | -1919.6208 | -0.7221533 | 1.6606E-26 | 6.86E-25 |
| ENSMUSG00000054196 | Cthrc1 | 1905.34202 | 1132.94912 | -772.39289 | -0.7225242 | 1.2006E-07 | 7.7695E-07 |
| ENSMUSG00000003500 | Impdh1 | 4823.85561 | 2909.52695 | -1914.3287 | -0.7229138 | 1.1116E-27 | 4.9714E-26 |
| ENSMUSG00000018068 | Ints2 | 4066.81798 | 2445.83582 | -1620.9822 | -0.7240264 | 3.2903E-18 | 7.1477E-17 |
| ENSMUSG00000053693 | Mast1 | 137.588737 | 79.4962516 | -58.092486 | -0.7244605 | 0.00037324 | 0.0013753 |
| ENSMUSG00000094683 | - | 52.8969036 | 30.1919058 | -22.704998 | -0.7244623 | 0.00155318 | 0.00503206 |
| ENSMUSG00000048578 | Mlec | 45322.8955 | 27208.3018 | -18114.594 | -0.724572 | 1.3519E-15 | 2.2909E-14 |
| ENSMUSG00000023832 | Acat2 | 3199.27333 | 1921.35386 | -1277.9195 | -0.7245981 | 1.972E-16 | 3.5997E-15 |
| ENSMUSG00000019214 | Chtf18 | 3146.16044 | 1890.03325 | -1256.1272 | -0.7247411 | 1.6081E-17 | 3.2338E-16 |
| ENSMUSG00000026032 | Ndufb3 | 3565.2189 | 2146.57459 | -1418.6443 | -0.7248001 | 8.4041E-26 | 3.2994E-24 |
| ENSMUSG00000097979 | - | 150.668149 | 87.8527055 | -62.815444 | -0.7250695 | 4.879E-05 | 0.00021202 |
| ENSMUSG00000037126 | Psd | 541.653068 | 322.060033 | -219.59304 | -0.725117 | 1.6892E-08 | 1.2402E-07 |
| ENSMUSG00000022131 | Gpr180 | 2837.25494 | 1709.97571 | -1127.2792 | -0.7252085 | 1.6234E-34 | 1.1321E-32 |
| ENSMUSG00000086688 | - | 108.007002 | 61.6290042 | -46.377998 | -0.7254867 | 0.00127487 | 0.00421717 |
| ENSMUSG00000063524 | Eno1 | 101774.61 | 60906.4968 | -40868.114 | -0.7258827 | 1.0706E-12 | 1.3148E-11 |
| ENSMUSG00000040048 | Ndufb10 | 5569.45278 | 3346.06822 | -2223.3846 | -0.7263724 | 1.3994E-20 | 3.691E-19 |
| ENSMUSG00000096472 | Cdkn2d | 2445.1969 | 1455.37915 | -989.81775 | -0.7271875 | 1.4999E-09 | 1.2631E-08 |
| ENSMUSG00000022385 | Gtse1 | 2399.25806 | 1441.20393 | -958.05413 | -0.7274237 | 4.0078E-24 | 1.381E-22 |
| ENSMUSG00000030082 | Sec61a1 | 39559.7271 | 23820.8642 | -15738.863 | -0.7283038 | 1.7272E-47 | 2.4171E-45 |
| ENSMUSG00000004980 | Hnrnpa2b1 | 122257.377 | 72882.54 | -49374.837 | -0.7284766 | 5.4556E-11 | 5.5016E-10 |

| | | | | | | | |
|---|---|---|---|---|---|---|---|
| ENSMUSG00000027546 | Atp9a | 53.8299114 | 28.8408658 | -24.989046 | -0.7285277 | 0.01138143 | 0.02973568 |
| ENSMUSG00000030978 | Rrm1 | 31458.2207 | 18851.6355 | -12606.585 | -0.728964 | 1.6226E-18 | 3.6164E-17 |
| ENSMUSG00000051107 | - | 2.70869086 | 0.32899174 | -2.3796991 | -0.7296511 | 0.01932645 | 0.04715736 |
| ENSMUSG00000025364 | Pa2g4 | 27570.579 | 16571.8016 | -10998.777 | -0.7299489 | 1.4573E-38 | 1.3035E-36 |
| ENSMUSG00000022802 | Lmln | 182.058478 | 104.628331 | -77.430148 | -0.7299697 | 0.0003627 | 0.0013408 |
| ENSMUSG00000013629 | Cad | 9331.05272 | 5574.84145 | -3756.2113 | -0.7303753 | 1.4122E-14 | 2.144E-13 |
| ENSMUSG00000026496 | Parp1 | 9924.10219 | 5953.56991 | -3970.5323 | -0.7305785 | 3.465E-26 | 1.3841E-24 |
| ENSMUSG00000042312 | S100a13 | 3132.78483 | 1864.23087 | -1268.554 | -0.7307803 | 3.6221E-11 | 3.7482E-10 |
| ENSMUSG00000057278 | Snrpg | 6532.15106 | 3895.90227 | -2636.2488 | -0.7309573 | 3.6194E-13 | 4.6918E-12 |
| ENSMUSG00000026605 | Cenpf | 7221.1676 | 4282.64058 | -2938.527 | -0.731092 | 3.8714E-09 | 3.0953E-08 |
| ENSMUSG00000042426 | Dhx29 | 4627.46237 | 2781.48826 | -1845.9741 | -0.7311126 | 6.9393E-43 | 7.8009E-41 |
| ENSMUSG00000005087 | Cd44 | 46791.3025 | 27791.3462 | -18999.956 | -0.7320523 | 2.641E-10 | 2.4685E-09 |
| ENSMUSG00000023456 | Tpi1 | 130747.865 | 78307.1298 | -52440.735 | -0.7324094 | 7.9924E-25 | 2.9152E-23 |
| ENSMUSG00000056209 | Npm3 | 4824.8925 | 2881.92869 | -1942.9638 | -0.732741 | 3.1022E-17 | 6.1246E-16 |
| ENSMUSG00000063904 | Dpp3 | 10776.0898 | 6449.04498 | -4327.0448 | -0.732936 | 3.1698E-23 | 1.0367E-21 |
| ENSMUSG00000003934 | Efnb3 | 112.044755 | 63.899736 | -48.145019 | -0.7338174 | 0.00050316 | 0.00180792 |
| ENSMUSG00000064797 | Gm24357 | 77.0894155 | 43.1451864 | -33.944229 | -0.7338678 | 0.00213504 | 0.00671592 |
| ENSMUSG00000028452 | Vcp | 31411.1919 | 18793.2076 | -12617.984 | -0.7340982 | 2.2936E-25 | 8.7866E-24 |
| ENSMUSG00000022557 | Bop1 | 12073.7307 | 7228.74183 | -4844.9889 | -0.7341875 | 3.921E-30 | 2.0736E-28 |
| ENSMUSG00000024800 | Rpp30 | 4278.70229 | 2550.82308 | -1727.8792 | -0.7344862 | 2.0993E-16 | 3.8252E-15 |
| ENSMUSG00000033257 | Ttll4 | 5247.91877 | 3114.76107 | -2133.1577 | -0.7345273 | 5.0069E-11 | 5.064E-10 |
| ENSMUSG00000084071 | - | 3.24679421 | 0.16153669 | -3.0852575 | -0.736388 | 0.01030348 | 0.02722014 |
| ENSMUSG00000040463 | Mybbp1a | 28459.9798 | 17043.2646 | -11416.715 | -0.7366929 | 8.4607E-55 | 1.7405E-52 |
| ENSMUSG00000097102 | - | 105.025932 | 59.3361092 | -45.689823 | -0.7369304 | 0.00096899 | 0.00328458 |
| ENSMUSG00000038252 | Ncapd2 | 18783.8172 | 11184.5168 | -7599.3005 | -0.7371048 | 2.5338E-17 | 5.0318E-16 |
| ENSMUSG00000071528 | Atp5md; Gm3 | 11939.5066 | 7094.46315 | -4845.0435 | -0.7371564 | 4.0164E-14 | 5.8105E-13 |
| ENSMUSG00000038217 | Tlcd2 | 94.3002727 | 53.1890246 | -41.111248 | -0.7375554 | 0.00109401 | 0.00366727 |
| ENSMUSG00000058064 | - | 69.6026901 | 39.5546647 | -30.048025 | -0.737993 | 0.00088652 | 0.00302847 |
| ENSMUSG00000073607 | Gm10548 | 25.5698085 | 12.8824899 | -12.687319 | -0.7384642 | 0.01793451 | 0.04422254 |
| ENSMUSG00000051341 | Zfp52 | 4238.82879 | 2516.21296 | -1722.6158 | -0.7385143 | 2.7088E-14 | 3.9832E-13 |
| ENSMUSG00000040078 | - | 2405.99429 | 1362.4889 | -1043.5054 | -0.7394418 | 0.00070018 | 0.00244054 |
| ENSMUSG00000035150 | Eif2s3x | 6660.23845 | 3980.29551 | -2679.9429 | -0.7399756 | 2.4913E-56 | 5.5708E-54 |
| ENSMUSG00000083725 | - | 19.8708823 | 10.2065535 | -9.6643288 | -0.7401417 | 0.01592872 | 0.03986929 |
| ENSMUSG00000086644 | - | 73.3900423 | 40.8634452 | -32.526597 | -0.7406214 | 0.0015176 | 0.00493249 |
| ENSMUSG00000027429 | Sec23b | 9583.54426 | 5719.29063 | -3864.2536 | -0.740753 | 1.8453E-44 | 2.1943E-42 |
| ENSMUSG00000083396 | - | 129.85767 | 74.6936242 | -55.164046 | -0.7409366 | 6.0737E-05 | 0.00026025 |
| ENSMUSG00000042367 | Gjb3 | 2866.17915 | 1689.45416 | -1176.725 | -0.7411273 | 5.966E-10 | 5.2765E-09 |
| ENSMUSG00000020307 | Cdc34 | 7606.37779 | 4499.75536 | -3106.6224 | -0.7411593 | 1.8784E-12 | 2.2454E-11 |
| ENSMUSG00000020849 | Ywhae | 58841.1025 | 34923.6517 | -23917.451 | -0.7411779 | 8.5294E-17 | 1.6247E-15 |
| ENSMUSG00000024644 | Cndp2 | 6300.66759 | 3754.24971 | -2546.4179 | -0.7413022 | 5.8652E-33 | 3.6453E-31 |
| ENSMUSG00000026667 | Uhmk1 | 6915.20667 | 4095.12272 | -2820.084 | -0.7413996 | 1.205E-13 | 1.6483E-12 |
| ENSMUSG00000042515 | Pwwp3b | 2312.50559 | 1361.34983 | -951.15577 | -0.7414394 | 1.8132E-09 | 1.5132E-08 |
| ENSMUSG00000062590 | Armc9 | 534.856057 | 314.848064 | -220.00799 | -0.741531 | 3.7758E-09 | 3.0236E-08 |
| ENSMUSG00000005410 | Mcm5 | 25218.1413 | 14998.6241 | -10219.517 | -0.7415605 | 4.7352E-23 | 1.5292E-21 |
| ENSMUSG00000014551 | Mrps25 | 5905.39042 | 3511.23108 | -2394.1593 | -0.7416517 | 1.0188E-22 | 3.2195E-21 |
| ENSMUSG00000039183 | Nubp2 | 3959.71824 | 2351.62868 | -1608.0896 | -0.7420817 | 3.9345E-20 | 1.003E-18 |
| ENSMUSG00000092212 | - | 25.5967832 | 12.820067 | -12.776716 | -0.7423109 | 0.01909187 | 0.04665138 |
| ENSMUSG00000041263 | Rusc1 | 1293.67803 | 761.531616 | -532.14642 | -0.7430963 | 3.4553E-10 | 3.1747E-09 |
| ENSMUSG00000098232 | - | 35.1131086 | 17.7126142 | -17.400494 | -0.7432495 | 0.0199035 | 0.04837031 |
| ENSMUSG00000015112 | Slc25a13 | 4476.37468 | 2647.829 | -1828.5457 | -0.7434961 | 3.2482E-14 | 4.7426E-13 |
| ENSMUSG00000025810 | Nrp1 | 22114.5277 | 13043.9837 | -9070.544 | -0.743755 | 1.4902E-11 | 1.6143E-10 |
| ENSMUSG00000030867 | Plk1 | 7210.04642 | 4244.52551 | -2965.5209 | -0.7438075 | 2.5977E-10 | 2.4323E-09 |
| ENSMUSG00000019942 | Cdk1 | 21370.5917 | 12699.6196 | -8670.972 | -0.7438377 | 8.1021E-27 | 3.4296E-25 |
| ENSMUSG00000032370 | Lactb | 1736.4326 | 1029.67055 | -706.76205 | -0.7466055 | 2.0509E-27 | 8.9961E-26 |
| ENSMUSG00000097080 | 1700086O06 | 301.973723 | 176.369834 | -125.60389 | -0.746901 | 4.3408E-09 | 3.4478E-08 |
| ENSMUSG00000001228 | Uhrf1 | 10022.2836 | 5920.56371 | -4101.7199 | -0.747749 | 4.9864E-17 | 9.6956E-16 |
| ENSMUSG00000006456 | Rbm14 | 3153.40592 | 1841.55754 | -1311.8484 | -0.7478212 | 3.8706E-08 | 2.7037E-07 |
| ENSMUSG00000086709 | - | 134.310813 | 76.4007789 | -57.910034 | -0.7482629 | 0.0001602 | 0.00063451 |
| ENSMUSG00000029594 | Rbm19 | 2153.70639 | 1277.45506 | -876.25133 | -0.749458 | 1.1134E-36 | 8.8436E-35 |
| ENSMUSG00000037621 | Atoh8 | 7067.20198 | 3814.9954 | -3252.2066 | -0.7494645 | 0.00488867 | 0.01412098 |
| ENSMUSG00000024966 | Stip1 | 44907.9244 | 26621.6474 | -18286.277 | -0.7499601 | 5.9911E-42 | 6.3859E-40 |
| ENSMUSG00000008373 | Prpf31 | 4839.47044 | 2866.10776 | -1973.3627 | -0.7501688 | 1.9419E-34 | 1.3361E-32 |
| ENSMUSG00000030359 | Pzp | 22.657977 | 11.950929 | -10.707048 | -0.7502074 | 0.00935826 | 0.02498936 |
| ENSMUSG00000097403 | 9230116N13 | 34.8513318 | 18.3963859 | -16.454946 | -0.7502483 | 0.00698752 | 0.01933647 |
| ENSMUSG00000051855 | Mest | 122.073271 | 66.5715699 | -55.501701 | -0.7511082 | 0.00298142 | 0.00905429 |
| ENSMUSG00000001025 | S100a6 | 144052.086 | 84823.6166 | -59228.469 | -0.7515518 | 3.9094E-16 | 6.9813E-15 |
| ENSMUSG00000087808 | - | 2.80483841 | 0.33117404 | -2.4736644 | -0.7517385 | 0.01692293 | 0.04203557 |
| ENSMUSG00000021823 | Vcl | 18600.7822 | 10967.873 | -7632.9091 | -0.7522763 | 6.4675E-20 | 1.6166E-18 |
| ENSMUSG00000082925 | - | 60.4037055 | 34.1278077 | -26.275898 | -0.7530766 | 0.00025858 | 0.00098564 |
| ENSMUSG00000028195 | Ccn1 | 83864.0389 | 48603.4732 | -35260.566 | -0.7536144 | 2.8625E-07 | 1.765E-06 |
| ENSMUSG00000024795 | Kif20b | 11315.182 | 6673.00669 | -4642.1754 | -0.7542777 | 1.046E-25 | 4.0986E-24 |
| ENSMUSG00000003541 | Ier3 | 52384.6945 | 28233.972 | -24150.723 | -0.7550444 | 0.00415118 | 0.01220844 |
| ENSMUSG00000041957 | Pkp2 | 3599.70547 | 2111.87586 | -1487.8296 | -0.7559051 | 2.3138E-15 | 3.8294E-14 |
| ENSMUSG00000063328 | - | 34.4037484 | 17.8675709 | -16.536177 | -0.7564718 | 0.01087877 | 0.02856388 |
| ENSMUSG00000065258 | Gm23969 | 39.1756032 | 21.1438197 | -18.031783 | -0.7567793 | 0.00453554 | 0.01320479 |

| | | | | | | | |
|---|---|---|---|---|---|---|---|
| ENSMUSG00000005103 | Wdr1 | 32382.9052 | 19116.823 | -13266.082 | -0.7571547 | 4.5461E-59 | 1.1405E-56 |
| ENSMUSG00000003411 | Rab3b | 2485.72609 | 1448.18655 | -1037.5395 | -0.7574416 | 2.5987E-10 | 2.4323E-09 |
| ENSMUSG00000021211 | Akr1c12 | 104.056597 | 58.4705586 | -45.586039 | -0.7584063 | 0.00026737 | 0.00101501 |
| ENSMUSG00000026784 | Pdss1 | 1429.0704 | 836.730099 | -592.3403 | -0.7589695 | 1.5765E-15 | 2.6541E-14 |
| ENSMUSG00000047104 | Pbp2 | 89.7497117 | 45.1408406 | -44.608871 | -0.7598106 | 0.01335085 | 0.03416516 |
| ENSMUSG00000090021 | - | 201.946271 | 103.423688 | -98.522583 | -0.7604361 | 0.01142804 | 0.0298423 |
| ENSMUSG00000025507 | Pidd1 | 574.824541 | 334.924931 | -239.89961 | -0.7605598 | 1.9345E-11 | 2.0716E-10 |
| ENSMUSG00000023004 | Tuba1b | 20619.0597 | 12043.1411 | -8575.9186 | -0.760827 | 2.4949E-14 | 3.6844E-13 |
| ENSMUSG00000032057 | Hoatz | 40.4231883 | 22.1816384 | -18.24155 | -0.7611495 | 0.00137441 | 0.00451091 |
| ENSMUSG00000070305 | Mpzl3 | 64.5931691 | 33.927174 | -30.665995 | -0.7617103 | 0.0073743 | 0.02027316 |
| ENSMUSG00000027242 | Wdr76 | 1917.52782 | 1115.89622 | -801.63161 | -0.7618808 | 9.974E-12 | 1.1032E-10 |
| ENSMUSG00000031584 | Gsr | 8045.86256 | 4720.2458 | -3325.6168 | -0.7619298 | 1.2638E-27 | 5.6033E-26 |
| ENSMUSG00000098091 | - | 101.149331 | 57.2140496 | -43.935281 | -0.7625533 | 7.5789E-05 | 0.00031813 |
| ENSMUSG00000098208 | - | 67.2370441 | 36.5586312 | -30.678413 | -0.7628853 | 0.00243938 | 0.00757135 |
| ENSMUSG00000025091 | Pnliprp2 | 2.39740822 | 0 | -2.3974082 | -0.7632041 | 0.00535557 | 0.01529228 |
| ENSMUSG00000024912 | Fosl1 | 6310.33449 | 3635.36546 | -2674.969 | -0.7633465 | 1.0793E-07 | 7.0421E-07 |
| ENSMUSG00000045294 | Insig1 | 2318.20624 | 1252.69074 | -1065.5155 | -0.76399 | 0.00256424 | 0.0079147 |
| ENSMUSG00000050234 | Gja4 | 47.6121184 | 25.7687232 | -21.843395 | -0.7644152 | 0.00244709 | 0.00758727 |
| ENSMUSG00000061482 | H4c4 | 24.3222234 | 12.2299988 | -12.092225 | -0.7646363 | 0.0116337 | 0.0303186 |
| ENSMUSG00000015451 | C4a | 73.4006655 | 40.6042859 | -32.79638 | -0.7649544 | 0.00073848 | 0.00256267 |
| ENSMUSG00000034659 | Tmem109 | 13604.0265 | 7948.04913 | -5655.9774 | -0.7654249 | 7.1615E-21 | 1.9488E-19 |
| ENSMUSG00000042489 | Clspn | 4607.88779 | 2677.40305 | -1930.4847 | -0.7656046 | 1.3483E-12 | 1.6383E-11 |
| ENSMUSG00000023505 | Cdca3 | 9903.88242 | 5786.7808 | -4117.1016 | -0.7656732 | 1.15E-21 | 3.3319E-20 |
| ENSMUSG00000027079 | Clp1 | 1714.51196 | 1001.29148 | -713.22048 | -0.7657258 | 1.2129E-21 | 3.5046E-20 |
| ENSMUSG00000038489 | Polr2l | 1601.32814 | 923.49705 | -677.83109 | -0.7658037 | 1.0236E-08 | 7.7672E-08 |
| ENSMUSG00000083019 | - | 18.9370343 | 9.18299028 | -9.7540441 | -0.765863 | 0.0180561 | 0.04451169 |
| ENSMUSG00000063171 | Rps4l | 6261.31352 | 3645.9954 | -2615.3181 | -0.7659701 | 2.6019E-15 | 4.2921E-14 |
| ENSMUSG00000017716 | Birc5 | 9928.22493 | 5795.16128 | -4133.0636 | -0.7664598 | 1.2169E-20 | 3.2344E-19 |
| ENSMUSG00000060288 | Ppih | 1663.18225 | 951.980354 | -711.2019 | -0.7672538 | 5.78E-07 | 3.4041E-06 |
| ENSMUSG00000032601 | Prkar2a | 7856.20146 | 4544.08133 | -3312.1201 | -0.7676238 | 1.811E-10 | 1.7272E-09 |
| ENSMUSG00000027176 | Cstf3 | 4605.97513 | 2693.57118 | -1912.4039 | -0.7678411 | 2.4777E-34 | 1.6878E-32 |
| ENSMUSG00000034187 | Nsf | 7554.78429 | 4424.72991 | -3130.0544 | -0.7679836 | 5.1278E-50 | 8.7181E-48 |
| ENSMUSG00000039114 | Nrn1 | 47471.6362 | 27795.8055 | -19675.831 | -0.7681105 | 1.6677E-48 | 2.4682E-46 |
| ENSMUSG00000026841 | Fibcd1 | 3.5758205 | 0.48461008 | -3.0912104 | -0.7681296 | 0.0172426 | 0.04271098 |
| ENSMUSG00000065208 | Gm24616 | 56.720583 | 30.2711406 | -26.449442 | -0.7681434 | 0.00327062 | 0.00984536 |
| ENSMUSG00000081473 | - | 2.73482549 | 0.3239963 | -2.4108292 | -0.7682106 | 0.0155487 | 0.03904637 |
| ENSMUSG00000025228 | Actr1a | 19953.2648 | 11656.0221 | -8297.2427 | -0.7684218 | 1.4298E-28 | 6.8722E-27 |
| ENSMUSG00000090206 | Tepp | 76.284439 | 42.7840154 | -33.500424 | -0.7687338 | 7.5257E-05 | 0.00031634 |
| ENSMUSG00000079478 | Znrd2 | 1699.96082 | 982.206454 | -717.75436 | -0.7689989 | 1.6165E-10 | 1.5525E-09 |
| ENSMUSG00000081865 | - | 84.9417168 | 47.0580846 | -37.883632 | -0.7691479 | 0.00026175 | 0.00099625 |
| ENSMUSG00000030342 | Cd9 | 91511.0467 | 53476.7513 | -38034.295 | -0.7692779 | 5.1174E-35 | 3.7199E-33 |
| ENSMUSG00000097971 | - | 3591.62025 | 2094.69855 | -1496.9217 | -0.7700478 | 3.4128E-25 | 1.2906E-23 |
| ENSMUSG00000054293 | P2ry10b | 4746.87525 | 2733.53567 | -2013.3396 | -0.7700597 | 3.0712E-09 | 2.4894E-08 |
| ENSMUSG00000078566 | Bnip3 | 16374.1416 | 8793.71511 | -7580.4265 | -0.7704541 | 0.00245549 | 0.00760873 |
| ENSMUSG00000004849 | Ap1s1 | 16323.8234 | 9515.94894 | -6807.8744 | -0.7705984 | 3.5172E-26 | 1.4022E-24 |
| ENSMUSG00000048076 | Arf1 | 42075.013 | 24579.8303 | -17495.183 | -0.7707435 | 2.6882E-42 | 2.9574E-40 |
| ENSMUSG00000038482 | Tfdp1 | 11715.8836 | 6789.89642 | -4925.9872 | -0.7717953 | 1.4552E-14 | 2.2011E-13 |
| ENSMUSG00000037894 | H2az1 | 43752.798 | 25163.2677 | -18589.53 | -0.7719034 | 2.6375E-09 | 2.154E-08 |
| ENSMUSG00000074457 | S100a16 | 7589.59241 | 4396.11313 | -3193.4793 | -0.7724514 | 1.1218E-14 | 1.7184E-13 |
| ENSMUSG00000063885 | - | 34.9190117 | 18.4631226 | -16.455889 | -0.7727458 | 0.00423907 | 0.01243316 |
| ENSMUSG00000020706 | Ftsj3 | 16824.8461 | 9806.44684 | -7018.3993 | -0.7729168 | 1.6095E-34 | 1.1262E-32 |
| ENSMUSG00000037541 | Shank2 | 318.22762 | 151.516866 | -166.71075 | -0.7732765 | 0.01842134 | 0.04528727 |
| ENSMUSG00000025499 | Hras | 5087.70564 | 2942.83742 | -2144.8682 | -0.7733011 | 8.4489E-14 | 1.172E-12 |
| ENSMUSG00000058400 | Qrfpr | 23.1792984 | 11.9206435 | -11.258655 | -0.773976 | 0.00731237 | 0.0201271 |
| ENSMUSG00000021453 | Gadd45g | 5533.09237 | 3204.68596 | -2328.4064 | -0.7757824 | 2.1518E-18 | 4.7396E-17 |
| ENSMUSG00000024409 | Psors1c2 | 2.92712058 | 0.32899174 | -2.5981288 | -0.7769992 | 0.01442544 | 0.03657836 |
| ENSMUSG00000064602 | - | 59.0563693 | 31.6856004 | -27.370769 | -0.7771365 | 0.00202299 | 0.00639375 |
| ENSMUSG00000060128 | - | 19.3089772 | 9.75992882 | -9.5490484 | -0.7774722 | 0.01008919 | 0.0267192 |
| ENSMUSG00000094852 | - | 15.3999958 | 7.20326947 | -8.1967263 | -0.7784466 | 0.0192475 | 0.04698144 |
| ENSMUSG00000074280 | - | 4367.68346 | 2518.23493 | -1849.4485 | -0.7789427 | 6.0162E-15 | 9.4622E-14 |
| ENSMUSG00000081188 | - | 19.6869782 | 9.57156345 | -10.115415 | -0.7804977 | 0.01503 | 0.03793829 |
| ENSMUSG00000024045 | Akap8 | 7850.5717 | 4552.26828 | -3298.3034 | -0.7808176 | 1.5746E-38 | 1.4023E-36 |
| ENSMUSG00000087249 | - | 176.486126 | 99.4494745 | -77.036651 | -0.7810998 | 1.737E-06 | 9.4935E-06 |
| ENSMUSG00000098030 | - | 3.12451203 | 0.48392287 | -2.6405892 | -0.781267 | 0.01809842 | 0.04457328 |
| ENSMUSG00000050896 | Rtn4rl2 | 3.12451203 | 0.48392287 | -2.6405892 | -0.781267 | 0.01809842 | 0.04457328 |
| ENSMUSG00000028820 | Sfpq | 23956.2131 | 13639.5412 | -10316.672 | -0.7813002 | 2.5249E-08 | 1.8092E-07 |
| ENSMUSG00000065820 | - | 2.92712058 | 0.29759609 | -2.6295245 | -0.7813836 | 0.01367739 | 0.0348793 |
| ENSMUSG00000077394 | Gm24339 | 22.6444897 | 10.3329462 | -12.311543 | -0.782046 | 0.02056101 | 0.04973907 |
| ENSMUSG00000015749 | Anp32e | 22390.101 | 12946.3681 | -9443.7329 | -0.7821317 | 1.8548E-26 | 7.6008E-25 |
| ENSMUSG00000031819 | Emc8 | 5437.85738 | 3142.526 | -2295.3314 | -0.7831377 | 5.9069E-27 | 2.5159E-25 |
| ENSMUSG00000072789 | Gm10420 | 85.9702194 | 47.4725014 | -38.497718 | -0.7834164 | 7.5192E-05 | 0.0003162 |
| ENSMUSG00000078134 | - | 95.8972695 | 53.3795138 | -42.517756 | -0.7838385 | 4.5045E-05 | 0.00019679 |
| ENSMUSG00000065649 | - | 31.9682111 | 15.0435131 | -16.924698 | -0.7839774 | 0.01606803 | 0.04017886 |
| ENSMUSG00000029580 | Actb | 150851.38 | 87129.9085 | -63721.472 | -0.7845546 | 1.9835E-29 | 1.0001E-27 |
| ENSMUSG00000027811 | 4930579G24 | 4894.60228 | 2812.9371 | -2081.6652 | -0.785012 | 2.3451E-16 | 4.2694E-15 |

| | | | | | | | |
|---|---|---|---|---|---|---|---|
| ENSMUSG00000024556 | Me2 | 9140.07457 | 5271.77734 | -3868.2972 | -0.785082 | 7.1268E-25 | 2.6088E-23 |
| ENSMUSG00000070490 | - | 115.583939 | 63.839534 | -51.744405 | -0.7852976 | 0.00010441 | 0.00042862 |
| ENSMUSG00000037010 | Apln | 2743.7706 | 1509.50696 | -1234.2636 | -0.7854535 | 0.00015506 | 0.00061571 |
| ENSMUSG00000019773 | Fbxo5 | 4201.39583 | 2423.7065 | -1777.6893 | -0.7857177 | 4.5429E-28 | 2.0815E-26 |
| ENSMUSG00000024516 | Sec11c | 7527.39157 | 4323.91371 | -3203.4779 | -0.7859673 | 7.7739E-17 | 1.4891E-15 |
| ENSMUSG00000036667 | Tcaf1 | 16373.4614 | 9383.04698 | -6990.4144 | -0.7859772 | 9.9695E-14 | 1.3765E-12 |
| ENSMUSG00000098192 | - | 103.165666 | 56.1376672 | -47.027999 | -0.7863145 | 0.00037898 | 0.00139421 |
| ENSMUSG00000029715 | Pop7 | 1382.72047 | 794.585903 | -588.13457 | -0.7866512 | 1.3474E-19 | 3.288E-18 |
| ENSMUSG00000084893 | - | 3.60705149 | 0.45913278 | -3.1479187 | -0.786674 | 0.01447405 | 0.0366883 |
| ENSMUSG00000097070 | - | 18.4857259 | 8.74930231 | -9.7364236 | -0.7880842 | 0.01494746 | 0.03773921 |
| ENSMUSG00000002105 | Slc39a13 | 8452.13962 | 4867.46245 | -3584.6772 | -0.7881183 | 4.6249E-28 | 2.1143E-26 |
| ENSMUSG00000083287 | - | 22.1036229 | 10.4120534 | -11.69157 | -0.7884748 | 0.0150386 | 0.03795534 |
| ENSMUSG00000031167 | Rbm3-ps; Rbı | 22236.8087 | 12720.1868 | -9516.6219 | -0.7898045 | 7.352E-15 | 1.1467E-13 |
| ENSMUSG00000073676 | Hspe1 | 13633.7361 | 7811.5969 | -5822.1392 | -0.7904053 | 8.8855E-18 | 1.8501E-16 |
| ENSMUSG00000064032 | - | 48.8049797 | 23.0238459 | -25.781134 | -0.7905648 | 0.01485444 | 0.03754122 |
| ENSMUSG00000052488 | Cherp | 8191.75449 | 4693.96015 | -3497.7943 | -0.7910184 | 1.0948E-18 | 2.4968E-17 |
| ENSMUSG00000097994 | - | 173.846795 | 97.7438692 | -76.102926 | -0.7912551 | 4.1597E-07 | 2.5036E-06 |
| ENSMUSG00000022602 | Arc | 53.8163025 | 26.5120396 | -27.304263 | -0.7915844 | 0.00928231 | 0.024822 |
| ENSMUSG00000084817 | - | 236.946888 | 133.742926 | -103.20396 | -0.791628 | 2.0497E-08 | 1.4884E-07 |
| ENSMUSG00000078452 | Raet1c; Raet | 177.237871 | 97.0899917 | -80.14788 | -0.7921017 | 0.00015085 | 0.00060038 |
| ENSMUSG00000038210 | Hoxa11 | 6816.09383 | 3905.81076 | -2910.2831 | -0.7922916 | 8.4374E-21 | 2.2839E-19 |
| ENSMUSG00000034825 | Nrip3 | 27.9675256 | 13.3289963 | -14.638529 | -0.7922995 | 0.01402338 | 0.03565996 |
| ENSMUSG00000063253 | Scoc | 7846.28107 | 4492.3528 | -3353.9283 | -0.792687 | 9.5388E-20 | 2.3473E-18 |
| ENSMUSG00000091635 | Vmn2r13 | 26.1358482 | 12.8169386 | -13.31891 | -0.7931019 | 0.00938295 | 0.02505205 |
| ENSMUSG00000026234 | Ncl | 82089.439 | 47139.009 | -34950.43 | -0.7943649 | 2.6948E-37 | 2.2087E-35 |
| ENSMUSG00000074140 | - | 260.448401 | 143.79425 | -116.65415 | -0.7978074 | 1.913E-05 | 8.8937E-05 |
| ENSMUSG00000083465 | - | 51.4129578 | 26.0611737 | -25.351784 | -0.7986584 | 0.0044992 | 0.01311164 |
| ENSMUSG00000005374 | Tbl2 | 5658.91374 | 3225.56252 | -2433.3512 | -0.7995007 | 1.4043E-20 | 3.6989E-19 |
| ENSMUSG00000018411 | Mapt | 97.0876763 | 52.0581617 | -45.029515 | -0.7998417 | 0.00050451 | 0.00181226 |
| ENSMUSG00000046223 | Plaur | 3614.12358 | 2025.54218 | -1588.5814 | -0.7998742 | 4.928E-08 | 3.3816E-07 |
| ENSMUSG00000081034 | - | 25.5571612 | 12.5978089 | -12.959352 | -0.8001037 | 0.00759795 | 0.02081569 |
| ENSMUSG00000002343 | Armc6 | 1887.23446 | 1075.7062 | -811.52825 | -0.8009598 | 2.1236E-24 | 7.5192E-23 |
| ENSMUSG00000080974 | Gm9009 | 35.1201283 | 16.6740798 | -18.446048 | -0.8031657 | 0.01222522 | 0.03166265 |
| ENSMUSG00000029326 | Enoph1 | 2661.90455 | 1510.86958 | -1151.035 | -0.8040555 | 4.2513E-19 | 1.0018E-17 |
| ENSMUSG00000009013 | Dynll1 | 16862.0014 | 9543.30402 | -7318.6974 | -0.8043953 | 5.6102E-15 | 8.8496E-14 |
| ENSMUSG00000046056 | Sbsn | 2419.54711 | 1347.21377 | -1072.3333 | -0.8044174 | 2.1849E-07 | 1.3695E-06 |
| ENSMUSG00000031657 | Heatr3 | 11627.3418 | 6597.53453 | -5029.8073 | -0.8045915 | 9.0442E-19 | 2.0788E-17 |
| ENSMUSG00000027329 | Spef1 | 518.066152 | 290.835547 | -227.23061 | -0.8056083 | 6.991E-11 | 6.9815E-10 |
| ENSMUSG00000024981 | Acsl5 | 16047.1618 | 9148.55232 | -6898.6094 | -0.8060514 | 2.287E-48 | 3.3367E-46 |
| ENSMUSG00000020737 | Jpt1 | 15701.7198 | 8887.79396 | -6813.9259 | -0.8086471 | 5.6253E-20 | 1.413E-18 |
| ENSMUSG00000074797 | Itpa | 5326.44206 | 3008.21787 | -2318.2242 | -0.8090835 | 1.2634E-16 | 2.3671E-15 |
| ENSMUSG00000062825 | Actg1 | 69037.3947 | 39285.3933 | -29752.001 | -0.8091102 | 7.6274E-54 | 1.4665E-51 |
| ENSMUSG00000020091 | Eif4ebp2 | 3048.37578 | 1730.5013 | -1317.8745 | -0.8091478 | 7.7008E-30 | 3.9904E-28 |
| ENSMUSG00000057696 | - | 56.2381652 | 29.2173929 | -27.020772 | -0.8098621 | 0.00134516 | 0.00442196 |
| ENSMUSG00000079297 | - | 239.384105 | 133.852554 | -105.53155 | -0.8100767 | 1.2957E-09 | 1.0987E-08 |
| ENSMUSG00000091387 | Gcnt4 | 5465.98594 | 3057.78191 | -2408.204 | -0.8111808 | 2.715E-10 | 2.533E-09 |
| ENSMUSG00000008475 | Arpc5 | 11917.3316 | 6751.91521 | -5165.4164 | -0.8125134 | 7.3757E-34 | 4.9214E-32 |
| ENSMUSG00000058258 | Idi1 | 1442.83605 | 791.269837 | -651.56622 | -0.8131662 | 3.4648E-06 | 1.8114E-05 |
| ENSMUSG00000021041 | - | 75.8574634 | 40.3952811 | -35.462182 | -0.8132397 | 0.00022208 | 0.00085909 |
| ENSMUSG00000097903 | - | 5.75150438 | 1.12401391 | -4.6274905 | -0.8135476 | 0.01749768 | 0.0432595 |
| ENSMUSG00000057788 | Ddx49 | 4079.76343 | 2309.41473 | -1770.3487 | -0.8141499 | 2.2696E-36 | 1.7619E-34 |
| ENSMUSG00000095545 | Gm4724 | 4.29123871 | 0.972269 | -3.3189697 | -0.8143257 | 0.02020238 | 0.04898672 |
| ENSMUSG00000065304 | Gm23245 | 28.9532123 | 13.2502061 | -15.703006 | -0.814739 | 0.01251872 | 0.0322766 |
| ENSMUSG00000022454 | Nell2 | 26.916183 | 11.9592988 | -14.956884 | -0.8149389 | 0.01508115 | 0.0380394 |
| ENSMUSG00000003604 | Aven | 1961.07433 | 1095.73979 | -865.33454 | -0.8152519 | 2.6599E-11 | 2.8004E-10 |
| ENSMUSG00000078784 | Rbis | 4492.39482 | 2526.5661 | -1965.8287 | -0.8157052 | 8.7895E-18 | 1.8339E-16 |
| ENSMUSG00000022947 | Cbr3 | 353.987267 | 190.20535 | -163.78192 | -0.8172925 | 7.3206E-05 | 0.00030867 |
| ENSMUSG00000055676 | - | 55.887139 | 29.1274118 | -26.759727 | -0.8175093 | 0.00077919 | 0.00269267 |
| ENSMUSG00000064984 | Snord73a | 30.0019131 | 15.1826 | -14.819313 | -0.8177938 | 0.00260839 | 0.0080353 |
| ENSMUSG00000030120 | Mlf2 | 29212.8907 | 16521.688 | -12691.203 | -0.8177983 | 5.5399E-54 | 1.0958E-51 |
| ENSMUSG00000027746 | Ufm1 | 7690.27571 | 4339.57239 | -3350.7033 | -0.8180499 | 2.2169E-33 | 1.4252E-31 |
| ENSMUSG00000018362 | Kpna2 | 23125.2176 | 13011.4283 | -10113.789 | -0.8180958 | 9.321E-22 | 2.7199E-20 |
| ENSMUSG00000082208 | - | 26.3873107 | 12.3375968 | -14.049714 | -0.8181399 | 0.01027134 | 0.0271422 |
| ENSMUSG00000026229 | Psmd1 | 23076.8986 | 13045.1531 | -10031.745 | -0.8185443 | 6.6177E-54 | 1.2843E-51 |
| ENSMUSG00000032815 | Fanca | 3447.49233 | 1940.07155 | -1507.4208 | -0.8191094 | 1.8736E-23 | 6.2168E-22 |
| ENSMUSG00000028776 | Tinagl1 | 7856.80654 | 4408.94476 | -3447.8618 | -0.8198084 | 6.792E-19 | 1.57E-17 |
| ENSMUSG00000081583 | Gm14769 | 33.6636884 | 16.7929404 | -16.870748 | -0.8203396 | 0.00273341 | 0.00837154 |
| ENSMUSG00000069083 | - | 4.43965551 | 0.96410564 | -3.4755499 | -0.820585 | 0.01912615 | 0.04671295 |
| ENSMUSG00000069272 | H2ac8 | 70.2292029 | 37.1247681 | -33.104435 | -0.8206582 | 0.00019034 | 0.00074386 |
| ENSMUSG00000032889 | - | 100.160967 | 48.395871 | -51.765096 | -0.8209169 | 0.00702542 | 0.0194309 |
| ENSMUSG00000030607 | Acan | 2917.63662 | 1550.06855 | -1367.5681 | -0.8225292 | 0.00015871 | 0.00062873 |
| ENSMUSG00000024925 | Rnaseh2c | 2968.49138 | 1663.17183 | -1305.3196 | -0.8227739 | 9.2888E-21 | 2.5044E-19 |
| ENSMUSG00000080021 | - | 64.3545763 | 30.2087279 | -34.145848 | -0.8229006 | 0.00856541 | 0.02315412 |
| ENSMUSG00000005470 | Asf1b | 6183.88411 | 3483.21081 | -2700.6733 | -0.8229417 | 2.5046E-49 | 4.0892E-47 |
| ENSMUSG00000038412 | Higd1a | 7040.34765 | 3901.47493 | -3138.8727 | -0.8240061 | 1.5093E-10 | 1.4549E-09 |

| | | | | | | | |
|---|---|---|---|---|---|---|---|
| ENSMUSG00000061390 | - | 40.2402459 | 18.6822442 | -21.558002 | -0.8240188 | 0.01016842 | 0.02690479 |
| ENSMUSG00000018821 | Avpi1 | 12993.5687 | 7207.8643 | -5785.7044 | -0.824302 | 4.2614E-11 | 4.368E-10 |
| ENSMUSG00000072974 | Gm4787 | 23.3454589 | 10.9038278 | -12.441631 | -0.8249055 | 0.00929684 | 0.02484471 |
| ENSMUSG00000040204 | Pclaf | 10062.1965 | 5631.57381 | -4430.6227 | -0.8249696 | 2.5151E-21 | 7.1073E-20 |
| ENSMUSG00000092517 | - | 4.12507825 | 0.49145135 | -3.6336269 | -0.8255459 | 0.01029983 | 0.02721398 |
| ENSMUSG00000025153 | Fasn | 17302.333 | 9603.56062 | -7698.7724 | -0.8256136 | 6.2053E-12 | 7.0179E-11 |
| ENSMUSG00000062687 | Cks1brt | 4.68092525 | 1.18429047 | -3.4966348 | -0.8258187 | 0.02024216 | 0.04906006 |
| ENSMUSG00000096952 | - | 63.3942497 | 33.0220262 | -30.372224 | -0.825841 | 0.00053352 | 0.00190648 |
| ENSMUSG00000024691 | Fam111a | 31180.3216 | 17492.8356 | -13687.486 | -0.8260104 | 4.8201E-32 | 2.8909E-30 |
| ENSMUSG00000057160 | - | 115.810107 | 63.2894055 | -52.520701 | -0.8264168 | 4.1124E-07 | 2.4773E-06 |
| ENSMUSG00000026675 | Hsd17b7 | 2482.41923 | 1385.3604 | -1097.0588 | -0.8265219 | 5.051E-18 | 1.0745E-16 |
| ENSMUSG00000082362 | - | 16.6147354 | 7.37608372 | -9.2386517 | -0.8268457 | 0.01347713 | 0.03444547 |
| ENSMUSG00000021943 | Gdf10 | 4.33841161 | 0.81670704 | -3.5217046 | -0.826962 | 0.01651661 | 0.04113057 |
| ENSMUSG00000020089 | Ppa1 | 15256.5297 | 8532.34168 | -6724.188 | -0.8272808 | 2.3733E-23 | 7.8117E-22 |
| ENSMUSG00000086522 | - | 44.9697153 | 21.5658982 | -23.403817 | -0.8281122 | 0.00554692 | 0.01577866 |
| ENSMUSG00000032202 | Rab27a | 178.657366 | 95.893325 | -82.764041 | -0.828136 | 1.4215E-05 | 6.759E-05 |
| ENSMUSG00000026520 | Pycr2 | 3461.06389 | 1932.20507 | -1528.8588 | -0.8288327 | 1.3683E-21 | 3.9204E-20 |
| ENSMUSG00000061613 | U2af1 | 9029.83682 | 5047.10897 | -3982.7278 | -0.8291085 | 2.9018E-26 | 1.1682E-24 |
| ENSMUSG00000075232 | Amd1 | 8118.72545 | 4544.3767 | -3574.3488 | -0.8293432 | 1.473E-32 | 8.965E-31 |
| ENSMUSG00000078812 | Eif5a | 90907.546 | 50822.4098 | -40085.136 | -0.8303963 | 5.4794E-30 | 2.8682E-28 |
| ENSMUSG00000067106 | - | 50.038647 | 24.1754057 | -25.863241 | -0.8305681 | 0.00455285 | 0.01324956 |
| ENSMUSG00000098412 | - | 2.92712058 | 0.17533831 | -2.7517823 | -0.8312901 | 0.00655707 | 0.01830401 |
| ENSMUSG00000037185 | Krt80 | 433.548425 | 193.984309 | -239.56412 | -0.8313991 | 0.01177743 | 0.03061863 |
| ENSMUSG00000059195 | - | 1164.28981 | 648.405558 | -515.88425 | -0.8316289 | 7.128E-21 | 1.9422E-19 |
| ENSMUSG00000047347 | Tdg-ps2; Tdg | 11.6090402 | 4.82296843 | -6.7860717 | -0.8321847 | 0.01588753 | 0.0397807 |
| ENSMUSG00000078453 | Abracl | 7417.62989 | 4128.27479 | -3289.3551 | -0.832987 | 6.4156E-22 | 1.9018E-20 |
| ENSMUSG00000081308 | - | 44.5540158 | 22.477219 | -22.076797 | -0.8330251 | 0.00126233 | 0.00417972 |
| ENSMUSG00000084413 | - | 51.897299 | 26.7835006 | -25.113798 | -0.833246 | 0.00063057 | 0.00221669 |
| ENSMUSG00000040751 | Lat2 | 76.5153944 | 40.2097165 | -36.305678 | -0.8332737 | 0.00014201 | 0.00056861 |
| ENSMUSG00000083262 | - | 7.2528849 | 1.73527506 | -5.5176098 | -0.8334154 | 0.01904614 | 0.04655623 |
| ENSMUSG00000090114 | - | 3.05449912 | 0.16153669 | -2.8929624 | -0.8341521 | 0.00590285 | 0.01667355 |
| ENSMUSG00000036114 | Rpp25l | 1347.64487 | 743.147354 | -604.49752 | -0.8353563 | 1.6678E-12 | 2.0088E-11 |
| ENSMUSG00000029649 | Pomp | 11081.3176 | 6170.75584 | -4910.5617 | -0.8356195 | 1.1792E-29 | 6.0045E-28 |
| ENSMUSG00000097819 | - | 125.060829 | 64.1893309 | -60.871499 | -0.8356373 | 0.0008109 | 0.00279287 |
| ENSMUSG00000089365 | - | 65.2953223 | 33.7324808 | -31.562841 | -0.8360432 | 0.00027419 | 0.00103878 |
| ENSMUSG00000014226 | Cacybp | 17894.4229 | 9945.47748 | -7948.9454 | -0.8361041 | 1.0487E-23 | 3.5193E-22 |
| ENSMUSG00000024989 | Cep55 | 8551.84469 | 4766.97539 | -3784.8693 | -0.8364022 | 4.2075E-39 | 3.9523E-37 |
| ENSMUSG00000019235 | Rps6kl1 | 36.8227261 | 17.6252818 | -19.197444 | -0.8367864 | 0.00463836 | 0.01346635 |
| ENSMUSG00000097388 | - | 163.965956 | 88.6570402 | -75.308916 | -0.8371212 | 9.731E-07 | 5.527E-06 |
| ENSMUSG00000029910 | Mad2l1 | 13526.4834 | 7541.46577 | -5985.0177 | -0.837442 | 5.4376E-48 | 7.7682E-46 |
| ENSMUSG00000001416 | Cct3 | 48670.5279 | 27093.2222 | -21577.306 | -0.838374 | 2.0451E-38 | 1.7902E-36 |
| ENSMUSG00000028044 | Cks1b | 14331.7606 | 7944.32041 | -6387.4402 | -0.8385447 | 1.3027E-21 | 3.748E-20 |
| ENSMUSG00000097965 | - | 3.61969878 | 0.48299996 | -3.1366988 | -0.8402336 | 0.01140254 | 0.0297863 |
| ENSMUSG00000081170 | - | 27.4281517 | 12.3508496 | -15.077302 | -0.8405979 | 0.00891757 | 0.02396503 |
| ENSMUSG00000065041 | Gm24044 | 93.7620477 | 49.7863571 | -43.975691 | -0.8415665 | 2.3392E-05 | 0.00010717 |
| ENSMUSG00000051220 | Ercc6l | 6734.22898 | 3731.0014 | -3003.2276 | -0.8415757 | 3.0257E-26 | 1.2133E-24 |
| ENSMUSG00000081251 | - | 9.2167283 | 3.46780477 | -5.7489235 | -0.8418403 | 0.01912255 | 0.0467097 |
| ENSMUSG00000088712 | Gm22728 | 6.53417208 | 1.860634 | -4.6735381 | -0.8420931 | 0.02021849 | 0.04902002 |
| ENSMUSG00000047986 | Palm3 | 4.17225115 | 0.622497 | -3.5497542 | -0.8422 | 0.01227054 | 0.03174417 |
| ENSMUSG00000033520 | Idi2 | 4.11062922 | 0.80481395 | -3.3058153 | -0.8424606 | 0.01538294 | 0.0386726 |
| ENSMUSG00000049287 | Iba57 | 558.988561 | 302.613919 | -256.37464 | -0.8432689 | 6.3477E-08 | 4.2829E-07 |
| ENSMUSG00000056962 | Jmjd6 | 6942.53353 | 3839.37951 | -3103.154 | -0.8435821 | 3.5667E-25 | 1.3439E-23 |
| ENSMUSG00000018932 | Map2k3 | 9681.5923 | 5357.24268 | -4324.3496 | -0.8436275 | 4.7984E-27 | 2.0522E-25 |
| ENSMUSG00000031227 | Magee1 | 3474.81411 | 1927.21211 | -1547.602 | -0.8441468 | 3.2533E-46 | 4.2092E-44 |
| ENSMUSG00000020358 | Hnrnpab | 44668.7225 | 24753.4334 | -19915.289 | -0.8442158 | 8.878E-36 | 6.7146E-34 |
| ENSMUSG00000096243 | - | 12.0375086 | 3.70480445 | -8.3327042 | -0.8442649 | 0.02019363 | 0.04897705 |
| ENSMUSG00000081564 | - | 22.7794191 | 8.44695075 | -14.332468 | -0.8454911 | 0.01845278 | 0.04535378 |
| ENSMUSG00000078184 | - | 122.939991 | 63.8518103 | -59.088181 | -0.8456254 | 0.00016673 | 0.00065871 |
| ENSMUSG00000072919 | Noxred1 | 24.9970579 | 11.0101332 | -13.986925 | -0.8458998 | 0.01046781 | 0.02760818 |
| ENSMUSG00000002477 | Snrpd1 | 13563.3372 | 7511.99669 | -6051.3406 | -0.845955 | 3.8915E-42 | 4.2583E-40 |
| ENSMUSG00000030263 | Irag2 | 14.4484042 | 5.93871431 | -8.5096899 | -0.8463079 | 0.01408198 | 0.0358001 |
| ENSMUSG00000081126 | - | 45.6155511 | 22.934201 | -22.68135 | -0.846411 | 0.00119507 | 0.00397622 |
| ENSMUSG00000040434 | Large2 | 66.0628881 | 34.4500437 | -31.612844 | -0.8468047 | 0.00010829 | 0.00044317 |
| ENSMUSG00000098113 | - | 48.4950683 | 24.515966 | -23.979102 | -0.8469919 | 0.00104198 | 0.00351059 |
| ENSMUSG00000045983 | Eif4g1 | 69829.2253 | 38570.7845 | -31258.441 | -0.8471024 | 1.7156E-29 | 8.6929E-28 |
| ENSMUSG00000042699 | Dhx9 | 23061.0424 | 12132.7345 | -10928.308 | -0.8472467 | 3.3494E-05 | 0.00014959 |
| ENSMUSG00000028441 | 1110017D15 | 8.48962449 | 3.07110729 | -5.4185172 | -0.8476035 | 0.01877076 | 0.04600883 |
| ENSMUSG00000024902 | Mrpl11 | 4133.88451 | 2279.01264 | -1854.8719 | -0.8480524 | 7.5263E-26 | 2.9661E-24 |
| ENSMUSG00000027654 | Fam83d | 2773.80504 | 1528.35998 | -1245.4451 | -0.8484766 | 6.6085E-25 | 2.4277E-23 |
| ENSMUSG00000077493 | - | 23.4559338 | 9.6831739 | -13.77276 | -0.8486551 | 0.01437144 | 0.03646389 |
| ENSMUSG00000028587 | Orc1 | 1383.41225 | 754.584206 | -628.82804 | -0.8488529 | 3.5512E-12 | 4.1111E-11 |
| ENSMUSG00000021048 | Mthfd1 | 8822.36463 | 4886.62068 | -3935.7439 | -0.8490537 | 1.0769E-78 | 5.9878E-76 |
| ENSMUSG00000077565 | Gm23201 | 52.715454 | 26.1036538 | -26.6118 | -0.8492923 | 0.00153118 | 0.00496994 |
| ENSMUSG00000078776 | 9530053A07 | 14.7107122 | 5.00395351 | -9.7067587 | -0.8506617 | 0.01906215 | 0.04658982 |
| ENSMUSG00000069939 | - | 24.4944419 | 11.8320748 | -12.662367 | -0.8516091 | 0.00319411 | 0.0096401 |

| | | | | | | | |
|---|---|---|---|---|---|---|---|
| ENSMUSG00000002984 | Tomm40 | 5067.96173 | 2788.99956 | -2278.9622 | -0.8517843 | 4.0283E-29 | 1.9778E-27 |
| ENSMUSG00000082136 | - | 3.93937241 | 0.63918643 | -3.300186 | -0.8521797 | 0.01261068 | 0.03247301 |
| ENSMUSG00000059355 | Wdr83os | 2182.67408 | 1195.25288 | -987.42119 | -0.8524799 | 2.587E-18 | 5.6798E-17 |
| ENSMUSG00000024590 | Lmnb1 | 8616.94203 | 4700.01321 | -3916.9288 | -0.8537799 | 1.2309E-14 | 1.8772E-13 |
| ENSMUSG00000091625 | Lsm5 | 4383.06447 | 2359.74737 | -2023.3171 | -0.8539191 | 1.1305E-08 | 8.5093E-08 |
| ENSMUSG00000066944 | - | 17.6284801 | 5.45597531 | -12.172505 | -0.8540839 | 0.01883439 | 0.0461373 |
| ENSMUSG00000021037 | Ahsa1 | 15148.9161 | 8341.47246 | -6807.4437 | -0.8549889 | 3.1771E-47 | 4.3573E-45 |
| ENSMUSG00000081390 | - | 4.45739916 | 0.82298017 | -3.634419 | -0.8553953 | 0.01329439 | 0.03403761 |
| ENSMUSG00000020116 | Pno1 | 7193.00224 | 3947.86934 | -3245.1329 | -0.8558686 | 1.9909E-29 | 1.0014E-27 |
| ENSMUSG00000044244 | Il20rb | 78.1918784 | 39.8410281 | -38.35085 | -0.8565384 | 0.00032064 | 0.001198 |
| ENSMUSG00000082262 | - | 17.0921785 | 7.69221811 | -9.3999604 | -0.8569905 | 0.00739944 | 0.02033685 |
| ENSMUSG00000026895 | Ndufa8 | 6625.38512 | 3268.57147 | -3356.8136 | -0.8571059 | 0.00136712 | 0.00448985 |
| ENSMUSG00000069439 | - | 9.48233093 | 3.28436986 | -6.1979611 | -0.8572309 | 0.01765083 | 0.04359623 |
| ENSMUSG00000019577 | Pdk4 | 409.062683 | 215.740656 | -193.32203 | -0.8580508 | 3.7338E-06 | 1.9426E-05 |
| ENSMUSG00000051223 | Bzw1 | 41776.8081 | 22990.9458 | -18785.862 | -0.8586569 | 1.2153E-90 | 8.9291E-88 |
| ENSMUSG00000015176 | Nolc1 | 19970.0577 | 10955.844 | -9014.2136 | -0.8590386 | 5.5551E-39 | 5.171E-37 |
| ENSMUSG00000078087 | - | 4.96607326 | 0.99485848 | -3.9712148 | -0.8593271 | 0.01365823 | 0.03483476 |
| ENSMUSG00000021427 | Ssr1 | 32151.8705 | 17614.2085 | -14537.662 | -0.8596339 | 2.9321E-33 | 1.8617E-31 |
| ENSMUSG00000059854 | Hydin | 4.8470857 | 1.14168895 | -3.7053967 | -0.8600802 | 0.01596868 | 0.03995472 |
| ENSMUSG00000038064 | Cldn22 | 5.46965099 | 1.20379304 | -4.2658579 | -0.8614068 | 0.01454186 | 0.03684175 |
| ENSMUSG00000090066 | 1110002E22 | 31.8804545 | 12.9362867 | -18.944168 | -0.8616621 | 0.01238503 | 0.03200012 |
| ENSMUSG00000081455 | - | 28.6471476 | 13.5957686 | -15.051379 | -0.8625644 | 0.0030426 | 0.00922374 |
| ENSMUSG00000044026 | Slc35g1 | 1108.39079 | 597.935875 | -510.45492 | -0.8631471 | 5.0493E-12 | 5.7644E-11 |
| ENSMUSG00000088059 | - | 52.9427052 | 25.4527023 | -27.490003 | -0.8639863 | 0.00226662 | 0.00708355 |
| ENSMUSG00000083899 | - | 1086.5654 | 583.625535 | -502.93986 | -0.8640391 | 2.4958E-10 | 2.3434E-09 |
| ENSMUSG00000058927 | - | 116.732082 | 60.9982368 | -55.733845 | -0.865461 | 2.9161E-06 | 1.5433E-05 |
| ENSMUSG00000075307 | Klhl41 | 22.7041882 | 10.1433588 | -12.560829 | -0.8666658 | 0.00679252 | 0.01887554 |
| ENSMUSG00000038843 | Gcnt1 | 3035.66902 | 1632.54829 | -1403.1207 | -0.8687152 | 1.6532E-12 | 1.9935E-11 |
| ENSMUSG00000083093 | - | 10.307384 | 3.4953948 | -6.8119892 | -0.8687867 | 0.01638534 | 0.0408383 |
| ENSMUSG00000024791 | Cdca5 | 3714.97843 | 2012.70025 | -1702.2782 | -0.8690717 | 1.119E-20 | 2.9858E-19 |
| ENSMUSG00000026617 | Bpnt1 | 2288.66676 | 1246.23755 | -1042.4292 | -0.8690945 | 6.4824E-44 | 7.4086E-42 |
| ENSMUSG00000071343 | - | 85.0979934 | 42.759852 | -42.338141 | -0.8696596 | 0.00036452 | 0.00134654 |
| ENSMUSG00000070527 | Mkrn3 | 33.6267083 | 15.6447261 | -17.981982 | -0.8701415 | 0.00382607 | 0.01135129 |
| ENSMUSG00000027239 | Mdk | 12.644132 | 5.33520003 | -7.308932 | -0.8701714 | 0.00982906 | 0.02612447 |
| ENSMUSG00000016319 | Slc25a5 | 49113.4027 | 26486.0652 | -22627.338 | -0.8712602 | 3.5001E-16 | 6.2721E-15 |
| ENSMUSG00000082144 | - | 7.50680192 | 2.0836216 | -5.4231803 | -0.8726549 | 0.01620007 | 0.0404206 |
| ENSMUSG00000081600 | - | 114.588469 | 59.1702944 | -55.418175 | -0.8732171 | 1.4592E-05 | 6.9183E-05 |
| ENSMUSG00000022997 | Wnt1 | 4.22122579 | 0.61278622 | -3.6084396 | -0.8733372 | 0.00968231 | 0.02576105 |
| ENSMUSG00000027715 | Ccna2 | 20768.5157 | 11232.438 | -9536.0778 | -0.8734506 | 5.0802E-23 | 1.633E-21 |
| ENSMUSG00000066245 | - | 51.4857342 | 24.8453083 | -26.640426 | -0.8738987 | 0.00128812 | 0.00425418 |
| ENSMUSG00000030672 | Mylpf | 123.360068 | 63.4527548 | -59.907314 | -0.8739122 | 1.7919E-05 | 8.3646E-05 |
| ENSMUSG00000071042 | Rasgrp3 | 68.1629315 | 26.663835 | -41.499097 | -0.8741044 | 0.01248133 | 0.0322004 |
| ENSMUSG00000043091 | Tuba1c | 12420.5007 | 6711.75562 | -5708.745 | -0.8752372 | 2.9534E-24 | 1.0333E-22 |
| ENSMUSG00000081067 | - | 13.9658647 | 5.83825238 | -8.1276123 | -0.8757298 | 0.00854014 | 0.02309251 |
| ENSMUSG00000082148 | - | 2.78709476 | 0 | -2.7870948 | -0.8758034 | 0.00244976 | 0.00759441 |
| ENSMUSG00000068263 | Efcc1 | 49.9535322 | 22.8693661 | -27.084166 | -0.8759079 | 0.00424507 | 0.01244544 |
| ENSMUSG00000025727 | A930017K11 | 4.32576432 | 0.78312908 | -3.5426352 | -0.8761496 | 0.01167141 | 0.03039509 |
| ENSMUSG00000062038 | - | 14.8515781 | 6.06254635 | -8.7890317 | -0.8764777 | 0.01070901 | 0.02818652 |
| ENSMUSG00000006517 | Mvd | 1866.80914 | 986.744642 | -880.0645 | -0.8768116 | 1.2017E-08 | 9.0227E-08 |
| ENSMUSG00000047547 | Cltb | 6090.26047 | 3239.40014 | -2850.8603 | -0.8790338 | 4.376E-11 | 4.4654E-10 |
| ENSMUSG00000008601 | Rab25 | 12.960511 | 4.90729402 | -8.053217 | -0.8798387 | 0.01288475 | 0.03309177 |
| ENSMUSG00000074622 | Mafb | 1397.17229 | 744.221682 | -652.9506 | -0.8806266 | 3.2452E-12 | 3.7803E-11 |
| ENSMUSG00000082778 | - | 57.1932386 | 28.2392377 | -28.954001 | -0.8817919 | 0.00050267 | 0.00180704 |
| ENSMUSG00000072934 | Trpc5os | 4.68421987 | 0.76643965 | -3.9177802 | -0.8818998 | 0.01070096 | 0.02816892 |
| ENSMUSG00000054999 | Naaladl1 | 79.6531714 | 37.221384 | -42.431787 | -0.8821441 | 0.00254638 | 0.00786431 |
| ENSMUSG00000004951 | Hspb1 | 16687.62 | 8527.62304 | -8159.997 | -0.8822278 | 2.2342E-05 | 0.00010275 |
| ENSMUSG00000080979 | - | 3.72933367 | 0.3239963 | -3.4053374 | -0.8823951 | 0.0058606 | 0.01657468 |
| ENSMUSG00000073408 | Mucl3 | 9.70256239 | 3.14858125 | -6.5539811 | -0.8823977 | 0.01493662 | 0.03771717 |
| ENSMUSG00000030735 | - | 381.157002 | 156.12925 | -225.02775 | -0.8842905 | 0.00928641 | 0.02482651 |
| ENSMUSG00000002341 | Ncan | 4.4480465 | 0.82262539 | -3.6254211 | -0.8853578 | 0.01106362 | 0.02899372 |
| ENSMUSG00000025574 | Tk1 | 8634.31818 | 4624.69669 | -4009.6215 | -0.8867004 | 4.2475E-23 | 1.3782E-21 |
| ENSMUSG00000097569 | - | 4.12837287 | 0.78721991 | -3.341153 | -0.8869509 | 0.01163402 | 0.0303186 |
| ENSMUSG00000021061 | Sptb | 87.7093527 | 42.2978596 | -45.411493 | -0.8889274 | 0.00083596 | 0.00287103 |
| ENSMUSG00000083686 | - | 7.60804583 | 2.45427465 | -5.1537712 | -0.8890289 | 0.0143009 | 0.0363073 |
| ENSMUSG00000095524 | Gm15682 | 6.88423664 | 1.89709573 | -4.9871409 | -0.8902826 | 0.01431117 | 0.03632889 |
| ENSMUSG00000041423 | Paqr6 | 3.35739078 | 0.31611304 | -3.0412777 | -0.8906809 | 0.00613021 | 0.01724946 |
| ENSMUSG00000049916 | 2610318N02 | 109.085973 | 50.339213 | -58.74676 | -0.8911354 | 0.00249724 | 0.00772996 |
| ENSMUSG00000065738 | Gm24494 | 108.742088 | 55.8225541 | -52.919533 | -0.8914395 | 5.5397E-06 | 2.8028E-05 |
| ENSMUSG00000027533 | Fabp5 | 16454.8454 | 8753.98694 | -7700.8585 | -0.8914912 | 1.0572E-17 | 2.1748E-16 |
| ENSMUSG00000056612 | Ppp1r14b | 14698.8454 | 7875.4219 | -6823.4235 | -0.8915382 | 3.8634E-36 | 2.9767E-34 |
| ENSMUSG00000036856 | Wnt4 | 743.154562 | 282.933839 | -460.22072 | -0.8917294 | 0.01116658 | 0.02922632 |
| ENSMUSG00000020869 | Lrrc59 | 33230.2087 | 17823.6126 | -15406.596 | -0.8923598 | 9.1979E-49 | 1.3913E-46 |
| ENSMUSG00000038074 | Fkbp14 | 3743.36373 | 1977.69599 | -1765.6677 | -0.8933251 | 3.848E-13 | 4.9755E-12 |
| ENSMUSG00000079051 | Vinac1 | 13.9009482 | 5.71731449 | -8.1836337 | -0.893415 | 0.00734894 | 0.02021155 |
| ENSMUSG00000080365 | Gm25776 | 5.00059887 | 0.95150704 | -4.0490918 | -0.893632 | 0.01078867 | 0.02837077 |

| | | | | | | | |
|---|---|---|---|---|---|---|---|
| ENSMUSG00000077440 | Gm23130 | 172.91444 | 84.9538587 | -87.960581 | -0.8938131 | 0.00023673 | 0.00091064 |
| ENSMUSG00000027074 | Slc43a3 | 3371.1734 | 1802.54904 | -1568.6244 | -0.8938254 | 2.4816E-33 | 1.5855E-31 |
| ENSMUSG00000035745 | Grin3b | 11.6570531 | 4.10035414 | -7.556699 | -0.8941058 | 0.01251129 | 0.03226711 |
| ENSMUSG00000022296 | Baalc | 167.040001 | 86.1805469 | -80.859454 | -0.8952751 | 8.3242E-07 | 4.7888E-06 |
| ENSMUSG00000028069 | Gpatch4 | 4106.07113 | 2193.95359 | -1912.1175 | -0.8960332 | 7.6139E-39 | 6.9926E-37 |
| ENSMUSG00000044367 | Slc16a13 | 840.813582 | 443.434228 | -397.37935 | -0.896142 | 1.4898E-13 | 2.013E-12 |
| ENSMUSG00000080811 | - | 14.7335522 | 5.05469384 | -9.6788584 | -0.8965462 | 0.01277929 | 0.03285374 |
| ENSMUSG00000087461 | C230014O12 | 56.3933585 | 24.8471271 | -31.546231 | -0.8968865 | 0.00427313 | 0.0125259 |
| ENSMUSG00000036078 | Sigmar1 | 5973.43234 | 3183.85448 | -2789.5779 | -0.9013188 | 4.8876E-49 | 7.6754E-47 |
| ENSMUSG00000097467 | - | 305.507022 | 157.755591 | -147.75143 | -0.9014497 | 6.5285E-08 | 4.3962E-07 |
| ENSMUSG00000090628 | - | 16.6403388 | 5.01949822 | -11.620841 | -0.9018506 | 0.01314452 | 0.03367066 |
| ENSMUSG00000040998 | Npnt | 13994.9859 | 7410.51568 | -6584.4702 | -0.9019398 | 4.6705E-22 | 1.4006E-20 |
| ENSMUSG00000060512 | 0610040J01F | 3428.33648 | 1814.46065 | -1613.8758 | -0.9037278 | 3.7488E-24 | 1.3005E-22 |
| ENSMUSG00000073680 | Tmem88b | 10.7815325 | 4.19447839 | -6.5870541 | -0.90428 | 0.00921532 | 0.02466532 |
| ENSMUSG00000006575 | Rundc3a | 406.05275 | 212.123914 | -193.92884 | -0.904835 | 2.8875E-12 | 3.3963E-11 |
| ENSMUSG00000039748 | Exo1 | 2940.99607 | 1560.93345 | -1380.0626 | -0.9064625 | 3.8007E-44 | 4.4679E-42 |
| ENSMUSG00000034192 | Lsm3 | 6834.75215 | 3615.60091 | -3219.1512 | -0.9083316 | 1.3489E-32 | 8.3084E-31 |
| ENSMUSG00000081745 | - | 15.0415402 | 4.6931907 | -10.34835 | -0.909553 | 0.0123029 | 0.03180388 |
| ENSMUSG00000045799 | - | 940.866047 | 485.981147 | -454.8849 | -0.9104951 | 1.1256E-09 | 9.6086E-09 |
| ENSMUSG00000043177 | - | 154.966376 | 49.5326452 | -105.43373 | -0.9112231 | 0.01198191 | 0.03110593 |
| ENSMUSG00000003779 | Kif20a | 16126.021 | 8519.83329 | -7606.1877 | -0.9112586 | 2.1704E-36 | 1.6912E-34 |
| ENSMUSG00000083761 | - | 13.8646208 | 4.25243149 | -9.6121893 | -0.9119505 | 0.01196507 | 0.03107112 |
| ENSMUSG00000087213 | 2810408I11R | 574.626102 | 300.609698 | -274.0164 | -0.9136176 | 3.9738E-17 | 7.7782E-16 |
| ENSMUSG00000080818 | - | 10.1538709 | 3.52520804 | -6.6286628 | -0.914016 | 0.01042998 | 0.02753311 |
| ENSMUSG00000062421 | Arf2 | 7797.56248 | 4107.48826 | -3690.0742 | -0.9141027 | 5.1845E-32 | 3.0915E-30 |
| ENSMUSG00000031375 | Bgn | 28765.0254 | 15047.8702 | -13717.155 | -0.9143161 | 9.457E-18 | 1.9592E-16 |
| ENSMUSG00000097424 | 1700030M09 | 13.8207425 | 5.07324765 | -8.7474949 | -0.9143784 | 0.00870046 | 0.02346124 |
| ENSMUSG00000034206 | Polq | 3095.83508 | 1602.38434 | -1493.4507 | -0.9145223 | 4.1876E-11 | 4.2988E-10 |
| ENSMUSG00000046057 | - | 121.482058 | 59.4452726 | -62.036786 | -0.9157951 | 5.5213E-05 | 0.00023782 |
| ENSMUSG00000021270 | Hsp90aa1 | 122658.496 | 64422.9508 | -58235.545 | -0.9158421 | 1.7263E-26 | 7.1026E-25 |
| ENSMUSG00000064620 | Gm22303 | 13.6629731 | 4.43760123 | -9.2253718 | -0.9170415 | 0.01101538 | 0.02888572 |
| ENSMUSG00000020354 | Sgcd | 351.791141 | 171.745681 | -180.04546 | -0.9172244 | 7.2791E-05 | 0.00030711 |
| ENSMUSG00000051379 | Flrt3 | 1599.39363 | 837.013183 | -762.38045 | -0.9176046 | 2.9779E-21 | 8.369E-20 |
| ENSMUSG00000088632 | - | 7.51189828 | 1.98957836 | -5.5223199 | -0.9177067 | 0.01154179 | 0.03010202 |
| ENSMUSG00000014782 | Plekhg4 | 249.443939 | 126.725869 | -122.71807 | -0.9179006 | 1.1655E-07 | 7.5637E-07 |
| ENSMUSG00000088789 | - | 26.523733 | 10.4288912 | -16.094842 | -0.9199437 | 0.00664787 | 0.01852615 |
| ENSMUSG00000042660 | Wdr55 | 4766.30854 | 2466.52205 | -2299.7865 | -0.9201338 | 6.8014E-13 | 8.5161E-12 |
| ENSMUSG00000038374 | Rbm8a | 14245.2911 | 7462.61176 | -6782.6793 | -0.9216234 | 9.6777E-32 | 5.721E-30 |
| ENSMUSG00000066878 | - | 218.0839 | 112.497566 | -105.58633 | -0.9222863 | 8.9048E-12 | 9.9182E-11 |
| ENSMUSG00000022136 | Dnajc3 | 12386.4925 | 6519.99816 | -5866.4944 | -0.9224077 | 2.222E-98 | 2.2856E-95 |
| ENSMUSG00000042793 | Lgr6 | 3211.53098 | 1658.74916 | -1552.7818 | -0.9230947 | 4.6699E-13 | 5.9708E-12 |
| ENSMUSG00000088323 | Gm25663 | 15.754195 | 6.3346969 | -9.4194981 | -0.9241394 | 0.00564505 | 0.01603342 |
| ENSMUSG00000049246 | - | 20.2689598 | 8.04490776 | -12.224052 | -0.9248738 | 0.00666954 | 0.01858401 |
| ENSMUSG00000048109 | Rbm15 | 1517.54389 | 793.129778 | -724.41411 | -0.9249261 | 2.1838E-31 | 1.2514E-29 |
| ENSMUSG00000083245 | - | 29.3844442 | 13.6897646 | -15.69468 | -0.9253028 | 0.00070648 | 0.00245916 |
| ENSMUSG00000036492 | Rnf39 | 20.097703 | 5.87816246 | -14.219541 | -0.9264805 | 0.01082548 | 0.02843848 |
| ENSMUSG00000078592 | - | 42.2317168 | 19.0907762 | -23.140941 | -0.9265125 | 0.00112266 | 0.00375656 |
| ENSMUSG00000046295 | Ankle1 | 1004.56257 | 517.447948 | -487.11462 | -0.9266517 | 2.4269E-13 | 3.2045E-12 |
| ENSMUSG00000097252 | - | 516.846524 | 266.566429 | -250.28009 | -0.9283579 | 2.0354E-14 | 3.0189E-13 |
| ENSMUSG00000082424 | - | 55.9165682 | 25.9730081 | -29.94356 | -0.9286436 | 0.00045184 | 0.0016408 |
| ENSMUSG00000083261 | - | 61.3053607 | 28.546932 | -32.758429 | -0.9293184 | 0.00051401 | 0.00184347 |
| ENSMUSG00000065362 | Gm24411 | 32.2592955 | 14.3426265 | -17.916669 | -0.9300749 | 0.0016064 | 0.00518219 |
| ENSMUSG00000027171 | Prrg4 | 601.944017 | 302.076707 | -299.86731 | -0.9307797 | 2.1414E-07 | 1.3435E-06 |
| ENSMUSG00000050550 | - | 103.686931 | 46.348203 | -57.338728 | -0.9318689 | 0.00147267 | 0.00479896 |
| ENSMUSG00000077734 | - | 70.5374998 | 31.0562935 | -39.481206 | -0.9322674 | 0.00196392 | 0.00622701 |
| ENSMUSG00000056228 | Cars2 | 1464.44162 | 763.370068 | -701.07156 | -0.9326475 | 1.0495E-48 | 1.576E-46 |
| ENSMUSG00000024660 | Incenp | 14890.4734 | 7733.6271 | -7156.8463 | -0.933211 | 1.0858E-30 | 6.0699E-29 |
| ENSMUSG00000000948 | - | 632.301728 | 325.576532 | -306.7252 | -0.9333639 | 8.3594E-17 | 1.5953E-15 |
| ENSMUSG00000029409 | U90926 | 366.686532 | 106.705514 | -259.98102 | -0.9342883 | 0.01019464 | 0.02696031 |
| ENSMUSG00000020080 | Hkdc1 | 14.8786744 | 5.63952083 | -9.2391535 | -0.9349054 | 0.00618093 | 0.01737316 |
| ENSMUSG00000043807 | Ly6g5b | 11.4167451 | 3.71505758 | -7.7016875 | -0.9351432 | 0.00921323 | 0.02466291 |
| ENSMUSG00000042745 | Id1 | 9554.92407 | 4878.82359 | -4676.1005 | -0.9351447 | 4.4809E-12 | 5.144E-11 |
| ENSMUSG00000043618 | - | 221.617214 | 110.671328 | -110.94589 | -0.9352565 | 1.95E-07 | 1.2324E-06 |
| ENSMUSG00000090467 | - | 13.4688762 | 3.19616784 | -10.272708 | -0.9370396 | 0.0095135 | 0.02535783 |
| ENSMUSG00000064907 | - | 4.00938532 | 0.35067661 | -3.6587087 | -0.9375893 | 0.0040943 | 0.01206532 |
| ENSMUSG00000071658 | Gng3 | 15.6674001 | 5.48055526 | -10.186845 | -0.9382176 | 0.00789409 | 0.02152097 |
| ENSMUSG00000056201 | Cfl1 | 140604.188 | 72674.5181 | -67929.67 | -0.9385923 | 1.1802E-27 | 5.2665E-26 |
| ENSMUSG00000042734 | Ttc9 | 1218.96424 | 610.769227 | -608.19501 | -0.9388397 | 3.7192E-08 | 2.6024E-07 |
| ENSMUSG00000039055 | Eme1 | 1412.2699 | 722.600485 | -689.66942 | -0.940259 | 1.0965E-15 | 1.872E-14 |
| ENSMUSG00000081244 | - | 3.47967295 | 0.33779791 | -3.141875 | -0.9405668 | 0.00450036 | 0.01311164 |
| ENSMUSG00000065725 | Gm26165 | 8.41025892 | 2.57177269 | -5.8384862 | -0.9408554 | 0.00948333 | 0.02529379 |
| ENSMUSG00000043445 | Pgp | 3308.16336 | 1702.92148 | -1605.2419 | -0.9411788 | 8.3061E-23 | 2.641E-21 |
| ENSMUSG00000079852 | Klra18; Klra4 | 10235.4059 | 5202.50141 | -5032.9045 | -0.9421498 | 1.8529E-12 | 2.2162E-11 |
| ENSMUSG00000033597 | Caskin1 | 46.4728976 | 19.7009692 | -26.771928 | -0.942716 | 0.00235506 | 0.00733177 |
| ENSMUSG00000032477 | Cdc25a | 3884.36724 | 1999.91496 | -1884.4523 | -0.9430078 | 1.1673E-25 | 4.5568E-24 |

| | | | | | | | |
|---|---|---|---|---|---|---|---|
| ENSMUSG00000026202 | Tuba4a | 3713.59148 | 1780.72924 | -1932.8622 | -0.9454237 | 3.1703E-05 | 0.00014206 |
| ENSMUSG00000079426 | Arpc4 | 13809.5056 | 7115.11078 | -6694.3949 | -0.9454501 | 9.9501E-34 | 6.5818E-32 |
| ENSMUSG00000032575 | Manf | 14096.6144 | 7215.20722 | -6881.4072 | -0.9454545 | 3.0122E-19 | 7.1473E-18 |
| ENSMUSG00000086021 | - | 145.37432 | 72.2908287 | -73.083492 | -0.9457861 | 9.1696E-08 | 6.0441E-07 |
| ENSMUSG00000080914 | - | 19.6488491 | 7.41811833 | -12.230731 | -0.9469492 | 0.00559273 | 0.01589798 |
| ENSMUSG00000032401 | Lctl | 222.466722 | 108.569794 | -113.89693 | -0.9492965 | 2.2495E-06 | 1.2067E-05 |
| ENSMUSG00000049539 | H1f1 | 60.9764561 | 26.1097566 | -34.866699 | -0.9493399 | 0.00175433 | 0.00561713 |
| ENSMUSG00000050503 | Fbxl22 | 27.1928184 | 11.9340054 | -15.258813 | -0.9493607 | 0.00143203 | 0.00467614 |
| ENSMUSG00000026955 | Sapcd2 | 839.013601 | 428.662652 | -410.35095 | -0.9505136 | 4.2712E-24 | 1.4694E-22 |
| ENSMUSG00000038845 | Phb | 7766.0917 | 3996.83652 | -3769.2552 | -0.950873 | 8.0845E-52 | 1.4462E-49 |
| ENSMUSG00000081892 | - | 38.2596205 | 16.7972385 | -21.462382 | -0.950874 | 0.00103072 | 0.0034755 |
| ENSMUSG00000074800 | - | 2132.54438 | 553.708324 | -1578.8361 | -0.9513358 | 0.00877601 | 0.02363401 |
| ENSMUSG00000097711 | Gm5523 | 62.6845807 | 28.4649283 | -34.219652 | -0.9531233 | 0.00038318 | 0.00140845 |
| ENSMUSG00000030031 | Kbtbd8 | 412.498356 | 202.885564 | -209.61279 | -0.9532694 | 3.2695E-07 | 2E-06 |
| ENSMUSG00000069266 | H4c2 | 5.56909315 | 0.97755652 | -4.5915366 | -0.9533795 | 0.00698322 | 0.01932977 |
| ENSMUSG00000038803 | Ost4 | 2442.11652 | 1244.79671 | -1197.3198 | -0.9537469 | 1.602E-22 | 4.986E-21 |
| ENSMUSG00000080929 | - | 3.88710314 | 0.50151691 | -3.3855862 | -0.9539821 | 0.00527136 | 0.01508031 |
| ENSMUSG00000084299 | - | 4.33841161 | 0.47764973 | -3.8607619 | -0.9544192 | 0.00464576 | 0.01348562 |
| ENSMUSG00000083659 | - | 7.63088584 | 2.25184892 | -5.3790369 | -0.9546282 | 0.00857412 | 0.02317219 |
| ENSMUSG00000064853 | Gm23442 | 6.31394061 | 1.27521531 | -5.0387253 | -0.9550815 | 0.00774649 | 0.02118595 |
| ENSMUSG00000025980 | Hspd1 | 60719.2267 | 30910.957 | -29808.27 | -0.9560755 | 1.6833E-22 | 5.2073E-21 |
| ENSMUSG00000081614 | - | 7.69909701 | 1.75625829 | -5.9428387 | -0.9570829 | 0.00815602 | 0.02216162 |
| ENSMUSG00000054161 | Fam83e | 11.148379 | 3.82380804 | -7.324571 | -0.9571953 | 0.0067479 | 0.01877438 |
| ENSMUSG00000001056 | Nhp2 | 9507.67116 | 4840.06094 | -4667.6102 | -0.9580335 | 3.5904E-25 | 1.3503E-23 |
| ENSMUSG00000050677 | Ccdc96 | 130.131441 | 62.239247 | -67.892194 | -0.9584071 | 1.1857E-05 | 5.7018E-05 |
| ENSMUSG00000014704 | Hoxa2 | 171.946906 | 85.6105407 | -86.336366 | -0.9588922 | 5.9038E-10 | 5.2328E-09 |
| ENSMUSG00000074445 | Sprr2a3 | 7.21655755 | 1.25732287 | -5.9592347 | -0.9589473 | 0.00668967 | 0.01863001 |
| ENSMUSG00000064208 | - | 21.6192817 | 7.27673765 | -14.342544 | -0.9591883 | 0.00688849 | 0.01911128 |
| ENSMUSG00000080801 | - | 9.61480583 | 2.82664817 | -6.7881577 | -0.9616628 | 0.00798463 | 0.02174475 |
| ENSMUSG00000030587 | 2200002D01 | 9.96157577 | 3.40291622 | -6.5586595 | -0.9620866 | 0.00668088 | 0.01861056 |
| ENSMUSG00000031919 | Tmed6 | 7.45453266 | 1.84388813 | -5.6106445 | -0.9631804 | 0.00794711 | 0.02165977 |
| ENSMUSG00000020277 | Pfkl | 22280.4819 | 11333.1631 | -10947.319 | -0.9632185 | 2.2092E-33 | 1.4247E-31 |
| ENSMUSG00000027133 | Nop10 | 4136.90144 | 2096.26142 | -2040.64 | -0.9636388 | 1.4381E-24 | 5.1543E-23 |
| ENSMUSG00000078915 | - | 45.8816849 | 20.3634698 | -25.518215 | -0.9641452 | 0.00049771 | 0.00179193 |
| ENSMUSG00000004933 | Matk | 38.0248184 | 16.9695421 | -21.055276 | -0.9641553 | 0.00051983 | 0.00186208 |
| ENSMUSG00000093992 | - | 5.18450298 | 0.83332179 | -4.3511812 | -0.9645179 | 0.00577146 | 0.01635406 |
| ENSMUSG00000093916 | - | 111.370451 | 52.8510479 | -58.519403 | -0.9645345 | 1.4828E-05 | 7.019E-05 |
| ENSMUSG00000049721 | Gal3st1 | 439.30789 | 221.6529 | -217.65499 | -0.9654264 | 2.7623E-19 | 6.5924E-18 |
| ENSMUSG00000077711 | - | 26.9768432 | 11.7776539 | -15.199189 | -0.9655263 | 0.00062872 | 0.00221136 |
| ENSMUSG00000063882 | Uqcrh | 7374.04156 | 3738.1499 | -3635.8917 | -0.966244 | 1.0143E-29 | 5.1775E-28 |
| ENSMUSG00000049932 | H2ax | 4098.23878 | 2081.46895 | -2016.7698 | -0.9663145 | 5.4523E-37 | 4.4159E-35 |
| ENSMUSG00000077563 | - | 215.824043 | 105.452164 | -110.37188 | -0.9670491 | 5.2731E-08 | 3.5943E-07 |
| ENSMUSG00000000325 | - | 9.1695554 | 1.63710572 | -7.5324497 | -0.9671668 | 0.00601957 | 0.01697527 |
| ENSMUSG00000092116 | - | 56.5586789 | 26.3584327 | -30.200246 | -0.9678337 | 6.5564E-05 | 0.00027971 |
| ENSMUSG00000096544 | - | 1291.22127 | 644.598057 | -646.62321 | -0.9690009 | 1.821E-13 | 2.4358E-12 |
| ENSMUSG00000033170 | Card10 | 1451.97599 | 724.875779 | -727.10021 | -0.9712982 | 9.1984E-15 | 1.4185E-13 |
| ENSMUSG00000030890 | Ilk | 46.6082575 | 21.7144285 | -24.893829 | -0.9713459 | 6.4467E-05 | 0.00027532 |
| ENSMUSG00000038816 | Ctnnal1 | 3390.0849 | 1689.83497 | -1700.2499 | -0.9718646 | 5.0031E-14 | 7.1425E-13 |
| ENSMUSG00000087569 | - | 8.18163647 | 1.62439051 | -6.557246 | -0.9720447 | 0.00657264 | 0.01834134 |
| ENSMUSG00000018909 | Arrb1 | 6365.78557 | 3208.23395 | -3157.5516 | -0.972199 | 1.1809E-25 | 4.6012E-24 |
| ENSMUSG00000024790 | Sac3d1 | 1638.94666 | 821.193009 | -817.75365 | -0.9726268 | 6.8782E-18 | 1.4468E-16 |
| ENSMUSG00000062075 | Lmnb2 | 3899.72772 | 1975.45699 | -1924.2707 | -0.9728977 | 5.9725E-52 | 1.0778E-49 |
| ENSMUSG00000025757 | Hspa4l | 40393.1533 | 20311.7306 | -20081.423 | -0.9740163 | 7.0791E-24 | 2.4032E-22 |
| ENSMUSG00000092819 | Gm23639 | 27.0071126 | 11.3977281 | -15.609384 | -0.9743587 | 0.00125223 | 0.00415096 |
| ENSMUSG00000042988 | Notum | 2534.95109 | 1253.99978 | -1280.9513 | -0.9784428 | 8.7581E-13 | 1.0841E-11 |
| ENSMUSG00000098111 | - | 169.349919 | 72.697268 | -96.652651 | -0.9789816 | 0.00079984 | 0.00275848 |
| ENSMUSG00000026238 | Ptma | 89885.4395 | 44655.9989 | -45229.441 | -0.9816951 | 1.3191E-16 | 2.458E-15 |
| ENSMUSG00000097152 | - | 10.5841411 | 3.34546072 | -7.2386803 | -0.9826882 | 0.00611507 | 0.01721389 |
| ENSMUSG00000025196 | Cpn1 | 28.2811412 | 12.173534 | -16.107607 | -0.9833567 | 0.00088174 | 0.00301365 |
| ENSMUSG00000028410 | Dnaja1 | 17569.5248 | 8824.47032 | -8745.0544 | -0.9841809 | 3.4237E-45 | 4.1924E-43 |
| ENSMUSG00000064380 | Gm26448 | 27.4079535 | 10.8794696 | -16.528484 | -0.9858403 | 0.00202129 | 0.00639034 |
| ENSMUSG00000050195 | Scd4 | 16.8127797 | 4.86252118 | -11.950258 | -0.9859686 | 0.00641921 | 0.0179619 |
| ENSMUSG00000031104 | Rab33a | 25.1530257 | 10.2211035 | -14.931922 | -0.9877647 | 0.00136452 | 0.00448275 |
| ENSMUSG00000069008 | - | 44.7625967 | 4.37892514 | -40.383672 | -0.9881617 | 0.00262683 | 0.00807411 |
| ENSMUSG00000018770 | Atp5g3 | 20987.0072 | 10435.8522 | -10551.155 | -0.9888968 | 1.9775E-23 | 6.53E-22 |
| ENSMUSG00000049299 | Trappc1 | 3633.69342 | 1804.64609 | -1829.0473 | -0.9889904 | 5.2339E-22 | 1.565E-20 |
| ENSMUSG00000093157 | Mir3098 | 20.5279732 | 6.84038325 | -13.68759 | -0.9896934 | 0.00514765 | 0.01477365 |
| ENSMUSG00000094234 | - | 89.1584781 | 19.5890613 | -69.569417 | -0.9923178 | 0.00591976 | 0.01671671 |
| ENSMUSG00000065676 | - | 31.5885956 | 13.6853936 | -17.903202 | -0.9923757 | 0.0004858 | 0.00175177 |
| ENSMUSG00000006567 | Atp7b | 42.2535951 | 17.4582324 | -24.795363 | -0.9940468 | 0.00106592 | 0.00358125 |
| ENSMUSG00000065701 | Rny1 | 52.069087 | 22.7268626 | -29.342224 | -0.996075 | 0.00023174 | 0.00089311 |
| ENSMUSG00000028370 | Pappa | 650.957046 | 311.121071 | -339.83598 | -0.9969874 | 2.2741E-08 | 1.6438E-07 |
| ENSMUSG00000022221 | Ripk3 | 119.994884 | 55.3125887 | -64.682295 | -0.9970438 | 1.1975E-05 | 5.7557E-05 |
| ENSMUSG00000048616 | Nog | 179.6731 | 59.4153493 | -120.25775 | -0.9984353 | 0.00483308 | 0.01398594 |
| ENSMUSG00000074252 | Gm10654 | 24.8841284 | 7.79476054 | -17.089368 | -0.9988417 | 0.005385 | 0.01536906 |

| | | | | | | | |
|---|---|---|---|---|---|---|---|
| ENSMUSG00000029161 | Cgref1 | 2110.09992 | 1042.01512 | -1068.0848 | -0.9989974 | 1.0482E-23 | 3.5193E-22 |
| ENSMUSG00000038068 | Rnf144b | 307.964681 | 142.461648 | -165.50303 | -1.0000399 | 5.9173E-06 | 2.9807E-05 |
| ENSMUSG00000015812 | Gnrh1 | 55.0035362 | 25.1581191 | -29.845417 | -1.0015038 | 2.4529E-05 | 0.00011204 |
| ENSMUSG00000028360 | Slc44a5 | 4.2683987 | 0.47293439 | -3.7954643 | -1.0046707 | 0.00335976 | 0.0100945 |
| ENSMUSG00000071470 | Ccnb1ip1 | 234.427319 | 108.864432 | -125.56289 | -1.006415 | 1.1707E-06 | 6.5682E-06 |
| ENSMUSG00000022893 | Adamts1 | 28788.0726 | 14208.5553 | -14579.517 | -1.006754 | 1.0531E-37 | 8.9154E-36 |
| ENSMUSG00000092827 | Mir3091 | 158.9216 | 75.5214134 | -83.400187 | -1.0068211 | 1.11E-08 | 8.3704E-08 |
| ENSMUSG00000065847 | Gm25188 | 6.16552381 | 1.17766661 | -4.9878572 | -1.0075679 | 0.00496998 | 0.0143277 |
| ENSMUSG00000050100 | Hmx2 | 66.2982214 | 28.9959814 | -37.30224 | -1.0077118 | 0.00011541 | 0.00046966 |
| ENSMUSG00000029352 | Crybb3 | 8.79155447 | 1.80452927 | -6.9870252 | -1.0077252 | 0.00523733 | 0.01499964 |
| ENSMUSG00000074978 | - | 22.6865662 | 7.90035738 | -14.786209 | -1.0078307 | 0.003541 | 0.01058645 |
| ENSMUSG00000012705 | Retn | 5.05712444 | 0.64327726 | -4.4138472 | -1.0083086 | 0.00360394 | 0.01075277 |
| ENSMUSG00000079501 | - | 29.4740025 | 11.7003682 | -17.773634 | -1.0084165 | 0.00137996 | 0.00452623 |
| ENSMUSG00000090602 | - | 119.814517 | 56.4359905 | -63.378527 | -1.0085658 | 8.557E-08 | 5.6658E-07 |
| ENSMUSG00000041358 | Nutm1 | 22.4647203 | 8.7730265 | -13.691694 | -1.0096665 | 0.00172147 | 0.00552396 |
| ENSMUSG00000065509 | Mir421 | 8.01547601 | 2.21791618 | -5.7975598 | -1.0097296 | 0.00531099 | 0.0151789 |
| ENSMUSG00000018451 | 6330403K07 | 12.8676581 | 3.92652923 | -8.9411289 | -1.0097369 | 0.00486985 | 0.01407453 |
| ENSMUSG00000042215 | Bag2 | 3362.33937 | 1657.20569 | -1705.1337 | -1.0098612 | 5.8665E-42 | 6.2857E-40 |
| ENSMUSG00000055831 | - | 56.8060067 | 24.9089146 | -31.897092 | -1.0111266 | 7.9294E-05 | 0.00033176 |
| ENSMUSG00000061186 | Sfmbt2 | 244.473752 | 116.46165 | -128.0121 | -1.0127712 | 1.0697E-09 | 9.1538E-09 |
| ENSMUSG00000058838 | - | 36.5106034 | 14.036374 | -22.474229 | -1.0139755 | 0.0015767 | 0.00509678 |
| ENSMUSG00000091623 | - | 45.2180048 | 19.6213127 | -25.596692 | -1.015342 | 0.0001213 | 0.00049153 |
| ENSMUSG00000048562 | Sp8 | 3249.84196 | 1548.62815 | -1701.2138 | -1.01588 | 1.7318E-10 | 1.6563E-09 |
| ENSMUSG00000086925 | - | 29.9445475 | 12.3452334 | -17.599314 | -1.0166118 | 0.00054602 | 0.00194574 |
| ENSMUSG00000032593 | Amigo3 | 201.846513 | 96.2506455 | -105.59587 | -1.0169401 | 6.2436E-11 | 6.2704E-10 |
| ENSMUSG00000094732 | 1500015L24 | 14.4273659 | 4.83696608 | -9.5903998 | -1.0208839 | 0.0034731 | 0.01039556 |
| ENSMUSG00000017499 | Cdc6 | 4367.27822 | 2112.26059 | -2255.0176 | -1.021244 | 4.2826E-19 | 1.0057E-17 |
| ENSMUSG00000038605 | Samd10 | 318.064151 | 152.378946 | -165.6852 | -1.0218967 | 4.0975E-14 | 5.9236E-13 |
| ENSMUSG00000017861 | Mybl2 | 3782.04626 | 1849.13351 | -1932.9128 | -1.0219828 | 2.3597E-46 | 3.0723E-44 |
| ENSMUSG00000048617 | Rtbdn | 17.2634353 | 6.0103767 | -11.253059 | -1.0227792 | 0.00318432 | 0.0096137 |
| ENSMUSG00000021670 | Hmgcr | 10652.5434 | 5127.73621 | -5524.8072 | -1.0238305 | 8.7221E-17 | 1.6568E-15 |
| ENSMUSG00000020897 | Aurkb | 5793.59231 | 2831.1402 | -2962.4521 | -1.0243973 | 1.0467E-56 | 2.3661E-54 |
| ENSMUSG00000048878 | Hexim1 | 2581.37419 | 1251.20223 | -1330.172 | -1.0263426 | 4.3528E-27 | 1.8656E-25 |
| ENSMUSG00000058809 | - | 6417.05498 | 3102.29652 | -3314.7585 | -1.027946 | 2.9988E-24 | 1.0474E-22 |
| ENSMUSG00000026069 | Il1rl1 | 50856.0352 | 23420.698 | -27435.337 | -1.0286244 | 2.2547E-07 | 1.4111E-06 |
| ENSMUSG00000027774 | Gfm1 | 9461.94436 | 4624.75862 | -4837.1857 | -1.0289796 | 3.373E-121 | 9.912E-118 |
| ENSMUSG00000085119 | - | 28.939725 | 11.7099575 | -17.229767 | -1.0317448 | 0.00051031 | 0.00183084 |
| ENSMUSG00000025792 | Slc25a10 | 3836.62921 | 1862.70727 | -1973.9219 | -1.0323079 | 2.5086E-48 | 3.6089E-46 |
| ENSMUSG00000025185 | Loxl4 | 3937.34178 | 1911.13536 | -2026.2064 | -1.0327337 | 3.2998E-45 | 4.0649E-43 |
| ENSMUSG00000082810 | - | 8.91713127 | 1.98185605 | -6.9352752 | -1.033215 | 0.00441179 | 0.01288828 |
| ENSMUSG00000021485 | Mxd3 | 761.549257 | 360.968922 | -400.58034 | -1.0341903 | 8.9775E-14 | 1.2428E-12 |
| ENSMUSG00000037364 | Srrt | 12183.7865 | 5904.87416 | -6278.9123 | -1.0348403 | 2.3305E-48 | 3.3762E-46 |
| ENSMUSG00000085898 | - | 6.06937626 | 0.61461374 | -5.4547625 | -1.0369705 | 0.00234174 | 0.00729252 |
| ENSMUSG00000031683 | Lsm6 | 7482.883 | 3594.20608 | -3888.6769 | -1.0369756 | 1.5032E-24 | 5.3781E-23 |
| ENSMUSG00000013415 | Igf2bp1 | 94.8886212 | 42.2110465 | -52.677575 | -1.0378668 | 8.5561E-06 | 4.2074E-05 |
| ENSMUSG00000047428 | Dlk2 | 2361.62424 | 1129.91986 | -1231.7044 | -1.038398 | 6.961E-21 | 1.8992E-19 |
| ENSMUSG00000092730 | Snora24 | 92.6771412 | 42.2629457 | -50.414195 | -1.0406422 | 2.2818E-07 | 1.4277E-06 |
| ENSMUSG00000027489 | Necab3 | 67.9510911 | 30.0329752 | -37.918116 | -1.041156 | 8.4017E-06 | 4.133E-05 |
| ENSMUSG00000002068 | Ccne1 | 3761.04774 | 1810.20053 | -1950.8472 | -1.041833 | 3.8712E-38 | 3.3322E-36 |
| ENSMUSG00000022790 | Igsf11 | 72.950541 | 29.3310763 | -43.619465 | -1.0427703 | 0.00040775 | 0.0014907 |
| ENSMUSG00000084899 | - | 48.2700493 | 18.7529379 | -29.517111 | -1.0433244 | 0.00080463 | 0.00277408 |
| ENSMUSG00000080875 | - | 334.940167 | 158.080395 | -176.85977 | -1.0441535 | 4.5102E-15 | 7.2261E-14 |
| ENSMUSG00000063586 | - | 25.6383286 | 10.1092467 | -15.529082 | -1.0451295 | 0.00062873 | 0.00221136 |
| ENSMUSG00000079429 | Mroh2a | 595.393938 | 273.416765 | -321.97717 | -1.0454582 | 1.4061E-08 | 1.0424E-07 |
| ENSMUSG00000092086 | - | 37.5378354 | 13.2764997 | -24.261336 | -1.0478277 | 0.00164732 | 0.00530174 |
| ENSMUSG00000031987 | Egln1 | 19041.6863 | 9074.43015 | -9967.2561 | -1.0485896 | 1.209E-25 | 4.7016E-24 |
| ENSMUSG00000095512 | - | 17.7273911 | 6.65398418 | -11.073407 | -1.0488952 | 0.00105945 | 0.00356243 |
| ENSMUSG00000039095 | En2 | 306.111879 | 142.063532 | -164.04835 | -1.049558 | 1.2236E-10 | 1.1964E-09 |
| ENSMUSG00000021208 | Ifi27l2b | 71.0296142 | 29.8244867 | -41.205127 | -1.049584 | 7.7151E-05 | 0.00032312 |
| ENSMUSG00000056888 | Glipr1 | 36.1033604 | 12.5930641 | -23.510296 | -1.050634 | 0.00193348 | 0.00614009 |
| ENSMUSG00000029581 | Fscn1 | 20834.91 | 10020.893 | -10814.017 | -1.0521535 | 1.158E-129 | 4.766E-126 |
| ENSMUSG00000055416 | - | 5.09165005 | 0.66897195 | -4.4226781 | -1.0539496 | 0.00264473 | 0.00812534 |
| ENSMUSG00000077285 | - | 16.8154215 | 5.79581998 | -11.019601 | -1.0576177 | 0.00182176 | 0.00581763 |
| ENSMUSG00000034120 | Srsf2 | 17260.1187 | 8181.10764 | -9079.0111 | -1.0590218 | 9.4591E-30 | 4.8406E-28 |
| ENSMUSG00000090173 | Fbxw10 | 7.66031509 | 1.96052191 | -5.6997932 | -1.0595905 | 0.00350696 | 0.0104908 |
| ENSMUSG00000032193 | Ldlr | 4303.36012 | 1858.30986 | -2445.0503 | -1.0603183 | 1.1676E-05 | 5.6199E-05 |
| ENSMUSG00000074650 | - | 10.013845 | 3.05624539 | -6.9575996 | -1.0607314 | 0.00273324 | 0.00837154 |
| ENSMUSG00000080486 | - | 157.262759 | 69.2953623 | -87.967397 | -1.061342 | 1.6425E-06 | 9.0201E-06 |
| ENSMUSG00000066362 | - | 27.12442 | 8.77196614 | -18.352454 | -1.0627682 | 0.00225284 | 0.00704873 |
| ENSMUSG00000064563 | Snora7a | 153.433123 | 70.1775986 | -83.255524 | -1.0628985 | 4.3205E-10 | 3.9155E-09 |
| ENSMUSG00000063694 | Cycs | 11614.9062 | 5487.70398 | -6127.2023 | -1.0639338 | 1.0304E-30 | 5.7758E-29 |
| ENSMUSG00000028780 | Sema3c | 22773.5446 | 10555.6118 | -12217.933 | -1.0642731 | 1.4189E-13 | 1.9268E-12 |
| ENSMUSG00000033854 | Kcnk10 | 33.853529 | 11.5013737 | -22.352155 | -1.0646762 | 0.00172797 | 0.00554308 |
| ENSMUSG00000046179 | E2f8 | 1407.45623 | 651.773829 | -755.6824 | -1.0650309 | 1.4735E-13 | 1.9929E-12 |
| ENSMUSG00000032085 | Tagln | 132.939265 | 61.1012456 | -71.83802 | -1.0662199 | 7.4881E-12 | 8.4178E-11 |

| | | | | | | | |
|---|---|---|---|---|---|---|---|
| ENSMUSG00000037601 | Nme1 | 7648.61198 | 3610.00273 | -4038.6092 | -1.0668771 | 1.4421E-33 | 9.4183E-32 |
| ENSMUSG00000085985 | - | 13.6064475 | 3.96058227 | -9.6458652 | -1.0677824 | 0.00286937 | 0.00874628 |
| ENSMUSG00000026971 | Itgb6 | 134.558141 | 53.3819015 | -81.176239 | -1.0708387 | 0.00022506 | 0.00086964 |
| ENSMUSG00000083836 | - | 99.8338785 | 44.7686891 | -55.065189 | -1.071048 | 3.8575E-08 | 2.6962E-07 |
| ENSMUSG00000039620 | Trmt9b | 75.1922904 | 32.6744964 | -42.517794 | -1.0720139 | 2.2859E-06 | 1.225E-05 |
| ENSMUSG00000058672 | Tubb2a | 10161.2009 | 4799.06995 | -5362.131 | -1.0725454 | 6.1518E-56 | 1.3322E-53 |
| ENSMUSG00000061488 | - | 16.9858382 | 4.77619579 | -12.209642 | -1.074979 | 0.00278206 | 0.00850663 |
| ENSMUSG00000097138 | - | 16.4822605 | 5.33434288 | -11.147918 | -1.0784572 | 0.00177388 | 0.00567532 |
| ENSMUSG00000083391 | - | 47.3932582 | 18.9047509 | -28.488507 | -1.0800516 | 0.00015057 | 0.00059939 |
| ENSMUSG00000083460 | - | 28.999854 | 10.8766224 | -18.123232 | -1.0806978 | 0.0004668 | 0.00168919 |
| ENSMUSG00000083311 | - | 12.4304898 | 2.92273732 | -9.5077525 | -1.081926 | 0.00291639 | 0.00887384 |
| ENSMUSG00000043102 | Qrfp | 9.91769749 | 2.90448218 | -7.0132153 | -1.0821998 | 0.00236456 | 0.00735689 |
| ENSMUSG00000035049 | Rrp12 | 4059.92292 | 1891.1692 | -2168.7537 | -1.0839852 | 5.0222E-31 | 2.8462E-29 |
| ENSMUSG00000054932 | Afp | 235.334602 | 106.656086 | -128.67852 | -1.0847672 | 1.6165E-12 | 1.9515E-11 |
| ENSMUSG00000033272 | Slc35a4 | 10634.4931 | 4995.92993 | -5638.5632 | -1.0854929 | 5.708E-124 | 1.957E-120 |
| ENSMUSG00000097755 | - | 289.659887 | 130.280694 | -159.37919 | -1.0897305 | 1.3899E-11 | 1.5105E-10 |
| ENSMUSG00000093178 | - | 78.8041294 | 32.7773389 | -46.02679 | -1.0898675 | 1.5566E-05 | 7.3429E-05 |
| ENSMUSG00000052861 | Dnah6 | 58.1952977 | 24.1030533 | -34.092244 | -1.0904281 | 2.3191E-05 | 0.0001063 |
| ENSMUSG00000031972 | Acta1 | 24.4445056 | 9.0068601 | -15.437645 | -1.0922021 | 0.00046225 | 0.00167418 |
| ENSMUSG00000033777 | Tlr13 | 4.89935497 | 0.64327726 | -4.2560777 | -1.0931292 | 0.00200126 | 0.00633236 |
| ENSMUSG00000044744 | - | 52.8597361 | 21.52536 | -31.334376 | -1.0950863 | 3.0502E-05 | 0.00013722 |
| ENSMUSG00000024761 | - | 151.339459 | 16.0986063 | -135.24085 | -1.0961449 | 0.00127887 | 0.00422836 |
| ENSMUSG00000032060 | Cryab | 22073.7942 | 9477.89789 | -12595.896 | -1.096173 | 6.7567E-07 | 3.9354E-06 |
| ENSMUSG00000062077 | Trim54 | 319.159306 | 139.316081 | -179.84323 | -1.0975523 | 4.0502E-08 | 2.8215E-07 |
| ENSMUSG00000041247 | Lamp3 | 13.1122224 | 4.16733452 | -8.9448879 | -1.0979008 | 0.00159068 | 0.00513631 |
| ENSMUSG00000044217 | Aqp5 | 20050.1337 | 9307.43357 | -10742.7 | -1.0980392 | 7.6965E-63 | 2.2619E-60 |
| ENSMUSG00000058101 | 1700084J12F | 4.45739916 | 0 | -4.4573992 | -1.0986162 | 0.00038928 | 0.00142904 |
| ENSMUSG00000029372 | Ppbp | 3582.01445 | 1528.10385 | -2053.9106 | -1.0991626 | 1.0111E-06 | 5.7332E-06 |
| ENSMUSG00000038188 | Scarf1 | 10.7586925 | 2.99098375 | -7.7677087 | -1.1019263 | 0.00200386 | 0.00633913 |
| ENSMUSG00000027220 | Syt13 | 969.339469 | 442.590172 | -526.7493 | -1.1026472 | 1.6355E-22 | 5.0747E-21 |
| ENSMUSG00000008482 | Rnf151 | 80.865922 | 34.2473335 | -46.618589 | -1.1045421 | 2.2316E-06 | 1.1974E-05 |
| ENSMUSG00000035964 | Tmem59l | 35.1448707 | 11.2316743 | -23.913196 | -1.1047577 | 0.00126981 | 0.00420111 |
| ENSMUSG00000051716 | Apon | 23.1709074 | 8.1073343 | -15.063573 | -1.1052226 | 0.00062461 | 0.00219913 |
| ENSMUSG00000097035 | - | 76.05942 | 31.7474482 | -44.311972 | -1.1064631 | 4.1885E-06 | 2.1595E-05 |
| ENSMUSG00000067161 | - | 93.0895461 | 39.1978494 | -53.891697 | -1.1068743 | 1.9042E-06 | 1.0355E-05 |
| ENSMUSG00000025192 | Entpd7 | 1095.06626 | 488.095285 | -606.97097 | -1.1087314 | 1.9208E-12 | 2.2947E-11 |
| ENSMUSG00000089617 | - | 13.7675116 | 3.03630803 | -10.731204 | -1.1087443 | 0.00229883 | 0.00717407 |
| ENSMUSG00000082746 | - | 15.1704116 | 4.72707277 | -10.443339 | -1.1120649 | 0.00123487 | 0.0040987 |
| ENSMUSG00000032315 | Cyp1a1 | 121.656797 | 13.0292848 | -108.62751 | -1.1121377 | 0.00114543 | 0.00382467 |
| ENSMUSG00000030500 | Slc17a6 | 7.91423211 | 1.64566827 | -6.2685638 | -1.1145383 | 0.0021754 | 0.00682511 |
| ENSMUSG00000038600 | Atp6v0a4 | 232.148002 | 103.438678 | -128.70932 | -1.1147328 | 1.5744E-13 | 2.1224E-12 |
| ENSMUSG00000096755 | - | 16.7235302 | 4.52033388 | -12.203196 | -1.1157721 | 0.0019124 | 0.00607783 |
| ENSMUSG00000042816 | Gpr151 | 11.6402712 | 3.11585558 | -8.5244156 | -1.1201089 | 0.00174859 | 0.00560049 |
| ENSMUSG00000028527 | Ak4 | 5906.44856 | 2479.89435 | -3426.5542 | -1.1201663 | 6.0632E-07 | 3.5587E-06 |
| ENSMUSG00000005483 | Dnajb1 | 9740.95889 | 4455.50332 | -5285.4556 | -1.1205144 | 6.1756E-77 | 2.9545E-74 |
| ENSMUSG00000075312 | - | 195.711474 | 55.2805056 | -140.43097 | -1.12188 | 0.00164402 | 0.00529276 |
| ENSMUSG00000022174 | Dad1 | 6455.88595 | 2931.54601 | -3524.3399 | -1.1220393 | 1.1665E-37 | 9.835E-36 |
| ENSMUSG00000084215 | - | 56.7669159 | 22.6396352 | -34.127281 | -1.1235606 | 1.3347E-05 | 6.3691E-05 |
| ENSMUSG00000049357 | Brd8dc | 48.7289087 | 19.6023247 | -29.126584 | -1.1259954 | 8.1031E-06 | 3.9985E-05 |
| ENSMUSG00000037278 | Tmem97 | 4167.45506 | 1884.16501 | -2283.29 | -1.1289474 | 2.0338E-39 | 1.9461E-37 |
| ENSMUSG00000021250 | Fos | 2200.73197 | 955.116285 | -1245.6157 | -1.1292485 | 1.4878E-10 | 1.437E-09 |
| ENSMUSG00000027368 | Dusp2 | 102.796178 | 33.8533421 | -68.942835 | -1.1312809 | 0.00070142 | 0.00244404 |
| ENSMUSG00000038167 | Plekhg6 | 72.1359895 | 30.2413289 | -41.894661 | -1.1350936 | 1.3291E-07 | 8.5715E-07 |
| ENSMUSG00000022176 | Rem2 | 980.052475 | 439.281765 | -540.77071 | -1.1357329 | 2.0274E-31 | 1.165E-29 |
| ENSMUSG00000096280 | - | 4.49192478 | 0.30640226 | -4.1855225 | -1.1375619 | 0.00085369 | 0.00292505 |
| ENSMUSG00000027612 | Mmp24 | 18.3205271 | 5.69352832 | -12.626999 | -1.1394804 | 0.00085225 | 0.00292109 |
| ENSMUSG00000029868 | Trpv6 | 27.0747925 | 9.6732531 | -17.401539 | -1.1406437 | 0.00018832 | 0.00073675 |
| ENSMUSG00000028613 | Lrp8 | 2723.93779 | 1137.13041 | -1586.8074 | -1.141353 | 9.3214E-08 | 6.1383E-07 |
| ENSMUSG00000071414 | - | 60.0071208 | 24.0814791 | -35.925642 | -1.1414014 | 3.6668E-06 | 1.9102E-05 |
| ENSMUSG00000024754 | Cemip2 | 10906.5365 | 4545.28217 | -6361.2544 | -1.1414383 | 1.1836E-07 | 7.6688E-07 |
| ENSMUSG00000063229 | Ldha | 141608.179 | 63430.7695 | -78177.41 | -1.1415006 | 1.415E-38 | 1.2712E-36 |
| ENSMUSG00000051723 | - | 7.68135336 | 1.28430951 | -6.3970439 | -1.1433847 | 0.00157225 | 0.00508478 |
| ENSMUSG00000098077 | - | 10.2061401 | 2.45479045 | -7.7513497 | -1.1457111 | 0.00154274 | 0.00500194 |
| ENSMUSG00000000248 | Clec2g | 30.0331441 | 9.49167365 | -20.54147 | -1.1465638 | 0.00070724 | 0.0024614 |
| ENSMUSG00000071516 | H2ac13 | 30.5069837 | 10.3713826 | -20.135601 | -1.1469293 | 0.00033017 | 0.0012304 |
| ENSMUSG00000038354 | Ankrd35 | 430.472357 | 183.229197 | -247.24316 | -1.1495726 | 2.9858E-10 | 2.7706E-09 |
| ENSMUSG00000096726 | - | 46.3937752 | 15.6362134 | -30.757562 | -1.1549516 | 0.00033595 | 0.00124977 |
| ENSMUSG00000091747 | - | 16043.7797 | 7096.55376 | -8947.226 | -1.156576 | 1.7605E-34 | 1.2154E-32 |
| ENSMUSG00000039563 | 2210406O10 | 74.0516983 | 29.9472203 | -44.104478 | -1.1578957 | 5.2085E-07 | 3.0829E-06 |
| ENSMUSG00000073674 | - | 236.229755 | 101.121372 | -135.10838 | -1.1610089 | 5.1876E-13 | 6.5917E-12 |
| ENSMUSG00000052384 | Nrros | 57.1527766 | 20.2944554 | -36.858321 | -1.161606 | 0.00013102 | 0.00052779 |
| ENSMUSG00000032122 | Slc37a2 | 647.38165 | 284.630927 | -362.75072 | -1.1619403 | 9.0384E-30 | 4.6485E-28 |
| ENSMUSG00000020019 | Ntn4 | 2728.14114 | 1208.51626 | -1519.6249 | -1.1663139 | 2.848E-80 | 1.6275E-77 |
| ENSMUSG00000022686 | B3gnt5 | 4.39068087 | 0 | -4.3906809 | -1.1664889 | 0.00024006 | 0.00092186 |
| ENSMUSG00000038963 | Slco4a1 | 4923.58175 | 2105.80026 | -2817.7815 | -1.1669463 | 5.7755E-14 | 8.1883E-13 |

| | | | | | | | |
|---|---|---|---|---|---|---|---|
| ENSMUSG00000071192 | Wfikkn1 | 41.3374909 | 15.4287884 | -25.908702 | -1.1670005 | 2.4728E-05 | 0.00011279 |
| ENSMUSG00000036752 | Tubb4b | 18394.5776 | 8102.25829 | -10292.319 | -1.1683294 | 1.7453E-48 | 2.5646E-46 |
| ENSMUSG00000097697 | 4833412C05 | 6.50294108 | 0.80389104 | -5.69905 | -1.1725604 | 0.00099816 | 0.00337511 |
| ENSMUSG00000024621 | Csf1r | 139.893393 | 51.0788269 | -88.814566 | -1.1754331 | 4.5584E-05 | 0.00019901 |
| ENSMUSG00000083840 | - | 10.6490576 | 2.55506496 | -8.0939926 | -1.1772766 | 0.00106501 | 0.00357881 |
| ENSMUSG00000028212 | Ccne2 | 4406.33109 | 1933.85919 | -2472.4719 | -1.1780611 | 9.0042E-75 | 4.1163E-72 |
| ENSMUSG00000057388 | Mrpl18 | 6487.32298 | 2831.96293 | -3655.3601 | -1.1780852 | 5.701E-41 | 5.8935E-39 |
| ENSMUSG00000058914 | C1qtnf3 | 307.626007 | 118.659442 | -188.96657 | -1.1784038 | 4.1532E-06 | 2.143E-05 |
| ENSMUSG00000091844 | Gm8251 | 28.4944745 | 7.8816706 | -20.612804 | -1.1786982 | 0.00080747 | 0.00278245 |
| ENSMUSG00000088219 | - | 120.61002 | 49.4378911 | -71.172128 | -1.1806712 | 6.7171E-09 | 5.2204E-08 |
| ENSMUSG00000060096 | - | 44.4190863 | 16.9819931 | -27.437093 | -1.1814796 | 4.1851E-06 | 2.1583E-05 |
| ENSMUSG00000064634 | Gm22620 | 71.3976097 | 27.1383829 | -44.259227 | -1.1825187 | 7.222E-06 | 3.5869E-05 |
| ENSMUSG00000032053 | Pou2af1 | 13.8682243 | 3.40373578 | -10.464489 | -1.1882433 | 0.00096235 | 0.0032653 |
| ENSMUSG00000097189 | - | 20.8080248 | 6.70159944 | -14.106425 | -1.1903569 | 0.0002696 | 0.0010229 |
| ENSMUSG00000085165 | - | 8.1605982 | 1.59983612 | -6.5607621 | -1.1939059 | 0.0010273 | 0.00346453 |
| ENSMUSG00000022325 | Pop1 | 1948.49765 | 841.348251 | -1107.1494 | -1.1952273 | 1.5245E-46 | 2.0234E-44 |
| ENSMUSG00000007888 | Crlf1 | 3752.83702 | 1617.50539 | -2135.3316 | -1.1982122 | 2.3414E-46 | 3.068E-44 |
| ENSMUSG00000097739 | - | 21.3816155 | 6.66473702 | -14.716878 | -1.2023014 | 0.00023939 | 0.00091947 |
| ENSMUSG00000058743 | Kcnj14 | 10.9687312 | 2.48861456 | -8.4801167 | -1.2046794 | 0.00082207 | 0.00282754 |
| ENSMUSG00000085341 | - | 13.8192497 | 3.62757039 | -10.191679 | -1.2059873 | 0.00063408 | 0.00222673 |
| ENSMUSG00000011752 | Pgam1 | 25687.1563 | 10985.1363 | -14702.02 | -1.2090694 | 1.829E-47 | 2.5253E-45 |
| ENSMUSG00000001988 | Npas1 | 35.2209417 | 12.9268412 | -22.294101 | -1.2109605 | 6.6103E-06 | 3.3038E-05 |
| ENSMUSG00000091866 | - | 23.2995356 | 2.46611135 | -20.833424 | -1.2114428 | 0.00053465 | 0.00190951 |
| ENSMUSG00000030584 | Dpf1 | 402.974414 | 164.659613 | -238.3148 | -1.2119617 | 4.7512E-12 | 5.4361E-11 |
| ENSMUSG00000030170 | Wnt5b | 3866.37 | 1642.7131 | -2223.6569 | -1.2122089 | 1.6332E-35 | 1.1999E-33 |
| ENSMUSG00000072082 | Ccnf | 4328.99954 | 1850.63014 | -2478.3694 | -1.2156137 | 5.4625E-78 | 2.9572E-75 |
| ENSMUSG00000092056 | - | 32.9678156 | 11.5980675 | -21.369748 | -1.2204362 | 2.006E-05 | 9.3031E-05 |
| ENSMUSG00000044814 | Olfr543 | 8.13446357 | 1.49285674 | -6.6416068 | -1.2234796 | 0.00076693 | 0.00265386 |
| ENSMUSG00000039391 | Ccdc81 | 15.8731826 | 3.74902435 | -12.124158 | -1.2239067 | 0.00063642 | 0.00223392 |
| ENSMUSG00000082433 | - | 11.1871609 | 2.41865078 | -8.7685102 | -1.2271808 | 0.00068337 | 0.0023872 |
| ENSMUSG00000020583 | Matn3 | 33.1436376 | 10.6004347 | -22.543203 | -1.230618 | 0.00012231 | 0.00049542 |
| ENSMUSG00000045903 | Npas4 | 19.1615221 | 4.8907133 | -14.270809 | -1.2339042 | 0.00046896 | 0.00169641 |
| ENSMUSG00000081093 | - | 24.8029611 | 7.26447061 | -17.538491 | -1.2398129 | 0.0002069 | 0.00080445 |
| ENSMUSG00000048776 | Pthlh | 542.333055 | 221.361052 | -320.972 | -1.2417438 | 9.2825E-19 | 2.1312E-17 |
| ENSMUSG00000064340 | - | 16.301651 | 4.50255348 | -11.799098 | -1.2443809 | 0.00027045 | 0.00102574 |
| ENSMUSG00000049233 | Apoo-ps | 249.629162 | 49.8782837 | -199.75088 | -1.2469859 | 0.00060209 | 0.00212712 |
| ENSMUSG00000036875 | Dna2 | 1134.19627 | 455.7849 | -678.41137 | -1.2481384 | 7.6679E-15 | 1.1932E-13 |
| ENSMUSG00000048520 | Fbxl13 | 12.6086447 | 3.15992978 | -9.4487149 | -1.2541311 | 0.00040024 | 0.00146665 |
| ENSMUSG00000073672 | - | 38.1929022 | 13.3094478 | -24.883454 | -1.2552418 | 4.5903E-06 | 2.3502E-05 |
| ENSMUSG00000095159 | - | 47.1235209 | 16.7005465 | -30.422974 | -1.2562955 | 2.8764E-06 | 1.5235E-05 |
| ENSMUSG00000074662 | - | 7.48905827 | 1.17147448 | -6.3175838 | -1.2564613 | 0.00053923 | 0.00192354 |
| ENSMUSG00000022498 | Txndc11 | 3549.32605 | 1470.55802 | -2078.768 | -1.2578029 | 1.4716E-66 | 5.2196E-64 |
| ENSMUSG00000097254 | C430042M11 | 122.884306 | 47.3266614 | -75.557644 | -1.2613721 | 3.4095E-10 | 3.1368E-09 |
| ENSMUSG00000025961 | 4933402D24 | 18.2495525 | 3.48546036 | -14.764092 | -1.2689294 | 0.000476 | 0.00171945 |
| ENSMUSG00000077391 | Gm24336 | 15.0499312 | 3.29820557 | -11.751726 | -1.2762498 | 0.00037872 | 0.00139376 |
| ENSMUSG00000064168 | - | 41.3327034 | 14.349675 | -26.983028 | -1.2772305 | 2.1235E-06 | 1.1454E-05 |
| ENSMUSG00000059430 | Actg2 | 166.47887 | 65.3408039 | -101.13807 | -1.2777575 | 1.7548E-15 | 2.9301E-14 |
| ENSMUSG00000090101 | Snhg9 | 65.2784187 | 23.6458106 | -41.632608 | -1.2821172 | 9.6424E-08 | 6.3375E-07 |
| ENSMUSG00000030491 | Tdrd12 | 27.1709401 | 7.88026219 | -19.290678 | -1.2838477 | 8.3539E-05 | 0.00034845 |
| ENSMUSG00000034981 | Parm1 | 5341.14082 | 2148.07774 | -3193.0631 | -1.285688 | 8.5739E-34 | 5.6898E-32 |
| ENSMUSG00000082632 | - | 11.3812578 | 2.12728342 | -9.2539744 | -1.2858791 | 0.00040021 | 0.00146665 |
| ENSMUSG00000005649 | Cabp5 | 13.3171648 | 2.72434753 | -10.592817 | -1.2876246 | 0.00035861 | 0.00132828 |
| ENSMUSG00000040666 | Sh3bgr | 250.968625 | 96.2193731 | -154.74925 | -1.2900616 | 1.6847E-12 | 2.028E-11 |
| ENSMUSG00000020649 | Rrm2 | 57894.8124 | 23540.6754 | -34354.137 | -1.2906559 | 1.345E-119 | 3.459E-116 |
| ENSMUSG00000082967 | - | 17.3932684 | 3.80500845 | -13.58826 | -1.297994 | 0.00029594 | 0.0011136 |
| ENSMUSG00000031125 | 3830403N18 | 39.4674621 | 12.7895803 | -26.677882 | -1.3015529 | 6.4169E-06 | 3.2143E-05 |
| ENSMUSG00000028671 | Gale | 1335.94895 | 529.031272 | -806.91768 | -1.3131336 | 6.1229E-45 | 7.3661E-43 |
| ENSMUSG00000095042 | - | 60.7111414 | 16.4660333 | -44.245108 | -1.3151944 | 9.386E-05 | 0.00038859 |
| ENSMUSG00000001642 | Akr1b3 | 48540.1074 | 19295.1935 | -29244.914 | -1.3162989 | 3.3678E-67 | 1.2372E-64 |
| ENSMUSG00000022797 | Tfrc | 40737.1012 | 16071.3469 | -24665.754 | -1.3253709 | 1.6659E-61 | 4.6946E-59 |
| ENSMUSG00000019987 | Arg1 | 75.714983 | 25.9381427 | -49.77684 | -1.3256071 | 2.4927E-07 | 1.5497E-06 |
| ENSMUSG00000038271 | Iffo1 | 63.0357702 | 16.5746794 | -46.461115 | -1.3282342 | 9.1922E-05 | 0.0003811 |
| ENSMUSG00000029503 | P2rx2 | 15.8632987 | 3.06412864 | -12.79917 | -1.3337641 | 0.00022868 | 0.00088229 |
| ENSMUSG00000043782 | Bicdl2 | 13.4118195 | 2.69354456 | -10.718275 | -1.3342343 | 0.00021958 | 0.00084991 |
| ENSMUSG00000068617 | Efcab1 | 168.879229 | 61.005938 | -107.87329 | -1.3358992 | 2.0889E-10 | 1.9784E-09 |
| ENSMUSG00000056055 | Sag | 17.0381075 | 4.40205426 | -12.636053 | -1.3361089 | 7.917E-05 | 0.0003313 |
| ENSMUSG00000069302 | H2ac12 | 19.0593165 | 4.661434 | -14.397883 | -1.3366539 | 0.00011277 | 0.00046031 |
| ENSMUSG00000088544 | Mir1945 | 8.81439448 | 0.96564513 | -7.8487493 | -1.3409702 | 0.00019168 | 0.00074868 |
| ENSMUSG00000022582 | Ly6g | 207.703962 | 69.2877412 | -138.41622 | -1.347287 | 2.317E-07 | 1.447E-06 |
| ENSMUSG00000027907 | S100a11 | 16249.6855 | 6231.32589 | -10018.36 | -1.3601277 | 2.4272E-49 | 3.9947E-47 |
| ENSMUSG00000045005 | Fzd5 | 3059.86698 | 1162.31136 | -1897.5556 | -1.3732017 | 8.1711E-49 | 1.2452E-46 |
| ENSMUSG00000076999 | - | 30.8294208 | 5.74080232 | -25.088618 | -1.3737014 | 0.0001427 | 0.00057057 |
| ENSMUSG00000020793 | Galr2 | 11.125539 | 1.42668644 | -9.6988526 | -1.3848178 | 0.00013731 | 0.00055138 |
| ENSMUSG00000085022 | Gm5860 | 33.9791058 | 8.69182736 | -25.287278 | -1.3936894 | 2.8473E-05 | 0.00012864 |
| ENSMUSG00000062704 | - | 43.74245 | 11.7438364 | -31.998614 | -1.3981401 | 1.2313E-05 | 5.8991E-05 |

| | | | | | | | |
|---|---|---|---|---|---|---|---|
| ENSMUSG00000005481 | Ddx39a | 14957.214 | 5597.34334 | -9359.8706 | -1.4074063 | 7.512E-112 | 1.405E-108 |
| ENSMUSG00000062417 | - | 82.0023585 | 25.5799367 | -56.422422 | -1.4107022 | 1.3044E-07 | 8.4197E-07 |
| ENSMUSG00000048988 | Elfn1 | 26.5204384 | 6.69731205 | -19.823126 | -1.4191868 | 1.4354E-05 | 6.8163E-05 |
| ENSMUSG00000047139 | Cd24a | 68995.4298 | 25116.7042 | -43878.726 | -1.4243626 | 6.6355E-39 | 6.1489E-37 |
| ENSMUSG00000088025 | - | 11.9870411 | 1.14766369 | -10.839377 | -1.4246874 | 7.4477E-05 | 0.00031351 |
| ENSMUSG00000083120 | - | 24.4901856 | 6.55908766 | -17.931098 | -1.4259654 | 6.6428E-06 | 3.3193E-05 |
| ENSMUSG00000063628 | - | 130.00842 | 42.6767801 | -87.33164 | -1.4283435 | 5.2394E-10 | 4.6802E-09 |
| ENSMUSG00000083907 | - | 93.5876179 | 31.4151232 | -62.172495 | -1.4336127 | 2.4669E-12 | 2.9233E-11 |
| ENSMUSG00000055271 | - | 30.4890177 | 7.57612318 | -22.912895 | -1.4468202 | 1.1777E-05 | 5.6673E-05 |
| ENSMUSG00000029657 | Hsph1 | 28817.224 | 10417.0288 | -18400.195 | -1.4478065 | 1.8464E-65 | 6.1266E-63 |
| ENSMUSG00000024742 | Fen1 | 569.465931 | 200.750306 | -368.71563 | -1.454439 | 2.0042E-29 | 1.0056E-27 |
| ENSMUSG00000035711 | Dok3 | 76.6339723 | 24.6638758 | -51.970097 | -1.4559649 | 1.9255E-10 | 1.8313E-09 |
| ENSMUSG00000078238 | - | 519.429344 | 181.058056 | -338.37129 | -1.4725071 | 4.5321E-33 | 2.8425E-31 |
| ENSMUSG00000004371 | Il11 | 403.267953 | 135.114429 | -268.15352 | -1.5113948 | 1.1105E-24 | 4.0149E-23 |
| ENSMUSG00000017832 | Hspb9 | 16.4822605 | 2.47372083 | -14.00854 | -1.5134131 | 2.9054E-05 | 0.00013116 |
| ENSMUSG00000026576 | Atp1b1 | 9747.49342 | 3325.00254 | -6422.4909 | -1.5144618 | 5.77E-42 | 6.2147E-40 |
| ENSMUSG00000083346 | - | 17.4927106 | 2.81280996 | -14.679901 | -1.5239253 | 2.1194E-05 | 9.7956E-05 |
| ENSMUSG00000069378 | Prdm6 | 599.099484 | 190.686405 | -408.41308 | -1.5328874 | 6.4538E-16 | 1.1252E-14 |
| ENSMUSG00000042607 | Asb4 | 24.5812368 | 5.34347588 | -19.237761 | -1.5362361 | 5.3559E-06 | 2.7158E-05 |
| ENSMUSG00000024512 | Dynap | 80.2745878 | 19.7931669 | -60.481421 | -1.5580528 | 4.5307E-07 | 2.7142E-06 |
| ENSMUSG00000071552 | Tigit | 2584.31808 | 848.322951 | -1735.9951 | -1.5583312 | 1.5223E-35 | 1.1265E-33 |
| ENSMUSG00000019851 | Perp | 623.770335 | 198.640222 | -425.13011 | -1.5650084 | 4.7723E-22 | 1.429E-20 |
| ENSMUSG00000093798 | - | 159.436211 | 49.3429127 | -110.0933 | -1.5793405 | 1.6602E-18 | 3.6963E-17 |
| ENSMUSG00000054215 | Sprr2k | 193.626317 | 44.6845151 | -148.9418 | -1.5886759 | 5.7943E-07 | 3.4116E-06 |
| ENSMUSG00000003849 | Nqo1 | 39.2853597 | 10.2602768 | -29.025083 | -1.5997432 | 6.651E-09 | 5.1749E-08 |
| ENSMUSG00000089997 | 1810020O05 | 31.9302692 | 6.24866619 | -25.681603 | -1.6045488 | 2.3751E-06 | 1.2701E-05 |
| ENSMUSG00000091971 | Hspa1a | 308.902268 | 96.488729 | -212.41354 | -1.6065149 | 4.3345E-28 | 1.9904E-26 |
| ENSMUSG00000015656 | Hspa8 | 156371.408 | 50260.7497 | -106110.66 | -1.6214758 | 1.201E-113 | 2.746E-110 |
| ENSMUSG00000051726 | Kcnf1 | 1844.23283 | 590.824974 | -1253.4079 | -1.6250206 | 3.333E-109 | 4.572E-106 |
| ENSMUSG00000033967 | Rnf225 | 132.591534 | 34.7840036 | -97.80753 | -1.6316853 | 6.0701E-10 | 5.3663E-09 |
| ENSMUSG00000082087 | - | 77.589577 | 20.9269494 | -56.662628 | -1.6419882 | 4.0182E-11 | 4.1331E-10 |
| ENSMUSG00000057163 | Prss2 | 26.4200346 | 4.64534131 | -21.774693 | -1.6779396 | 1.0869E-06 | 6.124E-06 |
| ENSMUSG00000021822 | Plau | 4285.70442 | 1284.13382 | -3001.5706 | -1.6892381 | 2.9477E-44 | 3.4851E-42 |
| ENSMUSG00000085715 | Tsix | 8.94093296 | 0 | -8.940933 | -1.7148317 | 7.9724E-07 | 4.6041E-06 |
| ENSMUSG00000031253 | Srpx2 | 165.035695 | 42.5422954 | -122.4934 | -1.7281771 | 1.7704E-13 | 2.3727E-12 |
| ENSMUSG00000084822 | - | 25.7926818 | 4.40178453 | -21.390897 | -1.7867135 | 1.0042E-07 | 6.5832E-07 |
| ENSMUSG00000071203 | Naip5 | 30.9222737 | 5.29966343 | -25.62261 | -1.812565 | 4.4639E-08 | 3.0878E-07 |
| ENSMUSG00000053613 | - | 251.163097 | 60.7588112 | -190.40429 | -1.8293198 | 8.0658E-16 | 1.3955E-14 |
| ENSMUSG00000023336 | Wfdc1 | 109.446883 | 24.241927 | -85.204956 | -1.9085864 | 2.5728E-15 | 4.2478E-14 |
| ENSMUSG00000090877 | Hspa1b | 2191.44827 | 560.23207 | -1631.2162 | -1.9364264 | 4.639E-102 | 5.613E-99 |
| ENSMUSG00000027375 | Mal | 3128.60195 | 660.868141 | -2467.7338 | -1.9946942 | 5.1129E-18 | 1.0866E-16 |
| ENSMUSG00000041986 | Elmod1 | 116.844667 | 23.5154439 | -93.329223 | -2.0058822 | 9.8203E-16 | 1.6835E-14 |
| ENSMUSG00000048455 | Sprr1b | 211.851658 | 38.8592206 | -172.99244 | -2.2157229 | 1.8976E-25 | 7.2829E-24 |
| ENSMUSG00000092837 | Rpph1 | 237.237785 | 30.0453193 | -207.19247 | -2.5719266 | 2.2756E-24 | 8.0298E-23 |
| ENSMUSG00000033737 | Fndc3c1 | 814.878673 | 34.5313669 | -780.34731 | -4.2919331 | 4.492E-136 | 2.31E-132 |
